# Characterization of novel pollen-expressed transcripts reveals their potential roles in pollen heat stress response in *Arabidopsis thaliana*

**DOI:** 10.1101/2020.08.20.258764

**Authors:** Nicholas Rutley, Laetitia Poidevin, Tirza Doniger, Richard Tillet, Abhishek Rath, Javier Forment, Gilad Luria, Karen Schlauch, Alejandro Ferrando, Jeffery Harper, Gad Miller

## Abstract

The male gametophyte is the most heat-sensitive of all plant tissues. In recent years, long noncoding RNAs (lncRNAs) have emerged as important components of cellular regulatory networks involved in most biological processes, including response to stress. While examining RNAseq datasets of developing and germinating *Arabidopsis thaliana* pollen exposed to heat stress (HS), we identified 66 novel and 246 recently-annotated intergenic expressed loci (XLOCs) of unknown function, with the majority encoding lncRNAs. Comparison to HS in cauline leaves and other RNAseq experiments, indicated 74% of the 312 XLOCs are pollen-specific, and at least 42% are HS-responsive. Phylogenetic analysis revealed 96% of the genes evolved recently in *Brassicaceae*. We found that 50 genes are putative targets of microRNAs, and that 30% of the XLOCs contain small open reading frames (ORFs) with homology to protein sequences. Finally, RNAseq of ribosome-protected RNA fragments together with predictions of periodic footprint of the ribosome P-sites indicated that 23 of these ORFs are likely to be translated. Our findings indicate that many of the 312 unknown genes might be functional, and play significant role in pollen biology, including the HS response.

## INTRODUCTION

High temperatures have profound harmful effects on plants’ reproduction, often causing severe damages to complete loss of crops (1-3). The male gametophyte is thought to be the most sensitive to heat stress (HS) compared to all other organs and tissues in most plants, including the female gametophyte (4-6). The male gametophyte exists as a short cell lineage, beginning with the completion of the pollen mother cell meiosis within the anther, producing four haploid microspores. These microspores undergo an asymmetric cell division (pollen mitosis I) to produce a vegetative cell and generative cell. The generative cell divides once more (pollen mitosis II), giving rise to two identical sperm cells, followed by a maturation stage of the pollen in preparation for dehiscence. At pollination, pollen lands on a receptive stigma and then grows a tube that is guided towards the ovule. In species that underwent one round of pollen mitosis in the anther, the mitotic division of the generative cell takes place during pollen tube growth. Upon arrival at the ovule, the sperm cells are released from the pollen tube and go on to fertilize the egg and central cell (7). These delicate stages of pollen developmental are extremely vulnerable to HS, with even a short moderate- or mild chronic heat stress causing increase pollen abortion, reduced fitness, tube growth arrest, or even tube rupture (8,9). Also, the expression of many heat shock proteins (HSPs) and heat shock transcription factors (HSFs) is poor in pollen compared to other plant cells (6,10). These findings have contributed to the perception that pollen lack a robust HS response (HSR).

RNAseq experiments in pollen from different species indicated that HS has a profound impact on pollen gene expression and suggested pollen may have a different HSRs than other types of plant cells (6,11-14). The first RNAseq reports in Arabidopsis and maize mature pollen identified a significant number of novel transcribed loci, including transcripts with homologies to known proteins and long noncoding RNAs (lncRNAs) (15,16). Interestingly, the reproductive tissues of maize, including pollen and embryo sac, had more examples of lncRNA expression than any other tissues characterized (15). LncRNAs are transcripts exceeding 200 nucleotides in length that lack open reading frames longer than 100 amino acids. This somewhat arbitrary limit distinguishes lncRNAs from small non-coding RNAs such as microRNAs (miRNAs), small interfering RNAs (siRNAs), small nucleolar RNAs (snoRNAs), and other short RNAs (17). LncRNAs are generally polyadenylated and often have highly tissue-specific expression (18). LncRNAs mainly include intergenic ncRNAs (lincRNAs), intronic ncRNAs (incRNAs), and natural antisense transcripts (NATs) that overlap with coding regions. They are associated with a broad range of biological processes, including plant development and stress response, influencing gene expression by acting as molecular scaffolds, decoys or target mimics of microRNAs (miRNAs), and small interfering RNA precursors (18,19). As molecular scaffolds, lncRNAs may bind both DNA and protein recruiting regulatory components such as chromatin modulators to specific gene loci. As decoys, some lncRNAs may bind transcription factors to prevent them from interacting with DNA to induce target gene expression. Functional analyses of several plant lncRNAs demonstrated their profound involvement in plant development and physiology, including that of the male gametophyte (18,19). For example, a single polymorphism change in the sequence of long-day– specific male-fertility–associated RNA (LDMAR) in rice alters the secondary structure of the RNA molecule and it ability to function, causing defective anthers and pollen grains resulting in male sterility (20).

Many lncRNAs contain small open reading frames (smORFs) that can be translated into small polypeptides (>100 amino acids), also known as micropeptides or microproteins, with growing evidence supporting they are biologically functional, regulating target genes in *cis* or *trans* (19,21). The pollen-specific Zm908 lncRNA gene in maize encodes microproteins that interact with profilin 1, sequestering it from binding actin filaments. Overexpression of Zm908 caused developmental defects in maize pollen development and reduced germination. (22). The potential involvement of lncRNAs in cytoplasmic male sterility (CMS) has been recently suggested, however conclusive evidence directly implicating lncRNAs with CMS is still missing (23). An extensive data survey of RNAseq experiments in several plants, including wheat, maize, rice, and Arabidopsis, showed that a relatively large proportion of lncRNAs are responsive to abiotic stresses (24-26). The expression of HSFB2a, which is essential for the fertility of both the female and male gametophytes in Arabidopsis, is controlled by heat stress-induced NAT (27). The study of lncRNAs in plants is still an emerging area of research with the function of most lncRNAs awaiting discovery (18). Thus, research is needed to identify additional functional examples of lncRNAs involved in the HSR in both vegetative tissues and male gametophyte.

The latest complete *Arabidopsis thaliana* reference genome annotation, Araport11v4, significantly expanded the number of genes compared with the earlier TAIR10 annotation, including a massive increase in the number of long noncoding RNAs (lincRNAs and NATs) from 259 to 3559 genes. The updated annotation version also included 508 intergenic novel transcribed regions (28,29) and resulted from an assembly of tissue-specific RNAseq libraries from 113 datasets, which also included pollen. However, the transcriptome coverage and number of expressed genes in these pollen datasets were relatively small, including only 6,301 pollen expressed genes (29).

In this study, we report on the identification of 312 uncharacterized polyadenylated expressed loci (XLOCs) from three independent RNAseq experiments with Arabidopsis pollen exposed to HS. Sixty-six of those pollen XLOCs are entirely novel. Ninety two percent of the 312 XLOCs encode for lncRNAs, and 73% were not present in other sporophytic tissues RNAseq datasets, suggesting they are pollen-specific. Also, most of the XLOCs show differential expression in pollen matured under HS. Phylogenetic classification revealed that more than half of these genes are *A. thaliana*-specific, but 9 are highly conserved in eudicots. We identified 50 of the XLOCs as putative targets of microRNAs, suggesting potential involvement in gene expression regulation. Many of the identified lncRNAs contain smORFs for potential microproteins with homology to protein sequences. Lastly, ribosome elongating footprint analysis of ribosome protected RNAseq data showed that 23 of the ORF-containing transcripts are very likely to be translated. Our findings support the perception that pollen has a unique HSR and provide multiple targets for functional analyses of new pollen-specific genes with the potential of profoundly impacting pollen development and physiology.

## MATERIALS AND METHODS

### Plant material, growth conditions and pollen collection

*Arabidopsis thaliana* (Col-0) were grown in 16:8 light regime and at 21°C for five weeks until flowering had established. Pollen was harvested from open flowers using a customized pollen vacuum wand (30). The customized pollen vacuum wand consisted of nylon mesh filters (Membrane Solutions, USA) of three grades; 80 µm, 40µm, and 10 µm. Pollen deposited on the 10-micron filter was washed off using 0.3 M mannitol, pelleted, frozen in liquid nitrogen, and stored at −80C until required. After vacuuming, plants were then moved to a heat stress regime during the light photoperiod, consisting of a stepwise increase in temperature from the beginning of the light period from 22°C to 37°C over 6 hours, holding at 37°C for 2 hours, then a decrease to 16°C over 6 hours. Plants were maintained at 16°C during the night (Figure S1). This heat stress regime was repeated for 3 days, and pollen was collected on the morning of the 4^th^ day. In parallel to pollen sampling, cauline leaves representing control and heat stress conditions were collected from the same plants. Cauline leaf control samples were collected on the same day as pollen control samples, while heat-stressed cauline leaf samples were collected at the heat stress maximum (38°C) on the 3^rd^ day of heat stress.

### RNA isolation, library preparation, and sequencing

For RNA-Sequencing: Total RNA was extracted from pollen and cauline leaf samples using RNA was extracted using the TRizolTM reagent (Life Technologies, Carlsbad, CA) according to the manufacturer’s recommendations). RNA was shipped to Beijing Genomics Institute (BGI, China) for multiplexed Illumina HiSeq 2000 paired-end sequencing. Sequencing was performed at 2 × 50 bp for all samples.

### Sequence QC and novel gene prediction

Sequence pairs were trimmed and filtered using Trimmomatic v. 032 (31). After filtering sequences to remove multiplexing barcodes and sequencing adapters using NGS QC Toolkit (v.2.3), sequence pairs were aligned to TAIR10 *A. thaliana* reference genome using the spliced aligner tool Tophat2 (V.2.0.13) and Bowtie 2 (v.2.2.4).

Unannotated transcripts (i.e., transcripts not mapping to the TAIR10 reference genome) were identified and mapped using the in-house services of the Nevada INBRE Bioinformatics Core at the University of Reno (USA). Briefly, genome alignments generated by Tophat2 were processed using the Cufflinks package (v.2.2.1) reference annotation-based transcript (RABT) assembly method. Predictions from the independent Cufflinks runs were combined using the cuffmerge command to produce a single set of predicted transcripts.

Generating sequence alignments; sequence pairs were aligned to the *A. thaliana* genome using HISAT spliced read alignment tool (v.0.1.6; (32)). Genomic coordinates of the putative novel transcripts were combined with the exon coordinates of all known TAIR10 genes (33), as annotated in Ensembl build 27 and supplied to HISAT. Novel genes were then assigned intergenic XLOCs identifiers. For expression quantification, the number of read-pairs aligned to each gene was counted using the featureCounts tool from the subread package (v. 0.30; (34)). Read pairs were counted just once per pair, summarized to gene loci. Ambiguous or multiply-aligned read pairs were excluded from count totals. Reads were then normalized to Reads Per Million reads (RPM).

To determine which of the unannotated transcripts had then been annotated in the Araport11 genome annotation, we used IntersectBed from the Bedtools suite (Quinlan and Hall 2010) to compare Araport11 with our novel transcripts. Those transcripts that overlapped with an Araport11 annotated gene were considered annotated in Araport 11.

The bioinformatics protocols for the GP_HS libraries followed the same procedures as described (35) with a few changes. First, during the cleaning steps the non-coding RNAs were not bioinformatically removed. The GTF file used for the htseq-count included the identified XLOCs both in forward and reverse orientation. Finally, the htseq-count used the ‘nonunique all’ function to attribute a read to both RNAs when the mapping overlapped two RNAs.

### RNA-seq data analysis

Transcript counts were filtered to exclude those with < 10 counts in all samples. Filtered count data were then normalized via the median ratio method (20979621). Differential gene expression between control and HS conditions was examined using DESeq2 [68]. Comparisons were considered using simple contrasts. Fold changes in gene expression were transformed to log_2_ fold change values. Correction for multiple testing was performed within each comparison to adjust for the false discovery rate. XLOCs with ≥ 1-log_2_ fold changes and adjusted p-value ≤ 0.05 were considered to meet the standard significance threshold for this study.

### Validation of RNA-Seq data by quantitative PCR (Q-PCR)

For validation of gene expression using qRT-PCR: Total RNA was extracted from Col-0 wild type control inflorescences and following the heat stress regime using Trizol reagent (Thermo Fisher Scientific). First strand cDNA synthesis was performed using qScript Flex cDNA synthesis kit (Quantabio). qRT-PCR was performed in CFX96Connect (Bio-Rad). The gene HTR5 (AT4G40040) served as a reference gene. Relative normalized expressed was calculated using 2-^ΔΔCq^ method. Primer sequences are listed in Table S1.

### Phylostratigraphic analysis

To understand the conservation of these novel transcripts, we performed megablast at the NCBI using the novel transcripts as the query and requiring an E-value of <=1e-05. We ran the search twice using the nr/nt database once excluding hits within Brassicaceae and second time only searching within Brassicaceae but eliminating hits to Arabidopsis thaliana. We then extracted the full taxonomic lineage for each hit and were then able to assign hits to a given level in the phylogenetic tree.

### Other bioinformatics tools used

CANTATAdb (http://cantata.amu.edu.pl/) was used to identify conserved lncRNAs (36). psRNATarget V2 (2017 release) (http://plantgrn.noble.org/psRNATarget/) (37) was used to identify putative miRNA targets among novel genes. Hits were filtered using an expectation threshold of ≤ 3. IVG (v.2.4.3.) was used to visualize mapped reads to the Arabidopsis genome, importing Araport11 .bed files.

## RESULTS

### Mapping and identification of novel pollen-expressed transcripts

In a transcriptome experiment designed to identify differentially expressed genes that are pollen-specific and responsive to temperature stress, we compared the transcriptome of maturing pollen and cauline leaves exposed to heat stress (HS) (Rutley et al. in preparation; stress regime plot in Figure S1). Paired-end RNAseq data were generated from poly-adenylated RNA from Arabidopsis (Col-0) mature pollen (MP) and cauline leaves grown under control (22°C) or exposed to heat stress cycle for three days, referred to herein as MP_HS dataset and CL_dataset, respectively. The analysis of the RNAseq filtered raw data was performed using an in-house *de novo* assembly pipeline with the TAIR10 genome build as reference (Figure 1A; Materials and methods). We also similarly analyzed recently published RNAseq data from mature pollen developed under a diurnal cycle of hot day and cold nights and control conditions (13), referred to herein as MP_ Hot/Cold dataset. While interrogating the three RNAseq datasets, we assembled 400 transcriptional units (TUs, assigned with TCONS identifiers) originating from 312 intergenic expressed loci (XLOCs) that did not overlap with any loci annotated in TAIR10, with 41 of the XLOCs having two or more splice variants TCONS. (Table S2, S3). About 80% (246/312) of these transcripts were later annotated in the Araport11 database as expressed genes mostly without any other functional annotations (i.e., “unknown genes”) (v1.10.4, release 06/2016; (29)), referred herein as ‘Araport recent’ genes. The remaining 66 genes are yet to be annotated (Figure 1B, Table S3). The average depth of the MP_HS, MP_Hot/Cold, and CL_HS datasets was 63.7, 30.7, and 60.7 million reads, respectively (Table S4). Notwithstanding, while reads for all 312 XLOCs with overlapping coordinates were present in the MP datasets (all but one in MP_Cold/Hot dataset), 190 were absent in the CL dataset, suggesting these transcripts are enriched in pollen (Figure 2, Table 1, Table S4).

**Table 1:** Summary of novel genes from three independent pollen HS RNAseq experiments. Clear reads refer to the filtered raw data after removing adapter sequences, contamination, and low-quality reads. PEXs, pollen expressed XLOC genes, hPEXs, high PEX DEXs, differentially expressed XLOC genes, PSXs, pollen-specific XLOC gene.

| | Total<br>clean<br>reads<br>(million) | Mean<br>reads per<br>sample<br>(million) | PEXs<br>(rpm >1) | hPEXs<br>(rpm > 10) | DEXs<br>(Log2 ≥ 1,<br>P ≤ 0.05) | PSXs | Pollen-specific<br>DEX |
| --- | --- | --- | --- | --- | --- | --- | --- |
| MP_HS | 382.0 | 63.7 | 312/312 | 269 | 37/312 | 230/312 | 21/230 |
| MP_Hot/<br>Cold | 184.0* | 30.7* | 311/312 | 213 | 96/311 | 229/311 | 73/229 |
| GP_HS | 609.7 | 101.6 | 61/312 | 9 | 12/61 | 30/61 | 10/30 |

**Figure 1:**
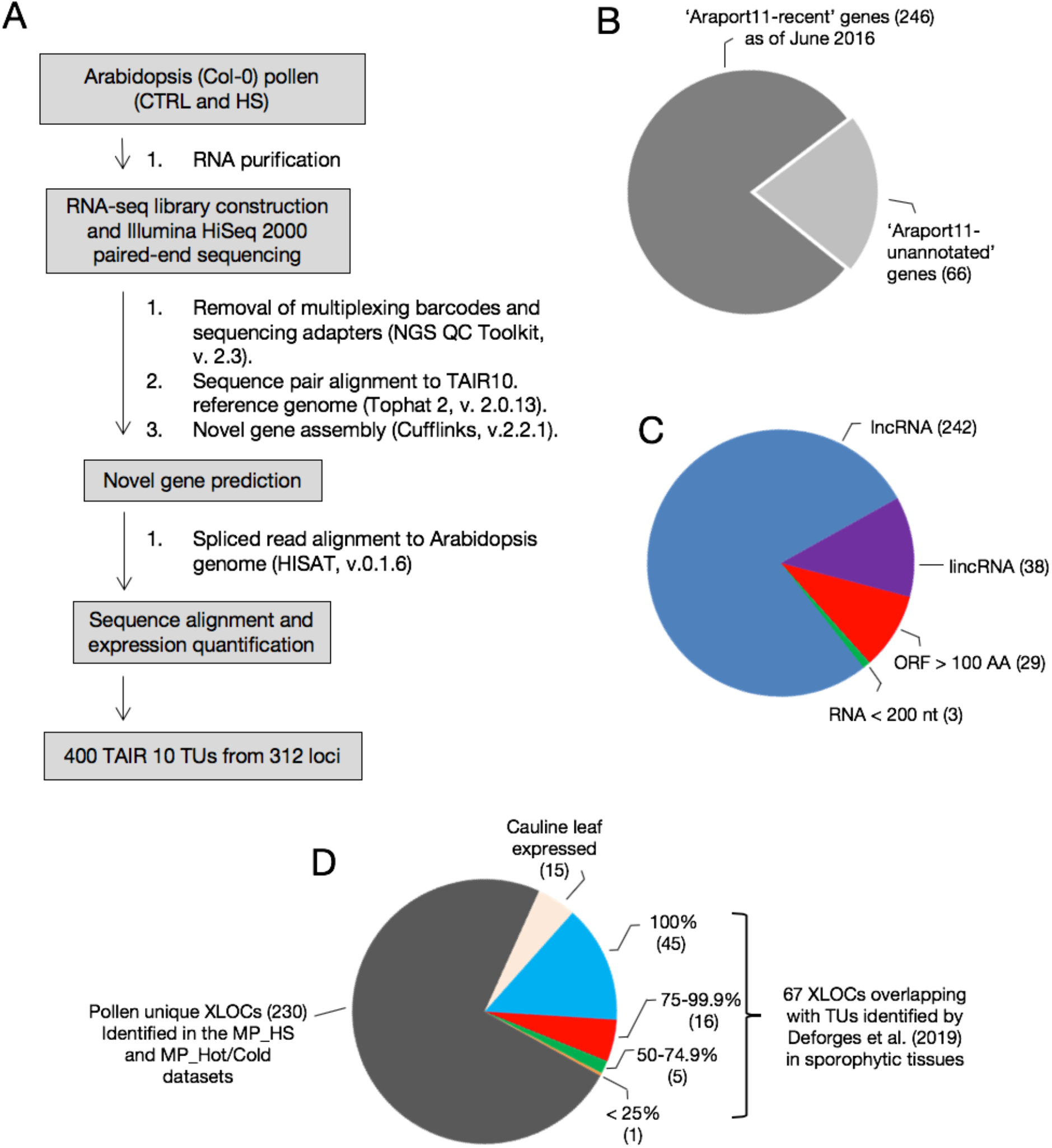
Identification and characterization of novel transcriptional units (TUs). A. The workflow pipeline for the identification of TUs in MP_HS and MP_Hot/Cold RNA-seq data. B. Pie chart of the distribution between expressed loci (XLOCs) annotated at the previous update of Araport11 (‘Araport-recent’) and XLOCs that remained as unannotated Araport11 genome annotation (v1.10.4, release 06/2016). C. Breakdown of XLOCs by class: long noncoding RNA (lncRNA), long intergenic noncoding RNA (lincRNA), Open reading frame (ORF) > 100 amino acids, and RNA length < 200 nucleotides. D. 230 TUs identified in pollen libraries from MP_HS, and MP_Hot/Cold that were not found among MP_HS cauline leaf libraries or sporophytic lncRNAs from Deforges et al. (2019).

**Figure 2:**
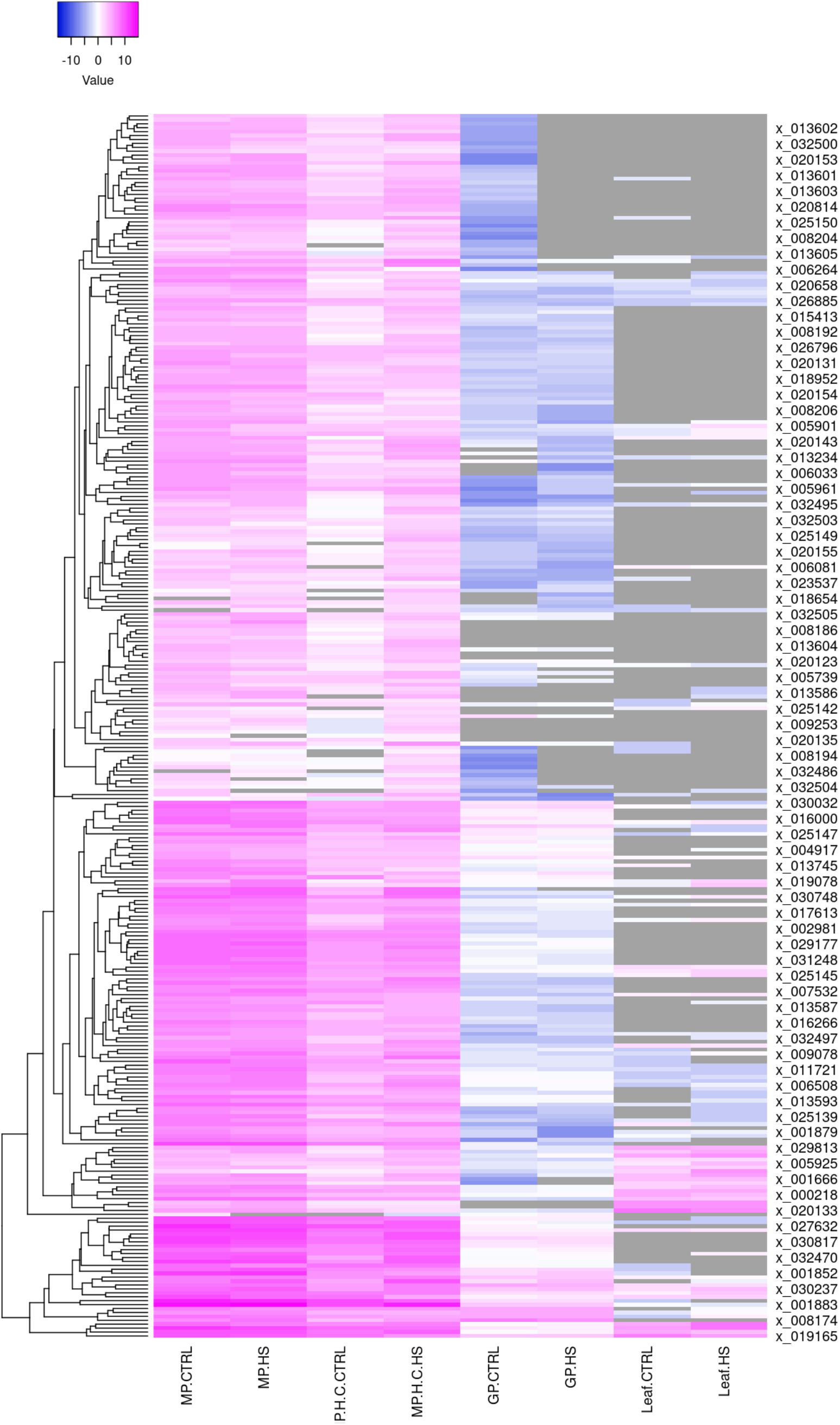
Heatmap of the expression abundance of XLOCs in the four RNAseq datasets under control and HS conditions.

### The majority of the newly discovered XLOC genes encode pollen-specific long noncoding RNAs

Based on the criteria of RNA longer than 200 nucleotides or having ORF(s) shorter than 100 amino acids (18), 286 of the XLOCs were categorized as encoding long noncoding RNAs (lncRNAs) and 29 XLOCs contain ORFs ranging between 100 to 379 amino acids, suggesting they may be protein coding genes (Figure 1C and Table S3). Indeed 19/29 overlap with gene models in Araport11 (Table S3). Three remaining TUs transcribe RNA shorter than 200 nucleotides, and so do not meet the criteria of lncRNAs (Figure 1C).

A subclass of lncRNAs are referred to as long intergenic noncoding RNAs (lincRNAs), defined as being located 500 or more bp away from annotated protein-coding genes, not encoding transposable elements (TEs), and not overlapping with natural antisense transcripts (NATs; (19). As the MP_HS and MP_ Hot/Cold RNAseq datasets are non-strand-specific, we employed only the two former criteria mentioned above, searching for lincRNAs. Among the 286 lncRNAs, 65% are located > 500 bp from the nearest gene model and 51% show overlaps with TEs (Table S3). Only 38 XLOCs (13.3%) meet two of these additional requirements for long intergenic noncoding RNA (referred to here as putative lincRNA; Figure 1C).

Similarly, Deforges and co-authors recently identified 862 new lncRNA genes in RNAseq experiments in Arabidopsis seedlings that had no prior annotation in TAIR10, with about half them later independently annotated in Araport11 v1.10.4 database release before their publication (38). The dataset of Deforges et al. 2019 included RNAseq from whole Arabidopsis seedlings or roots and shoots from 12 experimental conditions, including high or low phosphate concentrations, and treatments with the plant hormones auxin (indole acetic acid, IAA), abscisic acid (ABA), methyl-jasmonate (MeJA) or the ethylene precursor 1-aminocyclopropane-1-carboxylic acid (ACC). We, therefore, compared the coordinates of the newly identified TUs encoding genes in our pollen datasets with those of the genes identified in the Deforges et al. (2019) datasets. Of the 312 XLOCs identified in the MP_HS dataset, 14.4% and 7.1% overlapped completely or partially, respectively, whereas 230 of the pollen-identified expressed XLOC loci were not present within the sporophytic RNAseq datasets of (38) or our cauline leaves libraries (Figure 1D, Table S5). Thus, this comparison between the pollen and sporophytic RNAseq datasets suggested that 78.5% of the identified XLOCs may exclusively express in pollen.

### The majority of the TUs are pollen-specific

We interrogated yet another pollen RNAseq dataset from an HS experiment in which pollen was germinated *in vitro* for 5 hours at 24°C or 35°C for 5 hours (GP_HS dataset;(35)). The GP_HS libraries are strand-specific, adding a higher order of resolution for producing gene models, providing us with the ability to exclude a potential expression of overlapping TEs on the opposite strand (Table S6). The average sequencing depth of the GP_HS sequencing was 101.6 million reads per sample (Table 1, Table S4). Yet, 75 of the 312 XLOCs were either absent or had a negligible abundance of mean read per million (RPM) lower than 0.05 in both the control and HS samples (Figure 2, Table S5). Thus, given the depth of the RNAseqs, the differences in the presence of the XLOCs between the MP and GP experiments reflect the differences in the physiological and developmental phases of mature dry versus hydrated germinating pollen. Moreover, all the 312 XLOCs detected in mature pollen had the expression level >1 RPM, whereas in the GP_HS, only 61 had > 1 RPM (Figure 3A, Table 1, Table S5).

**Figure 3:**
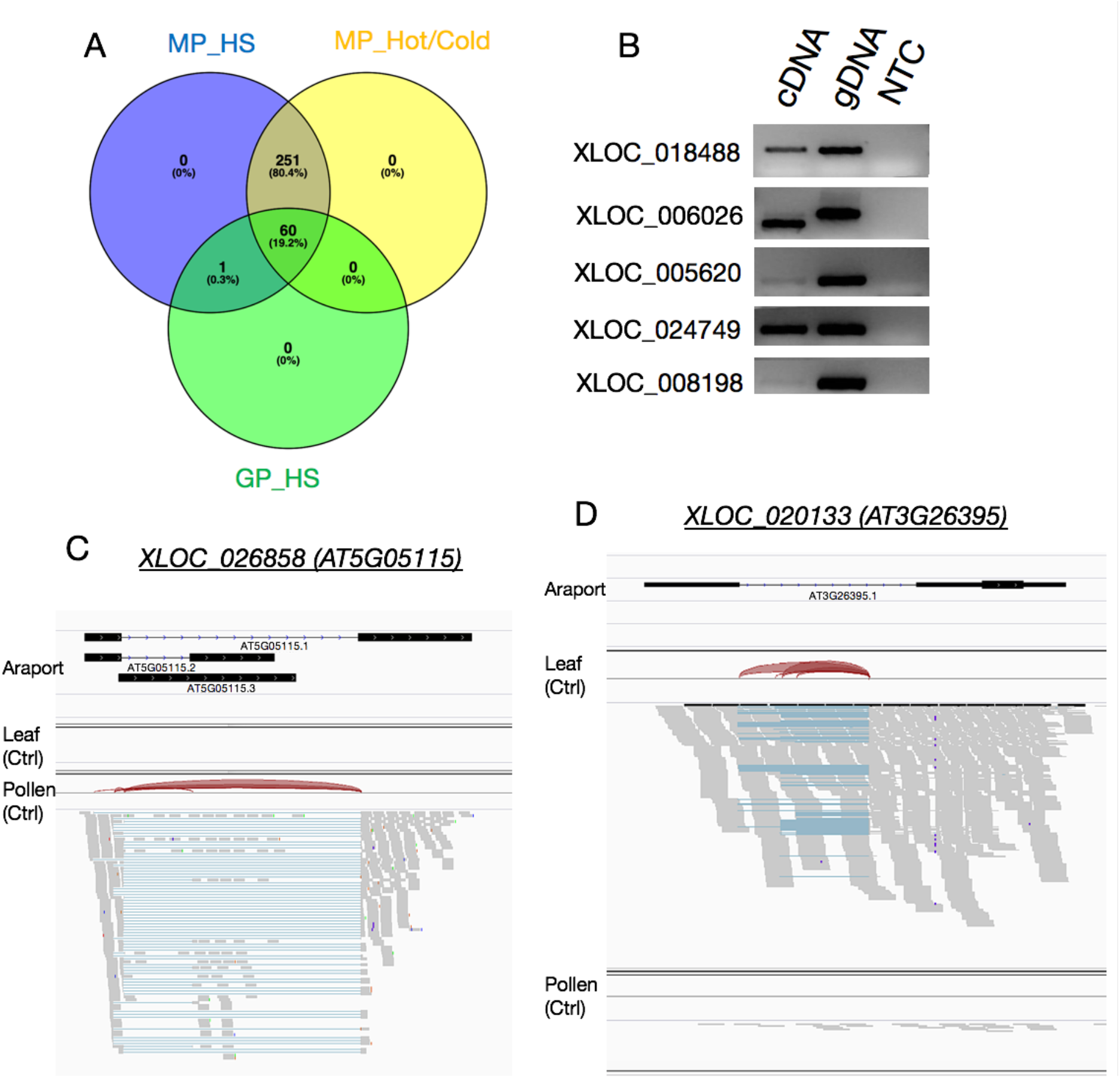
Comparison between the XLOCs expression between the three pollen RNAseq experiments and validation of expression. A. Venn diagram of the presence of pollen expressed XLOCs (PEXs; RPM > 1) in the three pollen datasets. C and D. Visualization from IGV of pollen-specific (C) and cauline leaf-specific (D) XLOCs. Reads are shown aligned onto the reference Araport11 reference genome. Splice junctions are represented by brown arcs from the beginning to the end of the junction. gDNA, genomic DNA. NTC, no template control.

Therefore, we set the threshold indicating <u>p</u>ollen <u>e</u>xpressed <u>X</u>LOC genes (PEXs) to 1 RPM. We verified the expression of TUs in the RNAseq datasets using endpoint PCRs for five selected stress-responsive PEX with cDNA pooled from inflorescences grown under control and heat stress (Figure 3B).

Among pollen expressed genes, 269 and 213 from MP_HS and MP_Hot/Cold datasets, respectively, were shown higher than 10 mean RPM. All of these genes had none or low expression level (RPM < 1) in cauline leaves (Table 1, Table S5). As an example, using the Integrative Genome Viewer (IGV) tool, we verified that XLOC_0265858 lacks detectable reads in cauline leaves (Figure 3C). In contrast, as a non-pollen-exclusive example, we show IGV of XLOC_020133, one of only 8 PEXs with higher expression in cauline leaves compared to pollen (Figure 3D). Thus, we defined a total of 230 pollen-specific XLOCs (PSXs) as those genes not present and with RPM < 1 in either the CL_HS or Deforges et al. 2019 (38) RNAseq datasets. Interestingly, out of 61 PEXs in germinating pollen only 30 are pollen-specific (Table 1).

### Conservation among land plants

To search for conserved genes among the 312 XLOCs, we used CANTATAdb 2.0, a database of putative lncRNAs predicted from hundreds of RNAseq libraries from 39 species covering a broad diversity of land plant species (39). Query entry for each of the 312 XLOC main TCONs identified homologs for only 30 genes in *A. lyrata* and an additional four genes in three species of other *Brassicacaea* family members (Table S7). All the identified conserved genes were expressed in both MP libraries, and 6 were also expressed in germinating pollen. In addition, around half (18/34) were pollen-specific. No putative lncRNA homologs were identified from more evolutionarily diverse species, suggesting that the majority of TUs encoding loci identified here originated within the *Brassicaceae* lineage (Table S7). The relatively low number of lncRNA homologs hits from CANTATAdb 2.0 may result from the libraries used to construct the database, as it is curated from transcriptomic studies not including pure pollen or pollen-enriched samples.

Therefore, we conducted a more comprehensive search for homologous genes using BLASTn megablast within the standard databases nucleotide collection (nr/nt). We then used the BLASTn results in a phylostratigraphic approach to determine the phylogenetic origin of each XLOC, assigning each XLOC to a phylostratum according to the oldest phylogenetic node to which the XLOC can be traced (Figure 4). We found that the vast majority of XLOCs (301 of 312) evolved within the Brassicaceae family, including all novel yet unannotated TUs. Eleven PEXs matched homologous sequence in earlier divergent species, including 4 XLOCs assigned to the Magnoliopsida node, which have homologs in monocots (XLOC_013590, XLOC_008192, XLOC_006026, and XLOC_030237). Among the 9 phylogenetically oldest PEXs, belonging to Eudicotyladones and Magnoliopsida, 6 are pollen-specific.

**Figure 4.**
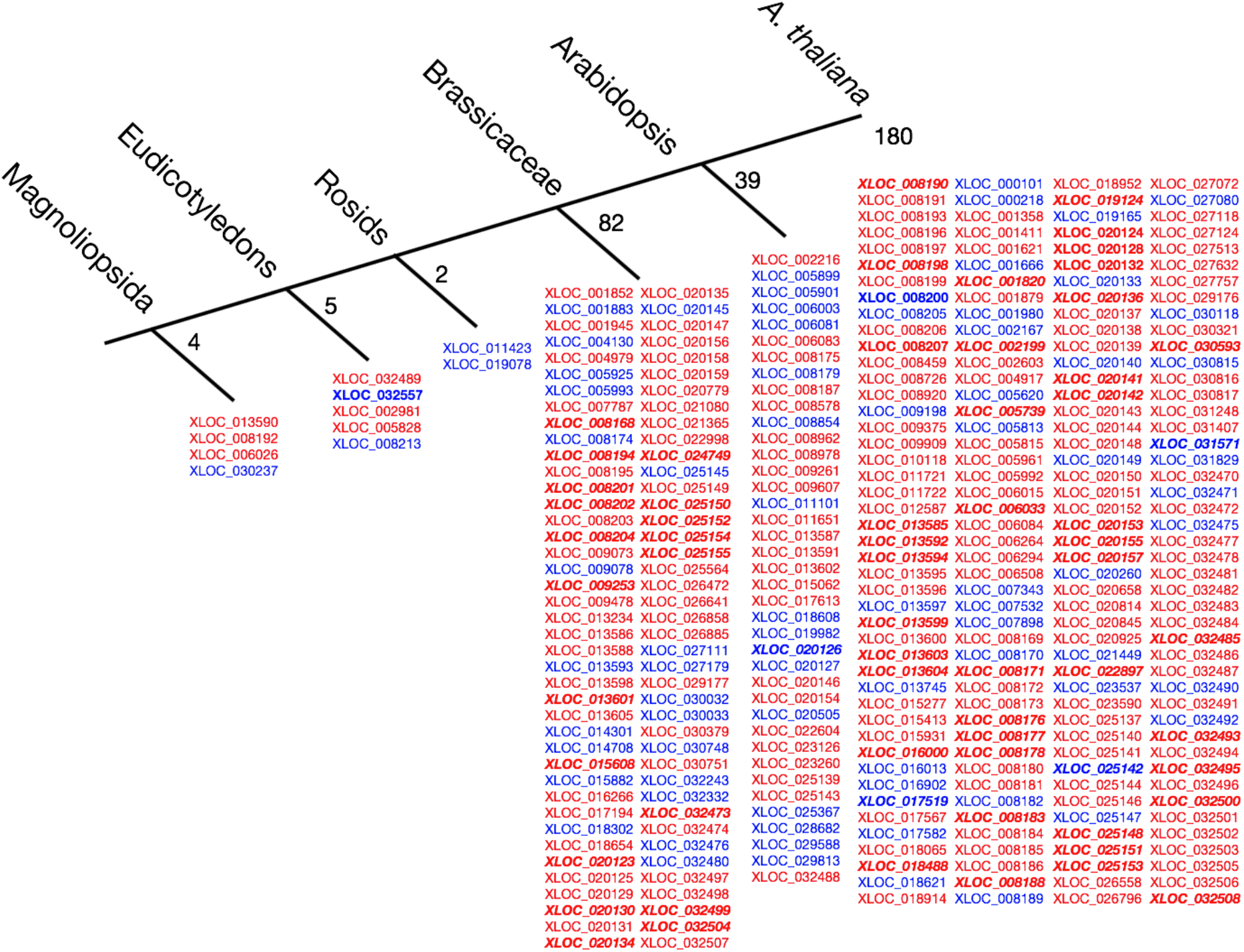
Phylostratigraphy of the 312 PEXs. Each XLOC was assigned to a phylostratum corresponding to the oldest phylogenetic node to which the gene could be traced. In red, pollen-specific XLOCs. In blue, XLOCs also expressed in sporophytic tissue in Deforges et al. (2019) or cauline leaf from MP_HS. In bold italics, novel unannotated genes.

### Heat stress-driven differential expression in developing pollen

The induction of HSPs and HSFs in cauline leaves was several folds higher than in the three pollen datasets and the abundance (RPM) of the transcripts was 1-2 order of magnitude higher, in most cases (Figure S2), in line with previous reports indicating that the heat stress response (HSR) in the male gametophyte is not robust as in vegetative tissues (6,10). Yet, there were significant differences in the level of induction or lack of induction for some HSPs in pollen (Figure S2), which likely resulted from differences in the experimental setups (see materials and methods).

In each of the three independent pollen datasets, the majority of XLOCs were present in both control and heat stress conditions, having RPM values > 1 (Figure 5A-C). Only 10, 34 and 19 XLOCs in the MP_HS, MP_Hot/Cold and GP_HS, respectively, were specifically expressed in either one of the conditions with MP_HS. We then explored differential expression among the XLOCs in each of the three pollen datasets, comparing between control and HS conditions, using the criteria of a log fold change threshold of ≥ 1 or ≤ −1 and significance of adjusted p-value < 0.05. In the experiments in which the HS occurred during the pollen development, 37 and 97 of the <u>X</u>LOCs were deemed <u>d</u>ifferentially <u>e</u>xpressed (DEXs) in the MP_HS and MP_Hot/Cold datasets, respectively (Figure 5D,E). Nevertheless, a potentially higher percentage of the PEXs may be heat stress-responsive as an additional 12 and 11 genes passed the significance threshold for being differentially expressed in the MP_HS and MP_Hot/Cold datasets, respectively (Figure 5D, E; Table S5). Additionally, of the 55 and 98 PEXs in the MP_HS and MP_Hot/Cold datasets, respectively, showing an average fold change ≥ 2 but with significance above the adjusted p-value threshold (Figure 5D,E), 24 were common to both (Figure S3). In germinated pollen, 12 PEXs showed clear differential expression under the *in vitro* heat stress treatment, and an additional 5 passed only the fold change or significant threshold suggesting potential stress-responsiveness (Figure 5F). HS cycle of 22-37°C/16° day/night (MP_HS) resulted in similar numbers of up- and down-regulated genes (Figure 5G), whereas pollen developed under a hot day (peaking at 40°C at noon) and cold nights (1°C) had far more upregulated genes than down-regulated genes (Figure 5H). The differences in the pattern of DEXs (Figures 5G-I, 6A) potentially reflect the difference between the heat stress regimes and the physiological phase of the pollen during the experiment. Consequently, there was a relatively small overlap among the DEGs between the three HS experiments (Figure 6A, 6B, Figures S4-S6), suggesting the experimental setups impacted the level and the direction of expression in most of the HS-responsive XLOCs. Yet, 42% of the PEXs showed significantly responded to HS across all three experiments (130 DEXs). Only one DEX, XLOC_006026, was common to all three HS experiments and changed in the same direction in all the three experiments, suggesting that it might be a part of the core heat stress response in pollen. An additional 10 DEGs were common to both mature pollen datasets (Figure 6B, 6C). Nine of these 10 DEGs were showing similar expression trend following heat stress in the mature pollen datasets (Figure 6B). Of the four DEX from GP_HS that were also differentially expressed in one of the other datasets, three showed a similar response trend to heat stress.

**Figure 5:**
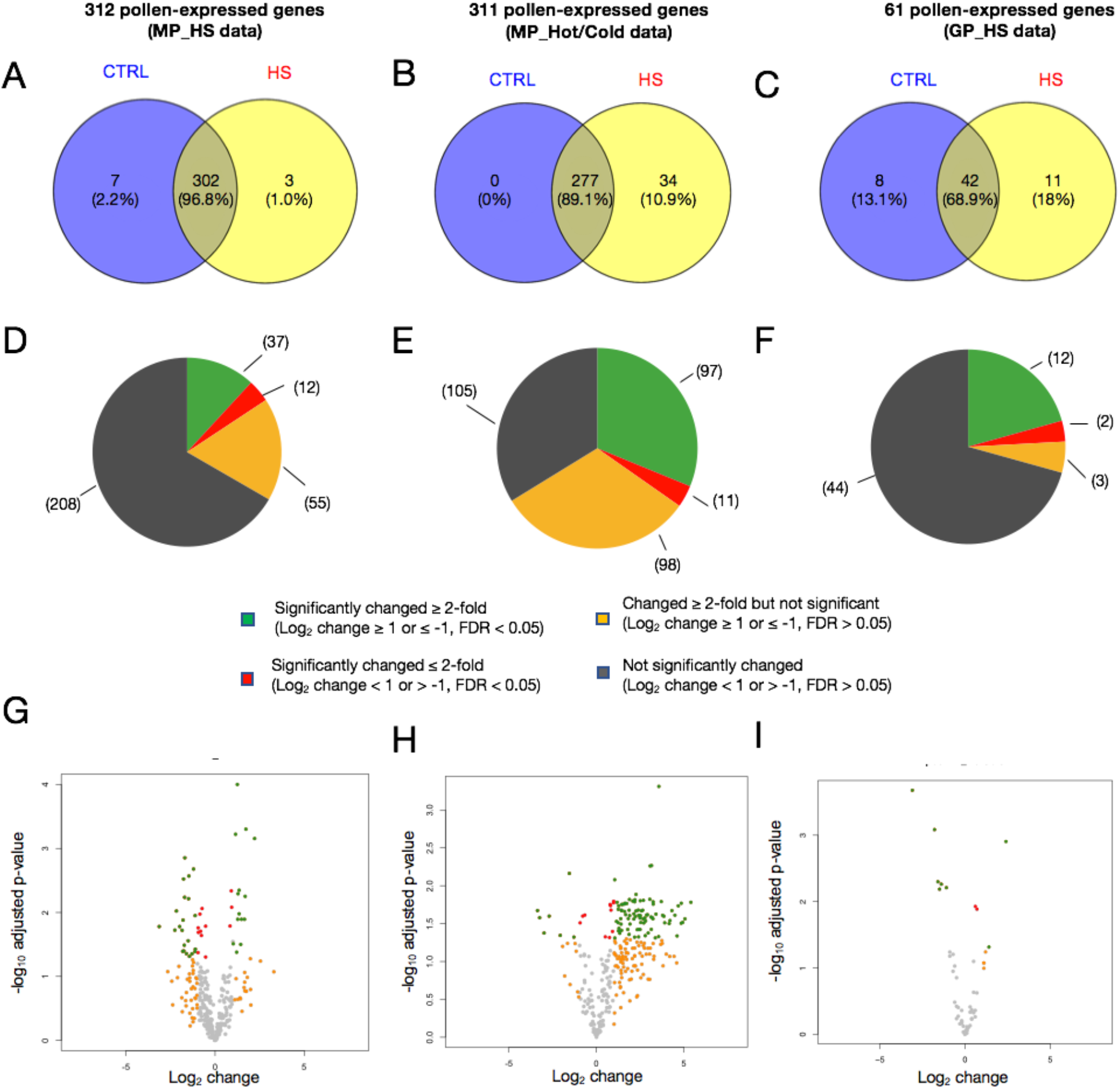
Differential expression of the PEXs in pollen RNA-seq libraries. A-C. Comparison between control and HS conditions of the PEXs (RPM > 1) in each of the pollen experiment. D-F. Pie charts showing the distribution of HS-responsive and not-responsive PEXs in each of the experiments. G-I. Volcano plots showing the dynamic range of the differential expression of the PEXs.

**Figure 6:**
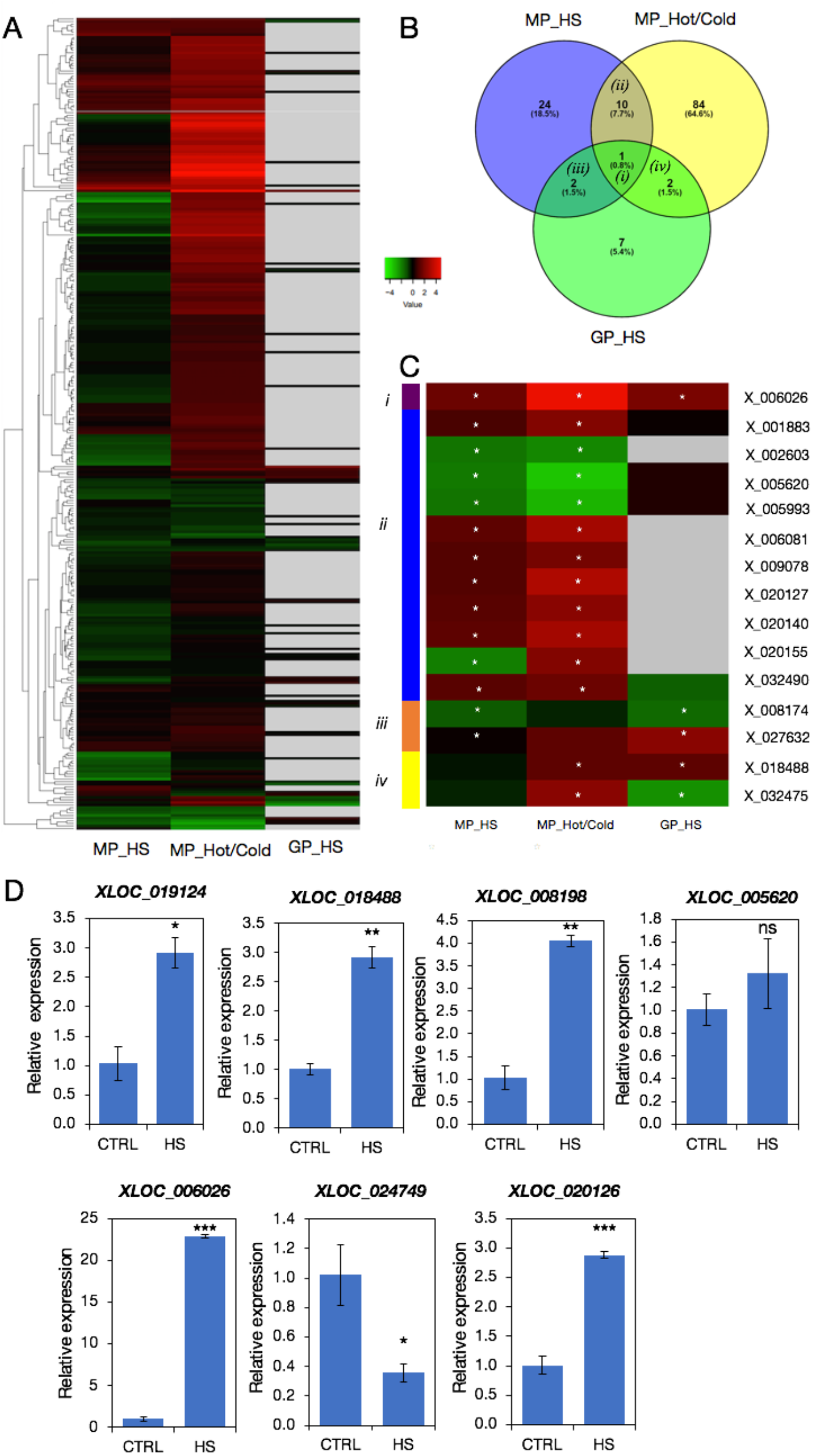
Comparison of differential expressed XLOCs (DEXs) in the three pollen experiments. A. Heatmap of log_2_ changes for all 312 XLOCs in MP_HS, MP_Hot/Cold, and GP_HS libraries. B. Venn diagram showing overlap of DEXs among the three pollen RNAseq datasets. C. Heatmap of the DEXs present in two or more of the RNAseq datasets (sections *i-iv* in B). Asterisks indicate DEXs with fold change ≥2 and adjusted p-value <0.05. D. Real-time PCR validation of differential expression of seven PEXs using cDNA from control and heat stress inflorescences. T-test p-value <0.05 and <0.01, are indicated with ^*^ and ^**^, respectively.

We further conducted quantitative real-time PCRs to validate differential expression during HS for 6 PSXs and one PEX that is not pollen-specific, XLOC_005620. To this end, we used cDNA from inflorescence of wild type Col-0 plants grown under control conditions or exposed to three days HS regime as in the MP_HS experiment (Figure S1). We show that 5 of 6 PSXs were upregulated in response to heat stress (Figure 6D). Among the six PSXs, the induction of *XLOC_006026*, which increased in all three experiments (Table S8), was the most prominent, with a 23-fold increase (Figure 6D). In comparison, *XLOC_019124, XLOC_ 018488, XLOC_008198*, and *XLOC_020126*, which were induced in the MP_Hot/Cold experiment (Table S8), only increased 3 to 4-fold during HS in the inflorescences (Figure 6D). In contrast, *XLOC_005620*, which significantly decreased in both mature pollen HS datasets (Figure 6C), did not change in the qRT-PCR experiment (Figure 6D), which might be due to non-gametophyte-specific expression. The real-time PCR results validate the DEXs identified in the datasets from three pollen experiments and support the notion that some of the PEXs induced during HS may play a role in the stress acclimation of the male gametophyte.

### XLOCs potentially regulated by miRNAs

A major mechanism of gene expression regulation is mediated through small RNAs, including microRNAs (miRNAs), which can impact mRNA degradation and translational repression. Similar to mRNA, lncRNAs can be targets of miRNAs, and act as miRNA decoys, sequestering specific miRNAs (40). To identify PEXs potentially targeted by miRNA binding, we used the psRNATarget tool employing the confidence cut-off threshold of 3.0 (Dai et al., 2018). We identified 50 PEXs as putative targets of one or more Arabidopsis miRNAs, the majority of which were predicted to be processed by cleavage rather than translational inhibition (Table S9; Figure 7A). Several miRNAs have been characterized as pollen-expressed under control conditions (41). While our RNA-seq libraries are unlikely to contain mature miRNAs, we looked for reads mapping to the pollen-expressed primary(pri)- miRNAs in the MP_HS dataset. We found several reads mapping to miR447a, an *A. thaliana*-specific microRNA (42), in the control treatment but non following HS. Correspondingly, while the expression of pri-miRNA447a decreased in HS, expression of its putative target XLOC_032495 (TCONS_00050386) increased in both MP_HS and MP_Hot/Cold (Figure 5B). miRNA447a is downregulated in cold-imbibed seeds vs. dry seeds (43) and hypoxia treated roots vs. control (44), pointing to it being a stress-regulated, and possibly stress-regulating, miRNA.

**Figure 7:**
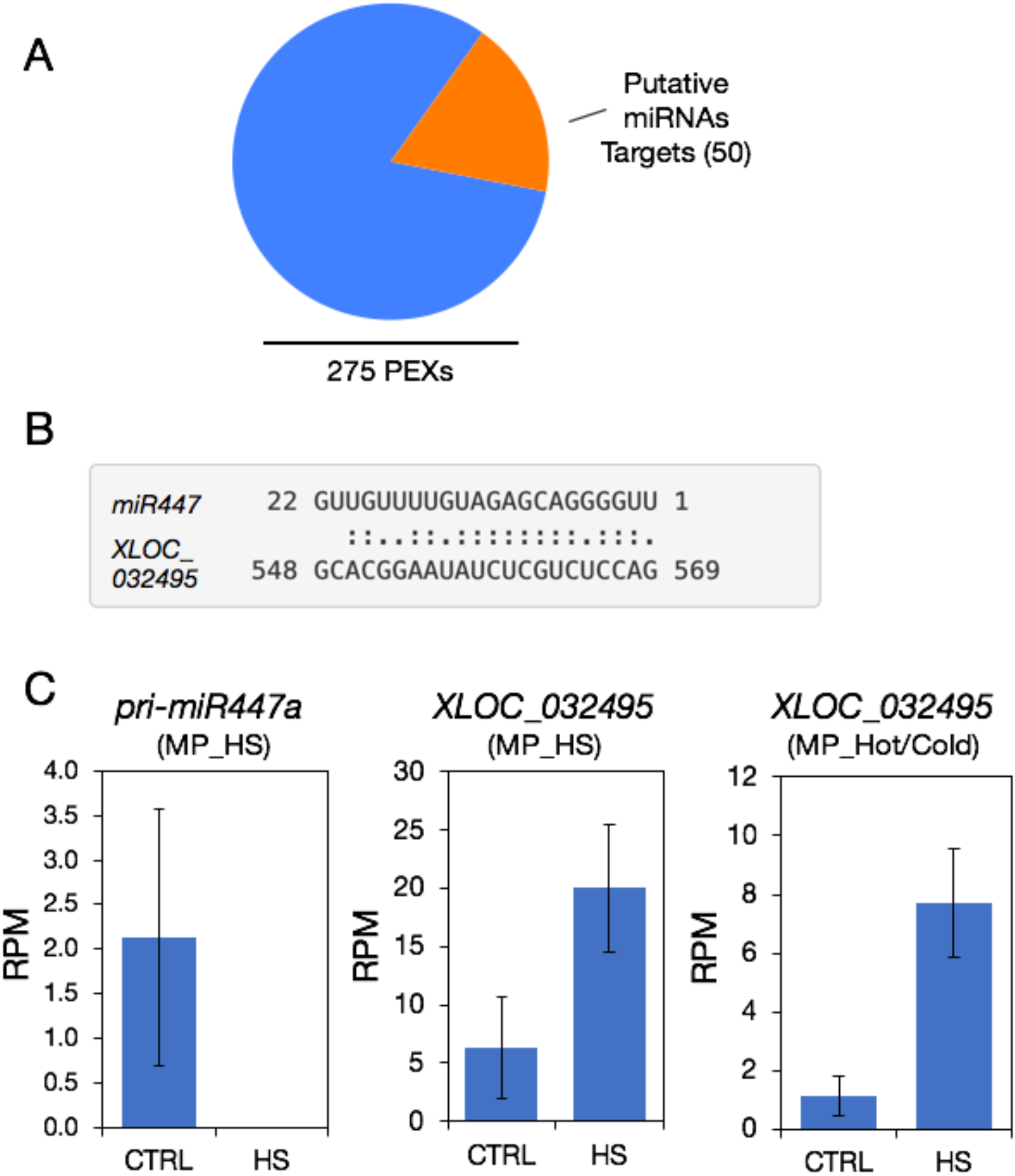
Prediction of miRNA targets among PEGs. A. Pie chart showing the proportion of PEGs predicted to be targets of miRNAs according to the psRNATarget tool. B. Pairwise alignment result of XLOC_032495 sequence predicted to be the target of miR447 by the psTARGET tool C. Expression of XLOC_032495 and its predicted targeting pri-miR477 in the mature pollen datasets.

### Periodic movement in ribosome-protected mRNA fragments (RPFs) indicates translation potential for some XLOCs

Ribosome profiling, also known as ribosome sequencing (Riboseq), is a powerful method for identifying transcripts engaged with ribosomes and allows for the likelihood prediction of transcripts undergoing *in vivo* translation. Parallel to the RNAseq of the GP_HS experiment, sequencing of ribosome protected fragments (RPF) was performed from the same *in vitro* HS experiment of *in vitro* germinated pollen (35). As the germinated pollen sequencing libraries are strand-oriented, we determined the coding strand for each of these XLOC genes (Table S6). We found consistent alignments for a total of 75 RPFs (RPF-GP_HS dataset) to 73 XLOCs within the RNA (GP_HS dataset), where XLOC_020146 and XLOC_005993 had RPF transcripts from both strands (Table S6). We focused on the 45 XLOCS with RPF reads values ≥ 1 RPM in either control and HS (35°C) or both, including XLOC_020146 plus and minus transcripts. (Table S10).

To gain further insight into the translational profiles revealed by the RPF values, we used the RiboWave v1.0 tool (45), a pipeline able to denoise the original RPF signal by extracting the <u>p</u>eriodic footprint of the <u>P</u>-sites (PF P-sites) of actively elongating ribosomes. The PF P-site values were decomposed into the 3 different frames, to determine which frame is likely translated by the ribosome. One limitation of the XLOCs annotations is that no clear definition of exons has been established that allows the discrimination between coding and untranslated regions (UTRs). Therefore, we could not exploit the statistical significance of translational prediction of the RiboWave algorithm. However, visual inspection of the denoised PF P-site tracking allowed us to define a minimum number of 5 PF P-sites in the same frame as a proxy of an increased likelihood for translation. From the initial 45 XLOCs, 23 XLOCs passed this filter and could be considered as very likely translated (Figure 8A, Table S10).

**Figure 8:**
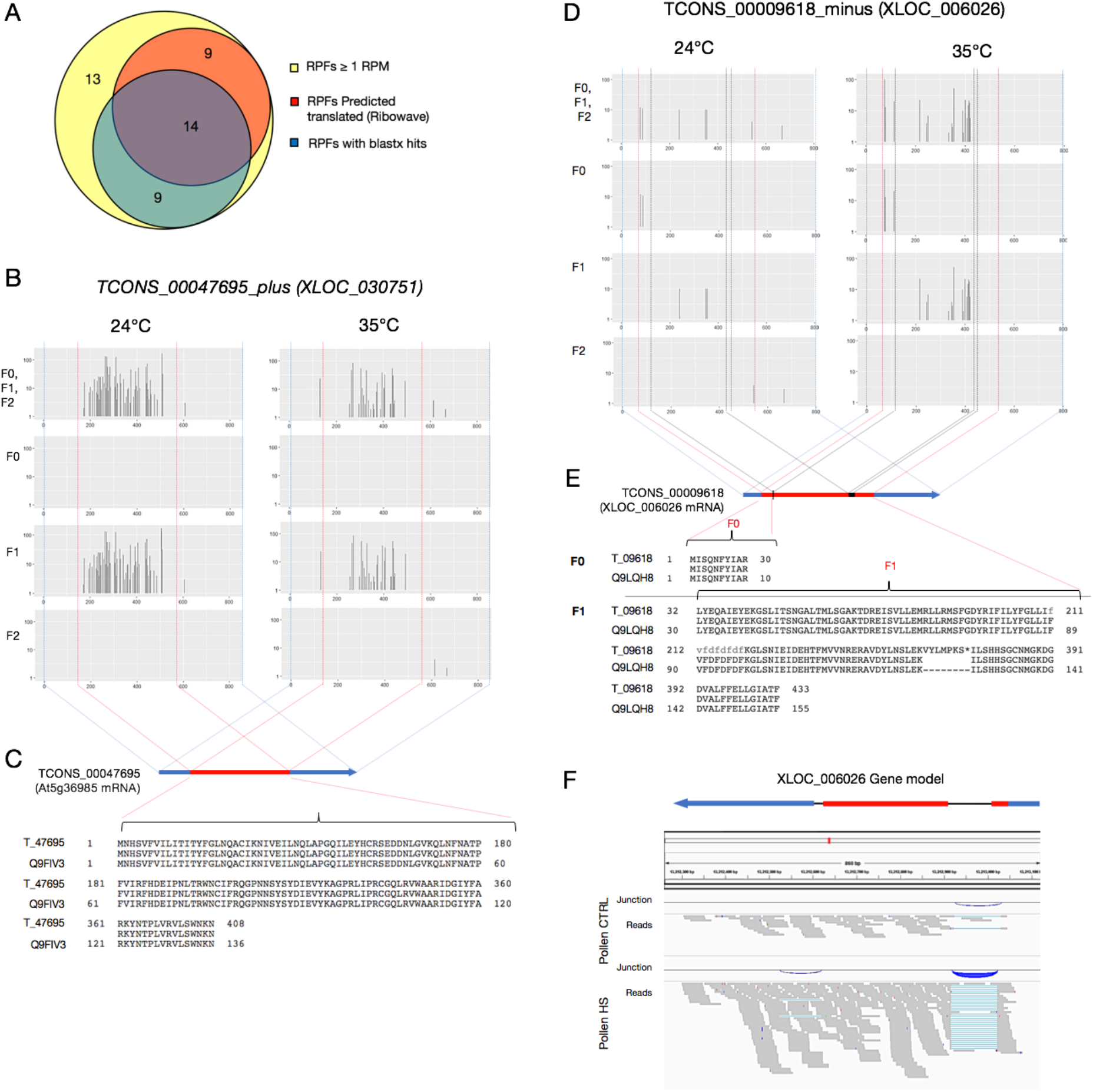
Periodic movement in ribosome-protected mRNA. A. Venn comparison of significant RPFs (RPM > 1) showing overlaps between RPFs predicted to move on ribosomes and those with ORFs showing homology to protein sequences. B-F characterization of the HS-induced PEXs that are predicted to be translated. B. Periodic footprint of the P-sites (PF P-sites) plot along the three frames (F0-F2) of the transcripts of XLOC_030751, generated by the RiboWave algorithm, indicating the activity of elongating ribosomes. D. PF P-sites plot along the three frames (F0-F2) of the transcripts of XLOC_006026. C and E show the mRNA model of XLOC_030751 (At5g36985) and XLOC_006026, respectively. Blue boxes – UTRs, red boxes, CDSs. The black box in E represents the unspliced intron-2. The dotted blue, red, and black dotted vertical lines in B and D indicate the transcripts’ borders, intron-exon junctions, and intron-2 borders, respectively. F. XLOC_006026 gene model corresponding to the IGV plot of the RNAseq reads in control and HS mature pollen, generated from the MP_HS dataset. Splice junctions are represented by blue arcs.

Additionally, a blastx (nr) for all 312 XLOCs (limited to *Arabidopsis thaliana*) identified partial hits (50-96% identity) or full match hits (≥ 97% identity) with sequences of predicted proteins (E value ≤ 1e-7) for l16 (37%) of the genes, of which 92 (30%) are lncRNAs (table S11). The proportion of blastx hits positives within the 75 RPFs XLOCs was 51%, and 60% (14 of 23) of the RPFs with the minimum number of 5 PF P-sites (Figure 8a, Table S10, S11), corresponded with the increased likelihood of these XLOCs being translated. Interestingly, 6 of these 14 XLOCs encode for predicted UniProt proteins, the putative products for two of which, XLOC_030571 and XLOC_032470, are homologs of pollen-specific self-incompatibility proteins.

In figure 8B and 8D, we present the PF P-site periodicity plots for two of these TUs, TCONS_00047695 (*XLOC_030571*) and TCONS_00009618 (*XLOC_006026*), both pollen-specific and HS-induced expressed loci (Table S5).

XLOC_030751, annotated in Araport11 as unknown gene AT5G36985, is intronless with an ORF (DOC S3) coding for self-incompatibility protein homolog 25 (SPH25) (Figure 8C). The periodicity plot for *XLOC_030571* clearly shows that only one reading frame (F1) is being translated, with the PF-P sites concentrated within the CDS along the transcript.

In contrast, the periodicity plot of *XLOC_006026* (Figures 3, 4, and 6), revealed PF-P sites in two frames, F0 and F1 (Figure 8D), suggesting that the TU contain an unprocessed intron. The blastx search identified a near-perfect identity with a phosphopyruvate carboxykinase (PEPCK) protein. The pairwise alignment of the translated XLOC_006026 mRNA with the protein sequence resulted in two ways split, with the first part showing 100% identity with the first ten amino acids. The second part is identical with the other 145 amino acids, except for seven extra amino acids and a stop codon in the C-terminus (Figure 8E), corresponding with the two frames being read (Figure 8D). Inspection of the XLOC _006026 CDS revealed two premature stop codons at +37 and +347; the first originated by what seems like an inaccurate exon1-intron1 site splicing event leading to frameshift, and the 2^nd^ by the inclusion of intron 2 (Doc S4, Figure 8F). Yet, although the vast majority of XLOC _006026 transcripts seem to be incorrectly processed, IGV output shows that few transcripts may still be fully and correctly spliced to produce the full-length protein-coding sequence CDS (Figure 8F).

## DISCUSSION

### Discovery of novel transcripts in Arabidopsis pollen

Novel genes that might only be expressed in a restricted context, such as a single cell type alone or in combination with stress, are only slowly being incorporated into reference genome annotations. The reason for overlooking these genes is partly because RNAseq studies tend to focus on known genes and exclude sequence reads that do not match an existing reference genome. Therefore, it is not surprising that even twenty years after the first draft of the Arabidopsis genome was published (46), additional new genes are still being discovered. The recent Araport11 annotation update released in 2016 added more than 600 and 5000 novel protein-coding and non-coding RNAs, respectively, relying predominantly on RNAseqs from sporophytic tissues (29). Comparison between microarray datasets of flower, roots, and leaves identified 32% of ∼3,700 lincRNAs in Arabidopsis as showing preferential expression in one of these organs (19). Epigenetic reprogramming taking place during plant sexual reproduction involves the overall reduction of DNA methylation in the germline during gametogenesis, allowing the potential expression of otherwise silent genetic elements, including TEs, endogenous protein coding genes, and introduced transgenes (47). It is therefore likely that the 312 PEX genes (pollen expressed intergenic e<u>x</u>pressed loci) reported herein have gone unnoticed for such a long time is due, in part, for their expression being mostly restricted to pollen (Figures 1-3), rather than the depth of the RNA sequencing (Table S4).

Compared with sporophytic tissues, such as root or leaf, which are composed of multiple cell types with different specialized functions, the male gametophyte is simple, containing a haploid vegetative nucleus and two sperm cells in tricellular pollen. This feature is advantageous for obtaining a large quantity of uniform cell-type-specific for any omics profiling. In comparison, obtaining a uniform sporophytic single-cell population from homogenized tissue for transcriptomics requires the isolation of tag-labeled cells by fluorescent activated cell sorter (FACS). Moreover, RNA extracted from the enriched isolated cells needs to be amplified to construct a library, which is biased towards transcripts with relatively high abundance, and transcripts with low copy number are often omitted (48).

In recent years RNAseq studies have added many new genes to the genome annotations of Arabidopsis and human, organisms completely sequenced two decades ago. Genomic studies based on various transcriptomic platforms identified thousands of lncRNAs in diverse animal and plant genomes, including over 58,000 in the human genome (49). As lncRNAs are more tissue-specific and expressed at lower levels than protein-coding mRNAs (49), it is plausible that future studies will identify many more yet unidentified cell-type-specific lncRNA loci in the Arabidopsis genome.

## THE MAJORITY OF THE LNCRNAS PEX GENES REFLECT THE AGE OF THE ARABIDOPSIS POLLEN TRANSCRIPTOME

Our finding that the vast majority of the 312 PEX genes belong to the Brassicaceae family (Figure 4), 180 of which are *A. thaliaana* specific, suggests these genes evolved relatively recently. These findings are in agreement with the feature of lncRNAs evolving more rapidly compared with protein-coding genes (18,49,50). The transcriptome of the male gametophyte appears to be enriched for recently evolved genes (i.e., lineage-specific genes and orphan genes lacking homologs in other lineages). These young genes include short peptides, intergenic transcripts, long noncoding RNAs (lncRNAs), and *de novo* genes at their transitory stages, also known as proto-genes (51). Similarly, 180 and 39 of the PEXs are species-specific and taxon-specific orphan genes, respectively (Figure 4), indicating they are young genes that recently evolved in the *Brassicaceae* lineage. The emergence of *de novo* genes from non-genic regions is relatively frequent in eukaryotes and may be part of a progressive evolutionary process that starts with the expression of intergenic regions to proto-genes and, finally, functional genes (51).

A survey of RNAseq datasets from 15 diverse flowering plant species indicated that transcription from unannotated intergenic regions is quite frequent in plants, but a large part of it is due to random non-functional transcriptional noise (52). Yet, predictions based on a computational analysis in Arabidopsis indicated that 38% of intergenic transcribed regions and 40% of the annotated ncRNAs have similar features to protein-coding or RNA genes and are likely functional (52). Since the majority of the 312 XLOC genes are relatively abundant (RPM > 10), expressed exclusively in developing/maturing pollen, and HS-responsive, we postulate that most, if not all, are functional genes. The function of these young genes may be diverse, including direct or indirect gene expression control. For example, the 50 PEXs we identified having miRNA targets may function as target mimics and molecular sponges that inhibit the action from a subset of miRNAs. Furthermore, 30% of the lncRNA PEXs contain smORFs that matches protein sequences. Indication of interaction with ribosomes, for those present in the riboseq dataset, and the demonstration of movement along the ribosome (active PF-P footprint) indicated that some of the lncRNAs produce micro-proteins and proteins that may be functional (DOC S2, Figure 8A, Table S11). Functional studies in a broad range of species from yeast to humans demonstrate that microproteins can functionally impact development and physiology (21). In the context of pollen, there are several examples of microproteins having a profound impact on pollen growth and fertilization, including self-incompatibility, tube growth, and sperm release inside the ovule (22,53-55).

The occurrence of XLOC_030751 and XLOC_032470 encoding for self-incompatibility related protein homologs among the 23 PEXs PF-P sites-positive very likely to be translated (Figure 8a, Table S10) lend support for the plausibility that some the ORF containing PEXs are functional. Interestingly, the expression of both self-incompatibility related-genes is relatively abundant in mature pollen, and they responded to the stress regime in the MP_Hot/Cold experiment (Table S5). Direct experimental evidence at the protein level is needed to determine their actual presence and whether they function in pollen–pistil interactions. Yet, they might be involved in switching from selfing to outcrossing mode of mating, which is employed in about half of the species in the Brassicaceae family (56).

### Potential involvement of PEX in pollen HSR

The limited HSR of pollen compared to cauline leaves indicated by the activation of *HSPs* (Figure S2) may account, at least in part, for its increased thermosensitivity, but also suggests pollen has distinct requirements for coping with high temperatures. A major conceptual difference between pollen and sporophytic cells is that the latter can respond to HS by reducing metabolism into ‘survival mode’, waiting for better growth conditions to resume growth or other physiological activities, whereas developing pollen and pollen tubes have a limiting window of time to properly mature and fertilize an ovule (13). We found that the expression of a large proportion (∼42%) of the 312 PEXs significantly changed ≥ 2-fold at least in one of the pollen HS experiments datasets (Figure 6B) and many of the other 58% are potentially HS-responsive (Figure 5E-F). A more comprehensive survey of the expression pattern of all these potential DEXs at different stages of pollen development along with different time points during an HS is required to determine their relevance to pollen HSR. However, given that the majority of the PEX specifically express in pollen (i.e., PSXs), it is likely that some of them either function in the pollen HSR, or are at least responsive to regulatory pathways that are controlling the HSR.

Several of the PSXs that might function at the core pollen HSR include XLOC_006026, which was induced in all three pollen experiments, and others that were increased in two of the pollen datasets; for example XLOC_027632, XLOC_002603 (Figure 6C), XLOC_009073 and XLOC_009261 (Figure S4), XLOC_008196, XLOC_008185 (Figure S5), XLOC_008726 (Figure S5). XLOC_006026 is intriguing as it is one of the most conserved PEXs that might code for a functional PEPCK, an enzyme involved in malate metabolism and gluconeogenesis. The closest PEPCK homolog of XLOC_006026 is AtPKC1 (AT4G37870), with 55.6% identity, was shown to function in malate metabolism in stomatal closure and drought tolerance in young Arabidopsis plants (57). PEPCK catalyses the reversible decarboxylation of oxaloacetate to yield phosphoenolpyruvate (PEP) and CO_2_ at the expense of ATP. Since PEP is a precursor for either the glucose and shikimate biosynthesis pathways or to pyruvate, it is situated at an important crossroads in plant metabolism, lying between organic and amino acids, lipids and sugars (58). Increased demand for carbohydrates during HS was suggested to contribute to the enhanced thermosensitivity of pollen (4). HS was shown to deplete the level of accumulated starch and soluble sugars in developing pollen, whereas HS tolerant tomato genotypes were better able to maintain pollen starch and sugar levels than sensitive genotypes (59-62). Therefore, the induction of PEPCK activity might help pollen accumulate specific sugars and better tolerate HS. It would therefore be highly interesting to test the potential impact of XLOC_006026 on pollen development and activity both during favourable conditions and elevated temperatures.

### Concluding remarks

It was recently suggested that lncRNAs make suitable environmental sensors or effectors to help plants adapt to changing environments, as their expression is extremely responsive to stresses and they evolve rapidly compared with protein-coding genes (18). However, functional studies of the involvement of lncRNAs in pollen development and physiology, let alone pollen acclimation to stress, are still at their earliest stages. The large proportion of genes encoding lncRNAs added to the Arabidopsis genome annotation in the previous Araport11 update, together with the thousands that have been identified in many other plant species in RNAseq experiments, raise the question about whether they have a function. Finding whether these lncRNA play a significant role and how do they perform their function is currently a major challenge in plant biology. Our identification in pollen and characterization of the novel and ‘Araport recent’ PEXs genes significantly provides a foundation for understanding potential functions for some of these lncRNAs in pollen-development and HSR.

## ACCESSION NUMBERS

The datasets of MP_HS and CL_HS have been deposited with the accession number PRJNA657848 in the NCBI Sequence Read Archive, https://www.ncbi.nlm.nih.gov/sra/.

The datasets of MP_Hot/Cold are available with accession number SRP110833 in the NCBI Sequence Read Archive, https://www.ncbi.nlm.nih.gov/sra/.

The Raw sequences derived from the Riboprofiling analysis and table containing counts and normalized TPM values for all A. thaliana genes have been deposited in the Gene Expression Omnibus database (http://www.ncbi.nlm.nih.gov/geo/) with the accession number GSE145795.

## SUPPLEMENTARY DATA

Table S1 – Realtime PCR primers

Table S2 – List of 400 TCONS and 312 XLOCs

Table S3 – Information about the 312 XLOCs

Table S4 – Number of clean reads for each RNAseq library

Table S5 – XLOCs expression and deferential expression values of all datasets

Table S6 –XLOC expression in the GP_HS RNAseq and Riboseq datasets

Table S7 – Identified conserved lncRNAs in the CANTATAdb

Table S8 – psRNATargets prediction

Table S9 – Log_2_ fold change of XLOCs tested by realtime PCRs

Table S10 – 44 XLOCs with RPM> 1 in the Riboseq data

Table 11 – BlastX results summary for all 312 XLOCs

Figure S1 – MP_HS RNAseq experiment regime plot

Figure S2 – Expression and log_2_ fold change level of HSPs and HSPs in pollen vs cauline leaves

Figure S3 – Venn diagrams showing overlap between XLOCs represented in the yellow and red sections in the pie charts in Figure 5D-F.

Figure S4 – Heatmap of specific DEXs in the MP_HS experiment

Figure S5 – Heatmap of specific DEXs in the MP_Hot/Cold experiment

Figure S6 - Heatmap of specific DEXs in the GP_HS experiment

Doc S1 – BLSTX of all 312 XLOCs against AT protein database

Doc S2 – 23 periodicity plots

Doc S3 – XLOC_030751 sequences

Doc S4 – XLOC_006026 sequences

## ACKNOWLEDGEMENT

We thank the Core Facility unit of the Faculty of Life Sciences in Bar Ilan University for providing qPCR services.

