## Supplemental Doc 2 for "Characterization of novel pollen-expressed transcripts reveals their potential roles in pollen heat stress response in *Arabidopsis thaliana*" (from another version of this preprint: version 2, 2020-08-26)

**TCONS_00003292_plus**

24°C

Raw P-sites | PF P-sites

F0, F1, F2

F0

F1

F2

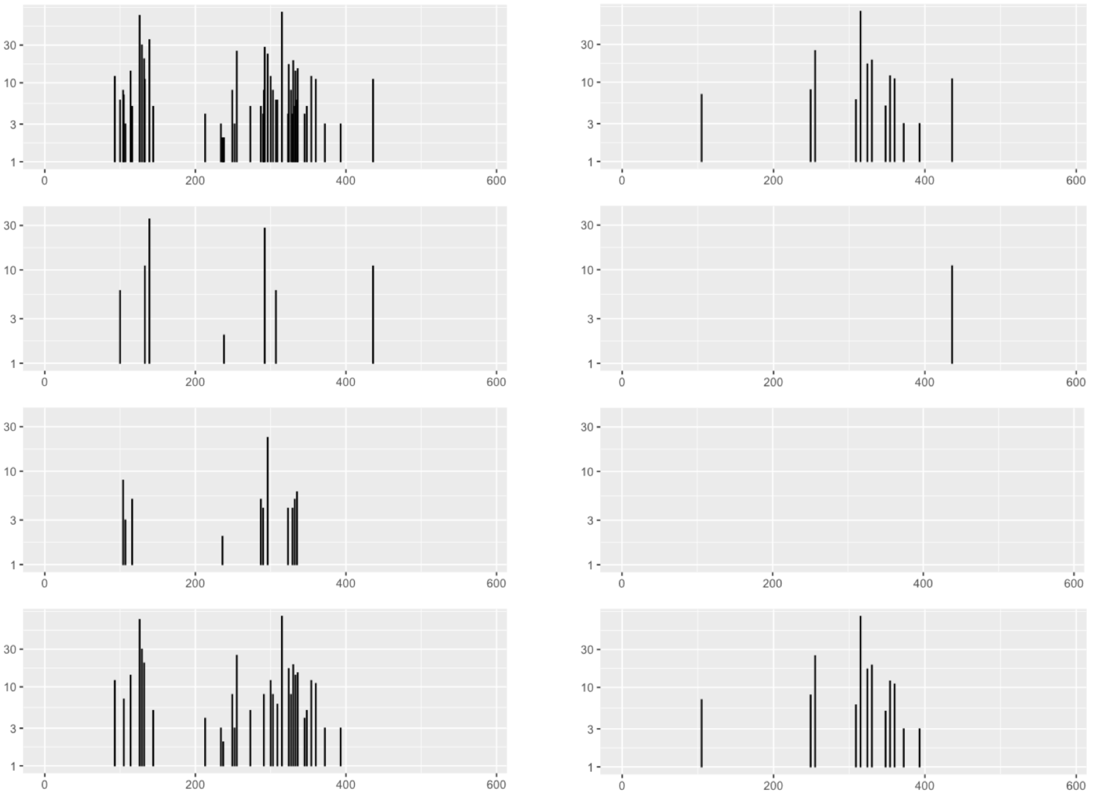

**TCONS_00013674_plus**

24°C

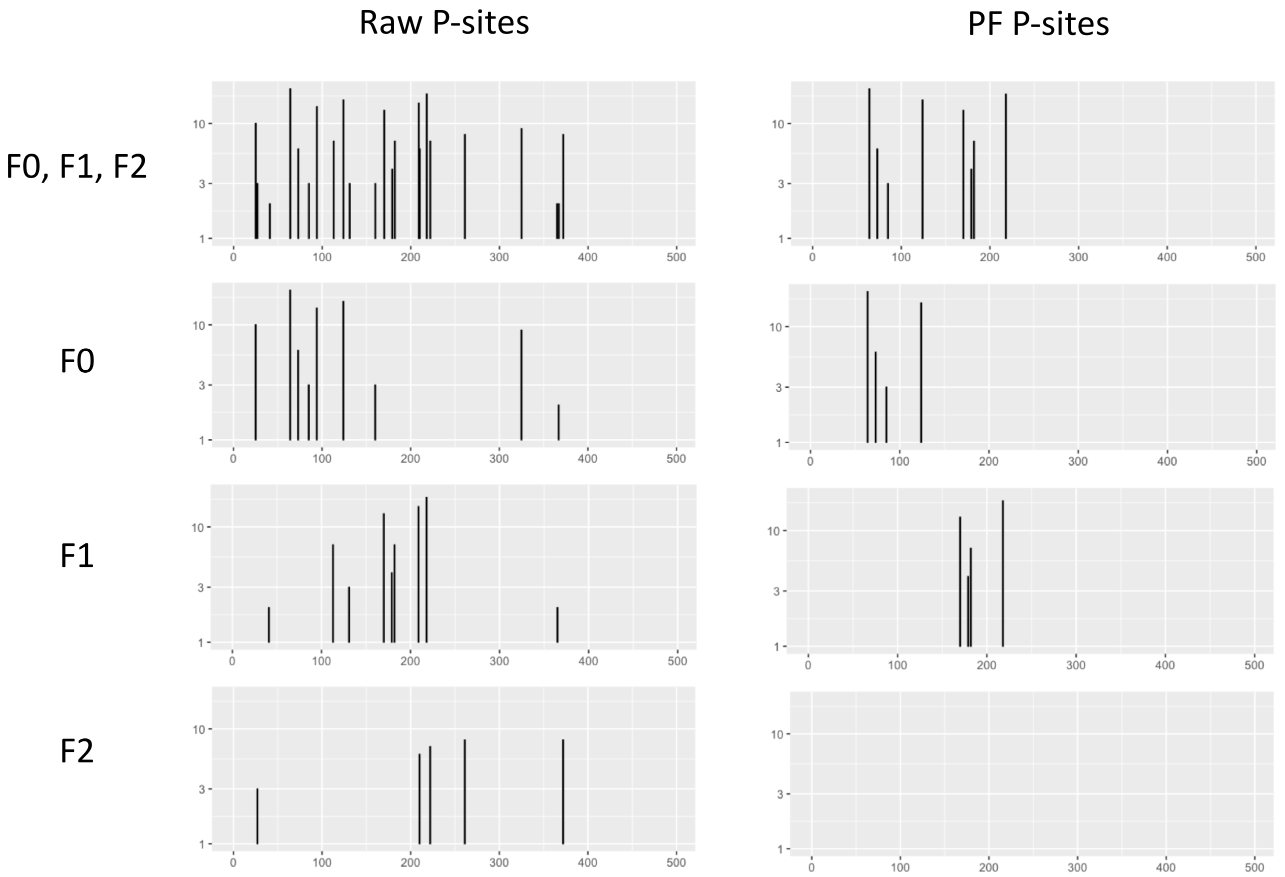

**TCONS_00014294_plus**

35°C

Raw P-sites

PF P-sites

F0, F1, F2

F0

F1

F2

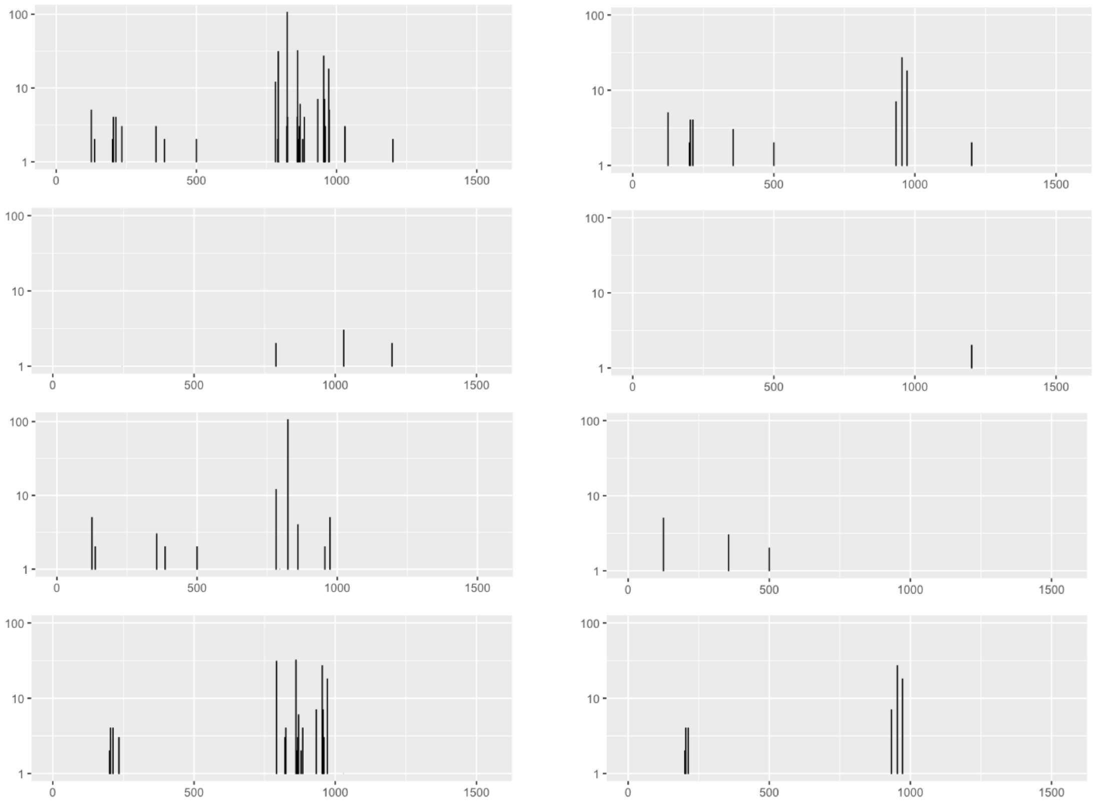

### TCONS_00021461_plus

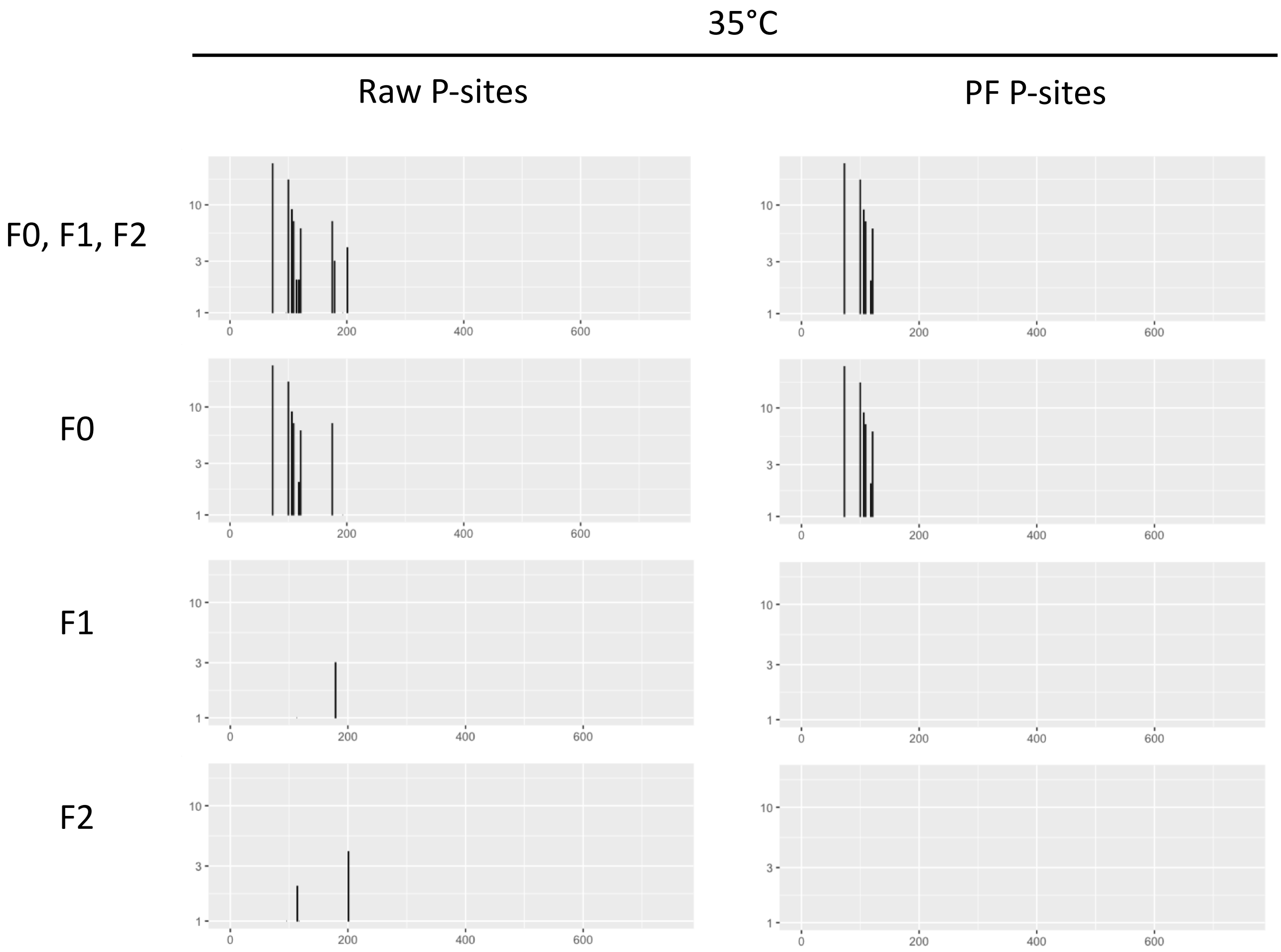

**TCONS_00024073_plus**

35°C

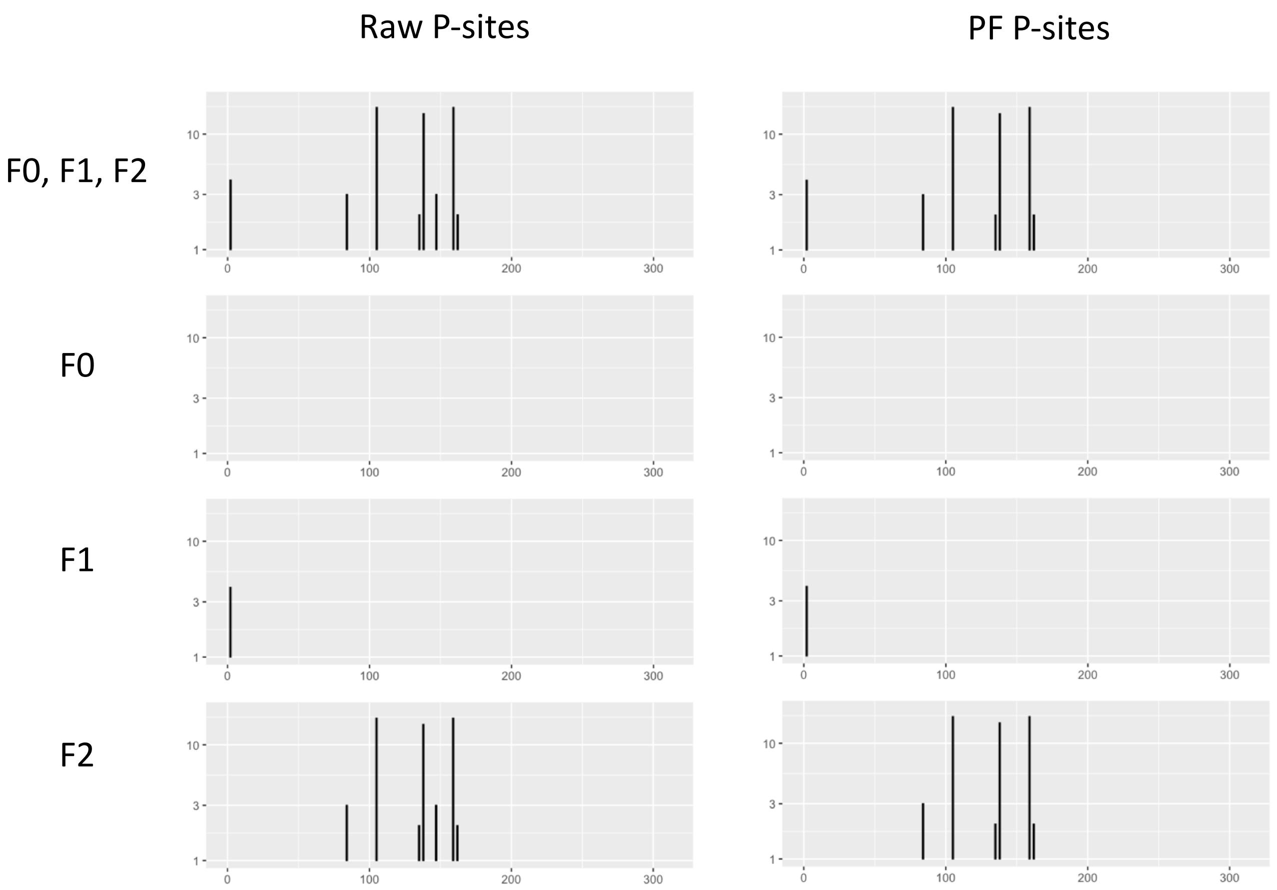

#### TCONS_00026848_plus

24°C

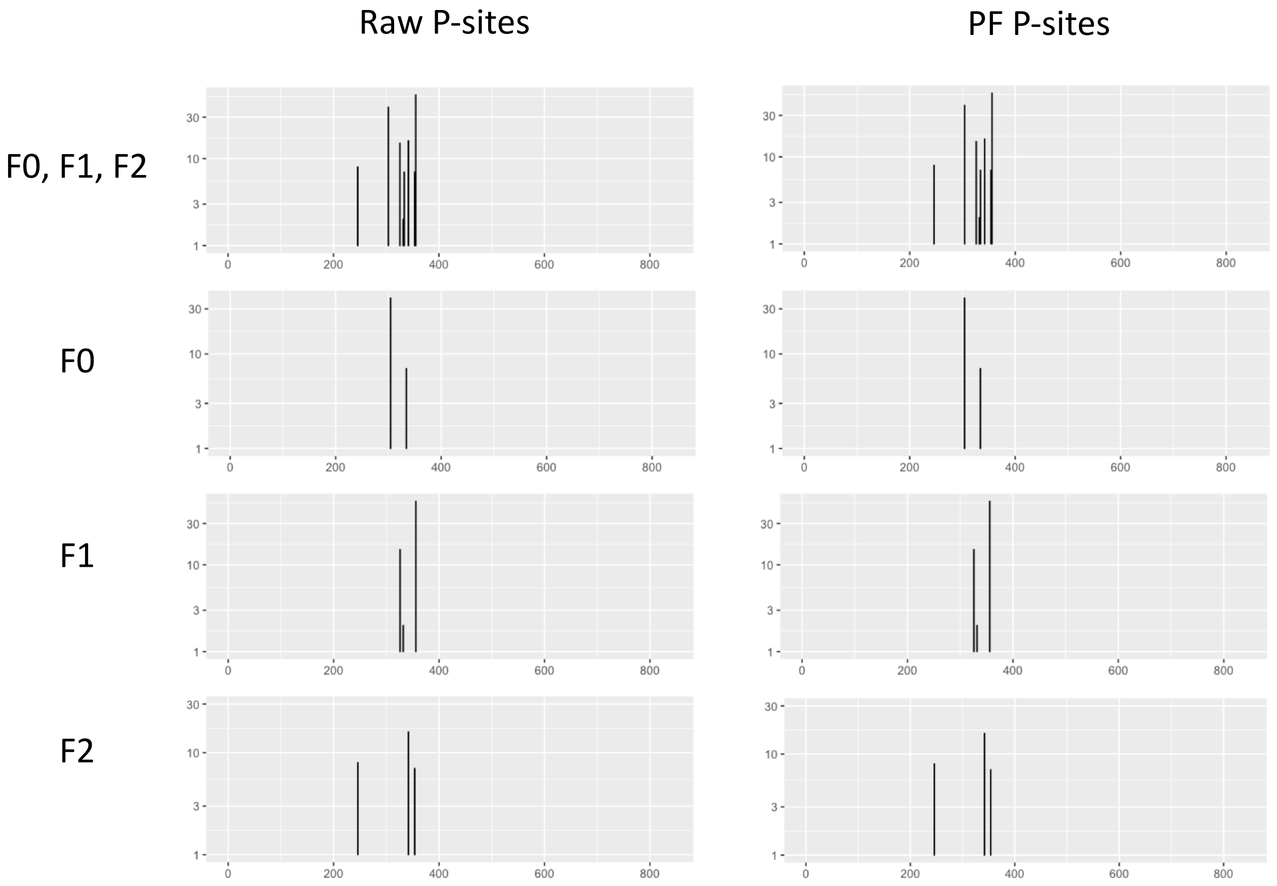

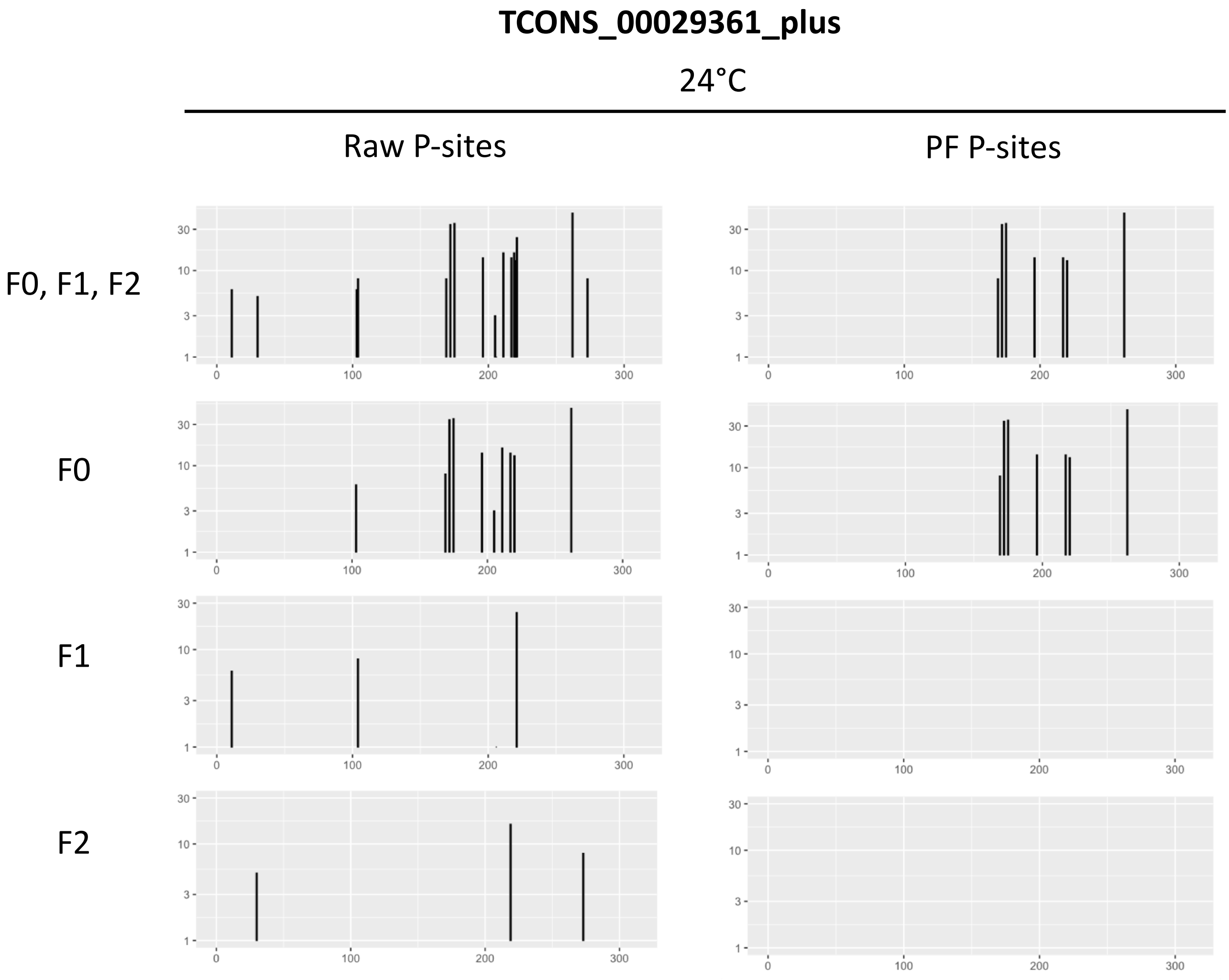
TCONS_00029361_plus
24°C
Raw P-sites
PF P-sites
F0, F1, F2
F0
F1
F2
30
10
3
1
0
100
200
300

**TCONS_00038372_plus**

35°C

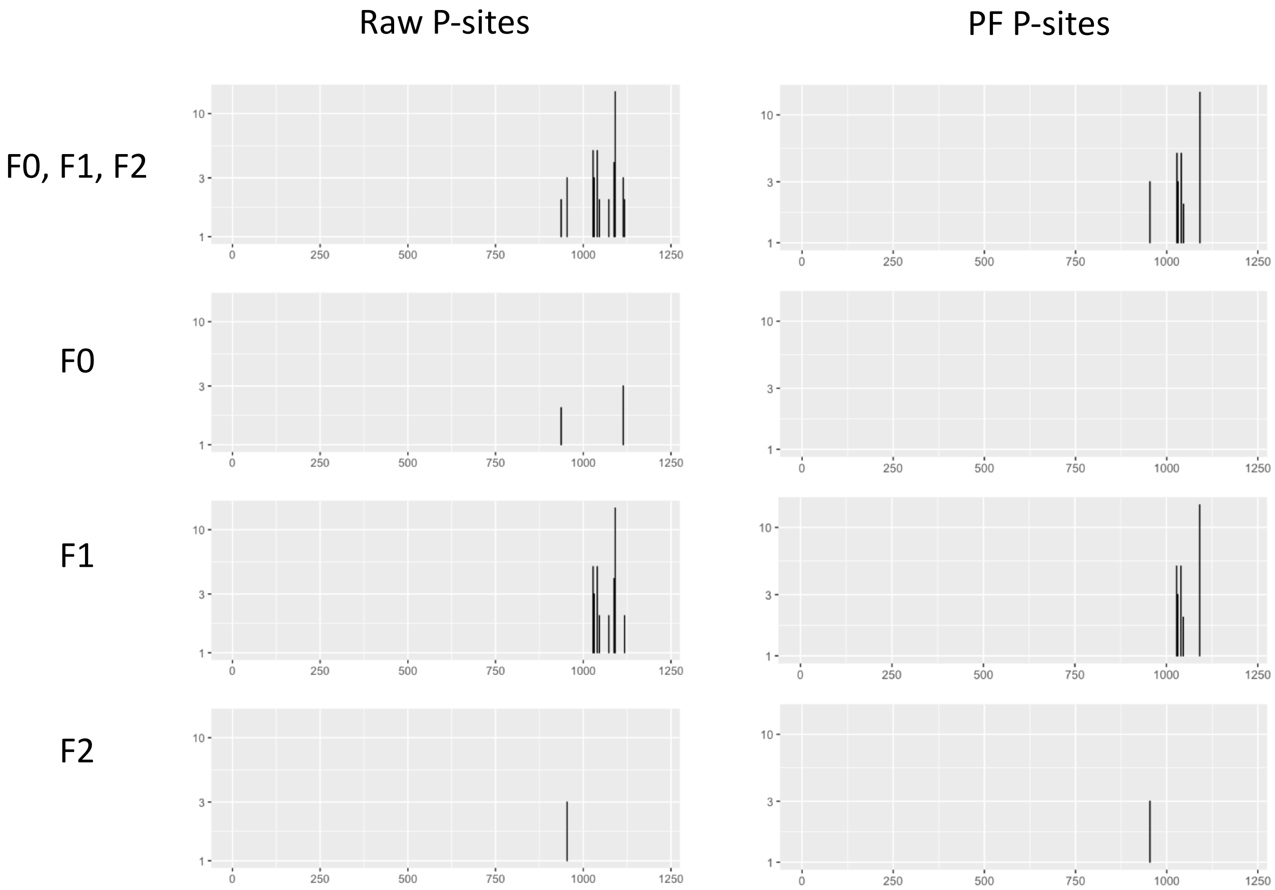

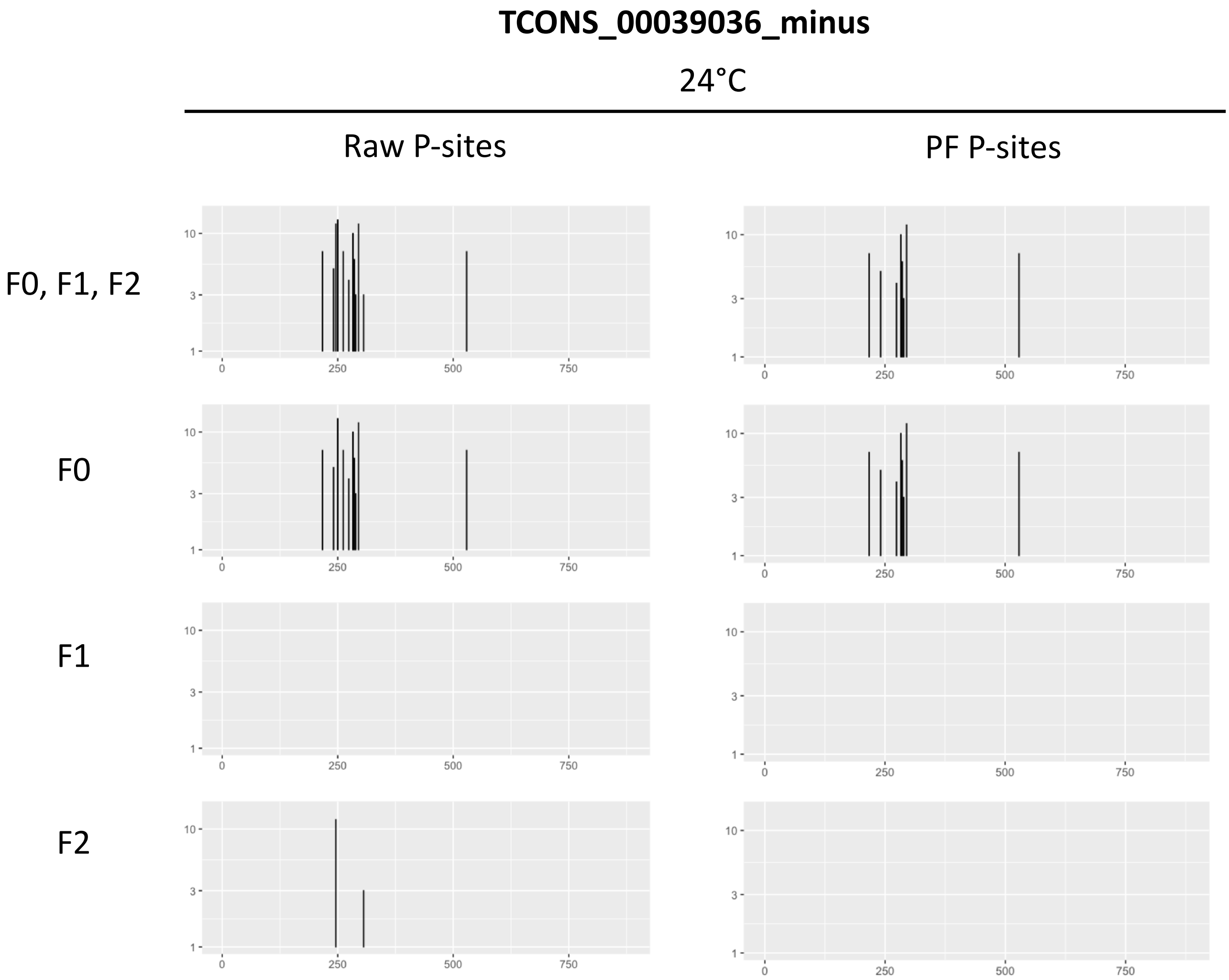
TCONS_00039036_minus
24°C
Raw P-sites
PF P-sites
F0, F1, F2
F0
F1
F2
10
3
1
0
250
500
750

**TCONS_00042679_plus**

24°C

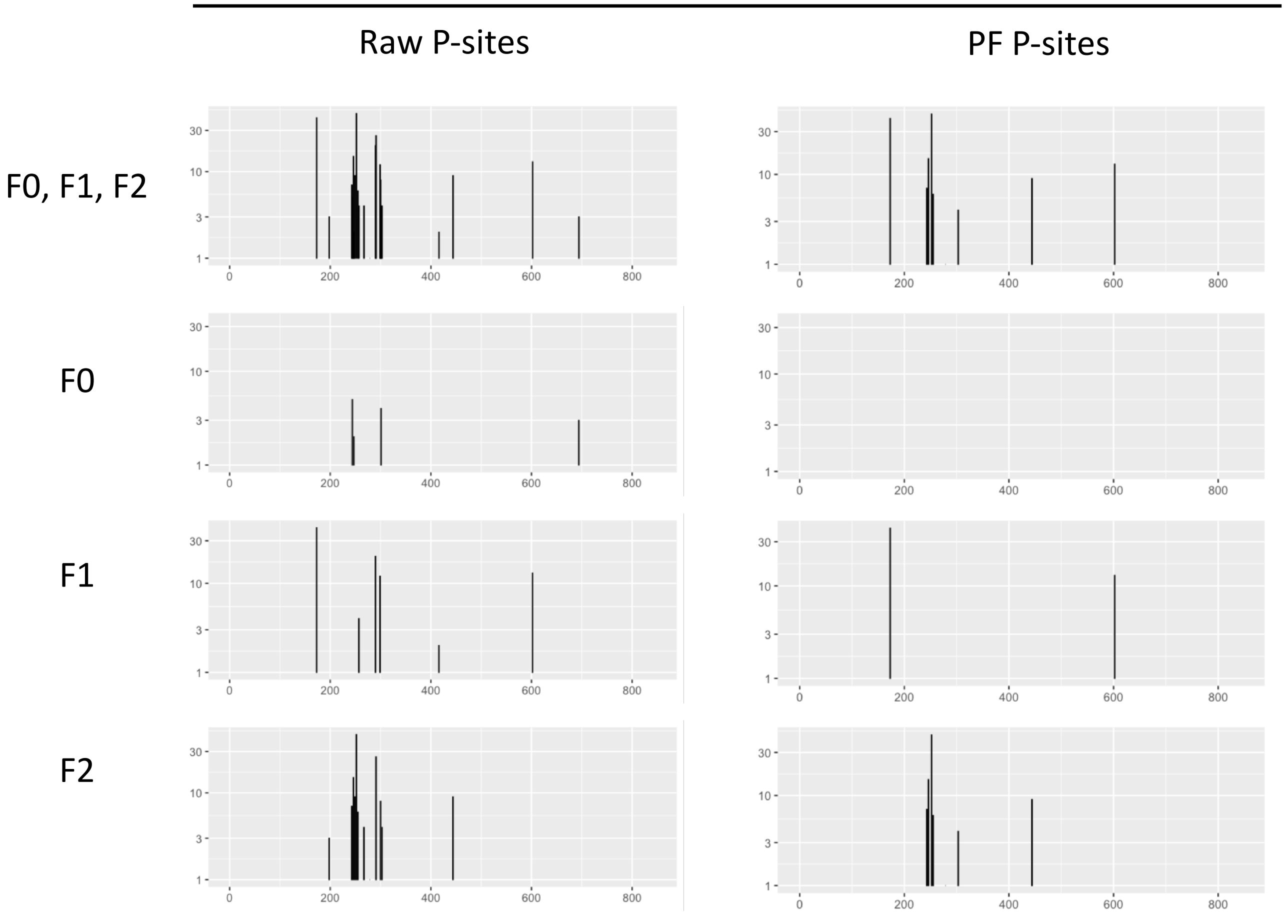

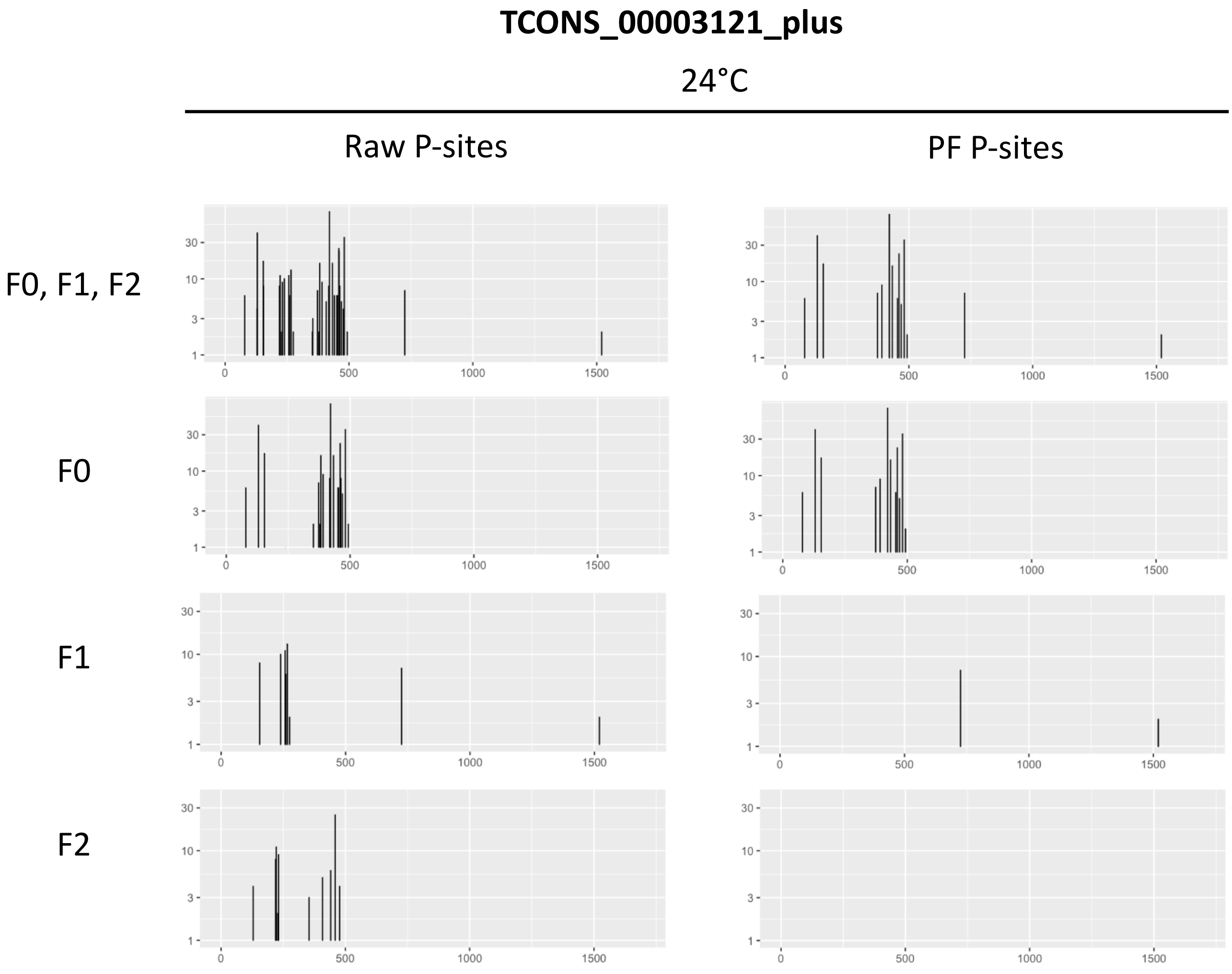
TCONS_00003121_plus
24°C
Raw P-sites
PF P-sites
F0, F1, F2
F0
F1
F2
30
10
3
1
0
500
1000
1500

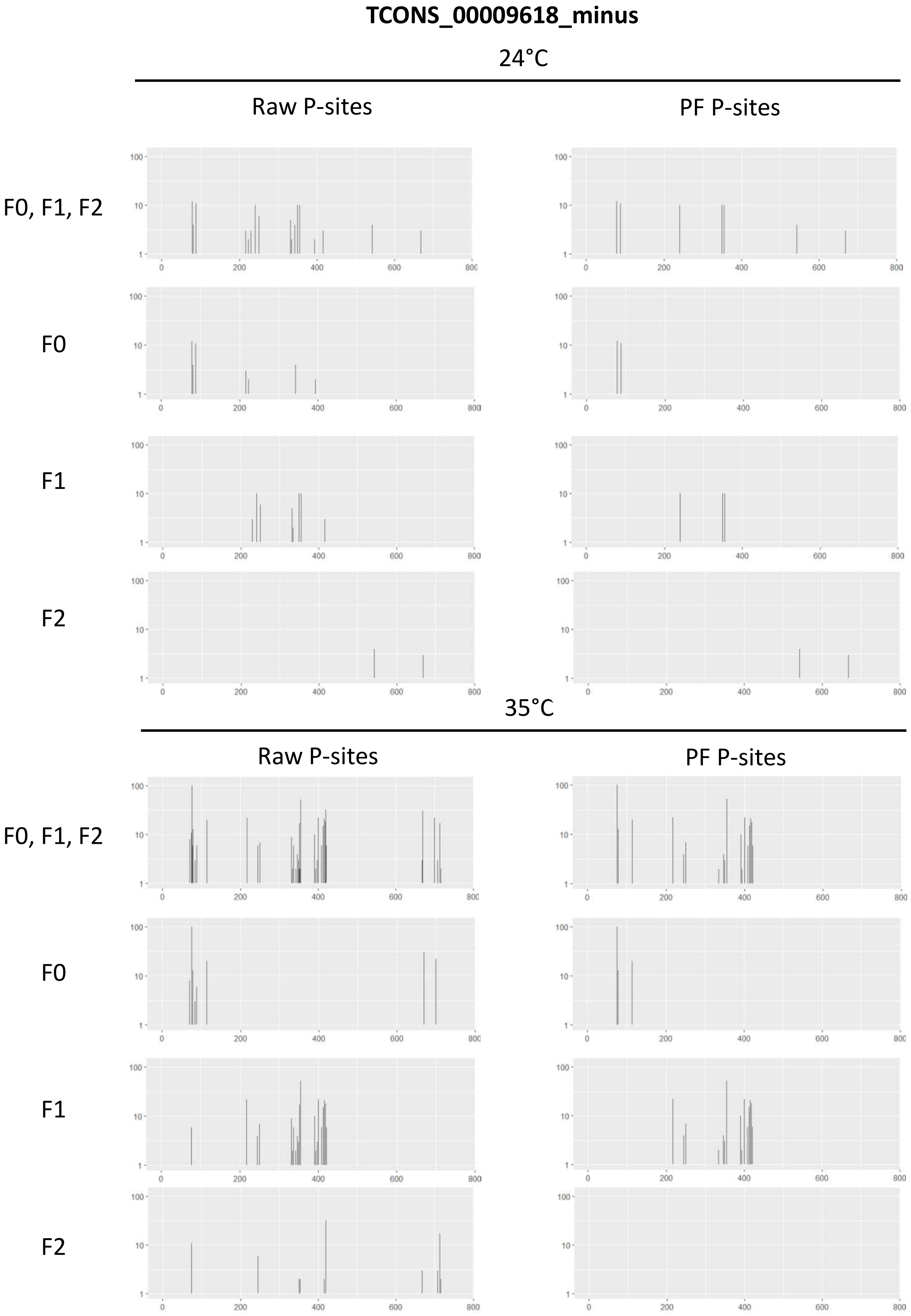
TCONS_00009618_minus
24°C
Raw P-sites
PF P-sites
F0, F1, F2
F0
F1
F2
35°C
Raw P-sites
PF P-sites
F0, F1, F2
F0
F1
F2

#### TCONS_00012928_minus

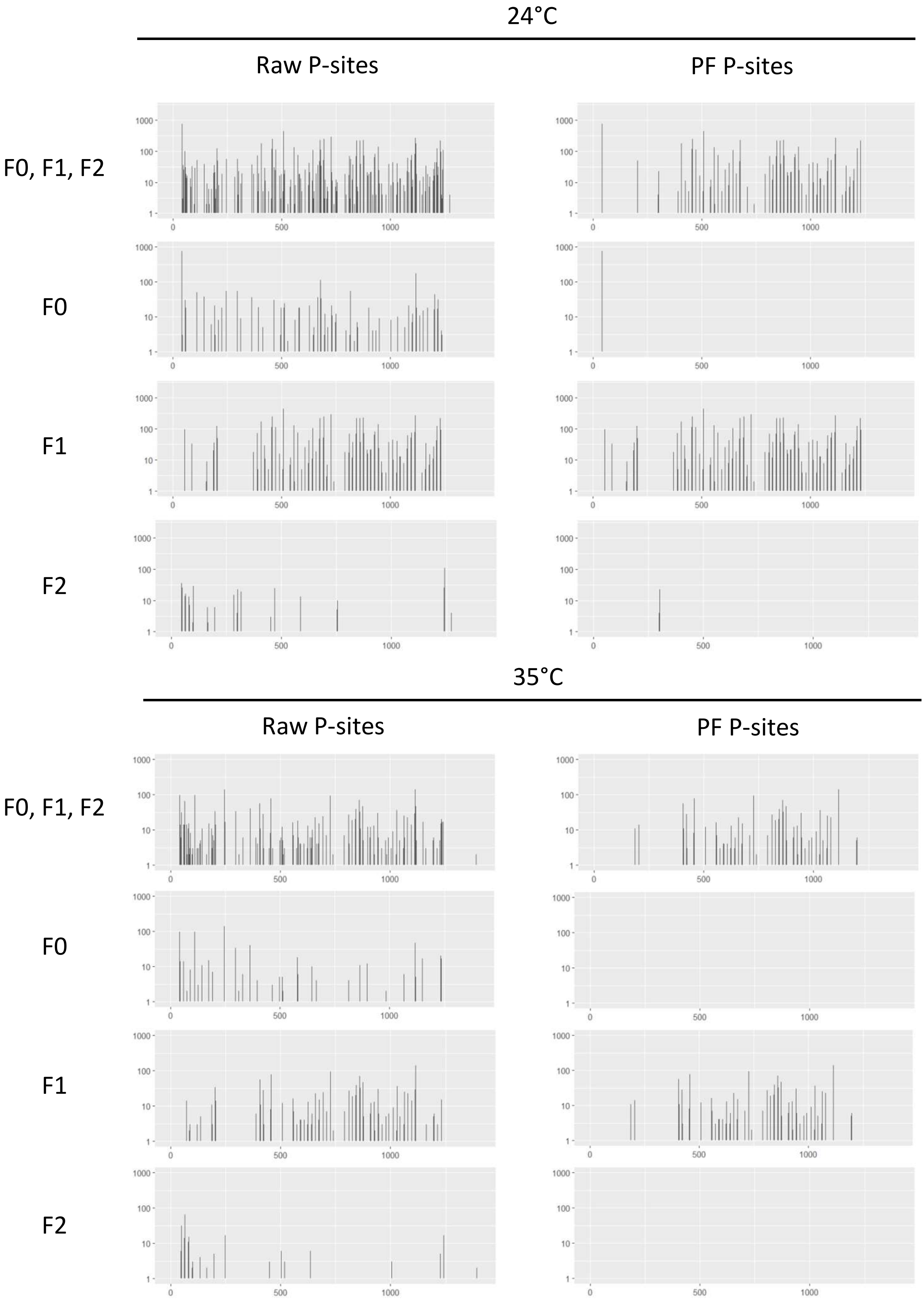

**TCONS_00013972_plus**

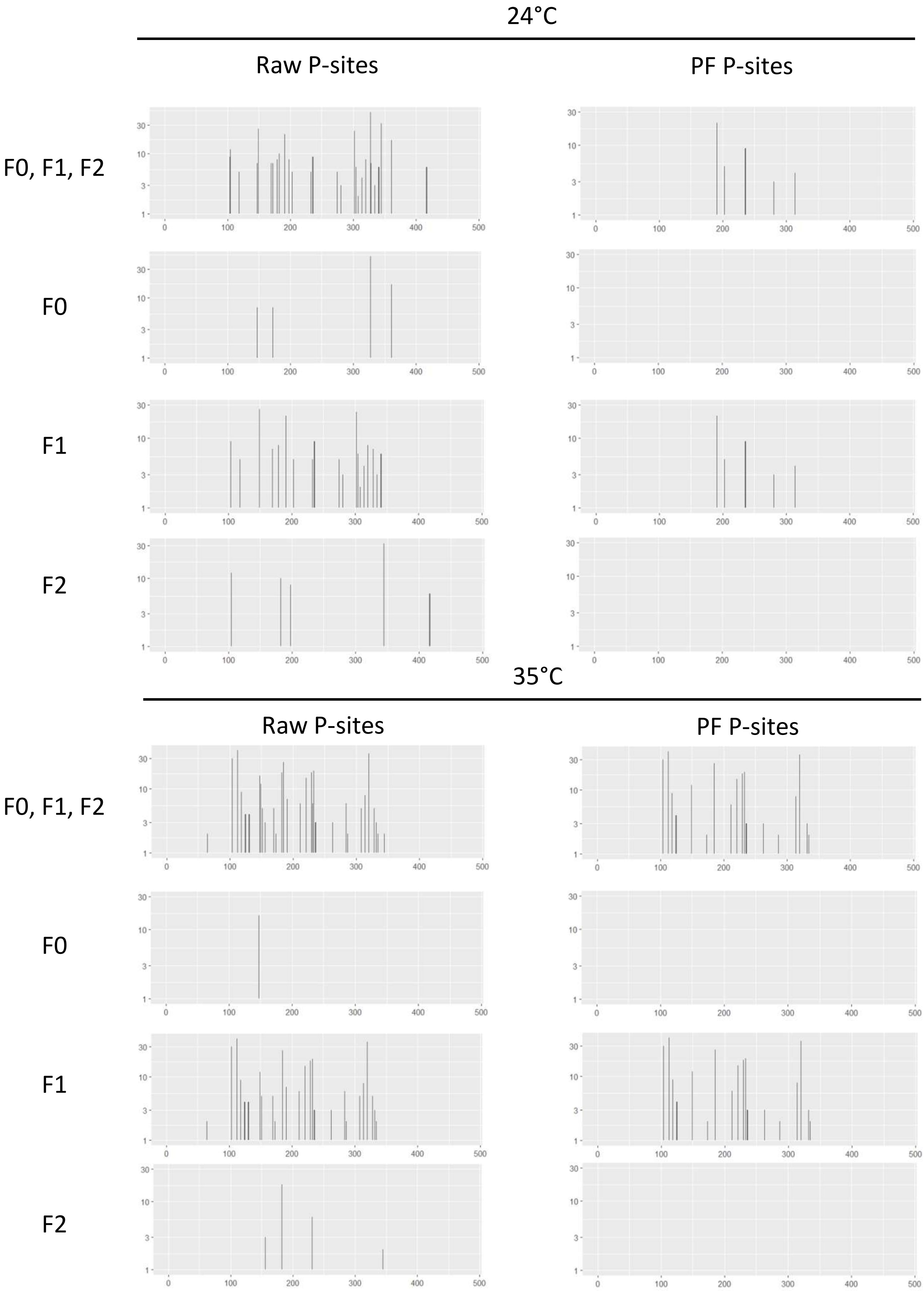

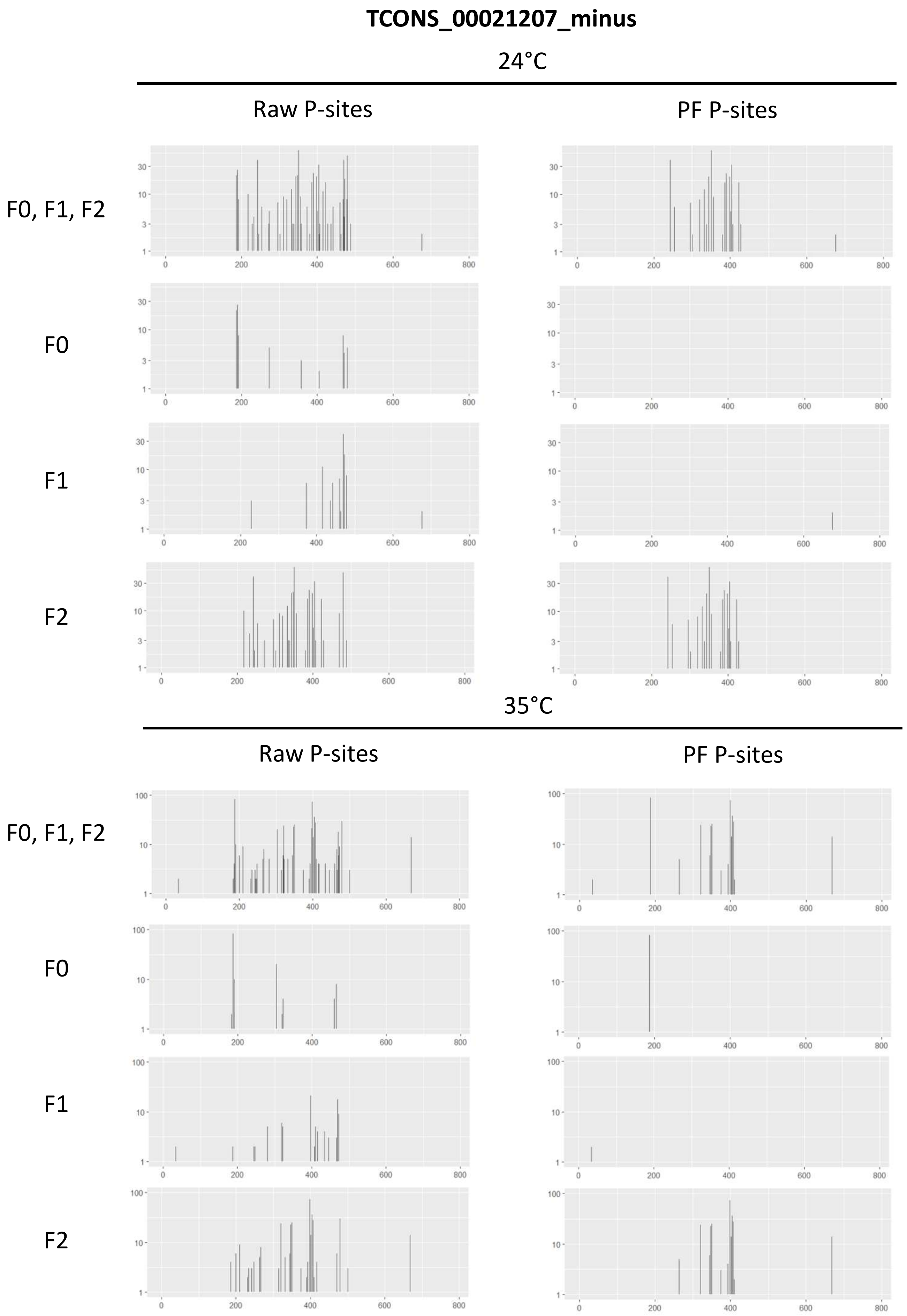
TCONS_00021207_minus
24°C
Raw P-sites
PF P-sites
F0, F1, F2
F0
F1
F2
35°C
Raw P-sites
PF P-sites
F0, F1, F2
F0
F1
F2

#### TCONS_00026364_minus

24°C

| Raw P-sites | PF P-sites |
| --- | --- |

F0, F1, F2

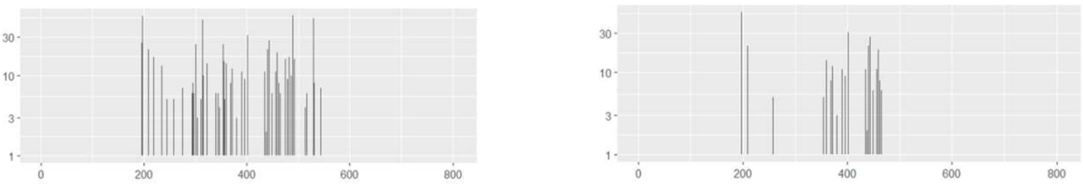


F0

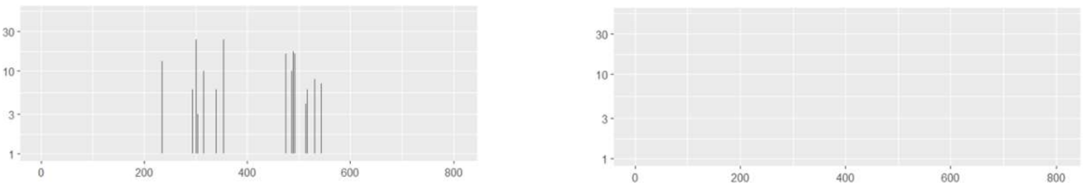

F1

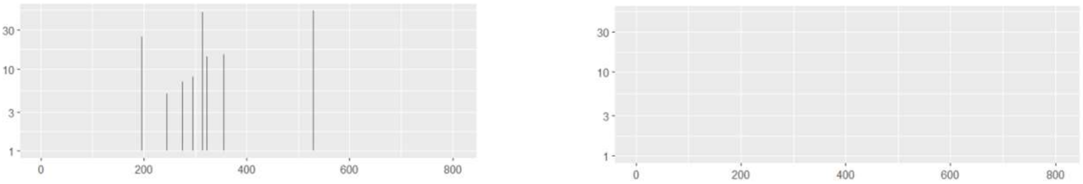

F2

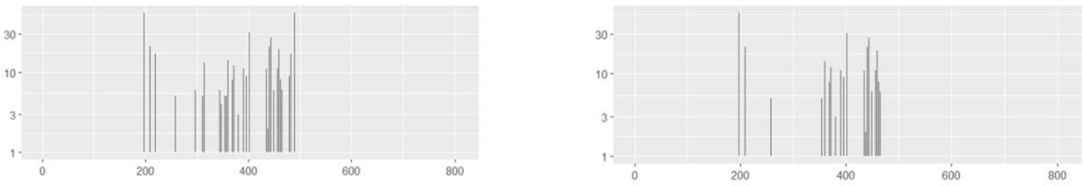

35°C

| Raw P-sites | PF P-sites |
| --- | --- |

F0, F1, F2

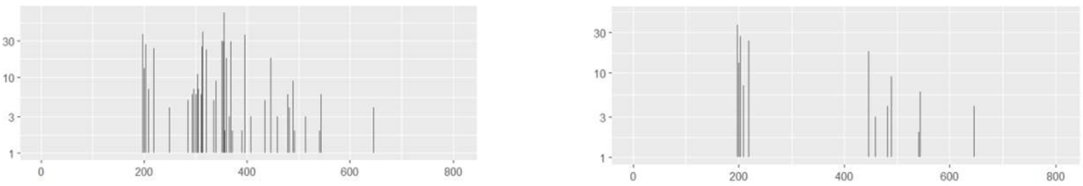

F0

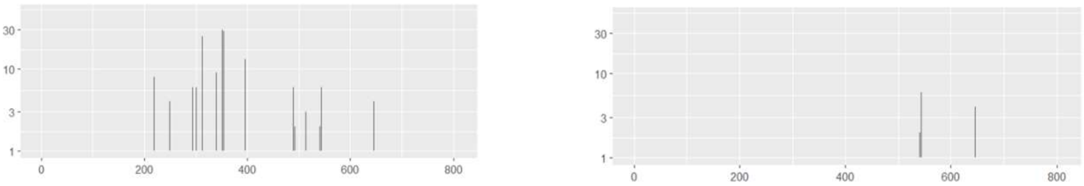

F1

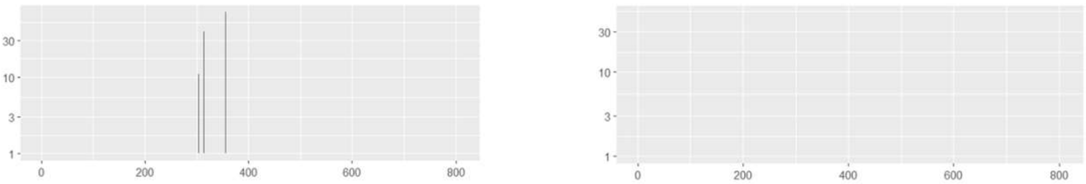

F2

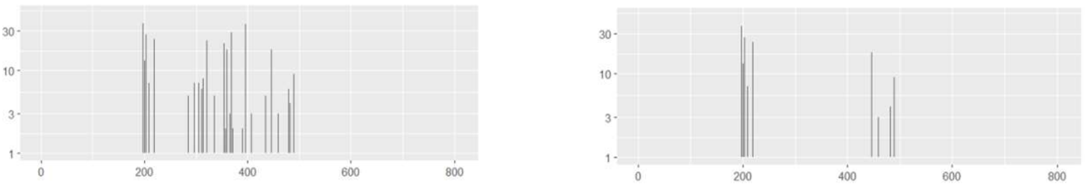

**TCONS_00031274_plus**

24°C

Raw P-sites | PF P-sites

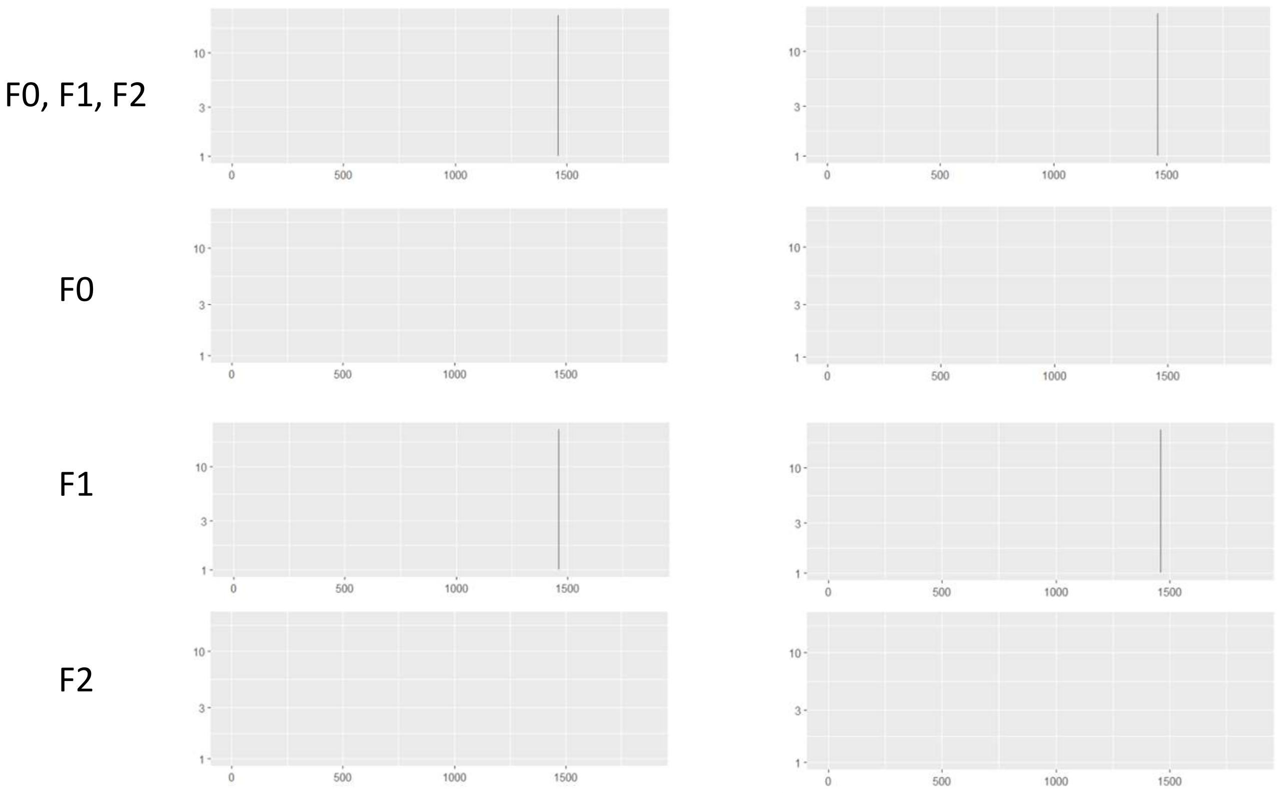


35°C

Raw P-sites | PF P-sites

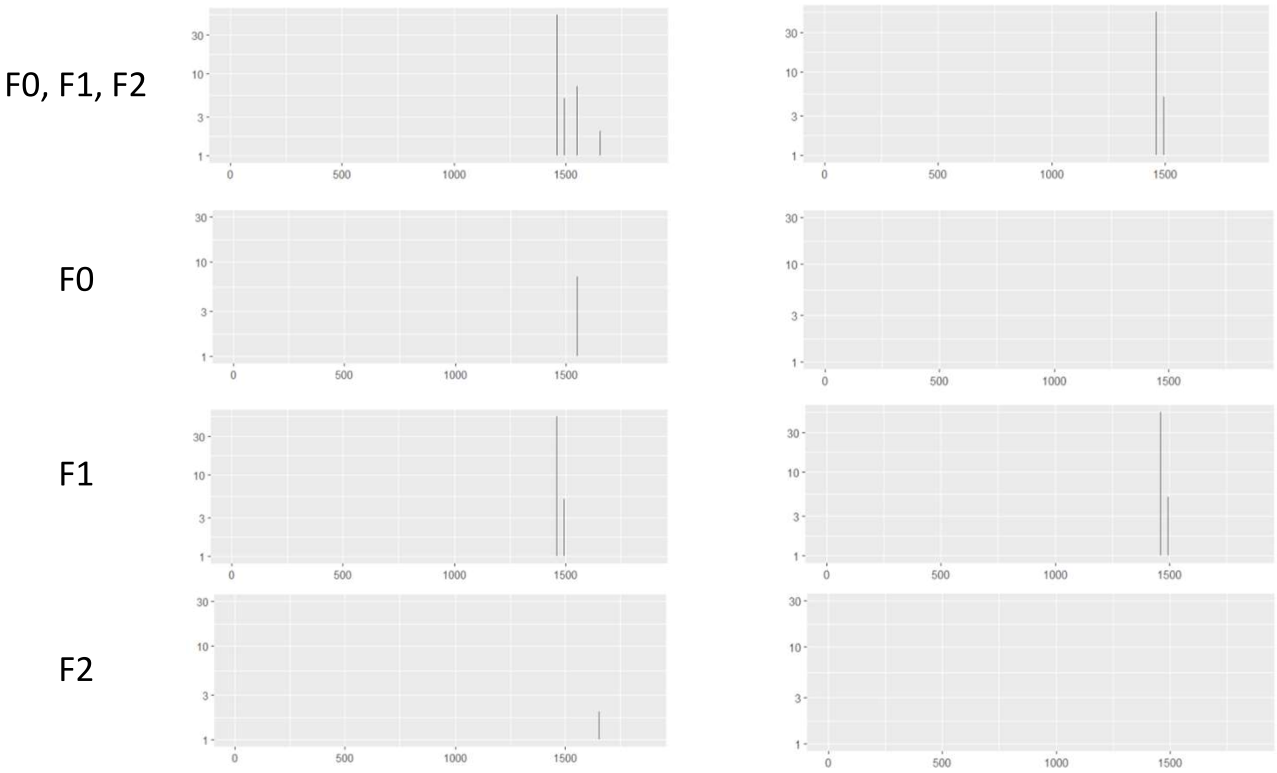

#### TCONS_00032246_minus

24°C

Raw P-sites | PF P-sites

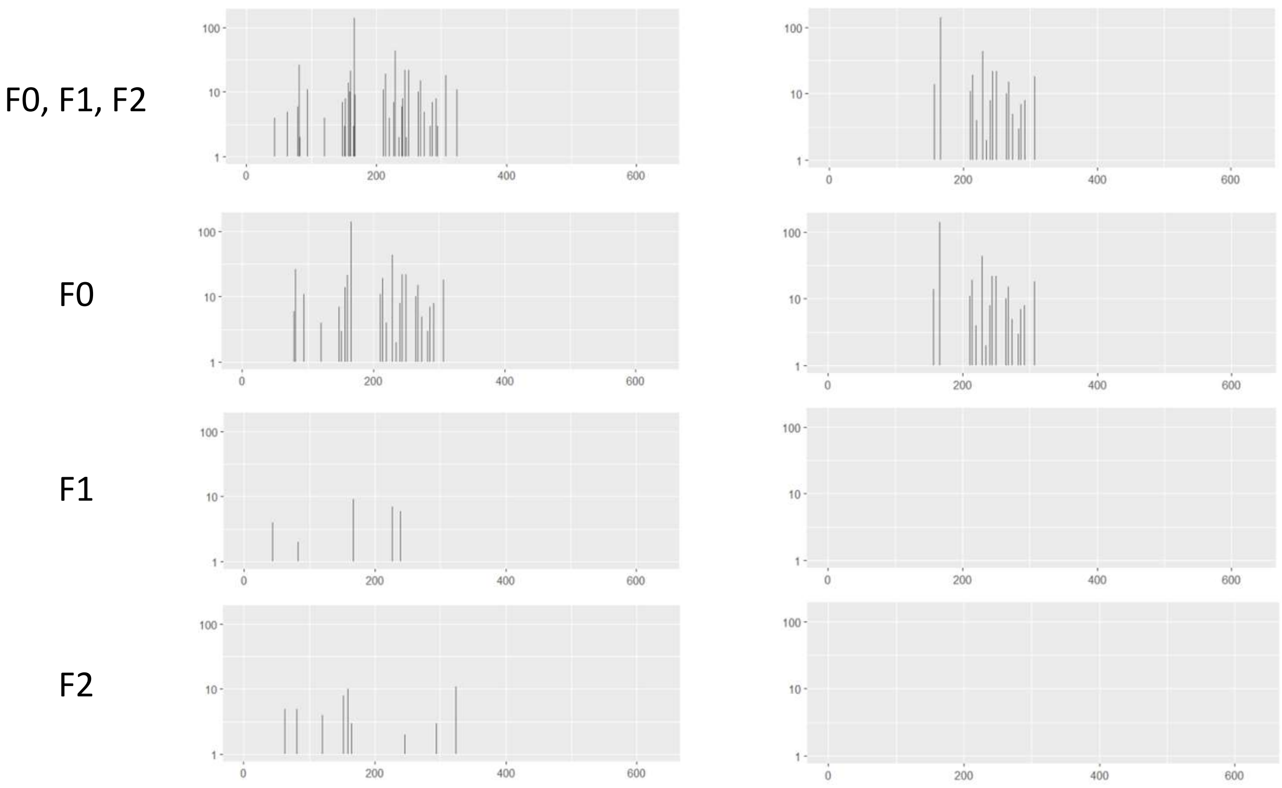


35°C

Raw P-sites | PF P-sites

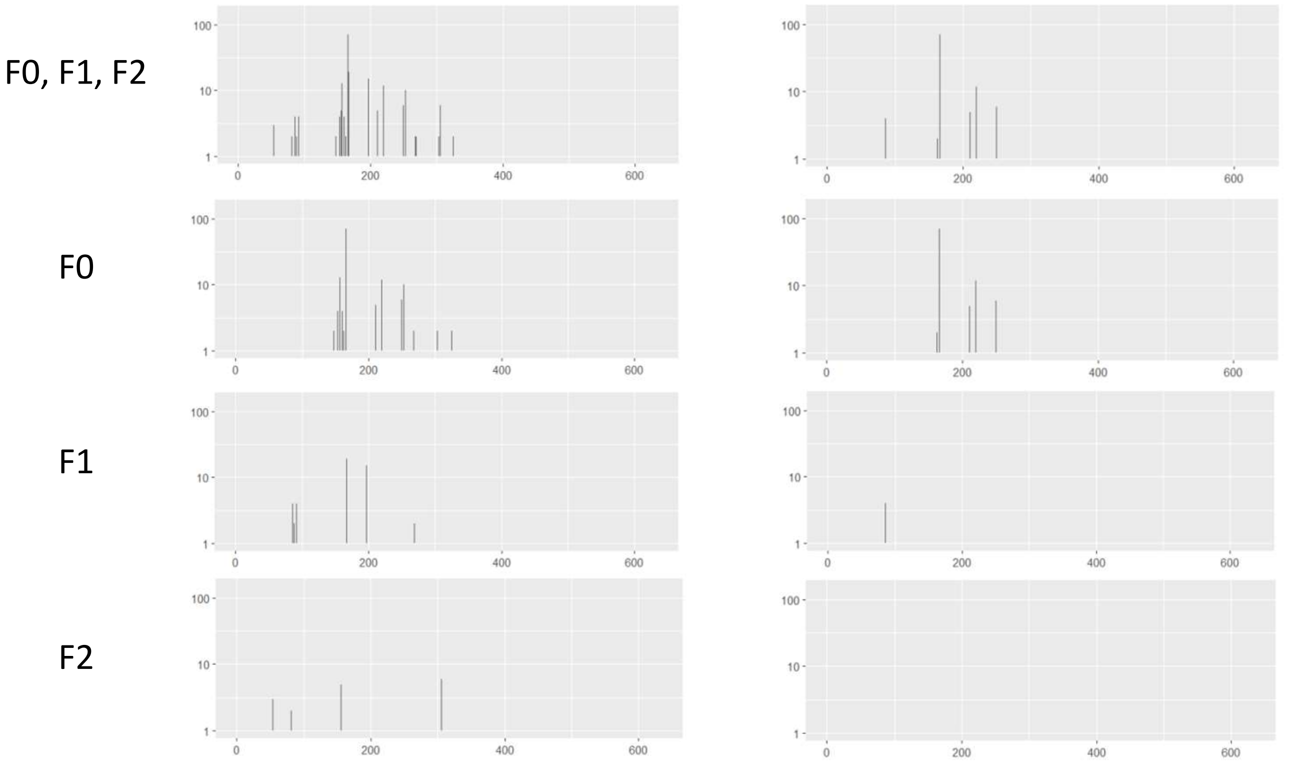

**TCONS_00035135_plus**

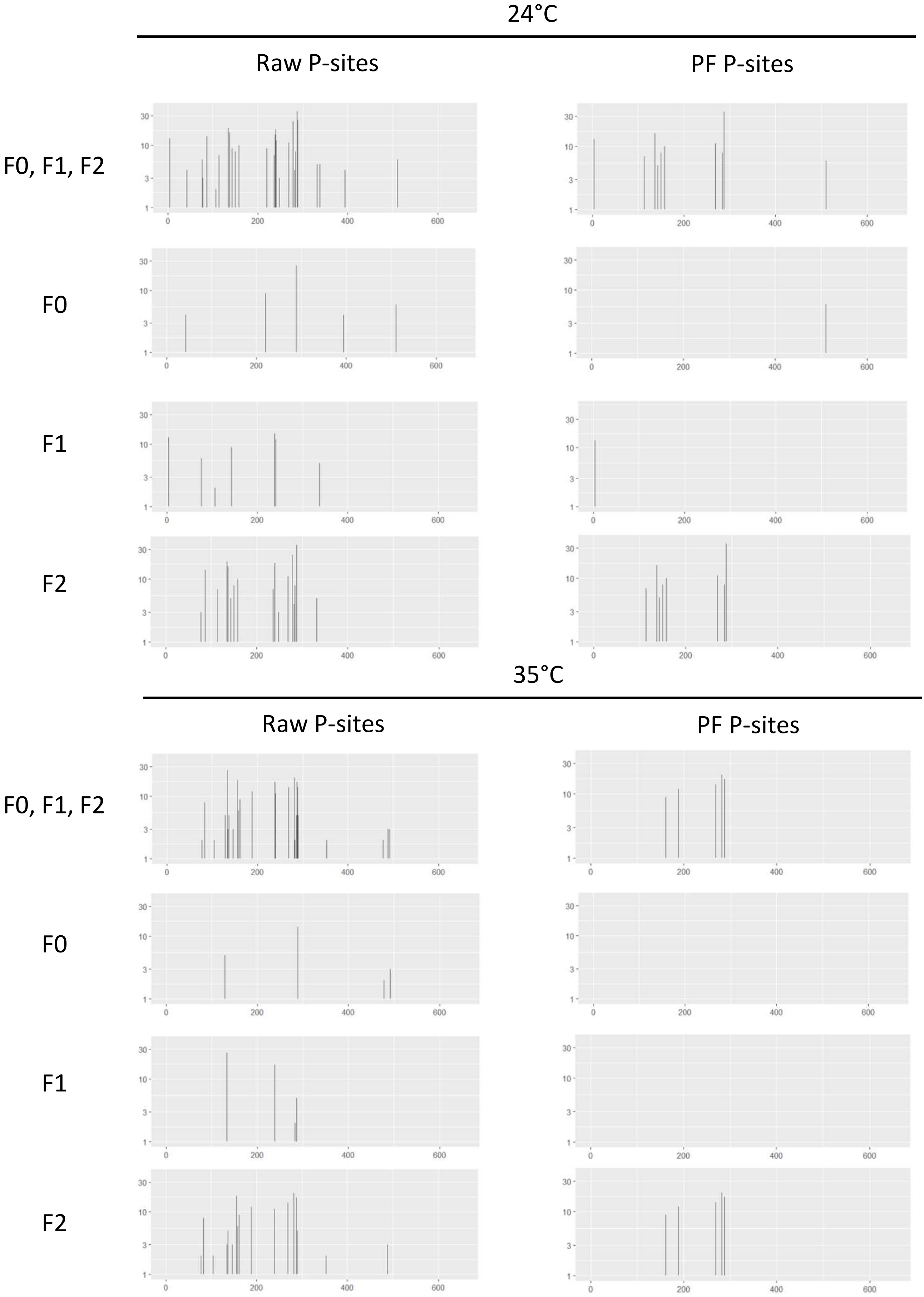

**TCONS_00047195_minus**

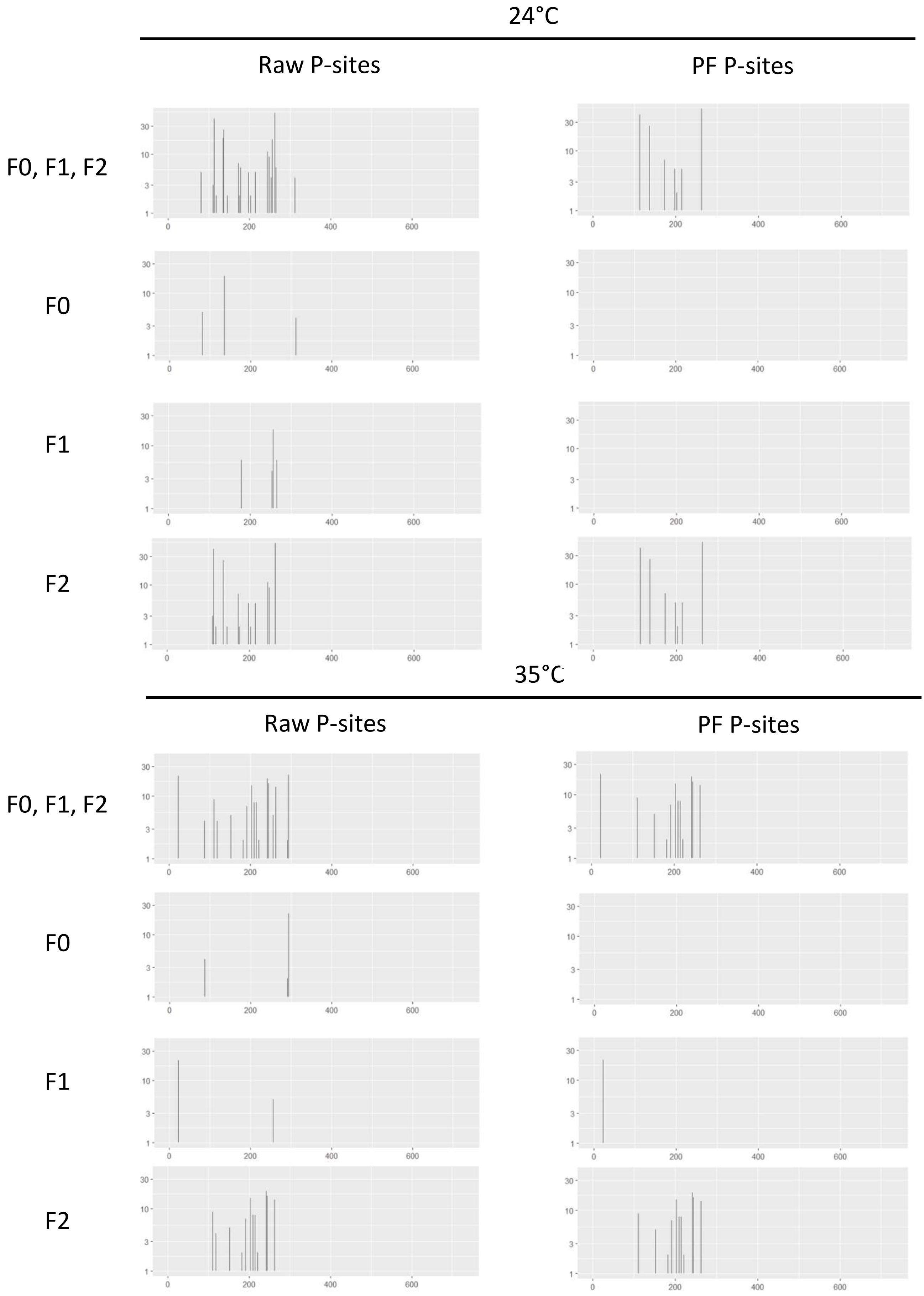

**TCONS_00047695_plus**

24°C

Raw P-sites | PF P-sites

F0, F1, F2

F0

F1

F2

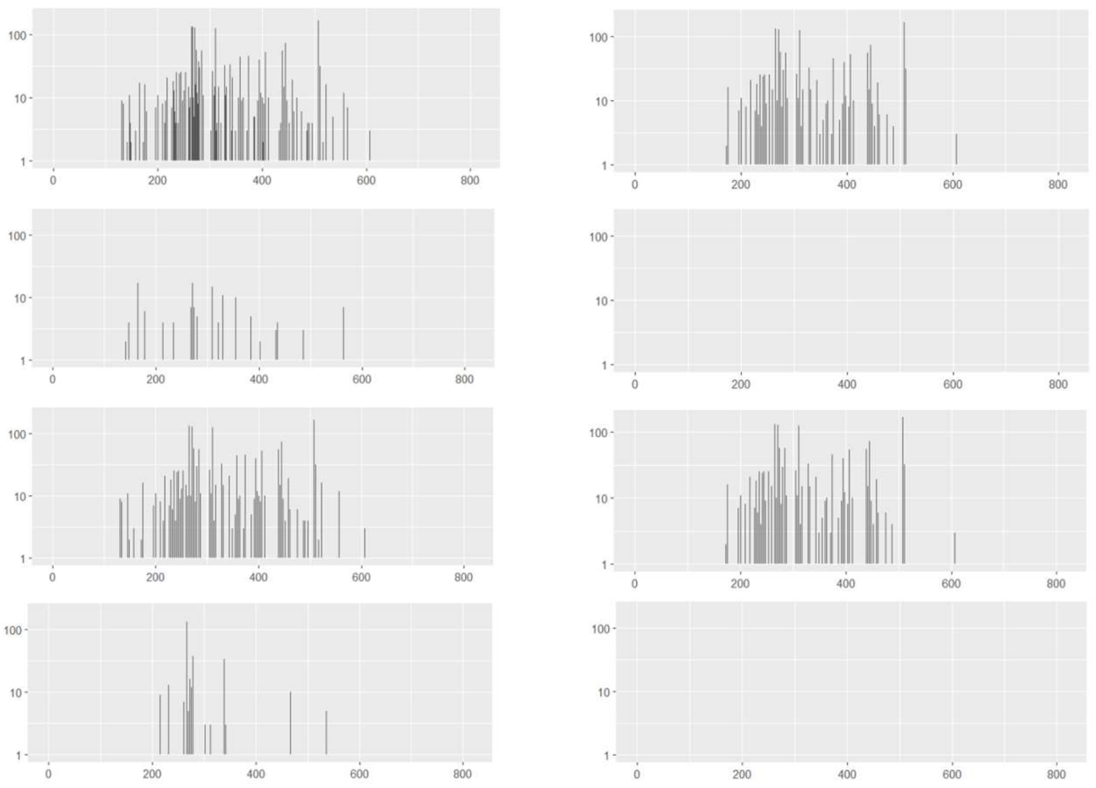


35°C

Raw P-sites | PF P-sites

F0, F1, F2

F0

F1

F2

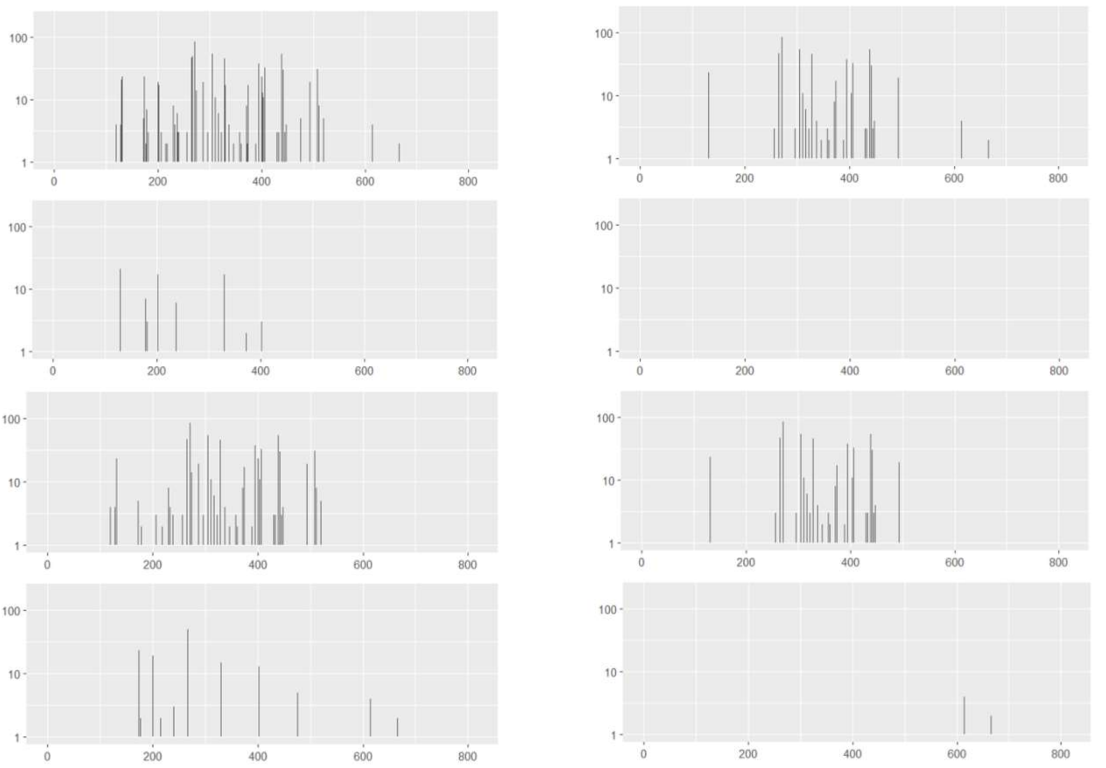

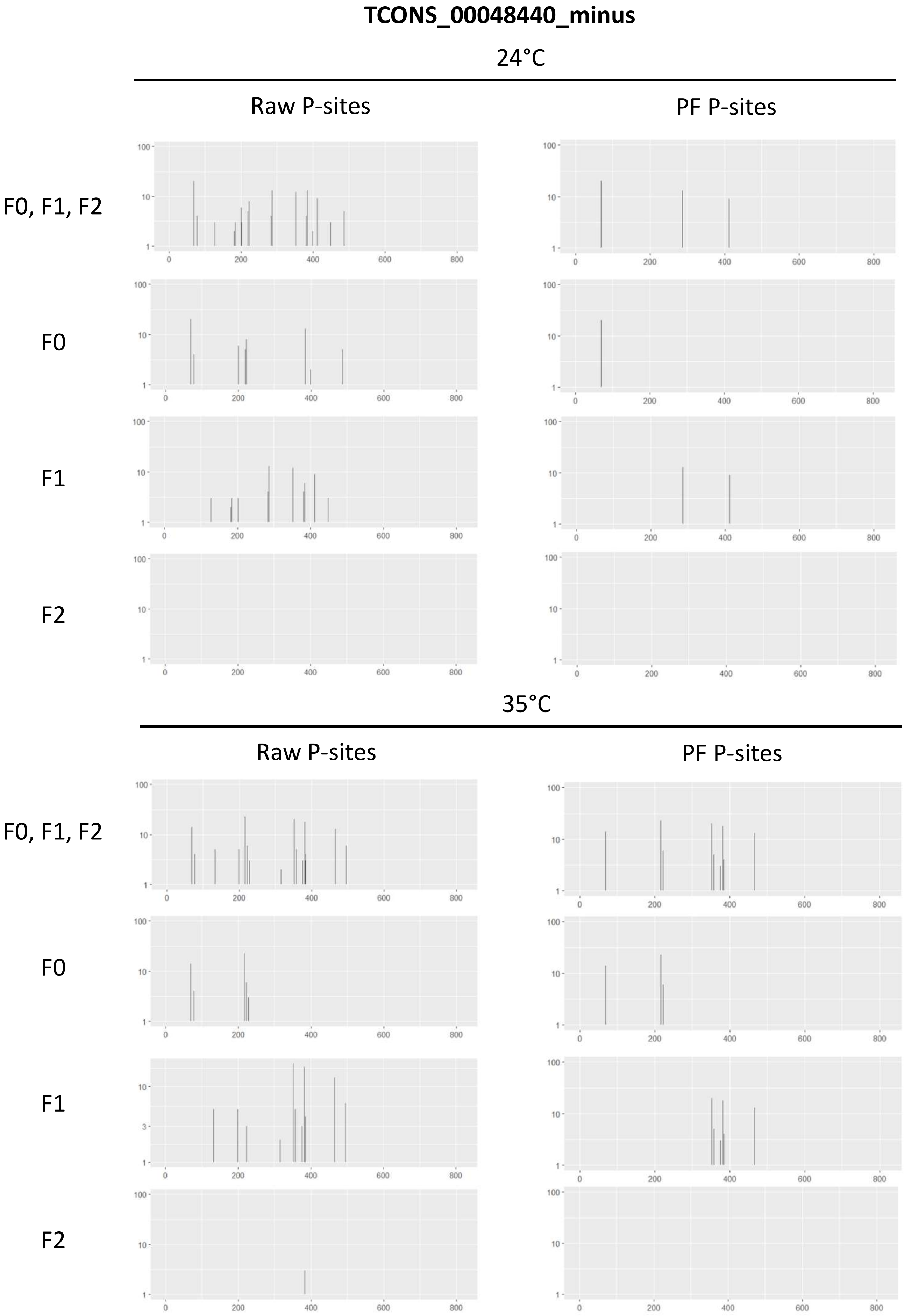

TCONS_00048440_minus
24°C
Raw P-sites
PF P-sites
F0, F1, F2
F0
F1
F2
35°C
Raw P-sites
PF P-sites
F0, F1, F2
F0
F1
F2

**TCONS_00050361_plus**

24°C

Raw P-sites | PF P-sites

F0, F1, F2

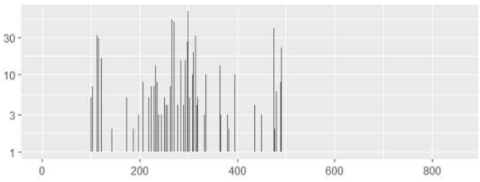


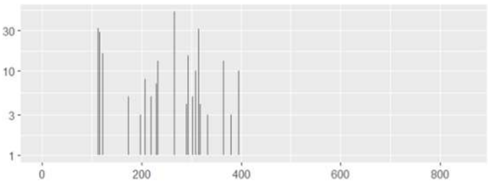


F0

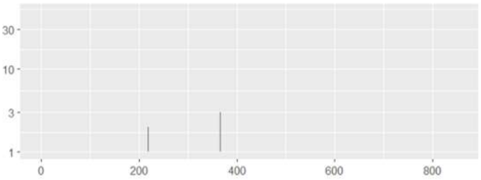


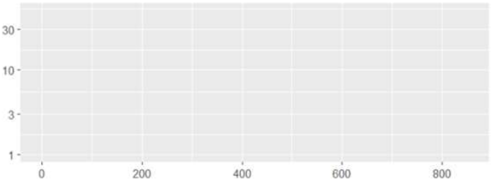


F1

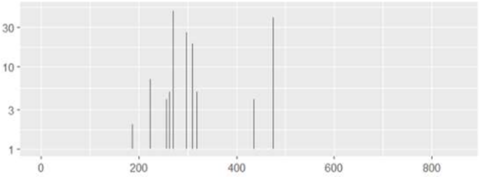


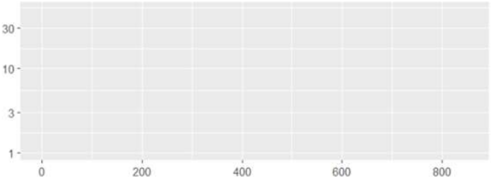


F2

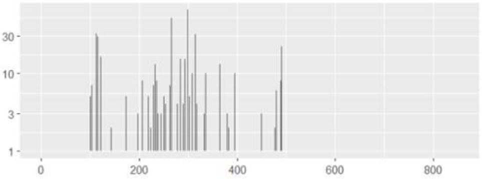


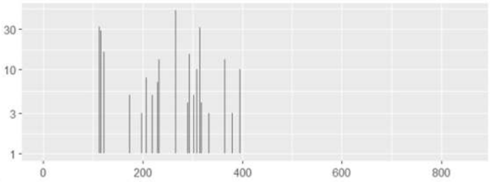


35°C

Raw P-sites | PF P-sites

F0, F1, F2

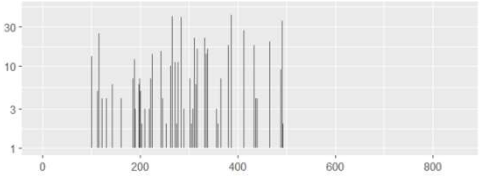


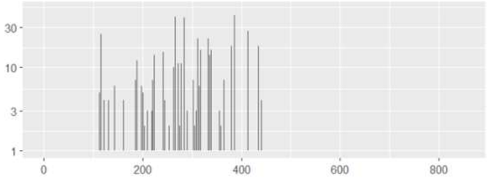


F0

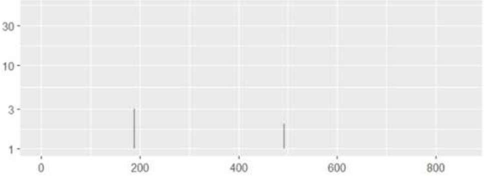


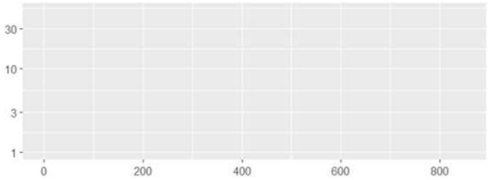


F1

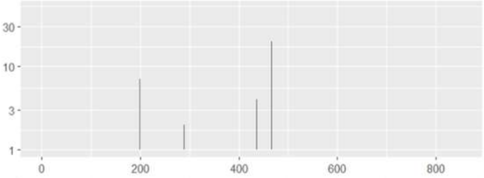


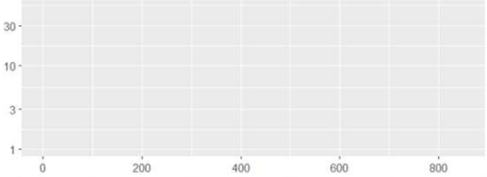


F2

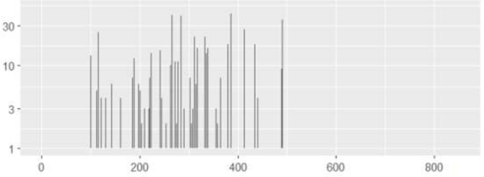


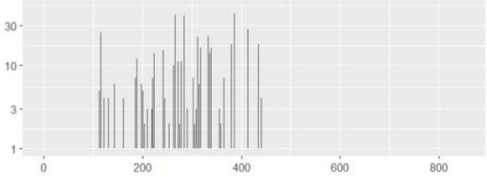

#### TCONS_00050361_plus

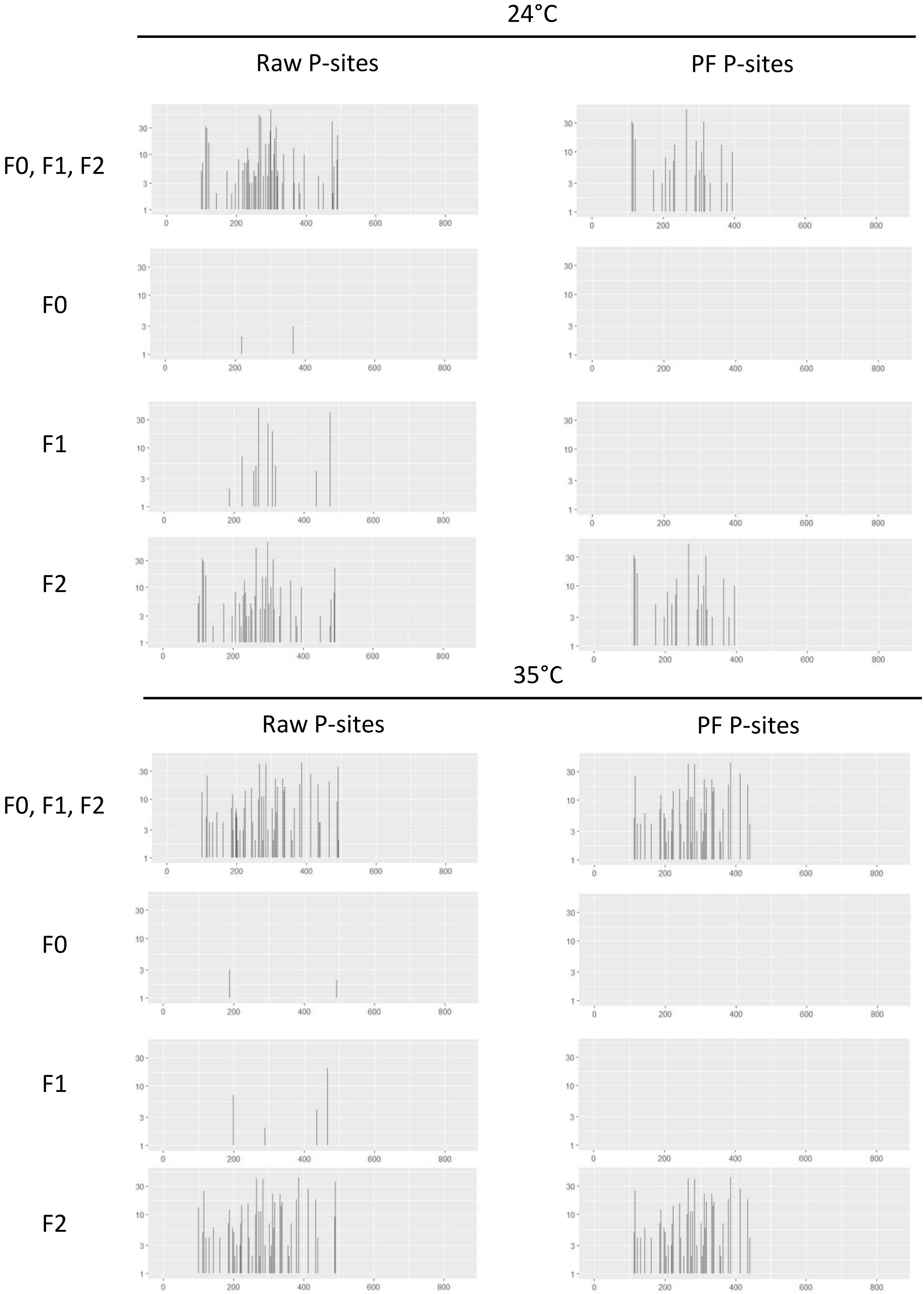
