## Supplemental Doc 4 for "Characterization of novel pollen-expressed transcripts reveals their potential roles in pollen heat stress response in *Arabidopsis thaliana*" (from another version of this preprint: version 2, 2020-08-26)

XLOC_006026 genomic fragment

```
Ch1: 13222206 -
AACGTCTAAATGATCCACTATAAATAACCATCTCATATCGAACAATTACCACAACCACAATTTGCCTCAGTCTAAATGATCTCTCAGAATTTTTATATAGCAAGGTATTTCTTATGTATAGCTCTAGAGATGTTTTGCTATACAGAATATTTATCAGTGCTTGTGTGACGAATTTAATAGATTgagtttgtgattttgttgtttgtagAGCTTTATGAGCAAGCTATTGAGTATGAGAAAGGTTCGTTAATCACTTCTAATGGAGCTTTGACGATGCTTTCTGGTGCTAAGACTGATCGAGAGATAAGCGTGTTGTTAGAGATGCGATTACTGAGGATGAGCTTTGGCGATTACAGAATTTTTATCctctattttggtttattgatttttgtttttgattttgattttgattttaaggGTTTGTCTAATATTGAAATCGATGAACATACTTTCATGGTGGTGAACAGAGAAAGAGCTGTTGATTACTTGAATTCCTTGGAAAAGGTTTACCTTATGCCTAAAAGTTAGATTCTTTCCCATCATTCTGGATGCAATATGGGAAAAGATGGAGATGTTGCCCTCTTCTTTGAACTTTTAGGTATTGCAACTTTTTAAGCCGACGTGAACgctgaaaaaaaagagaaacaccGACGTGAACGCAGAGAGAAGATTACTTGCGCAACCATCGAcggaaagaaagagaaacaaaacctttcttcttcttttttttttctatatttgtctagaaaaaaaaaattaacgttTCTTTTGGATGTAAAGAATCTTTTCTATTTAGATGTTTTGGCacaaaaaacgaagaaaaaaactataattttgtaatttattttattttagtttatgatTAAg     - 13221341
```

In black – exons, Red – introns. Orange – part of the coding sequence of the Q9LQH8 protein that is included in the extended first intron (Orange and red sequence) spliced to give the sequence of TCONS_00009618.

**GT** and **ag** – the alternative donor and acceptor splice sites of the extended intron removed in TCONS_00009618.

**GT** and **AG** – indicate the correct donor and acceptor sites of intron 1 and 2.

TCONS_00009618 (XLOC_006026)

AACGTCTAAATGATCCACTATAAATAACCATCTCATATCGAACAATTACCACAACCACAATTTGCCTCAGTCTAA**ATG**ATCTCTCAGAATTTTTATATAGCAAGA**G**CTTTA**TGA**GCAAGCTATTGAGTATGAGAAAGGTTCGTTAATCACTTCTAATGGAGCTTTGACGATGCTTTCTGGTGCTAAGACTGATCGAGAGATAAGCGTGTTGTTAGAGATGCGATTACTGAGGATGAGCTTTGGCGATTACAGAATTTTTATCCTCTATTTTGGTTTATTGATTTTTGTTTTTGATTTTGATTTTGATTTTAAGGGTTTGTCTAATATTGAAATCGATGAACATACTTTCATGGTGGTGAACAGAGAAAGAGCTGTTGATTACTTGAATTCCTTGGAAAAG**GT**TTACCTTATGCCTAAAAGTT**AG**ATTCTTTCCCATCATTCTGGATGCAATATGGGAAAAGATGGAGATGTTGCCCTCTTCTTTGAACTTTTAGGTATTGCAACTTTT**TAA**GCCGACGTGAACGCTGAAAAAAAAGAGAAACACCGACGTGAACGCAGAGAGAAGATTACTTGCGCAACCATCGACGGAAAGAAAGAGAAACAAAACCTTTCTTCTTCTTTTTTTTTTCTATATTTGTCTAGAAAAAAAAAATTAACGTTTCTTTTGGATGTAAAGAATCTTTTCTATTTAGATGTTTTGGCACAAAAAACGAAGAAAAAAACTATAATTTTGTAATTTATTTTATTTTAGTTTATGATTAAG

Black – UTRs, AG – the acceptor site left out from the incorrect splicing of intron 1 that caused a frame shift. Red sequence – unspliced intron 2. TGA – early stop codon caused by the frame shist.

Pairwise alignment of XLOC_006026 and TCONS_00009618:

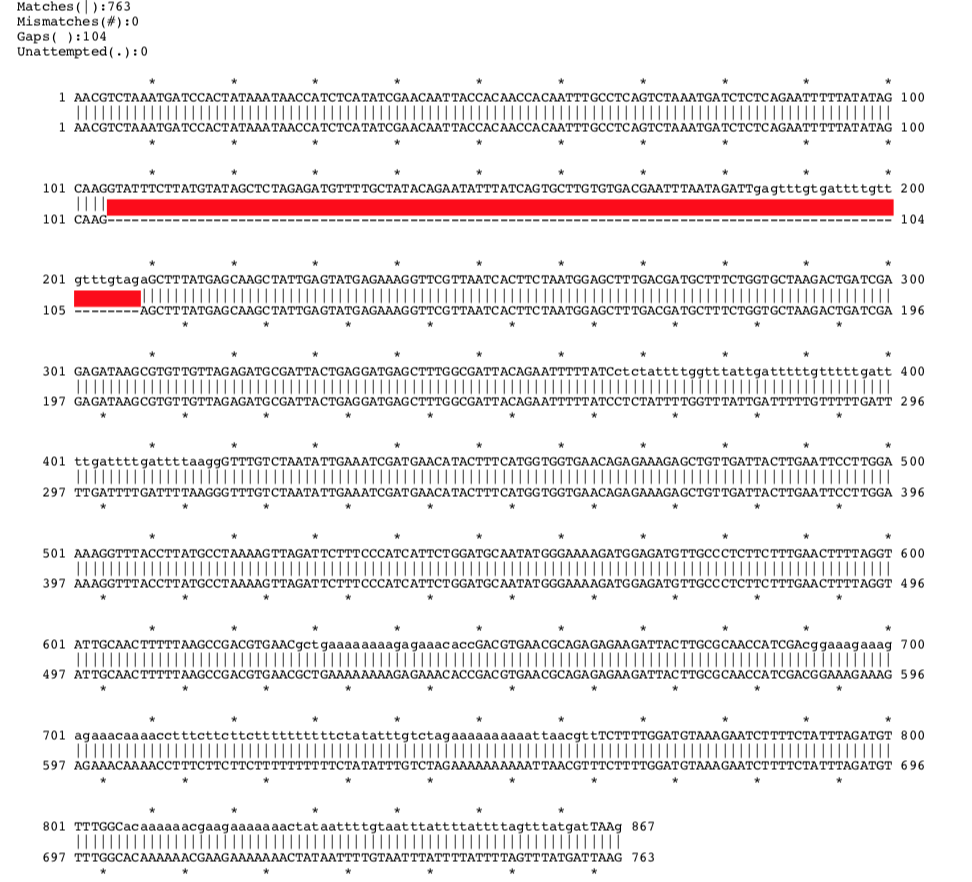

Position 107 to 508 with stop at 422-424

**F15O4.6/Q9LQH8 protein sequence from UnipProt:**

>tr|Q9LQH8|Q9LQH8_ARATH F15O4.6 OS=Arabidopsis thaliana OX=3702 PE=4 SV=1

MISQNFYIARYFLCIALEMFCYTEYLSVLLYEQAIEYEKGSLITSNGALTMLSGAKTDRE

ISVLLEMRLLRMSFGDYRIFILYFGLLIFVFDFDFDFKGLSNIEIDEHTFMVVNRERAVD

YLNSLEKILSHHSGCNMGKDGDVALFFELLGIATF

In red – the polypeptide part of the protein coded by 3’ half of exon 1 (see above in orange) that is excluded from TCONS_00009618
