## Supplemental Doc 1 for "Characterization of novel pollen-expressed transcripts reveals their potential roles in pollen heat stress response in *Arabidopsis thaliana*" (from another version of this preprint: version 2, 2020-08-26)

```
BLSTX of all 312 XLOCs against AT protein database


Job Title:312 sequences (XLOC_000101)
Program:
Database: nr All non-redundant GenBank CDS translations+PDB+SwissProt+PIR+PRF excluding environmental samples from WGS projects

Query #1: XLOC_000101 Query ID: lcl|Query_76535 Length: 713

No significant similarity found.


Query #2: XLOC_000218 Query ID: lcl|Query_76536 Length: 1006

No significant similarity found.


Query #3: XLOC_001358 Query ID: lcl|Query_76537 Length: 491

No significant similarity found.


Query #4: XLOC_001411 Query ID: lcl|Query_76538 Length: 798

No significant similarity found.


Query #5: XLOC_001621 Query ID: lcl|Query_76539 Length: 547

No significant similarity found.


Query #6: XLOC_001666 Query ID: lcl|Query_76540 Length: 1606

Sequences producing significant alignments:
                                                                  Max    Total Query   E   Per.
Description                                                       Score  Score cover Value Ident  Accession
AT5g14840/T9L3_140 [Arabidopsis thaliana]                         124    124   15%   8e-34 79.52  AAL16245.1
unnamed protein product [Arabidopsis thaliana]                    80.1   147   30%   2e-28 74.07  CAA0402639.1
At3g06433 [Arabidopsis thaliana]                                  108    108   12%   6e-28 83.58  AAS49072.1
hypothetical protein AXX17_AT1G32540 [Arabidopsis thaliana]       100    100   9%    2e-25 96.15  OAP19103.1
putative protein [Arabidopsis thaliana]                           97.8   164   49%   5e-21 43.09  CAC01883.1
hypothetical protein AXX17_AT3G54560 [Arabidopsis thaliana]       88.2   88.2  23%   3e-18 43.56  OAP02832.1
unnamed protein product [Arabidopsis thaliana]                    70.5   197   20%   6e-14 51.24  VYS49654.1
hypothetical protein AT1G24851 [Arabidopsis thaliana]             68.2   325   22%   5e-12 59.22  NP_173877.1
hypothetical protein AT1G25025 [Arabidopsis thaliana]             68.2   325   22%   5e-12 59.22  NP_173883.1
hypothetical protein AT4G20690 [Arabidopsis thaliana]             54.7   54.7  8%    2e-08 63.83  NP_193801.1


Alignments:
>AT5g14840/T9L3_140 [Arabidopsis thaliana]
Sequence ID: AAL16245.1 Length: 83
>putative protein [Arabidopsis thaliana]
Sequence ID: AAM20551.1 Length: 83 >putative protein [Arabidopsis thaliana]
Sequence ID: AAM91297.1 Length: 83
Range 1: 1 to 83

Score:124 bits(312), Expect:8e-34,
Method:Compositional matrix adjust.,
Identities:66/83(80%), Positives:70/83(84%), Gaps:0/83(0%)

Query  792  INYVWLDPFQRVRRQLFTRFSYKTNKLYSSSDIKIVFRPQARLSSTIGSFILRLYSLKRS  613
            +NYVWLDPF RVRRQ FTRFSYKTNKLYSSSDIKIVFRPQ RL STI SF+L L SLK+S
Sbjct  1    MNYVWLDPFLRVRRQPFTRFSYKTNKLYSSSDIKIVFRPQVRLLSTISSFLLHLCSLKKS  60

Query  612  SFVIRPICLIAGSLLKERFFFFP  544
            SFVIRPI L  G +L ERFFF P
Sbjct  61   SFVIRPIRLSRGLVLIERFFFLP  83


>unnamed protein product [Arabidopsis thaliana]
Sequence ID: CAA0402639.1 Length: 170
Range 1: 112 to 165

Score:80.1 bits(196), Expect:2e-28,
Method:Compositional matrix adjust.,
Identities:40/54(74%), Positives:42/54(77%), Gaps:0/54(0%)

Query  664  FIYNRLVHTPSILVKEIILCNPSNLPNRGLVIKRKIFLLSLEIPISYLCDSCIN  503
            FIYNRLV TPS+LV+EIILCNPSN P     I RKI L SLEIP SY CDSCIN
Sbjct  112  FIYNRLVLTPSVLVEEIILCNPSNSPKSRARINRKILLPSLEIPFSYFCDSCIN  165


Range 2: 11 to 112

Score:67.8 bits(164), Expect:2e-28,
Method:Compositional matrix adjust.,
Identities:55/108(51%), Positives:63/108(58%), Gaps:6/108(5%)

Query  986  HSPVPSSSFTCPSEVSRRLARPK*LTYSHVSRSCSAHHLPAGSSRNCLTQHTTYLQ*WPS  807
            H    SSS   PSEVSR+ AR K L  S   RS S    PAG S+  +  H   L+
Sbjct  11   HLETKSSSSAHPSEVSRQPARLKQLLVSSPVRSIS----PAGPSQ--VANHLARLKQLII  64

Query  806  SFITI*TTYGLIPFKGYVGSFLQGSAIKRINSILPRTSRSSSGLKLAF  663
            S ++  ++        YVGS LQGSAIKRINSILPRTSRSSSGLK AF
Sbjct  65   SPVSSSSSAHPSEVPRYVGSLLQGSAIKRINSILPRTSRSSSGLKFAF  112


>At3g06433 [Arabidopsis thaliana]
Sequence ID: AAS49072.1 Length: 101
Range 1: 1 to 67

Score:108 bits(271), Expect:6e-28,
Method:Compositional matrix adjust.,
Identities:56/67(84%), Positives:59/67(88%), Gaps:0/67(0%)

Query  792  INYVWLDPFQRVRRQLFTRFSYKTNKLYSSSDIKIVFRPQARLSSTIGSFILRLYSLKRS  613
            +NYVWLDPF RVRRQ FTRFSYKTNKLYSSSDIK VFRPQARLSSTI SF+L   SLK+S
Sbjct  1    MNYVWLDPFLRVRRQPFTRFSYKTNKLYSSSDIKFVFRPQARLSSTISSFLLHQCSLKKS  60

Query  612  SFVIRPI  592
            SF IRPI
Sbjct  61   SFEIRPI  67


>hypothetical protein AXX17_AT1G32540 [Arabidopsis thaliana]
Sequence ID: OAP19103.1 Length: 52
Range 1: 1 to 52

Score:100 bits(249), Expect:2e-25,
Method:Compositional matrix adjust.,
Identities:50/52(96%), Positives:50/52(96%), Gaps:0/52(0%)

Query  115  MGDSISLLLHRFSSFSCGKIGGSSELDMKVDLSQTHIDCSGFHFLNSTHHAW  270
            MGDSISLLLHRFSSFSC KIGGSSELDMKVDLSQTH DCSGFHFLNSTHHAW
Sbjct  1    MGDSISLLLHRFSSFSCDKIGGSSELDMKVDLSQTHSDCSGFHFLNSTHHAW  52


>putative protein, partial [Arabidopsis thaliana]
Sequence ID: CAC01883.1 Length: 472
Range 1: 44 to 192

Score:97.8 bits(242), Expect:5e-21,
Method:Compositional matrix adjust.,
Identities:81/188(43%), Positives:97/188(51%), Gaps:39/188(20%)

Query  1066  SACPSQTARQLARLKQLASSPDRSISSTRPSRVARLLVRPKCLADLPARSNLPTHTFQEA  887
             ++ PS+ ARQL RLK L S    S SS  PS+ AR L RPK LA  P  S+L +   Q A
Sbjct  44    ASGPSKAARQLTRLKHLVSRSVSSSSSAHPSQAARQLTRPKYLASRPVSSSLSSRPSQAA  103

Query  886   AQLITFPPDRLETA*LNIRHTFSNGLARLLRYKLRMA*SLSKGT*AAFYKVQL*NE*TLF  707
              QL T P   +  + L+   T  + ++R
Sbjct  104   RQL-TRPKYLVCPSILSSSFTRPSEVSR--------------------------------  130

Query  706   FLGHQDRLPASSSPFIYNRLVHTPSILVKEIILCNPSNLPNRGLVIKRKIFLLSLEIPIS  527
                   RL   +SPFIYN+LV TPS+LV+EIILCNPSN P     I RKIFL SLEIP S
Sbjct  131   ------RLARPNSPFIYNQLVLTPSVLVEEIILCNPSNSPKSRARINRKIFLPSLEIPFS  184

Query  526   YLCDSCIN  503
             YLCDSCIN
Sbjct  185   YLCDSCIN  192


Range 2: 36 to 136

Score:66.6 bits(161), Expect:6e-11,
Method:Compositional matrix adjust.,
Identities:65/129(50%), Positives:74/129(57%), Gaps:28/129(21%)

Query  1304  AEYYKYGIACPVKRIWIRLDTQRQRALIHLPVSNSSSACPSQTARQLARLEQLVSLPVSN  1125
             AE YKYGIA                            + PS+ ARQL RL+ LVS  VS+
Sbjct  36    AEGYKYGIA----------------------------SGPSKAARQLTRLKHLVSRSVSS  67

Query  1124  SSSACPSRTARQLARLEQlvslsvsnssSACPSQTARQFARSKYLVHSPVPSSSFTCPSE  945
             SSSA PS+ ARQL R + L S  VS+S S+ PSQ ARQ  R KYLV   + SSSFT PSE
Sbjct  68    SSSAHPSQAARQLTRPKYLASRPVSSSLSSRPSQAARQLTRPKYLVCPSILSSSFTRPSE  127

Query  944   VSRRLARPK  918
             VSRRLARP
Sbjct  128   VSRRLARPN  136


>hypothetical protein AXX17_AT3G54560 [Arabidopsis thaliana]
Sequence ID: OAP02832.1 Length: 358
Range 1: 195 to 357

Score:88.2 bits(217), Expect:3e-18,
Method:Compositional matrix adjust.,
Identities:71/163(44%), Positives:84/163(51%), Gaps:37/163(22%)

Query  1234  KELSSTCPSQTARQLARLKQLVNLPVSNSSSACPSRTARQLARLEQLVSLPVSNSSSAC-  1058
             + +SS  P Q A   AR KQLV+ PV + S A PS+ A Q A L+QLVS PV + S A
Sbjct  195   RSISSAGPPQVAHPFARRKQLVSSPVRSISPAGPSQVAYQFAGLKQLVSSPVRSISPAGP  254

Query  1057  -------PSQTARQLARLKQLASSPDR---------------------------SISSTR  980
                    PSQ ARQL R K LAS                               S S TR
Sbjct  255   SSSSSVRPSQAARQLTRPKYLASRRSLSSSSSVRPSQAARQLTRPKYLVCPSILSSSFTR  314

Query  979   PSRVARLLVRPKCLADLPAR--SNLPTHTFQEAAQLITFPPDR  857
             PS V+R L RPKCLA+LP    S+LPT T +EA QL+T+PPD
Sbjct  315   PSEVSRQLARPKCLANLPVHSTSSLPTRTSREAIQLVTYPPDH  357


>unnamed protein product [Arabidopsis thaliana]
Sequence ID: VYS49654.1 Length: 131
Range 1: 11 to 131

Score:70.5 bits(171), Expect:6e-14,
Method:Compositional matrix adjust.,
Identities:62/121(51%), Positives:71/121(58%), Gaps:20/121(16%)

Query  1220  HLPVSNSSSACPSQTARQLARLEQLVS--------------------LPVSNSSSACPSR  1101
             HL + +SSSA PSQ ARQL RL+ LVS                     PVS++ SA PS+
Sbjct  11    HLEIKSSSSALPSQPARQLTRLKHLVSSSARQSQAARQLTRPKYLASRPVSSTLSARPSQ  70

Query  1100  TARQLARLEQlvslsvsnssSACPSQTARQFARSKYLVHSPVPSSSFTCPSEVSRRLARP  921
              ARQL R + L S  VS+S SA  SQ  RQ  R KYLV   + SSSFT PSEVSRRL RP
Sbjct  71    AARQLTRPKYLASRPVSSSLSARSSQADRQLTRPKYLVCPSILSSSFTRPSEVSRRLDRP  130

Query  920   K  918
             K
Sbjct  131   K  131


Range 2: 6 to 107

Score:65.1 bits(157), Expect:6e-12,
Method:Compositional matrix adjust.,
Identities:57/108(53%), Positives:64/108(59%), Gaps:9/108(8%)

Query  1252  GLTRRDKEL---SSTCPSQTARQLARLKQLVNLPVSNSSSACPSRTARQLARLEQLVSLP  1082
             G+ R   E+   SS  PSQ ARQL RLK LV      SSSA  S+ ARQL R + L S P
Sbjct  6     GIPRGHLEIKSSSSALPSQPARQLTRLKHLV------SSSARQSQAARQLTRPKYLASRP  59

Query  1081  VSNSSSACPSQTARQLARLKQLASSPDRSISSTRPSRVARLLVRPKCL  938
             VS++ SA PSQ ARQL R K LAS P  S  S R S+  R L RPK L
Sbjct  60    VSSTLSARPSQAARQLTRPKYLASRPVSSSLSARSSQADRQLTRPKYL  107


Range 3: 11 to 85

Score:62.0 bits(149), Expect:6e-11,
Method:Compositional matrix adjust.,
Identities:45/81(56%), Positives:51/81(62%), Gaps:6/81(7%)

Query  1168  NLPVSNSSSACPSRTARQLARLEQLVSLPVSNSSSACPSQTARQLARLKQLASSPDRSIS  989
             +L + +SSSA PS+ ARQL RL+ LVS      SSA  SQ ARQL R K LAS P  S
Sbjct  11    HLEIKSSSSALPSQPARQLTRLKHLVS------SSARQSQAARQLTRPKYLASRPVSSTL  64

Query  988   STRPSRVARLLVRPKCLADLP  926
             S RPS+ AR L RPK LA  P
Sbjct  65    SARPSQAARQLTRPKYLASRP  85


>hypothetical protein AT1G24851 [Arabidopsis thaliana]
Sequence ID: NP_173877.1 Length: 258
>F5A9.15 [Arabidopsis thaliana]
Sequence ID: AAG03125.1 Length: 258 >hypothetical protein AT1G24851 [Arabidopsis thaliana]
Sequence ID: AEE30570.1 Length: 258
Range 1: 156 to 258

Score:68.2 bits(165), Expect:5e-12,
Method:Compositional matrix adjust.,
Identities:61/103(59%), Positives:71/103(68%), Gaps:0/103(0%)

Query  1226  LIHLPVSNSSSACPSQTARQLARLEQLVSLPVSNSSSACPSRTARQLARLEQlvslsvsn  1047
             L+  P+ +   A PSQTARQL R +   S  VS+SSSA PS+ ARQL RL+QLVS  VS+
Sbjct  156   LVSSPLRSIPPAGPSQTARQLTRPKYPASRFVSSSSSAHPSQAARQLTRLKQLVSSPVSS  215

Query  1046  ssSACPSQTARQFARSKYLVHSPVPSSSFTCPSEVSRRLARPK  918
             SSSA P Q ARQ  R KYLV   + S+SFT PSEVS R ARPK
Sbjct  216   SSSAHPYQAARQLTRPKYLVCPSILSNSFTRPSEVSLRPARPK  258


Range 2: 122 to 228

Score:67.8 bits(164), Expect:5e-12,
Method:Compositional matrix adjust.,
Identities:55/107(51%), Positives:64/107(59%), Gaps:0/107(0%)

Query  1198  RQLARLKQLVNLPVSNSSSACPSRTARQLARLEQLVSLPVSNSSSACPSQTARQLARLKQ  1019
             RQL RLKQLV+ P+ +   A PS+ ARQL RL QLVS P+ +   A PSQTARQL R K
Sbjct  122   RQLTRLKQLVSSPLRSIPPAGPSQAARQLTRLNQLVSSPLRSIPPAGPSQTARQLTRPKY  181

Query  1018  LASSPDRSISSTRPSRVARLLVRPKCLADLPARSNLPTHTFQEAAQL  878
              AS    S SS  PS+ AR L R K L   P  S+   H +Q A QL
Sbjct  182   PASRFVSSSSSAHPSQAARQLTRLKQLVSSPVSSSSSAHPYQAARQL  228


Range 3: 136 to 234

Score:67.0 bits(162), Expect:1e-11,
Method:Compositional matrix adjust.,
Identities:55/99(56%), Positives:64/99(64%), Gaps:0/99(0%)

Query  1234  KELSSTCPSQTARQLARLKQLVNLPVSNSSSACPSRTARQLARLEQLVSLPVSNSSSACP  1055
             + +    PSQ ARQL RL QLV+ P+ +   A PS+TARQL R +   S  VS+SSSA P
Sbjct  136   RSIPPAGPSQAARQLTRLNQLVSSPLRSIPPAGPSQTARQLTRPKYPASRFVSSSSSAHP  195

Query  1054  SQTARQLARLKQLASSPDRSISSTRPSRVARLLVRPKCL  938
             SQ ARQL RLKQL SSP  S SS  P + AR L RPK L
Sbjct  196   SQAARQLTRLKQLVSSPVSSSSSAHPYQAARQLTRPKYL  234


Range 4: 91 to 202

Score:66.2 bits(160), Expect:2e-11,
Method:Compositional matrix adjust.,
Identities:60/112(54%), Positives:68/112(60%), Gaps:0/112(0%)

Query  1213  PSQTARQLARLKQLVNLPVSNSSSACPSRTARQLARLEQLVSLPVSNSSSACPSQTARQL  1034
             PSQ ARQL R K   +  VS+SSSA PS+  RQL RL+QLVS P+ +   A PSQ ARQL
Sbjct  91    PSQAARQLTRPKYPASRSVSSSSSAHPSQADRQLTRLKQLVSSPLRSIPPAGPSQAARQL  150

Query  1033  ARLKQLASSPDRSISSTRPSRVARLLVRPKCLADLPARSNLPTHTFQEAAQL  878
              RL QL SSP RSI    PS+ AR L RPK  A     S+   H  Q A QL
Sbjct  151   TRLNQLVSSPLRSIPPAGPSQTARQLTRPKYPASRFVSSSSSAHPSQAARQL  202


Range 5: 122 to 220

Score:56.2 bits(134), Expect:4e-08,
Method:Compositional matrix adjust.,
Identities:52/99(53%), Positives:61/99(61%), Gaps:0/99(0%)

Query  1172  RQLARLEQLVSLPVSNSSSACPSRTARQLARLEQlvslsvsnssSACPSQTARQFARSKY  993
             RQL RL+QLVS P+ +   A PS+ ARQL RL QLVS  + +   A PSQTARQ  R KY
Sbjct  122   RQLTRLKQLVSSPLRSIPPAGPSQAARQLTRLNQLVSSPLRSIPPAGPSQTARQLTRPKY  181

Query  992   LVHSPVPSSSFTCPSEVSRRLARPK*LTYSHVSRSCSAH  876
                  V SSS   PS+ +R+L R K L  S VS S SAH
Sbjct  182   PASRFVSSSSSAHPSQAARQLTRLKQLVSSPVSSSSSAH  220


>hypothetical protein AT1G25025 [Arabidopsis thaliana]
Sequence ID: NP_173883.1 Length: 258
>hypothetical protein AT1G25112 [Arabidopsis thaliana]
Sequence ID: NP_173887.1 Length: 258 >hypothetical protein AT1G25180 [Arabidopsis thaliana]
Sequence ID: NP_173891.1 Length: 258 >F5A9.9 [Arabidopsis thaliana]
Sequence ID: AAG03127.1 Length: 258 >F5A9.5 [Arabidopsis thaliana]
Sequence ID: AAG03129.1 Length: 258 >F5A9.1 [Arabidopsis thaliana]
Sequence ID: AAG03131.1 Length: 258 >unknown protein [Arabidopsis thaliana]
Sequence ID: AAG28803.1 Length: 258 >At1g25025 [Arabidopsis thaliana]
Sequence ID: ABK59689.1 Length: 258
Range 1: 156 to 258

Score:68.2 bits(165), Expect:5e-12,
Method:Compositional matrix adjust.,
Identities:61/103(59%), Positives:71/103(68%), Gaps:0/103(0%)

Query  1226  LIHLPVSNSSSACPSQTARQLARLEQLVSLPVSNSSSACPSRTARQLARLEQlvslsvsn  1047
             L+  P+ +   A PSQTARQL R +   S  VS+SSSA PS+ ARQL RL+QLVS  VS+
Sbjct  156   LVSSPLRSIPPAGPSQTARQLTRPKYPASRFVSSSSSAHPSQAARQLTRLKQLVSSPVSS  215

Query  1046  ssSACPSQTARQFARSKYLVHSPVPSSSFTCPSEVSRRLARPK  918
             SSSA P Q ARQ  R KYLV   + S+SFT PSEVS R ARPK
Sbjct  216   SSSAHPYQAARQLTRPKYLVCPSILSNSFTRPSEVSLRPARPK  258


Range 2: 122 to 228

Score:67.8 bits(164), Expect:6e-12,
Method:Compositional matrix adjust.,
Identities:55/107(51%), Positives:64/107(59%), Gaps:0/107(0%)

Query  1198  RQLARLKQLVNLPVSNSSSACPSRTARQLARLEQLVSLPVSNSSSACPSQTARQLARLKQ  1019
             RQL RLKQLV+ P+ +   A PS+ ARQL RL QLVS P+ +   A PSQTARQL R K
Sbjct  122   RQLTRLKQLVSSPLRSIPPAGPSQAARQLTRLNQLVSSPLRSIPPAGPSQTARQLTRPKY  181

Query  1018  LASSPDRSISSTRPSRVARLLVRPKCLADLPARSNLPTHTFQEAAQL  878
              AS    S SS  PS+ AR L R K L   P  S+   H +Q A QL
Sbjct  182   PASRFVSSSSSAHPSQAARQLTRLKQLVSSPVSSSSSAHPYQAARQL  228


Range 3: 136 to 234

Score:67.0 bits(162), Expect:1e-11,
Method:Compositional matrix adjust.,
Identities:55/99(56%), Positives:64/99(64%), Gaps:0/99(0%)

Query  1234  KELSSTCPSQTARQLARLKQLVNLPVSNSSSACPSRTARQLARLEQLVSLPVSNSSSACP  1055
             + +    PSQ ARQL RL QLV+ P+ +   A PS+TARQL R +   S  VS+SSSA P
Sbjct  136   RSIPPAGPSQAARQLTRLNQLVSSPLRSIPPAGPSQTARQLTRPKYPASRFVSSSSSAHP  195

Query  1054  SQTARQLARLKQLASSPDRSISSTRPSRVARLLVRPKCL  938
             SQ ARQL RLKQL SSP  S SS  P + AR L RPK L
Sbjct  196   SQAARQLTRLKQLVSSPVSSSSSAHPYQAARQLTRPKYL  234


Range 4: 91 to 202

Score:66.2 bits(160), Expect:2e-11,
Method:Compositional matrix adjust.,
Identities:60/112(54%), Positives:68/112(60%), Gaps:0/112(0%)

Query  1213  PSQTARQLARLKQLVNLPVSNSSSACPSRTARQLARLEQLVSLPVSNSSSACPSQTARQL  1034
             PSQ ARQL R K   +  VS+SSSA PS+  RQL RL+QLVS P+ +   A PSQ ARQL
Sbjct  91    PSQAARQLTRPKYPASRSVSSSSSAHPSQADRQLTRLKQLVSSPLRSIPPAGPSQAARQL  150

Query  1033  ARLKQLASSPDRSISSTRPSRVARLLVRPKCLADLPARSNLPTHTFQEAAQL  878
              RL QL SSP RSI    PS+ AR L RPK  A     S+   H  Q A QL
Sbjct  151   TRLNQLVSSPLRSIPPAGPSQTARQLTRPKYPASRFVSSSSSAHPSQAARQL  202


Range 5: 122 to 220

Score:56.2 bits(134), Expect:5e-08,
Method:Compositional matrix adjust.,
Identities:52/99(53%), Positives:61/99(61%), Gaps:0/99(0%)

Query  1172  RQLARLEQLVSLPVSNSSSACPSRTARQLARLEQlvslsvsnssSACPSQTARQFARSKY  993
             RQL RL+QLVS P+ +   A PS+ ARQL RL QLVS  + +   A PSQTARQ  R KY
Sbjct  122   RQLTRLKQLVSSPLRSIPPAGPSQAARQLTRLNQLVSSPLRSIPPAGPSQTARQLTRPKY  181

Query  992   LVHSPVPSSSFTCPSEVSRRLARPK*LTYSHVSRSCSAH  876
                  V SSS   PS+ +R+L R K L  S VS S SAH
Sbjct  182   PASRFVSSSSSAHPSQAARQLTRLKQLVSSPVSSSSSAH  220


>hypothetical protein AT4G20690 [Arabidopsis thaliana]
Sequence ID: NP_193801.1 Length: 138
>hypothetical protein AT4G20690 [Arabidopsis thaliana]
Sequence ID: AEE84354.1 Length: 138 >putative protein [Arabidopsis thaliana]
Sequence ID: CAB45835.1 Length: 138 >putative protein [Arabidopsis thaliana]
Sequence ID: CAB79069.1 Length: 138
Range 1: 67 to 113

Score:54.7 bits(130), Expect:2e-08,
Method:Compositional matrix adjust.,
Identities:30/47(64%), Positives:35/47(74%), Gaps:0/47(0%)

Query  690  IVFRPQARLSSTIGSFILRLYSLKRSSFVIRPICLIAGSLLKERFFF  550
            IVFRPQARLSSTIGSF+L L  LK+SSFVIRPI L  G ++  +
Sbjct  67   IVFRPQARLSSTIGSFLLYLCLLKKSSFVIRPIRLSRGLVINRKILL  113


Query #7: XLOC_001820 Query ID: lcl|Query_76541 Length: 462

No significant similarity found.


Query #8: XLOC_001852 Query ID: lcl|Query_76542 Length: 1702

Sequences producing significant alignments:
                                                                  Max    Total Query   E   Per.
Description                                                       Score  Score cover Value Ident  Accession
unnamed protein product [Arabidopsis thaliana]                    73.6   73.6  8%    4e-15 74.00  CAA0270315.1
unnamed protein product [Arabidopsis thaliana]                    73.6   73.6  8%    4e-15 74.00  VYS48141.1


Alignments:
>unnamed protein product [Arabidopsis thaliana]
Sequence ID: CAA0270315.1 Length: 113
Range 1: 55 to 104

Score:73.6 bits(179), Expect:4e-15,
Method:Compositional matrix adjust.,
Identities:37/50(74%), Positives:41/50(82%), Gaps:3/50(6%)

Query  1419  SLATLEKLC---KFFKHLQEEGKLLGRACDRP*LSIKCKERKCGCSTSYR  1279
             SLATLEKLC   KFFK + EEGKLL RACDR   SIKCKE+KCGCST+Y+
Sbjct  55    SLATLEKLCNHQKFFKQMHEEGKLLRRACDRSDFSIKCKEKKCGCSTNYQ  104


>unnamed protein product [Arabidopsis thaliana]
Sequence ID: VYS48141.1 Length: 113
Range 1: 55 to 104

Score:73.6 bits(179), Expect:4e-15,
Method:Compositional matrix adjust.,
Identities:37/50(74%), Positives:41/50(82%), Gaps:3/50(6%)

Query  1419  SLATLEKLC---KFFKHLQEEGKLLGRACDRP*LSIKCKERKCGCSTSYR  1279
             SLATLEKLC   KFFK + EEGKLL RACDR   SIKCKE+KCGCST+Y+
Sbjct  55    SLATLEKLCNHQKFFKQMHEEGKLLRRACDRSDFSIKCKEKKCGCSTNYQ  104


Query #9: XLOC_001879 Query ID: lcl|Query_76543 Length: 1651

Sequences producing significant alignments:
                                                                  Max    Total Query   E   Per.
Description                                                       Score  Score cover Value Ident  Accession
putative non-LTR retroelement reverse transcriptase [Arabidops... 100    100   12%   3e-21 66.67  AAD17398.1
non-LTR retroelement reverse transcriptase [Arabidopsis thaliana] 87.4   87.4  10%   1e-18 72.41  NP_179830.1
At3g42803 [Arabidopsis thaliana]                                  74.7   74.7  8%    3e-15 71.43  AAT41836.1
unnamed protein product [Arabidopsis thaliana]                    64.7   64.7  12%   2e-10 47.83  BAA97156.1
reverse transcriptase, putative; 16838-20266 [Arabidopsis...      65.5   65.5  12%   2e-10 50.72  AAG51783.1


Alignments:
>putative non-LTR retroelement reverse transcriptase [Arabidopsis thaliana]
Sequence ID: AAD17398.1 Length: 1225
Range 1: 978 to 1046

Score:100 bits(249), Expect:3e-21,
Method:Compositional matrix adjust.,
Identities:46/69(67%), Positives:56/69(81%), Gaps:0/69(0%)

Query  693   FFSITPSRQI*ALSPIPSSWNIFPRNSLFYNFDFIFWRGREFRLGEEVLKLFPWIIWYIW  514
              F   PSRQI ALSPIPSS  IFPRNSLFYNFDF+  RG+EF + E+++++FPWI+WYIW
Sbjct  978   LFLCPPSRQIWALSPIPSSEYIFPRNSLFYNFDFLLSRGKEFDIAEDIMEIFPWILWYIW  1037

Query  513   ISRNHFVFE  487
              SRN F+FE
Sbjct  1038  KSRNRFIFE  1046


>non-LTR retroelement reverse transcriptase [Arabidopsis thaliana]
Sequence ID: NP_179830.1 Length: 256
>putative non-LTR retroelement reverse transcriptase [Arabidopsis thaliana]
Sequence ID: AAD22356.1 Length: 256 >non-LTR retroelement reverse transcriptase [Arabidopsis thaliana]
Sequence ID: AEC07306.1 Length: 256
Range 1: 104 to 161

Score:87.4 bits(215), Expect:1e-18,
Method:Compositional matrix adjust.,
Identities:42/58(72%), Positives:47/58(81%), Gaps:0/58(0%)

Query  651  PIPSSWNIFPRNSLFYNFDFIFWRGREFRLGEEVLKLFPWIIWYIWISRNHFVFEK*R  478
            P     NIFPRNSLFYNFDF+FWR REF +GE+VL+LFPWIIWYIW S+N FVFE  R
Sbjct  104  PFQGPGNIFPRNSLFYNFDFLFWRDREFGIGEKVLELFPWIIWYIWKSKNRFVFENFR  161


>At3g42803 [Arabidopsis thaliana]
Sequence ID: AAT41836.1 Length: 146
Range 1: 98 to 146

Score:74.7 bits(182), Expect:3e-15,
Method:Compositional matrix adjust.,
Identities:35/49(71%), Positives:39/49(79%), Gaps:0/49(0%)

Query  693  FFSITPSRQI*ALSPIPSSWNIFPRNSLFYNFDFIFWRGREFRLGEEVL  547
             F   PSRQ  ALS IP+  NIFPRNSLFYNFDF+FWRGR+F +GEEVL
Sbjct  98   LFHCLPSRQTWALSLIPTLGNIFPRNSLFYNFDFLFWRGRKFGIGEEVL  146


>unnamed protein product [Arabidopsis thaliana]
Sequence ID: BAA97156.1 Length: 329
Range 1: 65 to 133

Score:64.7 bits(156), Expect:2e-10,
Method:Compositional matrix adjust.,
Identities:33/69(48%), Positives:46/69(66%), Gaps:0/69(0%)

Query  693  FFSITPSRQI*ALSPIPSSWNIFPRNSLFYNFDFIFWRGREFRLGEEVLKLFPWIIWYIW  514
             F   P+RQ+ ALS IPSS + F  +S++ N D+++WR  E    EE L++FPWI+WYIW
Sbjct  65   IFECPPARQVWALSGIPSSPSRFLSSSIYNNLDYLYWRANEIGACEESLRVFPWIMWYIW  124

Query  513  ISRNHFVFE  487
             +RN   FE
Sbjct  125  KARNRKNFE  133


>reverse transcriptase, putative; 16838-20266 [Arabidopsis thaliana]
Sequence ID: AAG51783.1 Length: 1142
Range 1: 880 to 944

Score:65.5 bits(158), Expect:2e-10,
Method:Compositional matrix adjust.,
Identities:35/69(51%), Positives:43/69(62%), Gaps:4/69(5%)

Query  693  FFSITPSRQI*ALSPIPSSWNIFPRNSLFYNFDFIFWRGREFRLGEEVLKLFPWIIWYIW  514
             F   P+RQI ALS IP++  IFP NS+F N D +FWR      G +    +PWIIWYIW
Sbjct  880  LFQCHPARQIWALSQIPTAPGIFPSNSIFTNLDHLFWR---IPSGVDSAP-YPWIIWYIW  935

Query  513  ISRNHFVFE  487
             +RN  VFE
Sbjct  936  KARNEKVFE  944


Query #10: XLOC_001883 Query ID: lcl|Query_76544 Length: 422

Sequences producing significant alignments:
                                                                  Max    Total Query   E   Per.
Description                                                       Score  Score cover Value Ident  Accession
Mannose-binding lectin superfamily protein [Arabidopsis thaliana] 66.6   66.6  27%   4e-13 74.36  NP_175623.2
unnamed protein product [Arabidopsis thaliana]                    64.3   64.3  27%   3e-12 71.79  CAA0288322.1
unnamed protein product [Arabidopsis thaliana]                    58.5   58.5  22%   1e-11 90.32  CAA0405760.1
MATE efflux family protein [Arabidopsis thaliana]                 53.1   53.1  21%   7e-09 76.67  NP_178495.1


Alignments:
>Mannose-binding lectin superfamily protein [Arabidopsis thaliana]
Sequence ID: NP_175623.2 Length: 615
>Mannose-binding lectin superfamily protein [Arabidopsis thaliana]
Sequence ID: AEE32756.1 Length: 615
Range 1: 1 to 39

Score:66.6 bits(161), Expect:4e-13,
Method:Composition-based stats.,
Identities:29/39(74%), Positives:38/39(97%), Gaps:0/39(0%)

Query  70   IYNYSRSPGFYAEKILKVSIKRMEKEKEVSSVLDLIRVG  186
            +YNYSRSPGFYA+KILK+SIKRM+K+ E+SSV+DLI++G
Sbjct  1    MYNYSRSPGFYAKKILKMSIKRMKKKNEMSSVVDLIKIG  39


>unnamed protein product [Arabidopsis thaliana]
Sequence ID: CAA0288322.1 Length: 614
Range 1: 1 to 39

Score:64.3 bits(155), Expect:3e-12,
Method:Composition-based stats.,
Identities:28/39(72%), Positives:37/39(94%), Gaps:0/39(0%)

Query  70   IYNYSRSPGFYAEKILKVSIKRMEKEKEVSSVLDLIRVG  186
            +YNYS SPGFYA+KILK+SIKRM+K+ E+SSV+DLI++G
Sbjct  1    MYNYSHSPGFYAKKILKMSIKRMKKKNEMSSVVDLIKIG  39


>unnamed protein product [Arabidopsis thaliana]
Sequence ID: CAA0405760.1 Length: 84
Range 1: 50 to 80

Score:58.5 bits(140), Expect:1e-11,
Method:Compositional matrix adjust.,
Identities:28/31(90%), Positives:30/31(96%), Gaps:0/31(0%)

Query  94   GFYAEKILKVSIKRMEKEKEVSSVLDLIRVG  186
            GFYAEKILKVSIKRMEKE +VSSV+DLIRVG
Sbjct  50   GFYAEKILKVSIKRMEKENDVSSVMDLIRVG  80


>MATE efflux family protein [Arabidopsis thaliana]
Sequence ID: NP_178495.1 Length: 171
>predicted protein [Arabidopsis thaliana]
Sequence ID: AAM15444.1 Length: 171 >MATE efflux family protein [Arabidopsis thaliana]
Sequence ID: AEC05795.1 Length: 171
Range 1: 3 to 32

Score:53.1 bits(126), Expect:7e-09,
Method:Compositional matrix adjust.,
Identities:23/30(77%), Positives:28/30(93%), Gaps:0/30(0%)

Query  64   TTIYNYSRSPGFYAEKILKVSIKRMEKEKE  153
             +++NYSRSPGFY EKILKVSIKR+EK+KE
Sbjct  3    ASVHNYSRSPGFYVEKILKVSIKRIEKKKE  32


Query #11: XLOC_001945 Query ID: lcl|Query_76545 Length: 669

Sequences producing significant alignments:
                                                                  Max    Total Query   E   Per.
Description                                                       Score  Score cover Value Ident  Accession
putative MuDR-A-like transposon protein [Arabidopsis thaliana]    100    100   48%   7e-24 46.15  AAB81881.1
putative Mutator-like transposase; 12516-14947 [Arabidopsis...    97.8   97.8  48%   7e-23 45.45  AAG52094.1
mutator transposase MUDRA, putative; 66518-63770 [Arabidopsis...  75.5   75.5  53%   3e-15 38.24  AAG51216.1


Alignments:
>putative MuDR-A-like transposon protein [Arabidopsis thaliana]
Sequence ID: AAB81881.1 Length: 761
>putative MuDR-A-like transposon protein [Arabidopsis thaliana]
Sequence ID: CAB77993.1 Length: 761
Range 1: 617 to 759

Score:100 bits(250), Expect:7e-24,
Method:Compositional matrix adjust.,
Identities:66/143(46%), Positives:82/143(57%), Gaps:35/143(24%)

Query  110  KIWMTSTYRLVTSPPEPILPGKKKTNRQKT-YPRIKGKHESPKKKTIK----ETLGRKGR  274
            K W+ S+Y LVT+PPEPILPG+KK   +K  + RIKGK+ESPKKK  K    E LG+KG+
Sbjct  617  KYWLNSSYGLVTAPPEPILPGRKKEKSKKEKFARIKGKNESPKKKKRKKNEVEKLGKKGK  676

Query  275  IIHCSSCGEAGHNATRCNLYPKEKTKRQR------TTEVE--------------------  376
            IIHC SCGEAGHNA RC  +PKEK  R+R        E+E
Sbjct  677  IIHCKSCGEAGHNALRCKKFPKEKVPRKRKSKNDGNDEIEKKAKKAKNDKKDTNEKDSKK  736

Query  377  ----PSSSQVPYTISLTQPSQTS  433
                   + +P TI +TQPSQ +
Sbjct  737  EKKSKKETHMPDTIFITQPSQAT  759


>putative Mutator-like transposase; 12516-14947 [Arabidopsis thaliana]
Sequence ID: AAG52094.1 Length: 761
Range 1: 617 to 759

Score:97.8 bits(242), Expect:7e-23,
Method:Compositional matrix adjust.,
Identities:65/143(45%), Positives:81/143(56%), Gaps:35/143(24%)

Query  110  KIWMTSTYRLVTSPPEPILPGKKKTNRQKT-YPRIKGKHESPKKKTIK----ETLGRKGR  274
            K W+ S+Y LVT+PPEPILPG+KK   +K  + RIKGK+ESPKKK  K    E LG+KG+
Sbjct  617  KYWLNSSYGLVTAPPEPILPGRKKEKSKKGKFARIKGKNESPKKKKRKKNEVEKLGKKGK  676

Query  275  IIHCSSCGEAGHNATRCNLYPKEKTKRQR------TTEVE--------------------  376
            IIHC SCGEAGHNA  C  +PKEK  R+R        E+E
Sbjct  677  IIHCKSCGEAGHNALGCKKFPKEKVPRKRKSKNDGNDEIEKKAKKAKNDKKDTDEKNSKK  736

Query  377  ----PSSSQVPYTISLTQPSQTS  433
                   + +P TI +TQPSQ +
Sbjct  737  EKKSKKETHMPDTIFITQPSQAT  759


>mutator transposase MUDRA, putative; 66518-63770 [Arabidopsis thaliana]
Sequence ID: AAG51216.1 Length: 826
Range 1: 666 to 800

Score:75.5 bits(184), Expect:3e-15,
Method:Composition-based stats.,
Identities:52/136(38%), Positives:78/136(57%), Gaps:18/136(13%)

Query  68   YSDAKKSHFSCSP----AKIWMTSTYRLVTSP-PEPILPGKKKTNRQKTYPRIKGKHESP  232
            Y+ A +  +   P     +++  +   L+T+P  +P+ PG+KK   +K Y RIKG +ESP
Sbjct  666  YTIAWREQYDTGPDPVRGQMYWPTGLGLITAPLQDPVPPGRKK-GEKKNYHRIKGPNESP  724

Query  233  KKKTIKE-TLGRKGRIIHCSSCGEAGHNATRCNLYPKEKTKRQRTT-EVEPSSSQVPY--  400
            KKK  +   +G+KG ++HC SCGEAGHNA  C  +PKEK  R++   E++  +SQ
Sbjct  725  KKKGPQRLKVGKKGVVMHCKSCGEAGHNAAGCKKFPKEKKGRKKKNQEIKAGTSQPTMEL  784

Query  401  --------TISLTQPS  424
                    TI+LTQ S
Sbjct  785  QETTHGADTITLTQRS  800


Query #12: XLOC_001980 Query ID: lcl|Query_76546 Length: 584

No significant similarity found.


Query #13: XLOC_002167 Query ID: lcl|Query_76547 Length: 444

No significant similarity found.


Query #14: XLOC_002199 Query ID: lcl|Query_76548 Length: 492

Sequences producing significant alignments:
                                                                  Max    Total Query   E   Per.
Description                                                       Score  Score cover Value Ident  Accession
putative non-LTR retroelement reverse transcriptase [Arabidops... 53.9   53.9  29%   2e-08 51.92  AAC33226.1


Alignments:
>putative non-LTR retroelement reverse transcriptase [Arabidopsis thaliana]
Sequence ID: AAC33226.1 Length: 1529
Range 1: 1380 to 1431

Score:53.9 bits(128), Expect:2e-08,
Method:Compositional matrix adjust.,
Identities:27/52(52%), Positives:35/52(67%), Gaps:3/52(5%)

Query  294   AAGDRVNKWNTRQQVDCT---NIEETRDPLLFSCPYSSILSKDLAQRLLTTS  148
             + GDR+  WN+ Q V CT   N EETRD L FSC Y+S + + L QRLL+T+
Sbjct  1380  STGDRIKAWNSGQLVTCTLCNNAEETRDHLFFSCQYTSYVWEALTQRLLSTN  1431


Query #15: XLOC_002216 Query ID: lcl|Query_76549 Length: 2774

Sequences producing significant alignments:
                                                                  Max    Total Query   E   Per.
Description                                                       Score  Score cover Value Ident  Accession
unnamed protein product [Arabidopsis thaliana]                    178    178   12%   6e-46 76.92  VYS48234.1
Hypothetical protein [Arabidopsis thaliana]                       177    177   12%   9e-46 76.07  AAF63113.1
unnamed protein product [Arabidopsis thaliana]                    161    161   11%   1e-45 98.17  CAA0272879.1
unnamed protein product [Arabidopsis thaliana]                    159    159   11%   6e-42 74.29  CAA0384298.1
hypothetical protein AT1G43815 [Arabidopsis thaliana]             126    126   9%    1e-33 100.00 NP_001320857.1
Similar to gi|4263548 T6A13.8 putative En/Spm transposon prote... 86.3   149   16%   1e-28 50.56  AAF06088.1
hypothetical protein [Arabidopsis thaliana]                       86.3   143   16%   5e-27 50.56  AAG50940.1
F27J15.14 [Arabidopsis thaliana]                                  78.2   138   16%   1e-25 50.59  AAF69717.1
unknown protein [Arabidopsis thaliana]                            77.4   134   15%   3e-24 48.31  AAG51466.1
Similar to reverse transcriptase [Arabidopsis thaliana]           108    108   8%    1e-23 71.60  AAF63129.1
En/Spm transposon protein-like [Arabidopsis thaliana]             78.6   130   16%   4e-23 47.78  BAA97097.1
unnamed protein product [Arabidopsis thaliana]                    94.0   94.0  7%    2e-22 91.18  VYS48240.1
hypothetical protein AXX17_AT4G09730 [Arabidopsis thaliana]       90.5   205   24%   2e-21 49.65  OAP00305.1
transposon protein-like [Arabidopsis thaliana]                    81.6   123   12%   5e-21 50.56  BAB01786.1
F15O4.28 [Arabidopsis thaliana]                                   80.1   122   13%   8e-21 49.44  AAF79375.1
hypothetical protein AXX17_AT4G07440 [Arabidopsis thaliana]       75.1   122   11%   1e-20 65.67  OAO99482.1
putative PttA-like transposon protein [Arabidopsis thaliana]      81.3   122   12%   1e-20 50.00  AAD14504.1
T32E20.13 [Arabidopsis thaliana]                                  71.2   119   12%   6e-20 47.06  AAF79806.1
unnamed protein product [Arabidopsis thaliana]                    75.1   116   12%   7e-19 47.19  BAB09492.1
En/Spm-like transposon protein [Arabidopsis thaliana]             68.2   111   13%   2e-17 45.35  AAF18624.1
hypothetical protein AXX17_AT3G35660 [Arabidopsis thaliana]       73.9   73.9  7%    4e-15 55.38  OAP06507.1
putative protein [Arabidopsis thaliana]                           80.9   80.9  11%   6e-15 39.42  CAB86692.1
hypothetical protein AXX17_AT2G07280 [Arabidopsis thaliana]       57.4   100   13%   5e-14 37.63  OAP08596.1
hypothetical protein AXX17_AT5G29850 [Arabidopsis thaliana]       61.2   99.4  12%   6e-14 40.45  OAO90463.1
En/Spm transposon protein (mosaic-like protein) [Arabidopsis...   76.3   76.3  8%    9e-14 48.75  AAD15362.1
hypothetical protein [Arabidopsis thaliana]                       72.8   72.8  7%    1e-13 49.25  AAO86862.1
hypothetical protein At2g10070 [Arabidopsis thaliana]             72.8   72.8  7%    1e-13 49.25  AAY82259.1
hypothetical protein AXX17_AT3G33080 [Arabidopsis thaliana]       67.0   67.0  5%    8e-13 65.38  OAP06944.1
En/Spm-like transposon protein [Arabidopsis thaliana]             64.7   64.7  5%    1e-11 63.46  AAD28691.1
putative PttA-like transposon protein [Arabidopsis thaliana]      68.2   68.2  8%    4e-11 46.25  AAD28675.1
putative protein [Arabidopsis thaliana]                           62.0   62.0  5%    6e-11 62.00  CAB86451.1
contains similarity to Petunia PTTA' (GB:AF009516) [Arabidopsi... 64.3   64.3  6%    5e-10 52.38  AAD17347.1
hypothetical protein [Arabidopsis thaliana]                       60.1   60.1  5%    8e-10 55.77  AAO73427.1
hypothetical protein [Arabidopsis thaliana]                       62.0   62.0  7%    9e-10 40.30  AAG50554.1
hypothetical protein AXX17_AT2G07250 [Arabidopsis thaliana]       42.7   85.1  9%    1e-09 50.00  OAP07941.1
En/Spm-like transposon protein [Arabidopsis thaliana]             60.1   60.1  5%    3e-09 55.77  AAC97240.1
hypothetical protein AXX17_AT4G06200 [Arabidopsis thaliana]       62.0   62.0  9%    3e-09 39.78  OAP00993.1
F9C16.29 [Arabidopsis thaliana]                                   62.0   62.0  7%    4e-09 40.30  AAF79676.1
putative protein [Arabidopsis thaliana]                           57.4   57.4  6%    4e-08 50.00  CAB86680.1


Alignments:
>unnamed protein product [Arabidopsis thaliana]
Sequence ID: VYS48234.1 Length: 697
Range 1: 581 to 697

Score:178 bits(451), Expect:6e-46,
Method:Compositional matrix adjust.,
Identities:90/117(77%), Positives:96/117(82%), Gaps:0/117(0%)

Query  2233  VLLDSTWRFLAQGMDRLEFGGSKEGLKLYSAYILGWRPSGAHLKNKESTRPRHSAGGRVM  2412
             VLLDSTWRFLAQGMDRLEFGG KEG ++YSAY+LGWRPSGAH+KNKE TRPRHSAGGRV+
Sbjct  581   VLLDSTWRFLAQGMDRLEFGGPKEGFEVYSAYVLGWRPSGAHVKNKEPTRPRHSAGGRVV  640

Query  2413  HLRKKRSCSTKPLDQDPSDVTKRKSLDFSTRPRPEWCSADFPILFFSQLGHPRIAYK  2583
             HLR KRS STKPL   PS VTK K L FSTRPR E   AD  IL +SQL  PRIAYK
Sbjct  641   HLRMKRSHSTKPLGLGPSCVTKTKPLGFSTRPRAERSFADLSILSYSQLRQPRIAYK  697


>Hypothetical protein [Arabidopsis thaliana]
Sequence ID: AAF63113.1 Length: 668
Range 1: 552 to 668

Score:177 bits(449), Expect:9e-46,
Method:Compositional matrix adjust.,
Identities:89/117(76%), Positives:96/117(82%), Gaps:0/117(0%)

Query  2233  VLLDSTWRFLAQGMDRLEFGGSKEGLKLYSAYILGWRPSGAHLKNKESTRPRHSAGGRVM  2412
             VLLDSTWRFLAQGMDRLEFGG KEG ++YSAY+LGWRPSGAH+KNKE TRPRHSAGGRV+
Sbjct  552   VLLDSTWRFLAQGMDRLEFGGPKEGFEVYSAYVLGWRPSGAHVKNKEPTRPRHSAGGRVV  611

Query  2413  HLRKKRSCSTKPLDQDPSDVTKRKSLDFSTRPRPEWCSADFPILFFSQLGHPRIAYK  2583
             HLR KRS STK L   PS +TK K L FSTRPR E   AD  IL +SQLG PRIAYK
Sbjct  612   HLRMKRSHSTKLLGLGPSCMTKTKPLGFSTRPRAERSCADLSILSYSQLGQPRIAYK  668


>unnamed protein product [Arabidopsis thaliana]
Sequence ID: CAA0272879.1 Length: 109
Range 1: 1 to 109

Score:161 bits(407), Expect:1e-45,
Method:Compositional matrix adjust.,
Identities:107/109(98%), Positives:108/109(99%), Gaps:0/109(0%)

Query  541  MWRICSSSSIHVETTSISGSTGSVQSVSCDSSTGSVQPVSNLEstmgsvksstssLCITS  720
            MWRICSSSSIHVETTSI GSTGSVQSVSCDSSTGSVQPVSNLESTMGSVKSSTSSLCITS
Sbjct  1    MWRICSSSSIHVETTSICGSTGSVQSVSCDSSTGSVQPVSNLESTMGSVKSSTSSLCITS  60

Query  721  RLQAQNSRaeeeedeeaeSRKMLKGIQLSHKMCLLLYMSCFSCQVVRNI  867
            +LQAQNSRAEEEEDEEAESRKMLKGIQLSHKMCLLLYMSCFSCQVVRNI
Sbjct  61   QLQAQNSRAEEEEDEEAESRKMLKGIQLSHKMCLLLYMSCFSCQVVRNI  109


>unnamed protein product [Arabidopsis thaliana]
Sequence ID: CAA0384298.1 Length: 384
Range 1: 3 to 107

Score:159 bits(403), Expect:6e-42,
Method:Compositional matrix adjust.,
Identities:78/105(74%), Positives:85/105(80%), Gaps:0/105(0%)

Query  2230  QVLLDSTWRFLAQGMDRLEFGGSKEGLKLYSAYILGWRPSGAHLKNKESTRPRHSAGGRV  2409
             +VLLDSTWRFLAQGMDRLEFGG KEG ++YSAY+L WRPSGAH+KNKE TRPRHSAGGRV
Sbjct  3     KVLLDSTWRFLAQGMDRLEFGGPKEGFEVYSAYVLDWRPSGAHVKNKEPTRPRHSAGGRV  62

Query  2410  MHLRKKRSCSTKPLDQDPSDVTKRKSLDFSTRPRPEWCSADFPIL  2544
             +HLR KRS STKPL  +PS VTK K L FSTRPR E   AD
Sbjct  63    VHLRMKRSYSTKPLGLEPSCVTKTKPLGFSTRPRAERSCADLSFF  107


>hypothetical protein AT1G43815 [Arabidopsis thaliana]
Sequence ID: NP_001320857.1 Length: 91
>hypothetical protein AT1G43815 [Arabidopsis thaliana]
Sequence ID: NP_001320858.1 Length: 91 >hypothetical protein AT1G43815 [Arabidopsis thaliana]
Sequence ID: ANM58423.1 Length: 91 >hypothetical protein AT1G43815 [Arabidopsis thaliana]
Sequence ID: ANM58424.1 Length: 91
Range 1: 1 to 90

Score:126 bits(316), Expect:1e-33,
Method:Compositional matrix adjust.,
Identities:90/90(100%), Positives:90/90(100%), Gaps:0/90(0%)

Query  541  MWRICSSSSIHVETTSISGSTGSVQSVSCDSSTGSVQPVSNLEstmgsvksstssLCITS  720
            MWRICSSSSIHVETTSISGSTGSVQSVSCDSSTGSVQPVSNLESTMGSVKSSTSSLCITS
Sbjct  1    MWRICSSSSIHVETTSISGSTGSVQSVSCDSSTGSVQPVSNLESTMGSVKSSTSSLCITS  60

Query  721  RLQAQNSRaeeeedeeaeSRKMLKGIQLSH  810
            RLQAQNSRAEEEEDEEAESRKMLKGIQLSH
Sbjct  61   RLQAQNSRAEEEEDEEAESRKMLKGIQLSH  90


>Similar to gi|4263548 T6A13.8 putative En/Spm transposon protein homolog (mosaic protein) from Arabidopsis thaliana chromosome II sequence gb|AC006250 [Arabidopsis thaliana]
Sequence ID: AAF06088.1 Length: 497
Range 1: 196 to 282

Score:86.3 bits(212), Expect:1e-28,
Method:Compositional matrix adjust.,
Identities:45/89(51%), Positives:56/89(62%), Gaps:5/89(5%)

Query  1007  KRHIYFLEFAT---WHPSLTGVIEDKFYKIC*LRMKDMVSKACSKRARDQPPWIEKTFWK  1177
             KR   FLEFA    W P +TG ++  F +I   R+KDMVS A  +R R+QPPWI +T W
Sbjct  196   KRERLFLEFAKTHHWDPLITGTVQYYFNEIVKRRLKDMVSTA--RRTREQPPWIGETLWG  253

Query  1178  EMCEYWDI*EAIANSSTTSTARMSDRNVI  1264
              MC+YWD  EA   S T S AR+SDRN +
Sbjct  254   TMCDYWDTEEAQKRSKTYSKARLSDRNGL  282


Range 2: 130 to 196

Score:62.8 bits(151), Expect:1e-28,
Method:Compositional matrix adjust.,
Identities:28/67(42%), Positives:42/67(62%), Gaps:1/67(1%)

Query  813   DVLAPLHELLLMPSREKYMTVISPTFELETLWFGHDNGN-LIRKIIKIFTNTFDGSFYSW  989
             D    L   L +P+RE++  V+SP  + +T WF  D G+ L+RKI +IFT  F+  +Y+W
Sbjct  130   DSAMALQSTLSVPNREQWCCVLSPILKPKTEWFTRDRGSKLVRKITRIFTQKFNAPYYNW  189

Query  990   RCVPTEK  1010
              CVP +K
Sbjct  190   SCVPLDK  196


>hypothetical protein [Arabidopsis thaliana]
Sequence ID: AAG50940.1 Length: 438
Range 1: 208 to 294

Score:86.3 bits(212), Expect:5e-27,
Method:Compositional matrix adjust.,
Identities:45/89(51%), Positives:56/89(62%), Gaps:5/89(5%)

Query  1007  KRHIYFLEFAT---WHPSLTGVIEDKFYKIC*LRMKDMVSKACSKRARDQPPWIEKTFWK  1177
             KR   FLEFA    W P +TG ++  F +I   R+KDMVS A  +R R+QPPWI +T W
Sbjct  208   KRERLFLEFAKTHHWDPLITGTVQYYFNEIVKRRLKDMVSTA--RRTREQPPWIGETLWG  265

Query  1178  EMCEYWDI*EAIANSSTTSTARMSDRNVI  1264
              MC+YWD  EA   S T S AR+SDRN +
Sbjct  266   TMCDYWDTEEAQKRSKTYSKARLSDRNGL  294


Range 2: 142 to 208

Score:57.4 bits(137), Expect:5e-27,
Method:Compositional matrix adjust.,
Identities:27/67(40%), Positives:41/67(61%), Gaps:1/67(1%)

Query  813   DVLAPLHELLLMPSREKYMTVISPTFELETLWFGHDNGN-LIRKIIKIFTNTFDGSFYSW  989
             D    L   L +P+RE++  V+SP  + +T WF  D G+ L+RKI +IFT  F+  +Y+W
Sbjct  142   DSALALQSTLSVPNREQWCCVLSPIPKPKTEWFTRDRGSKLVRKITQIFTQKFNAPYYNW  201

Query  990   RCVPTEK  1010
               VP +K
Sbjct  202   SSVPLDK  208


>F27J15.14 [Arabidopsis thaliana]
Sequence ID: AAF69717.1 Length: 1526
Range 1: 1229 to 1311

Score:78.2 bits(191), Expect:1e-25,
Method:Compositional matrix adjust.,
Identities:43/85(51%), Positives:51/85(60%), Gaps:5/85(5%)

Query  1019  YFLEFAT---WHPSLTGVIEDKFYKIC*LRMKDMVSKACSKRARDQPPWIEKTFWKEMCE  1189
             YFLEFA    W P +TG ++  F +IC  RMK MVS    + +R +P WI KT WKEM
Sbjct  1229  YFLEFAKTHHWDPLITGTVQFYFNEICLRRMKGMVSTV--RTSRKKPKWIGKTLWKEMTA  1286

Query  1190  YWDI*EAIANSSTTSTARMSDRNVI  1264
             YWD  EA   S   S ARMSDRN +
Sbjct  1287  YWDTEEAQERSQIYSNARMSDRNGL  1311


Range 2: 1159 to 1225

Score:60.5 bits(145), Expect:1e-25,
Method:Compositional matrix adjust.,
Identities:28/67(42%), Positives:41/67(61%), Gaps:1/67(1%)

Query  813   DVLAPLHELLLMPSREKY-MTVISPTFELETLWFGHDNGNLIRKIIKIFTNTFDGSFYSW  989
             D L  L++LL++P R+KY +  I+  F     +    N  L+R I ++FTN FDG +YSW
Sbjct  1159  DTLQSLNDLLMLPERDKYFLPYITACFSCVWRFTRDTNSRLVRNITRVFTNKFDGPYYSW  1218

Query  990   RCVPTEK  1010
              CVP E+
Sbjct  1219  TCVPQER  1225


>unknown protein [Arabidopsis thaliana]
Sequence ID: AAG51466.1 Length: 477
Range 1: 259 to 345

Score:77.4 bits(189), Expect:3e-24,
Method:Compositional matrix adjust.,
Identities:43/89(48%), Positives:53/89(59%), Gaps:5/89(5%)

Query  1007  KRHIYFLEFAT---WHPSLTGVIEDKFYKIC*LRMKDMVSKACSKRARDQPPWIEKTFWK  1177
             KR   FLEFA    W P + G ++  F +I   R+KD+VS A  +  R+QPPWI +T W
Sbjct  259   KRERLFLEFAKTHHWDPLIIGTVQYYFNEIVKRRLKDIVSTA--RTTREQPPWIGETLWG  316

Query  1178  EMCEYWDI*EAIANSSTTSTARMSDRNVI  1264
              MC YWD  EA   S T S AR+SDRN I
Sbjct  317   TMCAYWDKEEAQKRSRTYSKARLSDRNGI  345


Range 2: 203 to 259

Score:57.0 bits(136), Expect:3e-24,
Method:Compositional matrix adjust.,
Identities:27/57(47%), Positives:35/57(61%), Gaps:1/57(1%)

Query  843   LMPSREKYMTVISPTFELETLWFGHDNGN-LIRKIIKIFTNTFDGSFYSWRCVPTEK  1010
             L+P RE +  V+SP    +T WF  D G+ L+RKI +IF   FD  FY+W CVP  K
Sbjct  203   LVPGREAWTCVLSPIPRPKTEWFTRDRGSRLVRKITRIFLQKFDAPFYNWSCVPIAK  259


>Similar to reverse transcriptase [Arabidopsis thaliana]
Sequence ID: AAF63129.1 Length: 602
Range 1: 522 to 602

Score:108 bits(270), Expect:1e-23,
Method:Compositional matrix adjust.,
Identities:58/81(72%), Positives:62/81(76%), Gaps:0/81(0%)

Query  2341  RPSGAHLKNKESTRPRHSAGGRVMHLRKKRSCSTKPLDQDPSDVTKRKSLDFSTRPRPEW  2520
             RPSGAH+KNKE TRPRHSAGGRV+HLR KRS STK L   PS +TK K L FSTRPR E
Sbjct  522   RPSGAHVKNKEPTRPRHSAGGRVVHLRMKRSHSTKLLGLGPSCMTKTKPLGFSTRPRAER  581

Query  2521  CSADFPILFFSQLGHPRIAYK  2583
               AD  IL +SQLG PRIAYK
Sbjct  582   SCADLSILSYSQLGQPRIAYK  602


>En/Spm transposon protein-like [Arabidopsis thaliana]
Sequence ID: BAA97097.1 Length: 504
Range 1: 203 to 290

Score:78.6 bits(192), Expect:4e-23,
Method:Compositional matrix adjust.,
Identities:43/90(48%), Positives:52/90(57%), Gaps:5/90(5%)

Query  1007  KRHIYFLEFAT---WHPSLTGVIEDKFYKIC*LRMKDMVSKACSKRARDQPPWIEKTFWK  1177
             KR   FLEFA    W P +TG ++  F +I   R+KDMVS A  +  R+QPPWI +T W
Sbjct  203   KRERLFLEFAKTHHWDPLITGTVQYYFNEIVKRRLKDMVSTA--RTTREQPPWIGETMWG  260

Query  1178  EMCEYWDI*EAIANSSTTSTARMSDRNVIC  1267
              MC YWD   A   S T S A +SDRN I
Sbjct  261   TMCAYWDTEAAQKRSQTYSKAHLSDRNGIS  290


Range 2: 132 to 199

Score:52.4 bits(124), Expect:4e-23,
Method:Compositional matrix adjust.,
Identities:27/68(40%), Positives:39/68(57%), Gaps:1/68(1%)

Query  798  STLP*DVLAPLHELLLMPSREKYMTVISPTFELETLWFGHDNGN-LIRKIIKIFTNTFDG  974
            S L  D +  L+++L +P RE +  V+S     +T  F  D G+ L+RKI +IF   FD
Sbjct  132  SELQEDSVVALNDILFVPGREAWCCVLSSIPRPKTKRFTRDRGSRLVRKITRIFLQKFDA  191

Query  975  SFYSWRCV  998
             FY+W CV
Sbjct  192  PFYNWSCV  199


>unnamed protein product [Arabidopsis thaliana]
Sequence ID: VYS48240.1 Length: 80
Range 1: 1 to 68

Score:94.0 bits(232), Expect:2e-22,
Method:Compositional matrix adjust.,
Identities:62/68(91%), Positives:64/68(94%), Gaps:0/68(0%)

Query  541  MWRICSSSSIHVETTSISGSTGSVQSVSCDSSTGSVQPVSNLEstmgsvksstssLCITS  720
            MWRICSSSSIHVETTSISGSTGSVQSVSCDSSTGSVQPVSNLESTMGSVKSSTSSLCITS
Sbjct  1    MWRICSSSSIHVETTSISGSTGSVQSVSCDSSTGSVQPVSNLESTMGSVKSSTSSLCITS  60

Query  721  RLQAQNSR  744
            +L+    R
Sbjct  61   QLKTHVQR  68


>hypothetical protein AXX17_AT4G09730 [Arabidopsis thaliana]
Sequence ID: OAP00305.1 Length: 324
Range 1: 41 to 170

Score:90.5 bits(223), Expect:2e-21,
Method:Compositional matrix adjust.,
Identities:70/141(50%), Positives:85/141(60%), Gaps:11/141(7%)

Query  308  LNRRPSGLPS*YEFMSANCQAPLQDYDRNSIAQPPTGPSLKDYPPSTHLFQSCEgsprgs  487
            LNR P GLPS Y+F   + Q PLQD DRN IAQ PTG +++DYPP T LFQS E    GS
Sbjct  41   LNRTPRGLPSQYDFRPPDRQPPLQDPDRNPIAQAPTGATVRDYPPPTQLFQSGE----GS  96

Query  488  vsprasgsprgsgsTQPQCGGSVHRLASTSKQPPSAVQREAFNQSPATVQREVSNQSPIL  667
                 S   R SGSTQP+ GGSVHRL   S Q  + V      +S A +QRE   Q+P++
Sbjct  97   PCGSGSIPFRASGSTQPRFGGSVHRLPCRSNQTTAPV------KSSAPIQREAPIQTPVV  150

Query  668  NQQWEASNPPPRASVSHHGSK  730
            +QQ    +P P  S SHH S+
Sbjct  151  DQQRPLPHPRPSVS-SHHSSQ  170


Range 2: 20 to 35

Score:34.3 bits(77), Expect:2e-21,
Method:Compositional matrix adjust.,
Identities:14/16(88%), Positives:15/16(93%), Gaps:0/16(0%)

Query  243  KAREEDEGDRRFPQID  290
            KAREE EGDRRFPQI+
Sbjct  20   KAREEKEGDRRFPQIE  35


Range 3: 195 to 261

Score:80.5 bits(197), Expect:2e-15,
Method:Compositional matrix adjust.,
Identities:41/67(61%), Positives:49/67(73%), Gaps:0/67(0%)

Query  1251  TVMLSVTNERGVYLRIRSLGRYINKKRKYIQNSSSFTSLQSQLEDANRKIEEEATLQARR  1430
             T+  ++TN+RG Y  + SL  YIN KRKY   SSSFT+LQSQLE AN +IEE+ATLQA R
Sbjct  195   TLSSTITNDRGEYYGVGSLRSYINGKRKYQGTSSSFTTLQSQLEKANHEIEEQATLQAHR  254

Query  1431  EVETLQV  1451
             E E L V
Sbjct  255   EAEALWV  261


>transposon protein-like [Arabidopsis thaliana]
Sequence ID: BAB01786.1 Length: 269
Range 1: 47 to 133

Score:81.6 bits(200), Expect:5e-21,
Method:Compositional matrix adjust.,
Identities:45/89(51%), Positives:53/89(59%), Gaps:5/89(5%)

Query  1007  KRHIYFLEFAT---WHPSLTGVIEDKFYKIC*LRMKDMVSKACSKRARDQPPWIEKTFWK  1177
             KR   FLEFA    W P LTG ++  F +I   R+KDMVS A  +  R+QPPWI +T W
Sbjct  47    KRERLFLEFAKTHHWDPLLTGTVQYYFNEIVKRRLKDMVSTA--RTTREQPPWIGETLWG  104

Query  1178  EMCEYWDI*EAIANSSTTSTARMSDRNVI  1264
              MC YWD   A   S T S AR+SDRN I
Sbjct  105   TMCAYWDTEAAQKRSRTYSKARLSDRNGI  133


Range 2: 13 to 47

Score:42.0 bits(97), Expect:5e-21,
Method:Compositional matrix adjust.,
Identities:18/35(51%), Positives:23/35(65%), Gaps:1/35(2%)

Query  909   FGHDNGN-LIRKIIKIFTNTFDGSFYSWRCVPTEK  1010
             F  D G+ L+RKI +IF   FD  FY+W CVP +K
Sbjct  13    FTRDRGSRLVRKITRIFLQKFDAPFYNWSCVPVDK  47


>F15O4.28 [Arabidopsis thaliana]
Sequence ID: AAF79375.1 Length: 570
Range 1: 269 to 355

Score:80.1 bits(196), Expect:8e-21,
Method:Compositional matrix adjust.,
Identities:44/89(49%), Positives:53/89(59%), Gaps:5/89(5%)

Query  1007  KRHIYFLEFAT---WHPSLTGVIEDKFYKIC*LRMKDMVSKACSKRARDQPPWIEKTFWK  1177
             KR   FLEFA    W P +TG ++  F +I   R+KDMVS A  +  R+QPPWI +T W
Sbjct  269   KRERLFLEFAKTHHWDPLITGTVQYYFNEIVKRRLKDMVSTA--RTTREQPPWIGETLWG  326

Query  1178  EMCEYWDI*EAIANSSTTSTARMSDRNVI  1264
              MC YWD   A   S T S AR+SDRN I
Sbjct  327   TMCAYWDTEAAQKRSRTYSKARLSDRNGI  355


Range 2: 226 to 269

Score:42.7 bits(99), Expect:8e-21,
Method:Compositional matrix adjust.,
Identities:20/44(45%), Positives:28/44(63%), Gaps:2/44(4%)

Query  885   TFELETLW-FGHDNGN-LIRKIIKIFTNTFDGSFYSWRCVPTEK  1010
             +F +  +W F  D G+ L+RKI +IF   FD  FY+W CVP +K
Sbjct  226   SFFVACVWRFTRDRGSRLVRKITRIFLQKFDAPFYNWSCVPVDK  269


>hypothetical protein AXX17_AT4G07440 [Arabidopsis thaliana]
Sequence ID: OAO99482.1 Length: 318
Range 1: 59 to 120

Score:75.1 bits(183), Expect:1e-20,
Method:Compositional matrix adjust.,
Identities:44/67(66%), Positives:48/67(71%), Gaps:5/67(7%)

Query  530  TQPQCGGSVHRLASTSKQPPSAVQREAFNQSPATVQREVSNQSPILNQQWEASNPPPRAS  709
            TQP+  GS+H LAST KQ P AVQ EA NQSPA VQRE SN  P+L    EASNPPPR
Sbjct  59   TQPRSRGSIHHLASTLKQSPEAVQPEASNQSPAAVQREASN--PLLQ---EASNPPPRVF  113

Query  710  VSHHGSK  730
            VSHH S+
Sbjct  114  VSHHSSQ  120


Range 2: 142 to 177

Score:47.0 bits(110), Expect:1e-20,
Method:Compositional matrix adjust.,
Identities:23/36(64%), Positives:27/36(75%), Gaps:0/36(0%)

Query  792  RYSTLP*DVLAPLHELLLMPSREKYMTVISPTFELE  899
            R ST+P D+LA L+ELL  PSREKY TVISP   L+
Sbjct  142  RESTIPEDLLATLYELLAQPSREKYTTVISPKLSLK  177


>putative PttA-like transposon protein [Arabidopsis thaliana]
Sequence ID: AAD14504.1 Length: 451
Range 1: 155 to 236

Score:81.3 bits(199), Expect:1e-20,
Method:Compositional matrix adjust.,
Identities:43/86(50%), Positives:54/86(62%), Gaps:4/86(4%)

Query  1007  KRHIYFLEFATWHPSLTGVIEDKFYKIC*LRMKDMVSKACSKRARDQPPWIEKTFWKEMC  1186
             KR   FLEFA   P +TG ++  F +I   R+KDMVS   ++R R+QPPWI +T W  MC
Sbjct  155   KRERLFLEFA--DPLITGTVQYYFNEIVKRRLKDMVS--TTRRTREQPPWIGETLWGTMC  210

Query  1187  EYWDI*EAIANSSTTSTARMSDRNVI  1264
              YWD  EA   S T S AR+SDRN +
Sbjct  211   AYWDTEEAHKRSKTYSKARLSDRNGL  236


Range 2: 121 to 155

Score:41.2 bits(95), Expect:1e-20,
Method:Compositional matrix adjust.,
Identities:16/35(46%), Positives:24/35(68%), Gaps:1/35(2%)

Query  909   FGHDNGN-LIRKIIKIFTNTFDGSFYSWRCVPTEK  1010
             F  D G+ L+RKI ++FT  F+  +Y+W CVP +K
Sbjct  121   FTRDRGSKLVRKITRVFTQKFNAPYYNWSCVPLDK  155


>T32E20.13 [Arabidopsis thaliana]
Sequence ID: AAF79806.1 Length: 1335
Range 1: 1004 to 1086

Score:71.2 bits(173), Expect:6e-20,
Method:Compositional matrix adjust.,
Identities:40/85(47%), Positives:51/85(60%), Gaps:5/85(5%)

Query  1019  YFLEFAT---WHPSLTGVIEDKFYKIC*LRMKDMVSKACSKRARDQPPWIEKTFWKEMCE  1189
             YF+EFA    W P +T  ++  F +IC  RMK MVS A  + ++++P WI KT WK M
Sbjct  1004  YFIEFAKTHHWDPLITWTVQFYFNEICLRRMKGMVSTA--RTSQNKPKWIGKTLWKTMTA  1061

Query  1190  YWDI*EAIANSSTTSTARMSDRNVI  1264
             YWD  EA   S   S ARMSDRN +
Sbjct  1062  YWDTEEAQERSQICSNARMSDRNGL  1086


Range 2: 971 to 1000

Score:48.5 bits(114), Expect:6e-20,
Method:Composition-based stats.,
Identities:18/30(60%), Positives:22/30(73%), Gaps:0/30(0%)

Query  921   NGNLIRKIIKIFTNTFDGSFYSWRCVPTEK  1010
             N  L+R I ++FTN FDG FYSW CVP E+
Sbjct  971   NSRLVRNITRVFTNKFDGPFYSWTCVPQER  1000


>unnamed protein product [Arabidopsis thaliana]
Sequence ID: BAB09492.1 Length: 305
Range 1: 47 to 133

Score:75.1 bits(183), Expect:7e-19,
Method:Compositional matrix adjust.,
Identities:42/89(47%), Positives:52/89(58%), Gaps:5/89(5%)

Query  1007  KRHIYFLEFAT---WHPSLTGVIEDKFYKIC*LRMKDMVSKACSKRARDQPPWIEKTFWK  1177
             KR   FL+FA    W P +T  ++  F +I   R+KDMVS A  +  R+QPPWI +T W
Sbjct  47    KRERLFLKFAKTHHWDPLITRTVQYYFNEIVKRRLKDMVSTA--RTTREQPPWIGETLWG  104

Query  1178  EMCEYWDI*EAIANSSTTSTARMSDRNVI  1264
              MC YWD   A   S T S AR+SDRN I
Sbjct  105   TMCAYWDTEAAQKRSRTYSKARLSDRNGI  133


Range 2: 13 to 47

Score:41.2 bits(95), Expect:7e-19,
Method:Compositional matrix adjust.,
Identities:17/35(49%), Positives:23/35(65%), Gaps:1/35(2%)

Query  909   FGHDNGN-LIRKIIKIFTNTFDGSFYSWRCVPTEK  1010
             F  D G+ L+RKI +IF   FD  FY+W C+P +K
Sbjct  13    FTRDRGSRLVRKITRIFLQKFDAPFYNWSCMPVDK  47


>En/Spm-like transposon protein [Arabidopsis thaliana]
Sequence ID: AAF18624.1 Length: 330
Range 1: 47 to 114

Score:68.2 bits(165), Expect:2e-17,
Method:Compositional matrix adjust.,
Identities:39/86(45%), Positives:46/86(53%), Gaps:18/86(20%)

Query  1007  KRHIYFLEFATWHPSLTGVIEDKFYKIC*LRMKDMVSKACSKRARDQPPWIEKTFWKEMC  1186
             KR   FLEFA                +C  R+KDMVS A  +R R+QPPWI +T W  MC
Sbjct  47    KRERLFLEFA----------------VCKRRLKDMVSTA--RRTREQPPWIGETLWAVMC  88

Query  1187  EYWDI*EAIANSSTTSTARMSDRNVI  1264
              YWD   A   S T S AR+SDRN +
Sbjct  89    AYWDTEPAQKRSRTYSKARLSDRNGL  114


Range 2: 4 to 47

Score:43.1 bits(100), Expect:2e-17,
Method:Compositional matrix adjust.,
Identities:20/44(45%), Positives:29/44(65%), Gaps:2/44(4%)

Query  885   TFELETLW-FGHDNGN-LIRKIIKIFTNTFDGSFYSWRCVPTEK  1010
             +F +  LW F  D G+ L+RKI +IFT  F+  +Y+W CVP +K
Sbjct  4     SFIVSCLWRFTRDRGSKLVRKITRIFTQKFNAPYYNWSCVPVDK  47


>hypothetical protein AXX17_AT3G35660 [Arabidopsis thaliana]
Sequence ID: OAP06507.1 Length: 107
Range 1: 6 to 70

Score:73.9 bits(180), Expect:4e-15,
Method:Compositional matrix adjust.,
Identities:36/65(55%), Positives:51/65(78%), Gaps:0/65(0%)

Query  1257  MLSVTNERGVYLRIRSLGRYINKKRKYIQNSSSFTSLQSQLEDANRKIEEEATLQARREV  1436
             M +VTN+ G Y ++R +G Y N+KRKY ++S SFTSLQ+QL++AN K++E+ TLQA RE
Sbjct  6     MSTVTNDIGEYSKVRRVGNYTNEKRKYPKSSFSFTSLQTQLKEANCKMKEQETLQAEREE  65

Query  1437  ETLQV  1451
             E LQ+
Sbjct  66    EALQI  70


>putative protein [Arabidopsis thaliana]
Sequence ID: CAB86692.1 Length: 565
Range 1: 249 to 349

Score:80.9 bits(198), Expect:6e-15,
Method:Compositional matrix adjust.,
Identities:41/104(39%), Positives:62/104(59%), Gaps:3/104(2%)

Query  953   IYKHV*WVILQLEMCTY*KRHIYFLEFATWHPSLTGVIEDKFYKIC*LRMKDMVSKACSK  1132
             ++K++    +   +C + K   YFL    W P +TG+++  F ++C  R+KDMVS   ++
Sbjct  249   LHKNLMLPTITGHVCQWIKERGYFLSL-RWDPLITGIVQYHFNQVCKRRLKDMVS--TTR  305

Query  1133  RARDQPPWIEKTFWKEMCEYWDI*EAIANSSTTSTARMSDRNVI  1264
             R R+QPPWI +T W  MC YWD  +A   S T S AR+SD N +
Sbjct  306   RTREQPPWIGETLWAVMCAYWDTEQAQKWSQTYSKARLSDHNGL  349


>hypothetical protein AXX17_AT2G07280 [Arabidopsis thaliana]
Sequence ID: OAP08596.1 Length: 483
Range 1: 184 to 274

Score:57.4 bits(137), Expect:5e-14,
Method:Compositional matrix adjust.,
Identities:35/93(38%), Positives:51/93(54%), Gaps:6/93(6%)

Query  1019  YFLEFAT---WHPSLTGVIEDKFYKIC*LRMKDMVSKACSKRARDQPPWIEKTFWKEMCE  1189
             YF  FA    W   +T ++++    I   RMK +VS+A  K++ +QPPWI  T WK+M
Sbjct  184   YFRNFARKYYWDQGVTELVKEGSLVIAKKRMKGIVSQA--KKSGEQPPWIGDTLWKQMWV  241

Query  1190  YWDI*EAIANSSTTSTARMSDRNVI-CHKRERG  1285
             +W+  +AI  S T S  R S+R  +  HK   G
Sbjct  242   HWNTEDAIQKSETASNCRNSNRGGLGVHKHLAG  274


Range 2: 142 to 177

Score:42.7 bits(99), Expect:5e-14,
Method:Compositional matrix adjust.,
Identities:18/36(50%), Positives:21/36(58%), Gaps:0/36(0%)

Query  894   LETLWFGHDNGNLIRKIIKIFTNTFDGSFYSWRCVP  1001
             +ETLWF   NG L R I  IF   FDG + SW+  P
Sbjct  142   VETLWFNRHNGKLSRVISGIFRRKFDGPYISWKVTP  177


>hypothetical protein AXX17_AT5G29850 [Arabidopsis thaliana]
Sequence ID: OAO90463.1 Length: 535
Range 1: 265 to 351

Score:61.2 bits(147), Expect:6e-14,
Method:Compositional matrix adjust.,
Identities:36/89(40%), Positives:51/89(57%), Gaps:6/89(6%)

Query  1019  YFLEFAT---WHPSLTGVIEDKFYKIC*LRMKDMVSKACSKRARDQPPWIEKTFWKEMCE  1189
             YF  FA    W   +T ++++ F  I   RMK +VS+A  K++ +QPPWI  T WK+M
Sbjct  265   YFRNFARKYYWDEGITELVKEGFLVIAKKRMKGIVSQA--KKSGEQPPWIRDTLWKQMWV  322

Query  1190  YWDI*EAIANSSTTSTARMSDRNVI-CHK  1273
             +W+  +A   S TTS  R SDR  +  HK
Sbjct  323   HWNTEDAQLKSETTSNCRNSDRGGLGVHK  351


Range 2: 227 to 258

Score:38.1 bits(87), Expect:6e-14,
Method:Compositional matrix adjust.,
Identities:15/32(47%), Positives:18/32(56%), Gaps:0/32(0%)

Query  906   WFGHDNGNLIRKIIKIFTNTFDGSFYSWRCVP  1001
             WF   NG L R I  IF   FDG ++SW+  P
Sbjct  227   WFNRHNGKLSRVISSIFRRKFDGPYFSWKVTP  258


>En/Spm transposon protein (mosaic-like protein) [Arabidopsis thaliana]
Sequence ID: AAD15362.1 Length: 400
Range 1: 110 to 185

Score:76.3 bits(186), Expect:9e-14,
Method:Compositional matrix adjust.,
Identities:39/80(49%), Positives:50/80(62%), Gaps:4/80(5%)

Query  1025  LEFATWHPSLTGVIEDKFYKIC*LRMKDMVSKACSKRARDQPPWIEKTFWKEMCEYWDI*  1204
             L+F    P  TG ++  F +I   R+KDMVS A  +R R+QPPWI +T W  MC+YWD
Sbjct  110   LQFPNLKP--TGTVQYYFNEIVKRRLKDMVSTA--RRTREQPPWIGETLWGTMCDYWDTE  165

Query  1205  EAIANSSTTSTARMSDRNVI  1264
             EA   S T S AR+SDRN +
Sbjct  166   EAQKRSKTYSKARLSDRNGL  185


>hypothetical protein [Arabidopsis thaliana]
Sequence ID: AAO86862.1 Length: 208
>hypothetical protein At2g10070 [Arabidopsis thaliana]
Sequence ID: AAV63871.1 Length: 208
Range 1: 119 to 185

Score:72.8 bits(177), Expect:1e-13,
Method:Compositional matrix adjust.,
Identities:33/67(49%), Positives:46/67(68%), Gaps:1/67(1%)

Query  813   DVLAPLHELLLMPSREKYMTVISPTFELETLWFGHD-NGNLIRKIIKIFTNTFDGSFYSW  989
             D L  L+++L +P RE + TV+SPT   +T WF  D N +L+RK  K++TN FDG FYSW
Sbjct  119   DSLRALNDVLQVPGREAWTTVLSPTLMPKTTWFTRDTNSSLVRKTTKVWTNKFDGLFYSW  178

Query  990   RCVPTEK  1010
              CVP ++
Sbjct  179   SCVPQDR  185


>hypothetical protein At2g10070 [Arabidopsis thaliana]
Sequence ID: AAY82259.1 Length: 208
Range 1: 119 to 185

Score:72.8 bits(177), Expect:1e-13,
Method:Compositional matrix adjust.,
Identities:33/67(49%), Positives:46/67(68%), Gaps:1/67(1%)

Query  813   DVLAPLHELLLMPSREKYMTVISPTFELETLWFGHD-NGNLIRKIIKIFTNTFDGSFYSW  989
             D L  L+++L +P RE + TV+SPT   +T WF  D N +L+RK  K++TN FDG FYSW
Sbjct  119   DSLRALNDVLQVPGREAWTTVLSPTLMPKTTWFTRDTNSSLVRKTTKVWTNKFDGLFYSW  178

Query  990   RCVPTEK  1010
              CVP ++
Sbjct  179   SCVPQDR  185


>hypothetical protein AXX17_AT3G33080 [Arabidopsis thaliana]
Sequence ID: OAP06944.1 Length: 95
Range 1: 37 to 88

Score:67.0 bits(162), Expect:8e-13,
Method:Compositional matrix adjust.,
Identities:34/52(65%), Positives:40/52(76%), Gaps:0/52(0%)

Query  1296  IRSLGRYINKKRKYIQNSSSFTSLQSQLEDANRKIEEEATLQARREVETLQV  1451
              R LG Y N+KRKY   SSSFTSLQ+QLE+AN KI+E+ TLQA RE E LQ+
Sbjct  37    FRRLGCYTNQKRKYSGGSSSFTSLQTQLEEANYKIKEQKTLQAEREAEALQI  88


>En/Spm-like transposon protein [Arabidopsis thaliana]
Sequence ID: AAD28691.1 Length: 129
Range 1: 14 to 65

Score:64.7 bits(156), Expect:1e-11,
Method:Compositional matrix adjust.,
Identities:33/52(63%), Positives:40/52(76%), Gaps:0/52(0%)

Query  1302  SLGRYINKKRKYIQNSSSFTSLQSQLEDANRKIEEEATLQARREVETLQVRT  1457
             S    IN KRKY ++S +FTSLQS LE+ANRKIEE+ATLQA RE E L+V+
Sbjct  14    SFREVINGKRKYPRSSYAFTSLQSHLEEANRKIEEQATLQAHREAEALRVKA  65


>putative PttA-like transposon protein [Arabidopsis thaliana]
Sequence ID: AAD28675.1 Length: 420
Range 1: 158 to 233

Score:68.2 bits(165), Expect:4e-11,
Method:Compositional matrix adjust.,
Identities:37/80(46%), Positives:46/80(57%), Gaps:4/80(5%)

Query  1025  LEFATWHPSLTGVIEDKFYKIC*LRMKDMVSKACSKRARDQPPWIEKTFWKEMCEYWDI*  1204
             L F    PS  G ++  F +I   R+KDMVS    +R R+QP WI +T W  MC YWD
Sbjct  158   LRFPNQKPS--GTVQYYFNEIVKRRLKDMVS--TDRRTREQPSWIGETLWGTMCAYWDTE  213

Query  1205  EAIANSSTTSTARMSDRNVI  1264
             EA   S T S AR+SDRN +
Sbjct  214   EAKKRSKTYSKARLSDRNGL  233


>putative protein [Arabidopsis thaliana]
Sequence ID: CAB86451.1 Length: 112
Range 1: 27 to 76

Score:62.0 bits(149), Expect:6e-11,
Method:Compositional matrix adjust.,
Identities:31/50(62%), Positives:41/50(82%), Gaps:0/50(0%)

Query  1302  SLGRYINKKRKYIQNSSSFTSLQSQLEDANRKIEEEATLQARREVETLQV  1451
             S G Y N+KRKY ++SSSFTSLQ+QL++AN K++E+ TLQA RE E LQ+
Sbjct  27    SDGNYTNEKRKYPKSSSSFTSLQTQLKEANCKMKEQETLQAEREEEALQI  76


>contains similarity to Petunia PTTA' (GB:AF009516) [Arabidopsis thaliana]
Sequence ID: AAD17347.1 Length: 357
>putative transposon protein [Arabidopsis thaliana]
Sequence ID: CAB80814.1 Length: 357
Range 1: 162 to 222

Score:64.3 bits(155), Expect:5e-10,
Method:Compositional matrix adjust.,
Identities:33/63(52%), Positives:39/63(61%), Gaps:2/63(3%)

Query  1076  FYKIC*LRMKDMVSKACSKRARDQPPWIEKTFWKEMCEYWDI*EAIANSSTTSTARMSDR  1255
             F +I   R+KDMVS A  +  R+QPPWI +T W  MC YWD   A   S T S AR+SDR
Sbjct  162   FNEIVKRRLKDMVSTA--RTTREQPPWIGETLWGTMCAYWDTEAAQKRSRTYSKARLSDR  219

Query  1256  NVI  1264
             N I
Sbjct  220   NGI  222


>hypothetical protein [Arabidopsis thaliana]
Sequence ID: AAO73427.1 Length: 151
Range 1: 1 to 50

Score:60.1 bits(144), Expect:8e-10,
Method:Compositional matrix adjust.,
Identities:29/52(56%), Positives:35/52(67%), Gaps:2/52(3%)

Query  1109  MVSKACSKRARDQPPWIEKTFWKEMCEYWDI*EAIANSSTTSTARMSDRNVI  1264
             MVS A  +R R+QPPWI +T W  MC+YWD  EA   S T S AR+SDRN +
Sbjct  1     MVSTA--RRTREQPPWIGETLWGTMCDYWDTEEAQKRSKTYSKARLSDRNGL  50


>hypothetical protein [Arabidopsis thaliana]
Sequence ID: AAG50554.1 Length: 251
Range 1: 140 to 206

Score:62.0 bits(149), Expect:9e-10,
Method:Compositional matrix adjust.,
Identities:27/67(40%), Positives:42/67(62%), Gaps:1/67(1%)

Query  813   DVLAPLHELLLMPSREKYMTVISPTFELETLWFGHDNGN-LIRKIIKIFTNTFDGSFYSW  989
             D    L +++ +P+RE++  V+SP    +T WF  D G+ L+RKI +IFT  F+  +Y+W
Sbjct  140   DSAVALQDIMSVPNREQWTCVLSPIPRPKTEWFTRDRGSKLVRKITRIFTQKFNAPYYNW  199

Query  990   RCVPTEK  1010
              CVP  K
Sbjct  200   SCVPLAK  206


>hypothetical protein AXX17_AT2G07250 [Arabidopsis thaliana]
Sequence ID: OAP07941.1 Length: 407
Range 1: 142 to 177

Score:42.7 bits(99), Expect:1e-09,
Method:Compositional matrix adjust.,
Identities:18/36(50%), Positives:21/36(58%), Gaps:0/36(0%)

Query  894   LETLWFGHDNGNLIRKIIKIFTNTFDGSFYSWRCVP  1001
             +ETLWF   NG L R I  IF   FDG + SW+  P
Sbjct  142   VETLWFNRHNGKLSRVISGIFRRKFDGPYISWKVTP  177


Range 2: 184 to 239

Score:42.4 bits(98), Expect:1e-09,
Method:Compositional matrix adjust.,
Identities:23/58(40%), Positives:34/58(58%), Gaps:5/58(8%)

Query  1019  YFLEFAT---WHPSLTGVIEDKFYKIC*LRMKDMVSKACSKRARDQPPWIEKTFWKEM  1183
             YF  FA    W   +T ++++    I   RMK +VS+A  K++ +QPPWI  T WK+M
Sbjct  184   YFRNFARKYYWDQGVTELVKEGSLVIAKKRMKGIVSQA--KKSGEQPPWIGDTLWKQM  239


>En/Spm-like transposon protein [Arabidopsis thaliana]
Sequence ID: AAC97240.1 Length: 224
Range 1: 1 to 50

Score:60.1 bits(144), Expect:3e-09,
Method:Compositional matrix adjust.,
Identities:29/52(56%), Positives:35/52(67%), Gaps:2/52(3%)

Query  1109  MVSKACSKRARDQPPWIEKTFWKEMCEYWDI*EAIANSSTTSTARMSDRNVI  1264
             MVS A  +R R+QPPWI +T W  MC+YWD  EA   S T S AR+SDRN +
Sbjct  1     MVSTA--RRTREQPPWIGETLWGTMCDYWDTEEAQKRSKTYSKARLSDRNGL  50


>hypothetical protein AXX17_AT4G06200 [Arabidopsis thaliana]
Sequence ID: OAP00993.1 Length: 438
Range 1: 175 to 265

Score:62.0 bits(149), Expect:3e-09,
Method:Compositional matrix adjust.,
Identities:37/93(40%), Positives:52/93(55%), Gaps:6/93(6%)

Query  1019  YFLEFAT---WHPSLTGVIEDKFYKIC*LRMKDMVSKACSKRARDQPPWIEKTFWKEMCE  1189
             YF  FA    W   +T ++++ F  I   RMK +VS+A  K++ +QPPWI  T WK+M
Sbjct  175   YFRNFARKYYWDEGITELVKESFLVIAKKRMKGIVSQA--KKSGEQPPWIRDTLWKQMWV  232

Query  1190  YWDI*EAIANSSTTSTARMSDRNVI-CHKRERG  1285
             +W+  +A   S TTS  R SDR  +  HK   G
Sbjct  233   HWNTEDAQLKSETTSNCRNSDRGGLGVHKHLAG  265


>F9C16.29 [Arabidopsis thaliana]
Sequence ID: AAF79676.1 Length: 489
Range 1: 140 to 206

Score:62.0 bits(149), Expect:4e-09,
Method:Compositional matrix adjust.,
Identities:27/67(40%), Positives:42/67(62%), Gaps:1/67(1%)

Query  813   DVLAPLHELLLMPSREKYMTVISPTFELETLWFGHDNGN-LIRKIIKIFTNTFDGSFYSW  989
             D    L +++ +P+RE++  V+SP    +T WF  D G+ L+RKI +IFT  F+  +Y+W
Sbjct  140   DSAVALQDIMSVPNREQWTCVLSPIPRPKTEWFTRDRGSKLVRKITRIFTQKFNAPYYNW  199

Query  990   RCVPTEK  1010
              CVP  K
Sbjct  200   SCVPLAK  206


>putative protein [Arabidopsis thaliana]
Sequence ID: CAB86680.1 Length: 276
Range 1: 27 to 88

Score:57.4 bits(137), Expect:4e-08,
Method:Compositional matrix adjust.,
Identities:32/64(50%), Positives:40/64(62%), Gaps:3/64(4%)

Query  1076  FYKIC*LRMKDMVSKACSKRARDQPPWI-EKTFWKEMCEYWDI*EAIANSSTTSTARMSD  1252
             F  IC  RMKDM+S    + +++ P WI +   W+ MC YWD  EAIA S T S ARMSD
Sbjct  27    FEDICQRRMKDMISTV--RTSQECPKWIIDSHIWETMCAYWDTEEAIAKSLTYSKARMSD  84

Query  1253  RNVI  1264
             RN +
Sbjct  85    RNGL  88


Query #16: XLOC_002603 Query ID: lcl|Query_76550 Length: 380

No significant similarity found.


Query #17: XLOC_002981 Query ID: lcl|Query_76551 Length: 900

No significant similarity found.


Query #18: XLOC_004130 Query ID: lcl|Query_76552 Length: 643

Sequences producing significant alignments:
                                                                  Max    Total Query    E    Per.
Description                                                       Score  Score cover Value  Ident  Accession
hypothetical protein AT1G80625 [Arabidopsis thaliana]             331    331   93%   2e-116 99.00  NP_001322100.1
hypothetical protein AT3G19790 [Arabidopsis thaliana]             71.2   71.2  28%   7e-15  55.74  NP_001078190.1
unnamed protein product [Arabidopsis thaliana]                    68.9   68.9  28%   5e-14  54.10  VYS57954.1
Hypothetical protein [Arabidopsis thaliana]                       65.9   65.9  30%   8e-13  40.00  AAD25556.1
hypothetical protein [Arabidopsis thaliana]                       62.8   62.8  28%   5e-11  40.32  CAA22150.1
unnamed protein product [Arabidopsis thaliana]                    61.2   61.2  31%   1e-10  42.65  VYS50968.1
hypothetical protein AXX17_AT2G16000 [Arabidopsis thaliana]       57.4   57.4  28%   2e-10  38.71  OAP10783.1
hypothetical protein AXX17_AT5G29290 [Arabidopsis thaliana]       52.0   52.0  31%   1e-08  35.29  OAO93174.1


Alignments:
>hypothetical protein AT1G80625 [Arabidopsis thaliana]
Sequence ID: NP_001322100.1 Length: 199
>hypothetical protein AT1G80625 [Arabidopsis thaliana]
Sequence ID: ANM59765.1 Length: 199 >hypothetical protein AXX17_AT1G75480 [Arabidopsis thaliana]
Sequence ID: OAP17711.1 Length: 199 >unnamed protein product [Arabidopsis thaliana]
Sequence ID: CAA0344675.1 Length: 199 >unnamed protein product [Arabidopsis thaliana]
Sequence ID: VYS51644.1 Length: 199
Range 1: 1 to 199

Score:331 bits(848), Expect:2e-116,
Method:Compositional matrix adjust.,
Identities:199/201(99%), Positives:199/201(99%), Gaps:2/201(0%)

Query  9    MAFHLLRPKQRFDLNPFLIRIFstsssslpsqlpsNFFTKKPLVDRARDSGNVFDRFAIN  188
            MAFHLLRPKQRFDLNPFLIRIFSTSSSSLPSQLPSNFFTKKPLVDRARDSGNVFDRFAIN
Sbjct  1    MAFHLLRPKQRFDLNPFLIRIFSTSSSSLPSQLPSNFFTKKPLVDRARDSGNVFDRFAIN  60

Query  189  RYRREKPF*DEIQSEVLITeqpkeekrrrpgrpkkkqLVEKKPPLDEKKSRAASIARKTF  368
            RYRREKP  DEIQSEVLITEQPKEEKRRRPGRPKKKQLVEKKPPLDEKKSRAASIARKTF
Sbjct  61   RYRREKP--DEIQSEVLITEQPKEEKRRRPGRPKKKQLVEKKPPLDEKKSRAASIARKTF  118

Query  369  TAFGSSLQNALKNHPDYICSVTHAMDVLNGLPRVKKWLPLYRASMDHLMADVAHRQAFLT  548
            TAFGSSLQNALKNHPDYICSVTHAMDVLNGLPRVKKWLPLYRASMDHLMADVAHRQAFLT
Sbjct  119  TAFGSSLQNALKNHPDYICSVTHAMDVLNGLPRVKKWLPLYRASMDHLMADVAHRQAFLT  178

Query  549  VSDPEDMIRYLRYKTQKMKRE  611
            VSDPEDMIRYLRYKTQKMKRE
Sbjct  179  VSDPEDMIRYLRYKTQKMKRE  199


>hypothetical protein AT3G19790 [Arabidopsis thaliana]
Sequence ID: NP_001078190.1 Length: 167
>hypothetical protein AT3G19790 [Arabidopsis thaliana]
Sequence ID: NP_566647.1 Length: 167 >At3g19790 [Arabidopsis thaliana]
Sequence ID: AAS99662.1 Length: 167 >At3g19790 [Arabidopsis thaliana]
Sequence ID: AAT41800.1 Length: 167 >hypothetical protein AT3G19790 [Arabidopsis thaliana]
Sequence ID: AEE76288.1 Length: 167 >hypothetical protein AT3G19790 [Arabidopsis thaliana]
Sequence ID: AEE76289.1 Length: 167 >hypothetical protein AXX17_AT3G21050 [Arabidopsis thaliana]
Sequence ID: OAP03283.1 Length: 167
Range 1: 107 to 167

Score:71.2 bits(173), Expect:7e-15,
Method:Compositional matrix adjust.,
Identities:34/61(56%), Positives:40/61(65%), Gaps:0/61(0%)

Query  429  VTHAMDVLNGLPRVKKWLPLYRASMDHLMADVAHRQAFLTVSDPEDMIRYLRYKTQKMKR  608
            V  +M  L+ LP V+ W PLYRAS+ H  ADV HR+AFL  SDPED I YL  KT K +
Sbjct  107  VFDSMKALHDLPGVRMWSPLYRASLTHFQADVTHREAFLAFSDPEDKICYLENKTGKKRD  166

Query  609  E  611
            E
Sbjct  167  E  167


>unnamed protein product [Arabidopsis thaliana]
Sequence ID: VYS57954.1 Length: 167
Range 1: 107 to 167

Score:68.9 bits(167), Expect:5e-14,
Method:Compositional matrix adjust.,
Identities:33/61(54%), Positives:40/61(65%), Gaps:0/61(0%)

Query  429  VTHAMDVLNGLPRVKKWLPLYRASMDHLMADVAHRQAFLTVSDPEDMIRYLRYKTQKMKR  608
            V  +M  L+ LP V+ W PLYRAS+ +  ADV HR+AFL  SDPED I YL  KT K +
Sbjct  107  VFDSMKALHDLPGVRMWSPLYRASLTYFQADVTHREAFLAFSDPEDKICYLENKTGKKRD  166

Query  609  E  611
            E
Sbjct  167  E  167


>Hypothetical protein [Arabidopsis thaliana]
Sequence ID: AAD25556.1 Length: 189
Range 1: 119 to 183

Score:65.9 bits(159), Expect:8e-13,
Method:Compositional matrix adjust.,
Identities:26/65(40%), Positives:43/65(66%), Gaps:0/65(0%)

Query  399  LKNHPDYICSVTHAMDVLNGLPRVKKWLPLYRASMDHLMADVAHRQAFLTVSDPEDMIRY  578
            ++ HP++ C    AM +L+ LP ++ W PLY+A++ HL  D+A+RQ FL   D E+ + Y
Sbjct  119  IEMHPEFSCCQLRAMQILHSLPAIRVWSPLYKAAIQHLKKDIANRQTFLFYEDDENKVLY  178

Query  579  LRYKT  593
            L ++T
Sbjct  179  LEFET  183


>hypothetical protein [Arabidopsis thaliana]
Sequence ID: CAA22150.1 Length: 457
>hypothetical protein [Arabidopsis thaliana]
Sequence ID: CAB79200.1 Length: 457
Range 1: 247 to 308

Score:62.8 bits(151), Expect:5e-11,
Method:Compositional matrix adjust.,
Identities:25/62(40%), Positives:39/62(62%), Gaps:0/62(0%)

Query  408  HPDYICSVTHAMDVLNGLPRVKKWLPLYRASMDHLMADVAHRQAFLTVSDPEDMIRYLRY  587
            HP + C    AM +L+ LP ++ W PLY+A++ HL  D+ +RQ FL   D E+ + YL +
Sbjct  247  HPKFSCCQLRAMQILHSLPAIRVWSPLYKAAIQHLKKDITNRQTFLFYEDDENKVLYLEF  306

Query  588  KT  593
            +T
Sbjct  307  ET  308


>unnamed protein product [Arabidopsis thaliana]
Sequence ID: VYS50968.1 Length: 291
Range 1: 222 to 287

Score:61.2 bits(147), Expect:1e-10,
Method:Compositional matrix adjust.,
Identities:29/68(43%), Positives:44/68(64%), Gaps:2/68(2%)

Query  399  LKNHPDYICSVTHAMDVLNGLPRVKKWLPLYRASMDHLMADVAHRQAFLTVSDPEDMIRY  578
            L++  +Y  SV   + +LNGLP V+ W P Y+A+++HL+AD A RQAF+     ED I Y
Sbjct  222  LESEKEY--SVKSVIQMLNGLPGVRMWSPFYKAAVEHLVADEASRQAFVAYPRDEDKIEY  279

Query  579  LRYKTQKM  602
            L  +T++
Sbjct  280  LELRTRRF  287


>hypothetical protein AXX17_AT2G16000 [Arabidopsis thaliana]
Sequence ID: OAP10783.1 Length: 96
Range 1: 27 to 88

Score:57.4 bits(137), Expect:2e-10,
Method:Compositional matrix adjust.,
Identities:24/62(39%), Positives:38/62(61%), Gaps:0/62(0%)

Query  414  DYICSVTHAMDVLNGLPRVKKWLPLYRASMDHLMADVAHRQAFLTVSDPEDMIRYLRYKT  593
            ++  SV   ++ LNGLP V+ W P ++ S+DHL+AD A RQ F+     E   R+L + T
Sbjct  27   EHASSVKSVLETLNGLPGVRMWSPFHKTSIDHLIADEASRQGFIAFPRAEHKSRFLEFMT  86

Query  594  QK  599
            ++
Sbjct  87   RR  88


>hypothetical protein AXX17_AT5G29290 [Arabidopsis thaliana]
Sequence ID: OAO93174.1 Length: 76
Range 1: 7 to 72

Score:52.0 bits(123), Expect:1e-08,
Method:Compositional matrix adjust.,
Identities:24/68(35%), Positives:41/68(60%), Gaps:2/68(2%)

Query  399  LKNHPDYICSVTHAMDVLNGLPRVKKWLPLYRASMDHLMADVAHRQAFLTVSDPEDMIRY  578
            L++  +Y  SV   +  LNGLP V+ W P ++A++++L+ D A R  F+T    ED I +
Sbjct  7    LESENEY--SVKSVLQKLNGLPGVRMWSPFHKATVENLIVDEASRHCFITFPRAEDKIEF  64

Query  579  LRYKTQKM  602
            L  +T++
Sbjct  65   LELRTRRF  72


Query #19: XLOC_004917 Query ID: lcl|Query_76553 Length: 405

No significant similarity found.


Query #20: XLOC_004979 Query ID: lcl|Query_76554 Length: 1177

Sequences producing significant alignments:
                                                                  Max    Total Query   E   Per.
Description                                                       Score  Score cover Value Ident  Accession
hypothetical protein AXX17_AT3G51120 [Arabidopsis thaliana]       60.5   60.5  6%    1e-09 96.30  OAP06540.1
RNA-binding (RRM/RBD/RNP motifs) family protein [Arabidopsis...   60.5   60.5  6%    1e-09 96.30  NP_001030873.1
putative protein [Arabidopsis thaliana]                           60.5   60.5  6%    1e-09 96.30  CAB88054.1


Alignments:
>hypothetical protein AXX17_AT3G51120 [Arabidopsis thaliana]
Sequence ID: OAP06540.1 Length: 257
>unnamed protein product [Arabidopsis thaliana]
Sequence ID: VYS60593.1 Length: 257
Range 1: 53 to 79

Score:60.5 bits(145), Expect:1e-09,
Method:Compositional matrix adjust.,
Identities:26/27(96%), Positives:27/27(100%), Gaps:0/27(0%)

Query  316  YLSRIPPHMDHVRLRHILAQFGELGRI  236
            YLSRIPPHMDHVRLRHILAQ+GELGRI
Sbjct  53   YLSRIPPHMDHVRLRHILAQYGELGRI  79


>RNA-binding (RRM/RBD/RNP motifs) family protein [Arabidopsis thaliana]
Sequence ID: NP_001030873.1 Length: 257
>RNA-binding (RRM/RBD/RNP motifs) family protein [Arabidopsis thaliana]
Sequence ID: NP_001326046.1 Length: 257 >RNA-binding (RRM/RBD/RNP motifs) family protein [Arabidopsis thaliana]
Sequence ID: NP_001326047.1 Length: 257 >RNA-binding (RRM/RBD/RNP motifs) family protein [Arabidopsis thaliana]
Sequence ID: NP_001326048.1 Length: 257 >RNA-binding (RRM/RBD/RNP motifs) family protein [Arabidopsis thaliana]
Sequence ID: NP_191210.2 Length: 257 >At3g56510 [Arabidopsis thaliana]
Sequence ID: AAR23731.1 Length: 257 >At3g56510 [Arabidopsis thaliana]
Sequence ID: AAS68115.1 Length: 257 >RNA-binding (RRM/RBD/RNP motifs) family protein [Arabidopsis thaliana]
Sequence ID: AEE79529.1 Length: 257 >RNA-binding (RRM/RBD/RNP motifs) family protein [Arabidopsis thaliana]
Sequence ID: AEE79530.1 Length: 257 >RNA-binding (RRM/RBD/RNP motifs) family protein [Arabidopsis thaliana]
Sequence ID: ANM63992.1 Length: 257
Range 1: 53 to 79

Score:60.5 bits(145), Expect:1e-09,
Method:Compositional matrix adjust.,
Identities:26/27(96%), Positives:27/27(100%), Gaps:0/27(0%)

Query  316  YLSRIPPHMDHVRLRHILAQFGELGRI  236
            YLSRIPPHMDHVRLRHILAQ+GELGRI
Sbjct  53   YLSRIPPHMDHVRLRHILAQYGELGRI  79


>putative protein [Arabidopsis thaliana]
Sequence ID: CAB88054.1 Length: 266
Range 1: 53 to 79

Score:60.5 bits(145), Expect:1e-09,
Method:Compositional matrix adjust.,
Identities:26/27(96%), Positives:27/27(100%), Gaps:0/27(0%)

Query  316  YLSRIPPHMDHVRLRHILAQFGELGRI  236
            YLSRIPPHMDHVRLRHILAQ+GELGRI
Sbjct  53   YLSRIPPHMDHVRLRHILAQYGELGRI  79


Query #21: XLOC_005620 Query ID: lcl|Query_76555 Length: 572

No significant similarity found.


Query #22: XLOC_005739 Query ID: lcl|Query_76556 Length: 1044

No significant similarity found.


Query #23: XLOC_005813 Query ID: lcl|Query_76557 Length: 1015

Sequences producing significant alignments:
                                                                  Max    Total Query   E   Per.
Description                                                       Score  Score cover Value Ident  Accession
unnamed protein product [Arabidopsis thaliana]                    146    146   21%   9e-44 93.24  CAA0261202.1
hypothetical protein AXX17_AT1G32620 [Arabidopsis thaliana]       109    109   15%   8e-30 100.00 OAP15554.1
unnamed protein product [Arabidopsis thaliana]                    114    114   15%   3e-27 98.08  VYS47756.1


Alignments:
>unnamed protein product [Arabidopsis thaliana]
Sequence ID: CAA0261202.1 Length: 76
Range 1: 3 to 76

Score:146 bits(369), Expect:9e-44,
Method:Compositional matrix adjust.,
Identities:69/74(93%), Positives:71/74(95%), Gaps:0/74(0%)

Query  398  LHSRSTEGTKIFIGHWPIDGGKSELQWCLAIPTVELVCLDAWRRGVKRPPRVSALNHYCV  219
            L SRSTEGTKIFIGHWPIDGGKSELQWCLAIPTVELVCLDA RRGVKRPPRVSA+NHYC
Sbjct  3    LDSRSTEGTKIFIGHWPIDGGKSELQWCLAIPTVELVCLDARRRGVKRPPRVSAINHYCG  62

Query  218  NVLLYFQTIWVGHE  177
            NVLLYFQTIWVGH+
Sbjct  63   NVLLYFQTIWVGHQ  76


>hypothetical protein AXX17_AT1G32620 [Arabidopsis thaliana]
Sequence ID: OAP15554.1 Length: 52
Range 1: 1 to 52

Score:109 bits(273), Expect:8e-30,
Method:Compositional matrix adjust.,
Identities:52/52(100%), Positives:52/52(100%), Gaps:0/52(0%)

Query  550  MKMLFLKQFGGNKENYEEGGDVDKRLRWRSKLVTAREPDKTGDKSLTHSQNY  705
            MKMLFLKQFGGNKENYEEGGDVDKRLRWRSKLVTAREPDKTGDKSLTHSQNY
Sbjct  1    MKMLFLKQFGGNKENYEEGGDVDKRLRWRSKLVTAREPDKTGDKSLTHSQNY  52


>unnamed protein product [Arabidopsis thaliana]
Sequence ID: VYS47756.1 Length: 697
Range 1: 3 to 54

Score:114 bits(284), Expect:3e-27,
Method:Compositional matrix adjust.,
Identities:51/52(98%), Positives:51/52(98%), Gaps:0/52(0%)

Query  398  LHSRSTEGTKIFIGHWPIDGGKSELQWCLAIPTVELVCLDAWRRGVKRPPRV  243
            L SRSTEGTKIFIGHWPIDGGKSELQWCLAIPTVELVCLDAWRRGVKRPPRV
Sbjct  3    LDSRSTEGTKIFIGHWPIDGGKSELQWCLAIPTVELVCLDAWRRGVKRPPRV  54


Query #24: XLOC_005815 Query ID: lcl|Query_76558 Length: 764

Sequences producing significant alignments:
                                                                  Max    Total Query   E   Per.
Description                                                       Score  Score cover Value Ident  Accession
hypothetical protein AXX17_AT1G33220 [Arabidopsis thaliana]       38.9   76.2  27%   9e-08 48.94  OAP19270.1
F-box and associated interaction domains-containing protein...    38.9   76.2  27%   1e-07 48.94  NP_174521.1


Alignments:
>hypothetical protein AXX17_AT1G33220 [Arabidopsis thaliana]
Sequence ID: OAP19270.1 Length: 380
Range 1: 41 to 78

Score:38.9 bits(89), Expect:9e-08,
Method:Compositional matrix adjust.,
Identities:23/47(49%), Positives:27/47(57%), Gaps:12/47(25%)

Query  665  RAFINNQKMTFQFI---*PKIYLVSVNPKIVLI*CKSCNNVSEIVLN  534
            + FINN KMTFQF+     KIY VSVNPK+          V E+ LN
Sbjct  41   KTFINNHKMTFQFVLSTRSKIYSVSVNPKV---------EVRELTLN  78


Range 2: 15 to 43

Score:37.4 bits(85), Expect:9e-08,
Method:Compositional matrix adjust.,
Identities:15/29(52%), Positives:20/29(68%), Gaps:0/29(0%)

Query  744  IWWKSSPSYIVQFRIVCKRWNALFQDKGF  658
            I  +  P+ + +F+ VCKRWNALF DK F
Sbjct  15   ILSRVPPTSLDRFKTVCKRWNALFNDKTF  43


>F-box and associated interaction domains-containing protein [Arabidopsis thaliana]
Sequence ID: NP_174521.1 Length: 380
>F-box and associated interaction domains-containing protein [Arabidopsis thaliana]
Sequence ID: AEE31486.1 Length: 380
Range 1: 41 to 78

Score:38.9 bits(89), Expect:1e-07,
Method:Compositional matrix adjust.,
Identities:23/47(49%), Positives:27/47(57%), Gaps:12/47(25%)

Query  665  RAFINNQKMTFQFI---*PKIYLVSVNPKIVLI*CKSCNNVSEIVLN  534
            + FINN KMTFQF+     KIY VSVNPK+          V E+ LN
Sbjct  41   KTFINNHKMTFQFVLSTRSKIYSVSVNPKV---------EVRELTLN  78


Range 2: 15 to 43

Score:37.4 bits(85), Expect:1e-07,
Method:Compositional matrix adjust.,
Identities:15/29(52%), Positives:20/29(68%), Gaps:0/29(0%)

Query  744  IWWKSSPSYIVQFRIVCKRWNALFQDKGF  658
            I  +  P+ + +F+ VCKRWNALF DK F
Sbjct  15   ILSRVPPTSLDRFKTVCKRWNALFNDKTF  43


Query #25: XLOC_005828 Query ID: lcl|Query_76559 Length: 810

No significant similarity found.


Query #26: XLOC_005899 Query ID: lcl|Query_76560 Length: 637

No significant similarity found.


Query #27: XLOC_005901 Query ID: lcl|Query_76561 Length: 620

No significant similarity found.


Query #28: XLOC_005925 Query ID: lcl|Query_76562 Length: 1889

No significant similarity found.


Query #29: XLOC_005961 Query ID: lcl|Query_76563 Length: 553

Sequences producing significant alignments:
                                                                  Max    Total Query   E   Per.
Description                                                       Score  Score cover Value Ident  Accession
unnamed protein product [Arabidopsis thaliana]                    56.6   56.6  34%   2e-10 53.12  VYS52357.1
unnamed protein product [Arabidopsis thaliana]                    56.2   56.2  16%   2e-10 86.67  CAA0384287.1
transmembrane protein [Arabidopsis thaliana]                      55.8   55.8  16%   3e-10 86.67  NP_001326023.1
hypothetical protein AT2G14265 [Arabidopsis thaliana]             54.3   54.3  22%   3e-09 60.00  NP_001325174.1


Alignments:
>unnamed protein product [Arabidopsis thaliana]
Sequence ID: VYS52357.1 Length: 87
Range 1: 34 to 87

Score:56.6 bits(135), Expect:2e-10,
Method:Compositional matrix adjust.,
Identities:34/64(53%), Positives:37/64(57%), Gaps:10/64(15%)

Query  2    HGTLVSLQNTVNRISCYY*HV*RDLHFQDLMDMKLWFLYKPRLIESPVIMDMFAETSTSE  181
            HGTLVSLQNT    S            + L      F  KPRLIES VIMD+F ETSTS+
Sbjct  34   HGTLVSLQNTFEETSTS----------KILWTWNFGFFTKPRLIESSVIMDLFVETSTSK  83

Query  182  TLWT  193
            TLWT
Sbjct  84   TLWT  87


>unnamed protein product [Arabidopsis thaliana]
Sequence ID: CAA0384287.1 Length: 74
>unnamed protein product [Arabidopsis thaliana]
Sequence ID: VYS59150.1 Length: 74
Range 1: 15 to 44

Score:56.2 bits(134), Expect:2e-10,
Method:Compositional matrix adjust.,
Identities:26/30(87%), Positives:27/30(90%), Gaps:0/30(0%)

Query  68   RDLHFQDLMDMKLWFLYKPRLIESPVIMDM  157
            RDLH QDL DMKLW LYKPRLI+SPVIMDM
Sbjct  15   RDLHLQDLTDMKLWSLYKPRLIKSPVIMDM  44


>transmembrane protein [Arabidopsis thaliana]
Sequence ID: NP_001326023.1 Length: 74
>transmembrane protein [Arabidopsis thaliana]
Sequence ID: ANM63965.1 Length: 74
Range 1: 15 to 44

Score:55.8 bits(133), Expect:3e-10,
Method:Compositional matrix adjust.,
Identities:26/30(87%), Positives:27/30(90%), Gaps:0/30(0%)

Query  68   RDLHFQDLMDMKLWFLYKPRLIESPVIMDM  157
            RDLH QDL DMKLW LYKPRLI+SPVIMDM
Sbjct  15   RDLHLQDLTDMKLWSLYKPRLIKSPVIMDM  44


>hypothetical protein AT2G14265 [Arabidopsis thaliana]
Sequence ID: NP_001325174.1 Length: 111
>hypothetical protein AT2G14265 [Arabidopsis thaliana]
Sequence ID: ANM63061.1 Length: 111
Range 1: 62 to 111

Score:54.3 bits(129), Expect:3e-09,
Method:Compositional matrix adjust.,
Identities:30/50(60%), Positives:32/50(64%), Gaps:9/50(18%)

Query  91   YGHETLVSLQTTVNRISCYYGH---------VCRDIHL*DLMDIKLLPNS  213
            YGH TLVSLQ T   +   YGH         VCRD+HL DLMDIKLLPNS
Sbjct  62   YGHGTLVSLQNTRPPLPRSYGHGTLVSLQKHVCRDLHLQDLMDIKLLPNS  111


Query #30: XLOC_005992 Query ID: lcl|Query_76564 Length: 952

Sequences producing significant alignments:
                                                                  Max    Total Query   E   Per.
Description                                                       Score  Score cover Value Ident  Accession
unnamed protein product [Arabidopsis thaliana]                    88.2   88.2  23%   2e-21 65.79  VYS52439.1
hypothetical protein AT2G14265 [Arabidopsis thaliana]             88.2   88.2  29%   4e-21 57.89  NP_001325174.1
hypothetical protein AT1G62895 [Arabidopsis thaliana]             75.9   75.9  24%   7e-17 57.69  NP_001321703.1
transmembrane protein [Arabidopsis thaliana]                      59.3   150   34%   2e-14 84.38  NP_001326023.1
unnamed protein product [Arabidopsis thaliana]                    59.3   110   23%   6e-11 84.38  CAA0384287.1
unnamed protein product [Arabidopsis thaliana]                    57.8   57.8  24%   3e-10 48.28  VYS52357.1
unnamed protein product [Arabidopsis thaliana]                    41.6   77.4  16%   6e-08 73.33  CAA0360812.1
unnamed protein product [Arabidopsis thaliana]                    41.2   77.0  16%   7e-08 73.33  VYS52323.1


Alignments:
>unnamed protein product [Arabidopsis thaliana]
Sequence ID: VYS52439.1 Length: 86
Range 1: 11 to 63

Score:88.2 bits(217), Expect:2e-21,
Method:Compositional matrix adjust.,
Identities:50/76(66%), Positives:50/76(65%), Gaps:23/76(30%)

Query  171  FSKKLLLLQHWRLDLIFALSHVSNSFHLRSIEIPVNRISYYHGHAI*NGYKSVKSLQKPP  350
            FSKKLLLLQ WRLDLIFALSHVSNSFHLRSIEIP                       KPP
Sbjct  11   FSKKLLLLQQWRLDLIFALSHVSNSFHLRSIEIP-----------------------KPP  47

Query  351  PLIPYGHGTLVSLQNT  398
            PL PYGH TLVSLQNT
Sbjct  48   PLRPYGHETLVSLQNT  63


>hypothetical protein AT2G14265 [Arabidopsis thaliana]
Sequence ID: NP_001325174.1 Length: 111
>hypothetical protein AT2G14265 [Arabidopsis thaliana]
Sequence ID: ANM63061.1 Length: 111
Range 1: 29 to 107

Score:88.2 bits(217), Expect:4e-21,
Method:Compositional matrix adjust.,
Identities:55/95(58%), Positives:61/95(64%), Gaps:18/95(18%)

Query  195  QHWRLDLIFALSHVSNSFHLRSIEIPVNRISYYHGHAI*NGYKSVKSLQKPPPLIP--YG  368
            Q WRL+LI ALSHVSNSFHL SIEIP    S  +GH       ++ SLQ   P +P  YG
Sbjct  29   QQWRLELISALSHVSNSFHLCSIEIPRPLPSKPYGHG------TLVSLQNTRPPLPRSYG  82

Query  369  HGTLVSLQNTVNRISCYY*HVCRDLHLQDLMDMKL  473
            HGTLVSLQ           HVCRDLHLQDLMD+KL
Sbjct  83   HGTLVSLQK----------HVCRDLHLQDLMDIKL  107


>hypothetical protein AT1G62895 [Arabidopsis thaliana]
Sequence ID: NP_001321703.1 Length: 86
>hypothetical protein AT1G62895 [Arabidopsis thaliana]
Sequence ID: ANM59338.1 Length: 86
Range 1: 19 to 73

Score:75.9 bits(185), Expect:7e-17,
Method:Compositional matrix adjust.,
Identities:45/78(58%), Positives:45/78(57%), Gaps:23/78(29%)

Query  195  QHWRLDLIFALSHVSNSFHLRSIEIPVNRISYYHGHAI*NGYKSVKSLQKPPPLIPYGHG  374
            Q WRLDLIFALSHVSNSFHLRSIEIP                       KPPPL PYGH
Sbjct  19   QQWRLDLIFALSHVSNSFHLRSIEIP-----------------------KPPPLRPYGHE  55

Query  375  TLVSLQNTVNRISCYY*H  428
            TLVSLQNT    S  Y H
Sbjct  56   TLVSLQNTRPPPSRPYGH  73


>transmembrane protein [Arabidopsis thaliana]
Sequence ID: NP_001326023.1 Length: 74
>transmembrane protein [Arabidopsis thaliana]
Sequence ID: ANM63965.1 Length: 74
Range 1: 13 to 44

Score:59.3 bits(142), Expect:2e-14,
Method:Compositional matrix adjust.,
Identities:27/32(84%), Positives:29/32(90%), Gaps:0/32(0%)

Query  429  VCRDLHLQDLMDMKLWFFYKPQLIESPVIMDM  524
            VCRDLHLQDL DMKLW  YKP+LI+SPVIMDM
Sbjct  13   VCRDLHLQDLTDMKLWSLYKPRLIKSPVIMDM  44


Range 2: 40 to 74

Score:40.0 bits(92), Expect:2e-14,
Method:Compositional matrix adjust.,
Identities:19/35(54%), Positives:22/35(62%), Gaps:0/35(0%)

Query  586  VIMDTKKLFVDFFYSLQRSPPLRPYEHRNFTKFLR  690
            VIMD +         LQR+PP RPY +R FTKFLR
Sbjct  40   VIMDMRPPPPRPCGQLQRAPPPRPYGYRTFTKFLR  74


Range 3: 2 to 44

Score:51.2 bits(121), Expect:5e-08,
Method:Compositional matrix adjust.,
Identities:25/43(58%), Positives:33/43(76%), Gaps:0/43(0%)

Query  301  MLYRMDINQ*KVCRNLHL*YLMDMELWFLYKTRLIESPVIIDM  429
            +L+R++     VCR+LHL  L DM+LW LYK RLI+SPVI+DM
Sbjct  2    LLHRLEQMGGSVCRDLHLQDLTDMKLWSLYKPRLIKSPVIMDM  44


>unnamed protein product [Arabidopsis thaliana]
Sequence ID: CAA0384287.1 Length: 74
>unnamed protein product [Arabidopsis thaliana]
Sequence ID: VYS59150.1 Length: 74
Range 1: 13 to 44

Score:59.3 bits(142), Expect:6e-11,
Method:Compositional matrix adjust.,
Identities:27/32(84%), Positives:29/32(90%), Gaps:0/32(0%)

Query  429  VCRDLHLQDLMDMKLWFFYKPQLIESPVIMDM  524
            VCRDLHLQDL DMKLW  YKP+LI+SPVIMDM
Sbjct  13   VCRDLHLQDLTDMKLWSLYKPRLIKSPVIMDM  44


Range 2: 2 to 44

Score:50.8 bits(120), Expect:7e-08,
Method:Compositional matrix adjust.,
Identities:25/43(58%), Positives:33/43(76%), Gaps:0/43(0%)

Query  301  MLYRMDINQ*KVCRNLHL*YLMDMELWFLYKTRLIESPVIIDM  429
            +L+R++     VCR+LHL  L DM+LW LYK RLI+SPVI+DM
Sbjct  2    LLHRLEQMGGSVCRDLHLQDLTDMKLWSLYKPRLIKSPVIMDM  44


>unnamed protein product [Arabidopsis thaliana]
Sequence ID: VYS52357.1 Length: 87
Range 1: 1 to 87

Score:57.8 bits(138), Expect:3e-10,
Method:Compositional matrix adjust.,
Identities:42/87(48%), Positives:48/87(55%), Gaps:9/87(10%)

Query  232  MYQTRFIYVASRFQLIESPIIMDM--LYRMDINQ*KVC--RNLHL*YLMDMELW-----F  384
            M QTRF YVASRF LI+ P+IMDM  L         +   +N          LW     F
Sbjct  1    MCQTRFTYVASRFPLIKFPVIMDMRPLPSKPYGHGTLVSLQNTFEETSTSKILWTWNFGF  60

Query  385  LYKTRLIESPVIIDMFAETSTFKTLWT  465
              K RLIES VI+D+F ETST KTLWT
Sbjct  61   FTKPRLIESSVIMDLFVETSTSKTLWT  87


>unnamed protein product [Arabidopsis thaliana]
Sequence ID: CAA0360812.1 Length: 61
Range 1: 4 to 33

Score:41.6 bits(96), Expect:6e-08,
Method:Compositional matrix adjust.,
Identities:22/30(73%), Positives:23/30(76%), Gaps:0/30(0%)

Query  433  AETSTFKTLWT*NFGFFTNHS*SNLLLLWT  522
            +ETSTFKTLWT NFGFFTNH  S    LWT
Sbjct  4    SETSTFKTLWTWNFGFFTNHETSTSKALWT  33


Range 2: 38 to 61

Score:35.8 bits(81), Expect:6e-08,
Method:Compositional matrix adjust.,
Identities:17/24(71%), Positives:20/24(83%), Gaps:2/24(8%)

Query  620  FFT--VCRDLHL*DLMNIEILPNS  685
            FFT  VC+DL L DLM+IE+LPNS
Sbjct  38   FFTKHVCKDLRLQDLMDIELLPNS  61


>unnamed protein product [Arabidopsis thaliana]
Sequence ID: VYS52323.1 Length: 61
Range 1: 4 to 33

Score:41.2 bits(95), Expect:7e-08,
Method:Compositional matrix adjust.,
Identities:22/30(73%), Positives:23/30(76%), Gaps:0/30(0%)

Query  433  AETSTFKTLWT*NFGFFTNHS*SNLLLLWT  522
            +ETSTFKTLWT NFGFFTNH  S    LWT
Sbjct  4    SETSTFKTLWTWNFGFFTNHETSTSKALWT  33


Range 2: 38 to 61

Score:35.8 bits(81), Expect:7e-08,
Method:Compositional matrix adjust.,
Identities:17/24(71%), Positives:20/24(83%), Gaps:2/24(8%)

Query  620  FFT--VCRDLHL*DLMNIEILPNS  685
            FFT  VC+DL L DLM+IE+LPNS
Sbjct  38   FFTNHVCKDLRLQDLMDIELLPNS  61


Query #31: XLOC_005993 Query ID: lcl|Query_76565 Length: 1202

Sequences producing significant alignments:
                                                                  Max    Total Query   E   Per.
Description                                                       Score  Score cover Value Ident  Accession
unnamed protein product [Arabidopsis thaliana]                    166    405   55%   3e-73 89.22  CAA0266934.1
hypothetical protein AXX17_AT1G35970 [Arabidopsis thaliana]       159    214   28%   2e-47 97.75  OAP13408.1
hypothetical protein AXX17_AT2G04180 [Arabidopsis thaliana]       112    112   17%   2e-28 74.29  OAP11213.1
hypothetical protein AXX17_AT1G48910 [Arabidopsis thaliana]       70.5   113   26%   2e-18 46.58  OAP18984.1
Hypothetical protein [Arabidopsis thaliana]                       70.1   113   26%   2e-18 46.58  AAD25621.1
hypothetical protein AXX17_AT2G00260 [Arabidopsis thaliana]       65.1   104   23%   7e-16 38.67  OAP08664.1
putative helicase [Arabidopsis thaliana]                          59.7   98.2  26%   5e-14 36.05  AAD32757.1
putative helicase [Arabidopsis thaliana]                          46.2   84.3  27%   7e-10 29.55  AAD25596.1
hypothetical protein [Arabidopsis thaliana]                       45.4   83.9  26%   8e-10 30.34  AAG51081.1
unnamed protein product [Arabidopsis thaliana]                    61.2   61.2  17%   2e-09 41.43  VYS62284.1
unnamed protein product [Arabidopsis thaliana]                    60.8   60.8  17%   2e-09 41.43  CAA0408802.1
hypothetical protein, 5' partial; 93859-91015 [Arabidopsis...     45.4   82.8  26%   2e-09 30.34  AAG52315.1
unnamed protein product [Arabidopsis thaliana]                    60.8   60.8  17%   2e-09 41.43  CAA0408793.1
helicase-like protein [Arabidopsis thaliana]                      47.4   78.9  30%   2e-08 29.35  BAB02793.1
contains similarity to C. elegans predicted proteins...           49.3   77.4  27%   9e-08 30.11  AAD17353.1


Alignments:
>unnamed protein product [Arabidopsis thaliana]
Sequence ID: CAA0266934.1 Length: 270
Range 1: 169 to 270

Score:166 bits(419), Expect:3e-73,
Method:Compositional matrix adjust.,
Identities:91/102(89%), Positives:94/102(92%), Gaps:3/102(2%)

Query  634  VEDISFDIHYS---RPAVQHLSLHLPDQQPLIFDANQSLDSVISRDGVDRTMFTEWMKIN  804
            V + +F  H +   RPAVQHLSLHLPDQQPLIFDANQSLDSVISRDGVDRTMFTEWMKIN
Sbjct  169  VRNYNFKAHMNVECRPAVQHLSLHLPDQQPLIFDANQSLDSVISRDGVDRTMFTEWMKIN  228

Query  805  QSDEEAKSMTYVQFPTRFVWNtttkkwtkrkQGFKKFEALDI  930
            QSDEEAKSMTYVQFPTRFVWNTTTKKWTKRKQGFKKFEALDI
Sbjct  229  QSDEEAKSMTYVQFPTRFVWNTTTKKWTKRKQGFKKFEALDI  270


Range 2: 104 to 168

Score:130 bits(328), Expect:3e-73,
Method:Compositional matrix adjust.,
Identities:65/66(98%), Positives:65/66(98%), Gaps:1/66(1%)

Query  333  DQTNPPKITFNIILTCILERKYNKNIPSNDVSYQQVKKSQPLLVSSGSETKEQQVINMPS  512
            DQTNPPKITFNIILTCILERKYNKNIPSNDVSYQQVKKSQPLLVSSGSETKE QVINMPS
Sbjct  104  DQTNPPKITFNIILTCILERKYNKNIPSNDVSYQQVKKSQPLLVSSGSETKE-QVINMPS  162

Query  513  HQYKPY  530
            HQYKPY
Sbjct  163  HQYKPY  168


Range 3: 1 to 57

Score:108 bits(269), Expect:3e-26,
Method:Compositional matrix adjust.,
Identities:52/57(91%), Positives:55/57(96%), Gaps:0/57(0%)

Query  152  MISMLLLPQTVIVEPTTNVKLFSQTKRFWTVLKSDIAPLVTNMLGCKLLELVNKGDR  322
            MISMLLLPQTVIVEPTTNVKLFSQTKRFWTVLKSD APLVTN+LGC+LLELVNKG+
Sbjct  1    MISMLLLPQTVIVEPTTNVKLFSQTKRFWTVLKSDTAPLVTNILGCELLELVNKGEE  57


>hypothetical protein AXX17_AT1G35970 [Arabidopsis thaliana]
Sequence ID: OAP13408.1 Length: 112
Range 1: 24 to 112

Score:159 bits(402), Expect:2e-47,
Method:Compositional matrix adjust.,
Identities:87/89(98%), Positives:89/89(100%), Gaps:0/89(0%)

Query  664  SRPAVQHLSLHLPDQQPLIFDANQSLDSVISRDGVDRTMFTEWMKINQSDEEAKSMTYVQ  843
            ++PAVQHLSLHLPDQQPLIFDANQSLDSVISRDGVDRTMFTEWMKINQSDEEAKSMTYVQ
Sbjct  24   NKPAVQHLSLHLPDQQPLIFDANQSLDSVISRDGVDRTMFTEWMKINQSDEEAKSMTYVQ  83

Query  844  FPTRFVWNtttkkwtkrkQGFKKFEALDI  930
            FPTRFVWNTTTKKWTKRKQGFKKFEALDI
Sbjct  84   FPTRFVWNTTTKKWTKRKQGFKKFEALDI  112


Range 2: 1 to 24

Score:54.7 bits(130), Expect:9e-09,
Method:Compositional matrix adjust.,
Identities:23/24(96%), Positives:23/24(95%), Gaps:0/24(0%)

Query  418  MMCPTNKLRKVNHY*FHPVLKPKN  489
            MMCPTNKLRKVNHY FHPVLKPKN
Sbjct  1    MMCPTNKLRKVNHYWFHPVLKPKN  24


>hypothetical protein AXX17_AT2G04180 [Arabidopsis thaliana]
Sequence ID: OAP11213.1 Length: 215
Range 1: 61 to 130

Score:112 bits(280), Expect:2e-28,
Method:Compositional matrix adjust.,
Identities:52/70(74%), Positives:59/70(84%), Gaps:0/70(0%)

Query  79   EASVLPNRLPNFQDQMVLGNLYGINDFDVVASSNRYRRTNHKCKIVFTDKTFLDSVKERH  258
            EAS+ P RLP F+DQ+VLGNLY INDFDVV SS RYRRTNH+ KIVFTDKTFL+SVKERH
Sbjct  61   EASIHPTRLPKFKDQLVLGNLYNINDFDVVISSKRYRRTNHEWKIVFTDKTFLESVKERH  120

Query  259  CSIGYEYVRM  288
              I YE+ R+
Sbjct  121  YCISYEFFRL  130


>hypothetical protein AXX17_AT1G48910 [Arabidopsis thaliana]
Sequence ID: OAP18984.1 Length: 1295
Range 1: 550 to 622

Score:70.5 bits(171), Expect:2e-18,
Method:Composition-based stats.,
Identities:34/73(47%), Positives:51/73(69%), Gaps:0/73(0%)

Query  649  FDIHYSRPAVQHLSLHLPDQQPLIFDANQSLDSVISRDGVDRTMFTEWMKINQSDEEAKS  828
            F+IH+  P VQ L LHLP +Q  IF+  ++L++V  R G +RTM TE+ ++N+  E+A+
Sbjct  550  FNIHHHNPPVQRLPLHLPGEQSTIFEEEENLENVEYRYGHERTMLTEYFELNKICEDARK  609

Query  829  MTYVQFPTRFVWN  867
            + YVQ PT FVW+
Sbjct  610  LKYVQVPTMFVWD  622


Range 2: 509 to 548

Score:43.1 bits(100), Expect:2e-18,
Method:Composition-based stats.,
Identities:19/40(48%), Positives:26/40(65%), Gaps:5/40(12%)

Query  542  GSVVKTKGESNYVDE-----IKKFQECRYISACEAMWRIF  646
            G+ V  +G  N  +E     I ++ +CRY+SACEAMWRIF
Sbjct  509  GNSVNGQGSGNGFEEKPRNEINEYLDCRYLSACEAMWRIF  548


>Hypothetical protein [Arabidopsis thaliana]
Sequence ID: AAD25621.1 Length: 1250
Range 1: 564 to 636

Score:70.1 bits(170), Expect:2e-18,
Method:Composition-based stats.,
Identities:34/73(47%), Positives:51/73(69%), Gaps:0/73(0%)

Query  649  FDIHYSRPAVQHLSLHLPDQQPLIFDANQSLDSVISRDGVDRTMFTEWMKINQSDEEAKS  828
            F+IH+  P VQ L LHLP +Q  IF+  ++L++V  R G +RTM TE+ ++N+  E+A+
Sbjct  564  FNIHHHNPPVQRLPLHLPGEQSTIFEEEENLENVEYRYGHERTMLTEYFELNKICEDARK  623

Query  829  MTYVQFPTRFVWN  867
            + YVQ PT FVW+
Sbjct  624  LKYVQVPTMFVWD  636


Range 2: 523 to 562

Score:43.1 bits(100), Expect:2e-18,
Method:Composition-based stats.,
Identities:19/40(48%), Positives:26/40(65%), Gaps:5/40(12%)

Query  542  GSVVKTKGESNYVDE-----IKKFQECRYISACEAMWRIF  646
            G+ V  +G  N  +E     I ++ +CRY+SACEAMWRIF
Sbjct  523  GNSVNGQGSGNGFEEKPRNEINEYLDCRYLSACEAMWRIF  562


>hypothetical protein AXX17_AT2G00260 [Arabidopsis thaliana]
Sequence ID: OAP08664.1 Length: 2020
Range 1: 644 to 715

Score:65.1 bits(157), Expect:7e-16,
Method:Composition-based stats.,
Identities:29/75(39%), Positives:50/75(66%), Gaps:6/75(8%)

Query  649  FDIHYSRPAVQHLSLHLPDQQPLIFDANQSLDSVISRDGVDRTMFTEWMKINQ---SDEE  819
            FD+H+S  +VQ LS+HLP +Q +I+D ++    V++++    + F E++K N+    D E
Sbjct  644  FDLHHSSTSVQRLSIHLPGEQKIIYDDDE---DVLNKEENQTSQFLEFLKTNKKIAEDPE  700

Query  820  AKSMTYVQFPTRFVW  864
            A+  TY++FP+ FVW
Sbjct  701  ARKFTYIEFPSHFVW  715


Range 2: 621 to 642

Score:39.3 bits(90), Expect:7e-16,
Method:Composition-based stats.,
Identities:14/22(64%), Positives:19/22(86%), Gaps:0/22(0%)

Query  581  DEIKKFQECRYISACEAMWRIF  646
            +EIK++ +CRYIS CEA W+IF
Sbjct  621  NEIKRWYDCRYISPCEAAWQIF  642


>putative helicase [Arabidopsis thaliana]
Sequence ID: AAD32757.1 Length: 1241
Range 1: 617 to 702

Score:59.7 bits(143), Expect:5e-14,
Method:Composition-based stats.,
Identities:31/86(36%), Positives:47/86(54%), Gaps:0/86(0%)

Query  649  FDIHYSRPAVQHLSLHLPDQQPLIFDANQSLDSVISRDGVDRTMFTEWMKINQSDEEAKS  828
            F I Y    V  LS HLP +QPL F+  Q++D +  +   +  MF  ++K+NQ  E A+
Sbjct  617  FPIQYRTTPVMKLSYHLPGKQPLCFEDTQNIDELSEKKANEDFMFIGFLKLNQECEFARQ  676

Query  829  MTYVQFPTRFVWNtttkkwtkrkQGF  906
              Y + P  F W+   K+W  R++GF
Sbjct  677  FIYTEIPPYFTWDGQNKQWKLRERGF  702


Range 2: 595 to 617

Score:38.5 bits(88), Expect:5e-14,
Method:Composition-based stats.,
Identities:14/23(61%), Positives:19/23(82%), Gaps:0/23(0%)

Query  584  EIKKFQECRYISACEAMWRIFRL  652
            EIK + +CRY+SA EA+WRIF+
Sbjct  595  EIKDYFDCRYVSASEAVWRIFKF  617


>putative helicase [Arabidopsis thaliana]
Sequence ID: AAD25596.1 Length: 1219
Range 1: 477 to 553

Score:46.2 bits(108), Expect:7e-10,
Method:Composition-based stats.,
Identities:26/88(30%), Positives:49/88(55%), Gaps:11/88(12%)

Query  643  ISFDIHYSRPAVQHLSLHLPDQQPLIFDANQSLDSVISRDGVDRTMFTEWMKINQSDEEA  822
            + + IHY   +V  LS HLP +Q + F  ++ +++V+++  +D ++           + A
Sbjct  477  LKYPIHYRSTSVMKLSFHLPGEQYIYFKGDEEVETVLNKADLDGSI-----------QIA  525

Query  823  KSMTYVQFPTRFVWNtttkkwtkrkQGF  906
            + +TY   PTRF ++   KK+  RK+GF
Sbjct  526  RKLTYPNIPTRFTYDPKEKKFNLRKKGF  553


Range 2: 456 to 478

Score:38.1 bits(87), Expect:7e-10,
Method:Composition-based stats.,
Identities:13/23(57%), Positives:18/23(78%), Gaps:0/23(0%)

Query  581  DEIKKFQECRYISACEAMWRIFR  649
            +E++ F  CRY+SACEA WRI +
Sbjct  456  NEVEDFFNCRYVSACEAAWRILK  478


>hypothetical protein [Arabidopsis thaliana]
Sequence ID: AAG51081.1 Length: 1678
Range 1: 1004 to 1091

Score:45.4 bits(106), Expect:8e-10,
Method:Compositional matrix adjust.,
Identities:27/89(30%), Positives:44/89(49%), Gaps:7/89(7%)

Query  658   HYSRPAVQHLSLHLPDQQPLIFDANQSLDSVISRDGVDRTMFTEWMKIN------QSDEE  819
             H S P VQ LS H+  +QP  FD   +++ V+ R     + F  W+ +N      ++ +
Sbjct  1004  HISTP-VQKLSFHVEGKQPAYFDPKSNIEDVLERVANVDSQFMAWLTLNRRNAVGKNGKR  1062

Query  820   AKSMTYVQFPTRFVWNtttkkwtkrkQGF  906
             A+   Y + P  F W+   K + KR +GF
Sbjct  1063  ARECLYAEIPAYFTWDGENKSFKKRTRGF  1091


Range 2: 977 to 999

Score:38.5 bits(88), Expect:8e-10,
Method:Composition-based stats.,
Identities:14/23(61%), Positives:20/23(86%), Gaps:0/23(0%)

Query  581  DEIKKFQECRYISACEAMWRIFR  649
            +EIK + +CRY+SA EA+WRIF+
Sbjct  977  NEIKDWFDCRYVSASEAVWRIFK  999


>unnamed protein product [Arabidopsis thaliana]
Sequence ID: VYS62284.1 Length: 461
Range 1: 65 to 134

Score:61.2 bits(147), Expect:2e-09,
Method:Compositional matrix adjust.,
Identities:29/70(41%), Positives:42/70(60%), Gaps:0/70(0%)

Query  79   EASVLPNRLPNFQDQMVLGNLYGINDFDVVASSNRYRRTNHKCKIVFTDKTFLDSVKERH  258
            + S+ P RL  F+ +++ GN++ I+DFDV    N YR TNH  KI F D T ++ V   H
Sbjct  65   QGSIHPRRLHKFKPKLIEGNIFSIHDFDVKKQQNSYRLTNHDYKIFFNDHTSMNVVDNIH  124

Query  259  CSIGYEYVRM  288
              I +EY R+
Sbjct  125  NDIPHEYFRI  134


>unnamed protein product [Arabidopsis thaliana]
Sequence ID: CAA0408802.1 Length: 466
Range 1: 65 to 134

Score:60.8 bits(146), Expect:2e-09,
Method:Compositional matrix adjust.,
Identities:29/70(41%), Positives:42/70(60%), Gaps:0/70(0%)

Query  79   EASVLPNRLPNFQDQMVLGNLYGINDFDVVASSNRYRRTNHKCKIVFTDKTFLDSVKERH  258
            + S+ P RL  F+ +++ GN++ I+DFDV    N YR TNH  KI F D T ++ V   H
Sbjct  65   QGSIHPRRLHKFKPKLIEGNIFSIHDFDVKKQQNSYRLTNHDYKIFFNDHTSMNVVDNIH  124

Query  259  CSIGYEYVRM  288
              I +EY R+
Sbjct  125  NDIPHEYFRI  134


>hypothetical protein, 5' partial; 93859-91015, partial [Arabidopsis thaliana]
Sequence ID: AAG52315.1 Length: 729
Range 1: 92 to 179

Score:45.4 bits(106), Expect:2e-09,
Method:Compositional matrix adjust.,
Identities:27/89(30%), Positives:44/89(49%), Gaps:7/89(7%)

Query  658  HYSRPAVQHLSLHLPDQQPLIFDANQSLDSVISRDGVDRTMFTEWMKIN------QSDEE  819
            H S P VQ LS H+  +QP  FD   +++ V+ R     + F  W+ +N      ++ +
Sbjct  92   HISTP-VQKLSFHVEGKQPAYFDPKSNIEDVLERVANVDSQFMAWLTLNRRNAVGKNGKR  150

Query  820  AKSMTYVQFPTRFVWNtttkkwtkrkQGF  906
            A+   Y + P  F W+   K + KR +GF
Sbjct  151  ARECLYAEIPAYFTWDGENKSFKKRTRGF  179


Range 2: 65 to 87

Score:37.4 bits(85), Expect:2e-09,
Method:Compositional matrix adjust.,
Identities:14/23(61%), Positives:20/23(86%), Gaps:0/23(0%)

Query  581  DEIKKFQECRYISACEAMWRIFR  649
            +EIK + +CRY+SA EA+WRIF+
Sbjct  65   NEIKDWFDCRYVSASEAVWRIFK  87


>unnamed protein product [Arabidopsis thaliana]
Sequence ID: CAA0408793.1 Length: 499
Range 1: 65 to 134

Score:60.8 bits(146), Expect:2e-09,
Method:Compositional matrix adjust.,
Identities:29/70(41%), Positives:42/70(60%), Gaps:0/70(0%)

Query  79   EASVLPNRLPNFQDQMVLGNLYGINDFDVVASSNRYRRTNHKCKIVFTDKTFLDSVKERH  258
            + S+ P RL  F+ +++ GN++ I+DFDV    N YR TNH  KI F D T ++ V   H
Sbjct  65   QGSIHPRRLHKFKPKLIEGNIFSIHDFDVKKQQNSYRLTNHDYKIFFNDHTSMNVVDNIH  124

Query  259  CSIGYEYVRM  288
              I +EY R+
Sbjct  125  NDIPHEYFRI  134


>helicase-like protein [Arabidopsis thaliana]
Sequence ID: BAB02793.1 Length: 1428
Range 1: 667 to 758

Score:47.4 bits(111), Expect:2e-08,
Method:Composition-based stats.,
Identities:27/92(29%), Positives:44/92(47%), Gaps:6/92(6%)

Query  649  FDIHYSRPAVQHLSLHLPDQQPLIFDANQSLDSVISRDGVDRTMFTEWMKINQSD-----  813
            F I +    VQ LS H   +QP  FDA   +  V+ R   + + F  W+ +N+ +
Sbjct  667  FPIQHRSTPVQKLSFHDKGKQPAYFDAKAKMADVLERVSNEDSQFLAWLTLNRKNAVGKN  726

Query  814  -EEAKSMTYVQFPTRFVWNtttkkwtkrkQGF  906
             + A+   Y + P  F W+   K++ KR +GF
Sbjct  727  GKRARDCLYAEIPAYFTWDGENKQFKKRTRGF  758


Range 2: 621 to 667

Score:31.6 bits(70), Expect:2e-08,
Method:Compositional matrix adjust.,
Identities:17/47(36%), Positives:25/47(53%), Gaps:9/47(19%)

Query  539  SGSVVKTKGESNYVDEIKKFQEC---------RYISACEAMWRIFRL  652
            +  VV++       DEIK + +C         RY+SA EA+WRIF+
Sbjct  621  TAPVVESDTTEKKKDEIKDWFDCSSYISFSPARYVSASEAIWRIFKF  667


>contains similarity to C. elegans predicted proteins (GB:AF025472, AF04064 and AF026212) [Arabidopsis thaliana]
Sequence ID: AAD17353.1 Length: 448
>hypothetical protein [Arabidopsis thaliana]
Sequence ID: CAB77939.1 Length: 448
Range 1: 328 to 420

Score:49.3 bits(116), Expect:9e-08,
Method:Compositional matrix adjust.,
Identities:28/93(30%), Positives:46/93(49%), Gaps:6/93(6%)

Query  646  SFDIHYSRPAVQHLSLHLPDQQPLIFDANQSLDSVISRDGVDRTMFTEWMKINQSD----  813
             F I +    VQ LS H+  +QP  FDA   +  V+ R   + + F  W+ +N+ +
Sbjct  328  KFPIQHRSTPVQKLSFHVEGKQPAYFDAKAKMVDVLERVSNEDSQFMVWLTLNKKNVVGK  387

Query  814  --EEAKSMTYVQFPTRFVWNtttkkwtkrkQGF  906
              + A++  Y + PT F W+   K + KR +GF
Sbjct  388  NGKRARNCLYAEIPTYFTWDGENKPFKKRTRGF  420


Range 2: 306 to 329

Score:28.1 bits(61), Expect:9e-08,
Method:Compositional matrix adjust.,
Identities:12/24(50%), Positives:17/24(70%), Gaps:0/24(0%)

Query  581  DEIKKFQECRYISACEAMWRIFRL  652
            D IK + +  Y+SA EA+WRIF+
Sbjct  306  DGIKDWFDYIYVSASEAIWRIFKF  329


Query #32: XLOC_006003 Query ID: lcl|Query_76566 Length: 1543

Sequences producing significant alignments:
                                                                  Max    Total Query   E   Per.
Description                                                       Score  Score cover Value Ident  Accession
hypothetical protein AXX17_AT1G36110 [Arabidopsis thaliana]       83.6   83.6  16%   1e-18 59.30  OAP19536.1
unnamed protein product [Arabidopsis thaliana]                    81.6   81.6  16%   1e-16 58.14  CAA0267218.1
unnamed protein product [Arabidopsis thaliana]                    79.7   79.7  16%   4e-16 58.14  VYS48040.1
unknown [Arabidopsis thaliana]                                    77.4   77.4  15%   7e-16 58.33  ABK28139.1
hypothetical protein At1g35365 [Arabidopsis thaliana]             77.4   77.4  15%   7e-16 58.33  ABE65397.1
unnamed protein product [Arabidopsis thaliana]                    77.0   77.0  15%   1e-15 58.33  CAA0267271.1
unnamed protein product [Arabidopsis thaliana]                    76.3   76.3  15%   8e-15 57.14  VYS48041.1
RING/U-box superfamily protein [Arabidopsis thaliana]             71.6   71.6  10%   2e-12 62.50  NP_180109.2
hypothetical protein [Arabidopsis thaliana]                       71.2   71.2  10%   2e-12 62.50  AAD20698.1
unnamed protein product [Arabidopsis thaliana]                    65.5   65.5  14%   3e-11 51.95  CAA0266139.1
hypothetical protein AXX17_AT1G35540 [Arabidopsis thaliana]       65.5   65.5  14%   3e-11 51.95  OAP15306.1
unnamed protein product [Arabidopsis thaliana]                    65.1   65.1  14%   4e-11 51.95  VYS47999.1
hypothetical protein (DUF1184) [Arabidopsis thaliana]             60.1   60.1  16%   2e-09 51.72  NP_189428.1
hypothetical protein AXX17_AT3G30440 [Arabidopsis thaliana]       59.7   59.7  16%   2e-09 51.72  OAP01379.1
hypothetical protein (DUF1184) [Arabidopsis thaliana]             60.1   60.1  16%   2e-09 51.72  NP_001325896.1


Alignments:
>hypothetical protein AXX17_AT1G36110 [Arabidopsis thaliana]
Sequence ID: OAP19536.1 Length: 122
Range 1: 35 to 118

Score:83.6 bits(205), Expect:1e-18,
Method:Compositional matrix adjust.,
Identities:51/86(59%), Positives:61/86(70%), Gaps:4/86(4%)

Query  697  EKQKDEAELLGVE--LFMSEAMFLLSDDIYSMLLFFYKIILKYTGNNERKNPTQVEKILI  524
            E +KDEA+ LGVE  LF++EAMFLLSDD+ SMLLF    +LK  GN     P  V ++L
Sbjct  35   EMKKDEAKRLGVEFSLFVAEAMFLLSDDLRSMLLFCL-WLLKDAGNKGLTAPVVVGRLLC  93

Query  523  NVRLYVFETNIKPKNGVSQADGKSTR  446
             V L+VFET IKPKNGV QADGKS +
Sbjct  94   -VVLHVFETYIKPKNGVYQADGKSNQ  118


>unnamed protein product [Arabidopsis thaliana]
Sequence ID: CAA0267218.1 Length: 241
Range 1: 35 to 118

Score:81.6 bits(200), Expect:1e-16,
Method:Compositional matrix adjust.,
Identities:50/86(58%), Positives:61/86(70%), Gaps:4/86(4%)

Query  697  EKQKDEAELLGVE--LFMSEAMFLLSDDIYSMLLFFYKIILKYTGNNERKNPTQVEKILI  524
            E +K+EA+ LGVE  LF++EAMFLLSDD+ SMLLF    +LK  GN     P  V ++L
Sbjct  35   EMKKNEAKRLGVEFSLFVAEAMFLLSDDLRSMLLFCL-WLLKDAGNKGLTAPVVVGRLLC  93

Query  523  NVRLYVFETNIKPKNGVSQADGKSTR  446
             V L+VFET IKPKNGV QADGKS +
Sbjct  94   -VVLHVFETYIKPKNGVYQADGKSNQ  118


>unnamed protein product [Arabidopsis thaliana]
Sequence ID: VYS48040.1 Length: 239
Range 1: 35 to 116

Score:79.7 bits(195), Expect:4e-16,
Method:Compositional matrix adjust.,
Identities:50/86(58%), Positives:59/86(68%), Gaps:6/86(6%)

Query  697  EKQKDEAELLGVE--LFMSEAMFLLSDDIYSMLLFFYKIILKYTGNNERKNPTQVEKILI  524
            E +KDEA+ LGVE  LF++EAMFLLSDD+ SMLLF    +LK  GN     P      L+
Sbjct  35   EMKKDEAKRLGVEFSLFVAEAMFLLSDDLRSMLLFCL-WLLKDAGNKGLTAPVVR---LL  90

Query  523  NVRLYVFETNIKPKNGVSQADGKSTR  446
             V L+VFET IKPKNGV QADGKS +
Sbjct  91   CVVLHVFETYIKPKNGVYQADGKSNQ  116


>unknown, partial [Arabidopsis thaliana]
Sequence ID: ABK28139.1 Length: 180
Range 1: 57 to 138

Score:77.4 bits(189), Expect:7e-16,
Method:Compositional matrix adjust.,
Identities:49/84(58%), Positives:58/84(69%), Gaps:4/84(4%)

Query  697  EKQKDEAELLGVEL--FMSEAMFLLSDDIYSMLLFFYKIILKYTGNNERKNPTQVEKILI  524
            EKQK+EA  LGVEL  FM+EAMF+LSDD+ SMLL    +I+KY G  +  N   V   L
Sbjct  57   EKQKEEAIRLGVELSLFMAEAMFILSDDLRSMLLLC-PLIVKYAGF-KYTNDGPVVGRLF  114

Query  523  NVRLYVFETNIKPKNGVSQADGKS  452
             V L+VFET IKPKNGV Q +GKS
Sbjct  115  CVMLHVFETYIKPKNGVYQVNGKS  138


>hypothetical protein At1g35365 [Arabidopsis thaliana]
Sequence ID: ABE65397.1 Length: 179
Range 1: 57 to 138

Score:77.4 bits(189), Expect:7e-16,
Method:Compositional matrix adjust.,
Identities:49/84(58%), Positives:58/84(69%), Gaps:4/84(4%)

Query  697  EKQKDEAELLGVEL--FMSEAMFLLSDDIYSMLLFFYKIILKYTGNNERKNPTQVEKILI  524
            EKQK+EA  LGVEL  FM+EAMF+LSDD+ SMLL    +I+KY G  +  N   V   L
Sbjct  57   EKQKEEAIRLGVELSLFMAEAMFILSDDLRSMLLLC-PLIVKYAGF-KYTNDGPVVGRLF  114

Query  523  NVRLYVFETNIKPKNGVSQADGKS  452
             V L+VFET IKPKNGV Q +GKS
Sbjct  115  CVMLHVFETYIKPKNGVYQVNGKS  138


>unnamed protein product [Arabidopsis thaliana]
Sequence ID: CAA0267271.1 Length: 186
Range 1: 45 to 126

Score:77.0 bits(188), Expect:1e-15,
Method:Compositional matrix adjust.,
Identities:49/84(58%), Positives:58/84(69%), Gaps:4/84(4%)

Query  697  EKQKDEAELLGVEL--FMSEAMFLLSDDIYSMLLFFYKIILKYTGNNERKNPTQVEKILI  524
            EKQK+EA  LGVEL  FM+EAMF+LSDD+ SMLL    +I+KY G  +  N   V   L
Sbjct  45   EKQKEEAIRLGVELSLFMAEAMFILSDDLRSMLLLC-PLIVKYAGF-KYTNDGPVVGRLF  102

Query  523  NVRLYVFETNIKPKNGVSQADGKS  452
             V L+VFET IKPKNGV Q +GKS
Sbjct  103  CVMLHVFETYIKPKNGVYQVNGKS  126


>unnamed protein product [Arabidopsis thaliana]
Sequence ID: VYS48041.1 Length: 250
Range 1: 45 to 126

Score:76.3 bits(186), Expect:8e-15,
Method:Compositional matrix adjust.,
Identities:48/84(57%), Positives:58/84(69%), Gaps:4/84(4%)

Query  697  EKQKDEAELLGVEL--FMSEAMFLLSDDIYSMLLFFYKIILKYTGNNERKNPTQVEKILI  524
            EKQK+EA  LGVEL  F++EAMF+LSDD+ SMLL    +I+KY G  +  N   V   L
Sbjct  45   EKQKEEAIRLGVELSLFVAEAMFILSDDLRSMLLLC-PLIVKYAGY-KYTNDGPVVGRLF  102

Query  523  NVRLYVFETNIKPKNGVSQADGKS  452
             V LYVFET IKPKNGV + DG+S
Sbjct  103  CVMLYVFETYIKPKNGVYEFDGRS  126


>RING/U-box superfamily protein [Arabidopsis thaliana]
Sequence ID: NP_180109.2 Length: 603
>RING/U-box superfamily protein [Arabidopsis thaliana]
Sequence ID: AEC07696.1 Length: 603
Range 1: 83 to 138

Score:71.6 bits(174), Expect:2e-12,
Method:Compositional matrix adjust.,
Identities:35/56(63%), Positives:43/56(76%), Gaps:0/56(0%)

Query  755  QRPKIAVAGNCRYFWSLNQRQANEQLLVDVAGFSRQRQTNEQGQLTLIPSFLKRSR  922
            Q+ +  +AG+ RYFWSLNQRQANEQLLV VAGFSRQRQ NEQG  ++I   +  S+
Sbjct  83   QKLRSLIAGDYRYFWSLNQRQANEQLLVVVAGFSRQRQANEQGLNSIITGDIPNSK  138


>hypothetical protein [Arabidopsis thaliana]
Sequence ID: AAD20698.1 Length: 546
>hypothetical protein [Arabidopsis thaliana]
Sequence ID: AAM15373.1 Length: 546
Range 1: 83 to 138

Score:71.2 bits(173), Expect:2e-12,
Method:Compositional matrix adjust.,
Identities:35/56(63%), Positives:43/56(76%), Gaps:0/56(0%)

Query  755  QRPKIAVAGNCRYFWSLNQRQANEQLLVDVAGFSRQRQTNEQGQLTLIPSFLKRSR  922
            Q+ +  +AG+ RYFWSLNQRQANEQLLV VAGFSRQRQ NEQG  ++I   +  S+
Sbjct  83   QKLRSLIAGDYRYFWSLNQRQANEQLLVVVAGFSRQRQANEQGLNSIITGDIPNSK  138


>unnamed protein product [Arabidopsis thaliana]
Sequence ID: CAA0266139.1 Length: 233
Range 1: 42 to 115

Score:65.5 bits(158), Expect:3e-11,
Method:Compositional matrix adjust.,
Identities:40/77(52%), Positives:52/77(67%), Gaps:5/77(6%)

Query  670  LGVEL--FMSEAMFLLSDDIYSMLLFFYKIILKYTGNNERKNPTQVEKILINVRLYVFET  497
            LGVEL  F++EAMFLLSDDI SML+F +  + KY G      P  V ++   ++ YVFET
Sbjct  42   LGVELSLFVAEAMFLLSDDIRSMLVFCF-WLFKYAGKRNFDGPV-VGRVFYVIQ-YVFET  98

Query  496  NIKPKNGVSQADGKSTR  446
             IKPKNGV +  GKS++
Sbjct  99   YIKPKNGVYKDGGKSSQ  115


>hypothetical protein AXX17_AT1G35540 [Arabidopsis thaliana]
Sequence ID: OAP15306.1 Length: 234
Range 1: 43 to 116

Score:65.5 bits(158), Expect:3e-11,
Method:Compositional matrix adjust.,
Identities:40/77(52%), Positives:52/77(67%), Gaps:5/77(6%)

Query  670  LGVEL--FMSEAMFLLSDDIYSMLLFFYKIILKYTGNNERKNPTQVEKILINVRLYVFET  497
            LGVEL  F++EAMFLLSDDI SML+F +  + KY G      P  V ++   ++ YVFET
Sbjct  43   LGVELSLFVAEAMFLLSDDIRSMLVFCF-WLFKYAGKRNFDGPV-VGRVFYVIQ-YVFET  99

Query  496  NIKPKNGVSQADGKSTR  446
             IKPKNGV +  GKS++
Sbjct  100  YIKPKNGVYKDGGKSSQ  116


>unnamed protein product [Arabidopsis thaliana]
Sequence ID: VYS47999.1 Length: 248
Range 1: 57 to 130

Score:65.1 bits(157), Expect:4e-11,
Method:Compositional matrix adjust.,
Identities:40/77(52%), Positives:52/77(67%), Gaps:5/77(6%)

Query  670  LGVEL--FMSEAMFLLSDDIYSMLLFFYKIILKYTGNNERKNPTQVEKILINVRLYVFET  497
            LGVEL  F++EAMFLLSDDI SML+F +  + KY G      P  V ++   ++ YVFET
Sbjct  57   LGVELSLFVAEAMFLLSDDIRSMLVFCF-WLFKYAGKRNFDGPV-VGRVFYVIQ-YVFET  113

Query  496  NIKPKNGVSQADGKSTR  446
             IKPKNGV +  GKS++
Sbjct  114  YIKPKNGVYKDGGKSSQ  130


>hypothetical protein (DUF1184) [Arabidopsis thaliana]
Sequence ID: NP_189428.1 Length: 244
>hypothetical protein (DUF1184) [Arabidopsis thaliana]
Sequence ID: AEE77377.1 Length: 244
Range 1: 39 to 121

Score:60.1 bits(144), Expect:2e-09,
Method:Compositional matrix adjust.,
Identities:45/87(52%), Positives:52/87(59%), Gaps:7/87(8%)

Query  697  EKQKDEAELLGVEL--FMSEAMFLLSDDIYSMLLFFYKIILKYTGNNERKNPTQ-VEKIL  527
            E QKDEA  LGVEL  F++E M LLSD++ SMLLF   ++     N E    T  V   L
Sbjct  39   EMQKDEAIRLGVELSLFVAETMVLLSDNLRSMLLFCLWVL----TNAECTYFTDPVVGRL  94

Query  526  INVRLYVFETNIKPKNGVSQADGKSTR  446
              V LYV ET IKPKNGV  AD KS +
Sbjct  95   HCVMLYVLETYIKPKNGVYLADVKSNQ  121


>hypothetical protein AXX17_AT3G30440 [Arabidopsis thaliana]
Sequence ID: OAP01379.1 Length: 244
>unnamed protein product [Arabidopsis thaliana]
Sequence ID: CAA0383935.1 Length: 244 >unnamed protein product [Arabidopsis thaliana]
Sequence ID: VYS58851.1 Length: 244
Range 1: 39 to 121

Score:59.7 bits(143), Expect:2e-09,
Method:Compositional matrix adjust.,
Identities:45/87(52%), Positives:52/87(59%), Gaps:7/87(8%)

Query  697  EKQKDEAELLGVEL--FMSEAMFLLSDDIYSMLLFFYKIILKYTGNNERKNPTQ-VEKIL  527
            E QKDEA  LGVEL  F++E M LLSD++ SMLLF   ++     N E    T  V   L
Sbjct  39   EMQKDEAIRLGVELSLFVAETMVLLSDNLRSMLLFCLWVL----TNAECTYFTDPVVGRL  94

Query  526  INVRLYVFETNIKPKNGVSQADGKSTR  446
              V LYV ET IKPKNGV  AD KS +
Sbjct  95   HCVMLYVLETYIKPKNGVYLADVKSNQ  121


>hypothetical protein (DUF1184) [Arabidopsis thaliana]
Sequence ID: NP_001325896.1 Length: 256
>hypothetical protein (DUF1184) [Arabidopsis thaliana]
Sequence ID: NP_001325897.1 Length: 256 >hypothetical protein (DUF1184) [Arabidopsis thaliana]
Sequence ID: ANM63827.1 Length: 256 >hypothetical protein (DUF1184) [Arabidopsis thaliana]
Sequence ID: ANM63828.1 Length: 256 >unnamed protein product [Arabidopsis thaliana]
Sequence ID: BAB02536.1 Length: 256
Range 1: 51 to 133

Score:60.1 bits(144), Expect:2e-09,
Method:Compositional matrix adjust.,
Identities:45/87(52%), Positives:52/87(59%), Gaps:7/87(8%)

Query  697  EKQKDEAELLGVEL--FMSEAMFLLSDDIYSMLLFFYKIILKYTGNNERKNPTQ-VEKIL  527
            E QKDEA  LGVEL  F++E M LLSD++ SMLLF   ++     N E    T  V   L
Sbjct  51   EMQKDEAIRLGVELSLFVAETMVLLSDNLRSMLLFCLWVL----TNAECTYFTDPVVGRL  106

Query  526  INVRLYVFETNIKPKNGVSQADGKSTR  446
              V LYV ET IKPKNGV  AD KS +
Sbjct  107  HCVMLYVLETYIKPKNGVYLADVKSNQ  133


Query #33: XLOC_006015 Query ID: lcl|Query_76567 Length: 899

No significant similarity found.


Query #34: XLOC_006026 Query ID: lcl|Query_76568 Length: 763

Sequences producing significant alignments:
                                                                  Max    Total Query   E   Per.
Description                                                       Score  Score cover Value Ident  Accession
F15O4.6 [Arabidopsis thaliana]                                    224    224   52%   2e-74 94.03  AAF79344.1
phosphoenolpyruvate carboxykinase 1 [Arabidopsis thaliana]        97.1   154   52%   3e-22 55.88  NP_195500.1
PEPCK [Arabidopsis thaliana]                                      97.1   154   52%   3e-22 55.88  OAP00328.1
phosphoenolpyruvate carboxykinase 2 [Arabidopsis thaliana]        86.3   143   52%   2e-18 50.00  NP_001331431.1
phosphoenolpyruvate carboxykinase 2 [Arabidopsis thaliana]        86.3   143   52%   2e-18 50.00  NP_001331433.1
unnamed protein product [Arabidopsis thaliana]                    86.3   143   52%   2e-18 50.00  VYS71520.1
PEPCK [Arabidopsis thaliana]                                      86.3   143   52%   2e-18 50.00  OAO90231.1
unnamed protein product [Arabidopsis thaliana]                    86.3   143   52%   2e-18 50.00  CAA0412274.1
phosphoenolpyruvate carboxykinase 2 [Arabidopsis thaliana]        85.9   142   52%   2e-18 50.00  NP_001331432.1
phosphoenolpyruvate carboxykinase (ATP) (EC 4.1.1.49)...          85.9   142   52%   2e-18 50.00  BAB10675.1
unnamed protein product [Arabidopsis thaliana]                    65.5   94.7  48%   3e-13 40.57  CAA0361730.1
phosphoenolpyruvate carboxykinase-like protein [Arabidopsis...    55.1   55.1  12%   5e-08 78.79  BAD94487.1


Alignments:
>F15O4.6 [Arabidopsis thaliana]
Sequence ID: AAF79344.1 Length: 155
Range 1: 30 to 155

Score:224 bits(572), Expect:2e-74,
Method:Compositional matrix adjust.,
Identities:126/134(94%), Positives:126/134(94%), Gaps:8/134(5%)

Query  107  LYEQAIEYEKGSLITSNGALTMLSGAKTDREISVLLEMRLLRMSFGDYRIFILYFGLLIf  286
            LYEQAIEYEKGSLITSNGALTMLSGAKTDREISVLLEMRLLRMSFGDYRIFILYFGLLIF
Sbjct  30   LYEQAIEYEKGSLITSNGALTMLSGAKTDREISVLLEMRLLRMSFGDYRIFILYFGLLIF  89

Query  287  vfdfdfdfKGLSNIEIDEHTFMVVNRERAVDYLNSLEKVYLMPKS*ILSHHSGCNMGKDG  466
            VFDFDFDFKGLSNIEIDEHTFMVVNRERAVDYLNSLEK        ILSHHSGCNMGKDG
Sbjct  90   VFDFDFDFKGLSNIEIDEHTFMVVNRERAVDYLNSLEK--------ILSHHSGCNMGKDG  141

Query  467  DVALFFELLGIATF  508
            DVALFFELLGIATF
Sbjct  142  DVALFFELLGIATF  155


>phosphoenolpyruvate carboxykinase 1 [Arabidopsis thaliana]
Sequence ID: NP_195500.1 Length: 671
>RecName: Full=Phosphoenolpyruvate carboxykinase (ATP); Short=PEP carboxykinase; Short=PEPCK [Arabidopsis thaliana]
Sequence ID: Q9T074.1 Length: 671 >AT4g37870/T28I19_150 [Arabidopsis thaliana]
Sequence ID: AAK50062.1 Length: 671 >AT4g37870/T28I19_150 [Arabidopsis thaliana]
Sequence ID: AAL77736.1 Length: 671 >phosphoenolpyruvate carboxykinase 1 [Arabidopsis thaliana]
Sequence ID: AEE86847.1 Length: 671 >unnamed protein product [Arabidopsis thaliana]
Sequence ID: CAA0397843.1 Length: 671 >phosphoenolpyruvate carboxykinase (ATP)-like protein [Arabidopsis thaliana]
Sequence ID: CAB38935.1 Length: 671
Range 1: 159 to 241

Score:97.1 bits(240), Expect:3e-22,
Method:Compositional matrix adjust.,
Identities:57/102(56%), Positives:68/102(66%), Gaps:19/102(18%)

Query  104  ELYEQAIEYEKGSLITSNGALTMLSGAKTDREISVLLEMRLLRMSFGDYRIFILYFGLLI  283
            ELYEQAI+YEKGS ITSNGAL  LSGAKT R      + R++R +  +     L++G
Sbjct  159  ELYEQAIKYEKGSFITSNGALATLSGAKTGR---APRDKRVVRDATTEDE---LWWG---  209

Query  284  fvfdfdfdfKGLSNIEIDEHTFMVVNRERAVDYLNSLEKVYL  409
                     KG  NIE+DEHTFM VNRERAVDYLNSLEKV++
Sbjct  210  ---------KGSPNIEMDEHTFM-VNRERAVDYLNSLEKVFV  241


Range 2: 342 to 374

Score:57.0 bits(136), Expect:1e-08,
Method:Compositional matrix adjust.,
Identities:26/33(79%), Positives:26/33(78%), Gaps:0/33(0%)

Query  404  YLMPKS*ILSHHSGCNMGKDGDVALFFELLGIA  502
            YLMPK  ILS HSGCNMGKDGDVALFF L G
Sbjct  342  YLMPKRRILSLHSGCNMGKDGDVALFFGLSGTG  374


>PEPCK [Arabidopsis thaliana]
Sequence ID: OAP00328.1 Length: 671
Range 1: 159 to 241

Score:97.1 bits(240), Expect:3e-22,
Method:Compositional matrix adjust.,
Identities:57/102(56%), Positives:68/102(66%), Gaps:19/102(18%)

Query  104  ELYEQAIEYEKGSLITSNGALTMLSGAKTDREISVLLEMRLLRMSFGDYRIFILYFGLLI  283
            ELYEQAI+YEKGS ITSNGAL  LSGAKT R      + R++R +  +     L++G
Sbjct  159  ELYEQAIKYEKGSFITSNGALATLSGAKTGR---APRDKRVVRDATTEDE---LWWG---  209

Query  284  fvfdfdfdfKGLSNIEIDEHTFMVVNRERAVDYLNSLEKVYL  409
                     KG  NIE+DEHTFM VNRERAVDYLNSLEKV++
Sbjct  210  ---------KGSPNIEMDEHTFM-VNRERAVDYLNSLEKVFV  241


Range 2: 342 to 374

Score:57.0 bits(136), Expect:1e-08,
Method:Compositional matrix adjust.,
Identities:26/33(79%), Positives:26/33(78%), Gaps:0/33(0%)

Query  404  YLMPKS*ILSHHSGCNMGKDGDVALFFELLGIA  502
            YLMPK  ILS HSGCNMGKDGDVALFF L G
Sbjct  342  YLMPKRRILSLHSGCNMGKDGDVALFFGLSGTG  374


>phosphoenolpyruvate carboxykinase 2 [Arabidopsis thaliana]
Sequence ID: NP_001331431.1 Length: 577
>phosphoenolpyruvate carboxykinase 2 [Arabidopsis thaliana]
Sequence ID: ANM69777.1 Length: 577
Range 1: 158 to 240

Score:86.3 bits(212), Expect:2e-18,
Method:Compositional matrix adjust.,
Identities:51/102(50%), Positives:65/102(63%), Gaps:19/102(18%)

Query  104  ELYEQAIEYEKGSLITSNGALTMLSGAKTDREISVLLEMRLLRMSFGDYRIFILYFGLLI  283
            ELYEQAI++EKGS +TS GAL  LSGAKT R      + R+++    D     L++G
Sbjct  158  ELYEQAIKFEKGSFVTSTGALATLSGAKTGRSPK---DKRVVK---DDTTEAELWWG---  208

Query  284  fvfdfdfdfKGLSNIEIDEHTFMVVNRERAVDYLNSLEKVYL  409
                     KG  NIE+DE TF+ VNRERAVDYLNSL+KV++
Sbjct  209  ---------KGSPNIEMDEKTFL-VNRERAVDYLNSLDKVFV  240


Range 2: 341 to 373

Score:57.4 bits(137), Expect:1e-08,
Method:Compositional matrix adjust.,
Identities:26/33(79%), Positives:26/33(78%), Gaps:0/33(0%)

Query  404  YLMPKS*ILSHHSGCNMGKDGDVALFFELLGIA  502
            YLMPK  ILS HSGCNMGKDGDVALFF L G
Sbjct  341  YLMPKRKILSLHSGCNMGKDGDVALFFGLSGTG  373


>phosphoenolpyruvate carboxykinase 2 [Arabidopsis thaliana]
Sequence ID: NP_001331433.1 Length: 670
>phosphoenolpyruvate carboxykinase 2 [Arabidopsis thaliana]
Sequence ID: NP_680468.1 Length: 670 >At5g65690 [Arabidopsis thaliana]
Sequence ID: ACI49792.1 Length: 670 >phosphoenolpyruvate carboxykinase 2 [Arabidopsis thaliana]
Sequence ID: AED98087.1 Length: 670 >phosphoenolpyruvate carboxykinase 2 [Arabidopsis thaliana]
Sequence ID: ANM69779.1 Length: 670
Range 1: 158 to 240

Score:86.3 bits(212), Expect:2e-18,
Method:Compositional matrix adjust.,
Identities:51/102(50%), Positives:65/102(63%), Gaps:19/102(18%)

Query  104  ELYEQAIEYEKGSLITSNGALTMLSGAKTDREISVLLEMRLLRMSFGDYRIFILYFGLLI  283
            ELYEQAI++EKGS +TS GAL  LSGAKT R      + R+++    D     L++G
Sbjct  158  ELYEQAIKFEKGSFVTSTGALATLSGAKTGRSPK---DKRVVK---DDTTEAELWWG---  208

Query  284  fvfdfdfdfKGLSNIEIDEHTFMVVNRERAVDYLNSLEKVYL  409
                     KG  NIE+DE TF+ VNRERAVDYLNSL+KV++
Sbjct  209  ---------KGSPNIEMDEKTFL-VNRERAVDYLNSLDKVFV  240


Range 2: 341 to 373

Score:57.4 bits(137), Expect:1e-08,
Method:Compositional matrix adjust.,
Identities:26/33(79%), Positives:26/33(78%), Gaps:0/33(0%)

Query  404  YLMPKS*ILSHHSGCNMGKDGDVALFFELLGIA  502
            YLMPK  ILS HSGCNMGKDGDVALFF L G
Sbjct  341  YLMPKRKILSLHSGCNMGKDGDVALFFGLSGTG  373


>unnamed protein product [Arabidopsis thaliana]
Sequence ID: VYS71520.1 Length: 670
Range 1: 158 to 240

Score:86.3 bits(212), Expect:2e-18,
Method:Compositional matrix adjust.,
Identities:51/102(50%), Positives:65/102(63%), Gaps:19/102(18%)

Query  104  ELYEQAIEYEKGSLITSNGALTMLSGAKTDREISVLLEMRLLRMSFGDYRIFILYFGLLI  283
            ELYEQAI++EKGS +TS GAL  LSGAKT R      + R+++    D     L++G
Sbjct  158  ELYEQAIKFEKGSFVTSTGALATLSGAKTGRSPK---DKRVVK---DDTTEAELWWG---  208

Query  284  fvfdfdfdfKGLSNIEIDEHTFMVVNRERAVDYLNSLEKVYL  409
                     KG  NIE+DE TF+ VNRERAVDYLNSL+KV++
Sbjct  209  ---------KGSPNIEMDEKTFL-VNRERAVDYLNSLDKVFV  240


Range 2: 341 to 373

Score:57.4 bits(137), Expect:1e-08,
Method:Compositional matrix adjust.,
Identities:26/33(79%), Positives:26/33(78%), Gaps:0/33(0%)

Query  404  YLMPKS*ILSHHSGCNMGKDGDVALFFELLGIA  502
            YLMPK  ILS HSGCNMGKDGDVALFF L G
Sbjct  341  YLMPKRKILSLHSGCNMGKDGDVALFFGLSGTG  373


>PEPCK [Arabidopsis thaliana]
Sequence ID: OAO90231.1 Length: 670
Range 1: 158 to 240

Score:86.3 bits(212), Expect:2e-18,
Method:Compositional matrix adjust.,
Identities:51/102(50%), Positives:65/102(63%), Gaps:19/102(18%)

Query  104  ELYEQAIEYEKGSLITSNGALTMLSGAKTDREISVLLEMRLLRMSFGDYRIFILYFGLLI  283
            ELYEQAI++EKGS +TS GAL  LSGAKT R      + R+++    D     L++G
Sbjct  158  ELYEQAIKFEKGSFVTSTGALATLSGAKTGRSPK---DKRVVK---DDTTEAELWWG---  208

Query  284  fvfdfdfdfKGLSNIEIDEHTFMVVNRERAVDYLNSLEKVYL  409
                     KG  NIE+DE TF+ VNRERAVDYLNSL+KV++
Sbjct  209  ---------KGSPNIEMDEKTFL-VNRERAVDYLNSLDKVFV  240


Range 2: 341 to 373

Score:57.4 bits(137), Expect:1e-08,
Method:Compositional matrix adjust.,
Identities:26/33(79%), Positives:26/33(78%), Gaps:0/33(0%)

Query  404  YLMPKS*ILSHHSGCNMGKDGDVALFFELLGIA  502
            YLMPK  ILS HSGCNMGKDGDVALFF L G
Sbjct  341  YLMPKRKILSLHSGCNMGKDGDVALFFGLSGTG  373


>unnamed protein product [Arabidopsis thaliana]
Sequence ID: CAA0412274.1 Length: 670
Range 1: 158 to 240

Score:86.3 bits(212), Expect:2e-18,
Method:Compositional matrix adjust.,
Identities:51/102(50%), Positives:65/102(63%), Gaps:19/102(18%)

Query  104  ELYEQAIEYEKGSLITSNGALTMLSGAKTDREISVLLEMRLLRMSFGDYRIFILYFGLLI  283
            ELYEQAI++EKGS +TS GAL  LSGAKT R      + R+++    D     L++G
Sbjct  158  ELYEQAIKFEKGSFVTSTGALATLSGAKTGRSPK---DKRVVK---DDTTEAELWWG---  208

Query  284  fvfdfdfdfKGLSNIEIDEHTFMVVNRERAVDYLNSLEKVYL  409
                     KG  NIE+DE TF+ VNRERAVDYLNSL+KV++
Sbjct  209  ---------KGSPNIEMDEKTFL-VNRERAVDYLNSLDKVFV  240


Range 2: 341 to 373

Score:57.4 bits(137), Expect:1e-08,
Method:Compositional matrix adjust.,
Identities:26/33(79%), Positives:26/33(78%), Gaps:0/33(0%)

Query  404  YLMPKS*ILSHHSGCNMGKDGDVALFFELLGIA  502
            YLMPK  ILS HSGCNMGKDGDVALFF L G
Sbjct  341  YLMPKRKILSLHSGCNMGKDGDVALFFGLSGTG  373


>phosphoenolpyruvate carboxykinase 2 [Arabidopsis thaliana]
Sequence ID: NP_001331432.1 Length: 701
>phosphoenolpyruvate carboxykinase 2 [Arabidopsis thaliana]
Sequence ID: ANM69778.1 Length: 701
Range 1: 158 to 240

Score:85.9 bits(211), Expect:2e-18,
Method:Compositional matrix adjust.,
Identities:51/102(50%), Positives:65/102(63%), Gaps:19/102(18%)

Query  104  ELYEQAIEYEKGSLITSNGALTMLSGAKTDREISVLLEMRLLRMSFGDYRIFILYFGLLI  283
            ELYEQAI++EKGS +TS GAL  LSGAKT R      + R+++    D     L++G
Sbjct  158  ELYEQAIKFEKGSFVTSTGALATLSGAKTGRSPK---DKRVVK---DDTTEAELWWG---  208

Query  284  fvfdfdfdfKGLSNIEIDEHTFMVVNRERAVDYLNSLEKVYL  409
                     KG  NIE+DE TF+ VNRERAVDYLNSL+KV++
Sbjct  209  ---------KGSPNIEMDEKTFL-VNRERAVDYLNSLDKVFV  240


Range 2: 341 to 373

Score:57.0 bits(136), Expect:1e-08,
Method:Compositional matrix adjust.,
Identities:26/33(79%), Positives:26/33(78%), Gaps:0/33(0%)

Query  404  YLMPKS*ILSHHSGCNMGKDGDVALFFELLGIA  502
            YLMPK  ILS HSGCNMGKDGDVALFF L G
Sbjct  341  YLMPKRKILSLHSGCNMGKDGDVALFFGLSGTG  373


>phosphoenolpyruvate carboxykinase (ATP) (EC 4.1.1.49) [Arabidopsis thaliana]
Sequence ID: BAB10675.1 Length: 628
>phosphoenolpyruvate carboxykinase (ATP) - like protein [Arabidopsis thaliana]
Sequence ID: CAA16690.1 Length: 628
Range 1: 120 to 202

Score:85.9 bits(211), Expect:2e-18,
Method:Compositional matrix adjust.,
Identities:51/102(50%), Positives:65/102(63%), Gaps:19/102(18%)

Query  104  ELYEQAIEYEKGSLITSNGALTMLSGAKTDREISVLLEMRLLRMSFGDYRIFILYFGLLI  283
            ELYEQAI++EKGS +TS GAL  LSGAKT R      + R+++    D     L++G
Sbjct  120  ELYEQAIKFEKGSFVTSTGALATLSGAKTGRSPK---DKRVVK---DDTTEAELWWG---  170

Query  284  fvfdfdfdfKGLSNIEIDEHTFMVVNRERAVDYLNSLEKVYL  409
                     KG  NIE+DE TF+ VNRERAVDYLNSL+KV++
Sbjct  171  ---------KGSPNIEMDEKTFL-VNRERAVDYLNSLDKVFV  202


Range 2: 303 to 335

Score:57.0 bits(136), Expect:1e-08,
Method:Compositional matrix adjust.,
Identities:26/33(79%), Positives:26/33(78%), Gaps:0/33(0%)

Query  404  YLMPKS*ILSHHSGCNMGKDGDVALFFELLGIA  502
            YLMPK  ILS HSGCNMGKDGDVALFF L G
Sbjct  303  YLMPKRKILSLHSGCNMGKDGDVALFFGLSGTG  335


>unnamed protein product [Arabidopsis thaliana]
Sequence ID: CAA0361730.1 Length: 551
Range 1: 363 to 433

Score:65.5 bits(158), Expect:3e-13,
Method:Compositional matrix adjust.,
Identities:43/106(41%), Positives:55/106(51%), Gaps:35/106(33%)

Query  92   FI*QELYEQAIEYEKGSLITSNGALTMLSGAKTDREISVLLEMRLLRMSFGDYRIFILYF  271
            F+  ELYEQAI+YEKGS ITSNGAL  LSGAKT R  +     R++R++   Y    L++
Sbjct  363  FLFVELYEQAIKYEKGSFITSNGALATLSGAKTGRAPT---HKRIVRVAATKYE---LWW  416

Query  272  GLLIfvfdfdfdfKGLSNIEIDEHTFMVVNRERAVDYLNSLEKVYL  409
            G                             +ERAVDYLN LE V++
Sbjct  417  G-----------------------------KERAVDYLNFLENVFV  433


Range 2: 465 to 481

Score:29.3 bits(64), Expect:3e-13,
Method:Compositional matrix adjust.,
Identities:11/17(65%), Positives:13/17(76%), Gaps:0/17(0%)

Query  463  WRCCPLL*TFRYCNFLS  513
            WR CPLL TF YCN ++
Sbjct  465  WRYCPLLWTFTYCNLIT  481


>phosphoenolpyruvate carboxykinase-like protein [Arabidopsis thaliana]
Sequence ID: BAD94487.1 Length: 455
Range 1: 126 to 158

Score:55.1 bits(131), Expect:5e-08,
Method:Compositional matrix adjust.,
Identities:26/33(79%), Positives:26/33(78%), Gaps:0/33(0%)

Query  404  YLMPKS*ILSHHSGCNMGKDGDVALFFELLGIA  502
            YLMPK  ILS HSGCNMGKDGDVALFF L G
Sbjct  126  YLMPKRRILSLHSGCNMGKDGDVALFFGLSGTG  158


Query #35: XLOC_006033 Query ID: lcl|Query_76569 Length: 836

Sequences producing significant alignments:
                                                                  Max    Total Query   E   Per.
Description                                                       Score  Score cover Value Ident  Accession
unknown protein [Arabidopsis thaliana]                            47.8   80.9  19%   4e-09 66.67  AAD17400.1


Alignments:
>unknown protein [Arabidopsis thaliana]
Sequence ID: AAD17400.1 Length: 524
Range 1: 434 to 472

Score:47.8 bits(112), Expect:4e-09,
Method:Compositional matrix adjust.,
Identities:26/39(67%), Positives:27/39(69%), Gaps:0/39(0%)

Query  319  GPFTISRPLVDFSGPTIPTVFTSFFQDSTVPSSKLLRRL  435
            G FT SRPLVD SGP IPTV   F +DSTVP SK  R L
Sbjct  434  GSFTGSRPLVDTSGPAIPTVSACFCRDSTVPPSKSRRHL  472


Range 2: 419 to 436

Score:33.1 bits(74), Expect:4e-09,
Method:Compositional matrix adjust.,
Identities:13/18(72%), Positives:16/18(88%), Gaps:0/18(0%)

Query  273  TQVQQTDLGRDPPRFGSF  326
             Q+QQ DLG+DPP+FGSF
Sbjct  419  VQLQQMDLGQDPPKFGSF  436


Query #36: XLOC_006081 Query ID: lcl|Query_76570 Length: 392

Sequences producing significant alignments:
                                                                  Max    Total Query   E   Per.
Description                                                       Score  Score cover Value Ident  Accession
unnamed protein product [Arabidopsis thaliana]                    93.6   93.6  38%   1e-25 88.00  CAA0269652.1
unnamed protein product [Arabidopsis thaliana]                    93.2   93.2  38%   2e-25 88.00  VYS48124.1
hypothetical protein AXX17_AT1G37330 [Arabidopsis thaliana]       92.4   92.4  38%   3e-25 90.00  OAP11900.1
transmembrane protein [Arabidopsis thaliana]                      91.3   91.3  38%   9e-25 86.00  NP_174875.1
unknown [Arabidopsis thaliana]                                    55.1   55.1  37%   1e-10 53.06  ABK28353.1
transmembrane protein [Arabidopsis thaliana]                      55.1   55.1  37%   1e-10 53.06  NP_001077669.1
hypothetical protein AXX17_AT1G37320 [Arabidopsis thaliana]       53.1   53.1  36%   8e-10 56.25  OAP14890.1


Alignments:
>unnamed protein product [Arabidopsis thaliana]
Sequence ID: CAA0269652.1 Length: 75
Range 1: 25 to 74

Score:93.6 bits(231), Expect:1e-25,
Method:Compositional matrix adjust.,
Identities:44/50(88%), Positives:47/50(94%), Gaps:0/50(0%)

Query  137  ARVQYASPSQRKKIGKEVWEQTLLSEIKIGASGSNSGRDPSCNNSCKPNR  286
            ARVQY SPSQRKKIGKEVW+QTLL ++KIGASGSNSGR PSCNNSCKPNR
Sbjct  25   ARVQYESPSQRKKIGKEVWDQTLLRDLKIGASGSNSGRAPSCNNSCKPNR  74


>unnamed protein product [Arabidopsis thaliana]
Sequence ID: VYS48124.1 Length: 75
Range 1: 25 to 74

Score:93.2 bits(230), Expect:2e-25,
Method:Compositional matrix adjust.,
Identities:44/50(88%), Positives:47/50(94%), Gaps:0/50(0%)

Query  137  ARVQYASPSQRKKIGKEVWEQTLLSEIKIGASGSNSGRDPSCNNSCKPNR  286
            ARVQY SPSQRKKIGKEVW+QTLL ++KIGASGSNSGR PSCNNSCKPNR
Sbjct  25   ARVQYESPSQRKKIGKEVWDQTLLRDLKIGASGSNSGRAPSCNNSCKPNR  74


>hypothetical protein AXX17_AT1G37330 [Arabidopsis thaliana]
Sequence ID: OAP11900.1 Length: 75
Range 1: 25 to 74

Score:92.4 bits(228), Expect:3e-25,
Method:Compositional matrix adjust.,
Identities:45/50(90%), Positives:46/50(92%), Gaps:0/50(0%)

Query  137  ARVQYASPSQRKKIGKEVWEQTLLSEIKIGASGSNSGRDPSCNNSCKPNR  286
            ARVQY SPSQRKKI KEVW+QTLL EIKIGASGSNSGR PSCNNSCKPNR
Sbjct  25   ARVQYESPSQRKKILKEVWDQTLLREIKIGASGSNSGRAPSCNNSCKPNR  74


>transmembrane protein [Arabidopsis thaliana]
Sequence ID: NP_174875.1 Length: 75
>unknown protein [Arabidopsis thaliana]
Sequence ID: AAG51257.1 Length: 75 >transmembrane protein [Arabidopsis thaliana]
Sequence ID: AEE31869.1 Length: 75
Range 1: 25 to 74

Score:91.3 bits(225), Expect:9e-25,
Method:Compositional matrix adjust.,
Identities:43/50(86%), Positives:46/50(92%), Gaps:0/50(0%)

Query  137  ARVQYASPSQRKKIGKEVWEQTLLSEIKIGASGSNSGRDPSCNNSCKPNR  286
            ARVQY SPSQR KIGKEVW+QTLL ++KIGASGSNSGR PSCNNSCKPNR
Sbjct  25   ARVQYESPSQRNKIGKEVWDQTLLRDLKIGASGSNSGRAPSCNNSCKPNR  74


>unknown, partial [Arabidopsis thaliana]
Sequence ID: ABK28353.1 Length: 75
Range 1: 25 to 73

Score:55.1 bits(131), Expect:1e-10,
Method:Compositional matrix adjust.,
Identities:26/49(53%), Positives:33/49(67%), Gaps:0/49(0%)

Query  137  ARVQYASPSQRKKIGKEVWEQTLLSEIKIGASGSNSGRDPSCNNSCKPN  283
            ARVQY SP  RK+IGK VW+Q + +EIKI   GS+S R  S ++   PN
Sbjct  25   ARVQYGSPVSRKEIGKGVWDQKVFNEIKIAVGGSDSVRAHSKDHKSNPN  73


>transmembrane protein [Arabidopsis thaliana]
Sequence ID: NP_001077669.1 Length: 74
>unknown protein [Arabidopsis thaliana]
Sequence ID: ABF59257.1 Length: 74 >transmembrane protein [Arabidopsis thaliana]
Sequence ID: AEE31868.1 Length: 74 >hypothetical protein AXX17_AT1G37300 [Arabidopsis thaliana]
Sequence ID: OAP12855.1 Length: 74 >unnamed protein product [Arabidopsis thaliana]
Sequence ID: CAA0269551.1 Length: 74 >unnamed protein product [Arabidopsis thaliana]
Sequence ID: VYS48121.1 Length: 74
Range 1: 25 to 73

Score:55.1 bits(131), Expect:1e-10,
Method:Compositional matrix adjust.,
Identities:26/49(53%), Positives:33/49(67%), Gaps:0/49(0%)

Query  137  ARVQYASPSQRKKIGKEVWEQTLLSEIKIGASGSNSGRDPSCNNSCKPN  283
            ARVQY SP  RK+IGK VW+Q + +EIKI   GS+S R  S ++   PN
Sbjct  25   ARVQYGSPVSRKEIGKGVWDQKVFNEIKIAVGGSDSVRAHSKDHKSNPN  73


>hypothetical protein AXX17_AT1G37320 [Arabidopsis thaliana]
Sequence ID: OAP14890.1 Length: 71
Range 1: 25 to 70

Score:53.1 bits(126), Expect:8e-10,
Method:Compositional matrix adjust.,
Identities:27/48(56%), Positives:32/48(66%), Gaps:2/48(4%)

Query  137  ARVQYASPSQRKKIGKEVWEQTLLSEIKIGASGSNSGRDPSCNNSCKP  280
            ARVQY SP  RK+IGKEVW+Q +  EIKI   GS+S   PS +  C P
Sbjct  25   ARVQYRSPGSRKEIGKEVWDQKIFHEIKIAVRGSDS--VPSHSRGCCP  70


Query #37: XLOC_006083 Query ID: lcl|Query_76571 Length: 880

Sequences producing significant alignments:
                                                                  Max    Total Query   E   Per.
Description                                                       Score  Score cover Value Ident  Accession
Serine/Threonine-kinase [Arabidopsis thaliana]                    116    116   18%   8e-33 100.00 NP_001322227.1
unnamed protein product [Arabidopsis thaliana]                    86.7   144   32%   8e-28 72.41  CAA0397273.1
AME3 [Arabidopsis thaliana]                                       87.8   87.8  19%   2e-21 72.41  OAO98285.1
AME3 [Arabidopsis thaliana]                                       86.7   119   24%   1e-20 72.41  OAO98284.1
protein kinase [Arabidopsis thaliana]                             87.0   118   24%   3e-20 72.41  BAF01108.1
Protein kinase superfamily protein [Arabidopsis thaliana]         87.0   118   24%   3e-20 72.41  NP_194992.1
Protein kinase superfamily protein [Arabidopsis thaliana]         86.7   118   24%   4e-20 72.41  NP_974666.1
Protein kinase superfamily protein [Arabidopsis thaliana]         87.0   87.0  19%   1e-18 72.41  NP_001031774.1
protein kinase [Arabidopsis thaliana]                             87.0   87.0  19%   1e-18 72.41  AAA57119.1


Alignments:
>Serine/Threonine-kinase [Arabidopsis thaliana]
Sequence ID: NP_001322227.1 Length: 55
>Serine/Threonine-kinase [Arabidopsis thaliana]
Sequence ID: ANM59905.1 Length: 55 >unnamed protein product [Arabidopsis thaliana]
Sequence ID: CAA0269588.1 Length: 55 >unnamed protein product [Arabidopsis thaliana]
Sequence ID: VYS48122.1 Length: 55
Range 1: 1 to 55

Score:116 bits(291), Expect:8e-33,
Method:Compositional matrix adjust.,
Identities:55/55(100%), Positives:55/55(100%), Gaps:0/55(0%)

Query  614  MTWDESLPEQEAKGVVIKGHGSNMMILSPPLREDDCDGHYVFSLRDNLISIIKVN  778
            MTWDESLPEQEAKGVVIKGHGSNMMILSPPLREDDCDGHYVFSLRDNLISIIKVN
Sbjct  1    MTWDESLPEQEAKGVVIKGHGSNMMILSPPLREDDCDGHYVFSLRDNLISIIKVN  55


>unnamed protein product [Arabidopsis thaliana]
Sequence ID: CAA0397273.1 Length: 434
Range 1: 50 to 107

Score:86.7 bits(213), Expect:8e-28,
Method:Compositional matrix adjust.,
Identities:42/58(72%), Positives:46/58(79%), Gaps:0/58(0%)

Query  602  KRPRMTWDESLPEQEAKGVVIKGHGSNMMILSPPLREDDCDGHYVFSLRDNLISIIKV  775
            KRPRMTWDE+  E EAK  VIKGHGS+  ILSPPLR+DD DGHYVFSLRDNL    K+
Sbjct  50   KRPRMTWDEAPAEPEAKRAVIKGHGSDGRILSPPLRDDDRDGHYVFSLRDNLTPRYKI  107


Range 2: 11 to 49

Score:57.4 bits(137), Expect:8e-28,
Method:Compositional matrix adjust.,
Identities:27/39(69%), Positives:30/39(76%), Gaps:0/39(0%)

Query  487  PSFSYRLSIRFLFGCVGD*FSVSVMIANGFENMDKERVK  603
            P   + + IRF  GCVGD FS SVMIANGFENMDKERV+
Sbjct  11   PQGIWPIGIRFRSGCVGDRFSASVMIANGFENMDKERVR  49


>AME3 [Arabidopsis thaliana]
Sequence ID: OAO98285.1 Length: 80
Range 1: 16 to 73

Score:87.8 bits(216), Expect:2e-21,
Method:Compositional matrix adjust.,
Identities:42/58(72%), Positives:46/58(79%), Gaps:0/58(0%)

Query  602  KRPRMTWDESLPEQEAKGVVIKGHGSNMMILSPPLREDDCDGHYVFSLRDNLISIIKV  775
            KRPRMTWDE+  E EAK  VIKGHGS+  ILSPPLR+DD DGHYVFSLRDNL    K+
Sbjct  16   KRPRMTWDEAPAEPEAKRAVIKGHGSDGRILSPPLRDDDRDGHYVFSLRDNLTPRYKI  73


>AME3 [Arabidopsis thaliana]
Sequence ID: OAO98284.1 Length: 173
Range 1: 16 to 73

Score:86.7 bits(213), Expect:1e-20,
Method:Compositional matrix adjust.,
Identities:42/58(72%), Positives:46/58(79%), Gaps:0/58(0%)

Query  602  KRPRMTWDESLPEQEAKGVVIKGHGSNMMILSPPLREDDCDGHYVFSLRDNLISIIKV  775
            KRPRMTWDE+  E EAK  VIKGHGS+  ILSPPLR+DD DGHYVFSLRDNL    K+
Sbjct  16   KRPRMTWDEAPAEPEAKRAVIKGHGSDGRILSPPLRDDDRDGHYVFSLRDNLTPRYKI  73


Range 2: 1 to 15

Score:33.1 bits(74), Expect:1e-20,
Method:Compositional matrix adjust.,
Identities:14/15(93%), Positives:15/15(100%), Gaps:0/15(0%)

Query  559  MIANGFENMDKERVK  603
            MIANGFENMDKERV+
Sbjct  1    MIANGFENMDKERVR  15


>protein kinase [Arabidopsis thaliana]
Sequence ID: BAF01108.1 Length: 400
Range 1: 16 to 73

Score:87.0 bits(214), Expect:3e-20,
Method:Compositional matrix adjust.,
Identities:42/58(72%), Positives:46/58(79%), Gaps:0/58(0%)

Query  602  KRPRMTWDESLPEQEAKGVVIKGHGSNMMILSPPLREDDCDGHYVFSLRDNLISIIKV  775
            KRPRMTWDE+  E EAK  VIKGHGS+  ILSPPLR+DD DGHYVFSLRDNL    K+
Sbjct  16   KRPRMTWDEAPAEPEAKRAVIKGHGSDGRILSPPLRDDDRDGHYVFSLRDNLTPRYKI  73


Range 2: 1 to 15

Score:31.6 bits(70), Expect:3e-20,
Method:Compositional matrix adjust.,
Identities:13/15(87%), Positives:15/15(100%), Gaps:0/15(0%)

Query  559  MIANGFENMDKERVK  603
            MIANGFE+MDKERV+
Sbjct  1    MIANGFESMDKERVR  15


>Protein kinase superfamily protein [Arabidopsis thaliana]
Sequence ID: NP_194992.1 Length: 400
>RecName: Full=Serine/threonine-protein kinase AFC3 [Arabidopsis thaliana]
Sequence ID: P51568.2 Length: 400 >At4g32660 [Arabidopsis thaliana]
Sequence ID: ABM06046.1 Length: 400 >Protein kinase superfamily protein [Arabidopsis thaliana]
Sequence ID: AEE86100.1 Length: 400 >protein kinase AME3 [Arabidopsis thaliana]
Sequence ID: CAA18595.1 Length: 400 >protein kinase AME3 [Arabidopsis thaliana]
Sequence ID: CAB79983.1 Length: 400 >unnamed protein product [Arabidopsis thaliana]
Sequence ID: VYS64675.1 Length: 400
Range 1: 16 to 73

Score:87.0 bits(214), Expect:3e-20,
Method:Compositional matrix adjust.,
Identities:42/58(72%), Positives:46/58(79%), Gaps:0/58(0%)

Query  602  KRPRMTWDESLPEQEAKGVVIKGHGSNMMILSPPLREDDCDGHYVFSLRDNLISIIKV  775
            KRPRMTWDE+  E EAK  VIKGHGS+  ILSPPLR+DD DGHYVFSLRDNL    K+
Sbjct  16   KRPRMTWDEAPAEPEAKRAVIKGHGSDGRILSPPLRDDDRDGHYVFSLRDNLTPRYKI  73


Range 2: 1 to 15

Score:31.6 bits(70), Expect:3e-20,
Method:Compositional matrix adjust.,
Identities:13/15(87%), Positives:15/15(100%), Gaps:0/15(0%)

Query  559  MIANGFENMDKERVK  603
            MIANGFE+MDKERV+
Sbjct  1    MIANGFESMDKERVR  15


>Protein kinase superfamily protein [Arabidopsis thaliana]
Sequence ID: NP_974666.1 Length: 356
>Protein kinase superfamily protein [Arabidopsis thaliana]
Sequence ID: AEE86099.1 Length: 356
Range 1: 16 to 73

Score:86.7 bits(213), Expect:4e-20,
Method:Compositional matrix adjust.,
Identities:42/58(72%), Positives:46/58(79%), Gaps:0/58(0%)

Query  602  KRPRMTWDESLPEQEAKGVVIKGHGSNMMILSPPLREDDCDGHYVFSLRDNLISIIKV  775
            KRPRMTWDE+  E EAK  VIKGHGS+  ILSPPLR+DD DGHYVFSLRDNL    K+
Sbjct  16   KRPRMTWDEAPAEPEAKRAVIKGHGSDGRILSPPLRDDDRDGHYVFSLRDNLTPRYKI  73


Range 2: 1 to 15

Score:31.6 bits(70), Expect:4e-20,
Method:Compositional matrix adjust.,
Identities:13/15(87%), Positives:15/15(100%), Gaps:0/15(0%)

Query  559  MIANGFENMDKERVK  603
            MIANGFE+MDKERV+
Sbjct  1    MIANGFESMDKERVR  15


>Protein kinase superfamily protein [Arabidopsis thaliana]
Sequence ID: NP_001031774.1 Length: 392
>Protein kinase superfamily protein [Arabidopsis thaliana]
Sequence ID: AEE86101.1 Length: 392
Range 1: 8 to 65

Score:87.0 bits(214), Expect:1e-18,
Method:Compositional matrix adjust.,
Identities:42/58(72%), Positives:46/58(79%), Gaps:0/58(0%)

Query  602  KRPRMTWDESLPEQEAKGVVIKGHGSNMMILSPPLREDDCDGHYVFSLRDNLISIIKV  775
            KRPRMTWDE+  E EAK  VIKGHGS+  ILSPPLR+DD DGHYVFSLRDNL    K+
Sbjct  8    KRPRMTWDEAPAEPEAKRAVIKGHGSDGRILSPPLRDDDRDGHYVFSLRDNLTPRYKI  65


>protein kinase, partial [Arabidopsis thaliana]
Sequence ID: AAA57119.1 Length: 395
Range 1: 11 to 68

Score:87.0 bits(214), Expect:1e-18,
Method:Compositional matrix adjust.,
Identities:42/58(72%), Positives:46/58(79%), Gaps:0/58(0%)

Query  602  KRPRMTWDESLPEQEAKGVVIKGHGSNMMILSPPLREDDCDGHYVFSLRDNLISIIKV  775
            KRPRMTWDE+  E EAK  VIKGHGS+  ILSPPLR+DD DGHYVFSLRDNL    K+
Sbjct  11   KRPRMTWDEAPAEPEAKRAVIKGHGSDGRILSPPLRDDDRDGHYVFSLRDNLTPRYKI  68


Query #38: XLOC_006084 Query ID: lcl|Query_76572 Length: 509

Sequences producing significant alignments:
                                                                  Max    Total Query   E   Per.
Description                                                       Score  Score cover Value Ident  Accession
transmembrane protein [Arabidopsis thaliana]                      55.8   92.8  31%   5e-13 75.76  NP_001326023.1
unnamed protein product [Arabidopsis thaliana]                    59.3   59.3  20%   2e-11 76.47  VYS51936.1
unnamed protein product [Arabidopsis thaliana]                    56.6   56.6  19%   9e-11 75.76  CAA0384287.1
hypothetical protein AXX17_AT5G63920 [Arabidopsis thaliana]       53.1   53.1  28%   5e-08 57.14  OAO93511.1


Alignments:
>transmembrane protein [Arabidopsis thaliana]
Sequence ID: NP_001326023.1 Length: 74
>transmembrane protein [Arabidopsis thaliana]
Sequence ID: ANM63965.1 Length: 74
Range 1: 12 to 44

Score:55.8 bits(133), Expect:5e-13,
Method:Compositional matrix adjust.,
Identities:25/33(76%), Positives:28/33(84%), Gaps:0/33(0%)

Query  25   DVCRDLHFQDLMDIELWFLYKPPLIESPVIMDM  123
             VCRDLH QDL D++LW LYKP LI+SPVIMDM
Sbjct  12   SVCRDLHLQDLTDMKLWSLYKPRLIKSPVIMDM  44


Range 2: 55 to 74

Score:37.0 bits(84), Expect:5e-13,
Method:Compositional matrix adjust.,
Identities:16/20(80%), Positives:17/20(85%), Gaps:0/20(0%)

Query  156  LQRPPPPRPYGHRNFIKFLR  215
            LQR PPPRPYG+R F KFLR
Sbjct  55   LQRAPPPRPYGYRTFTKFLR  74


>unnamed protein product [Arabidopsis thaliana]
Sequence ID: VYS51936.1 Length: 98
Range 1: 5 to 38

Score:59.3 bits(142), Expect:2e-11,
Method:Compositional matrix adjust.,
Identities:26/34(76%), Positives:30/34(88%), Gaps:0/34(0%)

Query  304  IDYGGGSPKIGGTDSANKWEKFLKQFDLDYLRNL  203
            IDYGGGSP IG T+S NKW+K L++FDLDYLRNL
Sbjct  5    IDYGGGSPDIGSTNSTNKWKKLLQKFDLDYLRNL  38


>unnamed protein product [Arabidopsis thaliana]
Sequence ID: CAA0384287.1 Length: 74
>unnamed protein product [Arabidopsis thaliana]
Sequence ID: VYS59150.1 Length: 74
Range 1: 12 to 44

Score:56.6 bits(135), Expect:9e-11,
Method:Compositional matrix adjust.,
Identities:25/33(76%), Positives:28/33(84%), Gaps:0/33(0%)

Query  25   DVCRDLHFQDLMDIELWFLYKPPLIESPVIMDM  123
             VCRDLH QDL D++LW LYKP LI+SPVIMDM
Sbjct  12   SVCRDLHLQDLTDMKLWSLYKPRLIKSPVIMDM  44


>hypothetical protein AXX17_AT5G63920 [Arabidopsis thaliana]
Sequence ID: OAO93511.1 Length: 728
Range 1: 43 to 86

Score:53.1 bits(126), Expect:5e-08,
Method:Composition-based stats.,
Identities:28/49(57%), Positives:31/49(63%), Gaps:5/49(10%)

Query  286  SPKIGGTDSANKWEKFLKQFDLDYLRNLIKFLCP*glggggLCKLFTDP  140
            SP+IGGTDSAN+WEK LK FDLD LRN    + P       LCKL   P
Sbjct  43   SPEIGGTDSANEWEKLLKPFDLDSLRNSFHKITP-----FQLCKLLELP  86


Query #39: XLOC_006264 Query ID: lcl|Query_76573 Length: 541

Sequences producing significant alignments:
                                                                  Max    Total Query   E   Per.
Description                                                       Score  Score cover Value Ident  Accession
unnamed protein product [Arabidopsis thaliana]                    57.0   97.0  36%   4e-14 97.67  CAA0272164.1


Alignments:
>unnamed protein product [Arabidopsis thaliana]
Sequence ID: CAA0272164.1 Length: 79
Range 1: 6 to 48

Score:57.0 bits(136), Expect:4e-14,
Method:Compositional matrix adjust.,
Identities:42/43(98%), Positives:43/43(100%), Gaps:0/43(0%)

Query  240  IHVLRDEREGLLGNGYNFFVGAldlrlldsrreesVTVTNQVE  368
            IHVLRDEREGLLGNGYNFFVGALDLRLLDSRREESVTVTN+VE
Sbjct  6    IHVLRDEREGLLGNGYNFFVGALDLRLLDSRREESVTVTNRVE  48


Range 2: 53 to 79

Score:40.0 bits(92), Expect:4e-14,
Method:Compositional matrix adjust.,
Identities:21/27(78%), Positives:22/27(81%), Gaps:0/27(0%)

Query  355  RIKLSDRTLN*ELGEEDLGTKHRFTSM  435
            +I  SD TLN ELGEEDL TKHRFTSM
Sbjct  53   KIFSSDITLNEELGEEDLETKHRFTSM  79


Query #40: XLOC_006294 Query ID: lcl|Query_76574 Length: 560

Sequences producing significant alignments:
                                                                  Max    Total Query   E   Per.
Description                                                       Score  Score cover Value Ident  Accession
hypothetical protein AXX17_AT5G28500 [Arabidopsis thaliana]       60.5   60.5  22%   2e-10 61.90  OAO92555.1
hypothetical protein AXX17_AT2G06050 [Arabidopsis thaliana]       59.7   59.7  22%   4e-10 61.90  OAP08384.1
hypothetical protein AXX17_AT1G38720 [Arabidopsis thaliana]       56.2   56.2  22%   8e-09 59.52  OAP14343.1
hypothetical protein AXX17_AT3G33420 [Arabidopsis thaliana]       53.5   53.5  17%   5e-08 75.00  OAP06548.1
F7F22.2 [Arabidopsis thaliana]                                    53.1   53.1  22%   8e-08 57.14  AAF24520.1


Alignments:
>hypothetical protein AXX17_AT5G28500 [Arabidopsis thaliana]
Sequence ID: OAO92555.1 Length: 845
Range 1: 291 to 332

Score:60.5 bits(145), Expect:2e-10,
Method:Composition-based stats.,
Identities:26/42(62%), Positives:33/42(78%), Gaps:0/42(0%)

Query  133  YVEPIIFLDVEMNPKEDLTWSDDEEDVRVDNIVMLVGEGITF  258
            YV+PII+ D  MNP EDL+WSDDE+DVRV+N+V +  EG  F
Sbjct  291  YVDPIIYPDSRMNPSEDLSWSDDEDDVRVENLVKMAEEGKAF  332


>hypothetical protein AXX17_AT2G06050 [Arabidopsis thaliana]
Sequence ID: OAP08384.1 Length: 730
Range 1: 103 to 144

Score:59.7 bits(143), Expect:4e-10,
Method:Composition-based stats.,
Identities:26/42(62%), Positives:33/42(78%), Gaps:0/42(0%)

Query  133  YVEPIIFLDVEMNPKEDLTWSDDEEDVRVDNIVMLVGEGITF  258
            +VEPIIF +  +NP EDLTWSDDE+D RVDN+V +  EG +F
Sbjct  103  HVEPIIFPNARLNPAEDLTWSDDEDDERVDNMVKMAKEGKSF  144


>hypothetical protein AXX17_AT1G38720 [Arabidopsis thaliana]
Sequence ID: OAP14343.1 Length: 783
Range 1: 209 to 250

Score:56.2 bits(134), Expect:8e-09,
Method:Composition-based stats.,
Identities:25/42(60%), Positives:33/42(78%), Gaps:0/42(0%)

Query  133  YVEPIIFLDVEMNPKEDLTWSDDEEDVRVDNIVMLVGEGITF  258
            +++PIIF D  MNP EDLTWS+D++DVRV+NIV +  EG  F
Sbjct  209  HLDPIIFPDSWMNPAEDLTWSNDKDDVRVENIVKMAEEGKPF  250


>hypothetical protein AXX17_AT3G33420 [Arabidopsis thaliana]
Sequence ID: OAP06548.1 Length: 962
Range 1: 285 to 316

Score:53.5 bits(127), Expect:5e-08,
Method:Composition-based stats.,
Identities:24/32(75%), Positives:28/32(87%), Gaps:0/32(0%)

Query  133  YVEPIIFLDVEMNPKEDLTWSDDEEDVRVDNI  228
            +V+PIIF D  MNP EDLTWSDDE+DVRV+NI
Sbjct  285  HVDPIIFPDSRMNPAEDLTWSDDEDDVRVENI  316


>F7F22.2 [Arabidopsis thaliana]
Sequence ID: AAF24520.1 Length: 815
Range 1: 158 to 199

Score:53.1 bits(126), Expect:8e-08,
Method:Composition-based stats.,
Identities:24/42(57%), Positives:32/42(76%), Gaps:0/42(0%)

Query  133  YVEPIIFLDVEMNPKEDLTWSDDEEDVRVDNIVMLVGEGITF  258
            +++PIIF D  MNP EDLTW +D++DVRV+NIV +  EG  F
Sbjct  158  HLDPIIFPDSWMNPAEDLTWLNDKDDVRVENIVKIAEEGKPF  199


Query #41: XLOC_006508 Query ID: lcl|Query_76575 Length: 769

No significant similarity found.


Query #42: XLOC_007343 Query ID: lcl|Query_76576 Length: 1073

No significant similarity found.


Query #43: XLOC_007532 Query ID: lcl|Query_76577 Length: 1278

No significant similarity found.


Query #44: XLOC_007787 Query ID: lcl|Query_76578 Length: 1078

Sequences producing significant alignments:
                                                                  Max    Total Query   E   Per.
Description                                                       Score  Score cover Value Ident  Accession
MATE efflux family protein [Arabidopsis thaliana]                 122    122   63%   3e-30 40.09  NP_200058.1
unnamed protein product [Arabidopsis thaliana]                    122    122   63%   3e-30 40.09  CAA0409331.1
unnamed protein product [Arabidopsis thaliana]                    120    120   56%   1e-29 38.28  CAA0374561.1
putative integral membrane protein [Arabidopsis thaliana]         119    119   56%   3e-29 38.28  AAM61608.1
MATE efflux family protein [Arabidopsis thaliana]                 119    119   56%   4e-29 38.28  NP_177511.1
MATE efflux family protein [Arabidopsis thaliana]                 118    118   53%   7e-29 40.31  NP_180983.4
hypothetical protein [Arabidopsis thaliana]                       118    118   53%   7e-29 40.31  AAC27412.1
hypothetical protein AXX17_AT2G30900 [Arabidopsis thaliana]       117    117   35%   2e-28 47.37  OAP08720.1
MATE efflux family protein [Arabidopsis thaliana]                 116    116   35%   2e-28 47.37  NP_001324516.1
unnamed protein product [Arabidopsis thaliana]                    112    112   35%   7e-27 45.86  CAA0261990.1
hypothetical protein AXX17_AT1G33050 [Arabidopsis thaliana]       112    112   35%   7e-27 45.86  OAP16194.1
MATE efflux family protein [Arabidopsis thaliana]                 74.7   74.7  35%   3e-14 34.59  NP_001323009.1
Chain A, Protein DETOXIFICATION 14 [Arabidopsis thaliana]         75.5   75.5  61%   3e-14 28.00  5Y50_A
MATE efflux family protein [Arabidopsis thaliana]                 75.1   75.1  61%   4e-14 28.44  NP_177270.1
hypothetical protein AXX17_AT1G15850 [Arabidopsis thaliana]       72.0   72.0  35%   5e-14 35.07  OAP17691.1
unnamed protein product [Arabidopsis thaliana]                    74.3   74.3  61%   6e-14 28.44  VYS50684.1
unnamed protein product [Arabidopsis thaliana]                    74.3   74.3  61%   7e-14 28.44  CAA0329013.1
MATE efflux family protein [Arabidopsis thaliana]                 73.9   73.9  35%   9e-14 34.59  NP_172967.2
Strong similarity to gi|4734005 F3L12.7 hypothetical protein...   73.6   73.6  35%   1e-13 34.59  AAD39645.1
hypothetical protein AXX17_AT1G65340 [Arabidopsis thaliana]       73.6   73.6  61%   1e-13 28.44  OAP18049.1
hypothetical protein AXX17_AT1G15840 [Arabidopsis thaliana]       72.8   72.8  35%   2e-13 33.83  OAP14688.1
unnamed protein product [Arabidopsis thaliana]                    70.9   70.9  30%   9e-13 40.00  VYS51993.1
MATE efflux family protein [Arabidopsis thaliana]                 70.9   70.9  30%   9e-13 40.00  NP_178491.1
unnamed protein product [Arabidopsis thaliana]                    70.5   70.5  35%   1e-12 35.07  CAA0206560.1
MATE efflux family protein [Arabidopsis thaliana]                 70.5   70.5  35%   1e-12 35.07  NP_172968.1
Strong similarity to gi|4734005 F3L12.7 hypothetical protein...   70.5   70.5  35%   1e-12 35.07  AAD39648.1
hypothetical protein AXX17_AT1G65340 [Arabidopsis thaliana]       69.7   69.7  35%   2e-12 34.09  OAP18050.1
hypothetical protein AXX17_AT2G03130 [Arabidopsis thaliana]       68.2   68.2  53%   3e-12 29.15  OAP10784.1
At1g66760/F4N21_11 [Arabidopsis thaliana]                         66.6   66.6  35%   8e-12 30.08  AAK97692.1
hypothetical protein [Arabidopsis thaliana]                       67.4   67.4  30%   1e-11 37.39  AAD28682.1
MATE efflux family protein [Arabidopsis thaliana]                 67.4   67.4  30%   1e-11 37.39  NP_001324928.1
MATE efflux family protein [Arabidopsis thaliana]                 67.4   67.4  30%   1e-11 37.39  NP_178499.2
MATE efflux family protein [Arabidopsis thaliana]                 67.4   67.4  35%   1e-11 34.31  NP_001117293.1
unnamed protein product [Arabidopsis thaliana]                    67.4   67.4  37%   1e-11 35.46  VYS51999.1
hypothetical protein AXX17_AT2G03120 [Arabidopsis thaliana]       67.0   67.0  37%   2e-11 34.75  OAP07829.1
hypothetical protein AXX17_AT2G03150 [Arabidopsis thaliana]       66.6   66.6  30%   2e-11 36.52  OAP09725.1
Strong similarity to gi|4734005 F3L12.7 hypothetical protein...   66.6   66.6  35%   3e-11 34.31  AAD39646.1
unnamed protein product [Arabidopsis thaliana]                    66.2   66.2  30%   3e-11 36.52  CAA0357024.1
unnamed protein product [Arabidopsis thaliana]                    66.2   66.2  30%   3e-11 36.52  VYS52000.1
MATE efflux family protein [Arabidopsis thaliana]                 66.2   66.2  35%   3e-11 34.31  NP_563964.1
hypothetical protein [Arabidopsis thaliana]                       65.9   65.9  30%   4e-11 36.52  AAD28683.1
MATE efflux family protein [Arabidopsis thaliana]                 65.9   65.9  30%   4e-11 36.52  NP_178498.2
MATE efflux family protein [Arabidopsis thaliana]                 65.5   65.5  35%   6e-11 30.08  NP_849854.1
putative MATE efflux family protein [Arabidopsis thaliana]        65.5   65.5  35%   6e-11 30.08  AAK25964.1
hypothetical protein AXX17_AT1G60540 [Arabidopsis thaliana]       65.5   65.5  35%   6e-11 30.08  OAP16837.1
hypothetical protein AXX17_AT2G03130 [Arabidopsis thaliana]       65.5   65.5  30%   6e-11 35.65  OAP10785.1
MATE efflux family protein [Arabidopsis thaliana]                 65.1   65.1  35%   7e-11 30.08  NP_564883.2
MATE efflux family protein [Arabidopsis thaliana]                 64.7   64.7  35%   8e-11 30.08  NP_001321857.1
hypothetical protein AXX17_AT2G03140 [Arabidopsis thaliana]       65.1   65.1  30%   1e-10 35.65  OAP11455.1
unnamed protein product [Arabidopsis thaliana]                    64.3   64.3  37%   1e-10 33.33  CAA0356956.1
unnamed protein product [Arabidopsis thaliana]                    63.9   63.9  37%   2e-10 33.33  CAA0357011.1
hypothetical protein AXX17_AT3G25120 [Arabidopsis thaliana]       63.5   63.5  43%   2e-10 30.62  OAP06402.1
unnamed protein product [Arabidopsis thaliana]                    63.5   63.5  43%   2e-10 30.62  VYS58370.1
MATE efflux family protein [Arabidopsis thaliana]                 63.5   63.5  43%   2e-10 30.62  NP_188997.1
MATE efflux family protein [Arabidopsis thaliana]                 62.4   62.4  37%   5e-10 32.62  NP_178497.2
MATE efflux family protein [Arabidopsis thaliana]                 62.4   62.4  37%   5e-10 32.62  NP_001324727.1
hypothetical protein [Arabidopsis thaliana]                       62.0   62.0  35%   8e-10 36.64  AAD28685.1
MATE efflux family protein [Arabidopsis thaliana]                 61.6   61.6  35%   9e-10 36.64  NP_178496.2
unnamed protein product [Arabidopsis thaliana]                    61.6   61.6  30%   1e-09 37.39  VYS51994.1
MATE efflux family protein [Arabidopsis thaliana]                 61.6   61.6  30%   1e-09 37.39  NP_178492.1
hypothetical protein AXX17_AT2G03100 [Arabidopsis thaliana]       61.6   61.6  30%   1e-09 37.39  OAP09049.1
hypothetical protein AXX17_AT2G03110 [Arabidopsis thaliana]       60.8   60.8  35%   1e-09 36.64  OAP07299.1
unnamed protein product [Arabidopsis thaliana]                    61.2   61.2  43%   1e-09 30.62  CAA0383444.1
MATE efflux family protein [Arabidopsis thaliana]                 56.2   56.2  33%   4e-08 29.03  NP_001321858.1


Alignments:
>MATE efflux family protein [Arabidopsis thaliana]
Sequence ID: NP_200058.1 Length: 486
>RecName: Full=Protein DETOXIFICATION 16; Short=AtDTX16; AltName: Full=Multidrug and toxic compound extrusion protein 16; Short=MATE protein 16 [Arabidopsis thaliana]
Sequence ID: Q9FHB6.1 Length: 486 >unknown protein [Arabidopsis thaliana]
Sequence ID: AAK76728.1 Length: 486 >unknown protein [Arabidopsis thaliana]
Sequence ID: AAL85047.1 Length: 486 >MATE efflux family protein [Arabidopsis thaliana]
Sequence ID: AED96215.1 Length: 486 >hypothetical protein AXX17_AT5G51330 [Arabidopsis thaliana]
Sequence ID: OAO95306.1 Length: 486 >unnamed protein product [Arabidopsis thaliana]
Sequence ID: VYS70114.1 Length: 486
Range 1: 109 to 312

Score:122 bits(305), Expect:3e-30,
Method:Compositional matrix adjust.,
Identities:93/232(40%), Positives:122/232(52%), Gaps:33/232(14%)

Query  278  ETRQFLYFYA*LPLSIIWINVSSILVFLDQDAIIA*TFGSYIVFLIPDLIGCVILKCFS-  454
            +   F+   A +PLSIIW N   +LVF  Q+  IA   GSY  F+IP +    +L+CF+
Sbjct  109  QRAMFVLTLASIPLSIIWANTEHLLVFFGQNKSIATLAGSYAKFMIPSIFAYGLLQCFNR  168

Query  455  -LRAKTECF--LSCS-VPVLLHVHACWGLVLKSGLGIREASQATSLSFWVNVVLLSIYVK  622
             L+A+   F  + CS V   LHV  CW LV KSGLG + A+ A S+S+W+NVVLL  YVK
Sbjct  169  FLQAQNNVFPVVFCSGVTTSLHVLLCWVLVFKSGLGFQGAALANSISYWLNVVLLFCYVK  228

Query  623  LSPSCAKTWTGFSPEALENVMVFLWSSICIHGLVRLVVHIYDSCLLVYIISLMLIIS*WN  802
             SPSC+ TWTGFS EAL +++ FL          RL V            +LM+ +  W+
Sbjct  229  FSPSCSLTWTGFSKEALRDILPFL----------RLAVP----------SALMVCLEMWS  268

Query  803  HNDLVIQENGLPLSLRLYFQNPVLETSVFPTWLCNHNNFW*RKLKESFASNT  958
               LV+    LP        NPVLETSV    L      W      S A++T
Sbjct  269  FELLVLLSGLLP--------NPVLETSVLSICLNTSGTMWMIPFGLSGAAST  312


>unnamed protein product [Arabidopsis thaliana]
Sequence ID: CAA0409331.1 Length: 486
Range 1: 109 to 312

Score:122 bits(305), Expect:3e-30,
Method:Compositional matrix adjust.,
Identities:93/232(40%), Positives:122/232(52%), Gaps:33/232(14%)

Query  278  ETRQFLYFYA*LPLSIIWINVSSILVFLDQDAIIA*TFGSYIVFLIPDLIGCVILKCFS-  454
            +   F+   A +PLSIIW N   +LVF  Q+  IA   GSY  F+IP +    +L+CF+
Sbjct  109  QRAMFVLTLASIPLSIIWANTEHLLVFFGQNKSIATLAGSYAKFMIPSIFAYGLLQCFNR  168

Query  455  -LRAKTECF--LSCS-VPVLLHVHACWGLVLKSGLGIREASQATSLSFWVNVVLLSIYVK  622
             L+A+   F  + CS V   LHV  CW LV KSGLG + A+ A S+S+W+NVVLL  YVK
Sbjct  169  FLQAQNNVFPVVFCSGVTTSLHVLLCWVLVFKSGLGFQGAALANSISYWLNVVLLFCYVK  228

Query  623  LSPSCAKTWTGFSPEALENVMVFLWSSICIHGLVRLVVHIYDSCLLVYIISLMLIIS*WN  802
             SPSC+ TWTGFS EAL +++ FL          RL V            +LM+ +  W+
Sbjct  229  FSPSCSLTWTGFSKEALRDILPFL----------RLAVP----------SALMVCLEMWS  268

Query  803  HNDLVIQENGLPLSLRLYFQNPVLETSVFPTWLCNHNNFW*RKLKESFASNT  958
               LV+    LP        NPVLETSV    L      W      S A++T
Sbjct  269  FELLVLLSGLLP--------NPVLETSVLSICLNTSGTMWMIPFGLSGAAST  312


>unnamed protein product [Arabidopsis thaliana]
Sequence ID: CAA0374561.1 Length: 457
Range 1: 121 to 301

Score:120 bits(300), Expect:1e-29,
Method:Compositional matrix adjust.,
Identities:80/209(38%), Positives:106/209(50%), Gaps:33/209(15%)

Query  311  LPLSIIWINVSSILVFLDQDAIIA*TFGSYIVFLIPDLIGCVILKCFS--LRAKTECF--  478
            +PLSI+W N    LVF  QD  IA   GSY  F+IP +    +L+C +  L+A+
Sbjct  121  VPLSIVWANTEHFLVFFGQDKSIAHLSGSYARFMIPSIFAYGLLQCLNRFLQAQNNVIPV  180

Query  479  -LSCSVPVLLHVHACWGLVLKSGLGIREASQATSLSFWVNVVLLSIYVKLSPSCAKTWTG  655
             +   V   LHV  CW LVLKSGLG R A+ A ++S+W+NV+LLS YVK SPSC+ TWTG
Sbjct  181  VICSGVTTSLHVIICWVLVLKSGLGFRGAAVANAISYWLNVILLSCYVKFSPSCSLTWTG  240

Query  656  FSPEALENVMVFLWSSICIHGLVRLVVHIYDSCLLVYIISLMLIIS*WNHNDLVIQENGL  835
            FS EA  +++ F+                     L    + M+ +  W+   LV+    L
Sbjct  241  FSKEARRDIIPFMK--------------------LAIPSAFMVCLEMWSFELLVLSSGLL  280

Query  836  PLSLRLYFQNPVLETSVFPTWLCNHNNFW  922
            P        NPVLETSV    L      W
Sbjct  281  P--------NPVLETSVLAICLNTSGTVW  301


>putative integral membrane protein [Arabidopsis thaliana]
Sequence ID: AAM61608.1 Length: 476
Range 1: 118 to 298

Score:119 bits(298), Expect:3e-29,
Method:Compositional matrix adjust.,
Identities:80/209(38%), Positives:108/209(51%), Gaps:33/209(15%)

Query  311  LPLSIIWINVSSILVFLDQDAIIA*TFGSYIVFLIPDLIGCVILKCFS--LRAKTECF--  478
            +PLSIIW N   ILV + QD  IA   GSY  ++IP L    +L+C +  L+A+   F
Sbjct  118  VPLSIIWANTEQILVLVHQDKSIASVAGSYAKYMIPSLFAYGLLQCINRFLQAQNNVFPV  177

Query  479  LSCS-VPVLLHVHACWGLVLKSGLGIREASQATSLSFWVNVVLLSIYVKLSPSCAKTWTG  655
              CS +   LH+  CW  VLK+GLG R A+ A S+S+W NV+LLS YVK SPSC+ +WTG
Sbjct  178  FVCSGITTCLHLLLCWLFVLKTGLGYRGAALAISVSYWFNVILLSCYVKFSPSCSHSWTG  237

Query  656  FSPEALENVMVFLWSSICIHGLVRLVVHIYDSCLLVYIISLMLIIS*WNHNDLVIQENGL  835
            FS EA +                     +YD   + +  ++M+ +  W+   LV+    L
Sbjct  238  FSKEAFQ--------------------ELYDFSKIAFPSAVMVCLELWSFELLVLASGLL  277

Query  836  PLSLRLYFQNPVLETSVFPTWLCNHNNFW  922
            P        NPVLETSV    L      W
Sbjct  278  P--------NPVLETSVLSICLNTSLTIW  298


>MATE efflux family protein [Arabidopsis thaliana]
Sequence ID: NP_177511.1 Length: 476
>RecName: Full=Protein DETOXIFICATION 17; Short=AtDTX17; AltName: Full=Multidrug and toxic compound extrusion protein 17; Short=MATE protein 17 [Arabidopsis thaliana]
Sequence ID: Q9C9U1.1 Length: 476 >putative integral membrane protein; 47574-45498 [Arabidopsis thaliana]
Sequence ID: AAG52084.1 Length: 476 >MATE efflux family protein [Arabidopsis thaliana]
Sequence ID: AEE35499.1 Length: 476 >hypothetical protein AXX17_AT1G67980 [Arabidopsis thaliana]
Sequence ID: OAP12794.1 Length: 476 >unnamed protein product [Arabidopsis thaliana]
Sequence ID: CAA0333356.1 Length: 476 >unnamed protein product [Arabidopsis thaliana]
Sequence ID: VYS50933.1 Length: 476
Range 1: 118 to 298

Score:119 bits(297), Expect:4e-29,
Method:Compositional matrix adjust.,
Identities:80/209(38%), Positives:108/209(51%), Gaps:33/209(15%)

Query  311  LPLSIIWINVSSILVFLDQDAIIA*TFGSYIVFLIPDLIGCVILKCFS--LRAKTECF--  478
            +PLSIIW N   ILV + QD  IA   GSY  ++IP L    +L+C +  L+A+   F
Sbjct  118  VPLSIIWANTEQILVLVHQDKSIASVAGSYAKYMIPSLFAYGLLQCINRFLQAQNNVFPV  177

Query  479  LSCS-VPVLLHVHACWGLVLKSGLGIREASQATSLSFWVNVVLLSIYVKLSPSCAKTWTG  655
              CS +   LH+  CW  VLK+GLG R A+ A S+S+W NV+LLS YVK SPSC+ +WTG
Sbjct  178  FVCSGITTCLHLLLCWLFVLKTGLGYRGAALAISVSYWFNVILLSCYVKFSPSCSHSWTG  237

Query  656  FSPEALENVMVFLWSSICIHGLVRLVVHIYDSCLLVYIISLMLIIS*WNHNDLVIQENGL  835
            FS EA +                     +YD   + +  ++M+ +  W+   LV+    L
Sbjct  238  FSKEAFQ--------------------ELYDFSKIAFPSAVMVCLELWSFELLVLASGLL  277

Query  836  PLSLRLYFQNPVLETSVFPTWLCNHNNFW  922
            P        NPVLETSV    L      W
Sbjct  278  P--------NPVLETSVLSICLNTSLTIW  298


>MATE efflux family protein [Arabidopsis thaliana]
Sequence ID: NP_180983.4 Length: 480
>RecName: Full=Protein DETOXIFICATION 15; Short=AtDTX15; AltName: Full=Multidrug and toxic compound extrusion protein 15; Short=MATE protein 15 [Arabidopsis thaliana]
Sequence ID: F4IHU9.1 Length: 480 >MATE efflux family protein [Arabidopsis thaliana]
Sequence ID: AEC08964.1 Length: 480
Range 1: 121 to 289

Score:118 bits(295), Expect:7e-29,
Method:Compositional matrix adjust.,
Identities:79/196(40%), Positives:107/196(54%), Gaps:32/196(16%)

Query  311  LPLSIIWINVSSILVFLDQDAIIA*TFGSYIVFLIPDLIGCVILKCFS--LRAKTECF--  478
            +PLSI+W N    LVF  QD  IA   GSY  F+IP +    +L+C +  L+A+
Sbjct  121  VPLSIVWANTEHFLVFFGQDKSIAHLSGSYARFMIPSIFAYGLLQCLNRFLQAQNNVIPV  180

Query  479  -LSCSVPVLLHVHACWGLVLKSGLGIREASQATSLSFWVNVVLLSIYVKLSPSCAKTWTG  655
             +   V   LHV  CW LVLKSGLG R A+ A ++S+W+NV+LLS YVK SPSC+ TWTG
Sbjct  181  VICSGVTTSLHVIICWVLVLKSGLGFRGAAVANAISYWLNVILLSCYVKFSPSCSLTWTG  240

Query  656  FSPEALENVMVFLWSSICIHGLVRLVVHIYDSCLLVYIISLMLIIS*WNHNDLVIQENGL  835
            FS EA  +++ F+          +LV+    S  +V  + +      W+   LV+    L
Sbjct  241  FSKEARRDIIPFM----------KLVI---PSAFMVCSLEM------WSFELLVLSSGLL  281

Query  836  PLSLRLYFQNPVLETS  883
            P        NPVLETS
Sbjct  282  P--------NPVLETS  289


>hypothetical protein [Arabidopsis thaliana]
Sequence ID: AAC27412.1 Length: 466
Range 1: 121 to 289

Score:118 bits(295), Expect:7e-29,
Method:Compositional matrix adjust.,
Identities:79/196(40%), Positives:107/196(54%), Gaps:32/196(16%)

Query  311  LPLSIIWINVSSILVFLDQDAIIA*TFGSYIVFLIPDLIGCVILKCFS--LRAKTECF--  478
            +PLSI+W N    LVF  QD  IA   GSY  F+IP +    +L+C +  L+A+
Sbjct  121  VPLSIVWANTEHFLVFFGQDKSIAHLSGSYARFMIPSIFAYGLLQCLNRFLQAQNNVIPV  180

Query  479  -LSCSVPVLLHVHACWGLVLKSGLGIREASQATSLSFWVNVVLLSIYVKLSPSCAKTWTG  655
             +   V   LHV  CW LVLKSGLG R A+ A ++S+W+NV+LLS YVK SPSC+ TWTG
Sbjct  181  VICSGVTTSLHVIICWVLVLKSGLGFRGAAVANAISYWLNVILLSCYVKFSPSCSLTWTG  240

Query  656  FSPEALENVMVFLWSSICIHGLVRLVVHIYDSCLLVYIISLMLIIS*WNHNDLVIQENGL  835
            FS EA  +++ F+          +LV+    S  +V  + +      W+   LV+    L
Sbjct  241  FSKEARRDIIPFM----------KLVI---PSAFMVCSLEM------WSFELLVLSSGLL  281

Query  836  PLSLRLYFQNPVLETS  883
            P        NPVLETS
Sbjct  282  P--------NPVLETS  289


>hypothetical protein AXX17_AT2G30900 [Arabidopsis thaliana]
Sequence ID: OAP08720.1 Length: 456
Range 1: 149 to 281

Score:117 bits(292), Expect:2e-28,
Method:Compositional matrix adjust.,
Identities:63/133(47%), Positives:84/133(63%), Gaps:5/133(3%)

Query  311  LPLSIIWINVSSILVFLDQDAIIA*TFGSYIVFLIPDLIGCVILKCFS--LRAKTECF--  478
            +PLSI+W N    LVF  QD  IA   GSY  F+IP +    +L+C +  L+A+
Sbjct  149  VPLSIVWANTEHFLVFFGQDKSIAHLSGSYARFMIPSIFAYGLLQCLNRFLQAQNNVIPV  208

Query  479  -LSCSVPVLLHVHACWGLVLKSGLGIREASQATSLSFWVNVVLLSIYVKLSPSCAKTWTG  655
             +   V   LHV  CW LVLKSGLG R A+ A ++S+W+NV+LLS YVK SPSC+ TWTG
Sbjct  209  VICSGVTTSLHVIICWVLVLKSGLGFRGAAVANAISYWLNVILLSCYVKFSPSCSLTWTG  268

Query  656  FSPEALENVMVFL  694
            FS EA  +++ F+
Sbjct  269  FSKEARRDIIPFM  281


>MATE efflux family protein [Arabidopsis thaliana]
Sequence ID: NP_001324516.1 Length: 438
>MATE efflux family protein [Arabidopsis thaliana]
Sequence ID: ANM62355.1 Length: 438 >unnamed protein product [Arabidopsis thaliana]
Sequence ID: VYS54421.1 Length: 438
Range 1: 121 to 253

Score:116 bits(291), Expect:2e-28,
Method:Compositional matrix adjust.,
Identities:63/133(47%), Positives:84/133(63%), Gaps:5/133(3%)

Query  311  LPLSIIWINVSSILVFLDQDAIIA*TFGSYIVFLIPDLIGCVILKCFS--LRAKTECF--  478
            +PLSI+W N    LVF  QD  IA   GSY  F+IP +    +L+C +  L+A+
Sbjct  121  VPLSIVWANTEHFLVFFGQDKSIAHLSGSYARFMIPSIFAYGLLQCLNRFLQAQNNVIPV  180

Query  479  -LSCSVPVLLHVHACWGLVLKSGLGIREASQATSLSFWVNVVLLSIYVKLSPSCAKTWTG  655
             +   V   LHV  CW LVLKSGLG R A+ A ++S+W+NV+LLS YVK SPSC+ TWTG
Sbjct  181  VICSGVTTSLHVIICWVLVLKSGLGFRGAAVANAISYWLNVILLSCYVKFSPSCSLTWTG  240

Query  656  FSPEALENVMVFL  694
            FS EA  +++ F+
Sbjct  241  FSKEARRDIIPFM  253


>unnamed protein product [Arabidopsis thaliana]
Sequence ID: CAA0261990.1 Length: 484
>unnamed protein product [Arabidopsis thaliana]
Sequence ID: VYS47790.1 Length: 484
Range 1: 127 to 259

Score:112 bits(281), Expect:7e-27,
Method:Compositional matrix adjust.,
Identities:61/133(46%), Positives:84/133(63%), Gaps:5/133(3%)

Query  311  LPLSIIWINVSSILVFLDQDAIIA*TFGSYIVFLIPDLIGCVILKCFSLRAKTE-----C  475
            +PLSI+W     ILV + Q+  IA   GSY  ++IP +    +L+C +   +T+
Sbjct  127  VPLSIVWFYTEDILVLVHQNKSIARLAGSYARYMIPSIYAYALLQCLNRFLQTQNNVFPV  186

Query  476  FLSCSVPVLLHVHACWGLVLKSGLGIREASQATSLSFWVNVVLLSIYVKLSPSCAKTWTG  655
            F+S  +   LH+  CW  VLKSGLG R A+ A S+S+WVNV+LLS YVK S SC++TWTG
Sbjct  187  FVSSGITTCLHLLLCWVFVLKSGLGHRGAALAISVSYWVNVILLSCYVKFSASCSQTWTG  246

Query  656  FSPEALENVMVFL  694
            FS EAL ++  FL
Sbjct  247  FSKEALSHIPAFL  259


>hypothetical protein AXX17_AT1G33050 [Arabidopsis thaliana]
Sequence ID: OAP16194.1 Length: 478
Range 1: 127 to 259

Score:112 bits(280), Expect:7e-27,
Method:Compositional matrix adjust.,
Identities:61/133(46%), Positives:84/133(63%), Gaps:5/133(3%)

Query  311  LPLSIIWINVSSILVFLDQDAIIA*TFGSYIVFLIPDLIGCVILKCFSLRAKTE-----C  475
            +PLSI+W     ILV + Q+  IA   GSY  ++IP +    +L+C +   +T+
Sbjct  127  VPLSIVWFYTEDILVLVHQNKSIARLAGSYARYMIPSIYAYALLQCLNRFLQTQNNVFPV  186

Query  476  FLSCSVPVLLHVHACWGLVLKSGLGIREASQATSLSFWVNVVLLSIYVKLSPSCAKTWTG  655
            F+S  +   LH+  CW  VLKSGLG R A+ A S+S+WVNV+LLS YVK S SC++TWTG
Sbjct  187  FVSSGITTCLHLLLCWVFVLKSGLGHRGAALAISVSYWVNVILLSCYVKFSASCSQTWTG  246

Query  656  FSPEALENVMVFL  694
            FS EAL ++  FL
Sbjct  247  FSKEALSHIPAFL  259


>MATE efflux family protein [Arabidopsis thaliana]
Sequence ID: NP_001323009.1 Length: 371
>MATE efflux family protein [Arabidopsis thaliana]
Sequence ID: ANM60745.1 Length: 371
Range 1: 9 to 141

Score:74.7 bits(182), Expect:3e-14,
Method:Compositional matrix adjust.,
Identities:46/133(35%), Positives:69/133(51%), Gaps:5/133(3%)

Query  311  LPLSIIWINVSSILVFLDQDAIIA*TFGSYIVFLIPDLIGCVILKCFSLRAKTECFLS--  484
            LPLS++W N+  ++V L QD  IA   G Y  +LIP L    +L+      K +  ++
Sbjct  9    LPLSLLWFNMGKLIVILGQDPAIAHEAGRYAAWLIPGLFAYAVLQPLIRYFKNQSLITPL  68

Query  485  ---CSVPVLLHVHACWGLVLKSGLGIREASQATSLSFWVNVVLLSIYVKLSPSCAKTWTG  655
                SV   +HV  CW LV KSGLG    + A SLS+W+  + L  ++  S +C++T
Sbjct  69   LVTSSVVFCIHVPLCWLLVYKSGLGHIGGALALSLSYWLYAIFLGSFMYYSSACSETRAP  128

Query  656  FSPEALENVMVFL  694
             + E  E V  F+
Sbjct  129  LTMEIFEGVREFI  141


>Chain A, Protein DETOXIFICATION 14 [Arabidopsis thaliana]
Sequence ID: 5Y50_A Length: 461
Range 1: 104 to 298

Score:75.5 bits(184), Expect:3e-14,
Method:Compositional matrix adjust.,
Identities:63/225(28%), Positives:99/225(44%), Gaps:35/225(15%)

Query  311  LPLSIIWINVSSILVFLDQDAIIA*TFGSYIVFLIPDLIGCVIL----KCFSLRAKTECF  478
            +PLS++W  +  IL  + QDA++A   G +  +LIP L G   L    + F  ++
Sbjct  104  IPLSLLWTYIGDILSLIGQDAMVAQEAGKFATWLIPALFGYATLQPLVRFFQAQSLILPL  163

Query  479  LSCSVPVL-LHVHACWGLVLKSGLGIREASQATSLSFWVNVVLLSIYVKLSPSCAKTWTG  655
            +  SV  L +H+  CW LV K GLG   A+ A  +S+W+NV +L +Y+  S SC+K+
Sbjct  164  VMSSVSSLCIHIVLCWSLVFKFGLGSLGAAIAIGVSYWLNVTVLGLYMTFSSSCSKSRAT  223

Query  656  FSPEALENVMVFLWSSICIHGLVRLVVHIYDSCLLVYIISLMLIIS*WNHNDLVIQENGL  835
             S    E +  F    I                      + M+ +  W+   LV+    L
Sbjct  224  ISMSLFEGMGEFFRFGIP--------------------SASMICLEWWSFEFLVLLSGIL  263

Query  836  PLSLRLYFQNPVLETSVFPTWLCNHNNFW*RKLKESFASNTGDRV  970
            P        NP LE SV    L   ++ +  ++ ES  +    RV
Sbjct  264  P--------NPKLEASVLSVCLSTQSSLY--QIPESLGAAASTRV  298


>MATE efflux family protein [Arabidopsis thaliana]
Sequence ID: NP_177270.1 Length: 485
>RecName: Full=Protein DETOXIFICATION 14; Short=AtDTX14; AltName: Full=Multidrug and toxic compound extrusion protein 14; Short=MATE protein 14 [Arabidopsis thaliana]
Sequence ID: Q9C994.1 Length: 485 >hypothetical protein; 49518-51504 [Arabidopsis thaliana]
Sequence ID: AAG51691.1 Length: 485 >MATE efflux family protein [Arabidopsis thaliana]
Sequence ID: AEE35165.1 Length: 485
Range 1: 120 to 314

Score:75.1 bits(183), Expect:4e-14,
Method:Compositional matrix adjust.,
Identities:64/225(28%), Positives:99/225(44%), Gaps:35/225(15%)

Query  311  LPLSIIWINVSSILVFLDQDAIIA*TFGSYIVFLIPDLIGCVIL----KCFSLRAKTECF  478
            +PLS++W  +  IL  + QDA++A   G +  +LIP L G   L    + F  ++
Sbjct  120  IPLSLLWTYIGDILSLIGQDAMVAQEAGKFATWLIPALFGYATLQPLVRFFQAQSLILPL  179

Query  479  LSCSVPVL-LHVHACWGLVLKSGLGIREASQATSLSFWVNVVLLSIYVKLSPSCAKTWTG  655
            +  SV  L +H+  CW LV K GLG   A+ A  +S+W+NV +L +Y+  S SC+K+
Sbjct  180  VMSSVSSLCIHIVLCWSLVFKFGLGSLGAAIAIGVSYWLNVTVLGLYMTFSSSCSKSRAT  239

Query  656  FSPEALENVMVFLWSSICIHGLVRLVVHIYDSCLLVYIISLMLIIS*WNHNDLVIQENGL  835
             S    E +  F    I    ++         CL             W+   LV+    L
Sbjct  240  ISMSLFEGMGEFFRFGIPSASMI---------CL-----------EWWSFEFLVLLSGIL  279

Query  836  PLSLRLYFQNPVLETSVFPTWLCNHNNFW*RKLKESFASNTGDRV  970
            P        NP LE SV    L   ++ +  ++ ES  +    RV
Sbjct  280  P--------NPKLEASVLSVCLSTQSSLY--QIPESLGAAASTRV  314


>hypothetical protein AXX17_AT1G15850 [Arabidopsis thaliana]
Sequence ID: OAP17691.1 Length: 200
Range 1: 39 to 171

Score:72.0 bits(175), Expect:5e-14,
Method:Compositional matrix adjust.,
Identities:47/134(35%), Positives:69/134(51%), Gaps:7/134(5%)

Query  311  LPLSIIWINVSSILVFLDQDAIIA*TFGSYIVFLIPDLIGCVILKCFSLRAKTECFL---  481
            LPLS++W N+  +LV L QD  IA   G +  +LIP L    +L+  +   K +  +
Sbjct  39   LPLSLLWFNMGKLLVILGQDPSIAHEAGRFAAWLIPGLFAYAVLQPLTRYFKNQSLITPL  98

Query  482  ---SCSVPVLLHVHACWGLVLKSGLGIREASQATSLSFWVNVVLLSIYVKLSPSCAKTWT  652
               SC V   LHV  CW LV KSGL     + A SLS+W+  + L  ++  S +C++T
Sbjct  99   LITSCVV-FCLHVPLCWLLVYKSGLDHIGGALALSLSYWLYAIFLGSFMYFSSACSETRA  157

Query  653  GFSPEALENVMVFL  694
              + E  E V  F+
Sbjct  158  PLTMEIFEGVREFI  171


>unnamed protein product [Arabidopsis thaliana]
Sequence ID: VYS50684.1 Length: 485
Range 1: 120 to 314

Score:74.3 bits(181), Expect:6e-14,
Method:Compositional matrix adjust.,
Identities:64/225(28%), Positives:99/225(44%), Gaps:35/225(15%)

Query  311  LPLSIIWINVSSILVFLDQDAIIA*TFGSYIVFLIPDLIGCVIL----KCFSLRAKTECF  478
            +PLS++W  +  IL  + QDA++A   G +  +LIP L G   L    + F  ++
Sbjct  120  IPLSLLWSYIGDILSLIGQDAMVAQEAGKFATWLIPALFGYATLQPLVRFFQAQSLILPL  179

Query  479  LSCSVPVL-LHVHACWGLVLKSGLGIREASQATSLSFWVNVVLLSIYVKLSPSCAKTWTG  655
            +  SV  L +H+  CW LV K GLG   A+ A  +S+W+NV +L +Y+  S SC+K+
Sbjct  180  VMSSVSSLCIHIVLCWSLVFKFGLGSLGAAIAIGVSYWLNVTVLGLYMTFSSSCSKSRAT  239

Query  656  FSPEALENVMVFLWSSICIHGLVRLVVHIYDSCLLVYIISLMLIIS*WNHNDLVIQENGL  835
             S    E +  F    I    ++         CL             W+   LV+    L
Sbjct  240  ISMSLFEGMGEFFRFGIPSASMI---------CL-----------EWWSFEFLVLLSGIL  279

Query  836  PLSLRLYFQNPVLETSVFPTWLCNHNNFW*RKLKESFASNTGDRV  970
            P        NP LE SV    L   ++ +  ++ ES  +    RV
Sbjct  280  P--------NPKLEASVLSVCLSTQSSLY--QIPESLGAAASTRV  314


>unnamed protein product [Arabidopsis thaliana]
Sequence ID: CAA0329013.1 Length: 485
Range 1: 120 to 314

Score:74.3 bits(181), Expect:7e-14,
Method:Compositional matrix adjust.,
Identities:64/225(28%), Positives:99/225(44%), Gaps:35/225(15%)

Query  311  LPLSIIWINVSSILVFLDQDAIIA*TFGSYIVFLIPDLIGCVIL----KCFSLRAKTECF  478
            +PLS++W  +  IL  + QDA++A   G +  +LIP L G   L    + F  ++
Sbjct  120  IPLSLLWSYIGDILSLIGQDAMVAQEAGKFATWLIPALFGYATLQPLVRFFQAQSLILPL  179

Query  479  LSCSVPVL-LHVHACWGLVLKSGLGIREASQATSLSFWVNVVLLSIYVKLSPSCAKTWTG  655
            +  SV  L +H+  CW LV K GLG   A+ A  +S+W+NV +L +Y+  S SC+K+
Sbjct  180  VMSSVSSLCVHIVLCWSLVFKFGLGSLGAAIAIGVSYWLNVTVLGLYMTFSSSCSKSRAT  239

Query  656  FSPEALENVMVFLWSSICIHGLVRLVVHIYDSCLLVYIISLMLIIS*WNHNDLVIQENGL  835
             S    E +  F    I    ++         CL             W+   LV+    L
Sbjct  240  ISMSLFEGMGEFFRFGIPSASMI---------CL-----------EWWSFEFLVLLSGIL  279

Query  836  PLSLRLYFQNPVLETSVFPTWLCNHNNFW*RKLKESFASNTGDRV  970
            P        NP LE SV    L   ++ +  ++ ES  +    RV
Sbjct  280  P--------NPKLEASVLSVCLSTQSSLY--QIPESLGAAASTRV  314


>MATE efflux family protein [Arabidopsis thaliana]
Sequence ID: NP_172967.2 Length: 487
>RecName: Full=Protein DETOXIFICATION 10; Short=AtDTX10; AltName: Full=Multidrug and toxic compound extrusion protein 10; Short=MATE protein 10 [Arabidopsis thaliana]
Sequence ID: Q8VYL8.1 Length: 487 >unknown protein [Arabidopsis thaliana]
Sequence ID: AAL49848.1 Length: 487 >unknown protein [Arabidopsis thaliana]
Sequence ID: AAM51440.1 Length: 487 >MATE efflux family protein [Arabidopsis thaliana]
Sequence ID: AEE29274.1 Length: 487
Range 1: 125 to 257

Score:73.9 bits(180), Expect:9e-14,
Method:Compositional matrix adjust.,
Identities:46/133(35%), Positives:69/133(51%), Gaps:5/133(3%)

Query  311  LPLSIIWINVSSILVFLDQDAIIA*TFGSYIVFLIPDLIGCVILKCFSLRAKTECFLS--  484
            LPLS++W N+  ++V L QD  IA   G Y  +LIP L    +L+      K +  ++
Sbjct  125  LPLSLLWFNMGKLIVILGQDPAIAHEAGRYAAWLIPGLFAYAVLQPLIRYFKNQSLITPL  184

Query  485  ---CSVPVLLHVHACWGLVLKSGLGIREASQATSLSFWVNVVLLSIYVKLSPSCAKTWTG  655
                SV   +HV  CW LV KSGLG    + A SLS+W+  + L  ++  S +C++T
Sbjct  185  LVTSSVVFCIHVPLCWLLVYKSGLGHIGGALALSLSYWLYAIFLGSFMYYSSACSETRAP  244

Query  656  FSPEALENVMVFL  694
             + E  E V  F+
Sbjct  245  LTMEIFEGVREFI  257


>Strong similarity to gi|4734005 F3L12.7 hypothetical protein from Arabidopsis thaliana BAC gb|AC007178 [Arabidopsis thaliana]
Sequence ID: AAD39645.1 Length: 487
Range 1: 125 to 257

Score:73.6 bits(179), Expect:1e-13,
Method:Compositional matrix adjust.,
Identities:46/133(35%), Positives:69/133(51%), Gaps:5/133(3%)

Query  311  LPLSIIWINVSSILVFLDQDAIIA*TFGSYIVFLIPDLIGCVILKCFSLRAKTECFLS--  484
            LPLS++W N+  ++V L QD  IA   G Y  +LIP L    +L+      K +  ++
Sbjct  125  LPLSLLWFNMGKLIVILGQDPAIAHEAGRYAAWLIPGLFAYAVLQPLIRYFKNQSLITPL  184

Query  485  ---CSVPVLLHVHACWGLVLKSGLGIREASQATSLSFWVNVVLLSIYVKLSPSCAKTWTG  655
                SV   +HV  CW LV KSGLG    + A SLS+W+  + L  ++  S +C++T
Sbjct  185  LVTSSVVFCIHVPLCWLLVYKSGLGHIGGALALSLSYWLYAIFLGSFMYYSSACSETRAP  244

Query  656  FSPEALENVMVFL  694
             + E  E V  F+
Sbjct  245  LTMEIFEGVREFI  257


>hypothetical protein AXX17_AT1G65340 [Arabidopsis thaliana]
Sequence ID: OAP18049.1 Length: 485
Range 1: 120 to 314

Score:73.6 bits(179), Expect:1e-13,
Method:Compositional matrix adjust.,
Identities:64/225(28%), Positives:98/225(43%), Gaps:35/225(15%)

Query  311  LPLSIIWINVSSILVFLDQDAIIA*TFGSYIVFLIPDLIGCVIL----KCFSLRAKTECF  478
            +PLS++W  +  IL  + QDA++A   G +  +LIP L G   L    + F  ++
Sbjct  120  IPLSVLWSYIGDILSLIGQDAMVAQEAGKFATWLIPALFGYATLQPLVRFFQAQSLILPL  179

Query  479  LSCSVPVL-LHVHACWGLVLKSGLGIREASQATSLSFWVNVVLLSIYVKLSPSCAKTWTG  655
            +  SV  L +H+  CW LV K GLG   A+ A  +S+W+NV +L +Y+  S SC K+
Sbjct  180  VMSSVSSLCVHIVLCWSLVFKFGLGSLGAAIAIGVSYWLNVTVLGLYMTFSSSCRKSRAP  239

Query  656  FSPEALENVMVFLWSSICIHGLVRLVVHIYDSCLLVYIISLMLIIS*WNHNDLVIQENGL  835
             S    E +  F    I    ++         CL             W+   LV+    L
Sbjct  240  ISMSLFEGMGEFFRFGIPSASMI---------CL-----------EWWSFEFLVLLSGIL  279

Query  836  PLSLRLYFQNPVLETSVFPTWLCNHNNFW*RKLKESFASNTGDRV  970
            P        NP LE SV    L   ++ +  ++ ES  +    RV
Sbjct  280  P--------NPKLEASVLSVCLSTQSSLY--QIPESLGAAASTRV  314


>hypothetical protein AXX17_AT1G15840 [Arabidopsis thaliana]
Sequence ID: OAP14688.1 Length: 487
>unnamed protein product [Arabidopsis thaliana]
Sequence ID: CAA0206524.1 Length: 487 >unnamed protein product [Arabidopsis thaliana]
Sequence ID: VYS46143.1 Length: 487
Range 1: 125 to 257

Score:72.8 bits(177), Expect:2e-13,
Method:Compositional matrix adjust.,
Identities:45/133(34%), Positives:69/133(51%), Gaps:5/133(3%)

Query  311  LPLSIIWINVSSILVFLDQDAIIA*TFGSYIVFLIPDLIGCVILKCFSLRAKTECFLS--  484
            LPLS++W N+  ++V L QD  IA   G Y  +LIP L    +L+      K +  ++
Sbjct  125  LPLSLLWFNMGKLIVILGQDPAIAHEAGRYAAWLIPGLFAYAVLQPLIRYFKNQSLITPL  184

Query  485  ---CSVPVLLHVHACWGLVLKSGLGIREASQATSLSFWVNVVLLSIYVKLSPSCAKTWTG  655
                SV   +HV  CW LV KSGLG    + A SLS+W+  + L  ++  S +C++T
Sbjct  185  LVTSSVVFCIHVPLCWLLVYKSGLGHIGGALALSLSYWLYAIFLGSFMYYSSACSETRAP  244

Query  656  FSPEALENVMVFL  694
             + E  + V  F+
Sbjct  245  LTMEIFDGVREFI  257


>unnamed protein product [Arabidopsis thaliana]
Sequence ID: VYS51993.1 Length: 476
Range 1: 123 to 237

Score:70.9 bits(172), Expect:9e-13,
Method:Compositional matrix adjust.,
Identities:46/115(40%), Positives:62/115(53%), Gaps:5/115(4%)

Query  317  LSIIWINVSSILVFLDQDAIIA*TFGSYIVFLIPDLIGCVILKCFSLRAKTECFL-----  481
            +SI+W+ +  IL+ L QD  I+   GSY  +LIP L G  I+   S    T+  +
Sbjct  123  ISILWLYIEKILISLGQDPEISRIAGSYAFWLIPALFGQAIVIPLSRFLLTQGLVIPLLF  182

Query  482  SCSVPVLLHVHACWGLVLKSGLGIREASQATSLSFWVNVVLLSIYVKLSPSCAKT  646
            +    +L HV  CW LV   GLG    + ATS+SFW   V+LS YV+ S SC KT
Sbjct  183  TAVTTLLFHVLVCWTLVFLFGLGCNGPAMATSVSFWFYAVILSCYVRFSSSCEKT  237


>MATE efflux family protein [Arabidopsis thaliana]
Sequence ID: NP_178491.1 Length: 476
>RecName: Full=Protein DETOXIFICATION 1; Short=AtDTX1; AltName: Full=Multidrug and toxic compound extrusion protein 1; Short=MATE protein 1 [Arabidopsis thaliana]
Sequence ID: Q9SIA5.1 Length: 476 >hypothetical protein [Arabidopsis thaliana]
Sequence ID: AAD28687.1 Length: 476 >At2g04040 [Arabidopsis thaliana]
Sequence ID: AAY25434.1 Length: 476 >At2g04040 [Arabidopsis thaliana]
Sequence ID: ABH04478.1 Length: 476 >MATE efflux family protein [Arabidopsis thaliana]
Sequence ID: AEC05791.1 Length: 476 >TX1 [Arabidopsis thaliana]
Sequence ID: OAP07361.1 Length: 476
Range 1: 123 to 237

Score:70.9 bits(172), Expect:9e-13,
Method:Compositional matrix adjust.,
Identities:46/115(40%), Positives:62/115(53%), Gaps:5/115(4%)

Query  317  LSIIWINVSSILVFLDQDAIIA*TFGSYIVFLIPDLIGCVILKCFSLRAKTECFL-----  481
            +SI+W+ +  IL+ L QD  I+   GSY  +LIP L G  I+   S    T+  +
Sbjct  123  ISILWLYIEKILISLGQDPEISRIAGSYAFWLIPALFGQAIVIPLSRFLLTQGLVIPLLF  182

Query  482  SCSVPVLLHVHACWGLVLKSGLGIREASQATSLSFWVNVVLLSIYVKLSPSCAKT  646
            +    +L HV  CW LV   GLG    + ATS+SFW   V+LS YV+ S SC KT
Sbjct  183  TAVTTLLFHVLVCWTLVFLFGLGCNGPAMATSVSFWFYAVILSCYVRFSSSCEKT  237


>unnamed protein product [Arabidopsis thaliana]
Sequence ID: CAA0206560.1 Length: 487
Range 1: 125 to 257

Score:70.5 bits(171), Expect:1e-12,
Method:Compositional matrix adjust.,
Identities:47/134(35%), Positives:69/134(51%), Gaps:7/134(5%)

Query  311  LPLSIIWINVSSILVFLDQDAIIA*TFGSYIVFLIPDLIGCVILKCFSLRAKTECFL---  481
            LPLS++W N+  +LV L QD  IA   G +  +LIP L    +L+  +   K +  +
Sbjct  125  LPLSLLWFNMGKLLVILGQDPSIAHEAGRFAAWLIPGLFAYAVLQPLTRYFKNQSLITPL  184

Query  482  ---SCSVPVLLHVHACWGLVLKSGLGIREASQATSLSFWVNVVLLSIYVKLSPSCAKTWT  652
               SC V   LHV  CW LV KSGL     + A SLS+W+  + L  ++  S +C++T
Sbjct  185  LITSCVV-FCLHVPLCWLLVYKSGLDHIGGALALSLSYWLYAIFLGSFMYFSSACSETRA  243

Query  653  GFSPEALENVMVFL  694
              + E  E V  F+
Sbjct  244  PLTMEIFEGVREFI  257


>MATE efflux family protein [Arabidopsis thaliana]
Sequence ID: NP_172968.1 Length: 487
>RecName: Full=Protein DETOXIFICATION 11; Short=AtDTX11; AltName: Full=Multidrug and toxic compound extrusion protein 11; Short=MATE protein 11 [Arabidopsis thaliana]
Sequence ID: F4HZH9.1 Length: 487 >MATE efflux family protein [Arabidopsis thaliana]
Sequence ID: AEE29275.1 Length: 487 >unnamed protein product [Arabidopsis thaliana]
Sequence ID: VYS46144.1 Length: 487
Range 1: 125 to 257

Score:70.5 bits(171), Expect:1e-12,
Method:Compositional matrix adjust.,
Identities:47/134(35%), Positives:69/134(51%), Gaps:7/134(5%)

Query  311  LPLSIIWINVSSILVFLDQDAIIA*TFGSYIVFLIPDLIGCVILKCFSLRAKTECFL---  481
            LPLS++W N+  +LV L QD  IA   G +  +LIP L    +L+  +   K +  +
Sbjct  125  LPLSLLWFNMGKLLVILGQDPSIAHEAGRFAAWLIPGLFAYAVLQPLTRYFKNQSLITPL  184

Query  482  ---SCSVPVLLHVHACWGLVLKSGLGIREASQATSLSFWVNVVLLSIYVKLSPSCAKTWT  652
               SC V   LHV  CW LV KSGL     + A SLS+W+  + L  ++  S +C++T
Sbjct  185  LITSCVV-FCLHVPLCWLLVYKSGLDHIGGALALSLSYWLYAIFLGSFMYFSSACSETRA  243

Query  653  GFSPEALENVMVFL  694
              + E  E V  F+
Sbjct  244  PLTMEIFEGVREFI  257


>Strong similarity to gi|4734005 F3L12.7 hypothetical protein from Arabidopsis thaliana BAC gb|AC007178 [Arabidopsis thaliana]
Sequence ID: AAD39648.1 Length: 481
Range 1: 125 to 257

Score:70.5 bits(171), Expect:1e-12,
Method:Compositional matrix adjust.,
Identities:47/134(35%), Positives:69/134(51%), Gaps:7/134(5%)

Query  311  LPLSIIWINVSSILVFLDQDAIIA*TFGSYIVFLIPDLIGCVILKCFSLRAKTECFL---  481
            LPLS++W N+  +LV L QD  IA   G +  +LIP L    +L+  +   K +  +
Sbjct  125  LPLSLLWFNMGKLLVILGQDPSIAHEAGRFAAWLIPGLFAYAVLQPLTRYFKNQSLITPL  184

Query  482  ---SCSVPVLLHVHACWGLVLKSGLGIREASQATSLSFWVNVVLLSIYVKLSPSCAKTWT  652
               SC V   LHV  CW LV KSGL     + A SLS+W+  + L  ++  S +C++T
Sbjct  185  LITSCVV-FCLHVPLCWLLVYKSGLDHIGGALALSLSYWLYAIFLGSFMYFSSACSETRA  243

Query  653  GFSPEALENVMVFL  694
              + E  E V  F+
Sbjct  244  PLTMEIFEGVREFI  257


>hypothetical protein AXX17_AT1G65340 [Arabidopsis thaliana]
Sequence ID: OAP18050.1 Length: 514
Range 1: 120 to 251

Score:69.7 bits(169), Expect:2e-12,
Method:Compositional matrix adjust.,
Identities:45/132(34%), Positives:69/132(52%), Gaps:5/132(3%)

Query  311  LPLSIIWINVSSILVFLDQDAIIA*TFGSYIVFLIPDLIGCVIL----KCFSLRAKTECF  478
            +PLS++W  +  IL  + QDA++A   G +  +LIP L G   L    + F  ++
Sbjct  120  IPLSVLWSYIGDILSLIGQDAMVAQEAGKFATWLIPALFGYATLQPLVRFFQAQSLILPL  179

Query  479  LSCSVPVL-LHVHACWGLVLKSGLGIREASQATSLSFWVNVVLLSIYVKLSPSCAKTWTG  655
            +  SV  L +H+  CW LV K GLG   A+ A  +S+W+NV +L +Y+  S SC K+
Sbjct  180  VMSSVSSLCVHIVLCWSLVFKFGLGSLGAAIAIGVSYWLNVTVLGLYMTFSSSCRKSRAP  239

Query  656  FSPEALENVMVF  691
             S    E +  F
Sbjct  240  ISMSLFEGMGEF  251


>hypothetical protein AXX17_AT2G03130 [Arabidopsis thaliana]
Sequence ID: OAP10784.1 Length: 296
Range 1: 126 to 296

Score:68.2 bits(165), Expect:3e-12,
Method:Compositional matrix adjust.,
Identities:58/199(29%), Positives:90/199(45%), Gaps:33/199(16%)

Query  317  LSIIWINVSSILVFLDQDAIIA*TFGSYIVFLIPDLIGCVILKCFSLRAKTECF----LS  484
            +SI+W  +  +L+   QD  I+   GSY V LIP L+   + +  +   +T+      L
Sbjct  126  ISILWFYMDKLLISFGQDPDISKVAGSYAVCLIPALLAQAVQQPLTRFLQTQGLVLPLLY  185

Query  485  CSVPVLL-HVHACWGLVLKSGLGIREASQATSLSFWVNVVLLSIYVKLSPSCAKTWTGFS  661
            C++  LL H+  C  LV   GLG   A+ A  LS+W NV++L++YV+ S +C KT    S
Sbjct  186  CAITTLLFHIPVCLILVYAFGLGSNGAALAIGLSYWFNVLILALYVRFSSACEKTRGFVS  245

Query  662  PEALENVMVFLWSSICIHGLVRLVVHIYDSCLLVYIISLMLIIS*WNHNDLVIQENGLPL  841
             + + +V  F    I    +  +                      W+  +L+I  +GL
Sbjct  246  DDFVLSVKQFFQYGIPSAAMTTIE---------------------WSLFELLILSSGL--  282

Query  842  SLRLYFQNPVLETSVFPTW  898
                   NP LETSV   W
Sbjct  283  -----LPNPKLETSVLSIW  296


>At1g66760/F4N21_11 [Arabidopsis thaliana]
Sequence ID: AAK97692.1 Length: 265
>At1g66760/F4N21_11 [Arabidopsis thaliana]
Sequence ID: AAQ22646.1 Length: 265 >AT1G66760 [Arabidopsis thaliana]
Sequence ID: BAH56840.1 Length: 265
Range 1: 122 to 254

Score:66.6 bits(161), Expect:8e-12,
Method:Compositional matrix adjust.,
Identities:40/133(30%), Positives:69/133(51%), Gaps:5/133(3%)

Query  311  LPLSIIWINVSSILVFLDQDAIIA*TFGSYIVFLIPDLIGCVILKCFSLRAKTECF----  478
            +P+SI+W+ ++ IL+ L QD  IA   G Y ++L+P L G  +L+      +++
Sbjct  122  VPISILWMFMNQILLLLHQDPQIAELAGVYCLWLVPALFGYSVLESLVRYFQSQSLIYPM  181

Query  479  -LSCSVPVLLHVHACWGLVLKSGLGIREASQATSLSFWVNVVLLSIYVKLSPSCAKTWTG  655
             LS    +  HV  CW +V K   G + A+ +  +S+W+N V L +Y+K S  C +T
Sbjct  182  VLSSLAALSFHVPLCWLMVHKFDFGAKGAAASIGISYWLNAVFLWVYMKRSSRCVETRIY  241

Query  656  FSPEALENVMVFL  694
             S +   +  +F
Sbjct  242  MSKDVFVHTNIFF  254


>hypothetical protein [Arabidopsis thaliana]
Sequence ID: AAD28682.1 Length: 480
Range 1: 126 to 240

Score:67.4 bits(163), Expect:1e-11,
Method:Compositional matrix adjust.,
Identities:43/115(37%), Positives:64/115(55%), Gaps:5/115(4%)

Query  317  LSIIWINVSSILVFLDQDAIIA*TFGSYIVFLIPDLIGCVILKCFSLRAKTECF----LS  484
            +SI+W  +  + V L QD  I+   GSY V LIP L+   + +  +   +T+      L
Sbjct  126  ISILWFYMDKLFVSLGQDPDISKVAGSYAVCLIPALLAQAVQQPLTRFLQTQGLVLPLLY  185

Query  485  CSVPVLL-HVHACWGLVLKSGLGIREASQATSLSFWVNVVLLSIYVKLSPSCAKT  646
            C++  LL H+  C  LV   GLG   A+ A  LS+W NV++L++YV+ S SC KT
Sbjct  186  CAITTLLFHIPVCLILVYAFGLGSNGAALAIGLSYWFNVLILALYVRFSSSCEKT  240


>MATE efflux family protein [Arabidopsis thaliana]
Sequence ID: NP_001324928.1 Length: 471
>MATE efflux family protein [Arabidopsis thaliana]
Sequence ID: ANM62799.1 Length: 471
Range 1: 126 to 240

Score:67.4 bits(163), Expect:1e-11,
Method:Compositional matrix adjust.,
Identities:43/115(37%), Positives:64/115(55%), Gaps:5/115(4%)

Query  317  LSIIWINVSSILVFLDQDAIIA*TFGSYIVFLIPDLIGCVILKCFSLRAKTECF----LS  484
            +SI+W  +  + V L QD  I+   GSY V LIP L+   + +  +   +T+      L
Sbjct  126  ISILWFYMDKLFVSLGQDPDISKVAGSYAVCLIPALLAQAVQQPLTRFLQTQGLVLPLLY  185

Query  485  CSVPVLL-HVHACWGLVLKSGLGIREASQATSLSFWVNVVLLSIYVKLSPSCAKT  646
            C++  LL H+  C  LV   GLG   A+ A  LS+W NV++L++YV+ S SC KT
Sbjct  186  CAITTLLFHIPVCLILVYAFGLGSNGAALAIGLSYWFNVLILALYVRFSSSCEKT  240


>MATE efflux family protein [Arabidopsis thaliana]
Sequence ID: NP_178499.2 Length: 483
>RecName: Full=Protein DETOXIFICATION 6; Short=AtDTX6; AltName: Full=Multidrug and toxic compound extrusion protein 6; Short=MATE protein 6 [Arabidopsis thaliana]
Sequence ID: Q8RWF5.1 Length: 483 >unknown protein [Arabidopsis thaliana]
Sequence ID: AAM13125.1 Length: 483 >At2g04100 [Arabidopsis thaliana]
Sequence ID: AAP31968.1 Length: 483 >MATE efflux family protein [Arabidopsis thaliana]
Sequence ID: AEC05799.1 Length: 483 >unnamed protein product [Arabidopsis thaliana]
Sequence ID: CAA0357037.1 Length: 483 >unnamed protein product [Arabidopsis thaliana]
Sequence ID: VYS52001.1 Length: 483
Range 1: 126 to 240

Score:67.4 bits(163), Expect:1e-11,
Method:Compositional matrix adjust.,
Identities:43/115(37%), Positives:64/115(55%), Gaps:5/115(4%)

Query  317  LSIIWINVSSILVFLDQDAIIA*TFGSYIVFLIPDLIGCVILKCFSLRAKTECF----LS  484
            +SI+W  +  + V L QD  I+   GSY V LIP L+   + +  +   +T+      L
Sbjct  126  ISILWFYMDKLFVSLGQDPDISKVAGSYAVCLIPALLAQAVQQPLTRFLQTQGLVLPLLY  185

Query  485  CSVPVLL-HVHACWGLVLKSGLGIREASQATSLSFWVNVVLLSIYVKLSPSCAKT  646
            C++  LL H+  C  LV   GLG   A+ A  LS+W NV++L++YV+ S SC KT
Sbjct  186  CAITTLLFHIPVCLILVYAFGLGSNGAALAIGLSYWFNVLILALYVRFSSSCEKT  240


>MATE efflux family protein [Arabidopsis thaliana]
Sequence ID: NP_001117293.1 Length: 410
>MATE efflux family protein [Arabidopsis thaliana]
Sequence ID: AEE29279.1 Length: 410
Range 1: 9 to 140

Score:67.4 bits(163), Expect:1e-11,
Method:Compositional matrix adjust.,
Identities:47/137(34%), Positives:68/137(49%), Gaps:15/137(10%)

Query  311  LPLSIIWINVSSILVFLDQDAIIA*TFGSYIVFLIPDLIGCVILKCFSLRAK--------  466
            LPL++IW+N+ ++LVFL QD  IA   G Y   LIP L    +L+  +   +
Sbjct  9    LPLTLIWLNMETLLVFLGQDPSIAHEAGRYAACLIPGLFAYAVLQPLTRYFQNQSMITPL  68

Query  467  --TECFLSCSVPVLLHVHACWGLVLKSGLGIREASQATSLSFWVNVVLLSIYVKLSPSCA  640
              T CF+ C     LHV  CW LV KSGLG    + A S S  +  ++L   +  S +C+
Sbjct  69   LITSCFVFC-----LHVPLCWLLVYKSGLGNLGGALALSFSNCLYTIILGSLMCFSSACS  123

Query  641  KTWTGFSPEALENVMVF  691
            +T    S E  + +  F
Sbjct  124  ETRAPLSMEIFDGIGEF  140


>unnamed protein product [Arabidopsis thaliana]
Sequence ID: VYS51999.1 Length: 476
Range 1: 123 to 263

Score:67.4 bits(163), Expect:1e-11,
Method:Compositional matrix adjust.,
Identities:50/141(35%), Positives:70/141(49%), Gaps:5/141(3%)

Query  317  LSIIWINVSSILVFLDQDAIIA*TFGSYIVFLIPDLIGCVI---LKCFSLRAK--TECFL  481
            +SI+WI +  +L+ L QD  I+   GSY ++LIP L    I   L  F L
Sbjct  123  ISILWIYIEKLLISLGQDPNISRVAGSYSLWLIPALFAHAIFLPLTRFLLAQGLVIPLLY  182

Query  482  SCSVPVLLHVHACWGLVLKSGLGIREASQATSLSFWVNVVLLSIYVKLSPSCAKTWTGFS  661
            S    +L H+  CW LV   GLG   A+ A SLSFW   V+LS +V+ S SC KT    S
Sbjct  183  SAMTTLLFHIAVCWTLVFALGLGSNGAAIAISLSFWFYAVILSCHVRFSSSCEKTRGFVS  242

Query  662  PEALENVMVFLWSSICIHGLV  724
             + + +V  +    +   GL+
Sbjct  243  NDFMSSVKQYFQYGVPSAGLI  263


>hypothetical protein AXX17_AT2G03120 [Arabidopsis thaliana]
Sequence ID: OAP07829.1 Length: 476
Range 1: 123 to 263

Score:67.0 bits(162), Expect:2e-11,
Method:Compositional matrix adjust.,
Identities:49/141(35%), Positives:70/141(49%), Gaps:5/141(3%)

Query  317  LSIIWINVSSILVFLDQDAIIA*TFGSYIVFLIPDLIGCVI---LKCFSLRAK--TECFL  481
            +SI+WI +  +L+ L QD  I+   GSY ++L+P L    I   L  F L
Sbjct  123  ISILWIYIEKLLITLGQDPDISRVAGSYSLWLVPALFAHAIFLPLTRFLLAQGLVIPLLY  182

Query  482  SCSVPVLLHVHACWGLVLKSGLGIREASQATSLSFWVNVVLLSIYVKLSPSCAKTWTGFS  661
            S    +L H+  CW LV   GLG   A+ A SLSFW   V+LS +V+ S SC KT    S
Sbjct  183  SAMTTLLFHIAVCWTLVFALGLGSNGAAIAISLSFWFYAVILSCHVRFSSSCEKTRGFVS  242

Query  662  PEALENVMVFLWSSICIHGLV  724
             + + +V  +    +   GL+
Sbjct  243  NDFMSSVKQYFQYGVPSAGLI  263


>hypothetical protein AXX17_AT2G03150 [Arabidopsis thaliana]
Sequence ID: OAP09725.1 Length: 483
Range 1: 126 to 240

Score:66.6 bits(161), Expect:2e-11,
Method:Compositional matrix adjust.,
Identities:42/115(37%), Positives:64/115(55%), Gaps:5/115(4%)

Query  317  LSIIWINVSSILVFLDQDAIIA*TFGSYIVFLIPDLIGCVILKCFSLRAKTECF----LS  484
            +SI+W  +  + V L QD  I+   GSY V LIP L+   + +  +   +T+      L
Sbjct  126  ISILWFYMDKLFVSLGQDPDISKVAGSYAVCLIPALLAQAVQQPLTRFLQTQGLVLPLLY  185

Query  485  CSVPVLL-HVHACWGLVLKSGLGIREASQATSLSFWVNVVLLSIYVKLSPSCAKT  646
            C++  LL H+  C  LV   GLG   A+ A  LS+W NV++L++YV+ S +C KT
Sbjct  186  CAITTLLFHIPVCLILVYAFGLGSNGAALAIGLSYWFNVLILALYVRFSSACEKT  240


>Strong similarity to gi|4734005 F3L12.7 hypothetical protein from Arabidopsis thaliana BAC gb|AC007178 [Arabidopsis thaliana]
Sequence ID: AAD39646.1 Length: 480
Range 1: 129 to 260

Score:66.6 bits(161), Expect:3e-11,
Method:Compositional matrix adjust.,
Identities:47/137(34%), Positives:68/137(49%), Gaps:15/137(10%)

Query  311  LPLSIIWINVSSILVFLDQDAIIA*TFGSYIVFLIPDLIGCVILKCFSLRAK--------  466
            LPL++IW+N+ ++LVFL QD  IA   G Y   LIP L    +L+  +   +
Sbjct  129  LPLTLIWLNMETLLVFLGQDPSIAHEAGRYAACLIPGLFAYAVLQPLTRYFQNQSMITPL  188

Query  467  --TECFLSCSVPVLLHVHACWGLVLKSGLGIREASQATSLSFWVNVVLLSIYVKLSPSCA  640
              T CF+ C     LHV  CW LV KSGLG    + A S S  +  ++L   +  S +C+
Sbjct  189  LITSCFVFC-----LHVPLCWLLVYKSGLGNLGGALALSFSNCLYTIILGSLMCFSSACS  243

Query  641  KTWTGFSPEALENVMVF  691
            +T    S E  + +  F
Sbjct  244  ETRAPLSMEIFDGIGEF  260


>unnamed protein product [Arabidopsis thaliana]
Sequence ID: CAA0357024.1 Length: 486
Range 1: 126 to 240

Score:66.2 bits(160), Expect:3e-11,
Method:Compositional matrix adjust.,
Identities:42/115(37%), Positives:64/115(55%), Gaps:5/115(4%)

Query  317  LSIIWINVSSILVFLDQDAIIA*TFGSYIVFLIPDLIGCVILKCFSLRAKTECF----LS  484
            +SI+W  +  +L+   QD  I+   GSY V LIP L+   + +  +   +T+      L
Sbjct  126  ISILWFYMDKLLISFGQDPDISKVAGSYAVCLIPALLAQAVQQPLTRFLQTQGLVLPLLY  185

Query  485  CSVPVLL-HVHACWGLVLKSGLGIREASQATSLSFWVNVVLLSIYVKLSPSCAKT  646
            C++  LL H+  C  LV   GLG   A+ A  LS+W NV++L++YV+ S SC KT
Sbjct  186  CAITTLLFHIPVCLILVYAFGLGSNGAALAIGLSYWFNVLILALYVRFSSSCEKT  240


>unnamed protein product [Arabidopsis thaliana]
Sequence ID: VYS52000.1 Length: 477
Range 1: 126 to 240

Score:66.2 bits(160), Expect:3e-11,
Method:Compositional matrix adjust.,
Identities:42/115(37%), Positives:64/115(55%), Gaps:5/115(4%)

Query  317  LSIIWINVSSILVFLDQDAIIA*TFGSYIVFLIPDLIGCVILKCFSLRAKTECF----LS  484
            +SI+W  +  +L+   QD  I+   GSY V LIP L+   + +  +   +T+      L
Sbjct  126  ISILWFYMDKLLISFGQDPDISKVAGSYAVCLIPALLAQAVQQPLTRFLQTQGLVLPLLY  185

Query  485  CSVPVLL-HVHACWGLVLKSGLGIREASQATSLSFWVNVVLLSIYVKLSPSCAKT  646
            C++  LL H+  C  LV   GLG   A+ A  LS+W NV++L++YV+ S SC KT
Sbjct  186  CAITTLLFHIPVCLILVYAFGLGSNGAALAIGLSYWFNVLILALYVRFSSSCEKT  240


>MATE efflux family protein [Arabidopsis thaliana]
Sequence ID: NP_563964.1 Length: 482
>RecName: Full=Protein DETOXIFICATION 13; Short=AtDTX13; AltName: Full=Multidrug and toxic compound extrusion protein 13; Short=MATE protein 13 [Arabidopsis thaliana]
Sequence ID: Q94AL1.1 Length: 482 >unknown protein [Arabidopsis thaliana]
Sequence ID: AAK76631.1 Length: 482 >unknown protein [Arabidopsis thaliana]
Sequence ID: AAL85036.1 Length: 482 >MATE efflux family protein [Arabidopsis thaliana]
Sequence ID: AEE29278.1 Length: 482 >unnamed protein product [Arabidopsis thaliana]
Sequence ID: CAA0206660.1 Length: 482 >unnamed protein product [Arabidopsis thaliana]
Sequence ID: VYS46147.1 Length: 482
Range 1: 129 to 260

Score:66.2 bits(160), Expect:3e-11,
Method:Compositional matrix adjust.,
Identities:47/137(34%), Positives:68/137(49%), Gaps:15/137(10%)

Query  311  LPLSIIWINVSSILVFLDQDAIIA*TFGSYIVFLIPDLIGCVILKCFSLRAK--------  466
            LPL++IW+N+ ++LVFL QD  IA   G Y   LIP L    +L+  +   +
Sbjct  129  LPLTLIWLNMETLLVFLGQDPSIAHEAGRYAACLIPGLFAYAVLQPLTRYFQNQSMITPL  188

Query  467  --TECFLSCSVPVLLHVHACWGLVLKSGLGIREASQATSLSFWVNVVLLSIYVKLSPSCA  640
              T CF+ C     LHV  CW LV KSGLG    + A S S  +  ++L   +  S +C+
Sbjct  189  LITSCFVFC-----LHVPLCWLLVYKSGLGNLGGALALSFSNCLYTIILGSLMCFSSACS  243

Query  641  KTWTGFSPEALENVMVF  691
            +T    S E  + +  F
Sbjct  244  ETRAPLSMEIFDGIGEF  260


>hypothetical protein [Arabidopsis thaliana]
Sequence ID: AAD28683.1 Length: 483
Range 1: 126 to 240

Score:65.9 bits(159), Expect:4e-11,
Method:Compositional matrix adjust.,
Identities:42/115(37%), Positives:64/115(55%), Gaps:5/115(4%)

Query  317  LSIIWINVSSILVFLDQDAIIA*TFGSYIVFLIPDLIGCVILKCFSLRAKTECF----LS  484
            +SI+W  +  + V L QD  I+   GSY V LIP L+   + +  +   +T+      L
Sbjct  126  ISILWFYMDKLFVSLGQDPDISKVAGSYAVCLIPALLAQAVQQPLTRFLQTQGLVLPLLY  185

Query  485  CSVPVLL-HVHACWGLVLKSGLGIREASQATSLSFWVNVVLLSIYVKLSPSCAKT  646
            C++  LL H+  C  LV   GLG   A+ A  LS+W NV++L++YV+ S +C KT
Sbjct  186  CAITTLLFHIPVCLILVYAFGLGSNGAALAIGLSYWFNVLILALYVRFSSACEKT  240


>MATE efflux family protein [Arabidopsis thaliana]
Sequence ID: NP_178498.2 Length: 477
>RecName: Full=Protein DETOXIFICATION 5; Short=AtDTX5; AltName: Full=Multidrug and toxic compound extrusion protein 5; Short=MATE protein 5 [Arabidopsis thaliana]
Sequence ID: Q9SIA1.2 Length: 477 >MATE efflux family protein [Arabidopsis thaliana]
Sequence ID: AEC05798.1 Length: 477
Range 1: 126 to 240

Score:65.9 bits(159), Expect:4e-11,
Method:Compositional matrix adjust.,
Identities:42/115(37%), Positives:64/115(55%), Gaps:5/115(4%)

Query  317  LSIIWINVSSILVFLDQDAIIA*TFGSYIVFLIPDLIGCVILKCFSLRAKTECF----LS  484
            +SI+W  +  + V L QD  I+   GSY V LIP L+   + +  +   +T+      L
Sbjct  126  ISILWFYMDKLFVSLGQDPDISKVAGSYAVCLIPALLAQAVQQPLTRFLQTQGLVLPLLY  185

Query  485  CSVPVLL-HVHACWGLVLKSGLGIREASQATSLSFWVNVVLLSIYVKLSPSCAKT  646
            C++  LL H+  C  LV   GLG   A+ A  LS+W NV++L++YV+ S +C KT
Sbjct  186  CAITTLLFHIPVCLILVYAFGLGSNGAALAIGLSYWFNVLILALYVRFSSACEKT  240


>MATE efflux family protein [Arabidopsis thaliana]
Sequence ID: NP_849854.1 Length: 482
>RecName: Full=Protein DETOXIFICATION 9; Short=AtDTX9; AltName: Full=Multidrug and toxic compound extrusion protein 9; Short=MATE protein 9 [Arabidopsis thaliana]
Sequence ID: Q9C9M8.1 Length: 482 >MATE efflux family protein, putative [Arabidopsis thaliana]
Sequence ID: AAG60073.1 Length: 482 >MATE efflux family protein [Arabidopsis thaliana]
Sequence ID: AEE34553.1 Length: 482 >unnamed protein product [Arabidopsis thaliana]
Sequence ID: CAA0319956.1 Length: 482 >unnamed protein product [Arabidopsis thaliana]
Sequence ID: VYS50216.1 Length: 482
Range 1: 122 to 254

Score:65.5 bits(158), Expect:6e-11,
Method:Compositional matrix adjust.,
Identities:40/133(30%), Positives:69/133(51%), Gaps:5/133(3%)

Query  311  LPLSIIWINVSSILVFLDQDAIIA*TFGSYIVFLIPDLIGCVILKCFSLRAKTECF----  478
            +P+SI+W+ ++ IL+ L QD  IA   G Y ++L+P L G  +L+      +++
Sbjct  122  VPISILWMFMNQILLLLHQDPQIAELAGVYCLWLVPALFGYSVLESLVRYFQSQSLIYPM  181

Query  479  -LSCSVPVLLHVHACWGLVLKSGLGIREASQATSLSFWVNVVLLSIYVKLSPSCAKTWTG  655
             LS    +  HV  CW +V K   G + A+ +  +S+W+N V L +Y+K S  C +T
Sbjct  182  VLSSLAALSFHVPLCWLMVHKFDFGAKGAAASIGISYWLNAVFLWVYMKRSSRCVETRIY  241

Query  656  FSPEALENVMVFL  694
             S +   +  +F
Sbjct  242  MSKDVFVHTNIFF  254


>putative MATE efflux family protein [Arabidopsis thaliana]
Sequence ID: AAK25964.1 Length: 482
Range 1: 122 to 254

Score:65.5 bits(158), Expect:6e-11,
Method:Compositional matrix adjust.,
Identities:40/133(30%), Positives:69/133(51%), Gaps:5/133(3%)

Query  311  LPLSIIWINVSSILVFLDQDAIIA*TFGSYIVFLIPDLIGCVILKCFSLRAKTECF----  478
            +P+SI+W+ ++ IL+ L QD  IA   G Y ++L+P L G  +L+      +++
Sbjct  122  VPISILWMFMNQILLLLHQDPQIAELAGVYCLWLVPALFGYSVLESLVRYFQSQSLIYPM  181

Query  479  -LSCSVPVLLHVHACWGLVLKSGLGIREASQATSLSFWVNVVLLSIYVKLSPSCAKTWTG  655
             LS    +  HV  CW +V K   G + A+ +  +S+W+N V L +Y+K S  C +T
Sbjct  182  VLSSLAALSFHVPLCWLMVHKFDFGAKGAAASIGISYWLNAVFLWVYMKRSSRCVETRIY  241

Query  656  FSPEALENVMVFL  694
             S +   +  +F
Sbjct  242  MSKDVFVHTNIFF  254


>hypothetical protein AXX17_AT1G60540 [Arabidopsis thaliana]
Sequence ID: OAP16837.1 Length: 482
Range 1: 122 to 254

Score:65.5 bits(158), Expect:6e-11,
Method:Compositional matrix adjust.,
Identities:40/133(30%), Positives:69/133(51%), Gaps:5/133(3%)

Query  311  LPLSIIWINVSSILVFLDQDAIIA*TFGSYIVFLIPDLIGCVILKCFSLRAKTECF----  478
            +P+SI+W+ ++ IL+ L QD  IA   G Y ++L+P L G  +L+      +++
Sbjct  122  VPISILWMFMNQILLLLHQDPQIAELAGVYCLWLVPALFGYSVLESLVRYFQSQSLIYPM  181

Query  479  -LSCSVPVLLHVHACWGLVLKSGLGIREASQATSLSFWVNVVLLSIYVKLSPSCAKTWTG  655
             LS    +  HV  CW +V K   G + A+ +  +S+W+N V L +Y+K S  C +T
Sbjct  182  VLSSLAALSFHVPLCWLMVHKFDFGAKGAAASIGISYWLNAVFLWVYMKRSSRCVETRIY  241

Query  656  FSPEALENVMVFL  694
             S +   +  +F
Sbjct  242  MSKDVFVHTNIFF  254


>hypothetical protein AXX17_AT2G03130 [Arabidopsis thaliana]
Sequence ID: OAP10785.1 Length: 486
Range 1: 126 to 240

Score:65.5 bits(158), Expect:6e-11,
Method:Compositional matrix adjust.,
Identities:41/115(36%), Positives:64/115(55%), Gaps:5/115(4%)

Query  317  LSIIWINVSSILVFLDQDAIIA*TFGSYIVFLIPDLIGCVILKCFSLRAKTECF----LS  484
            +SI+W  +  +L+   QD  I+   GSY V LIP L+   + +  +   +T+      L
Sbjct  126  ISILWFYMDKLLISFGQDPDISKVAGSYAVCLIPALLAQAVQQPLTRFLQTQGLVLPLLY  185

Query  485  CSVPVLL-HVHACWGLVLKSGLGIREASQATSLSFWVNVVLLSIYVKLSPSCAKT  646
            C++  LL H+  C  LV   GLG   A+ A  LS+W NV++L++YV+ S +C KT
Sbjct  186  CAITTLLFHIPVCLILVYAFGLGSNGAALAIGLSYWFNVLILALYVRFSSACEKT  240


>MATE efflux family protein [Arabidopsis thaliana]
Sequence ID: NP_564883.2 Length: 466
>MATE efflux family protein [Arabidopsis thaliana]
Sequence ID: AEE34552.1 Length: 466
Range 1: 122 to 254

Score:65.1 bits(157), Expect:7e-11,
Method:Compositional matrix adjust.,
Identities:40/133(30%), Positives:69/133(51%), Gaps:5/133(3%)

Query  311  LPLSIIWINVSSILVFLDQDAIIA*TFGSYIVFLIPDLIGCVILKCFSLRAKTECF----  478
            +P+SI+W+ ++ IL+ L QD  IA   G Y ++L+P L G  +L+      +++
Sbjct  122  VPISILWMFMNQILLLLHQDPQIAELAGVYCLWLVPALFGYSVLESLVRYFQSQSLIYPM  181

Query  479  -LSCSVPVLLHVHACWGLVLKSGLGIREASQATSLSFWVNVVLLSIYVKLSPSCAKTWTG  655
             LS    +  HV  CW +V K   G + A+ +  +S+W+N V L +Y+K S  C +T
Sbjct  182  VLSSLAALSFHVPLCWLMVHKFDFGAKGAAASIGISYWLNAVFLWVYMKRSSRCVETRIY  241

Query  656  FSPEALENVMVFL  694
             S +   +  +F
Sbjct  242  MSKDVFVHTNIFF  254


>MATE efflux family protein [Arabidopsis thaliana]
Sequence ID: NP_001321857.1 Length: 393
>MATE efflux family protein [Arabidopsis thaliana]
Sequence ID: ANM59503.1 Length: 393
Range 1: 122 to 254

Score:64.7 bits(156), Expect:8e-11,
Method:Compositional matrix adjust.,
Identities:40/133(30%), Positives:69/133(51%), Gaps:5/133(3%)

Query  311  LPLSIIWINVSSILVFLDQDAIIA*TFGSYIVFLIPDLIGCVILKCFSLRAKTECF----  478
            +P+SI+W+ ++ IL+ L QD  IA   G Y ++L+P L G  +L+      +++
Sbjct  122  VPISILWMFMNQILLLLHQDPQIAELAGVYCLWLVPALFGYSVLESLVRYFQSQSLIYPM  181

Query  479  -LSCSVPVLLHVHACWGLVLKSGLGIREASQATSLSFWVNVVLLSIYVKLSPSCAKTWTG  655
             LS    +  HV  CW +V K   G + A+ +  +S+W+N V L +Y+K S  C +T
Sbjct  182  VLSSLAALSFHVPLCWLMVHKFDFGAKGAAASIGISYWLNAVFLWVYMKRSSRCVETRIY  241

Query  656  FSPEALENVMVFL  694
             S +   +  +F
Sbjct  242  MSKDVFVHTNIFF  254


>hypothetical protein AXX17_AT2G03140 [Arabidopsis thaliana]
Sequence ID: OAP11455.1 Length: 711
Range 1: 126 to 240

Score:65.1 bits(157), Expect:1e-10,
Method:Compositional matrix adjust.,
Identities:41/115(36%), Positives:64/115(55%), Gaps:5/115(4%)

Query  317  LSIIWINVSSILVFLDQDAIIA*TFGSYIVFLIPDLIGCVILKCFSLRAKTECF----LS  484
            +SI+W  +  +L+   QD  I+   GSY V LIP L+   + +  +   +T+      L
Sbjct  126  ISILWFYMDKLLISFGQDPDISKVAGSYAVCLIPALLAQAVQQPLTRFLQTQGLVLPLLY  185

Query  485  CSVPVLL-HVHACWGLVLKSGLGIREASQATSLSFWVNVVLLSIYVKLSPSCAKT  646
            C++  LL H+  C  LV   GLG   A+ A  LS+W NV++L++YV+ S +C KT
Sbjct  186  CAITTLLFHIPVCLILVYAFGLGSNGAALAIGLSYWFNVLILALYVRFSSACEKT  240


>unnamed protein product [Arabidopsis thaliana]
Sequence ID: CAA0356956.1 Length: 476
Range 1: 123 to 263

Score:64.3 bits(155), Expect:1e-10,
Method:Compositional matrix adjust.,
Identities:47/141(33%), Positives:69/141(48%), Gaps:5/141(3%)

Query  317  LSIIWINVSSILVFLDQDAIIA*TFGSYIVFLIPDLIGCVI---LKCFSLRAK--TECFL  481
            +SI+WI +  +L+ L QD  I+   GSY ++L+P L    I   L  F L
Sbjct  123  ISILWIYIEKLLITLGQDPDISRVAGSYSLWLVPALFAHAIFLPLTRFLLAQGLVISLLY  182

Query  482  SCSVPVLLHVHACWGLVLKSGLGIREASQATSLSFWVNVVLLSIYVKLSPSCAKTWTGFS  661
            S    +L H+  CW LV   GLG   A+ A SLSFW   V+LS +V+   SC KT    S
Sbjct  183  SAMTTLLFHIAVCWTLVFALGLGSNGAAIAISLSFWFYAVILSCHVRFFSSCEKTRGFVS  242

Query  662  PEALENVMVFLWSSICIHGLV  724
             + + ++  +    +   GL+
Sbjct  243  NDFMSSIKQYFQYGVPSAGLI  263


>unnamed protein product [Arabidopsis thaliana]
Sequence ID: CAA0357011.1 Length: 476
Range 1: 123 to 263

Score:63.9 bits(154), Expect:2e-10,
Method:Compositional matrix adjust.,
Identities:47/141(33%), Positives:69/141(48%), Gaps:5/141(3%)

Query  317  LSIIWINVSSILVFLDQDAIIA*TFGSYIVFLIPDLIGCVI---LKCFSLRAK--TECFL  481
            +SI+WI +  +L+ L QD  I+   GSY ++L+P L    I   L  F L
Sbjct  123  ISILWIYIEKLLITLGQDPDISRVAGSYSLWLVPALFAHAIFLPLTRFLLAQGLVISLLY  182

Query  482  SCSVPVLLHVHACWGLVLKSGLGIREASQATSLSFWVNVVLLSIYVKLSPSCAKTWTGFS  661
            S    +L H+  CW LV   GLG   A+ A SLSFW   V+LS +V+   SC KT    S
Sbjct  183  SAMTTLLFHIAVCWTLVFALGLGSNGAAIAISLSFWFYAVILSCHVRFFSSCEKTRGFVS  242

Query  662  PEALENVMVFLWSSICIHGLV  724
             + + ++  +    +   GL+
Sbjct  243  NDFMSSIKQYFQYGVPSAGLI  263


>hypothetical protein AXX17_AT3G25120 [Arabidopsis thaliana]
Sequence ID: OAP06402.1 Length: 469
Range 1: 127 to 284

Score:63.5 bits(153), Expect:2e-10,
Method:Compositional matrix adjust.,
Identities:49/160(31%), Positives:83/160(51%), Gaps:7/160(4%)

Query  317  LSIIWINVSSILVFLDQDAIIA*TFGSYIVFLIPDLIGCVILKCFSLRAKTECFLSCSV-  493
            ++I+W    S+ + L QD  I+     Y+ +L P L+    L+      +T+C ++  V
Sbjct  127  ITILWFFTESVFLLLRQDPSISKQAALYMKYLAPGLLAYGFLQNILRFCQTQCIVTPLVL  186

Query  494  ----PVLLHVHACWGLVLKSGLGIREASQATSLSFWVNVVLLSIYVKLSPSCAKTWTGFS  661
                P+++++   + LV  +GLG   A  ATS+S W+  V L  YV  S    +TWTGFS
Sbjct  187  FSFLPLVINIGTTYALVHLAGLGFIGAPIATSISLWIAFVSLGFYVICSDKFKETWTGFS  246

Query  662  PEALENVMVFLWSSICIHGLVRLVVHIYDSCLLVYIISLM  781
             E+  +V++ L  SI    +V L    ++  +LV++  LM
Sbjct  247  MESFHHVVLNLTLSIPSAAMVCLEYWAFE--ILVFLAGLM  284


>unnamed protein product [Arabidopsis thaliana]
Sequence ID: VYS58370.1 Length: 469
Range 1: 127 to 284

Score:63.5 bits(153), Expect:2e-10,
Method:Compositional matrix adjust.,
Identities:49/160(31%), Positives:83/160(51%), Gaps:7/160(4%)

Query  317  LSIIWINVSSILVFLDQDAIIA*TFGSYIVFLIPDLIGCVILKCFSLRAKTECFLSCSV-  493
            ++I+W    S+ + L QD  I+     Y+ +L P L+    L+      +T+C ++  V
Sbjct  127  ITILWFFTESVFLLLRQDPSISKQAALYMKYLAPGLLAYGFLQNILRFCQTQCIVTPLVL  186

Query  494  ----PVLLHVHACWGLVLKSGLGIREASQATSLSFWVNVVLLSIYVKLSPSCAKTWTGFS  661
                P+++++   + LV  +GLG   A  ATS+S W+  V L  YV  S    +TWTGFS
Sbjct  187  FSFLPLVINIGTTYALVHLAGLGFIGAPIATSISLWIAFVSLGFYVICSDKFKETWTGFS  246

Query  662  PEALENVMVFLWSSICIHGLVRLVVHIYDSCLLVYIISLM  781
             E+  +V++ L  SI    +V L    ++  +LV++  LM
Sbjct  247  MESFHHVVLNLTLSIPSAAMVCLEYWAFE--ILVFLAGLM  284


>MATE efflux family protein [Arabidopsis thaliana]
Sequence ID: NP_188997.1 Length: 469
>RecName: Full=Protein DETOXIFICATION 18; Short=AtDTX18; AltName: Full=Multidrug and toxic compound extrusion protein 18; Short=MATE protein 18; AltName: Full=Protein LIKE ALF5 [Arabidopsis thaliana]
Sequence ID: Q9LUH3.1 Length: 469 >unknown protein [Arabidopsis thaliana]
Sequence ID: AAL49789.1 Length: 469 >unknown protein [Arabidopsis thaliana]
Sequence ID: AAM20025.1 Length: 469 >MATE efflux family protein [Arabidopsis thaliana]
Sequence ID: AEE76776.1 Length: 469 >unnamed protein product [Arabidopsis thaliana]
Sequence ID: BAB02773.1 Length: 469
Range 1: 127 to 284

Score:63.5 bits(153), Expect:2e-10,
Method:Compositional matrix adjust.,
Identities:49/160(31%), Positives:83/160(51%), Gaps:7/160(4%)

Query  317  LSIIWINVSSILVFLDQDAIIA*TFGSYIVFLIPDLIGCVILKCFSLRAKTECFLSCSV-  493
            ++I+W    S+ + L QD  I+     Y+ +L P L+    L+      +T+C ++  V
Sbjct  127  ITILWFFTESVFLLLRQDPSISKQAALYMKYLAPGLLAYGFLQNILRFCQTQCIVTPLVL  186

Query  494  ----PVLLHVHACWGLVLKSGLGIREASQATSLSFWVNVVLLSIYVKLSPSCAKTWTGFS  661
                P+++++   + LV  +GLG   A  ATS+S W+  V L  YV  S    +TWTGFS
Sbjct  187  FSFLPLVINIGTTYALVHLAGLGFIGAPIATSISLWIAFVSLGFYVICSDKFKETWTGFS  246

Query  662  PEALENVMVFLWSSICIHGLVRLVVHIYDSCLLVYIISLM  781
             E+  +V++ L  SI    +V L    ++  +LV++  LM
Sbjct  247  MESFHHVVLNLTLSIPSAAMVCLEYWAFE--ILVFLAGLM  284


>MATE efflux family protein [Arabidopsis thaliana]
Sequence ID: NP_178497.2 Length: 476
>RecName: Full=Protein DETOXIFICATION 2; Short=AtDTX2; AltName: Full=Multidrug and toxic compound extrusion protein 2; Short=MATE protein 2 [Arabidopsis thaliana]
Sequence ID: Q8GXM8.1 Length: 476 >unknown protein [Arabidopsis thaliana]
Sequence ID: AAO63931.1 Length: 476 >MATE efflux family protein [Arabidopsis thaliana]
Sequence ID: AEC05797.1 Length: 476 >unknown protein [Arabidopsis thaliana]
Sequence ID: BAC42772.1 Length: 476
Range 1: 123 to 263

Score:62.4 bits(150), Expect:5e-10,
Method:Compositional matrix adjust.,
Identities:46/141(33%), Positives:69/141(48%), Gaps:5/141(3%)

Query  317  LSIIWINVSSILVFLDQDAIIA*TFGSYIVFLIPDLIGCVI---LKCFSLRAK--TECFL  481
            +SI+WI +  +L+ L Q+  I+   GSY ++L+P L    I   L  F L
Sbjct  123  ISILWIYIEKLLITLGQEPDISRVAGSYSLWLVPALFAHAIFLPLTRFLLAQGLVISLLY  182

Query  482  SCSVPVLLHVHACWGLVLKSGLGIREASQATSLSFWVNVVLLSIYVKLSPSCAKTWTGFS  661
            S    +L H+  CW LV   GLG   A+ A SLSFW   V+LS +V+   SC KT    S
Sbjct  183  SAMTTLLFHIAVCWTLVFALGLGSNGAAIAISLSFWFYAVILSCHVRFFSSCEKTRGFVS  242

Query  662  PEALENVMVFLWSSICIHGLV  724
             + + ++  +    +   GL+
Sbjct  243  NDFMSSIKQYFQYGVPSAGLI  263


>MATE efflux family protein [Arabidopsis thaliana]
Sequence ID: NP_001324727.1 Length: 469
>hypothetical protein [Arabidopsis thaliana]
Sequence ID: AAD28684.1 Length: 469 >MATE efflux family protein [Arabidopsis thaliana]
Sequence ID: ANM62579.1 Length: 469
Range 1: 123 to 263

Score:62.4 bits(150), Expect:5e-10,
Method:Compositional matrix adjust.,
Identities:46/141(33%), Positives:69/141(48%), Gaps:5/141(3%)

Query  317  LSIIWINVSSILVFLDQDAIIA*TFGSYIVFLIPDLIGCVI---LKCFSLRAK--TECFL  481
            +SI+WI +  +L+ L Q+  I+   GSY ++L+P L    I   L  F L
Sbjct  123  ISILWIYIEKLLITLGQEPDISRVAGSYSLWLVPALFAHAIFLPLTRFLLAQGLVISLLY  182

Query  482  SCSVPVLLHVHACWGLVLKSGLGIREASQATSLSFWVNVVLLSIYVKLSPSCAKTWTGFS  661
            S    +L H+  CW LV   GLG   A+ A SLSFW   V+LS +V+   SC KT    S
Sbjct  183  SAMTTLLFHIAVCWTLVFALGLGSNGAAIAISLSFWFYAVILSCHVRFFSSCEKTRGFVS  242

Query  662  PEALENVMVFLWSSICIHGLV  724
             + + ++  +    +   GL+
Sbjct  243  NDFMSSIKQYFQYGVPSAGLI  263


>hypothetical protein [Arabidopsis thaliana]
Sequence ID: AAD28685.1 Length: 476
Range 1: 123 to 253

Score:62.0 bits(149), Expect:8e-10,
Method:Compositional matrix adjust.,
Identities:48/131(37%), Positives:63/131(48%), Gaps:5/131(3%)

Query  317  LSIIWINVSSILVFLDQDAIIA*TFGSYIVFLIPDLIGCVI---LKCFSLRAKT--ECFL  481
            +SI+WI +  +L+ L QD  I+   GSY + LIP L    I   L  F L
Sbjct  123  ISILWIYMEKLLISLGQDPDISRVAGSYALRLIPTLFAHAIVLPLTRFLLAQGLVLPLLY  182

Query  482  SCSVPVLLHVHACWGLVLKSGLGIREASQATSLSFWVNVVLLSIYVKLSPSCAKTWTGFS  661
                 +L H+  CW LV   GLG   A+ A S+SFW   + LS YV+ S SC KT    S
Sbjct  183  FALTTLLFHIAVCWTLVSALGLGSNGAALAISVSFWFFAMTLSCYVRFSSSCEKTRRFVS  242

Query  662  PEALENVMVFL  694
             + L +V  F
Sbjct  243  QDFLSSVKQFF  253


>MATE efflux family protein [Arabidopsis thaliana]
Sequence ID: NP_178496.2 Length: 476
>RecName: Full=Protein DETOXIFICATION 4; Short=AtDTX4; AltName: Full=Multidrug and toxic compound extrusion protein 4; Short=MATE protein 4 [Arabidopsis thaliana]
Sequence ID: Q9SIA3.2 Length: 476 >MATE efflux family protein [Arabidopsis thaliana]
Sequence ID: AEC05796.1 Length: 476
Range 1: 123 to 253

Score:61.6 bits(148), Expect:9e-10,
Method:Compositional matrix adjust.,
Identities:48/131(37%), Positives:63/131(48%), Gaps:5/131(3%)

Query  317  LSIIWINVSSILVFLDQDAIIA*TFGSYIVFLIPDLIGCVI---LKCFSLRAKT--ECFL  481
            +SI+WI +  +L+ L QD  I+   GSY + LIP L    I   L  F L
Sbjct  123  ISILWIYMEKLLISLGQDPDISRVAGSYALRLIPTLFAHAIVLPLTRFLLAQGLVLPLLY  182

Query  482  SCSVPVLLHVHACWGLVLKSGLGIREASQATSLSFWVNVVLLSIYVKLSPSCAKTWTGFS  661
                 +L H+  CW LV   GLG   A+ A S+SFW   + LS YV+ S SC KT    S
Sbjct  183  FALTTLLFHIAVCWTLVSALGLGSNGAALAISVSFWFFAMTLSCYVRFSSSCEKTRRFVS  242

Query  662  PEALENVMVFL  694
             + L +V  F
Sbjct  243  QDFLSSVKQFF  253


>unnamed protein product [Arabidopsis thaliana]
Sequence ID: VYS51994.1 Length: 476
Range 1: 123 to 237

Score:61.6 bits(148), Expect:1e-09,
Method:Compositional matrix adjust.,
Identities:43/115(37%), Positives:59/115(51%), Gaps:5/115(4%)

Query  317  LSIIWINVSSILVFLDQDAIIA*TFGSYIVFLIPDLIG---CVILKCFSLRAK--TECFL  481
            +S++WI +  +L+ L QD  I+   GSY ++LIP L      + L  F L
Sbjct  123  ISVLWIYIEKLLISLGQDPDISRVAGSYALWLIPALFAHAFFIPLTRFLLAQGLVLPLLY  182

Query  482  SCSVPVLLHVHACWGLVLKSGLGIREASQATSLSFWVNVVLLSIYVKLSPSCAKT  646
                 +L H+  CW  V   GLG   A+ A S+SFW  VV+LS YV+ S SC KT
Sbjct  183  CTLTTLLFHIPVCWAFVYAFGLGSNGAAMAISVSFWFYVVILSCYVRYSSSCDKT  237


>MATE efflux family protein [Arabidopsis thaliana]
Sequence ID: NP_178492.1 Length: 476
>RecName: Full=Protein DETOXIFICATION 3; Short=AtDTX3; AltName: Full=Multidrug and toxic compound extrusion protein 3; Short=MATE protein 3 [Arabidopsis thaliana]
Sequence ID: Q9SIA4.1 Length: 476 >hypothetical protein [Arabidopsis thaliana]
Sequence ID: AAD28686.1 Length: 476 >MATE efflux family protein [Arabidopsis thaliana]
Sequence ID: AEC05792.1 Length: 476 >unnamed protein product [Arabidopsis thaliana]
Sequence ID: CAA0356969.1 Length: 476
Range 1: 123 to 237

Score:61.6 bits(148), Expect:1e-09,
Method:Compositional matrix adjust.,
Identities:43/115(37%), Positives:59/115(51%), Gaps:5/115(4%)

Query  317  LSIIWINVSSILVFLDQDAIIA*TFGSYIVFLIPDLIG---CVILKCFSLRAK--TECFL  481
            +S++WI +  +L+ L QD  I+   GSY ++LIP L      + L  F L
Sbjct  123  ISVLWIYIEKLLISLGQDPDISRVAGSYALWLIPALFAHAFFIPLTRFLLAQGLVLPLLY  182

Query  482  SCSVPVLLHVHACWGLVLKSGLGIREASQATSLSFWVNVVLLSIYVKLSPSCAKT  646
                 +L H+  CW  V   GLG   A+ A S+SFW  VV+LS YV+ S SC KT
Sbjct  183  CTLTTLLFHIPVCWAFVYAFGLGSNGAAMAISVSFWFYVVILSCYVRYSSSCDKT  237


>hypothetical protein AXX17_AT2G03100 [Arabidopsis thaliana]
Sequence ID: OAP09049.1 Length: 476
Range 1: 123 to 237

Score:61.6 bits(148), Expect:1e-09,
Method:Compositional matrix adjust.,
Identities:43/115(37%), Positives:59/115(51%), Gaps:5/115(4%)

Query  317  LSIIWINVSSILVFLDQDAIIA*TFGSYIVFLIPDLIG---CVILKCFSLRAK--TECFL  481
            +S++WI +  +L+ L QD  I+   GSY ++LIP L      + L  F L
Sbjct  123  ISVLWIYIEKLLISLGQDPDISRVAGSYALWLIPALFAHAFFIPLTRFLLAQGLVLPLLY  182

Query  482  SCSVPVLLHVHACWGLVLKSGLGIREASQATSLSFWVNVVLLSIYVKLSPSCAKT  646
                 +L H+  CW  V   GLG   A+ A S+SFW  VV+LS YV+ S SC KT
Sbjct  183  CTLTTLLFHIPVCWAFVYAFGLGSNGAAMAISVSFWFYVVILSCYVRYSSSCDKT  237


>hypothetical protein AXX17_AT2G03110 [Arabidopsis thaliana]
Sequence ID: OAP07299.1 Length: 346
Range 1: 123 to 253

Score:60.8 bits(146), Expect:1e-09,
Method:Compositional matrix adjust.,
Identities:48/131(37%), Positives:63/131(48%), Gaps:5/131(3%)

Query  317  LSIIWINVSSILVFLDQDAIIA*TFGSYIVFLIPDLIGCVI---LKCFSLRAKT--ECFL  481
            +SI+WI +  +L+ L QD  I+   GSY + LIP L    I   L  F L
Sbjct  123  ISILWIYMEKLLISLGQDPDISRVSGSYALRLIPTLFAHAIVLPLTRFLLAQGLVLPLLY  182

Query  482  SCSVPVLLHVHACWGLVLKSGLGIREASQATSLSFWVNVVLLSIYVKLSPSCAKTWTGFS  661
                 +L H+  CW LV   GLG   A+ A S+SFW   + LS YV+ S SC KT    S
Sbjct  183  FALTTLLFHIVVCWTLVSALGLGSNGAALAISVSFWFFAMTLSCYVRFSSSCEKTRRFVS  242

Query  662  PEALENVMVFL  694
             + L +V  F
Sbjct  243  QDFLSSVKQFF  253


>unnamed protein product [Arabidopsis thaliana]
Sequence ID: CAA0383444.1 Length: 469
Range 1: 127 to 284

Score:61.2 bits(147), Expect:1e-09,
Method:Compositional matrix adjust.,
Identities:49/160(31%), Positives:82/160(51%), Gaps:7/160(4%)

Query  317  LSIIWINVSSILVFLDQDAIIA*TFGSYIVFLIPDLIGCVILKCFSLRAKTECFLSCSV-  493
            ++I+W    S+ + L QD  I+     Y+  L P L+    L+      +T+C ++  V
Sbjct  127  ITILWFFTESVFLLLRQDPSISKQAALYMKSLAPGLLAYGFLQNILRFCQTQCIVTPLVL  186

Query  494  ----PVLLHVHACWGLVLKSGLGIREASQATSLSFWVNVVLLSIYVKLSPSCAKTWTGFS  661
                P+++++   + LV  +GLG   A  ATS+S W+  V L  YV  S    +TWTGFS
Sbjct  187  FSFLPLVINIGTTYALVHLAGLGFIGAPIATSISLWIAFVSLGFYVICSDKFKETWTGFS  246

Query  662  PEALENVMVFLWSSICIHGLVRLVVHIYDSCLLVYIISLM  781
             E+  +V++ L  SI    +V L    ++  +LV++  LM
Sbjct  247  MESFHHVVLNLTLSIPSAAMVCLEYWAFE--ILVFLAGLM  284


>MATE efflux family protein [Arabidopsis thaliana]
Sequence ID: NP_001321858.1 Length: 338
>MATE efflux family protein [Arabidopsis thaliana]
Sequence ID: ANM59504.1 Length: 338
Range 1: 3 to 126

Score:56.2 bits(134), Expect:4e-08,
Method:Compositional matrix adjust.,
Identities:36/124(29%), Positives:61/124(49%), Gaps:5/124(4%)

Query  338  VSSILVFLDQDAIIA*TFGSYIVFLIPDLIGCVILKCFSLRAKTECF-----LSCSVPVL  502
            ++ IL+ L QD  IA   G Y ++L+P L G  +L+      +++       LS    +
Sbjct  3    MNQILLLLHQDPQIAELAGVYCLWLVPALFGYSVLESLVRYFQSQSLIYPMVLSSLAALS  62

Query  503  LHVHACWGLVLKSGLGIREASQATSLSFWVNVVLLSIYVKLSPSCAKTWTGFSPEALENV  682
             HV  CW +V K   G + A+ +  +S+W+N V L +Y+K S  C +T    S +   +
Sbjct  63   FHVPLCWLMVHKFDFGAKGAAASIGISYWLNAVFLWVYMKRSSRCVETRIYMSKDVFVHT  122

Query  683  MVFL  694
             +F
Sbjct  123  NIFF  126


Query #45: XLOC_007898 Query ID: lcl|Query_76579 Length: 1232

No significant similarity found.


Query #46: XLOC_008168 Query ID: lcl|Query_76580 Length: 236

No significant similarity found.


Query #47: XLOC_008169 Query ID: lcl|Query_76581 Length: 460

No significant similarity found.


Query #48: XLOC_008170 Query ID: lcl|Query_76582 Length: 905

No significant similarity found.


Query #49: XLOC_008171 Query ID: lcl|Query_76583 Length: 691

No significant similarity found.


Query #50: XLOC_008172 Query ID: lcl|Query_76584 Length: 563

No significant similarity found.


Query #51: XLOC_008173 Query ID: lcl|Query_76585 Length: 488

Sequences producing significant alignments:
                                                                  Max    Total Query   E   Per.
Description                                                       Score  Score cover Value Ident  Accession
F16A14.16 [Arabidopsis thaliana]                                  102    102   29%   9e-29 100.00 AAF79400.1
hypothetical protein AXX17_AT2G02320 [Arabidopsis thaliana]       54.3   54.3  36%   2e-08 46.77  OAP07693.1
hypothetical protein AXX17_AT2G02370 [Arabidopsis thaliana]       52.4   52.4  36%   8e-08 46.77  OAP11671.1


Alignments:
>F16A14.16 [Arabidopsis thaliana]
Sequence ID: AAF79400.1 Length: 48
Range 1: 1 to 48

Score:102 bits(253), Expect:9e-29,
Method:Compositional matrix adjust.,
Identities:48/48(100%), Positives:48/48(100%), Gaps:0/48(0%)

Query  237  MCVTLQKWESTRIKRAHQSLHHQSPVTALTLNDDLCWINGACNLDPIT  380
            MCVTLQKWESTRIKRAHQSLHHQSPVTALTLNDDLCWINGACNLDPIT
Sbjct  1    MCVTLQKWESTRIKRAHQSLHHQSPVTALTLNDDLCWINGACNLDPIT  48


>hypothetical protein AXX17_AT2G02320 [Arabidopsis thaliana]
Sequence ID: OAP07693.1 Length: 566
Range 1: 197 to 258

Score:54.3 bits(129), Expect:2e-08,
Method:Composition-based stats.,
Identities:29/62(47%), Positives:38/62(61%), Gaps:2/62(3%)

Query  252  QKWESTRIKRAHQSLHHQSPVTALTLND--DLCWINGACNLDPIT*TFTLIDPTVPTSAP  425
            Q+WES R K+  Q +   + +T    +   DLC ING+C L+P T T TL+DPT  TSAP
Sbjct  197  QRWESRRTKQNPQVMAASAKITCDRSHTSYDLCSINGSCILNPKTGTLTLMDPTFTTSAP  256

Query  426  SV  431
             V
Sbjct  257  LV  258


>hypothetical protein AXX17_AT2G02370 [Arabidopsis thaliana]
Sequence ID: OAP11671.1 Length: 732
Range 1: 363 to 424

Score:52.4 bits(124), Expect:8e-08,
Method:Composition-based stats.,
Identities:29/62(47%), Positives:38/62(61%), Gaps:2/62(3%)

Query  252  QKWESTRIKRAHQSLHHQSPVTALTLND--DLCWINGACNLDPIT*TFTLIDPTVPTSAP  425
            Q+WES R K+  Q +   + +T    +   DLC ING+C L+P T T TL+DPT  TSAP
Sbjct  363  QRWESRRTKQNPQVMAASAKITCDRSHTSYDLCSINGSCILNPKTGTLTLMDPTFTTSAP  422

Query  426  SV  431
             V
Sbjct  423  LV  424


Query #52: XLOC_008174 Query ID: lcl|Query_76586 Length: 1407

Sequences producing significant alignments:
                                                                  Max    Total Query    E    Per.
Description                                                       Score  Score cover Value  Ident  Accession
F9L1.30 [Arabidopsis thaliana]                                    430    430   55%   3e-150 100.00 AAD39662.1
suppressor SRP40-like protein [Arabidopsis thaliana]              56.2   56.2  20%   2e-08  41.58  NP_001154490.1
suppressor SRP40-like protein [Arabidopsis thaliana]              57.0   57.0  20%   3e-08  41.58  NP_001318192.1
hypothetical protein [Arabidopsis thaliana]                       56.2   56.2  20%   3e-08  41.58  AAD20071.1
unnamed protein product [Arabidopsis thaliana]                    56.6   56.6  20%   4e-08  41.58  CAA0356427.1
hypothetical protein AXX17_AT2G02600 [Arabidopsis thaliana]       56.6   56.6  20%   4e-08  41.58  OAP10934.1


Alignments:
>F9L1.30 [Arabidopsis thaliana]
Sequence ID: AAD39662.1 Length: 266
>hypothetical protein AXX17_AT1G16100 [Arabidopsis thaliana]
Sequence ID: OAP16696.1 Length: 266
Range 1: 7 to 266

Score:430 bits(1105), Expect:3e-150,
Method:Compositional matrix adjust.,
Identities:260/260(100%), Positives:260/260(100%), Gaps:0/260(0%)

Query  944  WDDAVERSTKESKASSEDTSQQNWELSNASSSKGSDISGVTHVRMQPssssqgseiSGLT  765
            WDDAVERSTKESKASSEDTSQQNWELSNASSSKGSDISGVTHVRMQPSSSSQGSEISGLT
Sbjct  7    WDDAVERSTKESKASSEDTSQQNWELSNASSSKGSDISGVTHVRMQPSSSSQGSEISGLT  66

Query  764  HLRMQPAPSPPSRLRPVHLMNNGIANYDPGRIPASVFSKNGDVDWSEISNESLFSLKIDG  585
            HLRMQPAPSPPSRLRPVHLMNNGIANYDPGRIPASVFSKNGDVDWSEISNESLFSLKIDG
Sbjct  67   HLRMQPAPSPPSRLRPVHLMNNGIANYDPGRIPASVFSKNGDVDWSEISNESLFSLKIDG  126

Query  584  YRKSNAQGSRHSNLKPDEVLMSGEFLRYSPSLVVKPEDAKEKTSEVEEVKNNEETRLQSH  405
            YRKSNAQGSRHSNLKPDEVLMSGEFLRYSPSLVVKPEDAKEKTSEVEEVKNNEETRLQSH
Sbjct  127  YRKSNAQGSRHSNLKPDEVLMSGEFLRYSPSLVVKPEDAKEKTSEVEEVKNNEETRLQSH  186

Query  404  NHREKYPSFPDVPLSSIPNNFYCLYAPRTKPVSFSLPVGekksmkrkkrrknkkkknkke  225
            NHREKYPSFPDVPLSSIPNNFYCLYAPRTKPVSFSLPVGEKKSMKRKKRRKNKKKKNKKE
Sbjct  187  NHREKYPSFPDVPLSSIPNNFYCLYAPRTKPVSFSLPVGEKKSMKRKKRRKNKKKKNKKE  246

Query  224  kkkkKCCTCMWTWLCSVFSK  165
            KKKKKCCTCMWTWLCSVFSK
Sbjct  247  KKKKKCCTCMWTWLCSVFSK  266


>suppressor SRP40-like protein [Arabidopsis thaliana]
Sequence ID: NP_001154490.1 Length: 198
>suppressor SRP40-like protein [Arabidopsis thaliana]
Sequence ID: AEC05728.1 Length: 198
Range 1: 93 to 188

Score:56.2 bits(134), Expect:2e-08,
Method:Compositional matrix adjust.,
Identities:42/101(42%), Positives:52/101(51%), Gaps:11/101(10%)

Query  719  PVHLM---NNGIANYDPGRIPASVFSK---NGDVDWSEISNESLFSLKIDGYRKSNAQGS  558
            PV +M   NNG   YDP RIP+SVF +   N   +WS  SNESLFS+ +     +   G
Sbjct  93   PVQVMDRDNNG--KYDPNRIPSSVFERSKSNVPAEWSCTSNESLFSIHLGNNSFTGYGGD  150

Query  557  RHSNLKPDEVLMSGEFLRYSPSLVVKPEDAKEKTSEVEEVK  435
                +K  E+  SGE L YSP L + P    E    VEE K
Sbjct  151  L---MKSGELYKSGELLAYSPGLPMPPVPGSEPKPVVEEPK  188


>suppressor SRP40-like protein [Arabidopsis thaliana]
Sequence ID: NP_001318192.1 Length: 310
>suppressor SRP40-like protein [Arabidopsis thaliana]
Sequence ID: AEC05727.2 Length: 310
Range 1: 93 to 188

Score:57.0 bits(136), Expect:3e-08,
Method:Compositional matrix adjust.,
Identities:42/101(42%), Positives:52/101(51%), Gaps:11/101(10%)

Query  719  PVHLM---NNGIANYDPGRIPASVFSK---NGDVDWSEISNESLFSLKIDGYRKSNAQGS  558
            PV +M   NNG   YDP RIP+SVF +   N   +WS  SNESLFS+ +     +   G
Sbjct  93   PVQVMDRDNNG--KYDPNRIPSSVFERSKSNVPAEWSCTSNESLFSIHLGNNSFTGYGGD  150

Query  557  RHSNLKPDEVLMSGEFLRYSPSLVVKPEDAKEKTSEVEEVK  435
                +K  E+  SGE L YSP L + P    E    VEE K
Sbjct  151  L---MKSGELYKSGELLAYSPGLPMPPVPGSEPKPVVEEPK  188


>hypothetical protein [Arabidopsis thaliana]
Sequence ID: AAD20071.1 Length: 252
Range 1: 93 to 188

Score:56.2 bits(134), Expect:3e-08,
Method:Compositional matrix adjust.,
Identities:42/101(42%), Positives:52/101(51%), Gaps:11/101(10%)

Query  719  PVHLM---NNGIANYDPGRIPASVFSK---NGDVDWSEISNESLFSLKIDGYRKSNAQGS  558
            PV +M   NNG   YDP RIP+SVF +   N   +WS  SNESLFS+ +     +   G
Sbjct  93   PVQVMDRDNNG--KYDPNRIPSSVFERSKSNVPAEWSCTSNESLFSIHLGNNSFTGYGGD  150

Query  557  RHSNLKPDEVLMSGEFLRYSPSLVVKPEDAKEKTSEVEEVK  435
                +K  E+  SGE L YSP L + P    E    VEE K
Sbjct  151  L---MKSGELYKSGELLAYSPGLPMPPVPGSEPKPVVEEPK  188


>unnamed protein product [Arabidopsis thaliana]
Sequence ID: CAA0356427.1 Length: 313
Range 1: 94 to 189

Score:56.6 bits(135), Expect:4e-08,
Method:Compositional matrix adjust.,
Identities:42/101(42%), Positives:52/101(51%), Gaps:11/101(10%)

Query  719  PVHLM---NNGIANYDPGRIPASVFSK---NGDVDWSEISNESLFSLKIDGYRKSNAQGS  558
            PV +M   NNG   YDP RIP+SVF +   N   +WS  SNESLFS+ +     +   G
Sbjct  94   PVQVMDRDNNG--KYDPNRIPSSVFERSKSNVPAEWSCTSNESLFSIHLGNNSFTGYGGD  151

Query  557  RHSNLKPDEVLMSGEFLRYSPSLVVKPEDAKEKTSEVEEVK  435
                +K  E+  SGE L YSP L + P    E    VEE K
Sbjct  152  L---MKSGELYKSGELLAYSPGLPMPPVPGSEPKPVVEEPK  189


>hypothetical protein AXX17_AT2G02600 [Arabidopsis thaliana]
Sequence ID: OAP10934.1 Length: 313
>unnamed protein product [Arabidopsis thaliana]
Sequence ID: VYS51942.1 Length: 313
Range 1: 94 to 189

Score:56.6 bits(135), Expect:4e-08,
Method:Compositional matrix adjust.,
Identities:42/101(42%), Positives:52/101(51%), Gaps:11/101(10%)

Query  719  PVHLM---NNGIANYDPGRIPASVFSK---NGDVDWSEISNESLFSLKIDGYRKSNAQGS  558
            PV +M   NNG   YDP RIP+SVF +   N   +WS  SNESLFS+ +     +   G
Sbjct  94   PVQVMDRDNNG--KYDPNRIPSSVFERSKSNVPAEWSCTSNESLFSIHLGNNSFTGYGGD  151

Query  557  RHSNLKPDEVLMSGEFLRYSPSLVVKPEDAKEKTSEVEEVK  435
                +K  E+  SGE L YSP L + P    E    VEE K
Sbjct  152  L---MKSGELYKSGELLAYSPGLPMPPVPGSEPKPVVEEPK  189


Query #53: XLOC_008175 Query ID: lcl|Query_76587 Length: 620

No significant similarity found.


Query #54: XLOC_008176 Query ID: lcl|Query_76588 Length: 946

No significant similarity found.


Query #55: XLOC_008177 Query ID: lcl|Query_76589 Length: 260

No significant similarity found.


Query #56: XLOC_008178 Query ID: lcl|Query_76590 Length: 621

No significant similarity found.


Query #57: XLOC_008179 Query ID: lcl|Query_76591 Length: 683

Sequences producing significant alignments:
                                                                  Max    Total Query   E   Per.
Description                                                       Score  Score cover Value Ident  Accession
Hypothetical protein [Arabidopsis thaliana]                       53.9   53.9  10%   2e-08 100.00 AAC00574.1


Alignments:
>Hypothetical protein [Arabidopsis thaliana]
Sequence ID: AAC00574.1 Length: 178
Range 1: 1 to 25

Score:53.9 bits(128), Expect:2e-08,
Method:Compositional matrix adjust.,
Identities:25/25(100%), Positives:25/25(100%), Gaps:0/25(0%)

Query  608  MRRNLEGIGEKAPKLGLFRALHVMP  682
            MRRNLEGIGEKAPKLGLFRALHVMP
Sbjct  1    MRRNLEGIGEKAPKLGLFRALHVMP  25


Query #58: XLOC_008180 Query ID: lcl|Query_76592 Length: 616

No significant similarity found.


Query #59: XLOC_008181 Query ID: lcl|Query_76593 Length: 682

No significant similarity found.


Query #60: XLOC_008182 Query ID: lcl|Query_76594 Length: 148

No significant similarity found.


Query #61: XLOC_008183 Query ID: lcl|Query_76595 Length: 668

No significant similarity found.


Query #62: XLOC_008184 Query ID: lcl|Query_76596 Length: 481

Sequences producing significant alignments:
                                                                  Max    Total Query   E   Per.
Description                                                       Score  Score cover Value Ident  Accession
hypothetical protein AT1G35242 [Arabidopsis thaliana]             59.3   59.3  22%   5e-12 72.22  NP_001319149.1


Alignments:
>hypothetical protein AT1G35242 [Arabidopsis thaliana]
Sequence ID: NP_001319149.1 Length: 50
>hypothetical protein AT1G35242 [Arabidopsis thaliana]
Sequence ID: AEE31772.1 Length: 50 >unnamed protein product [Arabidopsis thaliana]
Sequence ID: CAA0267017.1 Length: 50 >unnamed protein product [Arabidopsis thaliana]
Sequence ID: VYS48030.1 Length: 50
Range 1: 15 to 50

Score:59.3 bits(142), Expect:5e-12,
Method:Compositional matrix adjust.,
Identities:26/36(72%), Positives:31/36(86%), Gaps:0/36(0%)

Query  349  DVNLHVSHGYWDVLGIRGGFPVVVVIWPYKGVWIII  242
            DVNLHVS  YWDV GIRG  PVVV++WPYKG+W+I+
Sbjct  15   DVNLHVSDAYWDVFGIRGRSPVVVMVWPYKGMWLIV  50


Query #63: XLOC_008185 Query ID: lcl|Query_76597 Length: 1022

Sequences producing significant alignments:
                                                                  Max    Total Query    E    Per.
Description                                                       Score  Score cover Value  Ident  Accession
F12K21.13 [Arabidopsis thaliana]                                  382    382   58%   1e-134 100.00 AAF79258.1
unnamed protein product [Arabidopsis thaliana]                    158    158   32%   1e-47  80.18  VYS61896.1
hypothetical protein [Arabidopsis thaliana]                       121    185   29%   4e-40  83.58  CAB80805.1
transmembrane protein [Arabidopsis thaliana]                      68.9   68.9  31%   2e-12  36.52  NP_179821.1


Alignments:
>F12K21.13 [Arabidopsis thaliana]
Sequence ID: AAF79258.1 Length: 199
Range 1: 1 to 199

Score:382 bits(981), Expect:1e-134,
Method:Compositional matrix adjust.,
Identities:199/199(100%), Positives:199/199(100%), Gaps:0/199(0%)

Query  687  MSIHLNRDVFPYLKFGFGGCSHQCNLRFSTDEISSNLWFGEVIWVFDPGINRQIFLSEGT  508
            MSIHLNRDVFPYLKFGFGGCSHQCNLRFSTDEISSNLWFGEVIWVFDPGINRQIFLSEGT
Sbjct  1    MSIHLNRDVFPYLKFGFGGCSHQCNLRFSTDEISSNLWFGEVIWVFDPGINRQIFLSEGT  60

Query  507  GIDDNFLLPKTWSNDGDVLVVQRsvysyswsddddVLAVQRSIYSSSMEKEIMITNRRRW  328
            GIDDNFLLPKTWSNDGDVLVVQRSVYSYSWSDDDDVLAVQRSIYSSSMEKEIMITNRRRW
Sbjct  61   GIDDNFLLPKTWSNDGDVLVVQRSVYSYSWSDDDDVLAVQRSIYSSSMEKEIMITNRRRW  120

Query  327  IECYKRHDRDSSQAIYRCVRLIKVMQKAFGSFIYKILAFYEYMTRGLNRFHLLPVRLPFG  148
            IECYKRHDRDSSQAIYRCVRLIKVMQKAFGSFIYKILAFYEYMTRGLNRFHLLPVRLPFG
Sbjct  121  IECYKRHDRDSSQAIYRCVRLIKVMQKAFGSFIYKILAFYEYMTRGLNRFHLLPVRLPFG  180

Query  147  KQGHFHIFIEIVYNFHYFI  91
            KQGHFHIFIEIVYNFHYFI
Sbjct  181  KQGHFHIFIEIVYNFHYFI  199


>unnamed protein product [Arabidopsis thaliana]
Sequence ID: VYS61896.1 Length: 115
Range 1: 1 to 111

Score:158 bits(399), Expect:1e-47,
Method:Compositional matrix adjust.,
Identities:89/111(80%), Positives:95/111(85%), Gaps:0/111(0%)

Query  687  MSIHLNRDVFPYLKFGFGGCSHQCNLRFSTDEISSNLWFGEVIWVFDPGINRQIFLSEGT  508
            MSIHLN  VFPYLKF FGG SH+ +LRFSTDE+SSNLWFGEVIWVFDPGINRQI LSEG
Sbjct  1    MSIHLNHGVFPYLKFVFGGFSHRRDLRFSTDEVSSNLWFGEVIWVFDPGINRQILLSEGI  60

Query  507  GIDDNFLLPKTWSNDGDVLVVQRsvysyswsddddVLAVQRSIYSSSMEKE  355
            GID+NFL   TWS DGDVL VQRSVYS SWS+D DVLAV+RSIYS SMEK
Sbjct  61   GIDNNFLFSNTWSGDGDVLAVQRSVYSCSWSNDGDVLAVKRSIYSGSMEKS  111


>hypothetical protein [Arabidopsis thaliana]
Sequence ID: CAB80805.1 Length: 528
Range 1: 35 to 101

Score:121 bits(304), Expect:4e-40,
Method:Compositional matrix adjust.,
Identities:56/67(84%), Positives:60/67(89%), Gaps:0/67(0%)

Query  363  EKEIMITNRRRWIECYKRHDRDSSQAIYRCVRLIKVMQKAFGSFIYKILAFYEYMTRGLN  184
            +K IM+TNRRRW+ECYK HDRD  QA Y CVRLIKVMQKAFGSFIYK+LAFYEYMTRGLN
Sbjct  35   KKVIMVTNRRRWVECYKIHDRDPFQATYPCVRLIKVMQKAFGSFIYKMLAFYEYMTRGLN  94

Query  183  RFHLLPV  163
            RFHLL V
Sbjct  95   RFHLLLV  101


Range 2: 1 to 36

Score:63.9 bits(154), Expect:4e-40,
Method:Compositional matrix adjust.,
Identities:28/36(78%), Positives:32/36(88%), Gaps:0/36(0%)

Query  464  MVMYSLSNDPFIHIHGPMMMMYLLSNDPFIQVLWKK  357
            M MYSLSNDPFIH+HG MM+MY LSND FI++LWKK
Sbjct  1    MAMYSLSNDPFIHVHGLMMVMYSLSNDLFIRILWKK  36


>transmembrane protein [Arabidopsis thaliana]
Sequence ID: NP_179821.1 Length: 358
>hypothetical protein [Arabidopsis thaliana]
Sequence ID: AAD22363.1 Length: 358 >hypothetical protein At2g22340 [Arabidopsis thaliana]
Sequence ID: ABE65458.1 Length: 358 >transmembrane protein [Arabidopsis thaliana]
Sequence ID: AEC07295.1 Length: 358
Range 1: 241 to 355

Score:68.9 bits(167), Expect:2e-12,
Method:Compositional matrix adjust.,
Identities:42/115(37%), Positives:62/115(53%), Gaps:7/115(6%)

Query  372  SSMEKEIMITNRRRWIECY-------KRHDRDSSQAIYRCVRLIKVMQKAFGSFIYKILA  214
            ++M+ EI ITNR+R + C+       KR + +S QA  +   L KVM+KAF SFIY ++A
Sbjct  241  TAMKGEIKITNRQRCVRCFQHIILSFKRQEAESYQATIQVFLLTKVMRKAFWSFIYMVIA  300

Query  213  FYEYMTRGLNRFHLLPVRLPFGKQGHFHIFIEIVYNFHYFI*WLYLNRFMYHVLL  49
             YEY+  G NRF+LL  +    +     +     Y F  F+  +    F+Y V L
Sbjct  301  NYEYVKEGRNRFYLLACKASDRQAWSIPLLYRNCYQFLLFVSMVVYEWFIYLVKL  355


Query #64: XLOC_008186 Query ID: lcl|Query_76598 Length: 404

No significant similarity found.


Query #65: XLOC_008187 Query ID: lcl|Query_76599 Length: 854

No significant similarity found.


Query #66: XLOC_008188 Query ID: lcl|Query_76600 Length: 653

No significant similarity found.


Query #67: XLOC_008189 Query ID: lcl|Query_76601 Length: 2315

Sequences producing significant alignments:
                                                                  Max    Total Query   E   Per.
Description                                                       Score  Score cover Value Ident  Accession
hypothetical protein AXX17_AT3G09830 [Arabidopsis thaliana]       75.1   75.1  16%   3e-15 35.92  OAP06532.1
unnamed protein product [Arabidopsis thaliana]                    68.2   68.2  7%    8e-13 62.12  VYS60963.1
unknown protein [Arabidopsis thaliana]                            67.4   67.4  5%    2e-12 77.27  ABF59358.1
unnamed protein product [Arabidopsis thaliana]                    60.1   60.1  7%    2e-10 55.00  CAA0393036.1


Alignments:
>hypothetical protein AXX17_AT3G09830 [Arabidopsis thaliana]
Sequence ID: OAP06532.1 Length: 140
Range 1: 9 to 140

Score:75.1 bits(183), Expect:3e-15,
Method:Compositional matrix adjust.,
Identities:51/142(36%), Positives:73/142(51%), Gaps:24/142(16%)

Query  1810  PRFRFPKKPTSSRSDLSTRSYHRFSSSDAIFLSKSPLSEQPTLKRNSW*SRKHNEVFYLS  1631
             P+  F KKPTS RS+LS RSYH FS+ D I L+   LSEQPT + NS   RK  +  Y +
Sbjct  9     PKVSFLKKPTSPRSELSIRSYHHFSALDTISLTNPQLSEQPTSESNSKRPRKRQDTIYAT  68

Query  1630  I--------------*R*EACPSNTINLTNIGAVY*ALALFPRNI*HL*KSQTRTARTIG  1493
                             R    PS  +++  +           +++ H+ KSQ RT    G
Sbjct  69    TESFEFYRDLSHLYRDRPHPIPSCLLSVITVSM---------KHMTHV-KSQIRTVGATG  118

Query  1492  VEDRFPAIKNTSRAHLHIVLKL  1427
             +E + P I+ TSR +L+I++KL
Sbjct  119   LESKLPTIQKTSRGYLNIIMKL  140


>unnamed protein product [Arabidopsis thaliana]
Sequence ID: VYS60963.1 Length: 145
Range 1: 2 to 67

Score:68.2 bits(165), Expect:8e-13,
Method:Compositional matrix adjust.,
Identities:41/66(62%), Positives:44/66(66%), Gaps:6/66(9%)

Query  1947  LRDINVYLKQVQSVTRPNSIKPVSFRIVFLETPDEISYDILIYR-----FLQDSDFQRS-  1786
             LRDINVYLKQVQ V  P+SIKPVSF IVFLE PDE+  DI IYR     FL+     RS
Sbjct  2     LRDINVYLKQVQPVIGPSSIKPVSFGIVFLENPDELGNDIPIYRSPRVSFLKKPTSPRSE  61

Query  1785  LPRLDP  1768
             LP   P
Sbjct  62    LPSTSP  67


>unknown protein [Arabidopsis thaliana]
Sequence ID: ABF59358.1 Length: 150
Range 1: 2 to 45

Score:67.4 bits(163), Expect:2e-12,
Method:Compositional matrix adjust.,
Identities:34/44(77%), Positives:36/44(81%), Gaps:0/44(0%)

Query  1947  LRDINVYLKQVQSVTRPNSIKPVSFRIVFLETPDEISYDILIYR  1816
             LRDINVYLKQVQ V  P+SIKPVSF IVFLE PDE+  DI IYR
Sbjct  2     LRDINVYLKQVQPVIGPSSIKPVSFGIVFLENPDELGNDIPIYR  45


>unnamed protein product [Arabidopsis thaliana]
Sequence ID: CAA0393036.1 Length: 105
Range 1: 29 to 88

Score:60.1 bits(144), Expect:2e-10,
Method:Compositional matrix adjust.,
Identities:33/60(55%), Positives:39/60(65%), Gaps:0/60(0%)

Query  1816  ISPRFRFPKKPTSSRSDLSTRSYHRFSSSDAIFLSKSPLSEQPTLKRNSW*SRKHNEVFY  1637
             I+PR  F KKPTS RS+LS RSYH FS+ D I L+   LSEQPT + NS   RK  +  Y
Sbjct  29    IAPRVSFLKKPTSPRSELSIRSYHHFSALDTISLTNPQLSEQPTYESNSKRPRKKQDTIY  88


Query #68: XLOC_008190 Query ID: lcl|Query_76602 Length: 540

No significant similarity found.


Query #69: XLOC_008191 Query ID: lcl|Query_76603 Length: 1004

No significant similarity found.


Query #70: XLOC_008192 Query ID: lcl|Query_76604 Length: 848

Sequences producing significant alignments:
                                                                  Max    Total Query   E   Per.
Description                                                       Score  Score cover Value Ident  Accession
hypothetical protein [Arabidopsis thaliana]                       100    191   44%   4e-42 72.31  AAD32760.1
unknown protein [Arabidopsis thaliana]                            66.2   129   39%   1e-23 58.00  AAK92813.1
unnamed protein product [Arabidopsis thaliana]                    63.5   116   41%   1e-19 54.72  VYS53256.1
F21J9.3 [Arabidopsis thaliana]                                    63.5   116   40%   2e-19 54.72  AAF97981.1
hypothetical protein [Arabidopsis thaliana]                       61.6   113   40%   1e-18 54.72  AAD28316.1
nuclease [Arabidopsis thaliana]                                   61.2   108   37%   3e-17 54.72  NP_178997.2
unknown [Arabidopsis thaliana]                                    61.2   108   37%   3e-17 54.72  ABK28489.1
unnamed protein product [Arabidopsis thaliana]                    56.2   102   39%   2e-15 50.94  BAB02105.1
hypothetical protein At1g43590 [Arabidopsis thaliana]             68.6   68.6  22%   2e-13 50.00  ABE65399.1
unknown [Arabidopsis thaliana]                                    68.6   68.6  22%   2e-13 50.00  ABK28141.1
unknown protein; 65731-67017 [Arabidopsis thaliana]               66.6   66.6  20%   6e-12 54.39  AAG51231.1
unnamed protein product [Arabidopsis thaliana]                    63.2   63.2  20%   2e-11 52.63  BAA98206.1
ORF-c, putative; 9726-10503 [Arabidopsis thaliana]                49.3   88.2  29%   3e-11 44.68  AAG12748.1


Alignments:
>hypothetical protein [Arabidopsis thaliana]
Sequence ID: AAD32760.1 Length: 222
Range 1: 66 to 130

Score:100 bits(249), Expect:4e-42,
Method:Compositional matrix adjust.,
Identities:47/65(72%), Positives:52/65(80%), Gaps:0/65(0%)

Query  388  DHSPVFDDVEQGNIPRVNFFVNKRPYNMAYYLVDGIYHFYPTFVKSIRLP*SEPDKLLHN  209
            D SPVFDD+EQGN PRVNFFV +R YNMAYYL DGIY  YPTFVKSIRLP SEPDKL
Sbjct  66   DRSPVFDDIEQGNTPRVNFFVYQRQYNMAYYLADGIYPSYPTFVKSIRLPQSEPDKLFVQ  125

Query  208  IKKNV  194
            +++
Sbjct  126  LQEGC  130


Range 2: 128 to 185

Score:91.3 bits(225), Expect:4e-42,
Method:Compositional matrix adjust.,
Identities:46/64(72%), Positives:51/64(79%), Gaps:6/64(9%)

Query  203  EECRKDIEHAFGVLHN*FKIIRKPARLRDISDLTIIMRSCIILHNMIVENEQETFAQHWT  24
            E CRKDIE AFGVLH  FKII       +I+DL IIMRSCIILHNMIVENE++T+AQHWT
Sbjct  128  EGCRKDIERAFGVLHARFKIIW------NIADLAIIMRSCIILHNMIVENERDTYAQHWT  181

Query  23   YYDQ  12
             YDQ
Sbjct  182  DYDQ  185


>unknown protein [Arabidopsis thaliana]
Sequence ID: AAK92813.1 Length: 415
>unknown protein [Arabidopsis thaliana]
Sequence ID: AAM45005.1 Length: 415 >unnamed protein product [Arabidopsis thaliana]
Sequence ID: BAB02160.1 Length: 415
Range 1: 249 to 298

Score:66.2 bits(160), Expect:1e-23,
Method:Compositional matrix adjust.,
Identities:29/50(58%), Positives:37/50(74%), Gaps:0/50(0%)

Query  388  DHSPVFDDVEQGNIPRVNFFVNKRPYNMAYYLVDGIYHFYPTFVKSIRLP  239
            D SP+FDD+ QG  P V + VN R Y++AYYL DGIY  + TF++SIRLP
Sbjct  249  DRSPIFDDILQGRAPNVKYKVNGREYHLAYYLTDGIYPKWATFIQSIRLP  298


Range 2: 305 to 365

Score:63.5 bits(153), Expect:1e-23,
Method:Compositional matrix adjust.,
Identities:31/61(51%), Positives:42/61(68%), Gaps:0/61(0%)

Query  221  VIAQH*EECRKDIEHAFGVLHN*FKIIRKPARLRDISDLTIIMRSCIILHNMIVENEQET  42
            + A H E  RKD+E AFGVL   F II+ PA + D   +  IM++CIILHNMIVE+E++
Sbjct  305  LFATHQEADRKDVERAFGVLQARFHIIKNPALVWDKEKIGNIMKACIILHNMIVEDERDG  364

Query  41   F  39
            +
Sbjct  365  Y  365


>unnamed protein product [Arabidopsis thaliana]
Sequence ID: VYS53256.1 Length: 443
Range 1: 329 to 381

Score:63.5 bits(153), Expect:1e-19,
Method:Compositional matrix adjust.,
Identities:29/53(55%), Positives:40/53(75%), Gaps:0/53(0%)

Query  203  EECRKDIEHAFGVLHN*FKIIRKPARLRDISDLTIIMRSCIILHNMIVENEQE  45
            E CRKD+E AFGVL + F I+  P+RL + + L  IM SCII+HNMI+E+E++
Sbjct  329  EACRKDVERAFGVLQSRFAIVAGPSRLWNKTVLHDIMTSCIIMHNMIIEDERD  381


Range 2: 266 to 332

Score:52.8 bits(125), Expect:1e-19,
Method:Compositional matrix adjust.,
Identities:27/70(39%), Positives:41/70(58%), Gaps:3/70(4%)

Query  397  LDTDHSPVFDDVEQGNIPRVNFFVNKRPYNMAYYLVDGIYHFYPTFVKSIRLP*SEPDKL  218
            L+T H  +F ++ +G  P  ++ +N +PYNM YYL DGIY  + T V++I  P   P K
Sbjct  266  LETSH--LFANLAEGTAPPASYVINGKPYNMGYYLADGIYPKWSTLVQTIHDP-RGPKKK  322

Query  217  LHNIKKNVGR  188
            L  +K+   R
Sbjct  323  LFAMKQEACR  332


>F21J9.3 [Arabidopsis thaliana]
Sequence ID: AAF97981.1 Length: 457
Range 1: 238 to 290

Score:63.5 bits(153), Expect:2e-19,
Method:Compositional matrix adjust.,
Identities:29/53(55%), Positives:40/53(75%), Gaps:0/53(0%)

Query  203  EECRKDIEHAFGVLHN*FKIIRKPARLRDISDLTIIMRSCIILHNMIVENEQE  45
            E CRKD+E AFGVL + F I+  P+RL + + L  IM SCII+HNMI+E+E++
Sbjct  238  EACRKDVERAFGVLQSRFAIVAGPSRLWNKTVLHDIMTSCIIMHNMIIEDERD  290


Range 2: 176 to 241

Score:52.8 bits(125), Expect:2e-19,
Method:Compositional matrix adjust.,
Identities:25/67(37%), Positives:41/67(61%), Gaps:1/67(1%)

Query  388  DHSPVFDDVEQGNIPRVNFFVNKRPYNMAYYLVDGIYHFYPTFVKSIRLP*SEPDKLLHN  209
            + S +F ++ +G  P  ++ +N++PYNM+YYL DGIY  + T V++I  P   P K L
Sbjct  176  EASHLFANLAEGTAPPASYVINEKPYNMSYYLADGIYPKWSTLVQTIHDP-RGPKKKLFA  234

Query  208  IKKNVGR  188
            +K+   R
Sbjct  235  MKQEACR  241


>hypothetical protein [Arabidopsis thaliana]
Sequence ID: AAD28316.1 Length: 244
Range 1: 159 to 211

Score:61.6 bits(148), Expect:1e-18,
Method:Compositional matrix adjust.,
Identities:29/53(55%), Positives:39/53(73%), Gaps:0/53(0%)

Query  203  EECRKDIEHAFGVLHN*FKIIRKPARLRDISDLTIIMRSCIILHNMIVENEQE  45
            E CRKD+E AFGVL   F I+  P+RL + + L  IM SCII+HNMI+E+E++
Sbjct  159  EACRKDVERAFGVLQLRFAIVAGPSRLWNKTVLHDIMTSCIIMHNMIIEDERD  211


Range 2: 97 to 162

Score:51.6 bits(122), Expect:1e-18,
Method:Compositional matrix adjust.,
Identities:25/67(37%), Positives:39/67(58%), Gaps:1/67(1%)

Query  388  DHSPVFDDVEQGNIPRVNFFVNKRPYNMAYYLVDGIYHFYPTFVKSIRLP*SEPDKLLHN  209
            + S +F ++ +G  P  ++ +N +PYNM YYL DGIY  + T V++I  P   P K L
Sbjct  97   EASHLFANLAEGTAPPASYVINGKPYNMGYYLADGIYPKWSTLVQTIHDP-RGPKKKLFA  155

Query  208  IKKNVGR  188
            +K+   R
Sbjct  156  MKQEACR  162


>nuclease [Arabidopsis thaliana]
Sequence ID: NP_178997.2 Length: 343
>hypothetical protein At2g13770 [Arabidopsis thaliana]
Sequence ID: ABE65808.1 Length: 343 >nuclease [Arabidopsis thaliana]
Sequence ID: AEC06258.1 Length: 343
Range 1: 258 to 310

Score:61.2 bits(147), Expect:3e-17,
Method:Compositional matrix adjust.,
Identities:29/53(55%), Positives:39/53(73%), Gaps:0/53(0%)

Query  203  EECRKDIEHAFGVLHN*FKIIRKPARLRDISDLTIIMRSCIILHNMIVENEQE  45
            E CRKD+E AFGVL   F I+  P+RL + + L  IM SCII+HNMI+E+E++
Sbjct  258  EACRKDVERAFGVLQLRFAIVAGPSRLWNKTVLHDIMTSCIIMHNMIIEDERD  310


Range 2: 206 to 261

Score:47.4 bits(111), Expect:3e-17,
Method:Compositional matrix adjust.,
Identities:23/57(40%), Positives:33/57(57%), Gaps:1/57(1%)

Query  358  QGNIPRVNFFVNKRPYNMAYYLVDGIYHFYPTFVKSIRLP*SEPDKLLHNIKKNVGR  188
            +G  P  ++ +N +PYNM YYL DGIY  + T V++I  P   P K L  +K+   R
Sbjct  206  EGTAPPASYVINGKPYNMGYYLADGIYPKWSTLVQTIHDP-RGPKKKLFAMKQEACR  261


>unknown, partial [Arabidopsis thaliana]
Sequence ID: ABK28489.1 Length: 344
Range 1: 258 to 310

Score:61.2 bits(147), Expect:3e-17,
Method:Compositional matrix adjust.,
Identities:29/53(55%), Positives:39/53(73%), Gaps:0/53(0%)

Query  203  EECRKDIEHAFGVLHN*FKIIRKPARLRDISDLTIIMRSCIILHNMIVENEQE  45
            E CRKD+E AFGVL   F I+  P+RL + + L  IM SCII+HNMI+E+E++
Sbjct  258  EACRKDVERAFGVLQLRFAIVAGPSRLWNKTVLHDIMTSCIIMHNMIIEDERD  310


Range 2: 206 to 261

Score:47.4 bits(111), Expect:3e-17,
Method:Compositional matrix adjust.,
Identities:23/57(40%), Positives:33/57(57%), Gaps:1/57(1%)

Query  358  QGNIPRVNFFVNKRPYNMAYYLVDGIYHFYPTFVKSIRLP*SEPDKLLHNIKKNVGR  188
            +G  P  ++ +N +PYNM YYL DGIY  + T V++I  P   P K L  +K+   R
Sbjct  206  EGTAPPASYVINGKPYNMGYYLADGIYPKWSTLVQTIHDP-RGPKKKLFAMKQEACR  261


>unnamed protein product [Arabidopsis thaliana]
Sequence ID: BAB02105.1 Length: 194
Range 1: 124 to 170

Score:56.2 bits(134), Expect:2e-15,
Method:Compositional matrix adjust.,
Identities:27/53(51%), Positives:38/53(71%), Gaps:6/53(11%)

Query  203  EECRKDIEHAFGVLHN*FKIIRKPARLRDISDLTIIMRSCIILHNMIVENEQE  45
            E CRKD+E AF VLH+ F I+ +P+RL +       M SCII+HNMI+E+E++
Sbjct  124  ETCRKDVEPAFEVLHSRFAIVAEPSRLWN------KMTSCIIMHNMIIEDERD  170


Range 2: 66 to 127

Score:46.2 bits(108), Expect:2e-15,
Method:Compositional matrix adjust.,
Identities:23/63(37%), Positives:34/63(53%), Gaps:1/63(1%)

Query  376  VFDDVEQGNIPRVNFFVNKRPYNMAYYLVDGIYHFYPTFVKSIRLP*SEPDKLLHNIKKN  197
            +F ++ +   P  ++  N +PYNM YYL DGIY  + T V++I  P   P K L  +K
Sbjct  66   LFANLAEDTAPPASYVSNGKPYNMGYYLADGIYSKWSTLVQTIHDP-RGPKKKLFAMKLE  124

Query  196  VGR  188
              R
Sbjct  125  TCR  127


>hypothetical protein At1g43590 [Arabidopsis thaliana]
Sequence ID: ABE65399.1 Length: 168
Range 1: 41 to 104

Score:68.6 bits(166), Expect:2e-13,
Method:Compositional matrix adjust.,
Identities:32/64(50%), Positives:45/64(70%), Gaps:0/64(0%)

Query  221  VIAQH*EECRKDIEHAFGVLHN*FKIIRKPARLRDISDLTIIMRSCIILHNMIVENEQET  42
            + A + E CRKD+E AFGVL   F I++ PA + D   +  IMR+CIILHNMIVE+E++
Sbjct  41   LFATNQEACRKDVERAFGVLQARFAIVKHPALIWDKIKIGNIMRACIILHNMIVEDERDG  100

Query  41   FAQH  30
            + Q+
Sbjct  101  YTQY  104


>unknown, partial [Arabidopsis thaliana]
Sequence ID: ABK28141.1 Length: 169
Range 1: 41 to 104

Score:68.6 bits(166), Expect:2e-13,
Method:Compositional matrix adjust.,
Identities:32/64(50%), Positives:45/64(70%), Gaps:0/64(0%)

Query  221  VIAQH*EECRKDIEHAFGVLHN*FKIIRKPARLRDISDLTIIMRSCIILHNMIVENEQET  42
            + A + E CRKD+E AFGVL   F I++ PA + D   +  IMR+CIILHNMIVE+E++
Sbjct  41   LFATNQEACRKDVERAFGVLQARFAIVKHPALIWDKIKIGNIMRACIILHNMIVEDERDG  100

Query  41   FAQH  30
            + Q+
Sbjct  101  YTQY  104


>unknown protein; 65731-67017 [Arabidopsis thaliana]
Sequence ID: AAG51231.1 Length: 333
Range 1: 212 to 268

Score:66.6 bits(161), Expect:6e-12,
Method:Compositional matrix adjust.,
Identities:31/57(54%), Positives:40/57(70%), Gaps:0/57(0%)

Query  203  EECRKDIEHAFGVLHN*FKIIRKPARLRDISDLTIIMRSCIILHNMIVENEQETFAQ  33
            E CRKD+E AFGVL   F II+ PA   D   +  IMR+CIILHNMIVE++++ + Q
Sbjct  212  EACRKDVERAFGVLQARFAIIKHPALFHDKVKIGNIMRACIILHNMIVEDKRDGYTQ  268


>unnamed protein product [Arabidopsis thaliana]
Sequence ID: BAA98206.1 Length: 183
Range 1: 60 to 116

Score:63.2 bits(152), Expect:2e-11,
Method:Compositional matrix adjust.,
Identities:30/57(53%), Positives:39/57(68%), Gaps:0/57(0%)

Query  203  EECRKDIEHAFGVLHN*FKIIRKPARLRDISDLTIIMRSCIILHNMIVENEQETFAQ  33
            E CRKD+E AFGVL   F I++ PA   D   +  IMR+ IILHNMIVE+E++ + Q
Sbjct  60   EACRKDVERAFGVLQARFAIVKHPALFHDKVKIRNIMRAYIILHNMIVEDERDGYTQ  116


>ORF-c, putative; 9726-10503 [Arabidopsis thaliana]
Sequence ID: AAG12748.1 Length: 161
Range 1: 66 to 112

Score:49.3 bits(116), Expect:3e-11,
Method:Compositional matrix adjust.,
Identities:21/47(45%), Positives:28/47(59%), Gaps:0/47(0%)

Query  388  DHSPVFDDVEQGNIPRVNFFVNKRPYNMAYYLVDGIYHFYPTFVKSI  248
            D SP+ DD+ QG  P VN+ VN   Y++ YYL D IY  +  F+  I
Sbjct  66   DRSPICDDILQGQAPTVNYLVNGNKYHLGYYLTDKIYLKWANFINPI  112


Range 2: 125 to 161

Score:38.9 bits(89), Expect:3e-11,
Method:Compositional matrix adjust.,
Identities:18/37(49%), Positives:23/37(62%), Gaps:0/37(0%)

Query  203  EECRKDIEHAFGVLHN*FKIIRKPARLRDISDLTIIM  93
            E CRKD+E AFGVL   F I++ PA + D   +  IM
Sbjct  125  EACRKDVERAFGVLQARFAIMKNPAPIWDKVKIKNIM  161


Query #71: XLOC_008193 Query ID: lcl|Query_76605 Length: 371

Sequences producing significant alignments:
                                                                  Max    Total Query   E   Per.
Description                                                       Score  Score cover Value Ident  Accession
F9C16.2 [Arabidopsis thaliana]                                    144    144   77%   2e-44 84.38  AAF79691.1


Alignments:
>F9C16.2 [Arabidopsis thaliana]
Sequence ID: AAF79691.1 Length: 192
Range 1: 112 to 192

Score:144 bits(364), Expect:2e-44,
Method:Compositional matrix adjust.,
Identities:81/96(84%), Positives:81/96(84%), Gaps:15/96(15%)

Query  320  RDKFVLLLLSGCYVLKTMVIEQCPGDSAGHFIVRKASLKMFNCVKN*ETDAKTTGGYWEI  141
            RDKFVLLLLSGCYVLKTMVIEQCPGDSAGHFI               ETDAKTTGGYWEI
Sbjct  112  RDKFVLLLLSGCYVLKTMVIEQCPGDSAGHFI---------------ETDAKTTGGYWEI  156

Query  140  GyssqyssqYVCYKFNYDFVNILMYPESRKDLNSCN  33
            GYSSQYSSQYVCYKFNYDFVNILMYPESRKDLNSCN
Sbjct  157  GYSSQYSSQYVCYKFNYDFVNILMYPESRKDLNSCN  192


Query #72: XLOC_008194 Query ID: lcl|Query_76606 Length: 545

No significant similarity found.


Query #73: XLOC_008195 Query ID: lcl|Query_76607 Length: 929

No significant similarity found.


Query #74: XLOC_008196 Query ID: lcl|Query_76608 Length: 361

No significant similarity found.


Query #75: XLOC_008197 Query ID: lcl|Query_76609 Length: 802

No significant similarity found.


Query #76: XLOC_008198 Query ID: lcl|Query_76610 Length: 569

No significant similarity found.


Query #77: XLOC_008199 Query ID: lcl|Query_76611 Length: 581

No significant similarity found.


Query #78: XLOC_008200 Query ID: lcl|Query_76612 Length: 636

No significant similarity found.


Query #79: XLOC_008201 Query ID: lcl|Query_76613 Length: 613

No significant similarity found.


Query #80: XLOC_008202 Query ID: lcl|Query_76614 Length: 487

No significant similarity found.


Query #81: XLOC_008203 Query ID: lcl|Query_76615 Length: 487

No significant similarity found.


Query #82: XLOC_008204 Query ID: lcl|Query_76616 Length: 825

No significant similarity found.


Query #83: XLOC_008205 Query ID: lcl|Query_76617 Length: 560

No significant similarity found.


Query #84: XLOC_008206 Query ID: lcl|Query_76618 Length: 748

No significant similarity found.


Query #85: XLOC_008207 Query ID: lcl|Query_76619 Length: 332

No significant similarity found.


Query #86: XLOC_008213 Query ID: lcl|Query_76620 Length: 5603

Sequences producing significant alignments:
                                                                  Max    Total Query   E   Per.
Description                                                       Score  Score cover Value Ident  Accession
unnamed protein product [Arabidopsis thaliana]                    144    375   9%    2e-38 97.22  CAA0393038.1
unnamed protein product [Arabidopsis thaliana]                    146    377   9%    2e-39 98.61  VYS61323.1
hypothetical protein AT2G01035 [Arabidopsis thaliana]             149    380   9%    2e-40 100.00 NP_001323474.1
hypothetical protein AXX17_AT3G09810 [Arabidopsis thaliana]       161    219   7%    3e-45 86.41  OAP02487.1
hypothetical protein AT5G28345 [Arabidopsis thaliana]             160    160   4%    5e-44 98.89  NP_001331379.1
unnamed protein product [Arabidopsis thaliana]                    151    151   5%    3e-42 93.68  VYS68170.1
hypothetical protein AXX17_AT4G23970 [Arabidopsis thaliana]       156    156   4%    1e-40 97.73  OAO98525.1
unnamed protein product [Arabidopsis thaliana]                    144    144   4%    7e-40 97.65  VYS61324.1
unnamed protein product [Arabidopsis thaliana]                    144    144   4%    1e-39 98.81  CAA0405308.1
unnamed protein product [Arabidopsis thaliana]                    95.9   196   4%    2e-24 87.69  CAA0384583.1
unnamed protein product [Arabidopsis thaliana]                    93.6   228   5%    2e-21 74.32  CAA0381910.1
unnamed protein product [Arabidopsis thaliana]                    81.6   81.6  2%    4e-17 90.70  VYS60963.1
unknown protein [Arabidopsis thaliana]                            81.6   81.6  2%    4e-17 90.70  ABF59358.1
unnamed protein product [Arabidopsis thaliana]                    63.5   63.5  2%    4e-11 68.75  CAA0395933.1
hypothetical protein AT5G10336 [Arabidopsis thaliana]             59.3   59.3  1%    2e-10 100.00 NP_001119203.1
unnamed protein product [Arabidopsis thaliana]                    61.2   61.2  2%    2e-10 66.67  CAA0393036.1
hypothetical protein AXX17_ATUG01520 [Arabidopsis thaliana]       57.0   110   4%    3e-09 61.90  OAO89355.1
hypothetical protein [Arabidopsis thaliana]                       60.5   60.5  2%    4e-08 55.77  AAC69120.1


Alignments:
>unnamed protein product [Arabidopsis thaliana]
Sequence ID: CAA0393038.1 Length: 181
Range 1: 43 to 108

Score:138 bits(347), Expect:1e-52,
Method:Compositional matrix adjust.,
Identities:66/66(100%), Positives:66/66(100%), Gaps:0/66(0%)

Query  1165  CNQRNADPHCTINQIRKQISSNPKEFSWRSAYQFEAPRLSFLERPTYTLTSNSTKSYHSI  986
             CNQRNADPHCTINQIRKQISSNPKEFSWRSAYQFEAPRLSFLERPTYTLTSNSTKSYHSI
Sbjct  43    CNQRNADPHCTINQIRKQISSNPKEFSWRSAYQFEAPRLSFLERPTYTLTSNSTKSYHSI  102

Query  985   SVSAAR  968
             SVSAAR
Sbjct  103   SVSAAR  108


Range 2: 1 to 42

Score:92.4 bits(228), Expect:1e-52,
Method:Compositional matrix adjust.,
Identities:42/42(100%), Positives:42/42(100%), Gaps:0/42(0%)

Query  1394  MCRKLHALLSSRYSKTKHWSKVTKPVDFEINFQEDSDELRGV  1269
             MCRKLHALLSSRYSKTKHWSKVTKPVDFEINFQEDSDELRGV
Sbjct  1     MCRKLHALLSSRYSKTKHWSKVTKPVDFEINFQEDSDELRGV  42


Range 3: 110 to 181

Score:144 bits(363), Expect:2e-38,
Method:Compositional matrix adjust.,
Identities:70/72(97%), Positives:70/72(97%), Gaps:0/72(0%)

Query  470  QDKPSSNLYPRSLVRSSDSLPCSLLARSFGNLSHCSAHQSGHFSARHSDHLGEHPTAPQQ  291
            QDKPSSNLYPRSLVRSSDSLPCSLLARSF NLSHCSAHQSGHFSARHSDHLGEHPTAPQQ
Sbjct  110  QDKPSSNLYPRSLVRSSDSLPCSLLARSFSNLSHCSAHQSGHFSARHSDHLGEHPTAPQQ  169

Query  290  LMRSIRLPNPQA  255
            LMRSIRLPN QA
Sbjct  170  LMRSIRLPNSQA  181


>unnamed protein product [Arabidopsis thaliana]
Sequence ID: VYS61323.1 Length: 181
Range 1: 43 to 108

Score:138 bits(347), Expect:1e-52,
Method:Compositional matrix adjust.,
Identities:66/66(100%), Positives:66/66(100%), Gaps:0/66(0%)

Query  1165  CNQRNADPHCTINQIRKQISSNPKEFSWRSAYQFEAPRLSFLERPTYTLTSNSTKSYHSI  986
             CNQRNADPHCTINQIRKQISSNPKEFSWRSAYQFEAPRLSFLERPTYTLTSNSTKSYHSI
Sbjct  43    CNQRNADPHCTINQIRKQISSNPKEFSWRSAYQFEAPRLSFLERPTYTLTSNSTKSYHSI  102

Query  985   SVSAAR  968
             SVSAAR
Sbjct  103   SVSAAR  108


Range 2: 1 to 42

Score:92.0 bits(227), Expect:1e-52,
Method:Compositional matrix adjust.,
Identities:42/42(100%), Positives:42/42(100%), Gaps:0/42(0%)

Query  1394  MCRKLHALLSSRYSKTKHWSKVTKPVDFEINFQEDSDELRGV  1269
             MCRKLHALLSSRYSKTKHWSKVTKPVDFEINFQEDSDELRGV
Sbjct  1     MCRKLHALLSSRYSKTKHWSKVTKPVDFEINFQEDSDELRGV  42


Range 3: 110 to 181

Score:146 bits(369), Expect:2e-39,
Method:Compositional matrix adjust.,
Identities:71/72(99%), Positives:71/72(98%), Gaps:0/72(0%)

Query  470  QDKPSSNLYPRSLVRSSDSLPCSLLARSFGNLSHCSAHQSGHFSARHSDHLGEHPTAPQQ  291
            QDKPSSNLYPRSLVRSSDSLPCSLLARSFGNLSHCSAHQSGHFSARHSDHLGEHPTAPQQ
Sbjct  110  QDKPSSNLYPRSLVRSSDSLPCSLLARSFGNLSHCSAHQSGHFSARHSDHLGEHPTAPQQ  169

Query  290  LMRSIRLPNPQA  255
            LMRSIRLPN QA
Sbjct  170  LMRSIRLPNSQA  181


>hypothetical protein AT2G01035 [Arabidopsis thaliana]
Sequence ID: NP_001323474.1 Length: 181
>hypothetical protein AT2G01045 [Arabidopsis thaliana]
Sequence ID: NP_001324766.1 Length: 181 >hypothetical protein AT2G01035 [Arabidopsis thaliana]
Sequence ID: ANM61244.1 Length: 181 >hypothetical protein AT2G01045 [Arabidopsis thaliana]
Sequence ID: ANM62620.1 Length: 181
Range 1: 43 to 108

Score:138 bits(347), Expect:2e-52,
Method:Compositional matrix adjust.,
Identities:66/66(100%), Positives:66/66(100%), Gaps:0/66(0%)

Query  1165  CNQRNADPHCTINQIRKQISSNPKEFSWRSAYQFEAPRLSFLERPTYTLTSNSTKSYHSI  986
             CNQRNADPHCTINQIRKQISSNPKEFSWRSAYQFEAPRLSFLERPTYTLTSNSTKSYHSI
Sbjct  43    CNQRNADPHCTINQIRKQISSNPKEFSWRSAYQFEAPRLSFLERPTYTLTSNSTKSYHSI  102

Query  985   SVSAAR  968
             SVSAAR
Sbjct  103   SVSAAR  108


Range 2: 1 to 42

Score:92.0 bits(227), Expect:2e-52,
Method:Compositional matrix adjust.,
Identities:42/42(100%), Positives:42/42(100%), Gaps:0/42(0%)

Query  1394  MCRKLHALLSSRYSKTKHWSKVTKPVDFEINFQEDSDELRGV  1269
             MCRKLHALLSSRYSKTKHWSKVTKPVDFEINFQEDSDELRGV
Sbjct  1     MCRKLHALLSSRYSKTKHWSKVTKPVDFEINFQEDSDELRGV  42


Range 3: 110 to 181

Score:149 bits(377), Expect:2e-40,
Method:Compositional matrix adjust.,
Identities:72/72(100%), Positives:72/72(100%), Gaps:0/72(0%)

Query  470  QDKPSSNLYPRSLVRSSDSLPCSLLARSFGNLSHCSAHQSGHFSARHSDHLGEHPTAPQQ  291
            QDKPSSNLYPRSLVRSSDSLPCSLLARSFGNLSHCSAHQSGHFSARHSDHLGEHPTAPQQ
Sbjct  110  QDKPSSNLYPRSLVRSSDSLPCSLLARSFGNLSHCSAHQSGHFSARHSDHLGEHPTAPQQ  169

Query  290  LMRSIRLPNPQA  255
            LMRSIRLPNPQA
Sbjct  170  LMRSIRLPNPQA  181


>hypothetical protein AXX17_AT3G09810 [Arabidopsis thaliana]
Sequence ID: OAP02487.1 Length: 130
>hypothetical protein AXX17_AT3G09890 [Arabidopsis thaliana]
Sequence ID: OAP03697.1 Length: 130
Range 1: 30 to 130

Score:161 bits(408), Expect:3e-45,
Method:Compositional matrix adjust.,
Identities:89/103(86%), Positives:92/103(89%), Gaps:2/103(1%)

Query  1980  RRAIAS*EGCRRAVSWANYEPEKLQTNCELGKVQTRC*LDELRARKATDEQGARKAADEL  2159
             RRAIAS EGCRRAVSWANYEPEKLQTNCELGK + +  LDELRARKATDEQGARK AD+L
Sbjct  30    RRAIASSEGCRRAVSWANYEPEKLQTNCELGKCRLK--LDELRARKATDEQGARKVADDL  87

Query  2160  LARRSADGLLARKPTDELGARKAADELLARRSADELLARRTAS  2288
             LA RSAD LLARK TDELG RKAADELLA RSADELLARRTAS
Sbjct  88    LAWRSADELLARKATDELGVRKAADELLAMRSADELLARRTAS  130


Range 2: 1 to 36

Score:57.8 bits(138), Expect:7e-09,
Method:Compositional matrix adjust.,
Identities:29/37(78%), Positives:31/37(83%), Gaps:1/37(2%)

Query  937   MKSRSYQIVITLQQRQKYCDNSLSNSTLRCRLVFLES  1047
             MKSRSY +VITLQQRQKYCDNSLSNSTL+CR     S
Sbjct  1     MKSRSY-LVITLQQRQKYCDNSLSNSTLKCRRAIASS  36


>hypothetical protein AT5G28345 [Arabidopsis thaliana]
Sequence ID: NP_001331379.1 Length: 194
>hypothetical protein AT5G28345 [Arabidopsis thaliana]
Sequence ID: ANM69720.1 Length: 194
Range 1: 105 to 194

Score:160 bits(405), Expect:5e-44,
Method:Compositional matrix adjust.,
Identities:89/90(99%), Positives:89/90(98%), Gaps:0/90(0%)

Query  524  LIATYVFFsssarlsrrSQDKPSSNLYPRSLVRSSDSLPCSLLARSFGNLSHCSAHQSGH  345
            LIATYVFFSSSARLSRRSQDKPSSNLYPRSLVRSSDSLPCSLLARSFGNLSHCSAHQSGH
Sbjct  105  LIATYVFFSSSARLSRRSQDKPSSNLYPRSLVRSSDSLPCSLLARSFGNLSHCSAHQSGH  164

Query  344  FSARHSDHLGEHPTAPQQLMRSIRLPNPQA  255
            FSARHSDHLGEHPTAPQQLMRSIRLPN QA
Sbjct  165  FSARHSDHLGEHPTAPQQLMRSIRLPNSQA  194


>unnamed protein product [Arabidopsis thaliana]
Sequence ID: VYS68170.1 Length: 94
Range 1: 1 to 94

Score:151 bits(382), Expect:3e-42,
Method:Compositional matrix adjust.,
Identities:89/95(94%), Positives:90/95(94%), Gaps:1/95(1%)

Query  539  MACLQLIATYVFFsssarlsrrSQDKPSSNLYPRSLVRSSDSLPCSLLARSFGNLSHCSA  360
            MACLQLIA  VFFSSSARLSRRSQDKP SNLYPRSLVRSSDSLPCSLLARSFGNLSHCSA
Sbjct  1    MACLQLIAN-VFFSSSARLSRRSQDKPFSNLYPRSLVRSSDSLPCSLLARSFGNLSHCSA  59

Query  359  HQSGHFSARHSDHLGEHPTAPQQLMRSIRLPNPQA  255
            HQSGHFSAR+SDHLGEHP APQQLMRSIRLPN QA
Sbjct  60   HQSGHFSARYSDHLGEHPIAPQQLMRSIRLPNSQA  94


>hypothetical protein AXX17_AT4G23970 [Arabidopsis thaliana]
Sequence ID: OAO98525.1 Length: 355
Range 1: 1 to 88

Score:156 bits(394), Expect:1e-40,
Method:Compositional matrix adjust.,
Identities:86/88(98%), Positives:87/88(98%), Gaps:0/88(0%)

Query  545  MIMACLQLIATYVFFsssarlsrrSQDKPSSNLYPRSLVRSSDSLPCSLLARSFGNLSHC  366
            MIMACLQLIATYVFFSSSARLSRRSQD+PSSNLYPRSLVRSSDSLPCSLLARSFGNLSHC
Sbjct  1    MIMACLQLIATYVFFSSSARLSRRSQDEPSSNLYPRSLVRSSDSLPCSLLARSFGNLSHC  60

Query  365  SAHQSGHFSARHSDHLGEHPTAPQQLMR  282
            SAHQSGHFS RHSDHLGEHPTAPQQLMR
Sbjct  61   SAHQSGHFSVRHSDHLGEHPTAPQQLMR  88


>unnamed protein product [Arabidopsis thaliana]
Sequence ID: VYS61324.1 Length: 86
Range 1: 2 to 86

Score:144 bits(363), Expect:7e-40,
Method:Compositional matrix adjust.,
Identities:83/85(98%), Positives:83/85(97%), Gaps:0/85(0%)

Query  509  VFFsssarlsrrSQDKPSSNLYPRSLVRSSDSLPCSLLARSFGNLSHCSAHQSGHFSARH  330
             FFSSSARLSRRSQDKPSSNLYPRSLVRSSDSLPCSLLARSFGNLSHCSAHQSGHFSARH
Sbjct  2    FFFSSSARLSRRSQDKPSSNLYPRSLVRSSDSLPCSLLARSFGNLSHCSAHQSGHFSARH  61

Query  329  SDHLGEHPTAPQQLMRSIRLPNPQA  255
            SDHLGEHPTAPQQLMRSIRLPN QA
Sbjct  62   SDHLGEHPTAPQQLMRSIRLPNSQA  86


>unnamed protein product [Arabidopsis thaliana]
Sequence ID: CAA0405308.1 Length: 93
Range 1: 10 to 93

Score:144 bits(363), Expect:1e-39,
Method:Compositional matrix adjust.,
Identities:83/84(99%), Positives:83/84(98%), Gaps:0/84(0%)

Query  506  FFsssarlsrrSQDKPSSNLYPRSLVRSSDSLPCSLLARSFGNLSHCSAHQSGHFSARHS  327
            FFSSSARLSRRSQDKPSSNLYPRSLVRSSDSLPCSLLARSFGNLSHCSAHQSGHFSARHS
Sbjct  10   FFSSSARLSRRSQDKPSSNLYPRSLVRSSDSLPCSLLARSFGNLSHCSAHQSGHFSARHS  69

Query  326  DHLGEHPTAPQQLMRSIRLPNPQA  255
            DHLGEHPTAPQQLMRSIRLPN QA
Sbjct  70   DHLGEHPTAPQQLMRSIRLPNSQA  93


>unnamed protein product [Arabidopsis thaliana]
Sequence ID: CAA0384583.1 Length: 106
Range 1: 43 to 106

Score:95.9 bits(237), Expect:2e-24,
Method:Compositional matrix adjust.,
Identities:57/65(88%), Positives:59/65(90%), Gaps:1/65(1%)

Query  2094  LDELRARKATDEQGARKAADELLARRSADGLLARKPTDELGARKAADELLARRSADELLA  2273
             LDELRARKATDEQGA+K ADELLARRSAD LLARK TDELGA KA DEL ARRSADELLA
Sbjct  43    LDELRARKATDEQGAKKTADELLARRSADELLARKATDELGA-KAVDELSARRSADELLA  101

Query  2274  RRTAS  2288
             RRT+S
Sbjct  102   RRTSS  106


Range 2: 21 to 44

Score:40.0 bits(92), Expect:2e-24,
Method:Compositional matrix adjust.,
Identities:20/24(83%), Positives:21/24(87%), Gaps:0/24(0%)

Query  1996  ARKAAAELLAGRTTSQRNCRRTVS  2067
             ARKAA ELLA RTTSQRNCRRT+
Sbjct  21    ARKAADELLARRTTSQRNCRRTLD  44


Range 3: 11 to 77

Score:60.5 bits(145), Expect:4e-10,
Method:Compositional matrix adjust.,
Identities:43/67(64%), Positives:47/67(70%), Gaps:10/67(14%)

Query  2109  ARKATDEQG-ARKAADELLARRSA---------DGLLARKPTDELGARKAADELLARRSA  2258
             A K TD +  ARKAADELLARR+          D L ARK TDE GA+K ADELLARRSA
Sbjct  11    ASKLTDPRAIARKAADELLARRTTSQRNCRRTLDELRARKATDEQGAKKTADELLARRSA  70

Query  2259  DELLARR  2279
             DELLAR+
Sbjct  71    DELLARK  77


>unnamed protein product [Arabidopsis thaliana]
Sequence ID: CAA0381910.1 Length: 128
Range 1: 65 to 127

Score:86.3 bits(212), Expect:4e-24,
Method:Compositional matrix adjust.,
Identities:52/63(83%), Positives:54/63(85%), Gaps:0/63(0%)

Query  2097  DELRARKATDEQGARKAADELLARRSADGLLARKPTDELGARKAADELLARRSADELLAR  2276
             DEL ARKATDE G R AADELLA RSAD LLARK TDEL A+KAADELLARRSAD+LLA
Sbjct  65    DELLARKATDELGVRMAADELLAMRSADELLARKATDELVAKKAADELLARRSADDLLAI  124

Query  2277  RTA  2285
             RTA
Sbjct  125   RTA  127


Range 2: 26 to 64

Score:48.9 bits(115), Expect:4e-24,
Method:Compositional matrix adjust.,
Identities:29/42(69%), Positives:32/42(76%), Gaps:3/42(7%)

Query  1982  AGNSKLGRLPPSC*LGELRARETADEL*ARKSAD*VLARRTA  2107
             AGNSKLGRLP S    ELRARE  DE  ARK+AD +LARR+A
Sbjct  26    AGNSKLGRLPASY---ELRAREATDEQGARKAADDLLARRSA  64


Range 3: 28 to 98

Score:93.6 bits(231), Expect:2e-21,
Method:Compositional matrix adjust.,
Identities:55/74(74%), Positives:61/74(82%), Gaps:3/74(4%)

Query  2058  NCELGKVQTRC*LDELRARKATDEQGARKAADELLARRSADGLLARKPTDELGARKAADE  2237
             N +LG++       ELRAR+ATDEQGARKAAD+LLARRSAD LLARK TDELG R AADE
Sbjct  28    NSKLGRLPASY---ELRAREATDEQGARKAADDLLARRSADELLARKATDELGVRMAADE  84

Query  2238  LLARRSADELLARR  2279
             LLA RSADELLAR+
Sbjct  85    LLAMRSADELLARK  98


>unnamed protein product [Arabidopsis thaliana]
Sequence ID: VYS60963.1 Length: 145
Range 1: 1 to 43

Score:81.6 bits(200), Expect:4e-17,
Method:Compositional matrix adjust.,
Identities:39/43(91%), Positives:42/43(97%), Gaps:0/43(0%)

Query  4305  MLRDINVYLKQVQPVMRPNSIKPVSFGIVFLENPDELGSDIPI  4177
             MLRDINVYLKQVQPV+ P+SIKPVSFGIVFLENPDELG+DIPI
Sbjct  1     MLRDINVYLKQVQPVIGPSSIKPVSFGIVFLENPDELGNDIPI  43


>unknown protein [Arabidopsis thaliana]
Sequence ID: ABF59358.1 Length: 150
Range 1: 1 to 43

Score:81.6 bits(200), Expect:4e-17,
Method:Compositional matrix adjust.,
Identities:39/43(91%), Positives:42/43(97%), Gaps:0/43(0%)

Query  4305  MLRDINVYLKQVQPVMRPNSIKPVSFGIVFLENPDELGSDIPI  4177
             MLRDINVYLKQVQPV+ P+SIKPVSFGIVFLENPDELG+DIPI
Sbjct  1     MLRDINVYLKQVQPVIGPSSIKPVSFGIVFLENPDELGNDIPI  43


>unnamed protein product [Arabidopsis thaliana]
Sequence ID: CAA0395933.1 Length: 105
Range 1: 1 to 48

Score:63.5 bits(153), Expect:4e-11,
Method:Compositional matrix adjust.,
Identities:33/48(69%), Positives:37/48(77%), Gaps:5/48(10%)

Query  4305  MLRDINVYLKQVQPVMRPNSIKPVSFGIV-----FLENPDELGSDIPI  4177
             MLRDINVYLKQVQPVMRPNSIKPVSFGI+     FL+ P    S++ I
Sbjct  1     MLRDINVYLKQVQPVMRPNSIKPVSFGIIAPRVSFLKKPTSPRSELSI  48


>hypothetical protein AT5G10336 [Arabidopsis thaliana]
Sequence ID: NP_001119203.1 Length: 40
>hypothetical protein AT5G10336 [Arabidopsis thaliana]
Sequence ID: AED91526.1 Length: 40
Range 1: 1 to 28

Score:59.3 bits(142), Expect:2e-10,
Method:Compositional matrix adjust.,
Identities:28/28(100%), Positives:28/28(100%), Gaps:0/28(0%)

Query  4260  MRPNSIKPVSFGIVFLENPDELGSDIPI  4177
             MRPNSIKPVSFGIVFLENPDELGSDIPI
Sbjct  1     MRPNSIKPVSFGIVFLENPDELGSDIPI  28


>unnamed protein product [Arabidopsis thaliana]
Sequence ID: CAA0393036.1 Length: 105
Range 1: 1 to 48

Score:61.2 bits(147), Expect:2e-10,
Method:Compositional matrix adjust.,
Identities:32/48(67%), Positives:36/48(75%), Gaps:5/48(10%)

Query  4305  MLRDINVYLKQVQPVMRPNSIKPVSFGIV-----FLENPDELGSDIPI  4177
             MLRDINVYLKQVQPVMRPNSIK VSFGI+     FL+ P    S++ I
Sbjct  1     MLRDINVYLKQVQPVMRPNSIKQVSFGIIAPRVSFLKKPTSPRSELSI  48


>hypothetical protein AXX17_ATUG01520 [Arabidopsis thaliana]
Sequence ID: OAO89355.1 Length: 87
>hypothetical protein AXX17_AT1G38100 [Arabidopsis thaliana]
Sequence ID: OAP14064.1 Length: 87
Range 1: 45 to 86

Score:57.0 bits(136), Expect:3e-09,
Method:Compositional matrix adjust.,
Identities:26/42(62%), Positives:32/42(76%), Gaps:0/42(0%)

Query  4954  VTCSRIAPEISFPGSLTMVQSDIYSESYHRICKADAVCFNTD  4829
             VTC RIAPEI  PG+LT+ +SDI ++SY R  + DAVCF TD
Sbjct  45    VTCPRIAPEIRVPGNLTVARSDISAQSYRRFSEVDAVCFFTD  86


Range 2: 3 to 58

Score:53.5 bits(127), Expect:8e-08,
Method:Composition-based stats.,
Identities:31/56(55%), Positives:36/56(64%), Gaps:8/56(14%)

Query  5082  GTQTTLFAYYSSKFSVNAHQAKIIQSVYLLGDFPSGSNEPGDV--------*RVPG  4939
             G + TLFAY+SSK+S+ AHQAK I+SVYL G   S  NEP DV         RVPG
Sbjct  3     GRKRTLFAYHSSKYSMYAHQAKTIRSVYLWGGHLSRFNEPCDVTCPRIAPEIRVPG  58


>hypothetical protein [Arabidopsis thaliana]
Sequence ID: AAC69120.1 Length: 893
Range 1: 819 to 870

Score:60.5 bits(145), Expect:4e-08,
Method:Compositional matrix adjust.,
Identities:29/52(56%), Positives:40/52(76%), Gaps:0/52(0%)

Query  1698  VVMNSDRQWSNKTTRKITIFSGIISNSHTKPQKKADHDLRLWFESYDRFSKT  1543
             V ++S+ + S K+T+KI  F G I+NSHTK Q+KADH L L FESY++F+KT
Sbjct  819   VCLDSNHKRSGKSTQKIKQFGGNINNSHTKLQEKADHGLDLRFESYNQFTKT  870


Query #87: XLOC_008459 Query ID: lcl|Query_76621 Length: 919

No significant similarity found.


Query #88: XLOC_008578 Query ID: lcl|Query_76622 Length: 1089

Sequences producing significant alignments:
                                                                  Max    Total Query   E   Per.
Description                                                       Score  Score cover Value Ident  Accession
hypothetical protein AT4G03740 [Arabidopsis thaliana]             110    225   42%   7e-27 79.41  NP_192283.1
unnamed protein product [Arabidopsis thaliana]                    88.2   122   42%   5e-21 40.74  VYS61898.1


Alignments:
>hypothetical protein AT4G03740 [Arabidopsis thaliana]
Sequence ID: NP_192283.1 Length: 345
>hypothetical protein [Arabidopsis thaliana]
Sequence ID: AAD15330.1 Length: 345 >hypothetical protein AT4G03740 [Arabidopsis thaliana]
Sequence ID: AEE82351.1 Length: 345 >hypothetical protein [Arabidopsis thaliana]
Sequence ID: CAB77859.1 Length: 345
Range 1: 72 to 138

Score:110 bits(276), Expect:7e-27,
Method:Compositional matrix adjust.,
Identities:54/68(79%), Positives:59/68(86%), Gaps:1/68(1%)

Query  685  LGSCPHTLCTCCVSQVCCIKTWGPHEHLEGKVGSRQGVSKRLEPRRMASLKRVLGIAGDC  864
            LGSCPHTL  C VSQVCCIK + PHEHLE KVGSR+ VSKRLEPR MASL+RVL +AGDC
Sbjct  72   LGSCPHTLWPCWVSQVCCIKIFRPHEHLERKVGSRR-VSKRLEPRHMASLQRVLSVAGDC  130

Query  865  LYRKNQLK  888
            LYR+NQLK
Sbjct  131  LYRENQLK  138


Range 2: 29 to 156

Score:80.5 bits(197), Expect:9e-19,
Method:Compositional matrix adjust.,
Identities:52/135(39%), Positives:66/135(48%), Gaps:9/135(6%)

Query  545  LGYSKNSISKSILWMLHGMLKYPYPAYNDMPGDDQELWFRNFVHEFNWDPVHT--PSVRV  718
             G+SKN I KSIL MLHGMLK+PYP YNDMP ++QE+WFR F       P HT  P
Sbjct  29   FGFSKNGIFKSILRMLHGMLKHPYPTYNDMPRNNQEIWFRTFA--LGSCP-HTLWPCWVS  85

Query  719  AFHKYAALKLGDHMNTWKGKWEVGKECPKGLNPGVWPRLKEY*ALPETACTAKTNSKKPE  898
                    +  +H+    G   V K     L P     L+   ++       +   K  E
Sbjct  86   QVCCIKIFRPHEHLERKVGSRRVSKR----LEPRHMASLQRVLSVAGDCLYRENQLKISE  141

Query  899  ERTWWQMYGGAQCMY  943
            +R WWQ Y GAQC +
Sbjct  142  KRMWWQRYDGAQCGF  156


Range 3: 8 to 28

Score:33.9 bits(76), Expect:9e-19,
Method:Compositional matrix adjust.,
Identities:17/22(77%), Positives:18/22(81%), Gaps:1/22(4%)

Query  402  QQTGQTLLRRLHPKRLE*ERTW  467
            QQ G+TLLRRLHPKRLE   TW
Sbjct  8    QQPGRTLLRRLHPKRLE-GSTW  28


>unnamed protein product [Arabidopsis thaliana]
Sequence ID: VYS61898.1 Length: 345
Range 1: 29 to 156

Score:88.2 bits(217), Expect:5e-21,
Method:Compositional matrix adjust.,
Identities:55/135(41%), Positives:68/135(50%), Gaps:9/135(6%)

Query  545  LGYSKNSISKSILWMLHGMLKYPYPAYNDMPGDDQELWFRNFVHEFNWDPVHT--PSVRV  718
             G+SKN ISKSIL MLHGMLK+PYP YNDMP D+QE+WFR F       P HT  P
Sbjct  29   FGFSKNGISKSILRMLHGMLKHPYPTYNDMPRDNQEIWFRTFA--LGSCP-HTLWPCWVS  85

Query  719  AFHKYAALKLGDHMNTWKGKWEVGKECPKGLNPGVWPRLKEY*ALPETACTAKTNSKKPE  898
              +     +  +H+    G   V K     L P     L+    +       +   KK E
Sbjct  86   QVYCIKIFRPHEHLERKVGSRRVSKR----LEPRHMASLQRVLGVAGDCLYRENQLKKSE  141

Query  899  ERTWWQMYGGAQCMY  943
            +R WWQ Y GAQC +
Sbjct  142  KRMWWQRYDGAQCGF  156


Range 2: 8 to 28

Score:33.9 bits(76), Expect:5e-21,
Method:Compositional matrix adjust.,
Identities:17/22(77%), Positives:18/22(81%), Gaps:1/22(4%)

Query  402  QQTGQTLLRRLHPKRLE*ERTW  467
            QQ G+TLLRRLHPKRLE   TW
Sbjct  8    QQPGRTLLRRLHPKRLE-GSTW  28


Query #89: XLOC_008726 Query ID: lcl|Query_76623 Length: 510

Sequences producing significant alignments:
                                                                  Max    Total Query   E   Per.
Description                                                       Score  Score cover Value Ident  Accession
hypothetical protein AXX17_AT2G24510 [Arabidopsis thaliana]       95.5   165   47%   1e-34 97.83  OAP11137.1
unnamed protein product [Arabidopsis thaliana]                    86.7   164   65%   4e-34 92.86  CAA0372893.1
hypothetical protein AXX17_AT2G24510 [Arabidopsis thaliana]       70.1   112   33%   6e-19 91.18  OAP11136.1


Alignments:
>hypothetical protein AXX17_AT2G24510 [Arabidopsis thaliana]
Sequence ID: OAP11137.1 Length: 102
Range 1: 57 to 102

Score:95.5 bits(236), Expect:1e-34,
Method:Compositional matrix adjust.,
Identities:45/46(98%), Positives:45/46(97%), Gaps:0/46(0%)

Query  254  GHMCSDSHLFHNILYFHNTKL*VVDDNTYRRPTLSLLVCHCKCILI  391
            GHMCSDSHLFHNILYFHNTKL VVDDNTYRRPTLSLLVCHCKCILI
Sbjct  57   GHMCSDSHLFHNILYFHNTKLQVVDDNTYRRPTLSLLVCHCKCILI  102


Range 2: 1 to 34

Score:70.1 bits(170), Expect:1e-34,
Method:Compositional matrix adjust.,
Identities:31/34(91%), Positives:33/34(97%), Gaps:0/34(0%)

Query  55   MLLYDQIGRDYRPEVVSNPLCSSSQPTVLNVDEF  156
            MLLY QIGRDYRPEVVSNPLCSSSQPT+LN+DEF
Sbjct  1    MLLYHQIGRDYRPEVVSNPLCSSSQPTMLNIDEF  34


>unnamed protein product [Arabidopsis thaliana]
Sequence ID: CAA0372893.1 Length: 110
Range 1: 2 to 43

Score:86.7 bits(213), Expect:4e-34,
Method:Compositional matrix adjust.,
Identities:39/42(93%), Positives:41/42(97%), Gaps:0/42(0%)

Query  31   IDYRRKIFMLLYDQIGRDYRPEVVSNPLCSSSQPTVLNVDEF  156
            IDYRRKIFMLLY QIGRDYRPEVVSNPLCSSSQPT+LN+DEF
Sbjct  2    IDYRRKIFMLLYHQIGRDYRPEVVSNPLCSSSQPTMLNIDEF  43


Range 2: 45 to 102

Score:77.4 bits(189), Expect:4e-34,
Method:Compositional matrix adjust.,
Identities:45/73(62%), Positives:49/73(67%), Gaps:15/73(20%)

Query  146  STSLVNLIGEDCESYRSLV*NHGPVS**SFMYFFFPGHMCSDSHLFHNILYFHNTKL*VV  325
            S  +VN IG  CE+   +V +H              GHMCSDSHLFHNILYFHNTKL VV
Sbjct  45   SKRIVNHIGVLCET---MVQSHDK------------GHMCSDSHLFHNILYFHNTKLQVV  89

Query  326  DDNTYRRPTLSLL  364
            DDNTYRRPTLSLL
Sbjct  90   DDNTYRRPTLSLL  102


>hypothetical protein AXX17_AT2G24510 [Arabidopsis thaliana]
Sequence ID: OAP11136.1 Length: 67
Range 1: 1 to 34

Score:70.1 bits(170), Expect:6e-19,
Method:Compositional matrix adjust.,
Identities:31/34(91%), Positives:33/34(97%), Gaps:0/34(0%)

Query  55   MLLYDQIGRDYRPEVVSNPLCSSSQPTVLNVDEF  156
            MLLY QIGRDYRPEVVSNPLCSSSQPT+LN+DEF
Sbjct  1    MLLYHQIGRDYRPEVVSNPLCSSSQPTMLNIDEF  34


Range 2: 36 to 58

Score:42.7 bits(99), Expect:6e-19,
Method:Compositional matrix adjust.,
Identities:19/23(83%), Positives:21/23(91%), Gaps:0/23(0%)

Query  168  SERIVNHIGVLFETMVQSHDKAL  236
            S+RIVNHIGVL ETMVQSHDK +
Sbjct  36   SKRIVNHIGVLCETMVQSHDKGM  58


Query #90: XLOC_008854 Query ID: lcl|Query_76624 Length: 550

No significant similarity found.


Query #91: XLOC_008920 Query ID: lcl|Query_76625 Length: 893

Sequences producing significant alignments:
                                                                  Max    Total Query   E   Per.
Description                                                       Score  Score cover Value Ident  Accession
hypothetical protein AT2G13105 [Arabidopsis thaliana]             272    272   45%   1e-92 100.00 NP_001323576.1
unnamed protein product [Arabidopsis thaliana]                    268    268   45%   3e-91 98.53  CAA0360418.1
unnamed protein product [Arabidopsis thaliana]                    266    266   45%   2e-90 97.79  VYS52285.1
hypothetical protein AXX17_ATUG00720 [Arabidopsis thaliana]       266    266   55%   2e-81 80.72  OAO89433.1
hypothetical protein AXX17_AT3G35420 [Arabidopsis thaliana]       174    174   35%   2e-54 85.71  OAP04245.1
hypothetical protein AXX17_AT2G08370 [Arabidopsis thaliana]       180    180   30%   4e-52 95.65  OAP09767.1
unnamed protein product [Arabidopsis thaliana]                    149    149   41%   4e-44 62.60  VYS61899.1
hypothetical protein AT4G03728 [Arabidopsis thaliana]             102    102   22%   4e-26 70.59  NP_001154205.1


Alignments:
>hypothetical protein AT2G13105 [Arabidopsis thaliana]
Sequence ID: NP_001323576.1 Length: 136
>hypothetical protein AT2G13105 [Arabidopsis thaliana]
Sequence ID: ANM61353.1 Length: 136
Range 1: 1 to 136

Score:272 bits(695), Expect:1e-92,
Method:Compositional matrix adjust.,
Identities:136/136(100%), Positives:136/136(100%), Gaps:0/136(0%)

Query  802  MSQNLDNELRIYSLEKTNEALKENNEALKEDMLVLKSGVGKVMDELATTRLAMNAIMKSL  623
            MSQNLDNELRIYSLEKTNEALKENNEALKEDMLVLKSGVGKVMDELATTRLAMNAIMKSL
Sbjct  1    MSQNLDNELRIYSLEKTNEALKENNEALKEDMLVLKSGVGKVMDELATTRLAMNAIMKSL  60

Query  622  GVTIIPPELVALARDAGLGVHGSAGGRDSSPAPIVSPAPTTSPAVSLAVSRAASPTVPST  443
            GVTIIPPELVALARDAGLGVHGSAGGRDSSPAPIVSPAPTTSPAVSLAVSRAASPTVPST
Sbjct  61   GVTIIPPELVALARDAGLGVHGSAGGRDSSPAPIVSPAPTTSPAVSLAVSRAASPTVPST  120

Query  442  RTQQSPLDAWCASLGI  395
            RTQQSPLDAWCASLGI
Sbjct  121  RTQQSPLDAWCASLGI  136


>unnamed protein product [Arabidopsis thaliana]
Sequence ID: CAA0360418.1 Length: 136
Range 1: 1 to 136

Score:268 bits(685), Expect:3e-91,
Method:Compositional matrix adjust.,
Identities:134/136(99%), Positives:135/136(99%), Gaps:0/136(0%)

Query  802  MSQNLDNELRIYSLEKTNEALKENNEALKEDMLVLKSGVGKVMDELATTRLAMNAIMKSL  623
            MSQNLDNELRIYSLEKTNEALKENNEALKEDMLVLKSGVGKVMDELATTRLA+NAIMKSL
Sbjct  1    MSQNLDNELRIYSLEKTNEALKENNEALKEDMLVLKSGVGKVMDELATTRLALNAIMKSL  60

Query  622  GVTIIPPELVALARDAGLGVHGSAGGRDSSPAPIVSPAPTTSPAVSLAVSRAASPTVPST  443
            GVTIIPPELVALARDAGLGVHGSAGGRD SPAPIVSPAPTTSPAVSLAVSRAASPTVPST
Sbjct  61   GVTIIPPELVALARDAGLGVHGSAGGRDGSPAPIVSPAPTTSPAVSLAVSRAASPTVPST  120

Query  442  RTQQSPLDAWCASLGI  395
            RTQQSPLDAWCASLGI
Sbjct  121  RTQQSPLDAWCASLGI  136


>unnamed protein product [Arabidopsis thaliana]
Sequence ID: VYS52285.1 Length: 136
Range 1: 1 to 136

Score:266 bits(680), Expect:2e-90,
Method:Compositional matrix adjust.,
Identities:133/136(98%), Positives:134/136(98%), Gaps:0/136(0%)

Query  802  MSQNLDNELRIYSLEKTNEALKENNEALKEDMLVLKSGVGKVMDELATTRLAMNAIMKSL  623
            MSQNLDNELRIYSLEKTNEALKENNEALKEDMLVLKSGVGKVMDELATTRLA+NAIMKSL
Sbjct  1    MSQNLDNELRIYSLEKTNEALKENNEALKEDMLVLKSGVGKVMDELATTRLALNAIMKSL  60

Query  622  GVTIIPPELVALARDAGLGVHGSAGGRDSSPAPIVSPAPTTSPAVSLAVSRAASPTVPST  443
            GVTIIPPELVALARDAGLGVHGSAGGRD SPAPIVSPAPTTSPAVSLAVSRAASPTVPST
Sbjct  61   GVTIIPPELVALARDAGLGVHGSAGGRDGSPAPIVSPAPTTSPAVSLAVSRAASPTVPST  120

Query  442  RTQQSPLDAWCASLGI  395
            R QQSPLDAWCASLGI
Sbjct  121  RKQQSPLDAWCASLGI  136


>hypothetical protein AXX17_ATUG00720 [Arabidopsis thaliana]
Sequence ID: OAO89433.1 Length: 967
Range 1: 812 to 967

Score:266 bits(679), Expect:2e-81,
Method:Composition-based stats.,
Identities:134/166(81%), Positives:141/166(84%), Gaps:10/166(6%)

Query  892  QRGQSKNGYIYG*GSVQYRDINRSEKPAVAMSQNLDNELRIYSLEKTNEALKENNEALKE  713
            +RGQSKNGYIYG GSVQYRD+NRSEK  VAMSQNLDNELRIY LEKTNEALK+NNEALKE
Sbjct  812  KRGQSKNGYIYGLGSVQYRDVNRSEKHVVAMSQNLDNELRIYDLEKTNEALKKNNEALKE  871

Query  712  DMLVLKSGVGKVMDELATTRLAMNAIMKSLGVTIIPPELVALARDAGLGVHGSAGGRDSS  533
            DML+LKSGVGKVMDELATTRLA+NAIMKSLGVTIIPPELVA A  AGLGVHGSAGG D S
Sbjct  872  DMLLLKSGVGKVMDELATTRLALNAIMKSLGVTIIPPELVAGAATAGLGVHGSAGGHDGS  931

Query  532  PAPIVSPAPTTSPAVSLAVSRAASPTVPSTRTQQSPLDAWCASLGI  395
            PAP  SP           VS AASPT+P TR QQSPLDAWCASLG+
Sbjct  932  PAPTTSP----------VVSPAASPTIPPTRAQQSPLDAWCASLGL  967


>hypothetical protein AXX17_AT3G35420 [Arabidopsis thaliana]
Sequence ID: OAP04245.1 Length: 105
Range 1: 1 to 105

Score:174 bits(440), Expect:2e-54,
Method:Compositional matrix adjust.,
Identities:90/105(86%), Positives:93/105(88%), Gaps:0/105(0%)

Query  709  MLVLKSGVGKVMDELATTRLAMNAIMKSLGVTIIPPELVALARDAGLGVHGSAGGRDSSP  530
            MLVLKSGVGKVMDEL TT+LA+NAIMKSL VTIIPPELVA A  AGLGVHGSAGGRD SP
Sbjct  1    MLVLKSGVGKVMDELVTTQLALNAIMKSLRVTIIPPELVARAATAGLGVHGSAGGRDGSP  60

Query  529  APIVSPAPTTSPAVSLAVSRAASPTVPSTRTQQSPLDAWCASLGI  395
            API SPAPTTSPAVS  VS   SPTVPST+TQQSPLDAWCASLGI
Sbjct  61   APIASPAPTTSPAVSPDVSHVGSPTVPSTKTQQSPLDAWCASLGI  105


>hypothetical protein AXX17_AT2G08370 [Arabidopsis thaliana]
Sequence ID: OAP09767.1 Length: 533
Range 1: 442 to 533

Score:180 bits(457), Expect:4e-52,
Method:Compositional matrix adjust.,
Identities:88/92(96%), Positives:91/92(98%), Gaps:0/92(0%)

Query  670  ELATTRLAMNAIMKSLGVTIIPPELVALARDAGLGVHGSAGGRDSSPAPIVSPAPTTSPA  491
            +LATTR+A+NAIMKSLGVTIIPPELVALARDAGLGVHGSAGGRD SPAPIVSPAPTTSPA
Sbjct  442  KLATTRIALNAIMKSLGVTIIPPELVALARDAGLGVHGSAGGRDGSPAPIVSPAPTTSPA  501

Query  490  VSLAVSRAASPTVPSTRTQQSPLDAWCASLGI  395
            VSLAVSRAASPTVPSTRTQQSPLDAWCASLGI
Sbjct  502  VSLAVSRAASPTVPSTRTQQSPLDAWCASLGI  533


>unnamed protein product [Arabidopsis thaliana]
Sequence ID: VYS61899.1 Length: 155
Range 1: 31 to 147

Score:149 bits(375), Expect:4e-44,
Method:Compositional matrix adjust.,
Identities:77/123(63%), Positives:90/123(73%), Gaps:6/123(4%)

Query  892  QRGQSKNGYIYG*GSVQYRDINRSEKPAVAMSQNLDNELRIYSLEKTNEALKENNEALKE  713
            +RGQ+K GYIYG GSVQYRDIN S+K A  MS NLDNE+R+  LEK N ALKE+NEALKE
Sbjct  31   KRGQNKKGYIYGLGSVQYRDINPSQKVATTMSHNLDNEMRMNGLEKNNHALKEHNEALKE  90

Query  712  DMLVLKSGVGKVMDELATTRLAMNAIMKSLGVTIIPPELVALARDAGLGVHGSAGGRDSS  533
            DM+VLK+GVG+VMDELAT RLA+NAIMKSL   ++     A  RD    +     GRD S
Sbjct  91   DMVVLKTGVGRVMDELATARLALNAIMKSLEFNLM---WTAARRDGSFAL---VAGRDGS  144

Query  532  PAP  524
            P P
Sbjct  145  PPP  147


>hypothetical protein AT4G03728 [Arabidopsis thaliana]
Sequence ID: NP_001154205.1 Length: 141
>hypothetical protein AT4G03728 [Arabidopsis thaliana]
Sequence ID: AEE82350.1 Length: 141
Range 1: 31 to 98

Score:102 bits(253), Expect:4e-26,
Method:Compositional matrix adjust.,
Identities:48/68(71%), Positives:57/68(83%), Gaps:0/68(0%)

Query  892  QRGQSKNGYIYG*GSVQYRDINRSEKPAVAMSQNLDNELRIYSLEKTNEALKENNEALKE  713
            +RGQ+K GYIYG GSVQYRDIN S+K A  MS NLDNE+R+  LEK N ALK++NEALKE
Sbjct  31   KRGQNKKGYIYGLGSVQYRDINPSQKVATTMSHNLDNEMRMNGLEKNNHALKKHNEALKE  90

Query  712  DMLVLKSG  689
            DM+VLK+G
Sbjct  91   DMVVLKTG  98


Query #92: XLOC_008962 Query ID: lcl|Query_76626 Length: 1432

No significant similarity found.


Query #93: XLOC_008978 Query ID: lcl|Query_76627 Length: 479

No significant similarity found.


Query #94: XLOC_009073 Query ID: lcl|Query_76628 Length: 1234

No significant similarity found.


Query #95: XLOC_009078 Query ID: lcl|Query_76629 Length: 391

Sequences producing significant alignments:
                                                                  Max    Total Query   E   Per.
Description                                                       Score  Score cover Value Ident  Accession
serine carboxypeptidase [Arabidopsis thaliana]                    93.6   93.6  46%   1e-22 67.21  BAA96893.1
unnamed protein product [Arabidopsis thaliana]                    66.6   123   62%   6e-15 48.86  CAA0405760.1
Mannose-binding lectin superfamily protein [Arabidopsis thaliana] 64.7   64.7  30%   2e-12 70.00  NP_175623.2
unnamed protein product [Arabidopsis thaliana]                    62.4   62.4  30%   1e-11 67.50  CAA0288322.1


Alignments:
>serine carboxypeptidase [Arabidopsis thaliana]
Sequence ID: BAA96893.1 Length: 512
Range 1: 440 to 500

Score:93.6 bits(231), Expect:1e-22,
Method:Composition-based stats.,
Identities:41/61(67%), Positives:51/61(83%), Gaps:1/61(1%)

Query  389  PLFQWLTNGLLVLLHQEVWGSNPSYCDLLANLQINGGKCAGDLQRCTIIT-VRQDSTQRR  213
            PL QWLT G L+LL Q+VWGS+P+YCDLL +LQINGGKCAGDLQRCTI+  +R+  T+R
Sbjct  440  PLGQWLTKGSLMLLQQDVWGSSPNYCDLLVDLQINGGKCAGDLQRCTIMDLIRKKKTKRS  499

Query  212  Y  210
            +
Sbjct  500  F  500


>unnamed protein product [Arabidopsis thaliana]
Sequence ID: CAA0405760.1 Length: 84
Range 1: 1 to 82

Score:66.6 bits(161), Expect:6e-15,
Method:Compositional matrix adjust.,
Identities:43/88(49%), Positives:50/88(56%), Gaps:12/88(13%)

Query  378  MVDQWFISASTPRGLGFESQLLRFISKFTD*WRQVCRRSPTMYDYNRSP------GFYAE  217
            MVDQ FI+ASTPRG G     LR+     D   Q+        D  R        GFYAE
Sbjct  1    MVDQGFINASTPRGRGSS---LRYCDLLAD--LQI-NGGKCAKDLQRCTIIAVHLGFYAE  54

Query  216  KILNMSIKRMEKEKKVSYVVNLIRIGRK  133
            KIL +SIKRMEKE  VS V++LIR+GR
Sbjct  55   KILKVSIKRMEKENDVSSVMDLIRVGRN  82


Range 2: 16 to 47

Score:57.0 bits(136), Expect:3e-11,
Method:Compositional matrix adjust.,
Identities:26/32(81%), Positives:28/32(87%), Gaps:0/32(0%)

Query  332  GSNPSYCDLLANLQINGGKCAGDLQRCTIITV  237
            GS+  YCDLLA+LQINGGKCA DLQRCTII V
Sbjct  16   GSSLRYCDLLADLQINGGKCAKDLQRCTIIAV  47


>Mannose-binding lectin superfamily protein [Arabidopsis thaliana]
Sequence ID: NP_175623.2 Length: 615
>Mannose-binding lectin superfamily protein [Arabidopsis thaliana]
Sequence ID: AEE32756.1 Length: 615
Range 1: 1 to 40

Score:64.7 bits(156), Expect:2e-12,
Method:Composition-based stats.,
Identities:28/40(70%), Positives:37/40(92%), Gaps:0/40(0%)

Query  255  MYDYNRSPGFYAEKILNMSIKRMEKEKKVSYVVNLIRIGR  136
            MY+Y+RSPGFYA+KIL MSIKRM+K+ ++S VV+LI+IGR
Sbjct  1    MYNYSRSPGFYAKKILKMSIKRMKKKNEMSSVVDLIKIGR  40


>unnamed protein product [Arabidopsis thaliana]
Sequence ID: CAA0288322.1 Length: 614
Range 1: 1 to 40

Score:62.4 bits(150), Expect:1e-11,
Method:Composition-based stats.,
Identities:27/40(68%), Positives:36/40(90%), Gaps:0/40(0%)

Query  255  MYDYNRSPGFYAEKILNMSIKRMEKEKKVSYVVNLIRIGR  136
            MY+Y+ SPGFYA+KIL MSIKRM+K+ ++S VV+LI+IGR
Sbjct  1    MYNYSHSPGFYAKKILKMSIKRMKKKNEMSSVVDLIKIGR  40


Query #96: XLOC_009198 Query ID: lcl|Query_76630 Length: 1548

No significant similarity found.


Query #97: XLOC_009253 Query ID: lcl|Query_76631 Length: 433

No significant similarity found.


Query #98: XLOC_009261 Query ID: lcl|Query_76632 Length: 1324

Sequences producing significant alignments:
                                                                  Max    Total Query   E   Per.
Description                                                       Score  Score cover Value Ident  Accession
unnamed protein product [Arabidopsis thaliana]                    137    137   17%   3e-38 77.92  CAA0264251.1
unnamed protein product [Arabidopsis thaliana]                    137    137   17%   7e-38 76.62  VYS66939.1
zinc ion binding protein [Arabidopsis thaliana]                   137    137   17%   8e-38 76.62  NP_197073.1
unknown [Arabidopsis thaliana]                                    137    137   17%   8e-38 76.62  ABK28697.1
unnamed protein product [Arabidopsis thaliana]                    136    136   17%   1e-37 76.62  CAA0365701.1
zinc ion binding protein [Arabidopsis thaliana]                   135    135   17%   2e-37 76.62  NP_001190673.2
unknown protein; 62986-63538 [Arabidopsis thaliana]               132    132   17%   3e-36 75.32  AAG51213.1
hypothetical protein [Arabidopsis thaliana]                       137    137   17%   8e-36 76.62  CAB80843.1
unnamed protein product [Arabidopsis thaliana]                    114    114   15%   1e-29 73.53  CAA0402823.1
unnamed protein product [Arabidopsis thaliana]                    79.7   79.7  18%   5e-17 42.35  BAB09034.1
unknown [Arabidopsis thaliana]                                    78.6   78.6  18%   1e-16 38.10  ABK28583.1
hypothetical protein At3g42810 [Arabidopsis thaliana]             78.6   78.6  18%   1e-16 38.10  ABE65983.1
hypothetical protein [Arabidopsis thaliana]                       76.6   76.6  18%   6e-16 40.48  CAA16722.1
putative protein [Arabidopsis thaliana]                           68.2   68.2  18%   3e-13 34.52  CAB68121.1
zinc ion-binding protein [Arabidopsis thaliana]                   63.5   63.5  17%   5e-12 36.36  NP_001327268.1
unnamed protein product [Arabidopsis thaliana]                    62.0   62.0  17%   7e-11 36.84  CAA0384158.1
Hypothetical protein T10I18.4 [Arabidopsis thaliana]              60.8   60.8  17%   2e-10 34.21  AAF88011.1
hypothetical protein AXX17_AT3G33460 [Arabidopsis thaliana]       63.2   63.2  16%   6e-10 40.28  OAP03513.1
unknown [Arabidopsis thaliana]                                    58.9   58.9  18%   1e-09 32.50  ABK28780.1
unknown protein [Arabidopsis thaliana]                            58.9   58.9  18%   1e-09 32.50  ABF59287.1
GRF zinc finger protein [Arabidopsis thaliana]                    57.8   57.8  12%   1e-09 40.32  NP_001324832.1
T12C24.25 [Arabidopsis thaliana]                                  58.2   58.2  17%   2e-09 34.21  AAF88091.1
GRF zinc finger protein [Arabidopsis thaliana]                    57.8   57.8  16%   2e-09 31.03  NP_001320823.1
hypothetical protein [Arabidopsis thaliana]                       56.2   56.2  11%   7e-09 46.94  ABF59281.1
unknown [Arabidopsis thaliana]                                    56.2   56.2  11%   7e-09 46.94  ABK28512.1
unnamed protein product [Arabidopsis thaliana]                    55.5   55.5  12%   1e-08 40.74  CAA0393920.1
Mutator-like transposase [Arabidopsis thaliana]                   58.5   58.5  17%   2e-08 43.42  AAD15524.1
unnamed protein product [Arabidopsis thaliana]                    54.3   54.3  16%   2e-08 31.71  VYS52155.1
hypothetical protein [Arabidopsis thaliana]                       55.1   55.1  12%   2e-08 40.68  AAC69115.1
zinc ion binding protein [Arabidopsis thaliana]                   53.9   53.9  12%   4e-08 40.74  NP_178758.1
zinc ion-binding protein [Arabidopsis thaliana]                   53.5   53.5  12%   4e-08 38.89  NP_001325185.1
unnamed protein product [Arabidopsis thaliana]                    53.9   53.9  11%   5e-08 44.90  BAB01424.1


Alignments:
>unnamed protein product [Arabidopsis thaliana]
Sequence ID: CAA0264251.1 Length: 160
Range 1: 19 to 95

Score:137 bits(346), Expect:3e-38,
Method:Compositional matrix adjust.,
Identities:60/77(78%), Positives:70/77(90%), Gaps:0/77(0%)

Query  373  PPGVPSKCWCGASIITYTSKTKENPSRRFYRCAIALQRENEDHLFKWVDEALLDEIRMVE  552
            PPGVPSKCWCG  IIT+TSKTKENP RRFYRCAIA++RENE+HLFKWVDEALLDEI+MV
Sbjct  19   PPGVPSKCWCGEEIITFTSKTKENPDRRFYRCAIAMKRENEEHLFKWVDEALLDEIKMVN  78

Query  553  ENCKRVEEDVNDLKVQM  603
            E CKRV E+++DLK+ +
Sbjct  79   EKCKRVAENISDLKMNV  95


>unnamed protein product [Arabidopsis thaliana]
Sequence ID: VYS66939.1 Length: 169
Range 1: 19 to 95

Score:137 bits(344), Expect:7e-38,
Method:Compositional matrix adjust.,
Identities:59/77(77%), Positives:70/77(90%), Gaps:0/77(0%)

Query  373  PPGVPSKCWCGASIITYTSKTKENPSRRFYRCAIALQRENEDHLFKWVDEALLDEIRMVE  552
            PPGVPSKCWCG  IIT+TSKTKENP RRFYRCAIA++RENE+HLFKWVDEALLDEI+MV
Sbjct  19   PPGVPSKCWCGEEIITFTSKTKENPYRRFYRCAIAMKRENEEHLFKWVDEALLDEIKMVN  78

Query  553  ENCKRVEEDVNDLKVQM  603
            E CKRV E+++DL++ +
Sbjct  79   EKCKRVAENISDLRMNV  95


>zinc ion binding protein [Arabidopsis thaliana]
Sequence ID: NP_197073.1 Length: 169
>hypothetical protein At5g15690 [Arabidopsis thaliana]
Sequence ID: ABE66159.1 Length: 169 >zinc ion binding protein [Arabidopsis thaliana]
Sequence ID: AED92192.1 Length: 169 >putative protein [Arabidopsis thaliana]
Sequence ID: CAC01768.1 Length: 169
Range 1: 19 to 95

Score:137 bits(344), Expect:8e-38,
Method:Compositional matrix adjust.,
Identities:59/77(77%), Positives:70/77(90%), Gaps:0/77(0%)

Query  373  PPGVPSKCWCGASIITYTSKTKENPSRRFYRCAIALQRENEDHLFKWVDEALLDEIRMVE  552
            PPGVPSKCWCG  IIT+TSKTKENP RRFYRCAIA++RENE+HLFKWVDEALLDEI+MV
Sbjct  19   PPGVPSKCWCGEEIITFTSKTKENPYRRFYRCAIAMKRENEEHLFKWVDEALLDEIKMVN  78

Query  553  ENCKRVEEDVNDLKVQM  603
            E CKRV E+++DL++ +
Sbjct  79   EKCKRVAENISDLRMNV  95


>unknown, partial [Arabidopsis thaliana]
Sequence ID: ABK28697.1 Length: 170
Range 1: 19 to 95

Score:137 bits(344), Expect:8e-38,
Method:Compositional matrix adjust.,
Identities:59/77(77%), Positives:70/77(90%), Gaps:0/77(0%)

Query  373  PPGVPSKCWCGASIITYTSKTKENPSRRFYRCAIALQRENEDHLFKWVDEALLDEIRMVE  552
            PPGVPSKCWCG  IIT+TSKTKENP RRFYRCAIA++RENE+HLFKWVDEALLDEI+MV
Sbjct  19   PPGVPSKCWCGEEIITFTSKTKENPYRRFYRCAIAMKRENEEHLFKWVDEALLDEIKMVN  78

Query  553  ENCKRVEEDVNDLKVQM  603
            E CKRV E+++DL++ +
Sbjct  79   EKCKRVAENISDLRMNV  95


>unnamed protein product [Arabidopsis thaliana]
Sequence ID: CAA0365701.1 Length: 160
>unnamed protein product [Arabidopsis thaliana]
Sequence ID: VYS49614.1 Length: 160 >unnamed protein product [Arabidopsis thaliana]
Sequence ID: VYS61826.1 Length: 160
Range 1: 19 to 95

Score:136 bits(342), Expect:1e-37,
Method:Compositional matrix adjust.,
Identities:59/77(77%), Positives:70/77(90%), Gaps:0/77(0%)

Query  373  PPGVPSKCWCGASIITYTSKTKENPSRRFYRCAIALQRENEDHLFKWVDEALLDEIRMVE  552
            PPGVPSKCWCG  IIT+TSKTKENP RRFYRCAIA++RENE+HLFKWVDEALLDEI+MV
Sbjct  19   PPGVPSKCWCGEEIITFTSKTKENPYRRFYRCAIAMKRENEEHLFKWVDEALLDEIKMVN  78

Query  553  ENCKRVEEDVNDLKVQM  603
            E CKRV E+++DL++ +
Sbjct  79   EKCKRVAENISDLRMNV  95


>zinc ion binding protein [Arabidopsis thaliana]
Sequence ID: NP_001190673.2 Length: 160
>RecName: Full=Uncharacterized protein At4g04775 [Arabidopsis thaliana]
Sequence ID: Q9ZS96.1 Length: 160 >T4B21.8 gene product [Arabidopsis thaliana]
Sequence ID: AAD03442.1 Length: 160 >zinc ion binding protein [Arabidopsis thaliana]
Sequence ID: AEE82424.2 Length: 160
Range 1: 19 to 95

Score:135 bits(340), Expect:2e-37,
Method:Compositional matrix adjust.,
Identities:59/77(77%), Positives:70/77(90%), Gaps:0/77(0%)

Query  373  PPGVPSKCWCGASIITYTSKTKENPSRRFYRCAIALQRENEDHLFKWVDEALLDEIRMVE  552
            PPGVPSKCWCG  IIT+TSKTKENP RRFYRCAIA++RENE+HLFKWVDEALLDEI+MV
Sbjct  19   PPGVPSKCWCGEEIITFTSKTKENPYRRFYRCAIAMKRENEEHLFKWVDEALLDEIKMVN  78

Query  553  ENCKRVEEDVNDLKVQM  603
            E CKRV E+++DL++ +
Sbjct  79   EKCKRVVENISDLRMNV  95


>unknown protein; 62986-63538 [Arabidopsis thaliana]
Sequence ID: AAG51213.1 Length: 160
Range 1: 19 to 95

Score:132 bits(332), Expect:3e-36,
Method:Compositional matrix adjust.,
Identities:58/77(75%), Positives:69/77(89%), Gaps:0/77(0%)

Query  373  PPGVPSKCWCGASIITYTSKTKENPSRRFYRCAIALQRENEDHLFKWVDEALLDEIRMVE  552
            PP VPSKCWCG  IIT+TSKTKENP RRFYRCAIA++RENE+HLFKWVDEALLDEI+MV
Sbjct  19   PPCVPSKCWCGEEIITFTSKTKENPYRRFYRCAIAMKRENEEHLFKWVDEALLDEIKMVN  78

Query  553  ENCKRVEEDVNDLKVQM  603
            E CKRV E+++DL++ +
Sbjct  79   EKCKRVAENISDLRMNV  95


>hypothetical protein [Arabidopsis thaliana]
Sequence ID: CAB80843.1 Length: 381
Range 1: 19 to 95

Score:137 bits(346), Expect:8e-36,
Method:Compositional matrix adjust.,
Identities:59/77(77%), Positives:70/77(90%), Gaps:0/77(0%)

Query  373  PPGVPSKCWCGASIITYTSKTKENPSRRFYRCAIALQRENEDHLFKWVDEALLDEIRMVE  552
            PPGVPSKCWCG  IIT+TSKTKENP RRFYRCAIA++RENE+HLFKWVDEALLDEI+MV
Sbjct  19   PPGVPSKCWCGEEIITFTSKTKENPYRRFYRCAIAMKRENEEHLFKWVDEALLDEIKMVN  78

Query  553  ENCKRVEEDVNDLKVQM  603
            E CKRV E+++DL++ +
Sbjct  79   EKCKRVVENISDLRMNV  95


>unnamed protein product [Arabidopsis thaliana]
Sequence ID: CAA0402823.1 Length: 158
Range 1: 17 to 84

Score:114 bits(286), Expect:1e-29,
Method:Compositional matrix adjust.,
Identities:50/68(74%), Positives:61/68(89%), Gaps:0/68(0%)

Query  400  CGASIITYTSKTKENPSRRFYRCAIALQRENEDHLFKWVDEALLDEIRMVEENCKRVEED  579
            CG  IIT+TSKTKENP RRFYRCAIA++RENE+HLFKWVDEALLDEI+MV E CKRV E+
Sbjct  17   CGEEIITFTSKTKENPYRRFYRCAIAMKRENEEHLFKWVDEALLDEIKMVNEKCKRVAEN  76

Query  580  VNDLKVQM  603
            ++DL++ +
Sbjct  77   ISDLRMNV  84


>unnamed protein product [Arabidopsis thaliana]
Sequence ID: BAB09034.1 Length: 154
Range 1: 22 to 106

Score:79.7 bits(195), Expect:5e-17,
Method:Compositional matrix adjust.,
Identities:36/85(42%), Positives:51/85(60%), Gaps:2/85(2%)

Query  379  GVPSKCWCGASIITYTSKTKENPSRRFYRCAIALQRE--NEDHLFKWVDEALLDEIRMVE  552
            GVP  CWCGA+I    SK+  NP RR+YRCA A   +  +++H FKWVDEALLDEI  +
Sbjct  22   GVPKMCWCGATISALMSKSASNPYRRYYRCAYAATHKLVDDNHTFKWVDEALLDEIERLT  81

Query  553  ENCKRVEEDVNDLKVQMAKRHRFCF  627
                 +EE ++ +  +  +  +  F
Sbjct  82   RRTAALEETMSQITTENMEHQKMVF  106


>unknown, partial [Arabidopsis thaliana]
Sequence ID: ABK28583.1 Length: 155
Range 1: 14 to 97

Score:78.6 bits(192), Expect:1e-16,
Method:Compositional matrix adjust.,
Identities:32/84(38%), Positives:51/84(60%), Gaps:2/84(2%)

Query  355  PTHHFSPPGVPSKCWCGASIITYTSKTKENPSRRFYRCAIALQR--ENEDHLFKWVDEAL  528
            P       GVP +CWCG  ++   SK+  NP RR+YRC    +    N++H FKWVDEAL
Sbjct  14   PRGRGKVVGVPKRCWCGEGVVALISKSDMNPCRRYYRCGFTAKHRLRNDEHTFKWVDEAL  73

Query  529  LDEIRMVEENCKRVEEDVNDLKVQ  600
            L+EI  + +  + +E+++ +L+ +
Sbjct  74   LNEIETLIDKTREIEQELKELRTE  97


>hypothetical protein At3g42810 [Arabidopsis thaliana]
Sequence ID: ABE65983.1 Length: 154
>putative protein [Arabidopsis thaliana]
Sequence ID: CAB87198.1 Length: 154
Range 1: 14 to 97

Score:78.6 bits(192), Expect:1e-16,
Method:Compositional matrix adjust.,
Identities:32/84(38%), Positives:51/84(60%), Gaps:2/84(2%)

Query  355  PTHHFSPPGVPSKCWCGASIITYTSKTKENPSRRFYRCAIALQR--ENEDHLFKWVDEAL  528
            P       GVP +CWCG  ++   SK+  NP RR+YRC    +    N++H FKWVDEAL
Sbjct  14   PRGRGKVVGVPKRCWCGEGVVALISKSDMNPCRRYYRCGFTAKHRLRNDEHTFKWVDEAL  73

Query  529  LDEIRMVEENCKRVEEDVNDLKVQ  600
            L+EI  + +  + +E+++ +L+ +
Sbjct  74   LNEIETLIDKTREIEQELKELRTE  97


>hypothetical protein [Arabidopsis thaliana]
Sequence ID: CAA16722.1 Length: 154
>hypothetical protein [Arabidopsis thaliana]
Sequence ID: CAB78844.1 Length: 154
Range 1: 14 to 97

Score:76.6 bits(187), Expect:6e-16,
Method:Compositional matrix adjust.,
Identities:34/84(40%), Positives:51/84(60%), Gaps:2/84(2%)

Query  355  PTHHFSPPGVPSKCWCGASIITYTSKTKENPSRRFYRCAI-ALQR-ENEDHLFKWVDEAL  528
            P       GVP +CWCG +++   SK+  NP RR+YRC + AL R  N++H FKWVD+AL
Sbjct  14   PRGRGRVVGVPKRCWCGEAVVVLISKSDSNPCRRYYRCGLAALNRLRNDEHTFKWVDDAL  73

Query  529  LDEIRMVEENCKRVEEDVNDLKVQ  600
            L+EI  +       E+ + +L ++
Sbjct  74   LNEIETLAAKNGEFEQKLKELTIE  97


>putative protein [Arabidopsis thaliana]
Sequence ID: CAB68121.1 Length: 134
Range 1: 14 to 97

Score:68.2 bits(165), Expect:3e-13,
Method:Compositional matrix adjust.,
Identities:29/84(35%), Positives:49/84(58%), Gaps:2/84(2%)

Query  355  PTHHFSPPGVPSKCWCGASIITYTSKTKENPSRRFYRCAIALQR--ENEDHLFKWVDEAL  528
            P       GVP +CWC   ++   SK+  NP +R+YRC  A +    N++H FKWVD+AL
Sbjct  14   PQGRGKVVGVPKRCWCREGVMALISKSDLNPCQRYYRCGFAAKHRLRNDEHTFKWVDKAL  73

Query  529  LDEIRMVEENCKRVEEDVNDLKVQ  600
            L+EI  + +  +  E+++ + + +
Sbjct  74   LNEIETLTDKTREFEQELKEHRTE  97


>zinc ion-binding protein [Arabidopsis thaliana]
Sequence ID: NP_001327268.1 Length: 91
>zinc ion-binding protein [Arabidopsis thaliana]
Sequence ID: ANM65290.1 Length: 91
Range 1: 17 to 89

Score:63.5 bits(153), Expect:5e-12,
Method:Compositional matrix adjust.,
Identities:28/77(36%), Positives:47/77(61%), Gaps:4/77(5%)

Query  379  GVPSKCWCGASIITYTSKTKENPSRRFYRCAIALQRENEDHLFKWVDEALLDEIRMVEEN  558
            G PSKC CG  ++ YTS +K+NP R ++RC        +DHLFKWV++ + +E+
Sbjct  17   GFPSKCRCGRDVVIYTSSSKKNPGRPYFRCPTY----QDDHLFKWVEDCVYEEVVDAIPR  72

Query  559  CKRVEEDVNDLKVQMAK  609
               ++ +VN+ K ++A+
Sbjct  73   ISIIDSEVNNAKSEVAR  89


>unnamed protein product [Arabidopsis thaliana]
Sequence ID: CAA0384158.1 Length: 150
Range 1: 17 to 88

Score:62.0 bits(149), Expect:7e-11,
Method:Compositional matrix adjust.,
Identities:28/76(37%), Positives:46/76(60%), Gaps:4/76(5%)

Query  379  GVPSKCWCGASIITYTSKTKENPSRRFYRCAIALQRENEDHLFKWVDEALLDEIRMVEEN  558
            G PSKC CG  ++ YTS +K+NP R ++RC        +DHLFKWV++ + +E+
Sbjct  17   GFPSKCRCGRDVVIYTSSSKKNPGRPYFRCPTY----QDDHLFKWVEDCVYEEVVDAIPR  72

Query  559  CKRVEEDVNDLKVQMA  606
               ++ +VN+ K ++A
Sbjct  73   ISIIDSEVNNAKSEVA  88


>Hypothetical protein T10I18.4 [Arabidopsis thaliana]
Sequence ID: AAF88011.1 Length: 156
Range 1: 21 to 92

Score:60.8 bits(146), Expect:2e-10,
Method:Compositional matrix adjust.,
Identities:26/76(34%), Positives:45/76(59%), Gaps:4/76(5%)

Query  379  GVPSKCWCGASIITYTSKTKENPSRRFYRCAIALQRENEDHLFKWVDEALLDEIRMVEEN  558
            G PSKC CG  ++ YTS +K NP R F+RC      + +DHLFKWV++ + +++
Sbjct  21   GFPSKCECGLDVVIYTSASKTNPGRPFFRCP----TKQDDHLFKWVEDGVYEKVADALPK  76

Query  559  CKRVEEDVNDLKVQMA  606
               ++ ++N+ K ++
Sbjct  77   FSIIDSEINNAKSEVT  92


>hypothetical protein AXX17_AT3G33460 [Arabidopsis thaliana]
Sequence ID: OAP03513.1 Length: 604
Range 1: 451 to 521

Score:63.2 bits(152), Expect:6e-10,
Method:Compositional matrix adjust.,
Identities:29/72(40%), Positives:47/72(65%), Gaps:1/72(1%)

Query  379  GVPSKCWCGASIITYTSKTKENPSRRFYRCAIALQRENEDHLFKWVDEALLDEIRMVEEN  558
            G+P KC CGA  +  TS+T +NP R F+ C    + +N  HLFKW D A+++EI+ VE +
Sbjct  451  GLPIKCRCGAGSVVKTSETMKNPGRLFHCCPYGSKVDN-GHLFKWTDIAMVEEIKEVECD  509

Query  559  CKRVEEDVNDLK  594
             ++++ ++  LK
Sbjct  510  VEKIQAEIGSLK  521


>unknown, partial [Arabidopsis thaliana]
Sequence ID: ABK28780.1 Length: 166
Range 1: 28 to 103

Score:58.9 bits(141), Expect:1e-09,
Method:Compositional matrix adjust.,
Identities:26/80(33%), Positives:45/80(56%), Gaps:4/80(5%)

Query  367  FSPPGVPSKCWCGASIITYTSKTKENPSRRFYRCAIALQRENEDHLFKWVDEALLDEIRM  546
            F   G P KC CG  ++ +TS T +NP R F+RC    +   +DHLFKWV++ + +E+
Sbjct  28   FKSRGFPVKCKCGLDVVMFTSSTAKNPGRPFFRC----KSCEDDHLFKWVEDCMYEEVID  83

Query  547  VEENCKRVEEDVNDLKVQMA  606
                   ++ ++ + K ++A
Sbjct  84   ALPKISSIDNEIINAKAEVA  103


>unknown protein [Arabidopsis thaliana]
Sequence ID: ABF59287.1 Length: 165
Range 1: 28 to 103

Score:58.9 bits(141), Expect:1e-09,
Method:Compositional matrix adjust.,
Identities:26/80(33%), Positives:45/80(56%), Gaps:4/80(5%)

Query  367  FSPPGVPSKCWCGASIITYTSKTKENPSRRFYRCAIALQRENEDHLFKWVDEALLDEIRM  546
            F   G P KC CG  ++ +TS T +NP R F+RC    +   +DHLFKWV++ + +E+
Sbjct  28   FKSRGFPVKCKCGLDVVMFTSSTAKNPGRPFFRC----KSCEDDHLFKWVEDCMYEEVID  83

Query  547  VEENCKRVEEDVNDLKVQMA  606
                   ++ ++ + K ++A
Sbjct  84   ALPKISSIDNEIINAKAEVA  103


>GRF zinc finger protein [Arabidopsis thaliana]
Sequence ID: NP_001324832.1 Length: 130
>GRF zinc finger protein [Arabidopsis thaliana]
Sequence ID: ANM62691.1 Length: 130
Range 1: 15 to 76

Score:57.8 bits(138), Expect:1e-09,
Method:Compositional matrix adjust.,
Identities:25/62(40%), Positives:37/62(59%), Gaps:5/62(8%)

Query  379  GVPSKCWCGASIITYTSKTKENPSRRFYRCAI-----ALQRENEDHLFKWVDEALLDEIR  543
            G P KCWCG  +  +TSK+ +NP R F+RC           + + HLFKWV++A+ +E+
Sbjct  15   GFPVKCWCGDDVTIFTSKSVDNPGRPFFRCETKRDPKTWTNKKDSHLFKWVEDAVYEEVE  74

Query  544  MV  549
             V
Sbjct  75   DV  76


>T12C24.25 [Arabidopsis thaliana]
Sequence ID: AAF88091.1 Length: 156
Range 1: 21 to 92

Score:58.2 bits(139), Expect:2e-09,
Method:Compositional matrix adjust.,
Identities:26/76(34%), Positives:43/76(56%), Gaps:4/76(5%)

Query  379  GVPSKCWCGASIITYTSKTKENPSRRFYRCAIALQRENEDHLFKWVDEALLDEIRMVEEN  558
            G+ SKC CG  ++ YTS +K NP R F+RC      + +DHLFKWV+  + +E+
Sbjct  21   GLASKCHCGLEVVIYTSASKSNPGRPFFRCPT----KQDDHLFKWVEYGVYEEVVEALPK  76

Query  559  CKRVEEDVNDLKVQMA  606
               ++ ++   K ++A
Sbjct  77   ISSIDSEIMKAKCEVA  92


>GRF zinc finger protein [Arabidopsis thaliana]
Sequence ID: NP_001320823.1 Length: 152
>GRF zinc finger protein [Arabidopsis thaliana]
Sequence ID: ANM58381.1 Length: 152
Range 1: 15 to 101

Score:57.8 bits(138), Expect:2e-09,
Method:Compositional matrix adjust.,
Identities:27/87(31%), Positives:46/87(52%), Gaps:12/87(13%)

Query  379  GVPSKCWCGASIITYTSKTKENPSRRFYRCAI-----ALQRENEDHLFKWVDEALLDEIR  543
            G P KCWCG  +  +TSK+ +NP R F+RC           + + HLFKWV++A+ +E+
Sbjct  15   GFPVKCWCGDDVTIFTSKSVDNPGRPFFRCETKRDPKTWTNKRDSHLFKWVEDAVYEEVE  74

Query  544  -------MVEENCKRVEEDVNDLKVQM  603
                   ++     + + + N+L V +
Sbjct  75   DVLPKFVIIANELNKAKSEANELNVMI  101


>hypothetical protein [Arabidopsis thaliana]
Sequence ID: ABF59281.1 Length: 156
Range 1: 21 to 65

Score:56.2 bits(134), Expect:7e-09,
Method:Compositional matrix adjust.,
Identities:23/49(47%), Positives:33/49(67%), Gaps:4/49(8%)

Query  379  GVPSKCWCGASIITYTSKTKENPSRRFYRCAIALQRENEDHLFKWVDEA  525
            G+PSKC CG  ++ YTS + +NP R ++RC   L    +DHLFKWV++
Sbjct  21   GLPSKCHCGLDVVIYTSSSAKNPGRPYFRCPTRL----DDHLFKWVEDG  65


>unknown, partial [Arabidopsis thaliana]
Sequence ID: ABK28512.1 Length: 157
Range 1: 21 to 65

Score:56.2 bits(134), Expect:7e-09,
Method:Compositional matrix adjust.,
Identities:23/49(47%), Positives:33/49(67%), Gaps:4/49(8%)

Query  379  GVPSKCWCGASIITYTSKTKENPSRRFYRCAIALQRENEDHLFKWVDEA  525
            G+PSKC CG  ++ YTS + +NP R ++RC   L    +DHLFKWV++
Sbjct  21   GLPSKCHCGLDVVIYTSSSAKNPGRPYFRCPTRL----DDHLFKWVEDG  65


>unnamed protein product [Arabidopsis thaliana]
Sequence ID: CAA0393920.1 Length: 143
Range 1: 7 to 59

Score:55.5 bits(132), Expect:1e-08,
Method:Compositional matrix adjust.,
Identities:22/54(41%), Positives:34/54(62%), Gaps:1/54(1%)

Query  379  GVPSKCWCGASIITYTSKTKENPSRRFYRCAIALQRENEDHLFKWVDEALLDEI  540
            G+PS+C CG  ++  TSK  +NP R FY C   ++     HLF+W DE +++E+
Sbjct  7    GIPSRCRCGEDVVLRTSKNVKNPGRLFYACRCGVE-NVRGHLFRWTDETMVEEM  59


>Mutator-like transposase [Arabidopsis thaliana]
Sequence ID: AAD15524.1 Length: 534
Range 1: 247 to 307

Score:58.5 bits(140), Expect:2e-08,
Method:Compositional matrix adjust.,
Identities:33/76(43%), Positives:41/76(53%), Gaps:15/76(19%)

Query  382  VPSKCWCGASIITYTSKTKENPSRRFYRCAIALQRENEDHLFKWVDEALLDEIRMVEENC  561
            VP  CWCGASI    SK+  NP RR+YRCA A   +        VDEA++DEI
Sbjct  247  VPKLCWCGASISALISKSATNPYRRYYRCAYAAGHK--------VDEAMIDEI-------  291

Query  562  KRVEEDVNDLKVQMAK  609
            KR+    N LK  M++
Sbjct  292  KRLARRTNALKQTMSQ  307


>unnamed protein product [Arabidopsis thaliana]
Sequence ID: VYS52155.1 Length: 134
Range 1: 7 to 87

Score:54.3 bits(129), Expect:2e-08,
Method:Compositional matrix adjust.,
Identities:26/82(32%), Positives:46/82(56%), Gaps:8/82(9%)

Query  379  GVPSKCWCGASIITYTSKTKENPSRRFYRCAIALQRENEDHLFKWVDEALLDE-------  537
            G+P +C CG  ++  TSKT +NP R F+ C   ++     HLF+W DE +++E
Sbjct  7    GIPIRCCCGEDVVLRTSKTVKNPGRLFFACRCGVENVR-GHLFRWTDETMVEEMEDILPK  65

Query  538  IRMVEENCKRVEEDVNDLKVQM  603
            I  +E +   ++++V  L+ +M
Sbjct  66   IEKIEGSSLTLQKEVQSLECEM  87


>hypothetical protein [Arabidopsis thaliana]
Sequence ID: AAC69115.1 Length: 159
Range 1: 21 to 79

Score:55.1 bits(131), Expect:2e-08,
Method:Compositional matrix adjust.,
Identities:24/59(41%), Positives:36/59(61%), Gaps:5/59(8%)

Query  379  GVPSKCWCGASIITYTSKTKENPSRRFYRCAI-----ALQRENEDHLFKWVDEALLDEI  540
            GVPSKC CG+ +  + SKT+EN  R F+RC           + + HLFKWV++ + +E+
Sbjct  21   GVPSKCLCGSEVTIFVSKTQENQGRPFFRCVRNRDVNTWTTKRDGHLFKWVEDPVFEEV  79


>zinc ion binding protein [Arabidopsis thaliana]
Sequence ID: NP_178758.1 Length: 143
>hypothetical protein [Arabidopsis thaliana]
Sequence ID: AAM15270.1 Length: 143 >hypothetical protein [Arabidopsis thaliana]
Sequence ID: AAM15486.1 Length: 143 >zinc ion binding protein [Arabidopsis thaliana]
Sequence ID: AEC06054.1 Length: 143 >unnamed protein product [Arabidopsis thaliana]
Sequence ID: VYS52200.1 Length: 143
Range 1: 7 to 59

Score:53.9 bits(128), Expect:4e-08,
Method:Compositional matrix adjust.,
Identities:22/54(41%), Positives:33/54(61%), Gaps:1/54(1%)

Query  379  GVPSKCWCGASIITYTSKTKENPSRRFYRCAIALQRENEDHLFKWVDEALLDEI  540
            G+PS+C CG  ++  TSKT +NP R FY C    +     HLF+  DE +++E+
Sbjct  7    GIPSRCRCGEDVVLRTSKTVKNPRRLFYACRCG-EENGSGHLFRLTDEVMVEEM  59


>zinc ion-binding protein [Arabidopsis thaliana]
Sequence ID: NP_001325185.1 Length: 134
>zinc ion-binding protein [Arabidopsis thaliana]
Sequence ID: ANM63073.1 Length: 134
Range 1: 7 to 59

Score:53.5 bits(127), Expect:4e-08,
Method:Compositional matrix adjust.,
Identities:21/54(39%), Positives:34/54(62%), Gaps:1/54(1%)

Query  379  GVPSKCWCGASIITYTSKTKENPSRRFYRCAIALQRENEDHLFKWVDEALLDEI  540
            G+P +C CG  ++  TSKT +NP R F+ C   ++     HLF+W DE +++E+
Sbjct  7    GIPIRCCCGEDVVLRTSKTVKNPGRLFFACRCGVENVR-VHLFRWTDETMVEEM  59


>unnamed protein product [Arabidopsis thaliana]
Sequence ID: BAB01424.1 Length: 156
Range 1: 21 to 65

Score:53.9 bits(128), Expect:5e-08,
Method:Compositional matrix adjust.,
Identities:22/49(45%), Positives:31/49(63%), Gaps:4/49(8%)

Query  379  GVPSKCWCGASIITYTSKTKENPSRRFYRCAIALQRENEDHLFKWVDEA  525
            G+PSKC CG  ++ YTS   +NP R ++RC        +DHLFKWV++
Sbjct  21   GLPSKCHCGLDVVIYTSTYAKNPGRPYFRCPT----RKDDHLFKWVEDG  65


Query #99: XLOC_009375 Query ID: lcl|Query_76633 Length: 958

No significant similarity found.


Query #100: XLOC_009478 Query ID: lcl|Query_76634 Length: 613

No significant similarity found.


Query #101: XLOC_009607 Query ID: lcl|Query_33104 Length: 993

Sequences producing significant alignments:
                                                                  Max    Total Query   E   Per.
Description                                                       Score  Score cover Value Ident  Accession
hypothetical protein AXX17_AT5G30130 [Arabidopsis thaliana]       116    116   30%   7e-28 63.46  OAO92217.1
hypothetical protein AXX17_AT4G06230 [Arabidopsis thaliana]       60.5   108   30%   4e-17 53.97  OAP00463.1
sequence-specific DNA binding transcription factor [Arabidopsi... 70.9   70.9  16%   4e-13 58.93  NP_180011.1
unnamed protein product [Arabidopsis thaliana]                    70.9   70.9  16%   4e-13 58.93  VYS53356.1
unnamed protein product [Arabidopsis thaliana]                    69.7   69.7  17%   8e-13 55.93  CAA0370777.1


Alignments:
>hypothetical protein AXX17_AT5G30130 [Arabidopsis thaliana]
Sequence ID: OAO92217.1 Length: 2863
Range 1: 2760 to 2863

Score:116 bits(290), Expect:7e-28,
Method:Compositional matrix adjust.,
Identities:66/104(63%), Positives:75/104(72%), Gaps:2/104(1%)

Query  648   TLELQETCNISSTSKSDNEIQDQDQDIKDVEELEDIDQATQEGL*KYLYFKEPTSKRAQD  469
             TLELQET N S TS+SD E QDQ+QDIKD+EELEDIDQATQE L K  YFKEP SKR+QD
Sbjct  2760  TLELQETNNSSVTSQSDRESQDQEQDIKDMEELEDIDQATQEELQKDPYFKEPASKRSQD  2819

Query  468   VISEVSQRNLQEVTQKTSTRPKSHPCKVYS--ILLCRSKHPTNG  343
             +I EV +  LQEV+Q T   PK+H  KVYS  I+      P NG
Sbjct  2820  LIPEVPEEVLQEVSQATCIGPKTHLGKVYSVYIIFAGPHSPDNG  2863


>hypothetical protein AXX17_AT4G06230 [Arabidopsis thaliana]
Sequence ID: OAP00463.1 Length: 825
Range 1: 28 to 82

Score:60.5 bits(145), Expect:4e-17,
Method:Compositional matrix adjust.,
Identities:34/63(54%), Positives:41/63(65%), Gaps:8/63(12%)

Query  875  LQWILVSLTG*SFGIYRVTFPIAEEELAEAFSQISLERTTPLSMKSHLTGLEASRSQEKT  696
            + W +   TG     Y +   IAEEEL EAF+Q+SLERTTP SMK  LTG   SR+QEKT
Sbjct  28   VNWFVAPHTG-----YDL---IAEEELVEAFTQMSLERTTPPSMKYRLTGSRTSRNQEKT  79

Query  695  LDF  687
            L+
Sbjct  80   LEL  82


Range 2: 79 to 116

Score:47.8 bits(112), Expect:4e-17,
Method:Compositional matrix adjust.,
Identities:26/38(68%), Positives:29/38(76%), Gaps:0/38(0%)

Query  648  TLELQETCNISSTSKSDNEIQDQDQDIKDVEELEDIDQ  535
            TLELQET + S TS SD E QDQ+QDIKD+EE EDI
Sbjct  79   TLELQETNDSSVTSTSDRESQDQEQDIKDMEEFEDIPH  116


>sequence-specific DNA binding transcription factor [Arabidopsis thaliana]
Sequence ID: NP_180011.1 Length: 322
>hypothetical protein [Arabidopsis thaliana]
Sequence ID: AAD18107.1 Length: 322 >sequence-specific DNA binding transcription factor [Arabidopsis thaliana]
Sequence ID: AEC07564.1 Length: 322
Range 1: 178 to 233

Score:70.9 bits(172), Expect:4e-13,
Method:Compositional matrix adjust.,
Identities:33/56(59%), Positives:39/56(69%), Gaps:0/56(0%)

Query  984  GACNVNESGLMTRSTTSGLKRKAGKDIRPLCRLYKSASVDTSFIDWIKLWNLPSHV  817
            GA N NESGL   +TTSG+K K  +DI PLCR YK A++D+SFI WI  WNL   V
Sbjct  178  GASNANESGLKIGATTSGVKTKVDEDIEPLCRHYKCATMDSSFIHWITHWNLLDQV  233


>unnamed protein product [Arabidopsis thaliana]
Sequence ID: VYS53356.1 Length: 322
Range 1: 178 to 233

Score:70.9 bits(172), Expect:4e-13,
Method:Compositional matrix adjust.,
Identities:33/56(59%), Positives:39/56(69%), Gaps:0/56(0%)

Query  984  GACNVNESGLMTRSTTSGLKRKAGKDIRPLCRLYKSASVDTSFIDWIKLWNLPSHV  817
            GA N NESGL   +TTSG+K K  +DI PLCR YK A++D+SFI WI  WNL   V
Sbjct  178  GASNANESGLKIGATTSGVKTKVDEDIEPLCRHYKCATMDSSFIHWITHWNLLDQV  233


>unnamed protein product [Arabidopsis thaliana]
Sequence ID: CAA0370777.1 Length: 288
Range 1: 147 to 205

Score:69.7 bits(169), Expect:8e-13,
Method:Compositional matrix adjust.,
Identities:33/59(56%), Positives:39/59(66%), Gaps:0/59(0%)

Query  993  NSEGACNVNESGLMTRSTTSGLKRKAGKDIRPLCRLYKSASVDTSFIDWIKLWNLPSHV  817
            N  GA N NESGL   +TTSG+K K  +DI PLCR YK A++D+SFI WI  WN    V
Sbjct  147  NLGGASNANESGLKIGATTSGVKTKVDEDIEPLCRHYKCATMDSSFIHWITHWNFLDQV  205


Query #102: XLOC_009909 Query ID: lcl|Query_33105 Length: 795

Sequences producing significant alignments:
                                                                  Max    Total Query   E   Per.
Description                                                       Score  Score cover Value Ident  Accession
transmembrane protein [Arabidopsis thaliana]                      67.4   67.4  33%   5e-14 42.22  NP_001326023.1
hypothetical protein AT2G13845 [Arabidopsis thaliana]             43.1   86.3  21%   1e-10 50.00  NP_001324079.1
unnamed protein product [Arabidopsis thaliana]                    55.8   55.8  26%   1e-09 50.68  VYS52357.1
hypothetical protein AT2G14265 [Arabidopsis thaliana]             55.8   55.8  16%   2e-09 60.38  NP_001325174.1
unnamed protein product [Arabidopsis thaliana]                    52.8   52.8  33%   1e-08 42.22  CAA0384287.1


Alignments:
>transmembrane protein [Arabidopsis thaliana]
Sequence ID: NP_001326023.1 Length: 74
>transmembrane protein [Arabidopsis thaliana]
Sequence ID: ANM63965.1 Length: 74
Range 1: 4 to 74

Score:67.4 bits(163), Expect:5e-14,
Method:Compositional matrix adjust.,
Identities:38/90(42%), Positives:47/90(52%), Gaps:19/90(21%)

Query  340  HDIQNGYRTVKRDLYLQNLMDMELWFLYKTRLIESPVIIDMFEETSTSKTLWTWNFGFFT  519
            H ++    +V RDL+LQ+L DM+LW LYK RLI+SPVI+DM
Sbjct  4    HRLEQMGGSVCRDLHLQDLTDMKLWSLYKPRLIKSPVIMDMRPPPPRPCGQL--------  55

Query  520  NHG*SNLLLYGHETPPPIPYGHRTFTKFLR  609
                          PPP PYG+RTFTKFLR
Sbjct  56   -----------QRAPPPRPYGYRTFTKFLR  74


>hypothetical protein AT2G13845, partial [Arabidopsis thaliana]
Sequence ID: NP_001324079.1 Length: 143
>hypothetical protein AT2G13845, partial [Arabidopsis thaliana]
Sequence ID: ANM61889.1 Length: 143
Range 1: 74 to 111

Score:43.1 bits(100), Expect:1e-10,
Method:Compositional matrix adjust.,
Identities:19/38(50%), Positives:26/38(68%), Gaps:0/38(0%)

Query  317  IEFPVIMDMIYKMDIEQ*KETSTFKTLWTWNFGFFTKH  430
            I   +   ++++++     ETSTFKTLWTWNFGFFT H
Sbjct  74   IRHQLHHGLLHRLEQMGGSETSTFKTLWTWNFGFFTNH  111


Range 2: 111 to 130

Score:43.1 bits(100), Expect:1e-10,
Method:Compositional matrix adjust.,
Identities:18/20(90%), Positives:18/20(90%), Gaps:0/20(0%)

Query  466  EETSTSKTLWTWNFGFFTNH  525
             ETSTSK LWTWNFGFFTNH
Sbjct  111  HETSTSKALWTWNFGFFTNH  130


>unnamed protein product [Arabidopsis thaliana]
Sequence ID: VYS52357.1 Length: 87
Range 1: 25 to 87

Score:55.8 bits(133), Expect:1e-09,
Method:Compositional matrix adjust.,
Identities:37/73(51%), Positives:39/73(53%), Gaps:13/73(17%)

Query  375  RPLPSKPYGHGTLVSLQNTVNRISCYY*HV*RDLHFQDLMDMELWFLYKPRLIESPVI--  548
            RPLPSKPYGHGTLVSLQNT    S            + L      F  KPRLIES VI
Sbjct  25   RPLPSKPYGHGTLVSLQNTFEETSTS----------KILWTWNFGFFTKPRLIESSVIMD  74

Query  549  -WT*DPTSNTLWT  584
             +    TS TLWT
Sbjct  75   LFVETSTSKTLWT  87


>hypothetical protein AT2G14265 [Arabidopsis thaliana]
Sequence ID: NP_001325174.1 Length: 111
>hypothetical protein AT2G14265 [Arabidopsis thaliana]
Sequence ID: ANM63061.1 Length: 111
Range 1: 55 to 107

Score:55.8 bits(133), Expect:2e-09,
Method:Compositional matrix adjust.,
Identities:32/53(60%), Positives:35/53(66%), Gaps:9/53(16%)

Query  375  RPLPSKPYGHGTLVSLQNTVNRISCYY---------*HV*RDLHFQDLMDMEL  506
            RPLPSKPYGHGTLVSLQNT   +   Y          HV RDLH QDLMD++L
Sbjct  55   RPLPSKPYGHGTLVSLQNTRPPLPRSYGHGTLVSLQKHVCRDLHLQDLMDIKL  107


>unnamed protein product [Arabidopsis thaliana]
Sequence ID: CAA0384287.1 Length: 74
>unnamed protein product [Arabidopsis thaliana]
Sequence ID: VYS59150.1 Length: 74
Range 1: 4 to 74

Score:52.8 bits(125), Expect:1e-08,
Method:Compositional matrix adjust.,
Identities:38/90(42%), Positives:47/90(52%), Gaps:19/90(21%)

Query  340  HDIQNGYRTVKRDLYLQNLMDMELWFLYKTRLIESPVIIDMFEETSTSKTLWTWNFGFFT  519
            H ++    +V RDL+LQ+L DM+LW LYK RLI+SPVI+DM
Sbjct  4    HRLEQMGGSVCRDLHLQDLTDMKLWSLYKPRLIKSPVIMDMRPPPPRPCGQL--------  55

Query  520  NHG*SNLLLYGHETPPPIPYGHRTFTKFLR  609
                          PPP PYG+RTFTKFLR
Sbjct  56   -----------QRPPPPRPYGYRTFTKFLR  74


Query #103: XLOC_010118 Query ID: lcl|Query_33106 Length: 527

Sequences producing significant alignments:
                                                                  Max    Total Query   E   Per.
Description                                                       Score  Score cover Value Ident  Accession
unnamed protein product [Arabidopsis thaliana]                    55.1   55.1  20%   1e-08 72.22  CAA0397958.1
Galactose oxidase/kelch repeat superfamily protein [Arabidopsi... 55.1   55.1  20%   1e-08 72.22  NP_195605.1


Alignments:
>unnamed protein product [Arabidopsis thaliana]
Sequence ID: CAA0397958.1 Length: 370
Range 1: 333 to 368

Score:55.1 bits(131), Expect:1e-08,
Method:Compositional matrix adjust.,
Identities:26/36(72%), Positives:30/36(83%), Gaps:0/36(0%)

Query  527  AEIALERREGGEIWGKVEWYDLVLDGTSCIMKCLAV  420
            AEIA+ERREGGEIWGKVEW ++VL G   IM C+AV
Sbjct  333  AEIAVERREGGEIWGKVEWCNVVLSGNFHIMDCVAV  368


>Galactose oxidase/kelch repeat superfamily protein [Arabidopsis thaliana]
Sequence ID: NP_195605.1 Length: 370
>RecName: Full=F-box/kelch-repeat protein At4g38940 [Arabidopsis thaliana]
Sequence ID: Q9SVJ9.1 Length: 370 >unknown [Arabidopsis thaliana]
Sequence ID: AAM65521.1 Length: 370 >Galactose oxidase/kelch repeat superfamily protein [Arabidopsis thaliana]
Sequence ID: AEE86996.1 Length: 370 >hypothetical protein AXX17_AT4G44290 [Arabidopsis thaliana]
Sequence ID: OAO99011.1 Length: 370 >putative protein [Arabidopsis thaliana]
Sequence ID: CAB38814.1 Length: 370 >putative protein [Arabidopsis thaliana]
Sequence ID: CAB80557.1 Length: 370
Range 1: 333 to 368

Score:55.1 bits(131), Expect:1e-08,
Method:Compositional matrix adjust.,
Identities:26/36(72%), Positives:30/36(83%), Gaps:0/36(0%)

Query  527  AEIALERREGGEIWGKVEWYDLVLDGTSCIMKCLAV  420
            AEIA+ERREGGEIWGKVEW ++VL G   IM C+AV
Sbjct  333  AEIAVERREGGEIWGKVEWCNVVLSGNFHIMDCVAV  368


Query #104: XLOC_011101 Query ID: lcl|Query_33107 Length: 280

No significant similarity found.


Query #105: XLOC_011423 Query ID: lcl|Query_33108 Length: 478

No significant similarity found.


Query #106: XLOC_011651 Query ID: lcl|Query_33109 Length: 1967

No significant similarity found.


Query #107: XLOC_011721 Query ID: lcl|Query_33110 Length: 1343

Sequences producing significant alignments:
                                                                  Max    Total Query   E   Per.
Description                                                       Score  Score cover Value Ident  Accession
hypothetical protein AXX17_AT4G01860 [Arabidopsis thaliana]       119    226   48%   7e-46 76.62  OAO99509.1
F11O4.11 [Arabidopsis thaliana]                                   118    207   32%   2e-40 76.62  AAC62785.1
putative non-LTR retroelement reverse transcriptase [Arabidops... 116    200   32%   4e-38 70.13  AAD22286.1
hypothetical protein AXX17_AT4G21170 [Arabidopsis thaliana]       122    122   20%   8e-29 64.89  OAO96867.1
putative retroelement pol polyprotein [Arabidopsis thaliana]      111    111   17%   3e-27 70.13  AAM15063.1
hypothetical protein AXX17_AT3G15320 [Arabidopsis thaliana]       82.0   110   22%   1e-17 52.05  OAP05087.1
hypothetical protein AXX17_AT3G15320 [Arabidopsis thaliana]       81.6   110   22%   2e-17 52.05  OAP05088.1
unnamed protein product [Arabidopsis thaliana]                    81.6   81.6  16%   6e-16 50.00  BAB08713.1
hypothetical protein AXX17_AT5G32690 [Arabidopsis thaliana]       80.9   80.9  16%   2e-15 50.00  OAO94166.1
hypothetical protein At4g09700 [Arabidopsis thaliana]             75.1   102   21%   5e-15 49.32  ABE65517.1
unknown [Arabidopsis thaliana]                                    74.7   101   21%   5e-15 49.32  ABK28247.1
putative Ta11-like non-LTR retroelement protein [Arabidopsis...   66.6   66.6  24%   6e-11 34.86  AAD17399.1
unknown [Arabidopsis thaliana]                                    58.2   58.2  16%   1e-09 38.36  ABK28059.1
hypothetical protein [Arabidopsis thaliana]                       57.8   57.8  16%   1e-09 38.36  ABF59395.1
hypothetical protein [Arabidopsis thaliana]                       56.2   56.2  7%    1e-08 78.12  AAO37173.1
putative Ta11-like non-LTR retroelement protein [Arabidopsis...   56.6   56.6  24%   1e-08 32.46  AAD22298.1
unnamed protein product [Arabidopsis thaliana]                    58.9   58.9  16%   1e-08 42.47  CAA0360127.1


Alignments:
>hypothetical protein AXX17_AT4G01860 [Arabidopsis thaliana]
Sequence ID: OAO99509.1 Length: 560
Range 1: 96 to 172

Score:119 bits(297), Expect:7e-46,
Method:Compositional matrix adjust.,
Identities:59/77(77%), Positives:65/77(84%), Gaps:0/77(0%)

Query  656  LSNRWNFKGKAIGSDLGNGCf*fifeieENLEKVLANRPYHFDQWMVILQRWEPIISSTF  477
            LSNRWN KGKA GSDLG GCF F FE EE+L+KVL NRPYHF QWMVILQRW+P+IS +F
Sbjct  96   LSNRWNLKGKATGSDLGRGCFQFRFEYEEDLQKVLDNRPYHFGQWMVILQRWKPVISPSF  155

Query  476  PSLIPF*VELQGLPKHY  426
            PS IPF +ELQGLP HY
Sbjct  156  PSEIPFWIELQGLPMHY  172


Range 2: 228 to 372

Score:71.6 bits(174), Expect:7e-46,
Method:Compositional matrix adjust.,
Identities:53/148(36%), Positives:74/148(50%), Gaps:34/148(22%)

Query  352  LDYKNLKKHCTHCHRLSHGMDNCPGLEQEKVVSATKGNIPPARVPKP*FLYSQDskl*ki  173
            L+YKNLK HC HC RLSH   +CPGL++ + VS T     P + P+    Y++ +K  K
Sbjct  228  LEYKNLKSHCHHCQRLSHAEADCPGLKKVEAVSKTSKLTSPEQPPQ---NYAERNKGSKQ  284

Query  172  spsis*LQTVAY------------------------TDH-------STSSRARRPPLERT  86
            +P  S  + ++                         +D+       S +SR RRPPLER
Sbjct  285  NPRASHDKLLSSHSGLSRPLGDYNPYKRKSDGRNGRSDYRLRPGEMSETSRTRRPPLERE  344

Query  85   SCAGESSH*PLPIPTTEEVMDELREITV  2
                 S+  P+PIPT E +M ELRE+TV
Sbjct  345  VVTESSNPLPVPIPTKEAIMGELREVTV  372


Range 3: 175 to 197

Score:35.8 bits(81), Expect:7e-46,
Method:Compositional matrix adjust.,
Identities:13/23(57%), Positives:21/23(91%), Gaps:0/23(0%)

Query  420  QQLIYKIGEEIGEVLDHEITPSA  352
            ++++Y IG+E GEV+DHEI+P+A
Sbjct  175  KEMLYAIGKEAGEVVDHEISPAA  197


>F11O4.11 [Arabidopsis thaliana]
Sequence ID: AAC62785.1 Length: 577
>putative transposon protein [Arabidopsis thaliana]
Sequence ID: CAB77719.1 Length: 577
Range 1: 59 to 135

Score:118 bits(295), Expect:2e-40,
Method:Compositional matrix adjust.,
Identities:59/77(77%), Positives:65/77(84%), Gaps:0/77(0%)

Query  656  LSNRWNFKGKAIGSDLGNGCf*fifeieENLEKVLANRPYHFDQWMVILQRWEPIISSTF  477
            LSNRWN KGKA GSDLG GCF F FE EE+L+KVL NRPYHF QWMVILQRW+P+IS +F
Sbjct  59   LSNRWNLKGKATGSDLGRGCFQFRFEYEEDLQKVLDNRPYHFGQWMVILQRWKPVISPSF  118

Query  476  PSLIPF*VELQGLPKHY  426
            PS IPF +ELQGLP HY
Sbjct  119  PSEIPFWIELQGLPMHY  135


Range 2: 191 to 236

Score:53.9 bits(128), Expect:2e-40,
Method:Compositional matrix adjust.,
Identities:24/46(52%), Positives:31/46(67%), Gaps:0/46(0%)

Query  352  LDYKNLKKHCTHCHRLSHGMDNCPGLEQEKVVSATKGNIPPARVPK  215
            L+YKNLK HC HC RLSH   +CPGL++ + VS T    PP + P+
Sbjct  191  LEYKNLKSHCHHCQRLSHAEADCPGLKKVEAVSKTSKLTPPEQPPQ  236


Range 3: 138 to 160

Score:35.8 bits(81), Expect:2e-40,
Method:Compositional matrix adjust.,
Identities:13/23(57%), Positives:21/23(91%), Gaps:0/23(0%)

Query  420  QQLIYKIGEEIGEVLDHEITPSA  352
            ++++Y IG+E GEV+DHEI+P+A
Sbjct  138  KEMLYAIGKEAGEVVDHEISPAA  160


>putative non-LTR retroelement reverse transcriptase [Arabidopsis thaliana]
Sequence ID: AAD22286.1 Length: 1311
Range 1: 63 to 139

Score:116 bits(290), Expect:4e-38,
Method:Compositional matrix adjust.,
Identities:54/77(70%), Positives:63/77(81%), Gaps:0/77(0%)

Query  656  LSNRWNFKGKAIGSDLGNGCf*fifeieENLEKVLANRPYHFDQWMVILQRWEPIISSTF  477
            LSNRW  +GKA GSDLG G F   F+  E+L++VL NRPYHFDQWMVILQ+WEP+IS +F
Sbjct  63   LSNRWTLRGKATGSDLGQGVFQLKFDFSEDLQQVLDNRPYHFDQWMVILQKWEPVISPSF  122

Query  476  PSLIPF*VELQGLPKHY  426
            P LIPF +ELQGLPKHY
Sbjct  123  PCLIPFWIELQGLPKHY  139


Range 2: 195 to 234

Score:51.2 bits(121), Expect:4e-38,
Method:Composition-based stats.,
Identities:22/43(51%), Positives:29/43(67%), Gaps:3/43(6%)

Query  352  LDYKNLKKHCTHCHRLSHGMDNCPGLEQEKVVSATKGNIPPAR  224
            LDYKNLK HC HCHRL+H   +CPG+ ++   S T+ N  P +
Sbjct  195  LDYKNLKNHCRHCHRLTHEEKHCPGVAKK---SETQRNPSPVQ  234


Range 3: 136 to 163

Score:32.7 bits(73), Expect:4e-38,
Method:Composition-based stats.,
Identities:14/28(50%), Positives:19/28(67%), Gaps:0/28(0%)

Query  438  PKALLKQQLIYKIGEEIGEVLDHEITPS  355
            PK   K +++  IGEE+G V+D EIT S
Sbjct  136  PKHYWKPEMLKSIGEELGTVMDQEITSS  163


>hypothetical protein AXX17_AT4G21170 [Arabidopsis thaliana]
Sequence ID: OAO96867.1 Length: 1816
Range 1: 46 to 139

Score:122 bits(305), Expect:8e-29,
Method:Compositional matrix adjust.,
Identities:61/94(65%), Positives:71/94(75%), Gaps:0/94(0%)

Query  707  GKTDKSCCSKTMVSNIRLSNRWNFKGKAIGSDLGNGCf*fifeieENLEKVLANRPYHFD  528
            G+     C +       LSNRWN KGKA+GSDLG GCF F F+ EE+L  VL NRPYHFD
Sbjct  46   GRLTNPACQRLWALFPFLSNRWNLKGKALGSDLGKGCFQFKFDFEEDLLSVLKNRPYHFD  105

Query  527  QWMVILQRWEPIISSTFPSLIPF*VELQGLPKHY  426
            QWMVI+Q+WEPIIS +FPS+IPF +ELQGLPKHY
Sbjct  106  QWMVIIQKWEPIISDSFPSMIPFWIELQGLPKHY  139


>putative retroelement pol polyprotein [Arabidopsis thaliana]
Sequence ID: AAM15063.1 Length: 267
>putative retroelement pol polyprotein [Arabidopsis thaliana]
Sequence ID: AAM15139.1 Length: 267
Range 1: 54 to 130

Score:111 bits(277), Expect:3e-27,
Method:Compositional matrix adjust.,
Identities:54/77(70%), Positives:63/77(81%), Gaps:0/77(0%)

Query  656  LSNRWNFKGKAIGSDLGNGCf*fifeieENLEKVLANRPYHFDQWMVILQRWEPIISSTF  477
            LSN W  KGKAIGSDLG G F   F+  E+L++V+ NRPYHFDQWMVILQ+WEPIIS +F
Sbjct  54   LSNPWTLKGKAIGSDLGQGVFQLKFDFREDLQQVVDNRPYHFDQWMVILQKWEPIISPSF  113

Query  476  PSLIPF*VELQGLPKHY  426
            PSLI F +ELQGLPKH+
Sbjct  114  PSLISFWIELQGLPKHF  130


>hypothetical protein AXX17_AT3G15320 [Arabidopsis thaliana]
Sequence ID: OAP05087.1 Length: 1196
Range 1: 68 to 140

Score:82.0 bits(201), Expect:1e-17,
Method:Compositional matrix adjust.,
Identities:38/73(52%), Positives:52/73(71%), Gaps:0/73(0%)

Query  644  WNFKGKAIGSDLGNGCf*fifeieENLEKVLANRPYHFDQWMVILQRWEPIISSTFPSLI  465
            WN + + +G DLG+ CF F FE E +L+ VL   PYHF  WM+ILQ+WEP+IS TFP+ I
Sbjct  68   WNLESRVVGRDLGSECFQFRFENEADLQAVLRRGPYHFKNWMLILQQWEPVISDTFPAFI  127

Query  464  PF*VELQGLPKHY  426
             F V++ G+P H+
Sbjct  128  TFWVKIHGIPLHF  140


Range 2: 39 to 76

Score:28.5 bits(62), Expect:1e-17,
Method:Compositional matrix adjust.,
Identities:18/40(45%), Positives:22/40(55%), Gaps:12/40(30%)

Query  730  ASRTLTLMGRLTNPAAQR----------LWSLISGSRTVG  641
            A   LTL+GR+TNPA Q+          LW+L   SR VG
Sbjct  39   AENKLTLIGRVTNPAVQKPKAVAGYMPQLWNL--ESRVVG  76


>hypothetical protein AXX17_AT3G15320 [Arabidopsis thaliana]
Sequence ID: OAP05088.1 Length: 1697
Range 1: 68 to 140

Score:81.6 bits(200), Expect:2e-17,
Method:Compositional matrix adjust.,
Identities:38/73(52%), Positives:52/73(71%), Gaps:0/73(0%)

Query  644  WNFKGKAIGSDLGNGCf*fifeieENLEKVLANRPYHFDQWMVILQRWEPIISSTFPSLI  465
            WN + + +G DLG+ CF F FE E +L+ VL   PYHF  WM+ILQ+WEP+IS TFP+ I
Sbjct  68   WNLESRVVGRDLGSECFQFRFENEADLQAVLRRGPYHFKNWMLILQQWEPVISDTFPAFI  127

Query  464  PF*VELQGLPKHY  426
             F V++ G+P H+
Sbjct  128  TFWVKIHGIPLHF  140


Range 2: 39 to 76

Score:28.5 bits(62), Expect:2e-17,
Method:Compositional matrix adjust.,
Identities:18/40(45%), Positives:22/40(55%), Gaps:12/40(30%)

Query  730  ASRTLTLMGRLTNPAAQR----------LWSLISGSRTVG  641
            A   LTL+GR+TNPA Q+          LW+L   SR VG
Sbjct  39   AENKLTLIGRVTNPAVQKPKAVAGYMPQLWNL--ESRVVG  76


>unnamed protein product [Arabidopsis thaliana]
Sequence ID: BAB08713.1 Length: 439
Range 1: 63 to 138

Score:81.6 bits(200), Expect:6e-16,
Method:Compositional matrix adjust.,
Identities:38/76(50%), Positives:50/76(65%), Gaps:0/76(0%)

Query  656  LSNRWNFKGKAIGSDLGNGCf*fifeieENLEKVLANRPYHFDQWMVILQRWEPIISSTF  477
            L++ W F  +  G+DLGNG F F F  E++L + L  +PYHF QWM+I QRWEP I   F
Sbjct  63   LADHWKFSTRPTGADLGNGLFQFQFATEQDLARALKYQPYHFSQWMIIFQRWEPTIDKDF  122

Query  476  PSLIPF*VELQGLPKH  429
            PS IPF + +QG+  H
Sbjct  123  PSQIPFWINVQGVSIH  138


>hypothetical protein AXX17_AT5G32690 [Arabidopsis thaliana]
Sequence ID: OAO94166.1 Length: 1892
Range 1: 63 to 138

Score:80.9 bits(198), Expect:2e-15,
Method:Compositional matrix adjust.,
Identities:38/76(50%), Positives:50/76(65%), Gaps:0/76(0%)

Query  656  LSNRWNFKGKAIGSDLGNGCf*fifeieENLEKVLANRPYHFDQWMVILQRWEPIISSTF  477
            L++ W F  +  G+DLGNG F F F  E++L + L  +PYHF QWM+I QRWEP I   F
Sbjct  63   LADHWKFSTRPTGADLGNGLFQFQFATEQDLARALKYQPYHFSQWMIIFQRWEPTIDKDF  122

Query  476  PSLIPF*VELQGLPKH  429
            PS IPF + +QG+  H
Sbjct  123  PSQIPFWINVQGVSIH  138


>hypothetical protein At4g09700 [Arabidopsis thaliana]
Sequence ID: ABE65517.1 Length: 371
>putative protein [Arabidopsis thaliana]
Sequence ID: CAB39637.1 Length: 371 >putative protein [Arabidopsis thaliana]
Sequence ID: CAB78093.1 Length: 371
Range 1: 67 to 139

Score:75.1 bits(183), Expect:5e-15,
Method:Compositional matrix adjust.,
Identities:36/73(49%), Positives:49/73(67%), Gaps:0/73(0%)

Query  644  WNFKGKAIGSDLGNGCf*fifeieENLEKVLANRPYHFDQWMVILQRWEPIISSTFPSLI  465
            WN + + IG +LG   F F FE E +L+ VL   PYHF +WM ILQRWEPI+S  FP+ I
Sbjct  67   WNLENRVIGRELGPERFFFRFETEADLQLVLKKAPYHFKKWMFILQRWEPIVSEAFPAFI  126

Query  464  PF*VELQGLPKHY  426
            PF +++  +P H+
Sbjct  127  PFWIKVNDIPMHH  139


Range 2: 38 to 61

Score:26.9 bits(58), Expect:5e-15,
Method:Compositional matrix adjust.,
Identities:11/24(46%), Positives:15/24(62%), Gaps:0/24(0%)

Query  730  ASRTLTLMGRLTNPAAQRLWSLIS  659
            A    TL+GR+TNP  QR  +L+
Sbjct  38   AENKFTLLGRVTNPQIQRPRALVE  61


>unknown, partial [Arabidopsis thaliana]
Sequence ID: ABK28247.1 Length: 372
Range 1: 67 to 139

Score:74.7 bits(182), Expect:5e-15,
Method:Compositional matrix adjust.,
Identities:36/73(49%), Positives:49/73(67%), Gaps:0/73(0%)

Query  644  WNFKGKAIGSDLGNGCf*fifeieENLEKVLANRPYHFDQWMVILQRWEPIISSTFPSLI  465
            WN + + IG +LG   F F FE E +L+ VL   PYHF +WM ILQRWEPI+S  FP+ I
Sbjct  67   WNLENRVIGRELGPERFFFRFETEADLQLVLKKAPYHFKKWMFILQRWEPIVSEAFPAFI  126

Query  464  PF*VELQGLPKHY  426
            PF +++  +P H+
Sbjct  127  PFWIKVNDIPMHH  139


Range 2: 38 to 61

Score:26.9 bits(58), Expect:5e-15,
Method:Compositional matrix adjust.,
Identities:11/24(46%), Positives:15/24(62%), Gaps:0/24(0%)

Query  730  ASRTLTLMGRLTNPAAQRLWSLIS  659
            A    TL+GR+TNP  QR  +L+
Sbjct  38   AENKFTLLGRVTNPQIQRPRALVE  61


>putative Ta11-like non-LTR retroelement protein [Arabidopsis thaliana]
Sequence ID: AAD17399.1 Length: 589
Range 1: 45 to 146

Score:66.6 bits(161), Expect:6e-11,
Method:Compositional matrix adjust.,
Identities:38/109(35%), Positives:65/109(59%), Gaps:7/109(6%)

Query  752  LHSAMANRIPYSHSHGKTDKSCCSKTMVSNIRLSNRWNFKGKAIGSDLGNGCf*fifeie  573
            LH ++  R+   H  G++     +K ++S +   N W+ +G+  G  LG+  F F FE E
Sbjct  45   LHLSLVGRM--FHQGGRS-----TKALLSFLPKENIWDVEGRVRGVSLGDARFQFFFESE  97

Query  572  ENLEKVLANRPYHFDQWMVILQRWEPIISSTFPSLIPF*VELQGLPKHY  426
             +L+KVL  RP HF++W   L+RWEP + ++FP+++ F V  +G+P  +
Sbjct  98   VDLQKVLNKRPCHFNKWSFALERWEPHVGTSFPNIMTFWVRTEGIPAEF  146


>unknown, partial [Arabidopsis thaliana]
Sequence ID: ABK28059.1 Length: 134
Range 1: 61 to 133

Score:58.2 bits(139), Expect:1e-09,
Method:Composition-based stats.,
Identities:28/73(38%), Positives:46/73(63%), Gaps:0/73(0%)

Query  644  WNFKGKAIGSDLGNGCf*fifeieENLEKVLANRPYHFDQWMVILQRWEPIISSTFPSLI  465
            W  +G+ +G DLG G F   FE EE++ +V+    ++++ WM+ + RWEPII   +PS I
Sbjct  61   WKLEGQVVGRDLGLGTFRLDFEREEDILEVMKTETFNYNHWMLSIVRWEPIIHKDYPSAI  120

Query  464  PF*VELQGLPKHY  426
             F V + G+P+ +
Sbjct  121  TFWVRIVGVPRDF  133


>hypothetical protein [Arabidopsis thaliana]
Sequence ID: ABF59395.1 Length: 133
>unnamed protein product [Arabidopsis thaliana]
Sequence ID: BAB08530.1 Length: 133
Range 1: 61 to 133

Score:57.8 bits(138), Expect:1e-09,
Method:Composition-based stats.,
Identities:28/73(38%), Positives:46/73(63%), Gaps:0/73(0%)

Query  644  WNFKGKAIGSDLGNGCf*fifeieENLEKVLANRPYHFDQWMVILQRWEPIISSTFPSLI  465
            W  +G+ +G DLG G F   FE EE++ +V+    ++++ WM+ + RWEPII   +PS I
Sbjct  61   WKLEGQVVGRDLGLGTFRLDFEREEDILEVMKTETFNYNHWMLSIVRWEPIIHKDYPSAI  120

Query  464  PF*VELQGLPKHY  426
             F V + G+P+ +
Sbjct  121  TFWVRIVGVPRDF  133


>hypothetical protein [Arabidopsis thaliana]
Sequence ID: AAO37173.1 Length: 169
>hypothetical protein At2g06820 [Arabidopsis thaliana]
Sequence ID: AAV63870.1 Length: 169
Range 1: 1 to 32

Score:56.2 bits(134), Expect:1e-08,
Method:Compositional matrix adjust.,
Identities:25/32(78%), Positives:29/32(90%), Gaps:0/32(0%)

Query  521  MVILQRWEPIISSTFPSLIPF*VELQGLPKHY  426
            MVILQ+WEPIIS +FPSLI F +ELQGLPKH+
Sbjct  1    MVILQKWEPIISPSFPSLISFWIELQGLPKHF  32


>putative Ta11-like non-LTR retroelement protein [Arabidopsis thaliana]
Sequence ID: AAD22298.1 Length: 204
Range 1: 38 to 144

Score:56.6 bits(135), Expect:1e-08,
Method:Compositional matrix adjust.,
Identities:37/114(32%), Positives:60/114(52%), Gaps:13/114(11%)

Query  749  HSAMANRIPYS------HSHGKTDKSCCSKTMVSNIRLSNRWNFKGKAIGSDLGNGCf*f  588
            +SA+  R+  S      H  G++ ++  S        L N W+ +G+  G  LG+  F F
Sbjct  38   NSALLGRLQLSLVGRMFHQGGRSTEALLS-------FLPNIWDVEGRVRGVSLGDSRFQF  90

Query  587  ifeieENLEKVLANRPYHFDQWMVILQRWEPIISSTFPSLIPF*VELQGLPKHY  426
             FE E +L+KVL  RP HF +W   L+RW+  I  +FP  + F ++ +G+P  +
Sbjct  91   FFESETDLQKVLNKRPCHFSKWSFALERWKSHIGISFPDTMTFWIKTEGIPTEF  144


>unnamed protein product [Arabidopsis thaliana]
Sequence ID: CAA0360127.1 Length: 516
Range 1: 12 to 84

Score:58.9 bits(141), Expect:1e-08,
Method:Compositional matrix adjust.,
Identities:31/73(42%), Positives:45/73(61%), Gaps:0/73(0%)

Query  644  WNFKGKAIGSDLGNGCf*fifeieENLEKVLANRPYHFDQWMVILQRWEPIISSTFPSLI  465
            W  +GK +G+DLG G F F FE EE++ +VL   P+HFD WM+ L RW P +   +P  I
Sbjct  12   WQLEGKVVGADLGLGRFQFDFETEEDIIEVLKMEPFHFDHWMISLVRWSPSVHPDYPCSI  71

Query  464  PF*VELQGLPKHY  426
             F + + G+P  +
Sbjct  72   IFWIRVLGVPIQF  84


Query #108: XLOC_011722 Query ID: lcl|Query_33111 Length: 632

Sequences producing significant alignments:
                                                                  Max    Total Query   E   Per.
Description                                                       Score  Score cover Value Ident  Accession
unnamed protein product [Arabidopsis thaliana]                    207    207   48%   2e-68 99.03  VYS52415.1
unnamed protein product [Arabidopsis thaliana]                    203    203   48%   1e-66 96.12  CAA0361891.1
hypothetical protein AT2G14945 [Arabidopsis thaliana]             148    148   48%   2e-45 77.67  NP_001324754.1
hypothetical protein AXX17_AT4G21170 [Arabidopsis thaliana]       105    165   80%   1e-34 56.18  OAO96867.1
hypothetical protein AXX17_AT5G29230 [Arabidopsis thaliana]       101    101   50%   3e-26 46.30  OAO91562.1
hypothetical protein AXX17_AT4G06750 [Arabidopsis thaliana]       102    102   40%   1e-25 57.65  OAO98111.1
RNA-directed DNA polymerase-like protein [Arabidopsis thaliana]   102    102   48%   2e-24 47.57  CAB39638.1
hypothetical protein AXX17_AT3G37590 [Arabidopsis thaliana]       69.7   126   83%   9e-23 38.10  OAP06812.1
hypothetical protein AXX17_AT3G15320 [Arabidopsis thaliana]       96.3   153   87%   1e-22 42.57  OAP05086.1
hypothetical protein AXX17_AT3G15320 [Arabidopsis thaliana]       95.9   152   87%   3e-22 41.75  OAP05085.1
hypothetical protein AXX17_AT3G15320 [Arabidopsis thaliana]       95.9   152   87%   3e-22 41.75  OAP05088.1
Ribonuclease H-like superfamily protein [Arabidopsis thaliana]    91.7   91.7  48%   2e-21 47.17  NP_172471.1
Hypothetical protein [Arabidopsis thaliana]                       91.3   91.3  48%   4e-21 47.17  AAC34342.1
putative non-LTR retroelement reverse transcriptase [Arabidops... 88.2   88.2  46%   2e-19 45.10  AAD29058.1
hypothetical protein AXX17_AT2G05620 [Arabidopsis thaliana]       83.2   83.2  40%   3e-18 44.19  OAP08393.1
unnamed protein product [Arabidopsis thaliana]                    80.9   80.9  47%   2e-17 39.60  BAB02611.1
Beta-galactosidase related protein [Arabidopsis thaliana]         81.3   81.3  40%   2e-17 43.02  NP_671791.2
unnamed protein product [Arabidopsis thaliana]                    80.1   80.1  40%   4e-17 43.02  VYS52169.1
unnamed protein product [Arabidopsis thaliana]                    70.5   70.5  46%   1e-14 41.00  CAA0394294.1
non-LTR retroelement reverse transcriptase-like protein...        70.1   70.1  50%   2e-13 34.58  BAB08714.1
hypothetical protein AXX17_AT5G32690 [Arabidopsis thaliana]       69.3   69.3  50%   5e-13 34.58  OAO94166.1
putative non-LTR retroelement reverse transcriptase [Arabidops... 68.6   68.6  56%   7e-13 31.97  AAB84340.1
hypothetical protein AXX17_ATUG04250 [Arabidopsis thaliana]       67.8   67.8  22%   1e-12 65.96  OAO89120.1
Polynucleotidyl transferase, ribonuclease H-like superfamily...   64.7   64.7  38%   2e-12 40.24  NP_192687.1
putative non-LTR retroelement reverse transcriptase [Arabidops... 66.6   66.6  64%   3e-12 28.57  AAD24601.1
Ribonuclease H-like superfamily protein [Arabidopsis thaliana]    65.1   65.1  59%   7e-12 34.81  NP_189164.1
unnamed protein product [Arabidopsis thaliana]                    65.9   65.9  48%   7e-12 35.85  CAA0406188.1
hypothetical protein AXX17_AT2G11760 [Arabidopsis thaliana]       65.5   65.5  27%   1e-11 49.15  OAP10255.1
reverse transcriptase, putative; 16838-20266 [Arabidopsis...      63.2   63.2  48%   4e-11 32.73  AAG51783.1
putative non-LTR retroelement reverse transcriptase [Arabidops... 62.8   125   63%   6e-11 52.00  AAD15377.1
hypothetical protein AXX17_AT1G36150 [Arabidopsis thaliana]       62.8   62.8  28%   7e-11 42.62  OAP15291.1
unnamed protein product [Arabidopsis thaliana]                    62.4   62.4  28%   9e-11 42.62  VYS68395.1
hypothetical protein AXX17_AT3G32490 [Arabidopsis thaliana]       62.4   124   31%   1e-10 43.28  OAP03411.1
hypothetical protein At2g15750 [Arabidopsis thaliana]             59.3   59.3  34%   1e-10 41.77  ABE65445.1
unknown [Arabidopsis thaliana]                                    59.3   59.3  34%   1e-10 41.77  ABK28185.1
hypothetical protein AXX17_AT1G43730 [Arabidopsis thaliana]       62.0   62.0  48%   1e-10 32.41  OAP19763.1
hypothetical protein AXX17_AT3G37030 [Arabidopsis thaliana]       62.0   123   27%   2e-10 56.60  OAP06308.1
putative non-LTR retroelement reverse transcriptase [Arabidops... 61.6   61.6  68%   2e-10 28.38  AAD17395.1
hypothetical protein AXX17_AT2G06510 [Arabidopsis thaliana]       59.3   59.3  37%   2e-10 38.75  OAP07274.1
Very similar to retrotransposon reverse transcriptase...          61.6   61.6  63%   2e-10 30.61  AAF18538.1
hypothetical protein AXX17_AT5G33430 [Arabidopsis thaliana]       61.2   61.2  36%   2e-10 38.46  OAO95064.1
Ribonuclease H-like superfamily protein [Arabidopsis thaliana]    60.1   60.1  48%   6e-10 31.73  NP_194638.1
unnamed protein product [Arabidopsis thaliana]                    59.7   59.7  48%   6e-10 31.73  CAA0396872.1
putative non-LTR retroelement reverse transcriptase [Arabidops... 59.7   59.7  48%   9e-10 30.77  AAB82639.1
reverse transcriptase [Arabidopsis thaliana]                      59.3   59.3  48%   1e-09 33.64  AAA75254.1
RNA-directed DNA polymerase (reverse transcriptase)-related...    56.6   56.6  25%   1e-09 41.51  NP_683598.1
putative non-LTR retroelement reverse transcriptase [Arabidops... 56.2   56.2  48%   1e-08 32.43  AAD17398.1
unnamed protein product [Arabidopsis thaliana]                    56.2   56.2  37%   1e-08 40.00  CAA0247490.1
unnamed protein product [Arabidopsis thaliana]                    55.5   55.5  28%   2e-08 40.00  CAA0405280.1


Alignments:
>unnamed protein product, partial [Arabidopsis thaliana]
Sequence ID: VYS52415.1 Length: 150
Range 1: 1 to 103

Score:207 bits(528), Expect:2e-68,
Method:Compositional matrix adjust.,
Identities:102/103(99%), Positives:102/103(99%), Gaps:0/103(0%)

Query  324  IAQTIQGSRGPAFKPVGGRQIKSLEASIPALRDGIRSPDKEFLKGARSQTCLAKGHLKNK  503
            IAQTIQGSRGPAFKPVGGRQIKSLEASIPALRDGIRSPDKEFLKGARSQTCLAKGHLKNK
Sbjct  1    IAQTIQGSRGPAFKPVGGRQIKSLEASIPALRDGIRSPDKEFLKGARSQTCLAKGHLKNK  60

Query  504  CRTDSSFPHKVHKAVSFMCLVLRCSPKGRAPFISFQRKNLVFG  632
            CR DSSFPHKVHKAVSFMCLVLRCSPKGRAPFISFQRKNLVFG
Sbjct  61   CRADSSFPHKVHKAVSFMCLVLRCSPKGRAPFISFQRKNLVFG  103


>unnamed protein product, partial [Arabidopsis thaliana]
Sequence ID: CAA0361891.1 Length: 150
Range 1: 1 to 103

Score:203 bits(516), Expect:1e-66,
Method:Compositional matrix adjust.,
Identities:99/103(96%), Positives:102/103(99%), Gaps:0/103(0%)

Query  324  IAQTIQGSRGPAFKPVGGRQIKSLEASIPALRDGIRSPDKEFLKGARSQTCLAKGHLKNK  503
            IA+TIQGSRGPAFKPVGGRQIKSLEASIPALRDGI+SPDKEFLKGARSQTCLAKGHLKNK
Sbjct  1    IARTIQGSRGPAFKPVGGRQIKSLEASIPALRDGIQSPDKEFLKGARSQTCLAKGHLKNK  60

Query  504  CRTDSSFPHKVHKAVSFMCLVLRCSPKGRAPFISFQRKNLVFG  632
            CR DSSFPHKVHKAVSFMCLVLRCSPKG+APFISFQRKNLVFG
Sbjct  61   CRADSSFPHKVHKAVSFMCLVLRCSPKGKAPFISFQRKNLVFG  103


>hypothetical protein AT2G14945, partial [Arabidopsis thaliana]
Sequence ID: NP_001324754.1 Length: 132
>hypothetical protein AT2G14945, partial [Arabidopsis thaliana]
Sequence ID: ANM62608.1 Length: 132
Range 1: 1 to 80

Score:148 bits(374), Expect:2e-45,
Method:Compositional matrix adjust.,
Identities:80/103(78%), Positives:80/103(77%), Gaps:23/103(22%)

Query  324  IAQTIQGSRGPAFKPVGGRQIKSLEASIPALRDGIRSPDKEFLKGARSQTCLAKGHLKNK  503
            IAQTIQGSRGPAFKPVGGRQIKSLEASIPALRDGIRSPDKEFLKGARSQTCLAKGHLKNK
Sbjct  1    IAQTIQGSRGPAFKPVGGRQIKSLEASIPALRDGIRSPDKEFLKGARSQTCLAKGHLKNK  60

Query  504  CRTDSSFPHKVHKAVSFMCLVLRCSPKGRAPFISFQRKNLVFG  632
                                   CSPKGRAPFISFQRKNLVFG
Sbjct  61   -----------------------CSPKGRAPFISFQRKNLVFG  80


>hypothetical protein AXX17_AT4G21170 [Arabidopsis thaliana]
Sequence ID: OAO96867.1 Length: 1816
Range 1: 1561 to 1649

Score:105 bits(262), Expect:1e-34,
Method:Compositional matrix adjust.,
Identities:50/89(56%), Positives:62/89(69%), Gaps:0/89(0%)

Query  628   KTKFFLWKLMKGALPLGEHLRTRHINDTALCTLCGNEESVLHLFFKCPFAKQVWDLAPFK  449
             K KFFLWK + GALP+GE LR+R I   + C  C   E+ LHLFF CPFAK++W  APFK
Sbjct  1561  KIKFFLWKALAGALPVGELLRSRGIRAESSCPHCQAAETCLHLFFHCPFAKEIWAAAPFK  1620

Query  448   NSLSGDRIPSLRAGIEASKLLICLPPTGL  362
             ++++  RI S R+GIE SK    LPPTGL
Sbjct  1621  STINPLRISSFRSGIEGSKAWTTLPPTGL  1649


Range 2: 1675 to 1754

Score:60.5 bits(145), Expect:1e-34,
Method:Compositional matrix adjust.,
Identities:30/80(38%), Positives:49/80(61%), Gaps:0/80(0%)

Query  323   LCFTDALWSESSKSAGFGWIIKFRRSGFRSKGSASDSHIRSLVVAEAIAIQLALKQSLRL  144
             +CF+DA W   +K AGFGWI   + S    +G  ++ ++ S ++AEA+A+  A +Q+L L
Sbjct  1675  ICFSDAAWHSDTKMAGFGWIFLNQASKLEIQGFGTNLNVGSSLLAEALALYSAQQQALDL  1734

Query  143   GITYLAIVSDAKQVIETINS  84
             G   L   SD++ +I  +NS
Sbjct  1735  GFKKLHFASDSQLLISALNS  1754


>hypothetical protein AXX17_AT5G29230 [Arabidopsis thaliana]
Sequence ID: OAO91562.1 Length: 197
Range 1: 1 to 108

Score:101 bits(252), Expect:3e-26,
Method:Compositional matrix adjust.,
Identities:50/108(46%), Positives:65/108(60%), Gaps:2/108(1%)

Query  601  MKGALPLGEHLRTRHINDTALCTLCGNEESVLHLFFKCPFAKQVWDLAPFKNSLSGDRIP  422
            M+GALP GE+L  R +   A C  CG  E+  HLF KC FA  +WD APF+  L
Sbjct  1    MRGALPAGENLGLRGLRVEANCPFCGEAETPFHLFVKCQFATWIWDHAPFQKHLDPATFT  60

Query  421  SLRAGIEASKLLICLPPTGLNAGPLLPWIVWAISVS--QMLYGLKAPN  284
            SL+A IE +K   CLPP G+N GPL PWI+W+I ++  Q ++  K  N
Sbjct  61   SLQAFIEKAKTHTCLPPCGVNVGPLYPWIIWSIWITRNQRIFNDKRTN  108


>hypothetical protein AXX17_AT4G06750 [Arabidopsis thaliana]
Sequence ID: OAO98111.1 Length: 310
Range 1: 221 to 305

Score:102 bits(255), Expect:1e-25,
Method:Compositional matrix adjust.,
Identities:49/85(58%), Positives:59/85(69%), Gaps:0/85(0%)

Query  631  PKTKFFLWKLMKGALPLGEHLRTRHINDTALCTLCGNEESVLHLFFKCPFAKQVWDLAPF  452
            PK KFFLWK M+GALP+GE+L  R+IN  A C  CG  ES LHLFF C FA+ VW L PF
Sbjct  221  PKLKFFLWKAMRGALPVGENLIFRNINAAAQCPFCGENESTLHLFFTCRFARHVWLLNPF  280

Query  451  KNSLSGDRIPSLRAGIEASKLLICL  377
            +  L+ D+I S + GIE +K  ICL
Sbjct  281  ETRLNPDQIVSFKDGIEQTKGQICL  305


>RNA-directed DNA polymerase-like protein [Arabidopsis thaliana]
Sequence ID: CAB39638.1 Length: 1274
>RNA-directed DNA polymerase-like protein [Arabidopsis thaliana]
Sequence ID: CAB78094.1 Length: 1274
Range 1: 981 to 1083

Score:102 bits(253), Expect:2e-24,
Method:Compositional matrix adjust.,
Identities:49/103(48%), Positives:64/103(62%), Gaps:0/103(0%)

Query  631   PKTKFFLWKLMKGALPLGEHLRTRHINDTALCTLCGNEESVLHLFFKCPFAKQVWDLAPF  452
             PK K FLWK MKGALP+GE L  R+I     C  CG  ES LHL   CP+AK+VW+LAP
Sbjct  981   PKVKHFLWKAMKGALPVGEALSRRNIEAEVTCKRCGQTESSLHLMLLCPYAKKVWELAPV  1040

Query  451   KNSLSGDRIPSLRAGIEASKLLICLPPTGLNAGPLLPWIVWAI  323
               + S     S+   +  +K ++ LPPTGL + PL PW++W +
Sbjct  1041  LFNPSEATHSSVALLLVDAKRMVALPPTGLGSAPLYPWLLWHL  1083


>hypothetical protein AXX17_AT3G37590 [Arabidopsis thaliana]
Sequence ID: OAP06812.1 Length: 903
Range 1: 773 to 856

Score:69.7 bits(169), Expect:9e-23,
Method:Compositional matrix adjust.,
Identities:32/84(38%), Positives:54/84(64%), Gaps:0/84(0%)

Query  320  CFTDALWSESSKSAGFGWIIKFRRSGFRSKGSASDSHIRSLVVAEAIAIQLALKQSLRLG  141
            CFTDA W E S  AG GWI     +   S   +++++I   ++AE+ A+ LA++ +  LG
Sbjct  773  CFTDASWREESLEAGLGWIFVDHWNHTESHHQSAEANINYPLMAESTALLLAIQHASDLG  832

Query  140  ITYLAIVSDAKQVIETINSETPSM  69
            +  LAIVSD++Q+++ +N E+P +
Sbjct  833  LKKLAIVSDSQQLVKALNGESPPL  856


Range 2: 679 to 768

Score:56.6 bits(135), Expect:9e-23,
Method:Compositional matrix adjust.,
Identities:39/95(41%), Positives:53/95(55%), Gaps:8/95(8%)

Query  631  PKTKFFLWKLMKGALPLGEHLRTRHINDTALCTLCGNEESVLHLFFKCPFAKQVWDLAPF  452
            PK KF LWK MK ALP+GE+LR R+IN  A C  CG +ES   +F +    K +  L
Sbjct  679  PKLKFLLWKSMKNALPVGENLRIRNINPLANCPHCGEDESNKLIFEQ----KHISSLDLI  734

Query  451  KNSLSGDRIPSLRAGIEASKLLIC---LPPTGLNA  356
              S+S  +   L A I+AS   I    +PPT +++
Sbjct  735  SQSISQAK-EWLGAHIQASSSQIVKSWIPPTEIDS  768


>hypothetical protein AXX17_AT3G15320 [Arabidopsis thaliana]
Sequence ID: OAP05086.1 Length: 424
Range 1: 103 to 203

Score:96.3 bits(238), Expect:1e-22,
Method:Compositional matrix adjust.,
Identities:43/101(43%), Positives:64/101(63%), Gaps:0/101(0%)

Query  631  PKTKFFLWKLMKGALPLGEHLRTRHINDTALCTLCGNEESVLHLFFKCPFAKQVWDLAPF  452
            PK K FLWKL + ALP+G +L  R I +   C  CG  ES LH+   CP+A+++W++AP
Sbjct  103  PKLKMFLWKLKRRALPVGGNLVIRGITNDLTCKRCGGLESELHILLLCPYAQRLWEMAPL  162

Query  451  KNSLSGDRIPSLRAGIEASKLLICLPPTGLNAGPLLPWIVW  329
              S   DRI SL A + ++++ +CLPP+G++   +  WI W
Sbjct  163  LRSPRADRIESLAALLSSARVALCLPPSGISHTSIQHWIFW  203


Range 2: 288 to 373

Score:57.4 bits(137), Expect:3e-09,
Method:Compositional matrix adjust.,
Identities:35/86(41%), Positives:53/86(61%), Gaps:0/86(0%)

Query  335  CLGDLCFTDALWSESSKSAGFGWIIKFRRSGFRSKGSASDSHIRSLVVAEAIAIQLALKQ  156
            C   LCFTDA W+ SS S G GWI K        +GS+S  H  S + AEA+A++ AL +
Sbjct  288  CEAVLCFTDAAWNASSGSCGMGWIFKTHNQRVIHQGSSSRLHTPSALAAEALAVKAALNE  347

Query  155  SLRLGITYLAIVSDAKQVIETINSET  78
            +LR+ +T + + SD++ +I  + +ET
Sbjct  348  ALRMDLTSITVYSDSQVLISLLTTET  373


>hypothetical protein AXX17_AT3G15320 [Arabidopsis thaliana]
Sequence ID: OAP05085.1 Length: 1247
Range 1: 926 to 1028

Score:95.9 bits(237), Expect:3e-22,
Method:Compositional matrix adjust.,
Identities:43/103(42%), Positives:65/103(63%), Gaps:0/103(0%)

Query  631   PKTKFFLWKLMKGALPLGEHLRTRHINDTALCTLCGNEESVLHLFFKCPFAKQVWDLAPF  452
             PK K FLWKL + ALP+G +L  R I +   C  CG  ES LH+   CP+A+++W++AP
Sbjct  926   PKLKMFLWKLKRRALPVGGNLVIRGITNDLTCKRCGGLESELHILLLCPYAQRLWEMAPL  985

Query  451   KNSLSGDRIPSLRAGIEASKLLICLPPTGLNAGPLLPWIVWAI  323
               S   DRI SL A + ++++ +CLPP+G++   +  WI W +
Sbjct  986   LRSPRADRIESLAALLSSARVALCLPPSGISHTSIQHWIFWDL  1028


Range 2: 1111 to 1196

Score:56.6 bits(135), Expect:8e-09,
Method:Compositional matrix adjust.,
Identities:35/86(41%), Positives:53/86(61%), Gaps:0/86(0%)

Query  335   CLGDLCFTDALWSESSKSAGFGWIIKFRRSGFRSKGSASDSHIRSLVVAEAIAIQLALKQ  156
             C   LCFTDA W+ SS S G GWI K        +GS+S  H  S + AEA+A++ AL +
Sbjct  1111  CEAVLCFTDAAWNASSGSCGMGWIFKTHNQRVIHQGSSSRLHTPSALAAEALAVKAALNE  1170

Query  155   SLRLGITYLAIVSDAKQVIETINSET  78
             +LR+ +T + + SD++ +I  + +ET
Sbjct  1171  ALRMDLTSITVYSDSQVLISLLTTET  1196


>hypothetical protein AXX17_AT3G15320 [Arabidopsis thaliana]
Sequence ID: OAP05088.1 Length: 1697
Range 1: 1376 to 1478

Score:95.9 bits(237), Expect:3e-22,
Method:Compositional matrix adjust.,
Identities:43/103(42%), Positives:65/103(63%), Gaps:0/103(0%)

Query  631   PKTKFFLWKLMKGALPLGEHLRTRHINDTALCTLCGNEESVLHLFFKCPFAKQVWDLAPF  452
             PK K FLWKL + ALP+G +L  R I +   C  CG  ES LH+   CP+A+++W++AP
Sbjct  1376  PKLKMFLWKLKRRALPVGGNLVIRGITNDLTCKRCGGLESELHILLLCPYAQRLWEMAPL  1435

Query  451   KNSLSGDRIPSLRAGIEASKLLICLPPTGLNAGPLLPWIVWAI  323
               S   DRI SL A + ++++ +CLPP+G++   +  WI W +
Sbjct  1436  LRSPRADRIESLAALLSSARVALCLPPSGISHTSIQHWIFWDL  1478


Range 2: 1561 to 1646

Score:57.0 bits(136), Expect:7e-09,
Method:Compositional matrix adjust.,
Identities:35/86(41%), Positives:53/86(61%), Gaps:0/86(0%)

Query  335   CLGDLCFTDALWSESSKSAGFGWIIKFRRSGFRSKGSASDSHIRSLVVAEAIAIQLALKQ  156
             C   LCFTDA W+ SS S G GWI K        +GS+S  H  S + AEA+A++ AL +
Sbjct  1561  CEAVLCFTDAAWNASSGSCGMGWIFKTHNQRVIHQGSSSRLHTPSALAAEALAVKAALNE  1620

Query  155   SLRLGITYLAIVSDAKQVIETINSET  78
             +LR+ +T + + SD++ +I  + +ET
Sbjct  1621  ALRMDLTSITVYSDSQVLISLLTTET  1646


>Ribonuclease H-like superfamily protein [Arabidopsis thaliana]
Sequence ID: NP_172471.1 Length: 303
>Ribonuclease H-like superfamily protein [Arabidopsis thaliana]
Sequence ID: AEE28527.1 Length: 303
Range 1: 2 to 103

Score:91.7 bits(226), Expect:2e-21,
Method:Compositional matrix adjust.,
Identities:50/106(47%), Positives:58/106(54%), Gaps:7/106(6%)

Query  631  PKTKFFLWKLMKGALPLGEHLRTRHINDTALCTLCGNEESVLHLFFKCPFAKQVWDLAPF  452
            PK K FLWK   GALP+G  L  RHI+ T  C  CG  E+  H+ F C FA QVW+LAP
Sbjct  2    PKIKLFLWKAAAGALPVGAQLVRRHISSTWDCARCGAPETSTHMLFHCDFAAQVWNLAP-  60

Query  451  KNSLSGDRIPSLRAGIEASKLL---ICLPPTGLNAGPLLPWIVWAI  323
               L    IP     +EA  LL   I LPP G++   L PWI W I
Sbjct  61   ---LQLGHIPIGTPILEALNLLKKTIVLPPVGIHTATLFPWICWHI  103


>Hypothetical protein [Arabidopsis thaliana]
Sequence ID: AAC34342.1 Length: 374
Range 1: 2 to 103

Score:91.3 bits(225), Expect:4e-21,
Method:Compositional matrix adjust.,
Identities:50/106(47%), Positives:58/106(54%), Gaps:7/106(6%)

Query  631  PKTKFFLWKLMKGALPLGEHLRTRHINDTALCTLCGNEESVLHLFFKCPFAKQVWDLAPF  452
            PK K FLWK   GALP+G  L  RHI+ T  C  CG  E+  H+ F C FA QVW+LAP
Sbjct  2    PKIKLFLWKAAAGALPVGAQLVRRHISSTWDCARCGAPETSTHMLFHCDFAAQVWNLAP-  60

Query  451  KNSLSGDRIPSLRAGIEASKLL---ICLPPTGLNAGPLLPWIVWAI  323
               L    IP     +EA  LL   I LPP G++   L PWI W I
Sbjct  61   ---LQLGHIPIGTPILEALNLLKKTIVLPPVGIHTATLFPWICWHI  103


>putative non-LTR retroelement reverse transcriptase [Arabidopsis thaliana]
Sequence ID: AAD29058.1 Length: 1229
Range 1: 941 to 1038

Score:88.2 bits(217), Expect:2e-19,
Method:Compositional matrix adjust.,
Identities:46/102(45%), Positives:59/102(57%), Gaps:7/102(6%)

Query  631   PKTKFFLWKLMKGALPLGEHLRTRHINDTALCTLCGNEESVLHLFFKCPFAKQVWDLAPF  452
             PK K  +WK    ALP+G  L  RHI+ +A C  CG  ES  HLFF C FA QVW+LAP
Sbjct  941   PKIKHLMWKAAMEALPVGIQLVRRHISPSAACHRCGAPESTTHLFFHCEFAAQVWELAPL  1000

Query  451   KNSLSGDRIPSLRAGIEASKLL---ICLPPTGLNAGPLLPWI  335
             + +     +P   + ++A  LL   I LPPTG+ +  L PWI
Sbjct  1001  QETT----VPPGSSMLDALSLLKKAIILPPTGVTSAALFPWI  1038


>hypothetical protein AXX17_AT2G05620 [Arabidopsis thaliana]
Sequence ID: OAP08393.1 Length: 317
Range 1: 66 to 151

Score:83.2 bits(204), Expect:3e-18,
Method:Compositional matrix adjust.,
Identities:38/86(44%), Positives:50/86(58%), Gaps:0/86(0%)

Query  580  GEHLRTRHINDTALCTLCGNEESVLHLFFKCPFAKQVWDLAPFKNSLSGDRIPSLRAGIE  401
            G+ L  RHI   A C  CG  E++LH+ F C FA+ VW  APF    +     S+R G+
Sbjct  66   GQQLLDRHIAIDASCCRCGEPETILHMLFHCQFAQLVWSPAPFARHWASVGAGSVREGLI  125

Query  400  ASKLLICLPPTGLNAGPLLPWIVWAI  323
                L CLPPTG+ +GPL PWI W++
Sbjct  126  TGCKLACLPPTGIASGPLAPWICWSL  151


>unnamed protein product [Arabidopsis thaliana]
Sequence ID: BAB02611.1 Length: 332
Range 1: 139 to 239

Score:80.9 bits(198), Expect:2e-17,
Method:Compositional matrix adjust.,
Identities:40/101(40%), Positives:53/101(52%), Gaps:0/101(0%)

Query  631  PKTKFFLWKLMKGALPLGEHLRTRHINDTALCTLCGNEESVLHLFFKCPFAKQVWDLAPF  452
            P  + FL K  K ALP+G  L  R I     C  CG  E  LH+FF+CP A +VWDLA
Sbjct  139  PMIRMFLSKATKKALPIGTALSARGIVVETGCKRCGEIEDALHIFFRCPIAVKVWDLATL  198

Query  451  KNSLSGDRIPSLRAGIEASKLLICLPPTGLNAGPLLPWIVW  329
                    +  ++  +E  +  + LPP GL + PL PWI+W
Sbjct  199  LPKPRAGSVVYVKKILEKGRTCLSLPPLGLGSTPLYPWILW  239


>Beta-galactosidase related protein [Arabidopsis thaliana]
Sequence ID: NP_671791.2 Length: 408
>Beta-galactosidase related protein [Arabidopsis thaliana]
Sequence ID: AEC06020.2 Length: 408
Range 1: 157 to 242

Score:81.3 bits(199), Expect:2e-17,
Method:Compositional matrix adjust.,
Identities:37/86(43%), Positives:49/86(56%), Gaps:0/86(0%)

Query  580  GEHLRTRHINDTALCTLCGNEESVLHLFFKCPFAKQVWDLAPFKNSLSGDRIPSLRAGIE  401
            G+ L  RHI   A C  CG  E++LH+ F C FA+ VW   PF    +     S+R G+
Sbjct  157  GQQLLDRHIAIDASCCRCGEPETILHMLFHCQFAQLVWSPTPFARHWASVGAGSVREGLI  216

Query  400  ASKLLICLPPTGLNAGPLLPWIVWAI  323
                L CLPPTG+ +GPL PWI W++
Sbjct  217  TGCKLACLPPTGIASGPLAPWICWSL  242


>unnamed protein product [Arabidopsis thaliana]
Sequence ID: VYS52169.1 Length: 330
Range 1: 79 to 164

Score:80.1 bits(196), Expect:4e-17,
Method:Compositional matrix adjust.,
Identities:37/86(43%), Positives:49/86(56%), Gaps:0/86(0%)

Query  580  GEHLRTRHINDTALCTLCGNEESVLHLFFKCPFAKQVWDLAPFKNSLSGDRIPSLRAGIE  401
            G+ L  RHI   A C  C   E++LH+ F C FA+ VW  APF    +     S+R G+
Sbjct  79   GQQLLDRHIAIDASCCRCREPETILHMLFHCQFAQLVWSPAPFARHWASVGAGSVREGLI  138

Query  400  ASKLLICLPPTGLNAGPLLPWIVWAI  323
                L CLPPTG+ +GPL PWI W++
Sbjct  139  TGCKLACLPPTGIASGPLAPWICWSL  164


>unnamed protein product [Arabidopsis thaliana]
Sequence ID: CAA0394294.1 Length: 169
Range 1: 15 to 114

Score:70.5 bits(171), Expect:1e-14,
Method:Compositional matrix adjust.,
Identities:41/100(41%), Positives:61/100(61%), Gaps:2/100(2%)

Query  365  LKRRASAALDCLGDLC--FTDALWSESSKSAGFGWIIKFRRSGFRSKGSASDSHIRSLVV  192
            L  R+ AAL    D    FTDA W   +K  GFGW+I+           ++  ++RS ++
Sbjct  15   LPHRSFAALRPDLDRVSIFTDAAWKHETKDVGFGWVIRNCPELTALHHQSASRNVRSPLM  74

Query  191  AEAIAIQLALKQSLRLGITYLAIVSDAKQVIETINSETPS  72
            AEAIA+ LAL+ +  +GIT L++ SD++Q+I TI SE+PS
Sbjct  75   AEAIALFLALQYAQSIGITKLSMASDSQQLITTITSESPS  114


>non-LTR retroelement reverse transcriptase-like protein [Arabidopsis thaliana]
Sequence ID: BAB08714.1 Length: 1197
Range 1: 880 to 986

Score:70.1 bits(170), Expect:2e-13,
Method:Compositional matrix adjust.,
Identities:37/107(35%), Positives:53/107(49%), Gaps:0/107(0%)

Query  631  PKTKFFLWKLMKGALPLGEHLRTRHINDTALCTLCGNEESVLHLFFKCPFAKQVWDLAPF  452
            PK + FLWK + GALP+G  L  R  +   +C  CG  E+  H  F C F ++VW LAP
Sbjct  880  PKIRLFLWKAVSGALPVGSQLAIRIPSFDPVCCRCGELETSSHALFNCMFVQRVWRLAPL  939

Query  451  KNSLSGDRIPSLRAGIEASKLLICLPPTGLNAGPLLPWIVWAISVSQ  311
            +          ++ G+   K    LP  GL    L PWI W + +++
Sbjct  940  EGIDLFLVFTDVKLGLSWLKKKKTLPSVGLGYCALYPWICWTLWITR  986


>hypothetical protein AXX17_AT5G32690 [Arabidopsis thaliana]
Sequence ID: OAO94166.1 Length: 1892
Range 1: 1442 to 1548

Score:69.3 bits(168), Expect:5e-13,
Method:Compositional matrix adjust.,
Identities:37/107(35%), Positives:53/107(49%), Gaps:0/107(0%)

Query  631   PKTKFFLWKLMKGALPLGEHLRTRHINDTALCTLCGNEESVLHLFFKCPFAKQVWDLAPF  452
             PK + FLWK + GALP+G  L  R  +   +C  CG  E+  H  F C F ++VW LAP
Sbjct  1442  PKIRLFLWKAVSGALPVGSQLAIRIPSFDPVCCRCGELETSSHALFNCMFVQRVWRLAPL  1501

Query  451   KNSLSGDRIPSLRAGIEASKLLICLPPTGLNAGPLLPWIVWAISVSQ  311
             +          ++ G+   K    LP  GL    L PWI W + +++
Sbjct  1502  EGIDLFLVFTDVKLGLSWLKKKKTLPSVGLGYCALYPWICWTLWITR  1548


>putative non-LTR retroelement reverse transcriptase [Arabidopsis thaliana]
Sequence ID: AAB84340.1 Length: 1094
Range 1: 782 to 903

Score:68.6 bits(166), Expect:7e-13,
Method:Compositional matrix adjust.,
Identities:39/122(32%), Positives:64/122(52%), Gaps:3/122(2%)

Query  631  PKTKFFLWKLMKGALPLGEHLRTRHINDTALCTLC-GNEESVLHLFFKCPFAKQVWDLAP  455
            PK K FLWK++KGA+ + + LRTR +     C++C    E++ H+ F+CP A+QVW L P
Sbjct  782  PKIKVFLWKVLKGAVAVEDRLRTRGVLIEDGCSMCPEKNETLNHILFQCPLARQVWALTP  841

Query  454  FK--NSLSGDRIPSLRAGIEASKLLICLPPTGLNAGPLLPWIVWAISVSQMLYGLKAPNL  281
             +  N   GD I +    +  +     L P      P + WI+W     ++  G+ + +L
Sbjct  842  MQSPNHGFGDSIFTNVNHVIGNCHNTELSPHLRYVSPWIIWILWKNRNKRLFEGIGSVSL  901

Query  280  QV  275
             +
Sbjct  902  SI  903


>hypothetical protein AXX17_ATUG04250 [Arabidopsis thaliana]
Sequence ID: OAO89120.1 Length: 803
Range 1: 652 to 698

Score:67.8 bits(164), Expect:1e-12,
Method:Compositional matrix adjust.,
Identities:31/47(66%), Positives:36/47(76%), Gaps:0/47(0%)

Query  631  PKTKFFLWKLMKGALPLGEHLRTRHINDTALCTLCGNEESVLHLFFK  491
            PKTKFFLWKLM+GALPLGE+L+ RHI D A  TLC  EE+ L  F +
Sbjct  652  PKTKFFLWKLMRGALPLGENLKARHIIDVAGSTLCAQEETRLGAFLQ  698


>Polynucleotidyl transferase, ribonuclease H-like superfamily protein [Arabidopsis thaliana]
Sequence ID: NP_192687.1 Length: 170
>Polynucleotidyl transferase, ribonuclease H-like superfamily protein [Arabidopsis thaliana]
Sequence ID: AEE82756.1 Length: 170 >putative proteins [Arabidopsis thaliana]
Sequence ID: CAB78072.1 Length: 170
Range 1: 34 to 115

Score:64.7 bits(156), Expect:2e-12,
Method:Compositional matrix adjust.,
Identities:33/82(40%), Positives:52/82(63%), Gaps:0/82(0%)

Query  317  FTDALWSESSKSAGFGWIIKFRRSGFRSKGSASDSHIRSLVVAEAIAIQLALKQSLRLGI  138
            FTDA W   +K  GFGW+I+           ++  ++R  ++AEAIA+ LAL+ +  +GI
Sbjct  34   FTDAAWKHETKDVGFGWVIRNCPELTALHYQSAARNVRLPLMAEAIALFLALQYAQSIGI  93

Query  137  TYLAIVSDAKQVIETINSETPS  72
            T L++ SD++Q+I  I SE+PS
Sbjct  94   TKLSMASDSQQLITAITSESPS  115


>putative non-LTR retroelement reverse transcriptase [Arabidopsis thaliana]
Sequence ID: AAD24601.1 Length: 1319
Range 1: 1005 to 1143

Score:66.6 bits(161), Expect:3e-12,
Method:Compositional matrix adjust.,
Identities:40/140(29%), Positives:67/140(47%), Gaps:5/140(3%)

Query  631   PKTKFFLWKLMKGALPLGEHLRTRHINDTALCTLCGNE-ESVLHLFFKCPFAKQVWDLAP  455
             PK K F+WK +KGAL + + LR+R I     C  C  E E++ HL F+CPFA+QVW L+
Sbjct  1005  PKIKVFMWKALKGALAVEDRLRSRGIRTADGCLFCKEEIETINHLLFQCPFARQVWALSL  1064

Query  454   FKNSLSG---DRIPSLRAGIEASKLLICLPPTGLNAGPLLPWIVWAISVSQMLYGLKAPN  284
              +   +G       ++   I+ S+    +P       P L W +W      +  G    +
Sbjct  1065  IQAPATGFGTSIFSNINHVIQNSQNF-GIPRHMRTVSPWLLWEIWKNRNKTLFQGTGLTS  1123

Query  283   LQVSVGSSNSVDLVFAAKDQ  224
              ++   +    +L   A+++
Sbjct  1124  SEIVAKAYEECNLWINAQEK  1143


>Ribonuclease H-like superfamily protein [Arabidopsis thaliana]
Sequence ID: NP_189164.1 Length: 343
>Ribonuclease H-like superfamily protein [Arabidopsis thaliana]
Sequence ID: AEE77003.1 Length: 343 >reverse transcriptase-like protein [Arabidopsis thaliana]
Sequence ID: BAB02086.1 Length: 343 >unnamed protein product [Arabidopsis thaliana]
Sequence ID: CAA0383633.1 Length: 343
Range 1: 25 to 152

Score:65.1 bits(157), Expect:7e-12,
Method:Compositional matrix adjust.,
Identities:47/135(35%), Positives:70/135(51%), Gaps:16/135(11%)

Query  631  PKTKFFLWKLMKGALPLGEHLRTRHINDTALCTLCGNE-ESVLHLFFKCPFAKQVWDLAP  455
            PK K FLWKL+ GAL  G++L+ RHI +   C  C  E E+  HLFF C +A+QVW
Sbjct  25   PKIKHFLWKLLSGALATGDNLKRRHIRNHPQCHRCCQEDETSQHLFFDCFYAQQVW----  80

Query  454  FKNSLSGDRIPSLR-AGIE-ASKLLICLPPTGLNAGP----LLPWIVWAI--SVSQMLYG  299
                 SG     LR  GI   +K+ + L     N  P    L  WI+W +  S +Q+++
Sbjct  81   ---RASGIPHQELRTTGITMETKMELLLSSCLANRQPQLFNLAIWILWRLWKSRNQLVFQ  137

Query  298  LKAPNLQVSVGSSNS  254
             K+ + Q ++  + +
Sbjct  138  QKSISWQNTLQRARN  152


>unnamed protein product [Arabidopsis thaliana]
Sequence ID: CAA0406188.1 Length: 1836
Range 1: 1526 to 1628

Score:65.9 bits(159), Expect:7e-12,
Method:Compositional matrix adjust.,
Identities:38/106(36%), Positives:60/106(56%), Gaps:6/106(5%)

Query  631   PKTKFFLWKLMKGALPLGEHLRTRHINDTALCTLCGNE-ESVLHLFFKCPFAKQVWDLAP  455
             PK K FLWK++KGA+ + + LRTR I     C +C  E E++ H+ F+CP A+QVW L+
Sbjct  1526  PKIKTFLWKVLKGAVAVEDRLRTRGIRIYDGCLMCKEENETINHILFQCPLARQVWALSL  1585

Query  454   FKNSLSGDRIPSLRAGIEASKLLICLPPTGLNAGPLL--PWIVWAI  323
              ++ ++G    S+ A ++   +L       LN       PWI+W +
Sbjct  1586  LQSPVNG-YGSSIFANMD--HVLHYCQSLSLNNNVRFGSPWIIWVL  1628


>hypothetical protein AXX17_AT2G11760 [Arabidopsis thaliana]
Sequence ID: OAP10255.1 Length: 1587
Range 1: 1013 to 1071

Score:65.5 bits(158), Expect:1e-11,
Method:Compositional matrix adjust.,
Identities:29/59(49%), Positives:39/59(66%), Gaps:1/59(1%)

Query  631   PKTKFFLWKLMKGALPLGEHLRTRHINDTALCTLCGNE-ESVLHLFFKCPFAKQVWDLA  458
             PK K F+WK +KGAL + + LR+R I     C  C  E E++ HL F+CPFA+QVW L+
Sbjct  1013  PKIKVFMWKALKGALAVEDRLRSRGIQTADGCLFCKEEIETINHLLFQCPFARQVWALS  1071


>reverse transcriptase, putative; 16838-20266 [Arabidopsis thaliana]
Sequence ID: AAG51783.1 Length: 1142
Range 1: 835 to 934

Score:63.2 bits(152), Expect:4e-11,
Method:Compositional matrix adjust.,
Identities:36/110(33%), Positives:59/110(53%), Gaps:17/110(15%)

Query  631  PKTKFFLWKLMKGALPLGEHLRTRHINDTALCTLCG-NEESVLHLFFKCPFAKQVWDLAP  455
            PK + FLW+++ G +P+ E+LR R I     C  CG +EES+ H  F+C  A+Q+W L+
Sbjct  835  PKLRHFLWQILSGCVPVSENLRKRGILCDKGCVSCGASEESINHTLFQCHPARQIWALS-  893

Query  454  FKNSLSGDRIPSLRAGIEASKLLICLP------PTGLNAGPLLPWIVWAI  323
                    +IP+      ++ +   L       P+G+++ P  PWI+W I
Sbjct  894  --------QIPTAPGIFPSNSIFTNLDHLFWRIPSGVDSAP-YPWIIWYI  934


>putative non-LTR retroelement reverse transcriptase [Arabidopsis thaliana]
Sequence ID: AAD15377.1 Length: 1044
Range 1: 801 to 850

Score:62.8 bits(151), Expect:6e-11,
Method:Compositional matrix adjust.,
Identities:26/50(52%), Positives:36/50(72%), Gaps:0/50(0%)

Query  478  KQVWDLAPFKNSLSGDRIPSLRAGIEASKLLICLPPTGLNAGPLLPWIVW  329
            ++VWDL+PFK +L   RI S++ G+E SKLL+ LPP G+  G L  WI+W
Sbjct  801  EKVWDLSPFKTTLQASRITSMKQGLEVSKLLVTLPPIGIGQGQLPIWILW  850


Range 2: 911 to 994

Score:62.8 bits(151), Expect:7e-11,
Method:Compositional matrix adjust.,
Identities:30/84(36%), Positives:49/84(58%), Gaps:0/84(0%)

Query  320  CFTDALWSESSKSAGFGWIIKFRRSGFRSKGSASDSHIRSLVVAEAIAIQLALKQSLRLG  141
            C TDA W E +  AGFGW+     +   S   A+  +IRS ++A+A A+ LA++ +  LG
Sbjct  911  CSTDASWREETLQAGFGWVFVDHSNHLESHHKAAAMNIRSPLLAKASALSLAIQHAADLG  970

Query  140  ITYLAIVSDAKQVIETINSETPSM  69
               L + SD++Q+++ +N E   M
Sbjct  971  FKKLVVASDSQQLVKVLNGEPHPM  994


>hypothetical protein AXX17_AT1G36150 [Arabidopsis thaliana]
Sequence ID: OAP15291.1 Length: 1513
Range 1: 1197 to 1257

Score:62.8 bits(151), Expect:7e-11,
Method:Compositional matrix adjust.,
Identities:26/61(43%), Positives:38/61(62%), Gaps:1/61(1%)

Query  631   PKTKFFLWKLMKGALPLGEHLRTRHINDTALCTLCGNE-ESVLHLFFKCPFAKQVWDLAP  455
             PK K F W+ +   LP+ ++L+ R +   A+C LCG E E+V HL F+C   K++WDL P
Sbjct  1197  PKLKHFWWRSLHNGLPVADNLKKRGLRVDAICQLCGEEDETVNHLLFQCKLTKEIWDLTP  1256

Query  454   F  452

Sbjct  1257  I  1257


>unnamed protein product [Arabidopsis thaliana]
Sequence ID: VYS68395.1 Length: 2315
Range 1: 1999 to 2059

Score:62.4 bits(150), Expect:9e-11,
Method:Compositional matrix adjust.,
Identities:26/61(43%), Positives:37/61(60%), Gaps:1/61(1%)

Query  631   PKTKFFLWKLMKGALPLGEHLRTRHINDTALCTLCGNE-ESVLHLFFKCPFAKQVWDLAP  455
             PK K F W+     LP+ ++L+ R +   A+C LCG E E+V HL F+C   K++WDL P
Sbjct  1999  PKVKHFWWRSFHNGLPVADNLKKRGLRVDAICQLCGEEDETVNHLLFQCKLTKEIWDLTP  2058

Query  454   F  452

Sbjct  2059  I  2059


>hypothetical protein AXX17_AT3G32490 [Arabidopsis thaliana]
Sequence ID: OAP03411.1 Length: 2176
Range 1: 527 to 593

Score:62.4 bits(150), Expect:1e-10,
Method:Compositional matrix adjust.,
Identities:29/67(43%), Positives:43/67(64%), Gaps:1/67(1%)

Query  631  PKTKFFLWKLMKGALPLGEHLRTRHINDTALCTLCGNE-ESVLHLFFKCPFAKQVWDLAP  455
            P  K FLWK +KGA+ + + LRTR I     C++C  E E++ H+ F+CP A+QVW L+
Sbjct  527  PNIKVFLWKAVKGAVAVEDRLRTRGILIEDGCSMCPEENETINHILFQCPLARQVWALSL  586

Query  454  FKNSLSG  434
             ++  SG
Sbjct  587  LQSPFSG  593


Range 2: 1699 to 1765

Score:62.4 bits(150), Expect:1e-10,
Method:Compositional matrix adjust.,
Identities:29/67(43%), Positives:43/67(64%), Gaps:1/67(1%)

Query  631   PKTKFFLWKLMKGALPLGEHLRTRHINDTALCTLCGNE-ESVLHLFFKCPFAKQVWDLAP  455
             P  K FLWK +KGA+ + + LRTR I     C++C  E E++ H+ F+CP A+QVW L+
Sbjct  1699  PNIKVFLWKAVKGAVAVEDRLRTRGILIEDGCSMCPEENETINHILFQCPLARQVWALSL  1758

Query  454   FKNSLSG  434
              ++  SG
Sbjct  1759  LQSPFSG  1765


>hypothetical protein At2g15750 [Arabidopsis thaliana]
Sequence ID: ABE65445.1 Length: 153
Range 1: 1 to 79

Score:59.3 bits(142), Expect:1e-10,
Method:Compositional matrix adjust.,
Identities:33/79(42%), Positives:48/79(60%), Gaps:6/79(7%)

Query  511  VLHLFF----KCPFAKQVWDLAPFKNSLSGDRIPSLRAGIEASKLLICLPPTGLNAGPLL  344
            +++LFF       FA+++W LAPFK+ L    I S+  GI +S   ICLPP+GL  G L
Sbjct  1    MMNLFFIFSSSVLFAQKIWSLAPFKSPLICASILSVTEGIASSSKAICLPPSGLGGGRLS  60

Query  343  PWIVWAI--SVSQMLYGLK  293
            PW+ W I  S +Q+++  K
Sbjct  61   PWLFWTIWSSRNQLIFNKK  79


>unknown, partial [Arabidopsis thaliana]
Sequence ID: ABK28185.1 Length: 154
Range 1: 1 to 79

Score:59.3 bits(142), Expect:1e-10,
Method:Compositional matrix adjust.,
Identities:33/79(42%), Positives:48/79(60%), Gaps:6/79(7%)

Query  511  VLHLFF----KCPFAKQVWDLAPFKNSLSGDRIPSLRAGIEASKLLICLPPTGLNAGPLL  344
            +++LFF       FA+++W LAPFK+ L    I S+  GI +S   ICLPP+GL  G L
Sbjct  1    MMNLFFIFSSSVLFAQKIWSLAPFKSPLICASILSVTEGIASSSKAICLPPSGLGGGRLS  60

Query  343  PWIVWAI--SVSQMLYGLK  293
            PW+ W I  S +Q+++  K
Sbjct  61   PWLFWTIWSSRNQLIFNKK  79


>hypothetical protein AXX17_AT1G43730 [Arabidopsis thaliana]
Sequence ID: OAP19763.1 Length: 1033
Range 1: 442 to 544

Score:62.0 bits(149), Expect:1e-10,
Method:Compositional matrix adjust.,
Identities:35/108(32%), Positives:60/108(55%), Gaps:10/108(9%)

Query  631  PKTKFFLWKLMKGALPLGEHLRTRHINDTALCTLCGNE-ESVLHLFFKCPFAKQVWDLAP  455
            PK K FLW+L+ G++ + + LRTR + +   C LCG+E E++ H+ F+C  ++QVW ++
Sbjct  442  PKIKNFLWRLLNGSIAVEDRLRTRGLKNADGCLLCGSESETINHVMFQCSLSRQVWAMSL  501

Query  454  FKNSLSGDRIPSLRAGIEASKLLICLPPTGLNAGPLL----PWIVWAI  323
              + L+G         +  +   +    T  +  PLL    PWI+WA+
Sbjct  502  VPSPLNG-----FGDSLFTNLAHLFDMSTNPSLSPLLREISPWILWAL  544


>hypothetical protein AXX17_AT3G37030 [Arabidopsis thaliana]
Sequence ID: OAP06308.1 Length: 2295
Range 1: 2144 to 2196

Score:62.0 bits(149), Expect:2e-10,
Method:Compositional matrix adjust.,
Identities:30/53(57%), Positives:34/53(64%), Gaps:0/53(0%)

Query  631   PKTKFFLWKLMKGALPLGEHLRTRHINDTALCTLCGNEESVLHLFFKCPFAKQ  473
             PK K FLWK MK ALPLGE L  R+I     C  CG+ E+ LHL   CPFAK+
Sbjct  2144  PKVKHFLWKAMKSALPLGEALARRNIIADLNCKRCGSPETSLHLLLLCPFAKK  2196


Range 2: 906 to 963

Score:61.6 bits(148), Expect:2e-10,
Method:Compositional matrix adjust.,
Identities:26/58(45%), Positives:39/58(67%), Gaps:1/58(1%)

Query  628  KTKFFLWKLMKGALPLGEHLRTRHINDTALCTLCGNE-ESVLHLFFKCPFAKQVWDLA  458
            K K FLWK++KGA+ + + LRTR I     C +C  E E++ H+ F+CP A+QVW ++
Sbjct  906  KIKVFLWKVLKGAVAVEDRLRTRGIKAVDGCLMCNEENETINHILFQCPLARQVWAMS  963


>putative non-LTR retroelement reverse transcriptase [Arabidopsis thaliana]
Sequence ID: AAD17395.1 Length: 1138
Range 1: 829 to 976

Score:61.6 bits(148), Expect:2e-10,
Method:Compositional matrix adjust.,
Identities:42/148(28%), Positives:67/148(45%), Gaps:4/148(2%)

Query  631  PKTKFFLWKLMKGALPLGEHLRTRHINDTALCTLCGNE-ESVLHLFFKCPFAKQVWDLAP  455
            PK K FLW+++  ALP+ + +  R ++    C +CG+E ES  H+ F C  A+QVW L+
Sbjct  829  PKIKMFLWRILSAALPVADQIIRRGMSIDPRCQICGDEGESTNHVLFTCSMARQVWALSG  888

Query  454  FKNSLSGDRIPSLRAGIE---ASKLLICLPPTGLNAGPLLPWIVWAISVSQMLYGLKAPN  284
                  G +  S+ A I+     K +I +P     + P + W +W         G+
Sbjct  889  VPTPEFGFQNASIFANIQFLFELKKMILVPDLVKRSWPWVLWRLWKNRNKLFFDGITFCP  948

Query  283  LQVSVGSSNSVDLVFAAKDQHRIPISDP  200
            L   V         F A+ Q R+  S+
Sbjct  949  LNSIVKIQEDTLEWFQAQSQIRVSESEE  976


>hypothetical protein AXX17_AT2G06510 [Arabidopsis thaliana]
Sequence ID: OAP07274.1 Length: 163
Range 1: 30 to 109

Score:59.3 bits(142), Expect:2e-10,
Method:Compositional matrix adjust.,
Identities:31/80(39%), Positives:46/80(57%), Gaps:0/80(0%)

Query  320  CFTDALWSESSKSAGFGWIIKFRRSGFRSKGSASDSHIRSLVVAEAIAIQLALKQSLRLG  141
            CFTDA W E S  A FGWI     +   S   +  ++I S ++AEA AI LA++ +  LG
Sbjct  30   CFTDASWREESLEARFGWIFVDHLNHAESHHQSVATNIGSPLLAEATAINLAIQNAADLG  89

Query  140  ITYLAIVSDAKQVIETINSE  81
               L I SD++ +++ +N E
Sbjct  90   FKKLFIASDSQMLVKALNGE  109


>Very similar to retrotransposon reverse transcriptase [Arabidopsis thaliana]
Sequence ID: AAF18538.1 Length: 1231
Range 1: 901 to 1038

Score:61.6 bits(148), Expect:2e-10,
Method:Compositional matrix adjust.,
Identities:45/147(31%), Positives:69/147(46%), Gaps:22/147(14%)

Query  631   PKTKFFLWKLMKGALPLGEHLRTRHINDTALCTLCGNE-ESVLHLFFKCPFAKQVWDLAP  455
             PK K FLWK++ GA+P+ + L  R +   + C  CG E ES+ H+ F C F +QVW
Sbjct  901   PKIKVFLWKVLSGAIPVVDLLSYRGMKLDSRCQTCGCEGESIQHVLFSCSFPRQVW----  956

Query  454   FKNSLSGDRIPSLRAGIEASKLLICLPPTGLNAGPL---------LPWIVWAISVSQMLY  302
                ++S   +P L  G E   +   L    +N   L          PWI+W I  ++ L+
Sbjct  957   ---AMSNIHVPLL--GFECGSVYANLYHFLINRDNLKWPVELRRSFPWIIWRIWKNRNLF  1011

Query  301   ---GLKAPNLQVSVGSSNSVDLVFAAK  230
                G +   L+  +     V+  FAA+
Sbjct  1012  FFEGKRFTVLETILKVRKDVEDWFAAQ  1038


>hypothetical protein AXX17_AT5G33430 [Arabidopsis thaliana]
Sequence ID: OAO95064.1 Length: 1347
Range 1: 1244 to 1321

Score:61.2 bits(147), Expect:2e-10,
Method:Compositional matrix adjust.,
Identities:30/78(38%), Positives:44/78(56%), Gaps:2/78(2%)

Query  631   PKTKFFLWKLMKGALPLGEHLRTRHINDTALCTLCGNE-ESVLHLFFKCPFAKQVWDLAP  455
             PK K F W  +   LP+ ++L+ R +   A+  LCG E E+V HL F+C   K++WDL P
Sbjct  1244  PKVKHFWWHSLHNGLPVADNLKKRGLRVDAIYQLCGEEDETVNHLLFQCKLTKEIWDLTP  1303

Query  454   FKNSLSGDRI-PSLRAGI  404
                   G  + P+ +AGI
Sbjct  1304  ISTPSDGSWLSPTQQAGI  1321


>Ribonuclease H-like superfamily protein [Arabidopsis thaliana]
Sequence ID: NP_194638.1 Length: 575
>putative reverse transcriptase/RNA-dependent DNA polymerase [Arabidopsis thaliana]
Sequence ID: AAY78807.1 Length: 575 >Ribonuclease H-like superfamily protein [Arabidopsis thaliana]
Sequence ID: AEE85585.1 Length: 575 >putative protein [Arabidopsis thaliana]
Sequence ID: CAB43923.1 Length: 575 >putative protein [Arabidopsis thaliana]
Sequence ID: CAB79667.1 Length: 575
Range 1: 264 to 367

Score:60.1 bits(144), Expect:6e-10,
Method:Compositional matrix adjust.,
Identities:33/104(32%), Positives:53/104(50%), Gaps:1/104(0%)

Query  631  PKTKFFLWKLMKGALPLGEHLRTRHINDTALCTLCGN-EESVLHLFFKCPFAKQVWDLAP  455
            PK + FLWK +  +LP+   L  RH++  + C  C + +E+V HL FKC FA+  W ++
Sbjct  264  PKIQHFLWKCLSNSLPVAGALAYRHLSKESACIRCPSCKETVNHLLFKCTFARLTWAISS  323

Query  454  FKNSLSGDRIPSLRAGIEASKLLICLPPTGLNAGPLLPWIVWAI  323
                L G+   S+   +     L    P    A  L+PW++W +
Sbjct  324  IPIPLGGEWADSIYVNLYWVFNLGNGNPQWEKASQLVPWLLWRL  367


>unnamed protein product [Arabidopsis thaliana]
Sequence ID: CAA0396872.1 Length: 575
Range 1: 264 to 367

Score:59.7 bits(143), Expect:6e-10,
Method:Compositional matrix adjust.,
Identities:33/104(32%), Positives:53/104(50%), Gaps:1/104(0%)

Query  631  PKTKFFLWKLMKGALPLGEHLRTRHINDTALCTLCGN-EESVLHLFFKCPFAKQVWDLAP  455
            PK + FLWK +  +LP+   L  RH++  + C  C + +E+V HL FKC FA+  W ++
Sbjct  264  PKIQHFLWKCLSNSLPVAGALAYRHLSKESACIRCPSCKETVNHLLFKCTFARLTWAISS  323

Query  454  FKNSLSGDRIPSLRAGIEASKLLICLPPTGLNAGPLLPWIVWAI  323
                L G+   S+   +     L    P    A  L+PW++W +
Sbjct  324  IPIPLGGEWADSIYVNLYWVFNLGNGNPQWEKASQLVPWLLWRL  367


>putative non-LTR retroelement reverse transcriptase [Arabidopsis thaliana]
Sequence ID: AAB82639.1 Length: 1374
Range 1: 1061 to 1164

Score:59.7 bits(143), Expect:9e-10,
Method:Compositional matrix adjust.,
Identities:32/104(31%), Positives:49/104(47%), Gaps:1/104(0%)

Query  631   PKTKFFLWKLMKGALPLGEHLRTRHINDTALCTLC-GNEESVLHLFFKCPFAKQVWDLAP  455
             PK   FLW+ +   L +  +L  RH+     C  C  + E+V HL FKCPFA+  W ++P
Sbjct  1061  PKIHHFLWRCVNNCLSVASNLAYRHLAREKSCVRCPSHGETVNHLLFKCPFARLTWAISP  1120

Query  454   FKNSLSGDRIPSLRAGIEASKLLICLPPTGLNAGPLLPWIVWAI  323
                   G+   SL   +     +    P   +   L+PWI+W +
Sbjct  1121  LPAPPGGEWAESLFRNMHHVLSVHKSQPEESDHHALIPWILWRL  1164


>reverse transcriptase [Arabidopsis thaliana]
Sequence ID: AAA75254.1 Length: 1333
Range 1: 1055 to 1158

Score:59.3 bits(142), Expect:1e-09,
Method:Compositional matrix adjust.,
Identities:36/107(34%), Positives:54/107(50%), Gaps:7/107(6%)

Query  631   PKTKFFLWKLMKGALPLGEHLRTRHINDTALCTLCGNE-ESVLHLFFKCPFAKQVWDLAP  455
             PK K FLW+++  ALP+   +  R +     C +CG E ES+ H+ F C  A+QVW L+
Sbjct  1055  PKIKLFLWRILSSALPVAYQIIRRGMPIDPRCQVCGEEGESINHVLFTCSLARQVWALSG  1114

Query  454   FKNSLSGDRIPSLRAGIEASKLLICLPPTGL---NAGPLLPWIVWAI  323
                S  G +  S+ A I+    L+ L   GL         PW++W +
Sbjct  1115  VPTSQFGFQNSSIFANIQ---YLLELKGKGLIPEQIKKSWPWVLWRL  1158


>RNA-directed DNA polymerase (reverse transcriptase)-related family protein [Arabidopsis thaliana]
Sequence ID: NP_683598.1 Length: 163
>RNA-directed DNA polymerase (reverse transcriptase)-related family protein [Arabidopsis thaliana]
Sequence ID: AEE77224.1 Length: 163
Range 1: 16 to 68

Score:56.6 bits(135), Expect:1e-09,
Method:Compositional matrix adjust.,
Identities:22/53(42%), Positives:34/53(64%), Gaps:0/53(0%)

Query  631  PKTKFFLWKLMKGALPLGEHLRTRHINDTALCTLCGNEESVLHLFFKCPFAKQ  473
            PK K  +WK +  ALP+G  L +R+I+    CT C + E++ H+ F CPFA++
Sbjct  16   PKIKLLIWKALNNALPVGAQLLSRNISIEPFCTRCRDFETITHILFNCPFAQR  68


>putative non-LTR retroelement reverse transcriptase [Arabidopsis thaliana]
Sequence ID: AAD17398.1 Length: 1225
Range 1: 934 to 1036

Score:56.2 bits(134), Expect:1e-08,
Method:Compositional matrix adjust.,
Identities:36/111(32%), Positives:51/111(45%), Gaps:17/111(15%)

Query  628   KTKFFLWKLMKGALPLGEHLRTRHINDTALCTLCG-NEESVLHLFFKCPFAKQVWDLAPF  452
             K K F W+ + G L   + L +RHI    +C  CG  EES+ HL F CP ++Q+W L+P
Sbjct  934   KFKHFEWQCLSGCLATNQRLFSRHIGTEKVCPRCGAEEESINHLLFLCPPSRQIWALSPI  993

Query  451   --------KNSLSGDRIPSLRAGIEASKLLICLPPTGLNAGPLLPWIVWAI  323
                     +NSL  +    L  G E             +   + PWI+W I
Sbjct  994   PSSEYIFPRNSLFYNFDFLLSRGKEFD--------IAEDIMEIFPWILWYI  1036


>unnamed protein product [Arabidopsis thaliana]
Sequence ID: CAA0247490.1 Length: 865
Range 1: 58 to 140

Score:56.2 bits(134), Expect:1e-08,
Method:Compositional matrix adjust.,
Identities:34/85(40%), Positives:48/85(56%), Gaps:7/85(8%)

Query  320  CFTDALWSESSKSAGFGWIIKFRRSGFRS-----KGSASDSHIRSLVVAEAIAIQLALKQ  156
            CFTDA W+    +AG GW   FR +  RS     +G ++ SH+ S + AEA AI  AL Q
Sbjct  58   CFTDAAWNADWSTAGLGWY--FRSNTDRSDLVTEQGVSTHSHVFSALEAEAWAILAALDQ  115

Query  155  SLRLGITYLAIVSDAKQVIETINSE  81
            +L LG   + + SD   ++  +NSE
Sbjct  116  ALVLGFEEVQVSSDCLTLVNLLNSE  140


>unnamed protein product [Arabidopsis thaliana]
Sequence ID: CAA0405280.1 Length: 423
Range 1: 168 to 227

Score:55.5 bits(132), Expect:2e-08,
Method:Compositional matrix adjust.,
Identities:24/60(40%), Positives:35/60(58%), Gaps:1/60(1%)

Query  631  PKTKFFLWKLMKGALPLGEHLRTRHINDTALCTLCGN-EESVLHLFFKCPFAKQVWDLAP  455
            PK K F WK +  ALP GE L+ R I    +C  CG  +E+  HL ++C  +K++W+  P
Sbjct  168  PKIKHFWWKSLHDALPTGEILKRRKIVSDDMCIQCGEAQENTNHLLYQCRVSKEIWEQNP  227


Query #109: XLOC_012587 Query ID: lcl|Query_33112 Length: 1090

Sequences producing significant alignments:
                                                                  Max    Total Query   E   Per.
Description                                                       Score  Score cover Value Ident  Accession
unnamed protein product [Arabidopsis thaliana]                    73.9   73.9  21%   6e-16 54.43  VYS52357.1
hypothetical protein AT2G14265 [Arabidopsis thaliana]             73.2   126   42%   2e-15 51.09  NP_001325174.1
transmembrane protein [Arabidopsis thaliana]                      67.0   122   22%   2e-13 57.14  NP_001326023.1
unnamed protein product [Arabidopsis thaliana]                    67.0   67.0  18%   2e-13 54.55  VYS52439.1
hypothetical protein AT1G62895 [Arabidopsis thaliana]             59.3   59.3  12%   1e-10 60.78  NP_001321703.1
unnamed protein product [Arabidopsis thaliana]                    56.2   56.2  11%   7e-10 72.09  VYS52323.1
unnamed protein product [Arabidopsis thaliana]                    56.2   111   22%   1e-09 57.14  CAA0384287.1
unnamed protein product [Arabidopsis thaliana]                    54.7   54.7  11%   3e-09 69.77  CAA0360812.1


Alignments:
>unnamed protein product [Arabidopsis thaliana]
Sequence ID: VYS52357.1 Length: 87
Range 1: 1 to 63

Score:73.9 bits(180), Expect:6e-16,
Method:Compositional matrix adjust.,
Identities:43/79(54%), Positives:47/79(59%), Gaps:16/79(20%)

Query  703  MCQTRFIYVASRFRLIEFPVIMDMIYKMDIEQ*KGFRMDIFKWTRYYGHQTIVC*FFFRF  524
            MCQTRF YVASRF LI+FPVIMDM                   ++ YGH T+V      F
Sbjct  1    MCQTRFTYVASRFPLIKFPVIMDMRPLP---------------SKPYGHGTLVS-LQNTF  44

Query  523  AETSTFKTLWTWNFGFFTK  467
             ETST K LWTWNFGFFTK
Sbjct  45   EETSTSKILWTWNFGFFTK  63


>hypothetical protein AT2G14265 [Arabidopsis thaliana]
Sequence ID: NP_001325174.1 Length: 111
>hypothetical protein AT2G14265 [Arabidopsis thaliana]
Sequence ID: ANM63061.1 Length: 111
Range 1: 29 to 107

Score:73.2 bits(178), Expect:2e-15,
Method:Compositional matrix adjust.,
Identities:47/92(51%), Positives:54/92(58%), Gaps:19/92(20%)

Query  740  QQWRLDLIFALSHVSNSFHLCSIEIPVNRISCYYGH----DIQNGYRTVKRFPNGYI*MD  573
            QQWRL+LI ALSHVSNSFHLCSIEIP    S  YGH     +QN   T    P  Y
Sbjct  29   QQWRLELISALSHVSNSFHLCSIEIPRPLPSKPYGHGTLVSLQN---TRPPLPRSY----  81

Query  572  SLLWTPNHCLLIFLS--VCRDLYLQNLMDMEL  483
                   H  L+ L   VCRDL+LQ+LMD++L
Sbjct  82   ------GHGTLVSLQKHVCRDLHLQDLMDIKL  107


Range 2: 50 to 107

Score:53.5 bits(127), Expect:2e-08,
Method:Compositional matrix adjust.,
Identities:37/84(44%), Positives:43/84(51%), Gaps:26/84(30%)

Query  534  SFGLQRPLPSKPYGHGTLVSLQNTVNRISCYY*HGI*NEYR*VERFLNGYI*MDSLLWTP  355
            S  + RPLPSKPYGHGTLVSLQNT   +   Y HG                   +L+
Sbjct  50   SIEIPRPLPSKPYGHGTLVSLQNTRPPLPRSYGHG-------------------TLVSLQ  90

Query  354  KHCLLIFLSV*RDLHFQDLMDMEL  283
            KH       V RDLH QDLMD++L
Sbjct  91   KH-------VCRDLHLQDLMDIKL  107


>transmembrane protein [Arabidopsis thaliana]
Sequence ID: NP_001326023.1 Length: 74
>transmembrane protein [Arabidopsis thaliana]
Sequence ID: ANM63965.1 Length: 74
Range 1: 12 to 74

Score:67.0 bits(162), Expect:2e-13,
Method:Compositional matrix adjust.,
Identities:36/63(57%), Positives:39/63(61%), Gaps:15/63(23%)

Query  330  SV*RDLHFQDLMDMELWFLYKPRLIES---------------QFAETPPPIPYGHRTFTK  196
            SV RDLH QDL DM+LW LYKPRLI+S               Q    PPP PYG+RTFTK
Sbjct  12   SVCRDLHLQDLTDMKLWSLYKPRLIKSPVIMDMRPPPPRPCGQLQRAPPPRPYGYRTFTK  71

Query  195  FLR  187
            FLR
Sbjct  72   FLR  74


Range 2: 12 to 44

Score:55.5 bits(132), Expect:2e-09,
Method:Compositional matrix adjust.,
Identities:25/33(76%), Positives:30/33(90%), Gaps:0/33(0%)

Query  530  SVCRDLYLQNLMDMELWFLYKTRLIESPVIIDM  432
            SVCRDL+LQ+L DM+LW LYK RLI+SPVI+DM
Sbjct  12   SVCRDLHLQDLTDMKLWSLYKPRLIKSPVIMDM  44


>unnamed protein product [Arabidopsis thaliana]
Sequence ID: VYS52439.1 Length: 86
Range 1: 3 to 68

Score:67.0 bits(162), Expect:2e-13,
Method:Compositional matrix adjust.,
Identities:36/66(55%), Positives:43/66(65%), Gaps:0/66(0%)

Query  788  F*YKFEIQLSKKHLLLQQWRLDLIFALSHVSNSFHLCSIEIPVNRISCYYGHDIQNGYRT  609
            F ++ +   SKK LLLQQWRLDLIFALSHVSNSFHL SIEIP       YGH+     +
Sbjct  3    FFFQLKDWFSKKLLLLQQWRLDLIFALSHVSNSFHLRSIEIPKPPPLRPYGHETLVSLQN  62

Query  608  VKRFPN  591
             +  P+
Sbjct  63   TRPPPS  68


>hypothetical protein AT1G62895 [Arabidopsis thaliana]
Sequence ID: NP_001321703.1 Length: 86
>hypothetical protein AT1G62895 [Arabidopsis thaliana]
Sequence ID: ANM59338.1 Length: 86
Range 1: 19 to 69

Score:59.3 bits(142), Expect:1e-10,
Method:Compositional matrix adjust.,
Identities:31/51(61%), Positives:33/51(64%), Gaps:4/51(7%)

Query  740  QQWRLDLIFALSHVSNSFHLCSIEIPVNRISCYYGHD----IQNGYRTVKR  600
            QQWRLDLIFALSHVSNSFHL SIEIP       YGH+    +QN      R
Sbjct  19   QQWRLDLIFALSHVSNSFHLRSIEIPKPPPLRPYGHETLVSLQNTRPPPSR  69


>unnamed protein product [Arabidopsis thaliana]
Sequence ID: VYS52323.1 Length: 61
Range 1: 24 to 61

Score:56.2 bits(134), Expect:7e-10,
Method:Compositional matrix adjust.,
Identities:31/43(72%), Positives:32/43(74%), Gaps:5/43(11%)

Query  320  ETSTSKTLWTWNFGFFTNHG*SNLNLQRPHLQYLMDIELLPNS  192
            ETSTSK LWTWNFGFFTNH   +L LQ      LMDIELLPNS
Sbjct  24   ETSTSKALWTWNFGFFTNHVCKDLRLQD-----LMDIELLPNS  61


>unnamed protein product [Arabidopsis thaliana]
Sequence ID: CAA0384287.1 Length: 74
>unnamed protein product [Arabidopsis thaliana]
Sequence ID: VYS59150.1 Length: 74
Range 1: 12 to 74

Score:56.2 bits(134), Expect:1e-09,
Method:Compositional matrix adjust.,
Identities:36/63(57%), Positives:39/63(61%), Gaps:15/63(23%)

Query  330  SV*RDLHFQDLMDMELWFLYKPRLIES---------------QFAETPPPIPYGHRTFTK  196
            SV RDLH QDL DM+LW LYKPRLI+S               Q    PPP PYG+RTFTK
Sbjct  12   SVCRDLHLQDLTDMKLWSLYKPRLIKSPVIMDMRPPPPRPCGQLQRPPPPRPYGYRTFTK  71

Query  195  FLR  187
            FLR
Sbjct  72   FLR  74


Range 2: 12 to 44

Score:55.1 bits(131), Expect:3e-09,
Method:Compositional matrix adjust.,
Identities:25/33(76%), Positives:30/33(90%), Gaps:0/33(0%)

Query  530  SVCRDLYLQNLMDMELWFLYKTRLIESPVIIDM  432
            SVCRDL+LQ+L DM+LW LYK RLI+SPVI+DM
Sbjct  12   SVCRDLHLQDLTDMKLWSLYKPRLIKSPVIMDM  44


>unnamed protein product [Arabidopsis thaliana]
Sequence ID: CAA0360812.1 Length: 61
Range 1: 24 to 61

Score:54.7 bits(130), Expect:3e-09,
Method:Compositional matrix adjust.,
Identities:30/43(70%), Positives:31/43(72%), Gaps:5/43(11%)

Query  320  ETSTSKTLWTWNFGFFTNHG*SNLNLQRPHLQYLMDIELLPNS  192
            ETSTSK LWTWNFGFFT H   +L LQ      LMDIELLPNS
Sbjct  24   ETSTSKALWTWNFGFFTKHVCKDLRLQD-----LMDIELLPNS  61


Query #110: XLOC_013234 Query ID: lcl|Query_33113 Length: 617

No significant similarity found.


Query #111: XLOC_013585 Query ID: lcl|Query_33114 Length: 698

No significant similarity found.


Query #112: XLOC_013586 Query ID: lcl|Query_33115 Length: 485

Sequences producing significant alignments:
                                                                  Max    Total Query   E   Per.
Description                                                       Score  Score cover Value Ident  Accession
unnamed protein product [Arabidopsis thaliana]                    70.5   70.5  51%   5e-16 48.81  CAA0405760.1
Mannose-binding lectin superfamily protein [Arabidopsis thaliana] 75.5   75.5  26%   7e-16 74.42  NP_175623.2
serine carboxypeptidase [Arabidopsis thaliana]                    73.9   73.9  35%   2e-15 62.71  BAA96893.1
unnamed protein product [Arabidopsis thaliana]                    65.5   65.5  25%   3e-12 70.73  CAA0288322.1


Alignments:
>unnamed protein product [Arabidopsis thaliana]
Sequence ID: CAA0405760.1 Length: 84
Range 1: 1 to 84

Score:70.5 bits(171), Expect:5e-16,
Method:Compositional matrix adjust.,
Identities:41/84(49%), Positives:50/84(59%), Gaps:0/84(0%)

Query  7    MVDPGFINASTPRGIGFESQLIQFISRFVD*WRQVCRRSTMMYNYGRSPGFYAEKILKVS  186
            MVD GFINASTPRG G   +    ++       +  +            GFYAEKILKVS
Sbjct  1    MVDQGFINASTPRGRGSSLRYCDLLADLQINGGKCAKDLQRCTIIAVHLGFYAEKILKVS  60

Query  187  IKRVENKKEVSSVVDLIRVERNCP  258
            IKR+E + +VSSV+DLIRV RN P
Sbjct  61   IKRMEKENDVSSVMDLIRVGRNYP  84


>Mannose-binding lectin superfamily protein [Arabidopsis thaliana]
Sequence ID: NP_175623.2 Length: 615
>Mannose-binding lectin superfamily protein [Arabidopsis thaliana]
Sequence ID: AEE32756.1 Length: 615
Range 1: 1 to 43

Score:75.5 bits(184), Expect:7e-16,
Method:Composition-based stats.,
Identities:32/43(74%), Positives:39/43(90%), Gaps:0/43(0%)

Query  130  MYNYGRSPGFYAEKILKVSIKRVENKKEVSSVVDLIRVERNCP  258
            MYNY RSPGFYA+KILK+SIKR++ K E+SSVVDLI++ RNCP
Sbjct  1    MYNYSRSPGFYAKKILKMSIKRMKKKNEMSSVVDLIKIGRNCP  43


>serine carboxypeptidase [Arabidopsis thaliana]
Sequence ID: BAA96893.1 Length: 512
Range 1: 442 to 500

Score:73.9 bits(180), Expect:2e-15,
Method:Composition-based stats.,
Identities:37/59(63%), Positives:45/59(76%), Gaps:1/59(1%)

Query  2    AQWLTQGSLMLLHQEV*GSSPS*FNLLADL*INGGKCAEDLQ*CTIM-AVRQDSTQRRY  175
             QWLT+GSLMLL Q+V GSSP+  +LL DL INGGKCA DLQ CTIM  +R+  T+R +
Sbjct  442  GQWLTKGSLMLLQQDVWGSSPNYCDLLVDLQINGGKCAGDLQRCTIMDLIRKKKTKRSF  500


>unnamed protein product [Arabidopsis thaliana]
Sequence ID: CAA0288322.1 Length: 614
Range 1: 1 to 41

Score:65.5 bits(158), Expect:3e-12,
Method:Composition-based stats.,
Identities:29/41(71%), Positives:36/41(87%), Gaps:0/41(0%)

Query  130  MYNYGRSPGFYAEKILKVSIKRVENKKEVSSVVDLIRVERN  252
            MYNY  SPGFYA+KILK+SIKR++ K E+SSVVDLI++ RN
Sbjct  1    MYNYSHSPGFYAKKILKMSIKRMKKKNEMSSVVDLIKIGRN  41


Query #113: XLOC_013587 Query ID: lcl|Query_33116 Length: 1157

No significant similarity found.


Query #114: XLOC_013588 Query ID: lcl|Query_33117 Length: 803

Sequences producing significant alignments:
                                                                  Max    Total Query   E   Per.
Description                                                       Score  Score cover Value Ident  Accession
unnamed protein product [Arabidopsis thaliana]                    46.2   77.4  16%   4e-08 82.61  VYS47722.1


Alignments:
>unnamed protein product [Arabidopsis thaliana]
Sequence ID: VYS47722.1 Length: 424
Range 1: 396 to 418

Score:46.2 bits(108), Expect:4e-08,
Method:Compositional matrix adjust.,
Identities:19/23(83%), Positives:22/23(95%), Gaps:0/23(0%)

Query  501  VRRSMRHVREALKTCYYPYKSPH  433
            VRRS+RH++EALKTCY PYKSPH
Sbjct  396  VRRSVRHIQEALKTCYKPYKSPH  418


Range 2: 374 to 394

Score:31.2 bits(69), Expect:4e-08,
Method:Compositional matrix adjust.,
Identities:13/21(62%), Positives:13/21(61%), Gaps:0/21(0%)

Query  566  CNAPTAPTSGPHFLSRVHGPT  504
            CNAP  P SG   LSR HG T
Sbjct  374  CNAPIVPISGSRVLSRAHGST  394


Query #115: XLOC_013590 Query ID: lcl|Query_33118 Length: 524

No significant similarity found.


Query #116: XLOC_013591 Query ID: lcl|Query_33119 Length: 806

No significant similarity found.


Query #117: XLOC_013592 Query ID: lcl|Query_33120 Length: 943

No significant similarity found.


Query #118: XLOC_013593 Query ID: lcl|Query_33121 Length: 2236

No significant similarity found.


Query #119: XLOC_013594 Query ID: lcl|Query_33122 Length: 674

No significant similarity found.


Query #120: XLOC_013595 Query ID: lcl|Query_33123 Length: 1519

Sequences producing significant alignments:
                                                                  Max    Total Query   E   Per.
Description                                                       Score  Score cover Value Ident  Accession
putative protein [Arabidopsis thaliana]                           49.3   85.5  8%    5e-10 88.00  CAB39942.1
hypothetical protein [Arabidopsis thaliana]                       47.0   85.1  8%    5e-10 72.41  AAG51098.1
non-LTR retroelement reverse transcriptase-like protein...        43.5   77.8  8%    8e-08 78.26  BAB08270.1


Alignments:
>putative protein [Arabidopsis thaliana]
Sequence ID: CAB39942.1 Length: 473
>putative protein [Arabidopsis thaliana]
Sequence ID: CAB78214.1 Length: 473
Range 1: 411 to 435

Score:49.3 bits(116), Expect:5e-10,
Method:Compositional matrix adjust.,
Identities:22/25(88%), Positives:22/25(88%), Gaps:0/25(0%)

Query  77   FAVTIYTLWRERNERKHGASPNPSS  3
              VTIYTLWRERNERKHGASPN SS
Sbjct  411  LQVTIYTLWRERNERKHGASPNSSS  435


Range 2: 395 to 412

Score:36.2 bits(82), Expect:5e-10,
Method:Compositional matrix adjust.,
Identities:15/18(83%), Positives:15/18(83%), Gaps:0/18(0%)

Query  124  SNNWPDRSAGFLTRCILQ  71
            S NWPDR AGFL RCILQ
Sbjct  395  SRNWPDRIAGFLARCILQ  412


>hypothetical protein [Arabidopsis thaliana]
Sequence ID: AAG51098.1 Length: 504
Range 1: 470 to 498

Score:47.0 bits(110), Expect:5e-10,
Method:Compositional matrix adjust.,
Identities:21/29(72%), Positives:23/29(79%), Gaps:0/29(0%)

Query  87   PVAFCSDDLYSMARAKRAQTWGFPEPFKS  1
            P+ F  DDL+SMARAKRAQTW F E FKS
Sbjct  470  PLHFAGDDLHSMARAKRAQTWSFSEFFKS  498


Range 2: 456 to 474

Score:38.1 bits(87), Expect:5e-10,
Method:Compositional matrix adjust.,
Identities:15/19(79%), Positives:17/19(89%), Gaps:0/19(0%)

Query  128  KLKQLAGSQCRFPNPLHFA  72
            +L QLAGS CRFP+PLHFA
Sbjct  456  RLTQLAGSHCRFPSPLHFA  474


>non-LTR retroelement reverse transcriptase-like protein [Arabidopsis thaliana]
Sequence ID: BAB08270.1 Length: 489
Range 1: 428 to 450

Score:43.5 bits(101), Expect:8e-08,
Method:Compositional matrix adjust.,
Identities:18/23(78%), Positives:22/23(95%), Gaps:0/23(0%)

Query  71   VTIYTLWRERNERKHGASPNPSS  3
            V+I+T+WRERNERKHGAS NP+S
Sbjct  428  VSIHTIWRERNERKHGASLNPAS  450


Range 2: 410 to 427

Score:34.3 bits(77), Expect:8e-08,
Method:Compositional matrix adjust.,
Identities:13/18(72%), Positives:15/18(83%), Gaps:0/18(0%)

Query  124  SNNWPDRSAGFLTRCILQ  71
            S NW DR+AGFL RC+LQ
Sbjct  410  STNWHDRTAGFLARCLLQ  427


Query #121: XLOC_013596 Query ID: lcl|Query_33124 Length: 856

Sequences producing significant alignments:
                                                                  Max    Total Query   E   Per.
Description                                                       Score  Score cover Value Ident  Accession
hypothetical protein AXX17_AT2G11110 [Arabidopsis thaliana]       97.1   218   49%   2e-24 63.41  OAP09910.1


Alignments:
>hypothetical protein AXX17_AT2G11110 [Arabidopsis thaliana]
Sequence ID: OAP09910.1 Length: 137
Range 1: 1 to 57

Score:97.1 bits(240), Expect:2e-24,
Method:Compositional matrix adjust.,
Identities:52/82(63%), Positives:53/82(64%), Gaps:25/82(30%)

Query  216  MRNPSILGVCRIVCCCSVLWWMPMSFDHEWSVMLRYGPEPRPGMHDDA*FLLVFFVDSIR  395
            MRNPSILGVCRIVCCCSVLWWMPMSFDHEWSVMLR+
Sbjct  1    MRNPSILGVCRIVCCCSVLWWMPMSFDHEWSVMLRW------------------------  36

Query  396  CVGRQLTSPYQIQLATVCEEKQ  461
             VGRQLTSPYQIQLATV    Q
Sbjct  37   -VGRQLTSPYQIQLATVTRSNQ  57


Range 2: 72 to 111

Score:78.2 bits(191), Expect:4e-21,
Method:Compositional matrix adjust.,
Identities:38/40(95%), Positives:39/40(97%), Gaps:0/40(0%)

Query  736  SHLSQGELEFTYSPSKIEYLVLEETEHEILEVLRRITHLR  855
             HLSQGELEFTY+PSKIEYLVLEETEHEILEVLRRITHLR
Sbjct  72   CHLSQGELEFTYAPSKIEYLVLEETEHEILEVLRRITHLR  111


Range 3: 52 to 71

Score:43.5 bits(101), Expect:4e-21,
Method:Compositional matrix adjust.,
Identities:20/20(100%), Positives:20/20(100%), Gaps:0/20(0%)

Query  675  VTRSNQQVEGTRTLTPPVRK  734
            VTRSNQQVEGTRTLTPPVRK
Sbjct  52   VTRSNQQVEGTRTLTPPVRK  71


Query #122: XLOC_013597 Query ID: lcl|Query_33125 Length: 1363

No significant similarity found.


Query #123: XLOC_013598 Query ID: lcl|Query_33126 Length: 786

Sequences producing significant alignments:
                                                                  Max    Total Query   E   Per.
Description                                                       Score  Score cover Value Ident  Accession
unnamed protein product [Arabidopsis thaliana]                    72.0   72.0  18%   2e-13 77.55  CAA0395127.1
NOV [Arabidopsis thaliana]                                        71.2   71.2  18%   3e-13 75.51  OAP00397.1
unnamed protein product [Arabidopsis thaliana]                    71.2   71.2  18%   3e-13 75.51  VYS62565.1
Histidine kinase-, DNA gyrase B-, and HSP90-like ATPase family... 71.2   71.2  18%   3e-13 75.51  NP_193111.2
unnamed protein product [Arabidopsis thaliana]                    70.9   70.9  18%   4e-13 75.51  VYS62564.1
hypothetical protein [Arabidopsis thaliana]                       70.5   70.5  18%   4e-13 75.51  CAB36838.1
no vein-like protein [Arabidopsis thaliana]                       55.1   55.1  18%   7e-08 68.00  NP_001321617.1
hypothetical protein AXX17_AT1G08100 [Arabidopsis thaliana]       54.7   54.7  16%   7e-08 72.73  OAP13699.1
T23G18.16 [Arabidopsis thaliana]                                  54.7   54.7  16%   7e-08 72.73  AAF18242.1
no vein-like protein [Arabidopsis thaliana]                       54.7   54.7  18%   7e-08 68.00  NP_001321616.1
no vein-like protein [Arabidopsis thaliana]                       54.7   54.7  19%   7e-08 66.67  NP_172307.5
unnamed protein product [Arabidopsis thaliana]                    54.7   54.7  19%   8e-08 66.67  CAA0178752.1
unnamed protein product [Arabidopsis thaliana]                    54.7   54.7  18%   8e-08 68.00  CAA0178767.1


Alignments:
>unnamed protein product [Arabidopsis thaliana]
Sequence ID: CAA0395127.1 Length: 2845
Range 1: 481 to 529

Score:72.0 bits(175), Expect:2e-13,
Method:Compositional matrix adjust.,
Identities:38/49(78%), Positives:41/49(83%), Gaps:0/49(0%)

Query  503  KKRKG*RRNLEKSDSSKLLNRAPSK*QGHAKQEIPKSADDSDATRRFSM  357
            KKRKG RRN EKSDSSKLL + PSK QGHAKQEIPK ADDSDA + FS+
Sbjct  481  KKRKGERRNHEKSDSSKLLRKPPSKLQGHAKQEIPKLADDSDAKKVFSV  529


>NOV [Arabidopsis thaliana]
Sequence ID: OAP00397.1 Length: 2729
Range 1: 481 to 529

Score:71.2 bits(173), Expect:3e-13,
Method:Compositional matrix adjust.,
Identities:37/49(76%), Positives:42/49(85%), Gaps:0/49(0%)

Query  503  KKRKG*RRNLEKSDSSKLLNRAPSK*QGHAKQEIPKSADDSDATRRFSM  357
            KKRKG RRN EKSDSSKLL ++PSK QGHAKQEIPK ADDS+A + FS+
Sbjct  481  KKRKGERRNHEKSDSSKLLRKSPSKLQGHAKQEIPKLADDSEAKKVFSV  529


>unnamed protein product [Arabidopsis thaliana]
Sequence ID: VYS62565.1 Length: 2729
Range 1: 481 to 529

Score:71.2 bits(173), Expect:3e-13,
Method:Compositional matrix adjust.,
Identities:37/49(76%), Positives:42/49(85%), Gaps:0/49(0%)

Query  503  KKRKG*RRNLEKSDSSKLLNRAPSK*QGHAKQEIPKSADDSDATRRFSM  357
            KKRKG RRN EKSDSSKLL ++PSK QGHAKQEIPK ADDS+A + FS+
Sbjct  481  KKRKGERRNHEKSDSSKLLRKSPSKLQGHAKQEIPKLADDSEAKKVFSV  529


>Histidine kinase-, DNA gyrase B-, and HSP90-like ATPase family protein [Arabidopsis thaliana]
Sequence ID: NP_193111.2 Length: 2729
>RecName: Full=Protein NO VEIN; AltName: Full=Protein EMBRYO DEFECTIVE 2597 [Arabidopsis thaliana]
Sequence ID: F4JTS8.1 Length: 2729 >Histidine kinase-, DNA gyrase B-, and HSP90-like ATPase family protein [Arabidopsis thaliana]
Sequence ID: AEE83321.1 Length: 2729
Range 1: 481 to 529

Score:71.2 bits(173), Expect:3e-13,
Method:Compositional matrix adjust.,
Identities:37/49(76%), Positives:42/49(85%), Gaps:0/49(0%)

Query  503  KKRKG*RRNLEKSDSSKLLNRAPSK*QGHAKQEIPKSADDSDATRRFSM  357
            KKRKG RRN EKSDSSKLL ++PSK QGHAKQEIPK ADDS+A + FS+
Sbjct  481  KKRKGERRNHEKSDSSKLLRKSPSKLQGHAKQEIPKLADDSEAKKVFSV  529


>unnamed protein product [Arabidopsis thaliana]
Sequence ID: VYS62564.1 Length: 3000
Range 1: 481 to 529

Score:70.9 bits(172), Expect:4e-13,
Method:Compositional matrix adjust.,
Identities:37/49(76%), Positives:42/49(85%), Gaps:0/49(0%)

Query  503  KKRKG*RRNLEKSDSSKLLNRAPSK*QGHAKQEIPKSADDSDATRRFSM  357
            KKRKG RRN EKSDSSKLL ++PSK QGHAKQEIPK ADDS+A + FS+
Sbjct  481  KKRKGERRNHEKSDSSKLLRKSPSKLQGHAKQEIPKLADDSEAKKVFSV  529


>hypothetical protein [Arabidopsis thaliana]
Sequence ID: CAB36838.1 Length: 605
>hypothetical protein [Arabidopsis thaliana]
Sequence ID: CAB78416.1 Length: 605
Range 1: 481 to 529

Score:70.5 bits(171), Expect:4e-13,
Method:Compositional matrix adjust.,
Identities:37/49(76%), Positives:42/49(85%), Gaps:0/49(0%)

Query  503  KKRKG*RRNLEKSDSSKLLNRAPSK*QGHAKQEIPKSADDSDATRRFSM  357
            KKRKG RRN EKSDSSKLL ++PSK QGHAKQEIPK ADDS+A + FS+
Sbjct  481  KKRKGERRNHEKSDSSKLLRKSPSKLQGHAKQEIPKLADDSEAKKVFSV  529


>no vein-like protein [Arabidopsis thaliana]
Sequence ID: NP_001321617.1 Length: 656
>RecName: Full=Protein NO VEIN-LIKE [Arabidopsis thaliana]
Sequence ID: A0A1P8ARG1.1 Length: 656 >no vein-like protein [Arabidopsis thaliana]
Sequence ID: ANM59243.1 Length: 656
Range 1: 337 to 385

Score:55.1 bits(131), Expect:7e-08,
Method:Compositional matrix adjust.,
Identities:34/50(68%), Positives:38/50(76%), Gaps:2/50(4%)

Query  503  KKRKG*RRNLEKSDSSKLLNRAPSK*Q-GHAKQEIPKSADDSDATRRFSM  357
            KKRK   RN EKSDS KLL R PSK + GH KQ+IPKSADDSDA + FS+
Sbjct  337  KKRKAESRNHEKSDSPKLLRRGPSKLRRGHVKQKIPKSADDSDA-QIFSV  385


>hypothetical protein AXX17_AT1G08100 [Arabidopsis thaliana]
Sequence ID: OAP13699.1 Length: 477
Range 1: 359 to 402

Score:54.7 bits(130), Expect:7e-08,
Method:Compositional matrix adjust.,
Identities:32/44(73%), Positives:34/44(77%), Gaps:1/44(2%)

Query  503  KKRKG*RRNLEKSDSSKLLNRAPSK*Q-GHAKQEIPKSADDSDA  375
            KKRK   RN EKSDS KLL R PSK + GH KQ+IPKSADDSDA
Sbjct  359  KKRKAESRNHEKSDSPKLLRRGPSKLRRGHVKQKIPKSADDSDA  402


>T23G18.16 [Arabidopsis thaliana]
Sequence ID: AAF18242.1 Length: 512
Range 1: 394 to 437

Score:54.7 bits(130), Expect:7e-08,
Method:Compositional matrix adjust.,
Identities:32/44(73%), Positives:34/44(77%), Gaps:1/44(2%)

Query  503  KKRKG*RRNLEKSDSSKLLNRAPSK*Q-GHAKQEIPKSADDSDA  375
            KKRK   RN EKSDS KLL R PSK + GH KQ+IPKSADDSDA
Sbjct  394  KKRKAESRNHEKSDSPKLLRRGPSKLRRGHVKQKIPKSADDSDA  437


>no vein-like protein [Arabidopsis thaliana]
Sequence ID: NP_001321616.1 Length: 618
>no vein-like protein [Arabidopsis thaliana]
Sequence ID: ANM59242.1 Length: 618
Range 1: 337 to 385

Score:54.7 bits(130), Expect:7e-08,
Method:Compositional matrix adjust.,
Identities:34/50(68%), Positives:38/50(76%), Gaps:2/50(4%)

Query  503  KKRKG*RRNLEKSDSSKLLNRAPSK*Q-GHAKQEIPKSADDSDATRRFSM  357
            KKRK   RN EKSDS KLL R PSK + GH KQ+IPKSADDSDA + FS+
Sbjct  337  KKRKAESRNHEKSDSPKLLRRGPSKLRRGHVKQKIPKSADDSDA-QIFSV  385


>no vein-like protein [Arabidopsis thaliana]
Sequence ID: NP_172307.5 Length: 746
>no vein-like protein [Arabidopsis thaliana]
Sequence ID: AEE28271.1 Length: 746
Range 1: 391 to 440

Score:54.7 bits(130), Expect:7e-08,
Method:Compositional matrix adjust.,
Identities:34/51(67%), Positives:39/51(76%), Gaps:2/51(3%)

Query  503  KKRKG*RRNLEKSDSSKLLNRAPSK*Q-GHAKQEIPKSADDSDATRRFSMS  354
            KKRK   RN EKSDS KLL R PSK + GH KQ+IPKSADDSDA + FS++
Sbjct  391  KKRKAESRNHEKSDSPKLLRRGPSKLRRGHVKQKIPKSADDSDA-QIFSVN  440


>unnamed protein product [Arabidopsis thaliana]
Sequence ID: CAA0178752.1 Length: 746
Range 1: 391 to 440

Score:54.7 bits(130), Expect:8e-08,
Method:Compositional matrix adjust.,
Identities:34/51(67%), Positives:39/51(76%), Gaps:2/51(3%)

Query  503  KKRKG*RRNLEKSDSSKLLNRAPSK*Q-GHAKQEIPKSADDSDATRRFSMS  354
            KKRK   RN EKSDS KLL R PSK + GH KQ+IPKSADDSDA + FS++
Sbjct  391  KKRKAESRNHEKSDSPKLLRRGPSKLRRGHVKQKIPKSADDSDA-QIFSVN  440


>unnamed protein product [Arabidopsis thaliana]
Sequence ID: CAA0178767.1 Length: 954
Range 1: 337 to 385

Score:54.7 bits(130), Expect:8e-08,
Method:Compositional matrix adjust.,
Identities:34/50(68%), Positives:38/50(76%), Gaps:2/50(4%)

Query  503  KKRKG*RRNLEKSDSSKLLNRAPSK*Q-GHAKQEIPKSADDSDATRRFSM  357
            KKRK   RN EKSDS KLL R PSK + GH KQ+IPKSADDSDA + FS+
Sbjct  337  KKRKAESRNHEKSDSPKLLRRGPSKLRRGHVKQKIPKSADDSDA-QIFSV  385


Query #124: XLOC_013599 Query ID: lcl|Query_33127 Length: 668

No significant similarity found.


Query #125: XLOC_013600 Query ID: lcl|Query_33128 Length: 277

No significant similarity found.


Query #126: XLOC_013601 Query ID: lcl|Query_33129 Length: 705

No significant similarity found.


Query #127: XLOC_013602 Query ID: lcl|Query_33130 Length: 736

No significant similarity found.


Query #128: XLOC_013603 Query ID: lcl|Query_33131 Length: 670

No significant similarity found.


Query #129: XLOC_013604 Query ID: lcl|Query_33132 Length: 608

No significant similarity found.


Query #130: XLOC_013605 Query ID: lcl|Query_33133 Length: 501

No significant similarity found.


Query #131: XLOC_013745 Query ID: lcl|Query_33134 Length: 750

No significant similarity found.


Query #132: XLOC_014301 Query ID: lcl|Query_33135 Length: 390

Sequences producing significant alignments:
                                                                  Max    Total Query   E   Per.
Description                                                       Score  Score cover Value Ident  Accession
unnamed protein product [Arabidopsis thaliana]                    106    106   37%   1e-30 100.00 VYS57367.1
hypothetical protein AT3G14467 [Arabidopsis thaliana]             97.1   97.1  34%   4e-27 97.78  NP_001327719.1
unnamed protein product [Arabidopsis thaliana]                    97.1   97.1  34%   5e-27 97.78  CAA0382420.1


Alignments:
>unnamed protein product [Arabidopsis thaliana]
Sequence ID: VYS57367.1 Length: 84
Range 1: 36 to 84

Score:106 bits(264), Expect:1e-30,
Method:Compositional matrix adjust.,
Identities:49/49(100%), Positives:49/49(100%), Gaps:0/49(0%)

Query  184  RTFVQHLDQTRKPKRVWLHNIAIFYVPKKVQGKKWSCQSLGSTNKYHMT  330
            RTFVQHLDQTRKPKRVWLHNIAIFYVPKKVQGKKWSCQSLGSTNKYHMT
Sbjct  36   RTFVQHLDQTRKPKRVWLHNIAIFYVPKKVQGKKWSCQSLGSTNKYHMT  84


>hypothetical protein AT3G14467, partial [Arabidopsis thaliana]
Sequence ID: NP_001327719.1 Length: 63
>hypothetical protein AT3G14467, partial [Arabidopsis thaliana]
Sequence ID: ANM65773.1 Length: 63
Range 1: 19 to 63

Score:97.1 bits(240), Expect:4e-27,
Method:Compositional matrix adjust.,
Identities:44/45(98%), Positives:45/45(100%), Gaps:0/45(0%)

Query  196  QHLDQTRKPKRVWLHNIAIFYVPKKVQGKKWSCQSLGSTNKYHMT  330
            +HLDQTRKPKRVWLHNIAIFYVPKKVQGKKWSCQSLGSTNKYHMT
Sbjct  19   EHLDQTRKPKRVWLHNIAIFYVPKKVQGKKWSCQSLGSTNKYHMT  63


>unnamed protein product, partial [Arabidopsis thaliana]
Sequence ID: CAA0382420.1 Length: 63
Range 1: 19 to 63

Score:97.1 bits(240), Expect:5e-27,
Method:Compositional matrix adjust.,
Identities:44/45(98%), Positives:45/45(100%), Gaps:0/45(0%)

Query  196  QHLDQTRKPKRVWLHNIAIFYVPKKVQGKKWSCQSLGSTNKYHMT  330
            +HLDQTRKPKRVWLHNIAIFYVPKKVQGKKWSCQSLGSTNKYHMT
Sbjct  19   EHLDQTRKPKRVWLHNIAIFYVPKKVQGKKWSCQSLGSTNKYHMT  63


Query #133: XLOC_014708 Query ID: lcl|Query_33136 Length: 1002

Sequences producing significant alignments:
                                                                  Max    Total Query   E   Per.
Description                                                       Score  Score cover Value Ident  Accession
hypothetical protein AXX17_AT3G22760 [Arabidopsis thaliana]       144    144   20%   6e-43 100.00 OAP04024.1
hypothetical protein AXX17_AT3G22750 [Arabidopsis thaliana]       105    105   14%   3e-28 100.00 OAP06204.1
unnamed protein product [Arabidopsis thaliana]                    108    108   14%   1e-27 100.00 BAB01719.1
eukaryotic translation initiation factor 3C [Arabidopsis...       66.6   126   28%   3e-11 61.54  NP_001190102.1
eIF3c [Arabidopsis thaliana]                                      66.6   123   28%   3e-11 61.54  AAC83464.1
unnamed protein product [Arabidopsis thaliana]                    66.6   126   28%   3e-11 61.54  VYS60558.1
TIF3C1 [Arabidopsis thaliana]                                     66.6   126   28%   3e-11 61.54  OAP03790.1
unnamed protein product [Arabidopsis thaliana]                    66.6   126   28%   3e-11 61.54  CAA0386727.1
unnamed protein product [Arabidopsis thaliana]                    66.2   129   28%   4e-11 61.54  CAA0386725.1


Alignments:
>hypothetical protein AXX17_AT3G22760 [Arabidopsis thaliana]
Sequence ID: OAP04024.1 Length: 71
Range 1: 1 to 70

Score:144 bits(363), Expect:6e-43,
Method:Compositional matrix adjust.,
Identities:70/70(100%), Positives:70/70(100%), Gaps:0/70(0%)

Query  859  MTWSHNLSGGEVNACLSLTSSRRQKILATHLVFGIIRIAAMLGTSSIWHSQKINHFRPCS  680
            MTWSHNLSGGEVNACLSLTSSRRQKILATHLVFGIIRIAAMLGTSSIWHSQKINHFRPCS
Sbjct  1    MTWSHNLSGGEVNACLSLTSSRRQKILATHLVFGIIRIAAMLGTSSIWHSQKINHFRPCS  60

Query  679  SPSCHGQPQI  650
            SPSCHGQPQI
Sbjct  61   SPSCHGQPQI  70


>hypothetical protein AXX17_AT3G22750 [Arabidopsis thaliana]
Sequence ID: OAP06204.1 Length: 50
Range 1: 1 to 50

Score:105 bits(262), Expect:3e-28,
Method:Compositional matrix adjust.,
Identities:50/50(100%), Positives:50/50(100%), Gaps:0/50(0%)

Query  621  MELDQQNVQDICGCPWHEGELHGLKWLIFWLCHILEVPNMAAILMIPNTR  770
            MELDQQNVQDICGCPWHEGELHGLKWLIFWLCHILEVPNMAAILMIPNTR
Sbjct  1    MELDQQNVQDICGCPWHEGELHGLKWLIFWLCHILEVPNMAAILMIPNTR  50


>unnamed protein product [Arabidopsis thaliana]
Sequence ID: BAB01719.1 Length: 186
Range 1: 1 to 49

Score:108 bits(269), Expect:1e-27,
Method:Compositional matrix adjust.,
Identities:49/49(100%), Positives:49/49(100%), Gaps:0/49(0%)

Query  621  MELDQQNVQDICGCPWHEGELHGLKWLIFWLCHILEVPNMAAILMIPNT  767
            MELDQQNVQDICGCPWHEGELHGLKWLIFWLCHILEVPNMAAILMIPNT
Sbjct  1    MELDQQNVQDICGCPWHEGELHGLKWLIFWLCHILEVPNMAAILMIPNT  49


>eukaryotic translation initiation factor 3C [Arabidopsis thaliana]
Sequence ID: NP_001190102.1 Length: 900
>eukaryotic translation initiation factor 3C [Arabidopsis thaliana]
Sequence ID: NP_191174.1 Length: 900 >RecName: Full=Eukaryotic translation initiation factor 3 subunit C; Short=eIF3c; AltName: Full=Eukaryotic translation initiation factor 3 subunit 8; AltName: Full=eIF3 p110; AltName: Full=p105 [Arabidopsis thaliana]
Sequence ID: O49160.2 Length: 900 >putative eukaryotic translation initiation factor 3 subunit 8 [Arabidopsis thaliana]
Sequence ID: AAL36362.1 Length: 900 >putative eukaryotic translation initiation factor 3 subunit 8 [Arabidopsis thaliana]
Sequence ID: AAN71960.1 Length: 900 >eukaryotic translation initiation factor 3C [Arabidopsis thaliana]
Sequence ID: AEE79486.1 Length: 900 >eukaryotic translation initiation factor 3C [Arabidopsis thaliana]
Sequence ID: AEE79487.1 Length: 900 >PROBABLE EUKARYOTIC TRANSLATION INITIATION FACTOR 3 SUBUNIT 8 [Arabidopsis thaliana]
Sequence ID: CAB87414.1 Length: 900
Range 1: 623 to 674

Score:66.6 bits(161), Expect:3e-11,
Method:Composition-based stats.,
Identities:32/52(62%), Positives:39/52(75%), Gaps:0/52(0%)

Query  720  ILEVPNMAAILMIPNTR*VARIFCRLLEVSERQAFTSPPERLCDHVMTATRA  875
            +LEVPNMAA       R +++ F RLLE+SERQAFT+PPE + DHVM ATRA
Sbjct  623  LLEVPNMAANSHDAKRRVISKNFRRLLEISERQAFTAPPENVRDHVMAATRA  674


Range 2: 216 to 258

Score:60.1 bits(144), Expect:3e-09,
Method:Compositional matrix adjust.,
Identities:26/43(60%), Positives:34/43(79%), Gaps:0/43(0%)

Query  553  ERIMNKKDKFLHELYNKDPKEISWNWINKMFKIFVAAHGMKGN  681
            E++++KKDK L +L NKDPKEI+W+W+NK FK  VAA G KG
Sbjct  216  EKMLSKKDKLLEKLMNKDPKEITWDWVNKKFKEIVAARGKKGT  258


>eIF3c [Arabidopsis thaliana]
Sequence ID: AAC83464.1 Length: 900
Range 1: 623 to 674

Score:66.6 bits(161), Expect:3e-11,
Method:Composition-based stats.,
Identities:32/52(62%), Positives:39/52(75%), Gaps:0/52(0%)

Query  720  ILEVPNMAAILMIPNTR*VARIFCRLLEVSERQAFTSPPERLCDHVMTATRA  875
            +LEVPNMAA       R +++ F RLLE+SERQAFT+PPE + DHVM ATRA
Sbjct  623  LLEVPNMAANSHDAKRRVISKNFRRLLEISERQAFTAPPENVRDHVMAATRA  674


Range 2: 216 to 258

Score:57.0 bits(136), Expect:3e-08,
Method:Compositional matrix adjust.,
Identities:25/43(58%), Positives:33/43(76%), Gaps:0/43(0%)

Query  553  ERIMNKKDKFLHELYNKDPKEISWNWINKMFKIFVAAHGMKGN  681
            E++++KKDK L +L NKDPKEI+W+W+NK FK  VAA   KG
Sbjct  216  EKMLSKKDKLLEKLMNKDPKEITWDWVNKKFKEIVAAREKKGT  258


>unnamed protein product [Arabidopsis thaliana]
Sequence ID: VYS60558.1 Length: 900
Range 1: 623 to 674

Score:66.6 bits(161), Expect:3e-11,
Method:Composition-based stats.,
Identities:32/52(62%), Positives:39/52(75%), Gaps:0/52(0%)

Query  720  ILEVPNMAAILMIPNTR*VARIFCRLLEVSERQAFTSPPERLCDHVMTATRA  875
            +LEVPNMAA       R +++ F RLLE+SERQAFT+PPE + DHVM ATRA
Sbjct  623  LLEVPNMAANSHDAKRRVISKNFRRLLEISERQAFTAPPENVRDHVMAATRA  674


Range 2: 216 to 258

Score:60.1 bits(144), Expect:3e-09,
Method:Compositional matrix adjust.,
Identities:26/43(60%), Positives:34/43(79%), Gaps:0/43(0%)

Query  553  ERIMNKKDKFLHELYNKDPKEISWNWINKMFKIFVAAHGMKGN  681
            E++++KKDK L +L NKDPKEI+W+W+NK FK  VAA G KG
Sbjct  216  EKMLSKKDKLLEKLMNKDPKEITWDWVNKKFKEIVAARGKKGT  258


>TIF3C1 [Arabidopsis thaliana]
Sequence ID: OAP03790.1 Length: 901
Range 1: 624 to 675

Score:66.6 bits(161), Expect:3e-11,
Method:Composition-based stats.,
Identities:32/52(62%), Positives:39/52(75%), Gaps:0/52(0%)

Query  720  ILEVPNMAAILMIPNTR*VARIFCRLLEVSERQAFTSPPERLCDHVMTATRA  875
            +LEVPNMAA       R +++ F RLLE+SERQAFT+PPE + DHVM ATRA
Sbjct  624  LLEVPNMAANSHDAKRRVISKNFRRLLEISERQAFTAPPENVRDHVMAATRA  675


Range 2: 217 to 259

Score:60.1 bits(144), Expect:3e-09,
Method:Compositional matrix adjust.,
Identities:26/43(60%), Positives:34/43(79%), Gaps:0/43(0%)

Query  553  ERIMNKKDKFLHELYNKDPKEISWNWINKMFKIFVAAHGMKGN  681
            E++++KKDK L +L NKDPKEI+W+W+NK FK  VAA G KG
Sbjct  217  EKMLSKKDKLLEKLMNKDPKEITWDWVNKKFKEIVAARGKKGT  259


>unnamed protein product [Arabidopsis thaliana]
Sequence ID: CAA0386727.1 Length: 901
Range 1: 624 to 675

Score:66.6 bits(161), Expect:3e-11,
Method:Composition-based stats.,
Identities:32/52(62%), Positives:39/52(75%), Gaps:0/52(0%)

Query  720  ILEVPNMAAILMIPNTR*VARIFCRLLEVSERQAFTSPPERLCDHVMTATRA  875
            +LEVPNMAA       R +++ F RLLE+SERQAFT+PPE + DHVM ATRA
Sbjct  624  LLEVPNMAANSHDAKRRVISKNFRRLLEISERQAFTAPPENVRDHVMAATRA  675


Range 2: 217 to 259

Score:60.1 bits(144), Expect:3e-09,
Method:Compositional matrix adjust.,
Identities:26/43(60%), Positives:34/43(79%), Gaps:0/43(0%)

Query  553  ERIMNKKDKFLHELYNKDPKEISWNWINKMFKIFVAAHGMKGN  681
            E++++KKDK L +L NKDPKEI+W+W+NK FK  VAA G KG
Sbjct  217  EKMLSKKDKLLEKLMNKDPKEITWDWVNKKFKEIVAARGKKGT  259


>unnamed protein product [Arabidopsis thaliana]
Sequence ID: CAA0386725.1 Length: 1333
Range 1: 1056 to 1107

Score:66.2 bits(160), Expect:4e-11,
Method:Composition-based stats.,
Identities:32/52(62%), Positives:39/52(75%), Gaps:0/52(0%)

Query  720   ILEVPNMAAILMIPNTR*VARIFCRLLEVSERQAFTSPPERLCDHVMTATRA  875
             +LEVPNMAA       R +++ F RLLE+SERQAFT+PPE + DHVM ATRA
Sbjct  1056  LLEVPNMAANSHDAKRRVISKNFRRLLEISERQAFTAPPENVRDHVMAATRA  1107


Range 2: 649 to 690

Score:63.2 bits(152), Expect:3e-10,
Method:Composition-based stats.,
Identities:26/42(62%), Positives:34/42(80%), Gaps:0/42(0%)

Query  553  ERIMNKKDKFLHELYNKDPKEISWNWINKMFKIFVAAHGMKG  678
            E++++KKDK L +L NKDPKEI+W+W+NK FK  VAA G KG
Sbjct  649  EKMLSKKDKLLEKLMNKDPKEITWDWVNKKFKEIVAARGKKG  690


Query #134: XLOC_015062 Query ID: lcl|Query_33137 Length: 435

Sequences producing significant alignments:
                                                                  Max    Total Query   E   Per.
Description                                                       Score  Score cover Value Ident  Accession
unnamed protein product [Arabidopsis thaliana]                    83.2   129   71%   4e-24 54.17  BAB08713.1
hypothetical protein AXX17_AT5G32690 [Arabidopsis thaliana]       82.8   129   71%   6e-24 54.17  OAO94166.1
hypothetical protein AXX17_AT4G01860 [Arabidopsis thaliana]       68.2   102   67%   5e-16 50.00  OAO99509.1
F11O4.11 [Arabidopsis thaliana]                                   68.2   98.6  67%   7e-15 50.00  AAC62785.1
hypothetical protein AXX17_AT4G21170 [Arabidopsis thaliana]       64.3   64.3  56%   4e-12 41.46  OAO96867.1
putative non-LTR retroelement reverse transcriptase [Arabidops... 54.7   54.7  49%   8e-09 40.28  AAD22286.1
putative retroelement pol polyprotein [Arabidopsis thaliana]      53.5   53.5  52%   1e-08 39.47  AAM15063.1
hypothetical protein AXX17_AT3G15320 [Arabidopsis thaliana]       53.1   53.1  46%   2e-08 39.71  OAP05087.1
hypothetical protein AXX17_AT3G15320 [Arabidopsis thaliana]       53.1   53.1  46%   3e-08 39.71  OAP05088.1
unknown [Arabidopsis thaliana]                                    52.0   52.0  49%   6e-08 34.72  ABK28247.1
hypothetical protein At4g09700 [Arabidopsis thaliana]             52.0   52.0  49%   6e-08 34.72  ABE65517.1


Alignments:
>unnamed protein product [Arabidopsis thaliana]
Sequence ID: BAB08713.1 Length: 439
Range 1: 58 to 122

Score:83.2 bits(204), Expect:4e-24,
Method:Compositional matrix adjust.,
Identities:39/72(54%), Positives:50/72(69%), Gaps:7/72(9%)

Query  250  SLIPFLSDHWKTSSRLVGADLGNSLF*FQFANEPDIFANKPDLLKSLENQLFHFSQWMVI  71
            +++PFL+DHWK S+R  GADLGN LF FQFA E        DL ++L+ Q +HFSQWM+I
Sbjct  58   AVLPFLADHWKFSTRPTGADLGNGLFQFQFATE-------QDLARALKYQPYHFSQWMII  110

Query  70   LQRWEPEFPQGF  35
             QRWEP   + F
Sbjct  111  FQRWEPTIDKDF  122


Range 2: 26 to 56

Score:46.6 bits(109), Expect:4e-24,
Method:Compositional matrix adjust.,
Identities:22/31(71%), Positives:25/31(80%), Gaps:0/31(0%)

Query  345  VEVPGFDNSKLLRRHELTII*RLTNPAIEKV  253
            V+VP FDNS+LLRRH LT I RLTNPA + V
Sbjct  26   VKVPAFDNSELLRRHALTAIGRLTNPAEQDV  56


>hypothetical protein AXX17_AT5G32690 [Arabidopsis thaliana]
Sequence ID: OAO94166.1 Length: 1892
Range 1: 58 to 122

Score:82.8 bits(203), Expect:6e-24,
Method:Compositional matrix adjust.,
Identities:39/72(54%), Positives:50/72(69%), Gaps:7/72(9%)

Query  250  SLIPFLSDHWKTSSRLVGADLGNSLF*FQFANEPDIFANKPDLLKSLENQLFHFSQWMVI  71
            +++PFL+DHWK S+R  GADLGN LF FQFA E        DL ++L+ Q +HFSQWM+I
Sbjct  58   AVLPFLADHWKFSTRPTGADLGNGLFQFQFATE-------QDLARALKYQPYHFSQWMII  110

Query  70   LQRWEPEFPQGF  35
             QRWEP   + F
Sbjct  111  FQRWEPTIDKDF  122


Range 2: 26 to 56

Score:46.2 bits(108), Expect:6e-24,
Method:Compositional matrix adjust.,
Identities:22/31(71%), Positives:25/31(80%), Gaps:0/31(0%)

Query  345  VEVPGFDNSKLLRRHELTII*RLTNPAIEKV  253
            V+VP FDNS+LLRRH LT I RLTNPA + V
Sbjct  26   VKVPAFDNSELLRRHALTAIGRLTNPAEQDV  56


>hypothetical protein AXX17_AT4G01860 [Arabidopsis thaliana]
Sequence ID: OAO99509.1 Length: 560
Range 1: 91 to 149

Score:68.2 bits(165), Expect:5e-16,
Method:Compositional matrix adjust.,
Identities:33/66(50%), Positives:41/66(62%), Gaps:7/66(10%)

Query  250  SLIPFLSDHWKTSSRLVGADLGNSLF*FQFANEPDIFANKPDLLKSLENQLFHFSQWMVI  71
            SLIPFLS+ W    +  G+DLG   F F+F  E D       L K L+N+ +HF QWMVI
Sbjct  91   SLIPFLSNRWNLKGKATGSDLGRGCFQFRFEYEED-------LQKVLDNRPYHFGQWMVI  143

Query  70   LQRWEP  53
            LQRW+P
Sbjct  144  LQRWKP  149


Range 2: 58 to 89

Score:33.9 bits(76), Expect:5e-16,
Method:Compositional matrix adjust.,
Identities:13/32(41%), Positives:24/32(75%), Gaps:0/32(0%)

Query  348  RVEVPGFDNSKLLRRHELTII*RLTNPAIEKV  253
            R+  P  DNS+L++ + LT++ RLTNP+ +++
Sbjct  58   RIRAPTLDNSELIQENSLTLMGRLTNPSAQRL  89


>F11O4.11 [Arabidopsis thaliana]
Sequence ID: AAC62785.1 Length: 577
>putative transposon protein [Arabidopsis thaliana]
Sequence ID: CAB77719.1 Length: 577
Range 1: 54 to 112

Score:68.2 bits(165), Expect:7e-15,
Method:Compositional matrix adjust.,
Identities:33/66(50%), Positives:41/66(62%), Gaps:7/66(10%)

Query  250  SLIPFLSDHWKTSSRLVGADLGNSLF*FQFANEPDIFANKPDLLKSLENQLFHFSQWMVI  71
            SLIPFLS+ W    +  G+DLG   F F+F  E D       L K L+N+ +HF QWMVI
Sbjct  54   SLIPFLSNRWNLKGKATGSDLGRGCFQFRFEYEED-------LQKVLDNRPYHFGQWMVI  106

Query  70   LQRWEP  53
            LQRW+P
Sbjct  107  LQRWKP  112


Range 2: 21 to 52

Score:30.4 bits(67), Expect:7e-15,
Method:Compositional matrix adjust.,
Identities:12/32(38%), Positives:23/32(71%), Gaps:0/32(0%)

Query  348  RVEVPGFDNSKLLRRHELTII*RLTNPAIEKV  253
            R+  P  DNS+L++ + LT++  LTNP+ +++
Sbjct  21   RIRAPTLDNSELIQENSLTLMGILTNPSAQRL  52


>hypothetical protein AXX17_AT4G21170 [Arabidopsis thaliana]
Sequence ID: OAO96867.1 Length: 1816
Range 1: 49 to 122

Score:64.3 bits(155), Expect:4e-12,
Method:Compositional matrix adjust.,
Identities:34/82(41%), Positives:47/82(57%), Gaps:8/82(9%)

Query  280  THQSCNREGLSLIPFLSDHWKTSSRLVGADLGNSLF*FQFANEPDIFANKPDLLKSLENQ  101
            T+ +C R   +L PFLS+ W    + +G+DLG   F F+F  E D       LL  L+N+
Sbjct  49   TNPACQRL-WALFPFLSNRWNLKGKALGSDLGKGCFQFKFDFEED-------LLSVLKNR  100

Query  100  LFHFSQWMVILQRWEPEFPQGF  35
             +HF QWMVI+Q+WEP     F
Sbjct  101  PYHFDQWMVIIQKWEPIISDSF  122


>putative non-LTR retroelement reverse transcriptase [Arabidopsis thaliana]
Sequence ID: AAD22286.1 Length: 1311
Range 1: 58 to 122

Score:54.7 bits(130), Expect:8e-09,
Method:Compositional matrix adjust.,
Identities:29/72(40%), Positives:41/72(56%), Gaps:7/72(9%)

Query  250  SLIPFLSDHWKTSSRLVGADLGNSLF*FQFANEPDIFANKPDLLKSLENQLFHFSQWMVI  71
            +L  FLS+ W    +  G+DLG  +F  +F      F+   DL + L+N+ +HF QWMVI
Sbjct  58   ALFLFLSNRWTLRGKATGSDLGQGVFQLKFD-----FSE--DLQQVLDNRPYHFDQWMVI  110

Query  70   LQRWEPEFPQGF  35
            LQ+WEP     F
Sbjct  111  LQKWEPVISPSF  122


>putative retroelement pol polyprotein [Arabidopsis thaliana]
Sequence ID: AAM15063.1 Length: 267
>putative retroelement pol polyprotein [Arabidopsis thaliana]
Sequence ID: AAM15139.1 Length: 267
Range 1: 40 to 107

Score:53.5 bits(127), Expect:1e-08,
Method:Compositional matrix adjust.,
Identities:30/76(39%), Positives:45/76(59%), Gaps:8/76(10%)

Query  280  THQSCNREGLSLIPFLSDHWKTSSRLVGADLGNSLF*FQFANEPDIFANKPDLLKSLENQ  101
            T+QS  R   +L  FLS+ W    + +G+DLG  +F  +F         + DL + ++N+
Sbjct  40   TNQSGQRL-WALFLFLSNPWTLKGKAIGSDLGQGVFQLKFDF-------REDLQQVVDNR  91

Query  100  LFHFSQWMVILQRWEP  53
             +HF QWMVILQ+WEP
Sbjct  92   PYHFDQWMVILQKWEP  107


>hypothetical protein AXX17_AT3G15320 [Arabidopsis thaliana]
Sequence ID: OAP05087.1 Length: 1196
Range 1: 63 to 123

Score:53.1 bits(126), Expect:2e-08,
Method:Compositional matrix adjust.,
Identities:27/68(40%), Positives:36/68(52%), Gaps:7/68(10%)

Query  238  FLSDHWKTSSRLVGADLGNSLF*FQFANEPDIFANKPDLLKSLENQLFHFSQWMVILQRW  59
            ++   W   SR+VG DLG+  F F+F NE D+ A        L    +HF  WM+ILQ+W
Sbjct  63   YMPQLWNLESRVVGRDLGSECFQFRFENEADLQA-------VLRRGPYHFKNWMLILQQW  115

Query  58   EPEFPQGF  35
            EP     F
Sbjct  116  EPVISDTF  123


>hypothetical protein AXX17_AT3G15320 [Arabidopsis thaliana]
Sequence ID: OAP05088.1 Length: 1697
Range 1: 63 to 123

Score:53.1 bits(126), Expect:3e-08,
Method:Compositional matrix adjust.,
Identities:27/68(40%), Positives:36/68(52%), Gaps:7/68(10%)

Query  238  FLSDHWKTSSRLVGADLGNSLF*FQFANEPDIFANKPDLLKSLENQLFHFSQWMVILQRW  59
            ++   W   SR+VG DLG+  F F+F NE D+ A        L    +HF  WM+ILQ+W
Sbjct  63   YMPQLWNLESRVVGRDLGSECFQFRFENEADLQA-------VLRRGPYHFKNWMLILQQW  115

Query  58   EPEFPQGF  35
            EP     F
Sbjct  116  EPVISDTF  123


>unknown, partial [Arabidopsis thaliana]
Sequence ID: ABK28247.1 Length: 372
Range 1: 58 to 122

Score:52.0 bits(123), Expect:6e-08,
Method:Compositional matrix adjust.,
Identities:25/72(35%), Positives:39/72(54%), Gaps:7/72(9%)

Query  250  SLIPFLSDHWKTSSRLVGADLGNSLF*FQFANEPDIFANKPDLLKSLENQLFHFSQWMVI  71
            +L+ ++  +W   +R++G +LG   F F+F  E D       L   L+   +HF +WM I
Sbjct  58   ALVEYMVQYWNLENRVIGRELGPERFFFRFETEAD-------LQLVLKKAPYHFKKWMFI  110

Query  70   LQRWEPEFPQGF  35
            LQRWEP   + F
Sbjct  111  LQRWEPIVSEAF  122


>hypothetical protein At4g09700 [Arabidopsis thaliana]
Sequence ID: ABE65517.1 Length: 371
>putative protein [Arabidopsis thaliana]
Sequence ID: CAB39637.1 Length: 371 >putative protein [Arabidopsis thaliana]
Sequence ID: CAB78093.1 Length: 371
Range 1: 58 to 122

Score:52.0 bits(123), Expect:6e-08,
Method:Compositional matrix adjust.,
Identities:25/72(35%), Positives:39/72(54%), Gaps:7/72(9%)

Query  250  SLIPFLSDHWKTSSRLVGADLGNSLF*FQFANEPDIFANKPDLLKSLENQLFHFSQWMVI  71
            +L+ ++  +W   +R++G +LG   F F+F  E D       L   L+   +HF +WM I
Sbjct  58   ALVEYMVQYWNLENRVIGRELGPERFFFRFETEAD-------LQLVLKKAPYHFKKWMFI  110

Query  70   LQRWEPEFPQGF  35
            LQRWEP   + F
Sbjct  111  LQRWEPIVSEAF  122


Query #135: XLOC_015277 Query ID: lcl|Query_33138 Length: 373

No significant similarity found.


Query #136: XLOC_015413 Query ID: lcl|Query_33139 Length: 312

No significant similarity found.


Query #137: XLOC_015608 Query ID: lcl|Query_33140 Length: 843

Sequences producing significant alignments:
                                                                  Max    Total Query   E   Per.
Description                                                       Score  Score cover Value Ident  Accession
hypothetical protein AXX17_AT3G35670 [Arabidopsis thaliana]       186    186   31%   9e-60 100.00 OAP02798.1
GPI-anchored adhesin-like protein, putative (Protein of unknow... 52.4   82.8  28%   1e-09 50.94  NP_001321519.1
hypothetical protein AXX17_AT1G17520 [Arabidopsis thaliana]       52.4   82.8  28%   1e-09 50.94  OAP12735.1
hypothetical protein AXX17_AT1G17520 [Arabidopsis thaliana]       52.4   82.8  28%   1e-09 50.94  OAP12737.1
unnamed protein product [Arabidopsis thaliana]                    52.0   82.4  28%   1e-09 50.94  CAA0211909.1
GPI-anchored adhesin-like protein, putative (Protein of unknow... 52.0   82.4  28%   1e-09 50.94  NP_564005.2


Alignments:
>hypothetical protein AXX17_AT3G35670 [Arabidopsis thaliana]
Sequence ID: OAP02798.1 Length: 89
Range 1: 1 to 89

Score:186 bits(472), Expect:9e-60,
Method:Compositional matrix adjust.,
Identities:89/89(100%), Positives:89/89(100%), Gaps:0/89(0%)

Query  262  MRNGTLRESLKIFLKIHPIKVQDQEIISQHFFLIQEHNESFFVQVHVTIHFHIFYKYFYA  441
            MRNGTLRESLKIFLKIHPIKVQDQEIISQHFFLIQEHNESFFVQVHVTIHFHIFYKYFYA
Sbjct  1    MRNGTLRESLKIFLKIHPIKVQDQEIISQHFFLIQEHNESFFVQVHVTIHFHIFYKYFYA  60

Query  442  TKMKERNLEANWQRPCGVMIMLMSYRNRL  528
            TKMKERNLEANWQRPCGVMIMLMSYRNRL
Sbjct  61   TKMKERNLEANWQRPCGVMIMLMSYRNRL  89


>GPI-anchored adhesin-like protein, putative (Protein of unknown function, DUF547) [Arabidopsis thaliana]
Sequence ID: NP_001321519.1 Length: 403
>GPI-anchored adhesin-like protein, putative (Protein of unknown function, DUF547) [Arabidopsis thaliana]
Sequence ID: ANM59130.1 Length: 403
Range 1: 157 to 209

Score:52.4 bits(124), Expect:1e-09,
Method:Compositional matrix adjust.,
Identities:27/53(51%), Positives:35/53(66%), Gaps:0/53(0%)

Query  444  KDEGTQPRSKLAKTLWSYDNANELSE*IIRCIKNPAYTLMNLLMDKKSSDVLA  602
            KD   QPRSK+AK+L S+DNANELS+ +IRC++N   +L       KSS   A
Sbjct  157  KDHKAQPRSKVAKSLQSFDNANELSKEMIRCMRNIFVSLGETSAGSKSSQETA  209


Range 2: 108 to 133

Score:30.4 bits(67), Expect:1e-09,
Method:Compositional matrix adjust.,
Identities:13/26(50%), Positives:18/26(69%), Gaps:0/26(0%)

Query  328  DQEIISQHFFLIQEHNESFFVQVHVT  405
            +QEI+S HF LIQE NE    + ++T
Sbjct  108  EQEIMSLHFLLIQERNERKLAEYNLT  133


>hypothetical protein AXX17_AT1G17520 [Arabidopsis thaliana]
Sequence ID: OAP12735.1 Length: 529
Range 1: 157 to 209

Score:52.4 bits(124), Expect:1e-09,
Method:Compositional matrix adjust.,
Identities:27/53(51%), Positives:35/53(66%), Gaps:0/53(0%)

Query  444  KDEGTQPRSKLAKTLWSYDNANELSE*IIRCIKNPAYTLMNLLMDKKSSDVLA  602
            KD   QPRSK+AK+L S+DNANELS+ +IRC++N   +L       KSS   A
Sbjct  157  KDHKAQPRSKVAKSLQSFDNANELSKEMIRCMRNIFVSLGETSAGSKSSQETA  209


Range 2: 108 to 133

Score:30.4 bits(67), Expect:1e-09,
Method:Compositional matrix adjust.,
Identities:13/26(50%), Positives:18/26(69%), Gaps:0/26(0%)

Query  328  DQEIISQHFFLIQEHNESFFVQVHVT  405
            +QEI+S HF LIQE NE    + ++T
Sbjct  108  EQEIMSLHFLLIQERNERKLAEYNLT  133


>hypothetical protein AXX17_AT1G17520 [Arabidopsis thaliana]
Sequence ID: OAP12737.1 Length: 356
Range 1: 157 to 209

Score:52.4 bits(124), Expect:1e-09,
Method:Compositional matrix adjust.,
Identities:27/53(51%), Positives:35/53(66%), Gaps:0/53(0%)

Query  444  KDEGTQPRSKLAKTLWSYDNANELSE*IIRCIKNPAYTLMNLLMDKKSSDVLA  602
            KD   QPRSK+AK+L S+DNANELS+ +IRC++N   +L       KSS   A
Sbjct  157  KDHKAQPRSKVAKSLQSFDNANELSKEMIRCMRNIFVSLGETSAGSKSSQETA  209


Range 2: 108 to 133

Score:30.4 bits(67), Expect:1e-09,
Method:Compositional matrix adjust.,
Identities:13/26(50%), Positives:18/26(69%), Gaps:0/26(0%)

Query  328  DQEIISQHFFLIQEHNESFFVQVHVT  405
            +QEI+S HF LIQE NE    + ++T
Sbjct  108  EQEIMSLHFLLIQERNERKLAEYNLT  133


>unnamed protein product [Arabidopsis thaliana]
Sequence ID: CAA0211909.1 Length: 528
>unnamed protein product [Arabidopsis thaliana]
Sequence ID: VYS46303.1 Length: 528
Range 1: 156 to 208

Score:52.0 bits(123), Expect:1e-09,
Method:Compositional matrix adjust.,
Identities:27/53(51%), Positives:35/53(66%), Gaps:0/53(0%)

Query  444  KDEGTQPRSKLAKTLWSYDNANELSE*IIRCIKNPAYTLMNLLMDKKSSDVLA  602
            KD   QPRSK+AK+L S+DNANELS+ +IRC++N   +L       KSS   A
Sbjct  156  KDHKAQPRSKVAKSLQSFDNANELSKEMIRCMRNIFVSLGETSAGSKSSQETA  208


Range 2: 107 to 132

Score:30.4 bits(67), Expect:1e-09,
Method:Compositional matrix adjust.,
Identities:13/26(50%), Positives:18/26(69%), Gaps:0/26(0%)

Query  328  DQEIISQHFFLIQEHNESFFVQVHVT  405
            +QEI+S HF LIQE NE    + ++T
Sbjct  107  EQEIMSLHFLLIQERNERKLAEYNLT  132


>GPI-anchored adhesin-like protein, putative (Protein of unknown function, DUF547) [Arabidopsis thaliana]
Sequence ID: NP_564005.2 Length: 529
>At1g16750/F19K19_26 [Arabidopsis thaliana]
Sequence ID: AAM83216.1 Length: 529 >At1g16750/F19K19_26 [Arabidopsis thaliana]
Sequence ID: AAN46838.1 Length: 529 >GPI-anchored adhesin-like protein, putative (Protein of unknown function, DUF547) [Arabidopsis thaliana]
Sequence ID: AEE29492.1 Length: 529
Range 1: 157 to 209

Score:52.0 bits(123), Expect:1e-09,
Method:Compositional matrix adjust.,
Identities:27/53(51%), Positives:35/53(66%), Gaps:0/53(0%)

Query  444  KDEGTQPRSKLAKTLWSYDNANELSE*IIRCIKNPAYTLMNLLMDKKSSDVLA  602
            KD   QPRSK+AK+L S+DNANELS+ +IRC++N   +L       KSS   A
Sbjct  157  KDHKAQPRSKVAKSLQSFDNANELSKEMIRCMRNIFVSLGETSAGSKSSQETA  209


Range 2: 108 to 133

Score:30.4 bits(67), Expect:1e-09,
Method:Compositional matrix adjust.,
Identities:13/26(50%), Positives:18/26(69%), Gaps:0/26(0%)

Query  328  DQEIISQHFFLIQEHNESFFVQVHVT  405
            +QEI+S HF LIQE NE    + ++T
Sbjct  108  EQEIMSLHFLLIQERNERKLAEYNLT  133


Query #138: XLOC_015882 Query ID: lcl|Query_33141 Length: 1346

Sequences producing significant alignments:
                                                                  Max    Total Query   E   Per.
Description                                                       Score  Score cover Value Ident  Accession
hypothetical protein AXX17_AT3G40090 [Arabidopsis thaliana]       62.0   122   18%   5e-21 59.65  OAP01827.1
TOPLESS-related 2 [Arabidopsis thaliana]                          58.5   92.0  26%   4e-12 38.54  NP_188306.2
unnamed protein product [Arabidopsis thaliana]                    58.5   92.0  26%   5e-12 38.54  BAA95777.1
TOPLESS-related 3 [Arabidopsis thaliana]                          60.8   91.6  26%   6e-12 40.62  NP_001190409.1
TOPLESS-related 3 [Arabidopsis thaliana]                          60.5   91.3  26%   7e-12 40.62  NP_198055.3
unnamed protein product [Arabidopsis thaliana]                    60.5   91.3  26%   7e-12 40.62  VYS68046.1
TPR3 [Arabidopsis thaliana]                                       60.5   91.3  26%   7e-12 40.62  OAO94316.1
contains similarity to Pfam family PF0040 - WD domain, G-beta...  60.8   91.3  26%   7e-12 40.62  AAD48936.1
TPR1 [Arabidopsis thaliana]                                       57.8   89.3  26%   2e-11 37.76  OAP14430.1
TOPLESS-related 1 [Arabidopsis thaliana]                          57.8   89.3  26%   2e-11 37.76  NP_001319429.1
TOPLESS-related 1 [Arabidopsis thaliana]                          57.8   89.3  26%   2e-11 37.76  NP_178164.3
TPR1 [Arabidopsis thaliana]                                       57.8   89.3  26%   2e-11 37.76  OAP14431.1
hypothetical protein [Arabidopsis thaliana]                       57.8   89.3  26%   2e-11 37.76  BAE98958.1
unnamed protein product [Arabidopsis thaliana]                    57.8   89.3  26%   2e-11 37.76  VYS51629.1
WSIP1 [Arabidopsis thaliana]                                      57.8   88.6  26%   4e-11 37.76  OAP18986.1
unnamed protein product [Arabidopsis thaliana]                    57.8   88.6  26%   4e-11 37.76  CAA0208306.1
unknown protein; 52184-57536 [Arabidopsis thaliana]               57.0   88.6  26%   4e-11 37.76  AAF27128.1
Transducin family protein / WD-40 repeat family protein...        57.4   88.2  26%   4e-11 37.76  NP_001031050.1
Strong similarity to an unknown protein T21F11.18 gi|6730738...   57.4   88.2  26%   5e-11 37.76  AAF82145.1
Kinesin motor family protein [Arabidopsis thaliana]               64.7   64.7  13%   3e-10 68.85  NP_188650.2
hypothetical protein AXX17_AT3G21490 [Arabidopsis thaliana]       64.7   64.7  13%   3e-10 68.85  OAP05410.1
unnamed protein product [Arabidopsis thaliana]                    64.7   64.7  13%   3e-10 68.85  VYS57992.1
unnamed protein product [Arabidopsis thaliana]                    64.7   64.7  13%   3e-10 68.85  BAB01875.1
unnamed protein product [Arabidopsis thaliana]                    63.2   63.2  13%   9e-10 67.21  CAA0383058.1
Kinesin motor family protein [Arabidopsis thaliana]               61.6   61.6  13%   2e-09 68.85  NP_001327101.1
unnamed protein product [Arabidopsis thaliana]                    53.9   82.0  26%   3e-09 39.58  CAA0382578.1
WUS-interacting protein 2 [Arabidopsis thaliana]                  53.9   82.0  26%   3e-09 39.58  NP_001325883.1
WUS-interacting protein 2 [Arabidopsis thaliana]                  53.9   82.0  26%   3e-09 39.58  NP_001189905.1
WUS-interacting protein 2 [Arabidopsis thaliana]                  53.9   82.0  26%   3e-09 39.58  NP_188209.3
putative WD-repeat protein [Arabidopsis thaliana]                 53.1   79.3  27%   2e-08 38.61  BAF01694.1


Alignments:
>hypothetical protein AXX17_AT3G40090 [Arabidopsis thaliana]
Sequence ID: OAP01827.1 Length: 94
Range 1: 38 to 94

Score:62.0 bits(149), Expect:5e-21,
Method:Compositional matrix adjust.,
Identities:34/57(60%), Positives:38/57(66%), Gaps:6/57(10%)

Query  941  SRRADQVLAYGQRRSSEYNRCRRWDCG------STKKATF*LFHEGGF*DSSELNRF  789
            SR  DQVLAYGQRRSSEYNRC+RW  G        ++     FHEG F DS+ELNRF
Sbjct  38   SRWTDQVLAYGQRRSSEYNRCQRWTAGFFEIAVRQRRQPSSCFHEGWFYDSNELNRF  94


Range 2: 1 to 31

Score:60.1 bits(144), Expect:5e-21,
Method:Composition-based stats.,
Identities:29/31(94%), Positives:29/31(93%), Gaps:0/31(0%)

Query  1047  MQITEKKGIVKRIYSGLTNKLGGVVQFQSSK  955
             MQITEKKG VKRIYSGLTN LGGVVQFQSSK
Sbjct  1     MQITEKKGTVKRIYSGLTNNLGGVVQFQSSK  31


>TOPLESS-related 2 [Arabidopsis thaliana]
Sequence ID: NP_188306.2 Length: 1131
>RecName: Full=Topless-related protein 2 [Arabidopsis thaliana]
Sequence ID: Q9LRZ0.2 Length: 1131 >TOPLESS-related 2 [Arabidopsis thaliana]
Sequence ID: AEE75871.1 Length: 1131 >TPR2 [Arabidopsis thaliana]
Sequence ID: OAP03869.1 Length: 1131 >unnamed protein product [Arabidopsis thaliana]
Sequence ID: CAA0382680.1 Length: 1131 >unnamed protein product [Arabidopsis thaliana]
Sequence ID: VYS57623.1 Length: 1131
Range 1: 554 to 649

Score:58.5 bits(140), Expect:4e-12,
Method:Composition-based stats.,
Identities:37/96(39%), Positives:53/96(55%), Gaps:8/96(8%)

Query  1092  RLFCCGTSKLRDSPLMQITEKKGIVKRIYSGLTNKLGGVVQFQSSKKPILVKTG*SSFGI  913
             RLF CGTSK  DS L++  E +G +KR Y G   K  GVVQF +++   L     +
Sbjct  554   RLFSCGTSKEGDSFLVEWNESEGALKRTYLGFRKKSAGVVQFDTTRNRFLAVGEDNQIKF  613

Query  912   W---TASIF*VQQ----MPKVG-LRFDKEGNLLAVS  829
             W     ++  V +    +P +  LRF+K+GNLLAV+
Sbjct  614   WNMDNTNLLTVVEAEGGLPNLPRLRFNKDGNLLAVT  649


Range 2: 526 to 555

Score:33.5 bits(75), Expect:4e-12,
Method:Composition-based stats.,
Identities:15/30(50%), Positives:20/30(66%), Gaps:0/30(0%)

Query  1259  LYDTKCSKEN*KCPSG*CTTLLYNADGNRL  1170
             LYD   S+ +   P   CTT+LY+ADG+RL
Sbjct  526   LYDNVGSRVDYDAPGQWCTTMLYSADGSRL  555


>unnamed protein product [Arabidopsis thaliana]
Sequence ID: BAA95777.1 Length: 1161
Range 1: 554 to 649

Score:58.5 bits(140), Expect:5e-12,
Method:Composition-based stats.,
Identities:37/96(39%), Positives:53/96(55%), Gaps:8/96(8%)

Query  1092  RLFCCGTSKLRDSPLMQITEKKGIVKRIYSGLTNKLGGVVQFQSSKKPILVKTG*SSFGI  913
             RLF CGTSK  DS L++  E +G +KR Y G   K  GVVQF +++   L     +
Sbjct  554   RLFSCGTSKEGDSFLVEWNESEGALKRTYLGFRKKSAGVVQFDTTRNRFLAVGEDNQIKF  613

Query  912   W---TASIF*VQQ----MPKVG-LRFDKEGNLLAVS  829
             W     ++  V +    +P +  LRF+K+GNLLAV+
Sbjct  614   WNMDNTNLLTVVEAEGGLPNLPRLRFNKDGNLLAVT  649


Range 2: 526 to 555

Score:33.5 bits(75), Expect:5e-12,
Method:Composition-based stats.,
Identities:15/30(50%), Positives:20/30(66%), Gaps:0/30(0%)

Query  1259  LYDTKCSKEN*KCPSG*CTTLLYNADGNRL  1170
             LYD   S+ +   P   CTT+LY+ADG+RL
Sbjct  526   LYDNVGSRVDYDAPGQWCTTMLYSADGSRL  555


>TOPLESS-related 3 [Arabidopsis thaliana]
Sequence ID: NP_001190409.1 Length: 1134
>TOPLESS-related 3 [Arabidopsis thaliana]
Sequence ID: AED93643.1 Length: 1134
Range 1: 579 to 674

Score:60.8 bits(146), Expect:6e-12,
Method:Composition-based stats.,
Identities:39/96(41%), Positives:50/96(52%), Gaps:8/96(8%)

Query  1092  RLFCCGTSKLRDSPLMQITEKKGIVKRIYSGLTNKLGGVVQFQSSKKPILVKTG*SSFGI  913
             RLF CGTSK  DS L++  E +G +KR Y     KL GVVQF +SK   L
Sbjct  579   RLFSCGTSKDGDSFLVEWNESEGSIKRTYKEFQKKLAGVVQFDTSKNHFLAVGEDGQIKF  638

Query  912   WTASIF*VQQ-------MPKV-GLRFDKEGNLLAVS  829
             W  +   V         +P +  LRF+K+GNLLAV+
Sbjct  639   WDMNNINVLTSTDAEGGLPALPHLRFNKDGNLLAVT  674


Range 2: 551 to 580

Score:30.8 bits(68), Expect:6e-12,
Method:Composition-based stats.,
Identities:14/30(47%), Positives:18/30(60%), Gaps:0/30(0%)

Query  1259  LYDTKCSKEN*KCPSG*CTTLLYNADGNRL  1170
             LYD   S+ +   P   CT +LY+ADG RL
Sbjct  551   LYDNLGSRVDYDAPGKWCTRMLYSADGTRL  580


>TOPLESS-related 3 [Arabidopsis thaliana]
Sequence ID: NP_198055.3 Length: 1108
>RecName: Full=Topless-related protein 3 [Arabidopsis thaliana]
Sequence ID: Q84JM4.1 Length: 1108 >unknown protein [Arabidopsis thaliana]
Sequence ID: AAO42071.1 Length: 1108 >unknown protein [Arabidopsis thaliana]
Sequence ID: AAO50698.1 Length: 1108 >TOPLESS-related 3 [Arabidopsis thaliana]
Sequence ID: AED93642.1 Length: 1108 >unnamed protein product [Arabidopsis thaliana]
Sequence ID: CAA0405064.1 Length: 1108
Range 1: 553 to 648

Score:60.5 bits(145), Expect:7e-12,
Method:Compositional matrix adjust.,
Identities:39/96(41%), Positives:50/96(52%), Gaps:8/96(8%)

Query  1092  RLFCCGTSKLRDSPLMQITEKKGIVKRIYSGLTNKLGGVVQFQSSKKPILVKTG*SSFGI  913
             RLF CGTSK  DS L++  E +G +KR Y     KL GVVQF +SK   L
Sbjct  553   RLFSCGTSKDGDSFLVEWNESEGSIKRTYKEFQKKLAGVVQFDTSKNHFLAVGEDGQIKF  612

Query  912   WTASIF*VQQ-------MPKV-GLRFDKEGNLLAVS  829
             W  +   V         +P +  LRF+K+GNLLAV+
Sbjct  613   WDMNNINVLTSTDAEGGLPALPHLRFNKDGNLLAVT  648


Range 2: 525 to 554

Score:30.8 bits(68), Expect:7e-12,
Method:Composition-based stats.,
Identities:14/30(47%), Positives:18/30(60%), Gaps:0/30(0%)

Query  1259  LYDTKCSKEN*KCPSG*CTTLLYNADGNRL  1170
             LYD   S+ +   P   CT +LY+ADG RL
Sbjct  525   LYDNLGSRVDYDAPGKWCTRMLYSADGTRL  554


>unnamed protein product [Arabidopsis thaliana]
Sequence ID: VYS68046.1 Length: 1108
Range 1: 553 to 648

Score:60.5 bits(145), Expect:7e-12,
Method:Compositional matrix adjust.,
Identities:39/96(41%), Positives:50/96(52%), Gaps:8/96(8%)

Query  1092  RLFCCGTSKLRDSPLMQITEKKGIVKRIYSGLTNKLGGVVQFQSSKKPILVKTG*SSFGI  913
             RLF CGTSK  DS L++  E +G +KR Y     KL GVVQF +SK   L
Sbjct  553   RLFSCGTSKDGDSFLVEWNESEGSIKRTYKEFQKKLAGVVQFDTSKNHFLAVGEDGQIKF  612

Query  912   WTASIF*VQQ-------MPKV-GLRFDKEGNLLAVS  829
             W  +   V         +P +  LRF+K+GNLLAV+
Sbjct  613   WDMNNINVLTSTDAEGGLPALPHLRFNKDGNLLAVT  648


Range 2: 525 to 554

Score:30.8 bits(68), Expect:7e-12,
Method:Composition-based stats.,
Identities:14/30(47%), Positives:18/30(60%), Gaps:0/30(0%)

Query  1259  LYDTKCSKEN*KCPSG*CTTLLYNADGNRL  1170
             LYD   S+ +   P   CT +LY+ADG RL
Sbjct  525   LYDNLGSRVDYDAPGKWCTRMLYSADGTRL  554


>TPR3 [Arabidopsis thaliana]
Sequence ID: OAO94316.1 Length: 1108
Range 1: 553 to 648

Score:60.5 bits(145), Expect:7e-12,
Method:Compositional matrix adjust.,
Identities:39/96(41%), Positives:50/96(52%), Gaps:8/96(8%)

Query  1092  RLFCCGTSKLRDSPLMQITEKKGIVKRIYSGLTNKLGGVVQFQSSKKPILVKTG*SSFGI  913
             RLF CGTSK  DS L++  E +G +KR Y     KL GVVQF +SK   L
Sbjct  553   RLFSCGTSKDGDSFLVEWNESEGSIKRTYKEFQKKLAGVVQFDTSKNHFLAVGEDGQIKF  612

Query  912   WTASIF*VQQ-------MPKV-GLRFDKEGNLLAVS  829
             W  +   V         +P +  LRF+K+GNLLAV+
Sbjct  613   WDMNNINVLTSTDAEGGLPALPHLRFNKDGNLLAVT  648


Range 2: 525 to 554

Score:30.8 bits(68), Expect:7e-12,
Method:Composition-based stats.,
Identities:14/30(47%), Positives:18/30(60%), Gaps:0/30(0%)

Query  1259  LYDTKCSKEN*KCPSG*CTTLLYNADGNRL  1170
             LYD   S+ +   P   CT +LY+ADG RL
Sbjct  525   LYDNLGSRVDYDAPGKWCTRMLYSADGTRL  554


>contains similarity to Pfam family PF0040 - WD domain, G-beta repeat; score=10.8, E=3.2, N-2 [Arabidopsis thaliana]
Sequence ID: AAD48936.1 Length: 892
Range 1: 570 to 665

Score:60.8 bits(146), Expect:7e-12,
Method:Compositional matrix adjust.,
Identities:39/96(41%), Positives:50/96(52%), Gaps:8/96(8%)

Query  1092  RLFCCGTSKLRDSPLMQITEKKGIVKRIYSGLTNKLGGVVQFQSSKKPILVKTG*SSFGI  913
             RLF CGTSK  DS L++  E +G +KR Y     KL GVVQF +SK   L
Sbjct  570   RLFSCGTSKDGDSFLVEWNESEGSIKRTYKEFQKKLAGVVQFDTSKNHFLAVGEDGQIKF  629

Query  912   WTASIF*VQQ-------MPKV-GLRFDKEGNLLAVS  829
             W  +   V         +P +  LRF+K+GNLLAV+
Sbjct  630   WDMNNINVLTSTDAEGGLPALPHLRFNKDGNLLAVT  665


Range 2: 542 to 571

Score:30.4 bits(67), Expect:7e-12,
Method:Composition-based stats.,
Identities:14/30(47%), Positives:18/30(60%), Gaps:0/30(0%)

Query  1259  LYDTKCSKEN*KCPSG*CTTLLYNADGNRL  1170
             LYD   S+ +   P   CT +LY+ADG RL
Sbjct  542   LYDNLGSRVDYDAPGKWCTRMLYSADGTRL  571


>TPR1 [Arabidopsis thaliana]
Sequence ID: OAP14430.1 Length: 1120
Range 1: 564 to 659

Score:57.8 bits(138), Expect:2e-11,
Method:Composition-based stats.,
Identities:37/98(38%), Positives:50/98(51%), Gaps:12/98(12%)

Query  1092  RLFCCGTSKLRDSPLMQITEKKGIVKRIYSGLTNKLGGVVQFQSSKKPILVKTG*SSFGI  913
             RLF CGTSK  +S +++  E +G VKR Y G   +  GVVQF ++K   L      S
Sbjct  564   RLFSCGTSKDGESFIVEWNESEGAVKRTYQGFHKRSLGVVQFDTTKNRYLAAGDDFSIKF  623

Query  912   WTASIF----------*VQQMPKVGLRFDKEGNLLAVS  829
             W                +Q  P++  RF+KEG+LLAVS
Sbjct  624   WDMDTIQLLTAIDADGGLQASPRI--RFNKEGSLLAVS  659


Range 2: 536 to 565

Score:31.6 bits(70), Expect:2e-11,
Method:Composition-based stats.,
Identities:14/30(47%), Positives:19/30(63%), Gaps:0/30(0%)

Query  1259  LYDTKCSKEN*KCPSG*CTTLLYNADGNRL  1170
             LYD   S+ + + P   CTT+ Y+ADG RL
Sbjct  536   LYDNMGSRVDYEAPGRWCTTMAYSADGTRL  565


>TOPLESS-related 1 [Arabidopsis thaliana]
Sequence ID: NP_001319429.1 Length: 1120
>TOPLESS-related 1 [Arabidopsis thaliana]
Sequence ID: NP_849913.2 Length: 1120 >RecName: Full=Topless-related protein 1; AltName: Full=Protein MODIFIER OF SNC1 10 [Arabidopsis thaliana]
Sequence ID: Q0WV90.3 Length: 1120 >TOPLESS-related 1 [Arabidopsis thaliana]
Sequence ID: AEE36412.1 Length: 1120 >TOPLESS-related 1 [Arabidopsis thaliana]
Sequence ID: ANM60269.1 Length: 1120 >unnamed protein product [Arabidopsis thaliana]
Sequence ID: CAA0344431.1 Length: 1120
Range 1: 564 to 659

Score:57.8 bits(138), Expect:2e-11,
Method:Composition-based stats.,
Identities:37/98(38%), Positives:50/98(51%), Gaps:12/98(12%)

Query  1092  RLFCCGTSKLRDSPLMQITEKKGIVKRIYSGLTNKLGGVVQFQSSKKPILVKTG*SSFGI  913
             RLF CGTSK  +S +++  E +G VKR Y G   +  GVVQF ++K   L      S
Sbjct  564   RLFSCGTSKDGESFIVEWNESEGAVKRTYQGFHKRSLGVVQFDTTKNRYLAAGDDFSIKF  623

Query  912   WTASIF----------*VQQMPKVGLRFDKEGNLLAVS  829
             W                +Q  P++  RF+KEG+LLAVS
Sbjct  624   WDMDTIQLLTAIDADGGLQASPRI--RFNKEGSLLAVS  659


Range 2: 536 to 565

Score:31.6 bits(70), Expect:2e-11,
Method:Composition-based stats.,
Identities:14/30(47%), Positives:19/30(63%), Gaps:0/30(0%)

Query  1259  LYDTKCSKEN*KCPSG*CTTLLYNADGNRL  1170
             LYD   S+ + + P   CTT+ Y+ADG RL
Sbjct  536   LYDNMGSRVDYEAPGRWCTTMAYSADGTRL  565


>TOPLESS-related 1 [Arabidopsis thaliana]
Sequence ID: NP_178164.3 Length: 1119
>TOPLESS-related 1 [Arabidopsis thaliana]
Sequence ID: AEE36411.1 Length: 1119
Range 1: 563 to 658

Score:57.8 bits(138), Expect:2e-11,
Method:Composition-based stats.,
Identities:37/98(38%), Positives:50/98(51%), Gaps:12/98(12%)

Query  1092  RLFCCGTSKLRDSPLMQITEKKGIVKRIYSGLTNKLGGVVQFQSSKKPILVKTG*SSFGI  913
             RLF CGTSK  +S +++  E +G VKR Y G   +  GVVQF ++K   L      S
Sbjct  563   RLFSCGTSKDGESFIVEWNESEGAVKRTYQGFHKRSLGVVQFDTTKNRYLAAGDDFSIKF  622

Query  912   WTASIF----------*VQQMPKVGLRFDKEGNLLAVS  829
             W                +Q  P++  RF+KEG+LLAVS
Sbjct  623   WDMDTIQLLTAIDADGGLQASPRI--RFNKEGSLLAVS  658


Range 2: 535 to 564

Score:31.6 bits(70), Expect:2e-11,
Method:Composition-based stats.,
Identities:14/30(47%), Positives:19/30(63%), Gaps:0/30(0%)

Query  1259  LYDTKCSKEN*KCPSG*CTTLLYNADGNRL  1170
             LYD   S+ + + P   CTT+ Y+ADG RL
Sbjct  535   LYDNMGSRVDYEAPGRWCTTMAYSADGTRL  564


>TPR1 [Arabidopsis thaliana]
Sequence ID: OAP14431.1 Length: 1119
Range 1: 563 to 658

Score:57.8 bits(138), Expect:2e-11,
Method:Composition-based stats.,
Identities:37/98(38%), Positives:50/98(51%), Gaps:12/98(12%)

Query  1092  RLFCCGTSKLRDSPLMQITEKKGIVKRIYSGLTNKLGGVVQFQSSKKPILVKTG*SSFGI  913
             RLF CGTSK  +S +++  E +G VKR Y G   +  GVVQF ++K   L      S
Sbjct  563   RLFSCGTSKDGESFIVEWNESEGAVKRTYQGFHKRSLGVVQFDTTKNRYLAAGDDFSIKF  622

Query  912   WTASIF----------*VQQMPKVGLRFDKEGNLLAVS  829
             W                +Q  P++  RF+KEG+LLAVS
Sbjct  623   WDMDTIQLLTAIDADGGLQASPRI--RFNKEGSLLAVS  658


Range 2: 535 to 564

Score:31.6 bits(70), Expect:2e-11,
Method:Composition-based stats.,
Identities:14/30(47%), Positives:19/30(63%), Gaps:0/30(0%)

Query  1259  LYDTKCSKEN*KCPSG*CTTLLYNADGNRL  1170
             LYD   S+ + + P   CTT+ Y+ADG RL
Sbjct  535   LYDNMGSRVDYEAPGRWCTTMAYSADGTRL  564


>hypothetical protein [Arabidopsis thaliana]
Sequence ID: BAE98958.1 Length: 1119
Range 1: 563 to 658

Score:57.8 bits(138), Expect:2e-11,
Method:Composition-based stats.,
Identities:37/98(38%), Positives:50/98(51%), Gaps:12/98(12%)

Query  1092  RLFCCGTSKLRDSPLMQITEKKGIVKRIYSGLTNKLGGVVQFQSSKKPILVKTG*SSFGI  913
             RLF CGTSK  +S +++  E +G VKR Y G   +  GVVQF ++K   L      S
Sbjct  563   RLFSCGTSKDGESFIVEWNESEGAVKRTYQGFHKRSLGVVQFDTTKNRYLAAGDDFSIKF  622

Query  912   WTASIF----------*VQQMPKVGLRFDKEGNLLAVS  829
             W                +Q  P++  RF+KEG+LLAVS
Sbjct  623   WDMDTIQLLTAIDADGGLQASPRI--RFNKEGSLLAVS  658


Range 2: 535 to 564

Score:31.6 bits(70), Expect:2e-11,
Method:Composition-based stats.,
Identities:14/30(47%), Positives:19/30(63%), Gaps:0/30(0%)

Query  1259  LYDTKCSKEN*KCPSG*CTTLLYNADGNRL  1170
             LYD   S+ + + P   CTT+ Y+ADG RL
Sbjct  535   LYDNMGSRVDYEAPGRWCTTMAYSADGTRL  564


>unnamed protein product [Arabidopsis thaliana]
Sequence ID: VYS51629.1 Length: 1130
Range 1: 564 to 659

Score:57.8 bits(138), Expect:2e-11,
Method:Composition-based stats.,
Identities:37/98(38%), Positives:50/98(51%), Gaps:12/98(12%)

Query  1092  RLFCCGTSKLRDSPLMQITEKKGIVKRIYSGLTNKLGGVVQFQSSKKPILVKTG*SSFGI  913
             RLF CGTSK  +S +++  E +G VKR Y G   +  GVVQF ++K   L      S
Sbjct  564   RLFSCGTSKDGESFIVEWNESEGAVKRTYQGFHKRSLGVVQFDTTKNRYLAAGDDFSIKF  623

Query  912   WTASIF----------*VQQMPKVGLRFDKEGNLLAVS  829
             W                +Q  P++  RF+KEG+LLAVS
Sbjct  624   WDMDTIQLLTAIDADGGLQASPRI--RFNKEGSLLAVS  659


Range 2: 536 to 565

Score:31.6 bits(70), Expect:2e-11,
Method:Composition-based stats.,
Identities:14/30(47%), Positives:19/30(63%), Gaps:0/30(0%)

Query  1259  LYDTKCSKEN*KCPSG*CTTLLYNADGNRL  1170
             LYD   S+ + + P   CTT+ Y+ADG RL
Sbjct  536   LYDNMGSRVDYEAPGRWCTTMAYSADGTRL  565


>WSIP1 [Arabidopsis thaliana]
Sequence ID: OAP18986.1 Length: 1131
>unnamed protein product [Arabidopsis thaliana]
Sequence ID: VYS46204.1 Length: 1131
Range 1: 564 to 659

Score:57.8 bits(138), Expect:4e-11,
Method:Composition-based stats.,
Identities:37/98(38%), Positives:50/98(51%), Gaps:12/98(12%)

Query  1092  RLFCCGTSKLRDSPLMQITEKKGIVKRIYSGLTNKLGGVVQFQSSKKPILVKTG*SSFGI  913
             RLF CGTSK  +S +++  E +G VKR Y G   +  GVVQF ++K   L      S
Sbjct  564   RLFSCGTSKDGESFIVEWNESEGAVKRTYQGFHKRSLGVVQFDTTKNRYLAAGDDFSIKF  623

Query  912   WTASIF----------*VQQMPKVGLRFDKEGNLLAVS  829
             W                +Q  P++  RF+KEG+LLAVS
Sbjct  624   WDMDTVQLLTAIDGDGGLQASPRI--RFNKEGSLLAVS  659


Range 2: 536 to 565

Score:30.8 bits(68), Expect:4e-11,
Method:Composition-based stats.,
Identities:14/30(47%), Positives:18/30(60%), Gaps:0/30(0%)

Query  1259  LYDTKCSKEN*KCPSG*CTTLLYNADGNRL  1170
             LYD   S+ +   P   CTT+ Y+ADG RL
Sbjct  536   LYDNMGSRVDYDAPGRWCTTMAYSADGTRL  565


>unnamed protein product [Arabidopsis thaliana]
Sequence ID: CAA0208306.1 Length: 1129
Range 1: 564 to 659

Score:57.8 bits(138), Expect:4e-11,
Method:Composition-based stats.,
Identities:37/98(38%), Positives:50/98(51%), Gaps:12/98(12%)

Query  1092  RLFCCGTSKLRDSPLMQITEKKGIVKRIYSGLTNKLGGVVQFQSSKKPILVKTG*SSFGI  913
             RLF CGTSK  +S +++  E +G VKR Y G   +  GVVQF ++K   L      S
Sbjct  564   RLFSCGTSKDGESFIVEWNESEGAVKRTYQGFHKRSLGVVQFDTTKNRYLAAGDDFSIKF  623

Query  912   WTASIF----------*VQQMPKVGLRFDKEGNLLAVS  829
             W                +Q  P++  RF+KEG+LLAVS
Sbjct  624   WDMDTVQLLTAIDGDGGLQASPRI--RFNKEGSLLAVS  659


Range 2: 536 to 565

Score:30.8 bits(68), Expect:4e-11,
Method:Composition-based stats.,
Identities:14/30(47%), Positives:18/30(60%), Gaps:0/30(0%)

Query  1259  LYDTKCSKEN*KCPSG*CTTLLYNADGNRL  1170
             LYD   S+ +   P   CTT+ Y+ADG RL
Sbjct  536   LYDNMGSRVDYDAPGRWCTTMAYSADGTRL  565


>unknown protein; 52184-57536 [Arabidopsis thaliana]
Sequence ID: AAF27128.1 Length: 1073
Range 1: 470 to 565

Score:57.0 bits(136), Expect:4e-11,
Method:Compositional matrix adjust.,
Identities:37/98(38%), Positives:50/98(51%), Gaps:12/98(12%)

Query  1092  RLFCCGTSKLRDSPLMQITEKKGIVKRIYSGLTNKLGGVVQFQSSKKPILVKTG*SSFGI  913
             RLF CGTSK  +S +++  E +G VKR Y G   +  GVVQF ++K   L      S
Sbjct  470   RLFSCGTSKDGESFIVEWNESEGAVKRTYQGFHKRSLGVVQFDTTKNRYLAAGDDFSIKF  529

Query  912   WTASIF----------*VQQMPKVGLRFDKEGNLLAVS  829
             W                +Q  P++  RF+KEG+LLAVS
Sbjct  530   WDMDTIQLLTAIDADGGLQASPRI--RFNKEGSLLAVS  565


Range 2: 442 to 471

Score:31.6 bits(70), Expect:4e-11,
Method:Composition-based stats.,
Identities:14/30(47%), Positives:19/30(63%), Gaps:0/30(0%)

Query  1259  LYDTKCSKEN*KCPSG*CTTLLYNADGNRL  1170
             LYD   S+ + + P   CTT+ Y+ADG RL
Sbjct  442   LYDNMGSRVDYEAPGRWCTTMAYSADGTRL  471


>Transducin family protein / WD-40 repeat family protein [Arabidopsis thaliana]
Sequence ID: NP_001031050.1 Length: 1131
>Transducin family protein / WD-40 repeat family protein [Arabidopsis thaliana]
Sequence ID: NP_001031051.1 Length: 1131 >Transducin family protein / WD-40 repeat family protein [Arabidopsis thaliana]
Sequence ID: NP_563981.1 Length: 1131 >Transducin family protein / WD-40 repeat family protein [Arabidopsis thaliana]
Sequence ID: NP_849672.1 Length: 1131 >RecName: Full=Protein TOPLESS; AltName: Full=WUS-interacting protein 1 [Arabidopsis thaliana]
Sequence ID: Q94AI7.1 Length: 1131 >unknown protein [Arabidopsis thaliana]
Sequence ID: AAK76687.1 Length: 1131 >unknown protein [Arabidopsis thaliana]
Sequence ID: AAN13188.1 Length: 1131 >Transducin family protein / WD-40 repeat family protein [Arabidopsis thaliana]
Sequence ID: AEE29356.1 Length: 1131 >Transducin family protein / WD-40 repeat family protein [Arabidopsis thaliana]
Sequence ID: AEE29357.1 Length: 1131 >Transducin family protein / WD-40 repeat family protein [Arabidopsis thaliana]
Sequence ID: AEE29358.1 Length: 1131
Range 1: 564 to 659

Score:57.4 bits(137), Expect:4e-11,
Method:Composition-based stats.,
Identities:37/98(38%), Positives:50/98(51%), Gaps:12/98(12%)

Query  1092  RLFCCGTSKLRDSPLMQITEKKGIVKRIYSGLTNKLGGVVQFQSSKKPILVKTG*SSFGI  913
             RLF CGTSK  +S +++  E +G VKR Y G   +  GVVQF ++K   L      S
Sbjct  564   RLFSCGTSKDGESFIVEWNESEGAVKRTYQGFHKRSLGVVQFDTTKNRYLAAGDDFSIKF  623

Query  912   WTASIF----------*VQQMPKVGLRFDKEGNLLAVS  829
             W                +Q  P++  RF+KEG+LLAVS
Sbjct  624   WDMDAVQLLTAIDGDGGLQASPRI--RFNKEGSLLAVS  659


Range 2: 536 to 565

Score:30.8 bits(68), Expect:4e-11,
Method:Composition-based stats.,
Identities:14/30(47%), Positives:18/30(60%), Gaps:0/30(0%)

Query  1259  LYDTKCSKEN*KCPSG*CTTLLYNADGNRL  1170
             LYD   S+ +   P   CTT+ Y+ADG RL
Sbjct  536   LYDNMGSRVDYDAPGRWCTTMAYSADGTRL  565


>Strong similarity to an unknown protein T21F11.18 gi|6730738 from Arabidopsis thaliana BAC T21F11 gb|AC018849 and contains multiple WD PF|00400 domains. ESTs gb|Z34157, gb|AA006273, gb|AA605431, gb|W43588, gb|W43605, gb|Z34559, gb|R90037, gb|AI994125 come from this gene [Arabidopsis thaliana]
Sequence ID: AAF82145.1 Length: 1153
Range 1: 586 to 681

Score:57.4 bits(137), Expect:5e-11,
Method:Composition-based stats.,
Identities:37/98(38%), Positives:50/98(51%), Gaps:12/98(12%)

Query  1092  RLFCCGTSKLRDSPLMQITEKKGIVKRIYSGLTNKLGGVVQFQSSKKPILVKTG*SSFGI  913
             RLF CGTSK  +S +++  E +G VKR Y G   +  GVVQF ++K   L      S
Sbjct  586   RLFSCGTSKDGESFIVEWNESEGAVKRTYQGFHKRSLGVVQFDTTKNRYLAAGDDFSIKF  645

Query  912   WTASIF----------*VQQMPKVGLRFDKEGNLLAVS  829
             W                +Q  P++  RF+KEG+LLAVS
Sbjct  646   WDMDAVQLLTAIDGDGGLQASPRI--RFNKEGSLLAVS  681


Range 2: 558 to 587

Score:30.8 bits(68), Expect:5e-11,
Method:Composition-based stats.,
Identities:14/30(47%), Positives:18/30(60%), Gaps:0/30(0%)

Query  1259  LYDTKCSKEN*KCPSG*CTTLLYNADGNRL  1170
             LYD   S+ +   P   CTT+ Y+ADG RL
Sbjct  558   LYDNMGSRVDYDAPGRWCTTMAYSADGTRL  587


>Kinesin motor family protein [Arabidopsis thaliana]
Sequence ID: NP_188650.2 Length: 1114
>RecName: Full=Kinesin-like protein KIN-12F [Arabidopsis thaliana]
Sequence ID: F4JDI6.1 Length: 1114 >Kinesin motor family protein [Arabidopsis thaliana]
Sequence ID: AEE76340.1 Length: 1114
Range 1: 917 to 977

Score:64.7 bits(156), Expect:3e-10,
Method:Compositional matrix adjust.,
Identities:42/61(69%), Positives:46/61(75%), Gaps:0/61(0%)

Query  62   RDNRVEAQLkk*kqsseeseDALPIAVLGQAKFVEHYTELQEKYNELGLNHKATVE*TLE  241
            R  + EAQLK+ K SSEE EDAL  AVLG A+FVEHYTELQEKYN+L   HKATVE   E
Sbjct  917  RAEKAEAQLKQEKLSSEELEDALRRAVLGHARFVEHYTELQEKYNDLCSKHKATVEWITE  976

Query  242  L  244
            L
Sbjct  977  L  977


>hypothetical protein AXX17_AT3G21490 [Arabidopsis thaliana]
Sequence ID: OAP05410.1 Length: 1114
Range 1: 917 to 977

Score:64.7 bits(156), Expect:3e-10,
Method:Compositional matrix adjust.,
Identities:42/61(69%), Positives:46/61(75%), Gaps:0/61(0%)

Query  62   RDNRVEAQLkk*kqsseeseDALPIAVLGQAKFVEHYTELQEKYNELGLNHKATVE*TLE  241
            R  + EAQLK+ K SSEE EDAL  AVLG A+FVEHYTELQEKYN+L   HKATVE   E
Sbjct  917  RAEKAEAQLKQEKLSSEELEDALRRAVLGHARFVEHYTELQEKYNDLCSKHKATVEWITE  976

Query  242  L  244
            L
Sbjct  977  L  977


>unnamed protein product [Arabidopsis thaliana]
Sequence ID: VYS57992.1 Length: 1098
Range 1: 917 to 977

Score:64.7 bits(156), Expect:3e-10,
Method:Compositional matrix adjust.,
Identities:42/61(69%), Positives:46/61(75%), Gaps:0/61(0%)

Query  62   RDNRVEAQLkk*kqsseeseDALPIAVLGQAKFVEHYTELQEKYNELGLNHKATVE*TLE  241
            R  + EAQLK+ K SSEE EDAL  AVLG A+FVEHYTELQEKYN+L   HKATVE   E
Sbjct  917  RAEKAEAQLKQEKLSSEELEDALRRAVLGHARFVEHYTELQEKYNDLCSKHKATVEWITE  976

Query  242  L  244
            L
Sbjct  977  L  977


>unnamed protein product [Arabidopsis thaliana]
Sequence ID: BAB01875.1 Length: 1103
Range 1: 906 to 966

Score:64.7 bits(156), Expect:3e-10,
Method:Compositional matrix adjust.,
Identities:42/61(69%), Positives:46/61(75%), Gaps:0/61(0%)

Query  62   RDNRVEAQLkk*kqsseeseDALPIAVLGQAKFVEHYTELQEKYNELGLNHKATVE*TLE  241
            R  + EAQLK+ K SSEE EDAL  AVLG A+FVEHYTELQEKYN+L   HKATVE   E
Sbjct  906  RAEKAEAQLKQEKLSSEELEDALRRAVLGHARFVEHYTELQEKYNDLCSKHKATVEWITE  965

Query  242  L  244
            L
Sbjct  966  L  966


>unnamed protein product [Arabidopsis thaliana]
Sequence ID: CAA0383058.1 Length: 1114
Range 1: 917 to 977

Score:63.2 bits(152), Expect:9e-10,
Method:Compositional matrix adjust.,
Identities:41/61(67%), Positives:45/61(73%), Gaps:0/61(0%)

Query  62   RDNRVEAQLkk*kqsseeseDALPIAVLGQAKFVEHYTELQEKYNELGLNHKATVE*TLE  241
            R  + E QLK+ K SSEE EDAL  AVLG A+FVEHYTELQEKYN+L   HKATVE   E
Sbjct  917  RAEKAETQLKQEKLSSEELEDALRRAVLGHARFVEHYTELQEKYNDLCSKHKATVEWITE  976

Query  242  L  244
            L
Sbjct  977  L  977


>Kinesin motor family protein [Arabidopsis thaliana]
Sequence ID: NP_001327101.1 Length: 1061
>Kinesin motor family protein [Arabidopsis thaliana]
Sequence ID: ANM65106.1 Length: 1061
Range 1: 917 to 977

Score:61.6 bits(148), Expect:2e-09,
Method:Composition-based stats.,
Identities:42/61(69%), Positives:46/61(75%), Gaps:0/61(0%)

Query  62   RDNRVEAQLkk*kqsseeseDALPIAVLGQAKFVEHYTELQEKYNELGLNHKATVE*TLE  241
            R  + EAQLK+ K SSEE EDAL  AVLG A+FVEHYTELQEKYN+L   HKATVE   E
Sbjct  917  RAEKAEAQLKQEKLSSEELEDALRRAVLGHARFVEHYTELQEKYNDLCSKHKATVEWITE  976

Query  242  L  244
            L
Sbjct  977  L  977


>unnamed protein product [Arabidopsis thaliana]
Sequence ID: CAA0382578.1 Length: 1135
Range 1: 566 to 661

Score:53.9 bits(128), Expect:3e-09,
Method:Composition-based stats.,
Identities:38/96(40%), Positives:48/96(50%), Gaps:8/96(8%)

Query  1092  RLFCCGTSKLRDSPLMQITEKKGIVKRIYSGLTNKLGGVVQFQSSKKPILVKTG*SSFGI  913
             RLF CGTSK  +S +++  E +G VKR Y GL  +  GVVQF + K   LV
Sbjct  566   RLFSCGTSKEGESFIVEWNESEGAVKRTYLGLGKRSVGVVQFDTMKNKFLVAGDEFQVKF  625

Query  912   WTA-SIF*VQQMPKVG-------LRFDKEGNLLAVS  829
             W   S+  +      G       LR +KEG LLAVS
Sbjct  626   WDMDSVDLLSSTAAEGGLPSSPCLRINKEGTLLAVS  661


Range 2: 538 to 567

Score:28.1 bits(61), Expect:3e-09,
Method:Composition-based stats.,
Identities:13/30(43%), Positives:17/30(56%), Gaps:0/30(0%)

Query  1259  LYDTKCSKEN*KCPSG*CTTLLYNADGNRL  1170
             LYD   S+ +   P   CT++ Y ADG RL
Sbjct  538   LYDNMGSRVDYDAPGRSCTSMAYCADGTRL  567


>WUS-interacting protein 2 [Arabidopsis thaliana]
Sequence ID: NP_001325883.1 Length: 1135
>WUS-interacting protein 2 [Arabidopsis thaliana]
Sequence ID: NP_851003.2 Length: 1135 >RecName: Full=Topless-related protein 4; AltName: Full=WUS-interacting protein 2 [Arabidopsis thaliana]
Sequence ID: Q27GK7.2 Length: 1135 >WUS-interacting protein 2 [Arabidopsis thaliana]
Sequence ID: AEE75740.1 Length: 1135 >WUS-interacting protein 2 [Arabidopsis thaliana]
Sequence ID: ANM63812.1 Length: 1135 >WSIP2 [Arabidopsis thaliana]
Sequence ID: OAP05521.1 Length: 1135 >unnamed protein product [Arabidopsis thaliana]
Sequence ID: VYS57522.1 Length: 1135
Range 1: 566 to 661

Score:53.9 bits(128), Expect:3e-09,
Method:Composition-based stats.,
Identities:38/96(40%), Positives:48/96(50%), Gaps:8/96(8%)

Query  1092  RLFCCGTSKLRDSPLMQITEKKGIVKRIYSGLTNKLGGVVQFQSSKKPILVKTG*SSFGI  913
             RLF CGTSK  +S +++  E +G VKR Y GL  +  GVVQF + K   LV
Sbjct  566   RLFSCGTSKEGESFIVEWNESEGAVKRTYLGLGKRSVGVVQFDTMKNKFLVAGDEFQVKF  625

Query  912   WTA-SIF*VQQMPKVG-------LRFDKEGNLLAVS  829
             W   S+  +      G       LR +KEG LLAVS
Sbjct  626   WDMDSVDLLSSTAAEGGLPSSPCLRINKEGTLLAVS  661


Range 2: 538 to 567

Score:28.1 bits(61), Expect:3e-09,
Method:Composition-based stats.,
Identities:13/30(43%), Positives:17/30(56%), Gaps:0/30(0%)

Query  1259  LYDTKCSKEN*KCPSG*CTTLLYNADGNRL  1170
             LYD   S+ +   P   CT++ Y ADG RL
Sbjct  538   LYDNMGSRVDYDAPGRSCTSMAYCADGTRL  567


>WUS-interacting protein 2 [Arabidopsis thaliana]
Sequence ID: NP_001189905.1 Length: 1125
>WUS-interacting protein 2 [Arabidopsis thaliana]
Sequence ID: AEE75742.1 Length: 1125
Range 1: 556 to 651

Score:53.9 bits(128), Expect:3e-09,
Method:Composition-based stats.,
Identities:38/96(40%), Positives:48/96(50%), Gaps:8/96(8%)

Query  1092  RLFCCGTSKLRDSPLMQITEKKGIVKRIYSGLTNKLGGVVQFQSSKKPILVKTG*SSFGI  913
             RLF CGTSK  +S +++  E +G VKR Y GL  +  GVVQF + K   LV
Sbjct  556   RLFSCGTSKEGESFIVEWNESEGAVKRTYLGLGKRSVGVVQFDTMKNKFLVAGDEFQVKF  615

Query  912   WTA-SIF*VQQMPKVG-------LRFDKEGNLLAVS  829
             W   S+  +      G       LR +KEG LLAVS
Sbjct  616   WDMDSVDLLSSTAAEGGLPSSPCLRINKEGTLLAVS  651


Range 2: 528 to 557

Score:28.1 bits(61), Expect:3e-09,
Method:Composition-based stats.,
Identities:13/30(43%), Positives:17/30(56%), Gaps:0/30(0%)

Query  1259  LYDTKCSKEN*KCPSG*CTTLLYNADGNRL  1170
             LYD   S+ +   P   CT++ Y ADG RL
Sbjct  528   LYDNMGSRVDYDAPGRSCTSMAYCADGTRL  557


>WUS-interacting protein 2 [Arabidopsis thaliana]
Sequence ID: NP_188209.3 Length: 1137
>WUS-interacting protein 2 [Arabidopsis thaliana]
Sequence ID: AEE75741.1 Length: 1137
Range 1: 566 to 661

Score:53.9 bits(128), Expect:3e-09,
Method:Composition-based stats.,
Identities:38/96(40%), Positives:48/96(50%), Gaps:8/96(8%)

Query  1092  RLFCCGTSKLRDSPLMQITEKKGIVKRIYSGLTNKLGGVVQFQSSKKPILVKTG*SSFGI  913
             RLF CGTSK  +S +++  E +G VKR Y GL  +  GVVQF + K   LV
Sbjct  566   RLFSCGTSKEGESFIVEWNESEGAVKRTYLGLGKRSVGVVQFDTMKNKFLVAGDEFQVKF  625

Query  912   WTA-SIF*VQQMPKVG-------LRFDKEGNLLAVS  829
             W   S+  +      G       LR +KEG LLAVS
Sbjct  626   WDMDSVDLLSSTAAEGGLPSSPCLRINKEGTLLAVS  661


Range 2: 538 to 567

Score:28.1 bits(61), Expect:3e-09,
Method:Composition-based stats.,
Identities:13/30(43%), Positives:17/30(56%), Gaps:0/30(0%)

Query  1259  LYDTKCSKEN*KCPSG*CTTLLYNADGNRL  1170
             LYD   S+ +   P   CT++ Y ADG RL
Sbjct  538   LYDNMGSRVDYDAPGRSCTSMAYCADGTRL  567


>putative WD-repeat protein, partial [Arabidopsis thaliana]
Sequence ID: BAF01694.1 Length: 744
Range 1: 170 to 270

Score:53.1 bits(126), Expect:2e-08,
Method:Compositional matrix adjust.,
Identities:39/101(39%), Positives:49/101(48%), Gaps:8/101(7%)

Query  1107  CIFFGRLFCCGTSKLRDSPLMQITEKKGIVKRIYSGLTNKLGGVVQFQSSKKPILVKTG*  928
             C    RLF CGTSK  +S +++  E +G VKR Y GL  +  GVVQF + K   LV
Sbjct  170   CADGTRLFSCGTSKEGESFIVEWNESEGAVKRTYLGLGKRSVGVVQFDTMKNKFLVAGDE  229

Query  927   SSFGIWT-ASIF*VQQMPKVG-------LRFDKEGNLLAVS  829
                  W   S+  +      G       LR +KEG LLAVS
Sbjct  230   FQVKFWDMDSVDLLSSTAAEGGLPSSPCLRINKEGTLLAVS  270


Range 2: 147 to 176

Score:26.2 bits(56), Expect:2e-08,
Method:Compositional matrix adjust.,
Identities:13/30(43%), Positives:17/30(56%), Gaps:0/30(0%)

Query  1259  LYDTKCSKEN*KCPSG*CTTLLYNADGNRL  1170
             LYD   S+ +   P   CT++ Y ADG RL
Sbjct  147   LYDNMGSRVDYDAPGRSCTSMAYCADGTRL  176


Query #139: XLOC_015931 Query ID: lcl|Query_33142 Length: 878

Sequences producing significant alignments:
                                                                  Max    Total Query   E   Per.
Description                                                       Score  Score cover Value Ident  Accession
hypothetical protein; 81517-79153 [Arabidopsis thaliana]          78.2   140   34%   1e-26 71.43  AAG12669.1
putative non-LTR retroelement reverse transcriptase [Arabidops... 71.6   139   33%   2e-26 70.45  AAD22368.1
F1L3.4 [Arabidopsis thaliana]                                     73.9   139   33%   2e-26 64.15  AAF79490.1
Hypothetical protein [Arabidopsis thaliana]                       73.9   139   33%   2e-26 64.15  AAF97302.1
hypothetical protein At4g08860 [Arabidopsis thaliana]             71.6   133   35%   1e-24 62.75  ABE65516.1
unknown [Arabidopsis thaliana]                                    71.2   133   35%   2e-24 62.75  ABK28246.1
hypothetical protein AXX17_AT2G19110 [Arabidopsis thaliana]       73.2   127   34%   8e-23 68.75  OAP07776.1
F1E22.12 [Arabidopsis thaliana]                                   68.6   126   34%   1e-22 59.18  AAF23831.1
unknown [Arabidopsis thaliana]                                    70.1   125   33%   3e-22 71.74  ABK28199.1
putative reverse transcriptase [Arabidopsis thaliana]             70.5   125   33%   3e-22 71.74  AAD21515.1
RecName: Full=Putative ribonuclease H protein At1g65750...        68.6   125   34%   4e-22 59.18  P0C2F6.1
F23N19.5 [Arabidopsis thaliana]                                   70.9   124   33%   1e-21 67.39  AAF19536.1
F12A21.24 [Arabidopsis thaliana]                                  69.7   123   34%   1e-21 62.50  AAG28895.1
hypothetical protein At2g27870 [Arabidopsis thaliana]             67.4   122   33%   2e-21 69.57  ABE65462.1
hypothetical protein [Arabidopsis thaliana]                       61.2   122   33%   3e-21 59.09  AAC67202.1
putative non-LTR retroelement reverse transcriptase [Arabidops... 61.2   121   32%   3e-21 59.09  AAC26674.1
non-LTR retroelement reverse transcriptase-like [Arabidopsis...   61.6   121   33%   4e-21 59.09  BAB09815.1
F11A17.8 [Arabidopsis thaliana]                                   60.8   117   33%   5e-20 50.94  AAD49762.2
hypothetical protein At2g15045 [Arabidopsis thaliana]             65.9   115   34%   2e-19 55.10  AAT68735.1
putative non-LTR retroelement reverse transcriptase [Arabidops... 59.7   115   32%   2e-19 54.55  AAC63844.1
F9C16.13 [Arabidopsis thaliana]                                   55.8   109   27%   2e-17 62.86  AAF79665.1
hypothetical protein At1g26950 [Arabidopsis thaliana]             65.1   109   34%   2e-17 61.22  ABE65396.1
unknown [Arabidopsis thaliana]                                    65.1   109   34%   2e-17 61.22  ABK28138.1
Hypothetical protein [Arabidopsis thaliana]                       64.7   108   34%   2e-17 61.22  AAD14501.1
splicing factor like protein [Arabidopsis thaliana]               68.6   108   36%   2e-17 56.36  CAB10336.1
F27F5.12 [Arabidopsis thaliana]                                   48.9   94.3  28%   5e-13 70.97  AAF69156.1
F9C16.18 [Arabidopsis thaliana]                                   52.4   92.8  26%   1e-12 51.92  AAF79682.1
non-LTR retroelement reverse transcriptase-like protein...        68.2   68.2  21%   2e-12 49.25  BAB09192.1
unnamed protein product [Arabidopsis thaliana]                    65.1   65.1  21%   2e-11 47.76  CAA0407116.1
unnamed protein product [Arabidopsis thaliana]                    60.8   60.8  15%   4e-11 63.64  VYS68370.1
Polynucleotidyl transferase, ribonuclease H-like superfamily...   62.8   62.8  20%   8e-11 49.21  NP_680382.1
ftsh7 [Arabidopsis thaliana]                                      63.5   63.5  9%    2e-10 100.00 OAP03029.1
unnamed protein product [Arabidopsis thaliana]                    49.7   85.5  31%   2e-10 56.82  BAB01342.1


Alignments:
>hypothetical protein; 81517-79153 [Arabidopsis thaliana]
Sequence ID: AAG12669.1 Length: 256
Range 1: 73 to 121

Score:78.2 bits(191), Expect:1e-26,
Method:Compositional matrix adjust.,
Identities:35/49(71%), Positives:43/49(87%), Gaps:0/49(0%)

Query  241  KKWVTRLEVEGDSEIVVGFLKSGITNSHPFSFLIRLCYDFLSRGWIVRV  387
            +KWVTR+E+E DSEIVVGFLK+GI +SHP SFL RLCY F+S+ WIVR+
Sbjct  73   EKWVTRVELEVDSEIVVGFLKTGIGDSHPLSFLARLCYGFISKDWIVRI  121


Range 2: 123 to 175

Score:62.0 bits(149), Expect:1e-26,
Method:Compositional matrix adjust.,
Identities:28/53(53%), Positives:37/53(69%), Gaps:0/53(0%)

Query  392  HVHRETNRLTDVLTNYAFSLPLDFYLFEGRPDVVVSIVFDDANGSAYPHNVRI  550
            HV+RE NRL D L NYAF+LPL F+ F   PDVV S+  +D +G++ P  VR+
Sbjct  123  HVYREANRLADGLANYAFTLPLGFHFFHSSPDVVDSVRLEDTSGTSCPRLVRM  175


>putative non-LTR retroelement reverse transcriptase [Arabidopsis thaliana]
Sequence ID: AAD22368.1 Length: 321
Range 1: 224 to 267

Score:71.6 bits(174), Expect:2e-26,
Method:Compositional matrix adjust.,
Identities:31/44(70%), Positives:40/44(90%), Gaps:0/44(0%)

Query  256  RLEVEGDSEIVVGFLKSGITNSHPFSFLIRLCYDFLSRGWIVRV  387
            R+E+E DS++VVGFL +GI +SHP SFL+RLCYDFLS+GWIVR+
Sbjct  224  RVELEVDSKMVVGFLTTGIADSHPLSFLLRLCYDFLSKGWIVRI  267


Range 2: 269 to 321

Score:68.2 bits(165), Expect:2e-26,
Method:Compositional matrix adjust.,
Identities:31/53(58%), Positives:38/53(71%), Gaps:0/53(0%)

Query  392  HVHRETNRLTDVLTNYAFSLPLDFYLFEGRPDVVVSIVFDDANGSAYPHNVRI  550
            HV+RE NRL D L NYAFSL L  +L E RPDVV SI+ DD  G +YP +V++
Sbjct  269  HVYREANRLADGLANYAFSLSLGLHLLESRPDVVSSILLDDVAGVSYPRHVQV  321


>F1L3.4 [Arabidopsis thaliana]
Sequence ID: AAF79490.1 Length: 253
Range 1: 201 to 253

Score:73.9 bits(180), Expect:2e-26,
Method:Compositional matrix adjust.,
Identities:34/53(64%), Positives:40/53(75%), Gaps:0/53(0%)

Query  392  HVHRETNRLTDVLTNYAFSLPLDFYLFEGRPDVVVSIVFDDANGSAYPHNVRI  550
            HV+RE NRL D L NYAF LPL F+LF   PD+V+SIV DD  GSAYP NV++
Sbjct  201  HVYREANRLADGLANYAFFLPLGFHLFNSTPDIVMSIVHDDVAGSAYPRNVQV  253


Range 2: 154 to 199

Score:65.5 bits(158), Expect:2e-26,
Method:Compositional matrix adjust.,
Identities:29/46(63%), Positives:39/46(84%), Gaps:0/46(0%)

Query  250  VTRLEVEGDSEIVVGFLKSGITNSHPFSFLIRLCYDFLSRGWIVRV  387
            VT+LE+E DSE+VVGFL++GI +SHP SFL+RLC+  LS+ W VR+
Sbjct  154  VTQLEMEIDSEMVVGFLRTGIDDSHPLSFLVRLCHGLLSKDWSVRI  199


>Hypothetical protein [Arabidopsis thaliana]
Sequence ID: AAF97302.1 Length: 272
>hypothetical protein AT1G17390 [Arabidopsis thaliana]
Sequence ID: AAV68820.1 Length: 272
Range 1: 220 to 272

Score:73.9 bits(180), Expect:2e-26,
Method:Compositional matrix adjust.,
Identities:34/53(64%), Positives:40/53(75%), Gaps:0/53(0%)

Query  392  HVHRETNRLTDVLTNYAFSLPLDFYLFEGRPDVVVSIVFDDANGSAYPHNVRI  550
            HV+RE NRL D L NYAF LPL F+LF   PD+V+SIV DD  GSAYP NV++
Sbjct  220  HVYREANRLADGLANYAFFLPLGFHLFNSTPDIVMSIVHDDVAGSAYPRNVQV  272


Range 2: 173 to 218

Score:65.5 bits(158), Expect:2e-26,
Method:Compositional matrix adjust.,
Identities:29/46(63%), Positives:39/46(84%), Gaps:0/46(0%)

Query  250  VTRLEVEGDSEIVVGFLKSGITNSHPFSFLIRLCYDFLSRGWIVRV  387
            VT+LE+E DSE+VVGFL++GI +SHP SFL+RLC+  LS+ W VR+
Sbjct  173  VTQLEMEIDSEMVVGFLRTGIDDSHPLSFLVRLCHGLLSKDWSVRI  218


>hypothetical protein At4g08860 [Arabidopsis thaliana]
Sequence ID: ABE65516.1 Length: 106
Range 1: 4 to 52

Score:71.6 bits(174), Expect:1e-24,
Method:Compositional matrix adjust.,
Identities:32/51(63%), Positives:43/51(84%), Gaps:2/51(3%)

Query  235  RCKKWVTRLEVEGDSEIVVGFLKSGITNSHPFSFLIRLCYDFLSRGWIVRV  387
             CK  V++LE+E DSE+VVGFL++GI+ SHP SFL+R+CY F+SR WIVR+
Sbjct  4    ECK--VSQLELEVDSEVVVGFLRTGISESHPLSFLVRMCYGFISRDWIVRI  52


Range 2: 54 to 106

Score:62.0 bits(149), Expect:1e-24,
Method:Compositional matrix adjust.,
Identities:28/53(53%), Positives:36/53(67%), Gaps:0/53(0%)

Query  392  HVHRETNRLTDVLTNYAFSLPLDFYLFEGRPDVVVSIVFDDANGSAYPHNVRI  550
            H +RE NRL D L NYAFSLPL F  F   PD V S++ +D NG ++P +VR+
Sbjct  54   HKYREANRLADGLANYAFSLPLGFQFFYVCPDAVYSVMLEDRNGMSFPRHVRL  106


>unknown, partial [Arabidopsis thaliana]
Sequence ID: ABK28246.1 Length: 107
Range 1: 4 to 52

Score:71.2 bits(173), Expect:2e-24,
Method:Compositional matrix adjust.,
Identities:32/51(63%), Positives:43/51(84%), Gaps:2/51(3%)

Query  235  RCKKWVTRLEVEGDSEIVVGFLKSGITNSHPFSFLIRLCYDFLSRGWIVRV  387
             CK  V++LE+E DSE+VVGFL++GI+ SHP SFL+R+CY F+SR WIVR+
Sbjct  4    ECK--VSQLELEVDSEVVVGFLRTGISESHPLSFLVRMCYGFISRDWIVRI  52


Range 2: 54 to 106

Score:62.0 bits(149), Expect:2e-24,
Method:Compositional matrix adjust.,
Identities:28/53(53%), Positives:36/53(67%), Gaps:0/53(0%)

Query  392  HVHRETNRLTDVLTNYAFSLPLDFYLFEGRPDVVVSIVFDDANGSAYPHNVRI  550
            H +RE NRL D L NYAFSLPL F  F   PD V S++ +D NG ++P +VR+
Sbjct  54   HKYREANRLADGLANYAFSLPLGFQFFYVCPDAVYSVMLEDRNGMSFPRHVRL  106


>hypothetical protein AXX17_AT2G19110 [Arabidopsis thaliana]
Sequence ID: OAP07776.1 Length: 211
Range 1: 110 to 157

Score:73.2 bits(178), Expect:8e-23,
Method:Compositional matrix adjust.,
Identities:33/48(69%), Positives:42/48(87%), Gaps:0/48(0%)

Query  244  KWVTRLEVEGDSEIVVGFLKSGITNSHPFSFLIRLCYDFLSRGWIVRV  387
            K V RLE+E DSE+VVGFLK+GI++SHP SFL+RLC+ F+SR WIVR+
Sbjct  110  KRVERLELEVDSELVVGFLKTGISDSHPLSFLVRLCHGFISRDWIVRI  157


Range 2: 159 to 211

Score:54.3 bits(129), Expect:8e-23,
Method:Compositional matrix adjust.,
Identities:27/53(51%), Positives:34/53(64%), Gaps:0/53(0%)

Query  392  HVHRETNRLTDVLTNYAFSLPLDFYLFEGRPDVVVSIVFDDANGSAYPHNVRI  550
            H++RE+NRL D L NYA+SLPL F+ FE  P  V  IV DD   +    NVR+
Sbjct  159  HMYRESNRLADRLANYAYSLPLSFHAFEACPPCVALIVDDDIRRTTIFRNVRM  211


>F1E22.12 [Arabidopsis thaliana]
Sequence ID: AAF23831.1 Length: 1055
Range 1: 627 to 675

Score:68.6 bits(166), Expect:1e-22,
Method:Compositional matrix adjust.,
Identities:29/49(59%), Positives:42/49(85%), Gaps:0/49(0%)

Query  241  KKWVTRLEVEGDSEIVVGFLKSGITNSHPFSFLIRLCYDFLSRGWIVRV  387
            +K V R+E+E DSE++VGFLK+GI++SHP SFL+RLC+ FL + W+VR+
Sbjct  627  EKKVPRVELEVDSEVIVGFLKTGISDSHPLSFLVRLCHGFLQKDWLVRI  675


Range 2: 677 to 729

Score:58.2 bits(139), Expect:1e-22,
Method:Composition-based stats.,
Identities:27/53(51%), Positives:35/53(66%), Gaps:0/53(0%)

Query  392  HVHRETNRLTDVLTNYAFSLPLDFYLFEGRPDVVVSIVFDDANGSAYPHNVRI  550
            HV+RE NRL D L NYAFSL L F+ F+  PD + S++ +D  GS  P  VR+
Sbjct  677  HVYREANRLADGLANYAFSLSLGFHSFDLVPDAMSSLLREDTLGSTRPRRVRL  729


>unknown, partial [Arabidopsis thaliana]
Sequence ID: ABK28199.1 Length: 315
Range 1: 215 to 260

Score:70.1 bits(170), Expect:3e-22,
Method:Compositional matrix adjust.,
Identities:33/46(72%), Positives:38/46(82%), Gaps:0/46(0%)

Query  250  VTRLEVEGDSEIVVGFLKSGITNSHPFSFLIRLCYDFLSRGWIVRV  387
            VTRLE+E DSEIVVGFLK GI   HP SFL+RLC+DF+SR W VR+
Sbjct  215  VTRLEIEVDSEIVVGFLKIGINEVHPLSFLVRLCHDFISRDWRVRI  260


Range 2: 262 to 314

Score:55.1 bits(131), Expect:3e-22,
Method:Compositional matrix adjust.,
Identities:26/53(49%), Positives:33/53(62%), Gaps:0/53(0%)

Query  392  HVHRETNRLTDVLTNYAFSLPLDFYLFEGRPDVVVSIVFDDANGSAYPHNVRI  550
            HV+RE NRL D L NYAFSLPL F+     PD +  I+ DD +G+     VR+
Sbjct  262  HVYREANRLADGLANYAFSLPLGFHSLSLVPDSLRFILLDDTSGATVSRQVRM  314


>putative reverse transcriptase [Arabidopsis thaliana]
Sequence ID: AAD21515.1 Length: 314
>putative reverse transcriptase [Arabidopsis thaliana]
Sequence ID: AAM15081.1 Length: 314
Range 1: 215 to 260

Score:70.5 bits(171), Expect:3e-22,
Method:Compositional matrix adjust.,
Identities:33/46(72%), Positives:38/46(82%), Gaps:0/46(0%)

Query  250  VTRLEVEGDSEIVVGFLKSGITNSHPFSFLIRLCYDFLSRGWIVRV  387
            VTRLE+E DSEIVVGFLK GI   HP SFL+RLC+DF+SR W VR+
Sbjct  215  VTRLEIEVDSEIVVGFLKIGINEVHPLSFLVRLCHDFISRDWRVRI  260


Range 2: 262 to 314

Score:55.1 bits(131), Expect:3e-22,
Method:Compositional matrix adjust.,
Identities:26/53(49%), Positives:33/53(62%), Gaps:0/53(0%)

Query  392  HVHRETNRLTDVLTNYAFSLPLDFYLFEGRPDVVVSIVFDDANGSAYPHNVRI  550
            HV+RE NRL D L NYAFSLPL F+     PD +  I+ DD +G+     VR+
Sbjct  262  HVYREANRLADGLANYAFSLPLGFHSLSLVPDSLRFILLDDTSGATVSRQVRM  314


>RecName: Full=Putative ribonuclease H protein At1g65750 [Arabidopsis thaliana]
Sequence ID: P0C2F6.1 Length: 620
Range 1: 518 to 566

Score:68.6 bits(166), Expect:4e-22,
Method:Compositional matrix adjust.,
Identities:29/49(59%), Positives:42/49(85%), Gaps:0/49(0%)

Query  241  KKWVTRLEVEGDSEIVVGFLKSGITNSHPFSFLIRLCYDFLSRGWIVRV  387
            +K V R+E+E DSE++VGFLK+GI++SHP SFL+RLC+ FL + W+VR+
Sbjct  518  EKKVPRVELEVDSEVIVGFLKTGISDSHPLSFLVRLCHGFLQKDWLVRI  566


Range 2: 568 to 620

Score:56.6 bits(135), Expect:4e-22,
Method:Compositional matrix adjust.,
Identities:27/53(51%), Positives:35/53(66%), Gaps:0/53(0%)

Query  392  HVHRETNRLTDVLTNYAFSLPLDFYLFEGRPDVVVSIVFDDANGSAYPHNVRI  550
            HV+RE NRL D L NYAFSL L F+ F+  PD + S++ +D  GS  P  VR+
Sbjct  568  HVYREANRLADGLANYAFSLSLGFHSFDLVPDAMSSLLREDTLGSTRPRRVRL  620


>F23N19.5 [Arabidopsis thaliana]
Sequence ID: AAF19536.1 Length: 233
Range 1: 134 to 179

Score:70.9 bits(172), Expect:1e-21,
Method:Compositional matrix adjust.,
Identities:31/46(67%), Positives:41/46(89%), Gaps:0/46(0%)

Query  250  VTRLEVEGDSEIVVGFLKSGITNSHPFSFLIRLCYDFLSRGWIVRV  387
            +TRLE+E DSE+VVGFL++GI +SHP SFL+R+C+ FLSR WIVR+
Sbjct  134  ITRLELEVDSEMVVGFLRTGIGSSHPLSFLVRMCHGFLSRDWIVRI  179


Range 2: 181 to 233

Score:53.1 bits(126), Expect:1e-21,
Method:Compositional matrix adjust.,
Identities:26/53(49%), Positives:34/53(64%), Gaps:0/53(0%)

Query  392  HVHRETNRLTDVLTNYAFSLPLDFYLFEGRPDVVVSIVFDDANGSAYPHNVRI  550
            HV+RE NRL D L NYAF LPL ++ F   P+ + SI+ DD  G + P  VR+
Sbjct  181  HVYREANRLADGLANYAFDLPLGYHAFASPPNSLDSILRDDELGISVPRLVRM  233


>F12A21.24 [Arabidopsis thaliana]
Sequence ID: AAG28895.1 Length: 803
Range 1: 702 to 749

Score:69.7 bits(169), Expect:1e-21,
Method:Compositional matrix adjust.,
Identities:30/48(63%), Positives:40/48(83%), Gaps:0/48(0%)

Query  244  KWVTRLEVEGDSEIVVGFLKSGITNSHPFSFLIRLCYDFLSRGWIVRV  387
            K +TR+E+E DSE+VVGFLK+GI + HP SFL+RLC+  LS+ WIVR+
Sbjct  702  KALTRVELEVDSELVVGFLKTGIGDQHPLSFLVRLCHGLLSKDWIVRI  749


Range 2: 751 to 803

Score:53.5 bits(127), Expect:1e-21,
Method:Compositional matrix adjust.,
Identities:27/53(51%), Positives:34/53(64%), Gaps:0/53(0%)

Query  392  HVHRETNRLTDVLTNYAFSLPLDFYLFEGRPDVVVSIVFDDANGSAYPHNVRI  550
            HV+RE NRL D L NYAFSLPL F+     PD +  I+ +D+ GS  P  VR+
Sbjct  751  HVYREANRLADGLANYAFSLPLGFHSLIDVPDDLEVILHEDSLGSTRPRRVRL  803


>hypothetical protein At2g27870 [Arabidopsis thaliana]
Sequence ID: ABE65462.1 Length: 314
Range 1: 215 to 260

Score:67.4 bits(163), Expect:2e-21,
Method:Compositional matrix adjust.,
Identities:32/46(70%), Positives:37/46(80%), Gaps:0/46(0%)

Query  250  VTRLEVEGDSEIVVGFLKSGITNSHPFSFLIRLCYDFLSRGWIVRV  387
            VTRLE+E DSEIVVGFLK  I   HP SFL+RLC+DF+SR W VR+
Sbjct  215  VTRLEIEVDSEIVVGFLKIXINEVHPLSFLVRLCHDFISRDWRVRI  260


Range 2: 262 to 314

Score:55.1 bits(131), Expect:2e-21,
Method:Compositional matrix adjust.,
Identities:26/53(49%), Positives:33/53(62%), Gaps:0/53(0%)

Query  392  HVHRETNRLTDVLTNYAFSLPLDFYLFEGRPDVVVSIVFDDANGSAYPHNVRI  550
            HV+RE NRL D L NYAFSLPL F+     PD +  I+ DD +G+     VR+
Sbjct  262  HVYREANRLADGLANYAFSLPLGFHSLSLVPDSLRFILLDDTSGATVSRQVRM  314


>hypothetical protein [Arabidopsis thaliana]
Sequence ID: AAC67202.1 Length: 117
Range 1: 20 to 63

Score:61.2 bits(147), Expect:3e-21,
Method:Compositional matrix adjust.,
Identities:26/44(59%), Positives:35/44(79%), Gaps:0/44(0%)

Query  256  RLEVEGDSEIVVGFLKSGITNSHPFSFLIRLCYDFLSRGWIVRV  387
             LE+E DS +VV FLK+GI+ +HP SFL+RLCY  LS+ W+VR+
Sbjct  20   ELELEIDSALVVDFLKTGISETHPLSFLVRLCYGLLSKDWLVRI  63


Range 2: 65 to 117

Score:60.8 bits(146), Expect:3e-21,
Method:Compositional matrix adjust.,
Identities:25/53(47%), Positives:37/53(69%), Gaps:0/53(0%)

Query  392  HVHRETNRLTDVLTNYAFSLPLDFYLFEGRPDVVVSIVFDDANGSAYPHNVRI  550
            HV+RE N+L D L NYAFSLPL  +LF+  PD ++  + +D  G+A+P  +R+
Sbjct  65   HVYREANQLADGLANYAFSLPLGLHLFDSCPDCMLHCLAEDDRGAAHPRQIRV  117


>putative non-LTR retroelement reverse transcriptase [Arabidopsis thaliana]
Sequence ID: AAC26674.1 Length: 970
Range 1: 873 to 916

Score:61.2 bits(147), Expect:3e-21,
Method:Compositional matrix adjust.,
Identities:26/44(59%), Positives:35/44(79%), Gaps:0/44(0%)

Query  256  RLEVEGDSEIVVGFLKSGITNSHPFSFLIRLCYDFLSRGWIVRV  387
            R+E+  DSE+VVGFL +GI+ +HP SFL+RLC  F +R W+VRV
Sbjct  873  RVELNLDSELVVGFLSTGISKAHPLSFLVRLCQGFFTRDWLVRV  916


Range 2: 918 to 968

Score:60.5 bits(145), Expect:3e-21,
Method:Composition-based stats.,
Identities:27/51(53%), Positives:36/51(70%), Gaps:0/51(0%)

Query  392  HVHRETNRLTDVLTNYAFSLPLDFYLFEGRPDVVVSIVFDDANGSAYPHNV  544
            HV+RE NRL D L NYAF LPL F+ FE  P+ V+  + +DANG+++P  V
Sbjct  918  HVYREANRLADGLANYAFFLPLGFHCFEICPEDVLLFLVEDANGTSFPRAV  968


>non-LTR retroelement reverse transcriptase-like [Arabidopsis thaliana]
Sequence ID: BAB09815.1 Length: 676
Range 1: 579 to 622

Score:61.6 bits(148), Expect:4e-21,
Method:Compositional matrix adjust.,
Identities:26/44(59%), Positives:37/44(84%), Gaps:0/44(0%)

Query  256  RLEVEGDSEIVVGFLKSGITNSHPFSFLIRLCYDFLSRGWIVRV  387
            R+ +E DS +VVGFL+SGI +SHP +FL+RLC+ F+S+ WIVR+
Sbjct  579  RVRLEVDSALVVGFLQSGIGDSHPLAFLVRLCHGFISKDWIVRI  622


Range 2: 624 to 676

Score:60.1 bits(144), Expect:4e-21,
Method:Compositional matrix adjust.,
Identities:26/53(49%), Positives:37/53(69%), Gaps:0/53(0%)

Query  392  HVHRETNRLTDVLTNYAFSLPLDFYLFEGRPDVVVSIVFDDANGSAYPHNVRI  550
            HV+RE NRL D L NYAF+LP  F L +  P+ V SI+ +D  G+++P +VR+
Sbjct  624  HVYREANRLADGLANYAFTLPFGFLLLDSCPEHVSSILLEDVMGTSFPRHVRL  676


>F11A17.8 [Arabidopsis thaliana]
Sequence ID: AAD49762.2 Length: 803
Range 1: 54 to 106

Score:60.8 bits(146), Expect:5e-20,
Method:Compositional matrix adjust.,
Identities:27/53(51%), Positives:35/53(66%), Gaps:0/53(0%)

Query  392  HVHRETNRLTDVLTNYAFSLPLDFYLFEGRPDVVVSIVFDDANGSAYPHNVRI  550
            HV+RE NRL D L NY FSLPL F+ FE  P+ V  I+ +D  G+ +P  VR+
Sbjct  54   HVYREANRLADGLANYVFSLPLGFHSFECVPEHVRDILLEDTGGTCFPRQVRL  106


Range 2: 7 to 52

Score:57.0 bits(136), Expect:5e-20,
Method:Compositional matrix adjust.,
Identities:24/46(52%), Positives:36/46(78%), Gaps:0/46(0%)

Query  250  VTRLEVEGDSEIVVGFLKSGITNSHPFSFLIRLCYDFLSRGWIVRV  387
             +R+ +E DSE+VVG L +GI+ ++P SFL+RLC+ F+ R WIVR+
Sbjct  7    ASRVVIEVDSELVVGLLTTGISEANPLSFLVRLCHGFILRDWIVRI  52


>hypothetical protein At2g15045 [Arabidopsis thaliana]
Sequence ID: AAT68735.1 Length: 112
>hypothetical protein At2g15045 [Arabidopsis thaliana]
Sequence ID: AAV63873.1 Length: 112
Range 1: 10 to 58

Score:65.9 bits(159), Expect:2e-19,
Method:Compositional matrix adjust.,
Identities:27/49(55%), Positives:41/49(83%), Gaps:0/49(0%)

Query  241  KKWVTRLEVEGDSEIVVGFLKSGITNSHPFSFLIRLCYDFLSRGWIVRV  387
            +K  +R+E+E D+E+VVGFL++ ++ SHP SFL+RLCY F+S+ WIVR+
Sbjct  10   EKKASRVELEVDTEMVVGFLRTRVSESHPLSFLVRLCYGFISKDWIVRI  58


Range 2: 60 to 110

Score:49.7 bits(117), Expect:2e-19,
Method:Compositional matrix adjust.,
Identities:25/51(49%), Positives:31/51(60%), Gaps:0/51(0%)

Query  392  HVHRETNRLTDVLTNYAFSLPLDFYLFEGRPDVVVSIVFDDANGSAYPHNV  544
            HV+RE NRL D L NYAFSL + F  F+  PD V SI   D +G+    +V
Sbjct  60   HVYREANRLADGLANYAFSLSVCFDYFDSLPDCVCSISAADTSGATSSRHV  110


>putative non-LTR retroelement reverse transcriptase [Arabidopsis thaliana]
Sequence ID: AAC63844.1 Length: 1231
Range 1: 1134 to 1177

Score:59.7 bits(143), Expect:2e-19,
Method:Compositional matrix adjust.,
Identities:24/44(55%), Positives:36/44(81%), Gaps:0/44(0%)

Query  256   RLEVEGDSEIVVGFLKSGITNSHPFSFLIRLCYDFLSRGWIVRV  387
             R+E++ D ++VVGFL +G++N+HP SFL+RLC  F +R W+VRV
Sbjct  1134  RVELDLDCKLVVGFLSTGVSNAHPLSFLVRLCQGFFTRDWLVRV  1177


Range 2: 1179 to 1229

Score:55.8 bits(133), Expect:2e-19,
Method:Composition-based stats.,
Identities:24/51(47%), Positives:33/51(64%), Gaps:0/51(0%)

Query  392   HVHRETNRLTDVLTNYAFSLPLDFYLFEGRPDVVVSIVFDDANGSAYPHNV  544
             HV+RE NRL D L NYAF+LPL  + F+  P+ V  ++  D NG+ +P  V
Sbjct  1179  HVYREANRLADGLANYAFTLPLGLHCFDACPEGVRLLLLADVNGTEFPRAV  1229


>F9C16.13 [Arabidopsis thaliana]
Sequence ID: AAF79665.1 Length: 1172
Range 1: 263 to 297

Score:55.8 bits(133), Expect:2e-17,
Method:Compositional matrix adjust.,
Identities:22/35(63%), Positives:30/35(85%), Gaps:0/35(0%)

Query  283  IVVGFLKSGITNSHPFSFLIRLCYDFLSRGWIVRV  387
            +VVGFLK+G++  HP SFL+RLC+ FLS+ WIVR+
Sbjct  263  LVVGFLKTGVSERHPLSFLVRLCHGFLSKDWIVRI  297


Range 2: 299 to 344

Score:53.5 bits(127), Expect:2e-17,
Method:Composition-based stats.,
Identities:25/46(54%), Positives:30/46(65%), Gaps:0/46(0%)

Query  392  HVHRETNRLTDVLTNYAFSLPLDFYLFEGRPDVVVSIVFDDANGSA  529
            HV+RE NRL D L NYAF LP+  ++F   PDVV  +  DD  GSA
Sbjct  299  HVYREANRLADGLANYAFLLPIGIHVFVSVPDVVEPLFSDDVIGSA  344


>hypothetical protein At1g26950 [Arabidopsis thaliana]
Sequence ID: ABE65396.1 Length: 162
Range 1: 60 to 108

Score:65.1 bits(157), Expect:2e-17,
Method:Compositional matrix adjust.,
Identities:30/49(61%), Positives:39/49(79%), Gaps:0/49(0%)

Query  241  KKWVTRLEVEGDSEIVVGFLKSGITNSHPFSFLIRLCYDFLSRGWIVRV  387
            +K +TRLE+E DS++V GFL +GI +SH  SFL+RLCY F S+ WIVRV
Sbjct  60   EKGITRLELEVDSKLVAGFLTTGIEDSHLLSFLVRLCYGFSSKDWIVRV  108


Range 2: 110 to 160

Score:44.3 bits(103), Expect:2e-17,
Method:Compositional matrix adjust.,
Identities:25/51(49%), Positives:30/51(58%), Gaps:0/51(0%)

Query  392  HVHRETNRLTDVLTNYAFSLPLDFYLFEGRPDVVVSIVFDDANGSAYPHNV  544
            HV+RE NR  D L NYAFSL L F  F+     V SI+ +DA G A   +V
Sbjct  110  HVYREANRFADELANYAFSLLLGFLSFDSGLPFVDSIMREDAAGVAGARHV  160


>unknown, partial [Arabidopsis thaliana]
Sequence ID: ABK28138.1 Length: 163
Range 1: 60 to 108

Score:65.1 bits(157), Expect:2e-17,
Method:Compositional matrix adjust.,
Identities:30/49(61%), Positives:39/49(79%), Gaps:0/49(0%)

Query  241  KKWVTRLEVEGDSEIVVGFLKSGITNSHPFSFLIRLCYDFLSRGWIVRV  387
            +K +TRLE+E DS++V GFL +GI +SH  SFL+RLCY F S+ WIVRV
Sbjct  60   EKGITRLELEVDSKLVAGFLTTGIEDSHLLSFLVRLCYGFSSKDWIVRV  108


Range 2: 110 to 160

Score:44.3 bits(103), Expect:2e-17,
Method:Compositional matrix adjust.,
Identities:25/51(49%), Positives:30/51(58%), Gaps:0/51(0%)

Query  392  HVHRETNRLTDVLTNYAFSLPLDFYLFEGRPDVVVSIVFDDANGSAYPHNV  544
            HV+RE NR  D L NYAFSL L F  F+     V SI+ +DA G A   +V
Sbjct  110  HVYREANRFADELANYAFSLLLGFLSFDSGLPFVDSIMREDAAGVAGARHV  160


>Hypothetical protein [Arabidopsis thaliana]
Sequence ID: AAD14501.1 Length: 158
Range 1: 56 to 104

Score:64.7 bits(156), Expect:2e-17,
Method:Compositional matrix adjust.,
Identities:30/49(61%), Positives:39/49(79%), Gaps:0/49(0%)

Query  241  KKWVTRLEVEGDSEIVVGFLKSGITNSHPFSFLIRLCYDFLSRGWIVRV  387
            +K +TRLE+E DS++V GFL +GI +SH  SFL+RLCY F S+ WIVRV
Sbjct  56   EKGITRLELEVDSKLVAGFLTTGIEDSHLLSFLVRLCYGFSSKDWIVRV  104


Range 2: 106 to 156

Score:44.3 bits(103), Expect:2e-17,
Method:Compositional matrix adjust.,
Identities:25/51(49%), Positives:30/51(58%), Gaps:0/51(0%)

Query  392  HVHRETNRLTDVLTNYAFSLPLDFYLFEGRPDVVVSIVFDDANGSAYPHNV  544
            HV+RE NR  D L NYAFSL L F  F+     V SI+ +DA G A   +V
Sbjct  106  HVYREANRFADELANYAFSLLLGFLSFDSGLPFVDSIMREDAAGVAGARHV  156


>splicing factor like protein [Arabidopsis thaliana]
Sequence ID: CAB10336.1 Length: 559
>splicing factor like protein [Arabidopsis thaliana]
Sequence ID: CAB78600.1 Length: 559
Range 1: 74 to 128

Score:68.6 bits(166), Expect:2e-17,
Method:Compositional matrix adjust.,
Identities:31/55(56%), Positives:43/55(78%), Gaps:0/55(0%)

Query  223  FVRRRCKKWVTRLEVEGDSEIVVGFLKSGITNSHPFSFLIRLCYDFLSRGWIVRV  387
            FVR R +    R+E+E DSE+VVGFL+SGI+ +HP +FL+RLC+  LS+ W+VRV
Sbjct  74   FVRLRWQSCGGRVELEVDSELVVGFLQSGISEAHPLAFLVRLCHGLLSKDWLVRV  128


Range 2: 130 to 164

Score:40.4 bits(93), Expect:2e-17,
Method:Compositional matrix adjust.,
Identities:22/52(42%), Positives:27/52(51%), Gaps:17/52(32%)

Query  392  HVHRETNRLTDVLTNYAFSLPLDFYLFEGRPDVVVSIVFDDANGSAYPHNVR  547
            HV+RE NRL D L NYAFSL                  F+DA G+A+P + R
Sbjct  130  HVYREANRLADGLANYAFSLQ-----------------FEDATGTAFPRHGR  164


>F27F5.12 [Arabidopsis thaliana]
Sequence ID: AAF69156.1 Length: 205
Range 1: 119 to 149

Score:48.9 bits(115), Expect:5e-13,
Method:Compositional matrix adjust.,
Identities:22/31(71%), Positives:27/31(87%), Gaps:0/31(0%)

Query  250  VTRLEVEGDSEIVVGFLKSGITNSHPFSFLI  342
            +TRLEVE DSE+VVGFL  GI++SHP SF+I
Sbjct  119  ITRLEVEVDSELVVGFLSKGISDSHPLSFVI  149


Range 2: 154 to 205

Score:45.4 bits(106), Expect:5e-13,
Method:Compositional matrix adjust.,
Identities:25/53(47%), Positives:34/53(64%), Gaps:1/53(1%)

Query  392  HVHRETNRLTDVLTNYAFSLPLDFYLFEGRPDVVVSIVFDDANGSAYPHNVRI  550
            HV+RE NRL D L NYAF+LPL F++ +  P  V +I  DD    A+  +VR+
Sbjct  154  HVYREANRLADELENYAFTLPLTFHMLKVVPH-VETIFQDDDREIAFLRHVRL  205


>F9C16.18 [Arabidopsis thaliana]
Sequence ID: AAF79682.1 Length: 196
Range 1: 142 to 193

Score:52.4 bits(124), Expect:1e-12,
Method:Compositional matrix adjust.,
Identities:27/52(52%), Positives:32/52(61%), Gaps:0/52(0%)

Query  392  HVHRETNRLTDVLTNYAFSLPLDFYLFEGRPDVVVSIVFDDANGSAYPHNVR  547
            HV+RE NRL D L NYAF LP+  ++F   PDVV  +  DD  GSA    VR
Sbjct  142  HVYREANRLADGLANYAFLLPIGIHVFVSVPDVVEPLFSDDVIGSARLRRVR  193


Range 2: 114 to 140

Score:40.4 bits(93), Expect:1e-12,
Method:Compositional matrix adjust.,
Identities:15/27(56%), Positives:21/27(77%), Gaps:0/27(0%)

Query  307  GITNSHPFSFLIRLCYDFLSRGWIVRV  387
            G++  HP SFL+RLC+ FL + WIVR+
Sbjct  114  GVSERHPLSFLVRLCHGFLLKDWIVRI  140


>non-LTR retroelement reverse transcriptase-like protein [Arabidopsis thaliana]
Sequence ID: BAB09192.1 Length: 308
Range 1: 188 to 254

Score:68.2 bits(165), Expect:2e-12,
Method:Compositional matrix adjust.,
Identities:33/67(49%), Positives:49/67(73%), Gaps:3/67(4%)

Query  196  SNQHSE---TFLFVRRRCKKWVTRLEVEGDSEIVVGFLKSGITNSHPFSFLIRLCYDFLS  366
            S QH+E    +  +    +K V R+E+E DSE +VGFLK+GI++SHP SFL+RLC++FL
Sbjct  188  SAQHAELWGVYYGLYFAWEKKVPRVELEVDSEAIVGFLKTGISDSHPLSFLVRLCHNFLQ  247

Query  367  RGWIVRV  387
            + W+VR+
Sbjct  248  KDWLVRI  254


>unnamed protein product [Arabidopsis thaliana]
Sequence ID: CAA0407116.1 Length: 308
Range 1: 188 to 254

Score:65.1 bits(157), Expect:2e-11,
Method:Compositional matrix adjust.,
Identities:32/67(48%), Positives:48/67(71%), Gaps:3/67(4%)

Query  196  SNQHSE---TFLFVRRRCKKWVTRLEVEGDSEIVVGFLKSGITNSHPFSFLIRLCYDFLS  366
            S QH+E    +  +    +K V R+E+E DSE +VGFLK+ I++SHP SFL+RLC++FL
Sbjct  188  SAQHAELWGVYYGLYFAWEKKVPRVELEVDSEAIVGFLKTWISDSHPLSFLVRLCHNFLQ  247

Query  367  RGWIVRV  387
            + W+VR+
Sbjct  248  KDWLVRI  254


>unnamed protein product [Arabidopsis thaliana]
Sequence ID: VYS68370.1 Length: 111
Range 1: 64 to 107

Score:60.8 bits(146), Expect:4e-11,
Method:Compositional matrix adjust.,
Identities:28/44(64%), Positives:37/44(84%), Gaps:0/44(0%)

Query  256  RLEVEGDSEIVVGFLKSGITNSHPFSFLIRLCYDFLSRGWIVRV  387
            R+E+E DSE+VV FL  GI+++HP SFL+RLC+DFLSR W+V V
Sbjct  64   RVELEVDSELVVVFLTRGISDTHPLSFLVRLCHDFLSRDWLVCV  107


>Polynucleotidyl transferase, ribonuclease H-like superfamily protein [Arabidopsis thaliana]
Sequence ID: NP_680382.1 Length: 258
>Polynucleotidyl transferase, ribonuclease H-like superfamily protein [Arabidopsis thaliana]
Sequence ID: AED94885.1 Length: 258
Range 1: 188 to 250

Score:62.8 bits(151), Expect:8e-11,
Method:Compositional matrix adjust.,
Identities:31/63(49%), Positives:45/63(71%), Gaps:3/63(4%)

Query  196  SNQHSE---TFLFVRRRCKKWVTRLEVEGDSEIVVGFLKSGITNSHPFSFLIRLCYDFLS  366
            S QH+E    +  +    +K V R+E+E DSE +VGFLK+GI++SHP SFL+RLC++FL
Sbjct  188  SAQHAELWGVYYGLYFAWEKKVPRVELEVDSEAIVGFLKTGISDSHPLSFLVRLCHNFLQ  247

Query  367  RGW  375
            + W
Sbjct  248  KDW  250


>ftsh7 [Arabidopsis thaliana]
Sequence ID: OAP03029.1 Length: 851
Range 1: 18 to 45

Score:63.5 bits(153), Expect:2e-10,
Method:Compositional matrix adjust.,
Identities:28/28(100%), Positives:28/28(100%), Gaps:0/28(0%)

Query  458  DFYLFEGRPDVVVSIVFDDANGSAYPHN  541
            DFYLFEGRPDVVVSIVFDDANGSAYPHN
Sbjct  18   DFYLFEGRPDVVVSIVFDDANGSAYPHN  45


>unnamed protein product [Arabidopsis thaliana]
Sequence ID: BAB01342.1 Length: 138
Range 1: 86 to 129

Score:49.7 bits(117), Expect:2e-10,
Method:Compositional matrix adjust.,
Identities:25/44(57%), Positives:32/44(72%), Gaps:0/44(0%)

Query  392  HVHRETNRLTDVLTNYAFSLPLDFYLFEGRPDVVVSIVFDDANG  523
            HV+R+ NRLTD LTN+AFSL L F+ F   P  VVSI+ +D +G
Sbjct  86   HVYRKANRLTDGLTNHAFSLLLGFHAFASVPVEVVSILREDVDG  129


Range 2: 33 to 84

Score:35.8 bits(81), Expect:2e-10,
Method:Compositional matrix adjust.,
Identities:28/52(54%), Positives:37/52(71%), Gaps:3/52(5%)

Query  241  KKWVTRLEVEGDSEIVVGFLKSG---ITNSHPFSFLIRLCYDFLSRGWIVRV  387
            +K + RLEVE DSE+VVGFL +G   I  ++P S L+RLC+ FL +  IVRV
Sbjct  33   EKKIMRLEVEVDSEVVVGFLTTGIEEIEETNPLSILVRLCHGFLRKDSIVRV  84


Query #140: XLOC_016000 Query ID: lcl|Query_33143 Length: 882

No significant similarity found.


Query #141: XLOC_016013 Query ID: lcl|Query_33144 Length: 827

No significant similarity found.


Query #142: XLOC_016266 Query ID: lcl|Query_33145 Length: 817

Sequences producing significant alignments:
                                                                  Max    Total Query   E   Per.
Description                                                       Score  Score cover Value Ident  Accession
D-galactoside/L-rhamnose binding SUEL lectin protein...           116    116   53%   1e-31 48.78  NP_001319734.1
unnamed protein product [Arabidopsis thaliana]                    111    111   34%   1e-29 57.89  VYS60238.1
D-galactoside/L-rhamnose binding SUEL lectin protein...           109    109   34%   5e-29 57.89  NP_001118829.1
D-galactoside/L-rhamnose binding SUEL lectin protein...           110    110   34%   6e-29 57.89  NP_001326653.1
D-galactoside/L-rhamnose binding SUEL lectin protein...           94.0   94.0  30%   4e-23 59.52  NP_190876.1
unknown [Arabidopsis thaliana]                                    94.0   94.0  30%   4e-23 59.52  ABK28602.1
unnamed protein product [Arabidopsis thaliana]                    92.4   92.4  30%   1e-22 58.33  CAA0386054.1
unnamed protein product [Arabidopsis thaliana]                    91.7   91.7  30%   3e-22 58.33  VYS60239.1
unnamed protein product [Arabidopsis thaliana]                    89.7   89.7  29%   1e-21 57.50  CAA0386044.1
unknown [Arabidopsis thaliana]                                    89.4   89.4  30%   3e-21 57.14  AAM63449.1
D-galactoside/L-rhamnose binding SUEL lectin protein...           79.0   79.0  24%   1e-17 57.35  NP_190873.1
hypothetical protein AXX17_AT3G47460 [Arabidopsis thaliana]       79.7   79.7  24%   8e-17 57.35  OAP06208.1
hypothetical protein AXX17_AT3G47470 [Arabidopsis thaliana]       68.2   68.2  23%   6e-14 56.72  OAP04887.1
Putative membrane lipoprotein [Arabidopsis thaliana]              62.8   62.8  37%   5e-12 39.22  NP_190875.1
unnamed protein product [Arabidopsis thaliana]                    59.7   59.7  15%   9e-11 68.29  CAA0386050.1


Alignments:
>D-galactoside/L-rhamnose binding SUEL lectin protein [Arabidopsis thaliana]
Sequence ID: NP_001319734.1 Length: 162
>D-galactoside/L-rhamnose binding SUEL lectin protein [Arabidopsis thaliana]
Sequence ID: AEE79031.2 Length: 162 >unnamed protein product [Arabidopsis thaliana]
Sequence ID: CAA0386048.1 Length: 162
Range 1: 1 to 161

Score:116 bits(290), Expect:1e-31,
Method:Compositional matrix adjust.,
Identities:80/164(49%), Positives:100/164(60%), Gaps:21/164(12%)

Query  191  METTHSLHRHGFIvlllllsvsyvsCFYTRNIINGSYRL-------NPREY---------  322
            MET+H   RH FI+LLL++ +S VS   ++ I N S+         +P+
Sbjct  1    METSHYFRRHCFIILLLVMFLSSVSNLASK-INNSSFDARGTKINSDPKHISNSKSGGSP  59

Query  323  -RPNQLPIC-IDHPAPSVDLSPIFCGEGYVISEIKFADYGQPTGDCKNNTLKRGNCGAPA  496
             R  + P+C  ++P+    +   FC +GYV S IKFADYGQP G     TLKRGNCGAPA
Sbjct  60   QRDKEYPLCGSNNPSEDGIIYAPFCDKGYVFSRIKFADYGQPGGS-SCETLKRGNCGAPA  118

Query  497  TLRLVKKN*LGKEECWFPVTDEMFGPTHCKGPALRFVVSGTCKK  628
            TLRLVK+N LGKE C   +TDEMFGPTHCKGP + FV S  CKK
Sbjct  119  TLRLVKENCLGKERCRIYITDEMFGPTHCKGP-VSFVFSAICKK  161


>unnamed protein product [Arabidopsis thaliana]
Sequence ID: VYS60238.1 Length: 165
Range 1: 73 to 164

Score:111 bits(278), Expect:1e-29,
Method:Compositional matrix adjust.,
Identities:55/95(58%), Positives:67/95(70%), Gaps:3/95(3%)

Query  344  CIDHPAPSVDLSPIFCGEGYVISEIKFADYGQPTGDCKNNTLKRGNCGAPATLRLVKKN*  523
            C +H +    ++ IFC +GYVI+ + FADYG PTG C++   + GNCGAPATLRLVKKN
Sbjct  73   CTNHKSVRGPITRIFCQDGYVITNVNFADYGNPTGTCEH--FRHGNCGAPATLRLVKKNC  130

Query  524  LGKEECWFPVTDEMFGPTHCKGPALRFVVSGTCKK  628
            LGK +C F VTDEMFGP+HCKGP     V  TC K
Sbjct  131  LGKPKCVFLVTDEMFGPSHCKGPPT-LAVDATCTK  164


>D-galactoside/L-rhamnose binding SUEL lectin protein [Arabidopsis thaliana]
Sequence ID: NP_001118829.1 Length: 165
>D-galactoside/L-rhamnose binding SUEL lectin protein [Arabidopsis thaliana]
Sequence ID: AEE79033.1 Length: 165 >unnamed protein product [Arabidopsis thaliana]
Sequence ID: CAA0386052.1 Length: 165
Range 1: 73 to 164

Score:109 bits(273), Expect:5e-29,
Method:Compositional matrix adjust.,
Identities:55/95(58%), Positives:66/95(69%), Gaps:3/95(3%)

Query  344  CIDHPAPSVDLSPIFCGEGYVISEIKFADYGQPTGDCKNNTLKRGNCGAPATLRLVKKN*  523
            C +H +    ++ IFC +GYVI+ I FADYG PTG C++   + G CGAPATLRLVKKN
Sbjct  73   CTNHKSVRGPITRIFCQDGYVITNINFADYGNPTGTCEH--FRHGKCGAPATLRLVKKNC  130

Query  524  LGKEECWFPVTDEMFGPTHCKGPALRFVVSGTCKK  628
            LGK +C F VTDEMFGP+HCKGP     V  TC K
Sbjct  131  LGKPKCVFLVTDEMFGPSHCKGPPT-LAVDATCTK  164


>D-galactoside/L-rhamnose binding SUEL lectin protein [Arabidopsis thaliana]
Sequence ID: NP_001326653.1 Length: 195
>D-galactoside/L-rhamnose binding SUEL lectin protein [Arabidopsis thaliana]
Sequence ID: ANM64640.1 Length: 195
Range 1: 103 to 194

Score:110 bits(275), Expect:6e-29,
Method:Compositional matrix adjust.,
Identities:55/95(58%), Positives:66/95(69%), Gaps:3/95(3%)

Query  344  CIDHPAPSVDLSPIFCGEGYVISEIKFADYGQPTGDCKNNTLKRGNCGAPATLRLVKKN*  523
            C +H +    ++ IFC +GYVI+ I FADYG PTG C++   + G CGAPATLRLVKKN
Sbjct  103  CTNHKSVRGPITRIFCQDGYVITNINFADYGNPTGTCEH--FRHGKCGAPATLRLVKKNC  160

Query  524  LGKEECWFPVTDEMFGPTHCKGPALRFVVSGTCKK  628
            LGK +C F VTDEMFGP+HCKGP     V  TC K
Sbjct  161  LGKPKCVFLVTDEMFGPSHCKGPPT-LAVDATCTK  194


>D-galactoside/L-rhamnose binding SUEL lectin protein [Arabidopsis thaliana]
Sequence ID: NP_190876.1 Length: 155
>galactose-binding lectin family protein [Arabidopsis thaliana]
Sequence ID: ABE66016.1 Length: 155 >D-galactoside/L-rhamnose binding SUEL lectin protein [Arabidopsis thaliana]
Sequence ID: AEE79034.1 Length: 155 >putative protein [Arabidopsis thaliana]
Sequence ID: CAB64211.1 Length: 155
Range 1: 72 to 152

Score:94.0 bits(232), Expect:4e-23,
Method:Compositional matrix adjust.,
Identities:50/84(60%), Positives:60/84(71%), Gaps:4/84(4%)

Query  374  LSPIFCGE-GYVISEIKFADYGQPTGDCKNNTLKRGNCGAPATLRLVKKN*LGKEECWFP  550
            L+ I C E GYVI++I FADYG PTG C +   +RGNCGA AT+R+VKKN LGKE+C
Sbjct  72   LTRISCNEPGYVITKINFADYGNPTGTCGH--FRRGNCGARATMRIVKKNCLGKEKCHLL  129

Query  551  VTDEMFGPTHCKGPALRFVVSGTC  622
            VTDEMFGP+ CKG  +   V  TC
Sbjct  130  VTDEMFGPSKCKGAPM-LAVETTC  152


>unknown, partial [Arabidopsis thaliana]
Sequence ID: ABK28602.1 Length: 156
Range 1: 72 to 152

Score:94.0 bits(232), Expect:4e-23,
Method:Compositional matrix adjust.,
Identities:50/84(60%), Positives:60/84(71%), Gaps:4/84(4%)

Query  374  LSPIFCGE-GYVISEIKFADYGQPTGDCKNNTLKRGNCGAPATLRLVKKN*LGKEECWFP  550
            L+ I C E GYVI++I FADYG PTG C +   +RGNCGA AT+R+VKKN LGKE+C
Sbjct  72   LTRISCNEPGYVITKINFADYGNPTGTCGH--FRRGNCGARATMRIVKKNCLGKEKCHLL  129

Query  551  VTDEMFGPTHCKGPALRFVVSGTC  622
            VTDEMFGP+ CKG  +   V  TC
Sbjct  130  VTDEMFGPSKCKGAPM-LAVETTC  152


>unnamed protein product [Arabidopsis thaliana]
Sequence ID: CAA0386054.1 Length: 155
Range 1: 72 to 152

Score:92.4 bits(228), Expect:1e-22,
Method:Compositional matrix adjust.,
Identities:49/84(58%), Positives:60/84(71%), Gaps:4/84(4%)

Query  374  LSPIFCGE-GYVISEIKFADYGQPTGDCKNNTLKRGNCGAPATLRLVKKN*LGKEECWFP  550
            L+ I C E GYVI++I FADYG PTG C +   +RGNCGA AT+R+VKKN LGK++C
Sbjct  72   LTRISCNEPGYVITKINFADYGNPTGTCGH--FRRGNCGARATMRIVKKNCLGKKKCHLL  129

Query  551  VTDEMFGPTHCKGPALRFVVSGTC  622
            VTDEMFGP+ CKG  +   V  TC
Sbjct  130  VTDEMFGPSKCKGAPM-LAVETTC  152


>unnamed protein product [Arabidopsis thaliana]
Sequence ID: VYS60239.1 Length: 155
Range 1: 72 to 152

Score:91.7 bits(226), Expect:3e-22,
Method:Compositional matrix adjust.,
Identities:49/84(58%), Positives:59/84(70%), Gaps:4/84(4%)

Query  374  LSPIFCGE-GYVISEIKFADYGQPTGDCKNNTLKRGNCGAPATLRLVKKN*LGKEECWFP  550
            L+ I C E GYVI++I FADYG PTG C +   +R NCGA AT+R+VKKN LGKE+C
Sbjct  72   LTRISCNEPGYVITKINFADYGNPTGTCGH--FRRDNCGARATMRIVKKNCLGKEKCHLL  129

Query  551  VTDEMFGPTHCKGPALRFVVSGTC  622
            VTDEMFGP+ CKG  +   V  TC
Sbjct  130  VTDEMFGPSKCKGAPM-LAVETTC  152


>unnamed protein product [Arabidopsis thaliana]
Sequence ID: CAA0386044.1 Length: 154
>unnamed protein product [Arabidopsis thaliana]
Sequence ID: VYS60236.1 Length: 154
Range 1: 77 to 153

Score:89.7 bits(221), Expect:1e-21,
Method:Compositional matrix adjust.,
Identities:46/80(58%), Positives:56/80(70%), Gaps:3/80(3%)

Query  389  CGEGYVISEIKFADYGQPTGDCKNNTLKRGNCGAPATLRLVKKN*LGKEECWFPVTDEMF  568
            C +GYVIS+I +ADYGQ TG C     KRGNCGA  TL +VKK  L KE+C   V D++F
Sbjct  77   CEQGYVISKITYADYGQSTGSC--GKFKRGNCGASNTLNIVKKKCLRKEKCKLFVPDKIF  134

Query  569  GPTHCKGPALRFVVSGTCKK  628
            GP+HCKG AL  V+  TC+K
Sbjct  135  GPSHCKG-ALSLVIDATCRK  153


>unknown [Arabidopsis thaliana]
Sequence ID: AAM63449.1 Length: 155
Range 1: 72 to 152

Score:89.4 bits(220), Expect:3e-21,
Method:Compositional matrix adjust.,
Identities:48/84(57%), Positives:58/84(69%), Gaps:4/84(4%)

Query  374  LSPIFCGE-GYVISEIKFADYGQPTGDCKNNTLKRGNCGAPATLRLVKKN*LGKEECWFP  550
            L+ I C E  YVI++I FADYG PTG C +   +R NCGA AT+R+VKKN LGKE+C
Sbjct  72   LTRISCNEPEYVITKINFADYGNPTGTCGH--FRRDNCGARATMRIVKKNCLGKEKCHLL  129

Query  551  VTDEMFGPTHCKGPALRFVVSGTC  622
            VTDEMFGP+ CKG  +   V  TC
Sbjct  130  VTDEMFGPSKCKGAPM-LAVETTC  152


>D-galactoside/L-rhamnose binding SUEL lectin protein [Arabidopsis thaliana]
Sequence ID: NP_190873.1 Length: 142
>D-galactoside/L-rhamnose binding SUEL lectin protein [Arabidopsis thaliana]
Sequence ID: AEE79029.1 Length: 142 >putative protein [Arabidopsis thaliana]
Sequence ID: CAB86909.1 Length: 142
Range 1: 77 to 142

Score:79.0 bits(193), Expect:1e-17,
Method:Compositional matrix adjust.,
Identities:39/68(57%), Positives:47/68(69%), Gaps:2/68(2%)

Query  389  CGEGYVISEIKFADYGQPTGDCKNNTLKRGNCGAPATLRLVKKN*LGKEECWFPVTDEMF  568
            C +GYVIS+I +ADYGQ TG C     KRGNCGA  TL +V K  L KE+C   V D++F
Sbjct  77   CEQGYVISKITYADYGQSTGSC--GKFKRGNCGASNTLNIVNKKCLRKEKCKLFVPDKIF  134

Query  569  GPTHCKGP  592
            GP+HCKG
Sbjct  135  GPSHCKGA  142


>hypothetical protein AXX17_AT3G47460 [Arabidopsis thaliana]
Sequence ID: OAP06208.1 Length: 264
Range 1: 77 to 142

Score:79.7 bits(195), Expect:8e-17,
Method:Compositional matrix adjust.,
Identities:39/68(57%), Positives:47/68(69%), Gaps:2/68(2%)

Query  389  CGEGYVISEIKFADYGQPTGDCKNNTLKRGNCGAPATLRLVKKN*LGKEECWFPVTDEMF  568
            C +GYVIS+I +ADYGQ TG C     KRGNCGA  TL +V K  L KE+C   V D++F
Sbjct  77   CEQGYVISKITYADYGQSTGSC--GKFKRGNCGASNTLNIVNKKCLRKEKCKLFVPDKIF  134

Query  569  GPTHCKGP  592
            GP+HCKG
Sbjct  135  GPSHCKGA  142


>hypothetical protein AXX17_AT3G47470 [Arabidopsis thaliana]
Sequence ID: OAP04887.1 Length: 114
Range 1: 43 to 107

Score:68.2 bits(165), Expect:6e-14,
Method:Compositional matrix adjust.,
Identities:38/67(57%), Positives:47/67(70%), Gaps:4/67(5%)

Query  323  RPNQLPIC-IDHPAPSVDLSPIFCGEGYVISEIKFADYGQPTG-DCKNNTLKRGNCGAPA  496
            R  + P+C  ++P+    +   FC +GYV S IKFADYGQP G  C+  TLKRGNCGAPA
Sbjct  43   RDKEYPLCGSNNPSEDGIIYAPFCDKGYVFSRIKFADYGQPGGSSCE--TLKRGNCGAPA  100

Query  497  TLRLVKK  517
            TLRLVK+
Sbjct  101  TLRLVKE  107


>Putative membrane lipoprotein [Arabidopsis thaliana]
Sequence ID: NP_190875.1 Length: 105
>Putative membrane lipoprotein [Arabidopsis thaliana]
Sequence ID: AEE79032.1 Length: 105 >putative protein [Arabidopsis thaliana]
Sequence ID: CAB88332.1 Length: 105
Range 1: 8 to 98

Score:62.8 bits(151), Expect:5e-12,
Method:Compositional matrix adjust.,
Identities:40/102(39%), Positives:55/102(53%), Gaps:12/102(11%)

Query  215  RHGFIvlllllsvsyvsCFYTRNIINGSYRLNPREYRPNQLPICIDHPAPSVDLSPIFCG  394
            RHGFI++L+          +  +I   + +++      NQ+P+C         +    C
Sbjct  8    RHGFILVLV---------LFLTSISCLASKMDFSSKLGNQIPVCRHQNRSRAPVLMFDCK  58

Query  395  E-GYVISEIKFADYGQPTGDCKNNTLKRGNCGAPATLRLVKK  517
            E GYV S+I FADYG  +GDC N   +RG CGAP TLRLVKK
Sbjct  59   EKGYVFSKINFADYGHSSGDCGN--FRRGTCGAPDTLRLVKK  98


>unnamed protein product [Arabidopsis thaliana]
Sequence ID: CAA0386050.1 Length: 124
Range 1: 79 to 117

Score:59.7 bits(143), Expect:9e-11,
Method:Compositional matrix adjust.,
Identities:28/41(68%), Positives:32/41(78%), Gaps:2/41(4%)

Query  395  EGYVISEIKFADYGQPTGDCKNNTLKRGNCGAPATLRLVKK  517
            +GYV S+I FADYG  +GDC N   +RGNCGAP TLRLVKK
Sbjct  79   KGYVFSKINFADYGHSSGDCGN--FRRGNCGAPDTLRLVKK  117


Query #143: XLOC_016902 Query ID: lcl|Query_33146 Length: 806

Sequences producing significant alignments:
                                                                  Max    Total Query   E   Per.
Description                                                       Score  Score cover Value Ident  Accession
hypothetical protein AT3G02677 [Arabidopsis thaliana]             188    188   36%   1e-60 100.00 NP_001326758.1
hypothetical protein AXX17_AT3G01940 [Arabidopsis thaliana]       187    187   36%   3e-60 98.98  OAP02076.1
unnamed protein product [Arabidopsis thaliana]                    85.9   136   33%   2e-25 97.73  CAA0381121.1
unnamed protein product [Arabidopsis thaliana]                    85.9   135   33%   3e-25 97.73  VYS56084.1


Alignments:
>hypothetical protein AT3G02677 [Arabidopsis thaliana]
Sequence ID: NP_001326758.1 Length: 98
>hypothetical protein AT3G02677 [Arabidopsis thaliana]
Sequence ID: ANM64751.1 Length: 98
Range 1: 1 to 98

Score:188 bits(478), Expect:1e-60,
Method:Compositional matrix adjust.,
Identities:98/98(100%), Positives:98/98(100%), Gaps:0/98(0%)

Query  198  MQNLRPFVAKCYRGGRSIGNRQSRGFsssssPTKNEPSIWIKEVHSMEFWHLYVEKKISI  377
            MQNLRPFVAKCYRGGRSIGNRQSRGFSSSSSPTKNEPSIWIKEVHSMEFWHLYVEKKISI
Sbjct  1    MQNLRPFVAKCYRGGRSIGNRQSRGFSSSSSPTKNEPSIWIKEVHSMEFWHLYVEKKISI  60

Query  378  YIPARTKGSIHEGGTRSWSRVGRSCSRNKGKSGNFSRS  491
            YIPARTKGSIHEGGTRSWSRVGRSCSRNKGKSGNFSRS
Sbjct  61   YIPARTKGSIHEGGTRSWSRVGRSCSRNKGKSGNFSRS  98


>hypothetical protein AXX17_AT3G01940 [Arabidopsis thaliana]
Sequence ID: OAP02076.1 Length: 98
Range 1: 1 to 98

Score:187 bits(475), Expect:3e-60,
Method:Compositional matrix adjust.,
Identities:97/98(99%), Positives:98/98(100%), Gaps:0/98(0%)

Query  198  MQNLRPFVAKCYRGGRSIGNRQSRGFsssssPTKNEPSIWIKEVHSMEFWHLYVEKKISI  377
            MQNLRPFVAKCYRGGRSIGNRQSRGFSSSSSPTKNEPSIWIKEVHSMEFWHLYVEKKISI
Sbjct  1    MQNLRPFVAKCYRGGRSIGNRQSRGFSSSSSPTKNEPSIWIKEVHSMEFWHLYVEKKISI  60

Query  378  YIPARTKGSIHEGGTRSWSRVGRSCSRNKGKSGNFSRS  491
            YIPARTKGSIHEGGTRSWSRVGRSCSRN+GKSGNFSRS
Sbjct  61   YIPARTKGSIHEGGTRSWSRVGRSCSRNRGKSGNFSRS  98


>unnamed protein product [Arabidopsis thaliana]
Sequence ID: CAA0381121.1 Length: 100
Range 1: 1 to 44

Score:85.9 bits(211), Expect:2e-25,
Method:Compositional matrix adjust.,
Identities:43/44(98%), Positives:44/44(100%), Gaps:0/44(0%)

Query  198  MQNLRPFVAKCYRGGRSIGNRQSRGFsssssPTKNEPSIWIKEV  329
            MQNLRPFVAKCYRGGRSIGNRQSRGFSSSSSPTKNEPSIWIKE+
Sbjct  1    MQNLRPFVAKCYRGGRSIGNRQSRGFSSSSSPTKNEPSIWIKEL  44


Range 2: 41 to 85

Score:50.4 bits(119), Expect:2e-25,
Method:Compositional matrix adjust.,
Identities:41/45(91%), Positives:42/45(93%), Gaps:0/45(0%)

Query  379  IFQLGQKARYMRealdlgaeladlaaaTREKVAISVDLNLFKPSK  513
            I +LGQKARYMREALDLGAELADLAAAT EKVAIS DLNLFKPSK
Sbjct  41   IKELGQKARYMREALDLGAELADLAAATGEKVAISADLNLFKPSK  85


>unnamed protein product [Arabidopsis thaliana]
Sequence ID: VYS56084.1 Length: 100
Range 1: 1 to 44

Score:85.9 bits(211), Expect:3e-25,
Method:Compositional matrix adjust.,
Identities:43/44(98%), Positives:44/44(100%), Gaps:0/44(0%)

Query  198  MQNLRPFVAKCYRGGRSIGNRQSRGFsssssPTKNEPSIWIKEV  329
            MQNLRPFVAKCYRGGRSIGNRQSRGFSSSSSPTKNEPSIWIKE+
Sbjct  1    MQNLRPFVAKCYRGGRSIGNRQSRGFSSSSSPTKNEPSIWIKEL  44


Range 2: 41 to 85

Score:50.1 bits(118), Expect:3e-25,
Method:Compositional matrix adjust.,
Identities:41/45(91%), Positives:42/45(93%), Gaps:0/45(0%)

Query  379  IFQLGQKARYMRealdlgaeladlaaaTREKVAISVDLNLFKPSK  513
            I +LGQKARYMREALDLGAELADLAAAT EKVAISVDLNLFKP K
Sbjct  41   IKELGQKARYMREALDLGAELADLAAATGEKVAISVDLNLFKPKK  85


Query #144: XLOC_017194 Query ID: lcl|Query_33147 Length: 840

No significant similarity found.


Query #145: XLOC_017519 Query ID: lcl|Query_33148 Length: 967

No significant similarity found.


Query #146: XLOC_017567 Query ID: lcl|Query_33149 Length: 511

Sequences producing significant alignments:
                                                                  Max    Total Query   E   Per.
Description                                                       Score  Score cover Value Ident  Accession
unnamed protein product [Arabidopsis thaliana]                    155    155   52%   2e-46 95.56  VYS57313.1
unnamed protein product [Arabidopsis thaliana]                    152    152   52%   3e-45 94.44  CAA0382366.1


Alignments:
>unnamed protein product [Arabidopsis thaliana]
Sequence ID: VYS57313.1 Length: 314
Range 1: 1 to 90

Score:155 bits(391), Expect:2e-46,
Method:Compositional matrix adjust.,
Identities:86/90(96%), Positives:88/90(97%), Gaps:0/90(0%)

Query  148  IINFEDFPGGGEQHVPELHT*LYRRDCDEYIDQNNWDIVIAPGTRVRSRTMKRNKVVIFN  327
            +INFEDF GGGEQHVPELHT LYRRDCDEYIDQNNWDIVIAPGTRVRSRTMKRN+VVIFN
Sbjct  1    MINFEDFSGGGEQHVPELHTGLYRRDCDEYIDQNNWDIVIAPGTRVRSRTMKRNRVVIFN  60

Query  328  YTEVSLTVDGVrftrrtirrrqtWNLNFHI  417
            YTEVSLTVDGVRFTRRTIRRRQTWNLNFHI
Sbjct  61   YTEVSLTVDGVRFTRRTIRRRQTWNLNFHI  90


>unnamed protein product [Arabidopsis thaliana]
Sequence ID: CAA0382366.1 Length: 314
Range 1: 1 to 90

Score:152 bits(384), Expect:3e-45,
Method:Compositional matrix adjust.,
Identities:85/90(94%), Positives:87/90(96%), Gaps:0/90(0%)

Query  148  IINFEDFPGGGEQHVPELHT*LYRRDCDEYIDQNNWDIVIAPGTRVRSRTMKRNKVVIFN  327
            +INFEDF G GEQHVPELHT LYRRDCDEYIDQNNWDIVIAPGTRVRSRTMKRN+VVIFN
Sbjct  1    MINFEDFSGEGEQHVPELHTGLYRRDCDEYIDQNNWDIVIAPGTRVRSRTMKRNRVVIFN  60

Query  328  YTEVSLTVDGVrftrrtirrrqtWNLNFHI  417
            YTEVSLTVDGVRFTRRTIRRRQTWNLNFHI
Sbjct  61   YTEVSLTVDGVRFTRRTIRRRQTWNLNFHI  90


Query #147: XLOC_017582 Query ID: lcl|Query_33150 Length: 1307

No significant similarity found.


Query #148: XLOC_017613 Query ID: lcl|Query_33151 Length: 633

Sequences producing significant alignments:
                                                                  Max    Total Query   E   Per.
Description                                                       Score  Score cover Value Ident  Accession
putative protein [Arabidopsis thaliana]                           147    230   56%   1e-53 86.08  CAB77997.1
unnamed protein product [Arabidopsis thaliana]                    144    282   82%   3e-53 86.08  BAB01194.1
hypothetical protein AXX17_AT3G35140 [Arabidopsis thaliana]       126    126   48%   3e-37 62.04  OAP04479.1
Mutator-like transposase-like protein [Arabidopsis thaliana]      98.6   98.6  37%   8e-26 59.76  BAB02521.1
hypothetical protein [Arabidopsis thaliana]                       97.1   97.1  37%   3e-25 58.54  AAD25846.1
hypothetical protein [Arabidopsis thaliana]                       95.9   95.9  37%   9e-25 58.54  AAC23764.1
unknown [Arabidopsis thaliana]                                    95.9   95.9  37%   1e-24 58.54  ABK28509.1
hypothetical protein At3g32070 [Arabidopsis thaliana]             95.1   95.1  37%   2e-24 57.32  ABE65980.1
unknown [Arabidopsis thaliana]                                    95.1   95.1  37%   2e-24 57.32  ABK28581.1
hypothetical protein AXX17_ATUG02080 [Arabidopsis thaliana]       90.9   90.9  37%   3e-23 54.88  OAO89307.1
hypothetical protein At5g35792 [Arabidopsis thaliana]             91.7   91.7  37%   5e-23 56.10  ABE66193.1
unknown [Arabidopsis thaliana]                                    91.3   91.3  37%   5e-23 56.10  ABK28722.1
hypothetical protein AXX17_AT2G07580 [Arabidopsis thaliana]       84.3   84.3  19%   6e-21 95.12  OAP10464.1
hypothetical protein [Arabidopsis thaliana]                       79.3   79.3  30%   2e-18 58.46  AAD32920.1


Alignments:
>putative protein [Arabidopsis thaliana]
Sequence ID: CAB77997.1 Length: 848
>putative protein [Arabidopsis thaliana]
Sequence ID: CAB82108.1 Length: 848
Range 1: 721 to 799

Score:147 bits(372), Expect:1e-53,
Method:Compositional matrix adjust.,
Identities:68/79(86%), Positives:73/79(92%), Gaps:0/79(0%)

Query  68   MDRLCESDPWYDEMKSAKRIIQQLEEVAMMEGIPAICPCGGQILDKISENDGDKGKRYYK  247
            MDRLCESDPWY EMKSAKRI+QQLEEVAMMEGIP ICPCGG+ILD ISE DGDKGKR+Y+
Sbjct  721  MDRLCESDPWYGEMKSAKRIMQQLEEVAMMEGIPPICPCGGRILDIISEKDGDKGKRFYE  780

Query  248  CIVYKNDGLYIRKLWDKAM  304
            C  YKNDGL+IRKLWDKAM
Sbjct  781  CTNYKNDGLHIRKLWDKAM  799


Range 2: 808 to 847

Score:82.4 bits(202), Expect:1e-53,
Method:Compositional matrix adjust.,
Identities:36/40(90%), Positives:39/40(97%), Gaps:0/40(0%)

Query  310  EQVDNHHQKIQSLEYSNQEMRSELDEIQKKMGTLWRVCCG  429
            EQVDNHHQKIQSLEY NQ+MRS+LDEIQKKMGTLWR+CCG
Sbjct  808  EQVDNHHQKIQSLEYYNQQMRSKLDEIQKKMGTLWRICCG  847


>unnamed protein product [Arabidopsis thaliana]
Sequence ID: BAB01194.1 Length: 170
Range 1: 1 to 79

Score:144 bits(363), Expect:3e-53,
Method:Compositional matrix adjust.,
Identities:68/79(86%), Positives:73/79(92%), Gaps:0/79(0%)

Query  68   MDRLCESDPWYDEMKSAKRIIQQLEEVAMMEGIPAICPCGGQILDKISENDGDKGKRYYK  247
            MDRLCESDPWYDEMKSAKRI+QQLEEVAMME IP ICPCGG+ILD ISE DGDKGKRYY+
Sbjct  1    MDRLCESDPWYDEMKSAKRIMQQLEEVAMMEEIPIICPCGGRILDIISEKDGDKGKRYYE  60

Query  248  CIVYKNDGLYIRKLWDKAM  304
            C  YKNDGL+I+KLWDKAM
Sbjct  61   CTDYKNDGLHIQKLWDKAM  79


Range 2: 88 to 170

Score:84.7 bits(208), Expect:3e-53,
Method:Compositional matrix adjust.,
Identities:51/89(57%), Positives:60/89(67%), Gaps:12/89(13%)

Query  310  EQVDNHHQKIQSLEYSNQEMRSELDEIQKKMGTLWR-----VCCGK*GIIGFV*T*FLSL  474
            EQVDNHHQKIQSLEYSNQE+ SE DEIQKKMGTLWR     VC  +  +   +      L
Sbjct  88   EQVDNHHQKIQSLEYSNQEVLSEFDEIQKKMGTLWRVRNYWVCYYRYQVFHCI------L  141

Query  475  VCKTFIEYSLVFPVLS-SLSIPLYSQFYR  558
                + ++SLVF VLS +LS PLYS +YR
Sbjct  142  SIPLYSQFSLVFSVLSLNLSFPLYSLYYR  170


Range 3: 121 to 170

Score:53.1 bits(126), Expect:3e-08,
Method:Compositional matrix adjust.,
Identities:34/59(58%), Positives:38/59(64%), Gaps:10/59(16%)

Query  423  LWKVRNYWVCVNLISVFGL*DFY*VFPCISSI-IVIKYSFVFPVLSLNHSFPLYSLYYR  596
            LW+VRNYWVC            Y VF CI SI +  ++S VF VLSLN SFPLYSLYYR
Sbjct  121  LWRVRNYWVCYYR---------YQVFHCILSIPLYSQFSLVFSVLSLNLSFPLYSLYYR  170


>hypothetical protein AXX17_AT3G35140 [Arabidopsis thaliana]
Sequence ID: OAP04479.1 Length: 100
Range 1: 1 to 100

Score:126 bits(317), Expect:3e-37,
Method:Compositional matrix adjust.,
Identities:67/108(62%), Positives:78/108(72%), Gaps:14/108(12%)

Query  68   MDRLCESDPWYDEMKSAKRIIQQLEEVAMMEGIPAICPCGGQILDKISENDGDKGKRYYK  247
            MDRLCE+DPWYDEMK  K I++QLEEVAMM+GIPAICPCGG+IL        DK KRYY+
Sbjct  1    MDRLCENDPWYDEMKKVKWIMEQLEEVAMMKGIPAICPCGGRIL--------DKRKRYYE  52

Query  248  CIVYKNDGLYIRKLWDKAMG------*SKLITITKRFKAWSTLTKKCV  373
            C VYKNDGL+IRKLWDKAM         +     K+ ++  TLTKKCV
Sbjct  53   CTVYKNDGLHIRKLWDKAMVEEVNRLREQGYNHHKKIQSLETLTKKCV  100


>Mutator-like transposase-like protein [Arabidopsis thaliana]
Sequence ID: BAB02521.1 Length: 132
Range 1: 1 to 77

Score:98.6 bits(244), Expect:8e-26,
Method:Compositional matrix adjust.,
Identities:49/82(60%), Positives:60/82(73%), Gaps:8/82(9%)

Query  68   MDRLCESDPWYDEMKSAKRIIQQLEEVAMMEGIPAICPCGGQILDKISENDGDKGKRYYK  247
            MDRLCE DP+YD+MK AKR I+Q+E VAMMEGIP  CPCGG I+D   +      KRYY+
Sbjct  1    MDRLCERDPYYDDMKVAKRAIEQMEMVAMMEGIPKFCPCGGSIVDTRKDE-----KRYYQ  55

Query  248  CIVYKNDG---LYIRKLWDKAM  304
            C  +K+D    ++IRKLWDKAM
Sbjct  56   CEKFKDDRTDCMHIRKLWDKAM  77


>hypothetical protein [Arabidopsis thaliana]
Sequence ID: AAD25846.1 Length: 118
>hypothetical protein [Arabidopsis thaliana]
Sequence ID: AAM15462.1 Length: 118
Range 1: 1 to 77

Score:97.1 bits(240), Expect:3e-25,
Method:Compositional matrix adjust.,
Identities:48/82(59%), Positives:60/82(73%), Gaps:8/82(9%)

Query  68   MDRLCESDPWYDEMKSAKRIIQQLEEVAMMEGIPAICPCGGQILDKISENDGDKGKRYYK  247
            MDRLCE DP+YD+MK AKR I+Q+E VAMMEGIP  CPCGG I++   +      KRYY+
Sbjct  1    MDRLCERDPYYDDMKVAKRAIEQMEMVAMMEGIPKFCPCGGSIVETRKDE-----KRYYQ  55

Query  248  CIVYKNDG---LYIRKLWDKAM  304
            C  +K+D    ++IRKLWDKAM
Sbjct  56   CEKFKDDRTDCMHIRKLWDKAM  77


>hypothetical protein [Arabidopsis thaliana]
Sequence ID: AAC23764.1 Length: 132
>hypothetical protein At2g23490 [Arabidopsis thaliana]
Sequence ID: ABE65852.1 Length: 132
Range 1: 1 to 77

Score:95.9 bits(237), Expect:9e-25,
Method:Compositional matrix adjust.,
Identities:48/82(59%), Positives:59/82(71%), Gaps:8/82(9%)

Query  68   MDRLCESDPWYDEMKSAKRIIQQLEEVAMMEGIPAICPCGGQILDKISENDGDKGKRYYK  247
            MDRLCE DP+YD+MK AKR I Q+E VAMMEGIP  CPCGG I++   +      KRYY+
Sbjct  1    MDRLCERDPYYDDMKVAKRAIDQMEMVAMMEGIPKFCPCGGSIVETRKDE-----KRYYQ  55

Query  248  CIVYKNDG---LYIRKLWDKAM  304
            C  +K+D    ++IRKLWDKAM
Sbjct  56   CEKFKDDRTDCMHIRKLWDKAM  77


>unknown, partial [Arabidopsis thaliana]
Sequence ID: ABK28509.1 Length: 133
Range 1: 1 to 77

Score:95.9 bits(237), Expect:1e-24,
Method:Compositional matrix adjust.,
Identities:48/82(59%), Positives:59/82(71%), Gaps:8/82(9%)

Query  68   MDRLCESDPWYDEMKSAKRIIQQLEEVAMMEGIPAICPCGGQILDKISENDGDKGKRYYK  247
            MDRLCE DP+YD+MK AKR I Q+E VAMMEGIP  CPCGG I++   +      KRYY+
Sbjct  1    MDRLCERDPYYDDMKVAKRAIDQMEMVAMMEGIPKFCPCGGSIVETRKDE-----KRYYQ  55

Query  248  CIVYKNDG---LYIRKLWDKAM  304
            C  +K+D    ++IRKLWDKAM
Sbjct  56   CEKFKDDRTDCMHIRKLWDKAM  77


>hypothetical protein At3g32070 [Arabidopsis thaliana]
Sequence ID: ABE65980.1 Length: 132
>Mutator-like transposase-like protein [Arabidopsis thaliana]
Sequence ID: BAB01938.1 Length: 132
Range 1: 1 to 77

Score:95.1 bits(235), Expect:2e-24,
Method:Compositional matrix adjust.,
Identities:47/82(57%), Positives:59/82(71%), Gaps:8/82(9%)

Query  68   MDRLCESDPWYDEMKSAKRIIQQLEEVAMMEGIPAICPCGGQILDKISENDGDKGKRYYK  247
            MDR CE DP+YD++K AKR I+Q+E VAMMEGIP  CPCGG I+D   +      KRYY+
Sbjct  1    MDRECERDPYYDDLKVAKRAIEQMEMVAMMEGIPKFCPCGGSIVDTRKDE-----KRYYQ  55

Query  248  CIVYKNDG---LYIRKLWDKAM  304
            C  +K+D    ++IRKLWDKAM
Sbjct  56   CEKFKDDRTDLMHIRKLWDKAM  77


>unknown, partial [Arabidopsis thaliana]
Sequence ID: ABK28581.1 Length: 133
Range 1: 1 to 77

Score:95.1 bits(235), Expect:2e-24,
Method:Compositional matrix adjust.,
Identities:47/82(57%), Positives:59/82(71%), Gaps:8/82(9%)

Query  68   MDRLCESDPWYDEMKSAKRIIQQLEEVAMMEGIPAICPCGGQILDKISENDGDKGKRYYK  247
            MDR CE DP+YD++K AKR I+Q+E VAMMEGIP  CPCGG I+D   +      KRYY+
Sbjct  1    MDRECERDPYYDDLKVAKRAIEQMEMVAMMEGIPKFCPCGGSIVDTRKDE-----KRYYQ  55

Query  248  CIVYKNDG---LYIRKLWDKAM  304
            C  +K+D    ++IRKLWDKAM
Sbjct  56   CEKFKDDRTDLMHIRKLWDKAM  77


>hypothetical protein AXX17_ATUG02080 [Arabidopsis thaliana]
Sequence ID: OAO89307.1 Length: 105
Range 1: 1 to 77

Score:90.9 bits(224), Expect:3e-23,
Method:Compositional matrix adjust.,
Identities:45/82(55%), Positives:58/82(70%), Gaps:8/82(9%)

Query  68   MDRLCESDPWYDEMKSAKRIIQQLEEVAMMEGIPAICPCGGQILDKISENDGDKGKRYYK  247
            MDR  E DP+YD++K+AKR I+Q+E +AMMEGIP  CPCGG I+D   +      KRYY+
Sbjct  1    MDRESEKDPYYDDLKAAKRAIEQMEMIAMMEGIPKFCPCGGSIVDTRKDE-----KRYYQ  55

Query  248  CIVYK---NDGLYIRKLWDKAM  304
            C  +K   ND ++I KLWDKAM
Sbjct  56   CEKFKDDRNDLMHISKLWDKAM  77


>hypothetical protein At5g35792 [Arabidopsis thaliana]
Sequence ID: ABE66193.1 Length: 132
>mutator-like transposase [Arabidopsis thaliana]
Sequence ID: BAB09919.1 Length: 132
Range 1: 1 to 77

Score:91.7 bits(226), Expect:5e-23,
Method:Compositional matrix adjust.,
Identities:46/82(56%), Positives:59/82(71%), Gaps:8/82(9%)

Query  68   MDRLCESDPWYDEMKSAKRIIQQLEEVAMMEGIPAICPCGGQILDKISENDGDKGKRYYK  247
            MDRLCE DP+Y ++K AKR I+Q+E VAMMEGIP  CPCGG I+D   +      KRYY+
Sbjct  1    MDRLCERDPYYVDIKVAKRAIEQMEMVAMMEGIPKFCPCGGSIVDTRKDE-----KRYYQ  55

Query  248  CIVYKN---DGLYIRKLWDKAM  304
            C  +K+   D ++IRKLWDKA+
Sbjct  56   CEKFKDNRTDCMHIRKLWDKAI  77


>unknown, partial [Arabidopsis thaliana]
Sequence ID: ABK28722.1 Length: 133
Range 1: 1 to 77

Score:91.3 bits(225), Expect:5e-23,
Method:Compositional matrix adjust.,
Identities:46/82(56%), Positives:59/82(71%), Gaps:8/82(9%)

Query  68   MDRLCESDPWYDEMKSAKRIIQQLEEVAMMEGIPAICPCGGQILDKISENDGDKGKRYYK  247
            MDRLCE DP+Y ++K AKR I+Q+E VAMMEGIP  CPCGG I+D   +      KRYY+
Sbjct  1    MDRLCERDPYYVDIKVAKRAIEQMEMVAMMEGIPKFCPCGGSIVDTRKDE-----KRYYQ  55

Query  248  CIVYKN---DGLYIRKLWDKAM  304
            C  +K+   D ++IRKLWDKA+
Sbjct  56   CEKFKDNRTDCMHIRKLWDKAI  77


>hypothetical protein AXX17_AT2G07580 [Arabidopsis thaliana]
Sequence ID: OAP10464.1 Length: 81
Range 1: 24 to 64

Score:84.3 bits(207), Expect:6e-21,
Method:Compositional matrix adjust.,
Identities:39/41(95%), Positives:41/41(100%), Gaps:0/41(0%)

Query  310  EQVDNHHQKIQSLEYSNQEMRSELDEIQKKMGTLWRVCCGK  432
            EQVDNHHQKIQSLEYS+QEMRSELDEIQKKMGTLWRVCCG+
Sbjct  24   EQVDNHHQKIQSLEYSSQEMRSELDEIQKKMGTLWRVCCGE  64


>hypothetical protein [Arabidopsis thaliana]
Sequence ID: AAD32920.1 Length: 117
Range 1: 1 to 60

Score:79.3 bits(194), Expect:2e-18,
Method:Compositional matrix adjust.,
Identities:38/65(58%), Positives:45/65(69%), Gaps:5/65(7%)

Query  68   MDRLCESDPWYDEMKSAKRIIQQLEEVAMMEGIPAICPCGGQILDKISENDGDKGKRYYK  247
            MDRLCE DP+YD+MK  KR I Q+E VAMMEGIP  CPCGG I+D   +      KRYY+
Sbjct  1    MDRLCERDPYYDDMKVVKRAIAQMEMVAMMEGIPKFCPCGGSIVDTRKDE-----KRYYQ  55

Query  248  CIVYK  262
            C  +K
Sbjct  56   CEKFK  60


Query #149: XLOC_018065 Query ID: lcl|Query_33152 Length: 402

No significant similarity found.


Query #150: XLOC_018302 Query ID: lcl|Query_33153 Length: 802

No significant similarity found.


Query #151: XLOC_018488 Query ID: lcl|Query_33154 Length: 329

No significant similarity found.


Query #152: XLOC_018608 Query ID: lcl|Query_33155 Length: 563

No significant similarity found.


Query #153: XLOC_018621 Query ID: lcl|Query_33156 Length: 1514

Sequences producing significant alignments:
                                                                  Max    Total Query   E   Per.
Description                                                       Score  Score cover Value Ident  Accession
unnamed protein product [Arabidopsis thaliana]                    254    254   24%   5e-83 98.41  CAA0384222.1
unnamed protein product [Arabidopsis thaliana]                    106    106   13%   2e-27 100.00 VYS59110.1
hypothetical protein AT1G43720 [Arabidopsis thaliana]             98.2   98.2  26%   5e-22 39.26  NP_175038.1
unnamed protein product [Arabidopsis thaliana]                    98.2   98.2  26%   6e-22 39.26  CAA0272678.1
unnamed protein product [Arabidopsis thaliana]                    91.3   91.3  22%   6e-20 39.29  VYS59162.1
unnamed protein product [Arabidopsis thaliana]                    94.4   94.4  21%   1e-19 38.32  VYS59124.1
heat shock protein [Arabidopsis thaliana]                         90.1   90.1  24%   3e-19 38.10  NP_189873.1
unnamed protein product [Arabidopsis thaliana]                    88.2   88.2  25%   4e-19 36.64  CAA0384316.1
unnamed protein product [Arabidopsis thaliana]                    89.0   89.0  21%   9e-19 34.91  CAA0384260.1
Myb/SANT-like DNA-binding domain protein [Arabidopsis thaliana]   89.0   89.0  21%   1e-18 36.45  NP_683626.1
hypothetical protein AXX17_AT3G35980 [Arabidopsis thaliana]       85.1   85.1  25%   8e-18 36.64  OAP03107.1
unnamed protein product [Arabidopsis thaliana]                    85.9   85.9  20%   9e-18 37.62  VYS59084.1
unnamed protein product [Arabidopsis thaliana]                    84.7   84.7  20%   3e-17 37.62  BAB01018.1
unnamed protein product [Arabidopsis thaliana]                    77.0   77.0  17%   2e-14 42.39  CAA0268354.1
unnamed protein product [Arabidopsis thaliana]                    74.3   74.3  14%   6e-14 41.10  CAA0384198.1
heat shock protein [Arabidopsis thaliana]                         72.8   72.8  14%   4e-13 41.10  NP_189673.2


Alignments:
>unnamed protein product [Arabidopsis thaliana]
Sequence ID: CAA0384222.1 Length: 126
Range 1: 1 to 126

Score:254 bits(649), Expect:5e-83,
Method:Compositional matrix adjust.,
Identities:124/126(98%), Positives:124/126(98%), Gaps:0/126(0%)

Query  527  MIHNHHMNSHDRQHSRVDPSASRKNGSRESSGSWVPRRRKFFESTIQETMSSIRDFQRES  706
            MIHNHHMNSHDRQHSRVDPS SRKNGSRESSGSWVPRRRKFFESTIQETMSSIRDFQRES
Sbjct  1    MIHNHHMNSHDRQHSRVDPSGSRKNGSRESSGSWVPRRRKFFESTIQETMSSIRDFQRES  60

Query  707  FERLRPGAFDREDFTEFERAVDILESLEIRRYNDFYVQCLQLLKTDLFWRNYWMS*SRFQ  886
            FERLRPGAFDREDFTEFERAVDILESLEIRRYNDFYVQCLQLLKTDLFWRNYWMS SRFQ
Sbjct  61   FERLRPGAFDREDFTEFERAVDILESLEIRRYNDFYVQCLQLLKTDLFWRNYWMSSSRFQ  120

Query  887  TNEDRI  904
            TNEDRI
Sbjct  121  TNEDRI  126


>unnamed protein product [Arabidopsis thaliana]
Sequence ID: VYS59110.1 Length: 67
Range 1: 1 to 67

Score:106 bits(264), Expect:2e-27,
Method:Compositional matrix adjust.,
Identities:67/67(100%), Positives:67/67(100%), Gaps:0/67(0%)

Query  1164  MLGDPTCDLRDARCSTFIGARKCRKPVVDGSIRILGVLWLVFADILQAddgpddgsddrG  985
             MLGDPTCDLRDARCSTFIGARKCRKPVVDGSIRILGVLWLVFADILQADDGPDDGSDDRG
Sbjct  1     MLGDPTCDLRDARCSTFIGARKCRKPVVDGSIRILGVLWLVFADILQADDGPDDGSDDRG  60

Query  984   VQDPLGG  964
             VQDPLGG
Sbjct  61    VQDPLGG  67


>hypothetical protein AT1G43720 [Arabidopsis thaliana]
Sequence ID: NP_175038.1 Length: 314
>hypothetical protein AT1G43720 [Arabidopsis thaliana]
Sequence ID: AEE31990.1 Length: 314
Range 1: 68 to 201

Score:98.2 bits(243), Expect:5e-22,
Method:Compositional matrix adjust.,
Identities:53/135(39%), Positives:86/135(63%), Gaps:4/135(2%)

Query  557  DRQHSRVDPSASRKNGSRE---SSGSWVPRRRKFFESTIQETMSSIRDFQRESFERLRPG  727
            D   +R   SA R  GS      SGS    RR+ FE+TIQ++++  R+FQR+SF++LRPG
Sbjct  68   DESTARRGSSAQRSGGSSLVSIGSGSRRSHRRQSFETTIQDSITGFREFQRQSFQQLRPG  127

Query  728  AFDREDFTEFERAVDILESLEIRRYNDFYVQCLQLLKTDLFWRNYWMS*SRFQTNEDRI*  907
            AFD++D+ EF++A  I  +L + ++  F+  C+  LK  +FWR Y +  +   T+ED++
Sbjct  128  AFDQDDYDEFKKAEAIFIALNLPKHTRFHWACINALKELVFWRKYLIDITA-STDEDKVQ  186

Query  908  LLESLTGFTSQGENV  952
            LLE++TG +   ++V
Sbjct  187  LLEAMTGVSRNNQDV  201


>unnamed protein product [Arabidopsis thaliana]
Sequence ID: CAA0272678.1 Length: 315
Range 1: 69 to 202

Score:98.2 bits(243), Expect:6e-22,
Method:Compositional matrix adjust.,
Identities:53/135(39%), Positives:86/135(63%), Gaps:4/135(2%)

Query  557  DRQHSRVDPSASRKNGSRE---SSGSWVPRRRKFFESTIQETMSSIRDFQRESFERLRPG  727
            D   +R   SA R  GS      SGS    RR+ FE+TIQ++++  R+FQR+SF++LRPG
Sbjct  69   DESTARRGSSAQRSGGSSLVSIGSGSRRSHRRQSFETTIQDSITGFREFQRQSFQQLRPG  128

Query  728  AFDREDFTEFERAVDILESLEIRRYNDFYVQCLQLLKTDLFWRNYWMS*SRFQTNEDRI*  907
            AFD++D+ EF++A  I  +L + ++  F+  C+  LK  +FWR Y +  +   T+ED++
Sbjct  129  AFDQDDYDEFKKAEAIFIALNLPKHTRFHWACINALKELVFWRKYLIDITT-STDEDKVQ  187

Query  908  LLESLTGFTSQGENV  952
            LLE++TG +   ++V
Sbjct  188  LLEAMTGVSRNNQDV  202


>unnamed protein product [Arabidopsis thaliana]
Sequence ID: VYS59162.1 Length: 269
Range 1: 130 to 240

Score:91.3 bits(225), Expect:6e-20,
Method:Compositional matrix adjust.,
Identities:44/112(39%), Positives:73/112(65%), Gaps:1/112(0%)

Query  617  SGSWVPRRRKFFESTIQETMSSIRDFQRESFERLRPGAFDREDFTEFERAVDILESLEIR  796
            SGS   RR+++FE+T+ +TM+  R+FQ +S ++L P +FD+ DF E E  V I E++E+
Sbjct  130  SGSRGSRRKQYFETTLTDTMTGFREFQCQSLQQLHPNSFDQNDFEECETVVKIFEAMELP  189

Query  797  RYNDFYVQCLQLLKTDLFWRNYWMS*SRFQTNEDRI*LLESLTGFTSQGENV  952
                FY +C++  K D FWR Y+++ +  ++ ED+I  L+ LTG+T   + V
Sbjct  190  NDTKFYWECIRAFKEDEFWRKYFIARAD-KSFEDKIQFLQGLTGYTRDDKRV  240


>unnamed protein product [Arabidopsis thaliana]
Sequence ID: VYS59124.1 Length: 593
Range 1: 169 to 274

Score:94.4 bits(233), Expect:1e-19,
Method:Compositional matrix adjust.,
Identities:41/107(38%), Positives:73/107(68%), Gaps:1/107(0%)

Query  635  RRRKFFESTIQETMSSIRDFQRESFERLRPGAFDREDFTEFERAVDILESLEIRRYNDFY  814
            RR++ FE+T+ +T++ +R+FQR+S +++ P  FD +D+ EF+ A+ I ES+++    DFY
Sbjct  169  RRKQSFETTLTDTITGLREFQRQSLQQMCPNCFDEDDYNEFDMAMKIFESMDLPNDTDFY  228

Query  815  VQCLQLLKTDLFWRNYWMS*SRFQTNEDRI*LLESLTGFTSQGENVG  955
              C+   K ++FWR Y+++ S  +  ED++  L++LTG+T   E VG
Sbjct  229  WACIHAFKEEIFWRKYFINTSE-KPVEDKLKFLQALTGYTRDNEYVG  274


>heat shock protein [Arabidopsis thaliana]
Sequence ID: NP_189873.1 Length: 312
>heat shock protein [Arabidopsis thaliana]
Sequence ID: AEE77757.1 Length: 312 >hypothetical protein [Arabidopsis thaliana]
Sequence ID: CAB86674.1 Length: 312
Range 1: 82 to 203

Score:90.1 bits(222), Expect:3e-19,
Method:Compositional matrix adjust.,
Identities:48/126(38%), Positives:81/126(64%), Gaps:7/126(5%)

Query  584  SASRKNGSRESS---GSWVPRRRKFFESTIQETMSSIRDFQRESFERLRPGAFDREDFTE  754
            SA R  GS   S   GS   R+R+ FE+TI++ ++S R+FQR+SF    PGAFD++D+ E
Sbjct  82   SAQRSGGSSRVSIRSGSRGSRKRQSFETTIEDNITSFREFQRQSF---HPGAFDQDDYDE  138

Query  755  FERAVDILESLEIRRYNDFYVQCLQLLKTDLFWRNYWMS*SRFQTNEDRI*LLESLTGFT  934
            F++A  I   L++ ++  FY  C+  L+  ++WR Y++  +   T+ED++ LLE++TG +
Sbjct  139  FKKAEAIFIPLDLPKHTRFYWACINALEELVYWRKYFIDIAA-STDEDKVQLLEAMTGVS  197

Query  935  SQGENV  952
               ++V
Sbjct  198  PNNQDV  203


>unnamed protein product [Arabidopsis thaliana]
Sequence ID: CAA0384316.1 Length: 228
Range 1: 45 to 171

Score:88.2 bits(217), Expect:4e-19,
Method:Compositional matrix adjust.,
Identities:48/131(37%), Positives:82/131(62%), Gaps:7/131(5%)

Query  569  SRVDPSASRKNGSRESS---GSWVPRRRKFFESTIQETMSSIRDFQRESFERLRPGAFDR  739
            +R   SA R  GS   S   GS   R+R+ FE+TI++ ++  R+FQR+SF    PGAFD+
Sbjct  45   ARRKSSAQRSGGSSRVSIRSGSRGSRKRQSFETTIEDNITGFREFQRQSFH---PGAFDQ  101

Query  740  EDFTEFERAVDILESLEIRRYNDFYVQCLQLLKTDLFWRNYWMS*SRFQTNEDRI*LLES  919
            +D+ EF++A  I   L++ ++  FY  C+  L+  ++WR Y++  +   T+ED++ LLE+
Sbjct  102  DDYDEFKKAEAIFIPLDLPKHTIFYWACINALEELVYWRKYFIDIAA-STDEDKVQLLEA  160

Query  920  LTGFTSQGENV  952
            +TG +   ++V
Sbjct  161  MTGVSPNNQDV  171


>unnamed protein product [Arabidopsis thaliana]
Sequence ID: CAA0384260.1 Length: 309
Range 1: 176 to 280

Score:89.0 bits(219), Expect:9e-19,
Method:Compositional matrix adjust.,
Identities:37/106(35%), Positives:71/106(66%), Gaps:1/106(0%)

Query  635  RRRKFFESTIQETMSSIRDFQRESFERLRPGAFDREDFTEFERAVDILESLEIRRYNDFY  814
            RR++ FE+T+ + ++ +R+FQR+S +++RP  FD++D+ EF+ A+ + ES+ +    DF
Sbjct  176  RRKQLFETTLTDIITGLREFQRQSLQQMRPNCFDKDDYNEFDMAMKLFESMNLPNDTDFN  235

Query  815  VQCLQLLKTDLFWRNYWMS*SRFQTNEDRI*LLESLTGFTSQGENV  952
              C+   K ++FWR Y+++ +  +  ED++  L++LTG+T   E V
Sbjct  236  WACIHAFKAEMFWRKYFINRAE-RIVEDKLKFLQALTGYTRDNEYV  280


>Myb/SANT-like DNA-binding domain protein [Arabidopsis thaliana]
Sequence ID: NP_683626.1 Length: 330
>Myb/SANT-like DNA-binding domain protein [Arabidopsis thaliana]
Sequence ID: AEE77701.1 Length: 330
Range 1: 161 to 266

Score:89.0 bits(219), Expect:1e-18,
Method:Compositional matrix adjust.,
Identities:39/107(36%), Positives:72/107(67%), Gaps:1/107(0%)

Query  635  RRRKFFESTIQETMSSIRDFQRESFERLRPGAFDREDFTEFERAVDILESLEIRRYNDFY  814
            RR++ FE+T+ +T++ +R+FQR+S +++RP  FD++D+ +F+ A+ + ES+ +    DFY
Sbjct  161  RRKQSFETTLTDTITGLREFQRQSLQQMRPNCFDKDDYNKFDMAMKLFESMNLPNDTDFY  220

Query  815  VQCLQLLKTDLFWRNYWMS*SRFQTNEDRI*LLESLTGFTSQGENVG  955
              C+   K ++FWR Y+++ +  +T ED +  L++LT  T   E VG
Sbjct  221  WACIHAFKAEIFWRKYFINRAE-RTVEDNLKFLQALTRDTRDNEYVG  266


>hypothetical protein AXX17_AT3G35980 [Arabidopsis thaliana]
Sequence ID: OAP03107.1 Length: 264
Range 1: 45 to 171

Score:85.1 bits(209), Expect:8e-18,
Method:Compositional matrix adjust.,
Identities:48/131(37%), Positives:82/131(62%), Gaps:7/131(5%)

Query  569  SRVDPSASRKNGSRESS---GSWVPRRRKFFESTIQETMSSIRDFQRESFERLRPGAFDR  739
            +R + SA R  GS   S   GS   R+R+ FE+TI++ ++  R+FQR+SF    PGAFD+
Sbjct  45   ARRESSAQRCGGSSRVSIRSGSRGSRKRQSFETTIEDNITGFREFQRQSF---HPGAFDQ  101

Query  740  EDFTEFERAVDILESLEIRRYNDFYVQCLQLLKTDLFWRNYWMS*SRFQTNEDRI*LLES  919
            +D+ EF++A  I   L++ ++  FY  C+  L+  ++WR Y++  +   T+ED++ LLE+
Sbjct  102  DDYDEFKKAEAIFIPLDLSKHTRFYWACINALEELVYWRKYFIDIAA-STDEDKVQLLEA  160

Query  920  LTGFTSQGENV  952
            +TG     ++V
Sbjct  161  MTGVYPNNQDV  171


>unnamed protein product [Arabidopsis thaliana]
Sequence ID: VYS59084.1 Length: 313
Range 1: 212 to 311

Score:85.9 bits(211), Expect:9e-18,
Method:Compositional matrix adjust.,
Identities:38/101(38%), Positives:72/101(71%), Gaps:1/101(0%)

Query  650  FESTIQETMSSIRDFQRESFERLRPGAFDREDFTEFERAVDILESLEIRRYNDFYVQCLQ  829
            FE+TIQ++++S  +FQR+SF++LRPGAFD++D+ EF++A  I  +L++ ++  F+  C+
Sbjct  212  FETTIQDSITSFGEFQRQSFQQLRPGAFDQDDYDEFKKAEAIFIALDLPKHTRFHWACIN  271

Query  830  LLKTDLFWRNYWMS*SRFQTNEDRI*LLESLTGFTSQGENV  952
             LK  +FW  Y++  +    ++D++ LLE++TG +   ++V
Sbjct  272  ALKELVFWPKYFIDIAA-SPDKDKVQLLEAMTGVSRNNQDV  311


>unnamed protein product [Arabidopsis thaliana]
Sequence ID: BAB01018.1 Length: 313
Range 1: 212 to 311

Score:84.7 bits(208), Expect:3e-17,
Method:Compositional matrix adjust.,
Identities:38/101(38%), Positives:71/101(70%), Gaps:1/101(0%)

Query  650  FESTIQETMSSIRDFQRESFERLRPGAFDREDFTEFERAVDILESLEIRRYNDFYVQCLQ  829
            FE+TIQ++++S  +FQR+SF++LRP AFD++D+ EF++A  I  +L++ ++  F+  C+
Sbjct  212  FETTIQDSITSFGEFQRQSFQQLRPRAFDQDDYDEFKKAEAIFTALDLPKHTRFHWACIN  271

Query  830  LLKTDLFWRNYWMS*SRFQTNEDRI*LLESLTGFTSQGENV  952
             LK  +FW  Y++  +    ++D++ LLE++TG +   +NV
Sbjct  272  ALKELVFWPKYFIDIAA-SPDKDKVQLLEAMTGVSRNNQNV  311


>unnamed protein product [Arabidopsis thaliana]
Sequence ID: CAA0268354.1 Length: 450
Range 1: 217 to 308

Score:77.0 bits(188), Expect:2e-14,
Method:Compositional matrix adjust.,
Identities:39/92(42%), Positives:61/92(66%), Gaps:3/92(3%)

Query  584  SASRKNGSRE---SSGSWVPRRRKFFESTIQETMSSIRDFQRESFERLRPGAFDREDFTE  754
            SA R  GS      SGS   +RR+ FE+TIQ+ ++  R+FQR+SF++LRPGAFD++D+ E
Sbjct  217  SAQRSGGSSRVSIGSGSRGSQRRQSFETTIQDIITGFREFQRQSFQQLRPGAFDQDDYDE  276

Query  755  FERAVDILESLEIRRYNDFYVQCLQLLKTDLF  850
            F++A  I  +L + ++  F+  C+  LK  +F
Sbjct  277  FKKAEAIFIALNLPKHTRFHWACINALKELVF  308


>unnamed protein product [Arabidopsis thaliana]
Sequence ID: CAA0384198.1 Length: 292
Range 1: 175 to 247

Score:74.3 bits(181), Expect:6e-14,
Method:Compositional matrix adjust.,
Identities:30/73(41%), Positives:55/73(75%), Gaps:0/73(0%)

Query  650  FESTIQETMSSIRDFQRESFERLRPGAFDREDFTEFERAVDILESLEIRRYNDFYVQCLQ  829
            FE+TIQ++++S  +FQ++SF++LRPGAFD++D+ EF++A  I  +L++ ++  F+  C+
Sbjct  175  FETTIQDSITSFGEFQQQSFQQLRPGAFDQDDYDEFKKAEAIFIALDLPKHTRFHWACIN  234

Query  830  LLKTDLFWRNYWM  868
             LK  +FW  Y++
Sbjct  235  ALKELVFWPKYFI  247


>heat shock protein [Arabidopsis thaliana]
Sequence ID: NP_189673.2 Length: 397
>heat shock protein [Arabidopsis thaliana]
Sequence ID: AEE77648.1 Length: 397
Range 1: 212 to 284

Score:72.8 bits(177), Expect:4e-13,
Method:Compositional matrix adjust.,
Identities:30/73(41%), Positives:54/73(73%), Gaps:0/73(0%)

Query  650  FESTIQETMSSIRDFQRESFERLRPGAFDREDFTEFERAVDILESLEIRRYNDFYVQCLQ  829
            FE+TIQ++++S  +FQR+SF++LRP AFD++D+ EF++A  I  +L++ ++  F+  C+
Sbjct  212  FETTIQDSITSFGEFQRQSFQQLRPRAFDQDDYDEFKKAEAIFTALDLPKHTRFHWACIN  271

Query  830  LLKTDLFWRNYWM  868
             LK  +FW  Y++
Sbjct  272  ALKELVFWPKYFI  284


Query #154: XLOC_018654 Query ID: lcl|Query_33157 Length: 441

Sequences producing significant alignments:
                                                                  Max    Total Query   E   Per.
Description                                                       Score  Score cover Value Ident  Accession
Reticulon family protein [Arabidopsis thaliana]                   64.7   64.7  31%   3e-12 74.47  NP_200608.3
putative protein [Arabidopsis thaliana]                           64.7   64.7  31%   3e-12 74.47  BAD93974.1
unnamed protein product [Arabidopsis thaliana]                    64.7   64.7  31%   3e-12 74.47  VYS70713.1
Reticulon family protein [Arabidopsis thaliana]                   64.3   64.3  31%   3e-12 74.47  NP_001331115.1


Alignments:
>Reticulon family protein [Arabidopsis thaliana]
Sequence ID: NP_200608.3 Length: 487
>RecName: Full=Reticulon-like protein B21; Short=AtRTNLB21 [Arabidopsis thaliana]
Sequence ID: Q56X72.2 Length: 487 >Reticulon family protein [Arabidopsis thaliana]
Sequence ID: AED96983.1 Length: 487
Range 1: 211 to 257

Score:64.7 bits(156), Expect:3e-12,
Method:Compositional matrix adjust.,
Identities:35/47(74%), Positives:38/47(80%), Gaps:1/47(2%)

Query  303  DRSAE**DNCTQPQ-KLQKLVDLVLWRDVSRSTLVYGFRSFLIISSS  440
            DR AE  DN T  Q KLQ LVDLV+WRDVSRSTLV+GF +FLIISSS
Sbjct  211  DRFAEQEDNFTHSQSKLQSLVDLVMWRDVSRSTLVFGFGTFLIISSS  257


>putative protein [Arabidopsis thaliana]
Sequence ID: BAD93974.1 Length: 487
Range 1: 211 to 257

Score:64.7 bits(156), Expect:3e-12,
Method:Compositional matrix adjust.,
Identities:35/47(74%), Positives:38/47(80%), Gaps:1/47(2%)

Query  303  DRSAE**DNCTQPQ-KLQKLVDLVLWRDVSRSTLVYGFRSFLIISSS  440
            DR AE  DN T  Q KLQ LVDLV+WRDVSRSTLV+GF +FLIISSS
Sbjct  211  DRFAEQEDNFTHSQSKLQSLVDLVMWRDVSRSTLVFGFGTFLIISSS  257


>unnamed protein product [Arabidopsis thaliana]
Sequence ID: VYS70713.1 Length: 593
Range 1: 317 to 363

Score:64.7 bits(156), Expect:3e-12,
Method:Compositional matrix adjust.,
Identities:35/47(74%), Positives:38/47(80%), Gaps:1/47(2%)

Query  303  DRSAE**DNCTQPQ-KLQKLVDLVLWRDVSRSTLVYGFRSFLIISSS  440
            DR AE  DN T  Q KLQ LVDLV+WRDVSRSTLV+GF +FLIISSS
Sbjct  317  DRFAEQEDNFTHSQSKLQSLVDLVMWRDVSRSTLVFGFGTFLIISSS  363


>Reticulon family protein [Arabidopsis thaliana]
Sequence ID: NP_001331115.1 Length: 593
>Reticulon family protein [Arabidopsis thaliana]
Sequence ID: ANM69437.1 Length: 593 >hypothetical protein AXX17_AT5G57310 [Arabidopsis thaliana]
Sequence ID: OAO94395.1 Length: 593 >unnamed protein product [Arabidopsis thaliana]
Sequence ID: CAA0410561.1 Length: 593
Range 1: 317 to 363

Score:64.3 bits(155), Expect:3e-12,
Method:Compositional matrix adjust.,
Identities:35/47(74%), Positives:38/47(80%), Gaps:1/47(2%)

Query  303  DRSAE**DNCTQPQ-KLQKLVDLVLWRDVSRSTLVYGFRSFLIISSS  440
            DR AE  DN T  Q KLQ LVDLV+WRDVSRSTLV+GF +FLIISSS
Sbjct  317  DRFAEQEDNFTHSQSKLQSLVDLVMWRDVSRSTLVFGFGTFLIISSS  363


Query #155: XLOC_018914 Query ID: lcl|Query_33158 Length: 312

No significant similarity found.


Query #156: XLOC_018952 Query ID: lcl|Query_33159 Length: 555

Sequences producing significant alignments:
                                                                  Max    Total Query   E   Per.
Description                                                       Score  Score cover Value Ident  Accession
transmembrane protein [Arabidopsis thaliana]                      55.5   93.6  28%   4e-13 81.25  NP_001326023.1
hypothetical protein AT2G14265 [Arabidopsis thaliana]             61.6   61.6  25%   5e-12 68.75  NP_001325174.1
unnamed protein product [Arabidopsis thaliana]                    55.8   55.8  17%   3e-10 81.25  CAA0384287.1
unnamed protein product [Arabidopsis thaliana]                    46.6   82.8  23%   5e-10 78.57  VYS52357.1
unnamed protein product [Arabidopsis thaliana]                    42.0   81.6  26%   1e-09 73.91  VYS52323.1
unnamed protein product [Arabidopsis thaliana]                    41.6   81.2  26%   2e-09 85.00  CAA0360812.1


Alignments:
>transmembrane protein [Arabidopsis thaliana]
Sequence ID: NP_001326023.1 Length: 74
>transmembrane protein [Arabidopsis thaliana]
Sequence ID: ANM63965.1 Length: 74
Range 1: 13 to 44

Score:55.5 bits(132), Expect:4e-13,
Method:Compositional matrix adjust.,
Identities:26/32(81%), Positives:28/32(87%), Gaps:0/32(0%)

Query  12   VRRDLHFQDLMDMELWFLYKPRLIESPVIMDM  107
            V RDLH QDL DM+LW LYKPRLI+SPVIMDM
Sbjct  13   VCRDLHLQDLTDMKLWSLYKPRLIKSPVIMDM  44


Range 2: 55 to 74

Score:38.1 bits(87), Expect:4e-13,
Method:Compositional matrix adjust.,
Identities:17/20(85%), Positives:18/20(90%), Gaps:0/20(0%)

Query  109  LQTPPPPRPYGHRTFTKFLR  168
            LQ  PPPRPYG+RTFTKFLR
Sbjct  55   LQRAPPPRPYGYRTFTKFLR  74


>hypothetical protein AT2G14265 [Arabidopsis thaliana]
Sequence ID: NP_001325174.1 Length: 111
>hypothetical protein AT2G14265 [Arabidopsis thaliana]
Sequence ID: ANM63061.1 Length: 111
Range 1: 74 to 111

Score:61.6 bits(148), Expect:5e-12,
Method:Compositional matrix adjust.,
Identities:33/48(69%), Positives:34/48(70%), Gaps:10/48(20%)

Query  20   RPPLPRPYGHGTLVSL*TTVNRISCYYGHVCRHLHLQDLMDIELLPNS  163
            RPPLPR YGHGTLVSL            HVCR LHLQDLMDI+LLPNS
Sbjct  74   RPPLPRSYGHGTLVSL----------QKHVCRDLHLQDLMDIKLLPNS  111


>unnamed protein product [Arabidopsis thaliana]
Sequence ID: CAA0384287.1 Length: 74
>unnamed protein product [Arabidopsis thaliana]
Sequence ID: VYS59150.1 Length: 74
Range 1: 13 to 44

Score:55.8 bits(133), Expect:3e-10,
Method:Compositional matrix adjust.,
Identities:26/32(81%), Positives:28/32(87%), Gaps:0/32(0%)

Query  12   VRRDLHFQDLMDMELWFLYKPRLIESPVIMDM  107
            V RDLH QDL DM+LW LYKPRLI+SPVIMDM
Sbjct  13   VCRDLHLQDLTDMKLWSLYKPRLIKSPVIMDM  44


>unnamed protein product [Arabidopsis thaliana]
Sequence ID: VYS52357.1 Length: 87
Range 1: 60 to 87

Score:46.6 bits(109), Expect:5e-10,
Method:Compositional matrix adjust.,
Identities:22/28(79%), Positives:24/28(85%), Gaps:0/28(0%)

Query  60   FLYKPRLIESPVIMDMFADTSTSKTLWT  143
            F  KPRLIES VIMD+F +TSTSKTLWT
Sbjct  60   FFTKPRLIESSVIMDLFVETSTSKTLWT  87


Range 2: 45 to 61

Score:36.2 bits(82), Expect:5e-10,
Method:Compositional matrix adjust.,
Identities:16/17(94%), Positives:16/17(94%), Gaps:0/17(0%)

Query  16  EETSTSKTLWTWNFGFF  66
           EETSTSK LWTWNFGFF
Sbjct  45  EETSTSKILWTWNFGFF  61


>unnamed protein product [Arabidopsis thaliana]
Sequence ID: VYS52323.1 Length: 61
Range 1: 39 to 61

Score:42.0 bits(97), Expect:1e-09,
Method:Composition-based stats.,
Identities:17/23(74%), Positives:19/23(82%), Gaps:0/23(0%)

Query  95   YYGHVCRHLHLQDLMDIELLPNS  163
            +  HVC+ L LQDLMDIELLPNS
Sbjct  39   FTNHVCKDLRLQDLMDIELLPNS  61


Range 2: 5 to 33

Score:39.7 bits(91), Expect:1e-09,
Method:Compositional matrix adjust.,
Identities:21/29(72%), Positives:21/29(72%), Gaps:0/29(0%)

Query  19   ETSTSKTLWTWNFGFFINHG*SNLLLLWT  105
            ETST KTLWTWNFGFF NH  S    LWT
Sbjct  5    ETSTFKTLWTWNFGFFTNHETSTSKALWT  33


>unnamed protein product [Arabidopsis thaliana]
Sequence ID: CAA0360812.1 Length: 61
Range 1: 42 to 61

Score:41.6 bits(96), Expect:2e-09,
Method:Composition-based stats.,
Identities:17/20(85%), Positives:18/20(90%), Gaps:0/20(0%)

Query  104  HVCRHLHLQDLMDIELLPNS  163
            HVC+ L LQDLMDIELLPNS
Sbjct  42   HVCKDLRLQDLMDIELLPNS  61


Range 2: 5 to 33

Score:39.7 bits(91), Expect:2e-09,
Method:Compositional matrix adjust.,
Identities:21/29(72%), Positives:21/29(72%), Gaps:0/29(0%)

Query  19   ETSTSKTLWTWNFGFFINHG*SNLLLLWT  105
            ETST KTLWTWNFGFF NH  S    LWT
Sbjct  5    ETSTFKTLWTWNFGFFTNHETSTSKALWT  33


Query #157: XLOC_019078 Query ID: lcl|Query_33160 Length: 830

Sequences producing significant alignments:
                                                                  Max    Total Query   E   Per.
Description                                                       Score  Score cover Value Ident  Accession
unnamed protein product [Arabidopsis thaliana]                    62.0   62.0  16%   2e-10 54.35  VYS53547.1
unnamed protein product [Arabidopsis thaliana]                    61.2   61.2  16%   6e-10 54.35  CAA0371968.1
unnamed protein product [Arabidopsis thaliana]                    58.9   58.9  10%   8e-10 96.55  BAB08892.1
RING/U-box protein [Arabidopsis thaliana]                         58.9   58.9  16%   1e-09 58.70  NP_680100.1
hypothetical protein [Arabidopsis thaliana]                       59.3   59.3  16%   3e-09 52.17  AAC31224.1
RING/U-box protein with C6HC-type zinc finger [Arabidopsis...     58.9   58.9  16%   3e-09 52.17  NP_180182.2
unnamed protein product [Arabidopsis thaliana]                    58.2   58.2  21%   4e-09 41.67  VYS59439.1
hypothetical protein AXX17_AT2G21960 [Arabidopsis thaliana]       58.5   58.5  16%   5e-09 52.17  OAP09299.1
unnamed protein product [Arabidopsis thaliana]                    56.2   56.2  16%   7e-09 56.52  VYS59440.1
unnamed protein product [Arabidopsis thaliana]                    57.0   57.0  16%   1e-08 52.17  CAA0384584.1
zinc finger (C3HC4-type RING finger) family protein [Arabidops... 57.0   57.0  16%   2e-08 52.17  NP_190142.1
hypothetical protein AXX17_AT3G36360 [Arabidopsis thaliana]       54.7   54.7  14%   2e-08 62.50  OAP01632.1
unnamed protein product [Arabidopsis thaliana]                    55.5   55.5  16%   4e-08 52.17  VYS59441.1
RING/U-box protein with C6HC-type zinc finger [Arabidopsis...     55.1   55.1  19%   4e-08 43.64  NP_001325919.1
unnamed protein product [Arabidopsis thaliana]                    55.5   55.5  16%   4e-08 58.70  VYS59244.1
RING/U-box protein with C6HC-type zinc finger [Arabidopsis...     55.1   55.1  19%   5e-08 43.64  NP_190140.1
unnamed protein product [Arabidopsis thaliana]                    55.1   55.1  19%   5e-08 43.64  CAA0384581.1
RING/U-box protein with C6HC-type zinc finger domain-containin... 54.7   54.7  16%   6e-08 58.70  NP_189961.1
unnamed protein product [Arabidopsis thaliana]                    54.7   54.7  16%   8e-08 58.70  CAA0384375.1
hypothetical protein AXX17_AT3G39390 [Arabidopsis thaliana]       55.1   55.1  16%   8e-08 52.17  OAP03187.1
unnamed protein product [Arabidopsis thaliana]                    53.1   53.1  14%   9e-08 60.00  VYS59198.1
unnamed protein product [Arabidopsis thaliana]                    53.1   53.1  14%   9e-08 60.00  CAA0384328.1


Alignments:
>unnamed protein product [Arabidopsis thaliana]
Sequence ID: VYS53547.1 Length: 308
Range 1: 162 to 207

Score:62.0 bits(149), Expect:2e-10,
Method:Compositional matrix adjust.,
Identities:25/46(54%), Positives:35/46(76%), Gaps:0/46(0%)

Query  440  CVNPWFNTEQMLTVDFCRHQSCVECVKRYIEVKLVEGGVLSCPHYQ  577
            C+N   N +QM +VD C H  C ECVKR+IEV+L+EG +++CPHY+
Sbjct  162  CLNDDINADQMFSVDKCGHMFCSECVKRHIEVRLLEGSLITCPHYR  207


>unnamed protein product [Arabidopsis thaliana]
Sequence ID: CAA0371968.1 Length: 399
Range 1: 162 to 207

Score:61.2 bits(147), Expect:6e-10,
Method:Compositional matrix adjust.,
Identities:25/46(54%), Positives:35/46(76%), Gaps:0/46(0%)

Query  440  CVNPWFNTEQMLTVDFCRHQSCVECVKRYIEVKLVEGGVLSCPHYQ  577
            C+N   N +QM +VD C H  C ECVKR+IEV+L+EG +++CPHY+
Sbjct  162  CLNDDINADQMFSVDKCGHMFCSECVKRHIEVRLLEGSLITCPHYR  207


>unnamed protein product [Arabidopsis thaliana]
Sequence ID: BAB08892.1 Length: 187
Range 1: 9 to 37

Score:58.9 bits(141), Expect:8e-10,
Method:Compositional matrix adjust.,
Identities:28/29(97%), Positives:29/29(100%), Gaps:0/29(0%)

Query  89  SVLEDNRPKDILLILRRFWGDFLISTLSS  3
           S+LEDNRPKDILLILRRFWGDFLISTLSS
Sbjct  9   SILEDNRPKDILLILRRFWGDFLISTLSS  37


>RING/U-box protein [Arabidopsis thaliana]
Sequence ID: NP_680100.1 Length: 213
>RING domain protein [Arabidopsis thaliana]
Sequence ID: AAY63565.1 Length: 213 >RING/U-box protein [Arabidopsis thaliana]
Sequence ID: AEE78042.1 Length: 213 >unnamed protein product [Arabidopsis thaliana]
Sequence ID: CAA0384582.1 Length: 213
Range 1: 20 to 65

Score:58.9 bits(141), Expect:1e-09,
Method:Compositional matrix adjust.,
Identities:27/46(59%), Positives:34/46(73%), Gaps:0/46(0%)

Query  440  CVNPWFNTEQMLTVDFCRHQSCVECVKRYIEVKLVEGGVLSCPHYQ  577
            C N  F +EQM  V  C H+ C+EC+KRYIEV+L+EG VL CP+YQ
Sbjct  20   CFNDDFKSEQMYYVALCNHKFCLECMKRYIEVRLLEGTVLICPYYQ  65


>hypothetical protein [Arabidopsis thaliana]
Sequence ID: AAC31224.1 Length: 638
Range 1: 162 to 207

Score:59.3 bits(142), Expect:3e-09,
Method:Compositional matrix adjust.,
Identities:24/46(52%), Positives:34/46(73%), Gaps:0/46(0%)

Query  440  CVNPWFNTEQMLTVDFCRHQSCVECVKRYIEVKLVEGGVLSCPHYQ  577
            C+N   N +QM +VD   H  C ECVKR+IEV+L+EG +++CPHY+
Sbjct  162  CLNDDINADQMFSVDKSGHMCCSECVKRHIEVRLLEGSLITCPHYR  207


>RING/U-box protein with C6HC-type zinc finger [Arabidopsis thaliana]
Sequence ID: NP_180182.2 Length: 398
>RING/U-box protein with C6HC-type zinc finger [Arabidopsis thaliana]
Sequence ID: AEC07797.1 Length: 398
Range 1: 162 to 207

Score:58.9 bits(141), Expect:3e-09,
Method:Compositional matrix adjust.,
Identities:24/46(52%), Positives:34/46(73%), Gaps:0/46(0%)

Query  440  CVNPWFNTEQMLTVDFCRHQSCVECVKRYIEVKLVEGGVLSCPHYQ  577
            C+N   N +QM +VD   H  C ECVKR+IEV+L+EG +++CPHY+
Sbjct  162  CLNDDINADQMFSVDKSGHMCCSECVKRHIEVRLLEGSLITCPHYR  207


>unnamed protein product [Arabidopsis thaliana]
Sequence ID: VYS59439.1 Length: 290
Range 1: 95 to 154

Score:58.2 bits(139), Expect:4e-09,
Method:Compositional matrix adjust.,
Identities:25/60(42%), Positives:39/60(65%), Gaps:0/60(0%)

Query  440  CVNPWFNTEQMLTVDFCRHQSCVECVKRYIEVKLVEGGVLSCPHYQFFLWCRNGGFFFWC  619
            C++   N  QM  V  CRH+ C +C++R+IEV+L+EG V+ CPHY+     + GG+  +C
Sbjct  95   CLDDDINENQMFCVGKCRHRFCSDCMRRHIEVRLLEGSVMRCPHYRCKTTLKFGGWRIYC  154


>hypothetical protein AXX17_AT2G21960 [Arabidopsis thaliana]
Sequence ID: OAP09299.1 Length: 788
Range 1: 162 to 207

Score:58.5 bits(140), Expect:5e-09,
Method:Compositional matrix adjust.,
Identities:24/46(52%), Positives:34/46(73%), Gaps:0/46(0%)

Query  440  CVNPWFNTEQMLTVDFCRHQSCVECVKRYIEVKLVEGGVLSCPHYQ  577
            C+N   N +QM +VD   H  C ECVKR+IEV+L+EG +++CPHY+
Sbjct  162  CLNDDINADQMFSVDKSGHMCCSECVKRHIEVRLLEGSLITCPHYR  207


>unnamed protein product [Arabidopsis thaliana]
Sequence ID: VYS59440.1 Length: 187
Range 1: 20 to 65

Score:56.2 bits(134), Expect:7e-09,
Method:Compositional matrix adjust.,
Identities:26/46(57%), Positives:33/46(71%), Gaps:0/46(0%)

Query  440  CVNPWFNTEQMLTVDFCRHQSCVECVKRYIEVKLVEGGVLSCPHYQ  577
            C N  F +EQM  V  C H+ C+EC+KRYIEV+L+EG V  CP+YQ
Sbjct  20   CFNDDFKSEQMYYVALCNHKFCLECMKRYIEVRLLEGTVPICPYYQ  65


>unnamed protein product [Arabidopsis thaliana]
Sequence ID: CAA0384584.1 Length: 441
Range 1: 158 to 203

Score:57.0 bits(136), Expect:1e-08,
Method:Compositional matrix adjust.,
Identities:24/46(52%), Positives:31/46(67%), Gaps:0/46(0%)

Query  440  CVNPWFNTEQMLTVDFCRHQSCVECVKRYIEVKLVEGGVLSCPHYQ  577
            C N  F  E M +VD C HQ CVEC+ +YI+V+L+E   + CPHYQ
Sbjct  158  CFNDDFKAEHMFSVDLCGHQFCVECMTQYIKVRLLEESEMRCPHYQ  203


>zinc finger (C3HC4-type RING finger) family protein [Arabidopsis thaliana]
Sequence ID: NP_190142.1 Length: 503
>zinc finger (C3HC4-type RING finger) family protein [Arabidopsis thaliana]
Sequence ID: AEE78043.1 Length: 503 >putative protein [Arabidopsis thaliana]
Sequence ID: CAB75485.1 Length: 503
Range 1: 158 to 203

Score:57.0 bits(136), Expect:2e-08,
Method:Compositional matrix adjust.,
Identities:24/46(52%), Positives:31/46(67%), Gaps:0/46(0%)

Query  440  CVNPWFNTEQMLTVDFCRHQSCVECVKRYIEVKLVEGGVLSCPHYQ  577
            C N  F  E M +VD C HQ CVEC+ +YI+V+L+E   + CPHYQ
Sbjct  158  CFNDDFKAEHMFSVDLCGHQFCVECMTQYIKVRLLEESEMRCPHYQ  203


>hypothetical protein AXX17_AT3G36360 [Arabidopsis thaliana]
Sequence ID: OAP01632.1 Length: 199
Range 1: 68 to 107

Score:54.7 bits(130), Expect:2e-08,
Method:Compositional matrix adjust.,
Identities:25/40(63%), Positives:29/40(72%), Gaps:0/40(0%)

Query  458  NTEQMLTVDFCRHQSCVECVKRYIEVKLVEGGVLSCPHYQ  577
              EQM +V  C HQ CVECVK+YIE +L+EG V  CPHYQ
Sbjct  68   KAEQMYSVALCGHQFCVECVKQYIESRLLEGCVPRCPHYQ  107


>unnamed protein product [Arabidopsis thaliana]
Sequence ID: VYS59441.1 Length: 452
Range 1: 158 to 203

Score:55.5 bits(132), Expect:4e-08,
Method:Compositional matrix adjust.,
Identities:24/46(52%), Positives:31/46(67%), Gaps:0/46(0%)

Query  440  CVNPWFNTEQMLTVDFCRHQSCVECVKRYIEVKLVEGGVLSCPHYQ  577
            C N  F  E M +V  C HQ CVEC+K+YI+V+L+E   + CPHYQ
Sbjct  158  CFNDNFKAEHMFSVALCGHQFCVECMKQYIKVRLLEESEMRCPHYQ  203


>RING/U-box protein with C6HC-type zinc finger [Arabidopsis thaliana]
Sequence ID: NP_001325919.1 Length: 319
>RING/U-box protein with C6HC-type zinc finger [Arabidopsis thaliana]
Sequence ID: ANM63851.1 Length: 319
Range 1: 74 to 128

Score:55.1 bits(131), Expect:4e-08,
Method:Compositional matrix adjust.,
Identities:24/55(44%), Positives:36/55(65%), Gaps:0/55(0%)

Query  440  CVNPWFNTEQMLTVDFCRHQSCVECVKRYIEVKLVEGGVLSCPHYQFFLWCRNGG  604
            C++   N  QM  V  CRH+ C +C++R+IEV+L+EG V+ CPHY+     + GG
Sbjct  74   CLDDDINENQMFCVGKCRHRFCSDCMRRHIEVRLLEGSVMRCPHYRCKTTLKFGG  128


>unnamed protein product [Arabidopsis thaliana]
Sequence ID: VYS59244.1 Length: 359
Range 1: 156 to 201

Score:55.5 bits(132), Expect:4e-08,
Method:Compositional matrix adjust.,
Identities:27/46(59%), Positives:30/46(65%), Gaps:0/46(0%)

Query  440  CVNPWFNTEQMLTVDFCRHQSCVECVKRYIEVKLVEGGVLSCPHYQ  577
            C N     E+M +V  C HQ CVECVK YIEVKL+EGGV  C  YQ
Sbjct  156  CFNNVLEAEKMFSVAICGHQFCVECVKHYIEVKLLEGGVPRCLDYQ  201


>RING/U-box protein with C6HC-type zinc finger [Arabidopsis thaliana]
Sequence ID: NP_190140.1 Length: 348
>RING/U-box protein with C6HC-type zinc finger [Arabidopsis thaliana]
Sequence ID: AEE78041.1 Length: 348 >putative protein [Arabidopsis thaliana]
Sequence ID: CAB75483.1 Length: 348
Range 1: 103 to 157

Score:55.1 bits(131), Expect:5e-08,
Method:Compositional matrix adjust.,
Identities:24/55(44%), Positives:36/55(65%), Gaps:0/55(0%)

Query  440  CVNPWFNTEQMLTVDFCRHQSCVECVKRYIEVKLVEGGVLSCPHYQFFLWCRNGG  604
            C++   N  QM  V  CRH+ C +C++R+IEV+L+EG V+ CPHY+     + GG
Sbjct  103  CLDDDINENQMFCVGKCRHRFCSDCMRRHIEVRLLEGSVMRCPHYRCKTTLKFGG  157


>unnamed protein product [Arabidopsis thaliana]
Sequence ID: CAA0384581.1 Length: 352
Range 1: 132 to 186

Score:55.1 bits(131), Expect:5e-08,
Method:Compositional matrix adjust.,
Identities:24/55(44%), Positives:36/55(65%), Gaps:0/55(0%)

Query  440  CVNPWFNTEQMLTVDFCRHQSCVECVKRYIEVKLVEGGVLSCPHYQFFLWCRNGG  604
            C++   N  QM  V  CRH+ C +C++R+IEV+L+EG V+ CPHY+     + GG
Sbjct  132  CLDDDINENQMFCVGKCRHRFCSDCMRRHIEVRLLEGSVMRCPHYRCKTTLKFGG  186


>RING/U-box protein with C6HC-type zinc finger domain-containing protein [Arabidopsis thaliana]
Sequence ID: NP_189961.1 Length: 346
>RING/U-box protein with C6HC-type zinc finger domain-containing protein [Arabidopsis thaliana]
Sequence ID: AEE77826.1 Length: 346 >putative protein [Arabidopsis thaliana]
Sequence ID: CAB83147.1 Length: 346
Range 1: 158 to 203

Score:54.7 bits(130), Expect:6e-08,
Method:Compositional matrix adjust.,
Identities:27/46(59%), Positives:30/46(65%), Gaps:0/46(0%)

Query  440  CVNPWFNTEQMLTVDFCRHQSCVECVKRYIEVKLVEGGVLSCPHYQ  577
            C N     E+M +V  C HQ CVECVK YIEVKL+EGGV  C  YQ
Sbjct  158  CFNNVLEAEKMFSVAICGHQFCVECVKHYIEVKLLEGGVPRCLDYQ  203


>unnamed protein product [Arabidopsis thaliana]
Sequence ID: CAA0384375.1 Length: 533
Range 1: 158 to 203

Score:54.7 bits(130), Expect:8e-08,
Method:Compositional matrix adjust.,
Identities:27/46(59%), Positives:30/46(65%), Gaps:0/46(0%)

Query  440  CVNPWFNTEQMLTVDFCRHQSCVECVKRYIEVKLVEGGVLSCPHYQ  577
            C N     E+M +V  C HQ CVECVK YIEVKL+EGGV  C  YQ
Sbjct  158  CFNNVLEAEKMFSVAICGHQFCVECVKHYIEVKLLEGGVPRCLDYQ  203


>hypothetical protein AXX17_AT3G39390 [Arabidopsis thaliana]
Sequence ID: OAP03187.1 Length: 1167
Range 1: 491 to 536

Score:55.1 bits(131), Expect:8e-08,
Method:Compositional matrix adjust.,
Identities:24/46(52%), Positives:31/46(67%), Gaps:0/46(0%)

Query  440  CVNPWFNTEQMLTVDFCRHQSCVECVKRYIEVKLVEGGVLSCPHYQ  577
            C N  F  E M +V  C HQ CVEC+K+YI+V+L+E   + CPHYQ
Sbjct  491  CFNDNFKAEHMFSVALCGHQFCVECMKQYIKVRLLEESEMRCPHYQ  536


>unnamed protein product [Arabidopsis thaliana]
Sequence ID: VYS59198.1 Length: 199
Range 1: 68 to 107

Score:53.1 bits(126), Expect:9e-08,
Method:Compositional matrix adjust.,
Identities:24/40(60%), Positives:29/40(72%), Gaps:0/40(0%)

Query  458  NTEQMLTVDFCRHQSCVECVKRYIEVKLVEGGVLSCPHYQ  577
              EQM +V  C HQ CVECVK++IE +L+EG V  CPHYQ
Sbjct  68   KAEQMYSVALCGHQFCVECVKQHIESRLLEGCVPRCPHYQ  107


>unnamed protein product [Arabidopsis thaliana]
Sequence ID: CAA0384328.1 Length: 199
Range 1: 68 to 107

Score:53.1 bits(126), Expect:9e-08,
Method:Compositional matrix adjust.,
Identities:24/40(60%), Positives:29/40(72%), Gaps:0/40(0%)

Query  458  NTEQMLTVDFCRHQSCVECVKRYIEVKLVEGGVLSCPHYQ  577
              EQM +V  C HQ CVECVK++IE +L+EG V  CPHYQ
Sbjct  68   KAEQMYSVALCGHQFCVECVKQHIESRLLEGCVPRCPHYQ  107


Query #158: XLOC_019124 Query ID: lcl|Query_33161 Length: 489

Sequences producing significant alignments:
                                                                  Max    Total Query   E   Per.
Description                                                       Score  Score cover Value Ident  Accession
unnamed protein product [Arabidopsis thaliana]                    123    123   39%   4e-33 84.21  VYS59515.1
receptor protein kinase-like protein [Arabidopsis thaliana]       104    104   44%   2e-26 76.39  NP_190212.2
receptor protein kinase-like protein [Arabidopsis thaliana]       104    104   44%   3e-26 76.39  NP_001325612.1
unnamed protein product [Arabidopsis thaliana]                    101    101   44%   4e-25 73.61  CAA0384657.1
hypothetical protein AXX17_AT3G40140 [Arabidopsis thaliana]       99.4   99.4  44%   3e-24 72.22  OAP06583.1
unknown protein [Arabidopsis thaliana]                            78.2   78.2  44%   1e-17 60.27  AAM91676.1
unknown protein [Arabidopsis thaliana]                            78.2   78.2  44%   2e-17 60.27  AAL86354.1
kinase-like protein [Arabidopsis thaliana]                        77.8   77.8  44%   1e-16 60.27  NP_190213.2
unnamed protein product [Arabidopsis thaliana]                    77.0   77.0  44%   2e-16 60.27  CAA0384659.1
hypothetical protein AXX17_AT3G40170 [Arabidopsis thaliana]       75.5   75.5  44%   6e-16 58.90  OAP02871.1
putative protein (fragment) [Arabidopsis thaliana]                73.9   73.9  34%   2e-15 69.64  CAB90952.1
putative protein [Arabidopsis thaliana]                           66.2   66.2  25%   1e-12 80.49  CAB90954.1


Alignments:
>unnamed protein product [Arabidopsis thaliana]
Sequence ID: VYS59515.1 Length: 458
Range 1: 383 to 458

Score:123 bits(308), Expect:4e-33,
Method:Compositional matrix adjust.,
Identities:64/76(84%), Positives:64/76(84%), Gaps:12/76(15%)

Query  3    LILGVTFAFVFVILWSAFVVSILRKRQNAMQQSNKSPTMST------------EHGIRTG  146
            LILGVTFAFVFVILWSAFVVSILRKRQNAMQQSNKSPTMST            EHGIRTG
Sbjct  383  LILGVTFAFVFVILWSAFVVSILRKRQNAMQQSNKSPTMSTGHIGFWSLLCPQEHGIRTG  442

Query  147  TSPLFGQETVSDKIDS  194
            TSPLFGQETVSDKIDS
Sbjct  443  TSPLFGQETVSDKIDS  458


>receptor protein kinase-like protein [Arabidopsis thaliana]
Sequence ID: NP_190212.2 Length: 470
>receptor protein kinase-like protein [Arabidopsis thaliana]
Sequence ID: AEE78141.1 Length: 470 >putative protein [Arabidopsis thaliana]
Sequence ID: BAD43930.1 Length: 470
Range 1: 399 to 469

Score:104 bits(260), Expect:2e-26,
Method:Compositional matrix adjust.,
Identities:55/72(76%), Positives:58/72(80%), Gaps:1/72(1%)

Query  3    LILGVTFAFVFVILWSAFVVSILRKRQNAMQQSNKSPTMSTEHGIRTGTSPLFGQETVSD  182
            LILGVTFA VFVILWSAFVV+ILRKRQNA   SN +PT STE GI TGTSPLFGQE  SD
Sbjct  399  LILGVTFASVFVILWSAFVVNILRKRQNAKPVSNTAPTTSTERGIGTGTSPLFGQEMASD  458

Query  183  KIDS*YVVQDEH  218
              DS YVVQ+E
Sbjct  459  TNDS-YVVQNEQ  469


>receptor protein kinase-like protein [Arabidopsis thaliana]
Sequence ID: NP_001325612.1 Length: 478
>receptor protein kinase-like protein [Arabidopsis thaliana]
Sequence ID: ANM63530.1 Length: 478 >unnamed protein product [Arabidopsis thaliana]
Sequence ID: VYS59516.1 Length: 478
Range 1: 407 to 477

Score:104 bits(260), Expect:3e-26,
Method:Compositional matrix adjust.,
Identities:55/72(76%), Positives:58/72(80%), Gaps:1/72(1%)

Query  3    LILGVTFAFVFVILWSAFVVSILRKRQNAMQQSNKSPTMSTEHGIRTGTSPLFGQETVSD  182
            LILGVTFA VFVILWSAFVV+ILRKRQNA   SN +PT STE GI TGTSPLFGQE  SD
Sbjct  407  LILGVTFASVFVILWSAFVVNILRKRQNAKPVSNTAPTTSTERGIGTGTSPLFGQEMASD  466

Query  183  KIDS*YVVQDEH  218
              DS YVVQ+E
Sbjct  467  TNDS-YVVQNEQ  477


>unnamed protein product [Arabidopsis thaliana]
Sequence ID: CAA0384657.1 Length: 478
Range 1: 407 to 477

Score:101 bits(252), Expect:4e-25,
Method:Compositional matrix adjust.,
Identities:53/72(74%), Positives:57/72(79%), Gaps:1/72(1%)

Query  3    LILGVTFAFVFVILWSAFVVSILRKRQNAMQQSNKSPTMSTEHGIRTGTSPLFGQETVSD  182
            LILGVTFA VFV+LWSAFVV+ILRKRQNA   SN +PT STE GI TGTSPLFGQE  SD
Sbjct  407  LILGVTFASVFVLLWSAFVVNILRKRQNAKPVSNTAPTTSTERGIGTGTSPLFGQEMASD  466

Query  183  KIDS*YVVQDEH  218
            + DS YV Q E
Sbjct  467  RNDS-YVGQGEQ  477


>hypothetical protein AXX17_AT3G40140 [Arabidopsis thaliana]
Sequence ID: OAP06583.1 Length: 467
Range 1: 396 to 466

Score:99.4 bits(246), Expect:3e-24,
Method:Compositional matrix adjust.,
Identities:52/72(72%), Positives:57/72(79%), Gaps:1/72(1%)

Query  3    LILGVTFAFVFVILWSAFVVSILRKRQNAMQQSNKSPTMSTEHGIRTGTSPLFGQETVSD  182
            LILGVTFA VFV+LWSAFVV+ILRKRQNA   SN +PT+STE GI TGTSPLFGQE  SD
Sbjct  396  LILGVTFASVFVLLWSAFVVNILRKRQNAKPLSNTAPTISTERGIGTGTSPLFGQEMASD  455

Query  183  KIDS*YVVQDEH  218
            + D  YV Q E
Sbjct  456  RNDW-YVGQGEQ  466


>unknown protein [Arabidopsis thaliana]
Sequence ID: AAM91676.1 Length: 226
Range 1: 155 to 225

Score:78.2 bits(191), Expect:1e-17,
Method:Compositional matrix adjust.,
Identities:44/73(60%), Positives:50/73(68%), Gaps:3/73(4%)

Query  3    LILGVTFAFVFVILWSAFVVSILRKRQNAMQQSNKS-PTMSTEHGIRTGTSPLFGQETVS  179
            LILGVTFA  F +L SAF  +I +KRQNA  QSN + P  STEHG  TG SPL GQ+  S
Sbjct  155  LILGVTFASAFAVLASAFA-AIWKKRQNAKSQSNSTVPNTSTEHGTGTGMSPLVGQQFAS  213

Query  180  DKIDS*YVVQDEH  218
            D  DS YVV+D H
Sbjct  214  DMTDS-YVVEDGH  225


>unknown protein, partial [Arabidopsis thaliana]
Sequence ID: AAL86354.1 Length: 262
Range 1: 191 to 261

Score:78.2 bits(191), Expect:2e-17,
Method:Compositional matrix adjust.,
Identities:44/73(60%), Positives:50/73(68%), Gaps:3/73(4%)

Query  3    LILGVTFAFVFVILWSAFVVSILRKRQNAMQQSNKS-PTMSTEHGIRTGTSPLFGQETVS  179
            LILGVTFA  F +L SAF  +I +KRQNA  QSN + P  STEHG  TG SPL GQ+  S
Sbjct  191  LILGVTFASAFAVLASAFA-AIWKKRQNAKSQSNSTVPNTSTEHGTGTGMSPLVGQQFAS  249

Query  180  DKIDS*YVVQDEH  218
            D  DS YVV+D H
Sbjct  250  DMTDS-YVVEDGH  261


>kinase-like protein [Arabidopsis thaliana]
Sequence ID: NP_190213.2 Length: 471
>kinase-like protein [Arabidopsis thaliana]
Sequence ID: AEE78142.1 Length: 471 >unknown protein [Arabidopsis thaliana]
Sequence ID: BAC42269.1 Length: 471 >unnamed protein product [Arabidopsis thaliana]
Sequence ID: VYS59517.1 Length: 471
Range 1: 400 to 470

Score:77.8 bits(190), Expect:1e-16,
Method:Compositional matrix adjust.,
Identities:44/73(60%), Positives:50/73(68%), Gaps:3/73(4%)

Query  3    LILGVTFAFVFVILWSAFVVSILRKRQNAMQQSNKS-PTMSTEHGIRTGTSPLFGQETVS  179
            LILGVTFA  F +L SAF  +I +KRQNA  QSN + P  STEHG  TG SPL GQ+  S
Sbjct  400  LILGVTFASAFAVLASAFA-AIWKKRQNAKSQSNSTVPNTSTEHGTGTGMSPLVGQQFAS  458

Query  180  DKIDS*YVVQDEH  218
            D  DS YVV+D H
Sbjct  459  DMTDS-YVVEDGH  470


>unnamed protein product [Arabidopsis thaliana]
Sequence ID: CAA0384659.1 Length: 471
Range 1: 400 to 470

Score:77.0 bits(188), Expect:2e-16,
Method:Compositional matrix adjust.,
Identities:44/73(60%), Positives:50/73(68%), Gaps:3/73(4%)

Query  3    LILGVTFAFVFVILWSAFVVSILRKRQNAMQQSNKS-PTMSTEHGIRTGTSPLFGQETVS  179
            LILGVTFA  F +L SAF  +I +KRQNA  QSN + P  STEHG  TG SPL GQ+  S
Sbjct  400  LILGVTFASAFAVLASAFA-AIWKKRQNAKSQSNSTVPNTSTEHGTGTGMSPLVGQQFAS  458

Query  180  DKIDS*YVVQDEH  218
            D  DS YVV+D H
Sbjct  459  DMSDS-YVVEDGH  470


>hypothetical protein AXX17_AT3G40170 [Arabidopsis thaliana]
Sequence ID: OAP02871.1 Length: 470
Range 1: 399 to 469

Score:75.5 bits(184), Expect:6e-16,
Method:Compositional matrix adjust.,
Identities:43/73(59%), Positives:49/73(67%), Gaps:3/73(4%)

Query  3    LILGVTFAFVFVILWSAFVVSILRKRQNAMQQSNKS-PTMSTEHGIRTGTSPLFGQETVS  179
            LILGVTFA  F +L SAF  +I +KRQNA  QSN + P  STEHG  TG SPL GQ+  S
Sbjct  399  LILGVTFASAFAVLASAFA-AIWKKRQNAKSQSNSTVPNTSTEHGAGTGMSPLVGQQFAS  457

Query  180  DKIDS*YVVQDEH  218
            D  DS YV +D H
Sbjct  458  DMSDS-YVAEDGH  469


>putative protein (fragment), partial [Arabidopsis thaliana]
Sequence ID: CAB90952.1 Length: 410
Range 1: 355 to 409

Score:73.9 bits(180), Expect:2e-15,
Method:Compositional matrix adjust.,
Identities:39/56(70%), Positives:42/56(75%), Gaps:1/56(1%)

Query  51   AFVVSILRKRQNAMQQSNKSPTMSTEHGIRTGTSPLFGQETVSDKIDS*YVVQDEH  218
            AFVV+ILRKRQNA   SN +PT STE GI TGTSPLFGQE  SD+ DS YV Q E
Sbjct  355  AFVVNILRKRQNAKPVSNTAPTTSTERGIGTGTSPLFGQEMASDRNDS-YVGQGEQ  409


>putative protein [Arabidopsis thaliana]
Sequence ID: CAB90954.1 Length: 457
Range 1: 399 to 439

Score:66.2 bits(160), Expect:1e-12,
Method:Compositional matrix adjust.,
Identities:33/41(80%), Positives:35/41(85%), Gaps:0/41(0%)

Query  3    LILGVTFAFVFVILWSAFVVSILRKRQNAMQQSNKSPTMST  125
            LILGVTFA VFVILWSAFVV+ILRKRQNA   SN +PT ST
Sbjct  399  LILGVTFASVFVILWSAFVVNILRKRQNAKPVSNTAPTTST  439


Query #159: XLOC_019165 Query ID: lcl|Query_33162 Length: 669

Sequences producing significant alignments:
                                                                  Max    Total Query   E   Per.
Description                                                       Score  Score cover Value Ident  Accession
unnamed protein product [Arabidopsis thaliana]                    65.1   65.1  23%   1e-12 98.11  VYS59583.1
unnamed protein product [Arabidopsis thaliana]                    65.1   65.1  23%   3e-12 96.23  CAA0384715.1
unnamed protein product [Arabidopsis thaliana]                    50.4   50.4  24%   5e-08 74.07  CAA0301835.1
hypothetical protein AT2G46765 [Arabidopsis thaliana]             49.7   49.7  24%   9e-08 75.93  NP_001323720.1


Alignments:
>unnamed protein product [Arabidopsis thaliana]
Sequence ID: VYS59583.1 Length: 147
Range 1: 77 to 129

Score:65.1 bits(157), Expect:1e-12,
Method:Compositional matrix adjust.,
Identities:52/53(98%), Positives:53/53(100%), Gaps:0/53(0%)

Query  366  GRSNYGRGRFFSILQRLRCRSLSIEdldllpfflrlcllgpgMLIDVSAPFMI  524
            GRSNYGRGRFFSILQRLRCRSLSIEDLDLLPFFLRLCLLGPGMLIDVSAPFM+
Sbjct  77   GRSNYGRGRFFSILQRLRCRSLSIEDLDLLPFFLRLCLLGPGMLIDVSAPFMV  129


>unnamed protein product [Arabidopsis thaliana]
Sequence ID: CAA0384715.1 Length: 224
Range 1: 154 to 206

Score:65.1 bits(157), Expect:3e-12,
Method:Compositional matrix adjust.,
Identities:51/53(96%), Positives:52/53(98%), Gaps:0/53(0%)

Query  366  GRSNYGRGRFFSILQRLRCRSLSIEdldllpfflrlcllgpgMLIDVSAPFMI  524
            GRSNYGRGRFFSILQRLRCRSLSIEDLDLLPFF RLCLLGPGMLIDVSAPFM+
Sbjct  154  GRSNYGRGRFFSILQRLRCRSLSIEDLDLLPFFFRLCLLGPGMLIDVSAPFMV  206


>unnamed protein product [Arabidopsis thaliana]
Sequence ID: CAA0301835.1 Length: 74
Range 1: 16 to 69

Score:50.4 bits(119), Expect:5e-08,
Method:Compositional matrix adjust.,
Identities:40/54(74%), Positives:46/54(85%), Gaps:0/54(0%)

Query  352  QTKETnkqqnrkrkrfrlyrfRSENLEKKPLEENRIQITLNPQEIETRDTKTKK  191
            QTKETNKQQN + K FRL++ RSENLE++PLEENRIQITLNPQE ET +T  KK
Sbjct  16   QTKETNKQQNWESKGFRLHKSRSENLEQRPLEENRIQITLNPQENETIETTPKK  69


>hypothetical protein AT2G46765 [Arabidopsis thaliana]
Sequence ID: NP_001323720.1 Length: 74
>hypothetical protein AT2G46765 [Arabidopsis thaliana]
Sequence ID: ANM61504.1 Length: 74 >unnamed protein product [Arabidopsis thaliana]
Sequence ID: VYS55751.1 Length: 74
Range 1: 16 to 69

Score:49.7 bits(117), Expect:9e-08,
Method:Compositional matrix adjust.,
Identities:41/54(76%), Positives:46/54(85%), Gaps:0/54(0%)

Query  352  QTKETnkqqnrkrkrfrlyrfRSENLEKKPLEENRIQITLNPQEIETRDTKTKK  191
            QTKETNKQQN K K FRL+R RS NLE++PLEENRIQITLNPQE ET++  TKK
Sbjct  16   QTKETNKQQNWKSKGFRLHRSRSVNLEQRPLEENRIQITLNPQENETKEITTKK  69


Query #160: XLOC_019982 Query ID: lcl|Query_33163 Length: 949

No significant similarity found.


Query #161: XLOC_020123 Query ID: lcl|Query_33164 Length: 635

No significant similarity found.


Query #162: XLOC_020124 Query ID: lcl|Query_33165 Length: 517

No significant similarity found.


Query #163: XLOC_020125 Query ID: lcl|Query_33166 Length: 587

No significant similarity found.


Query #164: XLOC_020126 Query ID: lcl|Query_33167 Length: 564

No significant similarity found.


Query #165: XLOC_020127 Query ID: lcl|Query_33168 Length: 673

Sequences producing significant alignments:
                                                                  Max    Total Query   E   Per.
Description                                                       Score  Score cover Value Ident  Accession
unnamed protein product [Arabidopsis thaliana]                    62.4   62.4  48%   3e-11 30.00  VYS57115.1
Polynucleotidyl transferase, ribonuclease H-like superfamily...   62.4   62.4  48%   3e-11 30.00  NP_187847.1
unnamed protein product [Arabidopsis thaliana]                    57.8   57.8  48%   1e-09 32.50  CAA0382170.1
Polynucleotidyl transferase, ribonuclease H-like superfamily...   56.6   56.6  48%   4e-09 30.25  NP_187852.1
hypothetical protein AXX17_AT3G12490 [Arabidopsis thaliana]       56.2   56.2  48%   6e-09 31.67  OAP06054.1
unknown [Arabidopsis thaliana]                                    55.1   55.1  48%   1e-08 31.67  AAM66064.1
Polynucleotidyl transferase, ribonuclease H-like superfamily...   55.1   55.1  48%   1e-08 31.67  NP_566424.1


Alignments:
>unnamed protein product [Arabidopsis thaliana]
Sequence ID: VYS57115.1 Length: 230
Range 1: 106 to 223

Score:62.4 bits(150), Expect:3e-11,
Method:Compositional matrix adjust.,
Identities:36/120(30%), Positives:59/120(49%), Gaps:13/120(10%)

Query  203  IVKLRYCSRISRRPTRFLGSDLTTSPGIWNDRDSTKFFARSDVLKIGKFVDGRKFMRDIW  382
            I++L YC ++      FL    TT  G+WN +D+ K       L+IG+ +D R+++ D W
Sbjct  106  IIQLGYCDQVPNNLRSFLADPETTFVGVWNGQDAGKLARCCHQLEIGELLDIRRYVTDSW  165

Query  383  NRSY*QR-----------HQKRHISPANITTKSNWGSPKLGRKQILHATVDAYVSFKLAI  529
             RS  +            +Q   + P    + S+W +  L   QIL A++DAYV  +L +
Sbjct  166  GRSMSRSSFEEIVEECMGYQGVMLDPE--ISMSDWTAYDLDLDQILQASLDAYVCHQLGV  223


>Polynucleotidyl transferase, ribonuclease H-like superfamily protein [Arabidopsis thaliana]
Sequence ID: NP_187847.1 Length: 230
>hypothetical protein; 80835-81527 [Arabidopsis thaliana]
Sequence ID: AAG51022.1 Length: 230 >Polynucleotidyl transferase, ribonuclease H-like superfamily protein [Arabidopsis thaliana]
Sequence ID: AEE75194.1 Length: 230 >hypothetical protein AXX17_AT3G12430 [Arabidopsis thaliana]
Sequence ID: OAP01473.1 Length: 230 >unnamed protein product [Arabidopsis thaliana]
Sequence ID: CAA0382165.1 Length: 230 >unnamed protein product [Arabidopsis thaliana]
Sequence ID: BAB03148.1 Length: 230
Range 1: 106 to 223

Score:62.4 bits(150), Expect:3e-11,
Method:Compositional matrix adjust.,
Identities:36/120(30%), Positives:59/120(49%), Gaps:13/120(10%)

Query  203  IVKLRYCSRISRRPTRFLGSDLTTSPGIWNDRDSTKFFARSDVLKIGKFVDGRKFMRDIW  382
            I++L YC ++      FL    TT  G+WN +D+ K       L+IG+ +D R+++ D W
Sbjct  106  IIQLGYCDQVPNNLRSFLADPETTFVGVWNGQDAGKLARCCHQLEIGELLDIRRYVTDSW  165

Query  383  NRSY*QR-----------HQKRHISPANITTKSNWGSPKLGRKQILHATVDAYVSFKLAI  529
             RS  +            +Q   + P    + S+W +  L   QIL A++DAYV  +L +
Sbjct  166  GRSMRRSSFEEIVEECMGYQGVMLDPE--ISMSDWTAYDLDLDQILQASLDAYVCHQLGV  223


>unnamed protein product [Arabidopsis thaliana]
Sequence ID: CAA0382170.1 Length: 220
Range 1: 88 to 205

Score:57.8 bits(138), Expect:1e-09,
Method:Compositional matrix adjust.,
Identities:39/120(33%), Positives:61/120(50%), Gaps:13/120(10%)

Query  203  IVKLRYCSRISRRPTRFLGSDLTTSPGIWNDRDSTKFFARSDVLKIGKFVDGRKFMRD--  376
            I++L Y  R+ +    FL    TT  G+WN +D  K       ++IGK +D R F+RD
Sbjct  88   IIQLGYNYRLPKVLRTFLADPKTTFVGVWNGQDQKKLEKCRHRVEIGKLLDIRMFVRDSR  147

Query  377  ---IWNRSY*QRHQKR------HISPANITTKSNWGSPKLGRKQILHATVDAYVSFKLAI  529
               +   S+ Q  ++R       + PA     S+WG   L   Q+L A++++YV FKLA+
Sbjct  148  GARMCFCSFEQIVKERLGRVGVRLDPA--ICMSDWGVYNLNHYQVLQASIESYVCFKLAV  205


>Polynucleotidyl transferase, ribonuclease H-like superfamily protein [Arabidopsis thaliana]
Sequence ID: NP_187852.1 Length: 242
>hypothetical protein; 72371-73099 [Arabidopsis thaliana]
Sequence ID: AAG51003.1 Length: 242 >Polynucleotidyl transferase, ribonuclease H-like superfamily protein [Arabidopsis thaliana]
Sequence ID: AEE75198.1 Length: 242 >hypothetical protein AXX17_AT3G12480 [Arabidopsis thaliana]
Sequence ID: OAP02315.1 Length: 242 >unnamed protein product [Arabidopsis thaliana]
Sequence ID: CAA0382169.1 Length: 242 >unnamed protein product [Arabidopsis thaliana]
Sequence ID: VYS57120.1 Length: 242
Range 1: 114 to 232

Score:56.6 bits(135), Expect:4e-09,
Method:Compositional matrix adjust.,
Identities:36/119(30%), Positives:53/119(44%), Gaps:10/119(8%)

Query  203  IVKLRYCSRISRRPTRFLGSDLTTSPGIWNDRDSTKFFARSDVLKIGKFVDGRKFMRDIW  382
            I++LRYC R+ +    FL     T  GIWN +D+ K       L+I + +D R+F+ D
Sbjct  114  IIQLRYCERVPQVLRNFLADRDNTFVGIWNSQDAGKLERSRHQLEIAELMDLREFVSDSS  173

Query  383  NRSY*QRHQKRHISPANI----------TTKSNWGSPKLGRKQILHATVDAYVSFKLAI  529
             R     +    I   N+           + S+W    L   QIL A++D Y    LAI
Sbjct  174  GRRSMYNYSLEKIVEENLGYPGVRLDREVSMSDWRVYNLSYDQILQASIDVYACCSLAI  232


>hypothetical protein AXX17_AT3G12490 [Arabidopsis thaliana]
Sequence ID: OAP06054.1 Length: 220
Range 1: 88 to 205

Score:56.2 bits(134), Expect:6e-09,
Method:Compositional matrix adjust.,
Identities:38/120(32%), Positives:60/120(50%), Gaps:13/120(10%)

Query  203  IVKLRYCSRISRRPTRFLGSDLTTSPGIWNDRDSTKFFARSDVLKIGKFVDGRKFMRD--  376
            I++L Y   + +    FL    TT  G+WN +D  K       ++IGK +D R F+RD
Sbjct  88   IIQLGYIYGLPKVLRTFLADPKTTFVGVWNGQDQKKLEKCRHRVEIGKLLDIRMFVRDSR  147

Query  377  ---IWNRSY*QRHQKR------HISPANITTKSNWGSPKLGRKQILHATVDAYVSFKLAI  529
               +   S+ Q  ++R       + PA     S+WG   L   Q+L A++++YV FKLA+
Sbjct  148  GARMCFCSFEQIVKERLGRVGVRLDPA--ICMSDWGVYNLNHYQVLQASIESYVCFKLAV  205


>unknown [Arabidopsis thaliana]
Sequence ID: AAM66064.1 Length: 220
>unnamed protein product [Arabidopsis thaliana]
Sequence ID: VYS57121.1 Length: 220
Range 1: 88 to 205

Score:55.1 bits(131), Expect:1e-08,
Method:Compositional matrix adjust.,
Identities:38/120(32%), Positives:60/120(50%), Gaps:13/120(10%)

Query  203  IVKLRYCSRISRRPTRFLGSDLTTSPGIWNDRDSTKFFARSDVLKIGKFVDGRKFMRD--  376
            I++L Y   + +    FL    TT  G+WN +D  K       ++IGK +D R F+RD
Sbjct  88   IIQLGYNYGLPKVLRTFLADPKTTFVGVWNGQDQKKLEKCRHRVEIGKLLDIRMFVRDSR  147

Query  377  ---IWNRSY*QRHQKR------HISPANITTKSNWGSPKLGRKQILHATVDAYVSFKLAI  529
               +   S+ Q  ++R       + PA     S+WG   L   Q+L A++++YV FKLA+
Sbjct  148  GARMCFCSFEQIVKERLGRVGVRLDPA--ICMSDWGVYNLNHYQVLQASIESYVCFKLAV  205


>Polynucleotidyl transferase, ribonuclease H-like superfamily protein [Arabidopsis thaliana]
Sequence ID: NP_566424.1 Length: 220
>hypothetical protein; 71009-71671 [Arabidopsis thaliana]
Sequence ID: AAG51036.1 Length: 220 >At3g12470 [Arabidopsis thaliana]
Sequence ID: ABF74709.1 Length: 220 >Polynucleotidyl transferase, ribonuclease H-like superfamily protein [Arabidopsis thaliana]
Sequence ID: AEE75199.1 Length: 220 >unnamed protein product [Arabidopsis thaliana]
Sequence ID: BAB03154.1 Length: 220
Range 1: 88 to 205

Score:55.1 bits(131), Expect:1e-08,
Method:Compositional matrix adjust.,
Identities:38/120(32%), Positives:60/120(50%), Gaps:13/120(10%)

Query  203  IVKLRYCSRISRRPTRFLGSDLTTSPGIWNDRDSTKFFARSDVLKIGKFVDGRKFMRD--  376
            I++L Y   + +    FL    TT  G+WN +D  K       ++IGK +D R F+RD
Sbjct  88   IIQLGYNYGLPKVLRTFLADPKTTFVGVWNGQDQKKLEKCRHRVEIGKLLDIRMFVRDSR  147

Query  377  ---IWNRSY*QRHQKR------HISPANITTKSNWGSPKLGRKQILHATVDAYVSFKLAI  529
               +   S+ Q  ++R       + PA     S+WG   L   Q+L A++++YV FKLA+
Sbjct  148  GARMCFCSFEQIVKERLGRVGVRLDPA--ICMSDWGVYNLNHYQVLQASIESYVCFKLAV  205


Query #166: XLOC_020128 Query ID: lcl|Query_33169 Length: 570

No significant similarity found.


Query #167: XLOC_020129 Query ID: lcl|Query_33170 Length: 649

No significant similarity found.


Query #168: XLOC_020130 Query ID: lcl|Query_33171 Length: 384

No significant similarity found.


Query #169: XLOC_020131 Query ID: lcl|Query_33172 Length: 769

Sequences producing significant alignments:
                                                                  Max    Total Query   E   Per.
Description                                                       Score  Score cover Value Ident  Accession
F-box and associated interaction domains-containing protein...    112    112   47%   3e-28 52.89  NP_200864.1
F-box and associated interaction domains-containing protein...    112    112   47%   6e-28 52.89  NP_001330054.1
hypothetical protein AXX17_AT5G60010 [Arabidopsis thaliana]       111    111   47%   8e-28 52.07  OAO92930.1
unnamed protein product [Arabidopsis thaliana]                    110    110   47%   2e-27 52.89  CAA0411101.1
hypothetical protein AXX17_AT3G21160 [Arabidopsis thaliana]       108    108   47%   1e-26 49.59  OAP04329.1
F-box and associated interaction domains-containing protein...    106    106   46%   5e-26 49.17  NP_188622.1
unnamed protein product [Arabidopsis thaliana]                    105    105   47%   1e-25 50.41  VYS57980.1
F-box and associated interaction domains-containing protein...    103    103   47%   8e-25 49.59  NP_188638.1
unnamed protein product [Arabidopsis thaliana]                    103    103   47%   9e-25 49.59  CAA0383045.1
unnamed protein product [Arabidopsis thaliana]                    99.8   99.8  26%   5e-24 68.66  VYS57963.1
unnamed protein product [Arabidopsis thaliana]                    99.4   99.4  26%   1e-23 68.66  CAA0383027.1
F-box family protein [Arabidopsis thaliana]                       95.9   95.9  55%   5e-22 40.69  NP_188623.1
unnamed protein product [Arabidopsis thaliana]                    95.9   95.9  55%   8e-22 40.69  BAB01303.1
unnamed protein product [Arabidopsis thaliana]                    89.4   89.4  47%   5e-21 44.63  CAA0261612.1
F-box family protein [Arabidopsis thaliana]                       90.1   90.1  55%   9e-20 41.55  NP_174493.1
F-box family protein [Arabidopsis thaliana]                       83.2   83.2  43%   5e-19 46.28  ABE65746.1
unnamed protein product [Arabidopsis thaliana]                    87.4   87.4  47%   6e-19 44.63  VYS47772.1
RecName: Full=Putative F-box protein At3g13624 [Arabidopsis...    84.7   84.7  48%   2e-18 39.20  Q9LHN6.1
RecName: Full=F-box protein At1g59680 [Arabidopsis thaliana]      84.3   84.3  47%   4e-18 44.72  Q9LQ46.2
unnamed protein product [Arabidopsis thaliana]                    84.0   84.0  26%   5e-18 60.29  CAA0383028.1
hypothetical protein AXX17_AT3G13970 [Arabidopsis thaliana]       84.3   84.3  48%   7e-18 39.20  OAP04380.1
unnamed protein product [Arabidopsis thaliana]                    83.6   83.6  43%   7e-18 46.28  VYS50192.1
F-box and associated interaction domains-containing protein...    83.6   83.6  43%   7e-18 46.28  NP_176822.1
RecName: Full=F-box protein At1g66490 [Arabidopsis thaliana]      83.6   83.6  43%   8e-18 46.28  Q1PFG1.2
unnamed protein product [Arabidopsis thaliana]                    83.6   83.6  46%   8e-18 43.33  CAA0321408.1
F-box and associated interaction domains-containing protein...    83.6   83.6  46%   9e-18 43.33  NP_176912.1
F-box and associated interaction domains-containing protein...    82.0   82.0  49%   3e-17 40.16  NP_567289.1
unknown protein; 98896-95855 [Arabidopsis thaliana]               81.6   81.6  49%   5e-17 38.89  AAB65488.1
unnamed protein product [Arabidopsis thaliana]                    81.3   81.3  46%   5e-17 43.33  VYS50284.1
unnamed protein product [Arabidopsis thaliana]                    80.9   80.9  43%   6e-17 45.45  CAA0319417.1
hypothetical protein AXX17_AT1G11130 [Arabidopsis thaliana]       80.5   80.5  55%   9e-17 34.87  OAP16214.1
RecName: Full=F-box protein At1g10895 [Arabidopsis thaliana]      80.5   80.5  55%   9e-17 34.87  P0CB27.1
unnamed protein product [Arabidopsis thaliana]                    80.5   80.5  49%   9e-17 40.16  VYS61789.1
unnamed protein product [Arabidopsis thaliana]                    80.1   80.1  45%   1e-16 44.07  CAA0385286.1
unnamed protein product [Arabidopsis thaliana]                    80.1   80.1  46%   1e-16 41.86  CAA0395568.1
F-box family protein [Arabidopsis thaliana]                       80.1   80.1  45%   1e-16 44.07  NP_190520.2
unnamed protein product [Arabidopsis thaliana]                    80.1   80.1  45%   2e-16 44.07  VYS59865.1
putative protein [Arabidopsis thaliana]                           79.7   79.7  45%   3e-16 44.07  CAB62455.1
F-box family protein [Arabidopsis thaliana]                       79.0   79.0  44%   4e-16 43.97  NP_001117347.1
F-box associated ubiquitination effector family protein...        78.2   78.2  52%   4e-16 38.65  NP_001323259.1
putative F5A9.18 protein [Arabidopsis thaliana]                   79.0   79.0  44%   4e-16 43.97  AAL86342.1
unnamed protein product [Arabidopsis thaliana]                    79.0   79.0  44%   4e-16 43.97  VYS47126.1
hypothetical protein [Arabidopsis thaliana]                       79.0   79.0  44%   4e-16 43.97  BAD94008.1
F-box and associated interaction domains-containing protein...    78.6   78.6  48%   4e-16 38.28  NP_187980.1
unnamed protein product [Arabidopsis thaliana]                    78.6   78.6  48%   4e-16 37.80  CAA0382322.1
hypothetical protein AXX17_AT3G14080 [Arabidopsis thaliana]       78.6   78.6  48%   4e-16 37.80  OAP01484.1
unnamed protein product [Arabidopsis thaliana]                    78.6   78.6  46%   5e-16 40.60  VYS63000.1
hypothetical protein AXX17_AT3G43640 [Arabidopsis thaliana]       79.0   79.0  46%   6e-16 42.50  OAP03761.1
hypothetical protein [Arabidopsis thaliana]                       78.2   78.2  44%   8e-16 43.10  BAF00445.1
F-box and associated interaction domains-containing protein...    77.8   77.8  46%   8e-16 40.60  NP_193453.1
F-box and associated interaction domains-containing protein...    77.4   77.4  53%   1e-15 34.03  NP_564661.1
unnamed protein product [Arabidopsis thaliana]                    77.8   77.8  44%   1e-15 43.10  CAA0240873.1
F-box and associated interaction domains-containing protein...    75.1   75.1  46%   1e-15 43.70  NP_001321070.1
hypothetical protein AXX17_AT1G61370 [Arabidopsis thaliana]       73.9   73.9  46%   1e-15 41.18  OAP16804.1
Contains similarity to hypothetical protein gb|Z97343 from A....  75.5   75.5  46%   2e-15 42.02  AAC18790.1
T30E16.27 [Arabidopsis thaliana]                                  76.6   76.6  27%   2e-15 55.56  AAF79759.1
hypothetical protein AXX17_AT4G20250 [Arabidopsis thaliana]       77.0   77.0  46%   2e-15 39.85  OAP00162.1
unnamed protein product [Arabidopsis thaliana]                    75.5   75.5  46%   2e-15 43.70  CAA0321428.1
EDA1 [Arabidopsis thaliana]                                       76.6   76.6  47%   2e-15 38.93  OAP13995.1
hypothetical protein AXX17_AT3G43650 [Arabidopsis thaliana]       76.3   76.3  45%   3e-15 40.68  OAP01578.1
F-box and associated interaction domains-containing protein...    75.5   75.5  46%   6e-15 40.00  NP_190521.1
unnamed protein product [Arabidopsis thaliana]                    75.1   75.1  46%   6e-15 40.00  CAA0385288.1
RecName: Full=Probable F-box protein At1g67455 [Arabidopsis...    75.1   75.1  46%   7e-15 42.02  O64800.3
hypothetical protein AXX17_AT1G60230 [Arabidopsis thaliana]       74.7   74.7  43%   9e-15 44.63  OAP18488.1
unnamed protein product [Arabidopsis thaliana]                    73.2   73.2  46%   3e-14 39.17  VYS59866.1
unnamed protein product [Arabidopsis thaliana]                    72.4   72.4  47%   6e-14 39.20  BAB02540.1
F-box associated ubiquitination effector family protein...        71.2   71.2  26%   6e-14 53.62  NP_176177.1
unnamed protein product [Arabidopsis thaliana]                    72.4   72.4  47%   6e-14 39.20  CAA0382992.1
F-box family protein [Arabidopsis thaliana]                       72.0   72.0  47%   7e-14 39.20  NP_188590.2
hypothetical protein [Arabidopsis thaliana]                       70.5   70.5  42%   3e-13 39.34  CAB10501.1
RecName: Full=Putative F-box protein At3g13825 [Arabidopsis...    65.5   65.5  51%   2e-12 35.77  Q9LRW5.1
unnamed protein product [Arabidopsis thaliana]                    66.6   66.6  43%   5e-12 40.00  CAA0382341.1
unnamed protein product [Arabidopsis thaliana]                    66.6   66.6  43%   5e-12 40.00  VYS57289.1
F-box and associated interaction domains-containing protein...    66.2   66.2  43%   8e-12 39.13  NP_187999.1
F-box/associated interaction domain protein [Arabidopsis...       65.5   65.5  44%   1e-11 36.67  NP_188911.2
unnamed protein product [Arabidopsis thaliana]                    64.7   64.7  43%   2e-11 38.26  VYS58278.1
hypothetical protein AXX17_AT3G24470 [Arabidopsis thaliana]       64.3   64.3  44%   4e-11 35.83  OAP04928.1
unnamed protein product [Arabidopsis thaliana]                    64.3   64.3  44%   4e-11 35.00  VYS58276.1
unnamed protein product [Arabidopsis thaliana]                    64.3   64.3  44%   4e-11 35.00  CAA0383352.1
unnamed protein product [Arabidopsis thaliana]                    63.9   63.9  45%   5e-11 34.48  CAA0382340.1
F-box and associated interaction domains-containing protein...    63.9   63.9  45%   5e-11 34.48  NP_187998.1
unknown [Arabidopsis thaliana]                                    63.9   63.9  45%   5e-11 34.48  ABK28556.1
F-box associated ubiquitination effector family protein...        63.5   63.5  44%   6e-11 35.34  NP_172646.3
hypothetical protein AXX17_AT3G14270 [Arabidopsis thaliana]       62.4   62.4  45%   2e-10 33.62  OAP02955.1
hypothetical protein AXX17_AT3G18360 [Arabidopsis thaliana]       59.7   59.7  44%   2e-10 31.15  OAP01833.1
unnamed protein product [Arabidopsis thaliana]                    62.0   62.0  44%   2e-10 34.48  VYS45785.1
F-box and associated interaction domains-containing protein...    61.6   61.6  47%   3e-10 33.87  NP_566558.1
hypothetical protein AXX17_AT3G17610 [Arabidopsis thaliana]       60.8   60.8  47%   5e-10 33.87  OAP04275.1
F-box and associated interaction domains-containing protein...    58.2   58.2  23%   6e-10 45.00  NP_850216.1
unnamed protein product [Arabidopsis thaliana]                    56.2   56.2  27%   6e-10 40.00  VYS54341.1
F-box and associated interaction domains-containing protein...    59.7   59.7  42%   1e-09 38.05  NP_188912.1
unnamed protein product [Arabidopsis thaliana]                    59.7   59.7  42%   1e-09 38.05  CAA0383354.1
unnamed protein product [Arabidopsis thaliana]                    57.4   57.4  44%   1e-09 31.15  CAA0382744.1
F-box and associated interaction domains-containing protein...    58.2   58.2  23%   2e-09 50.82  NP_683479.1
unnamed protein product [Arabidopsis thaliana]                    58.9   58.9  46%   2e-09 33.33  CAA0383369.1
unnamed protein product [Arabidopsis thaliana]                    54.7   54.7  21%   3e-09 45.45  CAA0374401.1
unnamed protein product [Arabidopsis thaliana]                    58.2   58.2  47%   4e-09 33.59  CAA0364593.1
hypothetical protein [Arabidopsis thaliana]                       58.2   58.2  47%   4e-09 33.59  AAO37178.1
F-box and associated interaction domains-containing protein...    58.2   58.2  47%   4e-09 33.59  NP_179373.2
hypothetical protein AXX17_AT2G13160 [Arabidopsis thaliana]       58.2   58.2  47%   4e-09 33.59  OAP08881.1


Alignments:
>F-box and associated interaction domains-containing protein [Arabidopsis thaliana]
Sequence ID: NP_200864.1 Length: 374
>F-box and associated interaction domains-containing protein [Arabidopsis thaliana]
Sequence ID: AED97345.1 Length: 374
Range 1: 1 to 120

Score:112 bits(280), Expect:3e-28,
Method:Compositional matrix adjust.,
Identities:64/121(53%), Positives:72/121(59%), Gaps:1/121(0%)

Query  94   MTTISDISNYLVEKILSRAPITCLGAVRTTCKRWNALSKENTLCNGEARHQFLGFMTKKY  273
            MT +SD+S  LVE+IL R  IT LGAVR+TCK W  LSK   LC  E +HQFLGFM K Y
Sbjct  1    MTMMSDLSEDLVEEILCRVSITSLGAVRSTCKGWYVLSKTRVLCKAETKHQFLGFMKKNY  60

Query  274  KLCSMRFRSQWKI*rrrr*rfrGSNYKGVR*FSESC*DI*CVFQCDGLLLCVTNEDKTRL  453
            KLCSMRF             F   + K      +   DI  VFQCDGLLLCVT E+ TRL
Sbjct  61   KLCSMRFDLHGNFNEEGGEEFMNPSIKKSGNLLDQL-DICKVFQCDGLLLCVTKEENTRL  119

Query  454  A  456

Sbjct  120  V  120


>F-box and associated interaction domains-containing protein [Arabidopsis thaliana]
Sequence ID: NP_001330054.1 Length: 407
>RecName: Full=Putative F-box protein At5g60560 [Arabidopsis thaliana]
Sequence ID: Q9FKJ1.1 Length: 407 >F-box and associated interaction domains-containing protein [Arabidopsis thaliana]
Sequence ID: ANM68290.1 Length: 407 >unnamed protein product [Arabidopsis thaliana]
Sequence ID: BAB08239.1 Length: 407 >unnamed protein product [Arabidopsis thaliana]
Sequence ID: VYS70982.1 Length: 407
Range 1: 1 to 120

Score:112 bits(279), Expect:6e-28,
Method:Compositional matrix adjust.,
Identities:64/121(53%), Positives:72/121(59%), Gaps:1/121(0%)

Query  94   MTTISDISNYLVEKILSRAPITCLGAVRTTCKRWNALSKENTLCNGEARHQFLGFMTKKY  273
            MT +SD+S  LVE+IL R  IT LGAVR+TCK W  LSK   LC  E +HQFLGFM K Y
Sbjct  1    MTMMSDLSEDLVEEILCRVSITSLGAVRSTCKGWYVLSKTRVLCKAETKHQFLGFMKKNY  60

Query  274  KLCSMRFRSQWKI*rrrr*rfrGSNYKGVR*FSESC*DI*CVFQCDGLLLCVTNEDKTRL  453
            KLCSMRF             F   + K      +   DI  VFQCDGLLLCVT E+ TRL
Sbjct  61   KLCSMRFDLHGNFNEEGGEEFMNPSIKKSGNLLDQL-DICKVFQCDGLLLCVTKEENTRL  119

Query  454  A  456

Sbjct  120  V  120


>hypothetical protein AXX17_AT5G60010 [Arabidopsis thaliana]
Sequence ID: OAO92930.1 Length: 374
Range 1: 1 to 120

Score:111 bits(277), Expect:8e-28,
Method:Compositional matrix adjust.,
Identities:63/121(52%), Positives:72/121(59%), Gaps:1/121(0%)

Query  94   MTTISDISNYLVEKILSRAPITCLGAVRTTCKRWNALSKENTLCNGEARHQFLGFMTKKY  273
            MT +SD+S  LVE+IL R  IT LGAVR+TCK W  LSK   LC  E +HQFLGFM K Y
Sbjct  1    MTMMSDLSEDLVEEILCRVSITSLGAVRSTCKGWYVLSKTRVLCKAETKHQFLGFMKKNY  60

Query  274  KLCSMRFRSQWKI*rrrr*rfrGSNYKGVR*FSESC*DI*CVFQCDGLLLCVTNEDKTRL  453
            KLCSMRF             F   + K      +   DI  VFQCDGLLLCVT E+ +RL
Sbjct  61   KLCSMRFDLHGNFNEEGGEEFMNPSIKKSGNLLDQL-DICKVFQCDGLLLCVTKEENSRL  119

Query  454  A  456

Sbjct  120  V  120


>unnamed protein product [Arabidopsis thaliana]
Sequence ID: CAA0411101.1 Length: 407
Range 1: 1 to 120

Score:110 bits(275), Expect:2e-27,
Method:Compositional matrix adjust.,
Identities:64/121(53%), Positives:72/121(59%), Gaps:1/121(0%)

Query  94   MTTISDISNYLVEKILSRAPITCLGAVRTTCKRWNALSKENTLCNGEARHQFLGFMTKKY  273
            MT +SD+S  LVE+IL R  IT LGAVR+TCK W  LSK   LC  E +HQFLGFM K Y
Sbjct  1    MTMMSDLSEDLVEEILCRVSITSLGAVRSTCKGWYVLSKTRVLCKAETKHQFLGFMKKNY  60

Query  274  KLCSMRFRSQWKI*rrrr*rfrGSNYKGVR*FSESC*DI*CVFQCDGLLLCVTNEDKTRL  453
            KLCSMRF             F   + K      +   DI  VFQCDGLLLCVT E+ TRL
Sbjct  61   KLCSMRFDLHGNCNEEGGEEFMNPSIKKSGNLLDQL-DICKVFQCDGLLLCVTKEENTRL  119

Query  454  A  456

Sbjct  120  V  120


>hypothetical protein AXX17_AT3G21160 [Arabidopsis thaliana]
Sequence ID: OAP04329.1 Length: 389
Range 1: 1 to 109

Score:108 bits(269), Expect:1e-26,
Method:Compositional matrix adjust.,
Identities:60/121(50%), Positives:73/121(60%), Gaps:12/121(9%)

Query  94   MTTISDISNYLVEKILSRAPITCLGAVRTTCKRWNALSKENTLCNGEARHQFLGFMTKKY  273
            MT +SD++  LVE+ILSR PIT LGAVR+TCK WNALSKE  LC GE + QFLGFM   Y
Sbjct  1    MTMMSDLTQDLVEEILSRVPITSLGAVRSTCKGWNALSKERILCIGEPKQQFLGFMMLDY  60

Query  274  KLCSMRFRSQWKI*rrrr*rfrGSNYKGVR*FSESC*DI*CVFQCDGLLLCVTNEDKTRL  453
            +LCSMRF                 ++  +  +     +I  VF C GLLLCVT E  +RL
Sbjct  61   RLCSMRFNLH---------GILNEDFVSISMYQV---EISQVFYCAGLLLCVTREKSSRL  108

Query  454  A  456

Sbjct  109  V  109


>F-box and associated interaction domains-containing protein [Arabidopsis thaliana]
Sequence ID: NP_188622.1 Length: 389
>RecName: Full=F-box protein At3g19880 [Arabidopsis thaliana]
Sequence ID: Q9LT21.1 Length: 389 >F-box family protein [Arabidopsis thaliana]
Sequence ID: AAY78745.1 Length: 389 >F-box and associated interaction domains-containing protein [Arabidopsis thaliana]
Sequence ID: AEE76303.1 Length: 389 >unnamed protein product [Arabidopsis thaliana]
Sequence ID: BAB01302.1 Length: 389
Range 1: 1 to 108

Score:106 bits(265), Expect:5e-26,
Method:Compositional matrix adjust.,
Identities:59/120(49%), Positives:71/120(59%), Gaps:12/120(10%)

Query  94   MTTISDISNYLVEKILSRAPITCLGAVRTTCKRWNALSKENTLCNGEARHQFLGFMTKKY  273
            MT +SD++  LVE+ILSR PIT LGAVR+TCK WNALSKE  LC GE + QFLGFM   Y
Sbjct  1    MTMMSDLTQDLVEEILSRVPITSLGAVRSTCKGWNALSKERILCIGEPKQQFLGFMMLDY  60

Query  274  KLCSMRFRSQWKI*rrrr*rfrGSNYKGVR*FSESC*DI*CVFQCDGLLLCVTNEDKTRL  453
            +LCSMRF                 ++  +  +         VF C GLLLCVT E  +RL
Sbjct  61   RLCSMRFNLH---------GILNEDFVSISMYQVETSQ---VFYCAGLLLCVTREKSSRL  108


>unnamed protein product [Arabidopsis thaliana]
Sequence ID: VYS57980.1 Length: 402
Range 1: 1 to 120

Score:105 bits(262), Expect:1e-25,
Method:Compositional matrix adjust.,
Identities:61/121(50%), Positives:71/121(58%), Gaps:1/121(0%)

Query  94   MTTISDISNYLVEKILSRAPITCLGAVRTTCKRWNALSKENTLCNGEARHQFLGFMTKKY  273
            MT +SD+S  L+E+ILSR P T LGAVR+TCKRWN LSK+  LC  E       F+ KKY
Sbjct  1    MTMMSDLSQDLLEEILSRVPRTSLGAVRSTCKRWNTLSKDRILCKAEETRDQFRFIMKKY  60

Query  274  KLCSMRFRSQWKI*rrrr*rfrGSNYKGVR*FSESC*DI*CVFQCDGLLLCVTNEDKTRL  453
            KLCSMRF     +       F   + K +  F      I  VFQCDGLLLCVT ED  RL
Sbjct  61   KLCSMRFDLNGTLNEDGGTEFVDPSIKELGHFFNQV-KISKVFQCDGLLLCVTKEDNIRL  119

Query  454  A  456

Sbjct  120  V  120


>F-box and associated interaction domains-containing protein [Arabidopsis thaliana]
Sequence ID: NP_188638.1 Length: 402
>RecName: Full=Putative F-box protein At3g20030 [Arabidopsis thaliana]
Sequence ID: Q9LJZ8.1 Length: 402 >F-box and associated interaction domains-containing protein [Arabidopsis thaliana]
Sequence ID: AEE76322.1 Length: 402 >hypothetical protein AXX17_AT3G21370 [Arabidopsis thaliana]
Sequence ID: OAP04097.1 Length: 402 >unnamed protein product [Arabidopsis thaliana]
Sequence ID: BAB01860.1 Length: 402
Range 1: 1 to 120

Score:103 bits(256), Expect:8e-25,
Method:Compositional matrix adjust.,
Identities:60/121(50%), Positives:70/121(57%), Gaps:1/121(0%)

Query  94   MTTISDISNYLVEKILSRAPITCLGAVRTTCKRWNALSKENTLCNGEARHQFLGFMTKKY  273
            MT +SD+S  L+E+ILSR P T LGAVR+TCKRWN L K+  LC  E       F+ KKY
Sbjct  1    MTMMSDLSQDLLEEILSRVPRTSLGAVRSTCKRWNTLFKDRILCKAEETRDQFRFIMKKY  60

Query  274  KLCSMRFRSQWKI*rrrr*rfrGSNYKGVR*FSESC*DI*CVFQCDGLLLCVTNEDKTRL  453
            KLCSMRF     +       F   + K +  F      I  VFQCDGLLLCVT ED  RL
Sbjct  61   KLCSMRFDLNGTLNEDGGTEFVDPSIKELGHFFNQV-KISKVFQCDGLLLCVTKEDNIRL  119

Query  454  A  456

Sbjct  120  V  120


>unnamed protein product [Arabidopsis thaliana]
Sequence ID: CAA0383045.1 Length: 402
Range 1: 1 to 120

Score:103 bits(256), Expect:9e-25,
Method:Compositional matrix adjust.,
Identities:60/121(50%), Positives:70/121(57%), Gaps:1/121(0%)

Query  94   MTTISDISNYLVEKILSRAPITCLGAVRTTCKRWNALSKENTLCNGEARHQFLGFMTKKY  273
            MT +SD+S  L+E+ILSR P T LGAVR+TCKRWN L K+  LC  E       F+ KKY
Sbjct  1    MTMMSDLSQDLLEEILSRVPRTSLGAVRSTCKRWNTLFKDRILCKAEETRDQFRFIMKKY  60

Query  274  KLCSMRFRSQWKI*rrrr*rfrGSNYKGVR*FSESC*DI*CVFQCDGLLLCVTNEDKTRL  453
            KLCSMRF     +       F   + K +  F      I  VFQCDGLLLCVT ED  RL
Sbjct  61   KLCSMRFDLNGTLNEDGGTEFVDPSIKELGHFFNQV-KISKVFQCDGLLLCVTKEDNIRL  119

Query  454  A  456

Sbjct  120  V  120


>unnamed protein product [Arabidopsis thaliana]
Sequence ID: VYS57963.1 Length: 312
Range 1: 1 to 67

Score:99.8 bits(247), Expect:5e-24,
Method:Compositional matrix adjust.,
Identities:46/67(69%), Positives:53/67(79%), Gaps:0/67(0%)

Query  94   MTTISDISNYLVEKILSRAPITCLGAVRTTCKRWNALSKENTLCNGEARHQFLGFMTKKY  273
            MT +SD++  LVE+ILSR PIT LGAVR+TCK WNALSKE  LC GE + QFLGFM   Y
Sbjct  1    MTMMSDLTQDLVEEILSRVPITSLGAVRSTCKGWNALSKERILCIGEPKQQFLGFMMLDY  60

Query  274  KLCSMRF  294
            +LCSMRF
Sbjct  61   RLCSMRF  67


>unnamed protein product [Arabidopsis thaliana]
Sequence ID: CAA0383027.1 Length: 367
Range 1: 1 to 67

Score:99.4 bits(246), Expect:1e-23,
Method:Compositional matrix adjust.,
Identities:46/67(69%), Positives:53/67(79%), Gaps:0/67(0%)

Query  94   MTTISDISNYLVEKILSRAPITCLGAVRTTCKRWNALSKENTLCNGEARHQFLGFMTKKY  273
            MT +SD++  LVE+ILSR PIT LGAVR+TCK WNALSKE  LC GE + QFLGFM   Y
Sbjct  1    MTMMSDLTQDLVEEILSRVPITSLGAVRSTCKGWNALSKERILCIGEPKQQFLGFMMLDY  60

Query  274  KLCSMRF  294
            +LCSMRF
Sbjct  61   RLCSMRF  67


>F-box family protein [Arabidopsis thaliana]
Sequence ID: NP_188623.1 Length: 410
>RecName: Full=F-box protein At3g19890 [Arabidopsis thaliana]
Sequence ID: Q4PSN8.1 Length: 410 >F-box family protein [Arabidopsis thaliana]
Sequence ID: AAY78746.1 Length: 410 >F-box family protein [Arabidopsis thaliana]
Sequence ID: AEE76304.1 Length: 410 >hypothetical protein AXX17_AT3G21170 [Arabidopsis thaliana]
Sequence ID: OAP04868.1 Length: 410 >unnamed protein product [Arabidopsis thaliana]
Sequence ID: VYS57964.1 Length: 410
Range 1: 1 to 141

Score:95.9 bits(237), Expect:5e-22,
Method:Compositional matrix adjust.,
Identities:59/145(41%), Positives:85/145(58%), Gaps:8/145(5%)

Query  94   MTTISDISNYLVEKILSRAPITCLGAVRTTCKRWNALSKENTLCNGEARHQFLGFMTKKY  273
            MT ISD+S  LVE+ILS+APIT LGAVR+T K+WNALSK   L   EA+ QFLGFM   +
Sbjct  1    MTMISDLSKDLVEEILSKAPITSLGAVRSTHKQWNALSKGRLLYKAEAKDQFLGFMVMDH  60

Query  274  KLCSMRFRSQWKI*rrrr*rfrGSNYKGVR*FSE--SC*DI*CVFQCDGLLLCVTNEDKT  447
            +  SM F     +    +    G +   +R   +  +  DI  VFQCDGL+LCV +++ +
Sbjct  61   RFLSMIFH----LNGILKGDGEGFDRPSIREVGDIVNQIDISKVFQCDGLVLCVPSDNSS  116

Query  448  RLASSWY--GTHKLDILNPHKIAEI  516
             +  + Y   T  ++   PH  +++
Sbjct  117  VVVWNPYLGQTKWIEAREPHDESDM  141


>unnamed protein product [Arabidopsis thaliana]
Sequence ID: BAB01303.1 Length: 499
Range 1: 1 to 141

Score:95.9 bits(237), Expect:8e-22,
Method:Compositional matrix adjust.,
Identities:59/145(41%), Positives:85/145(58%), Gaps:8/145(5%)

Query  94   MTTISDISNYLVEKILSRAPITCLGAVRTTCKRWNALSKENTLCNGEARHQFLGFMTKKY  273
            MT ISD+S  LVE+ILS+APIT LGAVR+T K+WNALSK   L   EA+ QFLGFM   +
Sbjct  1    MTMISDLSKDLVEEILSKAPITSLGAVRSTHKQWNALSKGRLLYKAEAKDQFLGFMVMDH  60

Query  274  KLCSMRFRSQWKI*rrrr*rfrGSNYKGVR*FSE--SC*DI*CVFQCDGLLLCVTNEDKT  447
            +  SM F     +    +    G +   +R   +  +  DI  VFQCDGL+LCV +++ +
Sbjct  61   RFLSMIFH----LNGILKGDGEGFDRPSIREVGDIVNQIDISKVFQCDGLVLCVPSDNSS  116

Query  448  RLASSWY--GTHKLDILNPHKIAEI  516
             +  + Y   T  ++   PH  +++
Sbjct  117  VVVWNPYLGQTKWIEAREPHDESDM  141


>unnamed protein product [Arabidopsis thaliana]
Sequence ID: CAA0261612.1 Length: 197
Range 1: 1 to 120

Score:89.4 bits(220), Expect:5e-21,
Method:Compositional matrix adjust.,
Identities:54/121(45%), Positives:68/121(56%), Gaps:1/121(0%)

Query  94   MTTISDISNYLVEKILSRAPITCLGAVRTTCKRWNALSKENTLCNGEARHQFLGFMTKKY  273
            MT +SD+S  LVE+IL R PIT L AVR++CK WN LSK   LC  EAR+QFLGF    +
Sbjct  1    MTMMSDLSLDLVEEILCRVPITSLKAVRSSCKLWNVLSKNRILCKTEARNQFLGFTIMNH  60

Query  274  KLCSMRFRSQWKI*rrrr*rfrGSNYKGVR*FSESC*DI*CVFQCDGLLLCVTNEDKTRL  453
            +L SMRF             F   + K +        +I  VF C+GLLLCVT    ++L
Sbjct  61   RLYSMRFNLHGIGLDEDSEEFIDPSIKPIGNLLNQV-EISKVFSCEGLLLCVTRNHSSKL  119

Query  454  A  456

Sbjct  120  V  120


>F-box family protein [Arabidopsis thaliana]
Sequence ID: NP_174493.1 Length: 591
>RecName: Full=Putative F-box protein At1g32140 [Arabidopsis thaliana]
Sequence ID: Q9FVQ9.1 Length: 591 >hypothetical protein [Arabidopsis thaliana]
Sequence ID: AAG23440.1 Length: 591 >F-box family protein [Arabidopsis thaliana]
Sequence ID: AEE31440.1 Length: 591
Range 1: 1 to 130

Score:90.1 bits(222), Expect:9e-20,
Method:Compositional matrix adjust.,
Identities:59/142(42%), Positives:76/142(53%), Gaps:12/142(8%)

Query  94   MTTISDISNYLVEKILSRAPITCLGAVRTTCKRWNALSKENTLCNGEARHQFLGFMTKKY  273
            MT +SD+S  LVE+IL R PIT L AVR++CK WN LSK   LC  EAR+QFLGF    +
Sbjct  1    MTMMSDLSLDLVEEILCRVPITSLKAVRSSCKLWNVLSKNRILCKTEARNQFLGFTIMNH  60

Query  274  KLCSMRFRSQWKI*rrrr*rfrGSNYKGVR*FSESC*DI*CVFQCDGLLLCVTNEDKTRL  453
            +L SMRF             F   + K +        +I  VF C+GLLLCVT    ++L
Sbjct  61   RLYSMRFNLHGIGLNENSEEFIDPSIKPIGNLLNQV-EISKVFYCEGLLLCVTRNHSSKL  119

Query  454  ASSWYGTHKLDILNPHKIAEIR  519
                       + NP+ + EIR
Sbjct  120  V----------VWNPY-LGEIR  130


>F-box family protein [Arabidopsis thaliana]
Sequence ID: ABE65746.1 Length: 160
Range 1: 1 to 105

Score:83.2 bits(204), Expect:5e-19,
Method:Compositional matrix adjust.,
Identities:56/121(46%), Positives:66/121(54%), Gaps:25/121(20%)

Query  94   MTTISDISNYLVEKILSRAPITCLGAVRTTCKRWNALSKENTLCNGEARHQFLGFMTKKY  273
            M TISD+   LVE+ILSR P+T L AVR+TCK WNALSK  T   G+ R QFLGFM   +
Sbjct  1    MRTISDLPVALVEEILSRVPLTSLSAVRSTCKTWNALSK--TQIFGKTRQQFLGFMMIDF  58

Query  274  KLCSMRFRSQWKI*rrrr*rfrGSNYKGVR*FSESC---------*DI*CVFQCDGLLLC  426
             L S++F  Q            G NY+    F E            DI  VF C+GLLLC
Sbjct  59   GLYSIKFDLQ------------GLNYESD--FVEPSIKRVSILDQLDIFKVFHCEGLLLC  104

Query  427  V  429
            V
Sbjct  105  V  105


>unnamed protein product [Arabidopsis thaliana]
Sequence ID: VYS47772.1 Length: 515
Range 1: 1 to 120

Score:87.4 bits(215), Expect:6e-19,
Method:Compositional matrix adjust.,
Identities:54/121(45%), Positives:67/121(55%), Gaps:1/121(0%)

Query  94   MTTISDISNYLVEKILSRAPITCLGAVRTTCKRWNALSKENTLCNGEARHQFLGFMTKKY  273
            MT +SD+S  LVE+IL R PIT L AVR++CK WN LSK   LC  EAR+QFLGF
Sbjct  1    MTMMSDLSLDLVEEILCRVPITSLKAVRSSCKLWNVLSKNRILCKTEARNQFLGFTIMNP  60

Query  274  KLCSMRFRSQWKI*rrrr*rfrGSNYKGVR*FSESC*DI*CVFQCDGLLLCVTNEDKTRL  453
            +L SMRF             F   + K +        +I  VF C+GLLLCVT    ++L
Sbjct  61   RLYSMRFNLHGIGLDEDSEEFIDPSIKPIGNLLNQV-EISKVFSCEGLLLCVTRNHSSKL  119

Query  454  A  456

Sbjct  120  V  120


>RecName: Full=Putative F-box protein At3g13624 [Arabidopsis thaliana]
Sequence ID: Q9LHN6.1 Length: 366
>unnamed protein product [Arabidopsis thaliana]
Sequence ID: BAB02605.1 Length: 366
Range 1: 1 to 120

Score:84.7 bits(208), Expect:2e-18,
Method:Compositional matrix adjust.,
Identities:49/125(39%), Positives:68/125(54%), Gaps:5/125(4%)

Query  94   MTTISDISNYLVEKILSRAPITCLGAVRTTCKRWNALSKENTLCNGEARHQFLGFMTKKY  273
            MTTISD+   +VE+IL R P+T L AVR+ CK WN LSK   LC    + QFLGF+   Y
Sbjct  1    MTTISDLPEDVVEEILPRVPLTSLSAVRSICKTWNTLSKNRVLCKAAVKKQFLGFIMMDY  60

Query  274  KLCSMRFRSQWKI*rrrr*rfrGSNYKGVR*FSESC*DI*CVFQCDGLLLCVTNEDKTRL  453
            ++CSM+F  +              + K V    +   +I  V  CDGLLL VT ++ + +
Sbjct  61   RVCSMKFHLR---NDEGGDLVDDLSIKQVGILDQI--EISEVLHCDGLLLFVTKDNSSLV  115

Query  454  ASSWY  468
              + Y
Sbjct  116  VWNPY  120


>RecName: Full=F-box protein At1g59680 [Arabidopsis thaliana]
Sequence ID: Q9LQ46.2 Length: 364
Range 1: 1 to 108

Score:84.3 bits(207), Expect:4e-18,
Method:Compositional matrix adjust.,
Identities:55/123(45%), Positives:62/123(50%), Gaps:17/123(13%)

Query  94   MTTISDISNYLVEKILSRAPITCLGAVRTTCKRWNALSKENTLCNGE--ARHQFLGFMTK  267
            MTT+SD+S  LV +ILSR P+T L AVR TCK WN LSK       E  A  QFLGF
Sbjct  1    MTTMSDLSVDLVGEILSRVPLTSLSAVRCTCKSWNTLSKHQIFGKAELAATKQFLGFTVM  60

Query  268  KYKLCSMRFRSQWKI*rrrr*rfrGSNYKGVR*FSESC*DI*CVFQCDGLLLCVTNEDKT  447
             YK+CS+RF  Q            G  Y               VF  DGLLLCVT +D
Sbjct  61   DYKVCSLRFDLQGIRNDGDDFVDHGEVYD--------------VFHSDGLLLCVT-KDHW  105

Query  448  RLA  456
            RL
Sbjct  106  RLV  108


>unnamed protein product [Arabidopsis thaliana]
Sequence ID: CAA0383028.1 Length: 381
Range 1: 1 to 68

Score:84.0 bits(206), Expect:5e-18,
Method:Compositional matrix adjust.,
Identities:41/68(60%), Positives:49/68(72%), Gaps:0/68(0%)

Query  94   MTTISDISNYLVEKILSRAPITCLGAVRTTCKRWNALSKENTLCNGEARHQFLGFMTKKY  273
            MT ISD+S  LVE+ILS+APIT LGAVR+T K+WNALSK   L   EA+ QFLGFM   +
Sbjct  1    MTMISDLSKDLVEEILSKAPITSLGAVRSTHKQWNALSKGRLLYKAEAKDQFLGFMVMDH  60

Query  274  KLCSMRFR  297
            +  SM F
Sbjct  61   RFLSMTFH  68


>hypothetical protein AXX17_AT3G13970 [Arabidopsis thaliana]
Sequence ID: OAP04380.1 Length: 542
Range 1: 1 to 120

Score:84.3 bits(207), Expect:7e-18,
Method:Compositional matrix adjust.,
Identities:49/125(39%), Positives:68/125(54%), Gaps:5/125(4%)

Query  94   MTTISDISNYLVEKILSRAPITCLGAVRTTCKRWNALSKENTLCNGEARHQFLGFMTKKY  273
            MTTISD+   +VE+IL R P+T L AVR+ CK WN LSK   LC    + QFLGF+   Y
Sbjct  1    MTTISDLPEDVVEEILPRVPLTSLSAVRSICKTWNTLSKNRVLCKAAVKKQFLGFIMMDY  60

Query  274  KLCSMRFRSQWKI*rrrr*rfrGSNYKGVR*FSESC*DI*CVFQCDGLLLCVTNEDKTRL  453
            ++CSM+F  +              + K V    +   +I  V  CDGLLL VT ++ + +
Sbjct  61   RVCSMKFHLR---NDEGGDLVDDLSIKQVGILDQI--EISEVLHCDGLLLFVTKDNSSLV  115

Query  454  ASSWY  468
              + Y
Sbjct  116  VWNPY  120


>unnamed protein product [Arabidopsis thaliana]
Sequence ID: VYS50192.1 Length: 353
Range 1: 1 to 105

Score:83.6 bits(205), Expect:7e-18,
Method:Compositional matrix adjust.,
Identities:56/121(46%), Positives:66/121(54%), Gaps:25/121(20%)

Query  94   MTTISDISNYLVEKILSRAPITCLGAVRTTCKRWNALSKENTLCNGEARHQFLGFMTKKY  273
            M TISD+   LVE+ILSR P+T L AVR+TCK WNALSK  T   G+ R QFLGFM   +
Sbjct  1    MRTISDLPVALVEEILSRVPLTSLSAVRSTCKTWNALSK--TQIFGKTRQQFLGFMMIDF  58

Query  274  KLCSMRFRSQWKI*rrrr*rfrGSNYKGVR*FSESC---------*DI*CVFQCDGLLLC  426
             L S++F  Q            G NY+    F E            DI  VF C+GLLLC
Sbjct  59   GLYSIKFDLQ------------GLNYESD--FVEPSIKRVSILDQLDIFKVFHCEGLLLC  104

Query  427  V  429
            V
Sbjct  105  V  105


>F-box and associated interaction domains-containing protein [Arabidopsis thaliana]
Sequence ID: NP_176822.1 Length: 360
>hypothetical protein [Arabidopsis thaliana]
Sequence ID: AAG51152.1 Length: 360 >F-box and associated interaction domains-containing protein [Arabidopsis thaliana]
Sequence ID: AEE34515.1 Length: 360
Range 1: 1 to 105

Score:83.6 bits(205), Expect:7e-18,
Method:Compositional matrix adjust.,
Identities:56/121(46%), Positives:66/121(54%), Gaps:25/121(20%)

Query  94   MTTISDISNYLVEKILSRAPITCLGAVRTTCKRWNALSKENTLCNGEARHQFLGFMTKKY  273
            M TISD+   LVE+ILSR P+T L AVR+TCK WNALSK  T   G+ R QFLGFM   +
Sbjct  1    MRTISDLPVALVEEILSRVPLTSLSAVRSTCKTWNALSK--TQIFGKTRQQFLGFMMIDF  58

Query  274  KLCSMRFRSQWKI*rrrr*rfrGSNYKGVR*FSESC---------*DI*CVFQCDGLLLC  426
             L S++F  Q            G NY+    F E            DI  VF C+GLLLC
Sbjct  59   GLYSIKFDLQ------------GLNYESD--FVEPSIKRVSILDQLDIFKVFHCEGLLLC  104

Query  427  V  429
            V
Sbjct  105  V  105


>RecName: Full=F-box protein At1g66490 [Arabidopsis thaliana]
Sequence ID: Q1PFG1.2 Length: 370
Range 1: 1 to 105

Score:83.6 bits(205), Expect:8e-18,
Method:Compositional matrix adjust.,
Identities:56/121(46%), Positives:66/121(54%), Gaps:25/121(20%)

Query  94   MTTISDISNYLVEKILSRAPITCLGAVRTTCKRWNALSKENTLCNGEARHQFLGFMTKKY  273
            M TISD+   LVE+ILSR P+T L AVR+TCK WNALSK  T   G+ R QFLGFM   +
Sbjct  1    MRTISDLPVALVEEILSRVPLTSLSAVRSTCKTWNALSK--TQIFGKTRQQFLGFMMIDF  58

Query  274  KLCSMRFRSQWKI*rrrr*rfrGSNYKGVR*FSESC---------*DI*CVFQCDGLLLC  426
             L S++F  Q            G NY+    F E            DI  VF C+GLLLC
Sbjct  59   GLYSIKFDLQ------------GLNYESD--FVEPSIKRVSILDQLDIFKVFHCEGLLLC  104

Query  427  V  429
            V
Sbjct  105  V  105


>unnamed protein product [Arabidopsis thaliana]
Sequence ID: CAA0321408.1 Length: 398
Range 1: 5 to 120

Score:83.6 bits(205), Expect:8e-18,
Method:Compositional matrix adjust.,
Identities:52/120(43%), Positives:70/120(58%), Gaps:6/120(5%)

Query  103  ISDISNYLVEKILSRAPITCLGAVRTTCKRWNALSKENTLCNGEARHQFLGF--MTKKYK  276
            +SD+ N LVE+ILSR PIT LGAVR+TCKRWN LSK+  +C G+A  QF GF     +
Sbjct  5    MSDLPNDLVEEILSRVPITSLGAVRSTCKRWNGLSKDRIVCKGDANQQFTGFTRYVNEST  64

Query  277  LCSMRFRSQWKI*rrrr*rfrGSNYKGVR*FSESC*DI*CVFQCDGLLLCVTNEDKTRLA  456
            +CSMR     +I          S  K +  F E   ++  V+  DGLLL VT+E  +++
Sbjct  65   VCSMRLNLN-EIQNEDVELVELSINK-INKFIEL--ELFQVYYSDGLLLLVTDEVDSKIV  120


>F-box and associated interaction domains-containing protein [Arabidopsis thaliana]
Sequence ID: NP_176912.1 Length: 398
>RecName: Full=Putative F-box protein At1g67450 [Arabidopsis thaliana]
Sequence ID: O64801.1 Length: 398 >T1F15.9 [Arabidopsis thaliana]
Sequence ID: AAC18789.1 Length: 398 >F-box and associated interaction domains-containing protein [Arabidopsis thaliana]
Sequence ID: AEE34647.1 Length: 398
Range 1: 5 to 120

Score:83.6 bits(205), Expect:9e-18,
Method:Compositional matrix adjust.,
Identities:52/120(43%), Positives:70/120(58%), Gaps:6/120(5%)

Query  103  ISDISNYLVEKILSRAPITCLGAVRTTCKRWNALSKENTLCNGEARHQFLGF--MTKKYK  276
            +SD+ N LVE+ILSR PIT LGAVR+TCKRWN LSK+  +C G+A  QF GF     +
Sbjct  5    MSDLPNDLVEEILSRVPITSLGAVRSTCKRWNGLSKDRIVCKGDANQQFTGFTRYVNEST  64

Query  277  LCSMRFRSQWKI*rrrr*rfrGSNYKGVR*FSESC*DI*CVFQCDGLLLCVTNEDKTRLA  456
            +CSMR     +I          S  K +  F E   ++  V+  DGLLL VT+E  +++
Sbjct  65   VCSMRLDLN-EIQNEDVELVELSINK-INKFIEL--ELFQVYYSDGLLLLVTDEVDSKIV  120


>F-box and associated interaction domains-containing protein [Arabidopsis thaliana]
Sequence ID: NP_567289.1 Length: 391
>RecName: Full=F-box/kelch-repeat protein At4g05080 [Arabidopsis thaliana]
Sequence ID: Q9S9T0.1 Length: 391 >similar to a family of Arabidopsis thaliana predicted proteins; see GB:AC003952 [Arabidopsis thaliana]
Sequence ID: AAD48970.1 Length: 391 >F-box family protein [Arabidopsis thaliana]
Sequence ID: AAY78792.1 Length: 391 >F-box and associated interaction domains-containing protein [Arabidopsis thaliana]
Sequence ID: AEE82475.1 Length: 391 >unnamed protein product [Arabidopsis thaliana]
Sequence ID: CAA0393622.1 Length: 391 >AT4g05080 [Arabidopsis thaliana]
Sequence ID: CAB81050.1 Length: 391
Range 1: 1 to 125

Score:82.0 bits(201), Expect:3e-17,
Method:Compositional matrix adjust.,
Identities:51/127(40%), Positives:71/127(55%), Gaps:3/127(2%)

Query  94   MTTISDISNYLVEKILSRAPITCLGAVRTTCKRWNALSKENTLCNGEARHQF-LGFMTKK  270
            MT + D++  LV++ILSR PIT LGAVR+TCK WNALSK+  LC  + + QF  GFM
Sbjct  1    MTMMFDLTQDLVKEILSRVPITSLGAVRSTCKGWNALSKDRILCKAKPKQQFHQGFMLSD  60

Query  271  YKLCSMRFRSQWKI*rrrr*rfrGSNYKGVR*FSESC*DI*CVFQCDGLLLCVTNEDKTR  450
            Y+L SMRF                S  K +  F     +I  ++ C G+LLCVT + +
Sbjct  61   YRLRSMRFNISGTFKENGEEFVNLS-VKEIGNFLNKV-EISHMYYCGGILLCVTTDTRLV  118

Query  451  LASSWYG  471
            + + + G
Sbjct  119  IWNPYLG  125


>unknown protein; 98896-95855 [Arabidopsis thaliana]
Sequence ID: AAB65488.1 Length: 592
Range 1: 228 to 343

Score:81.6 bits(200), Expect:5e-17,
Method:Compositional matrix adjust.,
Identities:49/126(39%), Positives:70/126(55%), Gaps:10/126(7%)

Query  76   FFLFGTMTTISDISNYLVEKILSRAPITCLGAVRTTCKRWNALSKENTLCNGEARHQFLG  255
            F L   MTT+SD+   +V +IL R P+TCL  VR+ CK+WNALSK+     G+A+ QFLG
Sbjct  228  FALSSKMTTMSDLDEIMVAEILCRTPMTCLKTVRSVCKKWNALSKK-WFFFGKAK-QFLG  285

Query  256  FMTKKYKLCSMRFRSQWKI*rrrr*rfrGSNYKGVR*FSESC*DI*CVFQCDGLLLCVTN  435
            FM    ++CS+RF  +  +        + S    +        ++  +F  DGLLLC+
Sbjct  286  FMMMDSRVCSLRFDLRKDLVVEPPSIKQVSILDQI--------EVSKIFHSDGLLLCIIK  337

Query  436  EDKTRL  453
             D TRL
Sbjct  338  NDTTRL  343


>unnamed protein product [Arabidopsis thaliana]
Sequence ID: VYS50284.1 Length: 398
Range 1: 5 to 120

Score:81.3 bits(199), Expect:5e-17,
Method:Compositional matrix adjust.,
Identities:52/120(43%), Positives:69/120(57%), Gaps:6/120(5%)

Query  103  ISDISNYLVEKILSRAPITCLGAVRTTCKRWNALSKENTLCNGEARHQFLGF--MTKKYK  276
            +SD+ N LVE+ILSR PIT LGAVR+TCKRWN LSK+  +C G A  QF GF     +
Sbjct  5    MSDLPNDLVEEILSRVPITSLGAVRSTCKRWNGLSKDRIVCKGGANQQFTGFTRYVNEST  64

Query  277  LCSMRFRSQWKI*rrrr*rfrGSNYKGVR*FSESC*DI*CVFQCDGLLLCVTNEDKTRLA  456
            +CSM+     +I          S  K +  F E   ++  V+  DGLLL VT+E  ++L
Sbjct  65   VCSMKLDLN-EIQNEDVELVELSINK-INKFIEL--ELFQVYYSDGLLLLVTDEVDSKLV  120


>unnamed protein product [Arabidopsis thaliana]
Sequence ID: CAA0319417.1 Length: 375
Range 1: 1 to 105

Score:80.9 bits(198), Expect:6e-17,
Method:Compositional matrix adjust.,
Identities:55/121(45%), Positives:66/121(54%), Gaps:25/121(20%)

Query  94   MTTISDISNYLVEKILSRAPITCLGAVRTTCKRWNALSKENTLCNGEARHQFLGFMTKKY  273
            M TISD+   LVE+ILSR P+T L AV++TCK WNALSK  T   G+ R QFLGFM   +
Sbjct  1    MRTISDLPVALVEQILSRVPLTSLIAVQSTCKTWNALSK--TQIFGKTRQQFLGFMMIDF  58

Query  274  KLCSMRFRSQWKI*rrrr*rfrGSNYKGVR*FSESC---------*DI*CVFQCDGLLLC  426
             L S++F  Q            G NY+    F E            DI  VF C+GLLLC
Sbjct  59   GLYSIKFDLQ------------GLNYESD--FVEPSIKRVSILDQLDIFKVFHCEGLLLC  104

Query  427  V  429
            V
Sbjct  105  V  105


>hypothetical protein AXX17_AT1G11130 [Arabidopsis thaliana]
Sequence ID: OAP16214.1 Length: 359
Range 1: 1 to 142

Score:80.5 bits(197), Expect:9e-17,
Method:Compositional matrix adjust.,
Identities:53/152(35%), Positives:78/152(51%), Gaps:19/152(12%)

Query  94   MTTISDISNYLVEKILSRAPITCLGAVRTTCKRWNALSKENTLCNGEARHQFLGFMTKKY  273
            MTT+SD+   +V +IL R P+TCL  VR+ CK+WNALSK+     G+A+ QFLGFM
Sbjct  1    MTTMSDLDEIMVAEILCRTPMTCLKTVRSVCKKWNALSKK-WFFFGKAK-QFLGFMMMDS  58

Query  274  KLCSMRFRSQWKI*rrrr*rfrGSNYKGVR*FSESC*DI*CVFQCDGLLLCVTNEDKTRL  453
            ++CS+RF  +  +        + S    +        ++  +F  DGLLLC+   D TRL
Sbjct  59   RVCSLRFDLRKDLVVEPPSIKQVSILDQI--------EVSKIFHSDGLLLCIIKNDTTRL  110

Query  454  --------ASSWYGT-HKLDILNPHKIAEIRN  522
                     + W    H   IL+ + I   +N
Sbjct  111  LVWNPYLEQTRWIQPRHNFHILDCYAIGHDKN  142


>RecName: Full=F-box protein At1g10895 [Arabidopsis thaliana]
Sequence ID: P0CB27.1 Length: 359
Range 1: 1 to 142

Score:80.5 bits(197), Expect:9e-17,
Method:Compositional matrix adjust.,
Identities:53/152(35%), Positives:78/152(51%), Gaps:19/152(12%)

Query  94   MTTISDISNYLVEKILSRAPITCLGAVRTTCKRWNALSKENTLCNGEARHQFLGFMTKKY  273
            MTT+SD+   +V +IL R P+TCL  VR+ CK+WNALSK+     G+A+ QFLGFM
Sbjct  1    MTTMSDLDEIMVAEILCRTPMTCLKTVRSVCKKWNALSKK-WFFFGKAK-QFLGFMMMDS  58

Query  274  KLCSMRFRSQWKI*rrrr*rfrGSNYKGVR*FSESC*DI*CVFQCDGLLLCVTNEDKTRL  453
            ++CS+RF  +  +        + S    +        ++  +F  DGLLLC+   D TRL
Sbjct  59   RVCSLRFDLRKDLVVEPPSIKQVSILDQI--------EVSKIFHSDGLLLCIIKNDTTRL  110

Query  454  --------ASSWYGT-HKLDILNPHKIAEIRN  522
                     + W    H   IL+ + I   +N
Sbjct  111  LVWNPYLEQTRWIQPRHNFHILDCYAIGHDKN  142


>unnamed protein product [Arabidopsis thaliana]
Sequence ID: VYS61789.1 Length: 391
Range 1: 1 to 125

Score:80.5 bits(197), Expect:9e-17,
Method:Compositional matrix adjust.,
Identities:51/127(40%), Positives:71/127(55%), Gaps:3/127(2%)

Query  94   MTTISDISNYLVEKILSRAPITCLGAVRTTCKRWNALSKENTLCNGEARHQF-LGFMTKK  270
            MT + D++  LV++ILSR PIT LGAVR+TCK WNALSK+  LC  + + QF  GFM
Sbjct  1    MTMMFDLTQDLVKEILSRVPITSLGAVRSTCKGWNALSKDRILCKLKPKQQFHQGFMLSD  60

Query  271  YKLCSMRFRSQWKI*rrrr*rfrGSNYKGVR*FSESC*DI*CVFQCDGLLLCVTNEDKTR  450
            Y+L SMRF                S  K +  F     +I  ++ C G+LLCVT + +
Sbjct  61   YRLRSMRFNISGTFKENGEEFVNLS-VKEIGNFLNKV-EISHMYYCGGILLCVTTDTRLV  118

Query  451  LASSWYG  471
            + + + G
Sbjct  119  IWNPYLG  125


>unnamed protein product [Arabidopsis thaliana]
Sequence ID: CAA0385286.1 Length: 387
Range 1: 1 to 117

Score:80.1 bits(196), Expect:1e-16,
Method:Compositional matrix adjust.,
Identities:52/118(44%), Positives:69/118(58%), Gaps:3/118(2%)

Query  94   MTTISDISNYLVEKILSRAPITCLGAVRTTCKRWNALSKENTLC-NGEARHQFLGFMTKK  270
            MTTISD+S+ LV  ILSR P T L +VR+TCK+WNALSK         AR+QFL FM
Sbjct  1    MTTISDLSDDLVGDILSRVPFTSLISVRSTCKKWNALSKNQIFGRKTAARNQFLEFMILD  60

Query  271  YKLCSMRFRSQWKI*rrrr*rfrGSNYKGVR*FS-ESC*DI*CVFQCDGLLLCVTNED  441
             ++CS+R   Q  I    +  F   + K +   S +   +I  V+ CDGLLLC+  E+
Sbjct  61   SRVCSLRLDLQ-GIRNEDKEDFVDPSMKLISIPSNDDQVEISQVYHCDGLLLCIAKEN  117


>unnamed protein product [Arabidopsis thaliana]
Sequence ID: CAA0395568.1 Length: 381
Range 1: 1 to 114

Score:80.1 bits(196), Expect:1e-16,
Method:Compositional matrix adjust.,
Identities:54/129(42%), Positives:72/129(55%), Gaps:24/129(18%)

Query  94   MTTISDISNYLVEKILSRAPITCLGAVRTTCKRWNALSKENTLCNGEARHQFLGFMTKKY  273
            MTT+SD+S  LV +IL+R P+T L +VR TCK WNALSKE       AR+QF+GF
Sbjct  1    MTTMSDLSPDLVGEILTRVPMTSLISVRCTCKMWNALSKEGIFFKA-ARNQFMGFTMMDS  59

Query  274  KLCSMRFRSQWKI*rrrr*rfrGSNYKGVR*FSESC---------*DI*CVFQCDGLLLC  426
            ++CSM+F  Q            G+N +    F + C          ++  V QCDGLLLC
Sbjct  60   RVCSMKFDLQ----------GIGNNEQD---FVDPCIKQIAKLDQIEVSKVLQCDGLLLC  106

Query  427  VTNEDKTRL  453
            V  +D +RL
Sbjct  107  V-GKDNSRL  114


>F-box family protein [Arabidopsis thaliana]
Sequence ID: NP_190520.2 Length: 388
>RecName: Full=F-box protein At3g49510 [Arabidopsis thaliana]
Sequence ID: Q9CA02.1 Length: 388 >unknown protein; 74683-75849 [Arabidopsis thaliana]
Sequence ID: AAG52186.1 Length: 388 >At3g49510 [Arabidopsis thaliana]
Sequence ID: AAQ22620.1 Length: 388 >F-box family protein [Arabidopsis thaliana]
Sequence ID: AEE78551.2 Length: 388 >hypothetical protein [Arabidopsis thaliana]
Sequence ID: BAE99818.1 Length: 388
Range 1: 1 to 117

Score:80.1 bits(196), Expect:1e-16,
Method:Compositional matrix adjust.,
Identities:52/118(44%), Positives:69/118(58%), Gaps:3/118(2%)

Query  94   MTTISDISNYLVEKILSRAPITCLGAVRTTCKRWNALSKENTLC-NGEARHQFLGFMTKK  270
            MTTISD+S+ LV  ILSR P T L +VR+TCK+WNALSK         AR+QFL FM
Sbjct  1    MTTISDLSDDLVGDILSRVPFTSLISVRSTCKKWNALSKNQIFGRKTAARNQFLEFMILD  60

Query  271  YKLCSMRFRSQWKI*rrrr*rfrGSNYKGVR*FS-ESC*DI*CVFQCDGLLLCVTNED  441
             ++CS+R   Q  I    +  F   + K +   S +   +I  V+ CDGLLLC+  E+
Sbjct  61   SRVCSLRLDLQ-GIRNEDKEDFVDPSMKLISIPSNDDQVEISQVYHCDGLLLCIAKEN  117


>unnamed protein product [Arabidopsis thaliana]
Sequence ID: VYS59865.1 Length: 388
Range 1: 1 to 117

Score:80.1 bits(196), Expect:2e-16,
Method:Compositional matrix adjust.,
Identities:52/118(44%), Positives:69/118(58%), Gaps:3/118(2%)

Query  94   MTTISDISNYLVEKILSRAPITCLGAVRTTCKRWNALSKENTLC-NGEARHQFLGFMTKK  270
            MTTISD+S+ LV  ILSR P T L +VR+TCK+WNALSK         AR+QFL FM
Sbjct  1    MTTISDLSDDLVGDILSRVPFTSLISVRSTCKKWNALSKNQIFGRKTAARNQFLEFMILD  60

Query  271  YKLCSMRFRSQWKI*rrrr*rfrGSNYKGVR*FS-ESC*DI*CVFQCDGLLLCVTNED  441
             ++CS+R   Q  I    +  F   + K +   S +   +I  V+ CDGLLLC+  E+
Sbjct  61   SRVCSLRLDLQ-GIRNEDKEDFVDPSMKLISIPSNDDQVEISQVYHCDGLLLCIAKEN  117


>putative protein [Arabidopsis thaliana]
Sequence ID: CAB62455.1 Length: 662
Range 1: 1 to 117

Score:79.7 bits(195), Expect:3e-16,
Method:Compositional matrix adjust.,
Identities:52/118(44%), Positives:69/118(58%), Gaps:3/118(2%)

Query  94   MTTISDISNYLVEKILSRAPITCLGAVRTTCKRWNALSKENTLC-NGEARHQFLGFMTKK  270
            MTTISD+S+ LV  ILSR P T L +VR+TCK+WNALSK         AR+QFL FM
Sbjct  1    MTTISDLSDDLVGDILSRVPFTSLISVRSTCKKWNALSKNQIFGRKTAARNQFLEFMILD  60

Query  271  YKLCSMRFRSQWKI*rrrr*rfrGSNYKGVR*FS-ESC*DI*CVFQCDGLLLCVTNED  441
             ++CS+R   Q  I    +  F   + K +   S +   +I  V+ CDGLLLC+  E+
Sbjct  61   SRVCSLRLDLQ-GIRNEDKEDFVDPSMKLISIPSNDDQVEISQVYHCDGLLLCIAKEN  117


>F-box family protein [Arabidopsis thaliana]
Sequence ID: NP_001117347.1 Length: 433
>F-box family protein [Arabidopsis thaliana]
Sequence ID: NP_001117348.1 Length: 433 >F-box family protein [Arabidopsis thaliana]
Sequence ID: NP_001117350.1 Length: 433 >F-box family protein [Arabidopsis thaliana]
Sequence ID: NP_001117352.1 Length: 433 >F-box family protein [Arabidopsis thaliana]
Sequence ID: NP_001185091.1 Length: 433 >F-box family protein [Arabidopsis thaliana]
Sequence ID: NP_001319077.1 Length: 433 >RecName: Full=F-box/kelch-repeat protein At1g24800 [Arabidopsis thaliana]
Sequence ID: P0DI02.1 Length: 433 >RecName: Full=F-box/kelch-repeat protein At1g24881 [Arabidopsis thaliana]
Sequence ID: P0DI03.1 Length: 433 >RecName: Full=F-box/kelch-repeat protein At1g25055 [Arabidopsis thaliana]
Sequence ID: P0DI04.1 Length: 433 >RecName: Full=F-box/kelch-repeat protein At1g25150 [Arabidopsis thaliana]
Sequence ID: P0DI05.1 Length: 433 >RecName: Full=F-box/kelch-repeat protein At1g25211 [Arabidopsis thaliana]
Sequence ID: P0DI06.1 Length: 433 >unknown protein [Arabidopsis thaliana]
Sequence ID: AAN12922.1 Length: 433 >At1g24880 [Arabidopsis thaliana]
Sequence ID: AAQ55275.1 Length: 433 >F-box family protein [Arabidopsis thaliana]
Sequence ID: AEE30564.1 Length: 433 >F-box family protein [Arabidopsis thaliana]
Sequence ID: AEE30571.1 Length: 433 >F-box family protein [Arabidopsis thaliana]
Sequence ID: AEE30577.1 Length: 433
Range 1: 23 to 132

Score:79.0 bits(193), Expect:4e-16,
Method:Compositional matrix adjust.,
Identities:51/116(44%), Positives:70/116(60%), Gaps:7/116(6%)

Query  97   TTISDISNYLV-EKILSRAPITCLGAVRTTCKRWNALSKENTLCNGEARHQFLGFMTKKY  273
            T++ D+   LV EKIL+R PIT L AVR+TCK WNAL+K+  L  G+A  QFLGFMT
Sbjct  23   TSMCDLPPKLVGEKILTRIPITSLRAVRSTCKLWNALTKDRVL--GKAAAQFLGFMTMDS  80

Query  274  KLCSMRFRSQWKI*rrrr*rfrGSNYKGVR*FSESC*DI*CVFQCDGLLLCVTNED  441
            K+CS+RF    +  +         + K V   ++   +I  V+ CDGLLLCV  ++
Sbjct  81   KVCSVRF--HLRRSKEEEEDTMDLSIKQVDLLNQV--EISRVYHCDGLLLCVAKDN  132


>F-box associated ubiquitination effector family protein [Arabidopsis thaliana]
Sequence ID: NP_001323259.1 Length: 317
>F-box associated ubiquitination effector family protein [Arabidopsis thaliana]
Sequence ID: ANM61014.1 Length: 317
Range 1: 1 to 158

Score:78.2 bits(191), Expect:4e-16,
Method:Compositional matrix adjust.,
Identities:63/163(39%), Positives:79/163(48%), Gaps:33/163(20%)

Query  94   MTTISDISNYLVEKILSRAPITCLGAVRTTCKRWNALSKENTLCNGE--ARHQFLGFMTK  267
            MTT+SD+S  LV +ILSR P+T L AVR TCK WN LSK       E  A  QFLGF
Sbjct  1    MTTMSDLSVDLVGEILSRVPLTSLSAVRCTCKSWNTLSKHQIFGKAELAATKQFLGFTVM  60

Query  268  KYKLCSMRF----------------RSQWKI*rrrr*rfrGSNYKGVR*FSESC------  381
             YK+CS+RF                +S  KI  +   +  G+   G+   +  C
Sbjct  61   DYKVCSLRFDLQGIRNDGDDFVDHGKSGLKIRNKAVKKNHGN---GLHRKTGRCHIVERS  117

Query  382  ----*DI*CVFQCDGLLLCVTNEDKTRLASSWYGTHKLDILNP  498
                 ++  VF  DGLLLCVT +D  RL    + T K  IL P
Sbjct  118  ELNQVEVYDVFHSDGLLLCVT-KDHWRLVHV-HNTQKSMILTP  158


>putative F5A9.18 protein [Arabidopsis thaliana]
Sequence ID: AAL86342.1 Length: 433
Range 1: 23 to 132

Score:79.0 bits(193), Expect:4e-16,
Method:Compositional matrix adjust.,
Identities:51/116(44%), Positives:70/116(60%), Gaps:7/116(6%)

Query  97   TTISDISNYLV-EKILSRAPITCLGAVRTTCKRWNALSKENTLCNGEARHQFLGFMTKKY  273
            T++ D+   LV EKIL+R PIT L AVR+TCK WNAL+K+  L  G+A  QFLGFMT
Sbjct  23   TSMCDLPPKLVGEKILTRIPITSLRAVRSTCKLWNALTKDRVL--GKAAAQFLGFMTMDS  80

Query  274  KLCSMRFRSQWKI*rrrr*rfrGSNYKGVR*FSESC*DI*CVFQCDGLLLCVTNED  441
            K+CS+RF    +  +         + K V   ++   +I  V+ CDGLLLCV  ++
Sbjct  81   KVCSVRF--HLRRSKEEEEDTMDLSIKQVDLLNQV--EISRVYHCDGLLLCVAKDN  132


>unnamed protein product [Arabidopsis thaliana]
Sequence ID: VYS47126.1 Length: 433
Range 1: 23 to 132

Score:79.0 bits(193), Expect:4e-16,
Method:Compositional matrix adjust.,
Identities:51/116(44%), Positives:70/116(60%), Gaps:7/116(6%)

Query  97   TTISDISNYLV-EKILSRAPITCLGAVRTTCKRWNALSKENTLCNGEARHQFLGFMTKKY  273
            T++ D+   LV EKIL+R PIT L AVR+TCK WNAL+K+  L  G+A  QFLGFMT
Sbjct  23   TSMCDLPPKLVGEKILTRIPITSLRAVRSTCKLWNALTKDRVL--GKAAAQFLGFMTMDS  80

Query  274  KLCSMRFRSQWKI*rrrr*rfrGSNYKGVR*FSESC*DI*CVFQCDGLLLCVTNED  441
            K+CS+RF    +  +         + K V   ++   +I  V+ CDGLLLCV  ++
Sbjct  81   KVCSVRF--HLRRSKEEEEDTMDLSIKQVDLLNQV--EISRVYHCDGLLLCVAKDN  132


>hypothetical protein [Arabidopsis thaliana]
Sequence ID: BAD94008.1 Length: 433
>hypothetical protein [Arabidopsis thaliana]
Sequence ID: BAF01548.1 Length: 433
Range 1: 23 to 132

Score:79.0 bits(193), Expect:4e-16,
Method:Compositional matrix adjust.,
Identities:51/116(44%), Positives:70/116(60%), Gaps:7/116(6%)

Query  97   TTISDISNYLV-EKILSRAPITCLGAVRTTCKRWNALSKENTLCNGEARHQFLGFMTKKY  273
            T++ D+   LV EKIL+R PIT L AVR+TCK WNAL+K+  L  G+A  QFLGFMT
Sbjct  23   TSMCDLPPKLVGEKILTRIPITSLRAVRSTCKLWNALTKDRVL--GKAAAQFLGFMTMDS  80

Query  274  KLCSMRFRSQWKI*rrrr*rfrGSNYKGVR*FSESC*DI*CVFQCDGLLLCVTNED  441
            K+CS+RF    +  +         + K V   ++   +I  V+ CDGLLLCV  ++
Sbjct  81   KVCSVRF--HLRRSKEEEEDTMDLSIKQVDLLNQV--EISRVYHCDGLLLCVAKDN  132


>F-box and associated interaction domains-containing protein [Arabidopsis thaliana]
Sequence ID: NP_187980.1 Length: 395
>RecName: Full=F-box/kelch-repeat protein At3g13680 [Arabidopsis thaliana]
Sequence ID: Q9LID1.1 Length: 395 >F-box and associated interaction domains-containing protein [Arabidopsis thaliana]
Sequence ID: AEE75396.1 Length: 395 >unnamed protein product [Arabidopsis thaliana]
Sequence ID: BAB01916.1 Length: 395 >unnamed protein product [Arabidopsis thaliana]
Sequence ID: VYS57270.1 Length: 395
Range 1: 1 to 123

Score:78.6 bits(192), Expect:4e-16,
Method:Compositional matrix adjust.,
Identities:49/128(38%), Positives:75/128(58%), Gaps:8/128(6%)

Query  94   MTTISDISNYLVEKILSRAPITCLGAVRTTCKRWNALSKENTLCNGE---ARHQFLGFMT  264
            MTT+ D+   LVE+ILSR P+T L A+R+TC++WN+LSK + +C  +   A ++FLGFM
Sbjct  1    MTTMGDLPGDLVEEILSRVPLTSLRAIRSTCQKWNSLSK-SQICGRKATAAENKFLGFMM  59

Query  265  KKYKLCSMRFRSQWKI*rrrr*rfrGSNYKGVR*FSESC*DI*CVFQCDGLLLCVTNEDK  444
            K  ++CSM+F  Q    R         + K V    +   ++  VF CDGL+LC+  ++
Sbjct  60   KDSRVCSMKFDLQG--IRNDDGELVEPSIKQVSKLDQI--EVSQVFHCDGLVLCIIKDNT  115

Query  445  TRLASSWY  468
              L  + Y
Sbjct  116  GLLVWNPY  123


>unnamed protein product [Arabidopsis thaliana]
Sequence ID: CAA0382322.1 Length: 395
Range 1: 1 to 123

Score:78.6 bits(192), Expect:4e-16,
Method:Compositional matrix adjust.,
Identities:48/127(38%), Positives:71/127(55%), Gaps:6/127(4%)

Query  94   MTTISDISNYLVEKILSRAPITCLGAVRTTCKRWNALSKENTLCNG--EARHQFLGFMTK  267
            MTT+ D+   LVE+ILSR P+T L A+R+TC++WN+LSK          A ++FLGFM K
Sbjct  1    MTTMGDLPGDLVEEILSRVPLTSLRAIRSTCQKWNSLSKNQIFGRKATAAENKFLGFMMK  60

Query  268  KYKLCSMRFRSQWKI*rrrr*rfrGSNYKGVR*FSESC*DI*CVFQCDGLLLCVTNEDKT  447
              ++CSM+F  Q    R         + K V    +   ++  VF CDGL+LC+  ++
Sbjct  61   DSRVCSMKFDLQG--IRNDDGELVEPSIKQVSKLDQI--EVSQVFHCDGLVLCIIKDNTG  116

Query  448  RLASSWY  468
             L  + Y
Sbjct  117  LLVWNPY  123


>hypothetical protein AXX17_AT3G14080 [Arabidopsis thaliana]
Sequence ID: OAP01484.1 Length: 395
Range 1: 1 to 123

Score:78.6 bits(192), Expect:4e-16,
Method:Compositional matrix adjust.,
Identities:48/127(38%), Positives:71/127(55%), Gaps:6/127(4%)

Query  94   MTTISDISNYLVEKILSRAPITCLGAVRTTCKRWNALSKENTLCNG--EARHQFLGFMTK  267
            MTT+ D+   LVE+ILSR P+T L A+R+TC++WN+LSK          A ++FLGFM K
Sbjct  1    MTTMGDLPGDLVEEILSRVPLTSLRAIRSTCQKWNSLSKNQIFGRKATAAENKFLGFMMK  60

Query  268  KYKLCSMRFRSQWKI*rrrr*rfrGSNYKGVR*FSESC*DI*CVFQCDGLLLCVTNEDKT  447
              ++CSM+F  Q    R         + K V    +   ++  VF CDGL+LC+  ++
Sbjct  61   DSRVCSMKFDLQG--IRNDDGELVEPSIKQVSKLDQI--EVSQVFHCDGLVLCIIKDNTG  116

Query  448  RLASSWY  468
             L  + Y
Sbjct  117  LLVWNPY  123


>unnamed protein product [Arabidopsis thaliana]
Sequence ID: VYS63000.1 Length: 381
Range 1: 1 to 114

Score:78.6 bits(192), Expect:5e-16,
Method:Compositional matrix adjust.,
Identities:54/133(41%), Positives:71/133(53%), Gaps:32/133(24%)

Query  94   MTTISDISNYLVEKILSRAPITCLGAVRTTCKRWNALSKENTLCNGEARHQFLGFMTKKY  273
            MTT+SD+S  LV +IL+R P+T L +VR TCK WNALSKE       AR+QF+GF
Sbjct  1    MTTMSDLSPDLVGEILTRVPMTSLISVRCTCKMWNALSKEGIFFKA-ARNQFMGFTMMDS  59

Query  274  KLCSMRFRSQWKI*rrrr*rfrGSNYKGVR----*FSESC---------*DI*CVFQCDG  414
            ++CSM+F  Q                 G+R     F + C          ++  V QCDG
Sbjct  60   RVCSMKFDLQ-----------------GIRNNEQDFVDPCIKQIAKLDQIEVSKVLQCDG  102

Query  415  LLLCVTNEDKTRL  453
            LLLCV  +D +RL
Sbjct  103  LLLCV-GKDNSRL  114


>hypothetical protein AXX17_AT3G43640 [Arabidopsis thaliana]
Sequence ID: OAP03761.1 Length: 662
Range 1: 1 to 119

Score:79.0 bits(193), Expect:6e-16,
Method:Compositional matrix adjust.,
Identities:51/120(43%), Positives:69/120(57%), Gaps:3/120(2%)

Query  94   MTTISDISNYLVEKILSRAPITCLGAVRTTCKRWNALSKENTLC-NGEARHQFLGFMTKK  270
            MTTISD+S+  V  ILSR P T L +VR+TCK+WNALSK         AR+QFL FM
Sbjct  1    MTTISDLSDDFVGDILSRVPFTSLISVRSTCKKWNALSKNQIFGRKTAARNQFLEFMILD  60

Query  271  YKLCSMRFRSQWKI*rrrr*rfrGSNYKGVR*FS-ESC*DI*CVFQCDGLLLCVTNEDKT  447
             ++CS+R   Q  I    +  F   + K +   S +   +I  V+ CDGLLLC+  E+ +
Sbjct  61   SRVCSLRLDLQ-GIRNEDKEDFVDPSMKLISIPSNDDQVEISQVYHCDGLLLCIAKENSS  119


>hypothetical protein [Arabidopsis thaliana]
Sequence ID: BAF00445.1 Length: 433
Range 1: 23 to 132

Score:78.2 bits(191), Expect:8e-16,
Method:Compositional matrix adjust.,
Identities:50/116(43%), Positives:70/116(60%), Gaps:7/116(6%)

Query  97   TTISDISNYLV-EKILSRAPITCLGAVRTTCKRWNALSKENTLCNGEARHQFLGFMTKKY  273
            T++ D+   LV EKI++R PIT L AVR+TCK WNAL+K+  L  G+A  QFLGFMT
Sbjct  23   TSMCDLPPKLVGEKIITRIPITSLRAVRSTCKLWNALTKDRVL--GKAAAQFLGFMTMDS  80

Query  274  KLCSMRFRSQWKI*rrrr*rfrGSNYKGVR*FSESC*DI*CVFQCDGLLLCVTNED  441
            K+CS+RF    +  +         + K V   ++   +I  V+ CDGLLLCV  ++
Sbjct  81   KVCSVRF--HLRRSKEEEEDTMDLSIKQVDLLNQV--EISRVYHCDGLLLCVAKDN  132


>F-box and associated interaction domains-containing protein [Arabidopsis thaliana]
Sequence ID: NP_193453.1 Length: 381
>RecName: Full=Putative F-box protein At4g17200 [Arabidopsis thaliana]
Sequence ID: Q9M0M7.1 Length: 381 >F-box and associated interaction domains-containing protein [Arabidopsis thaliana]
Sequence ID: AEE83861.1 Length: 381 >putative protein [Arabidopsis thaliana]
Sequence ID: CAB78723.1 Length: 381
Range 1: 1 to 114

Score:77.8 bits(190), Expect:8e-16,
Method:Compositional matrix adjust.,
Identities:54/133(41%), Positives:70/133(52%), Gaps:32/133(24%)

Query  94   MTTISDISNYLVEKILSRAPITCLGAVRTTCKRWNALSKENTLCNGEARHQFLGFMTKKY  273
            MTT+SD+S  LV +IL+R P+T L +VR TCK WNALSKE       AR QF+GF
Sbjct  1    MTTMSDLSPDLVGEILTRVPMTSLISVRCTCKMWNALSKEGIFFKA-ARKQFMGFTMMDS  59

Query  274  KLCSMRFRSQWKI*rrrr*rfrGSNYKGVR----*FSESC---------*DI*CVFQCDG  414
            ++CSM+F  Q                 G+R     F + C          ++  V QCDG
Sbjct  60   RVCSMKFDLQ-----------------GIRNNEHDFVDPCIKQIAKLDQIEVSKVLQCDG  102

Query  415  LLLCVTNEDKTRL  453
            LLLCV  +D +RL
Sbjct  103  LLLCV-GKDNSRL  114


>F-box and associated interaction domains-containing protein [Arabidopsis thaliana]
Sequence ID: NP_564661.1 Length: 362
>RecName: Full=F-box protein At1g54550 [Arabidopsis thaliana]
Sequence ID: Q9SLH9.1 Length: 362 >Hypothetical protein [Arabidopsis thaliana]
Sequence ID: AAD25633.1 Length: 362 >F-box and associated interaction domains-containing protein [Arabidopsis thaliana]
Sequence ID: AEE33117.1 Length: 362 >hypothetical protein AXX17_AT1G49040 [Arabidopsis thaliana]
Sequence ID: OAP13347.1 Length: 362 >unnamed protein product [Arabidopsis thaliana]
Sequence ID: CAA0294790.1 Length: 362 >unnamed protein product [Arabidopsis thaliana]
Sequence ID: VYS49102.1 Length: 362
Range 1: 1 to 135

Score:77.4 bits(189), Expect:1e-15,
Method:Compositional matrix adjust.,
Identities:49/144(34%), Positives:77/144(53%), Gaps:16/144(11%)

Query  94   MTTISDISNYLVEKILSRAPITCLGAVRTTCKRWNALSKENTLC-NGEARHQFLGFMTKK  270
            M T++D+ + LV +I SR P+T L AVR+TCK+WNA+SK + L     A++QFL FM
Sbjct  1    MATVTDLPDDLVREIFSRVPLTSLRAVRSTCKKWNAISKYDILGKKAAAKNQFLEFMVTD  60

Query  271  YKLCSMRF-----RSQWKI*rrrr*rfrGSNYKGVR*FSESC*DI*CVFQCDGLLLCVTN  435
             ++CS+R      RS+  +      +    N       S+       V+ CDGLLLC+
Sbjct  61   SRVCSLRLDLQGIRSEEDLIDLSIKQISIPNKVDQVEISQ-------VYHCDGLLLCIAK  113

Query  436  EDKTRLA-SSWYGTHKLDILNPHK  504
            ++ + +  + + G  KL  + P K
Sbjct  114  DNSSVMVWNPYLGQTKL--IQPRK  135


>unnamed protein product [Arabidopsis thaliana]
Sequence ID: CAA0240873.1 Length: 433
Range 1: 23 to 132

Score:77.8 bits(190), Expect:1e-15,
Method:Compositional matrix adjust.,
Identities:50/116(43%), Positives:70/116(60%), Gaps:7/116(6%)

Query  97   TTISDISNYLV-EKILSRAPITCLGAVRTTCKRWNALSKENTLCNGEARHQFLGFMTKKY  273
            T++ D+   LV EKIL++ PIT L AVR+TCK WNAL+K+  L  G+A  QFLGFMT
Sbjct  23   TSMCDLPPKLVGEKILTKIPITSLRAVRSTCKLWNALTKDRVL--GKAAAQFLGFMTMDS  80

Query  274  KLCSMRFRSQWKI*rrrr*rfrGSNYKGVR*FSESC*DI*CVFQCDGLLLCVTNED  441
            K+CS+RF    +  +         + K V   ++   +I  V+ CDGLLLCV  ++
Sbjct  81   KVCSVRF--HLRRSKEEEEDTMDLSIKQVDLLNQV--EISRVYHCDGLLLCVAKDN  132


>F-box and associated interaction domains-containing protein [Arabidopsis thaliana]
Sequence ID: NP_001321070.1 Length: 208
>F-box and associated interaction domains-containing protein [Arabidopsis thaliana]
Sequence ID: ANM58652.1 Length: 208
Range 1: 3 to 112

Score:75.1 bits(183), Expect:1e-15,
Method:Compositional matrix adjust.,
Identities:52/119(44%), Positives:63/119(52%), Gaps:10/119(8%)

Query  103  ISDISNYLVEKILSRAPITCLGAVRTTCKRWNALSKENTLCNGEARHQFLGFMTKKYKLC  282
            ISD+   +VE+ILSR  I  LGA+R     WN LSK   +C  EAR QF GFM K  K+C
Sbjct  3    ISDLPEDMVEEILSRVSIISLGALR-----WNDLSKARVICKAEARQQFAGFMIKGSKVC  57

Query  283  SMRFRSQWKI*rrrr*rfrGSNYKGVR*FSESC*DI*CVFQCDGLLLCVTNE-DKTRLA  456
            SMRF     I          S  K +  F+    +I  VF CDGLLL ++ E   TRL
Sbjct  58   SMRFDLH-GIQNNNVEVVEPS-IKQIAKFNHV--EISQVFHCDGLLLMMSKEVSNTRLV  112


>hypothetical protein AXX17_AT1G61370 [Arabidopsis thaliana]
Sequence ID: OAP16804.1 Length: 159
Range 1: 3 to 112

Score:73.9 bits(180), Expect:1e-15,
Method:Compositional matrix adjust.,
Identities:49/119(41%), Positives:63/119(52%), Gaps:10/119(8%)

Query  103  ISDISNYLVEKILSRAPITCLGAVRTTCKRWNALSKENTLCNGEARHQFLGFMTKKYKLC  282
            IS++   +VE+ILSR  I  LGA+R     WN LSK   +C  EAR QF GFM K  K+C
Sbjct  3    ISNLPEDMVEEILSRVSIISLGALR-----WNDLSKARVICKAEARQQFAGFMIKGSKVC  57

Query  283  SMRFRSQWKI*rrrr*rfrGSNYKGVR*FSESC*DI*CVFQCDGLLLCVTNE-DKTRLA  456
            SMRF       +         + K +  F+    +I  VF CDGLLL ++ E   TRL
Sbjct  58   SMRFDLHG--IQNNNVEVVEPSIKQIAKFNHV--EISQVFHCDGLLLMMSKEVSNTRLV  112


>Contains similarity to hypothetical protein gb|Z97343 from A. thaliana [Arabidopsis thaliana]
Sequence ID: AAC18790.1 Length: 246
Range 1: 2 to 111

Score:75.5 bits(184), Expect:2e-15,
Method:Compositional matrix adjust.,
Identities:50/119(42%), Positives:63/119(52%), Gaps:10/119(8%)

Query  103  ISDISNYLVEKILSRAPITCLGAVRTTCKRWNALSKENTLCNGEARHQFLGFMTKKYKLC  282
            ISD+   +VE+ILSR  I  LGA+R     WN LSK   +C  EAR QF GFM K  K+C
Sbjct  2    ISDLPEDMVEEILSRVSIISLGALR-----WNDLSKARVICKAEARQQFAGFMIKGSKVC  56

Query  283  SMRFRSQWKI*rrrr*rfrGSNYKGVR*FSESC*DI*CVFQCDGLLLCVTNE-DKTRLA  456
            SMRF       +         + K +  F+    +I  VF CDGLLL ++ E   TRL
Sbjct  57   SMRFDLHG--IQNNNVEVVEPSIKQIAKFNHV--EISQVFHCDGLLLMMSKEVSNTRLV  111


>T30E16.27 [Arabidopsis thaliana]
Sequence ID: AAF79759.1 Length: 325
Range 1: 1 to 72

Score:76.6 bits(187), Expect:2e-15,
Method:Compositional matrix adjust.,
Identities:40/72(56%), Positives:46/72(63%), Gaps:2/72(2%)

Query  94   MTTISDISNYLVEKILSRAPITCLGAVRTTCKRWNALSKENTLCNGE--ARHQFLGFMTK  267
            MTT+SD+S  LV +ILSR P+T L AVR TCK WN LSK       E  A  QFLGF
Sbjct  1    MTTMSDLSVDLVGEILSRVPLTSLSAVRCTCKSWNTLSKHQIFGKAELAATKQFLGFTVM  60

Query  268  KYKLCSMRFRSQ  303
             YK+CS+RF  Q
Sbjct  61   DYKVCSLRFDLQ  72


>hypothetical protein AXX17_AT4G20250 [Arabidopsis thaliana]
Sequence ID: OAP00162.1 Length: 381
Range 1: 1 to 114

Score:77.0 bits(188), Expect:2e-15,
Method:Compositional matrix adjust.,
Identities:53/133(40%), Positives:70/133(52%), Gaps:32/133(24%)

Query  94   MTTISDISNYLVEKILSRAPITCLGAVRTTCKRWNALSKENTLCNGEARHQFLGFMTKKY  273
            MTT+SD+S  LV +IL+R P+T L +VR TCK WNALSKE       AR+QF+GF
Sbjct  1    MTTMSDLSPDLVGEILTRVPMTSLISVRCTCKMWNALSKEGIFFKA-ARNQFMGFTIMDS  59

Query  274  KLCSMRFRSQWKI*rrrr*rfrGSNYKGVR----*FSESC---------*DI*CVFQCDG  414
            ++CSM+F  Q                 G+R     F + C          ++  V  CDG
Sbjct  60   RVCSMKFDLQ-----------------GIRNNEQDFVDPCIKQIAKLDQIEVSKVLHCDG  102

Query  415  LLLCVTNEDKTRL  453
            LLLCV  +D +RL
Sbjct  103  LLLCV-GKDNSRL  114


>unnamed protein product [Arabidopsis thaliana]
Sequence ID: CAA0321428.1 Length: 265
Range 1: 3 to 112

Score:75.5 bits(184), Expect:2e-15,
Method:Compositional matrix adjust.,
Identities:52/119(44%), Positives:63/119(52%), Gaps:10/119(8%)

Query  103  ISDISNYLVEKILSRAPITCLGAVRTTCKRWNALSKENTLCNGEARHQFLGFMTKKYKLC  282
            ISD+   +VE+ILSR  I  LGA+R     WN LSK   +C  EAR QF GFM K  K+C
Sbjct  3    ISDLPEDMVEEILSRVSIISLGALR-----WNDLSKARVICKAEARQQFAGFMIKGSKVC  57

Query  283  SMRFRSQWKI*rrrr*rfrGSNYKGVR*FSESC*DI*CVFQCDGLLLCVTNE-DKTRLA  456
            SMRF     I          S  K +  F+    +I  VF CDGLLL ++ E   TRL
Sbjct  58   SMRFDLH-GIQNNNVEVVEPS-IKQIAKFNHV--EISQVFHCDGLLLMMSKEVSNTRLV  112


>EDA1 [Arabidopsis thaliana]
Sequence ID: OAP13995.1 Length: 380
>unnamed protein product [Arabidopsis thaliana]
Sequence ID: CAA0302856.1 Length: 380
Range 1: 1 to 145

Score:76.6 bits(187), Expect:2e-15,
Method:Compositional matrix adjust.,
Identities:58/149(39%), Positives:73/149(48%), Gaps:32/149(21%)

Query  94   MTTISDISNYLVEKILSRAPITCLGAVRTTCKRWNALSKENTLCNGE--ARHQFLGFMTK  267
            MTT+SD+S  LV +ILSR P+T L AVR TCK WN LSK       E  A  QFLGF
Sbjct  1    MTTMSDLSVDLVGEILSRVPLTSLSAVRCTCKSWNTLSKHQIFGKAELAATKQFLGFTVM  60

Query  268  KYKLCSMRF----------------RSQWKI*rrrr*rfrGSNYKGVR*FSESC------  381
             YK+CS+RF                +S  KI  +   +  G+   G+   +  C
Sbjct  61   DYKVCSLRFDLQGIRNDGDDFVDHGKSGLKIRNKAVKKNHGN---GLHRKTGRCHIVERS  117

Query  382  ----*DI*CVFQCDGLLLCVTNEDKTRLA  456
                 ++  VF  DGLLLCVT +D  RL
Sbjct  118  ELNQVEVYDVFHSDGLLLCVT-KDHWRLV  145


>hypothetical protein AXX17_AT3G43650 [Arabidopsis thaliana]
Sequence ID: OAP01578.1 Length: 389
Range 1: 1 to 117

Score:76.3 bits(186), Expect:3e-15,
Method:Compositional matrix adjust.,
Identities:48/118(41%), Positives:65/118(55%), Gaps:3/118(2%)

Query  94   MTTISDISNYLVEKILSRAPITCLGAVRTTCKRWNALSKENTLCNGE-ARHQFLGFMTKK  270
            MTTISD+ + LV+ I S  P T L AVR+TCK WNALSK         AR+QFL  M
Sbjct  1    MTTISDLPDDLVKDIFSWVPFTSLRAVRSTCKTWNALSKNQIFGKKSVARNQFLELMILD  60

Query  271  YKLCSMRFRSQWKI*rrrr*rfrGSNYKGVR*FS-ESC*DI*CVFQCDGLLLCVTNED  441
             ++CS+RF  Q KI           + K +   + +   +I  V+ CDGLLLC+  ++
Sbjct  61   SRVCSLRFDLQ-KIRIEDEEDLVDPSMKQISIPNIDDQVEISRVYHCDGLLLCIPKDN  117


>F-box and associated interaction domains-containing protein [Arabidopsis thaliana]
Sequence ID: NP_190521.1 Length: 388
>RecName: Full=Putative F-box protein At3g49520 [Arabidopsis thaliana]
Sequence ID: Q9SCK7.1 Length: 388 >F-box and associated interaction domains-containing protein [Arabidopsis thaliana]
Sequence ID: AEE78552.1 Length: 388 >putative protein [Arabidopsis thaliana]
Sequence ID: CAB62456.1 Length: 388
Range 1: 1 to 119

Score:75.5 bits(184), Expect:6e-15,
Method:Compositional matrix adjust.,
Identities:48/120(40%), Positives:66/120(55%), Gaps:3/120(2%)

Query  94   MTTISDISNYLVEKILSRAPITCLGAVRTTCKRWNALSKENTLCNGE-ARHQFLGFMTKK  270
            MTTISD+   LV++I S  P T L AVR+TCK WNALSK         AR+QFL  M
Sbjct  1    MTTISDLPYDLVKEIFSWVPFTSLRAVRSTCKTWNALSKNQIFGKKSVARNQFLELMILD  60

Query  271  YKLCSMRFRSQWKI*rrrr*rfrGSNYKGVR*F-SESC*DI*CVFQCDGLLLCVTNEDKT  447
             ++CS+RF  Q KI           + K +    ++   +I  V+ CDGLLLC+  ++ +
Sbjct  61   SRVCSLRFDLQ-KIRNEDEEDLVDPSMKQISIPNNDDQVEISRVYHCDGLLLCIPKDNSS  119


>unnamed protein product [Arabidopsis thaliana]
Sequence ID: CAA0385288.1 Length: 388
Range 1: 1 to 119

Score:75.1 bits(183), Expect:6e-15,
Method:Compositional matrix adjust.,
Identities:48/120(40%), Positives:66/120(55%), Gaps:3/120(2%)

Query  94   MTTISDISNYLVEKILSRAPITCLGAVRTTCKRWNALSKENTLCNGE-ARHQFLGFMTKK  270
            MTTISD+   LV++I S  P T L AVR+TCK WNALSK         AR+QFL  M
Sbjct  1    MTTISDLPYDLVKEIFSWVPFTSLRAVRSTCKTWNALSKNQIFGKKSVARNQFLELMILD  60

Query  271  YKLCSMRFRSQWKI*rrrr*rfrGSNYKGVR*F-SESC*DI*CVFQCDGLLLCVTNEDKT  447
             ++CS+RF  Q KI           + K +    ++   +I  V+ CDGLLLC+  ++ +
Sbjct  61   SRVCSLRFDLQ-KIRNEDEEDLVDPSMKQISIPNNDDQVEISRVYHCDGLLLCIPKDNSS  119


>RecName: Full=Probable F-box protein At1g67455 [Arabidopsis thaliana]
Sequence ID: O64800.3 Length: 343
Range 1: 3 to 112

Score:75.1 bits(183), Expect:7e-15,
Method:Compositional matrix adjust.,
Identities:50/119(42%), Positives:63/119(52%), Gaps:10/119(8%)

Query  103  ISDISNYLVEKILSRAPITCLGAVRTTCKRWNALSKENTLCNGEARHQFLGFMTKKYKLC  282
            ISD+   +VE+ILSR  I  LGA+R     WN LSK   +C  EAR QF GFM K  K+C
Sbjct  3    ISDLPEDMVEEILSRVSIISLGALR-----WNDLSKARVICKAEARQQFAGFMIKGSKVC  57

Query  283  SMRFRSQWKI*rrrr*rfrGSNYKGVR*FSESC*DI*CVFQCDGLLLCVTNE-DKTRLA  456
            SMRF       +         + K +  F+    +I  VF CDGLLL ++ E   TRL
Sbjct  58   SMRFDLHG--IQNNNVEVVEPSIKQIAKFNHV--EISQVFHCDGLLLMMSKEVSNTRLV  112


>hypothetical protein AXX17_AT1G60230 [Arabidopsis thaliana]
Sequence ID: OAP18488.1 Length: 353
Range 1: 1 to 105

Score:74.7 bits(182), Expect:9e-15,
Method:Compositional matrix adjust.,
Identities:54/121(45%), Positives:65/121(53%), Gaps:25/121(20%)

Query  94   MTTISDISNYLVEKILSRAPITCLGAVRTTCKRWNALSKENTLCNGEARHQFLGFMTKKY  273
            M TISD+   LVE+ILSR P+T L AV++TCK  NALSK  T   G+ R QFLGFM   +
Sbjct  1    MGTISDLPVALVEQILSRVPLTSLIAVQSTCKTCNALSK--TQIFGKTRQQFLGFMMIDF  58

Query  274  KLCSMRFRSQWKI*rrrr*rfrGSNYKGVR*FSESC---------*DI*CVFQCDGLLLC  426
             L S++F  Q            G NY+    F E            DI  VF C+GLLLC
Sbjct  59   GLYSIKFDLQ------------GLNYESD--FVEPSIKRVSILDQLDIFKVFHCEGLLLC  104

Query  427  V  429
            V
Sbjct  105  V  105


>unnamed protein product [Arabidopsis thaliana]
Sequence ID: VYS59866.1 Length: 388
Range 1: 1 to 119

Score:73.2 bits(178), Expect:3e-14,
Method:Compositional matrix adjust.,
Identities:47/120(39%), Positives:65/120(54%), Gaps:3/120(2%)

Query  94   MTTISDISNYLVEKILSRAPITCLGAVRTTCKRWNALSKENTLCNGE-ARHQFLGFMTKK  270
            MTTISD+   LV++I S  P T L AVR+TC  WNALSK         AR+QFL  M
Sbjct  1    MTTISDLPYDLVKEIFSWVPFTSLRAVRSTCTTWNALSKNQIFGKKSVARNQFLELMILD  60

Query  271  YKLCSMRFRSQWKI*rrrr*rfrGSNYKGVR*F-SESC*DI*CVFQCDGLLLCVTNEDKT  447
             ++CS+RF  Q KI           + K +    ++   +I  V+ CDGLLLC+  ++ +
Sbjct  61   SRVCSLRFDLQ-KIRNEDEEDLVDPSMKQISIPNNDDQVEISRVYHCDGLLLCIPKDNSS  119


>unnamed protein product [Arabidopsis thaliana]
Sequence ID: BAB02540.1 Length: 359
Range 1: 1 to 119

Score:72.4 bits(176), Expect:6e-14,
Method:Compositional matrix adjust.,
Identities:49/125(39%), Positives:65/125(52%), Gaps:10/125(8%)

Query  94   MTTISDISNYLVEKILSRAPITCLGAVRTTCKRWNALSKENTLC---NGEARHQFLGFMT  264
            MT +SDIS  L+E+ILSR PIT L AV++TCKRW  L  + +      G+  ++FL  MT
Sbjct  1    MTMMSDISQDLLEEILSRVPITSLRAVKSTCKRWKDLLNDPSFSKKYGGKRDNEFLAIMT  60

Query  265  KKYKLCSMRFRSQWKI*rrrr*rfrGSNYKGVR*FSESC*D-I*CVFQCDGLLLCVTNED  441
               +   M       +        +      ++   E   D I  VF CDGLLLC+TNED
Sbjct  61   SGSRASLM------SVNLHGPRDNKDLEDPFIKQIGELNQDQIFKVFHCDGLLLCITNED  114

Query  442  KTRLA  456
             TRL
Sbjct  115  NTRLV  119


>F-box associated ubiquitination effector family protein [Arabidopsis thaliana]
Sequence ID: NP_176177.1 Length: 264
>F-box associated ubiquitination effector family protein [Arabidopsis thaliana]
Sequence ID: AEE33603.1 Length: 264
Range 1: 1 to 69

Score:71.2 bits(173), Expect:6e-14,
Method:Compositional matrix adjust.,
Identities:37/69(54%), Positives:43/69(62%), Gaps:2/69(2%)

Query  103  ISDISNYLVEKILSRAPITCLGAVRTTCKRWNALSKENTLCNGE--ARHQFLGFMTKKYK  276
            +SD+S  LV +ILSR P+T L AVR TCK WN LSK       E  A  QFLGF    YK
Sbjct  1    MSDLSVDLVGEILSRVPLTSLSAVRCTCKSWNTLSKHQIFGKAELAATKQFLGFTVMDYK  60

Query  277  LCSMRFRSQ  303
            +CS+RF  Q
Sbjct  61   VCSLRFDLQ  69


>unnamed protein product [Arabidopsis thaliana]
Sequence ID: CAA0382992.1 Length: 359
Range 1: 1 to 119

Score:72.4 bits(176), Expect:6e-14,
Method:Compositional matrix adjust.,
Identities:49/125(39%), Positives:65/125(52%), Gaps:10/125(8%)

Query  94   MTTISDISNYLVEKILSRAPITCLGAVRTTCKRWNALSKENTLC---NGEARHQFLGFMT  264
            MT +SDIS  L+E+ILSR PIT L AV++TCKRW  L  + +      G+  ++FL  MT
Sbjct  1    MTMMSDISQDLLEEILSRVPITSLRAVKSTCKRWKDLLNDPSFSKKYGGKRDNEFLAIMT  60

Query  265  KKYKLCSMRFRSQWKI*rrrr*rfrGSNYKGVR*FSESC*D-I*CVFQCDGLLLCVTNED  441
               +   M       +        +      ++   E   D I  VF CDGLLLC+TNED
Sbjct  61   SGSRASLM------SVNLHGPRDNKDLEDPFIKQIGELNQDQIFKVFHCDGLLLCITNED  114

Query  442  KTRLA  456
             TRL
Sbjct  115  NTRLV  119


>F-box family protein [Arabidopsis thaliana]
Sequence ID: NP_188590.2 Length: 361
>RecName: Full=Putative F-box protein At3g19560 [Arabidopsis thaliana]
Sequence ID: Q9LJP0.2 Length: 361 >F-box family protein [Arabidopsis thaliana]
Sequence ID: AEE76258.1 Length: 361
Range 1: 3 to 121

Score:72.0 bits(175), Expect:7e-14,
Method:Compositional matrix adjust.,
Identities:49/125(39%), Positives:65/125(52%), Gaps:10/125(8%)

Query  94   MTTISDISNYLVEKILSRAPITCLGAVRTTCKRWNALSKENTLC---NGEARHQFLGFMT  264
            MT +SDIS  L+E+ILSR PIT L AV++TCKRW  L  + +      G+  ++FL  MT
Sbjct  3    MTMMSDISQDLLEEILSRVPITSLRAVKSTCKRWKDLLNDPSFSKKYGGKRDNEFLAIMT  62

Query  265  KKYKLCSMRFRSQWKI*rrrr*rfrGSNYKGVR*FSESC*D-I*CVFQCDGLLLCVTNED  441
               +   M       +        +      ++   E   D I  VF CDGLLLC+TNED
Sbjct  63   SGSRASLM------SVNLHGPRDNKDLEDPFIKQIGELNQDQIFKVFHCDGLLLCITNED  116

Query  442  KTRLA  456
             TRL
Sbjct  117  NTRLV  121


>hypothetical protein [Arabidopsis thaliana]
Sequence ID: CAB10501.1 Length: 364
Range 1: 1 to 104

Score:70.5 bits(171), Expect:3e-13,
Method:Compositional matrix adjust.,
Identities:48/122(39%), Positives:62/122(50%), Gaps:31/122(25%)

Query  103  ISDISNYLVEKILSRAPITCLGAVRTTCKRWNALSKENTLCNGEARHQFLGFMTKKYKLC  282
            +SD+S  LV +IL+R P+T L +VR TCK WNALSKE       AR QF+GF     ++C
Sbjct  1    MSDLSPDLVGEILTRVPMTSLISVRCTCKMWNALSKEGIFFKA-ARKQFMGFTMMDSRVC  59

Query  283  SMRFRSQWKI*rrrr*rfrGSNYKGVR----*FSESC---------*DI*CVFQCDGLLL  423
            SM+F  Q                 G+R     F + C          ++  V QCDGLLL
Sbjct  60   SMKFDLQ-----------------GIRNNEHDFVDPCIKQIAKLDQIEVSKVLQCDGLLL  102

Query  424  CV  429
            CV
Sbjct  103  CV  104


>RecName: Full=Putative F-box protein At3g13825 [Arabidopsis thaliana]
Sequence ID: Q9LRW5.1 Length: 172
>unnamed protein product [Arabidopsis thaliana]
Sequence ID: BAB02906.1 Length: 172
Range 1: 1 to 128

Score:65.5 bits(158), Expect:2e-12,
Method:Compositional matrix adjust.,
Identities:49/137(36%), Positives:67/137(48%), Gaps:14/137(10%)

Query  94   MTTISDISNYLVEKILSRAPITCLGAVRTTCKRWNALS----KEN-TLCNGEARHQFLGF  258
            MTT+S++S  LV +I SR P+  L  VR TC  WN LS     EN      +   QFLGF
Sbjct  1    MTTLSNLSVDLVGEIFSRVPLISLSEVRCTCTTWNTLSWNILSENYVFGKADTSKQFLGF  60

Query  259  MTKKYKLCSMRFRSQWKI*rrrr*rfrGSNYKGVR*FSESC*DI*CVFQCDGLLLCVTNE  438
            +    K+CS+R   Q          F   + K +    +   DI  +F CDGLLLCV
Sbjct  61   VVMNSKVCSLRLDLQ----GIHNNDFVDPSLKEINIVDQY--DISNIFHCDGLLLCVR--  112

Query  439  DKTRLASSWYGTHKLDI  489
               +  S ++  H+LD+
Sbjct  113  -WIQPRSKYHKFHRLDM  128


>unnamed protein product [Arabidopsis thaliana]
Sequence ID: CAA0382341.1 Length: 366
Range 1: 9 to 114

Score:66.6 bits(161), Expect:5e-12,
Method:Compositional matrix adjust.,
Identities:46/115(40%), Positives:66/115(57%), Gaps:12/115(10%)

Query  103  ISDISNYLVEKILSRAPITCLGAVRTTCKRWNALSKENTLCNGEA---RHQFLGFMTKKY  273
            +S +   LV++ILSR PIT L ++R+TCKRW A SK N L  G+A   +  +LGF+
Sbjct  9    MSTLPMVLVDEILSRVPITSLRSLRSTCKRWEAQSKTN-LVGGKATARKSSYLGFILIGN  67

Query  274  KLCSMRFRSQWKI*rrrr*rfrGSNYKGVR*FSESC*DI*CVFQCDGLLLCVTNE  438
            K+CSM      K+       F  ++   V  F +   +I  +F CDGLLLCV+N+
Sbjct  68   KICSM------KLDLNGGDDFVDTSVNQVSAFDDF--EISQLFHCDGLLLCVSNK  114


>unnamed protein product [Arabidopsis thaliana]
Sequence ID: VYS57289.1 Length: 377
Range 1: 9 to 114

Score:66.6 bits(161), Expect:5e-12,
Method:Compositional matrix adjust.,
Identities:46/115(40%), Positives:66/115(57%), Gaps:12/115(10%)

Query  103  ISDISNYLVEKILSRAPITCLGAVRTTCKRWNALSKENTLCNGEA---RHQFLGFMTKKY  273
            +S +   LV++ILSR PIT L ++R+TCKRW A SK N L  G+A   +  +LGF+
Sbjct  9    MSTLPMVLVDEILSRVPITSLRSLRSTCKRWEAQSKTN-LVGGKATARKSSYLGFILIGN  67

Query  274  KLCSMRFRSQWKI*rrrr*rfrGSNYKGVR*FSESC*DI*CVFQCDGLLLCVTNE  438
            K+CSM      K+       F  ++   V  F +   +I  +F CDGLLLCV+N+
Sbjct  68   KICSM------KLDLNGGDDFVDTSVNQVSAFDDF--EISQLFHCDGLLLCVSNK  114


>F-box and associated interaction domains-containing protein [Arabidopsis thaliana]
Sequence ID: NP_187999.1 Length: 377
>RecName: Full=Putative F-box protein At3g13830 [Arabidopsis thaliana]
Sequence ID: Q9LRW4.1 Length: 377 >F-box and associated interaction domains-containing protein [Arabidopsis thaliana]
Sequence ID: AEE75422.1 Length: 377 >hypothetical protein AXX17_AT3G14280 [Arabidopsis thaliana]
Sequence ID: OAP06874.1 Length: 377 >unnamed protein product [Arabidopsis thaliana]
Sequence ID: BAB02907.1 Length: 377
Range 1: 9 to 114

Score:66.2 bits(160), Expect:8e-12,
Method:Compositional matrix adjust.,
Identities:45/115(39%), Positives:66/115(57%), Gaps:12/115(10%)

Query  103  ISDISNYLVEKILSRAPITCLGAVRTTCKRWNALSKENTLCNGEA---RHQFLGFMTKKY  273
            +S +   LV++ILSR PIT L ++R+TCKRW A SK N L  G+A   +  +LGF+
Sbjct  9    MSTLPMVLVDEILSRVPITSLRSLRSTCKRWEAQSKTN-LVGGKATARKSSYLGFILIGN  67

Query  274  KLCSMRFRSQWKI*rrrr*rfrGSNYKGVR*FSESC*DI*CVFQCDGLLLCVTNE  438
            K+CSM      K+       F  ++   +  F +   +I  +F CDGLLLCV+N+
Sbjct  68   KICSM------KLDLNGGDDFVDTSVNQISAFDDF--EISQLFHCDGLLLCVSNK  114


>F-box/associated interaction domain protein [Arabidopsis thaliana]
Sequence ID: NP_188911.2 Length: 379
>RecName: Full=Probable F-box protein At3g22720 [Arabidopsis thaliana]
Sequence ID: Q9LUI9.1 Length: 379 >F-box/associated interaction domain protein [Arabidopsis thaliana]
Sequence ID: AEE76669.2 Length: 379 >unnamed protein product [Arabidopsis thaliana]
Sequence ID: BAB01247.1 Length: 379
Range 1: 2 to 113

Score:65.5 bits(158), Expect:1e-11,
Method:Compositional matrix adjust.,
Identities:44/120(37%), Positives:63/120(52%), Gaps:14/120(11%)

Query  103  ISDISNYLVEKILSRAPITCLGAVRTTCKRWNALSKENTLCN---GEARHQFLGFMTKKY  273
            +SD+   LVEKILSR P T L  +R+TC+RWNAL K+         +A  +    M K+Y
Sbjct  2    MSDLPLDLVEKILSRVPATSLKRLRSTCRRWNALLKDRRFTEKHFHKAPKESRVLMLKEY  61

Query  274  KLCSMRFRSQWKI*rrrr*rfrGSNYKGVR*FSESC*D---I*CVFQCDGLLLCVTNEDK  444
            ++        + I    +      ++KG   +S S  +   I  VF CDGLLLC TN+D+
Sbjct  62   RV--------FPISVNLKVTPPSVDFKGALGYSHSNSEQVVITEVFHCDGLLLCTTNDDR  113


>unnamed protein product [Arabidopsis thaliana]
Sequence ID: VYS58278.1 Length: 372
Range 1: 3 to 114

Score:64.7 bits(156), Expect:2e-11,
Method:Compositional matrix adjust.,
Identities:44/115(38%), Positives:57/115(49%), Gaps:6/115(5%)

Query  103  ISDISNYLVEKILSRAPITCLGAVRTTCKRWNALSKENTLCNGE---ARHQFLGFMTKKY  273
            +SD+S  LVE+ILSR P T L  +R+TCK WNAL K       +   A  + L  M K+Y
Sbjct  3    MSDLSLDLVEEILSRVPATSLKRLRSTCKLWNALFKNPGFTKKQFLKAPKESLVLMLKEY  62

Query  274  KLCSMRFRSQWKI*rrrr*rfrGSNYKGVR*FSESC*DI*CVFQCDGLLLCVTNE  438
             +C M   +   +           N K    +SE   DI   F CDGLLLC T +
Sbjct  63   SVCPM--IANLSVSAPSIEFKGALNLKNYPPYSEEV-DIHEAFHCDGLLLCTTTD  114


>hypothetical protein AXX17_AT3G24470 [Arabidopsis thaliana]
Sequence ID: OAP04928.1 Length: 379
Range 1: 2 to 113

Score:64.3 bits(155), Expect:4e-11,
Method:Compositional matrix adjust.,
Identities:43/120(36%), Positives:63/120(52%), Gaps:14/120(11%)

Query  103  ISDISNYLVEKILSRAPITCLGAVRTTCKRWNALSKENTLCN---GEARHQFLGFMTKKY  273
            +SD+   LVE+ILSR P T L  +R+TC+RWNAL K+         +A  +    M K+Y
Sbjct  2    MSDLPLDLVEEILSRVPATSLKRLRSTCRRWNALLKDRRFTEKHFHKAPKESRVLMLKEY  61

Query  274  KLCSMRFRSQWKI*rrrr*rfrGSNYKGVR*FSESC*D---I*CVFQCDGLLLCVTNEDK  444
            ++        + I    +      ++KG   +S S  +   I  VF CDGLLLC TN+D+
Sbjct  62   RV--------FPISVNLKVTPPSVDFKGALGYSHSNSEQVVITEVFHCDGLLLCTTNDDR  113


>unnamed protein product [Arabidopsis thaliana]
Sequence ID: VYS58276.1 Length: 379
Range 1: 2 to 113

Score:64.3 bits(155), Expect:4e-11,
Method:Compositional matrix adjust.,
Identities:42/120(35%), Positives:63/120(52%), Gaps:14/120(11%)

Query  103  ISDISNYLVEKILSRAPITCLGAVRTTCKRWNALSKENTLCN---GEARHQFLGFMTKKY  273
            +SD+   LVE+ILSR P   L  +R+TC+RWNAL K+         +A  + +  M K+Y
Sbjct  2    MSDLPLDLVEEILSRVPAASLKRLRSTCRRWNALLKDRRFTEKHFHKAPKESIVLMLKEY  61

Query  274  KLCSMRFRSQWKI*rrrr*rfrGSNYKGVR*FSESC*D---I*CVFQCDGLLLCVTNEDK  444
            ++        + I    +      ++KG   +S S  +   I  VF CDGLLLC TN+D+
Sbjct  62   RV--------FPISVNLKVTPPSVDFKGALGYSHSNSEQVVITEVFHCDGLLLCTTNDDR  113


>unnamed protein product [Arabidopsis thaliana]
Sequence ID: CAA0383352.1 Length: 379
Range 1: 2 to 113

Score:64.3 bits(155), Expect:4e-11,
Method:Compositional matrix adjust.,
Identities:42/120(35%), Positives:63/120(52%), Gaps:14/120(11%)

Query  103  ISDISNYLVEKILSRAPITCLGAVRTTCKRWNALSKENTLCN---GEARHQFLGFMTKKY  273
            +SD+   LVE+ILSR P   L  +R+TC+RWNAL K+         +A  + +  M K+Y
Sbjct  2    MSDLPLDLVEEILSRVPAASLKRLRSTCRRWNALLKDRRFTEKHFHKAPKESIVLMLKEY  61

Query  274  KLCSMRFRSQWKI*rrrr*rfrGSNYKGVR*FSESC*D---I*CVFQCDGLLLCVTNEDK  444
            ++        + I    +      ++KG   +S S  +   I  VF CDGLLLC TN+D+
Sbjct  62   RV--------FPISVNLKVTPPSVDFKGALGYSHSNSEQVVITEVFHCDGLLLCTTNDDR  113


>unnamed protein product [Arabidopsis thaliana]
Sequence ID: CAA0382340.1 Length: 415
Range 1: 1 to 108

Score:63.9 bits(154), Expect:5e-11,
Method:Compositional matrix adjust.,
Identities:40/116(34%), Positives:64/116(55%), Gaps:8/116(6%)

Query  94   MTTISDISNYLVEKILSRAPITCLGAVRTTCKRWNALSKENTLCNGEARHQFLGFMTKKY  273
            MTT+S++   ++E+ILSR P+T L  +R+TCK+WN LSK+  +     + Q  G M  K
Sbjct  1    MTTMSNLPAEVLEEILSRTPVTSLRTMRSTCKKWNNLSKKKIIPEAARKQQ--GLMLIKK  58

Query  274  KLCSMRFRSQWKI*rrrr*rfrGSNYKGVR*FSESC*DI*CVFQCDGLLLCVTNED  441
            K+CS+ F S  +I +        +     R       ++  +F CDG+LLCV  ++
Sbjct  59   KICSLSF-SLHEIHKDDYVVPCINQVDIPRNI-----EVEKIFHCDGILLCVIEDN  108


>F-box and associated interaction domains-containing protein [Arabidopsis thaliana]
Sequence ID: NP_187998.1 Length: 415
>RecName: Full=F-box protein At3g13820 [Arabidopsis thaliana]
Sequence ID: Q9LRW6.1 Length: 415 >F-box family protein [Arabidopsis thaliana]
Sequence ID: ABE65938.1 Length: 415 >F-box and associated interaction domains-containing protein [Arabidopsis thaliana]
Sequence ID: AEE75421.1 Length: 415 >unnamed protein product [Arabidopsis thaliana]
Sequence ID: BAB02905.1 Length: 415
Range 1: 1 to 108

Score:63.9 bits(154), Expect:5e-11,
Method:Compositional matrix adjust.,
Identities:40/116(34%), Positives:64/116(55%), Gaps:8/116(6%)

Query  94   MTTISDISNYLVEKILSRAPITCLGAVRTTCKRWNALSKENTLCNGEARHQFLGFMTKKY  273
            MTT+S++   ++E+ILSR P+T L  +R+TCK+WN LSK+  +     + Q  G M  K
Sbjct  1    MTTMSNLPAEVLEEILSRTPVTSLRTMRSTCKKWNNLSKKKIIPEAARKQQ--GLMLIKK  58

Query  274  KLCSMRFRSQWKI*rrrr*rfrGSNYKGVR*FSESC*DI*CVFQCDGLLLCVTNED  441
            K+CS+ F S  +I +        +     R       ++  +F CDG+LLCV  ++
Sbjct  59   KICSLSF-SLHEIHKDDYVVPCINQVDIPRNI-----EVEKIFHCDGILLCVIEDN  108


>unknown, partial [Arabidopsis thaliana]
Sequence ID: ABK28556.1 Length: 416
Range 1: 1 to 108

Score:63.9 bits(154), Expect:5e-11,
Method:Compositional matrix adjust.,
Identities:40/116(34%), Positives:64/116(55%), Gaps:8/116(6%)

Query  94   MTTISDISNYLVEKILSRAPITCLGAVRTTCKRWNALSKENTLCNGEARHQFLGFMTKKY  273
            MTT+S++   ++E+ILSR P+T L  +R+TCK+WN LSK+  +     + Q  G M  K
Sbjct  1    MTTMSNLPAEVLEEILSRTPVTSLRTMRSTCKKWNNLSKKKIIPEAARKQQ--GLMLIKK  58

Query  274  KLCSMRFRSQWKI*rrrr*rfrGSNYKGVR*FSESC*DI*CVFQCDGLLLCVTNED  441
            K+CS+ F S  +I +        +     R       ++  +F CDG+LLCV  ++
Sbjct  59   KICSLSF-SLHEIHKDDYVVPCINQVDIPRNI-----EVEKIFHCDGILLCVIEDN  108


>F-box associated ubiquitination effector family protein [Arabidopsis thaliana]
Sequence ID: NP_172646.3 Length: 377
>RecName: Full=F-box protein At1g11810 [Arabidopsis thaliana]
Sequence ID: Q9SA94.1 Length: 377 >EST gb|AA605362 comes from this gene [Arabidopsis thaliana]
Sequence ID: AAD30240.1 Length: 377 >At1g11810 [Arabidopsis thaliana]
Sequence ID: ABL66798.1 Length: 377 >F-box associated ubiquitination effector family protein [Arabidopsis thaliana]
Sequence ID: AEE28789.1 Length: 377 >hypothetical protein AXX17_AT1G12180 [Arabidopsis thaliana]
Sequence ID: OAP13460.1 Length: 377
Range 1: 3 to 109

Score:63.5 bits(153), Expect:6e-11,
Method:Compositional matrix adjust.,
Identities:41/116(35%), Positives:61/116(52%), Gaps:12/116(10%)

Query  97   TTISDISNYLVEKILSRAPITCLGAVRTTCKRWNALSKENTLCNGEA---RHQFLGFMTK  267
            TT+S +   LV++IL+R PIT L ++R+TCK+W A SK N L  G+A   +   +GF+
Sbjct  3    TTMSTLPVVLVDEILARVPITSLRSLRSTCKKWEASSKTN-LVGGKATARKSSHVGFILI  61

Query  268  KYKLCSMRFRSQWKI*rrrr*rfrGSNYKGVR*FSESC*DI*CVFQCDGLLLCVTN  435
              K+CSM+               + S + G          I  +F CDGLL C++N
Sbjct  62   GDKICSMKLDLNGGDDFVDTSVNQVSAFDGFA--------ISQLFHCDGLLFCISN  109


>hypothetical protein AXX17_AT3G14270 [Arabidopsis thaliana]
Sequence ID: OAP02955.1 Length: 418
>unnamed protein product [Arabidopsis thaliana]
Sequence ID: VYS57288.1 Length: 418
Range 1: 1 to 108

Score:62.4 bits(150), Expect:2e-10,
Method:Compositional matrix adjust.,
Identities:39/116(34%), Positives:63/116(54%), Gaps:8/116(6%)

Query  94   MTTISDISNYLVEKILSRAPITCLGAVRTTCKRWNALSKENTLCNGEARHQFLGFMTKKY  273
            M+T+S++   ++E+ILSR P+T L  +R+TCK+WN LSK+  +     + Q  G M  K
Sbjct  1    MSTMSNLPAEVLEEILSRTPVTSLRTMRSTCKKWNNLSKKKIIPEAARKQQ--GLMLIKK  58

Query  274  KLCSMRFRSQWKI*rrrr*rfrGSNYKGVR*FSESC*DI*CVFQCDGLLLCVTNED  441
            K+CS+ F S   I +        +     R       ++  +F CDG+LLCV  ++
Sbjct  59   KICSLSF-SLHDIHKDDYVVPCINQVDIPRNI-----EVEKIFHCDGILLCVIEDN  108


>hypothetical protein AXX17_AT3G18360 [Arabidopsis thaliana]
Sequence ID: OAP01833.1 Length: 158
>unnamed protein product [Arabidopsis thaliana]
Sequence ID: VYS57685.1 Length: 158
Range 1: 1 to 117

Score:59.7 bits(143), Expect:2e-10,
Method:Compositional matrix adjust.,
Identities:38/122(31%), Positives:60/122(49%), Gaps:12/122(9%)

Query  94   MTTISDISNYLVEKILSRAPITCLGAVRTTCKRWNALSKENTLCNGE-------ARHQFL  252
            MTT+SD+ + L E++LSR P+T L  VR TCK+WN LSK+ +            A  +F+
Sbjct  1    MTTLSDLPSDLAEEVLSRIPVTSLRGVRATCKKWNTLSKDRSFTRKHLAQAKAAAAREFM  60

Query  253  GFMTKKYKLCSMRFRSQWKI*rrrr*rfrGSNYKGVR*FSESC*DI*CVFQCDGLLLCVT  432
              M   +++  M       + +       G   K +     +  DI CV+ CD L+LC+
Sbjct  61   VVMVMDFQVYLMDI----NLHKDVDATINGQG-KLISLDDSNQVDISCVYHCDSLVLCIP  115

Query  433  NE  438
             +
Sbjct  116  KD  117


>unnamed protein product [Arabidopsis thaliana]
Sequence ID: VYS45785.1 Length: 377
Range 1: 3 to 109

Score:62.0 bits(149), Expect:2e-10,
Method:Compositional matrix adjust.,
Identities:40/116(34%), Positives:60/116(51%), Gaps:12/116(10%)

Query  97   TTISDISNYLVEKILSRAPITCLGAVRTTCKRWNALSKENTLCNGEA---RHQFLGFMTK  267
            TT+S +   LV++IL+R PIT L ++R+TCK+W   SK N L  G+A   +   +GF+
Sbjct  3    TTMSTLPVVLVDEILARVPITSLRSLRSTCKKWEGSSKTN-LVGGKATARKSSHVGFILI  61

Query  268  KYKLCSMRFRSQWKI*rrrr*rfrGSNYKGVR*FSESC*DI*CVFQCDGLLLCVTN  435
              K+CSM+               + S + G          I  +F CDGLL C++N
Sbjct  62   GDKICSMKLDLNGGDDFVDTSVNQVSAFDGFA--------ISQLFHCDGLLFCISN  109


>F-box and associated interaction domains-containing protein [Arabidopsis thaliana]
Sequence ID: NP_566558.1 Length: 391
>RecName: Full=F-box/kelch-repeat protein At3g16740 [Arabidopsis thaliana]
Sequence ID: Q9LUQ9.1 Length: 391 >At3g16740 [Arabidopsis thaliana]
Sequence ID: AAO44092.1 Length: 391 >F-box and associated interaction domains-containing protein [Arabidopsis thaliana]
Sequence ID: AEE75859.1 Length: 391 >unnamed protein product [Arabidopsis thaliana]
Sequence ID: CAA0382671.1 Length: 391 >unnamed protein product [Arabidopsis thaliana]
Sequence ID: VYS57614.1 Length: 391 >unnamed protein product [Arabidopsis thaliana]
Sequence ID: BAB02766.1 Length: 391
Range 1: 1 to 123

Score:61.6 bits(148), Expect:3e-10,
Method:Compositional matrix adjust.,
Identities:42/124(34%), Positives:61/124(49%), Gaps:4/124(3%)

Query  94   MTTISDISNYLVEKILSRAPITCLGAVRTTCKRWNALSKENTLCNGEARHQFLGFMTKKY  273
            M  ISD+   L E++LSR P+T + AVR TCK+WN LSK+ +      R        K+
Sbjct  1    MVQISDLPRDLTEEVLSRIPVTSMRAVRFTCKKWNTLSKDRSFTKKHLRGARAAAKKKQT  60

Query  274  KLCSMRFRSQWKI*rrrr*rfrGSNYKGVR*FS---ESC*DI*CVFQCDGLLLCVTNEDK  444
            K   +    Q+++          S  +  +  S   E   DI  +F C GLLLC+T +D
Sbjct  61   KEFQVIMMIQFRVYLYSVNLLNPSIERIGKLISLDVEDHVDISKIFHCGGLLLCIT-KDI  119

Query  445  TRLA  456
            +RL
Sbjct  120  SRLV  123


>hypothetical protein AXX17_AT3G17610 [Arabidopsis thaliana]
Sequence ID: OAP04275.1 Length: 391
Range 1: 1 to 123

Score:60.8 bits(146), Expect:5e-10,
Method:Compositional matrix adjust.,
Identities:42/124(34%), Positives:61/124(49%), Gaps:4/124(3%)

Query  94   MTTISDISNYLVEKILSRAPITCLGAVRTTCKRWNALSKENTLCNGEARHQFLGFMTKKY  273
            M  ISD+   L E++LSR P+T + AVR TCK+WN LSK+ +      R        K+
Sbjct  1    MVQISDLPRDLTEEVLSRIPVTSMRAVRFTCKKWNTLSKDRSFTKKHLRGARGAAKKKQT  60

Query  274  KLCSMRFRSQWKI*rrrr*rfrGSNYKGVR*FS---ESC*DI*CVFQCDGLLLCVTNEDK  444
            K   +    Q+++          S  +  +  S   E   DI  +F C GLLLC+T +D
Sbjct  61   KEFQVIMMIQFRVYLYSVNLLNPSIERIGKLISLDVEDHVDISKIFHCGGLLLCIT-KDI  119

Query  445  TRLA  456
            +RL
Sbjct  120  SRLV  123


>F-box and associated interaction domains-containing protein [Arabidopsis thaliana]
Sequence ID: NP_850216.1 Length: 150
>RecName: Full=Putative F-box protein At2g33655 [Arabidopsis thaliana]
Sequence ID: Q3EBP1.1 Length: 150 >F-box and associated interaction domains-containing protein [Arabidopsis thaliana]
Sequence ID: AEC08866.1 Length: 150
Range 1: 1 to 60

Score:58.2 bits(139), Expect:6e-10,
Method:Compositional matrix adjust.,
Identities:27/60(45%), Positives:35/60(58%), Gaps:0/60(0%)

Query  94   MTTISDISNYLVEKILSRAPITCLGAVRTTCKRWNALSKENTLCNGEARHQFLGFMTKKY  273
            M  +SD+   LVE+ILSR P+  +  VR TCK WNALSK  +        +FLG   KK+
Sbjct  1    MEKMSDLPRELVEEILSRVPVKSMREVRVTCKTWNALSKHISKAEAAREGEFLGIAKKKF  60


>unnamed protein product [Arabidopsis thaliana]
Sequence ID: VYS54341.1 Length: 86
Range 1: 1 to 70

Score:56.2 bits(134), Expect:6e-10,
Method:Compositional matrix adjust.,
Identities:28/70(40%), Positives:38/70(54%), Gaps:0/70(0%)

Query  94   MTTISDISNYLVEKILSRAPITCLGAVRTTCKRWNALSKENTLCNGEARHQFLGFMTKKY  273
            M  +SD+   LVE+ILSR P+  +  VR TCK WNALSK           +FLG    +
Sbjct  1    MEKMSDLPRELVEEILSRVPVKSMREVRLTCKTWNALSKHIGKAEAAREGEFLGIAKNQD  60

Query  274  KLCSMRFRSQ  303
            +L S +  +Q
Sbjct  61   QLASWQVVTQ  70


>F-box and associated interaction domains-containing protein [Arabidopsis thaliana]
Sequence ID: NP_188912.1 Length: 372
>RecName: Full=Putative F-box/kelch-repeat protein At3g22730 [Arabidopsis thaliana]
Sequence ID: Q9LUI8.1 Length: 372 >F-box and associated interaction domains-containing protein [Arabidopsis thaliana]
Sequence ID: AEE76671.1 Length: 372 >hypothetical protein AXX17_AT3G24480 [Arabidopsis thaliana]
Sequence ID: OAP06905.1 Length: 372 >unnamed protein product [Arabidopsis thaliana]
Sequence ID: BAB01248.1 Length: 372
Range 1: 3 to 112

Score:59.7 bits(143), Expect:1e-09,
Method:Compositional matrix adjust.,
Identities:43/113(38%), Positives:55/113(48%), Gaps:6/113(5%)

Query  103  ISDISNYLVEKILSRAPITCLGAVRTTCKRWNALSKENTLCNGE---ARHQFLGFMTKKY  273
            +SD+S  LVE+ILSR P T L  +R+TCK WNAL K       +   A  + L  M K+Y
Sbjct  3    MSDLSLDLVEEILSRVPATSLKRLRSTCKLWNALFKNPGFTKKQFLKAPKESLVLMLKEY  62

Query  274  KLCSMRFRSQWKI*rrrr*rfrGSNYKGVR*FSESC*DI*CVFQCDGLLLCVT  432
             +C M   +   +           N K    +SE   DI     CDGLLLC T
Sbjct  63   SVCPM--IANLSVSAPSIEFKGALNLKNYPPYSEEV-DIHEACHCDGLLLCTT  112


>unnamed protein product [Arabidopsis thaliana]
Sequence ID: CAA0383354.1 Length: 372
Range 1: 3 to 112

Score:59.7 bits(143), Expect:1e-09,
Method:Compositional matrix adjust.,
Identities:43/113(38%), Positives:55/113(48%), Gaps:6/113(5%)

Query  103  ISDISNYLVEKILSRAPITCLGAVRTTCKRWNALSKENTLCNGE---ARHQFLGFMTKKY  273
            +SD+S  LVE+ILSR P T L  +R+TCK WNAL K       +   A  + L  M K+Y
Sbjct  3    MSDLSLDLVEEILSRVPATSLKRLRSTCKLWNALFKNPGFTKKQFLKAPKESLVLMLKEY  62

Query  274  KLCSMRFRSQWKI*rrrr*rfrGSNYKGVR*FSESC*DI*CVFQCDGLLLCVT  432
             +C M   +   +           N K    +SE   DI     CDGLLLC T
Sbjct  63   SVCPM--IANLSVSAPSIEFKGALNLKNYPPYSEEV-DIHEACHCDGLLLCTT  112


>unnamed protein product [Arabidopsis thaliana]
Sequence ID: CAA0382744.1 Length: 158
Range 1: 1 to 117

Score:57.4 bits(137), Expect:1e-09,
Method:Compositional matrix adjust.,
Identities:38/122(31%), Positives:60/122(49%), Gaps:12/122(9%)

Query  94   MTTISDISNYLVEKILSRAPITCLGAVRTTCKRWNALSKENTLCNGE-------ARHQFL  252
            MTT+SD+ + L E++LSR P+T L  VR TCK+WN LSK+ +            A  +F+
Sbjct  1    MTTLSDLPSDLAEEVLSRIPVTSLRGVRATCKKWNTLSKDRSFTRKHLAQAKAAAAREFM  60

Query  253  GFMTKKYKLCSMRFRSQWKI*rrrr*rfrGSNYKGVR*FSESC*DI*CVFQCDGLLLCVT  432
              M   +++  M       + +       G   K +     +  DI CV+ CD L+LC+
Sbjct  61   VVMVMDFQVYLMGI----NLHKDVDATINGQG-KLISLDDSNQVDISCVYHCDCLVLCIP  115

Query  433  NE  438
             +
Sbjct  116  KD  117


>F-box and associated interaction domains-containing protein [Arabidopsis thaliana]
Sequence ID: NP_683479.1 Length: 216
>F-box and associated interaction domains-containing protein [Arabidopsis thaliana]
Sequence ID: AEE34648.1 Length: 216
Range 1: 3 to 58

Score:58.2 bits(139), Expect:2e-09,
Method:Compositional matrix adjust.,
Identities:31/61(51%), Positives:36/61(59%), Gaps:5/61(8%)

Query  103  ISDISNYLVEKILSRAPITCLGAVRTTCKRWNALSKENTLCNGEARHQFLGFMTKKYKLC  282
            ISD+   +VE+ILSR  I  LGA+R     WN LSK   +C  EAR QF GFM K  K
Sbjct  3    ISDLPEDMVEEILSRVSIISLGALR-----WNDLSKARVICKAEARQQFAGFMIKGSKEV  57

Query  283  S  285
            S
Sbjct  58   S  58


>unnamed protein product [Arabidopsis thaliana]
Sequence ID: CAA0383369.1 Length: 413
Range 1: 4 to 112

Score:58.9 bits(141), Expect:2e-09,
Method:Compositional matrix adjust.,
Identities:40/120(33%), Positives:57/120(47%), Gaps:12/120(10%)

Query  100  TISDISNYLVEKILSRAPITCLGAVRTTCKRWNALSKENTLCNGEARHQFLGFMTKKYKL  279
            TISD+  YL +KI SR P+  + A+R TCK W  L K  +L   E   Q +  M   Y L
Sbjct  4    TISDLPRYLKKKIFSRIPLRYVRALRLTCKEWETLIKSRSLKIDEEESQMVALM--DYNL  61

Query  280  CSMRFRSQWKI*rrrr*rfrGSNYKG-VR*FSESC*DI*CVFQCDGLLLCVTNEDKTRLA  456
            C M         +        +  KG +  F E    +  +F C+GLLLC+  +D T++
Sbjct  62   CLM--------SKSFNGGDPSTEIKGKLTCFDEQV-KVSMLFHCEGLLLCILKDDNTKVV  112


>unnamed protein product [Arabidopsis thaliana]
Sequence ID: CAA0374401.1 Length: 90
Range 1: 1 to 55

Score:54.7 bits(130), Expect:3e-09,
Method:Compositional matrix adjust.,
Identities:25/55(45%), Positives:32/55(58%), Gaps:0/55(0%)

Query  94   MTTISDISNYLVEKILSRAPITCLGAVRTTCKRWNALSKENTLCNGEARHQFLGF  258
            M  +SD+   LVE+ILSR P+  +  VR TCK WNALSK  +        +FLG
Sbjct  1    MEKMSDLPRELVEEILSRVPVKSMREVRVTCKTWNALSKHISKAEAAREGEFLGI  55


>unnamed protein product [Arabidopsis thaliana]
Sequence ID: CAA0364593.1 Length: 394
Range 1: 1 to 129

Score:58.2 bits(139), Expect:4e-09,
Method:Compositional matrix adjust.,
Identities:44/131(34%), Positives:66/131(50%), Gaps:12/131(9%)

Query  94   MTTISDISNYLVEKILSRAPITCLGAVRTTCKRWNALSKENTLCNG---EARHQFLGFMT  264
            M  +SD+   L+ +ILSR P+  L +VR TCK+WN LSK+ +       EA+ + L   +
Sbjct  1    MAIMSDLPRDLLAEILSRVPLASLRSVRFTCKKWNDLSKDRSFLKKQIVEAKKKQLK--S  58

Query  265  KKYKLCSMR-FRSQWKI*rrrr*rfrGSNYKG-VR*FSESC-----*DI*CVFQCDGLLL  423
            K++++  MR FR                  KG +  FS+        ++  VF CDGLLL
Sbjct  59   KEFEVIMMRNFRVYLTSVDLHNDVNPSFTPKGTLTSFSDDANQHQVDNVSSVFHCDGLLL  118

Query  424  CVTNEDKTRLA  456
            C+T +   RL
Sbjct  119  CITKDLNFRLV  129


>hypothetical protein [Arabidopsis thaliana]
Sequence ID: AAO37178.1 Length: 394
Range 1: 1 to 129

Score:58.2 bits(139), Expect:4e-09,
Method:Compositional matrix adjust.,
Identities:44/131(34%), Positives:66/131(50%), Gaps:12/131(9%)

Query  94   MTTISDISNYLVEKILSRAPITCLGAVRTTCKRWNALSKENTLCNG---EARHQFLGFMT  264
            M  +SD+   L+ +ILSR P+  L +VR TCK+WN LSK+ +       EA+ + L   +
Sbjct  1    MAIMSDLPRDLLAEILSRVPLASLRSVRFTCKKWNDLSKDRSFLKKQIVEAKKKQLK--S  58

Query  265  KKYKLCSMR-FRSQWKI*rrrr*rfrGSNYKG-VR*FSESC-----*DI*CVFQCDGLLL  423
            K++++  MR FR                  KG +  FS+        ++  VF CDGLLL
Sbjct  59   KEFEVIMMRNFRVYLTSVDLHNDVNPSFTPKGTLTSFSDDANQHQVDNVSSVFHCDGLLL  118

Query  424  CVTNEDKTRLA  456
            C+T +   RL
Sbjct  119  CITKDLNFRLV  129


>F-box and associated interaction domains-containing protein [Arabidopsis thaliana]
Sequence ID: NP_179373.2 Length: 394
>RecName: Full=F-box protein At2g17830 [Arabidopsis thaliana]
Sequence ID: Q84X17.2 Length: 394 >At2g17830 [Arabidopsis thaliana]
Sequence ID: ABJ17127.1 Length: 394 >F-box and associated interaction domains-containing protein [Arabidopsis thaliana]
Sequence ID: AEC06692.1 Length: 394
Range 1: 1 to 129

Score:58.2 bits(139), Expect:4e-09,
Method:Compositional matrix adjust.,
Identities:44/131(34%), Positives:66/131(50%), Gaps:12/131(9%)

Query  94   MTTISDISNYLVEKILSRAPITCLGAVRTTCKRWNALSKENTLCNG---EARHQFLGFMT  264
            M  +SD+   L+ +ILSR P+  L +VR TCK+WN LSK+ +       EA+ + L   +
Sbjct  1    MAIMSDLPRDLLAEILSRVPLASLRSVRFTCKKWNDLSKDRSFLKKQIVEAKKKQLK--S  58

Query  265  KKYKLCSMR-FRSQWKI*rrrr*rfrGSNYKG-VR*FSESC-----*DI*CVFQCDGLLL  423
            K++++  MR FR                  KG +  FS+        ++  VF CDGLLL
Sbjct  59   KEFEVIMMRNFRVYLTSVDLHNDVNPSFTPKGTLTSFSDDANQHQVDNVSSVFHCDGLLL  118

Query  424  CVTNEDKTRLA  456
            C+T +   RL
Sbjct  119  CITKDLNFRLV  129


>hypothetical protein AXX17_AT2G13160 [Arabidopsis thaliana]
Sequence ID: OAP08881.1 Length: 394
>unnamed protein product [Arabidopsis thaliana]
Sequence ID: VYS52680.1 Length: 394
Range 1: 1 to 129

Score:58.2 bits(139), Expect:4e-09,
Method:Compositional matrix adjust.,
Identities:44/131(34%), Positives:66/131(50%), Gaps:12/131(9%)

Query  94   MTTISDISNYLVEKILSRAPITCLGAVRTTCKRWNALSKENTLCNG---EARHQFLGFMT  264
            M  +SD+   L+ +ILSR P+  L +VR TCK+WN LSK+ +       EA+ + L   +
Sbjct  1    MAIMSDLPRDLLAEILSRVPLASLRSVRFTCKKWNDLSKDRSFLKKQIVEAKKKQLK--S  58

Query  265  KKYKLCSMR-FRSQWKI*rrrr*rfrGSNYKG-VR*FSESC-----*DI*CVFQCDGLLL  423
            K++++  MR FR                  KG +  FS+        ++  VF CDGLLL
Sbjct  59   KEFEVIMMRNFRVYLTSVDLHNDVNPSFTPKGTLTSFSDDANQHQVDNVSSVFHCDGLLL  118

Query  424  CVTNEDKTRLA  456
            C+T +   RL
Sbjct  119  CITKDLNFRLV  129


Query #170: XLOC_020132 Query ID: lcl|Query_33173 Length: 666

No significant similarity found.


Query #171: XLOC_020133 Query ID: lcl|Query_33174 Length: 648

Sequences producing significant alignments:
                                                                  Max    Total Query   E   Per.
Description                                                       Score  Score cover Value Ident  Accession
unknown protein [Arabidopsis thaliana]                            139    139   31%   3e-42 100.00 BAC43450.1
unnamed protein product [Arabidopsis thaliana]                    129    187   41%   5e-38 100.00 BAB02200.1
hypothetical protein AXX17_AT3G28620 [Arabidopsis thaliana]       111    111   29%   1e-31 92.06  OAP06325.1
hypothetical protein AT3G26395 [Arabidopsis thaliana]             52.8   52.8  11%   2e-09 100.00 NP_001326681.1
CYCLIN B1;5 [Arabidopsis thaliana]                                54.3   54.3  15%   5e-08 73.53  NP_001320189.1
CYCLIN B1;5 [Arabidopsis thaliana]                                54.3   54.3  15%   5e-08 73.53  NP_001319145.1
CYCLIN B1;5 [Arabidopsis thaliana]                                53.9   53.9  15%   6e-08 73.53  NP_001320191.1
CYCLIN B1;5 [Arabidopsis thaliana]                                53.9   53.9  15%   6e-08 73.53  NP_001319146.1
CYCLIN B1;5 [Arabidopsis thaliana]                                53.9   53.9  15%   6e-08 73.53  NP_001320187.1


Alignments:
>unknown protein [Arabidopsis thaliana]
Sequence ID: BAC43450.1 Length: 69
>unnamed protein product [Arabidopsis thaliana]
Sequence ID: BAD43318.1 Length: 69
Range 1: 1 to 69

Score:139 bits(349), Expect:3e-42,
Method:Compositional matrix adjust.,
Identities:69/69(100%), Positives:69/69(100%), Gaps:0/69(0%)

Query  377  MLCSLGVPFSVISVVNKKISCDVSTIFSSPDSGLLDGKVDVVKFLMMTYRHVLQLNSKVN  556
            MLCSLGVPFSVISVVNKKISCDVSTIFSSPDSGLLDGKVDVVKFLMMTYRHVLQLNSKVN
Sbjct  1    MLCSLGVPFSVISVVNKKISCDVSTIFSSPDSGLLDGKVDVVKFLMMTYRHVLQLNSKVN  60

Query  557  TCFRIRVQR  583
            TCFRIRVQR
Sbjct  61   TCFRIRVQR  69


>unnamed protein product [Arabidopsis thaliana]
Sequence ID: BAB02200.1 Length: 103
Range 1: 40 to 103

Score:129 bits(323), Expect:5e-38,
Method:Compositional matrix adjust.,
Identities:64/64(100%), Positives:64/64(100%), Gaps:0/64(0%)

Query  392  GVPFSVISVVNKKISCDVSTIFSSPDSGLLDGKVDVVKFLMMTYRHVLQLNSKVNTCFRI  571
            GVPFSVISVVNKKISCDVSTIFSSPDSGLLDGKVDVVKFLMMTYRHVLQLNSKVNTCFRI
Sbjct  40   GVPFSVISVVNKKISCDVSTIFSSPDSGLLDGKVDVVKFLMMTYRHVLQLNSKVNTCFRI  99

Query  572  RVQR  583
            RVQR
Sbjct  100  RVQR  103


Range 2: 16 to 41

Score:58.5 bits(140), Expect:8e-11,
Method:Compositional matrix adjust.,
Identities:25/26(96%), Positives:26/26(100%), Gaps:0/26(0%)

Query  2   VVFDVCRPEYFPVNSRPEWLGLLPGI  79
           VVFDVCRPEYFPVNSRPEWLGLLPG+
Sbjct  16  VVFDVCRPEYFPVNSRPEWLGLLPGV  41


>hypothetical protein AXX17_AT3G28620 [Arabidopsis thaliana]
Sequence ID: OAP06325.1 Length: 64
Range 1: 1 to 63

Score:111 bits(278), Expect:1e-31,
Method:Compositional matrix adjust.,
Identities:58/63(92%), Positives:58/63(92%), Gaps:0/63(0%)

Query  377  MLCSLGVPFSVISVVNKKISCDVSTIFSSPDSGLLDGKVDVVKFLMMTYRHVLQLNSKVN  556
            MLCSLGVPFSVISVVNKK SCDVSTIFSSPDSGLLDGKVDV KFLMMTYRHVLQLNSKV
Sbjct  1    MLCSLGVPFSVISVVNKKNSCDVSTIFSSPDSGLLDGKVDVGKFLMMTYRHVLQLNSKVI  60

Query  557  TCF  565
              F
Sbjct  61   RVF  63


>hypothetical protein AT3G26395 [Arabidopsis thaliana]
Sequence ID: NP_001326681.1 Length: 24
>hypothetical protein AT3G26395 [Arabidopsis thaliana]
Sequence ID: ANM64669.1 Length: 24 >unnamed protein product [Arabidopsis thaliana]
Sequence ID: CAA0383755.1 Length: 24 >unnamed protein product [Arabidopsis thaliana]
Sequence ID: VYS58677.1 Length: 24
Range 1: 1 to 24

Score:52.8 bits(125), Expect:2e-09,
Method:Compositional matrix adjust.,
Identities:24/24(100%), Positives:24/24(100%), Gaps:0/24(0%)

Query  512  MMTYRHVLQLNSKVNTCFRIRVQR  583
            MMTYRHVLQLNSKVNTCFRIRVQR
Sbjct  1    MMTYRHVLQLNSKVNTCFRIRVQR  24


>CYCLIN B1;5 [Arabidopsis thaliana]
Sequence ID: NP_001320189.1 Length: 486
>CYCLIN B1;5 [Arabidopsis thaliana]
Sequence ID: ANM57701.1 Length: 486
Range 1: 53 to 86

Score:54.3 bits(129), Expect:5e-08,
Method:Compositional matrix adjust.,
Identities:25/34(74%), Positives:30/34(88%), Gaps:0/34(0%)

Query  130  KLGLPDVNRLSPLSKLKDTGQKSEPLGTAIHRKI  29
            KL LPD NRLSPL++L+D GQ+S+ LGTAIHRKI
Sbjct  53   KLDLPDDNRLSPLTELEDIGQQSKTLGTAIHRKI  86


>CYCLIN B1;5 [Arabidopsis thaliana]
Sequence ID: NP_001319145.1 Length: 491
>CYCLIN B1;5 [Arabidopsis thaliana]
Sequence ID: NP_001320186.1 Length: 491 >CYCLIN B1;5 [Arabidopsis thaliana]
Sequence ID: NP_001320190.1 Length: 491 >CYCLIN B1;5 [Arabidopsis thaliana]
Sequence ID: NP_564446.3 Length: 491 >RecName: Full=Cyclin-B1-5; AltName: Full=Cyc3-At; AltName: Full=G2/mitotic-specific cyclin-B1-5; Short=CycB1;5 [Arabidopsis thaliana]
Sequence ID: Q39072.3 Length: 491 >CYCLIN B1;5 [Arabidopsis thaliana]
Sequence ID: AEE31715.1 Length: 491 >CYCLIN B1;5 [Arabidopsis thaliana]
Sequence ID: ANM57698.1 Length: 491 >CYCLIN B1;5 [Arabidopsis thaliana]
Sequence ID: ANM57702.1 Length: 491 >CYCLIN B1;5 [Arabidopsis thaliana]
Sequence ID: ANM57704.1 Length: 491
Range 1: 53 to 86

Score:54.3 bits(129), Expect:5e-08,
Method:Compositional matrix adjust.,
Identities:25/34(74%), Positives:30/34(88%), Gaps:0/34(0%)

Query  130  KLGLPDVNRLSPLSKLKDTGQKSEPLGTAIHRKI  29
            KL LPD NRLSPL++L+D GQ+S+ LGTAIHRKI
Sbjct  53   KLDLPDDNRLSPLTELEDIGQQSKTLGTAIHRKI  86


>CYCLIN B1;5 [Arabidopsis thaliana]
Sequence ID: NP_001320191.1 Length: 477
>CYCLIN B1;5 [Arabidopsis thaliana]
Sequence ID: ANM57703.1 Length: 477
Range 1: 53 to 86

Score:53.9 bits(128), Expect:6e-08,
Method:Compositional matrix adjust.,
Identities:25/34(74%), Positives:30/34(88%), Gaps:0/34(0%)

Query  130  KLGLPDVNRLSPLSKLKDTGQKSEPLGTAIHRKI  29
            KL LPD NRLSPL++L+D GQ+S+ LGTAIHRKI
Sbjct  53   KLDLPDDNRLSPLTELEDIGQQSKTLGTAIHRKI  86


>CYCLIN B1;5 [Arabidopsis thaliana]
Sequence ID: NP_001319146.1 Length: 483
>CYCLIN B1;5 [Arabidopsis thaliana]
Sequence ID: NP_001320188.1 Length: 483 >CYCLIN B1;5 [Arabidopsis thaliana]
Sequence ID: AEE31716.1 Length: 483 >CYCLIN B1;5 [Arabidopsis thaliana]
Sequence ID: ANM57700.1 Length: 483
Range 1: 53 to 86

Score:53.9 bits(128), Expect:6e-08,
Method:Compositional matrix adjust.,
Identities:25/34(74%), Positives:30/34(88%), Gaps:0/34(0%)

Query  130  KLGLPDVNRLSPLSKLKDTGQKSEPLGTAIHRKI  29
            KL LPD NRLSPL++L+D GQ+S+ LGTAIHRKI
Sbjct  53   KLDLPDDNRLSPLTELEDIGQQSKTLGTAIHRKI  86


>CYCLIN B1;5 [Arabidopsis thaliana]
Sequence ID: NP_001320187.1 Length: 498
>CYCLIN B1;5 [Arabidopsis thaliana]
Sequence ID: ANM57699.1 Length: 498
Range 1: 53 to 86

Score:53.9 bits(128), Expect:6e-08,
Method:Compositional matrix adjust.,
Identities:25/34(74%), Positives:30/34(88%), Gaps:0/34(0%)

Query  130  KLGLPDVNRLSPLSKLKDTGQKSEPLGTAIHRKI  29
            KL LPD NRLSPL++L+D GQ+S+ LGTAIHRKI
Sbjct  53   KLDLPDDNRLSPLTELEDIGQQSKTLGTAIHRKI  86


Query #172: XLOC_020134 Query ID: lcl|Query_33175 Length: 626

No significant similarity found.


Query #173: XLOC_020135 Query ID: lcl|Query_33176 Length: 714

Sequences producing significant alignments:
                                                                  Max    Total Query   E   Per.
Description                                                       Score  Score cover Value Ident  Accession
unnamed protein product [Arabidopsis thaliana]                    58.9   58.9  19%   3e-09 54.72  BAB01362.1


Alignments:
>unnamed protein product [Arabidopsis thaliana]
Sequence ID: BAB01362.1 Length: 711
Range 1: 642 to 688

Score:58.9 bits(141), Expect:3e-09,
Method:Compositional matrix adjust.,
Identities:29/53(55%), Positives:35/53(66%), Gaps:12/53(22%)

Query  502  LELSNGPSRDLWAYPYGQFFLWAKL------DGP*WTMVRKMSMPSKDHVHVG  362
            +ELSNGPSRD W YPY QFF   ++      +GP      KMSMP+KDHVH+G
Sbjct  642  IELSNGPSRDYWVYPYKQFFFMGQVGRFIMNNGP------KMSMPNKDHVHMG  688


Query #174: XLOC_020136 Query ID: lcl|Query_33177 Length: 608

No significant similarity found.


Query #175: XLOC_020137 Query ID: lcl|Query_33178 Length: 580

Sequences producing significant alignments:
                                                                  Max    Total Query   E   Per.
Description                                                       Score  Score cover Value Ident  Accession
F17F8.5 [Arabidopsis thaliana]                                    110    110   51%   2e-27 55.45  AAF98181.1
hypothetical protein [Arabidopsis thaliana]                       96.7   96.7  54%   3e-25 49.06  AAG51106.1
hypothetical protein At1G54955 [Arabidopsis thaliana]             95.5   95.5  54%   9e-25 48.11  AAT67565.1
T3P18.11 [Arabidopsis thaliana]                                   86.3   86.3  24%   1e-21 85.42  AAD43612.1
T2E6.13 [Arabidopsis thaliana]                                    84.3   84.3  52%   4e-20 46.08  AAF99801.1
non-LTR retroelement reverse transcriptase-like protein...        87.0   87.0  51%   2e-19 46.53  BAB09379.1
putative non-LTR retroelement reverse transcriptase [Arabidops... 86.3   86.3  50%   4e-19 46.94  AAC67331.1
hypothetical protein [Arabidopsis thaliana]                       83.2   83.2  52%   3e-18 46.08  AAG50886.1
putative non-LTR retroelement reverse transcriptase [Arabidops... 81.3   81.3  51%   2e-17 47.52  AAD12028.1
T23E23.16 [Arabidopsis thaliana]                                  80.9   80.9  51%   2e-17 47.52  AAF87143.1
putative protein [Arabidopsis thaliana]                           79.0   79.0  50%   1e-16 47.96  CAB39942.1
similar to reverse transcriptase (Pfam: transcript_fact.hmm,...   78.2   78.2  50%   2e-16 46.94  AAC13599.1
unnamed protein product [Arabidopsis thaliana]                    70.1   70.1  46%   3e-15 43.96  BAB10091.1
putative reverse transcriptase [Arabidopsis thaliana]             73.6   73.6  51%   7e-15 47.06  CAB45965.1
unnamed protein product [Arabidopsis thaliana]                    65.9   65.9  35%   6e-14 55.07  BAB10650.1
non-LTR retroelement reverse transcriptase-like protein...        67.4   67.4  51%   1e-12 40.59  BAB08270.1
putative non-LTR retroelement reverse transcriptase [Arabidops... 67.0   67.0  48%   2e-12 38.95  AAC63678.1
hypothetical protein AT1G21330 [Arabidopsis thaliana]             61.2   61.2  46%   2e-11 39.56  AAV68826.1
RNA-directed DNA polymerase (reverse transcriptase)-related...    61.6   61.6  51%   1e-10 36.63  NP_001327638.1
RNA-directed DNA polymerase (reverse transcriptase)-related...    61.6   61.6  51%   1e-10 36.63  NP_001327639.1
hypothetical protein [Arabidopsis thaliana]                       55.8   55.8  36%   2e-09 44.44  AAD17423.1
unnamed protein product [Arabidopsis thaliana]                    53.9   53.9  51%   6e-09 37.62  VYS61782.1
putative non-LTR reverse transcriptase [Arabidopsis thaliana]     53.1   53.1  51%   6e-08 36.63  BAE98403.1


Alignments:
>F17F8.5 [Arabidopsis thaliana]
Sequence ID: AAF98181.1 Length: 872
Range 1: 771 to 871

Score:110 bits(274), Expect:2e-27,
Method:Compositional matrix adjust.,
Identities:56/101(55%), Positives:67/101(66%), Gaps:1/101(0%)

Query  563  WAALAKGLWKSRFTTSWPQILAHVSAPHLDRVEGFLLLYAFQATVYTV-*rernerrNGP  387
            WAALAKG+WK+R++T W  +L H+S    DRVEGFL  Y FQAT+Y V       R +
Sbjct  771  WAALAKGIWKTRYSTRWSHLLTHISTHFQDRVEGFLTRYIFQATIYHVWRERNGRRHDAA  830

Query  386  PNSSAQLISWIDKQVKNQITTIRGMRDRRYDTTFQNWL*AR  264
            PN+ A +I WIDKQ +NQIT IR   DRRYD  FQ WL AR
Sbjct  831  PNTPATVIGWIDKQTRNQITIIRQSGDRRYDKAFQAWLRAR  871


>hypothetical protein [Arabidopsis thaliana]
Sequence ID: AAG51106.1 Length: 137
>hypothetical protein At1g54955 [Arabidopsis thaliana]
Sequence ID: AAX23786.1 Length: 137
Range 1: 32 to 137

Score:96.7 bits(239), Expect:3e-25,
Method:Compositional matrix adjust.,
Identities:52/106(49%), Positives:74/106(69%), Gaps:1/106(0%)

Query  575  IRNCWAALAKGLWKSRFTTSWPQILAHVSAPHLDRVEGFLLLYAFQATVYTV*rernerr  396
            +   WA +AKG++K +++T W Q+L HVS  +  ++E FL   AFQATVYT+  ERN RR
Sbjct  32   VSEIWAVIAKGIFKDKYSTDWSQLLDHVSNANQTQLESFLSRSAFQATVYTIWGERNGRR  91

Query  395  NG-PPNSSAQLISWIDKQVKNQITTIRGMRDRRYDTTFQNWL*ARV  261
            +G  P+S+  L +WIDK ++N+I+TIR M DRRYD  +Q WL +R
Sbjct  92   HGEEPHSTTLLTTWIDKHIRNKISTIRRMGDRRYDEAYQAWLISRT  137


>hypothetical protein At1G54955 [Arabidopsis thaliana]
Sequence ID: AAT67565.1 Length: 137
Range 1: 32 to 137

Score:95.5 bits(236), Expect:9e-25,
Method:Compositional matrix adjust.,
Identities:51/106(48%), Positives:74/106(69%), Gaps:1/106(0%)

Query  575  IRNCWAALAKGLWKSRFTTSWPQILAHVSAPHLDRVEGFLLLYAFQATVYTV*rernerr  396
            +   WA +AKG++K +++T W Q+L HVS  +  ++E +L   AFQATVYT+  ERN RR
Sbjct  32   VSEIWAVIAKGIFKDKYSTDWSQLLDHVSNANQTQLESYLSRSAFQATVYTIWGERNGRR  91

Query  395  NG-PPNSSAQLISWIDKQVKNQITTIRGMRDRRYDTTFQNWL*ARV  261
            +G  P+S+  L +WIDK ++N+I+TIR M DRRYD  +Q WL +R
Sbjct  92   HGEEPHSTTLLTTWIDKHIRNKISTIRRMGDRRYDEAYQAWLISRT  137


>T3P18.11 [Arabidopsis thaliana]
Sequence ID: AAD43612.1 Length: 103
Range 1: 38 to 85

Score:86.3 bits(212), Expect:1e-21,
Method:Compositional matrix adjust.,
Identities:41/48(85%), Positives:43/48(89%), Gaps:0/48(0%)

Query  563  WAALAKGLWKSRFTTSWPQILAHVSAPHLDRVEGFLLLYAFQATVYTV  420
            WAALAKGLWKSR+TTS PQILAHVSAP  DR+EGFLL Y FQATVYTV
Sbjct  38   WAALAKGLWKSRYTTSCPQILAHVSAPDQDRLEGFLLRYVFQATVYTV  85


>T2E6.13 [Arabidopsis thaliana]
Sequence ID: AAF99801.1 Length: 165
Range 1: 64 to 165

Score:84.3 bits(207), Expect:4e-20,
Method:Compositional matrix adjust.,
Identities:47/102(46%), Positives:67/102(65%), Gaps:1/102(0%)

Query  563  WAALAKGLWKSRFTTSWPQILAHVSAPHLDRVEGFLLLYAFQATVYTV*rernerrNG-P  387
            W A+AK + + RF+T W  I+ ++S    DR+  FL  Y FQ TV+TV +ERN+RR+G
Sbjct  64   WTAIAKNVLQHRFSTDWQAIVNYISETQTDRIRSFLSRYIFQLTVHTVWKERNDRRHGEE  123

Query  386  PNSSAQLISWIDKQVKNQITTIRGMRDRRYDTTFQNWL*ARV  261
            P +SA LISW+DKQ++NQ++ I    DRRY+   Q W   R+
Sbjct  124  PRTSANLISWMDKQIRNQLSIIISTGDRRYENGLQVWFSTRL  165


>non-LTR retroelement reverse transcriptase-like protein [Arabidopsis thaliana]
Sequence ID: BAB09379.1 Length: 1223
Range 1: 1122 to 1222

Score:87.0 bits(214), Expect:2e-19,
Method:Compositional matrix adjust.,
Identities:47/101(47%), Positives:71/101(70%), Gaps:1/101(0%)

Query  563   WAALAKGLWKSRFTTSWPQILAHVSAPHLDRVEGFLLLYAFQATVYTV*rernerrNG-P  387
             W  LA+G++K+++T+ W  I+  ++     RVE FL  Y FQAT+Y V RERN RR+G P
Sbjct  1122  WVDLARGIFKTQYTSHWQSIIEAITNSQHHRVEWFLRRYVFQATIYIVWRERNGRRHGEP  1181

Query  386   PNSSAQLISWIDKQVKNQITTIRGMRDRRYDTTFQNWL*AR  264
             PN+++QL+ WIDKQ++NQ+++I    D+RYD + Q W  +R
Sbjct  1182  PNTASQLVGWIDKQIRNQLSSICLKGDKRYDGSLQVWFQSR  1222


>putative non-LTR retroelement reverse transcriptase [Arabidopsis thaliana]
Sequence ID: AAC67331.1 Length: 1449
Range 1: 1348 to 1445

Score:86.3 bits(212), Expect:4e-19,
Method:Composition-based stats.,
Identities:46/98(47%), Positives:65/98(66%), Gaps:1/98(1%)

Query  563   WAALAKGLWKSRFTTSWPQILAHVSAPHLDRVEGFLLLYAFQATVYTV*rernerrNG-P  387
             W   AK ++KS ++T W  +L  +     DRVE F+  Y FQ +VY + RERN R++G
Sbjct  1348  WEMSAKSIYKSSYSTDWRHLLTKIQENWKDRVESFIARYIFQVSVYAIWRERNRRKHGRN  1407

Query  386   PNSSAQLISWIDKQVKNQITTIRGMRDRRYDTTFQNWL  273
             PN +AQL+ WIDKQ+++Q++ I+ M DRRYD  FQ WL
Sbjct  1408  PNQAAQLVRWIDKQIRDQLSAIKLMGDRRYDQGFQFWL  1445


>hypothetical protein [Arabidopsis thaliana]
Sequence ID: AAG50886.1 Length: 629
Range 1: 528 to 629

Score:83.2 bits(204), Expect:3e-18,
Method:Compositional matrix adjust.,
Identities:47/102(46%), Positives:67/102(65%), Gaps:1/102(0%)

Query  563  WAALAKGLWKSRFTTSWPQILAHVSAPHLDRVEGFLLLYAFQATVYTV*rernerrNG-P  387
            W A+AK + + RF+T W  I+ ++S    DR+  FL  Y FQ TV+TV +ERN+RR+G
Sbjct  528  WTAIAKNVLQHRFSTDWQTIVNYISETQTDRIRSFLSRYIFQLTVHTVWKERNDRRHGEE  587

Query  386  PNSSAQLISWIDKQVKNQITTIRGMRDRRYDTTFQNWL*ARV  261
            P +SA LISW+DKQ++NQ++ I    DRRY+   Q W   R+
Sbjct  588  PRTSANLISWMDKQIRNQLSIIISTGDRRYENGLQVWFSTRL  629


>putative non-LTR retroelement reverse transcriptase [Arabidopsis thaliana]
Sequence ID: AAD12028.1 Length: 1447
Range 1: 1346 to 1446

Score:81.3 bits(199), Expect:2e-17,
Method:Composition-based stats.,
Identities:48/101(48%), Positives:65/101(64%), Gaps:1/101(0%)

Query  563   WAALAKGLWKSRFTTSWPQILAHVSAPHLDRVEGFLLLYAFQATVYTV*rernerrNGPP  384
             W   AK +  +RFTT W  +L  VS+   +RVE FL+ YAFQA+VY++  ERN RR+G
Sbjct  1346  WTVTAKNILTTRFTTDWHLLLNTVSSLQQNRVENFLVRYAFQASVYSIWSERNRRRHGNT  1405

Query  383   -NSSAQLISWIDKQVKNQITTIRGMRDRRYDTTFQNWL*AR  264
              +S+ +LI WIDKQV+N +++IR   DR YD   Q W   R
Sbjct  1406  LHSATRLIGWIDKQVRNILSSIRIKGDRHYDAGLQMWFDTR  1446


>T23E23.16 [Arabidopsis thaliana]
Sequence ID: AAF87143.1 Length: 653
Range 1: 552 to 652

Score:80.9 bits(198), Expect:2e-17,
Method:Compositional matrix adjust.,
Identities:48/101(48%), Positives:65/101(64%), Gaps:1/101(0%)

Query  563  WAALAKGLWKSRFTTSWPQILAHVSAPHLDRVEGFLLLYAFQATVYTV*rernerrNGP-  387
            W  LAK ++K++F+T+W  IL  VS    +R E FL  Y FQAT++T+  ERN RR+G
Sbjct  552  WENLAKNIYKAKFSTNWSTILTSVSTTWRNRTESFLARYIFQATIHTIWHERNGRRHGER  611

Query  386  PNSSAQLISWIDKQVKNQITTIRGMRDRRYDTTFQNWL*AR  264
             NS+  LI W+DKQ++NQI+TI    D RYD   Q W  +R
Sbjct  612  SNSATHLIWWLDKQMRNQISTIAASGDHRYDKPLQLWFQSR  652


>putative protein [Arabidopsis thaliana]
Sequence ID: CAB39942.1 Length: 473
>putative protein [Arabidopsis thaliana]
Sequence ID: CAB78214.1 Length: 473
Range 1: 371 to 468

Score:79.0 bits(193), Expect:1e-16,
Method:Compositional matrix adjust.,
Identities:47/98(48%), Positives:64/98(65%), Gaps:1/98(1%)

Query  563  WAALAKGLWKSRFTTSWPQILAHVSAPHLDRVEGFLLLYAFQATVYTV*rernerrNG-P  387
            W  LAK ++ + F T W  I+ +VS    DR+ GFL     Q T+YT+ RERNER++G
Sbjct  371  WEPLAKTIYNTCFYTDWQTIINNVSRNWPDRIAGFLARCILQVTIYTLWRERNERKHGAS  430

Query  386  PNSSAQLISWIDKQVKNQITTIRGMRDRRYDTTFQNWL  273
            PNSS++LISWIDK ++N +  I+   DRR+D  FQ WL
Sbjct  431  PNSSSRLISWIDKHIRNHLMAIKQSGDRRFDRGFQVWL  468


>similar to reverse transcriptase (Pfam: transcript_fact.hmm, score: 72.31) [Arabidopsis thaliana]
Sequence ID: AAC13599.1 Length: 928
Range 1: 827 to 924

Score:78.2 bits(191), Expect:2e-16,
Method:Composition-based stats.,
Identities:46/98(47%), Positives:61/98(62%), Gaps:1/98(1%)

Query  563  WAALAKGLWKSRFTTSWPQILAHVSAPHLDRVEGFLLLYAFQATVYTV*rernerrNG-P  387
            W ALAK L +  +TT W  I+++VS    DRV  FL     QA+VYT+ RERN RR+G
Sbjct  827  WEALAKNLLQRSYTTDWQTIISYVSGQCHDRVSCFLARSVLQASVYTIWRERNGRRHGET  886

Query  386  PNSSAQLISWIDKQVKNQITTIRGMRDRRYDTTFQNWL  273
            PN +A+LI WIDK ++N ++ I    D+RYD   Q W
Sbjct  887  PNPAARLIQWIDKHIRNMLSVIHQKGDKRYDKGLQMWF  924


>unnamed protein product [Arabidopsis thaliana]
Sequence ID: BAB10091.1 Length: 116
Range 1: 26 to 116

Score:70.1 bits(170), Expect:3e-15,
Method:Compositional matrix adjust.,
Identities:40/91(44%), Positives:57/91(62%), Gaps:1/91(1%)

Query  563  WAALAKGLWKSRFTTSWPQILAHVSAPHLDRVEGFLLLYAFQATVYTV*rernerrNGP-  387
            W ALAK + K  +TT W  ++  VS    + +EGFL     QA VYT+ RERN RR+G
Sbjct  26   WEALAKNILKRSYTTDWHNLIISVSDQRHNTIEGFLARSVLQALVYTIWRERNVRRHGES  85

Query  386  PNSSAQLISWIDKQVKNQITTIRGMRDRRYD  294
            P  +A+ I WI+K ++N  + I+ MRD++YD
Sbjct  86   PKPAARFIQWINKHIRNTFSVIKRMRDKKYD  116


>putative reverse transcriptase [Arabidopsis thaliana]
Sequence ID: CAB45965.1 Length: 662
>putative reverse transcriptase [Arabidopsis thaliana]
Sequence ID: CAB78261.1 Length: 662
Range 1: 560 to 661

Score:73.6 bits(179), Expect:7e-15,
Method:Compositional matrix adjust.,
Identities:48/102(47%), Positives:64/102(62%), Gaps:2/102(1%)

Query  563  WAALAKGLWKSRFTTSWPQILAHVSAP-HLDRVEGFLLLYAFQATVYTV*rernerrNGP  387
            W ALA  ++ +++TT W  ++A  S     DR  GF+  Y FQA VYT+ RERN RR+G
Sbjct  560  WQALAWNIYGAKYTTHWNDLIAATSGHWQDDRTTGFIARYVFQAAVYTIWRERNNRRHGE  619

Query  386  -PNSSAQLISWIDKQVKNQITTIRGMRDRRYDTTFQNWL*AR  264
             PNS  +LI WIDKQV+++I+ +    DRRY T  Q W  AR
Sbjct  620  LPNSPVRLIRWIDKQVRDRISLLNTSGDRRYVTGLQVWFAAR  661


>unnamed protein product [Arabidopsis thaliana]
Sequence ID: BAB10650.1 Length: 76
Range 1: 4 to 72

Score:65.9 bits(159), Expect:6e-14,
Method:Compositional matrix adjust.,
Identities:38/69(55%), Positives:51/69(73%), Gaps:1/69(1%)

Query  476  DRVEGFLLLYAFQATVYTV*rernerrNGP-PNSSAQLISWIDKQVKNQITTIRGMRDRR  300
            +R+E FL  Y  QA+VYT+ RERN RR+G  PN +A+LI WIDKQ++N +++I+ M DRR
Sbjct  4    NRIERFLARYILQASVYTLWRERNGRRHGEDPNPAARLIRWIDKQMRNALSSIKEMGDRR  63

Query  299  YDTTFQNWL  273
            YDT  Q W
Sbjct  64   YDTGLQLWF  72


>non-LTR retroelement reverse transcriptase-like protein [Arabidopsis thaliana]
Sequence ID: BAB08270.1 Length: 489
Range 1: 386 to 486

Score:67.4 bits(163), Expect:1e-12,
Method:Compositional matrix adjust.,
Identities:41/101(41%), Positives:62/101(61%), Gaps:1/101(0%)

Query  563  WAALAKGLWKSRFTTSWPQILAHVSAPHLDRVEGFLLLYAFQATVYTV*rernerrNGPP  384
            W  LA  ++ + ++T W  I+  VS    DR  GFL     Q +++T+ RERNER++G
Sbjct  386  WEPLAATIYNTSYSTDWQTIINTVSTNWHDRTAGFLARCLLQVSIHTIWRERNERKHGAS  445

Query  383  -NSSAQLISWIDKQVKNQITTIRGMRDRRYDTTFQNWL*AR  264
             N +++L+ WIDK ++N + TI+   DRR+D  FQ WL AR
Sbjct  446  LNPASRLVRWIDKHIRNHLLTIKQSGDRRFDKGFQVWLQAR  486


>putative non-LTR retroelement reverse transcriptase [Arabidopsis thaliana]
Sequence ID: AAC63678.1 Length: 1216
Range 1: 844 to 938

Score:67.0 bits(162), Expect:2e-12,
Method:Composition-based stats.,
Identities:37/95(39%), Positives:61/95(64%), Gaps:1/95(1%)

Query  563  WAALAKGLWKSRFTTSWPQILAHVSAPHLDRVEGFLLLYAFQATVYTV*rernerrNGPP  384
            W ++AK ++K RF+T W  ++ ++S    DR++ FL  Y FQ +++++ RERN RR+G
Sbjct  844  WTSIAKNVYKDRFSTKWSAVVNYISDSQPDRIQSFLSRYTFQVSIHSIWRERNSRRHGEK  903

Query  383  NSSA-QLISWIDKQVKNQITTIRGMRDRRYDTTFQ  282
            + SA  LI  IDK ++NQ++TI+   D R +   Q
Sbjct  904  SRSASNLIRQIDKTIRNQLSTIKKKGDLRLEKGLQ  938


>hypothetical protein AT1G21330 [Arabidopsis thaliana]
Sequence ID: AAV68826.1 Length: 152
>hypothetical protein At1g21330 [Arabidopsis thaliana]
Sequence ID: AAX23753.1 Length: 152
Range 1: 51 to 141

Score:61.2 bits(147), Expect:2e-11,
Method:Compositional matrix adjust.,
Identities:36/91(40%), Positives:57/91(62%), Gaps:1/91(1%)

Query  563  WAALAKGLWKSRFTTSWPQILAHVSAPHLDRVEGFLLLYAFQATVYTV*rernerrNGP-  387
            W  L++ L   RF+T W  I+  ++   L +   FL+ Y FQ TV+++ +ERN  R+G
Sbjct  51   WTGLSRKLLSQRFSTDWDMIIKLITDTSLGKGCLFLVRYTFQLTVHSIWKERNGHRHGEA  110

Query  386  PNSSAQLISWIDKQVKNQITTIRGMRDRRYD  294
            P  S+QL   +DKQ +N+I++IR + DRRY+
Sbjct  111  PIPSSQLTRRLDKQTRNRISSIRELGDRRYE  141


>RNA-directed DNA polymerase (reverse transcriptase)-related family protein [Arabidopsis thaliana]
Sequence ID: NP_001327638.1 Length: 478
>RNA-directed DNA polymerase (reverse transcriptase)-related family protein [Arabidopsis thaliana]
Sequence ID: ANM65688.1 Length: 478 >hypothetical protein [Arabidopsis thaliana]
Sequence ID: BAD95408.1 Length: 478
Range 1: 378 to 478

Score:61.6 bits(148), Expect:1e-10,
Method:Compositional matrix adjust.,
Identities:37/101(37%), Positives:60/101(59%), Gaps:1/101(0%)

Query  563  WAALAKGLWKSRFTTSWPQILAHVSAPHLDRVEGFLLLYAFQATVYTV*rernerrNGP-  387
            W+ L + L    FT  W  IL  ++   L     FL  Y FQ T++++ +ERN RR+G
Sbjct  378  WSTLTRKLLSQHFTNRWEAILKLLTNKSLGHEVPFLTRYTFQLTLHSLWKERNGRRHGEV  437

Query  386  PNSSAQLISWIDKQVKNQITTIRGMRDRRYDTTFQNWL*AR  264
            P ++AQ++ ++DKQV+N+I++I+   DRRY+     W  +R
Sbjct  438  PQAAAQMVRFLDKQVRNRISSIQSQEDRRYNGCMTCWFGSR  478


>RNA-directed DNA polymerase (reverse transcriptase)-related family protein [Arabidopsis thaliana]
Sequence ID: NP_001327639.1 Length: 532
>RNA-directed DNA polymerase (reverse transcriptase)-related family protein [Arabidopsis thaliana]
Sequence ID: NP_001327640.1 Length: 532 >RNA-directed DNA polymerase (reverse transcriptase)-related family protein [Arabidopsis thaliana]
Sequence ID: ANM65689.1 Length: 532 >RNA-directed DNA polymerase (reverse transcriptase)-related family protein [Arabidopsis thaliana]
Sequence ID: ANM65690.1 Length: 532
Range 1: 432 to 532

Score:61.6 bits(148), Expect:1e-10,
Method:Compositional matrix adjust.,
Identities:37/101(37%), Positives:60/101(59%), Gaps:1/101(0%)

Query  563  WAALAKGLWKSRFTTSWPQILAHVSAPHLDRVEGFLLLYAFQATVYTV*rernerrNGP-  387
            W+ L + L    FT  W  IL  ++   L     FL  Y FQ T++++ +ERN RR+G
Sbjct  432  WSTLTRKLLSQHFTNRWEAILKLLTNKSLGHEVPFLTRYTFQLTLHSLWKERNGRRHGEV  491

Query  386  PNSSAQLISWIDKQVKNQITTIRGMRDRRYDTTFQNWL*AR  264
            P ++AQ++ ++DKQV+N+I++I+   DRRY+     W  +R
Sbjct  492  PQAAAQMVRFLDKQVRNRISSIQSQEDRRYNGCMTCWFGSR  532


>hypothetical protein [Arabidopsis thaliana]
Sequence ID: AAD17423.1 Length: 166
Range 1: 92 to 163

Score:55.8 bits(133), Expect:2e-09,
Method:Compositional matrix adjust.,
Identities:32/72(44%), Positives:49/72(68%), Gaps:1/72(1%)

Query  476  DRVEGFLLLYAFQATVYTV*rernerrNGPP-NSSAQLISWIDKQVKNQITTIRGMRDRR  300
            D+V+G L  Y  QAT+YT+ RERNER++G   NS+ +LI WID+ ++NQ++ ++   D R
Sbjct  92   DKVDGLLARYVLQATIYTIWRERNERKHGTGLNSANRLIKWIDRHIRNQLSVLKISGDHR  151

Query  299  YDTTFQNWL*AR  264
            Y+   Q W  +R
Sbjct  152  YNKGLQVWFTSR  163


>unnamed protein product [Arabidopsis thaliana]
Sequence ID: VYS61782.1 Length: 136
Range 1: 36 to 136

Score:53.9 bits(128), Expect:6e-09,
Method:Compositional matrix adjust.,
Identities:38/101(38%), Positives:57/101(56%), Gaps:1/101(0%)

Query  563  WAALAKGLWKSRFTTSWPQILAHVSAPHLDRVEGFLLLYAFQATVYTV*rernerrNGPP  384
            W AL   L   +FTT W  I+  ++   L     FL  Y FQ T++++ +ERN RR+G
Sbjct  36   WTALTCKLLTHKFTTYWDSIIKLLTDRSLGLERLFLTRYTFQLTLHSIWKERNGRRHGEM  95

Query  383  N-SSAQLISWIDKQVKNQITTIRGMRDRRYDTTFQNWL*AR  264
            + SSAQ I  +DK  +N++++I    D+RYD   Q W  +R
Sbjct  96   HASSAQQIRVLDKPFRNRVSSIGEQGDKRYDACMQVWFGSR  136


>putative non-LTR reverse transcriptase [Arabidopsis thaliana]
Sequence ID: BAE98403.1 Length: 278
Range 1: 177 to 277

Score:53.1 bits(126), Expect:6e-08,
Method:Compositional matrix adjust.,
Identities:37/101(37%), Positives:56/101(55%), Gaps:1/101(0%)

Query  563  WAALAKGLWKSRFTTSWPQILAHVSAPHLDRVEGFLLLYAFQATVYTV*rernerrNGPP  384
            W ALAKGL   R  +SW  I  H+       +  F L Y FQA ++++ RERN RR+G P
Sbjct  177  WIALAKGLLNGRNVSSWSLITPHLLDSSQPYLHVFTLRYTFQALIHSLWRERNGRRHGEP  236

Query  383  N-SSAQLISWIDKQVKNQITTIRGMRDRRYDTTFQNWL*AR  264
               +++L   IDK ++N+ +T++ M ++R     Q W   R
Sbjct  237  AIPASKLTKLIDKNIRNRFSTLQKMGNKRLQGGLQYWFQTR  277


Query #176: XLOC_020138 Query ID: lcl|Query_33179 Length: 374

No significant similarity found.


Query #177: XLOC_020139 Query ID: lcl|Query_33180 Length: 413

No significant similarity found.


Query #178: XLOC_020140 Query ID: lcl|Query_33181 Length: 1493

No significant similarity found.


Query #179: XLOC_020141 Query ID: lcl|Query_33182 Length: 487

No significant similarity found.


Query #180: XLOC_020142 Query ID: lcl|Query_33183 Length: 1125

No significant similarity found.


Query #181: XLOC_020143 Query ID: lcl|Query_33184 Length: 1538

Sequences producing significant alignments:
                                                                  Max    Total Query   E   Per.
Description                                                       Score  Score cover Value Ident  Accession
unnamed protein product [Arabidopsis thaliana]                    174    174   30%   2e-47 100.00 BAB01195.1
unnamed protein product [Arabidopsis thaliana]                    97.1   97.1  13%   2e-23 92.65  CAA0269411.1
unnamed protein product [Arabidopsis thaliana]                    97.1   97.1  13%   2e-23 92.65  VYS48116.1
hypothetical protein AT3G14670 [Arabidopsis thaliana]             93.6   93.6  26%   5e-21 83.33  NP_001118627.1
hypothetical protein AT3G14670 [Arabidopsis thaliana]             92.8   92.8  26%   1e-20 83.94  NP_001327682.1
hypothetical protein AT3G14670 [Arabidopsis thaliana]             92.4   92.4  26%   1e-20 83.94  NP_188085.1
unknown [Arabidopsis thaliana]                                    92.4   92.4  26%   2e-20 83.94  ABK28558.1
unnamed protein product [Arabidopsis thaliana]                    66.2   66.2  12%   3e-12 81.25  VYS52273.1
unknown protein [Arabidopsis thaliana]                            43.5   104   12%   5e-10 68.75  ABF59358.1
hypothetical protein AXX17_AT3G35460 [Arabidopsis thaliana]       52.4   52.4  12%   9e-08 63.16  OAP01399.1
unnamed protein product [Arabidopsis thaliana]                    43.5   77.8  10%   9e-08 68.75  VYS60963.1
unnamed protein product [Arabidopsis thaliana]                    52.4   52.4  12%   9e-08 59.49  VYS59161.1


Alignments:
>unnamed protein product [Arabidopsis thaliana]
Sequence ID: BAB01195.1 Length: 552
Range 1: 1 to 154

Score:174 bits(441), Expect:2e-47,
Method:Compositional matrix adjust.,
Identities:154/154(100%), Positives:154/154(100%), Gaps:0/154(0%)

Query  466  MAPRTIPPLKKRKRGVAGGTKPIDVVTKAtteppttteePSATEQNPVAIEGGVEEEPIP  287
            MAPRTIPPLKKRKRGVAGGTKPIDVVTKATTEPPTTTEEPSATEQNPVAIEGGVEEEPIP
Sbjct  1    MAPRTIPPLKKRKRGVAGGTKPIDVVTKATTEPPTTTEEPSATEQNPVAIEGGVEEEPIP  60

Query  286  TIIPTVVeegeqsdnneeensekdeneeseeeeseeeekeeeekeeeekeeEGNVTGGES  107
            TIIPTVVEEGEQSDNNEEENSEKDENEESEEEESEEEEKEEEEKEEEEKEEEGNVTGGES
Sbjct  61   TIIPTVVEEGEQSDNNEEENSEKDENEESEEEESEEEEKEEEEKEEEEKEEEGNVTGGES  120

Query  106  SNDSSTTLGEEYSSDKNMNDETAVENQVEIPASM  5
            SNDSSTTLGEEYSSDKNMNDETAVENQVEIPASM
Sbjct  121  SNDSSTTLGEEYSSDKNMNDETAVENQVEIPASM  154


>unnamed protein product [Arabidopsis thaliana]
Sequence ID: CAA0269411.1 Length: 116
Range 1: 1 to 68

Score:97.1 bits(240), Expect:2e-23,
Method:Compositional matrix adjust.,
Identities:63/68(93%), Positives:64/68(94%), Gaps:1/68(1%)

Query  466  MAPRTIPPLKKRKRGVAGGTKPIDVVTKAtteppttteePSATEQNPVAIEGG-VEEEPI  290
            MAPRTIPPLKKRKRGVAGGTKPID VTKAT EPPTTTE+PSATEQN VAIEGG VEEEPI
Sbjct  1    MAPRTIPPLKKRKRGVAGGTKPIDAVTKATKEPPTTTEKPSATEQNLVAIEGGRVEEEPI  60

Query  289  PTIIPTVV  266
            PTIIPTVV
Sbjct  61   PTIIPTVV  68


>unnamed protein product [Arabidopsis thaliana]
Sequence ID: VYS48116.1 Length: 116
Range 1: 1 to 68

Score:97.1 bits(240), Expect:2e-23,
Method:Compositional matrix adjust.,
Identities:63/68(93%), Positives:64/68(94%), Gaps:1/68(1%)

Query  466  MAPRTIPPLKKRKRGVAGGTKPIDVVTKAtteppttteePSATEQNPVAIEGG-VEEEPI  290
            MAPRTIPPLKKRKRGVAGGTKPID VTKAT EPPTTTE+PSATEQN VAIEGG VEEEPI
Sbjct  1    MAPRTIPPLKKRKRGVAGGTKPIDAVTKATKEPPTTTEKPSATEQNLVAIEGGRVEEEPI  60

Query  289  PTIIPTVV  266
            PTIIPTVV
Sbjct  61   PTIIPTVV  68


>hypothetical protein AT3G14670 [Arabidopsis thaliana]
Sequence ID: NP_001118627.1 Length: 231
>hypothetical protein AT3G14670 [Arabidopsis thaliana]
Sequence ID: AEE75554.1 Length: 231
Range 1: 19 to 156

Score:93.6 bits(231), Expect:5e-21,
Method:Compositional matrix adjust.,
Identities:115/138(83%), Positives:125/138(90%), Gaps:1/138(0%)

Query  412  GTKPIDVVTKAtteppttteePSATEQNPVAIEG-GVEEEPIPTIIPTVVeegeqsdnne  236
            GTKPID VTKATTEPP TTEEPSA++QNPV IEG GVEEE IPTII TVVEEGE+SDNNE
Sbjct  19   GTKPIDAVTKATTEPPMTTEEPSASKQNPVVIEGRGVEEEQIPTIITTVVEEGEKSDNNE  78

Query  235  eensekdeneeseeeeseeeekeeeekeeeekeeEGNVTGGESSNDSSTTLGEEYSSDKN  56
            EENSEKDE EESEEEESEEEEKEEEEKEEEEKEEEGNV GG SS+DSS TLG+E SSD+N
Sbjct  79   EENSEKDEKEESEEEESEEEEKEEEEKEEEEKEEEGNVAGGGSSDDSSRTLGKESSSDEN  138

Query  55   MNDETAVENQVEIPASMV  2
            M+DETAV  QV+IPA+M+
Sbjct  139  MDDETAVGKQVDIPAAMI  156


>hypothetical protein AT3G14670 [Arabidopsis thaliana]
Sequence ID: NP_001327682.1 Length: 237
>hypothetical protein AT3G14670 [Arabidopsis thaliana]
Sequence ID: ANM65735.1 Length: 237 >unnamed protein product [Arabidopsis thaliana]
Sequence ID: BAB02399.1 Length: 237
Range 1: 24 to 160

Score:92.8 bits(229), Expect:1e-20,
Method:Compositional matrix adjust.,
Identities:115/137(84%), Positives:124/137(90%), Gaps:1/137(0%)

Query  412  GTKPIDVVTKAtteppttteePSATEQNPVAIEG-GVEEEPIPTIIPTVVeegeqsdnne  236
            GTKPID VTKATTEPP TTEEPSA++QNPV IEG GVEEE IPTII TVVEEGE+SDNNE
Sbjct  24   GTKPIDAVTKATTEPPMTTEEPSASKQNPVVIEGRGVEEEQIPTIITTVVEEGEKSDNNE  83

Query  235  eensekdeneeseeeeseeeekeeeekeeeekeeEGNVTGGESSNDSSTTLGEEYSSDKN  56
            EENSEKDE EESEEEESEEEEKEEEEKEEEEKEEEGNV GG SS+DSS TLG+E SSD+N
Sbjct  84   EENSEKDEKEESEEEESEEEEKEEEEKEEEEKEEEGNVAGGGSSDDSSRTLGKESSSDEN  143

Query  55   MNDETAVENQVEIPASM  5
            M+DETAV  QV+IPA+M
Sbjct  144  MDDETAVGKQVDIPAAM  160


>hypothetical protein AT3G14670 [Arabidopsis thaliana]
Sequence ID: NP_188085.1 Length: 232
>hypothetical protein At3g14670 [Arabidopsis thaliana]
Sequence ID: ABE65941.1 Length: 232 >hypothetical protein AT3G14670 [Arabidopsis thaliana]
Sequence ID: AEE75553.1 Length: 232
Range 1: 19 to 155

Score:92.4 bits(228), Expect:1e-20,
Method:Compositional matrix adjust.,
Identities:115/137(84%), Positives:124/137(90%), Gaps:1/137(0%)

Query  412  GTKPIDVVTKAtteppttteePSATEQNPVAIEG-GVEEEPIPTIIPTVVeegeqsdnne  236
            GTKPID VTKATTEPP TTEEPSA++QNPV IEG GVEEE IPTII TVVEEGE+SDNNE
Sbjct  19   GTKPIDAVTKATTEPPMTTEEPSASKQNPVVIEGRGVEEEQIPTIITTVVEEGEKSDNNE  78

Query  235  eensekdeneeseeeeseeeekeeeekeeeekeeEGNVTGGESSNDSSTTLGEEYSSDKN  56
            EENSEKDE EESEEEESEEEEKEEEEKEEEEKEEEGNV GG SS+DSS TLG+E SSD+N
Sbjct  79   EENSEKDEKEESEEEESEEEEKEEEEKEEEEKEEEGNVAGGGSSDDSSRTLGKESSSDEN  138

Query  55   MNDETAVENQVEIPASM  5
            M+DETAV  QV+IPA+M
Sbjct  139  MDDETAVGKQVDIPAAM  155


>unknown, partial [Arabidopsis thaliana]
Sequence ID: ABK28558.1 Length: 233
Range 1: 19 to 155

Score:92.4 bits(228), Expect:2e-20,
Method:Compositional matrix adjust.,
Identities:115/137(84%), Positives:124/137(90%), Gaps:1/137(0%)

Query  412  GTKPIDVVTKAtteppttteePSATEQNPVAIEG-GVEEEPIPTIIPTVVeegeqsdnne  236
            GTKPID VTKATTEPP TTEEPSA++QNPV IEG GVEEE IPTII TVVEEGE+SDNNE
Sbjct  19   GTKPIDAVTKATTEPPMTTEEPSASKQNPVVIEGRGVEEEQIPTIITTVVEEGEKSDNNE  78

Query  235  eensekdeneeseeeeseeeekeeeekeeeekeeEGNVTGGESSNDSSTTLGEEYSSDKN  56
            EENSEKDE EESEEEESEEEEKEEEEKEEEEKEEEGNV GG SS+DSS TLG+E SSD+N
Sbjct  79   EENSEKDEKEESEEEESEEEEKEEEEKEEEEKEEEGNVAGGGSSDDSSRTLGKESSSDEN  138

Query  55   MNDETAVENQVEIPASM  5
            M+DETAV  QV+IPA+M
Sbjct  139  MDDETAVGKQVDIPAAM  155


>unnamed protein product [Arabidopsis thaliana]
Sequence ID: VYS52273.1 Length: 155
Range 1: 31 to 94

Score:66.2 bits(160), Expect:3e-12,
Method:Compositional matrix adjust.,
Identities:52/64(81%), Positives:58/64(90%), Gaps:0/64(0%)

Query  524  LIFSFYNPNFPAIFSNPNPNGaennssskeeeerssrrnkanRCCHEGNNGASDNDRGAI  345
            LIFSF+NP FP IFSNPNPNG ENN+SSKEEEE+SSRRNKANRCC++GN+GASD D GAI
Sbjct  31   LIFSFFNPKFPVIFSNPNPNGTENNASSKEEEEKSSRRNKANRCCYKGNDGASDKDEGAI  90

Query  344  SDGA  333
            SDG
Sbjct  91   SDGT  94


>unknown protein [Arabidopsis thaliana]
Sequence ID: ABF59358.1 Length: 150
Range 1: 1 to 32

Score:43.5 bits(101), Expect:5e-10,
Method:Compositional matrix adjust.,
Identities:22/32(69%), Positives:26/32(81%), Gaps:0/32(0%)

Query  1195  MMRYINVYLRQVQFVTRSNSIKPVSFGIVSLK  1290
             M+R INVYL+QVQ V   +SIKPVSFGIV L+
Sbjct  1     MLRDINVYLKQVQPVIGPSSIKPVSFGIVFLE  32


Range 2: 30 to 54

Score:34.3 bits(77), Expect:5e-10,
Method:Compositional matrix adjust.,
Identities:15/25(60%), Positives:18/25(72%), Gaps:0/25(0%)

Query  1281  FLETPDEISYDIPIYRFPRDSAFQR  1355
             FLE PDE+  DIPIYR PR S  ++
Sbjct  30    FLENPDELGNDIPIYRSPRVSFLKK  54


Range 3: 46 to 63

Score:26.6 bits(57), Expect:5e-10,
Method:Compositional matrix adjust.,
Identities:14/18(78%), Positives:14/18(77%), Gaps:0/18(0%)

Query  1331  SPRFSFPKKPTLPRSERS  1384
             SPR SF KKPT PRSE S
Sbjct  46    SPRVSFLKKPTSPRSEMS  63


>hypothetical protein AXX17_AT3G35460 [Arabidopsis thaliana]
Sequence ID: OAP01399.1 Length: 113
Range 1: 1 to 75

Score:52.4 bits(124), Expect:9e-08,
Method:Compositional matrix adjust.,
Identities:48/76(63%), Positives:53/76(69%), Gaps:15/76(19%)

Query  451  IPPLKKRKRGVAG-------------GTKPIDVVTKAtteppttteePSATEQNPVAIEG  311
            +PPLKKRKRG+                 KPI+ VTKATT+ PTT EEPS+TEQNP AIEG
Sbjct  1    MPPLKKRKRGLTKPIDGRTKETTEEPAAKPINAVTKATTKKPTTMEEPSSTEQNP-AIEG  59

Query  310  -GVEEEPIPTIIPTVV  266
             GVEEEPI TIIPTVV
Sbjct  60   RGVEEEPISTIIPTVV  75


>unnamed protein product [Arabidopsis thaliana]
Sequence ID: VYS60963.1 Length: 145
Range 1: 1 to 32

Score:43.5 bits(101), Expect:9e-08,
Method:Compositional matrix adjust.,
Identities:22/32(69%), Positives:26/32(81%), Gaps:0/32(0%)

Query  1195  MMRYINVYLRQVQFVTRSNSIKPVSFGIVSLK  1290
             M+R INVYL+QVQ V   +SIKPVSFGIV L+
Sbjct  1     MLRDINVYLKQVQPVIGPSSIKPVSFGIVFLE  32


Range 2: 30 to 54

Score:34.3 bits(77), Expect:9e-08,
Method:Compositional matrix adjust.,
Identities:15/25(60%), Positives:18/25(72%), Gaps:0/25(0%)

Query  1281  FLETPDEISYDIPIYRFPRDSAFQR  1355
             FLE PDE+  DIPIYR PR S  ++
Sbjct  30    FLENPDELGNDIPIYRSPRVSFLKK  54


>unnamed protein product [Arabidopsis thaliana]
Sequence ID: VYS59161.1 Length: 113
Range 1: 1 to 75

Score:52.4 bits(124), Expect:9e-08,
Method:Compositional matrix adjust.,
Identities:47/79(59%), Positives:51/79(64%), Gaps:21/79(26%)

Query  451  IPPLKKRKRGVAGGTKPIDVVTKAtteppttt----------------eePSATEQNPVA  320
            +PPLKKRKRG+   TKPID  TKATTE P                   EEPS+TEQNP A
Sbjct  1    MPPLKKRKRGL---TKPIDGRTKATTEEPAAKPINAVTKATTKKPTTTEEPSSTEQNP-A  56

Query  319  IEG-GVEEEPIPTIIPTVV  266
            IEG G+EEEPI TIIPTVV
Sbjct  57   IEGRGIEEEPISTIIPTVV  75


Query #182: XLOC_020144 Query ID: lcl|Query_33185 Length: 2048

Sequences producing significant alignments:
                                                                  Max    Total Query   E   Per.
Description                                                       Score  Score cover Value Ident  Accession
hypothetical protein AXX17_AT3G09830 [Arabidopsis thaliana]       69.7   69.7  16%   2e-13 35.48  OAP06532.1


Alignments:
>hypothetical protein AXX17_AT3G09830 [Arabidopsis thaliana]
Sequence ID: OAP06532.1 Length: 140
Range 1: 27 to 140

Score:69.7 bits(169), Expect:2e-13,
Method:Compositional matrix adjust.,
Identities:44/124(35%), Positives:64/124(51%), Gaps:24/124(19%)

Query  198  RSYHRFSSPDAIFLSNPPLSEQPTSKQNSRRSRKHNKAFYLSI*RYEAY-----------  344
            RSYH FS+ D I L+NP LSEQPTS+ NS+R RK     Y +   +E Y
Sbjct  27   RSYHHFSALDTISLTNPQLSEQPTSESNSKRPRKRQDTIYATTESFEFYRDLSHLYRDRP  86

Query  345  ---PSNTINLTNIEAGY*ALSLFPRNI*HLSISQTRTARTTGVEGRFPAIQNT*RAHLHI  515
               PS  +++  +           +++ H+  SQ RT   TG+E + P IQ T R +L+I
Sbjct  87   HPIPSCLLSVITVSM---------KHMTHVK-SQIRTVGATGLESKLPTIQKTSRGYLNI  136

Query  516  VLNL  527
            ++ L
Sbjct  137  IMKL  140


Query #183: XLOC_020145 Query ID: lcl|Query_33186 Length: 412

Sequences producing significant alignments:
                                                                  Max    Total Query   E   Per.
Description                                                       Score  Score cover Value Ident  Accession
unnamed protein product [Arabidopsis thaliana]                    132    132   44%   4e-39 100.00 BAB02224.1
TPX2 (targeting protein for Xklp2) protein family [Arabidopsis... 81.3   118   46%   5e-21 81.82  NP_199240.1
unnamed protein product [Arabidopsis thaliana]                    73.9   107   41%   1e-17 77.27  CAA0407451.1
unnamed protein product [Arabidopsis thaliana]                    63.5   63.5  40%   1e-12 56.67  BAB02622.1


Alignments:
>unnamed protein product [Arabidopsis thaliana]
Sequence ID: BAB02224.1 Length: 214
Range 1: 1 to 61

Score:132 bits(332), Expect:4e-39,
Method:Compositional matrix adjust.,
Identities:61/61(100%), Positives:61/61(100%), Gaps:0/61(0%)

Query  351  MLLETKITFVWEVCQLGFGPIDLPGSQKCRRPDYPLALVGKHPNSGQTVWWSFGKKSTLK  172
            MLLETKITFVWEVCQLGFGPIDLPGSQKCRRPDYPLALVGKHPNSGQTVWWSFGKKSTLK
Sbjct  1    MLLETKITFVWEVCQLGFGPIDLPGSQKCRRPDYPLALVGKHPNSGQTVWWSFGKKSTLK  60

Query  171  Y  169
            Y
Sbjct  61   Y  61


>TPX2 (targeting protein for Xklp2) protein family [Arabidopsis thaliana]
Sequence ID: NP_199240.1 Length: 309
>TPX2 (targeting protein for Xklp2) protein family [Arabidopsis thaliana]
Sequence ID: AED95084.1 Length: 309 >unnamed protein product [Arabidopsis thaliana]
Sequence ID: BAB10112.1 Length: 309
Range 1: 1 to 44

Score:81.3 bits(199), Expect:5e-21,
Method:Compositional matrix adjust.,
Identities:36/44(82%), Positives:37/44(84%), Gaps:0/44(0%)

Query  345  LETKITFVWEVCQLGFGPIDLPGSQKCRRPDYPLALVGKHPNSG  214
            +ETKITFVWEVCQLG GPI LPGSQKC RPDY LALVGKH   G
Sbjct  1    METKITFVWEVCQLGLGPISLPGSQKCMRPDYSLALVGKHSKLG  44


Range 2: 39 to 65

Score:37.7 bits(86), Expect:5e-21,
Method:Compositional matrix adjust.,
Identities:21/27(78%), Positives:23/27(85%), Gaps:1/27(3%)

Query  232  KASKLGTDSVVVLREKVHSEVLR-CVS  155
            K SKLGTDSVVVLR +VHS+VLR C S
Sbjct  39   KHSKLGTDSVVVLRGEVHSKVLRDCKS  65


>unnamed protein product [Arabidopsis thaliana]
Sequence ID: CAA0407451.1 Length: 209
Range 1: 1 to 44

Score:73.9 bits(180), Expect:1e-17,
Method:Compositional matrix adjust.,
Identities:34/44(77%), Positives:35/44(79%), Gaps:0/44(0%)

Query  345  LETKITFVWEVCQLGFGPIDLPGSQKCRRPDYPLALVGKHPNSG  214
            +ETKITFVWEVCQLG  PI L GSQKC RPDY LALVGKH   G
Sbjct  1    METKITFVWEVCQLGLDPISLLGSQKCMRPDYSLALVGKHSKLG  44


Range 2: 39 to 58

Score:33.9 bits(76), Expect:1e-17,
Method:Compositional matrix adjust.,
Identities:16/20(80%), Positives:18/20(90%), Gaps:0/20(0%)

Query  232  KASKLGTDSVVVLREKVHSE  173
            K SKLGTDSVVVLR ++HSE
Sbjct  39   KHSKLGTDSVVVLRGEIHSE  58


>unnamed protein product [Arabidopsis thaliana]
Sequence ID: BAB02622.1 Length: 211
Range 1: 120 to 178

Score:63.5 bits(153), Expect:1e-12,
Method:Compositional matrix adjust.,
Identities:34/60(57%), Positives:42/60(70%), Gaps:6/60(10%)

Query  153  VETHRSTSEWTFSRRTTTLSVPS-----LDAFRLMPGGSQAACISDYQVNQLGQSPTDRL  317
            VETHRST+EWT S  +  + +P       +AFRLMP  +Q AC+SDYQVNQ+ Q PTDRL
Sbjct  120  VETHRSTTEWT-SPDSGRIGLPHSLVRVWNAFRLMPRDNQTACLSDYQVNQVRQCPTDRL  178


Query #184: XLOC_020146 Query ID: lcl|Query_33187 Length: 1867

Sequences producing significant alignments:
                                                                  Max    Total Query   E   Per.
Description                                                       Score  Score cover Value Ident  Accession
hypothetical protein AXX17_AT5G30800 [Arabidopsis thaliana]       74.7   74.7  8%    2e-13 74.00  OAO94454.1
F7F22.14 [Arabidopsis thaliana]                                   59.3   59.3  5%    8e-09 81.82  AAF24528.1


Alignments:
>hypothetical protein AXX17_AT5G30800 [Arabidopsis thaliana]
Sequence ID: OAO94454.1 Length: 416
Range 1: 367 to 416

Score:74.7 bits(182), Expect:2e-13,
Method:Compositional matrix adjust.,
Identities:37/50(74%), Positives:42/50(84%), Gaps:0/50(0%)

Query  80   QVSACDSDLASYIEQSMAAGIANRCIVYHVSLLVVLRSVNTVDRTSLSSR  229
             VSACDSD AS IE+S+A GIAN+C +Y VSLLVVLRSVN +DRTSLS R
Sbjct  367  PVSACDSDFASSIEESVAVGIANKCSLYSVSLLVVLRSVNGMDRTSLSPR  416


>F7F22.14 [Arabidopsis thaliana]
Sequence ID: AAF24528.1 Length: 304
Range 1: 272 to 304

Score:59.3 bits(142), Expect:8e-09,
Method:Compositional matrix adjust.,
Identities:27/33(82%), Positives:30/33(90%), Gaps:0/33(0%)

Query  396  GLYMFEIHLEIMEHSGGKHRIYAVEEIRRLKTA  494
            GLYMFEIHLEIMEHSGGK RIYA E+I +LK+A
Sbjct  272  GLYMFEIHLEIMEHSGGKKRIYADEKIWKLKSA  304


Query #185: XLOC_020147 Query ID: lcl|Query_33188 Length: 424

Sequences producing significant alignments:
                                                                  Max    Total Query   E   Per.
Description                                                       Score  Score cover Value Ident  Accession
serine carboxypeptidase [Arabidopsis thaliana]                    84.0   84.0  41%   4e-19 61.67  BAA96893.1
unnamed protein product [Arabidopsis thaliana]                    67.8   67.8  59%   3e-15 45.24  CAA0405760.1
Mannose-binding lectin superfamily protein [Arabidopsis thaliana] 72.4   72.4  30%   4e-15 72.09  NP_175623.2
unnamed protein product [Arabidopsis thaliana]                    62.8   62.8  29%   1e-11 68.29  CAA0288322.1


Alignments:
>serine carboxypeptidase [Arabidopsis thaliana]
Sequence ID: BAA96893.1 Length: 512
Range 1: 441 to 500

Score:84.0 bits(206), Expect:4e-19,
Method:Composition-based stats.,
Identities:37/60(62%), Positives:47/60(78%), Gaps:1/60(1%)

Query  422  LAQWLTNGSLILLHREVWGSNPSYSDLLTDLHINRGKCVGDLQRCTII-AVRQDFTQRRY  246
            L QWLT GSL+LL ++VWGS+P+Y DLL DL IN GKC GDLQRCTI+  +R+  T+R +
Sbjct  441  LGQWLTKGSLMLLQQDVWGSSPNYCDLLVDLQINGGKCAGDLQRCTIMDLIRKKKTKRSF  500


>unnamed protein product [Arabidopsis thaliana]
Sequence ID: CAA0405760.1 Length: 84
Range 1: 1 to 84

Score:67.8 bits(164), Expect:3e-15,
Method:Compositional matrix adjust.,
Identities:38/84(45%), Positives:46/84(54%), Gaps:0/84(0%)

Query  414  MVDQWFINTSTPRGLGFESQLLRFINRFAY*SRQVCRRSTTMYNYSRSPGFYAEKILKIF  235
            MVDQ FIN STPRG G   +    +        +  +        +   GFYAEKILK+
Sbjct  1    MVDQGFINASTPRGRGSSLRYCDLLADLQINGGKCAKDLQRCTIIAVHLGFYAEKILKVS  60

Query  234  IKRMENEKEFSYVVDLIRVRRNCP  163
            IKRME E + S V+DLIRV RN P
Sbjct  61   IKRMEKENDVSSVMDLIRVGRNYP  84


>Mannose-binding lectin superfamily protein [Arabidopsis thaliana]
Sequence ID: NP_175623.2 Length: 615
>Mannose-binding lectin superfamily protein [Arabidopsis thaliana]
Sequence ID: AEE32756.1 Length: 615
Range 1: 1 to 43

Score:72.4 bits(176), Expect:4e-15,
Method:Composition-based stats.,
Identities:31/43(72%), Positives:37/43(86%), Gaps:0/43(0%)

Query  291  MYNYSRSPGFYAEKILKIFIKRMENEKEFSYVVDLIRVRRNCP  163
            MYNYSRSPGFYA+KILK+ IKRM+ + E S VVDLI++ RNCP
Sbjct  1    MYNYSRSPGFYAKKILKMSIKRMKKKNEMSSVVDLIKIGRNCP  43


>unnamed protein product [Arabidopsis thaliana]
Sequence ID: CAA0288322.1 Length: 614
Range 1: 1 to 41

Score:62.8 bits(151), Expect:1e-11,
Method:Composition-based stats.,
Identities:28/41(68%), Positives:34/41(82%), Gaps:0/41(0%)

Query  291  MYNYSRSPGFYAEKILKIFIKRMENEKEFSYVVDLIRVRRN  169
            MYNYS SPGFYA+KILK+ IKRM+ + E S VVDLI++ RN
Sbjct  1    MYNYSHSPGFYAKKILKMSIKRMKKKNEMSSVVDLIKIGRN  41


Query #186: XLOC_020148 Query ID: lcl|Query_33189 Length: 693

No significant similarity found.


Query #187: XLOC_020149 Query ID: lcl|Query_33190 Length: 529

No significant similarity found.


Query #188: XLOC_020150 Query ID: lcl|Query_33191 Length: 189

No significant similarity found.


Query #189: XLOC_020151 Query ID: lcl|Query_33192 Length: 558

No significant similarity found.


Query #190: XLOC_020152 Query ID: lcl|Query_33193 Length: 721

No significant similarity found.


Query #191: XLOC_020153 Query ID: lcl|Query_33194 Length: 756

No significant similarity found.


Query #192: XLOC_020154 Query ID: lcl|Query_33195 Length: 1060

Sequences producing significant alignments:
                                                                  Max    Total Query   E   Per.
Description                                                       Score  Score cover Value Ident  Accession
hypothetical protein AXX17_AT3G50110 [Arabidopsis thaliana]       140    140   20%   7e-41 97.18  OAP02241.1
unnamed protein product [Arabidopsis thaliana]                    139    139   20%   7e-41 97.18  CAA0386582.1
hypothetical protein AT3G55513 [Arabidopsis thaliana]             138    138   19%   3e-40 98.53  NP_001326971.1
hypothetical protein AXX17_AT3G33540 [Arabidopsis thaliana]       74.7   74.7  20%   2e-16 80.28  OAP03224.1


Alignments:
>hypothetical protein AXX17_AT3G50110 [Arabidopsis thaliana]
Sequence ID: OAP02241.1 Length: 87
Range 1: 1 to 71

Score:140 bits(352), Expect:7e-41,
Method:Compositional matrix adjust.,
Identities:69/71(97%), Positives:69/71(97%), Gaps:0/71(0%)

Query  867  MRPKLTCLKVDGEDGVVVLASASWCGSSFCLWSRVSLHWEHRRRLWPSSFTGSEQQRLRS  688
            MRPK TCLKVDGEDGVVVLASASWCGSSFCLWSRVSLH EHRRRLWPSSFTGSEQQRLRS
Sbjct  1    MRPKSTCLKVDGEDGVVVLASASWCGSSFCLWSRVSLHLEHRRRLWPSSFTGSEQQRLRS  60

Query  687  RYLGPSLVSHA  655
            RYLGPSLVSHA
Sbjct  61   RYLGPSLVSHA  71


>unnamed protein product [Arabidopsis thaliana]
Sequence ID: CAA0386582.1 Length: 81
>unnamed protein product [Arabidopsis thaliana]
Sequence ID: VYS60488.1 Length: 81
Range 1: 1 to 71

Score:139 bits(351), Expect:7e-41,
Method:Compositional matrix adjust.,
Identities:69/71(97%), Positives:69/71(97%), Gaps:0/71(0%)

Query  867  MRPKLTCLKVDGEDGVVVLASASWCGSSFCLWSRVSLHWEHRRRLWPSSFTGSEQQRLRS  688
            MRPK TCLKVDGEDGVVVLASASWCGSSFCLWSRVSLH EHRRRLWPSSFTGSEQQRLRS
Sbjct  1    MRPKSTCLKVDGEDGVVVLASASWCGSSFCLWSRVSLHLEHRRRLWPSSFTGSEQQRLRS  60

Query  687  RYLGPSLVSHA  655
            RYLGPSLVSHA
Sbjct  61   RYLGPSLVSHA  71


>hypothetical protein AT3G55513 [Arabidopsis thaliana]
Sequence ID: NP_001326971.1 Length: 87
>hypothetical protein AT3G55513 [Arabidopsis thaliana]
Sequence ID: ANM64971.1 Length: 87
Range 1: 1 to 68

Score:138 bits(348), Expect:3e-40,
Method:Compositional matrix adjust.,
Identities:67/68(99%), Positives:68/68(100%), Gaps:0/68(0%)

Query  867  MRPKLTCLKVDGEDGVVVLASASWCGSSFCLWSRVSLHWEHRRRLWPSSFTGSEQQRLRS  688
            MRPKLTCLKVDGEDGVVVLASASWCGSSFCLWSRVSLHWEHRRRLWPSSFTGSEQQRLRS
Sbjct  1    MRPKLTCLKVDGEDGVVVLASASWCGSSFCLWSRVSLHWEHRRRLWPSSFTGSEQQRLRS  60

Query  687  RYLGPSLV  664
            RYLGPSL+
Sbjct  61   RYLGPSLL  68


>hypothetical protein AXX17_AT3G33540 [Arabidopsis thaliana]
Sequence ID: OAP03224.1 Length: 71
Range 1: 1 to 71

Score:74.7 bits(182), Expect:2e-16,
Method:Compositional matrix adjust.,
Identities:57/71(80%), Positives:61/71(85%), Gaps:0/71(0%)

Query  827  MESSFSPLPHGVDLRFVSGHVFHCIgstgggsgllpsldlsnngSVLGTSVPPSSLMLPL  648
            M SSFSPLPHGVDLR VSGHVFHC+ STGGGS LLPS D+SNNGSVLG SVP SSL+LPL
Sbjct  1    MASSFSPLPHGVDLRLVSGHVFHCVWSTGGGSVLLPSPDVSNNGSVLGASVPHSSLILPL  60

Query  647  CVHVFWLFERF  615
            CV+VF   ERF
Sbjct  61   CVYVFCFIERF  71


Query #193: XLOC_020155 Query ID: lcl|Query_33196 Length: 371

No significant similarity found.


Query #194: XLOC_020156 Query ID: lcl|Query_33197 Length: 696

No significant similarity found.


Query #195: XLOC_020157 Query ID: lcl|Query_33198 Length: 741

No significant similarity found.


Query #196: XLOC_020158 Query ID: lcl|Query_33199 Length: 577

No significant similarity found.


Query #197: XLOC_020159 Query ID: lcl|Query_33200 Length: 670

No significant similarity found.


Query #198: XLOC_020260 Query ID: lcl|Query_33201 Length: 547

Sequences producing significant alignments:
                                                                  Max    Total Query   E   Per.
Description                                                       Score  Score cover Value Ident  Accession
hypothetical protein AT3G47965 [Arabidopsis thaliana]             80.1   80.1  24%   6e-20 86.36  NP_974401.1


Alignments:
>hypothetical protein AT3G47965 [Arabidopsis thaliana]
Sequence ID: NP_974401.1 Length: 44
>AT3g47961 [Arabidopsis thaliana]
Sequence ID: AAK32766.1 Length: 44 >AT3g47961/AT3g47961 [Arabidopsis thaliana]
Sequence ID: AAM91424.1 Length: 44 >hypothetical protein AT3G47965 [Arabidopsis thaliana]
Sequence ID: AEE78352.1 Length: 44
Range 1: 1 to 44

Score:80.1 bits(196), Expect:6e-20,
Method:Compositional matrix adjust.,
Identities:38/44(86%), Positives:41/44(93%), Gaps:0/44(0%)

Query  245  M*LSHEFLPRGLGRIEIVPTSIVEDTRFFKCMYCCEFYRLMSLV  376
            M LSHE LPRGLGRIE+VPTSIVE+T+ FKCMYCCEFYRLMSLV
Sbjct  1    MWLSHELLPRGLGRIEMVPTSIVEETKCFKCMYCCEFYRLMSLV  44


Query #199: XLOC_020505 Query ID: lcl|Query_33202 Length: 1489

Sequences producing significant alignments:
                                                                  Max    Total Query   E   Per.
Description                                                       Score  Score cover Value Ident  Accession
hypothetical protein AXX17_AT3G32650 [Arabidopsis thaliana]       84.3   84.3  16%   3e-19 53.57  OAP03023.1
K-box region protein (DUF1985) [Arabidopsis thaliana]             77.4   77.4  15%   2e-14 45.24  NP_001321227.1
K-box region protein (DUF1985) [Arabidopsis thaliana]             77.4   77.4  15%   2e-14 45.24  NP_174400.1
unnamed protein product [Arabidopsis thaliana]                    77.4   77.4  15%   3e-14 45.24  VYS47676.1
unknown protein [Arabidopsis thaliana]                            77.0   77.0  17%   5e-14 38.30  AAG50606.1
hypothetical protein AXX17_AT1G31740 [Arabidopsis thaliana]       76.3   76.3  15%   6e-14 44.05  OAP15120.1
putative protein [Arabidopsis thaliana]                           73.6   73.6  20%   7e-14 37.72  CAB86472.1
hypothetical protein AXX17_AT2G06070 [Arabidopsis thaliana]       75.1   75.1  17%   2e-13 39.36  OAP09613.1
hypothetical protein AXX17_AT1G38890 [Arabidopsis thaliana]       73.2   73.2  13%   4e-13 47.95  OAP18985.1
hypothetical protein [Arabidopsis thaliana]                       73.6   73.6  19%   4e-13 38.10  AAD28660.1
unknown protein; 55998-51558 [Arabidopsis thaliana]               72.8   72.8  13%   6e-13 47.95  AAG51237.1
putative protein [Arabidopsis thaliana]                           72.0   72.0  11%   1e-12 50.85  CAB81786.1
F7F22.7 [Arabidopsis thaliana]                                    68.6   68.6  23%   4e-12 30.08  AAF24523.1
unnamed protein product [Arabidopsis thaliana]                    69.3   69.3  16%   1e-11 41.46  BAB01835.1
hypothetical protein [Arabidopsis thaliana]                       69.3   69.3  16%   1e-11 41.46  AAC95211.1
Similar to a hypothetical protein At2g29240 gi|3980408 from...    69.3   69.3  16%   1e-11 41.46  AAF80659.1
F15O4.12 [Arabidopsis thaliana]                                   68.2   68.2  19%   2e-11 37.37  AAF79347.1
F22O13.23 [Arabidopsis thaliana]                                  68.2   68.2  16%   3e-11 41.46  AAF99775.1
putative protein [Arabidopsis thaliana]                           67.4   67.4  19%   5e-11 36.63  CAB86452.1
T9E19.2 gene product [Arabidopsis thaliana]                       66.2   66.2  12%   5e-11 44.26  AAC72876.1
Contains similarity to a putative protein T32A11_100 gi|741362... 67.0   67.0  11%   6e-11 50.88  AAF78272.1
transmembrane protein, putative (DUF1985) [Arabidopsis thaliana]  65.9   65.9  14%   7e-11 43.75  NP_174889.1
unnamed protein product [Arabidopsis thaliana]                    65.9   65.9  12%   9e-11 44.26  BAB10789.1
T32E20.31 [Arabidopsis thaliana]                                  65.5   65.5  13%   9e-11 47.76  AAF79792.1
unnamed protein product [Arabidopsis thaliana]                    66.2   66.2  18%   1e-10 36.73  BAB02447.1
hypothetical protein [Arabidopsis thaliana]                       66.2   66.2  12%   1e-10 44.26  AAD24651.1
hypothetical protein [Arabidopsis thaliana]                       66.2   66.2  12%   1e-10 44.26  AAD14458.1
hypothetical protein AXX17_AT1G37570 [Arabidopsis thaliana]       65.1   65.1  14%   1e-10 43.75  OAP14713.1
T22C5.24 [Arabidopsis thaliana]                                   65.9   65.9  12%   1e-10 44.26  AAF24952.1
unnamed protein product [Arabidopsis thaliana]                    64.7   64.7  13%   2e-10 47.76  VYS48136.1
unnamed protein product [Arabidopsis thaliana]                    64.7   64.7  13%   2e-10 47.76  CAA0270146.1
hypothetical protein [Arabidopsis thaliana]                       65.5   65.5  12%   2e-10 44.26  AAG50521.1
F21H2.5 [Arabidopsis thaliana]                                    65.5   65.5  12%   2e-10 44.26  AAD46015.1
hypothetical protein [Arabidopsis thaliana]                       65.5   65.5  12%   2e-10 44.26  AAG50582.1
unnamed protein product [Arabidopsis thaliana]                    64.7   64.7  12%   3e-10 44.26  BAB02940.1
putative protein [Arabidopsis thaliana]                           64.3   64.3  12%   4e-10 42.62  CAB81580.1
unnamed protein product [Arabidopsis thaliana]                    64.3   64.3  11%   4e-10 44.83  BAA98062.1
putative protein [Arabidopsis thaliana]                           63.5   63.5  12%   9e-10 42.62  CAB81070.1
transmembrane protein, putative (DUF1985) [Arabidopsis thaliana]  45.4   84.3  16%   1e-09 50.00  NP_178732.1
unnamed protein product [Arabidopsis thaliana]                    45.1   83.9  16%   1e-09 50.00  VYS52190.1
hypothetical protein AXX17_AT2G06010 [Arabidopsis thaliana]       45.1   83.9  16%   1e-09 50.00  OAP10666.1
putative protein [Arabidopsis thaliana]                           62.4   62.4  12%   2e-09 42.62  CAB86693.1
Strong similarity to gi|3252818 F26C24.9 hypothetical protein...  62.0   62.0  12%   2e-09 42.62  AAD12676.1
hypothetical protein [Arabidopsis thaliana]                       61.2   82.8  16%   2e-09 43.33  AAD29765.1
unnamed protein product [Arabidopsis thaliana]                    61.2   61.2  16%   3e-09 36.67  CAA0364214.1
hypothetical protein [Arabidopsis thaliana]                       61.2   61.2  12%   4e-09 40.98  AAC24188.1
hypothetical protein [Arabidopsis thaliana]                       60.5   60.5  12%   7e-09 39.34  AAD24662.1
T4B21.18 gene product [Arabidopsis thaliana]                      58.5   58.5  20%   7e-09 34.31  AAD03457.1
T24H24.2 gene product [Arabidopsis thaliana]                      58.9   58.9  16%   9e-09 37.35  AAC28200.1
hypothetical protein [Arabidopsis thaliana]                       59.3   59.3  20%   1e-08 34.31  AAD23020.1
hypothetical protein AXX17_AT2G12740 [Arabidopsis thaliana]       58.9   58.9  16%   2e-08 36.67  OAP08567.1
F27F5.16 [Arabidopsis thaliana]                                   59.3   59.3  12%   2e-08 39.34  AAF69170.1
unnamed protein product [Arabidopsis thaliana]                    56.2   56.2  15%   2e-08 40.96  BAA97088.1
hypothetical protein [Arabidopsis thaliana]                       59.3   59.3  12%   2e-08 39.34  AAC97230.2
unnamed protein product [Arabidopsis thaliana]                    58.5   58.5  16%   2e-08 37.35  BAB01421.1
CDS [Arabidopsis thaliana]                                        58.2   58.2  16%   3e-08 38.55  AAF31285.1
hypothetical protein AXX17_AT5G31140 [Arabidopsis thaliana]       56.6   56.6  17%   3e-08 35.63  OAO92309.1
hypothetical protein [Arabidopsis thaliana]                       57.8   57.8  16%   5e-08 36.25  AAD19785.1
unnamed protein product [Arabidopsis thaliana]                    55.1   55.1  9%    6e-08 50.00  CAA0405522.1
hypothetical protein (DUF1985) [Arabidopsis thaliana]             54.7   54.7  9%    7e-08 50.00  NP_001154746.1


Alignments:
>hypothetical protein AXX17_AT3G32650 [Arabidopsis thaliana]
Sequence ID: OAP03023.1 Length: 87
Range 1: 1 to 84

Score:84.3 bits(207), Expect:3e-19,
Method:Composition-based stats.,
Identities:45/84(54%), Positives:55/84(65%), Gaps:4/84(4%)

Query  555  METDEGITSMSSRVSEA----LIQTW*RAFCWKNQQLLQNSILQRIKKALPLEYKEVKSD  722
            METDEGITSMS  + +     L +        K     + ++L+RIKKA+PLEYKEVKSD
Sbjct  1    METDEGITSMSPDIDKEYPKRLFRHGREPLVGKVNNFCKTALLKRIKKAMPLEYKEVKSD  60

Query  723  PQFAHVFAIFESRLGYSARTVHSL  794
            P FAHVFAIFE+RLGYS    + L
Sbjct  61   PLFAHVFAIFENRLGYSVCIAYCL  84


>K-box region protein (DUF1985) [Arabidopsis thaliana]
Sequence ID: NP_001321227.1 Length: 623
>K-box region protein (DUF1985) [Arabidopsis thaliana]
Sequence ID: ANM58816.1 Length: 623
Range 1: 48 to 130

Score:77.4 bits(189), Expect:2e-14,
Method:Compositional matrix adjust.,
Identities:38/84(45%), Positives:56/84(66%), Gaps:6/84(7%)

Query  699  EYKEVKSDPQFAHVFAIFESRLGYSARTVHSLMCRQLVTRKKHELWFVFGNKPLRFSMQE  878
            E++ +KS  QF  +F    +R  +S + +H L+ RQ+VT+KKHELWFVFG  P+RFS++E
Sbjct  48   EWERIKSS-QFGKLFEFPVARCSHSGKLIHGLLSRQVVTKKKHELWFVFGGHPIRFSIRE  106

Query  879  FYAVTGLK-----YEDDFRHDLDS  935
            F+ VTGL+      ED+ +   DS
Sbjct  107  FHIVTGLRCGKLPTEDEVKKHQDS  130


>K-box region protein (DUF1985) [Arabidopsis thaliana]
Sequence ID: NP_174400.1 Length: 673
>EST gb|Z33866 comes from this gene [Arabidopsis thaliana]
Sequence ID: AAD21698.1 Length: 673 >K-box region protein (DUF1985) [Arabidopsis thaliana]
Sequence ID: AEE31318.1 Length: 673 >unnamed protein product [Arabidopsis thaliana]
Sequence ID: CAA0259238.1 Length: 673
Range 1: 48 to 130

Score:77.4 bits(189), Expect:2e-14,
Method:Compositional matrix adjust.,
Identities:38/84(45%), Positives:56/84(66%), Gaps:6/84(7%)

Query  699  EYKEVKSDPQFAHVFAIFESRLGYSARTVHSLMCRQLVTRKKHELWFVFGNKPLRFSMQE  878
            E++ +KS  QF  +F    +R  +S + +H L+ RQ+VT+KKHELWFVFG  P+RFS++E
Sbjct  48   EWERIKSS-QFGKLFEFPVARCSHSGKLIHGLLSRQVVTKKKHELWFVFGGHPIRFSIRE  106

Query  879  FYAVTGLK-----YEDDFRHDLDS  935
            F+ VTGL+      ED+ +   DS
Sbjct  107  FHIVTGLRCGKLPTEDEVKKHQDS  130


>unnamed protein product [Arabidopsis thaliana]
Sequence ID: VYS47676.1 Length: 673
Range 1: 48 to 130

Score:77.4 bits(189), Expect:3e-14,
Method:Compositional matrix adjust.,
Identities:38/84(45%), Positives:56/84(66%), Gaps:6/84(7%)

Query  699  EYKEVKSDPQFAHVFAIFESRLGYSARTVHSLMCRQLVTRKKHELWFVFGNKPLRFSMQE  878
            E++ +KS  QF  +F    +R  +S + +H L+ RQ+VT+KKHELWFVFG  P+RFS++E
Sbjct  48   EWERIKSS-QFGKLFEFPVARCSHSGKLIHGLLSRQVVTKKKHELWFVFGGHPIRFSIRE  106

Query  879  FYAVTGLK-----YEDDFRHDLDS  935
            F+ VTGL+      ED+ +   DS
Sbjct  107  FHIVTGLRCGKLPTEDEVKKHQDS  130


>unknown protein [Arabidopsis thaliana]
Sequence ID: AAG50606.1 Length: 1311
Range 1: 189 to 282

Score:77.0 bits(188), Expect:5e-14,
Method:Compositional matrix adjust.,
Identities:36/94(38%), Positives:60/94(63%), Gaps:5/94(5%)

Query  729  FAHVFAIFESRLGYSARTVHSLMCRQLVTRKKHELWFVFGNKPLRFSMQEFYAVTGL---  899
            F  +F +  SR+  S + +H+L+CRQL+T+KK+E W VFG  P+RFS+ EF +VTGL
Sbjct  189  FGPLFHLPVSRVATSGKVIHALLCRQLLTKKKYEFWTVFGGHPMRFSLLEFASVTGLPCG  248

Query  900  KYEDDFRHDLDS*RDD--KGFWSKLLKKGNTICI  995
            ++ D++  +     DD  K +W++L+    T+ +
Sbjct  249  EFPDEYDPEDSPVYDDGKKSYWNELIGPDKTVTL  282


>hypothetical protein AXX17_AT1G31740 [Arabidopsis thaliana]
Sequence ID: OAP15120.1 Length: 673
Range 1: 48 to 130

Score:76.3 bits(186), Expect:6e-14,
Method:Compositional matrix adjust.,
Identities:37/84(44%), Positives:56/84(66%), Gaps:6/84(7%)

Query  699  EYKEVKSDPQFAHVFAIFESRLGYSARTVHSLMCRQLVTRKKHELWFVFGNKPLRFSMQE  878
            E++ +K+  QF  +F    +R  +S + +H L+ RQ+VT+KKHELWFVFG  P+RFS++E
Sbjct  48   EWERIKNS-QFGKLFEFPVARCSHSGKLIHGLLSRQVVTKKKHELWFVFGGHPIRFSIRE  106

Query  879  FYAVTGLK-----YEDDFRHDLDS  935
            F+ VTGL+      ED+ +   DS
Sbjct  107  FHIVTGLRCGKLPTEDEVKKHQDS  130


>putative protein [Arabidopsis thaliana]
Sequence ID: CAB86472.1 Length: 264
Range 1: 2 to 109

Score:73.6 bits(179), Expect:7e-14,
Method:Compositional matrix adjust.,
Identities:43/114(38%), Positives:66/114(57%), Gaps:17/114(14%)

Query  771   SARTVHSLMCRQLVTRKKHELWFVFGNKPLRFSMQEFYAVTGL---KYEDDFRHDLDS*R  941
             S + +H+L+CRQLVT+KK+ELW VFG  P+RFS+ EF  VTGL   ++ +D+  D DS
Sbjct  2     SGKLMHALLCRQLVTKKKYELWTVFGGHPMRFSLTEFACVTGLPCAEFSEDYDPDDDSVF  61

Query  942   DD--KGFWSKLLKKGNTICIK------TFMDKQLSEVHMWDEVDRVRFVYLCVI  1079
              D  K +W +L+    T+ ++      T   K LS  H      R++  +L ++
Sbjct  62    VDGMKSYWDELIGPDKTVTLRDVSAMLTNKRKTLSSDH------RLKLAFLLIV  109


>hypothetical protein AXX17_AT2G06070 [Arabidopsis thaliana]
Sequence ID: OAP09613.1 Length: 1302
Range 1: 228 to 319

Score:75.1 bits(183), Expect:2e-13,
Method:Compositional matrix adjust.,
Identities:37/94(39%), Positives:54/94(57%), Gaps:9/94(9%)

Query  720  DPQFAHVFAIFESRLGYSARTVHSLMCRQLVTRKKHELWFVFGNKPLRFSMQEFYAVTGL  899
            D  F  +F++   R   S + +H+L+CRQLVT++KHE+W VFG  PL FS+ EF +VTGL
Sbjct  228  DSAFGPLFSLLVQRCSLSCKLIHALVCRQLVTKRKHEMWTVFGGHPLHFSLAEFASVTGL  287

Query  900  K-------YEDDFRHDLDS*RDDKGFWSKLLKKG  980
                    Y+ D+   +    DD  +W K + K
Sbjct  288  PCGDFSEDYDPDYEPKIPKEADD--YWLKKIGKS  319


>hypothetical protein AXX17_AT1G38890 [Arabidopsis thaliana]
Sequence ID: OAP18985.1 Length: 506
Range 1: 189 to 261

Score:73.2 bits(178), Expect:4e-13,
Method:Compositional matrix adjust.,
Identities:35/73(48%), Positives:49/73(67%), Gaps:7/73(9%)

Query  720  DPQFAHVFAIFESRLGYSARTVHSLMCRQLVTRKKHELWFVFGNKPLRFSMQEFYAVTGL  899
            D  F  +F++  S    S + VH+L+CRQLVT++K+E+W VFG +PLRFS+ EF +VTGL
Sbjct  189  DSCFGSLFSLRVSECPISGKLVHALLCRQLVTKQKYEMWTVFGGQPLRFSLVEFGSVTGL  248

Query  900  -------KYEDDF  917
                   +YE DF
Sbjct  249  PCGEFPEEYEPDF  261


>hypothetical protein [Arabidopsis thaliana]
Sequence ID: AAD28660.1 Length: 863
Range 1: 1 to 104

Score:73.6 bits(179), Expect:4e-13,
Method:Compositional matrix adjust.,
Identities:40/105(38%), Positives:58/105(55%), Gaps:7/105(6%)

Query  783   VHSLMCRQLVTRKKHELWFVFGNKPLRFSMQEFYAVTGLKYEDDFRHDLDS*RDD-----  947
             +H+L+CRQLVT+KK+ELW VFG  PLRFS+ EF  VTGL    +F  D D   D
Sbjct  1     MHALLCRQLVTKKKYELWTVFGGYPLRFSLPEFACVTGLPC-GEFSEDYDPDDDAVYVVG  59

Query  948   -KGFWSKLLKKGNTICIKTFMDKQLSEVHMWDEVDRVRFVYLCVI  1079
              K +W KL+    T+ +    D   ++        R++ V+L ++
Sbjct  60    MKSYWDKLIGPDKTVTLGDVSDMLTNKRKTLSSDHRLKLVFLLIV  104


>unknown protein; 55998-51558 [Arabidopsis thaliana]
Sequence ID: AAG51237.1 Length: 593
Range 1: 137 to 209

Score:72.8 bits(177), Expect:6e-13,
Method:Compositional matrix adjust.,
Identities:35/73(48%), Positives:49/73(67%), Gaps:7/73(9%)

Query  720  DPQFAHVFAIFESRLGYSARTVHSLMCRQLVTRKKHELWFVFGNKPLRFSMQEFYAVTGL  899
            D  F  +F++  S    S + VH+L+CRQLVT++K+E+W VFG +PLRFS+ EF +VTGL
Sbjct  137  DSCFGSLFSLRVSECPISGKLVHALLCRQLVTKQKYEMWTVFGGQPLRFSLVEFGSVTGL  196

Query  900  -------KYEDDF  917
                   +YE DF
Sbjct  197  PCGEFPEEYEPDF  209


>putative protein [Arabidopsis thaliana]
Sequence ID: CAB81786.1 Length: 702
Range 1: 81 to 139

Score:72.0 bits(175), Expect:1e-12,
Method:Compositional matrix adjust.,
Identities:30/59(51%), Positives:43/59(72%), Gaps:0/59(0%)

Query  726  QFAHVFAIFESRLGYSARTVHSLMCRQLVTRKKHELWFVFGNKPLRFSMQEFYAVTGLK  902
            QF  +F     +  +SA+ +H L+ RQLVT+K+HE WF+FG KPLRFS++EF+  TGL+
Sbjct  81   QFGKLFKYPVCQTAHSAKLIHGLLSRQLVTKKRHEFWFIFGGKPLRFSLREFHITTGLE  139


>F7F22.7 [Arabidopsis thaliana]
Sequence ID: AAF24523.1 Length: 268
Range 1: 54 to 175

Score:68.6 bits(166), Expect:4e-12,
Method:Compositional matrix adjust.,
Identities:37/123(30%), Positives:63/123(51%), Gaps:7/123(5%)

Query  729   FAHVFAIFESRLGYSARTVHSLMCRQLVTRKKHELWFVFGNKPLRFSMQEFYAVTGLKYE  908
             F  +F +  S+   S + +H+L+CRQLVT+K +ELW +FG  P+R S+ EF  VTGL
Sbjct  54    FGPLFHLPVSKCAMSGKLMHALLCRQLVTKKNYELWTLFGGHPMRLSLLEFAYVTGLPC-  112

Query  909   DDFRHDLDS*RDD------KGFWSKLLKKGNTICIKTFMDKQLSEVHMWDEVDRVRFVYL  1070
              +F  D D   D       K +W++L+    T+ +        ++        ++R  +L
Sbjct  113   GEFSEDYDPDDDPIFVVGMKSYWNELISLNKTVTLGDVYAMLTNKRKTLSSDHKLRLAFL  172

Query  1071  CVI  1079
              ++
Sbjct  173   LIV  175


>unnamed protein product [Arabidopsis thaliana]
Sequence ID: BAB01835.1 Length: 1015
>unnamed protein product [Arabidopsis thaliana]
Sequence ID: BAB10871.1 Length: 1015
Range 1: 29 to 109

Score:69.3 bits(168), Expect:1e-11,
Method:Compositional matrix adjust.,
Identities:34/82(41%), Positives:48/82(58%), Gaps:1/82(1%)

Query  663  SILQRIKKALPLEYKEVKSDPQFAHVFAIFESRLGYSARTVHSLMCRQLVTRKKHELWFV  842
            S++  I  AL  E  ++  + QF  +F   ES   YS + +H L+ RQLV RKK E+W +
Sbjct  29   SLIDDIANALDAEDMKILRESQFGKLFDFSES-AAYSGKLIHFLLTRQLVVRKKQEIWVI  87

Query  843  FGNKPLRFSMQEFYAVTGLKYE  908
            F   P+RFS+ EF  VTGL  +
Sbjct  88   FSGSPVRFSISEFQNVTGLNCD  109


>hypothetical protein [Arabidopsis thaliana]
Sequence ID: AAC95211.1 Length: 992
Range 1: 29 to 109

Score:69.3 bits(168), Expect:1e-11,
Method:Compositional matrix adjust.,
Identities:34/82(41%), Positives:48/82(58%), Gaps:1/82(1%)

Query  663  SILQRIKKALPLEYKEVKSDPQFAHVFAIFESRLGYSARTVHSLMCRQLVTRKKHELWFV  842
            S++  I  AL  E  ++  + QF  +F   ES   YS + +H L+ RQLV RKK E+W +
Sbjct  29   SLIDDIANALDAEDMKILRESQFGKLFDFSES-AAYSGKLIHFLLTRQLVVRKKQEIWVI  87

Query  843  FGNKPLRFSMQEFYAVTGLKYE  908
            F   P+RFS+ EF  VTGL  +
Sbjct  88   FSGSPVRFSISEFQNVTGLNCD  109


>Similar to a hypothetical protein At2g29240 gi|3980408 from Arabidopsis thaliana gb|AC004561 [Arabidopsis thaliana]
Sequence ID: AAF80659.1 Length: 992
Range 1: 28 to 108

Score:69.3 bits(168), Expect:1e-11,
Method:Compositional matrix adjust.,
Identities:34/82(41%), Positives:48/82(58%), Gaps:1/82(1%)

Query  663  SILQRIKKALPLEYKEVKSDPQFAHVFAIFESRLGYSARTVHSLMCRQLVTRKKHELWFV  842
            S++  I  AL  E  ++  + QF  +F   ES   YS + +H L+ RQLV RKK E+W +
Sbjct  28   SLIDDIANALDAEDMKILRESQFGKLFDFSES-AAYSGKLIHFLLTRQLVVRKKQEIWVI  86

Query  843  FGNKPLRFSMQEFYAVTGLKYE  908
            F   P+RFS+ EF  VTGL  +
Sbjct  87   FSGSPVRFSISEFQNVTGLNCD  108


>F15O4.12 [Arabidopsis thaliana]
Sequence ID: AAF79347.1 Length: 724
Range 1: 139 to 236

Score:68.2 bits(165), Expect:2e-11,
Method:Compositional matrix adjust.,
Identities:37/99(37%), Positives:56/99(56%), Gaps:5/99(5%)

Query  624  RAFCWKNQQLLQN----SILQRIKKALPLEYKEVKSDPQFAHVFAIFESRLGYSARTVHS  791
            R+    N + +Q+    S++  I  AL  E  ++  + QF  +F   ES   YSA+ ++
Sbjct  139  RSGSEPNSERVQSYSELSLIDDIANALDAEDMKILRESQFGKLFDFSES-AAYSAKLIYF  197

Query  792  LMCRQLVTRKKHELWFVFGNKPLRFSMQEFYAVTGLKYE  908
            L+ RQLV RKK E+W +F   P+RFS+ EF  VTGL  +
Sbjct  198  LLTRQLVVRKKQEIWVIFSGSPVRFSISEFQNVTGLNCD  236


>F22O13.23 [Arabidopsis thaliana]
Sequence ID: AAF99775.1 Length: 1014
Range 1: 28 to 108

Score:68.2 bits(165), Expect:3e-11,
Method:Compositional matrix adjust.,
Identities:34/82(41%), Positives:47/82(57%), Gaps:1/82(1%)

Query  663  SILQRIKKALPLEYKEVKSDPQFAHVFAIFESRLGYSARTVHSLMCRQLVTRKKHELWFV  842
            S++  I  AL  E  ++    QF  +F   ES   YS + +H L+ RQLV RKK E+W +
Sbjct  28   SLIDDIANALDAEDMKILRKSQFGKLFDFSES-AAYSGKLIHFLLTRQLVVRKKQEIWVI  86

Query  843  FGNKPLRFSMQEFYAVTGLKYE  908
            F   P+RFS+ EF  VTGL  +
Sbjct  87   FSGSPVRFSISEFQNVTGLNCD  108


>putative protein [Arabidopsis thaliana]
Sequence ID: CAB86452.1 Length: 1009
Range 1: 21 to 115

Score:67.4 bits(163), Expect:5e-11,
Method:Compositional matrix adjust.,
Identities:37/101(37%), Positives:57/101(56%), Gaps:12/101(11%)

Query  726   QFAHVFAIFESRLGYSARTVHSLMCRQLVTRKKHELWFVFGNKPLRFSMQEFYAVTGLKY  905
             QF  +F +  +R  +SA+  H ++ RQL+T++KHELW V+G  PLRFS++EF   TGL
Sbjct  21    QFKKLFELPVARCSHSAKLSHGMLARQLLTQRKHELWTVYGGYPLRFSLKEFQITTGLNC  80

Query  906   EDDFRHDLDS*RDDK------GFWSKLLKKGNTICIKTFMD  1010
             +   +   DS  +D         W++L  +    C+ T MD
Sbjct  81    D---KLPTDSEVEDHQDPAYLSVWNRLFGEK---CVVTVMD  115


>T9E19.2 gene product [Arabidopsis thaliana]
Sequence ID: AAC72876.1 Length: 374
Range 1: 28 to 88

Score:66.2 bits(160), Expect:5e-11,
Method:Compositional matrix adjust.,
Identities:27/61(44%), Positives:43/61(70%), Gaps:0/61(0%)

Query  720  DPQFAHVFAIFESRLGYSARTVHSLMCRQLVTRKKHELWFVFGNKPLRFSMQEFYAVTGL  899
            D  F  +F I +++  ++A+ V  L+CRQLVT+K +E+W VFG  P+RF ++EF  +TGL
Sbjct  28   DSPFGELFKIPQNKASFNAKLVLGLICRQLVTKKVNEMWIVFGGHPIRFGLREFSILTGL  87

Query  900  K  902
            +
Sbjct  88   E  88


>Contains similarity to a putative protein T32A11_100 gi|7413627 from Arabidopsis thaliana BAC T32A11 gb|AL138653 [Arabidopsis thaliana]
Sequence ID: AAF78272.1 Length: 1075
Range 1: 62 to 118

Score:67.0 bits(162), Expect:6e-11,
Method:Compositional matrix adjust.,
Identities:29/57(51%), Positives:40/57(70%), Gaps:0/57(0%)

Query  729  FAHVFAIFESRLGYSARTVHSLMCRQLVTRKKHELWFVFGNKPLRFSMQEFYAVTGL  899
            F  +F +  S+   S + +H+L+C QLVT+KK+ELW VFG  PLRF++ EF  VTGL
Sbjct  62   FGPLFHLPVSKCAMSGKLMHALLCTQLVTKKKYELWTVFGGYPLRFALPEFACVTGL  118


>transmembrane protein, putative (DUF1985) [Arabidopsis thaliana]
Sequence ID: NP_174889.1 Length: 439
>transmembrane protein, putative (DUF1985) [Arabidopsis thaliana]
Sequence ID: AEE31883.1 Length: 439
Range 1: 51 to 130

Score:65.9 bits(159), Expect:7e-11,
Method:Compositional matrix adjust.,
Identities:35/80(44%), Positives:52/80(65%), Gaps:9/80(11%)

Query  705  KEVKS--DPQFAHVFAIFESRLGYSARTVHSLMCRQLVTRKKHELWFVFGNKPLRFSMQE  878
            KEV++  D  F  +F++  S    S + VH+L+CRQLVT++K+EL  +FG +PLRFS+ E
Sbjct  51   KEVETILDSCFGSLFSLRVSECSISCKLVHALLCRQLVTKQKYELKMIFGGQPLRFSLVE  110

Query  879  FYAVTGL-------KYEDDF  917
            F  +TGL       +Y+ DF
Sbjct  111  FGYLTGLPCGEFPKEYDPDF  130


>unnamed protein product [Arabidopsis thaliana]
Sequence ID: BAB10789.1 Length: 442
Range 1: 100 to 160

Score:65.9 bits(159), Expect:9e-11,
Method:Compositional matrix adjust.,
Identities:27/61(44%), Positives:43/61(70%), Gaps:0/61(0%)

Query  720  DPQFAHVFAIFESRLGYSARTVHSLMCRQLVTRKKHELWFVFGNKPLRFSMQEFYAVTGL  899
            D  F  +F I +++  ++A+ V  L+CRQLVT+K +E+W VFG  P+RF ++EF  +TGL
Sbjct  100  DSPFGELFKIPKNKASFNAKLVLGLICRQLVTKKVNEMWIVFGGHPIRFGLREFSILTGL  159

Query  900  K  902
            +
Sbjct  160  E  160


>T32E20.31 [Arabidopsis thaliana]
Sequence ID: AAF79792.1 Length: 416
Range 1: 51 to 117

Score:65.5 bits(158), Expect:9e-11,
Method:Compositional matrix adjust.,
Identities:32/67(48%), Positives:47/67(70%), Gaps:2/67(2%)

Query  705  KEVKS--DPQFAHVFAIFESRLGYSARTVHSLMCRQLVTRKKHELWFVFGNKPLRFSMQE  878
            KEV++  D  F  +F++  S    S + VH+L+CRQLVT++K+EL  +FG +PLRFS+ E
Sbjct  51   KEVETILDSCFGSLFSLRVSECSISCKLVHALLCRQLVTKQKYELKMIFGGQPLRFSLVE  110

Query  879  FYAVTGL  899
            F  +TGL
Sbjct  111  FGYLTGL  117


>unnamed protein product [Arabidopsis thaliana]
Sequence ID: BAB02447.1 Length: 989
Range 1: 12 to 108

Score:66.2 bits(160), Expect:1e-10,
Method:Compositional matrix adjust.,
Identities:36/98(37%), Positives:54/98(55%), Gaps:5/98(5%)

Query  624  RAFCWKNQQLLQN----SILQRIKKALPLEYKEVKSDPQFAHVFAIFESRLGYSARTVHS  791
            RA    N + +Q+    S++  I  AL  +  ++    QF  +F I E    YS + +H
Sbjct  12   RAGSEPNGERVQSYNELSLIDDIANALDEDDMKILRKSQFGKLFDIPEG-AAYSGKLIHF  70

Query  792  LMCRQLVTRKKHELWFVFGNKPLRFSMQEFYAVTGLKY  905
            L+ RQLV  K++E+W +F   P+RFS+ EF  VTGL Y
Sbjct  71   LLTRQLVVSKENEIWVIFSGSPIRFSISEFERVTGLNY  108


>hypothetical protein [Arabidopsis thaliana]
Sequence ID: AAD24651.1 Length: 1472
Range 1: 100 to 160

Score:66.2 bits(160), Expect:1e-10,
Method:Composition-based stats.,
Identities:27/61(44%), Positives:43/61(70%), Gaps:0/61(0%)

Query  720  DPQFAHVFAIFESRLGYSARTVHSLMCRQLVTRKKHELWFVFGNKPLRFSMQEFYAVTGL  899
            D  F  +F I +++  ++A+ V  L+CRQLVT+K +E+W VFG  P+RF ++EF  +TGL
Sbjct  100  DSPFGELFKIPKNKASFNAKLVLGLICRQLVTKKVNEMWIVFGGHPIRFGLREFIILTGL  159

Query  900  K  902
            +
Sbjct  160  E  160


>hypothetical protein [Arabidopsis thaliana]
Sequence ID: AAD14458.1 Length: 1285
>hypothetical protein [Arabidopsis thaliana]
Sequence ID: CAB77815.1 Length: 1285
Range 1: 252 to 312

Score:66.2 bits(160), Expect:1e-10,
Method:Compositional matrix adjust.,
Identities:27/61(44%), Positives:43/61(70%), Gaps:0/61(0%)

Query  720  DPQFAHVFAIFESRLGYSARTVHSLMCRQLVTRKKHELWFVFGNKPLRFSMQEFYAVTGL  899
            D  F  +F I +++  ++A+ V  L+CRQLVT+K +E+W VFG  P+RF ++EF  +TGL
Sbjct  252  DSPFGELFKIPQNKASFNAKLVLGLICRQLVTKKVNEMWIVFGGHPIRFGLREFSILTGL  311

Query  900  K  902
            +
Sbjct  312  E  312


>hypothetical protein AXX17_AT1G37570 [Arabidopsis thaliana]
Sequence ID: OAP14713.1 Length: 439
Range 1: 51 to 130

Score:65.1 bits(157), Expect:1e-10,
Method:Compositional matrix adjust.,
Identities:35/80(44%), Positives:52/80(65%), Gaps:9/80(11%)

Query  705  KEVKS--DPQFAHVFAIFESRLGYSARTVHSLMCRQLVTRKKHELWFVFGNKPLRFSMQE  878
            KEV++  D  F  +F++  S    S + VH+L+CRQLVT++K+EL  +FG +PLRFS+ E
Sbjct  51   KEVETILDSCFGSLFSLRVSECSISCKLVHALLCRQLVTKQKYELKTIFGGQPLRFSLVE  110

Query  879  FYAVTGL-------KYEDDF  917
            F  +TGL       +Y+ DF
Sbjct  111  FGYLTGLPCGEFPKEYDPDF  130


>T22C5.24 [Arabidopsis thaliana]
Sequence ID: AAF24952.1 Length: 1444
Range 1: 100 to 160

Score:65.9 bits(159), Expect:1e-10,
Method:Composition-based stats.,
Identities:27/61(44%), Positives:43/61(70%), Gaps:0/61(0%)

Query  720  DPQFAHVFAIFESRLGYSARTVHSLMCRQLVTRKKHELWFVFGNKPLRFSMQEFYAVTGL  899
            D  F  +F I +++  ++A+ V  L+CRQLVT+K +E+W VFG  P+RF ++EF  +TGL
Sbjct  100  DSPFGKLFKIPKNKASFNAKLVLGLICRQLVTKKVNEMWIVFGGHPIRFGLREFSILTGL  159

Query  900  K  902
            +
Sbjct  160  E  160


>unnamed protein product [Arabidopsis thaliana]
Sequence ID: VYS48136.1 Length: 397
Range 1: 51 to 117

Score:64.7 bits(156), Expect:2e-10,
Method:Compositional matrix adjust.,
Identities:32/67(48%), Positives:47/67(70%), Gaps:2/67(2%)

Query  705  KEVKS--DPQFAHVFAIFESRLGYSARTVHSLMCRQLVTRKKHELWFVFGNKPLRFSMQE  878
            KEV++  D  F  +F++  S    S + VH+L+CRQLVT++K+EL  +FG +PLRFS+ E
Sbjct  51   KEVETILDSCFGSLFSLRVSECSISCKLVHALLCRQLVTKQKYELKTIFGGQPLRFSLVE  110

Query  879  FYAVTGL  899
            F  +TGL
Sbjct  111  FGYLTGL  117


>unnamed protein product [Arabidopsis thaliana]
Sequence ID: CAA0270146.1 Length: 429
Range 1: 51 to 117

Score:64.7 bits(156), Expect:2e-10,
Method:Compositional matrix adjust.,
Identities:32/67(48%), Positives:47/67(70%), Gaps:2/67(2%)

Query  705  KEVKS--DPQFAHVFAIFESRLGYSARTVHSLMCRQLVTRKKHELWFVFGNKPLRFSMQE  878
            KEV++  D  F  +F++  S    S + VH+L+CRQLVT++K+EL  +FG +PLRFS+ E
Sbjct  51   KEVETILDSCFGSLFSLRVSECSISCKLVHALLCRQLVTKQKYELKTIFGGQPLRFSLVE  110

Query  879  FYAVTGL  899
            F  +TGL
Sbjct  111  FGYLTGL  117


>hypothetical protein [Arabidopsis thaliana]
Sequence ID: AAG50521.1 Length: 1201
Range 1: 100 to 160

Score:65.5 bits(158), Expect:2e-10,
Method:Compositional matrix adjust.,
Identities:27/61(44%), Positives:43/61(70%), Gaps:0/61(0%)

Query  720  DPQFAHVFAIFESRLGYSARTVHSLMCRQLVTRKKHELWFVFGNKPLRFSMQEFYAVTGL  899
            D  F  +F I +++  ++A+ V  L+CRQLVT+K +E+W VFG  P+RF ++EF  +TGL
Sbjct  100  DSPFGKLFKIPKNKASFNAKLVLGLICRQLVTKKVNEMWIVFGGHPIRFGLREFSILTGL  159

Query  900  K  902
            +
Sbjct  160  E  160


>F21H2.5 [Arabidopsis thaliana]
Sequence ID: AAD46015.1 Length: 1132
Range 1: 100 to 160

Score:65.5 bits(158), Expect:2e-10,
Method:Compositional matrix adjust.,
Identities:27/61(44%), Positives:43/61(70%), Gaps:0/61(0%)

Query  720  DPQFAHVFAIFESRLGYSARTVHSLMCRQLVTRKKHELWFVFGNKPLRFSMQEFYAVTGL  899
            D  F  +F I +++  ++A+ V  L+CRQLVT+K +E+W VFG  P+RF ++EF  +TGL
Sbjct  100  DSPFGKLFKIPKNKASFNAKLVLGLICRQLVTKKVNEMWIVFGGHPIRFGLREFSILTGL  159

Query  900  K  902
            +
Sbjct  160  E  160


>hypothetical protein [Arabidopsis thaliana]
Sequence ID: AAG50582.1 Length: 1198
Range 1: 100 to 160

Score:65.5 bits(158), Expect:2e-10,
Method:Compositional matrix adjust.,
Identities:27/61(44%), Positives:43/61(70%), Gaps:0/61(0%)

Query  720  DPQFAHVFAIFESRLGYSARTVHSLMCRQLVTRKKHELWFVFGNKPLRFSMQEFYAVTGL  899
            D  F  +F I +++  ++A+ V  L+CRQLVT+K +E+W VFG  P+RF ++EF  +TGL
Sbjct  100  DSPFGKLFKIPKNKASFNAKLVLGLICRQLVTKKVNEMWIVFGGHPIRFGLREFSILTGL  159

Query  900  K  902
            +
Sbjct  160  E  160


>unnamed protein product [Arabidopsis thaliana]
Sequence ID: BAB02940.1 Length: 1119
Range 1: 100 to 160

Score:64.7 bits(156), Expect:3e-10,
Method:Compositional matrix adjust.,
Identities:27/61(44%), Positives:43/61(70%), Gaps:0/61(0%)

Query  720  DPQFAHVFAIFESRLGYSARTVHSLMCRQLVTRKKHELWFVFGNKPLRFSMQEFYAVTGL  899
            D  F  +F I +++  ++A+ V  L+CRQLVT+K +E+W VFG  P+RF ++EF  +TGL
Sbjct  100  DSPFGKLFKIPKNKASFNAKLVLGLICRQLVTKKVNEMWIVFGGHPIRFRLREFSILTGL  159

Query  900  K  902
            +
Sbjct  160  E  160


>putative protein [Arabidopsis thaliana]
Sequence ID: CAB81580.1 Length: 1113
Range 1: 100 to 160

Score:64.3 bits(155), Expect:4e-10,
Method:Compositional matrix adjust.,
Identities:26/61(43%), Positives:43/61(70%), Gaps:0/61(0%)

Query  720  DPQFAHVFAIFESRLGYSARTVHSLMCRQLVTRKKHELWFVFGNKPLRFSMQEFYAVTGL  899
            D  F  +F I +++  ++A+ V  L+CRQLVT+K +E+W VFG  P+RF ++EF  ++GL
Sbjct  100  DSPFGELFKIPKNKASFNAKLVLGLICRQLVTKKVNEMWIVFGGHPIRFGLREFSILSGL  159

Query  900  K  902
            +
Sbjct  160  E  160


>unnamed protein product [Arabidopsis thaliana]
Sequence ID: BAA98062.1 Length: 1444
Range 1: 103 to 160

Score:64.3 bits(155), Expect:4e-10,
Method:Composition-based stats.,
Identities:26/58(45%), Positives:42/58(72%), Gaps:0/58(0%)

Query  729  FAHVFAIFESRLGYSARTVHSLMCRQLVTRKKHELWFVFGNKPLRFSMQEFYAVTGLK  902
            F  +F I +++  ++A+ V  L+CRQLVT+K +E+W VFG  P+RF ++EF  +TGL+
Sbjct  103  FGKLFKIPKNKASFNAKLVLGLICRQLVTKKVNEMWIVFGGHPIRFGLREFSILTGLE  160


>putative protein [Arabidopsis thaliana]
Sequence ID: CAB81070.1 Length: 1312
Range 1: 100 to 160

Score:63.5 bits(153), Expect:9e-10,
Method:Compositional matrix adjust.,
Identities:26/61(43%), Positives:43/61(70%), Gaps:0/61(0%)

Query  720  DPQFAHVFAIFESRLGYSARTVHSLMCRQLVTRKKHELWFVFGNKPLRFSMQEFYAVTGL  899
            D  F  +F I +++  ++A+ V  L+CR+LVT+K +E+W VFG  P+RF ++EF  +TGL
Sbjct  100  DSPFGELFKIPKNKASFNAKLVLRLICRKLVTKKVNEMWIVFGGHPIRFGLREFSILTGL  159

Query  900  K  902
            +
Sbjct  160  E  160


>transmembrane protein, putative (DUF1985) [Arabidopsis thaliana]
Sequence ID: NP_178732.1 Length: 452
>hypothetical protein [Arabidopsis thaliana]
Sequence ID: AAC69122.1 Length: 452 >transmembrane protein, putative (DUF1985) [Arabidopsis thaliana]
Sequence ID: AEC06040.1 Length: 452
Range 1: 13 to 57

Score:45.4 bits(106), Expect:1e-09,
Method:Compositional matrix adjust.,
Identities:23/46(50%), Positives:30/46(65%), Gaps:2/46(4%)

Query  948   KGFWSKLLKKGNTICIKTFMDKQLSEVHMWDEV-DRVRFVYLCVIA  1082
             K FWS+LLK+   ICI+  M   + +V  W E  D +RFVY+CVIA
Sbjct  13    KDFWSRLLKRDGNICIRKLM-TDIEDVKKWSEKKDIIRFVYICVIA  57


Range 2: 59 to 95

Score:38.9 bits(89), Expect:1e-09,
Method:Compositional matrix adjust.,
Identities:17/37(46%), Positives:25/37(67%), Gaps:0/37(0%)

Query  1085  IVVARDEKKNIKVKYIKLVMDLEKLRAYHWGLHSFES  1195
             +V+A  E K I   YIK+VMDL  +R Y WGL ++++
Sbjct  59    LVMATSETKKIDHLYIKMVMDLHNVRNYPWGLKAYDN  95


>unnamed protein product [Arabidopsis thaliana]
Sequence ID: VYS52190.1 Length: 452
Range 1: 13 to 57

Score:45.1 bits(105), Expect:1e-09,
Method:Compositional matrix adjust.,
Identities:23/46(50%), Positives:30/46(65%), Gaps:2/46(4%)

Query  948   KGFWSKLLKKGNTICIKTFMDKQLSEVHMWDEV-DRVRFVYLCVIA  1082
             K FWS+LLK+   ICI+  M   + +V  W E  D +RFVY+CVIA
Sbjct  13    KDFWSRLLKRDGNICIRKLM-TDIEDVKKWSEKKDIIRFVYICVIA  57


Range 2: 59 to 95

Score:38.9 bits(89), Expect:1e-09,
Method:Compositional matrix adjust.,
Identities:17/37(46%), Positives:25/37(67%), Gaps:0/37(0%)

Query  1085  IVVARDEKKNIKVKYIKLVMDLEKLRAYHWGLHSFES  1195
             +V+A  E K I   YIK+VMDL  +R Y WGL ++++
Sbjct  59    LVMATSETKKIDHLYIKMVMDLHNVRNYPWGLKAYDN  95


>hypothetical protein AXX17_AT2G06010 [Arabidopsis thaliana]
Sequence ID: OAP10666.1 Length: 452
Range 1: 13 to 57

Score:45.1 bits(105), Expect:1e-09,
Method:Compositional matrix adjust.,
Identities:23/46(50%), Positives:30/46(65%), Gaps:2/46(4%)

Query  948   KGFWSKLLKKGNTICIKTFMDKQLSEVHMWDEV-DRVRFVYLCVIA  1082
             K FWS+LLK+   ICI+  M   + +V  W E  D +RFVY+CVIA
Sbjct  13    KDFWSRLLKRDGNICIRKLM-TDIEDVKKWSEKKDIIRFVYICVIA  57


Range 2: 59 to 95

Score:38.9 bits(89), Expect:1e-09,
Method:Compositional matrix adjust.,
Identities:17/37(46%), Positives:25/37(67%), Gaps:0/37(0%)

Query  1085  IVVARDEKKNIKVKYIKLVMDLEKLRAYHWGLHSFES  1195
             +V+A  E K I   YIK+VMDL  +R Y WGL ++++
Sbjct  59    LVMATSETKKIDHLYIKMVMDLHNVRNYPWGLKAYDN  95


>putative protein [Arabidopsis thaliana]
Sequence ID: CAB86693.1 Length: 1314
Range 1: 100 to 160

Score:62.4 bits(150), Expect:2e-09,
Method:Compositional matrix adjust.,
Identities:26/61(43%), Positives:42/61(68%), Gaps:0/61(0%)

Query  720  DPQFAHVFAIFESRLGYSARTVHSLMCRQLVTRKKHELWFVFGNKPLRFSMQEFYAVTGL  899
            D  F  +F I +++  ++A+ V  L+C QLVT+K +E+W VFG  P+RF ++EF  +TGL
Sbjct  100  DSPFGELFKIPKNKASFNAKLVLGLICCQLVTKKVNEMWIVFGGHPIRFGLREFSILTGL  159

Query  900  K  902
            +
Sbjct  160  E  160


>Strong similarity to gi|3252818 F26C24.9 hypothetical protein from Arabidopsis thaliana BAC gb|AC004705 [Arabidopsis thaliana]
Sequence ID: AAD12676.1 Length: 1305
Range 1: 100 to 160

Score:62.0 bits(149), Expect:2e-09,
Method:Compositional matrix adjust.,
Identities:26/61(43%), Positives:42/61(68%), Gaps:0/61(0%)

Query  720  DPQFAHVFAIFESRLGYSARTVHSLMCRQLVTRKKHELWFVFGNKPLRFSMQEFYAVTGL  899
            D  F  +F I +++  ++A+ V  L+CRQLVT+K +E+W VFG  P+RF ++EF  +T L
Sbjct  100  DSPFGELFKIPKNKASFNAKLVLGLICRQLVTKKVNEMWIVFGGHPIRFGLREFSILTRL  159

Query  900  K  902
            +
Sbjct  160  E  160


>hypothetical protein [Arabidopsis thaliana]
Sequence ID: AAD29765.1 Length: 917
>hypothetical protein [Arabidopsis thaliana]
Sequence ID: CAB77921.1 Length: 917
Range 1: 104 to 163

Score:61.2 bits(147), Expect:2e-09,
Method:Compositional matrix adjust.,
Identities:26/60(43%), Positives:38/60(63%), Gaps:0/60(0%)

Query  723  PQFAHVFAIFESRLGYSARTVHSLMCRQLVTRKKHELWFVFGNKPLRFSMQEFYAVTGLK  902
            P F+  F     +  +S + VH ++CR L TRKKHE+WFVF  +P RF ++EF  ++GL
Sbjct  104  PCFSPFFQFEVRKCSFSGKLVHQMLCRGLYTRKKHEIWFVFAGQPFRFLLREFAILSGLN  163


Range 2: 61 to 81

Score:21.6 bits(44), Expect:2e-09,
Method:Compositional matrix adjust.,
Identities:9/21(43%), Positives:12/21(57%), Gaps:0/21(0%)

Query  598  PKRLFRHGREPFVGKINNFYK  660
            PKRL+     P  G+IN + K
Sbjct  61   PKRLYATDCYPLNGRINTYSK  81


>unnamed protein product [Arabidopsis thaliana]
Sequence ID: CAA0364214.1 Length: 755
Range 1: 81 to 170

Score:61.2 bits(147), Expect:3e-09,
Method:Compositional matrix adjust.,
Identities:33/90(37%), Positives:49/90(54%), Gaps:6/90(6%)

Query  783   VHSLMCRQLVTRKKHELWFVFGNKPLRFSMQEFYAVTGL-----KYEDDFRHDLDS*RDD  947
             VH L+CRQL T + +  WFVFG +PLRFS+ EF  VTGL         + +  L     +
Sbjct  81    VHQLLCRQLYTDEPNATWFVFGGQPLRFSLMEFEEVTGLCCSEYPPPSEMKSVLSYPDGE  140

Query  948   KGFWSKLL-KKGNTICIKTFMDKQLSEVHM  1034
               +W KL+  K  ++ +K+ + +  SE  M
Sbjct  141   SPYWYKLIGGKLGSVTVKSLLLRLKSEPQM  170


>hypothetical protein [Arabidopsis thaliana]
Sequence ID: AAC24188.1 Length: 1756
Range 1: 100 to 160

Score:61.2 bits(147), Expect:4e-09,
Method:Composition-based stats.,
Identities:25/61(41%), Positives:41/61(67%), Gaps:0/61(0%)

Query  720  DPQFAHVFAIFESRLGYSARTVHSLMCRQLVTRKKHELWFVFGNKPLRFSMQEFYAVTGL  899
            D  F  +F I +++  ++A+ V  L+CRQ VT+K +E+W VFG  P+RF ++EF  +T L
Sbjct  100  DSSFGKLFKIPKNKASFNAKLVLGLICRQFVTKKVNEMWIVFGGHPIRFGLREFSILTRL  159

Query  900  K  902
            +
Sbjct  160  E  160


>hypothetical protein [Arabidopsis thaliana]
Sequence ID: AAD24662.1 Length: 1153
Range 1: 79 to 139

Score:60.5 bits(145), Expect:7e-09,
Method:Compositional matrix adjust.,
Identities:24/61(39%), Positives:40/61(65%), Gaps:0/61(0%)

Query  720  DPQFAHVFAIFESRLGYSARTVHSLMCRQLVTRKKHELWFVFGNKPLRFSMQEFYAVTGL  899
            D  F  +F I  ++  +S + V  L+CRQLVT+K++E+W VF   P+R  ++E+  +TGL
Sbjct  79   DSPFGDLFRIPTNKASFSGKLVLGLICRQLVTKKRYEMWMVFAGHPIRLGLREWALITGL  138

Query  900  K  902
            +
Sbjct  139  E  139


>T4B21.18 gene product [Arabidopsis thaliana]
Sequence ID: AAD03457.1 Length: 253
>hypothetical protein [Arabidopsis thaliana]
Sequence ID: CAB80847.1 Length: 253
Range 1: 37 to 133

Score:58.5 bits(140), Expect:7e-09,
Method:Compositional matrix adjust.,
Identities:35/102(34%), Positives:54/102(52%), Gaps:5/102(4%)

Query  663  SILQRIKKALPLEYKEVKSDPQFAHVFAIFESRLGYSARTVHSLMCRQLVTRKKHELWFV  842
            +I + I K +  E KE     +   +  + +S   +S +TVH L+CRQL   KK ELW +
Sbjct  37   AIRESIGKEIWAELKET----ELGLIAKLVDSHFLWSGKTVHYLLCRQLRILKK-ELWCI  91

Query  843  FGNKPLRFSMQEFYAVTGLKYEDDFRHDLDS*RDDKGFWSKL  968
               KP+RF + EF+ +TGL  +       +   D K F+S+L
Sbjct  92   VAGKPIRFGLNEFHHITGLNIDQLPTEKFEPEADYKAFFSEL  133


>T24H24.2 gene product [Arabidopsis thaliana]
Sequence ID: AAC28200.1 Length: 343
>hypothetical protein [Arabidopsis thaliana]
Sequence ID: CAB77867.1 Length: 343
Range 1: 33 to 113

Score:58.9 bits(141), Expect:9e-09,
Method:Compositional matrix adjust.,
Identities:31/83(37%), Positives:50/83(60%), Gaps:3/83(3%)

Query  666  ILQRIKKAL-PLEYKEVKSDPQFAHVFAIFESRLGYSARTVHSLMCRQLVTRKKHELWFV  842
            I+  ++KAL P+E+  +  D     +    E  L +S+  VH L+ R+++T    ELWFV
Sbjct  33   IVSTVRKALEPIEFNTL-MDTFLGPIVEFAEMDLAFSSHIVHHLLQRRILTNN-DELWFV  90

Query  843  FGNKPLRFSMQEFYAVTGLKYED  911
            F ++P+RFS++EF   TGL  +D
Sbjct  91   FADQPMRFSLREFIITTGLPMDD  113


>hypothetical protein [Arabidopsis thaliana]
Sequence ID: AAD23020.1 Length: 926
Range 1: 37 to 133

Score:59.3 bits(142), Expect:1e-08,
Method:Compositional matrix adjust.,
Identities:35/102(34%), Positives:55/102(53%), Gaps:5/102(4%)

Query  663  SILQRIKKALPLEYKEVKSDPQFAHVFAIFESRLGYSARTVHSLMCRQLVTRKKHELWFV  842
            +I + I + +  E KE     +   +  + +S   +S +TVH L+CRQL   KK ELW +
Sbjct  37   AIRESIGEEIWAELKET----ELGLIAKLVDSHFLWSGKTVHYLLCRQLRILKK-ELWCI  91

Query  843  FGNKPLRFSMQEFYAVTGLKYEDDFRHDLDS*RDDKGFWSKL  968
               KP+RF + EF+ +TGL  +D      +   D K F+S+L
Sbjct  92   VAGKPIRFGLNEFHHITGLNIDDLPTEKFEPEADYKAFFSEL  133


>hypothetical protein AXX17_AT2G12740 [Arabidopsis thaliana]
Sequence ID: OAP08567.1 Length: 847
Range 1: 81 to 170

Score:58.9 bits(141), Expect:2e-08,
Method:Compositional matrix adjust.,
Identities:33/90(37%), Positives:48/90(53%), Gaps:6/90(6%)

Query  783   VHSLMCRQLVTRKKHELWFVFGNKPLRFSMQEFYAVTGL-----KYEDDFRHDLDS*RDD  947
             VH L+CRQL T + +  WFVFG +PLRFS+ EF  VTGL         + +  L     +
Sbjct  81    VHQLLCRQLYTDEPNATWFVFGGQPLRFSLMEFEEVTGLCCSEYPPPSEMKSVLSYPDGE  140

Query  948   KGFWSKLL-KKGNTICIKTFMDKQLSEVHM  1034
               +W KL+  K   + +K+ + +  SE  M
Sbjct  141   SPYWYKLIGGKLGFVTVKSLLLRLKSEPQM  170


>F27F5.16 [Arabidopsis thaliana]
Sequence ID: AAF69170.1 Length: 1745
Range 1: 79 to 139

Score:59.3 bits(142), Expect:2e-08,
Method:Composition-based stats.,
Identities:24/61(39%), Positives:39/61(63%), Gaps:0/61(0%)

Query  720  DPQFAHVFAIFESRLGYSARTVHSLMCRQLVTRKKHELWFVFGNKPLRFSMQEFYAVTGL  899
            D  F  +F I  ++  +S + V  L+CRQLVT K++E+W VF   P+R  ++E+  +TGL
Sbjct  79   DSPFGDLFRIPTNKASFSGKLVLGLICRQLVTNKRYEMWMVFAGHPIRLGLREWALITGL  138

Query  900  K  902
            +
Sbjct  139  E  139


>unnamed protein product [Arabidopsis thaliana]
Sequence ID: BAA97088.1 Length: 200
Range 1: 43 to 125

Score:56.2 bits(134), Expect:2e-08,
Method:Compositional matrix adjust.,
Identities:34/83(41%), Positives:46/83(55%), Gaps:7/83(8%)

Query  705  KEVKSDPQFAHVFAIFESR----LGYSARTVHSLMCRQLVTRKKHELWFVFGNKPLRFSM  872
            KE+ SD  F    A    +    LG+S + V+ L+ RQLV+ KK ELWF   NK +RFS+
Sbjct  43   KEILSDSLFRLRKATGNQKGVNGLGFSGQLVYFLVVRQLVSSKKDELWFCINNKSMRFSL  102

Query  873  QEFYAVTGLKY---EDDFRHDLD  932
             E + VTGL     ED+   D +
Sbjct  103  TEVHLVTGLPCWIKEDEISVDFE  125


>hypothetical protein [Arabidopsis thaliana]
Sequence ID: AAC97230.2 Length: 1422
>hypothetical protein [Arabidopsis thaliana]
Sequence ID: AAM15356.1 Length: 1422
Range 1: 79 to 139

Score:59.3 bits(142), Expect:2e-08,
Method:Composition-based stats.,
Identities:24/61(39%), Positives:39/61(63%), Gaps:0/61(0%)

Query  720  DPQFAHVFAIFESRLGYSARTVHSLMCRQLVTRKKHELWFVFGNKPLRFSMQEFYAVTGL  899
            D  F  +F I  ++  +S + V  L+CRQLVT K++E+W VF   P+R  ++E+  +TGL
Sbjct  79   DSPFGDLFRIPTNKASFSGKLVLGLICRQLVTNKRYEMWMVFAGHPIRLGLREWALITGL  138

Query  900  K  902
            +
Sbjct  139  E  139


>unnamed protein product [Arabidopsis thaliana]
Sequence ID: BAB01421.1 Length: 733
Range 1: 33 to 113

Score:58.5 bits(140), Expect:2e-08,
Method:Compositional matrix adjust.,
Identities:31/83(37%), Positives:50/83(60%), Gaps:3/83(3%)

Query  666  ILQRIKKAL-PLEYKEVKSDPQFAHVFAIFESRLGYSARTVHSLMCRQLVTRKKHELWFV  842
            I+  ++KAL P+E+  +  D     +    E  L +S+  VH L+ R+++T    ELWFV
Sbjct  33   IVSTVRKALEPIEFNTL-MDTFLGPIVEFAEMDLAFSSHIVHHLLQRRILTNN-DELWFV  90

Query  843  FGNKPLRFSMQEFYAVTGLKYED  911
            F ++P+RFS++EF   TGL  +D
Sbjct  91   FADQPMRFSLREFIITTGLPMDD  113


>CDS [Arabidopsis thaliana]
Sequence ID: AAF31285.1 Length: 728
Range 1: 33 to 113

Score:58.2 bits(139), Expect:3e-08,
Method:Compositional matrix adjust.,
Identities:32/83(39%), Positives:50/83(60%), Gaps:3/83(3%)

Query  666  ILQRIKKAL-PLEYKEVKSDPQFAHVFAIFESRLGYSARTVHSLMCRQLVTRKKHELWFV  842
            I+  I+KAL P+E+  +  D     +    E  L +S+  VH L+ R+++T    ELWFV
Sbjct  33   IVSTIRKALEPIEFNTL-MDTFLGPIVEFAEMDLTFSSHIVHHLLQRRILTNN-DELWFV  90

Query  843  FGNKPLRFSMQEFYAVTGLKYED  911
            F ++P+RFS++EF   TGL  +D
Sbjct  91   FADQPMRFSLREFIITTGLPMDD  113


>hypothetical protein AXX17_AT5G31140 [Arabidopsis thaliana]
Sequence ID: OAO92309.1 Length: 248
Range 1: 17 to 99

Score:56.6 bits(135), Expect:3e-08,
Method:Compositional matrix adjust.,
Identities:31/87(36%), Positives:51/87(58%), Gaps:6/87(6%)

Query  645  QQLLQNSILQRIKKAL--PLEYKEVKSDPQFAHVFAIFESRLGYSARTVHSLMCRQLVTR  818
            +++L NS+  R+K     P+    +K+  + A       + LG+S + V  L+ RQLV+
Sbjct  17   KEILSNSLFSRLKTTFLGPI----IKAGLRKAGGNQKGPNGLGFSGQLVRFLVVRQLVSS  72

Query  819  KKHELWFVFGNKPLRFSMQEFYAVTGL  899
            ++  +WF   NKP+RFS+ EF+ VTGL
Sbjct  73   REDGIWFCINNKPMRFSLTEFHLVTGL  99


>hypothetical protein [Arabidopsis thaliana]
Sequence ID: AAD19785.1 Length: 833
Range 1: 29 to 107

Score:57.8 bits(138), Expect:5e-08,
Method:Compositional matrix adjust.,
Identities:29/80(36%), Positives:46/80(57%), Gaps:1/80(1%)

Query  663  SILQRIKKALPLEYKEVKSDPQFAHVFAIFESRLGYSARTVHSLMCRQLVTRKKHELWFV  842
            S++  I  AL  E  ++  + QF  +F I E    YS + +H L+ RQLV  K++++W +
Sbjct  29   SLIDNIADALDEEDMKILRESQFGKLFDIPEG-AAYSGKLIHFLLTRQLVVSKQNKIWVI  87

Query  843  FGNKPLRFSMQEFYAVTGLK  902
            F   P++FS+ EF  VT L
Sbjct  88   FSGSPIQFSISEFKRVTRLN  107


>unnamed protein product [Arabidopsis thaliana]
Sequence ID: CAA0405522.1 Length: 209
Range 1: 80 to 125

Score:55.1 bits(131), Expect:6e-08,
Method:Compositional matrix adjust.,
Identities:23/46(50%), Positives:33/46(71%), Gaps:0/46(0%)

Query  762  LGYSARTVHSLMCRQLVTRKKHELWFVFGNKPLRFSMQEFYAVTGL  899
            LG+S + V  L+ RQLV+ ++  +WF   NKP+RFS+ EF+ VTGL
Sbjct  80   LGFSGQLVRFLVVRQLVSSREDGIWFCINNKPMRFSLTEFHLVTGL  125


>hypothetical protein (DUF1985) [Arabidopsis thaliana]
Sequence ID: NP_001154746.1 Length: 209
>hypothetical protein (DUF1985) [Arabidopsis thaliana]
Sequence ID: AED93925.1 Length: 209
Range 1: 80 to 125

Score:54.7 bits(130), Expect:7e-08,
Method:Compositional matrix adjust.,
Identities:23/46(50%), Positives:33/46(71%), Gaps:0/46(0%)

Query  762  LGYSARTVHSLMCRQLVTRKKHELWFVFGNKPLRFSMQEFYAVTGL  899
            LG+S + V  L+ RQLV+ ++  +WF   NKP+RFS+ EF+ VTGL
Sbjct  80   LGFSGQLVRFLVVRQLVSSREDGIWFCINNKPMRFSLTEFHLVTGL  125


Query #200: XLOC_020658 Query ID: lcl|Query_33203 Length: 514

Sequences producing significant alignments:
                                                                  Max    Total Query   E   Per.
Description                                                       Score  Score cover Value Ident  Accession
hypothetical protein AXX17_ATUG01420 [Arabidopsis thaliana]       74.3   74.3  22%   1e-17 100.00 OAO89359.1
TSA1 [Arabidopsis thaliana]                                       60.5   60.5  21%   1e-10 83.78  OAP14298.1
unnamed protein product [Arabidopsis thaliana]                    58.5   58.5  21%   8e-10 81.08  VYS48873.1
unnamed protein product [Arabidopsis thaliana]                    58.2   58.2  21%   9e-10 81.08  CAA0289292.1
TSK-associating protein 1 [Arabidopsis thaliana]                  58.2   58.2  21%   9e-10 81.08  NP_849797.1
TSK-associating protein 1 [Arabidopsis thaliana]                  58.2   58.2  21%   1e-09 81.08  NP_564607.1
At1g52410/F19K6_14 [Arabidopsis thaliana]                         58.2   58.2  21%   1e-09 81.08  AAK82509.1
myosin-like protein; 12311-7712 [Arabidopsis thaliana]            57.8   57.8  21%   1e-09 81.08  AAG51543.1
putative protein [Arabidopsis thaliana]                           53.1   53.1  15%   5e-08 88.89  CAB91582.1


Alignments:
>hypothetical protein AXX17_ATUG01420 [Arabidopsis thaliana]
Sequence ID: OAO89359.1 Length: 58
Range 1: 21 to 58

Score:74.3 bits(181), Expect:1e-17,
Method:Compositional matrix adjust.,
Identities:38/38(100%), Positives:38/38(100%), Gaps:0/38(0%)

Query  1    FEAIESEFEAAMEGIEELKVSDSIGSGDDEEQCASGLL  114
            FEAIESEFEAAMEGIEELKVSDSIGSGDDEEQCASGLL
Sbjct  21   FEAIESEFEAAMEGIEELKVSDSIGSGDDEEQCASGLL  58


>TSA1 [Arabidopsis thaliana]
Sequence ID: OAP14298.1 Length: 757
Range 1: 223 to 259

Score:60.5 bits(145), Expect:1e-10,
Method:Compositional matrix adjust.,
Identities:31/37(84%), Positives:31/37(83%), Gaps:0/37(0%)

Query  1    FEAIESEFEAAMEGIEELKVSDSIGSGDDEEQCASGL  111
             EAIE EFEAAMEGIEELKVSDS GSGDDEEQ A  L
Sbjct  223  LEAIEREFEAAMEGIEELKVSDSTGSGDDEEQSAKRL  259


>unnamed protein product [Arabidopsis thaliana]
Sequence ID: VYS48873.1 Length: 755
Range 1: 219 to 255

Score:58.5 bits(140), Expect:8e-10,
Method:Compositional matrix adjust.,
Identities:30/37(81%), Positives:30/37(81%), Gaps:0/37(0%)

Query  1    FEAIESEFEAAMEGIEELKVSDSIGSGDDEEQCASGL  111
             EAIE EFEAAMEGIE LKVSDS GSGDDEEQ A  L
Sbjct  219  LEAIEREFEAAMEGIEALKVSDSTGSGDDEEQSAKRL  255


>unnamed protein product [Arabidopsis thaliana]
Sequence ID: CAA0289292.1 Length: 755
Range 1: 219 to 255

Score:58.2 bits(139), Expect:9e-10,
Method:Compositional matrix adjust.,
Identities:30/37(81%), Positives:30/37(81%), Gaps:0/37(0%)

Query  1    FEAIESEFEAAMEGIEELKVSDSIGSGDDEEQCASGL  111
             EAIE EFEAAMEGIE LKVSDS GSGDDEEQ A  L
Sbjct  219  LEAIEREFEAAMEGIEALKVSDSTGSGDDEEQSAKRL  255


>TSK-associating protein 1 [Arabidopsis thaliana]
Sequence ID: NP_849797.1 Length: 759
>RecName: Full=TSK-associating protein 1; Flags: Precursor [Arabidopsis thaliana]
Sequence ID: F4ICX9.1 Length: 759 >TSK-associating protein 1 [Arabidopsis thaliana]
Sequence ID: AEE32803.1 Length: 759
Range 1: 223 to 259

Score:58.2 bits(139), Expect:9e-10,
Method:Compositional matrix adjust.,
Identities:30/37(81%), Positives:30/37(81%), Gaps:0/37(0%)

Query  1    FEAIESEFEAAMEGIEELKVSDSIGSGDDEEQCASGL  111
             EAIE EFEAAMEGIE LKVSDS GSGDDEEQ A  L
Sbjct  223  LEAIEREFEAAMEGIEALKVSDSTGSGDDEEQSAKRL  259


>TSK-associating protein 1 [Arabidopsis thaliana]
Sequence ID: NP_564607.1 Length: 755
>TSK-associating protein 1 [Arabidopsis thaliana]
Sequence ID: AEE32802.1 Length: 755
Range 1: 223 to 259

Score:58.2 bits(139), Expect:1e-09,
Method:Compositional matrix adjust.,
Identities:30/37(81%), Positives:30/37(81%), Gaps:0/37(0%)

Query  1    FEAIESEFEAAMEGIEELKVSDSIGSGDDEEQCASGL  111
             EAIE EFEAAMEGIE LKVSDS GSGDDEEQ A  L
Sbjct  223  LEAIEREFEAAMEGIEALKVSDSTGSGDDEEQSAKRL  259


>At1g52410/F19K6_14 [Arabidopsis thaliana]
Sequence ID: AAK82509.1 Length: 755
>At1g52410/F19K6_14 [Arabidopsis thaliana]
Sequence ID: AAO64753.1 Length: 755
Range 1: 223 to 259

Score:58.2 bits(139), Expect:1e-09,
Method:Compositional matrix adjust.,
Identities:30/37(81%), Positives:30/37(81%), Gaps:0/37(0%)

Query  1    FEAIESEFEAAMEGIEELKVSDSIGSGDDEEQCASGL  111
             EAIE EFEAAMEGIE LKVSDS GSGDDEEQ A  L
Sbjct  223  LEAIEREFEAAMEGIEALKVSDSTGSGDDEEQSAKRL  259


>myosin-like protein; 12311-7712 [Arabidopsis thaliana]
Sequence ID: AAG51543.1 Length: 784
Range 1: 223 to 259

Score:57.8 bits(138), Expect:1e-09,
Method:Compositional matrix adjust.,
Identities:30/37(81%), Positives:30/37(81%), Gaps:0/37(0%)

Query  1    FEAIESEFEAAMEGIEELKVSDSIGSGDDEEQCASGL  111
             EAIE EFEAAMEGIE LKVSDS GSGDDEEQ A  L
Sbjct  223  LEAIEREFEAAMEGIEALKVSDSTGSGDDEEQSAKRL  259


>putative protein [Arabidopsis thaliana]
Sequence ID: CAB91582.1 Length: 448
Range 1: 1 to 27

Score:53.1 bits(126), Expect:5e-08,
Method:Compositional matrix adjust.,
Identities:24/27(89%), Positives:25/27(92%), Gaps:0/27(0%)

Query  246  IAATFAYLKDVRRYTTAWRVQVKVFHS  166
            +AATFAYLKDVR YTTAWRVQVKV HS
Sbjct  1    MAATFAYLKDVRPYTTAWRVQVKVLHS  27


Query #201: XLOC_020779 Query ID: lcl|Query_30692 Length: 4940

Sequences producing significant alignments:
                                                                  Max    Total Query   E   Per.
Description                                                       Score  Score cover Value Ident  Accession
hypothetical protein [Arabidopsis thaliana]                       198    333   12%   7e-57 88.46  AAD27899.1
hypothetical protein [Arabidopsis thaliana]                       188    394   21%   3e-53 67.15  AAD20429.1
hypothetical protein AXX17_AT5G29060 [Arabidopsis thaliana]       176    267   20%   3e-47 58.43  OAO90101.1
hypothetical protein AXX17_AT2G08070 [Arabidopsis thaliana]       134    201   12%   2e-30 58.39  OAP07569.1
T32E20.11 [Arabidopsis thaliana]                                  109    109   9%    9e-27 48.78  AAF79783.1
putative protein [Arabidopsis thaliana]                           106    106   8%    3e-23 60.00  CAB91583.1
hypothetical protein [Arabidopsis thaliana]                       94.7   94.7  9%    2e-19 50.00  AAD26889.1
putative protein [Arabidopsis thaliana]                           85.9   85.9  8%    2e-18 38.57  CAB87735.1
hypothetical protein AXX17_AT1G38610 [Arabidopsis thaliana]       87.4   87.4  9%    6e-17 47.77  OAP19085.1
F7F22.14 [Arabidopsis thaliana]                                   83.2   142   12%   3e-16 80.00  AAF24528.1
contains similarity to an Arabidopsis thaliana hypothetical...    74.7   147   10%   5e-14 80.95  AAD48941.1
En/Spm-like transposon protein [Arabidopsis thaliana]             76.6   76.6  4%    2e-13 53.62  AAD03363.1
unnamed protein product [Arabidopsis thaliana]                    74.7   74.7  7%    5e-13 55.83  CAA0394037.1
unnamed protein product [Arabidopsis thaliana]                    67.0   67.0  2%    1e-11 72.50  BAB02612.1
hypothetical protein AXX17_AT1G37440 [Arabidopsis thaliana]       71.2   71.2  3%    2e-11 62.96  OAP14491.1
F10A2.1 gene product [Arabidopsis thaliana]                       67.8   67.8  9%    5e-11 41.29  AAD29783.1
putative TNP2-like transposon protein [Arabidopsis thaliana]      65.9   65.9  2%    9e-10 63.83  AAD20646.1
unnamed protein product [Arabidopsis thaliana]                    64.3   64.3  6%    9e-10 54.39  VYS61969.1
putative transposon protein [Arabidopsis thaliana]                59.7   59.7  2%    9e-09 72.50  AAD27900.1


Alignments:
>hypothetical protein [Arabidopsis thaliana]
Sequence ID: AAD27899.1 Length: 226
>hypothetical protein [Arabidopsis thaliana]
Sequence ID: CAB81149.1 Length: 226
Range 1: 19 to 148

Score:198 bits(504), Expect:7e-57,
Method:Compositional matrix adjust.,
Identities:115/130(88%), Positives:117/130(90%), Gaps:0/130(0%)

Query  735   LFVEDVNYFGGRGYQNQRFKHQRGYKGVYGNVQPNYTqksqfqqpfqksssfsfTRNYDL  914
             LFVEDVNY GGRGYQNQR +HQRG  GVYGN QPNYT KSQFQQPFQ SSSFSFTRNYDL
Sbjct  19    LFVEDVNYVGGRGYQNQRLEHQRGNIGVYGNGQPNYTHKSQFQQPFQYSSSFSFTRNYDL  78

Query  915   ASYQAPPPPEPRSKIESMPEQILEGQKKILERPIPRWTEYRLTPYWHVALMESEDEFLNV  1094
             ASYQAPPPP  RSKIES+PEQILEGQKKILER IPRWTE RLTP WHVALMESEDEFL V
Sbjct  79    ASYQAPPPPAQRSKIESLPEQILEGQKKILERYIPRWTERRLTPSWHVALMESEDEFLKV  138

Query  1095  EESDEFDVVE  1124
             EESDEFDVVE
Sbjct  139   EESDEFDVVE  148


Range 2: 146 to 226

Score:135 bits(339), Expect:1e-34,
Method:Compositional matrix adjust.,
Identities:64/81(79%), Positives:68/81(83%), Gaps:0/81(0%)

Query  1874  VLWLCKLRVSRRALFRSFDDCAGKRSVPPIPESHPADAPLFLDRPHALTAYTPAWMMRGF  2053
             V+ LCKLRV RR L RSF DCAGKRSVPPIP+SHPADAPL LDRP+ALTAYTP WMMRG
Sbjct  146   VVELCKLRVFRRVLVRSFYDCAGKRSVPPIPDSHPADAPLVLDRPNALTAYTPTWMMRGL  205

Query  2054  SPRYLIPPSEPHDYRNPAKSC  2116
             SPRYL+PP EP D RNP KS
Sbjct  206   SPRYLLPPFEPPDSRNPLKSS  226


>hypothetical protein [Arabidopsis thaliana]
Sequence ID: AAD20429.1 Length: 378
Range 1: 70 to 206

Score:188 bits(478), Expect:3e-53,
Method:Compositional matrix adjust.,
Identities:92/137(67%), Positives:105/137(76%), Gaps:0/137(0%)

Query  951   SKIESMPEQILEGQKKILERPIPRWTEYRLTPYWHVALMESEDEFLNVEESDEFDVVEAV  1130
             SKIES+ EQILEGQ++ILERPIPRW E  LT  WH A MESE E + +EESD+F V E V
Sbjct  70    SKIESLLEQILEGQQRILERPIPRWNESSLTTSWHSASMESEYELVKIEESDDFVVAEVV  129

Query  1131  LTHTNPWCRLTPSHIYETSSCDEAKSTRLDIDLSNRSTHIYSVDRHHLGVD*HQYQTVPM  1310
                TNPWCR TPSHIYETS CDEA+S++LDIDL NRSTHI SVDRHHLGVD HQYQ   +
Sbjct  130   SNDTNPWCRSTPSHIYETSGCDEAESSKLDIDLQNRSTHIDSVDRHHLGVDRHQYQPEEV  189

Query  1311  STFDRK*SNEIDRAGSI  1361
               FDR  S +IDR  ++
Sbjct  190   PMFDRAESTKIDRIAAL  206


Range 2: 2 to 58

Score:43.9 bits(102), Expect:3e-53,
Method:Compositional matrix adjust.,
Identities:38/57(67%), Positives:43/57(75%), Gaps:0/57(0%)

Query  748  MSIILVEEVIRIRDLNTNEAIKVSMGMFnlttlrslsssnlsrKAVVSASQETTILL  918
            MSI+LV EV RIRD+ TN+A +V MGM  L TLR LSSSN  R A+VSASQ T ILL
Sbjct  2    MSIMLVAEVFRIRDMITNKATEVPMGMVQLATLRILSSSNRFRIAIVSASQGTMILL  58


Range 3: 201 to 362

Score:161 bits(407), Expect:4e-42,
Method:Compositional matrix adjust.,
Identities:108/166(65%), Positives:120/166(72%), Gaps:4/166(2%)

Query  1574  EHISELNYRHAKGRRVVARVSTNTTTSVDRHPSRHPSLVETKSFNIGTFQARITAKSLGT  1753
             + I+ LN +HAK  RVVA+VS +TTT VDR  S    LVE +S  I TFQA ITA S  T
Sbjct  201   DRIAALNCQHAKESRVVAQVSIDTTTRVDRQTS----LVEEESCPIATFQAGITAISSST  256

Query  1754  TVMSPTIplllhpppkrprnqsppLKPPYVINKSLKFIKTVLWLCKLRVSRRALFRSFDD  1933
             TV S   PL+LHPPPKR R  SPP KPP +INK+LKF KTVL L K RVSRRAL  S DD
Sbjct  257   TVTSHNAPLILHPPPKRLRYSSPPPKPPDIINKTLKFPKTVLRLSKPRVSRRALVLSVDD  316

Query  1934  CAGKRSVPPIPESHPADAPLFLDRPHALTAYTPAWMMRGFSPRYLI  2071
              A KRS+PPIPE +P DAPLFLDRPH L A TP WMMRG SPRYL+
Sbjct  317   YARKRSIPPIPEFYPVDAPLFLDRPHTLIASTPVWMMRGLSPRYLL  362


>hypothetical protein AXX17_AT5G29060 [Arabidopsis thaliana]
Sequence ID: OAO90101.1 Length: 384
Range 1: 121 to 247

Score:176 bits(446), Expect:3e-47,
Method:Compositional matrix adjust.,
Identities:97/166(58%), Positives:105/166(63%), Gaps:39/166(23%)

Query  897   TRNYDLASYQAPPPPEPRSKIESMPEQILEGQKKILERPIPRWTEYRLTPYWHVALMESE  1076
             TRNYDLASYQAPPPP P+SK+ES+ EQILEGQ++ILERPIPRW E  LT  WH A ME
Sbjct  121   TRNYDLASYQAPPPPTPKSKLESLLEQILEGQQRILERPIPRWNESSLTTSWHSASME--  178

Query  1077  DEFLNVEESDEFDVVEAVLTHTNPWCRLTPSHIYETSSCDEAKSTRLDIDLSNRSTHIYS  1256
                                                  SCDEA ST+LDIDL NRSTHI S
Sbjct  179   -------------------------------------SCDEAGSTKLDIDLQNRSTHIDS  201

Query  1257  VDRHHLGVD*HQYQTVPMSTFDRK*SNEIDRAGSIPDSSVDRHSPN  1394
             VDRHH GVD HQYQ   +STFD   S EIDRAGSI DSSVDRHSP+
Sbjct  202   VDRHHFGVDRHQYQPEEVSTFDGAESTEIDRAGSILDSSVDRHSPS  247


Range 2: 255 to 368

Score:90.9 bits(224), Expect:3e-18,
Method:Compositional matrix adjust.,
Identities:65/164(40%), Positives:79/164(48%), Gaps:50/164(30%)

Query  1580  ISELNYRHAKGRRVVARVSTNTTTSVDRHPSRHPSLVETKSFNIGTFQARITAKSLGTTV  1759
              + LN +HAK  RVVA VS +TTT VDRH S    LVE +S  I TFQ       +G T
Sbjct  255   FAALNCQHAKESRVVAHVSIDTTTRVDRHTS----LVEEESSPIATFQ-------VGITA  303

Query  1760  MSPTIplllhpppkrprnqsppLKPPYVINKSLKFIKTVLWLCKLRVSRRALFRSFDDCA  1939
             + P+  +  H               P +++   K +
Sbjct  304   IRPSTTVTSH-------------NAPLILHPPPKRL------------------------  326

Query  1940  GKRSVPPIPESHPADAPLFLDRPHALTAYTPAWMMRGFSPRYLI  2071
               RS+PPI E HPADAPLFLDRPH L A T  WMMRG SPRYL+
Sbjct  327   --RSIPPILEFHPADAPLFLDRPHTLIASTQVWMMRGLSPRYLL  368


>hypothetical protein AXX17_AT2G08070 [Arabidopsis thaliana]
Sequence ID: OAP07569.1 Length: 1891
Range 1: 1708 to 1864

Score:134 bits(336), Expect:2e-30,
Method:Compositional matrix adjust.,
Identities:94/161(58%), Positives:105/161(65%), Gaps:6/161(3%)

Query  1616  RVVARVSTNTTTSVDRHPSRHPSLVETKSFNIGTFQARITAKSLGTTVMSPTIplllhpp  1795
             R  A VS +TTT +DRH ++     E + F    F+   TA + G TV S T P LL PP
Sbjct  1708  RSQAMVSIDTTTCIDRHTTQ----AEPELFIRAGFRVEFTADTPGITVTSTTAPSLLDPP  1763

Query  1796  pkrprnqsppLKPPYVINKSLKFIKTVLWLCKLRVSRRALFRSFDDCAGKRSVPPIPESH  1975
              KR R  SPP K P  INKSLKF KTV    K RVSRRAL  SFDDCAGKRS+P I ESH
Sbjct  1764  HKRLRYSSPPPKTPDFINKSLKFPKTVSRFSKPRVSRRALVCSFDDCAGKRSIPSIHESH  1823

Query  1976  PADAPLFLDRPHALTAYTPAWMMRGFSPRYLI--PPSEPHD  2092
             PAD PLFLDRPHALTAYTPAWMMRG +   L+  P  +P D
Sbjct  1824  PADTPLFLDRPHALTAYTPAWMMRGSTQFRLLKNPDGKPTD  1864


Range 2: 1852 to 1890

Score:67.8 bits(164), Expect:2e-10,
Method:Compositional matrix adjust.,
Identities:32/39(82%), Positives:33/39(84%), Gaps:0/39(0%)

Query  2414  IRLLENPDCKPTDLRISWFPTQSIKAGANLTALPFTSSK  2530
              RLL+NPD KPTDL ISWFPTQS KAGANLTAL FTSS
Sbjct  1852  FRLLKNPDGKPTDLTISWFPTQSTKAGANLTALRFTSSN  1890


>T32E20.11 [Arabidopsis thaliana]
Sequence ID: AAF79783.1 Length: 146
Range 1: 20 to 146

Score:109 bits(272), Expect:9e-27,
Method:Composition-based stats.,
Identities:80/164(49%), Positives:92/164(56%), Gaps:37/164(22%)

Query  738   FVEDVNYFGGRGYQNQRFKHQRGYKGVYGNVQPNYTqksqfqqpfqksssfsfTRNYDLA  917
              VEDV Y   R + NQRF HQ+GYKG YGN Q +Y+Q SQFQQP Q ++SFSFTRN+DLA
Sbjct  20    LVEDVAYVRSRDFPNQRFDHQQGYKGSYGNGQTSYSQNSQFQQPLQNNNSFSFTRNFDLA  79

Query  918   SYQAPPPPEPRSKIESMPEQILEGQKKILERPIPRWTEYRLTPYWHVALMESEDEFLNVE  1097
             SYQ                         L  PIPRW E  LT  WH A MES  EF+  +
Sbjct  80    SYQD------------------------LTAPIPRWNESSLTTSWHAASMESGYEFMKDD  115

Query  1098  ESDEFDVVEAVLTHTNPWCRLTPSHIYETSSCDEAKSTRLDIDL  1229
             ESDEFDV + V T T             +  CDEAKS+RLD DL
Sbjct  116   ESDEFDVAKVVSTDT-------------SLGCDEAKSSRLDTDL  146


>putative protein [Arabidopsis thaliana]
Sequence ID: CAB91583.1 Length: 409
Range 1: 254 to 394

Score:106 bits(264), Expect:3e-23,
Method:Compositional matrix adjust.,
Identities:87/145(60%), Positives:96/145(66%), Gaps:4/145(2%)

Query  1631  VSTNTTTSVDRHPSRHPSLVETKSFNIGTFQARITAKSLGTTVMSPTIplllhpppkrpr  1810
             VS +TTT VDRHP++    VE +      F+   T  +  TTV SPT PLLL PPPKR R
Sbjct  254   VSIDTTTCVDRHPTQ----VEPEFLIRAGFRVEFTTDTPVTTVTSPTAPLLLDPPPKRLR  309

Query  1811  nqsppLKPPYVINKSLKFIKTVLWLCKLRVSRRALFRSFDDCAGKRSVPPIPESHPADAP  1990
               SPP KPP  INKSLKF KTVL L K RVSRRAL  SFDDC+G RS PPIP S  AD P
Sbjct  310   YSSPPPKPPDFINKSLKFPKTVLRLSKPRVSRRALILSFDDCSGNRSRPPIPLSCSADIP  369

Query  1991  LFLDRPHALTAYTPAWMMRGFSPRY  2065
              FL RPH  T YTP WMMRG + +Y
Sbjct  370   HFLGRPHTPTVYTPPWMMRGSTRKY  394


>hypothetical protein [Arabidopsis thaliana]
Sequence ID: AAD26889.1 Length: 419
Range 1: 276 to 403

Score:94.7 bits(234), Expect:2e-19,
Method:Compositional matrix adjust.,
Identities:76/152(50%), Positives:85/152(55%), Gaps:24/152(15%)

Query  1616  RVVARVSTNTTTSVDRHPSRHPSLVETKSFNIGTFQARITAKSLGTTVMSPTIplllhpp  1795
             R  A VS +TTT +DRH ++     E +      F+   TA + G TV SPT P LL PP
Sbjct  276   RSQAMVSIDTTTCIDRHTTQ----AEPEFLIRAGFRVEFTADTPGITVTSPTAPSLLDPP  331

Query  1796  pkrprnqsppLKPPYVINKSLKFIKTVLWLCKLRVSRRALFRSFDDCAGKRSVPPIPESH  1975
              KR RN  PP K P  INKSLKF KTV    K RVSRRAL  SFDDC
Sbjct  332   HKRLRNSFPPPKTPDFINKSLKFPKTVSRFSKPRVSRRALVCSFDDCG------------  379

Query  1976  PADAPLFLDRPHALTAYTPAWMMRGFSPRYLI  2071
                     DRPHALT+YTPAWMMRG S RYL+
Sbjct  380   --------DRPHALTSYTPAWMMRGLSLRYLL  403


>putative protein [Arabidopsis thaliana]
Sequence ID: CAB87735.1 Length: 170
Range 1: 20 to 125

Score:85.9 bits(211), Expect:2e-18,
Method:Compositional matrix adjust.,
Identities:54/140(39%), Positives:67/140(47%), Gaps:34/140(24%)

Query  738   FVEDVNYFGGRGYQNQRFKHQRGYKGVYGNVQPNYTqksqfqqpfqksssfsfTRNYDLA  917
              VE V+Y   RG+ NQRF HQ+GY+G YGN   +YTQ                      +
Sbjct  20    LVEVVDYVSSRGFPNQRFDHQQGYRGSYGNGPTSYTQN---------------------S  58

Query  918   SYQAPPPPEPRSKIESMPEQILEGQKKILERPIPRWTEYRLTPYWHVALMESEDEFLNVE  1097
              +Q P              Q    Q++ILERPIPRW E  LT   H   MESE E + +E
Sbjct  59    QFQQPL-------------QNSNSQQRILERPIPRWNESSLTTSRHSTSMESEYELVKIE  105

Query  1098  ESDEFDVVEAVLTHTNPWCR  1157
             ESD+F V EAV   T  + R
Sbjct  106   ESDDFAVAEAVSIDTIAYIR  125


>hypothetical protein AXX17_AT1G38610 [Arabidopsis thaliana]
Sequence ID: OAP19085.1 Length: 448
Range 1: 324 to 447

Score:87.4 bits(215), Expect:6e-17,
Method:Compositional matrix adjust.,
Identities:75/157(48%), Positives:83/157(52%), Gaps:33/157(21%)

Query  1634  STNTTTSVDRHPSRHPSLVETKSFNIGTFQARITAKSLGTTVMSPTIplllhpppkrprn  1813
             S +TTT VDRHP++     E + F   +F    T  + GT V SPT PLLL PPPKR R
Sbjct  324   SIDTTTYVDRHPTQ----AELEFFIRASFCVEFTVDTPGTIVKSPTAPLLLDPPPKRLRY  379

Query  1814  qsppLKPPYVINKSLKFIKTVLWLCKLRVSRRALFRSFDDCAGKRSVPPIPESHPADAPL  1993
              SPPLKP   INKS                             KRS PPIP    A  P
Sbjct  380   SSPPLKPLDFINKS-----------------------------KRSRPPIPLFCTAKVPH  410

Query  1994  FLDRPHALTAYTPAWMMRGFSPRYLIPPSEPHDYRNP  2104
             FL RPH   AYTP WMMRGFSPRY +PPS+P D RNP
Sbjct  411   FLCRPHTANAYTPPWMMRGFSPRYFLPPSDPPDSRNP  447


>F7F22.14 [Arabidopsis thaliana]
Sequence ID: AAF24528.1 Length: 304
Range 1: 229 to 278

Score:83.2 bits(204), Expect:3e-16,
Method:Compositional matrix adjust.,
Identities:40/50(80%), Positives:42/50(84%), Gaps:0/50(0%)

Query  2402  RFDSIRLLENPDCKPTDLRISWFPTQSIKAGANLTALPFTSSKRLVFIEL  2551
             RFDSIR LENPD K T+LRISWFPTQSIKAGANLTALPFTSS  L   E+
Sbjct  229   RFDSIRQLENPDGKSTNLRISWFPTQSIKAGANLTALPFTSSNGLYMFEI  278


Range 2: 71 to 216

Score:58.9 bits(141), Expect:3e-08,
Method:Compositional matrix adjust.,
Identities:67/159(42%), Positives:80/159(50%), Gaps:21/159(13%)

Query  1625  ARVSTNTTTSVDRHPSRHPSLVETKSFNIGT--------FQARITAKSLGTTVMSPTIpl  1780
             A VS +TTTS++RH   H   +     +I T        F+   TA +L T V SPT PL
Sbjct  71    AMVSIDTTTSINRH--HHQCRLTPPPVSIDTPHRLSQTGFRVEFTADTLSTIVTSPTAPL  128

Query  1781  llhpppkrprnqsppLKPPYVINKSLKFIKTVLWLCKLRVSRRALFRSFDDCAGKRSVPP  1960
             LL PP KR R  SPP KPP +INK+LK+                L R           PP
Sbjct  129   LLDPPLKRQRYSSPPPKPPDIINKTLKYPDLFYGSASHEFLVEGLSR-----------PP  177

Query  1961  IPESHPADAPLFLDRPHALTAYTPAWMMRGFSPRYLIPP  2077
             IP S PAD P FL R H  T YT   MMRG + +YL+PP
Sbjct  178   IPLSCPADIPHFLGRSHTPTTYTLPLMMRGSTRKYLLPP  216


>contains similarity to an Arabidopsis thaliana hypothetical protein, which contains similarity to retrotransposon Athila (GB:AF076275) [Arabidopsis thaliana]
Sequence ID: AAD48941.1 Length: 205
>AT4g07460 [Arabidopsis thaliana]
Sequence ID: CAB81116.1 Length: 205
Range 1: 108 to 149

Score:74.7 bits(182), Expect:5e-14,
Method:Compositional matrix adjust.,
Identities:34/42(81%), Positives:35/42(83%), Gaps:0/42(0%)

Query  1406  WTLIAPISLDFPPVAELVYTPEIPCLMPQQYTQEIQEERCKA  1531
             WT  APISLDFP VAELVYTPEIPC +PQQY QEIQEE CK
Sbjct  108   WTQTAPISLDFPLVAELVYTPEIPCPLPQQYMQEIQEEHCKV  149


Range 2: 19 to 102

Score:73.2 bits(178), Expect:2e-13,
Method:Compositional matrix adjust.,
Identities:51/138(37%), Positives:59/138(42%), Gaps:54/138(39%)

Query  735   LFVEDVNYFGGRGYQNQRFKHQRGYKGVYGNVQPNYTqksqfqqpfqksssfsfTRNYDL  914
             L VEDVNYFGGR +              YGN
Sbjct  19    LLVEDVNYFGGRCF--------------YGN-----------------------------  35

Query  915   ASYQAPPPPEPRSKIESMPEQILEGQKKILERPIPRWTEYRLTPYWHVALMESEDEFLNV  1094
                   P    +S+ +         QK++LERPIPRW E  LT  WH A MES  EF+ V
Sbjct  36    -----GPTTTHKSQFQQ------SCQKRMLERPIPRWNESSLTTSWHAASMESGYEFVKV  84

Query  1095  EESDEFDVVEAVLTHTNP  1148
             EESD+FDV EAV T T P
Sbjct  85    EESDKFDVAEAVSTDTIP  102


>En/Spm-like transposon protein [Arabidopsis thaliana]
Sequence ID: AAD03363.1 Length: 454
Range 1: 120 to 188

Score:76.6 bits(187), Expect:2e-13,
Method:Compositional matrix adjust.,
Identities:37/69(54%), Positives:45/69(65%), Gaps:3/69(4%)

Query  4313  FGMHLEIMERSGV---KQRSYAASRNLKTKNRIGVERHHPWCRATPIPDGNSKMQLIEVL  4483
              G H++    +G    K  + +   N +TK  +GVERHHPWCRATPIPD NSK QLI+VL
Sbjct  120   LGEHVDSFGDNGALWSKTDNLSYRSNPETKMHVGVERHHPWCRATPIPDDNSKTQLIDVL  179

Query  4484  KISKFDPYF  4510
             KISKF P
Sbjct  180   KISKFGPKL  188


>unnamed protein product [Arabidopsis thaliana]
Sequence ID: CAA0394037.1 Length: 384
Range 1: 34 to 149

Score:74.7 bits(182), Expect:5e-13,
Method:Compositional matrix adjust.,
Identities:67/120(56%), Positives:79/120(65%), Gaps:4/120(3%)

Query  1980  AGWDSGMGGTLLFPAQSSKERKSARRETRSLHNHKTVFINFKLLLMTYgglrggdwflgr  1801
             AG D GMGG LL P QSSKERK ARRETR L N KT+   FK+LLM  GGL GGD +L R
Sbjct  34    AGQDKGMGGLLLLPRQSSKERKIARRETRGLLNCKTISGKFKVLLMISGGLGGGDEYLCR  93

Query  1800  lggg*rsrgivGDMTVVPRLLAVIRA*KVPILNDFVSTSDGCRDGCRSTLVVVLVDTRAT  1621
              GGG RS   VGD+TVVP  ++ + + + P L   +  SD    GCRST VVVL++T A
Sbjct  94    FGGGSRSICAVGDVTVVPG-VSPVNSTRKPAL---IRNSDSAFVGCRSTQVVVLINTIAA  149


>unnamed protein product [Arabidopsis thaliana]
Sequence ID: BAB02612.1 Length: 187
Range 1: 135 to 174

Score:67.0 bits(162), Expect:1e-11,
Method:Compositional matrix adjust.,
Identities:29/40(73%), Positives:33/40(82%), Gaps:0/40(0%)

Query  4385  KTKNRIGVERHHPWCRATPIPDGNSKMQLIEVLKISKFDP  4504
             +TK  +GVERHHPWCRATPIPD NSK Q I+VLKIS + P
Sbjct  135   QTKIPVGVERHHPWCRATPIPDDNSKTQFIDVLKISIYGP  174


>hypothetical protein AXX17_AT1G37440 [Arabidopsis thaliana]
Sequence ID: OAP14491.1 Length: 952
Range 1: 898 to 951

Score:71.2 bits(173), Expect:2e-11,
Method:Compositional matrix adjust.,
Identities:34/54(63%), Positives:40/54(74%), Gaps:0/54(0%)

Query  1931  DCAGKRSVPPIPESHPADAPLFLDRPHALTAYTPAWMMRGFSPRYLIPPSEPHD  2092
             D   KRS+PPI ESHPAD P F  R HALTA+TP WMMR  S ++L+PPS+P D
Sbjct  898   DLNRKRSIPPILESHPADLPHFNGRSHALTAHTPPWMMRYSSKKHLLPPSDPPD  951


>F10A2.1 gene product [Arabidopsis thaliana]
Sequence ID: AAD29783.1 Length: 331
Range 1: 165 to 286

Score:67.8 bits(164), Expect:5e-11,
Method:Compositional matrix adjust.,
Identities:64/155(41%), Positives:75/155(48%), Gaps:33/155(21%)

Query  1586  ELNYRHAKGRRVVARVSTNTTTSVDRHPSRHPSLVETKSFNIGTFQARITAKSLGTTVMS  1765
             E ++R     R    VS  T T V+RHP R    + TK+    T Q +I A   G  V +
Sbjct  165   EASFRFDDENRSAELVSIETNTCVNRHPKRAEPELSTKA----TLQVKIIAAIRGPVVTT  220

Query  1766  PTIplllhpppkrprnqsppLKPPYVINKSLKFIKTVLWLCKLRVSRRALFRSFDDCAGK  1945
                PL LHPPPKR R  SPP KP  + NK                              K
Sbjct  221   TNAPLPLHPPPKRLRYSSPPPKPLDIFNKC-----------------------------K  251

Query  1946  RSVPPIPESHPADAPLFLDRPHALTAYTPAWMMRG  2050
             R +PPIPES+PAD P  LD+ HALTAYTPAW MRG
Sbjct  252   RCIPPIPESYPADVPHLLDQSHALTAYTPAWTMRG  286


>putative TNP2-like transposon protein [Arabidopsis thaliana]
Sequence ID: AAD20646.1 Length: 1040
Range 1: 2 to 48

Score:65.9 bits(159), Expect:9e-10,
Method:Compositional matrix adjust.,
Identities:30/47(64%), Positives:38/47(80%), Gaps:0/47(0%)

Query  2102  DFYNHVVQKVVLNTVEKNLSSSMLVYKLLAHEVYLGREEHQQGGTQV  1962
             ++++ V QKV +N  EK+LSSSML YK L HEVYL REEHQ+GGTQ+
Sbjct  2     EYWDQVDQKVAVNITEKDLSSSMLAYKQLTHEVYLEREEHQKGGTQL  48


>unnamed protein product [Arabidopsis thaliana]
Sequence ID: VYS61969.1 Length: 333
Range 1: 1 to 110

Score:64.3 bits(155), Expect:9e-10,
Method:Compositional matrix adjust.,
Identities:62/114(54%), Positives:74/114(64%), Gaps:4/114(3%)

Query  1962  MGGTLLFPAQSSKERKSARRETRSLHNHKTVFINFKLLLMTYgglrggdwflgrlggg*r  1783
             MGG LL P QSSKERK ARRETR L   KT+   FK+LLM  GGL GGD +L R GGG R
Sbjct  1     MGGLLLLPRQSSKERKIARRETRGLLYCKTISGKFKVLLMISGGLGGGDEYLCRFGGGSR  60

Query  1782  srgivGDMTVVPRLLAVIRA*KVPILNDFVSTSDGCRDGCRSTLVVVLVDTRAT  1621
             S   VGD+TVVP  ++ + + + P L   +  SD    GCRST VVVL++T A
Sbjct  61    SICAVGDVTVVPG-VSPVNSTRKPAL---IRNSDSAFVGCRSTQVVVLINTIAA  110


>putative transposon protein [Arabidopsis thaliana]
Sequence ID: AAD27900.1 Length: 221
>putative transposon protein [Arabidopsis thaliana]
Sequence ID: CAB81148.1 Length: 221
Range 1: 182 to 221

Score:59.7 bits(143), Expect:9e-09,
Method:Compositional matrix adjust.,
Identities:29/40(73%), Positives:34/40(85%), Gaps:0/40(0%)

Query  4839  MGVVTIKMKNPTDNRAREQPTKSSKSLMRNPNPNKKATQT  4720
             + + +IKMKNP DN A  +PTKSSKSLMRNPNPNKKA+QT
Sbjct  182   LALRSIKMKNPKDNLAMGKPTKSSKSLMRNPNPNKKASQT  221


Query #202: XLOC_020814 Query ID: lcl|Query_30693 Length: 875

Sequences producing significant alignments:
                                                                  Max    Total Query   E   Per.
Description                                                       Score  Score cover Value Ident  Accession
unnamed protein product [Arabidopsis thaliana]                    89.7   142   22%   3e-22 85.42  VYS52357.1
unnamed protein product [Arabidopsis thaliana]                    70.1   70.1  41%   7e-15 44.17  CAA0361231.1
hypothetical protein AT2G14265 [Arabidopsis thaliana]             62.8   173   34%   8e-12 54.79  NP_001325174.1
hypothetical protein AT2G13845 [Arabidopsis thaliana]             63.5   63.5  23%   9e-12 54.41  NP_001324079.1
unnamed protein product [Arabidopsis thaliana]                    59.7   59.7  20%   3e-11 60.00  VYS52323.1
unnamed protein product [Arabidopsis thaliana]                    57.8   57.8  20%   1e-10 58.33  CAA0360812.1
transmembrane protein [Arabidopsis thaliana]                      50.8   81.6  16%   3e-09 76.67  NP_001326023.1
unnamed protein product [Arabidopsis thaliana]                    50.4   50.4  10%   7e-08 76.67  CAA0384287.1


Alignments:
>unnamed protein product [Arabidopsis thaliana]
Sequence ID: VYS52357.1 Length: 87
Range 1: 1 to 48

Score:89.7 bits(221), Expect:3e-22,
Method:Compositional matrix adjust.,
Identities:41/48(85%), Positives:42/48(87%), Gaps:0/48(0%)

Query  334  MCQTRFIYVASRFRLIEFPVIMDMRPLPSKPYGHGTLVSLQNTVNRIS  477
            MCQTRF YVASRF LI+FPVIMDMRPLPSKPYGHGTLVSLQNT    S
Sbjct  1    MCQTRFTYVASRFPLIKFPVIMDMRPLPSKPYGHGTLVSLQNTFEETS  48


Range 2: 7 to 87

Score:52.4 bits(124), Expect:2e-08,
Method:Compositional matrix adjust.,
Identities:34/81(42%), Positives:44/81(54%), Gaps:14/81(17%)

Query  330  SHVSNSFHLCSIEIPVNR---ISCYYGH-----------ETSTFKTLWTWNFGFFTKHS*  467
            ++V++ F L    + ++     S  YGH           ETST K LWTWNFGFFTK
Sbjct  7    TYVASRFPLIKFPVIMDMRPLPSKPYGHGTLVSLQNTFEETSTSKILWTWNFGFFTKPRL  66

Query  468  SNLLLLLTCFEETSTSKTLWT  530
                +++  F ETSTSKTLWT
Sbjct  67   IESSVIMDLFVETSTSKTLWT  87


>unnamed protein product [Arabidopsis thaliana]
Sequence ID: CAA0361231.1 Length: 79
Range 1: 6 to 79

Score:70.1 bits(170), Expect:7e-15,
Method:Compositional matrix adjust.,
Identities:53/120(44%), Positives:56/120(46%), Gaps:46/120(38%)

Query  352  IYVASRFRLIEFPVIMDMRPLPSKPYGHGTLVSLQNTVNRISCYY*HVLKRPPLPRPYGH  531
            +Y   R R + F     +RPLPSKPYGHGTLVSLQNT             RPPLPR YGH
Sbjct  6    LYTNWRIRGLGF---FKIRPLPSKPYGHGTLVSLQNT-------------RPPLPRSYGH  49

Query  532  EILVYLQTTVNRISCYYGHGI*NEYESVKSFLNGYI*MDSLLWTPPLRPYGHKTFTKFLR  711
              LV LQ  V R                             L  P  RPYGHKTFTKFLR
Sbjct  50   GTLVSLQNHVCRD----------------------------LHLP--RPYGHKTFTKFLR  79


>hypothetical protein AT2G14265 [Arabidopsis thaliana]
Sequence ID: NP_001325174.1 Length: 111
>hypothetical protein AT2G14265 [Arabidopsis thaliana]
Sequence ID: ANM63061.1 Length: 111
Range 1: 36 to 66

Score:43.9 bits(102), Expect:3e-12,
Method:Compositional matrix adjust.,
Identities:23/31(74%), Positives:23/31(74%), Gaps:0/31(0%)

Query  318  IFALSHVSNSFHLCSIEIPVNRISCYYGHET  410
            I ALSHVSNSFHLCSIEIP    S  YGH T
Sbjct  36   ISALSHVSNSFHLCSIEIPRPLPSKPYGHGT  66


Range 2: 1 to 18

Score:38.9 bits(89), Expect:3e-12,
Method:Compositional matrix adjust.,
Identities:17/18(94%), Positives:17/18(94%), Gaps:0/18(0%)

Query  161  MAFSSLWTNWRIRGLGFF  214
            MAFSSL TNWRIRGLGFF
Sbjct  1    MAFSSLLTNWRIRGLGFF  18


Range 3: 95 to 107

Score:28.1 bits(61), Expect:3e-12,
Method:Compositional matrix adjust.,
Identities:10/13(77%), Positives:11/13(84%), Gaps:0/13(0%)

Query  500  RDLHFQDLMDMKF  538
            RDLH QDLMD+K
Sbjct  95   RDLHLQDLMDIKL  107


Range 4: 41 to 95

Score:62.8 bits(151), Expect:8e-12,
Method:Compositional matrix adjust.,
Identities:40/73(55%), Positives:43/73(58%), Gaps:20/73(27%)

Query  355  YVASRFRL--IEFPVIMDMRPLPSKPYGHGTLVSLQNTVNRISCYY*HVLKRPPLPRPYG  528
            +V++ F L  IE P     RPLPSKPYGHGTLVSLQNT             RPPLPR YG
Sbjct  41   HVSNSFHLCSIEIP-----RPLPSKPYGHGTLVSLQNT-------------RPPLPRSYG  82

Query  529  HEILVYLQTTVNR  567
            H  LV LQ  V R
Sbjct  83   HGTLVSLQKHVCR  95


>hypothetical protein AT2G13845, partial [Arabidopsis thaliana]
Sequence ID: NP_001324079.1 Length: 143
>hypothetical protein AT2G13845, partial [Arabidopsis thaliana]
Sequence ID: ANM61889.1 Length: 143
Range 1: 83 to 137

Score:63.5 bits(153), Expect:9e-12,
Method:Compositional matrix adjust.,
Identities:37/68(54%), Positives:42/68(61%), Gaps:13/68(19%)

Query  375  VNRISCYYGHETSTFKTLWTWNFGFFTKHS*SNLLLLLTCFEETSTSKTLWT*NFGLFTN  554
            ++R+    G ETSTFKTLWTWNFGFFT H             ETSTSK LWT NFG FTN
Sbjct  83   LHRLEQMGGSETSTFKTLWTWNFGFFTNH-------------ETSTSKALWTWNFGFFTN  129

Query  555  YG*SNLLL  578
            +   +L L
Sbjct  130  HVCKDLRL  137


>unnamed protein product [Arabidopsis thaliana]
Sequence ID: VYS52323.1 Length: 61
Range 1: 3 to 49

Score:59.7 bits(143), Expect:3e-11,
Method:Compositional matrix adjust.,
Identities:36/60(60%), Positives:38/60(63%), Gaps:13/60(21%)

Query  399  GHETSTFKTLWTWNFGFFTKHS*SNLLLLLTCFEETSTSKTLWT*NFGLFTNYG*SNLLL  578
            G ETSTFKTLWTWNFGFFT H             ETSTSK LWT NFG FTN+   +L L
Sbjct  3    GSETSTFKTLWTWNFGFFTNH-------------ETSTSKALWTWNFGFFTNHVCKDLRL  49


>unnamed protein product [Arabidopsis thaliana]
Sequence ID: CAA0360812.1 Length: 61
Range 1: 3 to 49

Score:57.8 bits(138), Expect:1e-10,
Method:Compositional matrix adjust.,
Identities:35/60(58%), Positives:37/60(61%), Gaps:13/60(21%)

Query  399  GHETSTFKTLWTWNFGFFTKHS*SNLLLLLTCFEETSTSKTLWT*NFGLFTNYG*SNLLL  578
            G ETSTFKTLWTWNFGFFT H             ETSTSK LWT NFG FT +   +L L
Sbjct  3    GSETSTFKTLWTWNFGFFTNH-------------ETSTSKALWTWNFGFFTKHVCKDLRL  49


>transmembrane protein [Arabidopsis thaliana]
Sequence ID: NP_001326023.1 Length: 74
>transmembrane protein [Arabidopsis thaliana]
Sequence ID: ANM63965.1 Length: 74
Range 1: 15 to 44

Score:50.8 bits(120), Expect:3e-09,
Method:Compositional matrix adjust.,
Identities:23/30(77%), Positives:25/30(83%), Gaps:0/30(0%)

Query  500  RDLHFQDLMDMKFWFIYKLRLIESPVIMDM  589
            RDLH QDL DMK W +YK RLI+SPVIMDM
Sbjct  15   RDLHLQDLTDMKLWSLYKPRLIKSPVIMDM  44


Range 2: 58 to 74

Score:30.8 bits(68), Expect:3e-09,
Method:Compositional matrix adjust.,
Identities:13/17(76%), Positives:15/17(88%), Gaps:0/17(0%)

Query  661  TPPLRPYGHKTFTKFLR  711
             PP RPYG++TFTKFLR
Sbjct  58   APPPRPYGYRTFTKFLR  74


>unnamed protein product [Arabidopsis thaliana]
Sequence ID: CAA0384287.1 Length: 74
>unnamed protein product [Arabidopsis thaliana]
Sequence ID: VYS59150.1 Length: 74
Range 1: 15 to 44

Score:50.4 bits(119), Expect:7e-08,
Method:Compositional matrix adjust.,
Identities:23/30(77%), Positives:25/30(83%), Gaps:0/30(0%)

Query  500  RDLHFQDLMDMKFWFIYKLRLIESPVIMDM  589
            RDLH QDL DMK W +YK RLI+SPVIMDM
Sbjct  15   RDLHLQDLTDMKLWSLYKPRLIKSPVIMDM  44


Query #203: XLOC_020845 Query ID: lcl|Query_30694 Length: 634

No significant similarity found.


Query #204: XLOC_020925G Query ID: lcl|Query_30695 Length: 508

No significant similarity found.


Query #205: XLOC_021080 Query ID: lcl|Query_30696 Length: 1859

No significant similarity found.


Query #206: XLOC_021365 Query ID: lcl|Query_30697 Length: 532

Sequences producing significant alignments:
                                                                  Max    Total Query   E   Per.
Description                                                       Score  Score cover Value Ident  Accession
kinase like protein [Arabidopsis thaliana]                        60.1   132   42%   7e-19 75.68  CAB10474.1
unnamed protein product [Arabidopsis thaliana]                    71.6   71.6  21%   8e-17 89.74  VYS62969.1
RPP4/RPP5-like protein [Arabidopsis thaliana]                     78.6   78.6  20%   1e-16 97.22  AXU93535.1
RPP4/RPP5-like protein [Arabidopsis thaliana]                     78.6   78.6  20%   1e-16 97.22  AXU93666.1
unnamed protein product [Arabidopsis thaliana]                    70.9   70.9  21%   2e-16 87.18  CAA0395531.1
hypothetical protein AT4G16915 [Arabidopsis thaliana]             75.5   75.5  23%   2e-16 85.37  NP_001154244.1
unnamed protein product [Arabidopsis thaliana]                    77.8   77.8  20%   2e-16 97.22  VYS62975.1
hypothetical protein AT4G16855 [Arabidopsis thaliana]             66.2   66.2  21%   1e-14 82.05  NP_001118996.1
hypothetical protein AXX17_AT4G20020 [Arabidopsis thaliana]       60.1   93.2  30%   4e-13 75.68  OAP01144.1
unnamed protein product [Arabidopsis thaliana]                    60.1   92.8  30%   5e-13 75.68  CAA0395546.1
Protein kinase superfamily protein [Arabidopsis thaliana]         60.1   92.8  30%   5e-13 75.68  NP_193430.3
unnamed protein product [Arabidopsis thaliana]                    60.1   92.8  30%   5e-13 75.68  VYS62979.1


Alignments:
>kinase like protein [Arabidopsis thaliana]
Sequence ID: CAB10474.1 Length: 803
>kinase like protein [Arabidopsis thaliana]
Sequence ID: CAB80968.1 Length: 803
Range 1: 628 to 664

Score:60.1 bits(144), Expect:7e-19,
Method:Compositional matrix adjust.,
Identities:28/37(76%), Positives:32/37(86%), Gaps:0/37(0%)

Query  471  RKELLNFLQETLSGPLPNHEVSSSYPTSMRKHVAAVP  361
            RK LL+FLQET+S P+PNHEVSS  PTSMRK VAA+P
Sbjct  628  RKALLDFLQETMSVPIPNHEVSSKAPTSMRKRVAALP  664


Range 2: 691 to 710

Score:39.7 bits(91), Expect:7e-19,
Method:Composition-based stats.,
Identities:17/21(81%), Positives:19/21(90%), Gaps:1/21(4%)

Query  247  LVCFRSLHRGPKMIDVWSDGD  185
            LVCFRSLH+GPK IDVWS G+
Sbjct  691  LVCFRSLHQGPK-IDVWSAGE  710


Range 3: 600 to 619

Score:32.7 bits(73), Expect:7e-19,
Method:Composition-based stats.,
Identities:15/20(75%), Positives:16/20(80%), Gaps:0/20(0%)

Query  524  GLTLAKDVTSIRNNPSAEGK  465
            GLT AKDVTS RNNPS E +
Sbjct  600  GLTSAKDVTSTRNNPSGEKR  619


>unnamed protein product [Arabidopsis thaliana]
Sequence ID: VYS62969.1 Length: 45
Range 1: 1 to 39

Score:71.6 bits(174), Expect:8e-17,
Method:Compositional matrix adjust.,
Identities:35/39(90%), Positives:37/39(94%), Gaps:0/39(0%)

Query  415  MVWKWTRQCLLQKV*KLFPSADGLFLMLVTSLAKVRPKT  531
            MVWKWTRQCLLQKV KLFPS +GLFLMLVTSLA+VRPKT
Sbjct  1    MVWKWTRQCLLQKVYKLFPSPEGLFLMLVTSLAEVRPKT  39


>RPP4/RPP5-like protein [Arabidopsis thaliana]
Sequence ID: AXU93535.1 Length: 1397
Range 1: 1362 to 1397

Score:78.6 bits(192), Expect:1e-16,
Method:Composition-based stats.,
Identities:35/36(97%), Positives:35/36(97%), Gaps:0/36(0%)

Query  450   LQETLSGPLPNHEVSSSYPTSMRKHVAAVPPSRCTL  343
             LQETLSGPLPNHEVSSSYPTSMRKHVAAVPP RCTL
Sbjct  1362  LQETLSGPLPNHEVSSSYPTSMRKHVAAVPPYRCTL  1397


>RPP4/RPP5-like protein [Arabidopsis thaliana]
Sequence ID: AXU93666.1 Length: 1259
Range 1: 1224 to 1259

Score:78.6 bits(192), Expect:1e-16,
Method:Composition-based stats.,
Identities:35/36(97%), Positives:35/36(97%), Gaps:0/36(0%)

Query  450   LQETLSGPLPNHEVSSSYPTSMRKHVAAVPPSRCTL  343
             LQETLSGPLPNHEVSSSYPTSMRKHVAAVPP RCTL
Sbjct  1224  LQETLSGPLPNHEVSSSYPTSMRKHVAAVPPYRCTL  1259


>unnamed protein product [Arabidopsis thaliana]
Sequence ID: CAA0395531.1 Length: 45
Range 1: 1 to 39

Score:70.9 bits(172), Expect:2e-16,
Method:Compositional matrix adjust.,
Identities:34/39(87%), Positives:36/39(92%), Gaps:0/39(0%)

Query  415  MVWKWTRQCLLQKV*KLFPSADGLFLMLVTSLAKVRPKT  531
            MVWKWTRQCLLQKV KLFPS +G FLMLVTSLA+VRPKT
Sbjct  1    MVWKWTRQCLLQKVYKLFPSPEGFFLMLVTSLAEVRPKT  39


>hypothetical protein AT4G16915 [Arabidopsis thaliana]
Sequence ID: NP_001154244.1 Length: 236
>hypothetical protein AT4G16915 [Arabidopsis thaliana]
Sequence ID: AEE83822.1 Length: 236
Range 1: 196 to 236

Score:75.5 bits(184), Expect:2e-16,
Method:Compositional matrix adjust.,
Identities:35/41(85%), Positives:38/41(92%), Gaps:0/41(0%)

Query  396  DSWMILHGLEVDQTMSPAESLKALSFS*WIVPYAGHIFGQS  518
            DSWMILHGLEVDQTMSP ESLKALSF+  IVPYAG+IFG+S
Sbjct  196  DSWMILHGLEVDQTMSPVESLKALSFTRRIVPYAGYIFGRS  236


>unnamed protein product [Arabidopsis thaliana]
Sequence ID: VYS62975.1 Length: 1534
Range 1: 1499 to 1534

Score:77.8 bits(190), Expect:2e-16,
Method:Composition-based stats.,
Identities:35/36(97%), Positives:35/36(97%), Gaps:0/36(0%)

Query  450   LQETLSGPLPNHEVSSSYPTSMRKHVAAVPPSRCTL  343
             LQETLSGPLPNHEVSSSYPTSMRKHVAAVPP RCTL
Sbjct  1499  LQETLSGPLPNHEVSSSYPTSMRKHVAAVPPCRCTL  1534


>hypothetical protein AT4G16855 [Arabidopsis thaliana]
Sequence ID: NP_001118996.1 Length: 45
>hypothetical protein AT4G16855 [Arabidopsis thaliana]
Sequence ID: NP_001319969.1 Length: 45 >hypothetical protein AT4G16855 [Arabidopsis thaliana]
Sequence ID: AEE83817.1 Length: 45 >hypothetical protein AT4G16855 [Arabidopsis thaliana]
Sequence ID: ANM67795.1 Length: 45
Range 1: 1 to 39

Score:66.2 bits(160), Expect:1e-14,
Method:Compositional matrix adjust.,
Identities:32/39(82%), Positives:34/39(87%), Gaps:0/39(0%)

Query  415  MVWKWTRQCLLQKV*KLFPSADGLFLMLVTSLAKVRPKT  531
            MVWKWTRQCLLQKV KL PS +G FLMLV SLA+VRPKT
Sbjct  1    MVWKWTRQCLLQKVYKLVPSPEGFFLMLVISLAEVRPKT  39


>hypothetical protein AXX17_AT4G20020 [Arabidopsis thaliana]
Sequence ID: OAP01144.1 Length: 435
Range 1: 199 to 235

Score:60.1 bits(144), Expect:4e-13,
Method:Compositional matrix adjust.,
Identities:28/37(76%), Positives:32/37(86%), Gaps:0/37(0%)

Query  471  RKELLNFLQETLSGPLPNHEVSSSYPTSMRKHVAAVP  361
            RK LL+FLQET+S P+PNHEVSS  PTSMRK VAA+P
Sbjct  199  RKALLDFLQETMSVPIPNHEVSSKAPTSMRKRVAALP  235


Range 2: 171 to 190

Score:33.1 bits(74), Expect:4e-13,
Method:Compositional matrix adjust.,
Identities:15/20(75%), Positives:16/20(80%), Gaps:0/20(0%)

Query  524  GLTLAKDVTSIRNNPSAEGK  465
            GLT AKDVTS RNNPS E +
Sbjct  171  GLTSAKDVTSTRNNPSGEKR  190


>unnamed protein product [Arabidopsis thaliana]
Sequence ID: CAA0395546.1 Length: 883
Range 1: 647 to 683

Score:60.1 bits(144), Expect:5e-13,
Method:Compositional matrix adjust.,
Identities:28/37(76%), Positives:32/37(86%), Gaps:0/37(0%)

Query  471  RKELLNFLQETLSGPLPNHEVSSSYPTSMRKHVAAVP  361
            RK LL+FLQET+S P+PNHEVSS  PTSMRK VAA+P
Sbjct  647  RKALLDFLQETMSVPIPNHEVSSKAPTSMRKRVAALP  683


Range 2: 619 to 638

Score:32.7 bits(73), Expect:5e-13,
Method:Composition-based stats.,
Identities:15/20(75%), Positives:16/20(80%), Gaps:0/20(0%)

Query  524  GLTLAKDVTSIRNNPSAEGK  465
            GLT AKDVTS RNNPS E +
Sbjct  619  GLTSAKDVTSTRNNPSGEKR  638


>Protein kinase superfamily protein [Arabidopsis thaliana]
Sequence ID: NP_193430.3 Length: 889
>Protein kinase superfamily protein [Arabidopsis thaliana]
Sequence ID: AEE83829.1 Length: 889 >kinase like protein [Arabidopsis thaliana]
Sequence ID: BAF00949.1 Length: 889
Range 1: 649 to 685

Score:60.1 bits(144), Expect:5e-13,
Method:Compositional matrix adjust.,
Identities:28/37(76%), Positives:32/37(86%), Gaps:0/37(0%)

Query  471  RKELLNFLQETLSGPLPNHEVSSSYPTSMRKHVAAVP  361
            RK LL+FLQET+S P+PNHEVSS  PTSMRK VAA+P
Sbjct  649  RKALLDFLQETMSVPIPNHEVSSKAPTSMRKRVAALP  685


Range 2: 621 to 640

Score:32.7 bits(73), Expect:5e-13,
Method:Composition-based stats.,
Identities:15/20(75%), Positives:16/20(80%), Gaps:0/20(0%)

Query  524  GLTLAKDVTSIRNNPSAEGK  465
            GLT AKDVTS RNNPS E +
Sbjct  621  GLTSAKDVTSTRNNPSGEKR  640


>unnamed protein product [Arabidopsis thaliana]
Sequence ID: VYS62979.1 Length: 889
Range 1: 649 to 685

Score:60.1 bits(144), Expect:5e-13,
Method:Compositional matrix adjust.,
Identities:28/37(76%), Positives:32/37(86%), Gaps:0/37(0%)

Query  471  RKELLNFLQETLSGPLPNHEVSSSYPTSMRKHVAAVP  361
            RK LL+FLQET+S P+PNHEVSS  PTSMRK VAA+P
Sbjct  649  RKALLDFLQETMSVPIPNHEVSSKAPTSMRKRVAALP  685


Range 2: 621 to 640

Score:32.7 bits(73), Expect:5e-13,
Method:Composition-based stats.,
Identities:15/20(75%), Positives:16/20(80%), Gaps:0/20(0%)

Query  524  GLTLAKDVTSIRNNPSAEGK  465
            GLT AKDVTS RNNPS E +
Sbjct  621  GLTSAKDVTSTRNNPSGEKR  640


Query #207: XLOC_021449 Query ID: lcl|Query_30698 Length: 537

Sequences producing significant alignments:
                                                                  Max    Total Query   E   Per.
Description                                                       Score  Score cover Value Ident  Accession
hypothetical protein AXX17_AT4G21650 [Arabidopsis thaliana]       57.0   57.0  26%   5e-11 63.46  OAO97871.1


Alignments:
>hypothetical protein AXX17_AT4G21650 [Arabidopsis thaliana]
Sequence ID: OAO97871.1 Length: 55
Range 1: 4 to 55

Score:57.0 bits(136), Expect:5e-11,
Method:Compositional matrix adjust.,
Identities:33/52(63%), Positives:35/52(67%), Gaps:5/52(9%)

Query  237  EDRTS-----EIFLQIELGT*LQIALGRLLHSTRGSRSAPRHPSGGKKHPRD  97
            EDRTS       +   +    LQIALGRLLHSTRGSRSAPRH  GGKKH RD
Sbjct  4    EDRTSTRIGPRRYFSRQNWELLQIALGRLLHSTRGSRSAPRHSYGGKKHSRD  55


Query #208: XLOC_022604 Query ID: lcl|Query_30699 Length: 655

Sequences producing significant alignments:
                                                                  Max    Total Query   E   Per.
Description                                                       Score  Score cover Value Ident  Accession
transmembrane protein [Arabidopsis thaliana]                      51.2   51.2  22%   2e-08 47.06  NP_001322992.1


Alignments:
>transmembrane protein [Arabidopsis thaliana]
Sequence ID: NP_001322992.1 Length: 68
>transmembrane protein [Arabidopsis thaliana]
Sequence ID: ANM60728.1 Length: 68 >unnamed protein product [Arabidopsis thaliana]
Sequence ID: CAA0295663.1 Length: 68 >unnamed protein product [Arabidopsis thaliana]
Sequence ID: VYS49138.1 Length: 68
Range 1: 1 to 51

Score:51.2 bits(121), Expect:2e-08,
Method:Compositional matrix adjust.,
Identities:24/51(47%), Positives:34/51(66%), Gaps:2/51(3%)

Query  84   MYKAVFWLITLIFLLFSGSSNTALARMAYETPNT--KEGQWEEKVIRGSKI  230
            M KA+ WLITL+FL+FS + N  LA   Y +P T  +EG W+ K+++  KI
Sbjct  1    MNKALVWLITLLFLIFSATPNRVLAHPPYISPKTRAREGVWDHKIMKVRKI  51


Query #209: XLOC_022897 Query ID: lcl|Query_30700 Length: 641

No significant similarity found.


Query 210: XLOC_022998 Query ID: lcl|Query_30701 Length: 797

No significant similarity found.


Query #211: XLOC_023126 Query ID: lcl|Query_30702 Length: 638

Sequences producing significant alignments:
                                                                  Max    Total Query   E   Per.
Description                                                       Score  Score cover Value Ident  Accession
unnamed protein product [Arabidopsis thaliana]                    80.9   80.9  22%   7e-19 83.33  BAB01826.1


Alignments:
>unnamed protein product [Arabidopsis thaliana]
Sequence ID: BAB01826.1 Length: 143
Range 1: 1 to 48

Score:80.9 bits(198), Expect:7e-19,
Method:Compositional matrix adjust.,
Identities:40/48(83%), Positives:43/48(89%), Gaps:1/48(2%)

Query  56   MSLMNPSNVNLATYVAFNDLQLGTHAQHVIGHIVRM*EAH-EIRTAND  196
            MSL+N SN+NLATYVAFNDLQLGT+ QHVIGHIVRM E H EIRTAND
Sbjct  1    MSLINLSNLNLATYVAFNDLQLGTYVQHVIGHIVRMWEVHTEIRTAND  48


Query #212: XLOC_023260 Query ID: lcl|Query_30703 Length: 568

Sequences producing significant alignments:
                                                                  Max    Total Query   E   Per.
Description                                                       Score  Score cover Value Ident  Accession
hypothetical protein [Arabidopsis thaliana]                       64.3   64.3  23%   3e-13 75.00  AAD15327.1


Alignments:
>hypothetical protein [Arabidopsis thaliana]
Sequence ID: AAD15327.1 Length: 87
>hypothetical protein [Arabidopsis thaliana]
Sequence ID: CAB77856.1 Length: 87
Range 1: 11 to 54

Score:64.3 bits(155), Expect:3e-13,
Method:Compositional matrix adjust.,
Identities:33/44(75%), Positives:37/44(84%), Gaps:0/44(0%)

Query  195  RILEEIITQQAAIQEDNNDLQNSAIVIALNVSSLILPGGNLEDI  326
            RILEEIITQ+ A+QEDN+DLQ SA V  LN+SSLILP G LEDI
Sbjct  11   RILEEIITQRVAMQEDNDDLQYSATVNTLNLSSLILPRGQLEDI  54


Query #213: XLOC_023537 Query ID: lcl|Query_30704 Length: 572

No significant similarity found.


Query #214: XLOC_023590 Query ID: lcl|Query_30705 Length: 508

No significant similarity found.


Query #215: XLOC_024749 Query ID: lcl|Query_30706 Length: 1214

No significant similarity found.


Query #216: XLOC_025137 Query ID: lcl|Query_30707 Length: 881

Sequences producing significant alignments:
                                                                  Max    Total Query   E   Per.
Description                                                       Score  Score cover Value Ident  Accession
hypothetical protein AXX17_AT1G11410 [Arabidopsis thaliana]       76.3   123   25%   1e-21 77.08  OAP16805.1
hypothetical protein [Arabidopsis thaliana]                       50.8   50.8  10%   3e-08 77.42  BAF00163.1


Alignments:
>hypothetical protein AXX17_AT1G11410 [Arabidopsis thaliana]
Sequence ID: OAP16805.1 Length: 80
Range 1: 28 to 75

Score:76.3 bits(186), Expect:1e-21,
Method:Compositional matrix adjust.,
Identities:37/48(77%), Positives:38/48(79%), Gaps:0/48(0%)

Query  544  QHLDLSSMKNRCPYAFISIARNFPSHGRVIKHPMNTFPSKNLQELNIF  687
            +HLDLS MKNR PYA IS ARN PSHGR IKHPMN  PSK L ELNIF
Sbjct  28   KHLDLSPMKNRRPYALISKARNLPSHGRKIKHPMNKDPSKTLPELNIF  75


Range 2: 1 to 34

Score:47.0 bits(110), Expect:1e-21,
Method:Compositional matrix adjust.,
Identities:24/34(71%), Positives:25/34(73%), Gaps:0/34(0%)

Query  464  IRIRIFLGGPLKHDVNPNSSYSSPINHNTLIYHP  565
            +RI I LGGP KHDVNPNSSYSSPIN   L   P
Sbjct  1    MRIGIILGGPSKHDVNPNSSYSSPINRKHLDLSP  34


>hypothetical protein [Arabidopsis thaliana]
Sequence ID: BAF00163.1 Length: 36
Range 1: 6 to 36

Score:50.8 bits(120), Expect:3e-08,
Method:Compositional matrix adjust.,
Identities:24/31(77%), Positives:27/31(87%), Gaps:0/31(0%)

Query  552  KVLWLIGEEYDELGFTSCFNGPPRNILILIS  460
            + L LIGEEY+ELGFTSCF+GPPR I ILIS
Sbjct  6    RCLRLIGEEYEELGFTSCFDGPPRIIPILIS  36


Query #217: XLOC_025139 Query ID: lcl|Query_30708 Length: 457

No significant similarity found.


Query #218: XLOC_025140 Query ID: lcl|Query_30709 Length: 969

No significant similarity found.


Query #219: XLOC_025141 Query ID: lcl|Query_30710 Length: 868

No significant similarity found.


Query #220: XLOC_025142 Query ID: lcl|Query_30711 Length: 368

No significant similarity found.


Query #221: XLOC_025143 Query ID: lcl|Query_30712 Length: 1087

No significant similarity found.


Query #222: XLOC_025144 Query ID: lcl|Query_30713 Length: 870

Sequences producing significant alignments:
                                                                  Max    Total Query   E   Per.
Description                                                       Score  Score cover Value Ident  Accession
unnamed protein product [Arabidopsis thaliana]                    56.6   56.6  12%   5e-09 80.00  BAB01478.1


Alignments:
>unnamed protein product [Arabidopsis thaliana]
Sequence ID: BAB01478.1 Length: 193
Range 1: 1 to 35

Score:56.6 bits(135), Expect:5e-09,
Method:Compositional matrix adjust.,
Identities:28/35(80%), Positives:31/35(88%), Gaps:0/35(0%)

Query  734  MFLLDECNLVILFKGKSIFKNGQHKRVLLKKGSKK  838
            MFLLDE NLVI+FKG+ IFKN Q+KRVLL KGSKK
Sbjct  1    MFLLDEGNLVIMFKGRDIFKNEQYKRVLLHKGSKK  35


Query #223: XLOC_025145 Query ID: lcl|Query_30714 Length: 2126

Sequences producing significant alignments:
                                                                  Max    Total Query   E   Per.
Description                                                       Score  Score cover Value Ident  Accession
hypothetical protein AXX17_AT4G12220 [Arabidopsis thaliana]       152    152   10%   2e-43 90.48  OAO99210.1
putative kinesin light chain [Arabidopsis thaliana]               73.6   73.6  6%    8e-13 75.00  AAM63491.1
Tetratricopeptide repeat (TPR)-like superfamily protein...        73.6   73.6  6%    8e-13 75.00  NP_192822.1
unnamed protein product [Arabidopsis thaliana]                    68.9   68.9  12%   6e-12 43.88  CAA0401921.1
unnamed protein product [Arabidopsis thaliana]                    68.6   68.6  12%   1e-11 44.44  VYS58807.1
RNA-directed DNA polymerase (reverse transcriptase)-related...    65.1   65.1  12%   4e-11 42.42  NP_178356.1
hypothetical protein AXX17_AT3G27760 [Arabidopsis thaliana]       62.0   62.0  12%   1e-10 40.82  OAP04261.1
unnamed protein product [Arabidopsis thaliana]                    61.6   61.6  12%   1e-10 40.82  CAA0383678.1
hypothetical protein AXX17_AT4G05670 [Arabidopsis thaliana]       62.0   62.0  16%   1e-09 33.33  OAO98474.1
unnamed protein product [Arabidopsis thaliana]                    62.0   62.0  16%   1e-09 33.33  VYS61841.1
hypothetical protein AXX17_AT2G03940 [Arabidopsis thaliana]       63.5   63.5  10%   1e-09 45.78  OAP09951.1
hypothetical protein AXX17_AT4G01230 [Arabidopsis thaliana]       62.4   62.4  12%   2e-09 40.40  OAO99387.1
hypothetical protein At4g04650 [Arabidopsis thaliana]             59.3   59.3  16%   8e-09 30.89  ABE65512.1
unknown [Arabidopsis thaliana]                                    59.3   59.3  16%   8e-09 30.89  ABK28243.1
unnamed protein product [Arabidopsis thaliana]                    59.3   59.3  16%   8e-09 30.89  CAA0393718.1
unnamed protein product [Arabidopsis thaliana]                    54.3   54.3  9%    2e-08 42.86  VYS47991.1
unnamed protein product [Arabidopsis thaliana]                    56.6   56.6  12%   2e-08 35.79  CAA0270646.1
unnamed protein product [Arabidopsis thaliana]                    57.0   78.9  9%    4e-08 50.77  CAA0205272.1
non-LTR retrolelement reverse transcriptase [Arabidopsis...       54.7   54.7  12%   4e-08 38.78  NP_683592.1
unnamed protein product [Arabidopsis thaliana]                    56.2   56.2  13%   1e-07 37.37  CAA0272723.1


Alignments:
>hypothetical protein AXX17_AT4G12220 [Arabidopsis thaliana]
Sequence ID: OAO99210.1 Length: 84
Range 1: 1 to 84

Score:152 bits(384), Expect:2e-43,
Method:Compositional matrix adjust.,
Identities:76/84(90%), Positives:76/84(90%), Gaps:7/84(8%)

Query  999  MIVLDFCNEN-------CMQSAIPFLPKRINGGLERKFEDIPYIFLGARVLETEARPFLS  841
            MIVLDFCNE        CMQSAIPFLPKRINGGLERKFEDIPYIFLGARVLETEARPFLS
Sbjct  1    MIVLDFCNERAVRPIDFCMQSAIPFLPKRINGGLERKFEDIPYIFLGARVLETEARPFLS  60

Query  840  RGLFIGRSCVWINQILEQHDPIFL  769
            RGLFIGRSCVWINQILEQHDPIFL
Sbjct  61   RGLFIGRSCVWINQILEQHDPIFL  84


>putative kinesin light chain [Arabidopsis thaliana]
Sequence ID: AAM63491.1 Length: 606
Range 1: 554 to 605

Score:73.6 bits(179), Expect:8e-13,
Method:Compositional matrix adjust.,
Identities:39/52(75%), Positives:39/52(75%), Gaps:6/52(11%)

Query  733  FEDEKKRLVELLKEDG------VVLLQNLIDPNARPPNKESSAKKWSSLGFK  870
            FEDEKKRL ELLKE G         LQNLIDPNARPP KESSAKKW SLGFK
Sbjct  554  FEDEKKRLAELLKEAGRSRNYKAKSLQNLIDPNARPPKKESSAKKWPSLGFK  605


>Tetratricopeptide repeat (TPR)-like superfamily protein [Arabidopsis thaliana]
Sequence ID: NP_192822.1 Length: 609
>RecName: Full=Protein KINESIN LIGHT CHAIN-RELATED 1 [Arabidopsis thaliana]
Sequence ID: O81629.1 Length: 609 >contains similarity to TPR domains (Pfam: TPR.hmm: score: 11.15) and kinesin motor domains (Pfam: kinesin2.hmm, score: 17.49, 20.52 and 10.94) [Arabidopsis thaliana]
Sequence ID: AAC33943.1 Length: 609 >unknown protein [Arabidopsis thaliana]
Sequence ID: AAL59992.1 Length: 609 >AT4g10840/F25I24_50 [Arabidopsis thaliana]
Sequence ID: AAL91265.1 Length: 609 >unknown protein [Arabidopsis thaliana]
Sequence ID: AAN41387.1 Length: 609 >Tetratricopeptide repeat (TPR)-like superfamily protein [Arabidopsis thaliana]
Sequence ID: AEE82933.1 Length: 609
Range 1: 557 to 608

Score:73.6 bits(179), Expect:8e-13,
Method:Compositional matrix adjust.,
Identities:39/52(75%), Positives:39/52(75%), Gaps:6/52(11%)

Query  733  FEDEKKRLVELLKEDG------VVLLQNLIDPNARPPNKESSAKKWSSLGFK  870
            FEDEKKRL ELLKE G         LQNLIDPNARPP KESSAKKW SLGFK
Sbjct  557  FEDEKKRLAELLKEAGRSRNYKAKSLQNLIDPNARPPKKESSAKKWPSLGFK  608


>unnamed protein product [Arabidopsis thaliana]
Sequence ID: CAA0401921.1 Length: 293
Range 1: 202 to 292

Score:68.9 bits(167), Expect:6e-12,
Method:Compositional matrix adjust.,
Identities:43/98(44%), Positives:57/98(58%), Gaps:14/98(14%)

Query  820   PPNKESSAKKWSSLGFKDPSPKKNVRYILKLAFQASIYSLWKERNSRLHTV-------LI  978
             PP       +W     K+PS  KNV  IL++A QA++Y+LW+ERN+RLHT        LI
Sbjct  202   PPPLFEDGVRW----LKNPSRDKNVTTILRIAHQATVYTLWRERNTRLHTSSSRPAAGLI  257

Query  979   TKIKNNH*VQT*SSLALSRVQENSPNVISFLVTWFELF  1092
              +IKN   ++T     LSR Q   PN +S+L TWF LF
Sbjct  258   VEIKNL--IRT-HLDPLSRAQRIGPNGLSYLATWFGLF  292


>unnamed protein product [Arabidopsis thaliana]
Sequence ID: VYS58807.1 Length: 327
Range 1: 236 to 327

Score:68.6 bits(166), Expect:1e-11,
Method:Compositional matrix adjust.,
Identities:44/99(44%), Positives:53/99(53%), Gaps:14/99(14%)

Query  820   PPNKESSAKKWSSLGFKDPSPKKNVRYILKLAFQASIYSLWKERNSRLH-------TVLI  978
             PP       +W     K+P   KNV  IL+LA  AS+Y++WKERN+RLH       T LI
Sbjct  236   PPLLFEDGIRW----LKNPCQDKNVTTILRLAHHASVYTIWKERNARLHASASRPATALI  291

Query  979   TKIKNNH*VQT*SSLALSRVQENSPNVISFLVTWFELFQ  1095
              +IK+   V       LSR Q   PN ISFL TWF LF
Sbjct  292   LEIKS---VIRCHLDPLSRAQRLGPNGISFLATWFGLFD  327


>RNA-directed DNA polymerase (reverse transcriptase)-related family protein [Arabidopsis thaliana]
Sequence ID: NP_178356.1 Length: 211
>putative non-LTR retroelement reverse transcriptase [Arabidopsis thaliana]
Sequence ID: AAC18934.1 Length: 211 >RNA-directed DNA polymerase (reverse transcriptase)-related family protein [Arabidopsis thaliana]
Sequence ID: AEC05591.1 Length: 211
Range 1: 120 to 211

Score:65.1 bits(157), Expect:4e-11,
Method:Compositional matrix adjust.,
Identities:42/99(42%), Positives:52/99(52%), Gaps:14/99(14%)

Query  820   PPNKESSAKKWSSLGFKDPSPKKNVRYILKLAFQASIYSLWKERNSRLH-------TVLI  978
             PP       +W     K+P   KNV  IL+L+  AS+Y++WKERN+RLH         LI
Sbjct  120   PPLLFEDGIRW----LKNPCQDKNVTTILRLSHHASVYTIWKERNARLHDSASRPAAALI  175

Query  979   TKIKNNH*VQT*SSLALSRVQENSPNVISFLVTWFELFQ  1095
              +IK+   V       LSR Q   PN ISFL TWF LF
Sbjct  176   LEIKS---VIRCHLDPLSRAQRLGPNRISFLATWFGLFD  211


>hypothetical protein AXX17_AT3G27760 [Arabidopsis thaliana]
Sequence ID: OAP04261.1 Length: 142
Range 1: 50 to 140

Score:62.0 bits(149), Expect:1e-10,
Method:Compositional matrix adjust.,
Identities:40/98(41%), Positives:50/98(51%), Gaps:14/98(14%)

Query  820   PPNKESSAKKWSSLGFKDPSPKKNVRYILKLAFQASIYSLWKERNSRLHTV-------LI  978
             PP+      +W     K+P   KNV  IL++A QAS+Y+LWKERNSRLH         +I
Sbjct  50    PPHLFGDGVRW----LKNPCHDKNVTMILRIAHQASVYTLWKERNSRLHNTTSRPAAGII  105

Query  979   TKIKNNH*VQT*SSLALSRVQENSPNVISFLVTWFELF  1092
              +IK    +       LSR Q       SFL TWF LF
Sbjct  106   LEIKA---LIRCHLDPLSRAQRLGHQGTSFLATWFGLF  140


>unnamed protein product [Arabidopsis thaliana]
Sequence ID: CAA0383678.1 Length: 135
>unnamed protein product [Arabidopsis thaliana]
Sequence ID: VYS58597.1 Length: 135
Range 1: 43 to 133

Score:61.6 bits(148), Expect:1e-10,
Method:Compositional matrix adjust.,
Identities:40/98(41%), Positives:50/98(51%), Gaps:14/98(14%)

Query  820   PPNKESSAKKWSSLGFKDPSPKKNVRYILKLAFQASIYSLWKERNSRLHTV-------LI  978
             PP+      +W     K+P   KNV  IL++A QAS+Y+LWKERNSRLH         +I
Sbjct  43    PPHLFGDGVRW----LKNPCHDKNVTMILRIAHQASVYTLWKERNSRLHNTTSRPAAGII  98

Query  979   TKIKNNH*VQT*SSLALSRVQENSPNVISFLVTWFELF  1092
              +IK    +       LSR Q       SFL TWF LF
Sbjct  99    LEIKA---LIRCHLDPLSRAQRLGHQGTSFLATWFGLF  133


>hypothetical protein AXX17_AT4G05670 [Arabidopsis thaliana]
Sequence ID: OAO98474.1 Length: 296
Range 1: 178 to 296

Score:62.0 bits(149), Expect:1e-09,
Method:Compositional matrix adjust.,
Identities:41/123(33%), Positives:60/123(48%), Gaps:8/123(6%)

Query  736   EDEKKRLVELLKEDGVVLLQNLIDPNARPPNKESSAKKWSSLGFKDPSPKKNVRYILKLA  915
             +D +  L    +  GVV        N  PP +      W       PS +KN+  I++LA
Sbjct  178   DDSRAHLFFECQFSGVVWRFFTASTNLNPPAQLMDCLNW----LLSPSREKNICLIIRLA  233

Query  916   FQASIYSLWKERNSRLHTVL---ITKIKNNH*VQT*SSL-ALSRVQENSPNVISFLVTWF  1083
             FQA +Y++W+ERN RLH+ +      I  +  +   + L  LSR   + PN +S L TWF
Sbjct  234   FQACVYAIWRERNQRLHSGVSRSTESILKDIQLTIRARLDPLSRSTAHQPNALSLLGTWF  293

Query  1084  ELF  1092
              LF
Sbjct  294   SLF  296


>unnamed protein product [Arabidopsis thaliana]
Sequence ID: VYS61841.1 Length: 296
Range 1: 178 to 296

Score:62.0 bits(149), Expect:1e-09,
Method:Compositional matrix adjust.,
Identities:41/123(33%), Positives:60/123(48%), Gaps:8/123(6%)

Query  736   EDEKKRLVELLKEDGVVLLQNLIDPNARPPNKESSAKKWSSLGFKDPSPKKNVRYILKLA  915
             +D +  L    +  GVV        N  PP +      W       PS +KN+  I++LA
Sbjct  178   DDSRAHLFFECQFSGVVWRFFTASTNLNPPAQLMDCLNW----LLSPSREKNICLIIRLA  233

Query  916   FQASIYSLWKERNSRLHTVL---ITKIKNNH*VQT*SSL-ALSRVQENSPNVISFLVTWF  1083
             FQA +Y++W+ERN RLH+ +      I  +  +   + L  LSR   + PN +S L TWF
Sbjct  234   FQACVYAIWRERNQRLHSGVSRSTESILKDIQLTIRARLDPLSRSTAHQPNALSLLGTWF  293

Query  1084  ELF  1092
              LF
Sbjct  294   SLF  296


>hypothetical protein AXX17_AT2G03940 [Arabidopsis thaliana]
Sequence ID: OAP09951.1 Length: 1646
Range 1: 1566 to 1645

Score:63.5 bits(153), Expect:1e-09,
Method:Compositional matrix adjust.,
Identities:38/83(46%), Positives:47/83(56%), Gaps:10/83(12%)

Query  865   FKDPSPKKNVRYILKLAFQASIYSLWKERNSRLH-------TVLITKIKNNH*VQT*SSL  1023
              K+P   KNV  IL+LA  AS+Y++WKERN+RLH         LI +IK+   V
Sbjct  1566  LKNPCQDKNVTTILRLAHHASVYTIWKERNARLHASASRPAAALILEIKS---VIRCHLD  1622

Query  1024  ALSRVQENSPNVISFLVTWFELF  1092
              LSR Q   PN + FL TWF LF
Sbjct  1623  PLSRAQRLGPNGVFFLATWFGLF  1645


>hypothetical protein AXX17_AT4G01230 [Arabidopsis thaliana]
Sequence ID: OAO99387.1 Length: 450
Range 1: 359 to 450

Score:62.4 bits(150), Expect:2e-09,
Method:Compositional matrix adjust.,
Identities:40/99(40%), Positives:52/99(52%), Gaps:14/99(14%)

Query  820   PPNKESSAKKWSSLGFKDPSPKKNVRYILKLAFQASIYSLWKERNSRLH-------TVLI  978
             PP       +W     K+P   KN+  IL+LA+ AS+Y +WKERNSRLH       + +I
Sbjct  359   PPVMFLDGVRW----LKNPCRDKNIALILRLAYHASVYYIWKERNSRLHNSSSKPASAII  414

Query  979   TKIKNNH*VQT*SSLALSRVQENSPNVISFLVTWFELFQ  1095
              +IKN   +       LSR Q  +P   S LVTWF  FQ
Sbjct  415   LEIKN---IIRCHLDPLSRAQMITPPASSLLVTWFGYFQ  450


>hypothetical protein At4g04650 [Arabidopsis thaliana]
Sequence ID: ABE65512.1 Length: 296
Range 1: 178 to 296

Score:59.3 bits(142), Expect:8e-09,
Method:Compositional matrix adjust.,
Identities:38/123(31%), Positives:59/123(47%), Gaps:8/123(6%)

Query  736   EDEKKRLVELLKEDGVVLLQNLIDPNARPPNKESSAKKWSSLGFKDPSPKKNVRYILKLA  915
             +D +  L    +  GVV        N  PP +      W       PS +KN+  I++LA
Sbjct  178   DDSRAHLFFECQFSGVVWRFFTASTNLNPPAQLMDCLNW----LLSPSREKNICLIIRLA  233

Query  916   FQASIYSLWKERNSRLHTVLI----TKIKNNH*VQT*SSLALSRVQENSPNVISFLVTWF  1083
             F + +Y++W+ERN RLH+ +     + +K+   +       LSR   + PN +S L TWF
Sbjct  234   FHSCVYAIWRERNQRLHSGVSRSTESILKDIQLIIRARLDPLSRSTAHQPNALSLLGTWF  293

Query  1084  ELF  1092
              LF
Sbjct  294   SLF  296


>unknown, partial [Arabidopsis thaliana]
Sequence ID: ABK28243.1 Length: 297
Range 1: 178 to 296

Score:59.3 bits(142), Expect:8e-09,
Method:Compositional matrix adjust.,
Identities:38/123(31%), Positives:59/123(47%), Gaps:8/123(6%)

Query  736   EDEKKRLVELLKEDGVVLLQNLIDPNARPPNKESSAKKWSSLGFKDPSPKKNVRYILKLA  915
             +D +  L    +  GVV        N  PP +      W       PS +KN+  I++LA
Sbjct  178   DDSRAHLFFECQFSGVVWRFFTASTNLNPPAQLMDCLNW----LLSPSREKNICLIIRLA  233

Query  916   FQASIYSLWKERNSRLHTVLI----TKIKNNH*VQT*SSLALSRVQENSPNVISFLVTWF  1083
             F + +Y++W+ERN RLH+ +     + +K+   +       LSR   + PN +S L TWF
Sbjct  234   FHSCVYAIWRERNQRLHSGVSRSTESILKDIQLIIRARLDPLSRSTAHQPNALSLLGTWF  293

Query  1084  ELF  1092
              LF
Sbjct  294   SLF  296


>unnamed protein product [Arabidopsis thaliana]
Sequence ID: CAA0393718.1 Length: 296
Range 1: 178 to 296

Score:59.3 bits(142), Expect:8e-09,
Method:Compositional matrix adjust.,
Identities:38/123(31%), Positives:59/123(47%), Gaps:8/123(6%)

Query  736   EDEKKRLVELLKEDGVVLLQNLIDPNARPPNKESSAKKWSSLGFKDPSPKKNVRYILKLA  915
             +D +  L    +  GVV        N  PP +      W       PS +KN+  I++LA
Sbjct  178   DDSRAHLFFECQFSGVVWRFFTASTNLNPPAQLMDCLNW----LLSPSREKNICLIIRLA  233

Query  916   FQASIYSLWKERNSRLHTVLI----TKIKNNH*VQT*SSLALSRVQENSPNVISFLVTWF  1083
             F + +Y++W+ERN RLH+ +     + +K+   +       LSR   + PN +S L TWF
Sbjct  234   FHSCVYAIWRERNQRLHSGVSRSTESILKDIQLIIRARLDPLSRSTAHQPNALSLLGTWF  293

Query  1084  ELF  1092
              LF
Sbjct  294   SLF  296


>unnamed protein product [Arabidopsis thaliana]
Sequence ID: VYS47991.1 Length: 89
Range 1: 16 to 89

Score:54.3 bits(129), Expect:2e-08,
Method:Compositional matrix adjust.,
Identities:33/77(43%), Positives:46/77(59%), Gaps:10/77(12%)

Query  886   KNVRYILKLAFQASIYSLWKERNSRLH-------TVLITKIKNNH*VQT*SSLALSRVQE  1044
             +N+  ILKL F AS+Y +W+ERN RLH       T +I +IK    V+T   + LSR Q
Sbjct  16    RNITLILKLIFPASLYFIWRERNQRLHSATSRPATSIIKEIKMI--VRT-RLVPLSRAQR  72

Query  1045  NSPNVISFLVTWFELFQ  1095
             ++P  ++ L TWF L Q
Sbjct  73    STPRALTLLATWFSLSQ  89


>unnamed protein product [Arabidopsis thaliana]
Sequence ID: CAA0270646.1 Length: 181
Range 1: 89 to 179

Score:56.6 bits(135), Expect:2e-08,
Method:Compositional matrix adjust.,
Identities:34/95(36%), Positives:50/95(52%), Gaps:8/95(8%)

Query  820   PPNKESSAKKWSSLGFKDPSPKKNVRYILKLAFQASIYSLWKERNSRLHTVLITK----I  987
             PPN   +   W     K  S  KN+  ++KL FQAS+Y +W+ERN R+H+ +       I
Sbjct  89    PPNNFGAILTW----LKTASTSKNLSLLIKLIFQASLYLIWRERNCRIHSNIFRNPPQII  144

Query  988   KNNH*VQT*SSLALSRVQENSPNVISFLVTWFELF  1092
             K    +       LSRV  ++ +  S ++TWFELF
Sbjct  145   KEIQVLLRAKLDPLSRVSHSNASRASIMITWFELF  179


>unnamed protein product [Arabidopsis thaliana]
Sequence ID: CAA0205272.1 Length: 1256
Range 1: 1131 to 1191

Score:57.0 bits(136), Expect:4e-08,
Method:Compositional matrix adjust.,
Identities:33/65(51%), Positives:41/65(63%), Gaps:11/65(16%)

Query  820   PPNKESSAKKWSSLGFKDPSPKKNVRYILKLAFQASIYSLWKERNSRLHT-------VLI  978
             PP    +  +W     K PS  KNV  ILKLAFQASIY++WKERN+RLHT        +I
Sbjct  1131  PPFLFENVVRW----LKVPSRNKNVSLILKLAFQASIYAIWKERNARLHTSSSRPPASII  1186

Query  979   TKIKN  993
             ++IKN
Sbjct  1187  SEIKN  1191


Range 2: 1190 to 1200

Score:21.9 bits(45), Expect:4e-08,
Method:Compositional matrix adjust.,
Identities:7/11(64%), Positives:10/11(90%), Gaps:0/11(0%)

Query  989   RTIIRCKLDPL  1021
             + II+C+LDPL
Sbjct  1190  KNIIKCRLDPL  1200


>non-LTR retrolelement reverse transcriptase [Arabidopsis thaliana]
Sequence ID: NP_683592.1 Length: 153
>non-LTR retrolelement reverse transcriptase [Arabidopsis thaliana]
Sequence ID: AEE77061.1 Length: 153
Range 1: 61 to 151

Score:54.7 bits(130), Expect:4e-08,
Method:Composition-based stats.,
Identities:38/98(39%), Positives:50/98(51%), Gaps:14/98(14%)

Query  820   PPNKESSAKKWSSLGFKDPSPKKNVRYILKLAFQASIYSLWKERNSRLHTV-------LI  978
             PP+      +W     K+P   KNV  IL++A QAS+Y+LWKERNSRL+         +I
Sbjct  61    PPHLFGDGVRW----LKNPCHDKNVTMILRIAHQASVYTLWKERNSRLNNATSRPAAGII  116

Query  979   TKIKNNH*VQT*SSLALSRVQENSPNVISFLVTWFELF  1092
              +IK    +       LSR Q       SFL TWF +F
Sbjct  117   LEIKA---LIRCHLDPLSRAQRLGHQGTSFLATWFGMF  151


>unnamed protein product [Arabidopsis thaliana]
Sequence ID: CAA0272723.1 Length: 322
Range 1: 227 to 321

Score:56.2 bits(134), Expect:1e-07,
Method:Compositional matrix adjust.,
Identities:37/99(37%), Positives:51/99(51%), Gaps:8/99(8%)

Query  811   NARPPNKESSAKKWSSLGFKDPSPKKNVRYILKLAFQASIYSLWKERNSRLHT----VLI  978
             N  PP +      W      +PS  KN   I++LAFQAS+Y++W+ERN  LHT    V
Sbjct  227   NLFPPAQLMYCLIW----LLNPSRDKNTTLIIRLAFQASVYAIWRERNLCLHTGVARVFD  282

Query  979   TKIKNNH*VQT*SSLALSRVQENSPNVISFLVTWFELFQ  1095
             + +K+           +SR+ E  P+  S L TWF LFQ
Sbjct  283   SVLKDIQLTIRARLDPISRITELHPSAPSVLRTWFSLFQ  321


Query #224: XLOC_025146 Query ID: lcl|Query_30715 Length: 537

Sequences producing significant alignments:
                                                                  Max    Total Query   E   Per.
Description                                                       Score  Score cover Value Ident  Accession
putative disease resistance response protein [Arabidopsis...      117    117   46%   3e-32 86.75  AAD29806.1
hypothetical protein AXX17_AT4G17640 [Arabidopsis thaliana]       55.5   55.5  37%   3e-10 100.00 OAO96812.1


Alignments:
>putative disease resistance response protein [Arabidopsis thaliana]
Sequence ID: AAD29806.1 Length: 276
Range 1: 194 to 276

Score:117 bits(294), Expect:3e-32,
Method:Compositional matrix adjust.,
Identities:72/83(87%), Positives:78/83(93%), Gaps:0/83(0%)

Query  510  KKQNKSKSGQETGEKAGQrrrrkarrnkKEDILLSGDKPATTAESSRSMPEAEPTNASKE  331
            KKQNKSKSGQETGEKAG+RR++KARRNKKE+ILLSGDKPATTA+SSRSMPEAEP NASKE
Sbjct  194  KKQNKSKSGQETGEKAGKRRQKKARRNKKENILLSGDKPATTAKSSRSMPEAEPVNASKE  253

Query  330  ASDARVEADQTLLIKPVIAGQKS  262
            ASDA  EADQT LIKPVI GQK+
Sbjct  254  ASDAGGEADQTPLIKPVIVGQKN  276


>hypothetical protein AXX17_AT4G17640 [Arabidopsis thaliana]
Sequence ID: OAO96812.1 Length: 68
Range 1: 1 to 68

Score:55.5 bits(132), Expect:3e-10,
Method:Compositional matrix adjust.,
Identities:68/68(100%), Positives:68/68(100%), Gaps:0/68(0%)

Query  278  MTGFIssvwsastlaslaslealvgsasGIDRLLSAVVAGLSPESKISSflfrraflrrl  457
            MTGFISSVWSASTLASLASLEALVGSASGIDRLLSAVVAGLSPESKISSFLFRRAFLRRL
Sbjct  1    MTGFISSVWSASTLASLASLEALVGSASGIDRLLSAVVAGLSPESKISSFLFRRAFLRRL  60

Query  458  WPAFSPVS  481
            WPAFSPVS
Sbjct  61   WPAFSPVS  68


Query #225: XLOC_025147 Query ID: lcl|Query_30716 Length: 164

No significant similarity found.


Query #226: XLOC_025148 Query ID: lcl|Query_30717 Length: 954

No significant similarity found.


Query #227: XLOC_025149 Query ID: lcl|Query_30718 Length: 489

No significant similarity found.


Query #228: XLOC_025150 Query ID: lcl|Query_30719 Length: 628

No significant similarity found.


Query #229: XLOC_025151 Query ID: lcl|Query_30720 Length: 475

No significant similarity found.


Query #230: XLOC_025152 Query ID: lcl|Query_30721 Length: 522

No significant similarity found.


Query #231: XLOC_025153 Query ID: lcl|Query_30722 Length: 444

No significant similarity found.


Query #232: XLOC_025154 Query ID: lcl|Query_30723 Length: 777

No significant similarity found.


Query #233: XLOC_025155G Query ID: lcl|Query_30724 Length: 934

No significant similarity found.


Query #234: XLOC_025367 Query ID: lcl|Query_30725 Length: 750

Sequences producing significant alignments:
                                                                  Max    Total Query   E   Per.
Description                                                       Score  Score cover Value Ident  Accession
hypothetical protein AXX17_AT5G04080 [Arabidopsis thaliana]       118    118   26%   7e-34 100.00 OAO95973.1
unnamed protein product [Arabidopsis thaliana]                    95.5   140   28%   7e-27 87.50  CAA0400657.1
hypothetical protein AT2G05185 [Arabidopsis thaliana]             57.0   57.0  14%   3e-10 77.14  NP_001318201.1
hypothetical protein AT5G22608 [Arabidopsis thaliana]             56.6   56.6  15%   4e-10 71.05  NP_001078613.1
hypothetical protein AT5G22608 [Arabidopsis thaliana]             56.6   56.6  15%   5e-10 71.05  NP_001190364.1


Alignments:
>hypothetical protein AXX17_AT5G04080 [Arabidopsis thaliana]
Sequence ID: OAO95973.1 Length: 67
Range 1: 2 to 67

Score:118 bits(296), Expect:7e-34,
Method:Compositional matrix adjust.,
Identities:66/66(100%), Positives:66/66(100%), Gaps:0/66(0%)

Query  252  CSGNLKARSFQRHKDHLLVPCGVIGIRPQSWLLADYVIPDLSWEISHKFQTInpnpnpss  73
            CSGNLKARSFQRHKDHLLVPCGVIGIRPQSWLLADYVIPDLSWEISHKFQTINPNPNPSS
Sbjct  2    CSGNLKARSFQRHKDHLLVPCGVIGIRPQSWLLADYVIPDLSWEISHKFQTINPNPNPSS  61

Query  72   PVASYV  55
            PVASYV
Sbjct  62   PVASYV  67


>unnamed protein product [Arabidopsis thaliana]
Sequence ID: CAA0400657.1 Length: 87
Range 1: 20 to 67

Score:95.5 bits(236), Expect:7e-27,
Method:Compositional matrix adjust.,
Identities:42/48(88%), Positives:46/48(95%), Gaps:0/48(0%)

Query  240  LKARSFQRHKDHLLVPCGVIGIRPQSWLLADYVIPDLSWEISHKFQTI  97
            + ARSFQRHKDHLLVPCGVIGIRPQSWLLADYVIPDLSWEISHK+ ++
Sbjct  20   VSARSFQRHKDHLLVPCGVIGIRPQSWLLADYVIPDLSWEISHKYASL  67


Range 2: 1 to 22

Score:45.4 bits(106), Expect:7e-27,
Method:Compositional matrix adjust.,
Identities:21/22(95%), Positives:21/22(95%), Gaps:0/22(0%)

Query  400  MIRQYYEESKNDEVAQGKKVSG  335
            MIRQYYEESKNDEVAQGKKVS
Sbjct  1    MIRQYYEESKNDEVAQGKKVSA  22


>hypothetical protein AT2G05185 [Arabidopsis thaliana]
Sequence ID: NP_001318201.1 Length: 82
>hypothetical protein AT5G22608 [Arabidopsis thaliana]
Sequence ID: NP_001318620.1 Length: 82 >hypothetical protein AT2G05185 [Arabidopsis thaliana]
Sequence ID: AEC05903.2 Length: 82 >hypothetical protein AT5G22608 [Arabidopsis thaliana]
Sequence ID: AED93050.2 Length: 82
Range 1: 41 to 75

Score:57.0 bits(136), Expect:3e-10,
Method:Compositional matrix adjust.,
Identities:27/35(77%), Positives:29/35(82%), Gaps:0/35(0%)

Query  255  RCSGNLKARSFQRHKDHLLVPCGVIGIRPQSWLLA  151
            RCSGNLKARSFQ HKDHLLVPC ++G   QSWL A
Sbjct  41   RCSGNLKARSFQGHKDHLLVPCWLMGNCFQSWLFA  75


>hypothetical protein AT5G22608 [Arabidopsis thaliana]
Sequence ID: NP_001078613.1 Length: 88
>hypothetical protein AT2G05185 [Arabidopsis thaliana]
Sequence ID: NP_973423.1 Length: 88 >At2g05185 [Arabidopsis thaliana]
Sequence ID: ABD57453.1 Length: 88 >hypothetical protein AT2G05185 [Arabidopsis thaliana]
Sequence ID: AEC05902.1 Length: 88 >hypothetical protein AT5G22608 [Arabidopsis thaliana]
Sequence ID: AED93048.1 Length: 88 >unnamed protein product [Arabidopsis thaliana]
Sequence ID: VYS61435.1 Length: 88
Range 1: 41 to 78

Score:56.6 bits(135), Expect:4e-10,
Method:Compositional matrix adjust.,
Identities:27/38(71%), Positives:30/38(78%), Gaps:0/38(0%)

Query  255  RCSGNLKARSFQRHKDHLLVPCGVIGIRPQSWLLADYV  142
            RCSGNLKARSFQ HKDHLLVPC ++G   QSWL A  +
Sbjct  41   RCSGNLKARSFQGHKDHLLVPCWLMGNCFQSWLFAGLL  78


>hypothetical protein AT5G22608 [Arabidopsis thaliana]
Sequence ID: NP_001190364.1 Length: 96
>hypothetical protein AT5G22608 [Arabidopsis thaliana]
Sequence ID: AED93049.1 Length: 96
Range 1: 49 to 86

Score:56.6 bits(135), Expect:5e-10,
Method:Compositional matrix adjust.,
Identities:27/38(71%), Positives:30/38(78%), Gaps:0/38(0%)

Query  255  RCSGNLKARSFQRHKDHLLVPCGVIGIRPQSWLLADYV  142
            RCSGNLKARSFQ HKDHLLVPC ++G   QSWL A  +
Sbjct  49   RCSGNLKARSFQGHKDHLLVPCWLMGNCFQSWLFAGLL  86


Query #235: XLOC_025564 Query ID: lcl|Query_30726 Length: 620

No significant similarity found.


Query #236: XLOC_026472 Query ID: lcl|Query_30727 Length: 784

Sequences producing significant alignments:
                                                                  Max    Total Query   E   Per.
Description                                                       Score  Score cover Value Ident  Accession
hypothetical protein AXX17_AT5G26840 [Arabidopsis thaliana]       59.3   59.3  10%   1e-10 100.00 OAO92616.1


Alignments:
>hypothetical protein AXX17_AT5G26840 [Arabidopsis thaliana]
Sequence ID: OAO92616.1 Length: 113
Range 1: 78 to 104

Score:59.3 bits(142), Expect:1e-10,
Method:Compositional matrix adjust.,
Identities:27/27(100%), Positives:27/27(100%), Gaps:0/27(0%)

Query  215  SMNDCDQQKKVTSSISPFRDEEEGDPY  295
            SMNDCDQQKKVTSSISPFRDEEEGDPY
Sbjct  78   SMNDCDQQKKVTSSISPFRDEEEGDPY  104


Query #237: XLOC_026558 Query ID: lcl|Query_30728 Length: 1070

No significant similarity found.


Query #238: XLOC_026641 Query ID: lcl|Query_30729 Length: 600

No significant similarity found.


Query #239: XLOC_026796 Query ID: lcl|Query_30730 Length: 499

No significant similarity found.


Query #240: XLOC_026858 Query ID: lcl|Query_30731 Length: 720

Sequences producing significant alignments:
                                                                  Max    Total Query   E   Per.
Description                                                       Score  Score cover Value Ident  Accession
unnamed protein product [Arabidopsis thaliana]                    102    102   23%   3e-25 82.14  VYS62574.1
Werner syndrome-like exonuclease [Arabidopsis thaliana]           97.4   97.4  23%   1e-23 78.57  NP_193123.2
Werner syndrome-like exonuclease [Arabidopsis thaliana]           97.4   97.4  23%   1e-23 78.57  NP_974543.1
unnamed protein product [Arabidopsis thaliana]                    97.4   97.4  23%   1e-23 78.57  CAA0395138.1
WRNEXO [Arabidopsis thaliana]                                     97.4   97.4  23%   1e-23 78.57  OAP01060.1
putative protein [Arabidopsis thaliana]                           97.8   97.8  23%   2e-23 78.57  CAB36851.1


Alignments:
>unnamed protein product [Arabidopsis thaliana]
Sequence ID: VYS62574.1 Length: 289
Range 1: 87 to 142

Score:102 bits(253), Expect:3e-25,
Method:Compositional matrix adjust.,
Identities:46/56(82%), Positives:50/56(89%), Gaps:0/56(0%)

Query  369  RNFPTMRFGGRILYSKTTIEVDKRGLQLLKVLETKRDESGKAFVGFDIEWKPRIRK  536
            RNFP MRFGGRILYSKT  EVDKR +QLLKVL+TKRDESG+AFVGFDIEW+P  RK
Sbjct  87   RNFPAMRFGGRILYSKTATEVDKRAMQLLKVLDTKRDESGRAFVGFDIEWRPSFRK  142


>Werner syndrome-like exonuclease [Arabidopsis thaliana]
Sequence ID: NP_193123.2 Length: 285
>Werner syndrome-like exonuclease [Arabidopsis thaliana]
Sequence ID: AEE83338.1 Length: 285 >exonuclease [Arabidopsis thaliana]
Sequence ID: CAC14871.1 Length: 285
Range 1: 87 to 142

Score:97.4 bits(241), Expect:1e-23,
Method:Compositional matrix adjust.,
Identities:44/56(79%), Positives:48/56(85%), Gaps:0/56(0%)

Query  369  RNFPTMRFGGRILYSKTTIEVDKRGLQLLKVLETKRDESGKAFVGFDIEWKPRIRK  536
            RNFP MRFGGRILYSKT  EVDKR +QL+KVL+TKRDESG AFVG DIEW+P  RK
Sbjct  87   RNFPAMRFGGRILYSKTATEVDKRAMQLIKVLDTKRDESGIAFVGLDIEWRPSFRK  142


>Werner syndrome-like exonuclease [Arabidopsis thaliana]
Sequence ID: NP_974543.1 Length: 288
>RecName: Full=Werner Syndrome-like exonuclease [Arabidopsis thaliana]
Sequence ID: Q84LH3.1 Length: 288 >Werner Syndrome-like exonuclease [Arabidopsis thaliana]
Sequence ID: AAO33765.1 Length: 288 >At4g13870 [Arabidopsis thaliana]
Sequence ID: AAR24686.1 Length: 288 >Werner syndrome-like exonuclease [Arabidopsis thaliana]
Sequence ID: AEE83339.1 Length: 288
Range 1: 87 to 142

Score:97.4 bits(241), Expect:1e-23,
Method:Compositional matrix adjust.,
Identities:44/56(79%), Positives:48/56(85%), Gaps:0/56(0%)

Query  369  RNFPTMRFGGRILYSKTTIEVDKRGLQLLKVLETKRDESGKAFVGFDIEWKPRIRK  536
            RNFP MRFGGRILYSKT  EVDKR +QL+KVL+TKRDESG AFVG DIEW+P  RK
Sbjct  87   RNFPAMRFGGRILYSKTATEVDKRAMQLIKVLDTKRDESGIAFVGLDIEWRPSFRK  142


>unnamed protein product [Arabidopsis thaliana]
Sequence ID: CAA0395138.1 Length: 288
Range 1: 87 to 142

Score:97.4 bits(241), Expect:1e-23,
Method:Compositional matrix adjust.,
Identities:44/56(79%), Positives:48/56(85%), Gaps:0/56(0%)

Query  369  RNFPTMRFGGRILYSKTTIEVDKRGLQLLKVLETKRDESGKAFVGFDIEWKPRIRK  536
            RNFP MRFGGRILYSKT  EVDKR +QL+KVL+TKRDESG AFVG DIEW+P  RK
Sbjct  87   RNFPAMRFGGRILYSKTATEVDKRAMQLIKVLDTKRDESGIAFVGLDIEWRPSFRK  142


>WRNEXO [Arabidopsis thaliana]
Sequence ID: OAP01060.1 Length: 288
Range 1: 87 to 142

Score:97.4 bits(241), Expect:1e-23,
Method:Compositional matrix adjust.,
Identities:44/56(79%), Positives:48/56(85%), Gaps:0/56(0%)

Query  369  RNFPTMRFGGRILYSKTTIEVDKRGLQLLKVLETKRDESGKAFVGFDIEWKPRIRK  536
            RNFP MRFGGRILYSKT  EVDKR +QL+KVL+TKRDESG AFVG DIEW+P  RK
Sbjct  87   RNFPAMRFGGRILYSKTATEVDKRAMQLIKVLDTKRDESGIAFVGLDIEWRPSFRK  142


>putative protein [Arabidopsis thaliana]
Sequence ID: CAB36851.1 Length: 313
>putative protein [Arabidopsis thaliana]
Sequence ID: CAB78429.1 Length: 313
Range 1: 87 to 142

Score:97.8 bits(242), Expect:2e-23,
Method:Compositional matrix adjust.,
Identities:44/56(79%), Positives:48/56(85%), Gaps:0/56(0%)

Query  369  RNFPTMRFGGRILYSKTTIEVDKRGLQLLKVLETKRDESGKAFVGFDIEWKPRIRK  536
            RNFP MRFGGRILYSKT  EVDKR +QL+KVL+TKRDESG AFVG DIEW+P  RK
Sbjct  87   RNFPAMRFGGRILYSKTATEVDKRAMQLIKVLDTKRDESGIAFVGLDIEWRPSFRK  142


Query #241: XLOC_026885 Query ID: lcl|Query_30732 Length: 889

Sequences producing significant alignments:
                                                                  Max    Total Query   E   Per.
Description                                                       Score  Score cover Value Ident  Accession
putative protein [Arabidopsis thaliana]                           285    285   82%   8e-90 75.92  CAB88131.1
unnamed protein product [Arabidopsis thaliana]                    263    263   77%   2e-87 75.65  VYS59265.1
unnamed protein product [Arabidopsis thaliana]                    194    194   82%   3e-58 62.86  CAA0268354.1
hypothetical protein [Arabidopsis thaliana]                       182    182   77%   3e-55 62.17  AAG51087.1
heat shock protein [Arabidopsis thaliana]                         181    181   77%   4e-55 62.17  NP_174818.2
unnamed protein product [Arabidopsis thaliana]                    169    169   65%   7e-50 70.62  BAB01018.1
unnamed protein product [Arabidopsis thaliana]                    167    167   65%   2e-49 71.13  VYS59084.1
heat shock protein [Arabidopsis thaliana]                         168    168   65%   1e-48 70.62  NP_189673.2
unnamed protein product [Arabidopsis thaliana]                    154    154   47%   3e-43 72.54  CAA0384397.1
Myb/SANT-like DNA-binding domain protein [Arabidopsis thaliana]   138    217   43%   7e-40 78.57  NP_198457.1
unnamed protein product [Arabidopsis thaliana]                    132    132   77%   4e-34 40.26  VYS59124.1
unnamed protein product [Arabidopsis thaliana]                    126    126   77%   1e-33 38.63  CAA0384260.1
unnamed protein product [Arabidopsis thaliana]                    126    126   37%   1e-33 71.82  CAA0384198.1
unnamed protein product [Arabidopsis thaliana]                    109    258   51%   1e-27 75.00  VYS68398.1
hypothetical protein AXX17_AT3G34300 [Arabidopsis thaliana]       111    111   34%   4e-29 64.08  OAP04121.1
hypothetical protein AXX17_AT5G33440 [Arabidopsis thaliana]       99.4   99.4  34%   4e-25 58.25  OAO94257.1
Myb/SANT-like DNA-binding domain protein [Arabidopsis thaliana]   94.0   94.0  77%   2e-21 32.03  NP_683626.1
hypothetical protein AXX17_AT3G35980 [Arabidopsis thaliana]       74.3   74.3  30%   1e-14 65.17  OAP03107.1
heat shock protein [Arabidopsis thaliana]                         74.7   74.7  30%   1e-14 64.84  NP_189873.1
unnamed protein product [Arabidopsis thaliana]                    73.2   73.2  30%   2e-14 66.29  CAA0384316.1
hypothetical protein AXX17_AT3G37590 [Arabidopsis thaliana]       69.3   69.3  45%   2e-12 68.66  OAP06812.1
hypothetical protein AT1G43720 [Arabidopsis thaliana]             67.0   67.0  45%   6e-12 52.99  NP_175038.1
unnamed protein product [Arabidopsis thaliana]                    60.5   60.5  45%   1e-09 52.59  CAA0272678.1


Alignments:
>putative protein [Arabidopsis thaliana]
Sequence ID: CAB88131.1 Length: 775
Range 1: 43 to 287

Score:285 bits(728), Expect:8e-90,
Method:Compositional matrix adjust.,
Identities:186/245(76%), Positives:210/245(85%), Gaps:0/245(0%)

Query  16   FKKKTLIQLFDEAIAMNNYTLKHLTSIGREYMVEKFN*AFDMNITYVFFKNKLDEFKKSY  195
             ++KTLIQLFDEAIAMNNYTLK+ T+IGREYMVE FN AF+MNITY FFKNKLDEFKKSY
Sbjct  43   IQEKTLIQLFDEAIAMNNYTLKNPTAIGREYMVENFNIAFNMNITYGFFKNKLDEFKKSY  102

Query  196  KRWKFLTHKTGITVDPETSMIFASDVWWREQEFGCELTKSFYRKPPEFWDVMKRCLVLDD  375
            KRWK LTHKTGITVDP TS I AS+ WW EQEFGC+LTKS  RKPP FWDVM+RCLVL D
Sbjct  103  KRWKALTHKTGITVDPNTSFINASNAWWTEQEFGCKLTKSLNRKPPVFWDVMQRCLVLHD  162

Query  376  VqsqpqhssrqrreqLINEQAIdeesdddsdadsGNVPATQVPETQEEEKVYRAAIDGDE  555
            VQSQ QH +RQRREQLI+E  +DEE +D SD+DSGN+P T+VPET EEE+VYR  ID DE
Sbjct  163  VQSQSQHFARQRREQLIHEHGVDEEGEDYSDSDSGNMPQTKVPETLEEEEVYRVTIDDDE  222

Query  556  TFQNSAPRNQQRIRPNLqstgrrgsssqrsgrsaqisVGSGSRGNRTRQSFKTTIQDTIA  735
             FQNSA R QQR RPN QST RRGSS+QRSG S+Q+S+GSGSRG+R RQSF+TTIQD+I
Sbjct  223  IFQNSAARRQQRGRPNFQSTARRGSSAQRSGGSSQVSIGSGSRGSRRRQSFETTIQDSII  282

Query  736  GYREF  750
            G+REF
Sbjct  283  GFREF  287


>unnamed protein product [Arabidopsis thaliana]
Sequence ID: VYS59265.1 Length: 264
Range 1: 1 to 230

Score:263 bits(673), Expect:2e-87,
Method:Compositional matrix adjust.,
Identities:174/230(76%), Positives:196/230(85%), Gaps:0/230(0%)

Query  61   MNNYTLKHLTSIGREYMVEKFN*AFDMNITYVFFKNKLDEFKKSYKRWKFLTHKTGITVD  240
            MNNYTLK+ T+IGREYMVE FN AF+MNITY FFKNKLDEFKKSYKRWK LTHKTGITVD
Sbjct  1    MNNYTLKNPTAIGREYMVENFNIAFNMNITYGFFKNKLDEFKKSYKRWKALTHKTGITVD  60

Query  241  PETSMIFASDVWWREQEFGCELTKSFYRKPPEFWDVMKRCLVLDDVqsqpqhssrqrreq  420
            P TS I AS+ WW EQEFGC+LTKS  RKPP FWDVM+RCLVL DVQSQ QH +RQRREQ
Sbjct  61   PNTSFINASNAWWTEQEFGCKLTKSLNRKPPVFWDVMQRCLVLHDVQSQSQHFARQRREQ  120

Query  421  LINEQAIdeesdddsdadsGNVPATQVPETQEEEKVYRAAIDGDETFQNSAPRNQQRIRP  600
            LI+E  +DEE +D SD+DSGN+P T+VPET EEE+VYR  ID DE FQNSA R QQR RP
Sbjct  121  LIHEHGVDEEGEDYSDSDSGNMPQTKVPETLEEEEVYRVTIDDDEIFQNSAARRQQRGRP  180

Query  601  NLqstgrrgsssqrsgrsaqisVGSGSRGNRTRQSFKTTIQDTIAGYREF  750
            N QST RRGSS+QRSG S+Q+S+GSGSRG+R RQSF+TTIQD+I G+REF
Sbjct  181  NFQSTARRGSSAQRSGGSSQVSIGSGSRGSRRRQSFETTIQDSIIGFREF  230


>unnamed protein product [Arabidopsis thaliana]
Sequence ID: CAA0268354.1 Length: 450
Range 1: 43 to 256

Score:194 bits(493), Expect:3e-58,
Method:Compositional matrix adjust.,
Identities:154/245(63%), Positives:179/245(73%), Gaps:31/245(12%)

Query  16   FKKKTLIQLFDEAIAMNNYTLKHLTSIGREYMVEKFN*AFDMNITYVFFKNKLDEFKKSY  195
              +KTLIQLFDEAIAMNNYTLK+ T+I                +T
Sbjct  43   IHEKTLIQLFDEAIAMNNYTLKNPTAI----------------VT---------------  71

Query  196  KRWKFLTHKTGITVDPETSMIFASDVWWREQEFGCELTKSFYRKPPEFWDVMKRCLVLDD  375
            +  K L  KTGITVDP+TS I ASD WW EQEFGC+LTKS  RKPP FWDVM+RCLVL D
Sbjct  72   RGGKLLRTKTGITVDPDTSFINASDAWWTEQEFGCKLTKSLNRKPPVFWDVMQRCLVLHD  131

Query  376  VqsqpqhssrqrreqLINEQAIdeesdddsdadsGNVPATQVPETQEEEKVYRAAIDGDE  555
            VQSQ QHS+RQRREQLI+E  +DEE +D SD+DSGN+P T+VPETQEEE+VYRA ID DE
Sbjct  132  VQSQSQHSARQRREQLIHEHGVDEEGEDYSDSDSGNMPQTEVPETQEEEEVYRATIDDDE  191

Query  556  TFQNSAPRNQQRIRPNLqstgrrgsssqrsgrsaqisVGSGSRGNRTRQSFKTTIQDTIA  735
             FQNSA R QQR RPN  ST RRGSS+QRSG S+++S+GSGSRG++ RQSF+TTIQD I
Sbjct  192  IFQNSAARRQQRGRPNFWSTARRGSSAQRSGGSSRVSIGSGSRGSQRRQSFETTIQDIIT  251

Query  736  GYREF  750
            G+REF
Sbjct  252  GFREF  256


>hypothetical protein [Arabidopsis thaliana]
Sequence ID: AAG51087.1 Length: 287
Range 1: 1 to 198

Score:182 bits(461), Expect:3e-55,
Method:Compositional matrix adjust.,
Identities:143/230(62%), Positives:167/230(72%), Gaps:32/230(13%)

Query  61   MNNYTLKHLTSIGREYMVEKFN*AFDMNITYVFFKNKLDEFKKSYKRWKFLTHKTGITVD  240
            MNNYTLK+ T+IGREYM+                                L  KTGITVD
Sbjct  1    MNNYTLKNPTAIGREYML--------------------------------LRTKTGITVD  28

Query  241  PETSMIFASDVWWREQEFGCELTKSFYRKPPEFWDVMKRCLVLDDVqsqpqhssrqrreq  420
            P+TS I ASD WW EQEFGC+LTKS  RKPP FWDVM+RCLV+ DVQSQ QHS+RQRREQ
Sbjct  29   PDTSFINASDAWWTEQEFGCKLTKSLNRKPPVFWDVMQRCLVVHDVQSQSQHSARQRREQ  88

Query  421  LINEQAIdeesdddsdadsGNVPATQVPETQEEEKVYRAAIDGDETFQNSAPRNQQRIRP  600
            LI+E  +DEE +D SD+DSGN+P T+VPETQEEE+VYR  ID DE FQNSA R QQR RP
Sbjct  89   LIHEHGVDEEGEDYSDSDSGNMPQTEVPETQEEEEVYRVTIDDDEIFQNSAARRQQRGRP  148

Query  601  NLqstgrrgsssqrsgrsaqisVGSGSRGNRTRQSFKTTIQDTIAGYREF  750
            N  ST RRGSS+QRSG S+++S+GSGSRG++ RQSF+TTIQD I G+REF
Sbjct  149  NFWSTARRGSSAQRSGGSSRVSIGSGSRGSQRRQSFETTIQDIITGFREF  198


>heat shock protein [Arabidopsis thaliana]
Sequence ID: NP_174818.2 Length: 287
>heat shock protein [Arabidopsis thaliana]
Sequence ID: AEE31829.1 Length: 287
Range 1: 1 to 198

Score:181 bits(460), Expect:4e-55,
Method:Compositional matrix adjust.,
Identities:143/230(62%), Positives:167/230(72%), Gaps:32/230(13%)

Query  61   MNNYTLKHLTSIGREYMVEKFN*AFDMNITYVFFKNKLDEFKKSYKRWKFLTHKTGITVD  240
            MNNYTLK+ T+IGREYM+                                L  KTGITVD
Sbjct  1    MNNYTLKNPTAIGREYML--------------------------------LRTKTGITVD  28

Query  241  PETSMIFASDVWWREQEFGCELTKSFYRKPPEFWDVMKRCLVLDDVqsqpqhssrqrreq  420
            P+TS I ASD WW EQEFGC+LTKS  RKPP FWDVM+RCLV+ DVQSQ QHS+RQRREQ
Sbjct  29   PDTSFINASDAWWTEQEFGCKLTKSLNRKPPVFWDVMQRCLVVHDVQSQSQHSARQRREQ  88

Query  421  LINEQAIdeesdddsdadsGNVPATQVPETQEEEKVYRAAIDGDETFQNSAPRNQQRIRP  600
            LI+E  +DEE +D SD+DSGN+P T+VPETQEEE+VYR  ID DE FQNSA R QQR RP
Sbjct  89   LIHEHGVDEEGEDYSDSDSGNMPQTEVPETQEEEEVYRVTIDDDEIFQNSAARRQQRGRP  148

Query  601  NLqstgrrgsssqrsgrsaqisVGSGSRGNRTRQSFKTTIQDTIAGYREF  750
            N  ST RRGSS+QRSG S+++S+GSGSRG++ RQSF+TTIQD I G+REF
Sbjct  149  NFWSTARRGSSAQRSGGSSRVSIGSGSRGSQRRQSFETTIQDIITGFREF  198


>unnamed protein product [Arabidopsis thaliana]
Sequence ID: BAB01018.1 Length: 313
Range 1: 33 to 226

Score:169 bits(427), Expect:7e-50,
Method:Compositional matrix adjust.,
Identities:137/194(71%), Positives:162/194(83%), Gaps:0/194(0%)

Query  169  KLDEFKKSYKRWKFLTHKTGITVDPETSMIFASDVWWREQEFGCELTKSFYRKPPEFWDV  348
            +LDEF+KSYK+WK LTHKTGITVDP+TS I ASD WW EQEFGC+LTK+  RKPP FWDV
Sbjct  33   RLDEFRKSYKKWKALTHKTGITVDPDTSFINASDAWWTEQEFGCKLTKNLNRKPPVFWDV  92

Query  349  MKRCLVLDDVqsqpqhssrqrreqLINEQAIdeesdddsdadsGNVPATQVPETQEEEKV  528
            M+RCLVL DVQSQ QH + QRREQLI+E  +DEE +D SD+DSGN+P T+VPETQEEE+V
Sbjct  93   MQRCLVLHDVQSQSQHFACQRREQLIHEHGVDEEGEDYSDSDSGNMPQTEVPETQEEEEV  152

Query  529  YRAAIDGDETFQNSAPRNQQRIRPNLqstgrrgsssqrsgrsaqisVGSGSRGNRTRQSF  708
            YR  ID DE FQNSA R QQR RPN QST RRGSS+QRSG ++++S+GSGS G+R RQSF
Sbjct  153  YRVTIDDDEIFQNSAARRQQRGRPNFQSTARRGSSAQRSGGTSRVSIGSGSGGSRRRQSF  212

Query  709  KTTIQDTIAGYREF  750
            +TTIQD+I  + EF
Sbjct  213  ETTIQDSITSFGEF  226


>unnamed protein product [Arabidopsis thaliana]
Sequence ID: VYS59084.1 Length: 313
Range 1: 33 to 226

Score:167 bits(424), Expect:2e-49,
Method:Compositional matrix adjust.,
Identities:138/194(71%), Positives:162/194(83%), Gaps:0/194(0%)

Query  169  KLDEFKKSYKRWKFLTHKTGITVDPETSMIFASDVWWREQEFGCELTKSFYRKPPEFWDV  348
            +LDEFKKSYK+WK LTHKTGITVDP+TS I ASD WW EQEFGC+LTK+  RKPP FWDV
Sbjct  33   RLDEFKKSYKKWKALTHKTGITVDPDTSFINASDAWWTEQEFGCKLTKNLNRKPPVFWDV  92

Query  349  MKRCLVLDDVqsqpqhssrqrreqLINEQAIdeesdddsdadsGNVPATQVPETQEEEKV  528
            M+RCLVL DVQSQ QHS+ QRREQLI+E  +DEE +D SD+DSGN+P T+VPETQEEE+V
Sbjct  93   MQRCLVLHDVQSQSQHSACQRREQLIHEHGVDEEGEDYSDSDSGNMPQTEVPETQEEEEV  152

Query  529  YRAAIDGDETFQNSAPRNQQRIRPNLqstgrrgsssqrsgrsaqisVGSGSRGNRTRQSF  708
            YR  ID  E FQNSA R QQR RPN QST RRGSS+QRSG ++++S+GSGS G+R RQSF
Sbjct  153  YRVTIDDGEIFQNSAARRQQRGRPNFQSTARRGSSAQRSGGTSRVSIGSGSGGSRRRQSF  212

Query  709  KTTIQDTIAGYREF  750
            +TTIQD+I  + EF
Sbjct  213  ETTIQDSITSFGEF  226


>heat shock protein [Arabidopsis thaliana]
Sequence ID: NP_189673.2 Length: 397
>heat shock protein [Arabidopsis thaliana]
Sequence ID: AEE77648.1 Length: 397
Range 1: 33 to 226

Score:168 bits(425), Expect:1e-48,
Method:Compositional matrix adjust.,
Identities:137/194(71%), Positives:162/194(83%), Gaps:0/194(0%)

Query  169  KLDEFKKSYKRWKFLTHKTGITVDPETSMIFASDVWWREQEFGCELTKSFYRKPPEFWDV  348
            +LDEF+KSYK+WK LTHKTGITVDP+TS I ASD WW EQEFGC+LTK+  RKPP FWDV
Sbjct  33   RLDEFRKSYKKWKALTHKTGITVDPDTSFINASDAWWTEQEFGCKLTKNLNRKPPVFWDV  92

Query  349  MKRCLVLDDVqsqpqhssrqrreqLINEQAIdeesdddsdadsGNVPATQVPETQEEEKV  528
            M+RCLVL DVQSQ QH + QRREQLI+E  +DEE +D SD+DSGN+P T+VPETQEEE+V
Sbjct  93   MQRCLVLHDVQSQSQHFACQRREQLIHEHGVDEEGEDYSDSDSGNMPQTEVPETQEEEEV  152

Query  529  YRAAIDGDETFQNSAPRNQQRIRPNLqstgrrgsssqrsgrsaqisVGSGSRGNRTRQSF  708
            YR  ID DE FQNSA R QQR RPN QST RRGSS+QRSG ++++S+GSGS G+R RQSF
Sbjct  153  YRVTIDDDEIFQNSAARRQQRGRPNFQSTARRGSSAQRSGGTSRVSIGSGSGGSRRRQSF  212

Query  709  KTTIQDTIAGYREF  750
            +TTIQD+I  + EF
Sbjct  213  ETTIQDSITSFGEF  226


>unnamed protein product [Arabidopsis thaliana]
Sequence ID: CAA0384397.1 Length: 419
Range 1: 33 to 174

Score:154 bits(390), Expect:3e-43,
Method:Compositional matrix adjust.,
Identities:103/142(73%), Positives:115/142(80%), Gaps:0/142(0%)

Query  169  KLDEFKKSYKRWKFLTHKTGITVDPETSMIFASDVWWREQEFGCELTKSFYRKPPEFWDV  348
            +LDEFKKSYKRWK LTHK GITVDP TS I ASD WW EQEFGC+LTKS  RKP  FWDV
Sbjct  33   QLDEFKKSYKRWKALTHKIGITVDPNTSFINASDAWWTEQEFGCKLTKSLNRKPLVFWDV  92

Query  349  MKRCLVLDDVqsqpqhssrqrreqLINEQAIdeesdddsdadsGNVPATQVPETQEEEKV  528
            M+RCLVL DVQSQ QH +RQRREQLI+E  +DEE +D SD+DSGN+P  +VPET EEE+V
Sbjct  93   MQRCLVLHDVQSQSQHFARQRREQLIHEHGVDEEGEDYSDSDSGNMPQIEVPETLEEEEV  152

Query  529  YRAAIDGDETFQNSAPRNQQRI  594
            YR  ID DE FQNSA R QQR+
Sbjct  153  YRVTIDDDEIFQNSAARRQQRV  174


>Myb/SANT-like DNA-binding domain protein [Arabidopsis thaliana]
Sequence ID: NP_198457.1 Length: 153
>Myb/SANT-like DNA-binding domain protein [Arabidopsis thaliana]
Sequence ID: AED94042.1 Length: 153 >unnamed protein product [Arabidopsis thaliana]
Sequence ID: BAA96882.1 Length: 153
Range 1: 1 to 84

Score:138 bits(347), Expect:7e-40,
Method:Compositional matrix adjust.,
Identities:66/84(79%), Positives:73/84(86%), Gaps:1/84(1%)

Query  61   MNNYTLKHLTSIGREYMVEKFN*AFDMNITYVFFKNKLDEFKKSYKRWKFLTHKTGITVD  240
            MNNYT+KH TSIGR+YMVEKFN  F+MN TYVFFKNK DE KKSYK WKFLTHKTGI+VD
Sbjct  1    MNNYTIKHPTSIGRDYMVEKFNQVFNMNRTYVFFKNKHDELKKSYKGWKFLTHKTGISVD  60

Query  241  PETSMIFASDVWWREQEF-GCELT  309
            P+TSMIFA DVWWRE+EF G E+
Sbjct  61   PKTSMIFAYDVWWREREFVGVEVV  84


Range 2: 79 to 123

Score:79.7 bits(195), Expect:1e-17,
Method:Compositional matrix adjust.,
Identities:41/45(91%), Positives:41/45(91%), Gaps:0/45(0%)

Query  617  VDVEVVHKDLGEAHKFLLEVVHEEIVHDNLLKQPYKTLLLVIENL  751
            V VEVVHKDLGEAHKFLLEVVHEEIV DNLLKQ YKTLLLVIEN
Sbjct  79   VGVEVVHKDLGEAHKFLLEVVHEEIVEDNLLKQLYKTLLLVIENF  123


>unnamed protein product [Arabidopsis thaliana]
Sequence ID: VYS59124.1 Length: 593
Range 1: 1 to 188

Score:132 bits(331), Expect:4e-34,
Method:Compositional matrix adjust.,
Identities:93/231(40%), Positives:131/231(56%), Gaps:44/231(19%)

Query  61   MNNYTLKHLTSIGREYMVEKFN*AFDMNITYVFFKNKLDEFKKSYKRWKFLTHK-TGITV  237
            M NY +K  T +GRE++++KFN  F++NI Y FFK KLD+ K+ YK++K L  K TGI+V
Sbjct  1    MGNYRIKDPTPLGREFILDKFNKKFNLNINYQFFKEKLDQLKRKYKKYKELMKKSTGISV  60

Query  238  DPETSMIFASDVWWREQEFGCELTKSFYRKPPEFWDVMKRCLVLDDVqsqpqhssrqrre  417
            D  TS+I AS+ WW+++E  C++ KSF R PPE WDVM+RC +L +V SQPQ S  QRRE
Sbjct  61   DTTTSVISASNSWWQKREV-CKIIKSFKRNPPEIWDVMQRCFILYEVHSQPQFSVNQRRE  119

Query  418  qLINEQAIdeesdddsdadsGNVPATQVPETQEEEKVYRAAIDGDETFQNSAPRNQQRIR  597
            QL+N+           D D G++P +++ +        R  +    + Q  A  +Q   R
Sbjct  120  QLMND---------GLDNDEGHIPTSRIQQ--------RGGLRRGSSSQRGAGNSQISTR  162

Query  598  PNLqstgrrgsssqrsgrsaqisVGSGSRGNRTRQSFKTTIQDTIAGYREF  750
                                       SRG+R +QSF+TT+ DTI G REF
Sbjct  163  -------------------------ISSRGSRRKQSFETTLTDTITGLREF  188


>unnamed protein product [Arabidopsis thaliana]
Sequence ID: CAA0384260.1 Length: 309
Range 1: 1 to 195

Score:126 bits(317), Expect:1e-33,
Method:Compositional matrix adjust.,
Identities:90/233(39%), Positives:126/233(54%), Gaps:41/233(17%)

Query  61   MNNYTLKHLTSIGREYMVEKFN*AFDMNITYVFFKNKLDEFKKSYKRWK-FLTHKTGITV  237
            M NY  K  T +GRE++++KFN  F++NI Y F K KLD+ K+ YK++K  + + TGI+V
Sbjct  1    MGNYRFKDPTLLGREFILDKFNKEFNLNINYQFVKEKLDQVKRKYKKYKELMKNSTGISV  60

Query  238  DPETSMIFASDVWWREQEFGCELTKSFYRKPPEFWDVMKRCLVLDDVqsqpqhssrqrre  417
            D  TS+I  S+ WW+E+E  C++ KSF RKPPE WDVM+RC +L DV SQPQ    QRRE
Sbjct  61   DTTTSVISTSNSWWQEREV-CKIIKSFKRKPPELWDVMQRCFILYDVHSQPQFFVNQRRE  119

Query  418  qLINEQAIdeesdddsdadsGNVPAT--QVPETQEEEKVYRAAIDGDETFQNSAPRNQQR  591
            QL+N+                + PA   Q+P ++ ++   R  +    + Q  A   Q
Sbjct  120  QLVND---------GWTMMKNSSPAAPFQIPTSRIQQ---RGGLRRGSSSQRGAGNFQIS  167

Query  592  IRPNLqstgrrgsssqrsgrsaqisVGSGSRGNRTRQSFKTTIQDTIAGYREF  750
             R                          GSRG+R +Q F+TT+ D I G REF
Sbjct  168  TR-------------------------IGSRGSRRKQLFETTLTDIITGLREF  195


>unnamed protein product [Arabidopsis thaliana]
Sequence ID: CAA0384198.1 Length: 292
Range 1: 33 to 142

Score:126 bits(316), Expect:1e-33,
Method:Compositional matrix adjust.,
Identities:79/110(72%), Positives:92/110(83%), Gaps:0/110(0%)

Query  169  KLDEFKKSYKRWKFLTHKTGITVDPETSMIFASDVWWREQEFGCELTKSFYRKPPEFWDV  348
            +LDEFKKSYK+WK LTHKTGITVDP+TS I ASD WW EQEFGC+LTK+  RK   FWDV
Sbjct  33   RLDEFKKSYKKWKALTHKTGITVDPDTSFINASDAWWTEQEFGCKLTKNLNRKSSVFWDV  92

Query  349  MKRCLVLDDVqsqpqhssrqrreqLINEQAIdeesdddsdadsGNVPATQ  498
            M+RCLVL DVQSQ QHS+ QRREQLI+E  +DEE +D SD+DSGN+P T+
Sbjct  93   MQRCLVLHDVQSQSQHSACQRREQLIHEHGVDEEGEDYSDSDSGNMPQTE  142


>unnamed protein product [Arabidopsis thaliana]
Sequence ID: VYS68398.1 Length: 264
Range 1: 102 to 149

Score:82.0 bits(201), Expect:3e-29,
Method:Compositional matrix adjust.,
Identities:42/48(88%), Positives:42/48(87%), Gaps:0/48(0%)

Query  599  LICSQQVDVEVVHKDLGEAHKFLLEVVHEEIVHDNLLKQPYKTLLLVI  742
            LI SQQV VEVVHKDLGEAHKFLLEVVHEEIV DNLLKQ YKTLLL
Sbjct  102  LIWSQQVGVEVVHKDLGEAHKFLLEVVHEEIVEDNLLKQLYKTLLLYF  149


Range 2: 64 to 101

Score:67.0 bits(162), Expect:3e-29,
Method:Compositional matrix adjust.,
Identities:31/38(82%), Positives:35/38(92%), Gaps:0/38(0%)

Query  484  VPATQVPETQEEEKVYRAAIDGDETFQNSAPRNQQRIR  597
            +PATQV +TQEEE+VYR AID +ETFQNSAPRNQQRIR
Sbjct  64   MPATQVSKTQEEEEVYRVAIDDEETFQNSAPRNQQRIR  101


Range 3: 1 to 68

Score:109 bits(273), Expect:1e-27,
Method:Compositional matrix adjust.,
Identities:51/68(75%), Positives:57/68(83%), Gaps:0/68(0%)

Query  109  MVEKFN*AFDMNITYVFFKNKLDEFKKSYKRWKFLTHKTGITVDPETSMIFASDVWWREQ  288
            MVEKFN  F+MN TYVFFKNK D+ KKSYK WKFLTHKTGI+VDP+TSMIFA DVWWRE+
Sbjct  1    MVEKFNQVFNMNRTYVFFKNKHDKLKKSYKGWKFLTHKTGISVDPKTSMIFAYDVWWRER  60

Query  289  EFGCELTK  312
            EF    T+
Sbjct  61   EFVMPATQ  68


>hypothetical protein AXX17_AT3G34300 [Arabidopsis thaliana]
Sequence ID: OAP04121.1 Length: 199
Range 1: 3 to 105

Score:111 bits(278), Expect:4e-29,
Method:Compositional matrix adjust.,
Identities:66/103(64%), Positives:77/103(74%), Gaps:0/103(0%)

Query  440  LMKKAMMTQTLIVVMCQQHKFPRHkkkkkCIVLLLMVMKRSKILLLETNNELGLICSQQV  619
            LMKK  +TQ + VV C + +F + KKKKKCIVL LM++K SKILLLE NNE+G I SQ +
Sbjct  3    LMKKVKITQIVTVVTCHKPRFLKRKKKKKCIVLPLMMVKYSKILLLEGNNEVGQIFSQPL  62

Query  620  DVEVVHKDLGEAHKFLLEVVHEEIVHDNLLKQPYKTLLLVIEN  748
            DVEVVHKDL E H+FLLEVV EE+  DN LKQ YKT  L  EN
Sbjct  63   DVEVVHKDLEEPHEFLLEVVQEEVGDDNPLKQQYKTASLASEN  105


>hypothetical protein AXX17_AT5G33440 [Arabidopsis thaliana]
Sequence ID: OAO94257.1 Length: 130
Range 1: 1 to 100

Score:99.4 bits(246), Expect:4e-25,
Method:Compositional matrix adjust.,
Identities:60/103(58%), Positives:66/103(64%), Gaps:3/103(2%)

Query  443  MKKAMMTQTLIVVMCQQHKFPRHkkkkkCIVLLLMVMKRSKILLLETNNELGLICSQQVD  622
            MKKAMMTQTL V   Q  K    ++  +  +      + S          + LI SQQV
Sbjct  1    MKKAMMTQTLTVPATQVSKTQEEEEVYRVAIDDDETFQNSAP---RNQQRIRLIWSQQVG  57

Query  623  VEVVHKDLGEAHKFLLEVVHEEIVHDNLLKQPYKTLLLVIENL  751
            VEVVHKDLGEAHKFLLEVVHEEIV DNLLKQ YKTLLLVIEN
Sbjct  58   VEVVHKDLGEAHKFLLEVVHEEIVEDNLLKQLYKTLLLVIENF  100


>Myb/SANT-like DNA-binding domain protein [Arabidopsis thaliana]
Sequence ID: NP_683626.1 Length: 330
>Myb/SANT-like DNA-binding domain protein [Arabidopsis thaliana]
Sequence ID: AEE77701.1 Length: 330
Range 1: 1 to 180

Score:94.0 bits(232), Expect:2e-21,
Method:Compositional matrix adjust.,
Identities:74/231(32%), Positives:114/231(49%), Gaps:52/231(22%)

Query  61   MNNYTLKHLTSIGREYMVEKFN*AFDMNITYVFFKNKLDEFKKSYKRWK-FLTHKTGITV  237
            M NY  K  T +GRE++++KFN  F++NI Y F K KLD+ K+ YK++K  + + TGI+V
Sbjct  1    MGNYRFKDPTILGREFILDKFNKEFNLNINYQFVKEKLDQLKRKYKKYKELMKNSTGISV  60

Query  238  DPETSMIFASDVWWREQEFGCELTKSFYRKPPEFWDVMKRCLVLDDVqsqpqhssrqrre  417
            D  TS+I  S+ WW+E+E             P+F    +R  +++D
Sbjct  61   DTTTSVISTSNSWWQERE-------------PQFSVNQRREQLVND--------------  93

Query  418  qLINEQAIdeesdddsdadsGNVPATQVPETQEEEKVYRAAIDGDETFQNSAPRNQQRIR  597
             L N++          +   G++  +QV ETQE E++  + I      +  +  + QR
Sbjct  94   GLDNDEG-----HVYFETYDGDMQDSQVLETQENEEIPTSRIQQRGGLRRGS--SSQRGA  146

Query  598  PNLqstgrrgsssqrsgrsaqisVGSGSRGNRTRQSFKTTIQDTIAGYREF  750
             N Q + R                  GSRG+R +QSF+TT+ DTI G REF
Sbjct  147  GNFQISTRI-----------------GSRGSRRKQSFETTLTDTITGLREF  180


>hypothetical protein AXX17_AT3G35980 [Arabidopsis thaliana]
Sequence ID: OAP03107.1 Length: 264
Range 1: 1 to 89

Score:74.3 bits(181), Expect:1e-14,
Method:Compositional matrix adjust.,
Identities:58/89(65%), Positives:71/89(79%), Gaps:0/89(0%)

Query  484  VPATQVPETQEEEKVYRAAIDGDETFQNSAPRNQQRIRPNLqstgrrgsssqrsgrsaqi  663
            +P T+VPE QEEE+VYR  ID DE FQ+ A R QQR RPNLQST RR SS+QR G S+++
Sbjct  1    MPQTEVPEMQEEEEVYRVTIDDDEIFQHCAARRQQRGRPNLQSTARRESSAQRCGGSSRV  60

Query  664  sVGSGSRGNRTRQSFKTTIQDTIAGYREF  750
            S+ SGSRG+R RQSF+TTI+D I G+REF
Sbjct  61   SIRSGSRGSRKRQSFETTIEDNITGFREF  89


>heat shock protein [Arabidopsis thaliana]
Sequence ID: NP_189873.1 Length: 312
>heat shock protein [Arabidopsis thaliana]
Sequence ID: AEE77757.1 Length: 312 >hypothetical protein [Arabidopsis thaliana]
Sequence ID: CAB86674.1 Length: 312
Range 1: 31 to 121

Score:74.7 bits(182), Expect:1e-14,
Method:Compositional matrix adjust.,
Identities:59/91(65%), Positives:73/91(80%), Gaps:0/91(0%)

Query  478  GNVPATQVPETQEEEKVYRAAIDGDETFQNSAPRNQQRIRPNLqstgrrgsssqrsgrsa  657
            GN+P T+VPE QEEE+VYR  ID DE FQ+ A R QQR RPNLQST +R SS+QRSG S+
Sbjct  31   GNMPQTEVPEMQEEEEVYRVTIDDDEIFQHCAARRQQRGRPNLQSTAKRESSAQRSGGSS  90

Query  658  qisVGSGSRGNRTRQSFKTTIQDTIAGYREF  750
            ++S+ SGSRG+R RQSF+TTI+D I  +REF
Sbjct  91   RVSIRSGSRGSRKRQSFETTIEDNITSFREF  121


>unnamed protein product [Arabidopsis thaliana]
Sequence ID: CAA0384316.1 Length: 228
Range 1: 1 to 89

Score:73.2 bits(178), Expect:2e-14,
Method:Compositional matrix adjust.,
Identities:59/89(66%), Positives:72/89(80%), Gaps:0/89(0%)

Query  484  VPATQVPETQEEEKVYRAAIDGDETFQNSAPRNQQRIRPNLqstgrrgsssqrsgrsaqi  663
            +P T+VPE QEEE+VYR  ID DE FQ+ A R QQR RPNLQST RR SS+QRSG S+++
Sbjct  1    MPQTEVPEMQEEEEVYRVTIDDDEIFQHCAARRQQRGRPNLQSTARRKSSAQRSGGSSRV  60

Query  664  sVGSGSRGNRTRQSFKTTIQDTIAGYREF  750
            S+ SGSRG+R RQSF+TTI+D I G+REF
Sbjct  61   SIRSGSRGSRKRQSFETTIEDNITGFREF  89


>hypothetical protein AXX17_AT3G37590 [Arabidopsis thaliana]
Sequence ID: OAP06812.1 Length: 903
Range 1: 1 to 134

Score:69.3 bits(168), Expect:2e-12,
Method:Compositional matrix adjust.,
Identities:92/134(69%), Positives:111/134(82%), Gaps:0/134(0%)

Query  349  MKRCLVLDDVqsqpqhssrqrreqLINEQAIdeesdddsdadsGNVPATQVPETQEEEKV  528
            M+RCLVL DVQSQ QH +RQRREQLI+E  +DEE +D SD+DSGN+P T+VPET EEE+V
Sbjct  1    MQRCLVLHDVQSQSQHFARQRREQLIHEHGVDEEGEDYSDSDSGNMPQTEVPETLEEEEV  60

Query  529  YRAAIDGDETFQNSAPRNQQRIRPNLqstgrrgsssqrsgrsaqisVGSGSRGNRTRQSF  708
            YR  ID DE FQNSA   QQR RPN QST +RGSS+QRSG S+++S+GSGSR +R RQSF
Sbjct  61   YRVTIDDDEIFQNSAAGRQQRGRPNFQSTAKRGSSAQRSGGSSRVSIGSGSRRSRRRQSF  120

Query  709  KTTIQDTIAGYREF  750
            +TTIQD+I G+REF
Sbjct  121  ETTIQDSITGFREF  134


>hypothetical protein AT1G43720 [Arabidopsis thaliana]
Sequence ID: NP_175038.1 Length: 314
>hypothetical protein AT1G43720 [Arabidopsis thaliana]
Sequence ID: AEE31990.1 Length: 314
Range 1: 1 to 116

Score:67.0 bits(162), Expect:6e-12,
Method:Compositional matrix adjust.,
Identities:71/134(53%), Positives:89/134(66%), Gaps:18/134(13%)

Query  349  MKRCLVLDDVqsqpqhssrqrreqLINEQAIdeesdddsdadsGNVPATQVPETQEEEKV  528
            M+RCLVL DVQSQ QHS+RQRREQLI+E  +DEE +D SD+DSGN+P  +VPETQEEE+V
Sbjct  1    MQRCLVLHDVQSQSQHSARQRREQLIHEHGVDEEGEDYSDSDSGNMPEAEVPETQEEEEV  60

Query  529  YRAAIDGDETFQNSAPRNQQRIRPNLqstgrrgsssqrsgrsaqisVGSGSRGNRTRQSF  708
            YR  ID DE+        Q+    +L                  +S+GSGSR +  RQSF
Sbjct  61   YRVTIDDDESTARRGSSAQRSGGSSL------------------VSIGSGSRRSHRRQSF  102

Query  709  KTTIQDTIAGYREF  750
            +TTIQD+I G+REF
Sbjct  103  ETTIQDSITGFREF  116


>unnamed protein product [Arabidopsis thaliana]
Sequence ID: CAA0272678.1 Length: 315
Range 1: 1 to 117

Score:60.5 bits(145), Expect:1e-09,
Method:Compositional matrix adjust.,
Identities:71/135(53%), Positives:88/135(65%), Gaps:19/135(14%)

Query  349  MKRCLVLDDVqsqpqhssrqrreqLINEQAIdeesdddsdadsGNVPATQVPETQEEEKV  528
            M+RCLVL DVQSQ QHS+RQRREQLI+E  +DEE +D SD+DSGN+P T+VPETQEEE+
Sbjct  1    MQRCLVLHDVQSQSQHSARQRREQLIHEHGVDEEGEDYSDSDSGNMPETEVPETQEEEEE  60

Query  529  -YRAAIDGDETFQNSAPRNQQRIRPNLqstgrrgsssqrsgrsaqisVGSGSRGNRTRQS  705
             YR  ID DE+        Q+    +L S                  +GSGSR +  RQS
Sbjct  61   VYRVTIDDDESTARRGSSAQRSGGSSLVS------------------IGSGSRRSHRRQS  102

Query  706  FKTTIQDTIAGYREF  750
            F+TTIQD+I G+REF
Sbjct  103  FETTIQDSITGFREF  117


Query #242: XLOC_027072 Query ID: lcl|Query_30733 Length: 508

No significant similarity found.


Query #243: XLOC_027080 Query ID: lcl|Query_30734 Length: 728

No significant similarity found.


Query #244: XLOC_027111 Query ID: lcl|Query_30735 Length: 742

Sequences producing significant alignments:
                                                                  Max    Total Query   E   Per.
Description                                                       Score  Score cover Value Ident  Accession
hypothetical protein AT4G10843 [Arabidopsis thaliana]             81.3   81.3  45%   4e-19 48.21  NP_001328696.1
hypothetical protein AT4G10843 [Arabidopsis thaliana]             82.4   82.4  45%   5e-19 48.21  NP_001328697.1
hypothetical protein AT4G10843 [Arabidopsis thaliana]             82.4   82.4  45%   6e-19 48.21  NP_001328695.1
unnamed protein product [Arabidopsis thaliana]                    75.5   147   29%   4e-16 92.31  CAA0405987.1
unnamed protein product [Arabidopsis thaliana]                    73.9   136   29%   1e-15 87.18  VYS62247.1
unnamed protein product [Arabidopsis thaliana]                    75.9   144   29%   1e-15 87.50  CAA0394571.1
hypothetical protein AT5G37793 [Arabidopsis thaliana]             71.6   139   33%   2e-15 97.96  NP_001331870.1
unnamed protein product [Arabidopsis thaliana]                    72.0   139   33%   7e-15 97.96  VYS68533.1
unnamed protein product [Arabidopsis thaliana]                    66.2   66.2  13%   8e-14 93.94  VYS67928.1
nucleoporin (DUF3414) [Arabidopsis thaliana]                      55.1   55.1  15%   5e-08 56.25  NP_195587.5
hypothetical protein AXX17_AT4G44080 [Arabidopsis thaliana]       55.1   55.1  15%   5e-08 56.25  OAO97877.1
unnamed protein product [Arabidopsis thaliana]                    55.1   55.1  15%   5e-08 56.25  CAA0397936.1


Alignments:
>hypothetical protein AT4G10843 [Arabidopsis thaliana]
Sequence ID: NP_001328696.1 Length: 105
>hypothetical protein AT4G10843 [Arabidopsis thaliana]
Sequence ID: NP_567365.4 Length: 105 >hypothetical protein At4g10845 [Arabidopsis thaliana]
Sequence ID: AAX23890.1 Length: 105 >hypothetical protein AT4G10843 [Arabidopsis thaliana]
Sequence ID: ANM66824.1 Length: 105 >hypothetical protein AT4G10843 [Arabidopsis thaliana]
Sequence ID: ANM66825.1 Length: 105
Range 1: 1 to 80

Score:81.3 bits(199), Expect:4e-19,
Method:Compositional matrix adjust.,
Identities:54/112(48%), Positives:64/112(57%), Gaps:32/112(28%)

Query  399  MKLRRKRWSWLIV*VFFA*kkkkkTV**SVISLL*AKIIKEMSSASLHPFFLVLIFLRNF  578
            MKLRRKRWSWL++                       K +K    +SL        F+  F
Sbjct  1    MKLRRKRWSWLVIE---------------------TKSLK----SSLPERRRRRRFI--F  33

Query  579  LLCED*L*YGDDIMDVSLFTF*TEETLIEDKSLSFCTCNGERWRRLFSFYKV  734
             LC+        ++DV+LFTF TEETLIEDKSLSFCTCNGERWRRLFSFYK+
Sbjct  34   ALCD-----SSFLLDVNLFTFWTEETLIEDKSLSFCTCNGERWRRLFSFYKL  80


>hypothetical protein AT4G10843 [Arabidopsis thaliana]
Sequence ID: NP_001328697.1 Length: 151
>hypothetical protein AT4G10843 [Arabidopsis thaliana]
Sequence ID: ANM66826.1 Length: 151
Range 1: 1 to 80

Score:82.4 bits(202), Expect:5e-19,
Method:Compositional matrix adjust.,
Identities:54/112(48%), Positives:64/112(57%), Gaps:32/112(28%)

Query  399  MKLRRKRWSWLIV*VFFA*kkkkkTV**SVISLL*AKIIKEMSSASLHPFFLVLIFLRNF  578
            MKLRRKRWSWL++                       K +K    +SL        F+  F
Sbjct  1    MKLRRKRWSWLVIE---------------------TKSLK----SSLPERRRRRRFI--F  33

Query  579  LLCED*L*YGDDIMDVSLFTF*TEETLIEDKSLSFCTCNGERWRRLFSFYKV  734
             LC+        ++DV+LFTF TEETLIEDKSLSFCTCNGERWRRLFSFYK+
Sbjct  34   ALCD-----SSFLLDVNLFTFWTEETLIEDKSLSFCTCNGERWRRLFSFYKL  80


>hypothetical protein AT4G10843 [Arabidopsis thaliana]
Sequence ID: NP_001328695.1 Length: 159
>hypothetical protein AT4G10843 [Arabidopsis thaliana]
Sequence ID: ANM66823.1 Length: 159
Range 1: 1 to 80

Score:82.4 bits(202), Expect:6e-19,
Method:Compositional matrix adjust.,
Identities:54/112(48%), Positives:63/112(56%), Gaps:32/112(28%)

Query  399  MKLRRKRWSWLIV*VFFA*kkkkkTV**SVISLL*AKIIKEMSSASLHPFFLVLIFLRNF  578
            MKLRRKRWSWL++                  SL           +SL        F+  F
Sbjct  1    MKLRRKRWSWLVI---------------ETKSL----------KSSLPERRRRRRFI--F  33

Query  579  LLCED*L*YGDDIMDVSLFTF*TEETLIEDKSLSFCTCNGERWRRLFSFYKV  734
             LC+        ++DV+LFTF TEETLIEDKSLSFCTCNGERWRRLFSFYK+
Sbjct  34   ALCD-----SSFLLDVNLFTFWTEETLIEDKSLSFCTCNGERWRRLFSFYKL  80


>unnamed protein product [Arabidopsis thaliana]
Sequence ID: CAA0405987.1 Length: 177
Range 1: 55 to 93

Score:75.5 bits(184), Expect:4e-16,
Method:Compositional matrix adjust.,
Identities:36/39(92%), Positives:37/39(94%), Gaps:0/39(0%)

Query  615  IMDVSLFTF*TEETLIEDKSLSFCTCNGERWRRLFSFYK  731
            I DV+LFTF TEETLIEDKSLSFCTCNGERWRRLFSFYK
Sbjct  55   IRDVNLFTFWTEETLIEDKSLSFCTCNGERWRRLFSFYK  93


Range 2: 1 to 34

Score:71.6 bits(174), Expect:9e-15,
Method:Compositional matrix adjust.,
Identities:34/34(100%), Positives:34/34(100%), Gaps:0/34(0%)

Query  332  MVVSSPTVEDGGRYGGRWRAVRDEVTEEALVVAD  433
            MVVSSPTVEDGGRYGGRWRAVRDEVTEEALVVAD
Sbjct  1    MVVSSPTVEDGGRYGGRWRAVRDEVTEEALVVAD  34


>unnamed protein product [Arabidopsis thaliana]
Sequence ID: VYS62247.1 Length: 165
Range 1: 69 to 107

Score:73.9 bits(180), Expect:1e-15,
Method:Compositional matrix adjust.,
Identities:34/39(87%), Positives:37/39(94%), Gaps:0/39(0%)

Query  615  IMDVSLFTF*TEETLIEDKSLSFCTCNGERWRRLFSFYK  731
            ++D +LFTF TEETLIEDKSLSFCTCNGERWRRLFSFYK
Sbjct  69   LLDANLFTFWTEETLIEDKSLSFCTCNGERWRRLFSFYK  107


Range 2: 1 to 33

Score:62.8 bits(151), Expect:1e-11,
Method:Compositional matrix adjust.,
Identities:30/33(91%), Positives:31/33(93%), Gaps:0/33(0%)

Query  332  MVVSSPTVEDGGRYGGRWRAVRDEVTEEALVVA  430
            MVVSSPTVEDGG YGGRWR VRDEVTEEA+VVA
Sbjct  1    MVVSSPTVEDGGTYGGRWRDVRDEVTEEAVVVA  33


>unnamed protein product [Arabidopsis thaliana]
Sequence ID: CAA0394571.1 Length: 269
Range 1: 68 to 107

Score:75.9 bits(185), Expect:1e-15,
Method:Compositional matrix adjust.,
Identities:35/40(88%), Positives:39/40(97%), Gaps:0/40(0%)

Query  615  IMDVSLFTF*TEETLIEDKSLSFCTCNGERWRRLFSFYKV  734
            ++DV+LFTF TEETLIEDKSLSFCTCNGERWRRLFSFYK+
Sbjct  68   LLDVNLFTFWTEETLIEDKSLSFCTCNGERWRRLFSFYKL  107


Range 2: 1 to 33

Score:68.9 bits(167), Expect:4e-13,
Method:Compositional matrix adjust.,
Identities:32/33(97%), Positives:33/33(100%), Gaps:0/33(0%)

Query  332  MVVSSPTVEDGGRYGGRWRAVRDEVTEEALVVA  430
            MVVSSPTVEDGGRYGGRWRAVRDEVTEEA+VVA
Sbjct  1    MVVSSPTVEDGGRYGGRWRAVRDEVTEEAVVVA  33


>hypothetical protein AT5G37793, partial [Arabidopsis thaliana]
Sequence ID: NP_001331870.1 Length: 100
>hypothetical protein AT5G37793, partial [Arabidopsis thaliana]
Sequence ID: ANM70244.1 Length: 100
Range 1: 1 to 49

Score:71.6 bits(174), Expect:2e-15,
Method:Compositional matrix adjust.,
Identities:48/49(98%), Positives:48/49(97%), Gaps:0/49(0%)

Query  332  MVVSSPTVEDGGRYGGRWRAVRDEVTEEALVVADrlsllrlkeeeeDGL  478
            MVVSSPTVEDGGRYGGRWRAVRDEVTEEALVVADRLSLLRLKEEEEDG
Sbjct  1    MVVSSPTVEDGGRYGGRWRAVRDEVTEEALVVADRLSLLRLKEEEEDGF  49


Range 2: 51 to 85

Score:67.8 bits(164), Expect:5e-14,
Method:Compositional matrix adjust.,
Identities:32/35(91%), Positives:32/35(91%), Gaps:0/35(0%)

Query  627  SLFTF*TEETLIEDKSLSFCTCNGERWRRLFSFYK  731
            SLF   TEETLIEDKSLSFCTCNGERWRRLFSFYK
Sbjct  51   SLFVIRTEETLIEDKSLSFCTCNGERWRRLFSFYK  85


>unnamed protein product [Arabidopsis thaliana]
Sequence ID: VYS68533.1 Length: 180
Range 1: 1 to 49

Score:72.0 bits(175), Expect:7e-15,
Method:Compositional matrix adjust.,
Identities:48/49(98%), Positives:48/49(97%), Gaps:0/49(0%)

Query  332  MVVSSPTVEDGGRYGGRWRAVRDEVTEEALVVADrlsllrlkeeeeDGL  478
            MVVSSPTVEDGGRYGGRWRAVRDEVTEEALVVADRLSLLRLKEEEEDG
Sbjct  1    MVVSSPTVEDGGRYGGRWRAVRDEVTEEALVVADRLSLLRLKEEEEDGF  49


Range 2: 51 to 85

Score:67.4 bits(163), Expect:4e-13,
Method:Compositional matrix adjust.,
Identities:32/35(91%), Positives:32/35(91%), Gaps:0/35(0%)

Query  627  SLFTF*TEETLIEDKSLSFCTCNGERWRRLFSFYK  731
            SLF   TEETLIEDKSLSFCTCNGERWRRLFSFYK
Sbjct  51   SLFVIRTEETLIEDKSLSFCTCNGERWRRLFSFYK  85


>unnamed protein product [Arabidopsis thaliana]
Sequence ID: VYS67928.1 Length: 74
Range 1: 1 to 33

Score:66.2 bits(160), Expect:8e-14,
Method:Compositional matrix adjust.,
Identities:31/33(94%), Positives:33/33(100%), Gaps:0/33(0%)

Query  332  MVVSSPTVEDGGRYGGRWRAVRDEVTEEALVVA  430
            MVV+SPTVEDGGRYGGRWRAVRDEVTEEA+VVA
Sbjct  1    MVVASPTVEDGGRYGGRWRAVRDEVTEEAVVVA  33


>nucleoporin (DUF3414) [Arabidopsis thaliana]
Sequence ID: NP_195587.5 Length: 1965
>nucleoporin (DUF3414) [Arabidopsis thaliana]
Sequence ID: AEE86971.1 Length: 1965 >unnamed protein product [Arabidopsis thaliana]
Sequence ID: VYS65328.1 Length: 1965
Range 1: 195 to 242

Score:55.1 bits(131), Expect:5e-08,
Method:Compositional matrix adjust.,
Identities:27/48(56%), Positives:32/48(66%), Gaps:10/48(20%)

Query  618  MDVSLFTF*TEETLIEDKSL----------SFCTCNGERWRRLFSFYK  731
            MDV+LFT   EETLIED  +          S+C+CNGERWR+L SFYK
Sbjct  195  MDVNLFTLWAEETLIEDNLILDILFLIYNESYCSCNGERWRKLCSFYK  242


>hypothetical protein AXX17_AT4G44080 [Arabidopsis thaliana]
Sequence ID: OAO97877.1 Length: 1968
Range 1: 195 to 242

Score:55.1 bits(131), Expect:5e-08,
Method:Compositional matrix adjust.,
Identities:27/48(56%), Positives:32/48(66%), Gaps:10/48(20%)

Query  618  MDVSLFTF*TEETLIEDKSL----------SFCTCNGERWRRLFSFYK  731
            MDV+LFT   EETLIED  +          S+C+CNGERWR+L SFYK
Sbjct  195  MDVNLFTLWAEETLIEDNLILDILFLIYNESYCSCNGERWRKLCSFYK  242


>unnamed protein product [Arabidopsis thaliana]
Sequence ID: CAA0397936.1 Length: 1968
Range 1: 195 to 242

Score:55.1 bits(131), Expect:5e-08,
Method:Compositional matrix adjust.,
Identities:27/48(56%), Positives:32/48(66%), Gaps:10/48(20%)

Query  618  MDVSLFTF*TEETLIEDKSL----------SFCTCNGERWRRLFSFYK  731
            MDV+LFT   EETLIED  +          S+C+CNGERWR+L SFYK
Sbjct  195  MDVNLFTLWAEETLIEDNLILDILFLIYNESYCSCNGERWRKLCSFYK  242


Query #245: XLOC_027118 Query ID: lcl|Query_30736 Length: 533

Sequences producing significant alignments:
                                                                  Max    Total Query   E   Per.
Description                                                       Score  Score cover Value Ident  Accession
hypothetical protein [Arabidopsis thaliana]                       79.0   79.0  33%   9e-17 61.67  AAD25573.1
hypothetical protein [Arabidopsis thaliana]                       77.0   77.0  33%   4e-16 60.00  AAD25576.1
putative protein [Arabidopsis thaliana]                           77.0   77.0  33%   4e-16 60.00  CAB51198.1
putative protein [Arabidopsis thaliana]                           76.6   76.6  33%   4e-16 60.00  CAB88532.1
T2L5.5 gene product [Arabidopsis thaliana]                        76.6   76.6  33%   5e-16 60.00  AAC62793.1
hypothetical protein [Arabidopsis thaliana]                       75.5   75.5  33%   1e-15 58.33  AAD25852.1
hypothetical protein [Arabidopsis thaliana]                       74.7   74.7  33%   2e-15 59.32  AAC97213.1
T24H24.3 gene product [Arabidopsis thaliana]                      74.7   74.7  33%   2e-15 58.33  AAC28201.1
hypothetical protein [Arabidopsis thaliana]                       73.2   73.2  33%   8e-15 58.33  AAD26891.2
Several near identical copies of this gene are pr> [Arabidopsi... 72.8   72.8  33%   9e-15 56.67  AAF31286.1


Alignments:
>hypothetical protein [Arabidopsis thaliana]
Sequence ID: AAD25573.1 Length: 775
Range 1: 706 to 765

Score:79.0 bits(193), Expect:9e-17,
Method:Compositional matrix adjust.,
Identities:37/60(62%), Positives:46/60(76%), Gaps:0/60(0%)

Query  197  HDIDVTPFSIRVLTSFLQTAKIKESDIYMMKIMECYTM*ASNSNLEENIIGNARKKSAVD  376
             D+DV+PFSI+VLT+F QT + +ES IYM+K MECY+M  S+SNLE NII N R K A D
Sbjct  706  EDMDVSPFSIKVLTTFPQTPRNEESGIYMLKFMECYSMYTSHSNLEGNIIQNVRNKLAAD  765


>hypothetical protein [Arabidopsis thaliana]
Sequence ID: AAD25576.1 Length: 792
Range 1: 723 to 782

Score:77.0 bits(188), Expect:4e-16,
Method:Compositional matrix adjust.,
Identities:36/60(60%), Positives:45/60(75%), Gaps:0/60(0%)

Query  197  HDIDVTPFSIRVLTSFLQTAKIKESDIYMMKIMECYTM*ASNSNLEENIIGNARKKSAVD  376
             D+DV+PFSI+VLT+F Q  + +ES IYM+K MECY+M  S+SNLE NII N R K A D
Sbjct  723  EDMDVSPFSIKVLTTFPQAPRNEESGIYMLKFMECYSMYTSHSNLEGNIIQNVRNKLAAD  782


>putative protein [Arabidopsis thaliana]
Sequence ID: CAB51198.1 Length: 820
Range 1: 751 to 810

Score:77.0 bits(188), Expect:4e-16,
Method:Compositional matrix adjust.,
Identities:36/60(60%), Positives:45/60(75%), Gaps:0/60(0%)

Query  197  HDIDVTPFSIRVLTSFLQTAKIKESDIYMMKIMECYTM*ASNSNLEENIIGNARKKSAVD  376
             D+DV+PFSI+VLT+F Q  + +ES IYM+K MECY+M  S+SNLE NII N R K A D
Sbjct  751  EDMDVSPFSIKVLTTFPQAPRNEESGIYMLKFMECYSMYTSHSNLEGNIIQNVRNKLAAD  810


>putative protein [Arabidopsis thaliana]
Sequence ID: CAB88532.1 Length: 742
Range 1: 673 to 732

Score:76.6 bits(187), Expect:4e-16,
Method:Compositional matrix adjust.,
Identities:36/60(60%), Positives:45/60(75%), Gaps:0/60(0%)

Query  197  HDIDVTPFSIRVLTSFLQTAKIKESDIYMMKIMECYTM*ASNSNLEENIIGNARKKSAVD  376
             D+DV+PFSI+VLT+F Q  + +ES IYM+K MECY+M  S+SNLE NII N R K A D
Sbjct  673  EDMDVSPFSIKVLTTFPQAPRNEESGIYMLKFMECYSMYTSHSNLEGNIIQNVRNKLAAD  732


>T2L5.5 gene product [Arabidopsis thaliana]
Sequence ID: AAC62793.1 Length: 696
Range 1: 627 to 686

Score:76.6 bits(187), Expect:5e-16,
Method:Compositional matrix adjust.,
Identities:36/60(60%), Positives:45/60(75%), Gaps:0/60(0%)

Query  197  HDIDVTPFSIRVLTSFLQTAKIKESDIYMMKIMECYTM*ASNSNLEENIIGNARKKSAVD  376
             D+DV+PFSI+VLT+F Q  + +ES IYM+K MECY+M  S+SNLE NII N R K A D
Sbjct  627  EDMDVSPFSIKVLTTFPQAPRNEESGIYMLKFMECYSMYTSHSNLEGNIIQNVRNKLAAD  686


>hypothetical protein [Arabidopsis thaliana]
Sequence ID: AAD25852.1 Length: 789
Range 1: 720 to 779

Score:75.5 bits(184), Expect:1e-15,
Method:Compositional matrix adjust.,
Identities:35/60(58%), Positives:44/60(73%), Gaps:0/60(0%)

Query  197  HDIDVTPFSIRVLTSFLQTAKIKESDIYMMKIMECYTM*ASNSNLEENIIGNARKKSAVD  376
             D+DV+PFSI+VLT+F Q  + +ES IYM+K MECY+M  S+SN E NII N R K A D
Sbjct  720  EDMDVSPFSIKVLTTFPQAPRNEESGIYMLKFMECYSMYTSHSNFEGNIIQNVRNKLAAD  779


>hypothetical protein [Arabidopsis thaliana]
Sequence ID: AAC97213.1 Length: 684
Range 1: 616 to 674

Score:74.7 bits(182), Expect:2e-15,
Method:Compositional matrix adjust.,
Identities:35/59(59%), Positives:44/59(74%), Gaps:0/59(0%)

Query  200  DIDVTPFSIRVLTSFLQTAKIKESDIYMMKIMECYTM*ASNSNLEENIIGNARKKSAVD  376
            D+DV+PFSI+VLT+F    + +ES IYM+K MECY+M  S+SNLE NII N R K A D
Sbjct  616  DMDVSPFSIKVLTTFPHAPRNEESGIYMLKFMECYSMYTSHSNLEGNIIQNVRNKLAAD  674


>T24H24.3 gene product [Arabidopsis thaliana]
Sequence ID: AAC28201.1 Length: 836
>hypothetical protein [Arabidopsis thaliana]
Sequence ID: CAB77869.1 Length: 836
Range 1: 767 to 826

Score:74.7 bits(182), Expect:2e-15,
Method:Compositional matrix adjust.,
Identities:35/60(58%), Positives:45/60(75%), Gaps:0/60(0%)

Query  197  HDIDVTPFSIRVLTSFLQTAKIKESDIYMMKIMECYTM*ASNSNLEENIIGNARKKSAVD  376
             ++DV+PFSI+VLT+F Q  + +ES IYM+K MECY+M  S+SNLE NII N R K A D
Sbjct  767  ENMDVSPFSIKVLTTFPQAPRNEESGIYMLKFMECYSMYTSHSNLEGNIIQNVRNKLAAD  826


>hypothetical protein [Arabidopsis thaliana]
Sequence ID: AAD26891.2 Length: 795
Range 1: 726 to 785

Score:73.2 bits(178), Expect:8e-15,
Method:Compositional matrix adjust.,
Identities:35/60(58%), Positives:44/60(73%), Gaps:0/60(0%)

Query  197  HDIDVTPFSIRVLTSFLQTAKIKESDIYMMKIMECYTM*ASNSNLEENIIGNARKKSAVD  376
             D+DV+PFSI+VLT+F Q  + +ES IYM+K MECY+M  S+SNLE NII N   K A D
Sbjct  726  EDMDVSPFSIKVLTTFPQAPRNEESRIYMLKFMECYSMYTSHSNLEGNIIQNVMNKLAAD  785


>Several near identical copies of this gene are pr&gt; [Arabidopsis thaliana]
Sequence ID: AAF31286.1 Length: 607
Range 1: 538 to 597

Score:72.8 bits(177), Expect:9e-15,
Method:Compositional matrix adjust.,
Identities:34/60(57%), Positives:43/60(71%), Gaps:0/60(0%)

Query  197  HDIDVTPFSIRVLTSFLQTAKIKESDIYMMKIMECYTM*ASNSNLEENIIGNARKKSAVD  376
             D+DV+PFSI+VLT+F Q  + +ES IYM+K MECY+M  S+SN E NII N R K   D
Sbjct  538  EDMDVSPFSIKVLTTFPQAPRNEESGIYMLKFMECYSMYTSHSNFEGNIIQNVRNKLVAD  597


Query #246: XLOC_027124 Query ID: lcl|Query_30737 Length: 416

No significant similarity found.


Query #247: XLOC_027179 Query ID: lcl|Query_30738 Length: 2029

Sequences producing significant alignments:
                                                                  Max    Total Query   E   Per.
Description                                                       Score  Score cover Value Ident  Accession
unnamed protein product [Arabidopsis thaliana]                    135    176   12%   5e-37 94.20  VYS68646.1
unnamed protein product [Arabidopsis thaliana]                    137    242   18%   1e-32 92.65  CAA0406188.1
hypothetical protein AT5G38747 [Arabidopsis thaliana]             122    239   25%   8e-32 91.94  NP_001332557.1
hypothetical protein AXX17_AT5G36120 [Arabidopsis thaliana]       87.4   87.4  6%    1e-20 95.24  OAO92383.1
unnamed protein product [Arabidopsis thaliana]                    89.7   89.7  9%    4e-19 68.75  BAD44378.1
H[+]-ATPase 2 [Arabidopsis thaliana]                              89.7   89.7  10%   9e-18 64.63  NP_194748.1
Chain A, Atpase 2, Plasma Membrane-type [Arabidopsis thaliana]    89.4   89.4  10%   1e-17 64.63  5KSD_A
H[+]-ATPase 2 [Arabidopsis thaliana]                              89.4   89.4  10%   1e-17 64.63  NP_001190870.1
unnamed protein product [Arabidopsis thaliana]                    88.6   88.6  9%    1e-17 68.75  CAA0371827.1
unnamed protein product [Arabidopsis thaliana]                    88.6   88.6  9%    1e-17 68.75  VYS53520.1
haloacid dehalogenase-like hydrolase family protein [Arabidops... 88.6   88.6  9%    1e-17 68.75  NP_850072.1
AT4G30190 [Arabidopsis thaliana]                                  86.7   86.7  10%   8e-17 63.41  BAH56991.1
H[+]-ATPase 1 [Arabidopsis thaliana]                              84.0   84.0  10%   7e-16 60.98  NP_001324051.1
plasma membrane proton pump H+ ATPase [Arabidopsis thaliana]      83.6   83.6  10%   7e-16 60.98  AAA32813.1
H[+]-ATPase 1 [Arabidopsis thaliana]                              83.6   83.6  10%   7e-16 60.98  NP_179486.1
H[+]-ATPase 3 [Arabidopsis thaliana]                              82.0   82.0  10%   2e-15 59.26  NP_001331203.1
H[+]-ATPase 3 [Arabidopsis thaliana]                              81.6   81.6  10%   3e-15 59.26  NP_001190559.1
AT5g57350/MJB24_16 [Arabidopsis thaliana]                         81.6   81.6  10%   3e-15 59.26  AAL09726.1
H[+]-ATPase 11 [Arabidopsis thaliana]                             79.7   79.7  10%   1e-14 58.02  NP_201073.1
unnamed protein product [Arabidopsis thaliana]                    79.0   79.0  12%   2e-14 51.02  CAA0384301.1
unnamed protein product [Arabidopsis thaliana]                    79.0   79.0  12%   2e-14 51.02  VYS59168.1
HA8 [Arabidopsis thaliana]                                        79.0   79.0  12%   2e-14 51.02  OAP07011.1
H[+]-ATPase 8 [Arabidopsis thaliana]                              78.6   78.6  12%   3e-14 51.02  NP_189850.1
hypothetical protein AXX17_AT4G13200 [Arabidopsis thaliana]       78.6   78.6  10%   3e-14 58.02  OAO97880.1
Theoretical protein with similarity to Swiss-Prot Accession...    77.8   77.8  12%   5e-14 50.00  AAA98916.1
aha9, 5' partial; 1-2403 [Arabidopsis thaliana]                   77.4   77.4  12%   5e-14 50.00  AAF27113.1
H[+]-ATPase 9 [Arabidopsis thaliana]                              77.4   77.4  12%   6e-14 50.00  NP_001319430.1
unnamed protein product [Arabidopsis thaliana]                    77.0   77.0  12%   9e-14 50.00  CAA0344746.1
H[+]-ATPase 9 [Arabidopsis thaliana]                              77.0   77.0  12%   9e-14 50.00  NP_001320419.1
H[+]-ATPase 9 [Arabidopsis thaliana]                              77.0   77.0  12%   1e-13 50.00  NP_001320420.1
H[+]-ATPase 9 [Arabidopsis thaliana]                              77.0   77.0  12%   1e-13 50.00  NP_178181.1
plasma membrane H+-ATPase [Arabidopsis thaliana]                  76.3   76.3  10%   1e-13 56.79  NP_001325337.1
unnamed protein product [Arabidopsis thaliana]                    76.3   76.3  12%   1e-13 49.48  CAA0393944.1
RecName: Full=ATPase 5, plasma membrane-type; AltName:...         75.9   75.9  10%   2e-13 56.79  Q9SJB3.3
plasma membrane H+-ATPase [Arabidopsis thaliana]                  75.9   75.9  10%   2e-13 56.79  NP_001318282.1
unnamed protein product [Arabidopsis thaliana]                    75.9   75.9  10%   2e-13 56.79  CAA0370878.1
HA5 [Arabidopsis thaliana]                                        75.9   75.9  10%   2e-13 56.79  OAP08952.1
plasma membrane H+-ATPase [Arabidopsis thaliana]                  75.9   75.9  10%   2e-13 56.79  NP_001325336.1
HA6 [Arabidopsis thaliana]                                        75.5   75.5  12%   3e-13 49.48  OAO89452.1
H[+]-ATPase 6 [Arabidopsis thaliana]                              75.5   75.5  12%   3e-13 49.48  NP_178762.1
unnamed protein product [Arabidopsis thaliana]                    75.5   75.5  12%   3e-13 49.48  VYS52201.1
H+-transporting ATPase-like protein [Arabidopsis thaliana]        73.6   73.6  13%   9e-13 46.60  CAB41144.1
H[+]-ATPase 4 [Arabidopsis thaliana]                              73.2   73.2  10%   1e-12 54.32  NP_001325714.1
Contains weak similarity to plasma membrane H+ ATPase from...     64.7   64.7  9%    2e-11 56.06  AAF78278.1
unnamed protein product [Arabidopsis thaliana]                    68.6   68.6  10%   3e-11 51.28  CAA0213975.1
autoinhibited H[+]-ATPase isoform 10 [Arabidopsis thaliana]       68.6   68.6  10%   3e-11 51.28  NP_173169.2
H[+]-ATPase 7 [Arabidopsis thaliana]                              64.7   64.7  8%    6e-10 53.85  NP_001327362.1
H[+]-ATPase 7 [Arabidopsis thaliana]                              64.7   64.7  8%    6e-10 53.85  NP_001190141.1
hypothetical protein AT1G44941 [Arabidopsis thaliana]             57.0   57.0  4%    8e-10 83.33  NP_001117439.1
unnamed protein product [Arabidopsis thaliana]                    57.0   57.0  4%    8e-10 83.33  CAA0274387.1
unnamed protein product [Arabidopsis thaliana]                    54.3   54.3  4%    6e-09 80.00  VYS48295.1
hypothetical protein AT1G44940 [Arabidopsis thaliana]             53.1   53.1  5%    3e-08 74.29  NP_001154402.1
unnamed protein product [Arabidopsis thaliana]                    53.1   53.1  5%    3e-08 74.29  VYS48292.1
hypothetical protein AT1G44940 [Arabidopsis thaliana]             53.1   53.1  5%    3e-08 74.29  NP_001154403.1


Alignments:
>unnamed protein product [Arabidopsis thaliana]
Sequence ID: VYS68646.1 Length: 107
Range 1: 10 to 78

Score:135 bits(340), Expect:5e-37,
Method:Compositional matrix adjust.,
Identities:65/69(94%), Positives:67/69(97%), Gaps:0/69(0%)

Query  795   SRQWKEVSKGNLNPDVCIETCLYSVEHGILLIVDRCLTLFDNPLVDSLHTTYNELKLQKE  974
             SRQWKEVSKGNLNPDVCIETCLYSVEHGILLIVDRCLTLFDNPLVDSLHTTYNELKLQKE
Sbjct  10    SRQWKEVSKGNLNPDVCIETCLYSVEHGILLIVDRCLTLFDNPLVDSLHTTYNELKLQKE  69

Query  975   CKNQT*KWV  1001
             CKNQ+ K +
Sbjct  70    CKNQSKKPI  78


Range 2: 90 to 107

Score:41.2 bits(95), Expect:5e-37,
Method:Compositional matrix adjust.,
Identities:18/18(100%), Positives:18/18(100%), Gaps:0/18(0%)

Query  989   LKMGVFGILNQCHKGDKR  1042
             LKMGVFGILNQCHKGDKR
Sbjct  90    LKMGVFGILNQCHKGDKR  107


>unnamed protein product [Arabidopsis thaliana]
Sequence ID: CAA0406188.1 Length: 1836
Range 1: 92 to 159

Score:137 bits(345), Expect:1e-32,
Method:Composition-based stats.,
Identities:63/68(93%), Positives:64/68(94%), Gaps:0/68(0%)

Query  792  FSRQWKEVSKGNLNPDVCIETCLYSVEHGILLIVDRCLTLFDNPLVDSLHTTYNELKLQK  971
              RQWKEVSKGNLNPDVC ETCLYSVEHGILLIVDRCLTLFDNPLVDSLHTTYNELKLQK
Sbjct  92   LGRQWKEVSKGNLNPDVCRETCLYSVEHGILLIVDRCLTLFDNPLVDSLHTTYNELKLQK  151

Query  972  ECKNQT*K  995
            ECKNQ+ K
Sbjct  152  ECKNQSKK  159


Range 2: 59 to 98

Score:74.7 bits(182), Expect:1e-15,
Method:Composition-based stats.,
Identities:36/41(88%), Positives:38/41(92%), Gaps:1/41(2%)

Query  420  RALTLGVNIKMITGQITSCKLGMGINMYPSSAFLGTHKW*E  542
            RALTLGVNIKMITGQ+TSCKLGMGINMYPSSAFLG  +W E
Sbjct  59   RALTLGVNIKMITGQVTSCKLGMGINMYPSSAFLG-RQWKE  98


Range 3: 49 to 61

Score:30.0 bits(66), Expect:1e-15,
Method:Composition-based stats.,
Identities:11/13(85%), Positives:12/13(92%), Gaps:0/13(0%)

Query  317  GGARENEGKPRCM  355
            GGARENEGKPR +
Sbjct  49   GGARENEGKPRAL  61


>hypothetical protein AT5G38747, partial [Arabidopsis thaliana]
Sequence ID: NP_001332557.1 Length: 153
>hypothetical protein AT5G38747, partial [Arabidopsis thaliana]
Sequence ID: ANM70994.1 Length: 153
Range 1: 72 to 133

Score:122 bits(306), Expect:8e-32,
Method:Compositional matrix adjust.,
Identities:57/62(92%), Positives:58/62(93%), Gaps:0/62(0%)

Query  792  FSRQWKEVSKGNLNPDVCIETCLYSVEHGILLIVDRCLTLFDNPLVDSLHTTYNELKLQK  971
              RQWKEVSKGNLNPDVCIETCLYSVEHGILLIVDRCLTLFDNPLVDSLHTTYNELKL+
Sbjct  72   LGRQWKEVSKGNLNPDVCIETCLYSVEHGILLIVDRCLTLFDNPLVDSLHTTYNELKLKN  131

Query  972  EC  977
             C
Sbjct  132  GC  133


Range 2: 110 to 153

Score:59.3 bits(142), Expect:1e-09,
Method:Compositional matrix adjust.,
Identities:29/44(66%), Positives:32/44(72%), Gaps:0/44(0%)

Query  931   TLFTQLTMSSSYRKNARTRLKNGCVRHSKSMSQRRQAVRTTKPH  1062
             TLF    + S +      +LKNGCVRHSKSMSQRRQAVRTTKPH
Sbjct  110   TLFDNPLVDSLHTTYNELKLKNGCVRHSKSMSQRRQAVRTTKPH  153


Range 3: 17 to 73

Score:58.2 bits(139), Expect:3e-09,
Method:Compositional matrix adjust.,
Identities:41/83(49%), Positives:45/83(54%), Gaps:26/83(31%)

Query  102  KIPRTLIKTRVQFSIEVARVDLWRCFPGF*SNVKLNIENMQRQKGSSGNQGGSLPLTYID  281
            +IPRTLIKTRVQFSIEVARVDLWR               ++R+ G           T ID
Sbjct  17   QIPRTLIKTRVQFSIEVARVDLWR---------------LERESGR---------FTSID  52

Query  282  SRRQ*SRCSPD*VVLEKTKESLG  350
              RQ        VVLEKTKESLG
Sbjct  53   LHRQQKAVVK--VVLEKTKESLG  73


>hypothetical protein AXX17_AT5G36120 [Arabidopsis thaliana]
Sequence ID: OAO92383.1 Length: 52
Range 1: 11 to 52

Score:87.4 bits(215), Expect:1e-20,
Method:Compositional matrix adjust.,
Identities:40/42(95%), Positives:41/42(97%), Gaps:0/42(0%)

Query  1026  TKETSGKNHQTPLTKMVSWKIWHVRDVENILVGASNDFLQRN  1151
             T+ETSGKNHQTP TKMVSWKIWHVRDVENILVGASNDFLQRN
Sbjct  11    TRETSGKNHQTPSTKMVSWKIWHVRDVENILVGASNDFLQRN  52


>unnamed protein product [Arabidopsis thaliana]
Sequence ID: BAD44378.1 Length: 266
Range 1: 70 to 133

Score:89.7 bits(221), Expect:4e-19,
Method:Compositional matrix adjust.,
Identities:44/64(69%), Positives:50/64(78%), Gaps:3/64(4%)

Query  786  LVFSRQWKEVSKGNLNPDVCIETCLYSVEHGILLIV---DRCLTLFDNPLVDSLHTTYNE  956
            LV+ RQ KEV +GNL+ DVC ETCLYS+EH I LI    DRCLTLFD+PLVDSLHT YNE
Sbjct  70   LVYGRQGKEVYRGNLDRDVCRETCLYSLEHRIPLIAFSQDRCLTLFDHPLVDSLHTIYNE  129

Query  957  LKLQ  968
             K +
Sbjct  130  PKAE  133


>H[+]-ATPase 2 [Arabidopsis thaliana]
Sequence ID: NP_194748.1 Length: 948
>RecName: Full=ATPase 2, plasma membrane-type; AltName: Full=Proton pump 2 [Arabidopsis thaliana]
Sequence ID: P19456.2 Length: 948 >H+-ATPase [Arabidopsis thaliana]
Sequence ID: AAA32751.1 Length: 948 >putative H+-transporting ATPase [Arabidopsis thaliana]
Sequence ID: AAK59580.1 Length: 948 >putative H+-transporting ATPase type 2 [Arabidopsis thaliana]
Sequence ID: AAN31920.1 Length: 948 >putative H+-transporting ATPase [Arabidopsis thaliana]
Sequence ID: AAN71968.1 Length: 948 >H[+]-ATPase 2 [Arabidopsis thaliana]
Sequence ID: AEE85731.1 Length: 948
Range 1: 464 to 542

Score:89.7 bits(221), Expect:9e-18,
Method:Compositional matrix adjust.,
Identities:53/82(65%), Positives:57/82(69%), Gaps:13/82(15%)

Query  318  VVLEKTKESLGAWCNIW*FVGLLP----PRHESTETIGRALTLGVNIKMIT------GQI  467
            VV EKTKES GA    W FVGLLP    PRH+S ETI RAL LGVN+KMIT      G+
Sbjct  464  VVPEKTKESPGA---PWEFVGLLPLFDPPRHDSAETIRRALNLGVNVKMITGDQLAIGKE  520

Query  468  TSCKLGMGINMYPSSAFLGTHK  533
            T  +LGMG NMYPSSA LGTHK
Sbjct  521  TGRRLGMGTNMYPSSALLGTHK  542


>Chain A, Atpase 2, Plasma Membrane-type [Arabidopsis thaliana]
Sequence ID: 5KSD_A Length: 833
>Chain B, Atpase 2, Plasma Membrane-type [Arabidopsis thaliana]
Sequence ID: 5KSD_B Length: 833
Range 1: 453 to 531

Score:89.4 bits(220), Expect:1e-17,
Method:Compositional matrix adjust.,
Identities:53/82(65%), Positives:57/82(69%), Gaps:13/82(15%)

Query  318  VVLEKTKESLGAWCNIW*FVGLLP----PRHESTETIGRALTLGVNIKMIT------GQI  467
            VV EKTKES GA    W FVGLLP    PRH+S ETI RAL LGVN+KMIT      G+
Sbjct  453  VVPEKTKESPGA---PWEFVGLLPLFDPPRHDSAETIRRALNLGVNVKMITGDQLAIGKE  509

Query  468  TSCKLGMGINMYPSSAFLGTHK  533
            T  +LGMG NMYPSSA LGTHK
Sbjct  510  TGRRLGMGTNMYPSSALLGTHK  531


>H[+]-ATPase 2 [Arabidopsis thaliana]
Sequence ID: NP_001190870.1 Length: 981
>H[+]-ATPase 2 [Arabidopsis thaliana]
Sequence ID: AEE85732.1 Length: 981
Range 1: 464 to 542

Score:89.4 bits(220), Expect:1e-17,
Method:Compositional matrix adjust.,
Identities:53/82(65%), Positives:57/82(69%), Gaps:13/82(15%)

Query  318  VVLEKTKESLGAWCNIW*FVGLLP----PRHESTETIGRALTLGVNIKMIT------GQI  467
            VV EKTKES GA    W FVGLLP    PRH+S ETI RAL LGVN+KMIT      G+
Sbjct  464  VVPEKTKESPGA---PWEFVGLLPLFDPPRHDSAETIRRALNLGVNVKMITGDQLAIGKE  520

Query  468  TSCKLGMGINMYPSSAFLGTHK  533
            T  +LGMG NMYPSSA LGTHK
Sbjct  521  TGRRLGMGTNMYPSSALLGTHK  542


>unnamed protein product [Arabidopsis thaliana]
Sequence ID: CAA0371827.1 Length: 584
Range 1: 388 to 451

Score:88.6 bits(218), Expect:1e-17,
Method:Compositional matrix adjust.,
Identities:44/64(69%), Positives:50/64(78%), Gaps:3/64(4%)

Query  786  LVFSRQWKEVSKGNLNPDVCIETCLYSVEHGILLIV---DRCLTLFDNPLVDSLHTTYNE  956
            LV+ RQ KEV +GNL+ DVC ETCLYS+EH I LI    DRCLTLFD+PLVDSLHT YNE
Sbjct  388  LVYGRQGKEVYRGNLDRDVCRETCLYSLEHRIPLIAFSQDRCLTLFDHPLVDSLHTIYNE  447

Query  957  LKLQ  968
             K +
Sbjct  448  PKAE  451


>unnamed protein product [Arabidopsis thaliana]
Sequence ID: VYS53520.1 Length: 584
Range 1: 388 to 451

Score:88.6 bits(218), Expect:1e-17,
Method:Compositional matrix adjust.,
Identities:44/64(69%), Positives:50/64(78%), Gaps:3/64(4%)

Query  786  LVFSRQWKEVSKGNLNPDVCIETCLYSVEHGILLIV---DRCLTLFDNPLVDSLHTTYNE  956
            LV+ RQ KEV +GNL+ DVC ETCLYS+EH I LI    DRCLTLFD+PLVDSLHT YNE
Sbjct  388  LVYGRQGKEVYRGNLDRDVCRETCLYSLEHRIPLIAFSQDRCLTLFDHPLVDSLHTIYNE  447

Query  957  LKLQ  968
             K +
Sbjct  448  PKAE  451


>haloacid dehalogenase-like hydrolase family protein [Arabidopsis thaliana]
Sequence ID: NP_850072.1 Length: 584
>RecName: Full=Endoribonuclease YBEY, chloroplastic; Flags: Precursor [Arabidopsis thaliana]
Sequence ID: Q8L5Z4.1 Length: 584 >unknown protein [Arabidopsis thaliana]
Sequence ID: AAM20655.1 Length: 584 >At2g25870 [Arabidopsis thaliana]
Sequence ID: ABO38782.1 Length: 584 >haloacid dehalogenase-like hydrolase family protein [Arabidopsis thaliana]
Sequence ID: AEC07765.1 Length: 584 >hypothetical protein AXX17_AT2G21710 [Arabidopsis thaliana]
Sequence ID: OAP07883.1 Length: 584 >unknown protein [Arabidopsis thaliana]
Sequence ID: BAC43355.1 Length: 584
Range 1: 388 to 451

Score:88.6 bits(218), Expect:1e-17,
Method:Compositional matrix adjust.,
Identities:44/64(69%), Positives:50/64(78%), Gaps:3/64(4%)

Query  786  LVFSRQWKEVSKGNLNPDVCIETCLYSVEHGILLIV---DRCLTLFDNPLVDSLHTTYNE  956
            LV+ RQ KEV +GNL+ DVC ETCLYS+EH I LI    DRCLTLFD+PLVDSLHT YNE
Sbjct  388  LVYGRQGKEVYRGNLDRDVCRETCLYSLEHRIPLIAFSQDRCLTLFDHPLVDSLHTIYNE  447

Query  957  LKLQ  968
             K +
Sbjct  448  PKAE  451


>AT4G30190 [Arabidopsis thaliana]
Sequence ID: BAH56991.1 Length: 816
Range 1: 340 to 418

Score:86.7 bits(213), Expect:8e-17,
Method:Compositional matrix adjust.,
Identities:52/82(63%), Positives:56/82(68%), Gaps:13/82(15%)

Query  318  VVLEKTKESLGAWCNIW*FVGLLP----PRHESTETIGRALTLGVNIKMIT------GQI  467
            VV EKTKES GA    W FVGLLP    PRH+S  TI RAL LGVN+KMIT      G+
Sbjct  340  VVPEKTKESPGA---PWEFVGLLPLFDPPRHDSAGTIRRALNLGVNVKMITGDQLAIGKE  396

Query  468  TSCKLGMGINMYPSSAFLGTHK  533
            T  +LGMG NMYPSSA LGTHK
Sbjct  397  TGRRLGMGTNMYPSSALLGTHK  418


>H[+]-ATPase 1 [Arabidopsis thaliana]
Sequence ID: NP_001324051.1 Length: 885
>H[+]-ATPase 1 [Arabidopsis thaliana]
Sequence ID: NP_001324052.1 Length: 885 >H[+]-ATPase 1 [Arabidopsis thaliana]
Sequence ID: ANM61856.1 Length: 885 >H[+]-ATPase 1 [Arabidopsis thaliana]
Sequence ID: ANM61857.1 Length: 885
Range 1: 400 to 478

Score:84.0 bits(206), Expect:7e-16,
Method:Compositional matrix adjust.,
Identities:50/82(61%), Positives:55/82(67%), Gaps:13/82(15%)

Query  318  VVLEKTKESLGAWCNIW*FVGLLP----PRHESTETIGRALTLGVNIKMIT------GQI  467
            VV EKTKES G     W FVGLLP    PRH+S ETI RAL LGVN+KMIT      G+
Sbjct  400  VVPEKTKESPGG---PWEFVGLLPLFDPPRHDSAETIRRALNLGVNVKMITGDQLAIGKE  456

Query  468  TSCKLGMGINMYPSSAFLGTHK  533
            T  +LGMG NMYPS+A LGT K
Sbjct  457  TGRRLGMGTNMYPSAALLGTDK  478


>plasma membrane proton pump H+ ATPase [Arabidopsis thaliana]
Sequence ID: AAA32813.1 Length: 949
Range 1: 464 to 542

Score:83.6 bits(205), Expect:7e-16,
Method:Compositional matrix adjust.,
Identities:50/82(61%), Positives:55/82(67%), Gaps:13/82(15%)

Query  318  VVLEKTKESLGAWCNIW*FVGLLP----PRHESTETIGRALTLGVNIKMIT------GQI  467
            VV EKTKES G     W FVGLLP    PRH+S ETI RAL LGVN+KMIT      G+
Sbjct  464  VVPEKTKESPGG---PWEFVGLLPLFDPPRHDSAETIRRALNLGVNVKMITGDQLAIGKE  520

Query  468  TSCKLGMGINMYPSSAFLGTHK  533
            T  +LGMG NMYPS+A LGT K
Sbjct  521  TGRRLGMGTNMYPSAALLGTDK  542


>H[+]-ATPase 1 [Arabidopsis thaliana]
Sequence ID: NP_179486.1 Length: 949
>RecName: Full=ATPase 1, plasma membrane-type; AltName: Full=Proton pump 1 [Arabidopsis thaliana]
Sequence ID: P20649.3 Length: 949 >plasma membrane proton ATPase (PMA) [Arabidopsis thaliana]
Sequence ID: AAC09030.1 Length: 949 >putative plasma membrane proton ATPase (PMA) [Arabidopsis thaliana]
Sequence ID: AAP40498.1 Length: 949 >H[+]-ATPase 1 [Arabidopsis thaliana]
Sequence ID: AEC06832.1 Length: 949 >PMA [Arabidopsis thaliana]
Sequence ID: OAP08431.1 Length: 949 >unnamed protein product [Arabidopsis thaliana]
Sequence ID: CAA0365797.1 Length: 949
Range 1: 464 to 542

Score:83.6 bits(205), Expect:7e-16,
Method:Compositional matrix adjust.,
Identities:50/82(61%), Positives:55/82(67%), Gaps:13/82(15%)

Query  318  VVLEKTKESLGAWCNIW*FVGLLP----PRHESTETIGRALTLGVNIKMIT------GQI  467
            VV EKTKES G     W FVGLLP    PRH+S ETI RAL LGVN+KMIT      G+
Sbjct  464  VVPEKTKESPGG---PWEFVGLLPLFDPPRHDSAETIRRALNLGVNVKMITGDQLAIGKE  520

Query  468  TSCKLGMGINMYPSSAFLGTHK  533
            T  +LGMG NMYPS+A LGT K
Sbjct  521  TGRRLGMGTNMYPSAALLGTDK  542


>H[+]-ATPase 3 [Arabidopsis thaliana]
Sequence ID: NP_001331203.1 Length: 687
>H[+]-ATPase 3 [Arabidopsis thaliana]
Sequence ID: ANM69533.1 Length: 687
Range 1: 204 to 281

Score:82.0 bits(201), Expect:2e-15,
Method:Compositional matrix adjust.,
Identities:48/81(59%), Positives:55/81(67%), Gaps:13/81(16%)

Query  321  VLEKTKESLGAWCNIW*FVGLLP----PRHESTETIGRALTLGVNIKMITG------QIT  470
            V EKTKES G+    W FVG+LP    PRH+S ETI RAL LGVN+KMITG      + T
Sbjct  204  VPEKTKESSGS---PWEFVGVLPLFDPPRHDSAETIRRALDLGVNVKMITGDQLAIAKET  260

Query  471  SCKLGMGINMYPSSAFLGTHK  533
              +LGMG NMYPSS+ LG HK
Sbjct  261  GRRLGMGSNMYPSSSLLGKHK  281


>H[+]-ATPase 3 [Arabidopsis thaliana]
Sequence ID: NP_001190559.1 Length: 949
>H[+]-ATPase 3 [Arabidopsis thaliana]
Sequence ID: NP_001331204.1 Length: 949 >H[+]-ATPase 3 [Arabidopsis thaliana]
Sequence ID: NP_200545.1 Length: 949 >RecName: Full=ATPase 3, plasma membrane-type; AltName: Full=Proton pump 3 [Arabidopsis thaliana]
Sequence ID: P20431.2 Length: 949 >ATPase [Arabidopsis thaliana]
Sequence ID: AAA32750.1 Length: 949 >putative plasma membrane proton pump ATPase 3 [Arabidopsis thaliana]
Sequence ID: AAL59975.1 Length: 949 >H[+]-ATPase 3 [Arabidopsis thaliana]
Sequence ID: AED96889.1 Length: 949 >H[+]-ATPase 3 [Arabidopsis thaliana]
Sequence ID: AED96890.1 Length: 949 >H[+]-ATPase 3 [Arabidopsis thaliana]
Sequence ID: ANM69534.1 Length: 949
Range 1: 466 to 543

Score:81.6 bits(200), Expect:3e-15,
Method:Compositional matrix adjust.,
Identities:48/81(59%), Positives:55/81(67%), Gaps:13/81(16%)

Query  321  VLEKTKESLGAWCNIW*FVGLLP----PRHESTETIGRALTLGVNIKMITG------QIT  470
            V EKTKES G+    W FVG+LP    PRH+S ETI RAL LGVN+KMITG      + T
Sbjct  466  VPEKTKESSGS---PWEFVGVLPLFDPPRHDSAETIRRALDLGVNVKMITGDQLAIAKET  522

Query  471  SCKLGMGINMYPSSAFLGTHK  533
              +LGMG NMYPSS+ LG HK
Sbjct  523  GRRLGMGSNMYPSSSLLGKHK  543


>AT5g57350/MJB24_16 [Arabidopsis thaliana]
Sequence ID: AAL09726.1 Length: 949
Range 1: 466 to 543

Score:81.6 bits(200), Expect:3e-15,
Method:Compositional matrix adjust.,
Identities:48/81(59%), Positives:55/81(67%), Gaps:13/81(16%)

Query  321  VLEKTKESLGAWCNIW*FVGLLP----PRHESTETIGRALTLGVNIKMITG------QIT  470
            V EKTKES G+    W FVG+LP    PRH+S ETI RAL LGVN+KMITG      + T
Sbjct  466  VPEKTKESSGS---PWEFVGVLPLFDPPRHDSAETIRRALDLGVNVKMITGDQLAIAKET  522

Query  471  SCKLGMGINMYPSSAFLGTHK  533
              +LGMG NMYPSS+ LG HK
Sbjct  523  GRRLGMGSNMYPSSSLLGKHK  543


>H[+]-ATPase 11 [Arabidopsis thaliana]
Sequence ID: NP_201073.1 Length: 956
>RecName: Full=ATPase 11, plasma membrane-type; AltName: Full=Proton pump 11 [Arabidopsis thaliana]
Sequence ID: Q9LV11.1 Length: 956 >AT5g62670/MRG21_9 [Arabidopsis thaliana]
Sequence ID: AAM78085.1 Length: 956 >At5g62670/MRG21_9 [Arabidopsis thaliana]
Sequence ID: AAR23718.1 Length: 956 >H[+]-ATPase 11 [Arabidopsis thaliana]
Sequence ID: AED97641.1 Length: 956 >HA11 [Arabidopsis thaliana]
Sequence ID: OAO93098.1 Length: 956 >unnamed protein product [Arabidopsis thaliana]
Sequence ID: CAA0411564.1 Length: 956
Range 1: 469 to 546

Score:79.7 bits(195), Expect:1e-14,
Method:Compositional matrix adjust.,
Identities:47/81(58%), Positives:53/81(65%), Gaps:13/81(16%)

Query  321  VLEKTKESLGAWCNIW*FVGLLP----PRHESTETIGRALTLGVNIKMIT------GQIT  470
            V E TKES G     W F+GL+P    PRH+S ETI RAL LGVN+KMIT      G+ T
Sbjct  469  VPEGTKESAGG---PWQFMGLMPLFDPPRHDSAETIRRALNLGVNVKMITGDQLAIGKET  525

Query  471  SCKLGMGINMYPSSAFLGTHK  533
              +LGMG NMYPSSA LG HK
Sbjct  526  GRRLGMGTNMYPSSALLGQHK  546


>unnamed protein product [Arabidopsis thaliana]
Sequence ID: CAA0384301.1 Length: 948
Range 1: 468 to 562

Score:79.0 bits(193), Expect:2e-14,
Method:Compositional matrix adjust.,
Identities:50/98(51%), Positives:59/98(60%), Gaps:14/98(14%)

Query  321  VLEKTKESLGAWCNIW*FVGLLP----PRHESTETIGRALTLGVNIKMITGQI------T  470
            V EKTKES G+    W FVGLLP    PRH+S ETI RAL LGVN+KMITG        T
Sbjct  468  VPEKTKESDGS---PWEFVGLLPLFDPPRHDSAETIRRALELGVNVKMITGDQLAIGIET  524

Query  471  SCKLGMGINMYPSSAFLGTHKW*EPTN-PMDEYVSQVS  581
              +LGMG NMYPS++ LG  K       P+DE + +
Sbjct  525  GRRLGMGTNMYPSTSLLGNSKDESLVGIPIDELIEKAD  562


>unnamed protein product [Arabidopsis thaliana]
Sequence ID: VYS59168.1 Length: 948
Range 1: 468 to 562

Score:79.0 bits(193), Expect:2e-14,
Method:Compositional matrix adjust.,
Identities:50/98(51%), Positives:59/98(60%), Gaps:14/98(14%)

Query  321  VLEKTKESLGAWCNIW*FVGLLP----PRHESTETIGRALTLGVNIKMITGQI------T  470
            V EKTKES G+    W FVGLLP    PRH+S ETI RAL LGVN+KMITG        T
Sbjct  468  VPEKTKESDGS---PWEFVGLLPLFDPPRHDSAETIRRALELGVNVKMITGDQLAIGIET  524

Query  471  SCKLGMGINMYPSSAFLGTHKW*EPTN-PMDEYVSQVS  581
              +LGMG NMYPS++ LG  K       P+DE + +
Sbjct  525  GRRLGMGTNMYPSTSLLGNSKDESLVGIPIDELIEKAD  562


>HA8 [Arabidopsis thaliana]
Sequence ID: OAP07011.1 Length: 948
Range 1: 468 to 562

Score:79.0 bits(193), Expect:2e-14,
Method:Compositional matrix adjust.,
Identities:50/98(51%), Positives:59/98(60%), Gaps:14/98(14%)

Query  321  VLEKTKESLGAWCNIW*FVGLLP----PRHESTETIGRALTLGVNIKMITGQI------T  470
            V EKTKES G+    W FVGLLP    PRH+S ETI RAL LGVN+KMITG        T
Sbjct  468  VPEKTKESDGS---PWEFVGLLPLFDPPRHDSAETIRRALELGVNVKMITGDQLAIGIET  524

Query  471  SCKLGMGINMYPSSAFLGTHKW*EPTN-PMDEYVSQVS  581
              +LGMG NMYPS++ LG  K       P+DE + +
Sbjct  525  GRRLGMGTNMYPSTSLLGNSKDESLVGIPIDELIEKAD  562


>H[+]-ATPase 8 [Arabidopsis thaliana]
Sequence ID: NP_189850.1 Length: 948
>RecName: Full=ATPase 8, plasma membrane-type; AltName: Full=Proton pump 8 [Arabidopsis thaliana]
Sequence ID: Q9M2A0.1 Length: 948 >H[+]-ATPase 8 [Arabidopsis thaliana]
Sequence ID: AEE77743.1 Length: 948 >plasma membrane H+-ATPase-like protein [Arabidopsis thaliana]
Sequence ID: CAB86447.1 Length: 948
Range 1: 468 to 562

Score:78.6 bits(192), Expect:3e-14,
Method:Compositional matrix adjust.,
Identities:50/98(51%), Positives:59/98(60%), Gaps:14/98(14%)

Query  321  VLEKTKESLGAWCNIW*FVGLLP----PRHESTETIGRALTLGVNIKMITGQI------T  470
            V EKTKES G+    W FVGLLP    PRH+S ETI RAL LGVN+KMITG        T
Sbjct  468  VPEKTKESDGS---PWEFVGLLPLFDPPRHDSAETIRRALELGVNVKMITGDQLAIGIET  524

Query  471  SCKLGMGINMYPSSAFLGTHKW*EPTN-PMDEYVSQVS  581
              +LGMG NMYPS++ LG  K       P+DE + +
Sbjct  525  GRRLGMGTNMYPSTSLLGNSKDESLVGIPIDELIEKAD  562


>hypothetical protein AXX17_AT4G13200 [Arabidopsis thaliana]
Sequence ID: OAO97880.1 Length: 905
Range 1: 468 to 545

Score:78.6 bits(192), Expect:3e-14,
Method:Compositional matrix adjust.,
Identities:47/81(58%), Positives:53/81(65%), Gaps:13/81(16%)

Query  321  VLEKTKESLGAWCNIW*FVGLLP----PRHESTETIGRALTLGVNIKMITG------QIT  470
            V EKTKES G     W FVGLLP    PRH+S ETI RAL LGVN+KMITG      + T
Sbjct  468  VPEKTKESSGG---PWEFVGLLPLFDPPRHDSAETIKRALDLGVNMKMITGDNLTIAKET  524

Query  471  SCKLGMGINMYPSSAFLGTHK  533
              +LGMG NMYP ++ LG HK
Sbjct  525  GRRLGMGTNMYPITSLLGNHK  545


>Theoretical protein with similarity to Swiss-Prot Accession Number P19456 plasma membrane ATPase 2 (proton pump) [Arabidopsis thaliana]
Sequence ID: AAA98916.1 Length: 859
Range 1: 401 to 495

Score:77.8 bits(190), Expect:5e-14,
Method:Compositional matrix adjust.,
Identities:49/98(50%), Positives:58/98(59%), Gaps:14/98(14%)

Query  321  VLEKTKESLGAWCNIW*FVGLLP----PRHESTETIGRALTLGVNIKMIT------GQIT  470
            V EK K S G     W F+GLLP    PRH+S ETI RAL LGVN+KMIT      G+ T
Sbjct  401  VSEKDKNSXG---EPWQFLGLLPLFDPPRHDSAETIRRALDLGVNVKMITGDQLAIGKET  457

Query  471  SCKLGMGINMYPSSAFLGTHKW*EPTN-PMDEYVSQVS  581
              +LGMG NMYPSSA LG  K     + P+DE + +
Sbjct  458  GRRLGMGTNMYPSSALLGQDKDESIASLPVDELIEKAD  495


>aha9, 5' partial; 1-2403, partial [Arabidopsis thaliana]
Sequence ID: AAF27113.1 Length: 612
Range 1: 128 to 222

Score:77.4 bits(189), Expect:5e-14,
Method:Compositional matrix adjust.,
Identities:49/98(50%), Positives:58/98(59%), Gaps:14/98(14%)

Query  321  VLEKTKESLGAWCNIW*FVGLLP----PRHESTETIGRALTLGVNIKMIT------GQIT  470
            V EK K S G     W F+GLLP    PRH+S ETI RAL LGVN+KMIT      G+ T
Sbjct  128  VSEKDKNSPG---EPWQFLGLLPLFDPPRHDSAETIRRALDLGVNVKMITGDQLAIGKET  184

Query  471  SCKLGMGINMYPSSAFLGTHKW*EPTN-PMDEYVSQVS  581
              +LGMG NMYPSSA LG  K     + P+DE + +
Sbjct  185  GRRLGMGTNMYPSSALLGQDKDESIASLPVDELIEKAD  222


>H[+]-ATPase 9 [Arabidopsis thaliana]
Sequence ID: NP_001319430.1 Length: 825
>H[+]-ATPase 9 [Arabidopsis thaliana]
Sequence ID: AEE36434.2 Length: 825
Range 1: 341 to 435

Score:77.4 bits(189), Expect:6e-14,
Method:Compositional matrix adjust.,
Identities:49/98(50%), Positives:58/98(59%), Gaps:14/98(14%)

Query  321  VLEKTKESLGAWCNIW*FVGLLP----PRHESTETIGRALTLGVNIKMIT------GQIT  470
            V EK K S G     W F+GLLP    PRH+S ETI RAL LGVN+KMIT      G+ T
Sbjct  341  VSEKDKNSPG---EPWQFLGLLPLFDPPRHDSAETIRRALDLGVNVKMITGDQLAIGKET  397

Query  471  SCKLGMGINMYPSSAFLGTHKW*EPTN-PMDEYVSQVS  581
              +LGMG NMYPSSA LG  K     + P+DE + +
Sbjct  398  GRRLGMGTNMYPSSALLGQDKDESIASLPVDELIEKAD  435


>unnamed protein product [Arabidopsis thaliana]
Sequence ID: CAA0344746.1 Length: 949
>unnamed protein product [Arabidopsis thaliana]
Sequence ID: VYS51648.1 Length: 949
Range 1: 470 to 564

Score:77.0 bits(188), Expect:9e-14,
Method:Compositional matrix adjust.,
Identities:49/98(50%), Positives:58/98(59%), Gaps:14/98(14%)

Query  321  VLEKTKESLGAWCNIW*FVGLLP----PRHESTETIGRALTLGVNIKMIT------GQIT  470
            V EK K S G     W F+GLLP    PRH+S ETI RAL LGVN+KMIT      G+ T
Sbjct  470  VSEKDKNSPG---EPWQFLGLLPLFDPPRHDSAETIRRALDLGVNVKMITGDQLAIGKET  526

Query  471  SCKLGMGINMYPSSAFLGTHKW*EPTN-PMDEYVSQVS  581
              +LGMG NMYPSSA LG  K     + P+DE + +
Sbjct  527  GRRLGMGTNMYPSSALLGQDKDESIASLPVDELIEKAD  564


>H[+]-ATPase 9 [Arabidopsis thaliana]
Sequence ID: NP_001320419.1 Length: 928
>H[+]-ATPase 9 [Arabidopsis thaliana]
Sequence ID: ANM57945.1 Length: 928
Range 1: 470 to 564

Score:77.0 bits(188), Expect:9e-14,
Method:Compositional matrix adjust.,
Identities:49/98(50%), Positives:58/98(59%), Gaps:14/98(14%)

Query  321  VLEKTKESLGAWCNIW*FVGLLP----PRHESTETIGRALTLGVNIKMIT------GQIT  470
            V EK K S G     W F+GLLP    PRH+S ETI RAL LGVN+KMIT      G+ T
Sbjct  470  VSEKDKNSPG---EPWQFLGLLPLFDPPRHDSAETIRRALDLGVNVKMITGDQLAIGKET  526

Query  471  SCKLGMGINMYPSSAFLGTHKW*EPTN-PMDEYVSQVS  581
              +LGMG NMYPSSA LG  K     + P+DE + +
Sbjct  527  GRRLGMGTNMYPSSALLGQDKDESIASLPVDELIEKAD  564


>H[+]-ATPase 9 [Arabidopsis thaliana]
Sequence ID: NP_001320420.1 Length: 927
>H[+]-ATPase 9 [Arabidopsis thaliana]
Sequence ID: ANM57946.1 Length: 927
Range 1: 470 to 564

Score:77.0 bits(188), Expect:1e-13,
Method:Compositional matrix adjust.,
Identities:49/98(50%), Positives:58/98(59%), Gaps:14/98(14%)

Query  321  VLEKTKESLGAWCNIW*FVGLLP----PRHESTETIGRALTLGVNIKMIT------GQIT  470
            V EK K S G     W F+GLLP    PRH+S ETI RAL LGVN+KMIT      G+ T
Sbjct  470  VSEKDKNSPG---EPWQFLGLLPLFDPPRHDSAETIRRALDLGVNVKMITGDQLAIGKET  526

Query  471  SCKLGMGINMYPSSAFLGTHKW*EPTN-PMDEYVSQVS  581
              +LGMG NMYPSSA LG  K     + P+DE + +
Sbjct  527  GRRLGMGTNMYPSSALLGQDKDESIASLPVDELIEKAD  564


>H[+]-ATPase 9 [Arabidopsis thaliana]
Sequence ID: NP_178181.1 Length: 954
>RecName: Full=ATPase 9, plasma membrane-type; AltName: Full=Proton pump 9 [Arabidopsis thaliana]
Sequence ID: Q42556.2 Length: 954 >Identical to gb|X73676 aha9 (ATAHA9) ATPase gene from Arabidopsis thaliana [Arabidopsis thaliana]
Sequence ID: AAF14653.1 Length: 954 >H[+]-ATPase 9 [Arabidopsis thaliana]
Sequence ID: AEE36433.1 Length: 954 >HA9 [Arabidopsis thaliana]
Sequence ID: OAP14596.1 Length: 954
Range 1: 470 to 564

Score:77.0 bits(188), Expect:1e-13,
Method:Compositional matrix adjust.,
Identities:49/98(50%), Positives:58/98(59%), Gaps:14/98(14%)

Query  321  VLEKTKESLGAWCNIW*FVGLLP----PRHESTETIGRALTLGVNIKMIT------GQIT  470
            V EK K S G     W F+GLLP    PRH+S ETI RAL LGVN+KMIT      G+ T
Sbjct  470  VSEKDKNSPG---EPWQFLGLLPLFDPPRHDSAETIRRALDLGVNVKMITGDQLAIGKET  526

Query  471  SCKLGMGINMYPSSAFLGTHKW*EPTN-PMDEYVSQVS  581
              +LGMG NMYPSSA LG  K     + P+DE + +
Sbjct  527  GRRLGMGTNMYPSSALLGQDKDESIASLPVDELIEKAD  564


>plasma membrane H+-ATPase [Arabidopsis thaliana]
Sequence ID: NP_001325337.1 Length: 688
>plasma membrane H+-ATPase [Arabidopsis thaliana]
Sequence ID: ANM63234.1 Length: 688
Range 1: 204 to 281

Score:76.3 bits(186), Expect:1e-13,
Method:Compositional matrix adjust.,
Identities:46/81(57%), Positives:52/81(64%), Gaps:13/81(16%)

Query  321  VLEKTKESLGAWCNIW*FVGLLP----PRHESTETIGRALTLGVNIKMIT------GQIT  470
            VLEK K++ G     W  VGLLP    PRH+S ETI RAL LGVN+KMIT      G+ T
Sbjct  204  VLEKKKDAPGG---PWQLVGLLPLFDPPRHDSAETIRRALNLGVNVKMITGDQLAIGKET  260

Query  471  SCKLGMGINMYPSSAFLGTHK  533
              +LGMG NMYPSSA LG  K
Sbjct  261  GRRLGMGTNMYPSSALLGQVK  281


>unnamed protein product [Arabidopsis thaliana]
Sequence ID: CAA0393944.1 Length: 949
Range 1: 468 to 561

Score:76.3 bits(186), Expect:1e-13,
Method:Compositional matrix adjust.,
Identities:48/97(49%), Positives:57/97(58%), Gaps:13/97(13%)

Query  321  VLEKTKESLGAWCNIW*FVGLLP----PRHESTETIGRALTLGVNIKMIT------GQIT  470
            V EK KES+G     W FVGLLP    PRH+S ETI RAL LGVN+KMIT      G+ T
Sbjct  468  VPEKDKESVG---TPWEFVGLLPLFDPPRHDSAETIRRALDLGVNVKMITGDQLAIGKET  524

Query  471  SCKLGMGINMYPSSAFLGTHKW*EPTNPMDEYVSQVS  581
              +LGMG NMYPSS+ L          P+DE + +
Sbjct  525  GRRLGMGTNMYPSSSLLENKDDTTGGVPVDELIEKAD  561


>RecName: Full=ATPase 5, plasma membrane-type; AltName: Full=Proton pump 5 [Arabidopsis thaliana]
Sequence ID: Q9SJB3.3 Length: 949
Range 1: 465 to 542

Score:75.9 bits(185), Expect:2e-13,
Method:Compositional matrix adjust.,
Identities:46/81(57%), Positives:52/81(64%), Gaps:13/81(16%)

Query  321  VLEKTKESLGAWCNIW*FVGLLP----PRHESTETIGRALTLGVNIKMIT------GQIT  470
            VLEK K++ G     W  VGLLP    PRH+S ETI RAL LGVN+KMIT      G+ T
Sbjct  465  VLEKKKDAPGG---PWQLVGLLPLFDPPRHDSAETIRRALNLGVNVKMITGDQLAIGKET  521

Query  471  SCKLGMGINMYPSSAFLGTHK  533
              +LGMG NMYPSSA LG  K
Sbjct  522  GRRLGMGTNMYPSSALLGQVK  542


>plasma membrane H+-ATPase [Arabidopsis thaliana]
Sequence ID: NP_001318282.1 Length: 931
>plasma membrane H+-ATPase [Arabidopsis thaliana]
Sequence ID: NP_180028.1 Length: 931 >putative plasma membrane proton ATPase [Arabidopsis thaliana]
Sequence ID: AAD23893.1 Length: 931 >plasma membrane H+-ATPase [Arabidopsis thaliana]
Sequence ID: AEC07588.1 Length: 931 >plasma membrane H+-ATPase [Arabidopsis thaliana]
Sequence ID: ANM63233.1 Length: 931
Range 1: 447 to 524

Score:75.9 bits(185), Expect:2e-13,
Method:Compositional matrix adjust.,
Identities:46/81(57%), Positives:52/81(64%), Gaps:13/81(16%)

Query  321  VLEKTKESLGAWCNIW*FVGLLP----PRHESTETIGRALTLGVNIKMIT------GQIT  470
            VLEK K++ G     W  VGLLP    PRH+S ETI RAL LGVN+KMIT      G+ T
Sbjct  447  VLEKKKDAPGG---PWQLVGLLPLFDPPRHDSAETIRRALNLGVNVKMITGDQLAIGKET  503

Query  471  SCKLGMGINMYPSSAFLGTHK  533
              +LGMG NMYPSSA LG  K
Sbjct  504  GRRLGMGTNMYPSSALLGQVK  524


>unnamed protein product [Arabidopsis thaliana]
Sequence ID: CAA0370878.1 Length: 956
>unnamed protein product [Arabidopsis thaliana]
Sequence ID: VYS53374.1 Length: 956
Range 1: 472 to 549

Score:75.9 bits(185), Expect:2e-13,
Method:Compositional matrix adjust.,
Identities:46/81(57%), Positives:52/81(64%), Gaps:13/81(16%)

Query  321  VLEKTKESLGAWCNIW*FVGLLP----PRHESTETIGRALTLGVNIKMIT------GQIT  470
            VLEK K++ G     W  VGLLP    PRH+S ETI RAL LGVN+KMIT      G+ T
Sbjct  472  VLEKKKDAPGG---PWQLVGLLPLFDPPRHDSAETIRRALNLGVNVKMITGDQLAIGKET  528

Query  471  SCKLGMGINMYPSSAFLGTHK  533
              +LGMG NMYPSSA LG  K
Sbjct  529  GRRLGMGTNMYPSSALLGQVK  549


>HA5 [Arabidopsis thaliana]
Sequence ID: OAP08952.1 Length: 956
Range 1: 472 to 549

Score:75.9 bits(185), Expect:2e-13,
Method:Compositional matrix adjust.,
Identities:46/81(57%), Positives:52/81(64%), Gaps:13/81(16%)

Query  321  VLEKTKESLGAWCNIW*FVGLLP----PRHESTETIGRALTLGVNIKMIT------GQIT  470
            VLEK K++ G     W  VGLLP    PRH+S ETI RAL LGVN+KMIT      G+ T
Sbjct  472  VLEKKKDAPGG---PWQLVGLLPLFDPPRHDSAETIRRALNLGVNVKMITGDQLAIGKET  528

Query  471  SCKLGMGINMYPSSAFLGTHK  533
              +LGMG NMYPSSA LG  K
Sbjct  529  GRRLGMGTNMYPSSALLGQVK  549


>plasma membrane H+-ATPase [Arabidopsis thaliana]
Sequence ID: NP_001325336.1 Length: 994
>plasma membrane H+-ATPase [Arabidopsis thaliana]
Sequence ID: ANM63232.1 Length: 994
Range 1: 510 to 587

Score:75.9 bits(185), Expect:2e-13,
Method:Compositional matrix adjust.,
Identities:46/81(57%), Positives:52/81(64%), Gaps:13/81(16%)

Query  321  VLEKTKESLGAWCNIW*FVGLLP----PRHESTETIGRALTLGVNIKMIT------GQIT  470
            VLEK K++ G     W  VGLLP    PRH+S ETI RAL LGVN+KMIT      G+ T
Sbjct  510  VLEKKKDAPGG---PWQLVGLLPLFDPPRHDSAETIRRALNLGVNVKMITGDQLAIGKET  566

Query  471  SCKLGMGINMYPSSAFLGTHK  533
              +LGMG NMYPSSA LG  K
Sbjct  567  GRRLGMGTNMYPSSALLGQVK  587


>HA6 [Arabidopsis thaliana]
Sequence ID: OAO89452.1 Length: 949
Range 1: 468 to 561

Score:75.5 bits(184), Expect:3e-13,
Method:Compositional matrix adjust.,
Identities:48/97(49%), Positives:56/97(57%), Gaps:13/97(13%)

Query  321  VLEKTKESLGAWCNIW*FVGLLP----PRHESTETIGRALTLGVNIKMIT------GQIT  470
            V EK KES G     W FVGLLP    PRH+S ETI RAL LGVN+KMIT      G+ T
Sbjct  468  VPEKDKESAG---TPWEFVGLLPLFDPPRHDSAETIRRALDLGVNVKMITGDQLAIGKET  524

Query  471  SCKLGMGINMYPSSAFLGTHKW*EPTNPMDEYVSQVS  581
              +LGMG NMYPSS+ L          P+DE + +
Sbjct  525  GRRLGMGTNMYPSSSLLENKDDTTGGVPVDELIEKAD  561


>H[+]-ATPase 6 [Arabidopsis thaliana]
Sequence ID: NP_178762.1 Length: 949
>RecName: Full=ATPase 6, plasma membrane-type; AltName: Full=Proton pump 6 [Arabidopsis thaliana]
Sequence ID: Q9SH76.1 Length: 949 >putative plasma membrane proton ATPase [Arabidopsis thaliana]
Sequence ID: AAD32758.1 Length: 949 >H[+]-ATPase 6 [Arabidopsis thaliana]
Sequence ID: AEC06056.1 Length: 949
Range 1: 468 to 561

Score:75.5 bits(184), Expect:3e-13,
Method:Compositional matrix adjust.,
Identities:48/97(49%), Positives:56/97(57%), Gaps:13/97(13%)

Query  321  VLEKTKESLGAWCNIW*FVGLLP----PRHESTETIGRALTLGVNIKMIT------GQIT  470
            V EK KES G     W FVGLLP    PRH+S ETI RAL LGVN+KMIT      G+ T
Sbjct  468  VPEKDKESAG---TPWEFVGLLPLFDPPRHDSAETIRRALDLGVNVKMITGDQLAIGKET  524

Query  471  SCKLGMGINMYPSSAFLGTHKW*EPTNPMDEYVSQVS  581
              +LGMG NMYPSS+ L          P+DE + +
Sbjct  525  GRRLGMGTNMYPSSSLLENKDDTTGGVPVDELIEKAD  561


>unnamed protein product [Arabidopsis thaliana]
Sequence ID: VYS52201.1 Length: 949
Range 1: 468 to 561

Score:75.5 bits(184), Expect:3e-13,
Method:Compositional matrix adjust.,
Identities:48/97(49%), Positives:56/97(57%), Gaps:13/97(13%)

Query  321  VLEKTKESLGAWCNIW*FVGLLP----PRHESTETIGRALTLGVNIKMIT------GQIT  470
            V EK KES G     W FVGLLP    PRH+S ETI RAL LGVN+KMIT      G+ T
Sbjct  468  VPEKDKESAG---TPWEFVGLLPLFDPPRHDSAETIRRALDLGVNVKMITGDQLAIGKET  524

Query  471  SCKLGMGINMYPSSAFLGTHKW*EPTNPMDEYVSQVS  581
              +LGMG NMYPSS+ L          P+DE + +
Sbjct  525  GRRLGMGTNMYPSSSLLENKDDTTGGVPVDELIEKAD  561


>H+-transporting ATPase-like protein [Arabidopsis thaliana]
Sequence ID: CAB41144.1 Length: 960
Range 1: 473 to 572

Score:73.6 bits(179), Expect:9e-13,
Method:Compositional matrix adjust.,
Identities:48/103(47%), Positives:59/103(57%), Gaps:14/103(13%)

Query  321  VLEKTKESLGAWCNIW*FVGLLP----PRHESTETIGRALTLGVNIKMIT------GQIT  470
            V E  K+S G     W FVGL+P    PRH+S ETI RAL LGV++KMIT      G+ T
Sbjct  473  VPEGRKDSAGG---PWQFVGLMPLFDPPRHDSAETIRRALNLGVSVKMITGDQLAIGKET  529

Query  471  SCKLGMGINMYPSSAFLGTHKW*EPTN-PMDEYVSQVSMVVSF  596
              +LGMG NMYPSSA LG +K       P+DE + +      F
Sbjct  530  GRRLGMGTNMYPSSALLGQNKDESIVALPVDELIEKADGFAGF  572


>H[+]-ATPase 4 [Arabidopsis thaliana]
Sequence ID: NP_001325714.1 Length: 960
>H[+]-ATPase 4 [Arabidopsis thaliana]
Sequence ID: NP_190378.2 Length: 960 >RecName: Full=ATPase 4, plasma membrane-type; AltName: Full=Proton pump 4 [Arabidopsis thaliana]
Sequence ID: Q9SU58.2 Length: 960 >H[+]-ATPase 4 [Arabidopsis thaliana]
Sequence ID: AEE78350.1 Length: 960 >H[+]-ATPase 4 [Arabidopsis thaliana]
Sequence ID: ANM63639.1 Length: 960 >HA4 [Arabidopsis thaliana]
Sequence ID: OAP05139.1 Length: 960 >unnamed protein product [Arabidopsis thaliana]
Sequence ID: CAA0384933.1 Length: 960 >unnamed protein product [Arabidopsis thaliana]
Sequence ID: VYS59692.1 Length: 960
Range 1: 473 to 550

Score:73.2 bits(178), Expect:1e-12,
Method:Compositional matrix adjust.,
Identities:44/81(54%), Positives:52/81(64%), Gaps:13/81(16%)

Query  321  VLEKTKESLGAWCNIW*FVGLLP----PRHESTETIGRALTLGVNIKMIT------GQIT  470
            V E  K+S G     W FVGL+P    PRH+S ETI RAL LGV++KMIT      G+ T
Sbjct  473  VPEGRKDSAGG---PWQFVGLMPLFDPPRHDSAETIRRALNLGVSVKMITGDQLAIGKET  529

Query  471  SCKLGMGINMYPSSAFLGTHK  533
              +LGMG NMYPSSA LG +K
Sbjct  530  GRRLGMGTNMYPSSALLGQNK  550


>Contains weak similarity to plasma membrane H+ ATPase from Phaseolus vulgaris gb|X94936. EST gb|T22206 comes from this gene [Arabidopsis thaliana]
Sequence ID: AAF78278.1 Length: 169
Range 1: 104 to 164

Score:64.7 bits(156), Expect:2e-11,
Method:Compositional matrix adjust.,
Identities:37/66(56%), Positives:43/66(65%), Gaps:6/66(9%)

Query  479  TWNGNKHVSIFG-FSWYAQVVGTNKPHG*ICLSSLNGCKFCHVRDMENIPGRESSEFLQR  655
            TWNGNKHVS+F  F  YAQ VGTNKPHG ICLS     K  H+RD+ENI    S++  +
Sbjct  104  TWNGNKHVSLFRLFKEYAQAVGTNKPHGRICLS-----KIWHIRDVENILVGASNDSSKE  158

Query  656  TDQIRK  673
             D I K
Sbjct  159  NDHIGK  164


>unnamed protein product [Arabidopsis thaliana]
Sequence ID: CAA0213975.1 Length: 947
Range 1: 485 to 562

Score:68.6 bits(166), Expect:3e-11,
Method:Compositional matrix adjust.,
Identities:40/78(51%), Positives:49/78(62%), Gaps:10/78(12%)

Query  366  W*FVGLLP----PRHESTETIGRALTLGVNIKMITG------QITSCKLGMGINMYPSSA  515
            W F GLLP    PRH+S ETI RAL+LGV +KMITG      + T  +LGMG NMYPSS+
Sbjct  485  WRFCGLLPLFDPPRHDSGETILRALSLGVCVKMITGDQLAIAKETGRRLGMGTNMYPSSS  544

Query  516  FLGTHKW*EPTNPMDEYV  569
             LG +       P+DE +
Sbjct  545  LLGHNNDEHEAIPVDELI  562


>autoinhibited H[+]-ATPase isoform 10 [Arabidopsis thaliana]
Sequence ID: NP_173169.2 Length: 947
>RecName: Full=ATPase 10, plasma membrane-type; AltName: Full=Proton pump 10 [Arabidopsis thaliana]
Sequence ID: Q43128.2 Length: 947 >plasma membrane H(+)-ATPase isoform AHA10 [Arabidopsis thaliana]
Sequence ID: AAB32310.2 Length: 947 >H+-transporting ATPase AHA10 [Arabidopsis thaliana]
Sequence ID: AAD50009.3 Length: 947 >autoinhibited H[+]-ATPase isoform 10 [Arabidopsis thaliana]
Sequence ID: AEE29565.1 Length: 947 >AHA10 [Arabidopsis thaliana]
Sequence ID: OAP13355.1 Length: 947 >unnamed protein product [Arabidopsis thaliana]
Sequence ID: VYS46360.1 Length: 947
Range 1: 485 to 562

Score:68.6 bits(166), Expect:3e-11,
Method:Compositional matrix adjust.,
Identities:40/78(51%), Positives:49/78(62%), Gaps:10/78(12%)

Query  366  W*FVGLLP----PRHESTETIGRALTLGVNIKMITG------QITSCKLGMGINMYPSSA  515
            W F GLLP    PRH+S ETI RAL+LGV +KMITG      + T  +LGMG NMYPSS+
Sbjct  485  WRFCGLLPLFDPPRHDSGETILRALSLGVCVKMITGDQLAIAKETGRRLGMGTNMYPSSS  544

Query  516  FLGTHKW*EPTNPMDEYV  569
             LG +       P+DE +
Sbjct  545  LLGHNNDEHEAIPVDELI  562


>H[+]-ATPase 7 [Arabidopsis thaliana]
Sequence ID: NP_001327362.1 Length: 961
>H[+]-ATPase 7 [Arabidopsis thaliana]
Sequence ID: NP_001327363.1 Length: 961 >H[+]-ATPase 7 [Arabidopsis thaliana]
Sequence ID: ANM65388.1 Length: 961 >H[+]-ATPase 7 [Arabidopsis thaliana]
Sequence ID: ANM65389.1 Length: 961 >unnamed protein product [Arabidopsis thaliana]
Sequence ID: VYS60981.1 Length: 961
Range 1: 480 to 544

Score:64.7 bits(156), Expect:6e-10,
Method:Compositional matrix adjust.,
Identities:35/65(54%), Positives:43/65(66%), Gaps:10/65(15%)

Query  366  W*FVGLLP----PRHESTETIGRALTLGVNIKMITG------QITSCKLGMGINMYPSSA  515
            W FV LLP    PRH+S +TI RAL LGV++KMITG      + T  +LGMG NMYPSS+
Sbjct  480  WDFVALLPLFDPPRHDSAQTIERALHLGVSVKMITGDQLAIAKETGRRLGMGTNMYPSSS  539

Query  516  FLGTH  530
             L  +
Sbjct  540  LLSDN  544


>H[+]-ATPase 7 [Arabidopsis thaliana]
Sequence ID: NP_001190141.1 Length: 961
>H[+]-ATPase 7 [Arabidopsis thaliana]
Sequence ID: NP_191592.5 Length: 961 >RecName: Full=ATPase 7, plasma membrane-type; AltName: Full=Proton pump 7 [Arabidopsis thaliana]
Sequence ID: Q9LY32.1 Length: 961 >H[+]-ATPase 7 [Arabidopsis thaliana]
Sequence ID: AEE80046.1 Length: 961 >H[+]-ATPase 7 [Arabidopsis thaliana]
Sequence ID: AEE80047.1 Length: 961 >HA7 [Arabidopsis thaliana]
Sequence ID: OAP01410.1 Length: 961 >unnamed protein product [Arabidopsis thaliana]
Sequence ID: CAA0387627.1 Length: 961 >plasma membrane H+-ATPase-like [Arabidopsis thaliana]
Sequence ID: CAB87870.1 Length: 961
Range 1: 480 to 544

Score:64.7 bits(156), Expect:6e-10,
Method:Compositional matrix adjust.,
Identities:35/65(54%), Positives:43/65(66%), Gaps:10/65(15%)

Query  366  W*FVGLLP----PRHESTETIGRALTLGVNIKMITG------QITSCKLGMGINMYPSSA  515
            W FV LLP    PRH+S +TI RAL LGV++KMITG      + T  +LGMG NMYPSS+
Sbjct  480  WDFVALLPLFDPPRHDSAQTIERALHLGVSVKMITGDQLAIAKETGRRLGMGTNMYPSSS  539

Query  516  FLGTH  530
             L  +
Sbjct  540  LLSDN  544


>hypothetical protein AT1G44941 [Arabidopsis thaliana]
Sequence ID: NP_001117439.1 Length: 60
>hypothetical protein AT1G44941 [Arabidopsis thaliana]
Sequence ID: AEE32067.1 Length: 60
Range 1: 30 to 59

Score:57.0 bits(136), Expect:8e-10,
Method:Compositional matrix adjust.,
Identities:25/30(83%), Positives:28/30(93%), Gaps:0/30(0%)

Query  1251  QVVPNGSWTLPLAVMLQACLPIQPRDTSLP  1162
             QVVPNGSWTLPL V+L+ACLPIQPRDT+ P
Sbjct  30    QVVPNGSWTLPLVVLLRACLPIQPRDTNFP  59


>unnamed protein product [Arabidopsis thaliana]
Sequence ID: CAA0274387.1 Length: 60
Range 1: 30 to 59

Score:57.0 bits(136), Expect:8e-10,
Method:Compositional matrix adjust.,
Identities:25/30(83%), Positives:28/30(93%), Gaps:0/30(0%)

Query  1251  QVVPNGSWTLPLAVMLQACLPIQPRDTSLP  1162
             QVVPNGSWTLPL V+L+ACLPIQPRDT+ P
Sbjct  30    QVVPNGSWTLPLVVLLRACLPIQPRDTNFP  59


>unnamed protein product [Arabidopsis thaliana]
Sequence ID: VYS48295.1 Length: 60
Range 1: 30 to 59

Score:54.3 bits(129), Expect:6e-09,
Method:Compositional matrix adjust.,
Identities:24/30(80%), Positives:27/30(90%), Gaps:0/30(0%)

Query  1251  QVVPNGSWTLPLAVMLQACLPIQPRDTSLP  1162
             QVVPNGSWTLPL V+L+ACLPIQ RDT+ P
Sbjct  30    QVVPNGSWTLPLVVLLRACLPIQSRDTNFP  59


>hypothetical protein AT1G44940 [Arabidopsis thaliana]
Sequence ID: NP_001154402.1 Length: 81
>hypothetical protein AT1G44940 [Arabidopsis thaliana]
Sequence ID: AEE32065.1 Length: 81
Range 1: 26 to 60

Score:53.1 bits(126), Expect:3e-08,
Method:Compositional matrix adjust.,
Identities:26/35(74%), Positives:28/35(80%), Gaps:1/35(2%)

Query  479  TWNGNKHVSIFG-FSWYAQVVGTNKPHG*ICLSSL  580
            TWNGNKHVS+F  F  YAQ VGTNKPHG ICLS +
Sbjct  26   TWNGNKHVSLFRLFKEYAQAVGTNKPHGRICLSKV  60


>unnamed protein product [Arabidopsis thaliana]
Sequence ID: VYS48292.1 Length: 86
Range 1: 31 to 65

Score:53.1 bits(126), Expect:3e-08,
Method:Compositional matrix adjust.,
Identities:26/35(74%), Positives:28/35(80%), Gaps:1/35(2%)

Query  479  TWNGNKHVSIFG-FSWYAQVVGTNKPHG*ICLSSL  580
            TWNGNKHVS+F  F  YAQ VGTNKPHG ICLS +
Sbjct  31   TWNGNKHVSLFRLFKEYAQAVGTNKPHGRICLSKV  65


>hypothetical protein AT1G44940 [Arabidopsis thaliana]
Sequence ID: NP_001154403.1 Length: 86
>hypothetical protein AT1G44940 [Arabidopsis thaliana]
Sequence ID: AEE32066.1 Length: 86
Range 1: 31 to 65

Score:53.1 bits(126), Expect:3e-08,
Method:Compositional matrix adjust.,
Identities:26/35(74%), Positives:28/35(80%), Gaps:1/35(2%)

Query  479  TWNGNKHVSIFG-FSWYAQVVGTNKPHG*ICLSSL  580
            TWNGNKHVS+F  F  YAQ VGTNKPHG ICLS +
Sbjct  31   TWNGNKHVSLFRLFKEYAQAVGTNKPHGRICLSKV  65


Query #248: XLOC_027513 Query ID: lcl|Query_30739 Length: 1052

No significant similarity found.


Query #249: XLOC_027632 Query ID: lcl|Query_30740 Length: 846

Sequences producing significant alignments:
                                                                  Max    Total Query   E   Per.
Description                                                       Score  Score cover Value Ident  Accession
unnamed protein product [Arabidopsis thaliana]                    89.7   157   22%   2e-22 93.48  CAA0364544.1
unnamed protein product [Arabidopsis thaliana]                    89.0   153   22%   3e-22 93.48  CAA0405880.1
hypothetical protein AT5G20460 [Arabidopsis thaliana]             59.3   116   19%   3e-11 60.00  NP_197546.1


Alignments:
>unnamed protein product [Arabidopsis thaliana]
Sequence ID: CAA0364544.1 Length: 71
Range 1: 26 to 71

Score:89.7 bits(221), Expect:2e-22,
Method:Compositional matrix adjust.,
Identities:43/46(93%), Positives:44/46(95%), Gaps:0/46(0%)

Query  370  WTALTAFVIGDVHYEWQIMDGTLWMMNHGWHIMDDASWIVHYGLIW  507
            + ALTAFVIGDVHY WQIMDGTLWMMNHGWHIMDDASWIVHYGLIW
Sbjct  26   FVALTAFVIGDVHYGWQIMDGTLWMMNHGWHIMDDASWIVHYGLIW  71


Range 2: 1 to 59

Score:67.8 bits(164), Expect:3e-14,
Method:Compositional matrix adjust.,
Identities:38/62(61%), Positives:45/62(72%), Gaps:5/62(8%)

Query  320  MEVFPMVTLDEGGSPCVGQLLRRLLLV--MSIMNGKLWMAHYG**IMDGILWMTHHGLYI  493
            MEVFPMVTLDEGGSPCVGQLL + + V   + + G +   HYG  IMDG LWM +HG +I
Sbjct  1    MEVFPMVTLDEGGSPCVGQLLAKDVFVALTAFVIGDV---HYGWQIMDGTLWMMNHGWHI  57

Query  494  MD  499
            MD
Sbjct  58   MD  59


>unnamed protein product [Arabidopsis thaliana]
Sequence ID: CAA0405880.1 Length: 67
Range 1: 22 to 67

Score:89.0 bits(219), Expect:3e-22,
Method:Compositional matrix adjust.,
Identities:43/46(93%), Positives:44/46(95%), Gaps:0/46(0%)

Query  370  WTALTAFVIGDVHYEWQIMDGTLWMMNHGWHIMDDASWIVHYGLIW  507
            + ALTAFVIGDVHY WQIMDGTLWMMNHGWHIMDDASWIVHYGLIW
Sbjct  22   FVALTAFVIGDVHYGWQIMDGTLWMMNHGWHIMDDASWIVHYGLIW  67


Range 2: 1 to 55

Score:64.3 bits(155), Expect:6e-13,
Method:Compositional matrix adjust.,
Identities:36/60(60%), Positives:40/60(66%), Gaps:5/60(8%)

Query  320  MEVFPMVTLDEGGSPCVGQLLRRLLLVMSIMNGKLWMAHYG**IMDGILWMTHHGLYIMD  499
            MEVFPMVTLDEGGSPCVGQ +   L    I +      HYG  IMDG LWM +HG +IMD
Sbjct  1    MEVFPMVTLDEGGSPCVGQDVFVALTAFVIGD-----VHYGWQIMDGTLWMMNHGWHIMD  55


>hypothetical protein AT5G20460 [Arabidopsis thaliana]
Sequence ID: NP_197546.1 Length: 50
>hypothetical protein AT5G20460 [Arabidopsis thaliana]
Sequence ID: AED92848.1 Length: 50 >unnamed protein product [Arabidopsis thaliana]
Sequence ID: VYS58295.1 Length: 50
Range 1: 1 to 50

Score:59.3 bits(142), Expect:3e-11,
Method:Compositional matrix adjust.,
Identities:33/55(60%), Positives:36/55(65%), Gaps:5/55(9%)

Query  320  MEVFPMVTLDEGGSPCVGQLLRRLLLVMSIMNGKLWMAHYG**IMDGILWMTHHG  484
            MEVFPMVTLDEGGSPCVGQ +   L    I +      HYG  IMDG LWM +HG
Sbjct  1    MEVFPMVTLDEGGSPCVGQDVFVALTAFVIGD-----VHYGWQIMDGTLWMMNHG  50


Range 2: 22 to 50

Score:57.4 bits(137), Expect:1e-10,
Method:Compositional matrix adjust.,
Identities:26/29(90%), Positives:27/29(93%), Gaps:0/29(0%)

Query  370  WTALTAFVIGDVHYEWQIMDGTLWMMNHG  456
            + ALTAFVIGDVHY WQIMDGTLWMMNHG
Sbjct  22   FVALTAFVIGDVHYGWQIMDGTLWMMNHG  50


Query #250: XLOC_027757 Query ID: lcl|Query_30741 Length: 791

Sequences producing significant alignments:
                                                                  Max    Total Query   E   Per.
Description                                                       Score  Score cover Value Ident  Accession
unnamed protein product [Arabidopsis thaliana]                    72.8   72.8  32%   5e-15 47.67  CAA0301581.1
unnamed protein product [Arabidopsis thaliana]                    73.2   73.2  32%   2e-14 47.67  VYS50968.1


Alignments:
>unnamed protein product [Arabidopsis thaliana]
Sequence ID: CAA0301581.1 Length: 181
Range 1: 4 to 89

Score:72.8 bits(177), Expect:5e-15,
Method:Compositional matrix adjust.,
Identities:41/86(48%), Positives:51/86(59%), Gaps:1/86(1%)

Query  456  DNWSDEEC*YFLQLCTAEE-KGNRRKTRLNITGKESIMR*FEEKFGKRYSWDKQFKNKYD  632
            DNWSDEE  YF QL   E  KGNR    +N TGK+ IMR FEE+F + Y     FKN+YD
Sbjct  4    DNWSDEETRYFFQLYADERRKGNRTSIGMNQTGKDCIMRKFEERFQRGYQKWNPFKNRYD  63

Query  633  VRRRTYTKVKNLLHNRTEITYDSV*R  710
              ++ Y   + L HNR  + YD + R
Sbjct  64   ACKKKYATFRLLTHNRPMLQYDDMGR  89


>unnamed protein product [Arabidopsis thaliana]
Sequence ID: VYS50968.1 Length: 291
Range 1: 4 to 89

Score:73.2 bits(178), Expect:2e-14,
Method:Compositional matrix adjust.,
Identities:41/86(48%), Positives:51/86(59%), Gaps:1/86(1%)

Query  456  DNWSDEEC*YFLQLCTAEE-KGNRRKTRLNITGKESIMR*FEEKFGKRYSWDKQFKNKYD  632
            DNWSDEE  YF QL   E  KGNR    +N TGK+ IMR FEE+F + Y     FKN+YD
Sbjct  4    DNWSDEETRYFFQLYADERRKGNRTSIGMNQTGKDCIMRKFEERFQRGYQKWNAFKNRYD  63

Query  633  VRRRTYTKVKNLLHNRTEITYDSV*R  710
              ++ Y   + L HNR  + YD + R
Sbjct  64   ACKKKYATFRLLTHNRPMLQYDDMGR  89


Query #251: XLOC_028682 Query ID: lcl|Query_30742 Length: 817

Sequences producing significant alignments:
                                                                  Max    Total Query   E   Per.
Description                                                       Score  Score cover Value Ident  Accession
ubiquinol oxidase [Arabidopsis thaliana]                          111    111   20%   4e-31 100.00 NP_001330488.1
ubiquinol oxidase [Arabidopsis thaliana]                          107    107   19%   2e-29 100.00 NP_001330489.1
hypothetical protein AXX17_AT5G64340 [Arabidopsis thaliana]       103    103   20%   4e-28 92.86  OAO90914.1
HSR3 [Arabidopsis thaliana]                                       73.6   121   36%   5e-21 57.69  OAP01931.1
alternative oxidase [Arabidopsis thaliana]                        86.3   86.3  37%   9e-19 51.85  CAA10364.1
F28K19.26 [Arabidopsis thaliana]                                  67.4   107   26%   6e-17 66.00  AAF17689.1
unnamed protein product [Arabidopsis thaliana]                    68.6   68.6  13%   8e-15 91.89  CAA0411774.1
hypothetical protein AT5G63625 [Arabidopsis thaliana]             70.5   70.5  13%   1e-14 91.89  NP_001331694.1
alternative oxidase 1A [Arabidopsis thaliana]                     74.3   74.3  37%   2e-14 48.28  NP_188876.1
oxidase [Arabidopsis thaliana]                                    72.8   72.8  22%   3e-14 65.57  AAA32870.1


Alignments:
>ubiquinol oxidase [Arabidopsis thaliana]
Sequence ID: NP_001330488.1 Length: 56
>ubiquinol oxidase [Arabidopsis thaliana]
Sequence ID: ANM68766.1 Length: 56 >hypothetical protein AXX17_AT5G65020 [Arabidopsis thaliana]
Sequence ID: OAO95958.1 Length: 56 >unnamed protein product [Arabidopsis thaliana]
Sequence ID: CAA0412022.1 Length: 56 >unnamed protein product [Arabidopsis thaliana]
Sequence ID: CAA0412161.1 Length: 56 >unnamed protein product [Arabidopsis thaliana]
Sequence ID: VYS71408.1 Length: 56
Range 1: 1 to 56

Score:111 bits(278), Expect:4e-31,
Method:Compositional matrix adjust.,
Identities:56/56(100%), Positives:56/56(100%), Gaps:0/56(0%)

Query  610  MRFASTITLGEKASTKEEDANQRKTEKESTGGDQGIANQRKTENESTGGDQGIASY  777
            MRFASTITLGEKASTKEEDANQRKTEKESTGGDQGIANQRKTENESTGGDQGIASY
Sbjct  1    MRFASTITLGEKASTKEEDANQRKTEKESTGGDQGIANQRKTENESTGGDQGIASY  56


>ubiquinol oxidase [Arabidopsis thaliana]
Sequence ID: NP_001330489.1 Length: 61
>ubiquinol oxidase [Arabidopsis thaliana]
Sequence ID: ANM68767.1 Length: 61
Range 1: 1 to 54

Score:107 bits(267), Expect:2e-29,
Method:Compositional matrix adjust.,
Identities:54/54(100%), Positives:54/54(100%), Gaps:0/54(0%)

Query  610  MRFASTITLGEKASTKEEDANQRKTEKESTGGDQGIANQRKTENESTGGDQGIA  771
            MRFASTITLGEKASTKEEDANQRKTEKESTGGDQGIANQRKTENESTGGDQGIA
Sbjct  1    MRFASTITLGEKASTKEEDANQRKTEKESTGGDQGIANQRKTENESTGGDQGIA  54


>hypothetical protein AXX17_AT5G64340 [Arabidopsis thaliana]
Sequence ID: OAO90914.1 Length: 56
Range 1: 1 to 56

Score:103 bits(258), Expect:4e-28,
Method:Compositional matrix adjust.,
Identities:52/56(93%), Positives:53/56(94%), Gaps:0/56(0%)

Query  610  MRFASTITLGEKASTKEEDANQRKTEKESTGGDQGIANQRKTENESTGGDQGIASY  777
            MRFA  ITLGEKASTKEEDANQRKTE ESTGGDQGIANQRKTENES+GGDQGIASY
Sbjct  1    MRFACAITLGEKASTKEEDANQRKTENESTGGDQGIANQRKTENESSGGDQGIASY  56


>HSR3 [Arabidopsis thaliana]
Sequence ID: OAP01931.1 Length: 354
>unnamed protein product [Arabidopsis thaliana]
Sequence ID: CAA0383312.1 Length: 354
Range 1: 41 to 109

Score:73.6 bits(179), Expect:5e-21,
Method:Compositional matrix adjust.,
Identities:45/78(58%), Positives:49/78(62%), Gaps:16/78(20%)

Query  571  LEPSCCISAWI-------GGMRFASTITLGEKASTKEEDANQRKTEKESTGGDQGIANQR  729
            L+P    SAWI       GGMRFASTITLGEK   KEEDANQ+KTE ESTGGD    N +
Sbjct  41   LKPGVT-SAWIWTRAPTIGGMRFASTITLGEKTPMKEEDANQKKTENESTGGDAAGGNNK  99

Query  730  KTENESTGGDQGIASY*G  783
                    GD+GIASY G
Sbjct  100  --------GDKGIASYWG  109


Range 2: 12 to 45

Score:47.8 bits(112), Expect:5e-21,
Method:Compositional matrix adjust.,
Identities:21/34(62%), Positives:27/34(79%), Gaps:0/34(0%)

Query  485  RRLLTATKPRLFSTIRTITSYEAFSASHLLNPHV  586
            + LL A  PRLFST+RT++S+EA SASH+L P V
Sbjct  12   KSLLMAAGPRLFSTVRTVSSHEALSASHILKPGV  45


>alternative oxidase [Arabidopsis thaliana]
Sequence ID: CAA10364.1 Length: 353
Range 1: 10 to 108

Score:86.3 bits(212), Expect:9e-19,
Method:Compositional matrix adjust.,
Identities:56/108(52%), Positives:64/108(59%), Gaps:16/108(14%)

Query  481  AKAAVNGD*TTFVLNDPYDYVLRGVLSKPSLEPSCCISAWI-------GGMRFASTITLG  639
            A+ AV+G  TTFVL+ PY      +     L+P    SAWI       GGMRFASTITLG
Sbjct  10   AQIAVSGGWTTFVLDGPYVSSHEALSRSHILKPGVT-SAWIWTRAPTIGGMRFASTITLG  68

Query  640  EKASTKEEDANQRKTEKESTGGDQGIANQRKTENESTGGDQGIASY*G  783
            EK   KEEDANQ+KTE ESTGGD    N +        GD+GIASY G
Sbjct  69   EKTPMKEEDANQKKTENESTGGDAAGGNNK--------GDKGIASYWG  108


>F28K19.26 [Arabidopsis thaliana]
Sequence ID: AAF17689.1 Length: 677
Range 1: 93 to 142

Score:67.4 bits(163), Expect:6e-17,
Method:Compositional matrix adjust.,
Identities:33/50(66%), Positives:41/50(82%), Gaps:0/50(0%)

Query  458  LLSVESVEPRRLLTATKPRLFSTIRTITSYEAFSASHLLNPHVAYRLGLE  607
            + ++  +E RR+LTA +PRL STIRTITS+ AFSAS LL P+VAYRLGLE
Sbjct  93   VFTLSRMEQRRMLTAARPRLSSTIRTITSHVAFSASRLLKPYVAYRLGLE  142


Range 2: 142 to 162

Score:40.0 bits(92), Expect:6e-17,
Method:Compositional matrix adjust.,
Identities:18/21(86%), Positives:19/21(90%), Gaps:0/21(0%)

Query  661  EDANQRKTEKESTGGDQGIAN  723
            EDANQRKTE ESTGGD+ IAN
Sbjct  142  EDANQRKTENESTGGDKEIAN  162


>unnamed protein product [Arabidopsis thaliana]
Sequence ID: CAA0411774.1 Length: 39
Range 1: 1 to 37

Score:68.6 bits(166), Expect:8e-15,
Method:Compositional matrix adjust.,
Identities:34/37(92%), Positives:35/37(94%), Gaps:0/37(0%)

Query  610  MRFASTITLGEKASTKEEDANQRKTEKESTGGDQGIA  720
            MRFA TITLGEKASTKEEDANQRKTE ESTGGD+GIA
Sbjct  1    MRFACTITLGEKASTKEEDANQRKTENESTGGDKGIA  37


>hypothetical protein AT5G63625 [Arabidopsis thaliana]
Sequence ID: NP_001331694.1 Length: 124
>hypothetical protein AT5G63625 [Arabidopsis thaliana]
Sequence ID: ANM70057.1 Length: 124 >unnamed protein product [Arabidopsis thaliana]
Sequence ID: VYS71299.1 Length: 124
Range 1: 1 to 37

Score:70.5 bits(171), Expect:1e-14,
Method:Compositional matrix adjust.,
Identities:34/37(92%), Positives:35/37(94%), Gaps:0/37(0%)

Query  610  MRFASTITLGEKASTKEEDANQRKTEKESTGGDQGIA  720
            MRFA TITLGEKASTKEEDANQRKTE ESTGGD+GIA
Sbjct  1    MRFACTITLGEKASTKEEDANQRKTENESTGGDKGIA  37


>alternative oxidase 1A [Arabidopsis thaliana]
Sequence ID: NP_188876.1 Length: 354
>RecName: Full=Ubiquinol oxidase 1a, mitochondrial; AltName: Full=Alternative oxidase 1a; Flags: Precursor [Arabidopsis thaliana]
Sequence ID: Q39219.2 Length: 354 >putative alternative oxidase 1a precursor [Arabidopsis thaliana]
Sequence ID: AAK43981.1 Length: 354 >putative alternative oxidase 1a precursor [Arabidopsis thaliana]
Sequence ID: AAL15234.1 Length: 354 >alternative oxidase 1A [Arabidopsis thaliana]
Sequence ID: AEE76627.1 Length: 354 >unnamed protein product [Arabidopsis thaliana]
Sequence ID: VYS58238.1 Length: 354 >alternative oxidase [Arabidopsis thaliana]
Sequence ID: BAA22625.1 Length: 354
Range 1: 6 to 109

Score:74.3 bits(181), Expect:2e-14,
Method:Compositional matrix adjust.,
Identities:56/116(48%), Positives:64/116(55%), Gaps:25/116(21%)

Query  475  GGAKAAVNGD*TTFVLNDPYDY-VLRGVLSKPSLEPSCCI-----SAWI-------GGMR  615
            GGAKAA     +  V   P  +  +R V S  +L  S  +     SAWI       GGMR
Sbjct  6    GGAKAAK----SLLVAAGPRLFSTVRTVSSHEALSASHILKPGVTSAWIWTRAPTIGGMR  61

Query  616  FASTITLGEKASTKEEDANQRKTEKESTGGDQGIANQRKTENESTGGDQGIASY*G  783
            FASTITLGEK   KEEDANQ+KTE ESTGGD    N +        GD+GIASY G
Sbjct  62   FASTITLGEKTPMKEEDANQKKTENESTGGDAAGGNNK--------GDKGIASYWG  109


>oxidase [Arabidopsis thaliana]
Sequence ID: AAA32870.1 Length: 305
Range 1: 8 to 60

Score:72.8 bits(177), Expect:3e-14,
Method:Compositional matrix adjust.,
Identities:40/61(66%), Positives:43/61(70%), Gaps:8/61(13%)

Query  601  IGGMRFASTITLGEKASTKEEDANQRKTEKESTGGDQGIANQRKTENESTGGDQGIASY*  780
            IGGMRFASTITLGEK   KEEDANQ+KTE ESTGGD    N +        GD+GIASY
Sbjct  8    IGGMRFASTITLGEKTPMKEEDANQKKTENESTGGDAAGGNNK--------GDKGIASYW  59

Query  781  G  783
            G
Sbjct  60   G  60


Query #252: XLOC_029176 Query ID: lcl|Query_30743 Length: 454

No significant similarity found.


Query #253: XLOC_029177 Query ID: lcl|Query_30744 Length: 922

No significant similarity found.


Query #254: XLOC_029588 Query ID: lcl|Query_30745 Length: 1586

Sequences producing significant alignments:
                                                                  Max    Total Query   E   Per.
Description                                                       Score  Score cover Value Ident  Accession
unnamed protein product [Arabidopsis thaliana]                    110    362   33%   5e-54 98.08  CAA0402789.1
unknown protein [Arabidopsis thaliana]                            168    168   15%   4e-50 100.00 ABF59405.1
unknown [Arabidopsis thaliana]                                    167    167   15%   5e-50 100.00 ABK28065.1
hypothetical protein AXX17_AT5G15050 [Arabidopsis thaliana]       110    253   23%   1e-41 92.73  OAO92530.1
hypothetical protein AXX17_AT5G15050 [Arabidopsis thaliana]       82.4   209   19%   6e-29 82.61  OAO92531.1
unnamed protein product [Arabidopsis thaliana]                    81.3   231   27%   9e-26 88.37  CAA0373951.1
unknown protein [Arabidopsis thaliana]                            81.3   138   17%   1e-25 88.37  ABF59334.1
transmembrane protein [Arabidopsis thaliana]                      79.0   79.0  7%    1e-17 94.74  NP_001332724.1
Hypothetical protein [Arabidopsis thaliana]                       75.9   75.9  12%   1e-13 52.11  AAD30632.1
hypothetical protein AXX17_AT2G27970 [Arabidopsis thaliana]       56.6   110   11%   8e-12 75.76  OAP09547.1
hypothetical protein AXX17_AT2G10350 [Arabidopsis thaliana]       67.8   67.8  11%   5e-11 47.76  OAP11405.1
putative retroelement pol polyprotein [Arabidopsis thaliana]      64.7   64.7  11%   4e-10 45.45  AAD03367.1


Alignments:
>unnamed protein product [Arabidopsis thaliana]
Sequence ID: CAA0402789.1 Length: 186
Range 1: 64 to 115

Score:110 bits(275), Expect:5e-54,
Method:Compositional matrix adjust.,
Identities:51/52(98%), Positives:52/52(100%), Gaps:0/52(0%)

Query  1062  REDSDIHNLAPKGGASRIGIAMAKNGDQTSTLDEIKKMMNDQQQCTISELLK  1217
             +EDSDIHNLAPKGGASRIGIAMAKNGDQTSTLDEIKKMMNDQQQCTISELLK
Sbjct  64    KEDSDIHNLAPKGGASRIGIAMAKNGDQTSTLDEIKKMMNDQQQCTISELLK  115


Range 2: 140 to 186

Score:82.8 bits(203), Expect:5e-54,
Method:Compositional matrix adjust.,
Identities:38/47(81%), Positives:41/47(87%), Gaps:0/47(0%)

Query  1301  YLCRPS*HSTPGTLRTSRSRGRLEFYDPIHPNPLSFSNYCKVSGPKV  1441
              +  P+  +TPGTLRTSRSRGRLEFYDPIHPNPLSFSNYCK SGPKV
Sbjct  140   IILMPTELTTPGTLRTSRSRGRLEFYDPIHPNPLSFSNYCKGSGPKV  186


Range 3: 118 to 148

Score:60.8 bits(146), Expect:5e-54,
Method:Compositional matrix adjust.,
Identities:30/31(97%), Positives:31/31(100%), Gaps:0/31(0%)

Query  1231  LRSVATLSLQPLPQVAPKSRQLIILMPTELT  1323
             +RSVATLSLQPLPQVAPKSRQLIILMPTELT
Sbjct  118   MRSVATLSLQPLPQVAPKSRQLIILMPTELT  148


Range 4: 45 to 64

Score:46.6 bits(109), Expect:5e-11,
Method:Compositional matrix adjust.,
Identities:19/20(95%), Positives:20/20(100%), Gaps:0/20(0%)

Query  319  QIGMIGAFGVCVVFNTPISK  378
            +IGMIGAFGVCVVFNTPISK
Sbjct  45   KIGMIGAFGVCVVFNTPISK  64


Range 5: 29 to 46

Score:32.3 bits(72), Expect:5e-11,
Method:Compositional matrix adjust.,
Identities:12/18(67%), Positives:16/18(88%), Gaps:0/18(0%)

Query  177  YVIFLRNHKEGIVYMSKM  230
            + + +RNHKEGIVYMSK+
Sbjct  29   FPLLIRNHKEGIVYMSKI  46


Range 6: 19 to 34

Score:29.3 bits(64), Expect:5e-11,
Method:Compositional matrix adjust.,
Identities:14/16(88%), Positives:14/16(87%), Gaps:0/16(0%)

Query  70   IKKSNVQTVSFPLLIR  117
            I  SNVQTVSFPLLIR
Sbjct  19   IITSNVQTVSFPLLIR  34


>unknown protein [Arabidopsis thaliana]
Sequence ID: ABF59405.1 Length: 81
Range 1: 1 to 81

Score:168 bits(425), Expect:4e-50,
Method:Compositional matrix adjust.,
Identities:81/81(100%), Positives:81/81(100%), Gaps:0/81(0%)

Query  1125  MAKNGDQTSTLDEIKKMMNDQQQCTISELLKKFISFEERCNTQLAALTASCSKITSTHNT  1304
             MAKNGDQTSTLDEIKKMMNDQQQCTISELLKKFISFEERCNTQLAALTASCSKITSTHNT
Sbjct  1     MAKNGDQTSTLDEIKKMMNDQQQCTISELLKKFISFEERCNTQLAALTASCSKITSTHNT  60

Query  1305  YADRVDTLLQGLCEPQDQEGV  1367
             YADRVDTLLQGLCEPQDQEGV
Sbjct  61    YADRVDTLLQGLCEPQDQEGV  81


>unknown, partial [Arabidopsis thaliana]
Sequence ID: ABK28065.1 Length: 82
Range 1: 1 to 81

Score:167 bits(424), Expect:5e-50,
Method:Compositional matrix adjust.,
Identities:81/81(100%), Positives:81/81(100%), Gaps:0/81(0%)

Query  1125  MAKNGDQTSTLDEIKKMMNDQQQCTISELLKKFISFEERCNTQLAALTASCSKITSTHNT  1304
             MAKNGDQTSTLDEIKKMMNDQQQCTISELLKKFISFEERCNTQLAALTASCSKITSTHNT
Sbjct  1     MAKNGDQTSTLDEIKKMMNDQQQCTISELLKKFISFEERCNTQLAALTASCSKITSTHNT  60

Query  1305  YADRVDTLLQGLCEPQDQEGV  1367
             YADRVDTLLQGLCEPQDQEGV
Sbjct  61    YADRVDTLLQGLCEPQDQEGV  81


>hypothetical protein AXX17_AT5G15050 [Arabidopsis thaliana]
Sequence ID: OAO92530.1 Length: 139
Range 1: 17 to 71

Score:110 bits(274), Expect:1e-41,
Method:Compositional matrix adjust.,
Identities:51/55(93%), Positives:53/55(96%), Gaps:0/55(0%)

Query  1062  REDSDIHNLAPKGGASRIGIAMAKNGDQTSTLDEIKKMMNDQQQCTISELLKKFI  1226
             +EDSDIHNLAPKGGASRIGIAMAKNGDQTSTLDEIKKMMNDQQQCTISELLK  +
Sbjct  17    KEDSDIHNLAPKGGASRIGIAMAKNGDQTSTLDEIKKMMNDQQQCTISELLKMMM  71


Range 2: 93 to 139

Score:82.0 bits(201), Expect:1e-41,
Method:Compositional matrix adjust.,
Identities:38/47(81%), Positives:41/47(87%), Gaps:0/47(0%)

Query  1301  YLCRPS*HSTPGTLRTSRSRGRLEFYDPIHPNPLSFSNYCKVSGPKV  1441
              +  P+  +TPGTLRTSRSRGRLEFYDPIHPNPLSFSNYCK SGPKV
Sbjct  93    IILMPTELTTPGTLRTSRSRGRLEFYDPIHPNPLSFSNYCKGSGPKV  139


Range 3: 71 to 134

Score:60.8 bits(146), Expect:2e-10,
Method:Compositional matrix adjust.,
Identities:37/68(54%), Positives:41/68(60%), Gaps:7/68(10%)

Query  1231  LRSVATLSLQPLPQVAPKSRQLIILMPTELTLYSRDFANLKIKRA---FRVLRPYPPEPS  1401
             +RSVATLSLQPLPQVAPKSRQLIILMPTELT        L+  R+        P  P P
Sbjct  71    MRSVATLSLQPLPQVAPKSRQLIILMPTELTTP----GTLRTSRSRGRLEFYDPIHPNPL  126

Query  1402  EFLKLLQG  1425
              F    +G
Sbjct  127   SFSNYCKG  134


>hypothetical protein AXX17_AT5G15050 [Arabidopsis thaliana]
Sequence ID: OAO92531.1 Length: 102
Range 1: 57 to 102

Score:82.4 bits(202), Expect:6e-29,
Method:Compositional matrix adjust.,
Identities:38/46(83%), Positives:41/46(89%), Gaps:0/46(0%)

Query  1304  LCRPS*HSTPGTLRTSRSRGRLEFYDPIHPNPLSFSNYCKVSGPKV  1441
             +  P+  +TPGTLRTSRSRGRLEFYDPIHPNPLSFSNYCK SGPKV
Sbjct  57    ILMPTELTTPGTLRTSRSRGRLEFYDPIHPNPLSFSNYCKGSGPKV  102


Range 2: 1 to 34

Score:67.0 bits(162), Expect:6e-29,
Method:Compositional matrix adjust.,
Identities:31/34(91%), Positives:32/34(94%), Gaps:0/34(0%)

Query  1125  MAKNGDQTSTLDEIKKMMNDQQQCTISELLKKFI  1226
             MAKNGDQTSTLDEIKKMMNDQQQCTISELLK  +
Sbjct  1     MAKNGDQTSTLDEIKKMMNDQQQCTISELLKMMM  34


Range 3: 34 to 97

Score:60.1 bits(144), Expect:1e-10,
Method:Compositional matrix adjust.,
Identities:37/68(54%), Positives:41/68(60%), Gaps:7/68(10%)

Query  1231  LRSVATLSLQPLPQVAPKSRQLIILMPTELTLYSRDFANLKIKRA---FRVLRPYPPEPS  1401
             +RSVATLSLQPLPQVAPKSRQLIILMPTELT        L+  R+        P  P P
Sbjct  34    MRSVATLSLQPLPQVAPKSRQLIILMPTELTTP----GTLRTSRSRGRLEFYDPIHPNPL  89

Query  1402  EFLKLLQG  1425
              F    +G
Sbjct  90    SFSNYCKG  97


>unnamed protein product [Arabidopsis thaliana]
Sequence ID: CAA0373951.1 Length: 198
Range 1: 86 to 128

Score:81.3 bits(199), Expect:9e-26,
Method:Compositional matrix adjust.,
Identities:38/43(88%), Positives:40/43(93%), Gaps:0/43(0%)

Query  1200  ISELLKKFISFEERCNTQLAALTASCSKITSTHNTYADRVDTL  1328
             I+ELL+KFISFEER NTQLAALTASCSKITS HNTYADRVDT
Sbjct  86    INELLEKFISFEERYNTQLAALTASCSKITSAHNTYADRVDTF  128


Range 2: 145 to 189

Score:57.4 bits(137), Expect:9e-26,
Method:Compositional matrix adjust.,
Identities:31/48(65%), Positives:36/48(75%), Gaps:4/48(8%)

Query  1375  LRPYPPEPSEFLK-LLQGLRSQSVTGGSYQQPPTAQILL*ASTTSGYN  1515
             LRPYPP+P  F   +LQGL SQSVTGGS QQPPTAQI    +T+ G+N
Sbjct  145   LRPYPPDPLSFSNYVLQGLSSQSVTGGSSQQPPTAQI---PTTSVGFN  189


Range 3: 18 to 48

Score:47.0 bits(110), Expect:4e-12,
Method:Compositional matrix adjust.,
Identities:25/39(64%), Positives:27/39(69%), Gaps:8/39(20%)

Query  114  QVSVSSAFQNKFQKTKKME*VYVIFLRNHKEGIVYMSKM  230
            +VSVS AFQNKF KTKKME        N KEGIVYM K+
Sbjct  18   KVSVSFAFQNKFHKTKKME--------NRKEGIVYMRKI  48


Range 4: 47 to 65

Score:45.4 bits(106), Expect:4e-12,
Method:Compositional matrix adjust.,
Identities:18/19(95%), Positives:19/19(100%), Gaps:0/19(0%)

Query  319  QIGMIGAFGVCVVFNTPIS  375
            +IGMIGAFGVCVVFNTPIS
Sbjct  47   KIGMIGAFGVCVVFNTPIS  65


>unknown protein [Arabidopsis thaliana]
Sequence ID: ABF59334.1 Length: 134
Range 1: 22 to 64

Score:81.3 bits(199), Expect:1e-25,
Method:Compositional matrix adjust.,
Identities:38/43(88%), Positives:40/43(93%), Gaps:0/43(0%)

Query  1200  ISELLKKFISFEERCNTQLAALTASCSKITSTHNTYADRVDTL  1328
             I+ELL+KFISFEER NTQLAALTASCSKITS HNTYADRVDT
Sbjct  22    INELLEKFISFEERYNTQLAALTASCSKITSAHNTYADRVDTF  64


Range 2: 81 to 125

Score:57.0 bits(136), Expect:1e-25,
Method:Compositional matrix adjust.,
Identities:31/48(65%), Positives:36/48(75%), Gaps:4/48(8%)

Query  1375  LRPYPPEPSEFLK-LLQGLRSQSVTGGSYQQPPTAQILL*ASTTSGYN  1515
             LRPYPP+P  F   +LQGL SQSVTGGS QQPPTAQI    +T+ G+N
Sbjct  81    LRPYPPDPLSFSNYVLQGLSSQSVTGGSSQQPPTAQI---PTTSVGFN  125


>transmembrane protein [Arabidopsis thaliana]
Sequence ID: NP_001332724.1 Length: 71
>transmembrane protein [Arabidopsis thaliana]
Sequence ID: NP_001332725.1 Length: 71 >transmembrane protein [Arabidopsis thaliana]
Sequence ID: ANM71176.1 Length: 71 >transmembrane protein [Arabidopsis thaliana]
Sequence ID: ANM71177.1 Length: 71
Range 1: 34 to 71

Score:79.0 bits(193), Expect:1e-17,
Method:Compositional matrix adjust.,
Identities:36/38(95%), Positives:38/38(100%), Gaps:0/38(0%)

Query  316  NQIGMIGAFGVCVVFNTPISKWINFSFINQTSIKHPRL  429
            ++IGMIGAFGVCVVFNTPISKWINFSFINQTSIKHPRL
Sbjct  34   SKIGMIGAFGVCVVFNTPISKWINFSFINQTSIKHPRL  71


>Hypothetical protein [Arabidopsis thaliana]
Sequence ID: AAD30632.1 Length: 1295
Range 1: 9 to 79

Score:75.9 bits(185), Expect:1e-13,
Method:Composition-based stats.,
Identities:37/71(52%), Positives:50/71(70%), Gaps:6/71(8%)

Query  1134  NGDQTSTLDEIKKMMN------DQQQCTISELLKKFISFEERCNTQLAALTASCSKITST  1295
             +G Q STLD IK+++       D+QQ  ++ L  KF+SF+ERCN QLAAL  + S+I ST
Sbjct  9     SGGQASTLDNIKQLLQQLSEKTDRQQLAVTTLSDKFVSFQERCNGQLAALAVNHSEIRST  68

Query  1296  HNTYADRVDTL  1328
             HN YA+RVD+L
Sbjct  69    HNAYAERVDSL  79


>hypothetical protein AXX17_AT2G27970 [Arabidopsis thaliana]
Sequence ID: OAP09547.1 Length: 77
Range 1: 37 to 69

Score:56.6 bits(135), Expect:8e-12,
Method:Compositional matrix adjust.,
Identities:25/33(76%), Positives:28/33(84%), Gaps:0/33(0%)

Query  316  NQIGMIGAFGVCVVFNTPISKWINFSFINQTSI  414
             +IGMIGAFGVCVVFNTPISKWINFSF  +  +
Sbjct  37   RKIGMIGAFGVCVVFNTPISKWINFSFRTKPQL  69


Range 2: 61 to 76

Score:27.7 bits(60), Expect:8e-12,
Method:Composition-based stats.,
Identities:13/16(81%), Positives:13/16(81%), Gaps:0/16(0%)

Query  389  FPL*TKPQLSTHVFRA  436
            F   TKPQLSTHVFRA
Sbjct  61   FSFRTKPQLSTHVFRA  76


Range 3: 22 to 40

Score:26.6 bits(57), Expect:8e-12,
Method:Compositional matrix adjust.,
Identities:10/19(53%), Positives:14/19(73%), Gaps:0/19(0%)

Query  177  YVIFLRNHKEGIVYMSKMN  233
            + + +RN KEGIVYM K+
Sbjct  22   FPLLIRNRKEGIVYMRKIG  40


>hypothetical protein AXX17_AT2G10350 [Arabidopsis thaliana]
Sequence ID: OAP11405.1 Length: 1283
Range 1: 9 to 75

Score:67.8 bits(164), Expect:5e-11,
Method:Composition-based stats.,
Identities:32/67(48%), Positives:47/67(70%), Gaps:6/67(8%)

Query  1134  NGDQTSTLDEIKKMMN------DQQQCTISELLKKFISFEERCNTQLAALTASCSKITST  1295
             +GDQ S LD+IK+++       D+QQ  ++ L  KF++F+E+CN QLAAL A+ S+I S
Sbjct  9     SGDQASILDDIKQLLQKLSKKTDRQQLAVTSLSNKFVTFQEQCNDQLAALAANHSEIRSA  68

Query  1296  HNTYADR  1316
             HN YA+R
Sbjct  69    HNAYAER  75


>putative retroelement pol polyprotein [Arabidopsis thaliana]
Sequence ID: AAD03367.1 Length: 1329
Range 1: 9 to 74

Score:64.7 bits(156), Expect:4e-10,
Method:Composition-based stats.,
Identities:30/66(45%), Positives:45/66(68%), Gaps:6/66(9%)

Query  1134  NGDQTSTLDEIKKMMN------DQQQCTISELLKKFISFEERCNTQLAALTASCSKITST  1295
             +GDQ STLD+IK+++       D+QQ  ++ L  KF++F+E+CN Q AAL  + S+I S
Sbjct  9     SGDQASTLDDIKQLLQQLSEKTDRQQLAVTSLSNKFVTFQEQCNGQHAALATNHSEIRSA  68

Query  1296  HNTYAD  1313
             HN YA+
Sbjct  69    HNAYAE  74


Query #255: XLOC_029813 Query ID: lcl|Query_30746 Length: 865

No significant similarity found.


Query #256: XLOC_030032 Query ID: lcl|Query_30747 Length: 1614

Sequences producing significant alignments:
                                                                  Max    Total Query    E    Per.
Description                                                       Score  Score cover Value  Ident  Accession
En/Spm-like transposon [Arabidopsis thaliana]                     781    781   72%   0.0    100.00 NP_001331299.1
unnamed protein product [Arabidopsis thaliana]                    778    926   88%   0.0    99.74  BAB10393.1
En/Spm-like transposon [Arabidopsis thaliana]                     777    777   72%   0.0    100.00 NP_001331300.1
En/Spm-like transposon [Arabidopsis thaliana]                     774    774   72%   0.0    100.00 NP_001331298.1
hypothetical protein AXX17_AT5G24050 [Arabidopsis thaliana]       759    759   72%   0.0    97.68  OAO94301.1
unnamed protein product [Arabidopsis thaliana]                    745    825   78%   0.0    93.69  CAA0404532.1
unnamed protein product [Arabidopsis thaliana]                    738    808   79%   0.0    95.10  VYS67775.1
unnamed protein product [Arabidopsis thaliana]                    499    617   90%   3e-169 64.22  BAB10394.1
gamma-irradiation and mitomycin c induced 1 [Arabidopsis...       499    617   90%   5e-169 64.22  NP_001330245.1
gamma-irradiation and mitomycin c induced 1 [Arabidopsis...       499    616   90%   5e-169 64.22  NP_197816.3
GMI1 [Arabidopsis thaliana]                                       501    615   90%   2e-168 64.45  OAO93531.1
unnamed protein product [Arabidopsis thaliana]                    497    612   90%   1e-167 63.98  CAA0404534.1
hypothetical protein AXX17_AT5G24030 [Arabidopsis thaliana]       455    455   52%   2e-156 81.11  OAO92751.1
At5g24280 [Arabidopsis thaliana]                                  176    239   28%   7e-56  65.60  AAZ74706.1
At5g24280 [Arabidopsis thaliana]                                  174    238   28%   2e-55  64.80  AAZ74709.1
At5g24280 [Arabidopsis thaliana]                                  174    236   28%   7e-55  64.80  AAZ74705.1
At5g24280 [Arabidopsis thaliana]                                  174    235   28%   8e-55  64.80  AAZ74715.1
At5g24280 [Arabidopsis thaliana]                                  170    234   28%   3e-54  64.00  AAZ74701.1
At5g24280 [Arabidopsis thaliana]                                  170    233   28%   4e-54  64.00  AAZ74703.1
putative protein [Arabidopsis thaliana]                           169    169   29%   8e-49  62.73  AAM97146.1
hypothetical protein AXX17_AT3G27420 [Arabidopsis thaliana]       169    169   23%   1e-48  70.63  OAP01651.1
unnamed protein product [Arabidopsis thaliana]                    159    159   29%   2e-44  56.74  BAB01319.1
defective in meristem silencing 3 [Arabidopsis thaliana]          120    120   40%   5e-29  32.52  NP_566916.1
Chain C, Protein DEFECTIVE IN MERISTEM SILENCING 3 [Arabidopsi... 120    120   40%   6e-29  32.52  6OIS_C
hypothetical protein [Arabidopsis thaliana]                       105    105   40%   1e-23  32.19  CAB66404.1
hypothetical protein AXX17_AT3G27430 [Arabidopsis thaliana]       78.6   78.6  9%    1e-16  72.00  OAP05396.1
hypothetical protein AXX17_AT3G27430 [Arabidopsis thaliana]       79.0   79.0  9%    3e-16  72.00  OAP05397.1


Alignments:
>En/Spm-like transposon [Arabidopsis thaliana]
Sequence ID: NP_001331299.1 Length: 807
>En/Spm-like transposon [Arabidopsis thaliana]
Sequence ID: ANM69636.1 Length: 807
Range 1: 420 to 807

Score:781 bits(2018), Expect:0.0,
Method:Compositional matrix adjust.,
Identities:388/388(100%), Positives:388/388(100%), Gaps:0/388(0%)

Query  2     KESQTERRELMLLTKLPECCVAGSNLTNLIFKVTDSDGVMDTSIHHDEKSGCFHTMsiet  181
             KESQTERRELMLLTKLPECCVAGSNLTNLIFKVTDSDGVMDTSIHHDEKSGCFHTMSIET
Sbjct  420   KESQTERRELMLLTKLPECCVAGSNLTNLIFKVTDSDGVMDTSIHHDEKSGCFHTMSIET  479

Query  182   dsssdeseiRYAFVHGSCKVPTLSLPEREGVFSFKVFHSRFPELHLSLKIQLTPAQILQR  361
             DSSSDESEIRYAFVHGSCKVPTLSLPEREGVFSFKVFHSRFPELHLSLKIQLTPAQILQR
Sbjct  480   DSSSDESEIRYAFVHGSCKVPTLSLPEREGVFSFKVFHSRFPELHLSLKIQLTPAQILQR  539

Query  362   DETSYSRMGLTPKSKMASTTYSPALSSQTGPSLRDVAQFTESFKENLIGYSEHRVEIDER  541
             DETSYSRMGLTPKSKMASTTYSPALSSQTGPSLRDVAQFTESFKENLIGYSEHRVEIDER
Sbjct  540   DETSYSRMGLTPKSKMASTTYSPALSSQTGPSLRDVAQFTESFKENLIGYSEHRVEIDER  599

Query  542   LHCLEAEQNQAKEELRTLQASLEPLGAMFPECLSTKESMMKQIEEKHHDTAASVFCCLYR  721
             LHCLEAEQNQAKEELRTLQASLEPLGAMFPECLSTKESMMKQIEEKHHDTAASVFCCLYR
Sbjct  600   LHCLEAEQNQAKEELRTLQASLEPLGAMFPECLSTKESMMKQIEEKHHDTAASVFCCLYR  659

Query  722   KSPPPQSLFLSKKGVFGLVALLGSVASTSLSRVLSEYLGEDMLLALVCKSAQIGPNNAEF  901
             KSPPPQSLFLSKKGVFGLVALLGSVASTSLSRVLSEYLGEDMLLALVCKSAQIGPNNAEF
Sbjct  660   KSPPPQSLFLSKKGVFGLVALLGSVASTSLSRVLSEYLGEDMLLALVCKSAQIGPNNAEF  719

Query  902   LRLQSNHRFHVLCLDAIRPWKDGLLENDPQKKLAMDDPKLPDGDPIPGFKGYAVNMIDLA  1081
             LRLQSNHRFHVLCLDAIRPWKDGLLENDPQKKLAMDDPKLPDGDPIPGFKGYAVNMIDLA
Sbjct  720   LRLQSNHRFHVLCLDAIRPWKDGLLENDPQKKLAMDDPKLPDGDPIPGFKGYAVNMIDLA  779

Query  1082  PEELTVQTYSGYGLRETLFYNLFGNLQV  1165
             PEELTVQTYSGYGLRETLFYNLFGNLQV
Sbjct  780   PEELTVQTYSGYGLRETLFYNLFGNLQV  807


>unnamed protein product [Arabidopsis thaliana]
Sequence ID: BAB10393.1 Length: 1335
Range 1: 853 to 1240

Score:778 bits(2009), Expect:0.0,
Method:Compositional matrix adjust.,
Identities:387/388(99%), Positives:388/388(100%), Gaps:0/388(0%)

Query  2     KESQTERRELMLLTKLPECCVAGSNLTNLIFKVTDSDGVMDTSIHHDEKSGCFHTMsiet  181
             KESQTERRELMLLTKLPECCVAGSNLTNLIFKVTDSDGVMDTSIHHDEKSGCFHTMSIET
Sbjct  853   KESQTERRELMLLTKLPECCVAGSNLTNLIFKVTDSDGVMDTSIHHDEKSGCFHTMSIET  912

Query  182   dsssdeseiRYAFVHGSCKVPTLSLPEREGVFSFKVFHSRFPELHLSLKIQLTPAQILQR  361
             DSSSDESEIRYAFVHGSCKVPTLSLPEREGVFSFKVFHSRFPELHLSLKIQLTPAQILQR
Sbjct  913   DSSSDESEIRYAFVHGSCKVPTLSLPEREGVFSFKVFHSRFPELHLSLKIQLTPAQILQR  972

Query  362   DETSYSRMGLTPKSKMASTTYSPALSSQTGPSLRDVAQFTESFKENLIGYSEHRVEIDER  541
             DETSYSRMGLTPKSKMASTTYSPALSSQTGPSLRDVAQFTESFKENLIGYSEHRVEIDER
Sbjct  973   DETSYSRMGLTPKSKMASTTYSPALSSQTGPSLRDVAQFTESFKENLIGYSEHRVEIDER  1032

Query  542   LHCLEAEQNQAKEELRTLQASLEPLGAMFPECLSTKESMMKQIEEKHHDTAASVFCCLYR  721
             LHCLEAEQNQAKEELRTLQASLEPLGAMFPECLSTKESMMKQIEEKHHDTAASVFCCLYR
Sbjct  1033  LHCLEAEQNQAKEELRTLQASLEPLGAMFPECLSTKESMMKQIEEKHHDTAASVFCCLYR  1092

Query  722   KSPPPQSLFLSKKGVFGLVALLGSVASTSLSRVLSEYLGEDMLLALVCKSAQIGPNNAEF  901
             KSPPPQSLFLSKKGVFGLVALLGSVASTSLSRVLSEYLGEDMLLALVCKSAQIGPNNAEF
Sbjct  1093  KSPPPQSLFLSKKGVFGLVALLGSVASTSLSRVLSEYLGEDMLLALVCKSAQIGPNNAEF  1152

Query  902   LRLQSNHRFHVLCLDAIRPWKDGLLENDPQKKLAMDDPKLPDGDPIPGFKGYAVNMIDLA  1081
             LRLQSNHRFHVLCLDAIRPWKDGLLENDPQKKLAMDDPKLPDGDPIPGFKGYAVNMIDLA
Sbjct  1153  LRLQSNHRFHVLCLDAIRPWKDGLLENDPQKKLAMDDPKLPDGDPIPGFKGYAVNMIDLA  1212

Query  1082  PEELTVQTYSGYGLRETLFYNLFGNLQV  1165
             PEELTVQTYSGYGLRETLFYNLFGNLQ+
Sbjct  1213  PEELTVQTYSGYGLRETLFYNLFGNLQL  1240


Range 2: 1251 to 1335

Score:147 bits(372), Expect:1e-36,
Method:Compositional matrix adjust.,
Identities:78/87(90%), Positives:78/87(89%), Gaps:3/87(3%)

Query  1237  FIAKDNGFIYS-GCSIPEFHFPITLKEDEEVKLRKMEAARDKVRMAAKKIEEEKCSMRKV  1413
             F A D   IY    SIPEFHFPITLKEDEEVKLRKMEAARDKVRMAAKKIEEEKCSMRKV
Sbjct  1251  FTADD--MIYGFWISIPEFHFPITLKEDEEVKLRKMEAARDKVRMAAKKIEEEKCSMRKV  1308

Query  1414  DKKMKKTNEKYHNVTNSLELIQSQSLE  1494
             DKKMKKTNEKYHNVTNSLELIQSQSLE
Sbjct  1309  DKKMKKTNEKYHNVTNSLELIQSQSLE  1335


>En/Spm-like transposon [Arabidopsis thaliana]
Sequence ID: NP_001331300.1 Length: 602
>En/Spm-like transposon [Arabidopsis thaliana]
Sequence ID: ANM69637.1 Length: 602
Range 1: 215 to 602

Score:777 bits(2007), Expect:0.0,
Method:Compositional matrix adjust.,
Identities:388/388(100%), Positives:388/388(100%), Gaps:0/388(0%)

Query  2     KESQTERRELMLLTKLPECCVAGSNLTNLIFKVTDSDGVMDTSIHHDEKSGCFHTMsiet  181
             KESQTERRELMLLTKLPECCVAGSNLTNLIFKVTDSDGVMDTSIHHDEKSGCFHTMSIET
Sbjct  215   KESQTERRELMLLTKLPECCVAGSNLTNLIFKVTDSDGVMDTSIHHDEKSGCFHTMSIET  274

Query  182   dsssdeseiRYAFVHGSCKVPTLSLPEREGVFSFKVFHSRFPELHLSLKIQLTPAQILQR  361
             DSSSDESEIRYAFVHGSCKVPTLSLPEREGVFSFKVFHSRFPELHLSLKIQLTPAQILQR
Sbjct  275   DSSSDESEIRYAFVHGSCKVPTLSLPEREGVFSFKVFHSRFPELHLSLKIQLTPAQILQR  334

Query  362   DETSYSRMGLTPKSKMASTTYSPALSSQTGPSLRDVAQFTESFKENLIGYSEHRVEIDER  541
             DETSYSRMGLTPKSKMASTTYSPALSSQTGPSLRDVAQFTESFKENLIGYSEHRVEIDER
Sbjct  335   DETSYSRMGLTPKSKMASTTYSPALSSQTGPSLRDVAQFTESFKENLIGYSEHRVEIDER  394

Query  542   LHCLEAEQNQAKEELRTLQASLEPLGAMFPECLSTKESMMKQIEEKHHDTAASVFCCLYR  721
             LHCLEAEQNQAKEELRTLQASLEPLGAMFPECLSTKESMMKQIEEKHHDTAASVFCCLYR
Sbjct  395   LHCLEAEQNQAKEELRTLQASLEPLGAMFPECLSTKESMMKQIEEKHHDTAASVFCCLYR  454

Query  722   KSPPPQSLFLSKKGVFGLVALLGSVASTSLSRVLSEYLGEDMLLALVCKSAQIGPNNAEF  901
             KSPPPQSLFLSKKGVFGLVALLGSVASTSLSRVLSEYLGEDMLLALVCKSAQIGPNNAEF
Sbjct  455   KSPPPQSLFLSKKGVFGLVALLGSVASTSLSRVLSEYLGEDMLLALVCKSAQIGPNNAEF  514

Query  902   LRLQSNHRFHVLCLDAIRPWKDGLLENDPQKKLAMDDPKLPDGDPIPGFKGYAVNMIDLA  1081
             LRLQSNHRFHVLCLDAIRPWKDGLLENDPQKKLAMDDPKLPDGDPIPGFKGYAVNMIDLA
Sbjct  515   LRLQSNHRFHVLCLDAIRPWKDGLLENDPQKKLAMDDPKLPDGDPIPGFKGYAVNMIDLA  574

Query  1082  PEELTVQTYSGYGLRETLFYNLFGNLQV  1165
             PEELTVQTYSGYGLRETLFYNLFGNLQV
Sbjct  575   PEELTVQTYSGYGLRETLFYNLFGNLQV  602


>En/Spm-like transposon [Arabidopsis thaliana]
Sequence ID: NP_001331298.1 Length: 430
>En/Spm-like transposon [Arabidopsis thaliana]
Sequence ID: NP_001331301.1 Length: 430 >En/Spm-like transposon [Arabidopsis thaliana]
Sequence ID: ANM69635.1 Length: 430 >En/Spm-like transposon [Arabidopsis thaliana]
Sequence ID: ANM69638.1 Length: 430
Range 1: 43 to 430

Score:774 bits(1998), Expect:0.0,
Method:Compositional matrix adjust.,
Identities:388/388(100%), Positives:388/388(100%), Gaps:0/388(0%)

Query  2     KESQTERRELMLLTKLPECCVAGSNLTNLIFKVTDSDGVMDTSIHHDEKSGCFHTMsiet  181
             KESQTERRELMLLTKLPECCVAGSNLTNLIFKVTDSDGVMDTSIHHDEKSGCFHTMSIET
Sbjct  43    KESQTERRELMLLTKLPECCVAGSNLTNLIFKVTDSDGVMDTSIHHDEKSGCFHTMSIET  102

Query  182   dsssdeseiRYAFVHGSCKVPTLSLPEREGVFSFKVFHSRFPELHLSLKIQLTPAQILQR  361
             DSSSDESEIRYAFVHGSCKVPTLSLPEREGVFSFKVFHSRFPELHLSLKIQLTPAQILQR
Sbjct  103   DSSSDESEIRYAFVHGSCKVPTLSLPEREGVFSFKVFHSRFPELHLSLKIQLTPAQILQR  162

Query  362   DETSYSRMGLTPKSKMASTTYSPALSSQTGPSLRDVAQFTESFKENLIGYSEHRVEIDER  541
             DETSYSRMGLTPKSKMASTTYSPALSSQTGPSLRDVAQFTESFKENLIGYSEHRVEIDER
Sbjct  163   DETSYSRMGLTPKSKMASTTYSPALSSQTGPSLRDVAQFTESFKENLIGYSEHRVEIDER  222

Query  542   LHCLEAEQNQAKEELRTLQASLEPLGAMFPECLSTKESMMKQIEEKHHDTAASVFCCLYR  721
             LHCLEAEQNQAKEELRTLQASLEPLGAMFPECLSTKESMMKQIEEKHHDTAASVFCCLYR
Sbjct  223   LHCLEAEQNQAKEELRTLQASLEPLGAMFPECLSTKESMMKQIEEKHHDTAASVFCCLYR  282

Query  722   KSPPPQSLFLSKKGVFGLVALLGSVASTSLSRVLSEYLGEDMLLALVCKSAQIGPNNAEF  901
             KSPPPQSLFLSKKGVFGLVALLGSVASTSLSRVLSEYLGEDMLLALVCKSAQIGPNNAEF
Sbjct  283   KSPPPQSLFLSKKGVFGLVALLGSVASTSLSRVLSEYLGEDMLLALVCKSAQIGPNNAEF  342

Query  902   LRLQSNHRFHVLCLDAIRPWKDGLLENDPQKKLAMDDPKLPDGDPIPGFKGYAVNMIDLA  1081
             LRLQSNHRFHVLCLDAIRPWKDGLLENDPQKKLAMDDPKLPDGDPIPGFKGYAVNMIDLA
Sbjct  343   LRLQSNHRFHVLCLDAIRPWKDGLLENDPQKKLAMDDPKLPDGDPIPGFKGYAVNMIDLA  402

Query  1082  PEELTVQTYSGYGLRETLFYNLFGNLQV  1165
             PEELTVQTYSGYGLRETLFYNLFGNLQV
Sbjct  403   PEELTVQTYSGYGLRETLFYNLFGNLQV  430


>hypothetical protein AXX17_AT5G24050 [Arabidopsis thaliana]
Sequence ID: OAO94301.1 Length: 1442
Range 1: 811 to 1198

Score:759 bits(1959), Expect:0.0,
Method:Compositional matrix adjust.,
Identities:379/388(98%), Positives:383/388(98%), Gaps:0/388(0%)

Query  2     KESQTERRELMLLTKLPECCVAGSNLTNLIFKVTDSDGVMDTSIHHDEKSGCFHTMsiet  181
             KESQTERRELMLLTKLPECCVAGSNLTNLIFKVTDSDGVMDTSIHHDEKSGCFHTMSIET
Sbjct  811   KESQTERRELMLLTKLPECCVAGSNLTNLIFKVTDSDGVMDTSIHHDEKSGCFHTMSIET  870

Query  182   dsssdeseiRYAFVHGSCKVPTLSLPEREGVFSFKVFHSRFPELHLSLKIQLTPAQILQR  361
             DSSSDESEIRYAFVHGSCKVPTLSLPE EGVFSFKVFHSRFPELHLSLKIQLTPAQILQR
Sbjct  871   DSSSDESEIRYAFVHGSCKVPTLSLPESEGVFSFKVFHSRFPELHLSLKIQLTPAQILQR  930

Query  362   DETSYSRMGLTPKSKMASTTYSPALSSQTGPSLRDVAQFTESFKENLIGYSEHRVEIDER  541
             DETSYSRMGLTPKSKMASTTYSPALSSQTGPSLRDVAQFTESFKENLIGYSEHRVEIDER
Sbjct  931   DETSYSRMGLTPKSKMASTTYSPALSSQTGPSLRDVAQFTESFKENLIGYSEHRVEIDER  990

Query  542   LHCLEAEQNQAKEELRTLQASLEPLGAMFPECLSTKESMMKQIEEKHHDTAASVFCCLYR  721
             LHCLEAEQNQAKEELRTLQASLEPLGAMFPECLSTKESMMKQIEEKHHDTAASVFCCLYR
Sbjct  991   LHCLEAEQNQAKEELRTLQASLEPLGAMFPECLSTKESMMKQIEEKHHDTAASVFCCLYR  1050

Query  722   KSPPPQSLFLSKKGVFGLVALLGSVASTSLSRVLSEYLGEDMLLALVCKSAQIGPNNAEF  901
             KSPPPQSLFLSKKGVFGLVALLGSVASTSLSRVLSEYLGEDMLLALVCKSAQIGPNNAEF
Sbjct  1051  KSPPPQSLFLSKKGVFGLVALLGSVASTSLSRVLSEYLGEDMLLALVCKSAQIGPNNAEF  1110

Query  902   LRLQSNHRFHVLCLDAIRPWKDGLLENDPQKKLAMDDPKLPDGDPIPGFKGYAVNMIDLA  1081
             LRLQSNHRFHVLCLDAIR WKDG+LENDPQKKLAMD PKLP+G+PI GFKGYAVNMIDLA
Sbjct  1111  LRLQSNHRFHVLCLDAIRAWKDGILENDPQKKLAMDVPKLPNGEPIRGFKGYAVNMIDLA  1170

Query  1082  PEELTVQTYSGYGLRETLFYNLFGNLQV  1165
             PEELT+QT SGYGLRETLFYNLFGNLQV
Sbjct  1171  PEELTIQTNSGYGLRETLFYNLFGNLQV  1198


>unnamed protein product, partial [Arabidopsis thaliana]
Sequence ID: CAA0404532.1 Length: 877
Range 1: 447 to 842

Score:745 bits(1923), Expect:0.0,
Method:Compositional matrix adjust.,
Identities:371/396(94%), Positives:381/396(96%), Gaps:0/396(0%)

Query  2     KESQTERRELMLLTKLPECCVAGSNLTNLIFKVTDSDGVMDTSIHHDEKSGCFHTMsiet  181
             KESQTERRELMLLTKLPECCVAGSNLTNLIFKVTDSDGVMDTSIHHDEKSGCFH MSIET
Sbjct  447   KESQTERRELMLLTKLPECCVAGSNLTNLIFKVTDSDGVMDTSIHHDEKSGCFHIMSIET  506

Query  182   dsssdeseiRYAFVHGSCKVPTLSLPEREGVFSFKVFHSRFPELHLSLKIQLTPAQILQR  361
             DSSSDES IRYAFVHGSCKVPTLSLPEREGVFSFKVFHSRFPELHLSLKIQLTPAQILQR
Sbjct  507   DSSSDESAIRYAFVHGSCKVPTLSLPEREGVFSFKVFHSRFPELHLSLKIQLTPAQILQR  566

Query  362   DETSYSRMGLTPKSKMASTTYSPALSSQTGPSLRDVAQFTESFKENLIGYSEHRVEIDER  541
             DE  YSR+ LTPKSKMAST  SP LSSQTGPSLRDVAQFTESF+ENLIGYSEHRVEIDER
Sbjct  567   DEIPYSRINLTPKSKMASTKNSPPLSSQTGPSLRDVAQFTESFQENLIGYSEHRVEIDER  626

Query  542   LHCLEAEQNQAKEELRTLQASLEPLGAMFPECLSTKESMMKQIEEKHHDTAASVFCCLYR  721
             L+CLEAEQNQAKEELRTLQASLEPLGA FPECLSTKESMMKQIEEKHHDTAASVFCCLYR
Sbjct  627   LNCLEAEQNQAKEELRTLQASLEPLGATFPECLSTKESMMKQIEEKHHDTAASVFCCLYR  686

Query  722   KSPPPQSLFLSKKGVFGLVALLGSVASTSLSRVLSEYLGEDMLLALVCKSAQIGPNNAEF  901
             KSPPPQSLFLSKK VFGLVALLGSVASTSLSRVLSEYLGEDM+LALVCKSAQIGPNNAE+
Sbjct  687   KSPPPQSLFLSKKRVFGLVALLGSVASTSLSRVLSEYLGEDMMLALVCKSAQIGPNNAEY  746

Query  902   LRLQSNHRFHVLCLDAIRPWKDGLLENDPQKKLAMDDPKLPDGDPIPGFKGYAVNMIDLA  1081
             LRLQSNHRFHVLCLDAIRPWKDGLLENDPQKKLAMD P+LPDGDPIPGFKGYAVNMIDLA
Sbjct  747   LRLQSNHRFHVLCLDAIRPWKDGLLENDPQKKLAMDGPELPDGDPIPGFKGYAVNMIDLA  806

Query  1082  PEELTVQTYSGYGLRETLFYNLFGNLQV*RPKSKLK  1189
             PEELT+QTYSGYGLRETLFYNLFGNLQV   + ++K
Sbjct  807   PEELTIQTYSGYGLRETLFYNLFGNLQVYETQKQVK  842


Range 2: 836 to 871

Score:79.7 bits(195), Expect:7e-15,
Method:Compositional matrix adjust.,
Identities:35/36(97%), Positives:36/36(100%), Gaps:0/36(0%)

Query  1168  ETQKQVKAALPHINGGGAVSLDGFIAKDNGFIYSGC  1275
             ETQKQVKAALPHINGGGAVSLDGFIAK+NGFIYSGC
Sbjct  836   ETQKQVKAALPHINGGGAVSLDGFIAKENGFIYSGC  871


>unnamed protein product [Arabidopsis thaliana]
Sequence ID: VYS67775.1 Length: 869
Range 1: 439 to 826

Score:738 bits(1906), Expect:0.0,
Method:Compositional matrix adjust.,
Identities:369/388(95%), Positives:375/388(96%), Gaps:0/388(0%)

Query  2     KESQTERRELMLLTKLPECCVAGSNLTNLIFKVTDSDGVMDTSIHHDEKSGCFHTMsiet  181
             KESQTERRELMLLTKLPECCVAGSNLTNLIFKVTDSDGVMDTSIHHDEKSGCFH MSIET
Sbjct  439   KESQTERRELMLLTKLPECCVAGSNLTNLIFKVTDSDGVMDTSIHHDEKSGCFHIMSIET  498

Query  182   dsssdeseiRYAFVHGSCKVPTLSLPEREGVFSFKVFHSRFPELHLSLKIQLTPAQILQR  361
             DSSSDES IRYAFVHGSCKVPTLSLPEREGVFSFKVFHSRFPELHLSLKIQLTPAQILQR
Sbjct  499   DSSSDESAIRYAFVHGSCKVPTLSLPEREGVFSFKVFHSRFPELHLSLKIQLTPAQILQR  558

Query  362   DETSYSRMGLTPKSKMASTTYSPALSSQTGPSLRDVAQFTESFKENLIGYSEHRVEIDER  541
             DE  YSR+ LTPKSKMAST  SP LSSQTGPSLRDVAQFTESF+ENLIGYSEHRVEIDER
Sbjct  559   DEIPYSRINLTPKSKMASTKNSPPLSSQTGPSLRDVAQFTESFQENLIGYSEHRVEIDER  618

Query  542   LHCLEAEQNQAKEELRTLQASLEPLGAMFPECLSTKESMMKQIEEKHHDTAASVFCCLYR  721
             L+CLEAEQNQAKEELRTLQASLEPLGA FPECLSTKESMMKQIEEKHHDTAASVFCCLYR
Sbjct  619   LNCLEAEQNQAKEELRTLQASLEPLGATFPECLSTKESMMKQIEEKHHDTAASVFCCLYR  678

Query  722   KSPPPQSLFLSKKGVFGLVALLGSVASTSLSRVLSEYLGEDMLLALVCKSAQIGPNNAEF  901
             KSPPPQSLFLSKKGVFGLVALLGSVASTSLSRVLSEYLGEDMLLALVCKSAQIGPNNAEF
Sbjct  679   KSPPPQSLFLSKKGVFGLVALLGSVASTSLSRVLSEYLGEDMLLALVCKSAQIGPNNAEF  738

Query  902   LRLQSNHRFHVLCLDAIRPWKDGLLENDPQKKLAMDDPKLPDGDPIPGFKGYAVNMIDLA  1081
             LRLQSNHRFHVLCLDAIRPWKDGLLENDPQKKLAM DP+LP+GDPIPGFK YAVNMIDL
Sbjct  739   LRLQSNHRFHVLCLDAIRPWKDGLLENDPQKKLAMVDPELPNGDPIPGFKDYAVNMIDLG  798

Query  1082  PEELTVQTYSGYGLRETLFYNLFGNLQV  1165
             PEELT+QT SGYGLRETLFYNLFGNLQV
Sbjct  799   PEELTIQTNSGYGLRETLFYNLFGNLQV  826


Range 2: 828 to 868

Score:70.1 bits(170), Expect:7e-12,
Method:Compositional matrix adjust.,
Identities:32/41(78%), Positives:34/41(82%), Gaps:0/41(0%)

Query  1168  ETQKQVKAALPHINGGGAVSLDGFIAKDNGFIYSGCSIPEF  1290
             ETQKQV+AALPHING GAVSL GFIA +NGFIYSGC    F
Sbjct  828   ETQKQVEAALPHINGCGAVSLHGFIAIENGFIYSGCRYLNF  868


>unnamed protein product [Arabidopsis thaliana]
Sequence ID: BAB10394.1 Length: 1634
Range 1: 1116 to 1531

Score:499 bits(1285), Expect:3e-169,
Method:Compositional matrix adjust.,
Identities:271/422(64%), Positives:326/422(77%), Gaps:40/422(9%)

Query  2     KESQTERRELMLLTKLPECCVAGSNLTNLIFKVTDSDGVMDTSIHHDEKSGCFHTMsiet  181
             KESQ + R+L L+T+LP+CC AG+NL NLIF+VT+ DG +DTSIHHDEKSGCFHTMSIE+
Sbjct  1116  KESQIDERQLRLVTELPDCCTAGTNLMNLIFQVTELDGSLDTSIHHDEKSGCFHTMSIES  1175

Query  182   dsssdeseiRYAFVHGSCKVPTLSLPEREGVFSFKVFHSRFPELHLSLKIQLTPAQILQR  361
             DSSS ES IRYAFVHGSCKV +LSLPE EGVFS +VFHSR+PEL +S+KIQ+T A   +R
Sbjct  1176  DSSSVESAIRYAFVHGSCKVSSLSLPENEGVFSCRVFHSRYPELQMSIKIQVTSAPTSER  1235

Query  362   DETSYSRMGLTPKSK-------------------------MASTTYSPALSSQTGPSLRD  466
             +E+ YS    TP SK                         +A  + S ALSS+T  SL D
Sbjct  1236  EESGYS----TPHSKTTPPPESGIPSITNPWPTPCSQFGVLAIRSSSLALSSET--SLMD  1289

Query  467   VAQFTESFKENLIGYSEHRVEIDERLHCLEAEQNQAKEELRTLQASLEPLGAMFPECLST  646
             +AQ+TE  KE +    E RVE++ERL CL+A++  A++E   LQASLEPLGA FPECLST
Sbjct  1290  MAQYTEDLKEKINIDEERRVELEERLKCLQAQREHAEQECSRLQASLEPLGAPFPECLST  1349

Query  647   KESMMKQIEEKHHDTAASVFCCLYRKSPPPQSLFLSKKGVFGLVALLGSVASTSLSRVLS  826
             KESMMKQIEEKHHDTAASVFCCLYRK+PPP+SLFLS+KG+FG+VALLGSVASTSLSRVLS
Sbjct  1350  KESMMKQIEEKHHDTAASVFCCLYRKAPPPRSLFLSQKGMFGVVALLGSVASTSLSRVLS  1409

Query  827   EYLGEDMLLALVCKSAQIGPNNAEFLRLQS---------NHRFHVLCLDAIRPWKDGLLE  979
             EYLG+D +L+LVCKS+Q GP + E+ + QS          +RF V+CLDA RPW++GL+
Sbjct  1410  EYLGKDTMLSLVCKSSQFGPKSDEYRKFQSEAASLGRSITNRFLVICLDATRPWRNGLVR  1469

Query  980   NDPQKKLAMDDPKLPDGDPIPGFKGYAVNMIDLAPEELTVQTYSGYGLRETLFYNLFGNL  1159
             NDPQK+LAMD+P LP+GDPIPGFKGYAVNMIDLA EEL +Q+ SGYGLRETLFY +F  L
Sbjct  1470  NDPQKRLAMDNPYLPNGDPIPGFKGYAVNMIDLASEELDIQSSSGYGLRETLFYGVFREL  1529

Query  1160  QV  1165
             QV
Sbjct  1530  QV  1531


Range 2: 1528 to 1629

Score:117 bits(294), Expect:3e-169,
Method:Composition-based stats.,
Identities:57/102(56%), Positives:72/102(70%), Gaps:0/102(0%)

Query  1153  EFTGIETQKQVKAALPHINGGGAVSLDGFIAKDNGFIYSGCSIPEFHFPITLKEDEEVKL  1332
             E    ET + ++AALPHINGG AVSLDG IA++NGFIYSGC  PE HFPIT+ E +E  L
Sbjct  1528  ELQVYETAEHLEAALPHINGGDAVSLDGVIARENGFIYSGCCTPEVHFPITVTERQEKAL  1587

Query  1333  RKMEAARDKVRMAAKKIEEEKCSMRKVDKKMKKTNEKYHNVT  1458
              ++E  RDK R   + + EE  S+R++ KK+KK NEKY N T
Sbjct  1588  VQLEITRDKKRKTEEMMTEENRSLRRLVKKLKKANEKYQNFT  1629


>gamma-irradiation and mitomycin c induced 1 [Arabidopsis thaliana]
Sequence ID: NP_001330245.1 Length: 1566
>gamma-irradiation and mitomycin c induced 1 [Arabidopsis thaliana]
Sequence ID: NP_001330246.1 Length: 1566 >gamma-irradiation and mitomycin c induced 1 [Arabidopsis thaliana]
Sequence ID: ANM68491.1 Length: 1566 >gamma-irradiation and mitomycin c induced 1 [Arabidopsis thaliana]
Sequence ID: ANM68492.1 Length: 1566
Range 1: 1048 to 1463

Score:499 bits(1286), Expect:5e-169,
Method:Compositional matrix adjust.,
Identities:271/422(64%), Positives:326/422(77%), Gaps:40/422(9%)

Query  2     KESQTERRELMLLTKLPECCVAGSNLTNLIFKVTDSDGVMDTSIHHDEKSGCFHTMsiet  181
             KESQ + R+L L+T+LP+CC AG+NL NLIF+VT+ DG +DTSIHHDEKSGCFHTMSIE+
Sbjct  1048  KESQIDERQLRLVTELPDCCTAGTNLMNLIFQVTELDGSLDTSIHHDEKSGCFHTMSIES  1107

Query  182   dsssdeseiRYAFVHGSCKVPTLSLPEREGVFSFKVFHSRFPELHLSLKIQLTPAQILQR  361
             DSSS ES IRYAFVHGSCKV +LSLPE EGVFS +VFHSR+PEL +S+KIQ+T A   +R
Sbjct  1108  DSSSVESAIRYAFVHGSCKVSSLSLPENEGVFSCRVFHSRYPELQMSIKIQVTSAPTSER  1167

Query  362   DETSYSRMGLTPKSK-------------------------MASTTYSPALSSQTGPSLRD  466
             +E+ YS    TP SK                         +A  + S ALSS+T  SL D
Sbjct  1168  EESGYS----TPHSKTTPPPESGIPSITNPWPTPCSQFGVLAIRSSSLALSSET--SLMD  1221

Query  467   VAQFTESFKENLIGYSEHRVEIDERLHCLEAEQNQAKEELRTLQASLEPLGAMFPECLST  646
             +AQ+TE  KE +    E RVE++ERL CL+A++  A++E   LQASLEPLGA FPECLST
Sbjct  1222  MAQYTEDLKEKINIDEERRVELEERLKCLQAQREHAEQECSRLQASLEPLGAPFPECLST  1281

Query  647   KESMMKQIEEKHHDTAASVFCCLYRKSPPPQSLFLSKKGVFGLVALLGSVASTSLSRVLS  826
             KESMMKQIEEKHHDTAASVFCCLYRK+PPP+SLFLS+KG+FG+VALLGSVASTSLSRVLS
Sbjct  1282  KESMMKQIEEKHHDTAASVFCCLYRKAPPPRSLFLSQKGMFGVVALLGSVASTSLSRVLS  1341

Query  827   EYLGEDMLLALVCKSAQIGPNNAEFLRLQS---------NHRFHVLCLDAIRPWKDGLLE  979
             EYLG+D +L+LVCKS+Q GP + E+ + QS          +RF V+CLDA RPW++GL+
Sbjct  1342  EYLGKDTMLSLVCKSSQFGPKSDEYRKFQSEAASLGRSITNRFLVICLDATRPWRNGLVR  1401

Query  980   NDPQKKLAMDDPKLPDGDPIPGFKGYAVNMIDLAPEELTVQTYSGYGLRETLFYNLFGNL  1159
             NDPQK+LAMD+P LP+GDPIPGFKGYAVNMIDLA EEL +Q+ SGYGLRETLFY +F  L
Sbjct  1402  NDPQKRLAMDNPYLPNGDPIPGFKGYAVNMIDLASEELDIQSSSGYGLRETLFYGVFREL  1461

Query  1160  QV  1165
             QV
Sbjct  1462  QV  1463


Range 2: 1460 to 1561

Score:117 bits(292), Expect:5e-169,
Method:Composition-based stats.,
Identities:57/102(56%), Positives:72/102(70%), Gaps:0/102(0%)

Query  1153  EFTGIETQKQVKAALPHINGGGAVSLDGFIAKDNGFIYSGCSIPEFHFPITLKEDEEVKL  1332
             E    ET + ++AALPHINGG AVSLDG IA++NGFIYSGC  PE HFPIT+ E +E  L
Sbjct  1460  ELQVYETAEHLEAALPHINGGDAVSLDGVIARENGFIYSGCCTPEVHFPITVTERQEKAL  1519

Query  1333  RKMEAARDKVRMAAKKIEEEKCSMRKVDKKMKKTNEKYHNVT  1458
              ++E  RDK R   + + EE  S+R++ KK+KK NEKY N T
Sbjct  1520  VQLEITRDKKRKTEEMMTEENRSLRRLVKKLKKANEKYQNFT  1561


>gamma-irradiation and mitomycin c induced 1 [Arabidopsis thaliana]
Sequence ID: NP_197816.3 Length: 1598
>RecName: Full=Structural maintenance of chromosomes flexible hinge domain-containing protein GMI1; Short=SMC hinge domain-containing protein GMI1; AltName: Full=Protein GAMMA-IRRADIATION AND MITOMYCIN C INDUCED 1 [Arabidopsis thaliana]
Sequence ID: F4KFS5.1 Length: 1598 >gamma-irradiation and mitomycin c induced 1 [Arabidopsis thaliana]
Sequence ID: AED93279.1 Length: 1598 >unnamed protein product [Arabidopsis thaliana]
Sequence ID: VYS67776.1 Length: 1598
Range 1: 1080 to 1495

Score:499 bits(1284), Expect:5e-169,
Method:Compositional matrix adjust.,
Identities:271/422(64%), Positives:326/422(77%), Gaps:40/422(9%)

Query  2     KESQTERRELMLLTKLPECCVAGSNLTNLIFKVTDSDGVMDTSIHHDEKSGCFHTMsiet  181
             KESQ + R+L L+T+LP+CC AG+NL NLIF+VT+ DG +DTSIHHDEKSGCFHTMSIE+
Sbjct  1080  KESQIDERQLRLVTELPDCCTAGTNLMNLIFQVTELDGSLDTSIHHDEKSGCFHTMSIES  1139

Query  182   dsssdeseiRYAFVHGSCKVPTLSLPEREGVFSFKVFHSRFPELHLSLKIQLTPAQILQR  361
             DSSS ES IRYAFVHGSCKV +LSLPE EGVFS +VFHSR+PEL +S+KIQ+T A   +R
Sbjct  1140  DSSSVESAIRYAFVHGSCKVSSLSLPENEGVFSCRVFHSRYPELQMSIKIQVTSAPTSER  1199

Query  362   DETSYSRMGLTPKSK-------------------------MASTTYSPALSSQTGPSLRD  466
             +E+ YS    TP SK                         +A  + S ALSS+T  SL D
Sbjct  1200  EESGYS----TPHSKTTPPPESGIPSITNPWPTPCSQFGVLAIRSSSLALSSET--SLMD  1253

Query  467   VAQFTESFKENLIGYSEHRVEIDERLHCLEAEQNQAKEELRTLQASLEPLGAMFPECLST  646
             +AQ+TE  KE +    E RVE++ERL CL+A++  A++E   LQASLEPLGA FPECLST
Sbjct  1254  MAQYTEDLKEKINIDEERRVELEERLKCLQAQREHAEQECSRLQASLEPLGAPFPECLST  1313

Query  647   KESMMKQIEEKHHDTAASVFCCLYRKSPPPQSLFLSKKGVFGLVALLGSVASTSLSRVLS  826
             KESMMKQIEEKHHDTAASVFCCLYRK+PPP+SLFLS+KG+FG+VALLGSVASTSLSRVLS
Sbjct  1314  KESMMKQIEEKHHDTAASVFCCLYRKAPPPRSLFLSQKGMFGVVALLGSVASTSLSRVLS  1373

Query  827   EYLGEDMLLALVCKSAQIGPNNAEFLRLQS---------NHRFHVLCLDAIRPWKDGLLE  979
             EYLG+D +L+LVCKS+Q GP + E+ + QS          +RF V+CLDA RPW++GL+
Sbjct  1374  EYLGKDTMLSLVCKSSQFGPKSDEYRKFQSEAASLGRSITNRFLVICLDATRPWRNGLVR  1433

Query  980   NDPQKKLAMDDPKLPDGDPIPGFKGYAVNMIDLAPEELTVQTYSGYGLRETLFYNLFGNL  1159
             NDPQK+LAMD+P LP+GDPIPGFKGYAVNMIDLA EEL +Q+ SGYGLRETLFY +F  L
Sbjct  1434  NDPQKRLAMDNPYLPNGDPIPGFKGYAVNMIDLASEELDIQSSSGYGLRETLFYGVFREL  1493

Query  1160  QV  1165
             QV
Sbjct  1494  QV  1495


Range 2: 1492 to 1593

Score:117 bits(293), Expect:5e-169,
Method:Composition-based stats.,
Identities:57/102(56%), Positives:72/102(70%), Gaps:0/102(0%)

Query  1153  EFTGIETQKQVKAALPHINGGGAVSLDGFIAKDNGFIYSGCSIPEFHFPITLKEDEEVKL  1332
             E    ET + ++AALPHINGG AVSLDG IA++NGFIYSGC  PE HFPIT+ E +E  L
Sbjct  1492  ELQVYETAEHLEAALPHINGGDAVSLDGVIARENGFIYSGCCTPEVHFPITVTERQEKAL  1551

Query  1333  RKMEAARDKVRMAAKKIEEEKCSMRKVDKKMKKTNEKYHNVT  1458
              ++E  RDK R   + + EE  S+R++ KK+KK NEKY N T
Sbjct  1552  VQLEITRDKKRKTEEMMTEENRSLRRLVKKLKKANEKYQNFT  1593


>GMI1 [Arabidopsis thaliana]
Sequence ID: OAO93531.1 Length: 1548
Range 1: 1030 to 1445

Score:501 bits(1290), Expect:2e-168,
Method:Compositional matrix adjust.,
Identities:272/422(64%), Positives:327/422(77%), Gaps:40/422(9%)

Query  2     KESQTERRELMLLTKLPECCVAGSNLTNLIFKVTDSDGVMDTSIHHDEKSGCFHTMsiet  181
             KESQ + R+L L+T+LP+CC AG+NL NLIF+VT+ DG +DTSIHHDEKSGCFHTMSIE+
Sbjct  1030  KESQIDERQLRLVTELPDCCTAGTNLMNLIFQVTELDGSLDTSIHHDEKSGCFHTMSIES  1089

Query  182   dsssdeseiRYAFVHGSCKVPTLSLPEREGVFSFKVFHSRFPELHLSLKIQLTPAQILQR  361
             DSSS ES IRYAFVHGSCKV +LSLPE EGVFS +VFHSR+PEL +S+KIQ+T A   +R
Sbjct  1090  DSSSVESAIRYAFVHGSCKVSSLSLPENEGVFSCRVFHSRYPELQMSIKIQVTSAPTSER  1149

Query  362   DETSYSRMGLTPKSK-------------------------MASTTYSPALSSQTGPSLRD  466
             +E+ YS    TP SK                         +A  + S ALSS+T  SL D
Sbjct  1150  EESGYS----TPHSKTTPPPESGIPSITNPWPTPCSQFGVLAIRSSSLALSSET--SLMD  1203

Query  467   VAQFTESFKENLIGYSEHRVEIDERLHCLEAEQNQAKEELRTLQASLEPLGAMFPECLST  646
             +AQ+TE  KE +    E RVE++ERL CL+A++  A++E   LQASLEPLGA FPECLST
Sbjct  1204  MAQYTEDLKEKINIDEERRVELEERLKCLQAQREHAEQECSRLQASLEPLGAPFPECLST  1263

Query  647   KESMMKQIEEKHHDTAASVFCCLYRKSPPPQSLFLSKKGVFGLVALLGSVASTSLSRVLS  826
             KESMMKQIEEKHHDTAASVFCCLYRK+PPP+SLFLS+KG+FG+VALLGSVASTSLSRVLS
Sbjct  1264  KESMMKQIEEKHHDTAASVFCCLYRKAPPPRSLFLSQKGMFGVVALLGSVASTSLSRVLS  1323

Query  827   EYLGEDMLLALVCKSAQIGPNNAEFLRLQS---------NHRFHVLCLDAIRPWKDGLLE  979
             EYLG+D +L+LVCKS+Q GP + E+ +LQS          +RF V+CLDA RPW++GL+
Sbjct  1324  EYLGKDTMLSLVCKSSQFGPKSDEYRKLQSEAASLGRSITNRFLVICLDATRPWRNGLVR  1383

Query  980   NDPQKKLAMDDPKLPDGDPIPGFKGYAVNMIDLAPEELTVQTYSGYGLRETLFYNLFGNL  1159
             NDPQK+LAMD+P LP+GDPIPGFKGYAVNMIDLA EEL +Q+ SGYGLRETLFY +F  L
Sbjct  1384  NDPQKRLAMDNPYLPNGDPIPGFKGYAVNMIDLASEELDIQSSSGYGLRETLFYGVFREL  1443

Query  1160  QV  1165
             QV
Sbjct  1444  QV  1445


Range 2: 1442 to 1543

Score:113 bits(283), Expect:2e-168,
Method:Composition-based stats.,
Identities:55/102(54%), Positives:72/102(70%), Gaps:0/102(0%)

Query  1153  EFTGIETQKQVKAALPHINGGGAVSLDGFIAKDNGFIYSGCSIPEFHFPITLKEDEEVKL  1332
             E    ET + ++AALPHINGG AVSLDG IA++NGFIYSGC  PE HFPIT+ E +E  L
Sbjct  1442  ELQVYETAEHLEAALPHINGGDAVSLDGVIARENGFIYSGCCTPEVHFPITVTERQEKAL  1501

Query  1333  RKMEAARDKVRMAAKKIEEEKCSMRKVDKKMKKTNEKYHNVT  1458
              ++E  +DK R   + + EE  S+R++ KK+KK NEKY + T
Sbjct  1502  VQLEITKDKKRKTEEMMTEENRSLRRLVKKLKKANEKYQHFT  1543


>unnamed protein product [Arabidopsis thaliana]
Sequence ID: CAA0404534.1 Length: 1566
Range 1: 1048 to 1463

Score:497 bits(1279), Expect:1e-167,
Method:Compositional matrix adjust.,
Identities:270/422(64%), Positives:324/422(76%), Gaps:40/422(9%)

Query  2     KESQTERRELMLLTKLPECCVAGSNLTNLIFKVTDSDGVMDTSIHHDEKSGCFHTMsiet  181
             KESQ + R+L L+T+LP+CC AG+NL NLIF+VT+ DG +DTSIHHDEKSGCFHTMSIE+
Sbjct  1048  KESQIDERQLRLVTELPDCCTAGTNLMNLIFQVTELDGSLDTSIHHDEKSGCFHTMSIES  1107

Query  182   dsssdeseiRYAFVHGSCKVPTLSLPEREGVFSFKVFHSRFPELHLSLKIQLTPAQILQR  361
             DSSS ES IRYAFVHGSCKV +LSLPE EGVFS +VFHSR+PEL +S+KIQ+T A   +R
Sbjct  1108  DSSSVESAIRYAFVHGSCKVSSLSLPENEGVFSCRVFHSRYPELQMSIKIQVTSAPTSER  1167

Query  362   DETSYSRMGLTPKSK-------------------------MASTTYSPALSSQTGPSLRD  466
             +E+ YS    TP SK                         +A  + S ALSS+T  SL D
Sbjct  1168  EESGYS----TPHSKTTPPPESGIPSITNPWPTPCSQFGVLAIRSSSLALSSET--SLMD  1221

Query  467   VAQFTESFKENLIGYSEHRVEIDERLHCLEAEQNQAKEELRTLQASLEPLGAMFPECLST  646
             +AQ+TE  KE +    E RVE++ERL CL+A+   A++E   LQASLEPLGA FPECLST
Sbjct  1222  MAQYTEDLKEKINIDEERRVELEERLKCLQAQSEHAEQECSRLQASLEPLGAPFPECLST  1281

Query  647   KESMMKQIEEKHHDTAASVFCCLYRKSPPPQSLFLSKKGVFGLVALLGSVASTSLSRVLS  826
             KESMMKQIEEKHHDTAASVFCCLYRK+PPP+SLFLS+KG+FG+VALLGS ASTSLSRVLS
Sbjct  1282  KESMMKQIEEKHHDTAASVFCCLYRKAPPPRSLFLSQKGMFGVVALLGSFASTSLSRVLS  1341

Query  827   EYLGEDMLLALVCKSAQIGPNNAEFLRLQS---------NHRFHVLCLDAIRPWKDGLLE  979
             EYLG+D +L+LVCKS+Q GP + E+ + QS          +RF V+CLDA RPW++GL+
Sbjct  1342  EYLGKDTMLSLVCKSSQFGPKSDEYRKFQSEAASLGRSITNRFLVICLDATRPWRNGLVR  1401

Query  980   NDPQKKLAMDDPKLPDGDPIPGFKGYAVNMIDLAPEELTVQTYSGYGLRETLFYNLFGNL  1159
             NDPQK+LAMD+P LP+GDPIPGFKGYAVNMIDLA EEL +Q+ SGYGLRETLFY +F  L
Sbjct  1402  NDPQKRLAMDNPYLPNGDPIPGFKGYAVNMIDLASEELDIQSSSGYGLRETLFYGVFREL  1461

Query  1160  QV  1165
             QV
Sbjct  1462  QV  1463


Range 2: 1460 to 1561

Score:114 bits(286), Expect:1e-167,
Method:Composition-based stats.,
Identities:56/102(55%), Positives:72/102(70%), Gaps:0/102(0%)

Query  1153  EFTGIETQKQVKAALPHINGGGAVSLDGFIAKDNGFIYSGCSIPEFHFPITLKEDEEVKL  1332
             E    +T + ++AALPHINGG AVSLDG IA++NGFIYSGC  PE HFPIT+ E +E  L
Sbjct  1460  ELQVYDTAEYLEAALPHINGGDAVSLDGVIARENGFIYSGCCTPEVHFPITVTERQEKAL  1519

Query  1333  RKMEAARDKVRMAAKKIEEEKCSMRKVDKKMKKTNEKYHNVT  1458
              ++E  RDK R   + + EE  S+R++ KK+KK NEKY N T
Sbjct  1520  VQLEITRDKKRKTEEMMTEENRSLRRLVKKLKKANEKYQNFT  1561


>hypothetical protein AXX17_AT5G24030 [Arabidopsis thaliana]
Sequence ID: OAO92751.1 Length: 457
Range 1: 153 to 457

Score:455 bits(1171), Expect:2e-156,
Method:Compositional matrix adjust.,
Identities:249/307(81%), Positives:257/307(83%), Gaps:28/307(9%)

Query  2    KESQTERRELMLLTKLPECCVAGSNLTNLIFKVTDSDGVMDTSIHHDEKSGCFHTMsiet  181
            KE+  ERREL LLTKLP+CC AGSNLTNLIFKVTDSDG MDT IHHDEKSGCFHTMSIET
Sbjct  153  KETLIERRELRLLTKLPDCCAAGSNLTNLIFKVTDSDGAMDTRIHHDEKSGCFHTMSIET  212

Query  182  dsssdeseiRYAFVHGSCKVPTLSLPEREGVFSFKVFHSRFPELHLSLKIQLTPAQILQR  361
            DSSSDES IRYAFVHGSCKVPTLSLPE EGV SFKVFHSRFPELHLSLKIQLTPAQ  +R
Sbjct  213  DSSSDESAIRYAFVHGSCKVPTLSLPESEGVLSFKVFHSRFPELHLSLKIQLTPAQTFER  272

Query  362  DE----TSYSRMGLTPKSKMASTTYSPA----------------------LSSQTGPSLR  463
            DE    T YSRM LTP+SKMASTT SP                       LSSQTG  L
Sbjct  273  DEIGCSTPYSRMSLTPQSKMASTTNSPVASTEQTPCSQFRVLAIKASPSTLSSQTG--LI  330

Query  464  DVAQFTESFKENLIGYSEHRVEIDERLHCLEAEQNQAKEELRTLQASLEPLGAMFPECLS  643
            D+AQFTES KE LI YSEHRVEIDERL CLEAEQNQAKEELRTLQASLEPLGA FPECLS
Sbjct  331  DMAQFTESLKEKLIIYSEHRVEIDERLKCLEAEQNQAKEELRTLQASLEPLGATFPECLS  390

Query  644  TKESMMKQIEEKHHDTAASVFCCLYRKSPPPQSLFLSKKGVFGLVALLGSVASTSLSRVL  823
            TKESMMKQIEEKHH TAASVFCCLYRK+PPP+SLFLSKKGVFGLVALLGSV+STSLSRVL
Sbjct  391  TKESMMKQIEEKHHITAASVFCCLYRKAPPPKSLFLSKKGVFGLVALLGSVSSTSLSRVL  450

Query  824  SEYLGED  844
            SEYLGED
Sbjct  451  SEYLGED  457


>At5g24280, partial [Arabidopsis thaliana]
Sequence ID: AAZ74706.1 Length: 162
>At5g24280, partial [Arabidopsis thaliana]
Sequence ID: AAZ74707.1 Length: 162 >At5g24280, partial [Arabidopsis thaliana]
Sequence ID: AAZ74708.1 Length: 162 >At5g24280, partial [Arabidopsis thaliana]
Sequence ID: AAZ74711.1 Length: 162
Range 1: 1 to 125

Score:176 bits(445), Expect:7e-56,
Method:Compositional matrix adjust.,
Identities:82/125(66%), Positives:100/125(80%), Gaps:9/125(7%)

Query  818   VLSEYLGEDMLLALVCKSAQIGPNNAEFLRLQS---------NHRFHVLCLDAIRPWKDG  970
             VLSEYLG+D +L+LVCKS+Q GP + E+ +LQS          +RF V+CLDA RPW++G
Sbjct  1     VLSEYLGKDTMLSLVCKSSQFGPKSDEYRKLQSEAASLGRSITNRFLVICLDATRPWRNG  60

Query  971   LLENDPQKKLAMDDPKLPDGDPIPGFKGYAVNMIDLAPEELTVQTYSGYGLRETLFYNLF  1150
             L+ NDPQK+LAMD+P LP+GDPIPGFKGYAVNMIDLA EEL +Q+ SGYGLRETLFY +F
Sbjct  61    LVRNDPQKRLAMDNPYLPNGDPIPGFKGYAVNMIDLASEELDIQSSSGYGLRETLFYGVF  120

Query  1151  GNLQV  1165
               LQV
Sbjct  121   RELQV  125


Range 2: 122 to 162

Score:63.5 bits(153), Expect:7e-56,
Method:Compositional matrix adjust.,
Identities:28/41(68%), Positives:33/41(80%), Gaps:0/41(0%)

Query  1153  EFTGIETQKQVKAALPHINGGGAVSLDGFIAKDNGFIYSGC  1275
             E    ET + ++AALPHINGG AVSLDG IA++NGFIYSGC
Sbjct  122   ELQVYETAEHLEAALPHINGGDAVSLDGVIARENGFIYSGC  162


>At5g24280, partial [Arabidopsis thaliana]
Sequence ID: AAZ74709.1 Length: 162
>At5g24280, partial [Arabidopsis thaliana]
Sequence ID: AAZ74714.1 Length: 162 >At5g24280, partial [Arabidopsis thaliana]
Sequence ID: AAZ74716.1 Length: 162
Range 1: 1 to 125

Score:174 bits(441), Expect:2e-55,
Method:Compositional matrix adjust.,
Identities:81/125(65%), Positives:99/125(79%), Gaps:9/125(7%)

Query  818   VLSEYLGEDMLLALVCKSAQIGPNNAEFLRLQS---------NHRFHVLCLDAIRPWKDG  970
             VLSEYLG+D +L+LVCKS+Q GP + E+ + QS          +RF V+CLDA RPW++G
Sbjct  1     VLSEYLGKDTMLSLVCKSSQFGPKSDEYRKFQSEAASLGRSITNRFLVICLDATRPWRNG  60

Query  971   LLENDPQKKLAMDDPKLPDGDPIPGFKGYAVNMIDLAPEELTVQTYSGYGLRETLFYNLF  1150
             L+ NDPQK+LAMD+P LP+GDPIPGFKGYAVNMIDLA EEL +Q+ SGYGLRETLFY +F
Sbjct  61    LVRNDPQKRLAMDNPYLPNGDPIPGFKGYAVNMIDLASEELDIQSSSGYGLRETLFYGVF  120

Query  1151  GNLQV  1165
               LQV
Sbjct  121   RELQV  125


Range 2: 122 to 162

Score:63.5 bits(153), Expect:2e-55,
Method:Compositional matrix adjust.,
Identities:28/41(68%), Positives:33/41(80%), Gaps:0/41(0%)

Query  1153  EFTGIETQKQVKAALPHINGGGAVSLDGFIAKDNGFIYSGC  1275
             E    ET + ++AALPHINGG AVSLDG IA++NGFIYSGC
Sbjct  122   ELQVYETAEHLEAALPHINGGDAVSLDGVIARENGFIYSGC  162


>At5g24280, partial [Arabidopsis thaliana]
Sequence ID: AAZ74705.1 Length: 162
>At5g24280, partial [Arabidopsis thaliana]
Sequence ID: AAZ74712.1 Length: 162 >At5g24280, partial [Arabidopsis thaliana]
Sequence ID: AAZ74713.1 Length: 162
Range 1: 1 to 125

Score:174 bits(440), Expect:7e-55,
Method:Compositional matrix adjust.,
Identities:81/125(65%), Positives:99/125(79%), Gaps:9/125(7%)

Query  818   VLSEYLGEDMLLALVCKSAQIGPNNAEFLRLQS---------NHRFHVLCLDAIRPWKDG  970
             VLSEYLG+D +L+LVCKS+Q GP + E+ + QS          +RF V+CLDA RPW++G
Sbjct  1     VLSEYLGKDTMLSLVCKSSQFGPKSDEYRKFQSEAASLGRSITNRFLVICLDATRPWRNG  60

Query  971   LLENDPQKKLAMDDPKLPDGDPIPGFKGYAVNMIDLAPEELTVQTYSGYGLRETLFYNLF  1150
             L+ NDPQK+LAMD+P LP+GDPIPGFKGYAVNMIDLA EEL +Q+ SGYGLRETLFY +F
Sbjct  61    LVRNDPQKRLAMDNPYLPNGDPIPGFKGYAVNMIDLASEELDIQSSSGYGLRETLFYGVF  120

Query  1151  GNLQV  1165
               LQV
Sbjct  121   RELQV  125


Range 2: 122 to 162

Score:62.0 bits(149), Expect:7e-55,
Method:Compositional matrix adjust.,
Identities:27/41(66%), Positives:33/41(80%), Gaps:0/41(0%)

Query  1153  EFTGIETQKQVKAALPHINGGGAVSLDGFIAKDNGFIYSGC  1275
             E    +T + ++AALPHINGG AVSLDG IA++NGFIYSGC
Sbjct  122   ELQVYDTAEHLEAALPHINGGDAVSLDGVIARENGFIYSGC  162


>At5g24280, partial [Arabidopsis thaliana]
Sequence ID: AAZ74715.1 Length: 162
Range 1: 1 to 125

Score:174 bits(440), Expect:8e-55,
Method:Compositional matrix adjust.,
Identities:81/125(65%), Positives:99/125(79%), Gaps:9/125(7%)

Query  818   VLSEYLGEDMLLALVCKSAQIGPNNAEFLRLQS---------NHRFHVLCLDAIRPWKDG  970
             VLSEYLG+D +L+LVCKS+Q GP + E+ + QS          +RF V+CLDA RPW++G
Sbjct  1     VLSEYLGKDTMLSLVCKSSQFGPKSDEYRKFQSEAASLGRSITNRFLVICLDATRPWRNG  60

Query  971   LLENDPQKKLAMDDPKLPDGDPIPGFKGYAVNMIDLAPEELTVQTYSGYGLRETLFYNLF  1150
             L+ NDPQK+LAMD+P LP+GDPIPGFKGYAVNMIDLA EEL +Q+ SGYGLRETLFY +F
Sbjct  61    LVRNDPQKRLAMDNPYLPNGDPIPGFKGYAVNMIDLASEELDIQSSSGYGLRETLFYGVF  120

Query  1151  GNLQV  1165
               LQV
Sbjct  121   RELQV  125


Range 2: 122 to 162

Score:61.6 bits(148), Expect:8e-55,
Method:Compositional matrix adjust.,
Identities:27/41(66%), Positives:33/41(80%), Gaps:0/41(0%)

Query  1153  EFTGIETQKQVKAALPHINGGGAVSLDGFIAKDNGFIYSGC  1275
             E    +T + ++AALPHINGG AVSLDG IA++NGFIYSGC
Sbjct  122   ELQVYDTAEYLEAALPHINGGDAVSLDGVIARENGFIYSGC  162


>At5g24280, partial [Arabidopsis thaliana]
Sequence ID: AAZ74701.1 Length: 162
>At5g24280, partial [Arabidopsis thaliana]
Sequence ID: AAZ74702.1 Length: 162 >At5g24280, partial [Arabidopsis thaliana]
Sequence ID: AAZ74704.1 Length: 162 >At5g24280, partial [Arabidopsis thaliana]
Sequence ID: AAZ74710.1 Length: 162
Range 1: 1 to 125

Score:170 bits(431), Expect:3e-54,
Method:Compositional matrix adjust.,
Identities:80/125(64%), Positives:98/125(78%), Gaps:9/125(7%)

Query  818   VLSEYLGEDMLLALVCKSAQIGPNNAEFLRLQS---------NHRFHVLCLDAIRPWKDG  970
             VLSEYLG+D +L+LVCKS+Q GP + E+ + QS          +RF V+CLDA RPW++G
Sbjct  1     VLSEYLGKDTMLSLVCKSSQFGPKSDEYRKFQSEAASLGRSITNRFLVICLDATRPWRNG  60

Query  971   LLENDPQKKLAMDDPKLPDGDPIPGFKGYAVNMIDLAPEELTVQTYSGYGLRETLFYNLF  1150
             L+ NDPQK+LAMD+P L +GDPIPGFKGYAVNMIDLA EEL +Q+ SGYGLRETLFY +F
Sbjct  61    LVRNDPQKRLAMDNPYLLNGDPIPGFKGYAVNMIDLASEELDIQSSSGYGLRETLFYGVF  120

Query  1151  GNLQV  1165
               LQV
Sbjct  121   RELQV  125


Range 2: 127 to 162

Score:63.5 bits(153), Expect:3e-54,
Method:Compositional matrix adjust.,
Identities:27/36(75%), Positives:32/36(88%), Gaps:0/36(0%)

Query  1168  ETQKQVKAALPHINGGGAVSLDGFIAKDNGFIYSGC  1275
             ET + ++AALPHINGG AVSLDG IA++NGFIYSGC
Sbjct  127   ETAEHLEAALPHINGGDAVSLDGVIARENGFIYSGC  162


>At5g24280, partial [Arabidopsis thaliana]
Sequence ID: AAZ74703.1 Length: 162
Range 1: 1 to 125

Score:170 bits(430), Expect:4e-54,
Method:Compositional matrix adjust.,
Identities:80/125(64%), Positives:97/125(77%), Gaps:9/125(7%)

Query  818   VLSEYLGEDMLLALVCKSAQIGPNNAEFLRLQS---------NHRFHVLCLDAIRPWKDG  970
             VLSEYLG+D +L+LVCKS Q GP + E+ + QS          +RF V+CLDA RPW++G
Sbjct  1     VLSEYLGKDTMLSLVCKSXQFGPKSDEYRKFQSEAASLGRSITNRFLVICLDATRPWRNG  60

Query  971   LLENDPQKKLAMDDPKLPDGDPIPGFKGYAVNMIDLAPEELTVQTYSGYGLRETLFYNLF  1150
             L+ NDPQK+LAMD+P L +GDPIPGFKGYAVNMIDLA EEL +Q+ SGYGLRETLFY +F
Sbjct  61    LVRNDPQKRLAMDNPYLLNGDPIPGFKGYAVNMIDLASEELDIQSSSGYGLRETLFYGVF  120

Query  1151  GNLQV  1165
               LQV
Sbjct  121   RELQV  125


Range 2: 122 to 162

Score:63.5 bits(153), Expect:4e-54,
Method:Compositional matrix adjust.,
Identities:28/41(68%), Positives:33/41(80%), Gaps:0/41(0%)

Query  1153  EFTGIETQKQVKAALPHINGGGAVSLDGFIAKDNGFIYSGC  1275
             E    ET + ++AALPHINGG AVSLDG IA++NGFIYSGC
Sbjct  122   ELQVYETAEHLEAALPHINGGDAVSLDGVIARENGFIYSGC  162


>putative protein [Arabidopsis thaliana]
Sequence ID: AAM97146.1 Length: 218
>At5g24280 [Arabidopsis thaliana]
Sequence ID: AAP13380.1 Length: 218
Range 1: 10 to 163

Score:169 bits(429), Expect:8e-49,
Method:Compositional matrix adjust.,
Identities:101/161(63%), Positives:119/161(73%), Gaps:9/161(5%)

Query  2    KESQTERRELMLLTKLPECCVAGSNLTNLIFKVTDSDGVMDTSIHHDEKSGCFHTMsiet  181
            KESQ E REL LLT+LP+CC AG+NL NLIF+V + DG +DTSIHHDEK GCFHTMSIE+
Sbjct  10   KESQIEERELRLLTELPDCCAAGTNLMNLIFQVMELDGSLDTSIHHDEKPGCFHTMSIES  69

Query  182  dsssdeseiRYAFVHGSCKVPTLSLPEREGVFSFKVFHSRFPELHLSLKIQLTPAQILQR  361
            DSSS ES IRYAFVHGSCKV +LSLPE EGVFS +VFHSR+PEL +S+KIQ+T A   +R
Sbjct  70   DSSSIESAIRYAFVHGSCKVSSLSLPENEGVFSCRVFHSRYPELQMSVKIQVTSAPTSER  129

Query  362  DETSYSRMGLTPKSKMASTTYSP--ALSSQTGPSLRDVAQF  478
            +E+ YS    TP S    TT  P   + S T P     +QF
Sbjct  130  EESGYS----TPHS---ITTPPPESGIPSITNPWQTPCSQF  163


>hypothetical protein AXX17_AT3G27420 [Arabidopsis thaliana]
Sequence ID: OAP01651.1 Length: 221
Range 1: 40 to 165

Score:169 bits(428), Expect:1e-48,
Method:Compositional matrix adjust.,
Identities:89/126(71%), Positives:106/126(84%), Gaps:0/126(0%)

Query  2    KESQTERRELMLLTKLPECCVAGSNLTNLIFKVTDSDGVMDTSIHHDEKSGCFHTMsiet  181
            KESQ E REL +LT+LP+CC AG+NL NLIF+VT+ DG +DTSIHHDEKSGC HTMSIE+
Sbjct  40   KESQIEERELRILTELPDCCAAGTNLMNLIFQVTELDGSLDTSIHHDEKSGCLHTMSIES  99

Query  182  dsssdeseiRYAFVHGSCKVPTLSLPEREGVFSFKVFHSRFPELHLSLKIQLTPAQILQR  361
            DSSS ES IRYAFVH SCKV +LSLPE EGVFS +VFHSR+PEL +S+KIQ+T A   +R
Sbjct  100  DSSSVESAIRYAFVHWSCKVSSLSLPENEGVFSCRVFHSRYPELQMSVKIQVTSAPTSER  159

Query  362  DETSYS  379
            +E+ YS
Sbjct  160  EESGYS  165


>unnamed protein product [Arabidopsis thaliana]
Sequence ID: BAB01319.1 Length: 258
Range 1: 60 to 230

Score:159 bits(402), Expect:2e-44,
Method:Compositional matrix adjust.,
Identities:101/178(57%), Positives:119/178(66%), Gaps:26/178(14%)

Query  2    KESQTERRELMLLT-----------------KLPECCVAGSNLTNLIFKVTDSDGVMDTS  130
            KESQ E REL LLT                 +LP+CC AG+NL NLIF+V + DG +DTS
Sbjct  60   KESQIEERELRLLTEELVSLLAFDPEIVCIDQLPDCCAAGTNLMNLIFQVMELDGSLDTS  119

Query  131  IHHDEKSGCFHTMsietdsssdeseiRYAFVHGSCKVPTLSLPEREGVFSFKVFHSRFPE  310
            IHHDEK GCFHTMSIE+DSSS ES IRYAFVHGSCKV +LSLPE EGVFS +VFHSR+PE
Sbjct  120  IHHDEKPGCFHTMSIESDSSSIESAIRYAFVHGSCKVSSLSLPENEGVFSCRVFHSRYPE  179

Query  311  LHLSLKIQLTPAQILQRDETSYSRMGLTPKSKMASTTYSP--ALSSQTGPSLRDVAQF  478
            L +S+KIQ+T A   +R+E+ YS    TP S    TT  P   + S T P     +QF
Sbjct  180  LQMSVKIQVTSAPTSEREESGYS----TPHSI---TTPPPESGIPSITNPWQTPCSQF  230


>defective in meristem silencing 3 [Arabidopsis thaliana]
Sequence ID: NP_566916.1 Length: 420
>RecName: Full=Protein DEFECTIVE IN MERISTEM SILENCING 3; AltName: Full=Protein INVOLVED IN DE NOVO 1 [Arabidopsis thaliana]
Sequence ID: Q94A79.1 Length: 420 >AT3g49250/F2K15_110 [Arabidopsis thaliana]
Sequence ID: AAK83630.1 Length: 420 >AT3g49250/F2K15_110 [Arabidopsis thaliana]
Sequence ID: AAL90947.1 Length: 420 >defective in meristem silencing 3 [Arabidopsis thaliana]
Sequence ID: AEE78515.1 Length: 420 >IDN1 [Arabidopsis thaliana]
Sequence ID: OAP02353.1 Length: 420 >unnamed protein product [Arabidopsis thaliana]
Sequence ID: CAA0385231.1 Length: 420
Range 1: 71 to 311

Score:120 bits(302), Expect:5e-29,
Method:Compositional matrix adjust.,
Identities:80/246(33%), Positives:123/246(50%), Gaps:33/246(13%)

Query  533   DERLHCLEAEQNQAKEELRTLQASLEPLGAMFPECLSTKESMMKQIEE------KHHDTA  694
             ++ L  L++++N+  E +  LQ  +  L +  P   S       Q E+      +H ++A
Sbjct  71    EDNLKFLKSQKNKMDEAIVDLQVHMSKLNSS-PTPRSENSDNSLQGEDINAQILRHENSA  129

Query  695   ASVFCCLYRKSPPPQSLFLSKKGVFGLVALLGSVASTSLSRVLSEYLGEDMLLALVCKSA  874
             A V   +        S  +  KGV G+VA LG V   +LS++LS YLG   +LA+VC++
Sbjct  130   AGVLSLVETLHGAQASQLMLTKGVVGVVAKLGKVNDENLSQILSNYLGTRSMLAVVCRNY  189

Query  875   QIGPNNAEFLRLQSNH----------------------RFHVLCLDAIRPWKDGLLENDP  988
             +    +   L    NH                       F  +CL+ +RP+    + +D
Sbjct  190   E----SVTALEAYDNHGNIDINAGLHCLGSSIGREIGDSFDAICLENLRPYVGQHIADDL  245

Query  989   QKKLAMDDPKLPDGDPIPGFKGYAVNMIDLAPEELTVQTYSGYGLRETLFYNLFGNLQV*  1168
             Q++L +  PKLP+G+  PGF G+AVNMI + P  L   T  GYGLRETLFYNLF  LQV
Sbjct  246   QRRLDLLKPKLPNGECPPGFLGFAVNMIQIDPAYLLCVTSYGYGLRETLFYNLFSRLQVY  305

Query  1169  RPKSKL  1186
             + ++ +
Sbjct  306   KTRADM  311


>Chain C, Protein DEFECTIVE IN MERISTEM SILENCING 3 [Arabidopsis thaliana]
Sequence ID: 6OIS_C Length: 449
>Chain D, Protein DEFECTIVE IN MERISTEM SILENCING 3 [Arabidopsis thaliana]
Sequence ID: 6OIS_D Length: 449 >Chain E, Protein DEFECTIVE IN MERISTEM SILENCING 3 [Arabidopsis thaliana]
Sequence ID: 6OIS_E Length: 449 >Chain F, Protein DEFECTIVE IN MERISTEM SILENCING 3 [Arabidopsis thaliana]
Sequence ID: 6OIS_F Length: 449 >Chain C, Protein DEFECTIVE IN MERISTEM SILENCING 3 [Arabidopsis thaliana]
Sequence ID: 6OIT_C Length: 449 >Chain D, Protein DEFECTIVE IN MERISTEM SILENCING 3 [Arabidopsis thaliana]
Sequence ID: 6OIT_D Length: 449 >Chain E, Protein DEFECTIVE IN MERISTEM SILENCING 3 [Arabidopsis thaliana]
Sequence ID: 6OIT_E Length: 449 >Chain F, Protein DEFECTIVE IN MERISTEM SILENCING 3 [Arabidopsis thaliana]
Sequence ID: 6OIT_F Length: 449
Range 1: 74 to 314

Score:120 bits(302), Expect:6e-29,
Method:Compositional matrix adjust.,
Identities:80/246(33%), Positives:123/246(50%), Gaps:33/246(13%)

Query  533   DERLHCLEAEQNQAKEELRTLQASLEPLGAMFPECLSTKESMMKQIEE------KHHDTA  694
             ++ L  L++++N+  E +  LQ  +  L +  P   S       Q E+      +H ++A
Sbjct  74    EDNLKFLKSQKNKMDEAIVDLQVHMSKLNSS-PTPRSENSDNSLQGEDINAQILRHENSA  132

Query  695   ASVFCCLYRKSPPPQSLFLSKKGVFGLVALLGSVASTSLSRVLSEYLGEDMLLALVCKSA  874
             A V   +        S  +  KGV G+VA LG V   +LS++LS YLG   +LA+VC++
Sbjct  133   AGVLSLVETLHGAQASQLMLTKGVVGVVAKLGKVNDENLSQILSNYLGTRSMLAVVCRNY  192

Query  875   QIGPNNAEFLRLQSNH----------------------RFHVLCLDAIRPWKDGLLENDP  988
             +    +   L    NH                       F  +CL+ +RP+    + +D
Sbjct  193   E----SVTALEAYDNHGNIDINAGLHCLGSSIGREIGDSFDAICLENLRPYVGQHIADDL  248

Query  989   QKKLAMDDPKLPDGDPIPGFKGYAVNMIDLAPEELTVQTYSGYGLRETLFYNLFGNLQV*  1168
             Q++L +  PKLP+G+  PGF G+AVNMI + P  L   T  GYGLRETLFYNLF  LQV
Sbjct  249   QRRLDLLKPKLPNGECPPGFLGFAVNMIQIDPAYLLCVTSYGYGLRETLFYNLFSRLQVY  308

Query  1169  RPKSKL  1186
             + ++ +
Sbjct  309   KTRADM  314


>hypothetical protein [Arabidopsis thaliana]
Sequence ID: CAB66404.1 Length: 422
Range 1: 67 to 290

Score:105 bits(261), Expect:1e-23,
Method:Compositional matrix adjust.,
Identities:75/233(32%), Positives:114/233(48%), Gaps:24/233(10%)

Query  533   DERLHCLEAEQNQAKEELRTLQASLEPLGAMFPECLSTKESMMKQIEE------KHHDTA  694
             ++ L  L++++N+  E +  LQ  +  L +  P   S       Q E+      +H ++A
Sbjct  67    EDNLKFLKSQKNKMDEAIVDLQVHMSKLNSS-PTPRSENSDNSLQGEDINAQILRHENSA  125

Query  695   ASVFCCLYRKSPPPQSLFLSKKGVFGLVALLGSVASTSLSR------VLSEYLGE---DM  847
             A V   +        S  +  KGV G+VA LG V   +LS+       L  Y      D+
Sbjct  126   AGVLSLVETLHGAQASQLMLTKGVVGVVAKLGKVNDENLSQNYESVTALEAYDNHGNIDI  185

Query  848   LLALVCKSAQIGPNNAEFLRLQSNHRFHVLCLDAIRPWKDGLLENDPQKKLAMDDPKLPD  1027
                L C  + IG         +    F  +CL+ +RP+    + +D Q++L +  PKLP+
Sbjct  186   NAGLHCLGSSIGR--------EIGDSFDAICLENLRPYVGQHIADDLQRRLDLLKPKLPN  237

Query  1028  GDPIPGFKGYAVNMIDLAPEELTVQTYSGYGLRETLFYNLFGNLQV*RPKSKL  1186
             G+  PGF G+AVNMI + P  L   T  GYGLRETLFYNLF  LQV + ++ +
Sbjct  238   GECPPGFLGFAVNMIQIDPAYLLCVTSYGYGLRETLFYNLFSRLQVYKTRADM  290


>hypothetical protein AXX17_AT3G27430 [Arabidopsis thaliana]
Sequence ID: OAP05396.1 Length: 129
Range 1: 40 to 89

Score:78.6 bits(192), Expect:1e-16,
Method:Compositional matrix adjust.,
Identities:36/50(72%), Positives:42/50(84%), Gaps:0/50(0%)

Query  2    KESQTERRELMLLTKLPECCVAGSNLTNLIFKVTDSDGVMDTSIHHDEKS  151
            KESQ E REL LLT+LP+CC AG+NL NLIF+VT+ DG +DTS HHDEKS
Sbjct  40   KESQIEERELRLLTELPDCCAAGTNLMNLIFQVTELDGSLDTSNHHDEKS  89


>hypothetical protein AXX17_AT3G27430 [Arabidopsis thaliana]
Sequence ID: OAP05397.1 Length: 182
Range 1: 40 to 89

Score:79.0 bits(193), Expect:3e-16,
Method:Compositional matrix adjust.,
Identities:36/50(72%), Positives:42/50(84%), Gaps:0/50(0%)

Query  2    KESQTERRELMLLTKLPECCVAGSNLTNLIFKVTDSDGVMDTSIHHDEKS  151
            KESQ E REL LLT+LP+CC AG+NL NLIF+VT+ DG +DTS HHDEKS
Sbjct  40   KESQIEERELRLLTELPDCCAAGTNLMNLIFQVTELDGSLDTSNHHDEKS  89


Query #257: XLOC_030033 Query ID: lcl|Query_30748 Length: 2194

Sequences producing significant alignments:
                                                                  Max    Total Query    E    Per.
Description                                                       Score  Score cover Value  Ident  Accession
unnamed protein product [Arabidopsis thaliana]                    645    1229  96%   0.0    100.00 BAB10393.1
En/Spm-like transposon [Arabidopsis thaliana]                     584    644   58%   0.0    93.78  NP_001331299.1
hypothetical protein AXX17_AT5G24050 [Arabidopsis thaliana]       583    1146  96%   0.0    93.78  OAO94301.1
unnamed protein product [Arabidopsis thaliana]                    559    672   62%   0.0    90.55  CAA0404532.1
unnamed protein product [Arabidopsis thaliana]                    554    647   60%   0.0    87.13  VYS67775.1
En/Spm-like transposon protein-like [Arabidopsis thaliana]        423    485   50%   1e-137 65.05  BAB01349.1
En/Spm-like transposon [Arabidopsis thaliana]                     402    402   30%   4e-131 89.64  NP_001331300.1
putative protein [Arabidopsis thaliana]                           296    296   40%   4e-89  47.27  CAB77996.1
hypothetical protein [Arabidopsis thaliana]                       289    289   40%   1e-86  46.08  AAC23763.1
unnamed protein product [Arabidopsis thaliana]                    273    273   42%   4e-81  44.03  BAB02522.1
RecName: Full=Uncharacterized protein At3g43530 [Arabidopsis...   258    318   57%   2e-79  47.28  Q9M237.1
unnamed protein product [Arabidopsis thaliana]                    249    249   39%   2e-72  45.36  BAB09148.1
putative protein [Arabidopsis thaliana]                           249    249   39%   3e-72  45.36  CAB78060.1
contains similarity to T. cruzi 3' end fragment (PID:g161956)...  248    248   41%   1e-71  42.17  AAB61080.1
F5J5.9 [Arabidopsis thaliana]                                     246    246   41%   5e-71  43.09  AAF18635.1
unknown protein [Arabidopsis thaliana]                            243    243   39%   6e-70  44.33  AAC17091.2
hypothetical protein AXX17_ATUG02420 [Arabidopsis thaliana]       241    241   39%   2e-69  44.33  OAO89237.1
T23E23.10 [Arabidopsis thaliana]                                  242    242   39%   6e-69  44.22  AAF87156.1
putative protein [Arabidopsis thaliana]                           236    236   39%   3e-67  43.64  CAB51205.1
unnamed protein product [Arabidopsis thaliana]                    233    233   39%   2e-66  43.64  BAB10960.1
hypothetical protein [Arabidopsis thaliana]                       234    234   41%   5e-66  42.39  AAD36941.1
hypothetical protein [Arabidopsis thaliana]                       220    220   23%   5e-64  60.92  AAD19761.1
hypothetical protein [Arabidopsis thaliana]                       208    208   33%   2e-60  43.35  AAD26890.1
contains similarity to several [Arabidopsis thaliana...           202    202   41%   6e-55  38.41  AAD38226.1
hypothetical protein [Arabidopsis thaliana]                       184    184   30%   2e-52  42.49  AAO92060.1
En/Spm-like transposon, putative (DUF1985) [Arabidopsis thaliana] 183    183   30%   2e-52  42.49  NP_178683.2
hypothetical protein At2g06420 [Arabidopsis thaliana]             183    183   30%   3e-52  42.49  AAY78684.1
hypothetical protein [Arabidopsis thaliana]                       186    186   30%   2e-50  44.05  AAD25589.1
putative protein [Arabidopsis thaliana]                           186    186   32%   4e-50  40.76  CAB86477.1
hypothetical protein AXX17_AT5G24030 [Arabidopsis thaliana]       181    181   16%   5e-49  72.50  OAO92751.1
putative protein [Arabidopsis thaliana]                           178    178   38%   1e-47  36.07  CAB83145.1
En/Spm-like transposon protein [Arabidopsis thaliana]             145    204   36%   3e-45  53.24  AAC26678.1
putative protein [Arabidopsis thaliana]                           174    174   41%   4e-45  35.48  CAB72487.1
hypothetical protein [Arabidopsis thaliana]                       164    164   31%   1e-43  40.17  AAD28050.1
unnamed protein product [Arabidopsis thaliana]                    158    158   16%   4e-39  63.28  BAB10394.1
hypothetical protein [Arabidopsis thaliana]                       150    150   39%   6e-38  32.43  AAD25845.1
hypothetical protein AXX17_AT2G07430 [Arabidopsis thaliana]       144    144   29%   4e-36  38.91  OAP10004.1
gamma-irradiation and mitomycin c induced 1 [Arabidopsis...       141    141   16%   1e-33  58.06  NP_001330245.1
gamma-irradiation and mitomycin c induced 1 [Arabidopsis...       140    140   16%   2e-33  58.06  NP_197816.3
GMI1 [Arabidopsis thaliana]                                       140    140   16%   2e-33  58.06  OAO93531.1
unnamed protein product [Arabidopsis thaliana]                    140    140   16%   2e-33  58.06  CAA0404534.1
F28J9.13 [Arabidopsis thaliana]                                   131    131   16%   6e-32  50.40  AAF06090.1
hypothetical protein AXX17_ATUG00940 [Arabidopsis thaliana]       130    190   31%   4e-31  37.78  OAO89416.1
putative protein [Arabidopsis thaliana]                           128    187   23%   1e-30  52.25  CAB43910.1
hypothetical protein [Arabidopsis thaliana]                       108    108   24%   3e-26  37.50  AAO73430.1
unnamed protein product [Arabidopsis thaliana]                    103    103   15%   1e-25  46.36  VYS48117.1
unnamed protein product [Arabidopsis thaliana]                    113    222   31%   1e-25  47.17  BAB01195.1
T25H8.3 gene product [Arabidopsis thaliana]                       106    106   20%   2e-24  37.41  AAD17348.1
hypothetical protein [Arabidopsis thaliana]                       107    107   24%   3e-24  36.22  AAD20103.1
hypothetical protein [Arabidopsis thaliana]                       100    177   29%   2e-21  50.57  AAB81878.1
unnamed protein product [Arabidopsis thaliana]                    80.1   80.1  11%   1e-17  45.65  VYS59032.1
hypothetical protein AXX17_ATUG01010 [Arabidopsis thaliana]       82.8   82.8  17%   1e-15  38.35  OAO89413.1
hypothetical protein AXX17_AT4G07660 [Arabidopsis thaliana]       65.1   65.1  10%   2e-12  36.84  OAO98863.1
hypothetical protein AXX17_ATUG02090 [Arabidopsis thaliana]       65.5   65.5  11%   4e-11  35.87  OAO89294.1
hypothetical protein AXX17_ATUG02100 [Arabidopsis thaliana]       66.6   66.6  9%    5e-11  41.33  OAO89302.1
putative protein [Arabidopsis thaliana]                           64.3   64.3  24%   1e-10  26.92  CAB86472.1
unnamed protein product [Arabidopsis thaliana]                    66.6   66.6  30%   1e-10  26.75  VYS47676.1
K-box region protein (DUF1985) [Arabidopsis thaliana]             65.5   65.5  30%   3e-10  26.34  NP_001321227.1
hypothetical protein AXX17_AT3G33420 [Arabidopsis thaliana]       65.9   65.9  25%   3e-10  30.05  OAP06548.1
hypothetical protein AXX17_AT1G31740 [Arabidopsis thaliana]       65.5   65.5  30%   3e-10  26.34  OAP15120.1
K-box region protein (DUF1985) [Arabidopsis thaliana]             65.5   65.5  30%   4e-10  26.34  NP_174400.1
unknown protein [Arabidopsis thaliana]                            65.1   65.1  23%   5e-10  28.29  AAG50606.1
putative protein [Arabidopsis thaliana]                           65.1   65.1  29%   5e-10  23.01  CAB81580.1
Contains similarity to a putative protein T32A11_100 gi|741362... 64.7   64.7  26%   6e-10  26.55  AAF78272.1
unnamed protein product [Arabidopsis thaliana]                    64.7   64.7  29%   7e-10  22.27  BAB02940.1
unnamed protein product [Arabidopsis thaliana]                    64.3   64.3  29%   9e-10  21.85  BAA98062.1
putative protein [Arabidopsis thaliana]                           64.3   64.3  27%   9e-10  23.15  CAB86452.1
hypothetical protein [Arabidopsis thaliana]                       63.9   63.9  19%   1e-09  32.89  AAD36942.1
T22C5.24 [Arabidopsis thaliana]                                   63.9   63.9  29%   1e-09  22.27  AAF24952.1
hypothetical protein [Arabidopsis thaliana]                       63.9   63.9  32%   1e-09  23.11  AAG50521.1
hypothetical protein [Arabidopsis thaliana]                       63.5   63.5  29%   2e-09  23.14  AAG50582.1
hypothetical protein [Arabidopsis thaliana]                       63.5   63.5  29%   2e-09  23.01  AAD14458.1
unnamed protein product [Arabidopsis thaliana]                    59.3   59.3  8%    3e-09  41.67  CAA0384296.1
F27F5.16 [Arabidopsis thaliana]                                   62.8   62.8  30%   3e-09  25.00  AAF69170.1
hypothetical protein [Arabidopsis thaliana]                       62.0   62.0  30%   5e-09  25.00  AAC97230.2
unnamed protein product [Arabidopsis thaliana]                    55.1   55.1  9%    5e-09  45.45  CAA0269437.1
hypothetical protein [Arabidopsis thaliana]                       60.5   60.5  31%   1e-08  23.01  AAC95211.1
unnamed protein product [Arabidopsis thaliana]                    60.5   60.5  31%   1e-08  23.01  BAB01835.1
Similar to a hypothetical protein At2g29240 gi|3980408 from...    60.5   60.5  31%   1e-08  23.01  AAF80659.1
hypothetical protein [Arabidopsis thaliana]                       59.7   59.7  9%    1e-08  43.06  AAD23703.1
En/Spm-like transposon [Arabidopsis thaliana]                     59.3   59.3  3%    2e-08  100.00 NP_001331298.1
putative protein [Arabidopsis thaliana]                           60.1   60.1  29%   2e-08  24.48  CAB86693.1
hypothetical protein [Arabidopsis thaliana]                       58.9   58.9  22%   5e-08  26.23  AAD26959.1
F22O13.23 [Arabidopsis thaliana]                                  58.5   58.5  31%   5e-08  22.59  AAF99775.1
putative protein [Arabidopsis thaliana]                           57.8   57.8  24%   8e-08  24.63  CAB81786.1
hypothetical protein [Arabidopsis thaliana]                       55.5   55.5  9%    9e-08  40.00  AAD20102.1


Alignments:
>unnamed protein product [Arabidopsis thaliana]
Sequence ID: BAB10393.1 Length: 1335
Range 1: 148 to 450

Score:645 bits(1665), Expect:0.0,
Method:Compositional matrix adjust.,
Identities:303/303(100%), Positives:303/303(100%), Gaps:0/303(0%)

Query  3    NEETKQISMDSEEETEPLRPLKMYFSPSDYLKPFKIGTKCYIHQVVNILESHLSSEEKKW  182
            NEETKQISMDSEEETEPLRPLKMYFSPSDYLKPFKIGTKCYIHQVVNILESHLSSEEKKW
Sbjct  148  NEETKQISMDSEEETEPLRPLKMYFSPSDYLKPFKIGTKCYIHQVVNILESHLSSEEKKW  207

Query  183  FWEHPQFKHFFHMHTDSNHKVMAMWMLLLRTACVDKKKECWFVVNGVPIRYSLQELALIS  362
            FWEHPQFKHFFHMHTDSNHKVMAMWMLLLRTACVDKKKECWFVVNGVPIRYSLQELALIS
Sbjct  208  FWEHPQFKHFFHMHTDSNHKVMAMWMLLLRTACVDKKKECWFVVNGVPIRYSLQELALIS  267

Query  363  GLYCHSYPKNYKAFGSLGFSGKHFGVGATVTYAKVKTKLLSMKKSSRERLKMAVLFFLCS  542
            GLYCHSYPKNYKAFGSLGFSGKHFGVGATVTYAKVKTKLLSMKKSSRERLKMAVLFFLCS
Sbjct  268  GLYCHSYPKNYKAFGSLGFSGKHFGVGATVTYAKVKTKLLSMKKSSRERLKMAVLFFLCS  327

Query  543  VIIGKRKTGEKAEAVEDFFLKSVEDLEWCKTFPWGRLAFDKNMKDIFHLMDHFDEELGAA  722
            VIIGKRKTGEKAEAVEDFFLKSVEDLEWCKTFPWGRLAFDKNMKDIFHLMDHFDEELGAA
Sbjct  328  VIIGKRKTGEKAEAVEDFFLKSVEDLEWCKTFPWGRLAFDKNMKDIFHLMDHFDEELGAA  387

Query  723  WVFPSFVMQLLAFETIPNLMNNFRKHVREADPECPRMCKMKFKPSSMKGFPISELYDTLG  902
            WVFPSFVMQLLAFETIPNLMNNFRKHVREADPECPRMCKMKFKPSSMKGFPISELYDTLG
Sbjct  388  WVFPSFVMQLLAFETIPNLMNNFRKHVREADPECPRMCKMKFKPSSMKGFPISELYDTLG  447

Query  903  TTK  911
            TTK
Sbjct  448  TTK  450


Range 2: 449 to 827

Score:583 bits(1502), Expect:0.0,
Method:Compositional matrix adjust.,
Identities:377/402(94%), Positives:379/402(94%), Gaps:23/402(5%)

Query  989   SQEIKSIMLPSPSEKELLKQIMddevghddvddMIVDGWTKRIIKENKPICFEELHIQDV  1168
             ++EIKSIMLPSPSEKELLKQIMDDEVGHDDVDDMIVDGWTKRIIKENKPICFEELHIQDV
Sbjct  449   TKEIKSIMLPSPSEKELLKQIMDDEVGHDDVDDMIVDGWTKRIIKENKPICFEELHIQDV  508

Query  1169  SSREIEANPknanvvgvrvgrkkvalkkvSEKGIVEQLQKQLEDGLNRIYMKIDDMDTRL  1348
             SSREIEANPKNANVVGVRVGRKKVALKKVSEKGIVEQLQKQLEDGLNRIYMKIDDMDTRL
Sbjct  509   SSREIEANPKNANVVGVRVGRKKVALKKVSEKGIVEQLQKQLEDGLNRIYMKIDDMDTRL  568

Query  1349  TYVESCVKNLkeakdtkgekedgekekeleendleedetetdkekekedessekRSKVER  1528
             TYVESCVKNLKEAKDTKGEKEDGEKEKELEENDLEEDETETDKEKEKEDESSEKRSKVER
Sbjct  569   TYVESCVKNLKEAKDTKGEKEDGEKEKELEENDLEEDETETDKEKEKEDESSEKRSKVER  628

Query  1529  MYGRKRTRTTTAKEQEEKGEEIQVGFQETSERGRKKAKLTWKKVDIKGVTADQGNSITTN  1708
             MYGRKRTRTTTAKEQEEKGEEIQVGFQETSERGRKKAKLTWKKVDIKGVTADQGNSITTN
Sbjct  629   MYGRKRTRTTTAKEQEEKGEEIQVGFQETSERGRKKAKLTWKKVDIKGVTADQGNSITTN  688

Query  1709  ERVVVEPSSKVGSWKLASNLKSTRQYNVQVGSSLPPCSISCVDEPFSVSVSCKVNPGPLK  1888
             ERVVVEPSSKVGSWKLASNLKSTRQYNVQVGSSLPPCSISCVDEPFSVSVSCKVNPGPLK
Sbjct  689   ERVVVEPSSKVGSWKLASNLKSTRQYNVQVGSSLPPCSISCVDEPFSVSVSCKVNPGPLK  748

Query  1889  HVVEKYPEALENLLPGSTVQNYILEVKVYLLSIE*LHQIKLCTIFFMQVFDGYNNHVAEG  2068
             HVVEKYPEALENLLPGSTVQNYILE                       VFDGYNNHVAEG
Sbjct  749   HVVEKYPEALENLLPGSTVQNYILE-----------------------VFDGYNNHVAEG  785

Query  2069  TNVLICIEGYCINDSMGLNQKVNSCGCIDLSGILQVTAGYGK  2194
             TNVLICIEGYCINDSMGLNQKVNSCGCIDLSGILQVTAGYGK
Sbjct  786   TNVLICIEGYCINDSMGLNQKVNSCGCIDLSGILQVTAGYGK  827


>En/Spm-like transposon [Arabidopsis thaliana]
Sequence ID: NP_001331299.1 Length: 807
>En/Spm-like transposon [Arabidopsis thaliana]
Sequence ID: ANM69636.1 Length: 807
Range 1: 26 to 404

Score:584 bits(1505), Expect:0.0,
Method:Compositional matrix adjust.,
Identities:377/402(94%), Positives:379/402(94%), Gaps:23/402(5%)

Query  989   SQEIKSIMLPSPSEKELLKQIMddevghddvddMIVDGWTKRIIKENKPICFEELHIQDV  1168
             ++EIKSIMLPSPSEKELLKQIMDDEVGHDDVDDMIVDGWTKRIIKENKPICFEELHIQDV
Sbjct  26    TKEIKSIMLPSPSEKELLKQIMDDEVGHDDVDDMIVDGWTKRIIKENKPICFEELHIQDV  85

Query  1169  SSREIEANPknanvvgvrvgrkkvalkkvSEKGIVEQLQKQLEDGLNRIYMKIDDMDTRL  1348
             SSREIEANPKNANVVGVRVGRKKVALKKVSEKGIVEQLQKQLEDGLNRIYMKIDDMDTRL
Sbjct  86    SSREIEANPKNANVVGVRVGRKKVALKKVSEKGIVEQLQKQLEDGLNRIYMKIDDMDTRL  145

Query  1349  TYVESCVKNLkeakdtkgekedgekekeleendleedetetdkekekedessekRSKVER  1528
             TYVESCVKNLKEAKDTKGEKEDGEKEKELEENDLEEDETETDKEKEKEDESSEKRSKVER
Sbjct  146   TYVESCVKNLKEAKDTKGEKEDGEKEKELEENDLEEDETETDKEKEKEDESSEKRSKVER  205

Query  1529  MYGRKRTRTTTAKEQEEKGEEIQVGFQETSERGRKKAKLTWKKVDIKGVTADQGNSITTN  1708
             MYGRKRTRTTTAKEQEEKGEEIQVGFQETSERGRKKAKLTWKKVDIKGVTADQGNSITTN
Sbjct  206   MYGRKRTRTTTAKEQEEKGEEIQVGFQETSERGRKKAKLTWKKVDIKGVTADQGNSITTN  265

Query  1709  ERVVVEPSSKVGSWKLASNLKSTRQYNVQVGSSLPPCSISCVDEPFSVSVSCKVNPGPLK  1888
             ERVVVEPSSKVGSWKLASNLKSTRQYNVQVGSSLPPCSISCVDEPFSVSVSCKVNPGPLK
Sbjct  266   ERVVVEPSSKVGSWKLASNLKSTRQYNVQVGSSLPPCSISCVDEPFSVSVSCKVNPGPLK  325

Query  1889  HVVEKYPEALENLLPGSTVQNYILEVKVYLLSIE*LHQIKLCTIFFMQVFDGYNNHVAEG  2068
             HVVEKYPEALENLLPGSTVQNYILE                       VFDGYNNHVAEG
Sbjct  326   HVVEKYPEALENLLPGSTVQNYILE-----------------------VFDGYNNHVAEG  362

Query  2069  TNVLICIEGYCINDSMGLNQKVNSCGCIDLSGILQVTAGYGK  2194
             TNVLICIEGYCINDSMGLNQKVNSCGCIDLSGILQVTAGYGK
Sbjct  363   TNVLICIEGYCINDSMGLNQKVNSCGCIDLSGILQVTAGYGK  404


Range 2: 1 to 27

Score:60.5 bits(145), Expect:1e-08,
Method:Compositional matrix adjust.,
Identities:27/27(100%), Positives:27/27(100%), Gaps:0/27(0%)

Query  831  MCKMKFKPSSMKGFPISELYDTLGTTK  911
            MCKMKFKPSSMKGFPISELYDTLGTTK
Sbjct  1    MCKMKFKPSSMKGFPISELYDTLGTTK  27


>hypothetical protein AXX17_AT5G24050 [Arabidopsis thaliana]
Sequence ID: OAO94301.1 Length: 1442
Range 1: 417 to 795

Score:583 bits(1503), Expect:0.0,
Method:Compositional matrix adjust.,
Identities:377/402(94%), Positives:379/402(94%), Gaps:23/402(5%)

Query  989   SQEIKSIMLPSPSEKELLKQIMddevghddvddMIVDGWTKRIIKENKPICFEELHIQDV  1168
             ++EIKSIMLPSPSEKELLKQIMDDEVGHDDVDDMIVDGWTKRIIKENKPICFEELHIQDV
Sbjct  417   TKEIKSIMLPSPSEKELLKQIMDDEVGHDDVDDMIVDGWTKRIIKENKPICFEELHIQDV  476

Query  1169  SSREIEANPknanvvgvrvgrkkvalkkvSEKGIVEQLQKQLEDGLNRIYMKIDDMDTRL  1348
             SSREIEANPKNANVVGVRVGRKKVALKKVSEKGIVEQLQKQLEDGLNRIYMKIDDMDTRL
Sbjct  477   SSREIEANPKNANVVGVRVGRKKVALKKVSEKGIVEQLQKQLEDGLNRIYMKIDDMDTRL  536

Query  1349  TYVESCVKNLkeakdtkgekedgekekeleendleedetetdkekekedessekRSKVER  1528
             TYVESCVKNLKEAKDTKGEKEDGEKEKELEENDLEEDETETDKEKEKEDESSEKRSKVER
Sbjct  537   TYVESCVKNLKEAKDTKGEKEDGEKEKELEENDLEEDETETDKEKEKEDESSEKRSKVER  596

Query  1529  MYGRKRTRTTTAKEQEEKGEEIQVGFQETSERGRKKAKLTWKKVDIKGVTADQGNSITTN  1708
             MYGRKRTRTTTAKEQEEKGEEIQVGFQETSERGRKKAKLTWKKVDIKGVTADQGNSITTN
Sbjct  597   MYGRKRTRTTTAKEQEEKGEEIQVGFQETSERGRKKAKLTWKKVDIKGVTADQGNSITTN  656

Query  1709  ERVVVEPSSKVGSWKLASNLKSTRQYNVQVGSSLPPCSISCVDEPFSVSVSCKVNPGPLK  1888
             ERVVVEPSSKVGSWKLASNLKSTRQYNVQVGSSLPPCSISCVDEPFSVSVSCKVNPGPLK
Sbjct  657   ERVVVEPSSKVGSWKLASNLKSTRQYNVQVGSSLPPCSISCVDEPFSVSVSCKVNPGPLK  716

Query  1889  HVVEKYPEALENLLPGSTVQNYILEVKVYLLSIE*LHQIKLCTIFFMQVFDGYNNHVAEG  2068
             HVVEKYPEALENLLPGSTVQNYILE                       VFDGYNNHVAEG
Sbjct  717   HVVEKYPEALENLLPGSTVQNYILE-----------------------VFDGYNNHVAEG  753

Query  2069  TNVLICIEGYCINDSMGLNQKVNSCGCIDLSGILQVTAGYGK  2194
             TNVLICIEGYCINDSMGLNQKVNSCGCIDLSGILQVTAGYGK
Sbjct  754   TNVLICIEGYCINDSMGLNQKVNSCGCIDLSGILQVTAGYGK  795


Range 2: 142 to 418

Score:562 bits(1449), Expect:0.0,
Method:Compositional matrix adjust.,
Identities:272/305(89%), Positives:274/305(89%), Gaps:30/305(9%)

Query  3    NEETKQISMDSEEETEPLRPLKMYFSPSDYLKPFKIGTKCYIHQVVNILESHLSSEEKKW  182
            NEETKQISMDSEEETEPLRPLK+                            HLSSEEKKW
Sbjct  142  NEETKQISMDSEEETEPLRPLKI----------------------------HLSSEEKKW  173

Query  183  FWEHPQFKHFFHMHTDSNHKVMAMWMLLLRTACVDKKKECWFVVNGVPIRYSLQELALIS  362
            FWEHPQFKHFFHMHTDSNHKVMAMWMLLLRTACVDKKKECWFVVNGVPIRYSLQELALIS
Sbjct  174  FWEHPQFKHFFHMHTDSNHKVMAMWMLLLRTACVDKKKECWFVVNGVPIRYSLQELALIS  233

Query  363  GLYCHSYPKNYKAFGSLGFSGKHFGVGATVTYAKVKTKLLSMKKSSRERLKMAVLFFLCS  542
            GLYCHSYPKNYKAFGSLGFSGKHFGVGATVTYAKVKTKLLSMKKSSRERLKMAVLFFLCS
Sbjct  234  GLYCHSYPKNYKAFGSLGFSGKHFGVGATVTYAKVKTKLLSMKKSSRERLKMAVLFFLCS  293

Query  543  VIIGKRKTGEKAEAVEDFFLKSVEDLEWCKTFPWGRLAFDKNMKDIFHLMDHFDEELGAA  722
            VIIGKRKTGEKAEAVEDFFLKSVEDLEWCKTFPWGRLAFDKNMKDIFHLMDHFDEELGAA
Sbjct  294  VIIGKRKTGEKAEAVEDFFLKSVEDLEWCKTFPWGRLAFDKNMKDIFHLMDHFDEELGAA  353

Query  723  WVFPSFVM--QLLAFETIPNLMNNFRKHVREADPECPRMCKMKFKPSSMKGFPISELYDT  896
            WVFPSFVM  +LLAFETIPNLMNNFRKHVREADPECPRMCKMKFKPSSMKGFPISELYDT
Sbjct  354  WVFPSFVMPLELLAFETIPNLMNNFRKHVREADPECPRMCKMKFKPSSMKGFPISELYDT  413

Query  897  LGTTK  911
            LG TK
Sbjct  414  LGATK  418


>unnamed protein product, partial [Arabidopsis thaliana]
Sequence ID: CAA0404532.1 Length: 877
Range 1: 53 to 431

Score:559 bits(1440), Expect:0.0,
Method:Compositional matrix adjust.,
Identities:364/402(91%), Positives:371/402(92%), Gaps:23/402(5%)

Query  989   SQEIKSIMLPSPSEKELLKQIMddevghddvddMIVDGWTKRIIKENKPICFEELHIQDV  1168
             ++EIKSIMLPSPSEKELLKQIMDDEVGHDDVDD+IVDGWTKRIIKENKPICFEELHIQDV
Sbjct  53    TKEIKSIMLPSPSEKELLKQIMDDEVGHDDVDDVIVDGWTKRIIKENKPICFEELHIQDV  112

Query  1169  SSREIEANPknanvvgvrvgrkkvalkkvSEKGIVEQLQKQLEDGLNRIYMKIDDMDTRL  1348
             SSREIEANPKNA+VVGVRVGRKKVALKKVSEKGI EQLQKQLEDG+ RI MKIDDMDTRL
Sbjct  113   SSREIEANPKNADVVGVRVGRKKVALKKVSEKGIFEQLQKQLEDGIKRIDMKIDDMDTRL  172

Query  1349  TYVESCVKNLkeakdtkgekedgekekeleendleedetetdkekekedessekRSKVER  1528
             TYVESCVKNLKEAKDTKGEKEDGEKEKELEENDL+EDETETDKEKEKEDESSEKRSKVER
Sbjct  173   TYVESCVKNLKEAKDTKGEKEDGEKEKELEENDLDEDETETDKEKEKEDESSEKRSKVER  232

Query  1529  MYGRKRTRTTTAKEQEEKGEEIQVGFQETSERGRKKAKLTWKKVDIKGVTADQGNSITTN  1708
             MYGRKRTRTTTAKEQEEKGEEIQVGFQETSERGRKKAKLTWKKVDIKGVTADQGNSITTN
Sbjct  233   MYGRKRTRTTTAKEQEEKGEEIQVGFQETSERGRKKAKLTWKKVDIKGVTADQGNSITTN  292

Query  1709  ERVVVEPSSKVGSWKLASNLKSTRQYNVQVGSSLPPCSISCVDEPFSVSVSCKVNPGPLK  1888
             +RVVVEPSS VGSWKLASNLKS RQYN QVGSSLPPCSISCVDEPFSVSVSCKVNPGPL
Sbjct  293   KRVVVEPSSNVGSWKLASNLKSMRQYNAQVGSSLPPCSISCVDEPFSVSVSCKVNPGPLT  352

Query  1889  HVVEKYPEALENLLPGSTVQNYILEVKVYLLSIE*LHQIKLCTIFFMQVFDGYNNHVAEG  2068
             HVVEKYPEALENLLPGSTVQNYILE                       VFDGYNNHVAEG
Sbjct  353   HVVEKYPEALENLLPGSTVQNYILE-----------------------VFDGYNNHVAEG  389

Query  2069  TNVLICIEGYCINDSMGLNQKVNSCGCIDLSGILQVTAGYGK  2194
             TNVLICIEGY INDSMGLNQKVNSCGCIDLSGILQVTAGYGK
Sbjct  390   TNVLICIEGYFINDSMGLNQKVNSCGCIDLSGILQVTAGYGK  431


Range 2: 1 to 54

Score:112 bits(281), Expect:7e-25,
Method:Compositional matrix adjust.,
Identities:51/54(94%), Positives:51/54(94%), Gaps:0/54(0%)

Query  750  LLAFETIPNLMNNFRKHVREADPECPRMCKMKFKPSSMKGFPISELYDTLGTTK  911
            LLAFETIPNLMN FR HVREADPECPRMCKMKFK SSMKGFPISELYDTLGTTK
Sbjct  1    LLAFETIPNLMNKFRNHVREADPECPRMCKMKFKASSMKGFPISELYDTLGTTK  54


>unnamed protein product [Arabidopsis thaliana]
Sequence ID: VYS67775.1 Length: 869
Range 1: 43 to 423

Score:554 bits(1427), Expect:0.0,
Method:Compositional matrix adjust.,
Identities:352/404(87%), Positives:363/404(89%), Gaps:25/404(6%)

Query  989   SQEIKSIMLPSPSEKELLKQIMddevghddvddMIVDGWTKRIIKENKPICFEELHIQDV  1168
             ++EIKSIMLPSPSEKELLKQIMDDEVGHDDVDD+IVDGWTKRIIKENKPICFEELHIQDV
Sbjct  43    TKEIKSIMLPSPSEKELLKQIMDDEVGHDDVDDVIVDGWTKRIIKENKPICFEELHIQDV  102

Query  1169  SSREIEANPknanvvgvrvgrkkvalkkvSEKGIVEQLQKQLEDGLNRIYMKIDDMDTRL  1348
             SSREIEANPKNA+VVGVRVGRKKVALKKVSEKGI EQLQKQLEDG+ RI MKIDDMDTRL
Sbjct  103   SSREIEANPKNADVVGVRVGRKKVALKKVSEKGIFEQLQKQLEDGIKRIDMKIDDMDTRL  162

Query  1349  TYVESCVKNLkeakdtkgeke--dgekekeleendleedetetdkekekedessekRSKV  1522
             TYVESCVKNLKEAKDTKGEKE  + EKE E  + D +E ET+ +KEKEKEDESSEKRSKV
Sbjct  163   TYVESCVKNLKEAKDTKGEKEDGEKEKELEENDLDEDETETDKEKEKEKEDESSEKRSKV  222

Query  1523  ERMYGRKRTRTTTAKEQEEKGEEIQVGFQETSERGRKKAKLTWKKVDIKGVTADQGNSIT  1702
             ERMYGRKRTRTTTAKEQEEKGEEIQVGFQETSERGRKKAKLTWKKVDIKGVTADQGNSIT
Sbjct  223   ERMYGRKRTRTTTAKEQEEKGEEIQVGFQETSERGRKKAKLTWKKVDIKGVTADQGNSIT  282

Query  1703  TNERVVVEPSSKVGSWKLASNLKSTRQYNVQVGSSLPPCSISCVDEPFSVSVSCKVNPGP  1882
             TN+RVVVEPSS VGSWKLASNLKS RQYN QVGSSLPPCSISCVDEPFSVSVSCKVNPGP
Sbjct  283   TNKRVVVEPSSNVGSWKLASNLKSMRQYNAQVGSSLPPCSISCVDEPFSVSVSCKVNPGP  342

Query  1883  LKHVVEKYPEALENLLPGSTVQNYILEVKVYLLSIE*LHQIKLCTIFFMQVFDGYNNHVA  2062
             L HVVEKYPEALENLLPGSTVQNYILE                       VFDGYNNHVA
Sbjct  343   LTHVVEKYPEALENLLPGSTVQNYILE-----------------------VFDGYNNHVA  379

Query  2063  EGTNVLICIEGYCINDSMGLNQKVNSCGCIDLSGILQVTAGYGK  2194
             EGTNVLICIEGY INDSMGLNQKVNSCGCIDLSGILQVTAGYGK
Sbjct  380   EGTNVLICIEGYFINDSMGLNQKVNSCGCIDLSGILQVTAGYGK  423


Range 2: 1 to 44

Score:92.8 bits(229), Expect:1e-18,
Method:Compositional matrix adjust.,
Identities:41/44(93%), Positives:41/44(93%), Gaps:0/44(0%)

Query  780  MNNFRKHVREADPECPRMCKMKFKPSSMKGFPISELYDTLGTTK  911
            MN FR HVREADPECPRMCKMKFK SSMKGFPISELYDTLGTTK
Sbjct  1    MNKFRNHVREADPECPRMCKMKFKASSMKGFPISELYDTLGTTK  44


>En/Spm-like transposon protein-like [Arabidopsis thaliana]
Sequence ID: BAB01349.1 Length: 714
Range 1: 39 to 347

Score:423 bits(1088), Expect:1e-137,
Method:Compositional matrix adjust.,
Identities:201/309(65%), Positives:237/309(76%), Gaps:6/309(1%)

Query  3    NEETKQIS----MDSEEETEPLRPLKMYFSPSDYLKPFKIGTKCYIHQVVNILESHLSSE  170
            +E T  IS     DS+EETE L+PLKMYF PS+Y KPFKI  KCY+H+ V +LE+HL
Sbjct  39   SEATNSISNDDRADSDEETEALQPLKMYFGPSEYTKPFKITAKCYLHKAVGLLETHLKES  98

Query  171  EKKWFWEHPQFKHFFHMHTDSNHKVMAMWMLLLRTACVDKKKECWFVVNGVPIRYSLQEL  350
            E KWF EHPQFKHFFHMH D NHKVM MW+L LRT C+DKKKE WF+VNGVPIRYSL+E
Sbjct  99   ELKWFLEHPQFKHFFHMHKDPNHKVMGMWLLFLRTTCLDKKKEVWFIVNGVPIRYSLEEF  158

Query  351  ALISGLYCHSYPKNYKAFGSLGFSGKHFGVGATVTYAKVKTKLLSMKKSSRERLKMAVLF  530
            AL+SGLYCH+YPK   + G   F+GK FG+GATV Y  V+ KLLSMKK S  RL++AVL+
Sbjct  159  ALMSGLYCHNYPKPLDSLGCTTFAGKMFGLGATVQYLDVENKLLSMKKPSDARLRVAVLY  218

Query  531  FLCSVIIGKRKTGEKAEAVEDFFLKSVEDLEWCKTFPWGRLAFDKNMKDIFHLMDHFDEE  710
            FLCSVI+GK KTG  A  VE FFL++V DLE CKTFPWGR AF+KN+KDIF+LM   +
Sbjct  219  FLCSVIVGKGKTGPNAPNVEKFFLRAVADLELCKTFPWGRFAFEKNVKDIFYLMKKCNGV  278

Query  711  LGAAWVFPSFVMQL--LAFETIPNLMNNFRKHVREADPECPRMCKMKFKPSSMKGFPISE  884
            +G   VFPSFVM L  LAFE IP L  NF + +  ADP+CPRMCKMKFK S+MKGFP+S+
Sbjct  279  VGPQKVFPSFVMPLEYLAFEAIPVLRENFCEDIESADPQCPRMCKMKFKSSTMKGFPMSD  338

Query  885  LYDTLGTTK  911
            +YD LGTTK
Sbjct  339  VYDKLGTTK  347


Range 2: 346 to 413

Score:62.0 bits(149), Expect:4e-09,
Method:Compositional matrix adjust.,
Identities:35/68(51%), Positives:50/68(73%), Gaps:0/68(0%)

Query  989   SQEIKSIMLPSPSEKELLKQIMddevghddvddMIVDGWTKRIIKENKPICFEELHIQDV  1168
             +++I+SI+ P+P EK LL++IMD E G +DVDD+I DGW KR++ E + ICFE L  +DV
Sbjct  346   TKDIQSILAPTPDEKLLLERIMDKECGVNDVDDLIADGWKKRLVDEERTICFEPLFNEDV  405

Query  1169  SSREIEAN  1192
             + R   AN
Sbjct  406   AHRSFVAN  413


>En/Spm-like transposon [Arabidopsis thaliana]
Sequence ID: NP_001331300.1 Length: 602
>En/Spm-like transposon [Arabidopsis thaliana]
Sequence ID: ANM69637.1 Length: 602
Range 1: 1 to 199

Score:402 bits(1034), Expect:4e-131,
Method:Compositional matrix adjust.,
Identities:199/222(90%), Positives:199/222(89%), Gaps:23/222(10%)

Query  1529  MYGRKRTRTTTAKEQEEKGEEIQVGFQETSERGRKKAKLTWKKVDIKGVTADQGNSITTN  1708
             MYGRKRTRTTTAKEQEEKGEEIQVGFQETSERGRKKAKLTWKKVDIKGVTADQGNSITTN
Sbjct  1     MYGRKRTRTTTAKEQEEKGEEIQVGFQETSERGRKKAKLTWKKVDIKGVTADQGNSITTN  60

Query  1709  ERVVVEPSSKVGSWKLASNLKSTRQYNVQVGSSLPPCSISCVDEPFSVSVSCKVNPGPLK  1888
             ERVVVEPSSKVGSWKLASNLKSTRQYNVQVGSSLPPCSISCVDEPFSVSVSCKVNPGPLK
Sbjct  61    ERVVVEPSSKVGSWKLASNLKSTRQYNVQVGSSLPPCSISCVDEPFSVSVSCKVNPGPLK  120

Query  1889  HVVEKYPEALENLLPGSTVQNYILEVKVYLLSIE*LHQIKLCTIFFMQVFDGYNNHVAEG  2068
             HVVEKYPEALENLLPGSTVQNYILE                       VFDGYNNHVAEG
Sbjct  121   HVVEKYPEALENLLPGSTVQNYILE-----------------------VFDGYNNHVAEG  157

Query  2069  TNVLICIEGYCINDSMGLNQKVNSCGCIDLSGILQVTAGYGK  2194
             TNVLICIEGYCINDSMGLNQKVNSCGCIDLSGILQVTAGYGK
Sbjct  158   TNVLICIEGYCINDSMGLNQKVNSCGCIDLSGILQVTAGYGK  199


>putative protein [Arabidopsis thaliana]
Sequence ID: CAB77996.1 Length: 715
>putative protein [Arabidopsis thaliana]
Sequence ID: CAB82107.1 Length: 715
Range 1: 172 to 481

Score:296 bits(758), Expect:4e-89,
Method:Compositional matrix adjust.,
Identities:147/311(47%), Positives:200/311(64%), Gaps:13/311(4%)

Query  36   EEETEPLRPLKMYFSPSDYLKPFKIGTKCYIHQVVNI---LESHLSSEEKKWFWEHPQFK  206
            +EETE + PL MYF PS+Y+K  K+ T+CYIH+++     LE  +S  E+ WF  HP F+
Sbjct  172  DEETEAVPPLSMYFPPSEYVKKIKLSTRCYIHELLTTFDKLEPPMSKSERLWFENHPSFQ  231

Query  207  HFFHMHTDSNHKVMAMWMLLLRTACVDKKKECWFVVNGVPIRYSLQELALISGLYCHSYP  386
            H FHM  D NH++M MWMLLLRTA +++KKE WF+VNGVPIRY + E ALIS   C +Y
Sbjct  232  HIFHMPRDPNHRLMGMWMLLLRTARIERKKEAWFIVNGVPIRYGILEHALISDFNCKNYK  291

Query  387  KNYKAFGSLGFSGKHFGVGATVTYAKVKTKLLSM---KKSSRERLKMAVLFFLCSVIIGK  557
              YK  G+L F  KHF     V    V+ KL+ M    + S+ERL+M VL+FL S+II
Sbjct  292  LGYKNTGNLDFKRKHFK-DTVVKREDVREKLIGMVPEGERSKERLRMMVLYFLSSIIIAP  350

Query  558  RKTGEKAEAVEDFFLKSVEDLEWCKTFPWGRLAFDKNMKDIFHLMDHFDEELGA----AW  725
             KTG+KA  V++FFLK + DL +C+ F WGR +FD  +  I H ++HF+  +       W
Sbjct  351  IKTGDKAPQVDEFFLKVMSDLTFCRNFQWGRYSFDYMLGTISHTVNHFNGSVTNNEKYIW  410

Query  726  VFPSFV--MQLLAFETIPNLMNNFRKHVREADPECPRMCKMKFKPSSMKGFPISELYDTL  899
              P F   M+LLAFE I  L   F + + EADP CPRMCK++FK + +KGFP+  +Y  L
Sbjct  411  PVPGFCLPMELLAFEAISQLREKFIEEIAEADPGCPRMCKVRFKKNHLKGFPLDTIYAEL  470

Query  900  GTTKVRNDLIV  932
            GTT+V + ++
Sbjct  471  GTTQVIDSIVT  481


>hypothetical protein [Arabidopsis thaliana]
Sequence ID: AAC23763.1 Length: 705
Range 1: 276 to 580

Score:289 bits(739), Expect:1e-86,
Method:Compositional matrix adjust.,
Identities:141/306(46%), Positives:200/306(65%), Gaps:10/306(3%)

Query  39   EETEPLRPLKMYFSPSDYLKPFKIGTKCYIHQVVNI---LESHLSSEEKKWFWEHPQFKH  209
            EE   ++PL MYF  S+Y K  K+ T+CYI +V+     LE  L++ EK +F EHP FKH
Sbjct  276  EEAMAMQPLGMYFPASEYTKKMKLATRCYISEVLKTFADLEHPLTNVEKNYFMEHPSFKH  335

Query  210  FFHMHTDSNHKVMAMWMLLLRTACVDKKKECWFVVNGVPIRYSLQELALISGLYCHSYPK  389
             +H+ +   HK+M MWML LRTA ++KKKE WFVVNGVPIRY ++E ALISG  C +YP
Sbjct  336  IYHLPSGYTHKLMGMWMLFLRTASIEKKKEVWFVVNGVPIRYGIREHALISGFNCKAYPA  395

Query  390  NYKAFGSLGFSGKHFGVGATVTYAKVKTKLLSMKKS-SRERLKMAVLFFLCSVIIGKRKT  566
            NY++ G++ F+ ++F  G  +    VKTKL+ M+ + S++RL+MAVL+FL S+I    KT
Sbjct  396  NYQSAGNMNFANRYFKTG-VIRREDVKTKLMEMEPARSKDRLRMAVLYFLTSIIAVPTKT  454

Query  567  GEKAEAVEDFFLKSVEDLEWCKTFPWGRLAFDKNMKDIFHLMDHFD---EELGAAWVFPS  737
            GE+A  ++DF +++  DL +CKTFPWGR +F+  +K I H +DHF+       + W  P
Sbjct  455  GERASPIDDFCVRAASDLTFCKTFPWGRYSFEYMLKSISHTLDHFNGVVPNTQSPWPVPG  514

Query  738  FVMQL--LAFETIPNLMNNFRKHVREADPECPRMCKMKFKPSSMKGFPISELYDTLGTTK  911
            F + L  LAFE IP+L   F +    +   CPRMCK+ FK + MKGF + ++   LGTT+
Sbjct  515  FCVPLEFLAFEAIPSLRERFIEEKEGSHAGCPRMCKVSFKRTEMKGFTLEQINHVLGTTE  574

Query  912  VRNDLI  929
            V   +I
Sbjct  575  VIESII  580


>unnamed protein product [Arabidopsis thaliana]
Sequence ID: BAB02522.1 Length: 672
Range 1: 208 to 521

Score:273 bits(699), Expect:4e-81,
Method:Compositional matrix adjust.,
Identities:140/318(44%), Positives:200/318(62%), Gaps:13/318(4%)

Query  3    NEETKQISMDSEEETEPLRPLKMYFSPSDYLKPFKIGTKCYIHQVVNI---LESHLSSEE  173
            +E+   + M   EE   ++PL MYF  S+Y K  K+ T+CYI +V+     LE  L+  E
Sbjct  208  DEDIANVDM---EEAMAMQPLGMYFPVSEYPKKMKLATRCYISEVLKTFADLEHPLTHVE  264

Query  174  KKWFWEHPQFKHFFHMHTDSNHKVMAMWMLLLRTACVDKKKECWFVVNGVPIRYSLQELA  353
            K +F EHP FKH +H+ +   HK+M MWML LRTA ++KKKE WFVVNGVPIRY ++E A
Sbjct  265  KNYFMEHPSFKHIYHLPSGYTHKLMRMWMLFLRTASIEKKKEVWFVVNGVPIRYGIREHA  324

Query  354  LISGLYCHSYPKNYKAFGSLGFSGKHFGVGATVTYAKVKTKLLSMKKS-SRERLKMAVLF  530
            LISG  C +YP +Y++ G++ F+ ++F  G  +    VKTKL+ M+ + S++RL+MA L+
Sbjct  325  LISGFNCKAYPASYQSAGNMNFANRYFKTG-VIRREDVKTKLMEMEPARSKDRLRMAALY  383

Query  531  FLCSVIIGKRKTGEKAEAVEDFFLKSVEDLEWCKTFPWGRLAFDKNMKDIFHLMDHFD--  704
            FL S+I+   KTGE+A  ++DF ++S  DL +CK FP GR +F+  +K I H +DHF+
Sbjct  384  FLTSIIVMPTKTGERASPIDDFCVRSASDLTFCKIFPSGRYSFEYMLKSISHKLDHFNGV  443

Query  705  -EELGAAWVFPSFVMQL--LAFETIPNLMNNFRKHVREADPECPRMCKMKFKPSSMKGFP  875
                 + W  P F + L  LAFE IP+L   F +    A   C RMCK+ FK   MKGF
Sbjct  444  VPNTQSPWPVPGFCVPLEFLAFEAIPSLRERFIEEKEGAHAGCQRMCKVNFKRIEMKGFT  503

Query  876  ISELYDTLGTTKVRNDLI  929
            + ++   LGTT+V   +I
Sbjct  504  LEQINHVLGTTEVIESII  521


>RecName: Full=Uncharacterized protein At3g43530 [Arabidopsis thaliana]
Sequence ID: Q9M237.1 Length: 615
>putative protein [Arabidopsis thaliana]
Sequence ID: CAB81793.1 Length: 615
Range 1: 47 to 263

Score:258 bits(659), Expect:2e-79,
Method:Compositional matrix adjust.,
Identities:139/294(47%), Positives:165/294(56%), Gaps:77/294(26%)

Query  30   DSEEETEPLRPLKMYFSPSDYLKPFKIGTKCYIHQVVNILESHLSSEEKKWFWEHPQFKH  209
            DS+EETE L+PLKMYF PSDY KPFKI  KCY+H+ V +LESHL   E KWF EHPQFKH
Sbjct  47   DSDEETEALQPLKMYFGPSDYTKPFKITAKCYLHKAVGLLESHLEESELKWFLEHPQFKH  106

Query  210  FFHMHTDSNHKVMAMWMLLLRTACVDKKKECWFVVNGVPIRYSLQELALISGLYCHSYPK  389
            FFHMH D NHKVM MW+L +RT C+DKK                      S +YC
Sbjct  107  FFHMHKDPNHKVMGMWLLFIRTTCLDKKG---------------------SMVYCQWR--  143

Query  390  NYKAFGSLGFSGKHFGVGATVTYAKVKTKLLSMKKSSRERLKMAVLFFLCSVIIGKRKTG  569
                      +   FG GAT+ Y  V+ KLLSMKK S  RL++ VL+FLC+   G
Sbjct  144  ----------TDPMFGPGATIQYPDVEKKLLSMKKPSEARLRVVVLYFLCNRWKG-----  188

Query  570  EKAEAVEDFFLKSVEDLEWCKTFPWGRLAFDKNMKDIFHLMDHFDEELGAAWVFPSFVMQ  749
                         V DLE CKTFPWG+ AF++N
Sbjct  189  ------------VVADLELCKTFPWGKNAFEEN---------------------------  209

Query  750  LLAFETIPNLMNNFRKHVREADPECPRMCKMKFKPSSMKGFPISELYDTLGTTK  911
             LAFE IP L  NF + +  ADP+CPRMCKMKFK S+MKGFP+S++YD LGTTK
Sbjct  210  YLAFEAIPVLRKNFCEDIESADPQCPRMCKMKFKSSTMKGFPMSDVYDKLGTTK  263


Range 2: 263 to 407

Score:60.5 bits(145), Expect:2e-79,
Method:Compositional matrix adjust.,
Identities:51/145(35%), Positives:80/145(55%), Gaps:19/145(13%)

Query  1001  KSIMLPSPSEKELLKQIMddevghddvddMIVDGWTKRIIKENKPICFEELHIQDVSSRE  1180
             KSI+ P+P E  LLK+IMD E G +DVDD+I DGW KR++ E + ICFE L  +DV+ +
Sbjct  263   KSILAPTPDENLLLKRIMDKECGVNDVDDLIADGWKKRLVDEERTICFEPLFNEDVAHQS  322

Query  1181  IEAN---------PknanvvgvrvgrkkvalkkvSEKGIVE---QLQKQLEDG-------  1303
               AN         P+ A V     G+   AL   S++G+ E   +++  +E+G
Sbjct  323   FVANNAPSTVVQAPRKAAVEKKGKGKTAAALTSPSDEGLTEVVNEMKNLMENGFKSMNKR  382

Query  1304  LNRIYMKIDDMDTRLTYVESCVKNL  1378
             +     K ++ D RL  +E+ +K++
Sbjct  383   MTNFSKKYEEQDKRLKLMETAIKSI  407


>unnamed protein product [Arabidopsis thaliana]
Sequence ID: BAB09148.1 Length: 628
Range 1: 140 to 424

Score:249 bits(635), Expect:2e-72,
Method:Compositional matrix adjust.,
Identities:132/291(45%), Positives:178/291(61%), Gaps:11/291(3%)

Query  45   TEPLRPLKMYFSPSDYLKPFKIGTKCYIHQVVNILESHLSSEEKKWFWEHPQFKHFFHMH  224
             +PL P  MYF P+ Y K  KIGT+C + Q V  +E+  +  E KWF  H QFKH FHM
Sbjct  140  NQPLPPEVMYFDPTTYTKVCKIGTRCQLVQTVEFIETLDA--ELKWFRNHDQFKHIFHMP  197

Query  225  TDSNHKVMAMWMLLLRTACVDKKKECWFVVNGVPIRYSLQELALISGLYCHSYPKNYKAF  404
             + NH +  MWML++RTA  +  +ECWFVVNGVPIRYS++E AL++GL C  YPKNYK
Sbjct  198  KEPNHMIQGMWMLMVRTAKTELARECWFVVNGVPIRYSIREHALLTGLNCREYPKNYKTL  257

Query  405  GSLGFSGKHFGVGATVTYAKVKTKLLSMK-KSSRERLKMAVLFFLCSVIIGKRKTGEKAE  581
            GSL F  K F     +    V+ KL   K + S  RLK+A+L FL  V+    K   K
Sbjct  258  GSLKFVEKLFKRTEDIKIKDVEEKLEEFKSEKSTARLKLAILLFLAKVMKADSKGDSK--  315

Query  582  AVEDFFLKSVEDLEWCKTFPWGRLAFDKNMKDIFHLMDHFDE--ELGAAWVFPSFV--MQ  749
             +E+F L+ V+++  C+TFPWGR +F++ M+ +  +M +     +  A   F  F+  ++
Sbjct  316  -IEEFLLRIVDNVRACETFPWGRFSFEQCMEGVQRVMKNMKGVVKQKAQTAFYGFITPLE  374

Query  750  LLAFETIPNLMNNFRKHVREADPECPRMCKMKFKPSSMKGFPISELYDTLG  902
            +LAFE IP L   FR+ VR AD ECPRMCK KF  S MKGF + E+ + LG
Sbjct  375  ILAFECIPQLGKRFREVVR-ADKECPRMCKHKFSESCMKGFTLEEINEALG  424


>putative protein [Arabidopsis thaliana]
Sequence ID: CAB78060.1 Length: 666
Range 1: 140 to 424

Score:249 bits(637), Expect:3e-72,
Method:Compositional matrix adjust.,
Identities:132/291(45%), Positives:179/291(61%), Gaps:11/291(3%)

Query  45   TEPLRPLKMYFSPSDYLKPFKIGTKCYIHQVVNILESHLSSEEKKWFWEHPQFKHFFHMH  224
             +PL P  MYF P+ Y K  KIGT+C + Q V  +E+  +  E KWF  H QFKH FHM
Sbjct  140  NQPLPPEVMYFDPTTYTKVCKIGTRCQLVQTVEFIETLDA--ELKWFRNHDQFKHIFHMP  197

Query  225  TDSNHKVMAMWMLLLRTACVDKKKECWFVVNGVPIRYSLQELALISGLYCHSYPKNYKAF  404
             + NH +  MWML++RTA  + ++ECWFVVNGVPIRYS++E AL++GL C  YPKNYK
Sbjct  198  KEPNHMIQGMWMLMVRTAKTELERECWFVVNGVPIRYSIREHALLTGLNCREYPKNYKTL  257

Query  405  GSLGFSGKHFGVGATVTYAKVKTKLLSMK-KSSRERLKMAVLFFLCSVIIGKRKTGEKAE  581
            GSL F  K F     +    V+ KL   K + S  RLK+A+L FL  V+    K   K
Sbjct  258  GSLKFVEKLFKRTEDIKIKDVEEKLEEFKSEKSTARLKLAILLFLAKVMKADSKGDSK--  315

Query  582  AVEDFFLKSVEDLEWCKTFPWGRLAFDKNMKDIFHLMDHFDEEL--GAAWVFPSFV--MQ  749
             +E+F L+ V+++  C+TFPWGR +F++ M+ +  +M +    +   A   F  F+  ++
Sbjct  316  -IEEFLLRIVDNVRACETFPWGRFSFEQCMEGVRRVMKNMKGVVKPKAQTAFYGFITPLE  374

Query  750  LLAFETIPNLMNNFRKHVREADPECPRMCKMKFKPSSMKGFPISELYDTLG  902
            +LAFE IP L   FR+ VR AD ECPRMCK KF  S MKGF + E+ + LG
Sbjct  375  ILAFECIPQLGKRFREAVR-ADKECPRMCKHKFSESCMKGFTLEEINEALG  424


>contains similarity to T. cruzi 3' end fragment (PID:g161956) [Arabidopsis thaliana]
Sequence ID: AAB61080.1 Length: 707
Range 1: 95 to 401

Score:248 bits(634), Expect:1e-71,
Method:Compositional matrix adjust.,
Identities:132/313(42%), Positives:184/313(58%), Gaps:17/313(5%)

Query  36   EEETEPLRPLKMYFSPSDYLKPFKIGTKCYIHQVVNILESHLSSEEKKWFWEHPQFKHFF  215
            ++E +P++P +M+FSPS+Y+K  KIGT+C + Q+V  LE     E+  WF  HPQF+H F
Sbjct  95   DDEEQPMQPERMFFSPSEYVKTCKIGTRCTVQQIVKYLEEF--KEDLPWFKAHPQFRHVF  152

Query  216  HMHTDSNHKVMAMWMLLLRTACVDKKKECWFVVNGVPIRYSLQELALISGLYCHSYPKNY  395
            HM  + NH    +WMLLLRTA  +  +ECWFVVNGVPIRYS++E AL+ G  CH YP+
Sbjct  153  HMPEEKNHMTQGLWMLLLRTAQTEMDRECWFVVNGVPIRYSIKEHALLCGFDCHDYPEEL  212

Query  396  KAF-----GSLGFSGKHFGVGATVTYAKVKTKLLSMKKSSR----ERLKMAVLFFLCSVI  548
            +         L F+ K F   + +    V+ KL  +KK       +R K+A+L FLC VI
Sbjct  213  QPSMKIVDADLKFAKKIFKKVSGIKIVDVEKKLDELKKCGEKKKTDRKKLAILLFLCKVI  272

Query  549  IGKRKTGEKAEAVEDFFLKSVEDLEWCKTFPWGRLAFDKNMKDIFHLMDHFDEELGAAWV  728
              K K       ++ FFLK V+D+  C+TFPWGR  FD  M+ I  +M     +
Sbjct  273  AAKSKADGN---IDKFFLKIVDDVRACETFPWGRFTFDGCMEGIKSVMKSMKGKAKLQTC  329

Query  729  FPSFV--MQLLAFETIPNLMNNFRKHVREADPECPRMCKMKFKPSSMKGFPISELYDTLG  902
            F  F+  +++L FE IP+L   FR  VR  + ECPRMCK KF  S MKGF + E+ + LG
Sbjct  330  FSGFILPLEILPFEAIPHLGQKFRDPVR-FENECPRMCKAKFSSSLMKGFSLEEINEELG  388

Query  903  TTKVRNDLIVYYY  941
            + KV + ++   Y
Sbjct  389  SVKVISSVLEPDY  401


>F5J5.9 [Arabidopsis thaliana]
Sequence ID: AAF18635.1 Length: 656
Range 1: 130 to 434

Score:246 bits(627), Expect:5e-71,
Method:Compositional matrix adjust.,
Identities:134/311(43%), Positives:182/311(58%), Gaps:15/311(4%)

Query  45   TEPLRPLKMYFSPSDYLKPFKIGTKCYIHQVVNILESHLSSEEKKWFWEHPQFKHFFHMH  224
             +PL P  MYF P+ Y K  KIGT+C + Q V  +E+     E KWF  H QFKH FHM
Sbjct  130  NQPLPPEVMYFDPTTYTKVCKIGTRCQLVQTVEFIETL--DAELKWFRNHDQFKHIFHMP  187

Query  225  TDSNHKVMAMWMLLLRTACVDKKKECWFVVNGVPIRYSLQELALISGLYCHSYPKNYKAF  404
             + NH +  MWML++RTA  +  +ECWFVVNGVPIRYS++E AL++GL C  YPKNYK
Sbjct  188  KEPNHMIQGMWMLMVRTAKTELARECWFVVNGVPIRYSIREHALLTGLNCREYPKNYKTL  247

Query  405  GSLGFSGKHFGVGATVTYAKVKTKLLSMK-KSSRERLKMAVLFFLCSVIIGKRKTGEKAE  581
            GSL F  K F     +    V+ KL   K + S  RLK+A+L FL  V+    K   K
Sbjct  248  GSLKFVEKLFKRTEDIKIKDVEEKLEEFKSEKSTARLKLAILLFLAKVMKADSKGDSK--  305

Query  582  AVEDFFLKSVEDLEWCKTFPWGRLAFDKNMKDIFHLMDHFDE--ELGAAWVFPSFVM---  746
             +++F L+ V+++  C+TFPWGR +F++ M+ +  +M +     +  A   F  F+
Sbjct  306  -IKEFLLRIVDNVRACETFPWGRFSFEQCMEGVRRVMKNMKGVVKQKAQTAFYGFITPLE  364

Query  747  ---QLLAFETIPNLMNNFRKHVREADPECPRMCKMKFKPSSMKGFPISELYDTLGTTKVR  917
               ++LAFE IP L   FR+ VR AD ECPRMCK KF  S MKGF + E+ + LG   V
Sbjct  365  VSSRILAFECIPQLGKRFREAVR-ADKECPRMCKHKFSESCMKGFTLEEINEALGDITVS  423

Query  918  NDLIVYYYLKF  950
              L+   ++ F
Sbjct  424  FVLLKSMFIIF  434


>unknown protein [Arabidopsis thaliana]
Sequence ID: AAC17091.2 Length: 652
Range 1: 140 to 424

Score:243 bits(619), Expect:6e-70,
Method:Compositional matrix adjust.,
Identities:129/291(44%), Positives:177/291(60%), Gaps:11/291(3%)

Query  45   TEPLRPLKMYFSPSDYLKPFKIGTKCYIHQVVNILESHLSSEEKKWFWEHPQFKHFFHMH  224
             +PL P  MYF P+ Y K  KIGT+C + Q V  +E+  +  E KWF  H QFKH FHM
Sbjct  140  NQPLPPEVMYFDPTTYTKVCKIGTRCQLVQTVEFIETLDA--ELKWFRNHDQFKHIFHMP  197

Query  225  TDSNHKVMAMWMLLLRTACVDKKKECWFVVNGVPIRYSLQELALISGLYCHSYPKNYKAF  404
             + NH +  MWML++RTA  + ++ECWFVVNGVPI YS++E AL++ L C  YPKNYK
Sbjct  198  KEPNHMIQGMWMLMVRTAKTELERECWFVVNGVPIGYSIREHALLTCLNCREYPKNYKTL  257

Query  405  GSLGFSGKHFGVGATVTYAKVKTKLLSMK-KSSRERLKMAVLFFLCSVIIGKRKTGEKAE  581
            GSL F  + F     +    V+ KL   K + S  RLK+A+L FL  V+    K   K
Sbjct  258  GSLKFVERLFKRTEDIKIKDVEEKLEEFKSEKSTARLKLAILLFLAKVMKADSKGDSK--  315

Query  582  AVEDFFLKSVEDLEWCKTFPWGRLAFDKNMKDIFHLMDHFDEEL--GAAWVFPSFV--MQ  749
             +E+F L+ V+++  C+TFPWGR +F++ M+ +  +M +    +   A   F  F+  ++
Sbjct  316  -IEEFLLRIVDNVRACETFPWGRFSFEQCMEGVRRVMKNMKGVVKPKAQTAFYGFITPLE  374

Query  750  LLAFETIPNLMNNFRKHVREADPECPRMCKMKFKPSSMKGFPISELYDTLG  902
            +LAFE IP L   FR+ VR AD ECPRMCK KF  S MKGF + E+ + LG
Sbjct  375  ILAFECIPQLGKRFREAVR-ADKECPRMCKHKFSESCMKGFTLEEINEALG  424


>hypothetical protein AXX17_ATUG02420 [Arabidopsis thaliana]
Sequence ID: OAO89237.1 Length: 665
Range 1: 138 to 422

Score:241 bits(616), Expect:2e-69,
Method:Compositional matrix adjust.,
Identities:129/291(44%), Positives:175/291(60%), Gaps:11/291(3%)

Query  45   TEPLRPLKMYFSPSDYLKPFKIGTKCYIHQVVNILESHLSSEEKKWFWEHPQFKHFFHMH  224
             +PL P  MYF P+ Y K  KIGT+C + Q V  +E+     E KWF  H QFKH FHM
Sbjct  138  NQPLPPEVMYFDPTTYTKVCKIGTRCQLVQTVEFIETL--DAELKWFRNHDQFKHIFHMP  195

Query  225  TDSNHKVMAMWMLLLRTACVDKKKECWFVVNGVPIRYSLQELALISGLYCHSYPKNYKAF  404
             + NH +  MWML++RTA  +  +ECWFVVNGVPIRYS++E AL++GL C  YPKNYK
Sbjct  196  KEPNHMIQGMWMLMVRTAKTELARECWFVVNGVPIRYSIKEHALLTGLNCREYPKNYKTL  255

Query  405  GSLGFSGKHFGVGATVTYAKVKTKLLSMK-KSSRERLKMAVLFFLCSVIIGKRKTGEKAE  581
            GSL F  K F     +    V+ KL   K + S  RLK+A+L FL  V+    K   K
Sbjct  256  GSLKFVEKLFKRTEDIKIKDVEEKLEEFKSEKSTARLKLAILLFLAKVMKADSKGDSK--  313

Query  582  AVEDFFLKSVEDLEWCKTFPWGRLAFDKNMKDIFHLMDHFDEEL--GAAWVFPSFV--MQ  749
             +E+  L+ V+++  C+TFPWGR +F++ M+ +  +M +  + +   A   F  F+  ++
Sbjct  314  -IEELLLRIVDNVRACETFPWGRFSFEQCMEGVQRVMKNMKDVVKPKAQTAFYGFITPLE  372

Query  750  LLAFETIPNLMNNFRKHVREADPECPRMCKMKFKPSSMKGFPISELYDTLG  902
            +LAFE IP L   FR+ V  AD ECPRMCK KF  S MK F + E+ + LG
Sbjct  373  ILAFECIPQLGKRFREVV-PADKECPRMCKHKFSESCMKRFTLEEINEALG  422


>T23E23.10 [Arabidopsis thaliana]
Sequence ID: AAF87156.1 Length: 762
Range 1: 121 to 405

Score:242 bits(618), Expect:6e-69,
Method:Compositional matrix adjust.,
Identities:130/294(44%), Positives:175/294(59%), Gaps:17/294(5%)

Query  45   TEPLRPLKMYFSPSDYLKPFKIGTKCYIHQVVNILESHLSSEEKKWFWEHPQFKHFFHMH  224
             +PL P  MYF P+ Y K  KIGT+C + Q V  +E+     E KWF  H QFKH FHM
Sbjct  121  NQPLPPEVMYFDPTTYTKVCKIGTRCQLVQTVEFIETL--DAELKWFRNHDQFKHIFHMP  178

Query  225  TDSNHKVMAMWMLLLRTACVDKKKECWFVVNGVPIRYSLQELALISGLYCHSYPKNYKAF  404
             + NH +  MWML++RTA  +  +ECWFVVNGVPIRYS++E AL++GL C  YPKNYK
Sbjct  179  KEPNHMIQGMWMLMVRTAKTELARECWFVVNGVPIRYSIREHALLTGLNCREYPKNYKTL  238

Query  405  GSLGFSGKHFGVGATVTYAKVKTKLLSMK-KSSRERLKMAVLFFLCSVIIGKRKTGEKAE  581
            GSL F  K F     +    V+ KL   K + S  RLK+ +L FL  V+    K   K
Sbjct  239  GSLKFVEKLFKRTEDIKIKDVEEKLEEFKSEKSTARLKLTILLFLAKVMKADSKGDSK--  296

Query  582  AVEDFFLKSVEDLEWCKTFPWGRLAFDKNMKDIFHLMDHFDEELG-----AAWVFPSFV-  743
             +E+  L+ V+++  C+TFPWGR +F++ M+ +  +M +    +G     A   F  F+
Sbjct  297  -IEELLLRIVDNVRACETFPWGRFSFEQCMEGVRRVMKNM---IGVVKPKAQTAFYGFIT  352

Query  744  -MQLLAFETIPNLMNNFRKHVREADPECPRMCKMKFKPSSMKGFPISELYDTLG  902
             +++LAFE IP L   FR+ V  AD ECPRMCK KF  S MKGF + E+ + LG
Sbjct  353  PLEILAFECIPQLGKRFREAV-PADKECPRMCKHKFSESCMKGFTLEEINEALG  405


>putative protein [Arabidopsis thaliana]
Sequence ID: CAB51205.1 Length: 664
Range 1: 140 to 424

Score:236 bits(601), Expect:3e-67,
Method:Compositional matrix adjust.,
Identities:127/291(44%), Positives:173/291(59%), Gaps:11/291(3%)

Query  45   TEPLRPLKMYFSPSDYLKPFKIGTKCYIHQVVNILESHLSSEEKKWFWEHPQFKHFFHMH  224
             +PL P  MYF P+ Y K  KIGT+C + Q V  +E+    EE KWF  H QFKH FHM
Sbjct  140  NQPLPPEVMYFDPTTYTKVCKIGTRCQLVQTVEFIETL--DEELKWFRNHDQFKHIFHMP  197

Query  225  TDSNHKVMAMWMLLLRTACVDKKKECWFVVNGVPIRYSLQELALISGLYCHSYPKNYKAF  404
             + NH +  MWML++RTA  +  +ECWFVVNGVPIRYS++E AL++GL C  YPKN K
Sbjct  198  KEPNHMIQGMWMLMVRTAKTELARECWFVVNGVPIRYSIREHALLTGLNCREYPKNNKTL  257

Query  405  GSLGFSGKHFGVGATVTYAKVKTKLLSMK-KSSRERLKMAVLFFLCSVIIGKRKTGEKAE  581
            GSL F  K F     +    V+ KL   K + S  RLK+A+L FL  V+    K   K
Sbjct  258  GSLKFVEKLFKRTEDIKIKDVEEKLEEFKSEKSTARLKLAILLFLAKVMKADSKGDSK--  315

Query  582  AVEDFFLKSVEDLEWCKTFPWGRLAFDKNMKDIFHLMDHFDEELGAA--WVFPSFV--MQ  749
             +E+  L+ V+++  C+TFPWGR +F++ M+ +  +M +    +       F  F+  ++
Sbjct  316  -IEELLLRIVDNVRACETFPWGRFSFEQCMEGVQRVMKNMKGVVKPKTHTAFYGFITPLK  374

Query  750  LLAFETIPNLMNNFRKHVREADPECPRMCKMKFKPSSMKGFPISELYDTLG  902
            +LAFE IP L   FR+ V  A+ EC RMCK KF  S MKGF + E+ + LG
Sbjct  375  ILAFECIPQLGKRFREAV-PANKECSRMCKHKFNESCMKGFSLEEINEALG  424


>unnamed protein product [Arabidopsis thaliana]
Sequence ID: BAB10960.1 Length: 628
Range 1: 140 to 424

Score:233 bits(593), Expect:2e-66,
Method:Compositional matrix adjust.,
Identities:127/291(44%), Positives:172/291(59%), Gaps:11/291(3%)

Query  45   TEPLRPLKMYFSPSDYLKPFKIGTKCYIHQVVNILESHLSSEEKKWFWEHPQFKHFFHMH  224
             +PL P  MYF P+ Y K  KIGT+C + Q V  +E+     E KWF  H QFKH FHM
Sbjct  140  NQPLPPEVMYFDPTTYTKVCKIGTRCQLVQTVEFIETL--DAELKWFKNHDQFKHIFHMP  197

Query  225  TDSNHKVMAMWMLLLRTACVDKKKECWFVVNGVPIRYSLQELALISGLYCHSYPKNYKAF  404
             + NH +  MWML++RTA  +  +ECWFVVNGVPIRY ++E AL++GL C  YPKNYK
Sbjct  198  KEPNHMIQGMWMLMVRTAKTELARECWFVVNGVPIRYLIKEHALLTGLNCREYPKNYKTL  257

Query  405  GSLGFSGKHFGVGATVTYAKVKTKLLSMK-KSSRERLKMAVLFFLCSVIIGKRKTGEKAE  581
            GSL F  K F     +    V+ KL   K + S  RLK+A+L FL  V+    K   K
Sbjct  258  GSLKFVEKLFKRTEDIKIKDVEEKLEEFKSEKSTARLKLAILLFLAKVMKADSKGDSK--  315

Query  582  AVEDFFLKSVEDLEWCKTFPWGRLAFDKNMKDIFHLMDHFDEEL--GAAWVFPSFV--MQ  749
             +E+  L+ V+++  C+TFPWGR +F++ M+ +  +M +    +   A   F  F+  ++
Sbjct  316  -IEELLLRIVDNIRACETFPWGRFSFEQCMEGVRRVMKNMKGVVKPKAQTAFYGFITLLE  374

Query  750  LLAFETIPNLMNNFRKHVREADPECPRMCKMKFKPSSMKGFPISELYDTLG  902
            +LAFE I  L   FR+ V  AD E PRMCK KF  S MKGF + E+ + LG
Sbjct  375  ILAFECILQLGKRFREAV-PADKERPRMCKHKFSESCMKGFTLEEINEALG  424


>hypothetical protein [Arabidopsis thaliana]
Sequence ID: AAD36941.1 Length: 735
>hypothetical protein [Arabidopsis thaliana]
Sequence ID: CAB77908.1 Length: 735
Range 1: 191 to 493

Score:234 bits(596), Expect:5e-66,
Method:Compositional matrix adjust.,
Identities:131/309(42%), Positives:178/309(57%), Gaps:11/309(3%)

Query  30   DSEEETEPLRPLKMYFSPSDYLKPFKIGTKCYIHQVVNILESHLSSEEKKWFWEHPQFKH  209
            D + +++PL P  M+F  S+Y K  KI ++C + Q V ++E     EE  WF  H QFKH
Sbjct  191  DDDLDSQPLPPQTMHFPFSEYGKVCKISSRCQVTQTVELIERKFK-EEAVWFKRHDQFKH  249

Query  210  FFHMHTDSNHKVMAMWMLLLRTACVDKKKECWFVVNGVPIRYSLQELALISGLYCHSYPK  389
             FHM  + NH+   MW+L+LRT   D  +E WFVVNGVPIRYS++E AL+SG  CHSYPK
Sbjct  250  IFHMPQEPNHQTQGMWVLMLRTVKTDLLEEAWFVVNGVPIRYSIKEHALLSGFDCHSYPK  309

Query  390  NYKAFGSLGFSGKHFGVGATVTYAKVKTKLLSMKKSSRERLKMAVLFFLCSVIIGKRKTG  569
                 G++ F  + F   + V    V  K   MK  S  RLK+ +L+FL  V+    K G
Sbjct  310  EMNTMGNINFVKRIFKKESGVKVVDVLAKCSKMKHGSN-RLKLVLLYFLVKVV----KAG  364

Query  570  EKAEA-VEDFFLKSVEDLEWCKTFPWGRLAFDKNMKDIFHLMDHFDE--ELGAAWVFPSF  740
             K +  +E+F L+ V DL  C+TFPWGR  F + M  I  +M + +   +  A   F  F
Sbjct  365  AKNDGNIEEFLLRIVGDLNACETFPWGRYIFLECMAGIRKMMKNMNGFVKPKAQPCFSGF  424

Query  741  V--MQLLAFETIPNLMNNFRKHVREADPECPRMCKMKFKPSSMKGFPISELYDTLGTTKV  914
            +  +++LA+E IP L   FR  VR A  +CPRMCK KFK  SMKG P+  +Y  LG TK
Sbjct  425  IVPLEILAYEAIPQLGLKFRVPVRSALSDCPRMCKHKFKECSMKGVPLEVIYKELGNTKD  484

Query  915  RNDLIVYYY  941
               ++V  Y
Sbjct  485  FPSILVPAY  493


>hypothetical protein [Arabidopsis thaliana]
Sequence ID: AAD19761.1 Length: 399
Range 1: 1 to 174

Score:220 bits(560), Expect:5e-64,
Method:Compositional matrix adjust.,
Identities:106/174(61%), Positives:131/174(75%), Gaps:0/174(0%)

Query  246  MAMWMLLLRTACVDKKKECWFVVNGVPIRYSLQELALISGLYCHSYPKNYKAFGSLGFSG  425
            M MW+L LRT+C++KK+E  F+VNGVPIRYSL+E AL+SGLYCHSY K + + GS  F G
Sbjct  1    MGMWLLFLRTSCLEKKREVPFIVNGVPIRYSLEEFALMSGLYCHSYTKPFNSLGSTTFVG  60

Query  426  KHFGVGATVTYAKVKTKLLSMKKSSRERLKMAVLFFLCSVIIGKRKTGEKAEAVEDFFLK  605
            K FG  AT+ Y  V+ KL+SMKK S  RL++AVL+FLCSVIIGK KTGE A +VE FFL+
Sbjct  61   KSFGPRATIQYPDVEEKLMSMKKPSEARLRVAVLYFLCSVIIGKGKTGENAPSVEKFFLR  120

Query  606  SVEDLEWCKTFPWGRLAFDKNMKDIFHLMDHFDEELGAAWVFPSFVMQLLAFET  767
             V DLE CK F WGR AFD+N+K+IF+LMD  +  +G   VFPSFVM L +  T
Sbjct  121  VVADLELCKIFSWGRYAFDENVKNIFYLMDQCNGVVGPQKVFPSFVMPLESILT  174


>hypothetical protein [Arabidopsis thaliana]
Sequence ID: AAD26890.1 Length: 349
>hypothetical protein [Arabidopsis thaliana]
Sequence ID: AAM15311.1 Length: 349
Range 1: 6 to 267

Score:208 bits(530), Expect:2e-60,
Method:Compositional matrix adjust.,
Identities:114/263(43%), Positives:156/263(59%), Gaps:16/263(6%)

Query  93   LKPFKIGTKCYIHQVVNILES---HLSSEEKKWFWEHPQFKHFFHMHTDSNHKVMAMWML  263
            +K  K+ T+ YIH+V+  L+     +S  E+ WF  H  F+H FHM  D NH++M MWML
Sbjct  6    VKKIKLSTRFYIHEVLTTLDKLQPAMSKSERAWFENHLSFQHIFHMPRDPNHRLMGMWML  65

Query  264  LLRTACVDKKKECWFVVNGVPIRYSLQELALISGLYCHSYP---KNYKAFGSLGFSGKHF  434
            LLRTA + +KKE WF+VNGV IRY + E ALISG  C +Y       K   +L F  KHF
Sbjct  66   LLRTAHIKRKKEPWFIVNGVSIRYGISEHALISGFNCKNYGFIHLGIKIRENLDFKKKHF  125

Query  435  GVGATVTYAKVKTKLLSM---KKSSRERLKMAVLFFLCSVIIGKRKTGEKAEAVEDFFLK  605
                 V +  V+ KL+ M    + S+ERL+M VL+FL ++II   KTG+KA  V++F LK
Sbjct  126  K-NIVVKHEDVREKLIGMVPLGERSKERLRMMVLYFLSNIIIAPIKTGDKAPQVDEFCLK  184

Query  606  SVEDLEWCKTFPWGRLAFDKNMKDIFHLMDHFDEELGA----AWVFPSFV--MQLLAFET  767
            +V DL +C+ F WGR +FD  + +I H ++HF+  +       W  P F   M+LLAFE
Sbjct  185  AVSDLTFCRNFQWGRYSFDYMLGNISHPVNHFNGSVTNNEKYIWPVPGFFLSMELLAFEA  244

Query  768  IPNLMNNFRKHVREADPECPRMC  836
            IP L     + +   D  CPRMC
Sbjct  245  IPQLRKKIIEEIAGPDQGCPRMC  267


>contains similarity to several [Arabidopsis thaliana hypothetical proteins including AC002983 and AF007271; my be a pseudogene [Arabidopsis thaliana]
Sequence ID: AAD38226.1 Length: 734
>hypothetical protein [Arabidopsis thaliana]
Sequence ID: CAB81122.1 Length: 734
Range 1: 156 to 401

Score:202 bits(515), Expect:6e-55,
Method:Compositional matrix adjust.,
Identities:116/302(38%), Positives:157/302(51%), Gaps:56/302(18%)

Query  36   EEETEPLRPLKMYFSPSDYLKPFKIGTKCYIHQVVNILESHLSSEEKKWFWEHPQFKHFF  215
            ++E +P++P +M+FSPS+Y+K  KIGT+C + Q V  LE     E+  WF  HPQF+H F
Sbjct  156  DDEEQPMQPKRMFFSPSEYVKTCKIGTRCTVQQTVKYLEEF--KEDLPWFKAHPQFRHVF  213

Query  216  HMHTDSNHKVMAMWMLLLRTACVDKKKECWFVVNGVPIRYSLQELALISGLYCHSYPKNY  395
            HM  + NHK   +WMLLLRTA  +  +ECWFVVNGVPIRYS++E AL+ G  CH YP+
Sbjct  214  HMPEEKNHKTQGLWMLLLRTAQTEMDRECWFVVNGVPIRYSIKEHALLCGFDCHDYPEEL  273

Query  396  KAFGSLGFSGKHFGVGATVTYAKVKTKLLSMKKSSRERLKMAVLFFLCSVIIGKRKTGEK  575
            +    +        V A + +AK                          VI  K K
Sbjct  274  QPSMKI--------VDADLKFAKK---------------------IFKKVIAAKSKADGN  304

Query  576  AEAVEDFFLKSVEDLEWCKTFPWGRLAFDKNMKDIFHLMDHFDEELGAAWVFPSFVMQLL  755
               ++ FFLK V+D+  C+TFPWG   FD  M+ I                       +L
Sbjct  305  ---IDKFFLKIVDDVRACETFPWGPFTFDGCMEGI---------------------KSIL  340

Query  756  AFETIPNLMNNFRKHVREADPECPRMCKMKFKPSSMKGFPISELYDTLGTTKVRNDLIVY  935
             FE IP+L   FR  VR  + ECP MCK KF  S MKGF + E+ + LG+ KV + ++
Sbjct  341  PFEAIPHLGQKFRDLVR-FENECPGMCKAKFSSSVMKGFSLEEINEELGSVKVISSVLES  399

Query  936  YY  941
             Y
Sbjct  400  DY  401


>hypothetical protein [Arabidopsis thaliana]
Sequence ID: AAO92060.1 Length: 249
Range 1: 6 to 237

Score:184 bits(466), Expect:2e-52,
Method:Compositional matrix adjust.,
Identities:99/233(42%), Positives:140/233(60%), Gaps:14/233(6%)

Query  93   LKPFKIGTKCYIHQVVNILES---HLSSEEKKWFWEHPQFKHFFHMHTDSNHKVMAMWML  263
            +K  K+ T+ YIH+V+  L+     +S  E+ WF  H  F+H FHM  D NH++M MWML
Sbjct  6    VKKIKLSTRFYIHEVLTTLDKLQPAMSKSERAWFENHLSFQHIFHMPRDPNHRLMGMWML  65

Query  264  LLRTACVDKKKECWFVVNGVPIRYSLQELALISGLYCHSYP---KNYKAFGSLGFSGKHF  434
            LLRTA + +KKE WF+VNGV IRY + E ALISG  C +Y       K   +L F  KHF
Sbjct  66   LLRTAHIKRKKEPWFIVNGVSIRYGISEHALISGFNCKNYGFIHLGIKIRENLDFKKKHF  125

Query  435  GVGATVTYAKVKTKLLSM---KKSSRERLKMAVLFFLCSVIIGKRKTGEKAEAVEDFFLK  605
                 V +  V+ KL+ M    + S+ERL+M VL+FL ++II   KTG+KA  V++F LK
Sbjct  126  K-NIVVKHEDVREKLIGMVPLGERSKERLRMMVLYFLSNIIIAPIKTGDKAPQVDEFCLK  184

Query  606  SVEDLEWCKTFPWGRLAFDKNMKDIFHLMDHFDEELGA----AWVFPSFVMQL  752
            +V DL +C+ F WGR +FD  + +I H ++HF+  +       W  P F + +
Sbjct  185  AVSDLTFCRNFQWGRYSFDHMLGNISHPVNHFNGSVTNNEKYIWPVPGFFLSM  237


>En/Spm-like transposon, putative (DUF1985) [Arabidopsis thaliana]
Sequence ID: NP_178683.2 Length: 249
>En/Spm-like transposon, putative (DUF1985) [Arabidopsis thaliana]
Sequence ID: AEC06007.1 Length: 249
Range 1: 6 to 237

Score:183 bits(465), Expect:2e-52,
Method:Compositional matrix adjust.,
Identities:99/233(42%), Positives:140/233(60%), Gaps:14/233(6%)

Query  93   LKPFKIGTKCYIHQVVNILES---HLSSEEKKWFWEHPQFKHFFHMHTDSNHKVMAMWML  263
            +K  K+ T+ YIH+V+  L+     +S  E+ WF  H  F+H FHM  D NH++M MWML
Sbjct  6    VKKIKLSTRFYIHEVLTTLDKLQPAMSKSERAWFENHLSFQHIFHMPRDPNHRLMGMWML  65

Query  264  LLRTACVDKKKECWFVVNGVPIRYSLQELALISGLYCHSYP---KNYKAFGSLGFSGKHF  434
            LLRTA + +KKE WF+VNGV IRY + E ALISG  C +Y       K   +L F  KHF
Sbjct  66   LLRTAHIKRKKEPWFIVNGVSIRYGISEHALISGFNCKNYGFIHLGIKIRENLDFKKKHF  125

Query  435  GVGATVTYAKVKTKLLSM---KKSSRERLKMAVLFFLCSVIIGKRKTGEKAEAVEDFFLK  605
                 V +  V+ KL+ M    + S+ERL+M VL+FL ++II   KTG+KA  V++F LK
Sbjct  126  K-NIVVKHEDVREKLIGMVPLGERSKERLRMMVLYFLSNIIIAPIKTGDKAPQVDEFCLK  184

Query  606  SVEDLEWCKTFPWGRLAFDKNMKDIFHLMDHFDEELGA----AWVFPSFVMQL  752
            +V DL +C+ F WGR +FD  + +I H ++HF+  +       W  P F + +
Sbjct  185  AVSDLTFCRNFQWGRYSFDYMLGNISHPVNHFNGSVTNNEKYIWPVPGFFLSM  237


>hypothetical protein At2g06420 [Arabidopsis thaliana]
Sequence ID: AAY78684.1 Length: 249
Range 1: 6 to 237

Score:183 bits(464), Expect:3e-52,
Method:Compositional matrix adjust.,
Identities:99/233(42%), Positives:140/233(60%), Gaps:14/233(6%)

Query  93   LKPFKIGTKCYIHQVVNILES---HLSSEEKKWFWEHPQFKHFFHMHTDSNHKVMAMWML  263
            +K  K+ T+ YIH+V+  L+     +S  E+ WF  H  F+H FHM  D NH++M MWML
Sbjct  6    VKKIKLSTRFYIHEVLTTLDKLQPAMSKSERAWFENHLSFQHIFHMPRDPNHRLMGMWML  65

Query  264  LLRTACVDKKKECWFVVNGVPIRYSLQELALISGLYCHSYP---KNYKAFGSLGFSGKHF  434
            LLRTA + +KKE WF+VNGV IRY + E ALISG  C +Y       K   +L F  KHF
Sbjct  66   LLRTAHIKRKKEPWFIVNGVSIRYGISEHALISGFNCKNYGFIHLGIKIRENLDFKKKHF  125

Query  435  GVGATVTYAKVKTKLLSM---KKSSRERLKMAVLFFLCSVIIGKRKTGEKAEAVEDFFLK  605
                 V +  V+ KL+ M    + S+ERL+M VL+FL ++II   KTG+KA  V++F LK
Sbjct  126  K-NIVVKHEDVREKLIGMVPLGERSKERLRMMVLYFLSNIIIAPIKTGDKAPQVDEFCLK  184

Query  606  SVEDLEWCKTFPWGRLAFDKNMKDIFHLMDHFDEELGA----AWVFPSFVMQL  752
            +V DL +C+ F WGR +FD  + +I H ++HF+  +       W  P F + +
Sbjct  185  AVSDLTFCRNFQWGRYSFDYMLGNISHPVNHFNGSVTNNEKYIWPVPGFFLSM  237


>hypothetical protein [Arabidopsis thaliana]
Sequence ID: AAD25589.1 Length: 535
Range 1: 137 to 346

Score:186 bits(473), Expect:2e-50,
Method:Compositional matrix adjust.,
Identities:100/227(44%), Positives:133/227(58%), Gaps:18/227(7%)

Query  234  NHKVMAMWMLLLRTACVDKKKECWFVVNGVPIRYSLQELALISGLYCHSYPKNYKAFGSL  413
            NH +  MWML++RTA  + ++ECWFVVNGVPIRYS++E AL++GL C  YPKNYK  GSL
Sbjct  137  NHMIQGMWMLMVRTAKTELERECWFVVNGVPIRYSIKEHALLTGLNCREYPKNYKTLGSL  196

Query  414  GFSGKHFGVGATVTYAKVKTKLLSMK-KSSRERLKMAVLFFLCSVIIGKRKTGEKAEAVE  590
             F  K F     +    V+ KL   K + S  RLK+A+L FL  V+    K   K   +E
Sbjct  197  KFVEKLFKRTEDIKIKDVEEKLEEFKSEKSTARLKLAILLFLAKVMKADSKRDSK---IE  253

Query  591  DFFLKSVEDLEWCKTFPWGRLAFDKNMKDIFHLMDHFDEELGAAWVFPSFVMQLLAFETI  770
            +  L+ V+++  C+TFPWGR +F++ M+             G   V  +    +LAFE I
Sbjct  254  ELLLRIVKNVRACETFPWGRFSFEQCME-------------GVQRVMKNMKGVILAFECI  300

Query  771  PNLMNNFRKHVREADPECPRMCKMKFKPSSMKGFPISELYDTLGTTK  911
            P L   FR+ V  AD ECPRMCK KF  S MKGF + E+ + LG  K
Sbjct  301  PQLGKRFREAV-PADKECPRMCKHKFNESCMKGFTLKEINEALGDIK  346


>putative protein [Arabidopsis thaliana]
Sequence ID: CAB86477.1 Length: 585
Range 1: 108 to 327

Score:186 bits(473), Expect:4e-50,
Method:Compositional matrix adjust.,
Identities:97/238(41%), Positives:139/238(58%), Gaps:18/238(7%)

Query  39   EETEPLRPLKMYFSPSDYLKPFKIGTKCYIHQVVNILESHLSSEEKKWFWEHPQFKHFFH  218
            E+T+PL P   YF  ++Y +  K+ ++   ++ +   +  L+ +EK WF  H QFKH +H
Sbjct  108  EDTQPLPPEMFYFKQTNYTECCKLSSRSEENRTMKEWDDILAEDEKNWFRTHLQFKHIWH  167

Query  219  MHTDSNHKVMAMWMLLLRTACVDKKKECWFVVNGVPIRYSLQELALISGLYCHSYPKNYK  398
            MH + NHK   MWMLLL TA ++    CWFVVNGVP+RYS +E  L+ G  C+ YP  YK
Sbjct  168  MHREENHKYTHMWMLLLCTAPMEIYSVCWFVVNGVPVRYSFREHGLLCGFDCYIYPPKYK  227

Query  399  AFGSLGFSGKHFGVGATVTYAKVKTKLLSMKKSSRERLKMAVLFFLCSVIIGKRKTGEKA  578
              GS                  V+ KL SM +   ER +MA+LFF+ +VI  K+K G
Sbjct  228  NLGSESIKA-------------VREKLHSMGECG-ERKQMAILFFVSTVITPKKKFG---  270

Query  579  EAVEDFFLKSVEDLEWCKTFPWGRLAFDKNMKDIFHLMDHFDEELGAAWVFPSFVMQL  752
              ++ F  + V+DL+ C+TFPWGR  FD N+K+IFH+M +F   +   W FP F++ L
Sbjct  271  -FIDAFIYRIVDDLDACETFPWGRYTFDDNIKNIFHMMKYFKGRVQQTWCFPGFLIPL  327


>hypothetical protein AXX17_AT5G24030 [Arabidopsis thaliana]
Sequence ID: OAO92751.1 Length: 457
Range 1: 41 to 137

Score:181 bits(458), Expect:5e-49,
Method:Compositional matrix adjust.,
Identities:87/120(73%), Positives:92/120(76%), Gaps:23/120(19%)

Query  1835  DEPFSVSVSCKVNPGPLKHVVEKYPEALENLLPGSTVQNYILEVKVYLLSIE*LHQIKLC  2014
             DEPFSVSV+CKVNPGPLKHV+E YPEALENLLPGSTVQ+YILEV
Sbjct  41    DEPFSVSVACKVNPGPLKHVLEMYPEALENLLPGSTVQDYILEV----------------  84

Query  2015  TIFFMQVFDGYNNHVAEGTNVLICIEGYCINDSMGLNQKVNSCGCIDLSGILQVTAGYGK  2194
                    FDGYNNHVAEGTNVLICIEG CI DSMG N+KVN CGC+DLSGILQVTAGYGK
Sbjct  85    -------FDGYNNHVAEGTNVLICIEGCCIKDSMGFNRKVNICGCVDLSGILQVTAGYGK  137


>putative protein [Arabidopsis thaliana]
Sequence ID: CAB83145.1 Length: 525
Range 1: 47 to 276

Score:178 bits(451), Expect:1e-47,
Method:Compositional matrix adjust.,
Identities:101/280(36%), Positives:141/280(50%), Gaps:50/280(17%)

Query  87   DYLKPFKIGTKCYIHQVVNILESHLSSEEKKWFWEHPQFKHFFHMHTDSNHKVMAMWMLL  266
            +Y K  K+GT+C + Q V  LE     E+  WF  H QF+H FHM  + NH    MWMLL
Sbjct  47   EYQKTCKVGTRCTVQQTVKYLEGF--KEDLPWFKAHSQFRHVFHMPEEKNHMTQGMWMLL  104

Query  267  LRTACVDKKKECWFVVNGVPIRYSLQELALISGLYCHSYPKNYKAFGSLGFSGKHFGVGA  446
            LRT+  +  ++CWFVVNGVPIRYS++E AL  G  C+ YPK  KA
Sbjct  105  LRTSQTEMDRKCWFVVNGVPIRYSIKEHALFCGFECYEYPKEKKA---------------  149

Query  447  TVTYAKVKTKLLSMKKSSRERLKMAVLFFLCSVIIGKRKTGEKAEAVEDFFLKSVEDLEW  626
                               +R K+++L FLC VI+ K K       ++ F LK ++D+
Sbjct  150  -------------------DRKKLSILLFLCKVIVAKSKVDGN---IDRFLLKIIDDVHA  187

Query  627  CKTFPWGRLAFDKNMKDIFHLMDHFDEELGAAWVFPSFVMQLLAFETIPNLMNNFRKHVR  806
            C+TFPW R  FD  M+ I  ++++ + +      F  F+  L        L   FR  +R
Sbjct  188  CETFPWDRFTFDGCMEGIKSIINNMNGKAKVETCFSGFIFPL-------ELGQKFRDPIR  240

Query  807  EADPECPRMCKMKFKPSSMKGFPISELYDTLGTTKVRNDL  926
              + ECPRMC  KF  S MK F + E+ + LG+    ND+
Sbjct  241  -FENECPRMCNSKFSNSVMKEFSLEEIIEELGSV---NDI  276


>En/Spm-like transposon protein [Arabidopsis thaliana]
Sequence ID: AAC26678.1 Length: 518
Range 1: 3 to 141

Score:145 bits(365), Expect:3e-45,
Method:Compositional matrix adjust.,
Identities:74/139(53%), Positives:94/139(67%), Gaps:4/139(2%)

Query  507  RLKMAVLFFLCSVIIGKRKTGEK--AEAVEDFFLKSVEDLEWCKTFPWGRLAFDKNMKDI  680
            R K A +     V++ +    E   A A E FFL++V DLE CKTFPWGR AF++N+KDI
Sbjct  3    RTKNAGVPLAAEVVVSQTINDEVVVASAAEKFFLRAVADLELCKTFPWGRFAFEENVKDI  62

Query  681  FHLMDHFDEELGAAWVFPSFVMQL--LAFETIPNLMNNFRKHVREADPECPRMCKMKFKP  854
            F+L+   +E +G   VFPSFVM L  LAFE IP L   F + +  ADP+C RMCK KFK
Sbjct  63   FYLIKKCNEVVGPQKVFPSFVMPLEYLAFEAIPVLRKIFCEDIESADPQCSRMCKRKFKS  122

Query  855  SSMKGFPISELYDTLGTTK  911
            S+MKGFP+S++YD LGTTK
Sbjct  123  STMKGFPMSDVYDKLGTTK  141


Range 2: 140 to 288

Score:59.3 bits(142), Expect:3e-45,
Method:Compositional matrix adjust.,
Identities:50/149(34%), Positives:82/149(55%), Gaps:19/149(12%)

Query  989   SQEIKSIMLPSPSEKELLKQIMddevghddvddMIVDGWTKRIIKENKPICFEELHIQDV  1168
             +++I+SI+  +P EK LL++IMD E   +DVDD+I DGW KR++ E + ICFE L  +DV
Sbjct  140   TKDIQSILASTPDEKLLLERIMDKECEVNDVDDLIADGWKKRLVDEERTICFEPLLNEDV  199

Query  1169  SSREIEAN---------PknanvvgvrvgrkkvalkkvSEKGIVE---QLQKQLEDGLNR  1312
             + R   AN         P+ A V     G+   AL   S+ G+ E   +++  +E+G
Sbjct  200   AHRSFVANKALSTVVKAPRKAAVEKKGKGKAAAALTSPSDGGLTEVVNEMKNLMENGFKS  259

Query  1313  IYMKIDDM-------DTRLTYVESCVKNL  1378
             +  ++ D        D RL  +E+ +K++
Sbjct  260   MNKRMKDFCKKYEEQDKRLKLMEAAIKSI  288


>putative protein [Arabidopsis thaliana]
Sequence ID: CAB72487.1 Length: 690
Range 1: 191 to 461

Score:174 bits(440), Expect:4e-45,
Method:Compositional matrix adjust.,
Identities:110/310(35%), Positives:156/310(50%), Gaps:45/310(14%)

Query  30   DSEEETEPLRPLKMYFSPSDYLKPFKIGTKCYIHQVVNILESHLSSEEKKWFWEHPQFKH  209
            D + +++PL    M+F  S+Y K  KI ++C + Q + ++E     E
Sbjct  191  DDDLDSQPLPQQTMHFPFSEYGKVCKISSRCQVTQTIELIERKFKEE-------------  237

Query  210  FFHMHTDSNHKVMAMWMLLLRTACVDKK-KECWFVVNGVPIRYSLQELALISGLYCHSYP  386
                                   C D   +E WFVVNGVPIRYS++E AL+SG  CHSYP
Sbjct  238  ---------------------AVCEDSLLEEAWFVVNGVPIRYSIKEHALLSGFDCHSYP  276

Query  387  KNYKAFGSLGFSGKHFGVGATVTYAKVKTKLLSMKKSSRERLKMAVLFFLCSVIIGKRKT  566
            K     G++ F  + F   + +    V  K   MK  S +RLK+ +L+FL  V+    K
Sbjct  277  KEMNTMGNINFVKRIFKKESGIKVVDVLAKCSKMKHGS-DRLKLVLLYFLVKVV----KA  331

Query  567  GEKAEA-VEDFFLKSVEDLEWCKTFPWGRLAFDKNMKDIFHLMDHFDE--ELGAAWVFPS  737
            G K +  +E+F L+ V DL  C+TFPWGR  F + M  I  +M + +   +  A   F
Sbjct  332  GAKNDGNIEEFLLRIVGDLNACETFPWGRYTFLECMAGIRKMMKNMNGFVKPKAQPCFSG  391

Query  738  FV--MQLLAFETIPNLMNNFRKHVREADPECPRMCKMKFKPSSMKGFPISELYDTLGTTK  911
            F+  +++L +E IP L   FR HVR A  +CPRMCK KFK  SMKG P+  +Y  LG TK
Sbjct  392  FIVPLEILPYEAIPQLGLKFRVHVRSALSDCPRMCKHKFKECSMKGVPLEVIYKELGNTK  451

Query  912  VRNDLIVYYY  941
                ++V  Y
Sbjct  452  DFPSILVPAY  461


>hypothetical protein [Arabidopsis thaliana]
Sequence ID: AAD28050.1 Length: 398
Range 1: 1 to 223

Score:164 bits(414), Expect:1e-43,
Method:Compositional matrix adjust.,
Identities:94/234(40%), Positives:135/234(57%), Gaps:15/234(6%)

Query  63   LKMYFSPSDYLKPFKIGTKCYIHQVVNILESHLSSEEKKWFWEHPQFKHFFHMHTDSNHK  242
            + ++F+P  Y K  KI T+C I     I+ S L   +KKWF  + QFKH +HM   S +K
Sbjct  1    MSLFFTPDGYEKILKISTRCTIGNTFAIIGSMLKENKKKWFRTNKQFKHIWHMDRHSKNK  60

Query  243  VMAMWMLLLRTACVDKKKECWFVVNGVPIRYSLQELALISGLYCHSYPKNYKA--FGSLG  416
            V  M MLL++TA   KK+ CWFVVN VPI YSL+E ALI+GL  H +  ++K   FGS
Sbjct  61   VHGMLMLLMQTASTQKKRVCWFVVNDVPICYSLREHALITGLDFHQFELDFKTRNFGSFD  120

Query  417  FSGKHFGVGATVTYAKVKTKLLSMKKS-SRERLKMAVLFFLCSVIIGKRKTGEKAEAVED  593
            F  K +G    V    V+  L SM+     ++L++AVL FLC+++ G+R+ G    ++
Sbjct  121  FVEKVYGT-QVVNVKDVEDMLKSMEDECDGDQLRVAVLLFLCAIVRGRRRFG----SIHS  175

Query  594  FFLKSVEDLEWCKTFPWGRLAFDKNMKDIFHLM-DHFDEELGAAWVFPSFVMQL  752
            F LK V      K FP G+  F+  M+ I HLM    D ++G  ++FP F++ L
Sbjct  176  FILKIV------KKFPSGQNTFEDTMEKIVHLMKKRLDGKVGIDYLFPGFIIPL  223


>unnamed protein product [Arabidopsis thaliana]
Sequence ID: BAB10394.1 Length: 1634
Range 1: 973 to 1100

Score:158 bits(400), Expect:4e-39,
Method:Compositional matrix adjust.,
Identities:81/128(63%), Positives:99/128(77%), Gaps:4/128(3%)

Query  1823  ISCVDEPFSVSVSCKVNPGPLKHVVEKYPEALENLLPGSTVQNYILEV-KVYLLSIE*LH  1999
             I  +D PFSVSV CKVNPGPLK V    P+ALENLLP STV+++ILEV  + L ++   +
Sbjct  973   IRAMDNPFSVSVPCKVNPGPLKRVAVNNPKALENLLPDSTVEDFILEVYNIGLCTLSNRN  1032

Query  2000  QIKL---CTIFFMQVFDGYNNHVAEGTNVLICIEGYCINDSMGLNQKVNSCGCIDLSGIL  2170
             Q+ L    TI FMQ+FDGYNNHVAEGT+VLI I+GY I D MG+N+KV+S GCI+LSGIL
Sbjct  1033  QLFLRIKLTIIFMQLFDGYNNHVAEGTDVLIHIDGYRIEDWMGINRKVDSRGCINLSGIL  1092

Query  2171  QVTAGYGK  2194
             +VT GYGK
Sbjct  1093  KVTEGYGK  1100


>hypothetical protein [Arabidopsis thaliana]
Sequence ID: AAD25845.1 Length: 564
>hypothetical protein [Arabidopsis thaliana]
Sequence ID: AAM15463.1 Length: 564
Range 1: 224 to 434

Score:150 bits(380), Expect:6e-38,
Method:Compositional matrix adjust.,
Identities:96/296(32%), Positives:137/296(46%), Gaps:89/296(30%)

Query  39   EETEPLRPLKMYFSPSDYLKPFKIGTKCYIHQVVNI---LESHLSSEEKKWFWEHPQFKH  209
            EE   ++PL MYF  S+Y K  K+ T+CYI +V+     LE  L++ E  +F EHP FKH
Sbjct  224  EEAMAMQPLGMYFPASEYTKKMKLATRCYISEVLKTFADLEHPLTNVENNYFMEHPSFKH  283

Query  210  FFHMHTDSNHKVMAMWMLLLRTACVDKKKECWFVVNGVPIRYSLQELALISGLYCHSYPK  389
             +H+ +   HK+M MWML LRT  ++KKKE W+   GV
Sbjct  284  IYHLPSGYTHKLMRMWMLFLRTTSIEKKKEVWYFKTGV----------------------  321

Query  390  NYKAFGSLGFSGKHFGVGATVTYAKVKTKLLSMKKS-SRERLKMAVLFFLCSVIIGKRKT  566
                                +    VKTKL+ M+ + S++RL+MAVL+FL S I+   KT
Sbjct  322  --------------------IRREDVKTKLMEMEPARSKDRLRMAVLYFLTSSIVVPTKT  361

Query  567  GEKAEAVEDFFLKSVEDLEWCKTFPWGRLAFDKNMKDIFHLMDHFDEELGAAWVFPSFVM  746
            GE+A  ++DF +++  DL    TF
Sbjct  362  GERASPIDDFCVRAASDL----TF------------------------------------  381

Query  747  QLLAFETIPNLMNNFRKHVREADPECPRMCKMKFKPSSMKGFPISELYDTLGTTKV  914
               +FE IP+L   F +    A   CPRMCK+ FK + MK F + ++   LGTT+V
Sbjct  382  ---SFEAIPSLRERFIEEKEGAHAGCPRMCKVNFKRTEMKRFTLQQINHVLGTTEV  434


>hypothetical protein AXX17_AT2G07430 [Arabidopsis thaliana]
Sequence ID: OAP10004.1 Length: 499
Range 1: 2 to 194

Score:144 bits(364), Expect:4e-36,
Method:Compositional matrix adjust.,
Identities:86/221(39%), Positives:121/221(54%), Gaps:34/221(15%)

Query  159  LSSEEKKWFWEHPQFKHFFHMHTDSNHKVMAMWMLLLRTACVDKKKECWFVVNGVPIRYS  338
            L   +KKWF  + QFKH +HM   S +KV  M MLL+RTA  +KK+ CWF+VN VPIRYS
Sbjct  2    LKENKKKWFRTNKQFKHIWHMDRHSKNKVHGMLMLLMRTASTEKKRVCWFMVNDVPIRYS  61

Query  339  LQELALISGLYCHSYPKNYKA--FGSLGFSGKHFGVGATVTYAKVKTKLLSMKKS-SRER  509
            L+E ALI+GL  H +  ++K   FGS  F  K +G    V    V+  L SM+      +
Sbjct  62   LREHALITGLDFHQFELDFKTRNFGSFDFVEKVYGT-QVVNVKDVEDMLKSMEDECDGHQ  120

Query  510  LKMAVLFFLCSVIIGKRKTGEKAEAVEDFFLKSVEDLEWCKTFPWGRLAFDKNMKDIFHL  689
            L++AVL FLC+++                           K FP GR  F+  M+ I HL
Sbjct  121  LRVAVLLFLCAIV---------------------------KKFPSGRNTFEDTMEKIVHL  153

Query  690  M-DHFDEELGAAWVFPSFV--MQLLAFETIPNLMNNFRKHV  803
            M    D ++G  ++FP F+  +++LAFE IP +   F+K+V
Sbjct  154  MKKRLDGKVGIEYLFPGFIIPLKVLAFECIPEMSKQFQKNV  194


>gamma-irradiation and mitomycin c induced 1 [Arabidopsis thaliana]
Sequence ID: NP_001330245.1 Length: 1566
>gamma-irradiation and mitomycin c induced 1 [Arabidopsis thaliana]
Sequence ID: NP_001330246.1 Length: 1566 >gamma-irradiation and mitomycin c induced 1 [Arabidopsis thaliana]
Sequence ID: ANM68491.1 Length: 1566 >gamma-irradiation and mitomycin c induced 1 [Arabidopsis thaliana]
Sequence ID: ANM68492.1 Length: 1566
Range 1: 932 to 1032

Score:141 bits(355), Expect:1e-33,
Method:Compositional matrix adjust.,
Identities:72/124(58%), Positives:85/124(68%), Gaps:23/124(18%)

Query  1823  ISCVDEPFSVSVSCKVNPGPLKHVVEKYPEALENLLPGSTVQNYILEVKVYLLSIE*LHQ  2002
             I  +D PFSVSV CKVNPGPLK V    P+ALENLLP STV+++ILE+
Sbjct  932   IRAMDNPFSVSVPCKVNPGPLKRVAVNNPKALENLLPDSTVEDFILEL------------  979

Query  2003  IKLCTIFFMQVFDGYNNHVAEGTNVLICIEGYCINDSMGLNQKVNSCGCIDLSGILQVTA  2182
                        FDGYNNHVAEGT+VLI I+GY I D MG+N+KV+S GCI+LSGIL+VT
Sbjct  980   -----------FDGYNNHVAEGTDVLIHIDGYRIEDWMGINRKVDSRGCINLSGILKVTE  1028

Query  2183  GYGK  2194
             GYGK
Sbjct  1029  GYGK  1032


>gamma-irradiation and mitomycin c induced 1 [Arabidopsis thaliana]
Sequence ID: NP_197816.3 Length: 1598
>RecName: Full=Structural maintenance of chromosomes flexible hinge domain-containing protein GMI1; Short=SMC hinge domain-containing protein GMI1; AltName: Full=Protein GAMMA-IRRADIATION AND MITOMYCIN C INDUCED 1 [Arabidopsis thaliana]
Sequence ID: F4KFS5.1 Length: 1598 >gamma-irradiation and mitomycin c induced 1 [Arabidopsis thaliana]
Sequence ID: AED93279.1 Length: 1598 >unnamed protein product [Arabidopsis thaliana]
Sequence ID: VYS67776.1 Length: 1598
Range 1: 964 to 1064

Score:140 bits(354), Expect:2e-33,
Method:Compositional matrix adjust.,
Identities:72/124(58%), Positives:85/124(68%), Gaps:23/124(18%)

Query  1823  ISCVDEPFSVSVSCKVNPGPLKHVVEKYPEALENLLPGSTVQNYILEVKVYLLSIE*LHQ  2002
             I  +D PFSVSV CKVNPGPLK V    P+ALENLLP STV+++ILE+
Sbjct  964   IRAMDNPFSVSVPCKVNPGPLKRVAVNNPKALENLLPDSTVEDFILEL------------  1011

Query  2003  IKLCTIFFMQVFDGYNNHVAEGTNVLICIEGYCINDSMGLNQKVNSCGCIDLSGILQVTA  2182
                        FDGYNNHVAEGT+VLI I+GY I D MG+N+KV+S GCI+LSGIL+VT
Sbjct  1012  -----------FDGYNNHVAEGTDVLIHIDGYRIEDWMGINRKVDSRGCINLSGILKVTE  1060

Query  2183  GYGK  2194
             GYGK
Sbjct  1061  GYGK  1064


>GMI1 [Arabidopsis thaliana]
Sequence ID: OAO93531.1 Length: 1548
Range 1: 914 to 1014

Score:140 bits(353), Expect:2e-33,
Method:Compositional matrix adjust.,
Identities:72/124(58%), Positives:85/124(68%), Gaps:23/124(18%)

Query  1823  ISCVDEPFSVSVSCKVNPGPLKHVVEKYPEALENLLPGSTVQNYILEVKVYLLSIE*LHQ  2002
             I  +D PFSVSV CKVNPGPLK V    P+ALENLLP STV+++ILE+
Sbjct  914   IRAMDNPFSVSVPCKVNPGPLKRVAVNNPKALENLLPDSTVEDFILEL------------  961

Query  2003  IKLCTIFFMQVFDGYNNHVAEGTNVLICIEGYCINDSMGLNQKVNSCGCIDLSGILQVTA  2182
                        FDGYNNHVAEGT+VLI I+GY I D MG+N+KV+S GCI+LSGIL+VT
Sbjct  962   -----------FDGYNNHVAEGTDVLIHIDGYRIEDWMGINRKVDSRGCINLSGILKVTE  1010

Query  2183  GYGK  2194
             GYGK
Sbjct  1011  GYGK  1014


>unnamed protein product [Arabidopsis thaliana]
Sequence ID: CAA0404534.1 Length: 1566
Range 1: 932 to 1032

Score:140 bits(353), Expect:2e-33,
Method:Compositional matrix adjust.,
Identities:72/124(58%), Positives:85/124(68%), Gaps:23/124(18%)

Query  1823  ISCVDEPFSVSVSCKVNPGPLKHVVEKYPEALENLLPGSTVQNYILEVKVYLLSIE*LHQ  2002
             I  +D PFSVSV CKVNPGPLK V    P+ALENLLP STV+++ILE+
Sbjct  932   IRAMDNPFSVSVPCKVNPGPLKRVAVNNPKALENLLPDSTVEDFILEL------------  979

Query  2003  IKLCTIFFMQVFDGYNNHVAEGTNVLICIEGYCINDSMGLNQKVNSCGCIDLSGILQVTA  2182
                        FDGYNNHVAEGT+VLI I+GY I D MG+N+KV+S GCI+LSGIL+VT
Sbjct  980   -----------FDGYNNHVAEGTDVLIHIDGYRIEDWMGINRKVDSRGCINLSGILKVTE  1028

Query  2183  GYGK  2194
             GYGK
Sbjct  1029  GYGK  1032


>F28J9.13 [Arabidopsis thaliana]
Sequence ID: AAF06090.1 Length: 467
Range 1: 3 to 127

Score:131 bits(330), Expect:6e-32,
Method:Compositional matrix adjust.,
Identities:63/125(50%), Positives:88/125(70%), Gaps:3/125(2%)

Query  573  KAEAVEDFFLKSVEDLEWCKTFPWGRLAFDKNMKDIFHLMDHFDEELGAAWVFPSFVMQ-  749
            K   VE FFL++V DL+ CKTFPWG+ AF +N+K+IF+LM   + ++ +  VFPS V+
Sbjct  3    KGTKVEKFFLRAVADLQLCKTFPWGKYAFAENLKNIFYLMAKCNGKVSSQKVFPSIVLPW  62

Query  750  --LLAFETIPNLMNNFRKHVREADPECPRMCKMKFKPSSMKGFPISELYDTLGTTKVRND  923
               LAFE+IP L NNF + +   DP+CP MCKM FK S+MKGF +S++YD LG TKV ++
Sbjct  63   RYYLAFESIPILRNNFCEDIESVDPQCPWMCKMMFKLSTMKGFSMSDVYDKLGKTKVSDN  122

Query  924  LIVYY  938
            + + Y
Sbjct  123  MFINY  127


>hypothetical protein AXX17_ATUG00940 [Arabidopsis thaliana]
Sequence ID: OAO89416.1 Length: 545
Range 1: 92 to 269

Score:130 bits(326), Expect:4e-31,
Method:Compositional matrix adjust.,
Identities:68/180(38%), Positives:101/180(56%), Gaps:11/180(6%)

Query  36   EEETEPLRPLKMYFSPSDYLKPFKIGTKCYIHQVVNILESHLSSEEKKWFWEHPQFKHFF  215
            ++E +P++P  M+F PS+Y K  K+GT+  + Q V  LE  +  E+  WF  H QF++ F
Sbjct  92   DDEEQPMQPQGMFFGPSEYQKTCKVGTRYTVQQTVKYLEGFI--EDLPWFKAHSQFRNVF  149

Query  216  HMHTDSNHKVMAMWMLLLRTACVDKKKECWFVVNGVPIRYSLQELALISGLYCHSYPKNY  395
            HM  + NH    M MLLLRT  ++  +ECWFVV G+P RYS++E AL+ G  CH Y K
Sbjct  150  HMPEEKNHMTQGMGMLLLRTTQIEMDRECWFVVIGLPTRYSIKEHALLCGFDCHEYLKEL  209

Query  396  KAF-----GSLGFSGKHFGVGATVTYAKVKTKLLSMK----KSSRERLKMAVLFFLCSVI  548
            +         L F+ K F     +    V+ K+  MK    +   +R K+A+L FLC ++
Sbjct  210  QPSIKILDAELKFAKKIFKKVFGIKIIDVEKKVDEMKNCGERKKADRKKLAILLFLCKIL  269


Range 2: 264 to 321

Score:60.5 bits(145), Expect:1e-08,
Method:Compositional matrix adjust.,
Identities:30/59(51%), Positives:39/59(66%), Gaps:1/59(1%)

Query  738  FVMQLLAFETIPNLMNNFRKHVREADPECPRMCKMKFKPSSMKGFPISELYDTLGTTKV  914
            F+ ++L FE IP+L   FR  VR  + ECPRMCK KF  S MKGF + E+ + LG+ KV
Sbjct  264  FLCKILPFEAIPHLGQKFRDPVR-FENECPRMCKAKFSSSVMKGFSLEEINEELGSVKV  321


>putative protein [Arabidopsis thaliana]
Sequence ID: CAB43910.1 Length: 493
>putative protein [Arabidopsis thaliana]
Sequence ID: CAB79654.1 Length: 493
Range 1: 108 to 216

Score:128 bits(321), Expect:1e-30,
Method:Compositional matrix adjust.,
Identities:58/111(52%), Positives:75/111(67%), Gaps:2/111(1%)

Query  45   TEPLRPLKMYFSPSDYLKPFKIGTKCYIHQVVNILESHLSSEEKKWFWEHPQFKHFFHMH  224
             +PL P  MYF P+ Y K  KIGT+C + Q V  +E+  +  E KWF  H QFKH FHM
Sbjct  108  NQPLPPEVMYFDPTTYTKVCKIGTRCQLVQTVEFIETLDA--ELKWFKNHDQFKHIFHMP  165

Query  225  TDSNHKVMAMWMLLLRTACVDKKKECWFVVNGVPIRYSLQELALISGLYCH  377
             + NH +  MWML++RTA  +  +ECWFVVNGVPIRYS++E AL++GL C
Sbjct  166  KEPNHMIQGMWMLMVRTAKTELARECWFVVNGVPIRYSIREHALLTGLNCR  216


Range 2: 232 to 294

Score:59.3 bits(142), Expect:2e-08,
Method:Compositional matrix adjust.,
Identities:32/64(50%), Positives:40/64(62%), Gaps:3/64(4%)

Query  717  AAWVFPSFV--MQLLAFETIPNLMNNFRKHVREADPECPRMCKMKFKPSSMKGFPISELY  890
            A   F  F+  +++LAFE IP L   FR+ V  AD ECPRMCK KF  S MKGF + E+
Sbjct  232  AQTAFYGFITPLEILAFECIPQLGKRFREAV-PADKECPRMCKHKFSESCMKGFTLEEIN  290

Query  891  DTLG  902
            + LG
Sbjct  291  EALG  294


>hypothetical protein [Arabidopsis thaliana]
Sequence ID: AAO73430.1 Length: 210
Range 1: 5 to 183

Score:108 bits(271), Expect:3e-26,
Method:Compositional matrix adjust.,
Identities:69/184(38%), Positives:100/184(54%), Gaps:10/184(5%)

Query  405  GSLGFSGKHFGVGATVTYAKVKTKLLSMKKSSRERLKMAVLFFLCSVIIGKRKTGEKAEA  584
            G++ F  + F   + +    V  K   MK  S +RLK+ +L+FL  V+    K G K +
Sbjct  5    GNINFVKRIFKKESGIKVVDVLAKYSKMKHGS-DRLKLVLLYFLVKVV----KAGAKNDG  59

Query  585  -VEDFFLKSVEDLEWCKTFPWGRLAFDKNMKDIFHLMDHFDE--ELGAAWVFPSFV--MQ  749
             +E+F L+ V DL  C+TFPWGR  F + M  I  +M + +   +  A   F  F+  ++
Sbjct  60   NIEEFLLRIVGDLNACETFPWGRYTFLECMAGIRKMMKNMNGFVKPKAQPCFSGFIVPLE  119

Query  750  LLAFETIPNLMNNFRKHVREADPECPRMCKMKFKPSSMKGFPISELYDTLGTTKVRNDLI  929
            +LA+E IP L   FR  VR A  +CPRMCK KFK  SMKG P+  +Y  LG +K    ++
Sbjct  120  ILAYEAIPQLGLKFRVLVRSALNDCPRMCKHKFKECSMKGVPLEVIYKELGNSKDFPSIL  179

Query  930  VYYY  941
            V  Y
Sbjct  180  VPAY  183


>unnamed protein product [Arabidopsis thaliana]
Sequence ID: VYS48117.1 Length: 115
Range 1: 1 to 109

Score:103 bits(258), Expect:1e-25,
Method:Compositional matrix adjust.,
Identities:51/110(46%), Positives:64/110(58%), Gaps:1/110(0%)

Query  159  LSSEEKKWFWEHPQFKHFFHMHTDSNHKVMAMWMLLLRTACVDKKKECWFVVNGVPIRYS  338
            +S  ++ WF  HP F+H FHM    NH++M MWM LLRTA ++ KKE WF+VN VPIRY
Sbjct  1    MSKSKRAWFENHPSFQHIFHMPRVPNHRLMGMWMFLLRTARIEMKKEAWFIVNSVPIRYG  60

Query  339  LQELALISGLYCHSYPKNYKAFGSLGFSGKHFGVGATVTYAKVKTKLLSM  488
            + E  LIS   C +Y   YK   +L F  KHF     V    V  KL+ M
Sbjct  61   ILEHTLISSFNCKNYTLGYKNTRNLDFKKKHFK-NTVVKREDVIEKLIGM  109


>unnamed protein product [Arabidopsis thaliana]
Sequence ID: BAB01195.1 Length: 552
Range 1: 153 to 256

Score:113 bits(283), Expect:1e-25,
Method:Compositional matrix adjust.,
Identities:50/106(47%), Positives:72/106(67%), Gaps:5/106(4%)

Query  24   SMDSEEETEPLRPLKMYFSPSDYLKPFKIGTKCYIHQVVNI---LESHLSSEEKKWFWEH  194
            SM+ +EET+ + PL MYF PS+Y+K  K+  +CYIH+V+     LE  +S  E++WF  H
Sbjct  153  SMEIDEETKAILPLSMYFPPSEYVKKIKLSIRCYIHEVLTTFDKLEPEMSKSEREWFQNH  212

Query  195  PQFKHFFHMHTDSNHKVMAMWMLLLRTACVDKKKECWFVVNGVPIR  332
            P F+H FHM  D NH++M MWMLL RTA +++KKE   ++   PI+
Sbjct  213  PSFQHIFHMPRDPNHRLMGMWMLLFRTARIERKKEACIII--APIK  256


Range 2: 250 to 382

Score:108 bits(270), Expect:6e-24,
Method:Compositional matrix adjust.,
Identities:56/133(42%), Positives:79/133(59%), Gaps:6/133(4%)

Query  543  VIIGKRKTGEKAEAVEDFFLKSVEDLEWCKTFPWGRLAFDKNMKDIFHLMDHFDEELGA-  719
            +II   KTG+KA  V++F LK+V DL +C+ F WGR  FD  +  I H ++HF+  +
Sbjct  250  IIIAPIKTGDKAPQVDEFCLKAVSDLTFCRNFQWGRYFFDYMLGTISHTVNHFNGSVTNN  309

Query  720  ---AWVFPSFV--MQLLAFETIPNLMNNFRKHVREADPECPRMCKMKFKPSSMKGFPISE  884
                W    F   M+LLAFE IP L   F + +  AD  CPRMCK++FK + +KGFP+
Sbjct  310  EKYVWSVLGFCLPMELLAFEAIPQLRETFMEDIAGADEGCPRMCKVRFKKNHLKGFPLDT  369

Query  885  LYDTLGTTKVRND  923
            +Y  LG T+ R +
Sbjct  370  IYAELGITQRRKE  382


>T25H8.3 gene product [Arabidopsis thaliana]
Sequence ID: AAD17348.1 Length: 301
>hypothetical protein [Arabidopsis thaliana]
Sequence ID: CAB80812.1 Length: 301
Range 1: 86 to 215

Score:106 bits(264), Expect:2e-24,
Method:Compositional matrix adjust.,
Identities:55/147(37%), Positives:83/147(56%), Gaps:17/147(11%)

Query  474  KLLSMKKSSRERLKMAVLFFLCSVIIGKRKTGEKAEAVEDFFLKSVEDLEWCKTFPWGRL  653
             L+ +   S++RL+M VL+FL S+I+   KTGEK   V++F L++  DL + KTFPW R
Sbjct  86   NLMDINNGSKDRLRMGVLYFLTSIIVVLTKTGEKGSPVDNFCLRAASDLTFSKTFPWRRY  145

Query  654  AFDKNMKDIFHLMDHFDEELGAAWVFPSFVMQLLAFETIPNLMNNFRKHVREADPECPRM  833
            +F+  +K I H +DHF+                 A E IP+L   F +    A   C RM
Sbjct  146  SFEFMLKPISHTLDHFN-----------------ALEAIPSLREQFIEKKEGAYTGCSRM  188

Query  834  CKMKFKPSSMKGFPISELYDTLGTTKV  914
            CK+ FK + +KGF + ++ + L TT+V
Sbjct  189  CKVNFKRTEIKGFTLEQINNVLDTTEV  215


>hypothetical protein [Arabidopsis thaliana]
Sequence ID: AAD20103.1 Length: 425
Range 1: 1 to 180

Score:107 bits(268), Expect:3e-24,
Method:Compositional matrix adjust.,
Identities:67/185(36%), Positives:101/185(54%), Gaps:10/185(5%)

Query  402  FGSLGFSGKHFGVGATVTYAKVKTKLLSMKKSSRERLKMAVLFFLCSVIIGKRKTGEKAE  581
             G++ F  + F   + +    V  K   MK  S +RLK+ +L+FL  V+    K G K +
Sbjct  1    MGNINFVKRIFKKESGIKVVDVLAKCSKMKHGS-DRLKLVLLYFLVKVV----KAGAKND  55

Query  582  A-VEDFFLKSVEDLEWCKTFPWGRLAFDKNMKDIFHLMDHFDE--ELGAAWVFPSFV--M  746
              +E+F L+ + D+  C+TFPWGR +F + M  I  +M + +   +  A   F  F+  +
Sbjct  56   GNIEEFLLRIIGDMNACETFPWGRYSFLECMAGIRKMMKNMNGFVKPKAQPCFSGFIVPL  115

Query  747  QLLAFETIPNLMNNFRKHVREADPECPRMCKMKFKPSSMKGFPISELYDTLGTTKVRNDL  926
            ++LA+E IP L   FR  VR A  +CPRMCK KFK  SMKG P+  +Y  LG +K    +
Sbjct  116  EILAYEAIPQLGLKFRVLVRSALNDCPRMCKHKFKECSMKGVPLEVIYKELGNSKDFPSI  175

Query  927  IVYYY  941
            +V  Y
Sbjct  176  LVPAY  180


>hypothetical protein [Arabidopsis thaliana]
Sequence ID: AAB81878.1 Length: 599
>hypothetical protein [Arabidopsis thaliana]
Sequence ID: CAB77985.1 Length: 599
Range 1: 143 to 227

Score:100 bits(250), Expect:2e-21,
Method:Compositional matrix adjust.,
Identities:44/87(51%), Positives:57/87(65%), Gaps:2/87(2%)

Query  54   LRPLKMYFSPSDYLKPFKIGTKCYIHQVVNILESHLSSEEKKWFWEHPQFKHFFHMHTDS  233
            ++P +M+F PS+Y+K  KIGT+C + Q V  LE     E+  WF  HPQF+H FHM  +
Sbjct  143  MQPERMFFGPSEYVKTCKIGTRCTVRQTVKYLEE--CKEDLPWFKAHPQFRHVFHMPEEK  200

Query  234  NHKVMAMWMLLLRTACVDKKKECWFVV  314
            NH    MWMLLLRTA  +  +ECWFVV
Sbjct  201  NHMTQGMWMLLLRTARTEMDRECWFVV  227


Range 2: 230 to 330

Score:77.0 bits(188), Expect:6e-14,
Method:Compositional matrix adjust.,
Identities:48/127(38%), Positives:64/127(50%), Gaps:27/127(21%)

Query  564  TGEKAEA-VEDFFLKSVEDLEWCKTFPWGRLAFDKNMKDIFHLMDHFDEELGAAWVFPSF  740
            T  KA+  ++ FFLK V+D+  C+T PWGR  FD  M+D
Sbjct  230  TKSKADGNIDKFFLKIVDDVRACETSPWGRFTFDGCMED---------------------  268

Query  741  VMQLLAFETIPNLMNNFRKHVREADPECPRMCKMKFKPSSMKGFPISELYDTLGTTKVRN  920
                L FE IP+L   FR  VR  + ECPRMCK KF  S MKGF + E+ + L + KV +
Sbjct  269  ----LPFEAIPHLGQKFRDPVR-FENECPRMCKAKFSSSVMKGFSLEEINEELESVKVIS  323

Query  921  DLIVYYY  941
             ++   Y
Sbjct  324  SVLESDY  330


>unnamed protein product [Arabidopsis thaliana]
Sequence ID: VYS59032.1 Length: 92
Range 1: 1 to 92

Score:80.1 bits(196), Expect:1e-17,
Method:Compositional matrix adjust.,
Identities:42/92(46%), Positives:56/92(60%), Gaps:6/92(6%)

Query  516  MAVLFFLCSVIIGKRKTGEKAEAVEDFFLKSVEDLEWCKTFPWGRLAFDKNMKDIFHLMD  695
            M VL+FL S+II   KTG+KA  V++F LK+V DL +C+ F WGR  FD  +  I H ++
Sbjct  1    MMVLYFLSSIIIAPIKTGDKAPQVDEFCLKAVSDLTFCRNFQWGRYFFDYMLGTISHTVN  60

Query  696  HFDEELGA----AWVFPSFV--MQLLAFETIP  773
            HF+  +       W  P F   M+LLAFE IP
Sbjct  61   HFNGSVTNNEKYVWSVPGFCLPMELLAFEAIP  92


>hypothetical protein AXX17_ATUG01010 [Arabidopsis thaliana]
Sequence ID: OAO89413.1 Length: 661
Range 1: 216 to 347

Score:82.8 bits(203), Expect:1e-15,
Method:Compositional matrix adjust.,
Identities:51/133(38%), Positives:73/133(54%), Gaps:7/133(5%)

Query  561  KTGEKAEA-VEDFFLKSVEDLEWCKTFPWGRLAFDKNMKDIFHLM---DHFDEELGAAWV  728
            K G K +  +E+F L+ + DL  C+TFPWG   F + M  I  +M   + F +     W
Sbjct  216  KAGAKNDGNIEEFLLRIIGDLNACETFPWGCYTFLECMAGIRKMMKNMNGFVKPKAQPW-  274

Query  729  FPSFV--MQLLAFETIPNLMNNFRKHVREADPECPRMCKMKFKPSSMKGFPISELYDTLG  902
            F  F+  +++LA+E IP L   FR  VR A  +CPR+CK KFK  S KG P+  +Y  LG
Sbjct  275  FSGFIVPLEILAYEAIPQLGLKFRVPVRSALNDCPRICKHKFKECSXKGVPLEVIYKELG  334

Query  903  TTKVRNDLIVYYY  941
             +K    ++V  Y
Sbjct  335  NSKDFPRILVPAY  347


>hypothetical protein AXX17_AT4G07660 [Arabidopsis thaliana]
Sequence ID: OAO98863.1 Length: 88
Range 1: 1 to 76

Score:65.1 bits(157), Expect:2e-12,
Method:Compositional matrix adjust.,
Identities:28/76(37%), Positives:47/76(61%), Gaps:0/76(0%)

Query  27   MDSEEETEPLRPLKMYFSPSDYLKPFKIGTKCYIHQVVNILESHLSSEEKKWFWEHPQFK  206
            MDS+E++ PL+P + +F+P +Y+K  K  T+C I   + ++   L+  EK WF ++ QFK
Sbjct  1    MDSDEDSMPLQPERFFFTPKEYVKTMKTYTRCNIAFTLAVIGQKLNDREKSWFIKNRQFK  60

Query  207  HFFHMHTDSNHKVMAM  254
            H +HM     +KV  +
Sbjct  61   HIWHMVRSDKNKVQGI  76


>hypothetical protein AXX17_ATUG02090 [Arabidopsis thaliana]
Sequence ID: OAO89294.1 Length: 221
Range 1: 2 to 93

Score:65.5 bits(158), Expect:4e-11,
Method:Compositional matrix adjust.,
Identities:33/92(36%), Positives:52/92(56%), Gaps:5/92(5%)

Query  669  MKDIFHLMDHFD---EELGAAWVFPSFV--MQLLAFETIPNLMNNFRKHVREADPECPRM  833
            +K I H +DHF+       + W  P     +++L FE IP+L   F +    AD  CPRM
Sbjct  2    LKSISHTLDHFNGIVPNTQSPWHVPGLCVPLEVLVFEVIPSLRERFIEEKEGADAGCPRM  61

Query  834  CKMKFKPSSMKGFPISELYDTLGTTKVRNDLI  929
            CK+ FK + MKGF + ++ + LGT+++   +I
Sbjct  62   CKVHFKRTEMKGFALEQINNVLGTSEIIESII  93


>hypothetical protein AXX17_ATUG02100 [Arabidopsis thaliana]
Sequence ID: OAO89302.1 Length: 337
Range 1: 263 to 337

Score:66.6 bits(161), Expect:5e-11,
Method:Compositional matrix adjust.,
Identities:31/75(41%), Positives:47/75(62%), Gaps:3/75(4%)

Query  39   EETEPLRPLKMYFSPSDYLKPFKIGTKCYIHQV---VNILESHLSSEEKKWFWEHPQFKH  209
            EET  ++PL MYF   +Y K  K+ T+CYI +V   ++ L+  L+  EK +F EHP FK+
Sbjct  263  EETMAMQPLSMYFLALEYTKKMKLATRCYISEVLKNIDDLKLALTDSEKNYFKEHPSFKY  322

Query  210  FFHMHTDSNHKVMAM  254
             +H+ +   HK+M M
Sbjct  323  IYHLPSGYTHKLMEM  337


>putative protein [Arabidopsis thaliana]
Sequence ID: CAB86472.1 Length: 264
Range 1: 6 to 208

Score:64.3 bits(155), Expect:1e-10,
Method:Compositional matrix adjust.,
Identities:56/208(27%), Positives:88/208(42%), Gaps:35/208(16%)

Query  252  MWMLLLRTACVDKKKECWFVVNGVPIRYSLQELALISGLYCHSYPKNYK-------AFGS  410
            M  LL R     KK E W V  G P+R+SL E A ++GL C  + ++Y          G
Sbjct  6    MHALLCRQLVTKKKYELWTVFGGHPMRFSLTEFACVTGLPCAEFSEDYDPDDDSVFVDGM  65

Query  411  LGFSGKHFGVGATVTYAKVKTKLLSMKK--SSRERLKMAVLFFLCSVIIGKRKTGEKAEA  584
              +  +  G   TVT   V   L + +K  SS  RLK+A L  +  V+I   +
Sbjct  66   KSYWDELIGPDKTVTLRDVSAMLTNKRKTLSSDHRLKLAFLLIVDGVLIASNQICRPTFK  125

Query  585  VEDFFLKSVEDLEWCKTFPWGRLAFDKNM---------------------KDIFHLMDHF  701
                +++ + DL+   +FPWG+ +F K +                       I  L++
Sbjct  126  ----YVEMLADLDKFLSFPWGKESFLKTVVGMRPNKRNLGKSTGKRQLTTDPIKSLVNQL  181

Query  702  DEELGAAWVFPSFVMQLLAFETIPNLMN  785
             ++      FP   +QL+AF  IP L++
Sbjct  182  QQKTFRLKGFP-LALQLIAFRNIPGLLD  208


>unnamed protein product [Arabidopsis thaliana]
Sequence ID: VYS47676.1 Length: 673
Range 1: 35 to 264

Score:66.6 bits(161), Expect:1e-10,
Method:Compositional matrix adjust.,
Identities:65/243(27%), Positives:107/243(44%), Gaps:32/243(13%)

Query  123  YIHQVVNILESHLSSEEKKWFWEH---PQFKHFFHMHTDS-NHKVMAMWMLLLRTACVDK  290
            Y+  + N+L+    SEE    WE     QF   F       +H    +  LL R     K
Sbjct  35   YLGTIANLLKG---SEE----WERIKSSQFGKLFEFPVARCSHSGKLIHGLLSRQVVTKK  87

Query  291  KKECWFVVNGVPIRYSLQELALISGLYCHSYP-----KNYKAFGSLGFSGKHFGVGATVT  455
            K E WFV  G PIR+S++E  +++GL C   P     K ++    L    + FG    VT
Sbjct  88   KHELWFVFGGHPIRFSIREFHIVTGLRCGKLPTEDEVKKHQDSKYLSVWNRLFGEKRMVT  147

Query  456  YAKVKTKLLSMKKSSRERLKMAVLFFLCSVIIGKRKTGEKAEAVEDFFLKSVEDLEWCKT  635
               V   L   K SS ++L +A++  L  V++   ++      V   F++ + D+++
Sbjct  148  IGDVLEMLQKKKLSSWKKLCLALIVILDGVVVCNDQS-----FVTLDFVEMLNDIDFFLE  202

Query  636  FPWGRLAFDKNMKD----------IFHLMDHFDEELGAAWVFPSFVMQLLAFETIPNLMN  785
            +PWGR AF   ++           +  L     ++  A + FP   +QL  FE+IP ++
Sbjct  203  YPWGRKAFLATIRRFGPPKDAPNPLGKLKKRLKQKTSACYGFP-LALQLQVFESIPVILE  261

Query  786  NFR  794
              +
Sbjct  262  RIK  264


>K-box region protein (DUF1985) [Arabidopsis thaliana]
Sequence ID: NP_001321227.1 Length: 623
>K-box region protein (DUF1985) [Arabidopsis thaliana]
Sequence ID: ANM58816.1 Length: 623
Range 1: 35 to 264

Score:65.5 bits(158), Expect:3e-10,
Method:Compositional matrix adjust.,
Identities:64/243(26%), Positives:107/243(44%), Gaps:32/243(13%)

Query  123  YIHQVVNILESHLSSEEKKWFWEH---PQFKHFFHMHTDS-NHKVMAMWMLLLRTACVDK  290
            Y+  + N+L+    SEE    WE     QF   F       +H    +  LL R     K
Sbjct  35   YLGTIANLLKG---SEE----WERIKSSQFGKLFEFPVARCSHSGKLIHGLLSRQVVTKK  87

Query  291  KKECWFVVNGVPIRYSLQELALISGLYCHSYP-----KNYKAFGSLGFSGKHFGVGATVT  455
            K E WFV  G PIR+S++E  +++GL C   P     K ++    L    + FG    VT
Sbjct  88   KHELWFVFGGHPIRFSIREFHIVTGLRCGKLPTEDEVKKHQDSKYLSVWNRLFGEKRMVT  147

Query  456  YAKVKTKLLSMKKSSRERLKMAVLFFLCSVIIGKRKTGEKAEAVEDFFLKSVEDLEWCKT  635
               V   L   K SS ++L +A++  +  V++   ++      V   F++ + D+++
Sbjct  148  IGDVLEMLQKKKLSSWKKLCLALIVIVDGVVVCNDQS-----FVTLDFVEMLNDIDFFLE  202

Query  636  FPWGRLAFDKNMKD----------IFHLMDHFDEELGAAWVFPSFVMQLLAFETIPNLMN  785
            +PWGR AF   ++           +  L     ++  A + FP   +QL  FE+IP ++
Sbjct  203  YPWGRKAFLATIRRFGPPKDAPNPLGKLKKRLKQKTSACYGFP-LALQLQVFESIPVILE  261

Query  786  NFR  794
              +
Sbjct  262  RIK  264


>hypothetical protein AXX17_AT3G33420 [Arabidopsis thaliana]
Sequence ID: OAP06548.1 Length: 962
Range 1: 46 to 240

Score:65.9 bits(159), Expect:3e-10,
Method:Compositional matrix adjust.,
Identities:61/203(30%), Positives:90/203(44%), Gaps:21/203(10%)

Query  210  FFHMHTDSNHKVMAMWMLLL--RTACVDKKKECWFVVNGVPIRYSLQELALISGLYCHSY  383
            F H+       V A + L L  R   V    E W V  G PIR+SL+E  +++GL C  Y
Sbjct  46   FHHIRFPQQTGVSASFGLFLLSRQLEVANSDEIWVVFAGTPIRFSLREFKIVTGLPCGKY  105

Query  384  PKNYKAFGSLGFSGKH-------FGVGATVTYAKVKTKLLSMKKSSRE-RLKMAVLFFLC  539
            PK  K     G  GK        FG+   VT  +V T L     + R+ RL+ A L  +
Sbjct  106  PKVQKN-KKRGTGGKQIPYYNTLFGLEEDVTVERVITMLTKRVVTDRDIRLRYACLALVD  164

Query  540  SVIIGKRKTGEKAEAVEDFFLKSVEDLEWCKTFPWGRLAFDKNMKDIFHLMDHFDEELGA  719
              ++    T    + +++    S EDL+   ++PWGRL+F+  M  I    +   E+L
Sbjct  165  GFLL---PTSHYPKIIKNHAEMS-EDLQGFLSYPWGRLSFEMMMTSI---KEREVEQLAT  217

Query  720  AWVFPS---FVMQLLAFETIPNL  779
              V      F +QL+  E  P +
Sbjct  218  TCVAVQGLLFALQLVVLEAPPAI  240


>hypothetical protein AXX17_AT1G31740 [Arabidopsis thaliana]
Sequence ID: OAP15120.1 Length: 673
Range 1: 35 to 264

Score:65.5 bits(158), Expect:3e-10,
Method:Compositional matrix adjust.,
Identities:64/243(26%), Positives:108/243(44%), Gaps:32/243(13%)

Query  123  YIHQVVNILESHLSSEEKKWFWE---HPQFKHFFHMHTDS-NHKVMAMWMLLLRTACVDK  290
            Y+  + N+L+    SEE    WE   + QF   F       +H    +  LL R     K
Sbjct  35   YLGTIANLLKG---SEE----WERIKNSQFGKLFEFPVARCSHSGKLIHGLLSRQVVTKK  87

Query  291  KKECWFVVNGVPIRYSLQELALISGLYCHSYP-----KNYKAFGSLGFSGKHFGVGATVT  455
            K E WFV  G PIR+S++E  +++GL C   P     K ++    L    + FG    VT
Sbjct  88   KHELWFVFGGHPIRFSIREFHIVTGLRCGKLPTEDEVKKHQDSKYLSVWNRLFGEKRMVT  147

Query  456  YAKVKTKLLSMKKSSRERLKMAVLFFLCSVIIGKRKTGEKAEAVEDFFLKSVEDLEWCKT  635
               V   L   K SS ++L +A++  +  V++   ++      V   F++ + D+++
Sbjct  148  IGDVLEMLQKKKLSSWKKLCLALIVIVDGVVVCNDQS-----FVTLDFVEILNDIDFFLE  202

Query  636  FPWGRLAFDKNMKD----------IFHLMDHFDEELGAAWVFPSFVMQLLAFETIPNLMN  785
            +PWGR AF   ++           +  L     ++  A + FP   +QL  FE+IP ++
Sbjct  203  YPWGRKAFLATIRRFGPPKDAPNPLGKLKKRLKQKTSACYGFP-LALQLQVFESIPVILE  261

Query  786  NFR  794
              +
Sbjct  262  RIK  264


>K-box region protein (DUF1985) [Arabidopsis thaliana]
Sequence ID: NP_174400.1 Length: 673
>EST gb|Z33866 comes from this gene [Arabidopsis thaliana]
Sequence ID: AAD21698.1 Length: 673 >K-box region protein (DUF1985) [Arabidopsis thaliana]
Sequence ID: AEE31318.1 Length: 673 >unnamed protein product [Arabidopsis thaliana]
Sequence ID: CAA0259238.1 Length: 673
Range 1: 35 to 264

Score:65.5 bits(158), Expect:4e-10,
Method:Compositional matrix adjust.,
Identities:64/243(26%), Positives:107/243(44%), Gaps:32/243(13%)

Query  123  YIHQVVNILESHLSSEEKKWFWEH---PQFKHFFHMHTDS-NHKVMAMWMLLLRTACVDK  290
            Y+  + N+L+    SEE    WE     QF   F       +H    +  LL R     K
Sbjct  35   YLGTIANLLKG---SEE----WERIKSSQFGKLFEFPVARCSHSGKLIHGLLSRQVVTKK  87

Query  291  KKECWFVVNGVPIRYSLQELALISGLYCHSYP-----KNYKAFGSLGFSGKHFGVGATVT  455
            K E WFV  G PIR+S++E  +++GL C   P     K ++    L    + FG    VT
Sbjct  88   KHELWFVFGGHPIRFSIREFHIVTGLRCGKLPTEDEVKKHQDSKYLSVWNRLFGEKRMVT  147

Query  456  YAKVKTKLLSMKKSSRERLKMAVLFFLCSVIIGKRKTGEKAEAVEDFFLKSVEDLEWCKT  635
               V   L   K SS ++L +A++  +  V++   ++      V   F++ + D+++
Sbjct  148  IGDVLEMLQKKKLSSWKKLCLALIVIVDGVVVCNDQS-----FVTLDFVEMLNDIDFFLE  202

Query  636  FPWGRLAFDKNMKD----------IFHLMDHFDEELGAAWVFPSFVMQLLAFETIPNLMN  785
            +PWGR AF   ++           +  L     ++  A + FP   +QL  FE+IP ++
Sbjct  203  YPWGRKAFLATIRRFGPPKDAPNPLGKLKKRLKQKTSACYGFP-LALQLQVFESIPVILE  261

Query  786  NFR  794
              +
Sbjct  262  RIK  264


>unknown protein [Arabidopsis thaliana]
Sequence ID: AAG50606.1 Length: 1311
Range 1: 210 to 409

Score:65.1 bits(157), Expect:5e-10,
Method:Compositional matrix adjust.,
Identities:58/205(28%), Positives:86/205(41%), Gaps:35/205(17%)

Query  261  LLLRTACVDKKKECWFVVNGVPIRYSLQELALISGLYCHSYPKNYKAFGSLGF-SGKH--  431
            LL R     KK E W V  G P+R+SL E A ++GL C  +P  Y    S  +  GK
Sbjct  210  LLCRQLLTKKKYEFWTVFGGHPMRFSLLEFASVTGLPCGEFPDEYDPEDSPVYDDGKKSY  269

Query  432  ----FGVGATVTYAKVKTKLLSMKK--SSRERLKMAVLFFLCSVIIGKRKTGEKAEAVED  593
                 G   TVT   +   L   +   SS  +LK+  L  +  V+I   +TG     +
Sbjct  270  WNELIGPDKTVTLGDISAMLTKKRSKLSSDHKLKLTFLLIVDGVLIASNQTGR----LTF  325

Query  594  FFLKSVEDLEWCKTFPWGRLAFDKNMK---------------------DIFHLMDHFDEE  710
             +++ + DLE    FPWGR +F K ++                      I +L+    +
Sbjct  326  KYVEMLADLEKFLLFPWGRESFMKTVEAMTPEKRILAKSTGKRQLTNDPIQNLVKQLQQI  385

Query  711  LGAAWVFPSFVMQLLAFETIPNLMN  785
                  FP   +QLLAF  IP +++
Sbjct  386  TFRLKGFP-HALQLLAFRNIPGMVD  409


>putative protein [Arabidopsis thaliana]
Sequence ID: CAB81580.1 Length: 1113
Range 1: 81 to 311

Score:65.1 bits(157), Expect:5e-10,
Method:Compositional matrix adjust.,
Identities:55/239(23%), Positives:102/239(42%), Gaps:28/239(11%)

Query  123  YIHQVVNILESHLSSEEKKWFWEHPQFKHFFHMHTD-SNHKVMAMWMLLLRTACVDKKKE  299
            YI  + N+L+      E ++  + P F   F +  + ++     +  L+ R     K  E
Sbjct  81   YISDIANVLKGK---PEMQFLLDSP-FGELFKIPKNKASFNAKLVLGLICRQLVTKKVNE  136

Query  300  CWFVVNGVPIRYSLQELALISGLYCHSYPKNYKAFGSLGFSGKHFGV---------GATV  452
             W V  G PIR+ L+E +++SGL C  YPK       +    +   V         G TV
Sbjct  137  MWIVFGGHPIRFGLREFSILSGLECGKYPKKKDVDDVISVKPECESVWKTLFDERFGDTV  196

Query  453  TYAKVKTKLLSMKKSSR--ERLKMAVLFFLCSVIIGKRKTGEKAEAVEDFFLKSVEDLEW  626
                     L  ++S    ++L ++++  +  V+                 ++  ++LE+
Sbjct  197  PTIADLVSWLQEEESMEGWKQLALSLIILVDGVVAAHSNPNRPTSKT----VEMTKNLEF  252

Query  627  CKTFPWGRLAFDKNMKDIFHLMDHFDEE------LGAAWVFPSF--VMQLLAFETIPNL  779
               +PWGR++F + +  I +    +D +      L  ++    F   +QLLAFETIP++
Sbjct  253  FCKYPWGRVSFTRTLGRIANFQTPYDAQQLIRGLLVGSYALHGFPLALQLLAFETIPSI  311


>Contains similarity to a putative protein T32A11_100 gi|7413627 from Arabidopsis thaliana BAC T32A11 gb|AL138653 [Arabidopsis thaliana]
Sequence ID: AAF78272.1 Length: 1075
Range 1: 62 to 282

Score:64.7 bits(156), Expect:6e-10,
Method:Compositional matrix adjust.,
Identities:60/226(27%), Positives:92/226(40%), Gaps:36/226(15%)

Query  201  FKHFFHMHTDSNHKVMAMWMLLLRTACVDKKK-ECWFVVNGVPIRYSLQELALISGLYCH  377
            F   FH+          +   LL T  V KKK E W V  G P+R++L E A ++GL C
Sbjct  62   FGPLFHLPVSKCAMSGKLMHALLCTQLVTKKKYELWTVFGGYPLRFALPEFACVTGLPCG  121

Query  378  SYPK-------NYKAFGSLGFSGKHFGVGATVTYAKVKTKLLSMKK--SSRERLKMAVLF  530
             + +            G   +  +  G   TVT   V   L + +K  SS  RLK+A L
Sbjct  122  EFSEDYDPDDDPVYVDGMKSYWDELIGPDKTVTLGDVSDMLTNKRKTLSSDHRLKLAFLL  181

Query  531  FLCSVIIGKRKTGEKAEAVEDFFLKSVEDLEWCKTFPWGRLAFDKNM-------------  671
             +  V+I   + G         +++ + DL+   +FPWGR +F K +
Sbjct  182  IVDGVLIASNQIGRPTFK----YVEMLADLDKFLSFPWGRESFLKTVVGMRPDKRNLGKS  237

Query  672  --------KDIFHLMDHFDEELGAAWVFPSFVMQLLAFETIPNLMN  785
                      I  L+    ++      FP   +QL+AF  IP L++
Sbjct  238  TGKRQLTTDPIKSLVSQLQQKTFRLKGFP-LALQLIAFRNIPGLLD  282


>unnamed protein product [Arabidopsis thaliana]
Sequence ID: BAB02940.1 Length: 1119
Range 1: 81 to 311

Score:64.7 bits(156), Expect:7e-10,
Method:Compositional matrix adjust.,
Identities:53/238(22%), Positives:100/238(42%), Gaps:26/238(10%)

Query  123  YIHQVVNILESHLSSEEKKWFWEHPQFKHFFHMHTDSNHKVMAMWMLLLRTACVDKKKEC  302
            YI  +  +L+      E ++  + P  K F      ++     +  L+ R     K  E
Sbjct  81   YISHIAKVLKG---KPEMQFLLDSPFGKLFKIPKNKASFNAKLVLGLICRQLVTKKVNEM  137

Query  303  WFVVNGVPIRYSLQELALISGLYCHSYPKNYKAFGSLGFSGK---------HFGVGATVT  455
            W V  G PIR+ L+E ++++GL C  YPK       +    +         H   G TV
Sbjct  138  WIVFGGHPIRFRLREFSILTGLECGKYPKKKDVDDVISVKPECESVWKTLFHERFGDTVP  197

Query  456  YAKVKTKLLSMKKSSR--ERLKMAVLFFLCSVIIGKRKTGEKAEAVEDFFLKSVEDLEWC  629
                    L  ++S    ++L ++++  +  V+                 +++ ++LE+
Sbjct  198  TIADLVSWLQEEESMEGWKQLALSLIILVDGVVAAHSNPNRPTSKT----VETTKNLEFF  253

Query  630  KTFPWGRLAFDKNMKDIFHLMDHFDEE------LGAAWVFPSF--VMQLLAFETIPNL  779
              +PWGR++F + +  I +    +D +      L  ++    F   +QLLAFETIP++
Sbjct  254  CKYPWGRVSFTRTLGRIANFQTPYDAQKLIRGILVGSYALHGFPLALQLLAFETIPSI  311


>unnamed protein product [Arabidopsis thaliana]
Sequence ID: BAA98062.1 Length: 1444
Range 1: 81 to 311

Score:64.3 bits(155), Expect:9e-10,
Method:Compositional matrix adjust.,
Identities:52/238(22%), Positives:102/238(42%), Gaps:26/238(10%)

Query  123  YIHQVVNILESHLSSEEKKWFWEHPQFKHFFHMHTDSNHKVMAMWMLLLRTACVDKKKEC  302
            YI  +  +L+      E ++    P  K F      ++     +  L+ R     K  E
Sbjct  81   YISDIAKVLKGK---PEMQFLLNSPFGKLFKIPKNKASFNAKLVLGLICRQLVTKKVNEM  137

Query  303  WFVVNGVPIRYSLQELALISGLYCHSYPKNYKAFGSLGFSGKHFGVGATVTYAKVKTKLL  482
            W V  G PIR+ L+E ++++GL C  YPK       +    +   V  T+ + +    +L
Sbjct  138  WIVFGGHPIRFGLREFSILTGLECGKYPKKKDVDDVISVKPECESVWNTLFHERFGNTVL  197

Query  483  SM--------KKSSRE---RLKMAVLFFLCSVIIGKRKTGEKAEAVEDFFLKSVEDLEWC  629
            ++        ++ S E   +L ++++  +  V+                 ++  ++LE+
Sbjct  198  TIADLVSWLQEEESMEGWKQLALSLIILVDGVVAAHSNPNRPTSKT----VEMTKNLEFF  253

Query  630  KTFPWGRLAFDKNMKDIFHLMDHFDEE------LGAAWVFPSF--VMQLLAFETIPNL  779
              +PWGR++F + +  I +    +D +      L  ++    F   +QLLAFETIP++
Sbjct  254  CKYPWGRVSFTRTLGRIANFQTPYDAQKLIRGILVGSYALHGFPLALQLLAFETIPSI  311


>putative protein [Arabidopsis thaliana]
Sequence ID: CAB86452.1 Length: 1009
Range 1: 21 to 216

Score:64.3 bits(155), Expect:9e-10,
Method:Compositional matrix adjust.,
Identities:50/216(23%), Positives:86/216(39%), Gaps:36/216(16%)

Query  198  QFKHFFHMHTDS-NHKVMAMWMLLLRTACVDKKKECWFVVNGVPIRYSLQELALISGLYC  374
            QFK  F +     +H       +L R     +K E W V  G P+R+SL+E  + +GL C
Sbjct  21   QFKKLFELPVARCSHSAKLSHGMLARQLLTQRKHELWTVYGGYPLRFSLKEFQITTGLNC  80

Query  375  HSYP-----KNYKAFGSLGFSGKHFGVGATVTYAKVKTKLLSMKKSSRERLKMAVLFFLC  539
               P     ++++    L    + FG    VT                    M V+  L
Sbjct  81   DKLPTDSEVEDHQDPAYLSVWNRLFGEKCVVTV-------------------MDVVEMLR  121

Query  540  SVIIGKRKTGEKAEAVEDFFLKSVEDLEWCKTFPWGRLAFDKNMK----------DIFHL  689
              +  + +   K   V   +++ + DLE+  ++PWGR+AF+  M+           I  L
Sbjct  122  GDVNAEPRNNRKQHRVSSSYVEMLHDLEFFMSYPWGRVAFESTMERFGPSTGENDPIAEL  181

Query  690  MDHFDEELGAAWVFPSFVMQLLAFETIPNLMNNFRK  797
                 ++    + FP   +QL    +IP L +  ++
Sbjct  182  KSRLSQKSSCCYGFP-LALQLQVLNSIPALCSRIKE  216


>hypothetical protein [Arabidopsis thaliana]
Sequence ID: AAD36942.1 Length: 963
>hypothetical protein [Arabidopsis thaliana]
Sequence ID: CAB77907.1 Length: 963
Range 1: 71 to 213

Score:63.9 bits(154), Expect:1e-09,
Method:Compositional matrix adjust.,
Identities:49/149(33%), Positives:74/149(49%), Gaps:15/149(10%)

Query  261  LLLRTACVDKKKECWFVVNGVPIRYSLQELALISGLYCHSYP------KNYKAFGSLGFS  422
            LL R   +DK  E W V  G P+R+SL+E  L++GL C  YP      K   A   + F
Sbjct  71   LLGRKLDIDKANEIWVVYAGTPVRFSLREFHLVTGLACGRYPDLPKKRKKGTAEKEIPFY  130

Query  423  GKHFGVGATVTYAKVKTKLLSMKKSSRE---RLKMAVLFFLCSVIIGKRKTGEKAEAVED  593
               F + + VT ++V T L   KK + +   R++ AVL  +   ++    T    + V++
Sbjct  131  STLFELESDVTVSRVITML--KKKVTDDPSLRIRYAVLALVDGYLL---PTSHYPKIVKE  185

Query  594  FFLKSVEDLEWCKTFPWGRLAFDKNMKDI  680
               + VE+L     +PWGRL F+  MK I
Sbjct  186  -HAEMVENLSSFLAYPWGRLTFEMTMKSI  213


>T22C5.24 [Arabidopsis thaliana]
Sequence ID: AAF24952.1 Length: 1444
Range 1: 81 to 311

Score:63.9 bits(154), Expect:1e-09,
Method:Compositional matrix adjust.,
Identities:53/238(22%), Positives:98/238(41%), Gaps:26/238(10%)

Query  123  YIHQVVNILESHLSSEEKKWFWEHPQFKHFFHMHTDSNHKVMAMWMLLLRTACVDKKKEC  302
            YI  +  +L+      E ++  + P  K F      ++     +  L+ R     K  E
Sbjct  81   YISDIAKVLKGK---PEMQFLLDSPFGKLFKIPKNKASFNAKLVLGLICRQLVTKKVNEM  137

Query  303  WFVVNGVPIRYSLQELALISGLYCHSYPKNYKAFGSLGFSGK---------HFGVGATVT  455
            W V  G PIR+ L+E ++++GL C  YPK       +    +         H   G TV
Sbjct  138  WIVFGGHPIRFGLREFSILTGLECGKYPKKKDVDDVISVKPECESVWNTLFHERFGDTVP  197

Query  456  YAKVKTKLLSMKKSSR--ERLKMAVLFFLCSVIIGKRKTGEKAEAVEDFFLKSVEDLEWC  629
                    L  ++S     +L ++++  +  V+                 ++  ++LE+
Sbjct  198  TIADLVSWLQEEESMEGWNQLALSLIILVDGVVAAHSNPNRPTSKT----VEMTKNLEFF  253

Query  630  KTFPWGRLAFDKNMKDIFHLMDHFDEE------LGAAWVFPSF--VMQLLAFETIPNL  779
              +PWGR++F + +  I +    +D +      L  ++    F   +QLLAFETIP++
Sbjct  254  CKYPWGRVSFTRTLGRIANFQTPYDAQKLIRGILVGSYALHGFPLALQLLAFETIPSI  311


>hypothetical protein [Arabidopsis thaliana]
Sequence ID: AAG50521.1 Length: 1201
Range 1: 81 to 333

Score:63.9 bits(154), Expect:1e-09,
Method:Compositional matrix adjust.,
Identities:61/264(23%), Positives:107/264(40%), Gaps:40/264(15%)

Query  123  YIHQVVNILESHLSSEEKKWFWEHPQFKHFFHMHTDSNHKVMAMWMLLLRTACVDKKKEC  302
            YI  +  +L+      E ++  + P  K F      ++     +  L+ R     K  E
Sbjct  81   YISDIAKVLKGK---PEMQFLLDSPFGKLFKIPKNKASFNAKLVLGLICRQLVTKKVNEM  137

Query  303  WFVVNGVPIRYSLQELALISGLYCHSYPKNYKAFGSLGFSGK---------HFGVGATVT  455
            W V  G PIR+ L+E ++++GL C  YPK       +    +         H   G TV
Sbjct  138  WIVFGGHPIRFGLREFSILTGLECGKYPKKKDVDDVISVKPECESVWNTLFHERFGDTVP  197

Query  456  YAKVKTKLLSMKKSSR--ERLKMAVLFFLCSVIIG----KRKTGEKAEAVEDFFLKSVED  617
                    L  ++S    ++L ++++  +  V+       R T +  E  ++
Sbjct  198  TIADLVSWLQEEESMEGWKQLALSLIILVDGVVAAHSNPNRPTSKTVEMTKNLEF-----  252

Query  618  LEWCKTFPWGRLAFDKNMKDIFHLMDHFDEE------LGAAWVFPSF--VMQLLAFETIP  773
              +CK +PWGR++F + +  I +    +D +      L  ++    F   +QLLAFETIP
Sbjct  253  --FCK-YPWGRVSFTRTLGRIANFQTPYDAQKLIRGILVGSYALHGFPLALQLLAFETIP  309

Query  774  NLMN------NFRKHVREADPECP  827
            ++             V+ AD  CP
Sbjct  310  SIAKLGPDDVEVNYIVKPADNVCP  333


>hypothetical protein [Arabidopsis thaliana]
Sequence ID: AAG50582.1 Length: 1198
Range 1: 81 to 311

Score:63.5 bits(153), Expect:2e-09,
Method:Compositional matrix adjust.,
Identities:56/242(23%), Positives:100/242(41%), Gaps:34/242(14%)

Query  123  YIHQVVNILESHLSSEEKKWFWEHPQFKHFFHMHTDSNHKVMAMWMLLLRTACVDKKKEC  302
            YI  +  +L+      E ++  + P  K F      ++     +  L+ R     K  E
Sbjct  81   YISDIAKVLKGK---PEMQFLLDSPFGKLFKIPKNKASFNAKLVLGLICRQLVTKKVNEM  137

Query  303  WFVVNGVPIRYSLQELALISGLYCHSYPKNYKAFGSLGFSGK---------HFGVGATVT  455
            W V  G PIR+ L+E ++++GL C  YPK       +    +         H   G TV
Sbjct  138  WIVFGGHPIRFGLREFSILTGLECGKYPKKKDVDDVISVKPECESVWNTLFHERFGDTVP  197

Query  456  YAKVKTKLLSMKKSSR--ERLKMAVLFFLCSVIIG----KRKTGEKAEAVEDFFLKSVED  617
                    L  ++S     +L ++++  +  V+       R T +  E  ++
Sbjct  198  TIADLVSWLQEEESMEGWNQLALSLIILVDGVVAAHSNPNRPTSKTVEMTKNLEF-----  252

Query  618  LEWCKTFPWGRLAFDKNMKDIFHLMDHFDEE------LGAAWVFPSF--VMQLLAFETIP  773
              +CK +PWGR++F + +  I +    +D +      L  ++    F   +QLLAFETIP
Sbjct  253  --FCK-YPWGRVSFTRTLGRIANFQTPYDAQKLIRGILVGSYALHGFPLALQLLAFETIP  309

Query  774  NL  779
            ++
Sbjct  310  SI  311


>hypothetical protein [Arabidopsis thaliana]
Sequence ID: AAD14458.1 Length: 1285
>hypothetical protein [Arabidopsis thaliana]
Sequence ID: CAB77815.1 Length: 1285
Range 1: 233 to 463

Score:63.5 bits(153), Expect:2e-09,
Method:Compositional matrix adjust.,
Identities:55/239(23%), Positives:101/239(42%), Gaps:28/239(11%)

Query  123  YIHQVVNILESHLSSEEKKWFWEHPQFKHFFHMHTDSNHKVMAMWMLLLRTACVDKKKEC  302
            YI  + N+L++     E ++  + P  + F      ++     +  L+ R     K  E
Sbjct  233  YISDIANVLKAK---PEMQFLLDSPFGELFKIPQNKASFNAKLVLGLICRQLVTKKVNEM  289

Query  303  WFVVNGVPIRYSLQELALISGLYCHSYPKNYKAFGSLGFSGKHFGV---------GATVT  455
            W V  G PIR+ L+E ++++GL C  YPK       +    +   V         G TV
Sbjct  290  WIVFGGHPIRFGLREFSILTGLECGKYPKKKDVEDVISVKPECESVWKTLFDERFGDTVP  349

Query  456  YAKVKTKLLSMKKSSR--ERLKMAVLFFLCSVIIGKRKTGEKAEAVEDFFLKSVEDLEWC  629
                    L  ++S    ++L ++++  +  V+                 ++  ++LE+
Sbjct  350  TIADLVSWLQEEESMEGWKQLALSLIILVDGVVAAHSNPNRPTSKT----VEMTKNLEFF  405

Query  630  KTFPWGRLAFDKNMKDIFHLMDHFDEE-------LG--AAWVFPSFVMQLLAFETIPNL  779
              +PWGR++F + +  I      +D +       +G  A   FP   +QLLAFETIP++
Sbjct  406  CKYPWGRVSFTRTLGRIAKFQTPYDAQQLIRGLLVGYYALHGFP-LALQLLAFETIPSI  463


>unnamed protein product [Arabidopsis thaliana]
Sequence ID: CAA0384296.1 Length: 197
Range 1: 92 to 151

Score:59.3 bits(142), Expect:3e-09,
Method:Compositional matrix adjust.,
Identities:25/60(42%), Positives:40/60(66%), Gaps:0/60(0%)

Query  30   DSEEETEPLRPLKMYFSPSDYLKPFKIGTKCYIHQVVNILESHLSSEEKKWFWEHPQFKH  209
            DS+E+  PL+P   +F+P +Y K  KI T+C I   + ++ + L++ EK+WF E+ QFKH
Sbjct  92   DSDEDFMPLQPESFFFTPDEYRKTLKIYTRCTIANTLLVIGAKLNNREKRWFIENQQFKH  151


>F27F5.16 [Arabidopsis thaliana]
Sequence ID: AAF69170.1 Length: 1745
Range 1: 60 to 291

Score:62.8 bits(151), Expect:3e-09,
Method:Compositional matrix adjust.,
Identities:60/240(25%), Positives:109/240(45%), Gaps:27/240(11%)

Query  123  YIHQVVNILESHLSSEEKKWFWEHPQFKHFFHMHTD-SNHKVMAMWMLLLRTACVDKKKE  299
            YI Q+   L+    + E  +  + P F   F + T+ ++     +  L+ R    +K+ E
Sbjct  60   YIGQIAAALKD---TPEMTFLLDSP-FGDLFRIPTNKASFSGKLVLGLICRQLVTNKRYE  115

Query  300  CWFVVNGVPIRYSLQELALISGLYCHSYPKNYKAFGSLGFSGKHFGVGATV--------T  455
             W V  G PIR  L+E ALI+GL C +YPK+      +        V AT+        T
Sbjct  116  MWMVFAGHPIRLGLREWALITGLECGTYPKDKDVESVMQREEGENTVWATLFGDDKAKPT  175

Query  456  YAKVKTKLLS-MKKSSRERLKMAVLFFLCSVIIGKRKTGEKAEAVEDFFLKSVEDLEWCK  632
              +++ +L+S     + ++L +A++  +  V+I  +    +     +  ++  ++L++
Sbjct  176  VEELRDRLISETDMPAWKKLALALIIIVDGVLICDKSPPLRP---NEMTVELTKNLDFFC  232

Query  633  TFPWGRLAFD---------KNMKDIFHLMDHFDEELGAAWVFPSFVMQLLAFETIPNLMN  785
             +PWGR +F          K   D+  L     +   A   FP   +QLLAF TIP++ +
Sbjct  233  KYPWGRTSFLLTLERIASFKGDTDVKKLRSGCKQHSYALHGFP-LGLQLLAFATIPSIAS  291


>hypothetical protein [Arabidopsis thaliana]
Sequence ID: AAC97230.2 Length: 1422
>hypothetical protein [Arabidopsis thaliana]
Sequence ID: AAM15356.1 Length: 1422
Range 1: 60 to 291

Score:62.0 bits(149), Expect:5e-09,
Method:Compositional matrix adjust.,
Identities:60/240(25%), Positives:109/240(45%), Gaps:27/240(11%)

Query  123  YIHQVVNILESHLSSEEKKWFWEHPQFKHFFHMHTD-SNHKVMAMWMLLLRTACVDKKKE  299
            YI Q+   L+    + E  +  + P F   F + T+ ++     +  L+ R    +K+ E
Sbjct  60   YIGQIAAALKD---TPEMTFLLDSP-FGDLFRIPTNKASFSGKLVLGLICRQLVTNKRYE  115

Query  300  CWFVVNGVPIRYSLQELALISGLYCHSYPKNYKAFGSLGFSGKHFGVGATV--------T  455
             W V  G PIR  L+E ALI+GL C +YPK+      +        V AT+        T
Sbjct  116  MWMVFAGHPIRLGLREWALITGLECGTYPKDKDVESVMQREEGENTVWATLFGDDKAKPT  175

Query  456  YAKVKTKLLS-MKKSSRERLKMAVLFFLCSVIIGKRKTGEKAEAVEDFFLKSVEDLEWCK  632
              +++ +L+S     + ++L +A++  +  V+I  +    +     +  ++  ++L++
Sbjct  176  VEELRDRLISETDMPAWKKLALALIIIVDGVLICDKSPPLRP---NEMTVELTKNLDFFC  232

Query  633  TFPWGRLAFD---------KNMKDIFHLMDHFDEELGAAWVFPSFVMQLLAFETIPNLMN  785
             +PWGR +F          K   D+  L     +   A   FP   +QLLAF TIP++ +
Sbjct  233  KYPWGRTSFLLTLERIASFKGDTDVKKLRSGCKQHSYALHGFP-LGLQLLAFATIPSIAS  291


>unnamed protein product [Arabidopsis thaliana]
Sequence ID: CAA0269437.1 Length: 72
Range 1: 2 to 66

Score:55.1 bits(131), Expect:5e-09,
Method:Compositional matrix adjust.,
Identities:30/66(45%), Positives:35/66(53%), Gaps:1/66(1%)

Query  291  KKECWFVVNGVPIRYSLQELALISGLYCHSYPKNYKAFGSLGFSGKHFGVGATVTYAKVK  470
            KKE WF+VN VPIRY + E  LIS   C +Y   YK   +L F  KHF     V    V
Sbjct  2    KKEAWFIVNSVPIRYGILEHTLISSFNCKNYTLGYKNTRNLDFKKKHFK-NTVVKREDVI  60

Query  471  TKLLSM  488
             KL+ M
Sbjct  61   EKLIGM  66


>hypothetical protein [Arabidopsis thaliana]
Sequence ID: AAC95211.1 Length: 992
Range 1: 30 to 259

Score:60.5 bits(145), Expect:1e-08,
Method:Compositional matrix adjust.,
Identities:55/239(23%), Positives:104/239(43%), Gaps:19/239(7%)

Query  135  VVNILESHLSSEEKKWFWEHPQFKHFFHMHTDSNHKVMAMWMLLLRTACVDKKKECWFVV  314
            +++ + + L +E+ K   E  QF   F     + +    +  LL R   V KK+E W +
Sbjct  30   LIDDIANALDAEDMKILRES-QFGKLFDFSESAAYSGKLIHFLLTRQLVVRKKQEIWVIF  88

Query  315  NGVPIRYSLQELALISGLYCHSYPKNYKAFGSLGFS-GKHF-------GVGATVTYAKVK  470
            +G P+R+S+ E   ++GL C   P+  +  G    + GK++        V       ++K
Sbjct  89   SGSPVRFSISEFQNVTGLNCDKPPRMKERKGKRKLAPGKYWYSLFDRSDVSVEWVVGRLK  148

Query  471  TKLLSMKKSSRERLKMAVLFFLCSVIIGKRKTGEKAEAVEDFFLKSVEDLEWCKTFPWGR  650
             +++   +    RL+ AVL  +  V+       + +    D      E+L+    FPWG
Sbjct  149  KRVV---QDQGIRLRYAVLALIDGVLCPTSGRSKISPVHADM----AENLDMFLNFPWGT  201

Query  651  LAFDKNMKDIF-HLMDHFDEELGAAWVFPSFVMQLLAFETIPNLMN-NFRKHVREADPE  821
            ++F   +K +     D    +      FP   +QLL  +++P +    F + V ++D E
Sbjct  202  MSFLMTLKSVTARGADKLMRKSITVQGFP-HALQLLLLQSVPRIRELQFPEAVEDSDSE  259


>unnamed protein product [Arabidopsis thaliana]
Sequence ID: BAB01835.1 Length: 1015
>unnamed protein product [Arabidopsis thaliana]
Sequence ID: BAB10871.1 Length: 1015
Range 1: 30 to 259

Score:60.5 bits(145), Expect:1e-08,
Method:Compositional matrix adjust.,
Identities:55/239(23%), Positives:104/239(43%), Gaps:19/239(7%)

Query  135  VVNILESHLSSEEKKWFWEHPQFKHFFHMHTDSNHKVMAMWMLLLRTACVDKKKECWFVV  314
            +++ + + L +E+ K   E  QF   F     + +    +  LL R   V KK+E W +
Sbjct  30   LIDDIANALDAEDMKILRES-QFGKLFDFSESAAYSGKLIHFLLTRQLVVRKKQEIWVIF  88

Query  315  NGVPIRYSLQELALISGLYCHSYPKNYKAFGSLGFS-GKHF-------GVGATVTYAKVK  470
            +G P+R+S+ E   ++GL C   P+  +  G    + GK++        V       ++K
Sbjct  89   SGSPVRFSISEFQNVTGLNCDKPPRMKERKGKRKLAPGKYWYSLFDRSDVSVEWVVGRLK  148

Query  471  TKLLSMKKSSRERLKMAVLFFLCSVIIGKRKTGEKAEAVEDFFLKSVEDLEWCKTFPWGR  650
             +++   +    RL+ AVL  +  V+       + +    D      E+L+    FPWG
Sbjct  149  KRVV---QDQGIRLRYAVLALIDGVLCPTSGRSKISPVHADM----AENLDMFLNFPWGT  201

Query  651  LAFDKNMKDIF-HLMDHFDEELGAAWVFPSFVMQLLAFETIPNLMN-NFRKHVREADPE  821
            ++F   +K +     D    +      FP   +QLL  +++P +    F + V ++D E
Sbjct  202  MSFLMTLKSVTARGADKLMRKSITVQGFP-HALQLLLLQSVPRIRELQFPEAVEDSDSE  259


>Similar to a hypothetical protein At2g29240 gi|3980408 from Arabidopsis thaliana gb|AC004561 [Arabidopsis thaliana]
Sequence ID: AAF80659.1 Length: 992
Range 1: 29 to 258

Score:60.5 bits(145), Expect:1e-08,
Method:Compositional matrix adjust.,
Identities:55/239(23%), Positives:104/239(43%), Gaps:19/239(7%)

Query  135  VVNILESHLSSEEKKWFWEHPQFKHFFHMHTDSNHKVMAMWMLLLRTACVDKKKECWFVV  314
            +++ + + L +E+ K   E  QF   F     + +    +  LL R   V KK+E W +
Sbjct  29   LIDDIANALDAEDMKILRES-QFGKLFDFSESAAYSGKLIHFLLTRQLVVRKKQEIWVIF  87

Query  315  NGVPIRYSLQELALISGLYCHSYPKNYKAFGSLGFS-GKHF-------GVGATVTYAKVK  470
            +G P+R+S+ E   ++GL C   P+  +  G    + GK++        V       ++K
Sbjct  88   SGSPVRFSISEFQNVTGLNCDKPPRMKERKGKRKLAPGKYWYSLFDRSDVSVEWVVGRLK  147

Query  471  TKLLSMKKSSRERLKMAVLFFLCSVIIGKRKTGEKAEAVEDFFLKSVEDLEWCKTFPWGR  650
             +++   +    RL+ AVL  +  V+       + +    D      E+L+    FPWG
Sbjct  148  KRVV---QDQGIRLRYAVLALIDGVLCPTSGRSKISPVHADM----AENLDMFLNFPWGT  200

Query  651  LAFDKNMKDIF-HLMDHFDEELGAAWVFPSFVMQLLAFETIPNLMN-NFRKHVREADPE  821
            ++F   +K +     D    +      FP   +QLL  +++P +    F + V ++D E
Sbjct  201  MSFLMTLKSVTARGADKLMRKSITVQGFP-HALQLLLLQSVPRIRELQFPEAVEDSDSE  258


>hypothetical protein [Arabidopsis thaliana]
Sequence ID: AAD23703.1 Length: 428
>hypothetical protein [Arabidopsis thaliana]
Sequence ID: AAM15236.1 Length: 428
Range 1: 212 to 282

Score:59.7 bits(143), Expect:1e-08,
Method:Compositional matrix adjust.,
Identities:31/72(43%), Positives:47/72(65%), Gaps:2/72(2%)

Query  738  FVMQLLAFETIPNLMNNFRKHVREADPECPRMCKMKFKPSSMKGFPISELYDTLGTTK-V  914
            F++++LA+E I  L   FR  VR A+ +CPRMCK KFK  SMKG P+  +Y  LG ++ +
Sbjct  212  FLVKILAYEAILQLQLKFRVPVR-AEKDCPRMCKHKFKECSMKGVPLEVIYKELGNSEDI  270

Query  915  RNDLIVYYYLKF  950
             + L+  Y+ K+
Sbjct  271  ASHLVPEYHEKY  282


>En/Spm-like transposon [Arabidopsis thaliana]
Sequence ID: NP_001331298.1 Length: 430
>En/Spm-like transposon [Arabidopsis thaliana]
Sequence ID: NP_001331301.1 Length: 430 >En/Spm-like transposon [Arabidopsis thaliana]
Sequence ID: ANM69635.1 Length: 430 >En/Spm-like transposon [Arabidopsis thaliana]
Sequence ID: ANM69638.1 Length: 430
Range 1: 1 to 27

Score:59.3 bits(142), Expect:2e-08,
Method:Compositional matrix adjust.,
Identities:27/27(100%), Positives:27/27(100%), Gaps:0/27(0%)

Query  2114  MGLNQKVNSCGCIDLSGILQVTAGYGK  2194
             MGLNQKVNSCGCIDLSGILQVTAGYGK
Sbjct  1     MGLNQKVNSCGCIDLSGILQVTAGYGK  27


>putative protein [Arabidopsis thaliana]
Sequence ID: CAB86693.1 Length: 1314
Range 1: 81 to 311

Score:60.1 bits(144), Expect:2e-08,
Method:Compositional matrix adjust.,
Identities:59/241(24%), Positives:102/241(42%), Gaps:32/241(13%)

Query  123  YIHQVVNILESHLSSEEKKWFWEHPQFKHFFHMHTDSNHKVMAMWMLLLRTACVDKK-KE  299
            YI  + N+L+      E ++  + P F   F +  +       + + L+    V KK  E
Sbjct  81   YISNIANVLKGK---PEMQFLLDSP-FGELFKIPKNKASFNAKLVLGLICCQLVTKKVNE  136

Query  300  CWFVVNGVPIRYSLQELALISGLYCHSYPKNYKAFGSLGFSGKHFGV---------GATV  452
             W V  G PIR+ L+E ++++GL C  YPK       +    +   V         G TV
Sbjct  137  MWIVFGGHPIRFGLREFSILTGLECGKYPKKKDVEDVISVKPECESVWKTLFDERFGDTV  196

Query  453  TYAKVKTKLLSMKKSSRERLKMAV-LFFLCSVIIGKRKTGEKAEAVEDFFLKSVE---DL  620
                     L  ++S     ++A+ L  L   ++       +A +      K+VE   +L
Sbjct  197  PTIADLVSWLQEEESMEGWKQLALSLIILVDGVVAAHSNPNRATS------KTVEMTKNL  250

Query  621  EWCKTFPWGRLAFDKNMKDIFHLMDHFDEE------LGAAWVFPSF--VMQLLAFETIPN  776
            E+   +PWGR++F + +  I      +D +      L  ++    F   + LLAFETIP+
Sbjct  251  EFFCKYPWGRVSFTRTLGRIAKFQTPYDAQQLIRGLLVGSYALHGFPLALHLLAFETIPS  310

Query  777  L  779
            +
Sbjct  311  I  311


>hypothetical protein [Arabidopsis thaliana]
Sequence ID: AAD26959.1 Length: 1218
Range 1: 100 to 279

Score:58.9 bits(141), Expect:5e-08,
Method:Compositional matrix adjust.,
Identities:48/183(26%), Positives:90/183(49%), Gaps:20/183(10%)

Query  288  KKKECWFVVNGVPIRYSLQELALISGLYCHSYPKNYKAFGSLGFSGKHFGVGATV-----  452
            K+ E W V  G PIR  L+E ALI+GL C +YPK+      +        V AT+
Sbjct  100  KRYEMWMVFAGHPIRLGLREWALITGLECGTYPKDKDVESVMQREEGENTVWATLFGDDK  159

Query  453  ---TYAKVKTKLLS-MKKSSRERLKMAVLFFLCSVIIGKRKTGEKAEAVEDFFLKSVEDL  620
               T  +++ +L+S     + ++L +A++  +  V+I  +    +     +  ++  ++L
Sbjct  160  ANPTVEELRDRLISETDMPAWKKLALALIIIVDGVLICDKSPPLRP---NEMTVELTKNL  216

Query  621  EWCKTFPWGRLAFDKNMKDIFHLMDHFD-EELGAAWVFPSFV-------MQLLAFETIPN  776
            ++   +PWGR +F   ++ I       D ++L +     S+V       +QLLAF+TIP+
Sbjct  217  DFSCKYPWGRTSFLLTLERIASFKGETDVKKLRSGCKQHSYVLHGFPLGLQLLAFKTIPS  276

Query  777  LMN  785
            + +
Sbjct  277  IAS  279


>F22O13.23 [Arabidopsis thaliana]
Sequence ID: AAF99775.1 Length: 1014
Range 1: 29 to 258

Score:58.5 bits(140), Expect:5e-08,
Method:Compositional matrix adjust.,
Identities:54/239(23%), Positives:104/239(43%), Gaps:19/239(7%)

Query  135  VVNILESHLSSEEKKWFWEHPQFKHFFHMHTDSNHKVMAMWMLLLRTACVDKKKECWFVV  314
            +++ + + L +E+ K   +  QF   F     + +    +  LL R   V KK+E W +
Sbjct  29   LIDDIANALDAEDMKILRKS-QFGKLFDFSESAAYSGKLIHFLLTRQLVVRKKQEIWVIF  87

Query  315  NGVPIRYSLQELALISGLYCHSYPKNYKAFGSLGFS-GKHF-------GVGATVTYAKVK  470
            +G P+R+S+ E   ++GL C   P+  +  G    + GK++        V       ++K
Sbjct  88   SGSPVRFSISEFQNVTGLNCDKPPRMKERKGKRKLAPGKYWYSLFDRSDVSVEWVVGRLK  147

Query  471  TKLLSMKKSSRERLKMAVLFFLCSVIIGKRKTGEKAEAVEDFFLKSVEDLEWCKTFPWGR  650
             +++   +    RL+ AVL  +  V+       + +    D      E+L+    FPWG
Sbjct  148  KRVV---QDQGIRLRYAVLALIDGVLCPTSGRSKISPVHADM----AENLDMFLNFPWGT  200

Query  651  LAFDKNMKDIF-HLMDHFDEELGAAWVFPSFVMQLLAFETIPNLMN-NFRKHVREADPE  821
            ++F   +K +     D    +      FP   +QLL  +++P +    F + V ++D E
Sbjct  201  MSFLMTLKSVTARGADKLMRKSITVQGFP-HALQLLLLQSVPRIRELQFPEAVEDSDSE  258


>putative protein [Arabidopsis thaliana]
Sequence ID: CAB81786.1 Length: 702
Range 1: 93 to 290

Score:57.8 bits(138), Expect:8e-08,
Method:Compositional matrix adjust.,
Identities:50/203(25%), Positives:82/203(40%), Gaps:28/203(13%)

Query  231  SNHKVMAMWMLLLRTACVDKKKECWFVVNGVPIRYSLQELALISGLYCHSYPKNYKAFGS  410
            + H    +  LL R     K+ E WF+  G P+R+SL+E  + +GL C   P   +
Sbjct  93   TAHSAKLIHGLLSRQLVTKKRHEFWFIFGGKPLRFSLREFHITTGLECRPIPSEEEILSH  152

Query  411  LGFSGKH-----FGVGATVTYAKVKTKLL-------SMKKSSRERLKMAVLFFLCSVIIG  554
                 K      FG    VT   V   L          K S  +R  + ++  +  VI+
Sbjct  153  QKIVSKPVWNSLFGSKKDVTVTDVLDMLGKDVKMPDGEKMSCWKRFCLLLILLVDGVIV-  211

Query  555  KRKTGEKAEAVEDFFLKSVEDLEWCKTFPWGRLAFDKNMKD-----------IFHLMDHF  701
                  K   +   ++K ++D+ +  ++PWGR+AF   M+            I  L
Sbjct  212  ---CSNKYLNITPEYVKMLDDVRFFLSYPWGRVAFKTTMERFGPPTLSDTDPITELKARL  268

Query  702  DEELGAAWVFPSFVMQLLAFETI  770
             ++    + FP   +Q+LAFE I
Sbjct  269  RQQSSCCYGFP-LALQMLAFEAI  290


>hypothetical protein [Arabidopsis thaliana]
Sequence ID: AAD20102.1 Length: 221
Range 1: 153 to 221

Score:55.5 bits(132), Expect:9e-08,
Method:Compositional matrix adjust.,
Identities:28/70(40%), Positives:41/70(58%), Gaps:1/70(1%)

Query  30   DSEEETEPLRPLKMYFSPSDYLKPFKIGTKCYIHQVVNILESHLSSEEKKWFWEHPQFKH  209
            D + +++PL P  M+F  S+Y K  KI ++C++ Q V ++E     EE  WF  H QFKH
Sbjct  153  DDDLDSQPLPPQTMHFLFSEYGKVCKISSRCHVTQTVELIERKF-KEEVVWFKRHDQFKH  211

Query  210  FFHMHTDSNH  239
             FHM  + N
Sbjct  212  IFHMPQEPNQ  221


Query #258: XLOC_030118 Query ID: lcl|Query_30749 Length: 1532

Sequences producing significant alignments:
                                                                  Max    Total Query   E   Per.
Description                                                       Score  Score cover Value Ident  Accession
hypothetical protein AT5G25755 [Arabidopsis thaliana]             58.5   58.5  9%    2e-10 63.27  NP_001332189.1


Alignments:
>hypothetical protein AT5G25755 [Arabidopsis thaliana]
Sequence ID: NP_001332189.1 Length: 62
>hypothetical protein AT5G25755 [Arabidopsis thaliana]
Sequence ID: NP_001332190.1 Length: 62 >hypothetical protein AT5G25755 [Arabidopsis thaliana]
Sequence ID: ANM70593.1 Length: 62 >hypothetical protein AT5G25755 [Arabidopsis thaliana]
Sequence ID: ANM70594.1 Length: 62
Range 1: 14 to 62

Score:58.5 bits(140), Expect:2e-10,
Method:Compositional matrix adjust.,
Identities:31/49(63%), Positives:38/49(77%), Gaps:2/49(4%)

Query  3    ALNLLKGRSLGKKE--QEKIMIS*NRVYWRELLIQKMLITKTSFTLVRQ  143
            A+ + +G+ LGKK   ++   +  NRVYWRELLIQKMLITKTSFTLVRQ
Sbjct  14   AIVMRQGKKLGKKRTREDHDQLKLNRVYWRELLIQKMLITKTSFTLVRQ  62


Query #259: XLOC_030237 Query ID: lcl|Query_30750 Length: 972

Sequences producing significant alignments:
                                                                  Max    Total Query   E   Per.
Description                                                       Score  Score cover Value Ident  Accession
F15O4.6 [Arabidopsis thaliana]                                    87.0   164   30%   1e-33 83.64  AAF79344.1
phosphoenolpyruvate carboxykinase 1 [Arabidopsis thaliana]        53.5   87.8  19%   4e-11 74.29  NP_195500.1
PEPCK [Arabidopsis thaliana]                                      53.5   87.8  19%   4e-11 74.29  OAP00328.1
phosphoenolpyruvate carboxykinase 2 [Arabidopsis thaliana]        49.3   81.2  17%   4e-09 65.71  NP_001331431.1
unnamed protein product [Arabidopsis thaliana]                    49.3   81.2  17%   4e-09 65.71  VYS71520.1
PEPCK [Arabidopsis thaliana]                                      49.3   81.2  17%   4e-09 65.71  OAO90231.1
phosphoenolpyruvate carboxykinase 2 [Arabidopsis thaliana]        49.3   81.2  17%   4e-09 65.71  NP_001331433.1
unnamed protein product [Arabidopsis thaliana]                    49.3   81.2  17%   4e-09 65.71  CAA0412274.1
phosphoenolpyruvate carboxykinase 2 [Arabidopsis thaliana]        49.3   80.9  17%   5e-09 65.71  NP_001331432.1
phosphoenolpyruvate carboxykinase (ATP) (EC 4.1.1.49)...          49.3   80.9  17%   5e-09 67.65  BAB10675.1
unnamed protein product [Arabidopsis thaliana]                    57.8   57.8  17%   1e-08 51.72  CAA0361730.1


Alignments:
>F15O4.6 [Arabidopsis thaliana]
Sequence ID: AAF79344.1 Length: 155
Range 1: 30 to 82

Score:87.0 bits(214), Expect:1e-33,
Method:Compositional matrix adjust.,
Identities:46/55(84%), Positives:50/55(90%), Gaps:2/55(3%)

Query  254  LYEQAIEYEKGSLITSNGALTMLSGAKTDREISVLLEMRLLKMSLSFGEEKKELL  418
            LYEQAIEYEKGSLITSNGALTMLSGAKTDREISVLLEMRLL+M  SFG+ +  +L
Sbjct  30   LYEQAIEYEKGSLITSNGALTMLSGAKTDREISVLLEMRLLRM--SFGDYRIFIL  82


Range 2: 115 to 155

Score:77.0 bits(188), Expect:1e-33,
Method:Compositional matrix adjust.,
Identities:39/49(80%), Positives:40/49(81%), Gaps:8/49(16%)

Query  403  KERAVDYLNSLEKVYLMPKR*ILSHHSRCNMGKDGDVALFFELLGIATF  549
            +ERAVDYLNSLEK        ILSHHS CNMGKDGDVALFFELLGIATF
Sbjct  115  RERAVDYLNSLEK--------ILSHHSGCNMGKDGDVALFFELLGIATF  155


>phosphoenolpyruvate carboxykinase 1 [Arabidopsis thaliana]
Sequence ID: NP_195500.1 Length: 671
>RecName: Full=Phosphoenolpyruvate carboxykinase (ATP); Short=PEP carboxykinase; Short=PEPCK [Arabidopsis thaliana]
Sequence ID: Q9T074.1 Length: 671 >AT4g37870/T28I19_150 [Arabidopsis thaliana]
Sequence ID: AAK50062.1 Length: 671 >AT4g37870/T28I19_150 [Arabidopsis thaliana]
Sequence ID: AAL77736.1 Length: 671 >phosphoenolpyruvate carboxykinase 1 [Arabidopsis thaliana]
Sequence ID: AEE86847.1 Length: 671 >unnamed protein product [Arabidopsis thaliana]
Sequence ID: CAA0397843.1 Length: 671 >phosphoenolpyruvate carboxykinase (ATP)-like protein [Arabidopsis thaliana]
Sequence ID: CAB38935.1 Length: 671
Range 1: 155 to 189

Score:53.5 bits(127), Expect:4e-11,
Method:Compositional matrix adjust.,
Identities:26/35(74%), Positives:29/35(82%), Gaps:0/35(0%)

Query  239  FSYSKLYEQAIEYEKGSLITSNGALTMLSGAKTDR  343
             S ++LYEQAI+YEKGS ITSNGAL  LSGAKT R
Sbjct  155  LSPAELYEQAIKYEKGSFITSNGALATLSGAKTGR  189


Range 2: 200 to 241

Score:34.3 bits(77), Expect:4e-11,
Method:Compositional matrix adjust.,
Identities:22/44(50%), Positives:24/44(54%), Gaps:18/44(40%)

Query  367  AITEDELELW*GK----------------ERAVDYLNSLEKVYL  450
            A TEDEL  W GK                ERAVDYLNSLEKV++
Sbjct  200  ATTEDEL--WWGKGSPNIEMDEHTFMVNRERAVDYLNSLEKVFV  241


>PEPCK [Arabidopsis thaliana]
Sequence ID: OAP00328.1 Length: 671
Range 1: 155 to 189

Score:53.5 bits(127), Expect:4e-11,
Method:Compositional matrix adjust.,
Identities:26/35(74%), Positives:29/35(82%), Gaps:0/35(0%)

Query  239  FSYSKLYEQAIEYEKGSLITSNGALTMLSGAKTDR  343
             S ++LYEQAI+YEKGS ITSNGAL  LSGAKT R
Sbjct  155  LSPAELYEQAIKYEKGSFITSNGALATLSGAKTGR  189


Range 2: 200 to 241

Score:34.3 bits(77), Expect:4e-11,
Method:Compositional matrix adjust.,
Identities:22/44(50%), Positives:24/44(54%), Gaps:18/44(40%)

Query  367  AITEDELELW*GK----------------ERAVDYLNSLEKVYL  450
            A TEDEL  W GK                ERAVDYLNSLEKV++
Sbjct  200  ATTEDEL--WWGKGSPNIEMDEHTFMVNRERAVDYLNSLEKVFV  241


>phosphoenolpyruvate carboxykinase 2 [Arabidopsis thaliana]
Sequence ID: NP_001331431.1 Length: 577
>phosphoenolpyruvate carboxykinase 2 [Arabidopsis thaliana]
Sequence ID: ANM69777.1 Length: 577
Range 1: 154 to 188

Score:49.3 bits(116), Expect:4e-09,
Method:Compositional matrix adjust.,
Identities:23/35(66%), Positives:28/35(80%), Gaps:0/35(0%)

Query  239  FSYSKLYEQAIEYEKGSLITSNGALTMLSGAKTDR  343
             S ++LYEQAI++EKGS +TS GAL  LSGAKT R
Sbjct  154  LSPAELYEQAIKFEKGSFVTSTGALATLSGAKTGR  188


Range 2: 202 to 240

Score:32.0 bits(71), Expect:4e-09,
Method:Compositional matrix adjust.,
Identities:18/39(46%), Positives:21/39(53%), Gaps:16/39(41%)

Query  382  ELELW*GK----------------ERAVDYLNSLEKVYL  450
            E ELW GK                ERAVDYLNSL+KV++
Sbjct  202  EAELWWGKGSPNIEMDEKTFLVNRERAVDYLNSLDKVFV  240


>unnamed protein product [Arabidopsis thaliana]
Sequence ID: VYS71520.1 Length: 670
Range 1: 154 to 188

Score:49.3 bits(116), Expect:4e-09,
Method:Compositional matrix adjust.,
Identities:23/35(66%), Positives:28/35(80%), Gaps:0/35(0%)

Query  239  FSYSKLYEQAIEYEKGSLITSNGALTMLSGAKTDR  343
             S ++LYEQAI++EKGS +TS GAL  LSGAKT R
Sbjct  154  LSPAELYEQAIKFEKGSFVTSTGALATLSGAKTGR  188


Range 2: 202 to 240

Score:32.0 bits(71), Expect:4e-09,
Method:Compositional matrix adjust.,
Identities:18/39(46%), Positives:21/39(53%), Gaps:16/39(41%)

Query  382  ELELW*GK----------------ERAVDYLNSLEKVYL  450
            E ELW GK                ERAVDYLNSL+KV++
Sbjct  202  EAELWWGKGSPNIEMDEKTFLVNRERAVDYLNSLDKVFV  240


>PEPCK [Arabidopsis thaliana]
Sequence ID: OAO90231.1 Length: 670
Range 1: 154 to 188

Score:49.3 bits(116), Expect:4e-09,
Method:Compositional matrix adjust.,
Identities:23/35(66%), Positives:28/35(80%), Gaps:0/35(0%)

Query  239  FSYSKLYEQAIEYEKGSLITSNGALTMLSGAKTDR  343
             S ++LYEQAI++EKGS +TS GAL  LSGAKT R
Sbjct  154  LSPAELYEQAIKFEKGSFVTSTGALATLSGAKTGR  188


Range 2: 202 to 240

Score:32.0 bits(71), Expect:4e-09,
Method:Compositional matrix adjust.,
Identities:18/39(46%), Positives:21/39(53%), Gaps:16/39(41%)

Query  382  ELELW*GK----------------ERAVDYLNSLEKVYL  450
            E ELW GK                ERAVDYLNSL+KV++
Sbjct  202  EAELWWGKGSPNIEMDEKTFLVNRERAVDYLNSLDKVFV  240


>phosphoenolpyruvate carboxykinase 2 [Arabidopsis thaliana]
Sequence ID: NP_001331433.1 Length: 670
>phosphoenolpyruvate carboxykinase 2 [Arabidopsis thaliana]
Sequence ID: NP_680468.1 Length: 670 >At5g65690 [Arabidopsis thaliana]
Sequence ID: ACI49792.1 Length: 670 >phosphoenolpyruvate carboxykinase 2 [Arabidopsis thaliana]
Sequence ID: AED98087.1 Length: 670 >phosphoenolpyruvate carboxykinase 2 [Arabidopsis thaliana]
Sequence ID: ANM69779.1 Length: 670
Range 1: 154 to 188

Score:49.3 bits(116), Expect:4e-09,
Method:Compositional matrix adjust.,
Identities:23/35(66%), Positives:28/35(80%), Gaps:0/35(0%)

Query  239  FSYSKLYEQAIEYEKGSLITSNGALTMLSGAKTDR  343
             S ++LYEQAI++EKGS +TS GAL  LSGAKT R
Sbjct  154  LSPAELYEQAIKFEKGSFVTSTGALATLSGAKTGR  188


Range 2: 202 to 240

Score:32.0 bits(71), Expect:4e-09,
Method:Compositional matrix adjust.,
Identities:18/39(46%), Positives:21/39(53%), Gaps:16/39(41%)

Query  382  ELELW*GK----------------ERAVDYLNSLEKVYL  450
            E ELW GK                ERAVDYLNSL+KV++
Sbjct  202  EAELWWGKGSPNIEMDEKTFLVNRERAVDYLNSLDKVFV  240


>unnamed protein product [Arabidopsis thaliana]
Sequence ID: CAA0412274.1 Length: 670
Range 1: 154 to 188

Score:49.3 bits(116), Expect:4e-09,
Method:Compositional matrix adjust.,
Identities:23/35(66%), Positives:28/35(80%), Gaps:0/35(0%)

Query  239  FSYSKLYEQAIEYEKGSLITSNGALTMLSGAKTDR  343
             S ++LYEQAI++EKGS +TS GAL  LSGAKT R
Sbjct  154  LSPAELYEQAIKFEKGSFVTSTGALATLSGAKTGR  188


Range 2: 202 to 240

Score:32.0 bits(71), Expect:4e-09,
Method:Compositional matrix adjust.,
Identities:18/39(46%), Positives:21/39(53%), Gaps:16/39(41%)

Query  382  ELELW*GK----------------ERAVDYLNSLEKVYL  450
            E ELW GK                ERAVDYLNSL+KV++
Sbjct  202  EAELWWGKGSPNIEMDEKTFLVNRERAVDYLNSLDKVFV  240


>phosphoenolpyruvate carboxykinase 2 [Arabidopsis thaliana]
Sequence ID: NP_001331432.1 Length: 701
>phosphoenolpyruvate carboxykinase 2 [Arabidopsis thaliana]
Sequence ID: ANM69778.1 Length: 701
Range 1: 154 to 188

Score:49.3 bits(116), Expect:5e-09,
Method:Compositional matrix adjust.,
Identities:23/35(66%), Positives:28/35(80%), Gaps:0/35(0%)

Query  239  FSYSKLYEQAIEYEKGSLITSNGALTMLSGAKTDR  343
             S ++LYEQAI++EKGS +TS GAL  LSGAKT R
Sbjct  154  LSPAELYEQAIKFEKGSFVTSTGALATLSGAKTGR  188


Range 2: 202 to 240

Score:31.6 bits(70), Expect:5e-09,
Method:Compositional matrix adjust.,
Identities:18/39(46%), Positives:21/39(53%), Gaps:16/39(41%)

Query  382  ELELW*GK----------------ERAVDYLNSLEKVYL  450
            E ELW GK                ERAVDYLNSL+KV++
Sbjct  202  EAELWWGKGSPNIEMDEKTFLVNRERAVDYLNSLDKVFV  240


>phosphoenolpyruvate carboxykinase (ATP) (EC 4.1.1.49) [Arabidopsis thaliana]
Sequence ID: BAB10675.1 Length: 628
>phosphoenolpyruvate carboxykinase (ATP) - like protein [Arabidopsis thaliana]
Sequence ID: CAA16690.1 Length: 628
Range 1: 117 to 150

Score:49.3 bits(116), Expect:5e-09,
Method:Compositional matrix adjust.,
Identities:23/34(68%), Positives:28/34(82%), Gaps:0/34(0%)

Query  242  SYSKLYEQAIEYEKGSLITSNGALTMLSGAKTDR  343
            S ++LYEQAI++EKGS +TS GAL  LSGAKT R
Sbjct  117  SPAELYEQAIKFEKGSFVTSTGALATLSGAKTGR  150


Range 2: 164 to 202

Score:31.6 bits(70), Expect:5e-09,
Method:Compositional matrix adjust.,
Identities:18/39(46%), Positives:21/39(53%), Gaps:16/39(41%)

Query  382  ELELW*GK----------------ERAVDYLNSLEKVYL  450
            E ELW GK                ERAVDYLNSL+KV++
Sbjct  164  EAELWWGKGSPNIEMDEKTFLVNRERAVDYLNSLDKVFV  202


>unnamed protein product [Arabidopsis thaliana]
Sequence ID: CAA0361730.1 Length: 551
Range 1: 363 to 420

Score:57.8 bits(138), Expect:1e-08,
Method:Compositional matrix adjust.,
Identities:30/58(52%), Positives:40/58(68%), Gaps:2/58(3%)

Query  239  FSYSKLYEQAIEYEKGSLITSNGALTMLSGAKTDREISV--LLEMRLLKMSLSFGEEK  406
            F + +LYEQAI+YEKGS ITSNGAL  LSGAKT R  +   ++ +   K  L +G+E+
Sbjct  363  FLFVELYEQAIKYEKGSFITSNGALATLSGAKTGRAPTHKRIVRVAATKYELWWGKER  420


Query #260: XLOC_030321 Query ID: lcl|Query_30751 Length: 864

Sequences producing significant alignments:
                                                                  Max    Total Query   E   Per.
Description                                                       Score  Score cover Value Ident  Accession
At1g43205 [Arabidopsis thaliana]                                  175    175   41%   1e-54 72.13  ACB15003.1
contains a short region of similarity to transposases...          177    177   41%   4e-50 72.13  AAB61075.1
hypothetical protein AXX17_ATUG00950 [Arabidopsis thaliana]       155    155   41%   3e-47 68.07  OAO89407.1
predicted transposon protein [Arabidopsis thaliana]               159    227   50%   3e-46 66.96  AAB81877.1
At4g08605 [Arabidopsis thaliana]                                  100    165   36%   5e-28 69.33  AAS76215.1
hypothetical protein AT1G23915 [Arabidopsis thaliana]             63.2   63.2  38%   1e-11 35.59  NP_001320415.1
putative protein [Arabidopsis thaliana]                           63.2   63.2  38%   1e-11 35.59  CAB51204.1
hypothetical protein AXX17_ATUG02410 [Arabidopsis thaliana]       62.8   62.8  38%   2e-11 34.75  OAO89215.1


Alignments:
>At1g43205 [Arabidopsis thaliana]
Sequence ID: ACB15003.1 Length: 137
Range 1: 16 to 137

Score:175 bits(443), Expect:1e-54,
Method:Compositional matrix adjust.,
Identities:88/122(72%), Positives:99/122(81%), Gaps:3/122(2%)

Query  258  DDLCKYNGEYGVPTNCFCGKHLDLEERIIDNQKKTFLKCPMSGQD---HIDKWWDLAVHE  428
            DD  KY+G++GVP  CFCGK L+L ER+I +QKKTFLKCPMSGQD   H+DK WDL VHE
Sbjct  16   DDPSKYDGDFGVPQICFCGKQLELVERLIGDQKKTFLKCPMSGQDDNYHVDKGWDLVVHE  75

Query  429  *CFCIDKRLGDHMQLIQNALKFGGDSNRGAIHQFCAEIEKLKDRLDKKDVEIVQLMDALG  608
             CFCIDKR G+H +LIQNA KFGGDSNR  I+Q  AEIE LKDRLDKKD EI +LMDALG
Sbjct  76   QCFCIDKRFGEHRELIQNAFKFGGDSNRLQINQIRAEIEDLKDRLDKKDAEITRLMDALG  135

Query  609  KK  614
            KK
Sbjct  136  KK  137


>contains a short region of similarity to transposases [Arabidopsis thaliana]
Sequence ID: AAB61075.1 Length: 716
Range 1: 595 to 716

Score:177 bits(449), Expect:4e-50,
Method:Compositional matrix adjust.,
Identities:88/122(72%), Positives:99/122(81%), Gaps:3/122(2%)

Query  258  DDLCKYNGEYGVPTNCFCGKHLDLEERIIDNQKKTFLKCPMSGQD---HIDKWWDLAVHE  428
            DD  KY+G++GVP  CFCGK L+L ER+I +QKKTFLKCPMSGQD   H+DK WDLAVHE
Sbjct  595  DDPSKYDGDFGVPQICFCGKQLELVERLIGDQKKTFLKCPMSGQDDNYHVDKGWDLAVHE  654

Query  429  *CFCIDKRLGDHMQLIQNALKFGGDSNRGAIHQFCAEIEKLKDRLDKKDVEIVQLMDALG  608
             CFCIDKR G+H +LIQNA KFGGDSNR  I+Q  AEIE LK RLDKKD EI +LMDALG
Sbjct  655  QCFCIDKRFGEHRELIQNAFKFGGDSNRLQINQIHAEIEDLKHRLDKKDAEIARLMDALG  714

Query  609  KK  614
            KK
Sbjct  715  KK  716


>hypothetical protein AXX17_ATUG00950 [Arabidopsis thaliana]
Sequence ID: OAO89407.1 Length: 130
Range 1: 17 to 130

Score:155 bits(393), Expect:3e-47,
Method:Compositional matrix adjust.,
Identities:81/119(68%), Positives:91/119(76%), Gaps:5/119(4%)

Query  258  DDLCKYNGEYGVPTNCFCGKHLDLEERIIDNQKKTFLKCPMSGQDHIDKWWDLAVHE*CF  437
            DD  KY+G++GVPTNCF GK L L ERII NQKKTFLKCPMSGQD      +  VHE CF
Sbjct  17   DDPYKYDGDFGVPTNCFGGKQLYLSERIIGNQKKTFLKCPMSGQDD-----NYQVHEHCF  71

Query  438  CIDKRLGDHMQLIQNALKFGGDSNRGAIHQFCAEIEKLKDRLDKKDVEIVQLMDALGKK  614
             IDK+ G+H +LIQNA K+GGDSNR  I+Q   EIE LKDRLDKKD EI +LMDALGKK
Sbjct  72   FIDKQFGEHRELIQNAFKYGGDSNRLQINQIRVEIEDLKDRLDKKDAEIARLMDALGKK  130


>predicted transposon protein [Arabidopsis thaliana]
Sequence ID: AAB81877.1 Length: 907
>predicted transposon protein [Arabidopsis thaliana]
Sequence ID: CAB77986.1 Length: 907
Range 1: 764 to 871

Score:159 bits(401), Expect:3e-46,
Method:Composition-based stats.,
Identities:77/115(67%), Positives:88/115(76%), Gaps:10/115(8%)

Query  258  DDLCKYNGEYGVPTNCFCGKHLDLEERIIDNQKKTFLKCPMSGQD---HIDKWWDLAVHE  428
            DDLCKY+G++GVPT CFC       E +I +QKKTFLKCPMSG D   H+DK WDLAVHE
Sbjct  764  DDLCKYDGDFGVPTICFC-------ECLIGDQKKTFLKCPMSGHDDNYHVDKCWDLAVHE  816

Query  429  *CFCIDKRLGDHMQLIQNALKFGGDSNRGAIHQFCAEIEKLKDRLDKKDVEIVQL  593
             CFCIDKR G+H +LIQNA K+GGDSNR  I+Q  AEIE LKD LDKKD EI +L
Sbjct  817  QCFCIDKRFGEHRKLIQNAFKYGGDSNRLQINQIRAEIEDLKDHLDKKDAEIARL  871


Range 2: 886 to 907

Score:37.7 bits(86), Expect:3e-46,
Method:Composition-based stats.,
Identities:18/22(82%), Positives:19/22(86%), Gaps:2/22(9%)

Query  644  IVFPF--VTSMLKFETMLNSFF  703
            IVFPF   TSMLKFETMLNS+F
Sbjct  886  IVFPFYVATSMLKFETMLNSYF  907


Range 3: 872 to 883

Score:30.4 bits(67), Expect:3e-46,
Method:Composition-based stats.,
Identities:11/12(92%), Positives:12/12(100%), Gaps:0/12(0%)

Query  595  WMHLERSKMGIE  630
            WMHLERSKMGI+
Sbjct  872  WMHLERSKMGID  883


>At4g08605 [Arabidopsis thaliana]
Sequence ID: AAS76215.1 Length: 111
>At4g08605 [Arabidopsis thaliana]
Sequence ID: AAT41802.1 Length: 111
Range 1: 1 to 75

Score:100 bits(249), Expect:5e-28,
Method:Compositional matrix adjust.,
Identities:52/75(69%), Positives:58/75(77%), Gaps:3/75(4%)

Query  378  MSGQD---HIDKWWDLAVHE*CFCIDKRLGDHMQLIQNALKFGGDSNRGAIHQFCAEIEK  548
            MSG D   H+DK WDLAVHE CFCIDKR G+H +LIQNA K+GGDSNR  I+Q  AEIE
Sbjct  1    MSGHDDNYHVDKCWDLAVHEQCFCIDKRFGEHRKLIQNAFKYGGDSNRLQINQIRAEIED  60

Query  549  LKDRLDKKDVEIVQL  593
            LKD LDKKD EI +L
Sbjct  61   LKDHLDKKDAEIARL  75


Range 2: 90 to 111

Score:36.2 bits(82), Expect:5e-28,
Method:Compositional matrix adjust.,
Identities:18/22(82%), Positives:19/22(86%), Gaps:2/22(9%)

Query  644  IVFPF--VTSMLKFETMLNSFF  703
            IVFPF   TSMLKFETMLNS+F
Sbjct  90   IVFPFYVATSMLKFETMLNSYF  111


Range 3: 76 to 87

Score:28.5 bits(62), Expect:5e-28,
Method:Compositional matrix adjust.,
Identities:11/12(92%), Positives:12/12(100%), Gaps:0/12(0%)

Query  595  WMHLERSKMGIE  630
            WMHLERSKMGI+
Sbjct  76   WMHLERSKMGID  87


>hypothetical protein AT1G23915 [Arabidopsis thaliana]
Sequence ID: NP_001320415.1 Length: 142
>hypothetical protein AT1G23915 [Arabidopsis thaliana]
Sequence ID: ANM57940.1 Length: 142 >unnamed protein product [Arabidopsis thaliana]
Sequence ID: CAA0384485.1 Length: 142 >unnamed protein product [Arabidopsis thaliana]
Sequence ID: VYS62923.1 Length: 142
Range 1: 23 to 140

Score:63.2 bits(152), Expect:1e-11,
Method:Compositional matrix adjust.,
Identities:42/118(36%), Positives:59/118(50%), Gaps:7/118(5%)

Query  276  NGEYGVPTNCFCGKHLDLEE-RIIDNQKKTFLKCPM------SGQDHIDKWWDLAVHE*C  434
            +  +GVP  CFCGK + LE  +    Q     KCPM        ++H+ +WWD AV E
Sbjct  23   DANWGVPEKCFCGKPVKLEVCKTGYRQGDRVYKCPMLKAEEYESRNHLIQWWDQAVTEEL  82

Query  435  FCIDKRLGDHMQLIQNALKFGGDSNRGAIHQFCAEIEKLKDRLDKKDVEIVQLMDALG  608
              ++ +L   ++  Q A     +S R  IH    EIE LK+RL  K+VEI +L   L
Sbjct  83   EKVNHKLDSALRHGQLACNHALESIRETIHLMQGEIETLKERLLAKEVEIDRLRALLS  140


>putative protein [Arabidopsis thaliana]
Sequence ID: CAB51204.1 Length: 142
Range 1: 23 to 140

Score:63.2 bits(152), Expect:1e-11,
Method:Compositional matrix adjust.,
Identities:42/118(36%), Positives:59/118(50%), Gaps:7/118(5%)

Query  276  NGEYGVPTNCFCGKHLDLEE-RIIDNQKKTFLKCPM------SGQDHIDKWWDLAVHE*C  434
            +  +GVP  CFCGK + LE  +    Q     KCPM        ++H+ +WWD AV E
Sbjct  23   DANWGVPEKCFCGKPVKLEVCKTGYRQGDRVYKCPMLKAEEYESRNHLIQWWDQAVTEEL  82

Query  435  FCIDKRLGDHMQLIQNALKFGGDSNRGAIHQFCAEIEKLKDRLDKKDVEIVQLMDALG  608
              ++ +L   ++  Q A     +S R  IH    EIE LK+RL  K+VEI +L   L
Sbjct  83   EKVNHKLDSTLRHGQLACNHALESIRETIHLMQGEIETLKERLLAKEVEIDRLKALLS  140


>hypothetical protein AXX17_ATUG02410 [Arabidopsis thaliana]
Sequence ID: OAO89215.1 Length: 142
Range 1: 23 to 140

Score:62.8 bits(151), Expect:2e-11,
Method:Compositional matrix adjust.,
Identities:41/118(35%), Positives:59/118(50%), Gaps:7/118(5%)

Query  276  NGEYGVPTNCFCGKHLDLEE-RIIDNQKKTFLKCPM------SGQDHIDKWWDLAVHE*C  434
            +  +GVP  CFCGK + LE  +    Q     KCPM        ++H+ +WWD AV E
Sbjct  23   DANWGVPEKCFCGKPVKLEVCKTGYRQGDRVYKCPMLKAEEYESRNHLIQWWDQAVTEEL  82

Query  435  FCIDKRLGDHMQLIQNALKFGGDSNRGAIHQFCAEIEKLKDRLDKKDVEIVQLMDALG  608
              ++ +L   ++  Q A     +S R  +H    EIE LK+RL  K+VEI +L   L
Sbjct  83   EKVNHKLDSALRHGQLACNHALESIRETVHLMQGEIETLKERLLAKEVEIDRLRALLS  140


Query #261: XLOC_030379 Query ID: lcl|Query_30752 Length: 2094

Sequences producing significant alignments:
                                                                  Max    Total Query   E   Per.
Description                                                       Score  Score cover Value Ident  Accession
unnamed protein product [Arabidopsis thaliana]                    137    137   11%   2e-36 82.28  BAB02612.1
En/Spm-like transposon protein [Arabidopsis thaliana]             123    123   12%   2e-29 70.45  AAD03363.1
hypothetical protein AXX17_ATUG04920 [Arabidopsis thaliana]       62.8   62.8  12%   3e-09 44.71  OAO89060.1


Alignments:
>unnamed protein product [Arabidopsis thaliana]
Sequence ID: BAB02612.1 Length: 187
Range 1: 108 to 186

Score:137 bits(345), Expect:2e-36,
Method:Compositional matrix adjust.,
Identities:65/79(82%), Positives:70/79(88%), Gaps:0/79(0%)

Query  733  SLHAWQAFGDYGAFWSKTDNLSCRSNSQTKIRVGVERHHPWCQETPIPDGDSKTQLIDVL  912
            SLH   AFGDYGAFWSKTDNLSCRS+SQTKI VGVERHHPWC+ TPIPD +SKTQ IDVL
Sbjct  108  SLHVCDAFGDYGAFWSKTDNLSCRSSSQTKIPVGVERHHPWCRATPIPDDNSKTQFIDVL  167

Query  913  KISKFGRELSLISKEGPSR  969
            KIS +G E+SLISKEGPSR
Sbjct  168  KISIYGPEVSLISKEGPSR  186


>En/Spm-like transposon protein [Arabidopsis thaliana]
Sequence ID: AAD03363.1 Length: 454
Range 1: 111 to 198

Score:123 bits(309), Expect:2e-29,
Method:Compositional matrix adjust.,
Identities:62/88(70%), Positives:69/88(78%), Gaps:1/88(1%)

Query  709  ESFIVFFRSL-HAWQAFGDYGAFWSKTDNLSCRSNSQTKIRVGVERHHPWCQETPIPDGD  885
            E   V F SL     +FGD GA WSKTDNLS RSN +TK+ VGVERHHPWC+ TPIPD +
Sbjct  111  EKLEVEFTSLGEHVDSFGDNGALWSKTDNLSYRSNPETKMHVGVERHHPWCRATPIPDDN  170

Query  886  SKTQLIDVLKISKFGRELSLISKEGPSR  969
            SKTQLIDVLKISKFG +LS +SKEGPSR
Sbjct  171  SKTQLIDVLKISKFGPKLSPMSKEGPSR  198


>hypothetical protein AXX17_ATUG04920 [Arabidopsis thaliana]
Sequence ID: OAO89060.1 Length: 2484
Range 1: 1333 to 1414

Score:62.8 bits(151), Expect:3e-09,
Method:Composition-based stats.,
Identities:38/85(45%), Positives:47/85(55%), Gaps:3/85(3%)

Query  712   SFIVFFRSLHAWQAFGDYGAFWSKTDNLSCRSNSQTKIRVGVERHHPWCQETPIPDGDSK  891
             S+    RSLH W AFGDYG F SKTDNLSC SNS+T+I V +   +      P   G S+
Sbjct  1333  SYTFSLRSLHVWDAFGDYGTFRSKTDNLSCDSNSETEIDVCISSLYTLVSMHPF-HGLSQ  1391

Query  892   TQLIDVLKISKFGRELSLISKEGPS  966
                +D   I +F   +S I  EG S
Sbjct  1392  EHPMD--HIERFEVLISSIKVEGVS  1414


Query #262: XLOC_030593 Query ID: lcl|Query_30753 Length: 1087

No significant similarity found.


Query #263: XLOC_030748 Query ID: lcl|Query_30754 Length: 1008

No significant similarity found.


Query #264: XLOC_030751 Query ID: lcl|Query_30755 Length: 816

Sequences producing significant alignments:
                                                                  Max    Total Query   E   Per.
Description                                                       Score  Score cover Value Ident  Accession
RecName: Full=S-protein homolog 25; Flags: Precursor...           283    283   50%   1e-97 100.00 Q9FIV3.1
hypothetical protein AXX17_AT5G33960 [Arabidopsis thaliana]       281    281   50%   1e-96 99.26  OAO93312.1
RecName: Full=S-protein homolog 30; Flags: Precursor...           221    221   47%   3e-73 82.31  Q1G373.1
unknown [Arabidopsis thaliana]                                    221    221   47%   3e-73 82.31  ABK28126.1
Plant self-incompatibility protein S1 family [Arabidopsis...      110    110   49%   1e-29 41.01  NP_175536.2
unnamed protein product [Arabidopsis thaliana]                    108    108   49%   7e-29 41.01  CAA0286036.1
Plant self-incompatibility protein S1 family [Arabidopsis...      105    105   49%   1e-27 40.29  NP_175535.2
unnamed protein product [Arabidopsis thaliana]                    105    105   49%   2e-27 42.14  VYS45498.1
hypothetical protein AXX17_AT1G09120 [Arabidopsis thaliana]       105    105   49%   2e-27 42.14  OAP14490.1
unnamed protein product [Arabidopsis thaliana]                    103    103   49%   4e-27 39.57  CAA0286008.1
Plant self-incompatibility protein S1 family [Arabidopsis...      101    101   46%   3e-26 41.27  NP_001119337.2
unnamed protein product [Arabidopsis thaliana]                    101    101   49%   3e-26 41.43  CAA0182200.1
Plant self-incompatibility protein S1 family [Arabidopsis...      101    101   49%   3e-26 41.43  NP_849622.1
Plant self-incompatibility protein S1 family [Arabidopsis...      101    101   44%   3e-26 44.35  NP_175534.1
unnamed protein product [Arabidopsis thaliana]                    100    100   44%   5e-26 44.35  CAA0286002.1
unnamed protein product [Arabidopsis thaliana]                    100    100   46%   1e-25 40.48  CAA0406330.1
Plant self-incompatibility protein S1 family [Arabidopsis...      98.2   98.2  50%   9e-25 37.59  NP_001189509.1
unnamed protein product [Arabidopsis thaliana]                    96.3   96.3  44%   3e-24 41.27  VYS48749.1
unknown protein [Arabidopsis thaliana]                            95.1   95.1  48%   1e-23 37.68  ABF59425.1
unknown [Arabidopsis thaliana]                                    95.1   95.1  48%   1e-23 37.68  ABK28085.1
hypothetical protein AXX17_AT1G60280 [Arabidopsis thaliana]       85.5   85.5  42%   2e-20 41.74  OAP17805.1
hypothetical protein AXX17_AT4G14990 [Arabidopsis thaliana]       85.1   85.1  41%   4e-20 38.05  OAO97504.1
Chain A, S-protein homolog 15 [Arabidopsis thaliana]              84.3   84.3  39%   6e-20 40.19  6G7G_A
unnamed protein product [Arabidopsis thaliana]                    79.3   79.3  44%   6e-18 36.67  VYS56903.1
plant self-incompatibility protein S1 family protein...           79.3   79.3  44%   6e-18 36.67  NP_001325522.1
F21J9.32 [Arabidopsis thaliana]                                   74.7   74.7  50%   3e-16 34.53  AAF97971.1
unnamed protein product [Arabidopsis thaliana]                    68.2   68.2  46%   1e-13 34.62  VYS53238.1
Plant self-incompatibility protein S1 family [Arabidopsis...      65.9   65.9  46%   8e-13 33.85  NP_001118369.1
Plant self-incompatibility protein S1 family [Arabidopsis...      63.2   63.2  47%   8e-12 28.15  NP_178663.1
unnamed protein product [Arabidopsis thaliana]                    63.2   63.2  47%   9e-12 28.15  VYS52148.1
hypothetical protein AXX17_AT2G05160 [Arabidopsis thaliana]       62.8   62.8  47%   1e-11 28.15  OAP10771.1
hypothetical protein AXX17_AT2G18790 [Arabidopsis thaliana]       60.5   60.5  46%   1e-10 34.09  OAP07766.1
Plant self-incompatibility protein S1 family [Arabidopsis...      60.1   60.1  46%   1e-10 34.85  NP_001118368.1
unnamed protein product [Arabidopsis thaliana]                    59.7   59.7  46%   2e-10 34.09  VYS53237.1
hypothetical protein AXX17_AT5G03760 [Arabidopsis thaliana]       57.8   57.8  40%   6e-10 33.33  OAO94980.1
unnamed protein product [Arabidopsis thaliana]                    57.8   57.8  40%   6e-10 33.33  VYS65835.1
unnamed protein product [Arabidopsis thaliana]                    58.2   58.2  46%   7e-10 32.31  CAA0370215.1
hypothetical protein AXX17_AT5G35760 [Arabidopsis thaliana]       57.0   57.0  45%   1e-09 35.43  OAO93026.1
Plant self-incompatibility protein S1 family [Arabidopsis...      57.0   57.0  45%   1e-09 35.43  NP_198660.1
Plant self-incompatibility protein S1 family [Arabidopsis...      55.1   55.1  35%   9e-09 34.69  NP_196055.1
unnamed protein product [Arabidopsis thaliana]                    53.9   53.9  45%   1e-08 34.65  VYS68614.1


Alignments:
>RecName: Full=S-protein homolog 25; Flags: Precursor [Arabidopsis thaliana]
Sequence ID: Q9FIV3.1 Length: 136
>unnamed protein product [Arabidopsis thaliana]
Sequence ID: BAB11641.1 Length: 136
Range 1: 1 to 136

Score:283 bits(725), Expect:1e-97,
Method:Compositional matrix adjust.,
Identities:136/136(100%), Positives:136/136(100%), Gaps:0/136(0%)

Query  686  MNHSVFVILITITYFGLNQACIKNIVEILNQLAPGQILEYHCRSEDDNLGVKQLNFNATP  507
            MNHSVFVILITITYFGLNQACIKNIVEILNQLAPGQILEYHCRSEDDNLGVKQLNFNATP
Sbjct  1    MNHSVFVILITITYFGLNQACIKNIVEILNQLAPGQILEYHCRSEDDNLGVKQLNFNATP  60

Query  506  FVIRFHDEIPNLTRWNCIFRQGPNNSYSYDIEVYKAGPRLIPRCGQLRVWAARIDGIYFA  327
            FVIRFHDEIPNLTRWNCIFRQGPNNSYSYDIEVYKAGPRLIPRCGQLRVWAARIDGIYFA
Sbjct  61   FVIRFHDEIPNLTRWNCIFRQGPNNSYSYDIEVYKAGPRLIPRCGQLRVWAARIDGIYFA  120

Query  326  RKYNTPLVRVLSWNKN  279
            RKYNTPLVRVLSWNKN
Sbjct  121  RKYNTPLVRVLSWNKN  136


>hypothetical protein AXX17_AT5G33960 [Arabidopsis thaliana]
Sequence ID: OAO93312.1 Length: 136
Range 1: 1 to 136

Score:281 bits(719), Expect:1e-96,
Method:Compositional matrix adjust.,
Identities:135/136(99%), Positives:135/136(99%), Gaps:0/136(0%)

Query  686  MNHSVFVILITITYFGLNQACIKNIVEILNQLAPGQILEYHCRSEDDNLGVKQLNFNATP  507
            MNHSVFVILITITYFGLNQAC KNIVEILNQLAPGQILEYHCRSEDDNLGVKQLNFNATP
Sbjct  1    MNHSVFVILITITYFGLNQACKKNIVEILNQLAPGQILEYHCRSEDDNLGVKQLNFNATP  60

Query  506  FVIRFHDEIPNLTRWNCIFRQGPNNSYSYDIEVYKAGPRLIPRCGQLRVWAARIDGIYFA  327
            FVIRFHDEIPNLTRWNCIFRQGPNNSYSYDIEVYKAGPRLIPRCGQLRVWAARIDGIYFA
Sbjct  61   FVIRFHDEIPNLTRWNCIFRQGPNNSYSYDIEVYKAGPRLIPRCGQLRVWAARIDGIYFA  120

Query  326  RKYNTPLVRVLSWNKN  279
            RKYNTPLVRVLSWNKN
Sbjct  121  RKYNTPLVRVLSWNKN  136


>RecName: Full=S-protein homolog 30; Flags: Precursor [Arabidopsis thaliana]
Sequence ID: Q1G373.1 Length: 130
>unknown protein [Arabidopsis thaliana]
Sequence ID: ABF59468.1 Length: 130
Range 1: 1 to 130

Score:221 bits(564), Expect:3e-73,
Method:Compositional matrix adjust.,
Identities:107/130(82%), Positives:116/130(89%), Gaps:2/130(1%)

Query  665  ILITITYFGLNQACIKNIVEILNQLAPGQILEYHCRSEDDNLGVKQLNFNATPFVIRFHD  486
            +L+TIT FGLNQACIKN V ILN LAPG+ILEYHC S  D+LGVK+L+FNATPF I+FHD
Sbjct  1    MLVTITCFGLNQACIKNHVVILNLLAPGRILEYHCYSNVDDLGVKRLDFNATPFTIKFHD  60

Query  485  EIPNLTRWNCIFRQGPNNS--YSYDIEVYKAGPRLIPRCGQLRVWAARIDGIYFARKYNT  312
            EIPNLT+WNCI RQGPNNS  YSYD+EVYKAGPRLIPRCGQLR WAARIDGIYFARKYNT
Sbjct  61   EIPNLTKWNCILRQGPNNSMEYSYDVEVYKAGPRLIPRCGQLRAWAARIDGIYFARKYNT  120

Query  311  PLVRVLSWNK  282
            PL RVL WNK
Sbjct  121  PLKRVLFWNK  130


>unknown, partial [Arabidopsis thaliana]
Sequence ID: ABK28126.1 Length: 131
Range 1: 1 to 130

Score:221 bits(563), Expect:3e-73,
Method:Compositional matrix adjust.,
Identities:107/130(82%), Positives:116/130(89%), Gaps:2/130(1%)

Query  665  ILITITYFGLNQACIKNIVEILNQLAPGQILEYHCRSEDDNLGVKQLNFNATPFVIRFHD  486
            +L+TIT FGLNQACIKN V ILN LAPG+ILEYHC S  D+LGVK+L+FNATPF I+FHD
Sbjct  1    MLVTITCFGLNQACIKNHVVILNLLAPGRILEYHCYSNVDDLGVKRLDFNATPFTIKFHD  60

Query  485  EIPNLTRWNCIFRQGPNNS--YSYDIEVYKAGPRLIPRCGQLRVWAARIDGIYFARKYNT  312
            EIPNLT+WNCI RQGPNNS  YSYD+EVYKAGPRLIPRCGQLR WAARIDGIYFARKYNT
Sbjct  61   EIPNLTKWNCILRQGPNNSMEYSYDVEVYKAGPRLIPRCGQLRAWAARIDGIYFARKYNT  120

Query  311  PLVRVLSWNK  282
            PL RVL WNK
Sbjct  121  PLKRVLFWNK  130


>Plant self-incompatibility protein S1 family [Arabidopsis thaliana]
Sequence ID: NP_175536.2 Length: 139
>Hypothetical protein [Arabidopsis thaliana]
Sequence ID: AAD30653.1 Length: 139 >At1g51250 [Arabidopsis thaliana]
Sequence ID: AAS76210.1 Length: 139 >At1g51250 [Arabidopsis thaliana]
Sequence ID: AAW70392.1 Length: 139 >Plant self-incompatibility protein S1 family [Arabidopsis thaliana]
Sequence ID: AEE32641.1 Length: 139 >hypothetical protein AXX17_AT1G45440 [Arabidopsis thaliana]
Sequence ID: OAP19848.1 Length: 139
Range 1: 1 to 137

Score:110 bits(275), Expect:1e-29,
Method:Compositional matrix adjust.,
Identities:57/139(41%), Positives:79/139(56%), Gaps:7/139(5%)

Query  686  MNHSVFVILITITYFGLNQACIKNIVEILNQLAPGQILEYHCRSEDDNLG---VKQLNFN  516
            M+H +F++L+T  YFG+N+AC KN V I N++ PG  L   CR             L +
Sbjct  1    MDHLIFLLLVTTMYFGINEACKKNHVVIHNEIGPGIDLNIACRQFSPKRTPSMFHTLKYK  60

Query  515  ATPFVIRFHD--EIPNLTRWNCIFRQGPNNSYSYDIEVYKAGPRLIPRCGQLRVWAARID  342
               ++I F D  ++P+  +W C+   G    Y +DIEVY+ G    PRCGQLR W AR D
Sbjct  61   DPFYIIEFADNNQLPHGEKWYCLISHGTRPKYWFDIEVYRQG--YYPRCGQLRSWIARKD  118

Query  341  GIYFARKYNTPLVRVLSWN  285
            GI+F RKY++P   VL W
Sbjct  119  GIWFTRKYHSPPGHVLDWK  137


>unnamed protein product [Arabidopsis thaliana]
Sequence ID: CAA0286036.1 Length: 139
Range 1: 1 to 137

Score:108 bits(270), Expect:7e-29,
Method:Compositional matrix adjust.,
Identities:57/139(41%), Positives:77/139(55%), Gaps:7/139(5%)

Query  686  MNHSVFVILITITYFGLNQACIKNIVEILNQLAPGQILEYHCRSEDDNLG---VKQLNFN  516
            M+H +F +L+T  YFG N+AC KN V I N++ PG  L   CR             L +
Sbjct  1    MDHLIFFLLVTTMYFGRNEACKKNHVVIHNEIGPGIDLNIACRQFSPKRTPSMFHTLKYK  60

Query  515  ATPFVIRFHD--EIPNLTRWNCIFRQGPNNSYSYDIEVYKAGPRLIPRCGQLRVWAARID  342
               ++I F D  ++P+  +W C+   G    Y +DIEVY+ G    PRCGQLR W AR D
Sbjct  61   DPFYIIEFADNNQLPHGEKWYCLISHGTRPKYWFDIEVYRQG--YYPRCGQLRSWIARKD  118

Query  341  GIYFARKYNTPLVRVLSWN  285
            GI+F RKY++P   VL W
Sbjct  119  GIWFTRKYHSPPGHVLDWK  137


>Plant self-incompatibility protein S1 family [Arabidopsis thaliana]
Sequence ID: NP_175535.2 Length: 139
>Hypothetical protein [Arabidopsis thaliana]
Sequence ID: AAD30637.1 Length: 139 >Plant self-incompatibility protein S1 family [Arabidopsis thaliana]
Sequence ID: AEE32640.2 Length: 139
Range 1: 1 to 137

Score:105 bits(262), Expect:1e-27,
Method:Compositional matrix adjust.,
Identities:56/139(40%), Positives:77/139(55%), Gaps:7/139(5%)

Query  686  MNHSVFVILITITYFGLNQACIKNIVEILNQLAPGQILEYHCRS---EDDNLGVKQLNFN  516
            M++ +  +L+T  YFG N+AC KN V I N+L PG  L   CR    E        L +
Sbjct  1    MDYLIMFLLVTTMYFGQNEACKKNHVVIHNELGPGIDLNIACRQFSIERTPSMFHTLKYK  60

Query  515  ATPFVIRFHD--EIPNLTRWNCIFRQGPNNSYSYDIEVYKAGPRLIPRCGQLRVWAARID  342
               ++I F D  ++P+  RW C+   G    Y YDIEVY+ G    P CGQLR W A+ D
Sbjct  61   DPFYIIEFEDNNQLPHGERWYCLLSHGTRPKYWYDIEVYRQG--YYPSCGQLRSWIAKQD  118

Query  341  GIYFARKYNTPLVRVLSWN  285
            GI+F R+Y++P   VL+W
Sbjct  119  GIWFTRRYDSPPGHVLNWK  137


>unnamed protein product [Arabidopsis thaliana]
Sequence ID: VYS45498.1 Length: 139
Range 1: 1 to 137

Score:105 bits(261), Expect:2e-27,
Method:Compositional matrix adjust.,
Identities:59/140(42%), Positives:74/140(52%), Gaps:8/140(5%)

Query  686  MNHSVFVILITITYFGLNQA-----CIKNIVEILNQLAPGQILEYHCRSEDDNLGVKQLN  522
            MN  +  + + +TYFGLN A     C KN V I N+L PG +L+YHC S D NL V  L
Sbjct  1    MNRFIIFMFVVVTYFGLNVAFDMFPCPKNKVLIRNELGPGLVLQYHCHSRDHNLDVANLQ  60

Query  521  FNATPFVIRFHDEIPNLTRWNCIFRQGPNNSYSYDIEVYKAGPRLIPRCGQLRVWAARID  342
            FN     I F D++   TRW+CI + G    Y  +   Y      + RCG +R W AR D
Sbjct  61   FNEYK-EIAFGDKLGKRTRWSCILKHGLYMRYYSEFIAYMMAN--VRRCGAIRNWIARKD  117

Query  341  GIYFARKYNTPLVRVLSWNK  282
            GIY  R  N P V   +WNK
Sbjct  118  GIYLIRNVNPPTVFRYTWNK  137


>hypothetical protein AXX17_AT1G09120 [Arabidopsis thaliana]
Sequence ID: OAP14490.1 Length: 139
Range 1: 1 to 137

Score:105 bits(261), Expect:2e-27,
Method:Compositional matrix adjust.,
Identities:59/140(42%), Positives:74/140(52%), Gaps:8/140(5%)

Query  686  MNHSVFVILITITYFGLNQA-----CIKNIVEILNQLAPGQILEYHCRSEDDNLGVKQLN  522
            MN  +  + + +TYFGLN A     C KN V I N+L PG +L+YHC S D NL V  L
Sbjct  1    MNRFIIFMFVVVTYFGLNVAFDMFPCPKNKVLIRNELGPGLVLQYHCHSRDHNLDVANLQ  60

Query  521  FNATPFVIRFHDEIPNLTRWNCIFRQGPNNSYSYDIEVYKAGPRLIPRCGQLRVWAARID  342
            FN     I F D++   TRW+CI + G    Y  +   Y      + RCG +R W AR D
Sbjct  61   FNEYK-EIAFGDKLGKRTRWSCILKHGLYMRYYSEFIAYMMAN--VRRCGAIRNWIARKD  117

Query  341  GIYFARKYNTPLVRVLSWNK  282
            GIY  R  N P V   +WNK
Sbjct  118  GIYLIRNVNPPAVFRYTWNK  137


>unnamed protein product [Arabidopsis thaliana]
Sequence ID: CAA0286008.1 Length: 139
Range 1: 1 to 137

Score:103 bits(258), Expect:4e-27,
Method:Compositional matrix adjust.,
Identities:55/139(40%), Positives:77/139(55%), Gaps:7/139(5%)

Query  686  MNHSVFVILITITYFGLNQACIKNIVEILNQLAPGQILEYHCRS---EDDNLGVKQLNFN  516
            M++ +  +L+T  YFG  +AC KN V I N+L+PG  L   CR    E        L +
Sbjct  1    MDYLIMFLLVTTMYFGQTEACKKNHVVIHNELSPGIDLNIACRQFSIERTPSMFHTLKYK  60

Query  515  ATPFVIRFHD--EIPNLTRWNCIFRQGPNNSYSYDIEVYKAGPRLIPRCGQLRVWAARID  342
               ++I F D  ++P+  RW C+   G    Y YDIEVY+ G    P CGQLR W A+ D
Sbjct  61   DPFYIIEFEDNNQLPHGERWYCLLSHGTRPKYWYDIEVYRQG--YYPSCGQLRSWIAKQD  118

Query  341  GIYFARKYNTPLVRVLSWN  285
            GI+F R+Y++P   VL+W
Sbjct  119  GIWFTRRYDSPPGHVLNWK  137


>Plant self-incompatibility protein S1 family [Arabidopsis thaliana]
Sequence ID: NP_001119337.2 Length: 132
>RecName: Full=S-protein homolog 15; Flags: Precursor [Arabidopsis thaliana]
Sequence ID: Q9FLY6.1 Length: 132 >Plant self-incompatibility protein S1 family [Arabidopsis thaliana]
Sequence ID: AED94440.1 Length: 132 >hypothetical protein AXX17_AT5G36920 [Arabidopsis thaliana]
Sequence ID: OAO90129.1 Length: 132 >unnamed protein product [Arabidopsis thaliana]
Sequence ID: VYS68726.1 Length: 132 >unnamed protein product [Arabidopsis thaliana]
Sequence ID: BAB11024.1 Length: 132
Range 1: 1 to 124

Score:101 bits(252), Expect:3e-26,
Method:Compositional matrix adjust.,
Identities:52/126(41%), Positives:76/126(60%), Gaps:2/126(1%)

Query  686  MNHSVFVILITITYFGLNQACIKNIVEILNQLAPGQILEYHCRSEDDNLGVKQLNFNATP  507
            M+  +F IL+T  YF  N+AC +  + I N L P +IL+YHCRS + N+GV+ LNF  T
Sbjct  1    MSRLIFFILVTAIYFVGNEACKEIEIVIKNTLGPSRILQYHCRSGNTNVGVQYLNFKGTR  60

Query  506  FVIRFHDEIPNLTRWNCIFRQGPNNSYSYDIEVYKAGPRLIPRCGQLRVWAARIDGIYFA  327
             +I+F D+    +RWNC+FRQG N  +  ++E Y+   +  P CG+    +AR+D IYF
Sbjct  61   -IIKFKDDGTERSRWNCLFRQGINMKFFTEVEAYRPDLKH-PLCGKRYELSARMDAIYFK  118

Query  326  RKYNTP  309
                 P
Sbjct  119  MDERPP  124


>unnamed protein product [Arabidopsis thaliana]
Sequence ID: CAA0182200.1 Length: 139
Range 1: 1 to 137

Score:101 bits(252), Expect:3e-26,
Method:Compositional matrix adjust.,
Identities:58/140(41%), Positives:72/140(51%), Gaps:8/140(5%)

Query  686  MNHSVFVILITITYFGLNQA-----CIKNIVEILNQLAPGQILEYHCRSEDDNLGVKQLN  522
            MN  +  + +  TYFGLN A     C K  V I N+L PG +L+YHC S D NL V  L
Sbjct  1    MNRFIIFMFVVATYFGLNVAFDMFPCPKTKVLIRNELGPGLVLQYHCHSRDHNLDVANLQ  60

Query  521  FNATPFVIRFHDEIPNLTRWNCIFRQGPNNSYSYDIEVYKAGPRLIPRCGQLRVWAARID  342
            FN     I F D++   TRW+CI + G    Y  +   Y      + RCG +R W AR D
Sbjct  61   FNEYK-EIAFGDKLGKRTRWSCILKHGLYMRYYSEFIAYMMAN--VRRCGAIRNWIARKD  117

Query  341  GIYFARKYNTPLVRVLSWNK  282
            GIY  R  N P V   +WNK
Sbjct  118  GIYLIRNVNPPAVFRYTWNK  137


>Plant self-incompatibility protein S1 family [Arabidopsis thaliana]
Sequence ID: NP_849622.1 Length: 139
>RecName: Full=S-protein homolog 14; Flags: Precursor [Arabidopsis thaliana]
Sequence ID: Q8GX02.1 Length: 139 >At1g09245 [Arabidopsis thaliana]
Sequence ID: AAO39885.1 Length: 139 >Plant self-incompatibility protein S1 family [Arabidopsis thaliana]
Sequence ID: AEE28418.1 Length: 139 >unknown protein [Arabidopsis thaliana]
Sequence ID: BAC43125.1 Length: 139
Range 1: 1 to 137

Score:101 bits(252), Expect:3e-26,
Method:Compositional matrix adjust.,
Identities:58/140(41%), Positives:73/140(52%), Gaps:8/140(5%)

Query  686  MNHSVFVILITITYFGLNQA-----CIKNIVEILNQLAPGQILEYHCRSEDDNLGVKQLN  522
            MN  +  + + +TYFGLN A     C KN V I N+L PG +L+YHC S D NL V  L
Sbjct  1    MNRFIIFMFVVVTYFGLNVAFDMFPCPKNKVLIRNELGPGLVLQYHCHSRDHNLDVANLQ  60

Query  521  FNATPFVIRFHDEIPNLTRWNCIFRQGPNNSYSYDIEVYKAGPRLIPRCGQLRVWAARID  342
            FN     I F D++   TRW+CI + G    Y  +   Y      + RCG +R W AR D
Sbjct  61   FNEYK-EIAFGDKLGKRTRWSCILKHGLYMRYYSEFIAYMMAN--VRRCGAIRNWIARKD  117

Query  341  GIYFARKYNTPLVRVLSWNK  282
             IY  R  N P V   +WNK
Sbjct  118  RIYLIRNVNPPAVFRYTWNK  137


>Plant self-incompatibility protein S1 family [Arabidopsis thaliana]
Sequence ID: NP_175534.1 Length: 125
>Plant self-incompatibility protein S1 family [Arabidopsis thaliana]
Sequence ID: AEE32639.1 Length: 125 >unnamed protein product [Arabidopsis thaliana]
Sequence ID: VYS48747.1 Length: 125
Range 1: 2 to 123

Score:101 bits(251), Expect:3e-26,
Method:Compositional matrix adjust.,
Identities:55/124(44%), Positives:71/124(57%), Gaps:6/124(4%)

Query  647  YFGLNQAC-IKNIVEILNQLAPGQILEYHCRS---EDDNLGVKQLNFNATPFVIRFHDEI  480
            +F LNQA   KN V I N+L+PG IL   CR    +D      +LNF    +VI F D
Sbjct  2    HFELNQATHCKNHVVIHNELSPGIILNIACRKGSIDDKPTRFHKLNFKDPSYVIEFEDNY  61

Query  479  PNLTRWNCIFRQGPNNSYSYDIEVYKAGPRLIPRCGQLRVWAARIDGIYFARKYNTPLVR  300
            PN   W C+   G    Y YDI+VY+ G  + P+C QLR+W AR+DGI+F R+Y +P
Sbjct  62   PNDETWYCLLSHGTRPKYWYDIQVYRQG--IQPKCNQLRLWIARMDGIWFTRRYESPPGH  119

Query  299  VLSW  288
            VL W
Sbjct  120  VLDW  123


>unnamed protein product [Arabidopsis thaliana]
Sequence ID: CAA0286002.1 Length: 125
Range 1: 2 to 123

Score:100 bits(250), Expect:5e-26,
Method:Compositional matrix adjust.,
Identities:55/124(44%), Positives:71/124(57%), Gaps:6/124(4%)

Query  647  YFGLNQAC-IKNIVEILNQLAPGQILEYHCRS---EDDNLGVKQLNFNATPFVIRFHDEI  480
            +F LNQA   KN V I N+L+PG IL   CR    +D      +LNF    +VI F D
Sbjct  2    HFELNQATHCKNHVIIHNELSPGIILNIACRKGSIDDKPTRFHKLNFKDPSYVIEFEDNY  61

Query  479  PNLTRWNCIFRQGPNNSYSYDIEVYKAGPRLIPRCGQLRVWAARIDGIYFARKYNTPLVR  300
            PN   W C+   G    Y YDI+VY+ G  + P+C QLR+W AR+DGI+F R+Y +P
Sbjct  62   PNDETWYCLLSHGTRPKYWYDIQVYRQG--IQPKCNQLRLWIARMDGIWFTRRYESPPGH  119

Query  299  VLSW  288
            VL W
Sbjct  120  VLDW  123


>unnamed protein product [Arabidopsis thaliana]
Sequence ID: CAA0406330.1 Length: 132
>unnamed protein product [Arabidopsis thaliana]
Sequence ID: CAA0406332.1 Length: 132
Range 1: 1 to 124

Score:100 bits(248), Expect:1e-25,
Method:Compositional matrix adjust.,
Identities:51/126(40%), Positives:75/126(59%), Gaps:2/126(1%)

Query  686  MNHSVFVILITITYFGLNQACIKNIVEILNQLAPGQILEYHCRSEDDNLGVKQLNFNATP  507
            M+  +F IL+T  YF  N+AC +  + I N L P +IL+YHCRS + N+GV+ LNF  T
Sbjct  1    MSRLIFFILVTAIYFVGNEACKEIEIVIKNTLGPSRILQYHCRSGNTNVGVQYLNFKGTR  60

Query  506  FVIRFHDEIPNLTRWNCIFRQGPNNSYSYDIEVYKAGPRLIPRCGQLRVWAARIDGIYFA  327
             +I+F D+    +RW C+FRQG N  +  ++E Y+   +  P CG+    +AR+D IYF
Sbjct  61   -IIKFKDDGTERSRWKCLFRQGINMKFFTEVEAYRPDLKH-PLCGKRYELSARMDAIYFK  118

Query  326  RKYNTP  309
                 P
Sbjct  119  MDERPP  124


>Plant self-incompatibility protein S1 family [Arabidopsis thaliana]
Sequence ID: NP_001189509.1 Length: 151
>Plant self-incompatibility protein S1 family [Arabidopsis thaliana]
Sequence ID: AEC05800.1 Length: 151
Range 1: 12 to 149

Score:98.2 bits(243), Expect:9e-25,
Method:Compositional matrix adjust.,
Identities:53/141(38%), Positives:77/141(54%), Gaps:8/141(5%)

Query  695  IQKMNHSVFVILITITYFGLNQAC-IKNIVEILNQLAPGQILEYHCRSE--DDNLGVKQL  525
            ++KMNH +   L+   +F  N+AC   N+VEI NQLAPG++L++HCR    + + GV+ L
Sbjct  12   LRKMNHLIIFTLVIAIHFSFNEACNTPNVVEIHNQLAPGKLLKHHCRGSINEQDKGVQYL  71

Query  524  NFNATPFVIRFHDEIPNLTR--WNCIFRQGPNNSYSYDIEVYKAGPRLIPRCGQLRVWAA  351
              N   F I F D      R  W C+   G    + ++++VY+A      RC Q R W A
Sbjct  72   KVNQN-FTIVFSDVSNRRERTVWTCMLNHGDKMEFHFNLQVYRAAA--TERCNQYRSWTA  128

Query  350  RIDGIYFARKYNTPLVRVLSW  288
            + DGI+F R  N P   V +W
Sbjct  129  KPDGIWFRRDRNKPSGHVRNW  149


>unnamed protein product [Arabidopsis thaliana]
Sequence ID: VYS48749.1 Length: 127
Range 1: 2 to 125

Score:96.3 bits(238), Expect:3e-24,
Method:Compositional matrix adjust.,
Identities:52/126(41%), Positives:70/126(55%), Gaps:7/126(5%)

Query  647  YFGLNQACIKNIVEILNQLAPGQILEYHCRS---EDDNLGVKQLNFNATPFVIRFHD--E  483
            YFG+N+AC KN V I N++ PG  L   CR    +        L +    ++I F D  +
Sbjct  2    YFGINEACKKNHVVIHNEIGPGIDLNIACRQFSPKRTPSMFHTLKYKDPFYIIEFADNNQ  61

Query  482  IPNLTRWNCIFRQGPNNSYSYDIEVYKAGPRLIPRCGQLRVWAARIDGIYFARKYNTPLV  303
            +P+  +W C+   G    Y +DIEVY+ G    PRCGQLR W AR DGI+F RKY++P
Sbjct  62   LPHGEKWYCLISHGTRPKYWFDIEVYRQG--YYPRCGQLRSWIARKDGIWFTRKYHSPPG  119

Query  302  RVLSWN  285
             VL W
Sbjct  120  HVLDWK  125


>unknown protein [Arabidopsis thaliana]
Sequence ID: ABF59425.1 Length: 137
>hypothetical protein AXX17_AT2G03180 [Arabidopsis thaliana]
Sequence ID: OAP08874.1 Length: 137 >unnamed protein product [Arabidopsis thaliana]
Sequence ID: CAA0357056.1 Length: 137 >unnamed protein product [Arabidopsis thaliana]
Sequence ID: VYS52002.1 Length: 137
Range 1: 1 to 135

Score:95.1 bits(235), Expect:1e-23,
Method:Compositional matrix adjust.,
Identities:52/138(38%), Positives:74/138(53%), Gaps:8/138(5%)

Query  686  MNHSVFVILITITYFGLNQAC-IKNIVEILNQLAPGQILEYHCRSE--DDNLGVKQLNFN  516
            MNH +   L+   +F  N+AC   N+VEI NQLAPG++L++HCR    + + GV+ L  N
Sbjct  1    MNHLIIFTLVIAIHFSFNEACNTPNVVEIHNQLAPGKLLKHHCRGSINEQDKGVQYLKVN  60

Query  515  ATPFVIRFHDEIPNLTR--WNCIFRQGPNNSYSYDIEVYKAGPRLIPRCGQLRVWAARID  342
               F I F D      R  W C+   G    + ++++VY+A      RC Q R W A+ D
Sbjct  61   QN-FTIVFSDVSNRRERTVWTCMLNHGDKMEFHFNLQVYRAAA--TERCNQYRSWTAKPD  117

Query  341  GIYFARKYNTPLVRVLSW  288
            GI+F R  N P   V +W
Sbjct  118  GIWFRRDRNKPSGHVRNW  135


>unknown, partial [Arabidopsis thaliana]
Sequence ID: ABK28085.1 Length: 138
Range 1: 1 to 135

Score:95.1 bits(235), Expect:1e-23,
Method:Compositional matrix adjust.,
Identities:52/138(38%), Positives:74/138(53%), Gaps:8/138(5%)

Query  686  MNHSVFVILITITYFGLNQAC-IKNIVEILNQLAPGQILEYHCRSE--DDNLGVKQLNFN  516
            MNH +   L+   +F  N+AC   N+VEI NQLAPG++L++HCR    + + GV+ L  N
Sbjct  1    MNHLIIFTLVIAIHFSFNEACNTPNVVEIHNQLAPGKLLKHHCRGSINEQDKGVQYLKVN  60

Query  515  ATPFVIRFHDEIPNLTR--WNCIFRQGPNNSYSYDIEVYKAGPRLIPRCGQLRVWAARID  342
               F I F D      R  W C+   G    + ++++VY+A      RC Q R W A+ D
Sbjct  61   QN-FTIVFSDVSNRRERTVWTCMLNHGDKMEFHFNLQVYRAAA--TERCNQYRSWTAKPD  117

Query  341  GIYFARKYNTPLVRVLSW  288
            GI+F R  N P   V +W
Sbjct  118  GIWFRRDRNKPSGHVRNW  135


>hypothetical protein AXX17_AT1G60280 [Arabidopsis thaliana]
Sequence ID: OAP17805.1 Length: 116
Range 1: 4 to 114

Score:85.5 bits(210), Expect:2e-20,
Method:Compositional matrix adjust.,
Identities:48/115(42%), Positives:60/115(52%), Gaps:4/115(3%)

Query  626  CIKNIVEILNQLAPGQILEYHCRSEDDNLGVKQLNFNATPFVIRFHDEIPNLTRWNCIFR  447
            C  N + I N+L PG +L+YHC S DDNL V+ L FN T   IR  D+I   TRW C+ +
Sbjct  4    CPSNYILITNELGPGLVLQYHCHSRDDNLNVENLQFNETK-KIRLADKIGKRTRWTCLLK  62

Query  446  QGPNNSYSYDIEVYKAGPRLIPRCGQLRVWAARIDGIYFARKYNTPLVRVLSWNK  282
             G    Y  +   Y+     + RCG  R W  R  GIY  R  N P V+   WNK
Sbjct  63   HGLYMRYYSEFIGYRMAN--VRRCGASRQWIVRKAGIYLTRNQNPPPVQHF-WNK  114


>hypothetical protein AXX17_AT4G14990 [Arabidopsis thaliana]
Sequence ID: OAO97504.1 Length: 127
Range 1: 1 to 111

Score:85.1 bits(209), Expect:4e-20,
Method:Compositional matrix adjust.,
Identities:43/113(38%), Positives:67/113(59%), Gaps:2/113(1%)

Query  665  ILITITYFGLNQACIKNIVEILNQLAPGQILEYHCRSEDDNLGVKQLNFNATPFVIRFHD  486
            +LI   ++G+N+AC +N + + N+L   +IL+YHC S   +LGV  L+FNA   +I+  D
Sbjct  1    MLIIAIFYGVNEACKENRIVLKNRLGHSKILQYHCHSPKVDLGVLYLDFNAIR-IIKVKD  59

Query  485  EIPNLTRWNCIFRQGPNNSYSYDIEVYKAGPRLIPRCGQLRVWAARIDGIYFA  327
            E  N+T+W+C+F+ G N  +   I+VY       P+C QLR W+     I F
Sbjct  60   EGVNITKWDCLFKHGINMRFHSYIQVYSQNT-FAPQCDQLRQWSFTKSAIGFT  111


>Chain A, S-protein homolog 15 [Arabidopsis thaliana]
Sequence ID: 6G7G_A Length: 115
Range 1: 3 to 107

Score:84.3 bits(207), Expect:6e-20,
Method:Compositional matrix adjust.,
Identities:43/107(40%), Positives:63/107(58%), Gaps:2/107(1%)

Query  629  ACIKNIVEILNQLAPGQILEYHCRSEDDNLGVKQLNFNATPFVIRFHDEIPNLTRWNCIF  450
             C +  + I N L P +IL+YHCRS + N+GV+ LNF  T  +I+F D+    +RWNC+F
Sbjct  3    GCKEIEIVIKNTLGPSRILQYHCRSGNTNVGVQYLNFKGTR-IIKFKDDGTERSRWNCLF  61

Query  449  RQGPNNSYSYDIEVYKAGPRLIPRCGQLRVWAARIDGIYFARKYNTP  309
            RQG N  +  ++E Y+   +  P CG+    +AR+D IYF      P
Sbjct  62   RQGINMKFFTEVEAYRPDLKH-PLCGKRYELSARMDAIYFKMDERPP  107


>unnamed protein product [Arabidopsis thaliana]
Sequence ID: VYS56903.1 Length: 131
Range 1: 1 to 116

Score:79.3 bits(194), Expect:6e-18,
Method:Compositional matrix adjust.,
Identities:44/120(37%), Positives:63/120(52%), Gaps:4/120(3%)

Query  686  MNHSVFVILITITYFGLNQACIKNIVEILNQLAPGQILEYHCRSEDDNLGVKQLNFNATP  507
            MN  +  +LI    FGLN+AC    +   N L+PG IL+ +C S + N     + F +
Sbjct  1    MNRLIAFLLIIALSFGLNKACEDCTIVFRNNLSPGIILKVNCESNNKNRVTGTVKFQSDT  60

Query  506  FVIRFHDEIPNLTRWNCIFRQGPNNSYSYDIEVYKAGPRLIPRCGQLRVWAARIDGIYFA  327
              I F +     T W+C+ +QG    YS     Y+ G   IPRCG+LRV+ A+ DGIY +
Sbjct  61   VRINFREAAFERTTWHCLVQQG---EYSQHFRAYR-GSAPIPRCGELRVYIAKRDGIYLS  116


>plant self-incompatibility protein S1 family protein [Arabidopsis thaliana]
Sequence ID: NP_001325522.1 Length: 131
>plant self-incompatibility protein S1 family protein [Arabidopsis thaliana]
Sequence ID: ANM63434.1 Length: 131 >hypothetical protein AXX17_AT3G10260 [Arabidopsis thaliana]
Sequence ID: OAP06845.1 Length: 131 >unnamed protein product [Arabidopsis thaliana]
Sequence ID: CAA0381949.1 Length: 131
Range 1: 1 to 116

Score:79.3 bits(194), Expect:6e-18,
Method:Compositional matrix adjust.,
Identities:44/120(37%), Positives:63/120(52%), Gaps:4/120(3%)

Query  686  MNHSVFVILITITYFGLNQACIKNIVEILNQLAPGQILEYHCRSEDDNLGVKQLNFNATP  507
            MN  +  +LI    FGLN+AC    +   N L+PG IL+ +C S + N     + F +
Sbjct  1    MNRLIAFLLIIALSFGLNKACEDCTIVFRNNLSPGIILKVNCESNNKNRVTGTVKFQSDT  60

Query  506  FVIRFHDEIPNLTRWNCIFRQGPNNSYSYDIEVYKAGPRLIPRCGQLRVWAARIDGIYFA  327
              I F +     T W+C+ +QG    YS     Y+ G   IPRCG+LRV+ A+ DGIY +
Sbjct  61   VRINFREAAFERTTWHCLVQQG---GYSQHFRAYR-GSAPIPRCGELRVYIAKRDGIYLS  116


>F21J9.32 [Arabidopsis thaliana]
Sequence ID: AAF97971.1 Length: 116
Range 1: 1 to 115

Score:74.7 bits(182), Expect:3e-16,
Method:Compositional matrix adjust.,
Identities:48/139(35%), Positives:64/139(46%), Gaps:27/139(19%)

Query  686  MNHSVFVILITITYFGL---NQACIKNIVEILNQLAPGQILEYHCRSEDDNLGVKQLNFN  516
            MN  +  + +  TYFGL     +C +N +EI+N+L PG             L +KQ
Sbjct  1    MNRFIIFMFVIATYFGLLVSVLSCKRNTIEIMNELGPG-------------LEIKQ----  43

Query  515  ATPFVIRFHDEIPNLTRWNCIFRQGPNNSYSYDIEVYKAGPRLIPRCGQLRVWAARIDGI  336
                 IRF D++   TRW C+ + G    Y  +   Y+AG   I RCG LR W  R DGI
Sbjct  44   -----IRFGDKLGRRTRWTCLLKHGLYMRYYKEFIAYRAGN--IDRCGALRRWVVRNDGI  96

Query  335  YFARKYNTPLVRVLSWNKN  279
            Y  R  N       +WNK
Sbjct  97   YLVRDRNPQPKFHHAWNKT  115


>unnamed protein product [Arabidopsis thaliana]
Sequence ID: VYS53238.1 Length: 143
Range 1: 1 to 128

Score:68.2 bits(165), Expect:1e-13,
Method:Compositional matrix adjust.,
Identities:45/130(35%), Positives:62/130(47%), Gaps:6/130(4%)

Query  686  MNHSVFVILITITYFGLNQACI--KNIVEILNQLAPGQILEYHCRSEDDNLGVKQLNFNA  513
            MN   F  +I +   GL+ A    KN V   N L    +L+ HC S+DD+LG   L
Sbjct  1    MNCFSFSFIIIVLCAGLSNAKFREKNSVVFKNSLGVKNVLKIHCTSKDDDLGYHYLRPGV  60

Query  512  TPFVIRFHDEIPNLTRWNCIFRQGPNNSYSY--DIEVYKAGPRLIPRCGQLRVWAARIDG  339
              +  RFHD +   T+++C   QG   +Y +  +   YK G  LI   G+  +W AR DG
Sbjct  61   QIYEFRFHDSVLK-TKFDCELWQGRGPTYKFYANFRAYKGGG-LIAHYGKKNIWEAREDG  118

Query  338  IYFARKYNTP  309
            IYF      P
Sbjct  119  IYFTHGKEIP  128


>Plant self-incompatibility protein S1 family [Arabidopsis thaliana]
Sequence ID: NP_001118369.1 Length: 143
>RecName: Full=S-protein homolog 11; Flags: Precursor [Arabidopsis thaliana]
Sequence ID: B3H4B5.1 Length: 143 >Plant self-incompatibility protein S1 family [Arabidopsis thaliana]
Sequence ID: AEC07420.1 Length: 143
Range 1: 1 to 128

Score:65.9 bits(159), Expect:8e-13,
Method:Compositional matrix adjust.,
Identities:44/130(34%), Positives:62/130(47%), Gaps:6/130(4%)

Query  686  MNHSVFVILITITYFGLNQACI--KNIVEILNQLAPGQILEYHCRSEDDNLGVKQLNFNA  513
            MN   F  +I +   G + A    KN V   N L    +L+ HC S+DD+LG   L
Sbjct  1    MNCFSFSFIIIVLCAGSSNAKFREKNSVVFKNSLGVKNVLKIHCTSKDDDLGYHYLRPGV  60

Query  512  TPFVIRFHDEIPNLTRWNCIFRQGPNNSYSY--DIEVYKAGPRLIPRCGQLRVWAARIDG  339
              +  RFHD +   T+++C   QG   +Y +  +   YK+G  LI   G+  +W AR DG
Sbjct  61   QIYEFRFHDSVLK-TKFDCELWQGRGPTYKFYANFRAYKSGG-LIAHYGKKNIWEAREDG  118

Query  338  IYFARKYNTP  309
            IYF      P
Sbjct  119  IYFTHGKEIP  128


>Plant self-incompatibility protein S1 family [Arabidopsis thaliana]
Sequence ID: NP_178663.1 Length: 135
>putative S1 self-incompatibility protein [Arabidopsis thaliana]
Sequence ID: AAD19763.1 Length: 135 >Plant self-incompatibility protein S1 family [Arabidopsis thaliana]
Sequence ID: AEC06001.1 Length: 135
Range 1: 3 to 135

Score:63.2 bits(152), Expect:8e-12,
Method:Compositional matrix adjust.,
Identities:38/135(28%), Positives:63/135(46%), Gaps:7/135(5%)

Query  677  SVFVILITITY-FGLNQACI---KNIVEILNQLAPG-QILEYHCRSEDDNLGVKQLNFNA  513
            ++FV+LI I    G N       KN +   N  +    +L  HC+S+DD+LG+  +   +
Sbjct  3    NLFVLLIIIALSVGSNNGSKLWPKNQLHFRNSFSRNYDVLTVHCKSKDDDLGIHTVA-RS  61

Query  512  TPFVIRFHDEIPNLTRWNCIFRQGPNNSYSYDIEVYKAGPRLIPRCGQLRVWAARIDGIY  333
              +  +F D +   T + C    G  + YS     YKA P  +   G +++W A  DGIY
Sbjct  62   YEYNFKFEDSVFGRTEFFCTLMHGVGSKYSVTFTAYKAKPAFVASTGVIKIWDALDDGIY  121

Query  332  FARKYNTPLVRVLSW  288
               + +   V++  W
Sbjct  122  LTDE-DHDFVKIYGW  135


>unnamed protein product [Arabidopsis thaliana]
Sequence ID: VYS52148.1 Length: 135
Range 1: 3 to 135

Score:63.2 bits(152), Expect:9e-12,
Method:Compositional matrix adjust.,
Identities:38/135(28%), Positives:63/135(46%), Gaps:7/135(5%)

Query  677  SVFVILITITY-FGLNQACI---KNIVEILNQLAPG-QILEYHCRSEDDNLGVKQLNFNA  513
            ++FV+LI I    G N       KN +   N  +    +L  HC+S+DD+LG+  +   +
Sbjct  3    NLFVLLIIIALSVGSNNGSKLWPKNQIHFRNSFSRNYDVLTVHCKSKDDDLGIHTVA-RS  61

Query  512  TPFVIRFHDEIPNLTRWNCIFRQGPNNSYSYDIEVYKAGPRLIPRCGQLRVWAARIDGIY  333
              +  +F D +   T + C    G  + YS     YKA P  +   G +++W A  DGIY
Sbjct  62   YEYNFKFDDSVFGRTEFFCTLMHGVGSKYSVTFTAYKAKPAFVASTGVIKIWDALDDGIY  121

Query  332  FARKYNTPLVRVLSW  288
               + +   V++  W
Sbjct  122  LTDE-DHDFVKMYGW  135


>hypothetical protein AXX17_AT2G05160 [Arabidopsis thaliana]
Sequence ID: OAP10771.1 Length: 135
Range 1: 3 to 135

Score:62.8 bits(151), Expect:1e-11,
Method:Compositional matrix adjust.,
Identities:38/135(28%), Positives:63/135(46%), Gaps:7/135(5%)

Query  677  SVFVILITITY-FGLNQACI---KNIVEILNQLAPG-QILEYHCRSEDDNLGVKQLNFNA  513
            ++FV+LI I    G N       KN +   N  +    +L  HC+S+DD+LG+  +   +
Sbjct  3    NLFVLLIIIALSVGSNNGSKLWPKNQLHFRNSFSRNYDVLTVHCKSKDDDLGIHTVA-RS  61

Query  512  TPFVIRFHDEIPNLTRWNCIFRQGPNNSYSYDIEVYKAGPRLIPRCGQLRVWAARIDGIY  333
              +  +F D +   T + C    G  + YS     YKA P  +   G +++W A  DGIY
Sbjct  62   YEYNFKFEDSVFGRTEFFCTLMHGVGSKYSVTFTAYKAKPAFVASTGVIKIWDALDDGIY  121

Query  332  FARKYNTPLVRVLSW  288
               + +   V++  W
Sbjct  122  LTDE-DHDFVKMYGW  135


>hypothetical protein AXX17_AT2G18790 [Arabidopsis thaliana]
Sequence ID: OAP07766.1 Length: 147
Range 1: 1 to 128

Score:60.5 bits(145), Expect:1e-10,
Method:Compositional matrix adjust.,
Identities:45/132(34%), Positives:62/132(46%), Gaps:10/132(7%)

Query  686  MNHSVFVILITITYFGLNQACI--KNIVEILNQLAPGQILEYHCRSEDDNLGVKQLNFNA  513
            MN   +  L+ I   GLN A    KN +   + L P ++L  HC SE D+     L
Sbjct  1    MNCFSYFFLVIILCAGLNNAKFNEKNFIIFKSSLGPKKLLRIHCTSEHDDTDYVYLRHGQ  60

Query  512  TPFVIRFHDEIPNLTRWNCIFRQGPNNSYSYDI----EVYKAGPRLIPRCGQLRVWAARI  345
            T +   FHD +   T ++C  +QG ++ Y+Y+       YK G  LI   G+   W AR
Sbjct  61   T-YAFSFHDSVLK-TIFDCELKQG-SSYYNYNFYARFRAYKGGG-LIVHYGKKNFWDARE  116

Query  344  DGIYFARKYNTP  309
            DGIYF     TP
Sbjct  117  DGIYFTHGKETP  128


>Plant self-incompatibility protein S1 family [Arabidopsis thaliana]
Sequence ID: NP_001118368.1 Length: 147
>RecName: Full=S-protein homolog 10; Flags: Precursor [Arabidopsis thaliana]
Sequence ID: B3H6H8.1 Length: 147 >Plant self-incompatibility protein S1 family [Arabidopsis thaliana]
Sequence ID: AEC07419.1 Length: 147
Range 1: 1 to 128

Score:60.1 bits(144), Expect:1e-10,
Method:Compositional matrix adjust.,
Identities:46/132(35%), Positives:62/132(46%), Gaps:10/132(7%)

Query  686  MNHSVFVILITITYFGLNQACI--KNIVEILNQLAPGQILEYHCRSEDDNLGVKQLNFNA  513
            MN   +  L+ I   GLN A    KN V   + L P ++L  HC SE D+     L
Sbjct  1    MNCFSYFFLVIILCAGLNNAKFNEKNSVIFKSSLGPKKLLRIHCTSEHDDTDYVYLRHGQ  60

Query  512  TPFVIRFHDEIPNLTRWNCIFRQGPNNSYSYDI----EVYKAGPRLIPRCGQLRVWAARI  345
            T +   FHD +   T ++C  +QG ++ Y+Y+       YK G  LI   G+   W AR
Sbjct  61   T-YAFSFHDSVLK-TIFDCELKQG-SSYYNYNFYARFRAYKGGG-LIVHYGKKNFWDARE  116

Query  344  DGIYFARKYNTP  309
            DGIYF     TP
Sbjct  117  DGIYFTHGKETP  128


>unnamed protein product [Arabidopsis thaliana]
Sequence ID: VYS53237.1 Length: 147
Range 1: 1 to 128

Score:59.7 bits(143), Expect:2e-10,
Method:Compositional matrix adjust.,
Identities:45/132(34%), Positives:62/132(46%), Gaps:10/132(7%)

Query  686  MNHSVFVILITITYFGLNQACI--KNIVEILNQLAPGQILEYHCRSEDDNLGVKQLNFNA  513
            MN   +  L+ I   GLN A    KN V   + L P ++L  HC SE D+     L
Sbjct  1    MNCFSYFFLVIILCAGLNNAKFNEKNSVIFKSSLGPKKLLRIHCTSEHDDTDYVYLRHGQ  60

Query  512  TPFVIRFHDEIPNLTRWNCIFRQGPNNSYSYDI----EVYKAGPRLIPRCGQLRVWAARI  345
            T +   FHD +   T ++C  +QG ++ Y+Y+       YK G  L+   G+   W AR
Sbjct  61   T-YAFSFHDSVLK-TIFDCELKQG-SSYYNYNFYARFRAYKGGG-LVVHYGKRNFWDARE  116

Query  344  DGIYFARKYNTP  309
            DGIYF     TP
Sbjct  117  DGIYFTHGKETP  128


>hypothetical protein AXX17_AT5G03760 [Arabidopsis thaliana]
Sequence ID: OAO94980.1 Length: 136
>unnamed protein product [Arabidopsis thaliana]
Sequence ID: CAA0400595.1 Length: 136
Range 1: 29 to 133

Score:57.8 bits(138), Expect:6e-10,
Method:Compositional matrix adjust.,
Identities:37/111(33%), Positives:52/111(46%), Gaps:8/111(7%)

Query  611  VEILNQLAPGQILEYHCRSEDDNLGVKQLNFNATPFVIRFHDEIPNLTRWNCIFRQGPNN  432
            V + NQL   ++L+ HCRS+DD+LG   L      +   F D I   T ++C   QGPN
Sbjct  29   VVLSNQLEHSKLLKVHCRSKDDDLGEHILKI-GQDYEFTFGDNIWQTTSFSCQMDQGPNF  87

Query  431  SYSYDIEVYKA--GPRLIPRCGQLRVWAARIDGIYFARKYNTPLVRVLSWN  285
             +  D   Y+      L   C     W AR DGIYF++    P  +   W+
Sbjct  88   KHHQDFVAYETSWSKALEASCK----WIAREDGIYFSQD-GVPPTKKYQWD  133


>unnamed protein product [Arabidopsis thaliana]
Sequence ID: VYS65835.1 Length: 136
Range 1: 29 to 133

Score:57.8 bits(138), Expect:6e-10,
Method:Compositional matrix adjust.,
Identities:37/111(33%), Positives:52/111(46%), Gaps:8/111(7%)

Query  611  VEILNQLAPGQILEYHCRSEDDNLGVKQLNFNATPFVIRFHDEIPNLTRWNCIFRQGPNN  432
            V + NQL   ++L+ HCRS+DD+LG   L      +   F D I   T ++C   QGPN
Sbjct  29   VVLSNQLEHSKLLKVHCRSKDDDLGEHILKI-GQDYEFTFGDNIWQTTSFSCQMDQGPNF  87

Query  431  SYSYDIEVYKA--GPRLIPRCGQLRVWAARIDGIYFARKYNTPLVRVLSWN  285
             +  D   Y+      L   C     W AR DGIYF++    P  +   W+
Sbjct  88   KHHQDFVAYETSWSKALEASCK----WIAREDGIYFSQD-GVPPTKKYQWD  133


>unnamed protein product [Arabidopsis thaliana]
Sequence ID: CAA0370215.1 Length: 143
Range 1: 1 to 128

Score:58.2 bits(139), Expect:7e-10,
Method:Compositional matrix adjust.,
Identities:42/130(32%), Positives:59/130(45%), Gaps:6/130(4%)

Query  686  MNHSVFVILITITYFGLNQACI--KNIVEILNQLAPGQILEYHCRSEDDNLGVKQLNFNA  513
            MN   F  +I +    L+ A    KN V   N L    +L+ H  S+DD+LG   L
Sbjct  1    MNCFSFSFIIIVLCARLSNAKFREKNSVVFKNSLGVKNVLKIHFTSKDDDLGYHYLRPGV  60

Query  512  TPFVIRFHDEIPNLTRWNCIFRQGPNNSYSY--DIEVYKAGPRLIPRCGQLRVWAARIDG  339
              +  RFHD +   T+++C   QG   +Y +  +   YK G  LI   G+  +W AR D
Sbjct  61   QIYEFRFHDSVLK-TKFDCELWQGRGPTYKFYANFRAYKGGG-LIAHYGKKNIWEAREDD  118

Query  338  IYFARKYNTP  309
            IYF      P
Sbjct  119  IYFTHGKEIP  128


>hypothetical protein AXX17_AT5G35760 [Arabidopsis thaliana]
Sequence ID: OAO93026.1 Length: 133
Range 1: 5 to 126

Score:57.0 bits(136), Expect:1e-09,
Method:Compositional matrix adjust.,
Identities:45/127(35%), Positives:60/127(47%), Gaps:9/127(7%)

Query  677  SVFVILITITYFGLNQACIK----NIVEILNQLAPGQILEYHCRSEDDNLGVKQLNFNAT  510
            S F+++I +   GL+ A +K    N V   + L    +L+ HC SED NLG   L    T
Sbjct  5    SCFLLVIGLC-IGLSNANLKWNEKNTVFFKSSLGRNNVLKIHCTSED-NLGFHFLRPGET  62

Query  509  PFVIRFHDEIPNLTRWNCIFRQGPNNSYSYDIEVYKAGPRLIPRCGQLRVWAARIDGIYF  330
             +   FHD I   + + C   QGPN  +      Y+ G  LI   G+   W AR DGIYF
Sbjct  63   -YDFSFHDSIVR-SDFYCELWQGPNFKFHASFMAYQGGG-LIVHYGKKNFWDAREDGIYF  119

Query  329  ARKYNTP  309
                 TP
Sbjct  120  THGKETP  126


>Plant self-incompatibility protein S1 family [Arabidopsis thaliana]
Sequence ID: NP_198660.1 Length: 133
>RecName: Full=S-protein homolog 9; Flags: Precursor [Arabidopsis thaliana]
Sequence ID: Q9FF19.1 Length: 133 >Plant self-incompatibility protein S1 family [Arabidopsis thaliana]
Sequence ID: AED94317.1 Length: 133 >unnamed protein product [Arabidopsis thaliana]
Sequence ID: BAB09356.1 Length: 133
Range 1: 5 to 126

Score:57.0 bits(136), Expect:1e-09,
Method:Compositional matrix adjust.,
Identities:45/127(35%), Positives:60/127(47%), Gaps:9/127(7%)

Query  677  SVFVILITITYFGLNQACI----KNIVEILNQLAPGQILEYHCRSEDDNLGVKQLNFNAT  510
            S F+++I +   GL+ A +    KN V   + L    +L+ HC SED NLG   L    T
Sbjct  5    SCFLLVIGLC-IGLSNANLIWNEKNTVFFKSSLGRNNVLKIHCTSED-NLGFHFLRPGET  62

Query  509  PFVIRFHDEIPNLTRWNCIFRQGPNNSYSYDIEVYKAGPRLIPRCGQLRVWAARIDGIYF  330
             +   FHD I   + + C   QGPN  +      Y+ G  LI   G+   W AR DGIYF
Sbjct  63   -YDFSFHDSIVR-SDFYCELWQGPNFKFHASFMAYQGGG-LIVHYGKKNFWDAREDGIYF  119

Query  329  ARKYNTP  309
                 TP
Sbjct  120  THGKETP  126


>Plant self-incompatibility protein S1 family [Arabidopsis thaliana]
Sequence ID: NP_196055.1 Length: 146
>Plant self-incompatibility protein S1 family [Arabidopsis thaliana]
Sequence ID: AED90731.1 Length: 146
Range 1: 29 to 121

Score:55.1 bits(131), Expect:9e-09,
Method:Compositional matrix adjust.,
Identities:34/98(35%), Positives:47/98(47%), Gaps:7/98(7%)

Query  611  VEILNQLAPGQILEYHCRSEDDNLGVKQLNFNATPFVIRFHDEIPNLTRWNCIFRQGPNN  432
            V + NQL   ++L+ HCRS+DD+LG   L      +   F D I   T ++C   QGPN
Sbjct  29   VVLSNQLEHSKLLKVHCRSKDDDLGEHILKI-GQDYEFTFGDNIWQTTSFSCQMDQGPNF  87

Query  431  SYSYDIEVYKA--GPRLIPRCGQLRVWAARIDGIYFAR  324
             +  D   Y+      L   C     W  R DGIYF++
Sbjct  88   KHHLDFVAYETSWSKALEASCK----WIGREDGIYFSQ  121


>unnamed protein product [Arabidopsis thaliana]
Sequence ID: VYS68614.1 Length: 133
Range 1: 5 to 126

Score:53.9 bits(128), Expect:1e-08,
Method:Compositional matrix adjust.,
Identities:44/127(35%), Positives:59/127(46%), Gaps:9/127(7%)

Query  677  SVFVILITITYFGLNQACIK----NIVEILNQLAPGQILEYHCRSEDDNLGVKQLNFNAT  510
            S F+++I +   GL+ A +K    N V   + L    +L+ HC SED NLG   L    T
Sbjct  5    SCFLLVIGLC-IGLSNANLKWNEKNTVFFKSSLGRNNVLKIHCTSED-NLGFHFLRPGET  62

Query  509  PFVIRFHDEIPNLTRWNCIFRQGPNNSYSYDIEVYKAGPRLIPRCGQLRVWAARIDGIYF  330
             +   FHD I   + + C   Q PN  +      Y+ G  LI   G+   W AR DGIYF
Sbjct  63   -YDFSFHDSIVR-SDFYCELWQWPNFKFHASFMAYQGGG-LIVHYGKKNFWDAREDGIYF  119

Query  329  ARKYNTP  309
                 TP
Sbjct  120  THGKETP  126


Query #265: XLOC_030815 Query ID: lcl|Query_30756 Length: 772

No significant similarity found.


Query #266: XLOC_030816 Query ID: lcl|Query_30757 Length: 641

Sequences producing significant alignments:
                                                                  Max    Total Query   E   Per.
Description                                                       Score  Score cover Value Ident  Accession
hypothetical protein AXX17_AT5G11300 [Arabidopsis thaliana]       61.6   61.6  44%   2e-10 36.46  OAO93304.1


Alignments:
>hypothetical protein AXX17_AT5G11300 [Arabidopsis thaliana]
Sequence ID: OAO93304.1 Length: 696
Range 1: 567 to 647

Score:61.6 bits(148), Expect:2e-10,
Method:Composition-based stats.,
Identities:35/96(36%), Positives:51/96(53%), Gaps:15/96(15%)

Query  322  GITTGISCFLCNQQSETRDHLSIYCDYNIQIWHQILRRLGQPGYKRNITDWSCLISWLSS  501
            G+   + C  CN   E+RDHL ++C++  +IWH ++RRLGQ        DWS LISWL S
Sbjct  567  GLLISLLCGFCNTDVESRDHLLLHCNFVEEIWHSMMRRLGQSPCI--FADWSSLISWLLS  624

Query  502  PR*SACHTDNNSKRQQQSD*DATWMIVQNTVVLFYK  609
               +  +  +N KR          + VQ T+ L +K
Sbjct  625  ---TTPNLSSNLKR----------IAVQATIYLLWK  647


Query #267: XLOC_030817 Query ID: lcl|Query_30758 Length: 1810

Sequences producing significant alignments:
                                                                  Max    Total Query   E   Per.
Description                                                       Score  Score cover Value Ident  Accession
hypothetical protein AXX17_AT4G08610 [Arabidopsis thaliana]       67.8   67.8  8%    3e-11 61.11  OAO97959.1
contains similarity to reverse transcriptase (Pfam: rvt.hmm,...   66.6   66.6  8%    1e-10 59.26  AAC26250.1


Alignments:
>hypothetical protein AXX17_AT4G08610 [Arabidopsis thaliana]
Sequence ID: OAO97959.1 Length: 568
Range 1: 294 to 347

Score:67.8 bits(164), Expect:3e-11,
Method:Compositional matrix adjust.,
Identities:33/54(61%), Positives:42/54(77%), Gaps:5/54(9%)

Query  2    KEFDFIRNEKE-----SSNGSVVTYLVLYIDNILLLGNNIPFLQSLEI*LESCL  148
            KEFDFIRNE+E      ++GS V +LVLY+D+ILLLGN+IP LQS++  LESC
Sbjct  294  KEFDFIRNEEEPCVYKKTSGSAVAFLVLYVDDILLLGNDIPLLQSVKTWLESCF  347


>contains similarity to reverse transcriptase (Pfam: rvt.hmm, score 19.29) [Arabidopsis thaliana]
Sequence ID: AAC26250.1 Length: 964
>putative retrotransposon protein [Arabidopsis thaliana]
Sequence ID: CAB80804.1 Length: 964
Range 1: 610 to 663

Score:66.6 bits(161), Expect:1e-10,
Method:Composition-based stats.,
Identities:32/54(59%), Positives:41/54(75%), Gaps:5/54(9%)

Query  2    KEFDFIRNEKE-----SSNGSVVTYLVLYIDNILLLGNNIPFLQSLEI*LESCL  148
            KEFDFIRNE+E      ++GS V +LVLY+D+ILLLGN+IP LQS++  L SC
Sbjct  610  KEFDFIRNEEEPCVYKKTSGSAVAFLVLYVDDILLLGNDIPLLQSVKTWLGSCF  663


Query #268: XLOC_031248 Query ID: lcl|Query_30759 Length: 818

Sequences producing significant alignments:
                                                                  Max    Total Query   E   Per.
Description                                                       Score  Score cover Value Ident  Accession
transmembrane protein [Arabidopsis thaliana]                      106    106   42%   1e-27 50.85  NP_001154549.2
unnamed protein product [Arabidopsis thaliana]                    105    105   42%   2e-27 50.00  CAA0374292.1


Alignments:
>transmembrane protein [Arabidopsis thaliana]
Sequence ID: NP_001154549.2 Length: 187
>transmembrane protein [Arabidopsis thaliana]
Sequence ID: AEC08794.2 Length: 187
Range 1: 52 to 161

Score:106 bits(265), Expect:1e-27,
Method:Compositional matrix adjust.,
Identities:60/118(51%), Positives:75/118(63%), Gaps:11/118(9%)

Query  164  WVLVSLQLVQTDLFWR---KWKSPWKKRQFGDDWIMTISNYLWKRMMRRFFVSCKQQWVL  334
            W   SL  + +DL        K PW  RQ+ ++ +         R +RR+F+  +Q+W L
Sbjct  52   WAFRSLTQILSDLICHHRPARKPPW--RQYVEEELEA------NRRIRRWFIPYQQKWGL  103

Query  335  NTTNTNDILFYMRVSINPKVLRCNRWIIAYDFQLHYRYFISKKIMIPKQESHIRLTKM  508
              T  NDI FYMRVSIN KVLRCNRWI AYDF++ YRYFI KK MIPKQESH +LT+M
Sbjct  104  RKTKINDISFYMRVSINSKVLRCNRWIAAYDFRMRYRYFIYKKKMIPKQESHTKLTEM  161


>unnamed protein product [Arabidopsis thaliana]
Sequence ID: CAA0374292.1 Length: 183
Range 1: 52 to 161

Score:105 bits(263), Expect:2e-27,
Method:Compositional matrix adjust.,
Identities:59/118(50%), Positives:75/118(63%), Gaps:11/118(9%)

Query  164  WVLVSLQLVQTDLFWR---KWKSPWKKRQFGDDWIMTISNYLWKRMMRRFFVSCKQQWVL  334
            W   SL  + +D+        K PW  RQ+ ++ +         R +RR+F+  +Q+W L
Sbjct  52   WAFRSLTQILSDMICHHRPARKPPW--RQYVEEELEA------NRRIRRWFIPYQQKWGL  103

Query  335  NTTNTNDILFYMRVSINPKVLRCNRWIIAYDFQLHYRYFISKKIMIPKQESHIRLTKM  508
              T  NDI FYMRVSIN KVLRCNRWI AYDF++ YRYFI KK MIPKQESH +LT+M
Sbjct  104  RKTKINDISFYMRVSINSKVLRCNRWIAAYDFRMRYRYFIYKKKMIPKQESHTKLTEM  161


Query #269: XLOC_031407 Query ID: lcl|Query_30760 Length: 881

No significant similarity found.


Query #270: XLOC_031571 Query ID: lcl|Query_30761 Length: 1056

Sequences producing significant alignments:
                                                                  Max    Total Query   E   Per.
Description                                                       Score  Score cover Value Ident  Accession
hypothetical protein AXX17_AT1G35380 [Arabidopsis thaliana]       69.7   69.7  9%    7e-15 97.14  OAP13075.1
transmembrane protein [Arabidopsis thaliana]                      58.2   58.2  7%    1e-09 92.59  NP_174704.1


Alignments:
>hypothetical protein AXX17_AT1G35380 [Arabidopsis thaliana]
Sequence ID: OAP13075.1 Length: 53
Range 1: 19 to 53

Score:69.7 bits(169), Expect:7e-15,
Method:Compositional matrix adjust.,
Identities:34/35(97%), Positives:34/35(97%), Gaps:0/35(0%)

Query  544  DSRPPTPFVTIEGAHLKGHNQSTIVTGASPASITL  648
            DSRPPTPFVTIEGAHLKG NQSTIVTGASPASITL
Sbjct  19   DSRPPTPFVTIEGAHLKGQNQSTIVTGASPASITL  53


>transmembrane protein [Arabidopsis thaliana]
Sequence ID: NP_174704.1 Length: 157
>hypothetical protein; 16456-17845 [Arabidopsis thaliana]
Sequence ID: AAG51906.1 Length: 157 >transmembrane protein [Arabidopsis thaliana]
Sequence ID: AEE31714.1 Length: 157
Range 1: 131 to 157

Score:58.2 bits(139), Expect:1e-09,
Method:Compositional matrix adjust.,
Identities:25/27(93%), Positives:26/27(96%), Gaps:0/27(0%)

Query  668  ADQLIFYNVIEAGLAPVTIVDWLCPLR  588
            +DQLIFYNVIEAGLAPVTIVDW CPLR
Sbjct  131  SDQLIFYNVIEAGLAPVTIVDWFCPLR  157


Query #271: XLOC_031829 Query ID: lcl|Query_30762 Length: 676

No significant similarity found.


Query #272: XLOC_032243 Query ID: lcl|Query_30763 Length: 1160

Sequences producing significant alignments:
                                                                  Max    Total Query   E   Per.
Description                                                       Score  Score cover Value Ident  Accession
hypothetical protein AT5G63625 [Arabidopsis thaliana]             100    178   21%   6e-25 100.00 NP_001331694.1
unnamed protein product [Arabidopsis thaliana]                    80.5   80.5  10%   9e-19 100.00 CAA0411774.1
alternative oxidase [Arabidopsis thaliana]                        84.3   84.3  21%   2e-17 54.00  CAA10364.1
hypothetical protein AXX17_AT5G64340 [Arabidopsis thaliana]       73.6   73.6  9%    4e-16 92.11  OAO90914.1
sucrose synthase 3 [Arabidopsis thaliana]                         79.0   79.0  9%    4e-15 97.37  NP_192137.1
T2H3.8 [Arabidopsis thaliana]                                     79.0   79.0  9%    4e-15 97.37  AAC28175.1
ubiquinol oxidase [Arabidopsis thaliana]                          70.5   70.5  9%    6e-15 89.47  NP_001330488.1
ubiquinol oxidase [Arabidopsis thaliana]                          70.5   70.5  9%    6e-15 89.47  NP_001330489.1
alternative oxidase 1A [Arabidopsis thaliana]                     76.6   76.6  25%   9e-15 46.67  NP_188876.1
HSR3 [Arabidopsis thaliana]                                       75.5   75.5  23%   3e-14 48.65  OAP01931.1
oxidase [Arabidopsis thaliana]                                    70.9   70.9  11%   5e-13 71.70  AAA32870.1
F28K19.26 [Arabidopsis thaliana]                                  48.5   90.9  12%   9e-12 82.76  AAF17689.1
sucrose synthase 2 [Arabidopsis thaliana]                         68.2   68.2  9%    1e-11 78.95  NP_199730.1
sucrose synthase 2 [Arabidopsis thaliana]                         68.2   68.2  9%    1e-11 78.95  NP_001331822.1
unnamed protein product [Arabidopsis thaliana]                    68.2   68.2  9%    1e-11 78.95  CAA0408606.1
unnamed protein product [Arabidopsis thaliana]                    68.2   68.2  9%    1e-11 78.95  VYS69780.1
SUS2 [Arabidopsis thaliana]                                       68.2   68.2  9%    1e-11 78.95  OAO95896.1
sucrose synthase 2 [Arabidopsis thaliana]                         68.2   68.2  9%    1e-11 78.95  NP_001331823.1
sucrose synthase [Arabidopsis thaliana]                           68.2   68.2  9%    1e-11 78.95  BAB10337.1
sucrose synthase [Arabidopsis thaliana]                           64.3   64.3  9%    2e-10 73.68  CAA43303.1


Alignments:
>hypothetical protein AT5G63625 [Arabidopsis thaliana]
Sequence ID: NP_001331694.1 Length: 124
>hypothetical protein AT5G63625 [Arabidopsis thaliana]
Sequence ID: ANM70057.1 Length: 124 >unnamed protein product [Arabidopsis thaliana]
Sequence ID: VYS71299.1 Length: 124
Range 1: 80 to 124

Score:100 bits(248), Expect:6e-25,
Method:Compositional matrix adjust.,
Identities:45/45(100%), Positives:45/45(100%), Gaps:0/45(0%)

Query  874   EAPSNLCYGTLQKTSDDSKTELHYQSKTNEKDVARLHCFHSQFSP  1008
             EAPSNLCYGTLQKTSDDSKTELHYQSKTNEKDVARLHCFHSQFSP
Sbjct  80    EAPSNLCYGTLQKTSDDSKTELHYQSKTNEKDVARLHCFHSQFSP  124


Range 2: 1 to 37

Score:78.6 bits(192), Expect:4e-17,
Method:Compositional matrix adjust.,
Identities:37/37(100%), Positives:37/37(100%), Gaps:0/37(0%)

Query  738  MRFACTITLGEKASTKEEDANQRKTENESTGGDKGIA  848
            MRFACTITLGEKASTKEEDANQRKTENESTGGDKGIA
Sbjct  1    MRFACTITLGEKASTKEEDANQRKTENESTGGDKGIA  37


>unnamed protein product [Arabidopsis thaliana]
Sequence ID: CAA0411774.1 Length: 39
Range 1: 1 to 39

Score:80.5 bits(197), Expect:9e-19,
Method:Compositional matrix adjust.,
Identities:39/39(100%), Positives:39/39(100%), Gaps:0/39(0%)

Query  738  MRFACTITLGEKASTKEEDANQRKTENESTGGDKGIASY  854
            MRFACTITLGEKASTKEEDANQRKTENESTGGDKGIASY
Sbjct  1    MRFACTITLGEKASTKEEDANQRKTENESTGGDKGIASY  39


>alternative oxidase [Arabidopsis thaliana]
Sequence ID: CAA10364.1 Length: 353
Range 1: 10 to 108

Score:84.3 bits(207), Expect:2e-17,
Method:Compositional matrix adjust.,
Identities:54/100(54%), Positives:60/100(60%), Gaps:17/100(17%)

Query  609  AKAAVNGD*TTFVLNDPYDYVLRGVLSKPSLEATCCISAWI-------GGMRFACTITLG  767
            A+ AV+G  TTFVL+ PY       LS+  +      SAWI       GGMRFA TITLG
Sbjct  10   AQIAVSGGWTTFVLDGPY-VSSHEALSRSHILKPGVTSAWIWTRAPTIGGMRFASTITLG  68

Query  768  EKASTKEEDANQRKTENESTG---------GDKGIASY*G  860
            EK   KEEDANQ+KTENESTG         GDKGIASY G
Sbjct  69   EKTPMKEEDANQKKTENESTGGDAAGGNNKGDKGIASYWG  108


>hypothetical protein AXX17_AT5G64340 [Arabidopsis thaliana]
Sequence ID: OAO90914.1 Length: 56
Range 1: 1 to 38

Score:73.6 bits(179), Expect:4e-16,
Method:Compositional matrix adjust.,
Identities:35/38(92%), Positives:37/38(97%), Gaps:0/38(0%)

Query  738  MRFACTITLGEKASTKEEDANQRKTENESTGGDKGIAS  851
            MRFAC ITLGEKASTKEEDANQRKTENESTGGD+GIA+
Sbjct  1    MRFACAITLGEKASTKEEDANQRKTENESTGGDQGIAN  38


>sucrose synthase 3 [Arabidopsis thaliana]
Sequence ID: NP_192137.1 Length: 809
>RecName: Full=Sucrose synthase 3; Short=AtSUS3; AltName: Full=Sucrose-UDP glucosyltransferase 3 [Arabidopsis thaliana]
Sequence ID: Q9M111.1 Length: 809 >putative sucrose synthetase [Arabidopsis thaliana]
Sequence ID: AAK93678.1 Length: 809 >AT4g02280/T2H3_8 [Arabidopsis thaliana]
Sequence ID: AAL09730.1 Length: 809 >putative sucrose synthetase [Arabidopsis thaliana]
Sequence ID: AAN13112.1 Length: 809 >sucrose synthase 3 [Arabidopsis thaliana]
Sequence ID: AEE82150.1 Length: 809 >glycosyltransferase, partial [Arabidopsis thaliana]
Sequence ID: AHL38709.1 Length: 809
Range 1: 143 to 180

Score:79.0 bits(193), Expect:4e-15,
Method:Compositional matrix adjust.,
Identities:37/38(97%), Positives:37/38(97%), Gaps:0/38(0%)

Query  985  NANVLRPSRSSSIGNGVQFLNRHLSSVMFRNKDCLEPL  872
            NANV RPSRSSSIGNGVQFLNRHLSSVMFRNKDCLEPL
Sbjct  143  NANVPRPSRSSSIGNGVQFLNRHLSSVMFRNKDCLEPL  180


>T2H3.8 [Arabidopsis thaliana]
Sequence ID: AAC28175.1 Length: 808
Range 1: 143 to 180

Score:79.0 bits(193), Expect:4e-15,
Method:Compositional matrix adjust.,
Identities:37/38(97%), Positives:37/38(97%), Gaps:0/38(0%)

Query  985  NANVLRPSRSSSIGNGVQFLNRHLSSVMFRNKDCLEPL  872
            NANV RPSRSSSIGNGVQFLNRHLSSVMFRNKDCLEPL
Sbjct  143  NANVPRPSRSSSIGNGVQFLNRHLSSVMFRNKDCLEPL  180


>ubiquinol oxidase [Arabidopsis thaliana]
Sequence ID: NP_001330488.1 Length: 56
>ubiquinol oxidase [Arabidopsis thaliana]
Sequence ID: ANM68766.1 Length: 56 >hypothetical protein AXX17_AT5G65020 [Arabidopsis thaliana]
Sequence ID: OAO95958.1 Length: 56 >unnamed protein product [Arabidopsis thaliana]
Sequence ID: CAA0412022.1 Length: 56 >unnamed protein product [Arabidopsis thaliana]
Sequence ID: CAA0412161.1 Length: 56 >unnamed protein product [Arabidopsis thaliana]
Sequence ID: VYS71408.1 Length: 56
Range 1: 1 to 38

Score:70.5 bits(171), Expect:6e-15,
Method:Compositional matrix adjust.,
Identities:34/38(89%), Positives:36/38(94%), Gaps:0/38(0%)

Query  738  MRFACTITLGEKASTKEEDANQRKTENESTGGDKGIAS  851
            MRFA TITLGEKASTKEEDANQRKTE ESTGGD+GIA+
Sbjct  1    MRFASTITLGEKASTKEEDANQRKTEKESTGGDQGIAN  38


>ubiquinol oxidase [Arabidopsis thaliana]
Sequence ID: NP_001330489.1 Length: 61
>ubiquinol oxidase [Arabidopsis thaliana]
Sequence ID: ANM68767.1 Length: 61
Range 1: 1 to 38

Score:70.5 bits(171), Expect:6e-15,
Method:Compositional matrix adjust.,
Identities:34/38(89%), Positives:36/38(94%), Gaps:0/38(0%)

Query  738  MRFACTITLGEKASTKEEDANQRKTENESTGGDKGIAS  851
            MRFA TITLGEKASTKEEDANQRKTE ESTGGD+GIA+
Sbjct  1    MRFASTITLGEKASTKEEDANQRKTEKESTGGDQGIAN  38


>alternative oxidase 1A [Arabidopsis thaliana]
Sequence ID: NP_188876.1 Length: 354
>RecName: Full=Ubiquinol oxidase 1a, mitochondrial; AltName: Full=Alternative oxidase 1a; Flags: Precursor [Arabidopsis thaliana]
Sequence ID: Q39219.2 Length: 354 >putative alternative oxidase 1a precursor [Arabidopsis thaliana]
Sequence ID: AAK43981.1 Length: 354 >putative alternative oxidase 1a precursor [Arabidopsis thaliana]
Sequence ID: AAL15234.1 Length: 354 >alternative oxidase 1A [Arabidopsis thaliana]
Sequence ID: AEE76627.1 Length: 354 >unnamed protein product [Arabidopsis thaliana]
Sequence ID: VYS58238.1 Length: 354 >alternative oxidase [Arabidopsis thaliana]
Sequence ID: BAA22625.1 Length: 354
Range 1: 3 to 118

Score:76.6 bits(187), Expect:9e-15,
Method:Compositional matrix adjust.,
Identities:56/120(47%), Positives:68/120(56%), Gaps:26/120(21%)

Query  594  MSRGGAKAAVNGD*TTFVLNDPYDY-VLRGVLSKPSLEATCCI-----SAWI-------G  734
            ++RGGAKAA     +  V   P  +  +R V S  +L A+  +     SAWI       G
Sbjct  3    ITRGGAKAAK----SLLVAAGPRLFSTVRTVSSHEALSASHILKPGVTSAWIWTRAPTIG  58

Query  735  GMRFACTITLGEKASTKEEDANQRKTENESTG---------GDKGIASY*GFTTFRGSKQ  887
            GMRFA TITLGEK   KEEDANQ+KTENESTG         GDKGIASY G    + +K+
Sbjct  59   GMRFASTITLGEKTPMKEEDANQKKTENESTGGDAAGGNNKGDKGIASYWGVEPNKITKE  118


>HSR3 [Arabidopsis thaliana]
Sequence ID: OAP01931.1 Length: 354
>unnamed protein product [Arabidopsis thaliana]
Sequence ID: CAA0383312.1 Length: 354
Range 1: 3 to 109

Score:75.5 bits(184), Expect:3e-14,
Method:Compositional matrix adjust.,
Identities:54/111(49%), Positives:64/111(57%), Gaps:26/111(23%)

Query  594  MSRGGAKAAVNGD*TTFVLNDPYDY-VLRGVLSKPSLEATCCI-----SAWI-------G  734
            ++RGGAKAA     +  +   P  +  +R V S  +L A+  +     SAWI       G
Sbjct  3    ITRGGAKAAK----SLLMAAGPRLFSTVRTVSSHEALSASHILKPGVTSAWIWTRAPTIG  58

Query  735  GMRFACTITLGEKASTKEEDANQRKTENESTG---------GDKGIASY*G  860
            GMRFA TITLGEK   KEEDANQ+KTENESTG         GDKGIASY G
Sbjct  59   GMRFASTITLGEKTPMKEEDANQKKTENESTGGDAAGGNNKGDKGIASYWG  109


>oxidase [Arabidopsis thaliana]
Sequence ID: AAA32870.1 Length: 305
Range 1: 8 to 60

Score:70.9 bits(172), Expect:5e-13,
Method:Compositional matrix adjust.,
Identities:38/53(72%), Positives:39/53(73%), Gaps:9/53(16%)

Query  729  IGGMRFACTITLGEKASTKEEDANQRKTENESTG---------GDKGIASY*G  860
            IGGMRFA TITLGEK   KEEDANQ+KTENESTG         GDKGIASY G
Sbjct  8    IGGMRFASTITLGEKTPMKEEDANQKKTENESTGGDAAGGNNKGDKGIASYWG  60


>F28K19.26 [Arabidopsis thaliana]
Sequence ID: AAF17689.1 Length: 677
Range 1: 114 to 142

Score:48.5 bits(114), Expect:9e-12,
Method:Compositional matrix adjust.,
Identities:24/29(83%), Positives:26/29(89%), Gaps:0/29(0%)

Query  649  STIRTITSYEAFLASHLLKPHVAYRLGLE  735
            STIRTITS+ AF AS LLKP+VAYRLGLE
Sbjct  114  STIRTITSHVAFSASRLLKPYVAYRLGLE  142


Range 2: 142 to 162

Score:42.4 bits(98), Expect:9e-12,
Method:Compositional matrix adjust.,
Identities:19/21(90%), Positives:20/21(95%), Gaps:0/21(0%)

Query  789  EDANQRKTENESTGGDKGIAS  851
            EDANQRKTENESTGGDK IA+
Sbjct  142  EDANQRKTENESTGGDKEIAN  162


>sucrose synthase 2 [Arabidopsis thaliana]
Sequence ID: NP_199730.1 Length: 807
>RecName: Full=Sucrose synthase 2; Short=AtSUS2; AltName: Full=Sucrose-UDP glucosyltransferase 2 [Arabidopsis thaliana]
Sequence ID: Q00917.3 Length: 807 >sucrose synthase 2 [Arabidopsis thaliana]
Sequence ID: AED95780.1 Length: 807 >glycosyltransferase, partial [Arabidopsis thaliana]
Sequence ID: AHL38586.1 Length: 807
Range 1: 140 to 177

Score:68.2 bits(165), Expect:1e-11,
Method:Compositional matrix adjust.,
Identities:30/38(79%), Positives:35/38(92%), Gaps:0/38(0%)

Query  985  NANVLRPSRSSSIGNGVQFLNRHLSSVMFRNKDCLEPL  872
            NA + RP+RSSSIGNGVQFLNRHLSS+MFRNK+ +EPL
Sbjct  140  NATLPRPTRSSSIGNGVQFLNRHLSSIMFRNKESMEPL  177


>sucrose synthase 2 [Arabidopsis thaliana]
Sequence ID: NP_001331822.1 Length: 741
>sucrose synthase 2 [Arabidopsis thaliana]
Sequence ID: ANM70192.1 Length: 741
Range 1: 74 to 111

Score:68.2 bits(165), Expect:1e-11,
Method:Compositional matrix adjust.,
Identities:30/38(79%), Positives:35/38(92%), Gaps:0/38(0%)

Query  985  NANVLRPSRSSSIGNGVQFLNRHLSSVMFRNKDCLEPL  872
            NA + RP+RSSSIGNGVQFLNRHLSS+MFRNK+ +EPL
Sbjct  74   NATLPRPTRSSSIGNGVQFLNRHLSSIMFRNKESMEPL  111


>unnamed protein product [Arabidopsis thaliana]
Sequence ID: CAA0408606.1 Length: 807
Range 1: 140 to 177

Score:68.2 bits(165), Expect:1e-11,
Method:Compositional matrix adjust.,
Identities:30/38(79%), Positives:35/38(92%), Gaps:0/38(0%)

Query  985  NANVLRPSRSSSIGNGVQFLNRHLSSVMFRNKDCLEPL  872
            NA + RP+RSSSIGNGVQFLNRHLSS+MFRNK+ +EPL
Sbjct  140  NATLPRPTRSSSIGNGVQFLNRHLSSIMFRNKESMEPL  177


>unnamed protein product [Arabidopsis thaliana]
Sequence ID: VYS69780.1 Length: 807
Range 1: 140 to 177

Score:68.2 bits(165), Expect:1e-11,
Method:Compositional matrix adjust.,
Identities:30/38(79%), Positives:35/38(92%), Gaps:0/38(0%)

Query  985  NANVLRPSRSSSIGNGVQFLNRHLSSVMFRNKDCLEPL  872
            NA + RP+RSSSIGNGVQFLNRHLSS+MFRNK+ +EPL
Sbjct  140  NATLPRPTRSSSIGNGVQFLNRHLSSIMFRNKESMEPL  177


>SUS2 [Arabidopsis thaliana]
Sequence ID: OAO95896.1 Length: 807
Range 1: 140 to 177

Score:68.2 bits(165), Expect:1e-11,
Method:Compositional matrix adjust.,
Identities:30/38(79%), Positives:35/38(92%), Gaps:0/38(0%)

Query  985  NANVLRPSRSSSIGNGVQFLNRHLSSVMFRNKDCLEPL  872
            NA + RP+RSSSIGNGVQFLNRHLSS+MFRNK+ +EPL
Sbjct  140  NATLPRPTRSSSIGNGVQFLNRHLSSIMFRNKESMEPL  177


>sucrose synthase 2 [Arabidopsis thaliana]
Sequence ID: NP_001331823.1 Length: 797
>sucrose synthase 2 [Arabidopsis thaliana]
Sequence ID: ANM70193.1 Length: 797
Range 1: 130 to 167

Score:68.2 bits(165), Expect:1e-11,
Method:Compositional matrix adjust.,
Identities:30/38(79%), Positives:35/38(92%), Gaps:0/38(0%)

Query  985  NANVLRPSRSSSIGNGVQFLNRHLSSVMFRNKDCLEPL  872
            NA + RP+RSSSIGNGVQFLNRHLSS+MFRNK+ +EPL
Sbjct  130  NATLPRPTRSSSIGNGVQFLNRHLSSIMFRNKESMEPL  167


>sucrose synthase [Arabidopsis thaliana]
Sequence ID: BAB10337.1 Length: 805
Range 1: 138 to 175

Score:68.2 bits(165), Expect:1e-11,
Method:Compositional matrix adjust.,
Identities:30/38(79%), Positives:35/38(92%), Gaps:0/38(0%)

Query  985  NANVLRPSRSSSIGNGVQFLNRHLSSVMFRNKDCLEPL  872
            NA + RP+RSSSIGNGVQFLNRHLSS+MFRNK+ +EPL
Sbjct  138  NATLPRPTRSSSIGNGVQFLNRHLSSIMFRNKESMEPL  175


>sucrose synthase [Arabidopsis thaliana]
Sequence ID: CAA43303.1 Length: 804
Range 1: 138 to 175

Score:64.3 bits(155), Expect:2e-10,
Method:Compositional matrix adjust.,
Identities:28/38(74%), Positives:34/38(89%), Gaps:0/38(0%)

Query  985  NANVLRPSRSSSIGNGVQFLNRHLSSVMFRNKDCLEPL  872
            NA + RP+RSSSIGNGVQ +NRHLSS+MFRNK+ +EPL
Sbjct  138  NATLPRPTRSSSIGNGVQLVNRHLSSIMFRNKESMEPL  175


Query #273: XLOC_032332 Query ID: lcl|Query_30764 Length: 454

Sequences producing significant alignments:
                                                                  Max    Total Query   E   Per.
Description                                                       Score  Score cover Value Ident  Accession
hypothetical protein AXX17_AT5G65150 [Arabidopsis thaliana]       92.4   92.4  62%   9e-25 61.05  OAO90605.1


Alignments:
>hypothetical protein AXX17_AT5G65150 [Arabidopsis thaliana]
Sequence ID: OAO90605.1 Length: 79
Range 1: 1 to 59

Score:92.4 bits(228), Expect:9e-25,
Method:Compositional matrix adjust.,
Identities:58/95(61%), Positives:58/95(61%), Gaps:36/95(37%)

Query  25   MAFWKPWTDPTISHPlllllllIVMGMRI*RSKDRDCH**CTNYRTEILFLGATIIIVGV  204
            MAFWKPWTDPTISHPLLLLLLLIV
Sbjct  1    MAFWKPWTDPTISHPLLLLLLLIV------------------------------------  24

Query  205  FGWAQGGLVFALDKVKKLDGLKEAEFFDHITSFCM  309
            FGWAQGG VFALDKVKKLDGLKEAEFFDHITSFCM
Sbjct  25   FGWAQGGRVFALDKVKKLDGLKEAEFFDHITSFCM  59


Query #274: XLOC_032470 Query ID: lcl|Query_30765 Length: 851

Sequences producing significant alignments:
                                                                  Max    Total Query   E   Per.
Description                                                       Score  Score cover Value Ident  Accession
hypothetical protein AXX17_AT5G03740 [Arabidopsis thaliana]       280    280   46%   4e-96 100.00 OAO93173.1
hypothetical protein AXX17_AT5G03760 [Arabidopsis thaliana]       68.9   68.9  45%   7e-14 32.59  OAO94980.1
unnamed protein product [Arabidopsis thaliana]                    66.2   66.2  44%   6e-13 32.31  VYS65835.1
Plant self-incompatibility protein S1 family [Arabidopsis...      63.5   63.5  45%   7e-12 31.58  NP_196055.1
hypothetical protein AXX17_AT5G03750 [Arabidopsis thaliana]       56.2   56.2  43%   2e-09 33.06  OAO95751.1
Plant self-incompatibility protein S1 family [Arabidopsis...      56.2   56.2  43%   2e-09 33.06  NP_001031830.1
hypothetical protein AXX17_AT2G05160 [Arabidopsis thaliana]       55.1   55.1  45%   7e-09 27.94  OAP10771.1
unnamed protein product [Arabidopsis thaliana]                    53.5   53.5  45%   2e-08 27.21  VYS52148.1
Plant self-incompatibility protein S1 family [Arabidopsis...      53.5   53.5  45%   2e-08 27.21  NP_178663.1
unnamed protein product [Arabidopsis thaliana]                    52.0   52.0  45%   8e-08 29.63  CAA0383810.1
Plant self-incompatibility protein S1 family [Arabidopsis...      52.4   52.4  42%   9e-08 32.23  NP_001330171.1
hypothetical protein AXX17_AT2G18790 [Arabidopsis thaliana]       52.4   52.4  39%   9e-08 33.05  OAP07766.1


Alignments:
>hypothetical protein AXX17_AT5G03740 [Arabidopsis thaliana]
Sequence ID: OAO93173.1 Length: 131
Range 1: 1 to 131

Score:280 bits(716), Expect:4e-96,
Method:Compositional matrix adjust.,
Identities:131/131(100%), Positives:131/131(100%), Gaps:0/131(0%)

Query  102  MKRLTIVLIVLVFYIGDTHGRNVLMIVNRLPKNATLKLHCYSGDDDFKTMYLRHNDPPQT  281
            MKRLTIVLIVLVFYIGDTHGRNVLMIVNRLPKNATLKLHCYSGDDDFKTMYLRHNDPPQT
Sbjct  1    MKRLTIVLIVLVFYIGDTHGRNVLMIVNRLPKNATLKLHCYSGDDDFKTMYLRHNDPPQT  60

Query  282  WRFKDAFFHETQFICNLHQGYHWAHHRSFIAYKSSMNTSQKNNANATWFAGEKGIYLSFN  461
            WRFKDAFFHETQFICNLHQGYHWAHHRSFIAYKSSMNTSQKNNANATWFAGEKGIYLSFN
Sbjct  61   WRFKDAFFHETQFICNLHQGYHWAHHRSFIAYKSSMNTSQKNNANATWFAGEKGIYLSFN  120

Query  462  QRTPEFMYIWM  494
            QRTPEFMYIWM
Sbjct  121  QRTPEFMYIWM  131


>hypothetical protein AXX17_AT5G03760 [Arabidopsis thaliana]
Sequence ID: OAO94980.1 Length: 136
>unnamed protein product [Arabidopsis thaliana]
Sequence ID: CAA0400595.1 Length: 136
Range 1: 1 to 132

Score:68.9 bits(167), Expect:7e-14,
Method:Compositional matrix adjust.,
Identities:44/135(33%), Positives:68/135(50%), Gaps:8/135(5%)

Query  102  MKRLTIVLIVLVFYIGDTH-----GRNVLMIVNRLPKNATLKLHCYSGDDDFKTMYLRHN  266
            M  L I  IV+   IG ++     G + +++ N+L  +  LK+HC S DDD     L+
Sbjct  1    MNNLFIFSIVIGLCIGVSNAFHEIGESKVVLSNQLEHSKLLKVHCRSKDDDLGEHILKIG  60

Query  267  DPPQTWRFKDAFFHETQFICNLHQGYHWAHHRSFIAYKSSMNTSQKNNANATWFAGEKGI  446
               + + F D  +  T F C + QG ++ HH+ F+AY++S   S+   A+  W A E GI
Sbjct  61   QDYE-FTFGDNIWQTTSFSCQMDQGPNFKHHQDFVAYETSW--SKALEASCKWIAREDGI  117

Query  447  YLSFNQRTPEFMYIW  491
            Y S +   P   Y W
Sbjct  118  YFSQDGVPPTKKYQW  132


>unnamed protein product [Arabidopsis thaliana]
Sequence ID: VYS65835.1 Length: 136
Range 1: 6 to 132

Score:66.2 bits(160), Expect:6e-13,
Method:Compositional matrix adjust.,
Identities:42/130(32%), Positives:66/130(50%), Gaps:8/130(6%)

Query  117  IVLIVLVFYIGDTH-----GRNVLMIVNRLPKNATLKLHCYSGDDDFKTMYLRHNDPPQT  281
            I  IV+   IG ++     G + +++ N+L  +  LK+HC S DDD     L+     +
Sbjct  6    IFSIVIGLCIGVSNAFHEIGESKVVLSNQLEHSKLLKVHCRSKDDDLGEHILKIGQDYE-  64

Query  282  WRFKDAFFHETQFICNLHQGYHWAHHRSFIAYKSSMNTSQKNNANATWFAGEKGIYLSFN  461
            + F D  +  T F C + QG ++ HH+ F+AY++S   S+   A+  W A E GIY S +
Sbjct  65   FTFGDNIWQTTSFSCQMDQGPNFKHHQDFVAYETSW--SKALEASCKWIAREDGIYFSQD  122

Query  462  QRTPEFMYIW  491
               P   Y W
Sbjct  123  GVPPTKKYQW  132


>Plant self-incompatibility protein S1 family [Arabidopsis thaliana]
Sequence ID: NP_196055.1 Length: 146
>Plant self-incompatibility protein S1 family [Arabidopsis thaliana]
Sequence ID: AED90731.1 Length: 146
Range 1: 1 to 130

Score:63.5 bits(153), Expect:7e-12,
Method:Compositional matrix adjust.,
Identities:42/133(32%), Positives:65/133(48%), Gaps:8/133(6%)

Query  102  MKRLTIVLIVLVFYIGDTH-----GRNVLMIVNRLPKNATLKLHCYSGDDDFKTMYLRHN  266
            M  L I  IV+   IG ++     G + +++ N+L  +  LK+HC S DDD     L+
Sbjct  1    MNNLFIFSIVIGLCIGVSNAFHEIGESKVVLSNQLEHSKLLKVHCRSKDDDLGEHILKIG  60

Query  267  DPPQTWRFKDAFFHETQFICNLHQGYHWAHHRSFIAYKSSMNTSQKNNANATWFAGEKGI  446
               + + F D  +  T F C + QG ++ HH  F+AY++S   S+   A+  W   E GI
Sbjct  61   QDYE-FTFGDNIWQTTSFSCQMDQGPNFKHHLDFVAYETSW--SKALEASCKWIGREDGI  117

Query  447  YLSFNQRTPEFMY  485
            Y S +   P   Y
Sbjct  118  YFSQDGVPPTKKY  130


>hypothetical protein AXX17_AT5G03750 [Arabidopsis thaliana]
Sequence ID: OAO95751.1 Length: 130
>unnamed protein product [Arabidopsis thaliana]
Sequence ID: CAA0400593.1 Length: 130 >unnamed protein product [Arabidopsis thaliana]
Sequence ID: VYS65834.1 Length: 130
Range 1: 1 to 120

Score:56.2 bits(134), Expect:2e-09,
Method:Compositional matrix adjust.,
Identities:41/124(33%), Positives:57/124(45%), Gaps:4/124(3%)

Query  102  MKRLTIVLIVLVFYIGDTHGRNVLMIVNRLPKNATLKLHCYSGDDDFKTMYLRHNDPPQT  281
            MK L I +I+L   I +T G + L++ N L  N  L + C S DD+     LR     +
Sbjct  1    MKNLFIFVILLSVCIRNTFGISTLLVKNEL-NNKVLGVRCRSKDDNLGDHILRVGQMTKN  59

Query  282  WRFKDAFFHETQFICNLHQGYHWAHHRSFIAYKSSMNTSQKNNANATWFAGEKGIYLSFN  461
              F D  +  T F CNL +G  +  H +F AY+S             W A E GIY + +
Sbjct  60   -NFDDNVWRRTLFWCNLWKGPDFKLHVAFDAYRSQWKADI--GPTYLWIAREDGIYYTQH  116

Query  462  QRTP  473
              TP
Sbjct  117  PETP  120


>Plant self-incompatibility protein S1 family [Arabidopsis thaliana]
Sequence ID: NP_001031830.1 Length: 130
>unknown protein [Arabidopsis thaliana]
Sequence ID: ABF59392.1 Length: 130 >Plant self-incompatibility protein S1 family [Arabidopsis thaliana]
Sequence ID: AED90730.1 Length: 130
Range 1: 1 to 120

Score:56.2 bits(134), Expect:2e-09,
Method:Compositional matrix adjust.,
Identities:41/124(33%), Positives:57/124(45%), Gaps:4/124(3%)

Query  102  MKRLTIVLIVLVFYIGDTHGRNVLMIVNRLPKNATLKLHCYSGDDDFKTMYLRHNDPPQT  281
            MK L I +I+L   I +T G + L++ N L  N  L + C S DD+     LR     +
Sbjct  1    MKNLFIFVILLSVCIRNTFGISTLLVKNEL-NNKVLGVRCRSKDDNLGDHILRVGQMTKN  59

Query  282  WRFKDAFFHETQFICNLHQGYHWAHHRSFIAYKSSMNTSQKNNANATWFAGEKGIYLSFN  461
              F D  +  T F CNL +G  +  H +F AY+S             W A E GIY + +
Sbjct  60   -NFDDNVWRRTLFWCNLWKGPDFKLHVAFDAYRSQWKADI--GPRYLWIAREDGIYYTQH  116

Query  462  QRTP  473
              TP
Sbjct  117  PETP  120


>hypothetical protein AXX17_AT2G05160 [Arabidopsis thaliana]
Sequence ID: OAP10771.1 Length: 135
Range 1: 1 to 135

Score:55.1 bits(131), Expect:7e-09,
Method:Compositional matrix adjust.,
Identities:38/136(28%), Positives:59/136(43%), Gaps:7/136(5%)

Query  102  MKRLTIVLIVLVFYIGDTHG-----RNVLMIVNRLPKN-ATLKLHCYSGDDDFKTMYLRH  263
            M  L ++LI++   +G  +G     +N L   N   +N   L +HC S DDD   ++
Sbjct  1    MNNLFVLLIIIALSVGSNNGSKLWPKNQLHFRNSFSRNYDVLTVHCKSKDDDLG-IHTVA  59

Query  264  NDPPQTWRFKDAFFHETQFICNLHQGYHWAHHRSFIAYKSSMNTSQKNNANATWFAGEKG  443
                  ++F+D+ F  T+F C L  G    +  +F AYK+             W A + G
Sbjct  60   RSYEYNFKFEDSVFGRTEFFCTLMHGVGSKYSVTFTAYKAKPAFVASTGVIKIWDALDDG  119

Query  444  IYLSFNQRTPEFMYIW  491
            IYL+        MY W
Sbjct  120  IYLTDEDHDFVKMYGW  135


>unnamed protein product [Arabidopsis thaliana]
Sequence ID: VYS52148.1 Length: 135
Range 1: 1 to 135

Score:53.5 bits(127), Expect:2e-08,
Method:Compositional matrix adjust.,
Identities:37/136(27%), Positives:58/136(42%), Gaps:7/136(5%)

Query  102  MKRLTIVLIVLVFYIGDTHG-----RNVLMIVNRLPKN-ATLKLHCYSGDDDFKTMYLRH  263
            M  L ++LI++   +G  +G     +N +   N   +N   L +HC S DDD   ++
Sbjct  1    MNNLFVLLIIIALSVGSNNGSKLWPKNQIHFRNSFSRNYDVLTVHCKSKDDDLG-IHTVA  59

Query  264  NDPPQTWRFKDAFFHETQFICNLHQGYHWAHHRSFIAYKSSMNTSQKNNANATWFAGEKG  443
                  ++F D+ F  T+F C L  G    +  +F AYK+             W A + G
Sbjct  60   RSYEYNFKFDDSVFGRTEFFCTLMHGVGSKYSVTFTAYKAKPAFVASTGVIKIWDALDDG  119

Query  444  IYLSFNQRTPEFMYIW  491
            IYL+        MY W
Sbjct  120  IYLTDEDHDFVKMYGW  135


>Plant self-incompatibility protein S1 family [Arabidopsis thaliana]
Sequence ID: NP_178663.1 Length: 135
>putative S1 self-incompatibility protein [Arabidopsis thaliana]
Sequence ID: AAD19763.1 Length: 135 >Plant self-incompatibility protein S1 family [Arabidopsis thaliana]
Sequence ID: AEC06001.1 Length: 135
Range 1: 1 to 135

Score:53.5 bits(127), Expect:2e-08,
Method:Compositional matrix adjust.,
Identities:37/136(27%), Positives:59/136(43%), Gaps:7/136(5%)

Query  102  MKRLTIVLIVLVFYIGDTHG-----RNVLMIVNRLPKN-ATLKLHCYSGDDDFKTMYLRH  263
            M  L ++LI++   +G  +G     +N L   N   +N   L +HC S DDD   ++
Sbjct  1    MNNLFVLLIIIALSVGSNNGSKLWPKNQLHFRNSFSRNYDVLTVHCKSKDDDLG-IHTVA  59

Query  264  NDPPQTWRFKDAFFHETQFICNLHQGYHWAHHRSFIAYKSSMNTSQKNNANATWFAGEKG  443
                  ++F+D+ F  T+F C L  G    +  +F AYK+             W A + G
Sbjct  60   RSYEYNFKFEDSVFGRTEFFCTLMHGVGSKYSVTFTAYKAKPAFVASTGVIKIWDALDDG  119

Query  444  IYLSFNQRTPEFMYIW  491
            IYL+        +Y W
Sbjct  120  IYLTDEDHDFVKIYGW  135


>unnamed protein product [Arabidopsis thaliana]
Sequence ID: CAA0383810.1 Length: 135
Range 1: 1 to 128

Score:52.0 bits(123), Expect:8e-08,
Method:Compositional matrix adjust.,
Identities:40/135(30%), Positives:60/135(44%), Gaps:14/135(10%)

Query  102  MKRLTIVLIVLVFYIGDTHGRNVLMIVNRLPKNATLKLHCYSGDDDFKTMYLRHNDPPQT  281
            MK L+I+L+     +G      +  I N L     L + CYS DD    + +    P
Sbjct  1    MKNLSILLVCSFCILGHVSSAGI-RIGNELKNKKLLWMRCYSKDDVIGPLII----PIGG  55

Query  282  WRFK---DAFFHETQFICNLHQGYHWAHHRSFIAYKSSMNTSQKNNANATWFAGEKGIYL  452
             RF       F  T+F+C L QG ++ HH++F A+K  + ++  +     W A E GIYL
Sbjct  56   HRFNYFGTNIFATTRFMCTLRQGPNYRHHQNFTAFK--LYSASDDGGVWDWRAREDGIYL  113

Query  453  SFNQR----TPEFMY  485
                      P FM+
Sbjct  114  KIKAERGVNRPVFMH  128


>Plant self-incompatibility protein S1 family [Arabidopsis thaliana]
Sequence ID: NP_001330171.1 Length: 160
>Plant self-incompatibility protein S1 family [Arabidopsis thaliana]
Sequence ID: ANM68413.1 Length: 160
Range 1: 34 to 150

Score:52.4 bits(124), Expect:9e-08,
Method:Compositional matrix adjust.,
Identities:39/121(32%), Positives:55/121(45%), Gaps:4/121(3%)

Query  111  LTIVLIVLVFYIGDTHGRNVLMIVNRLPKNATLKLHCYSGDDDFKTMYLRHNDPPQTWRF  290
            L I +I+L   I +T G + L++ N L  N  L + C S DD+     LR     +   F
Sbjct  34   LFIFVILLSVCIRNTFGISTLLVKNEL-NNKVLGVRCRSKDDNLGDHILRVGQMTKN-NF  91

Query  291  KDAFFHETQFICNLHQGYHWAHHRSFIAYKSSMNTSQKNNANATWFAGEKGIYLSFNQRT  470
             D  +  T F CNL +G  +  H +F AY+S             W A E GIY + +  T
Sbjct  92   DDNVWRRTLFWCNLWKGPDFKLHVAFDAYRSQWKADI--GPRYLWIAREDGIYYTQHPET  149

Query  471  P  473
            P
Sbjct  150  P  150


>hypothetical protein AXX17_AT2G18790 [Arabidopsis thaliana]
Sequence ID: OAP07766.1 Length: 147
Range 1: 25 to 135

Score:52.4 bits(124), Expect:9e-08,
Method:Compositional matrix adjust.,
Identities:39/118(33%), Positives:55/118(46%), Gaps:14/118(11%)

Query  162  RNVLMIVNRLPKNATLKLHCYSGDDDFKTMYLRHNDPPQTWRFKDAFFHE----TQFICN  329
            +N ++  + L     L++HC S  DD   +YLRH    QT+ F    FH+    T F C
Sbjct  25   KNFIIFKSSLGPKKLLRIHCTSEHDDTDYVYLRHG---QTYAFS---FHDSVLKTIFDCE  78

Query  330  LHQG---YHWAHHRSFIAYKSSMNTSQKNNANATWFAGEKGIYLSFNQRTPEFMYIWM  494
            L QG   Y++  +  F AYK           N  W A E GIY +  + TP+  Y W+
Sbjct  79   LKQGSSYYNYNFYARFRAYKGGGLIVHYGKKN-FWDAREDGIYFTHGKETPKLEYKWI  135


Query #275: XLOC_032471 Query ID: lcl|Query_30766 Length: 770

No significant similarity found.


Query #276: XLOC_032472 Query ID: lcl|Query_30767 Length: 553

No significant similarity found.


Query #277: XLOC_032473 Query ID: lcl|Query_30768 Length: 309

No significant similarity found.


Query #278: XLOC_032474 Query ID: lcl|Query_30769 Length: 627

No significant similarity found.


Query #279: XLOC_032475 Query ID: lcl|Query_30770 Length: 1400

No significant similarity found.


Query #280: XLOC_032476 Query ID: lcl|Query_30771 Length: 1819

No significant similarity found.


Query #281: XLOC_032477 Query ID: lcl|Query_30772 Length: 1392

No significant similarity found.


Query #282: XLOC_032478 Query ID: lcl|Query_30773 Length: 711

No significant similarity found.


Query #283: XLOC_032480 Query ID: lcl|Query_30774 Length: 1382

Sequences producing significant alignments:
                                                                  Max    Total Query   E   Per.
Description                                                       Score  Score cover Value Ident  Accession
unnamed protein product [Arabidopsis thaliana]                    76.3   76.3  8%    5e-14 82.93  CAA0385633.1
Tetratricopeptide repeat (TPR)-like superfamily protein...        73.6   73.6  8%    2e-13 80.49  NP_566944.1
unnamed protein product [Arabidopsis thaliana]                    73.6   73.6  8%    2e-13 80.49  VYS60035.1
crooked neck-like protein [Arabidopsis thaliana]                  73.9   73.9  8%    2e-13 80.49  CAB62633.1
cell cycle control crn (crooked neck) protein-like [Arabidopsi... 65.1   65.1  8%    2e-10 78.38  BAB10652.1
unnamed protein product [Arabidopsis thaliana]                    64.7   64.7  8%    2e-10 78.38  CAA0406845.1
cell cycle control crn (crooked neck) protein-like [Arabidopsi... 64.7   64.7  8%    2e-10 78.38  AAN72051.1
crooked neck protein, putative / cell cycle protein [Arabidops... 64.7   64.7  8%    2e-10 78.38  NP_198992.2
hypothetical protein AXX17_AT5G39580 [Arabidopsis thaliana]       64.7   64.7  8%    2e-10 78.38  OAO91098.1
crooked neck protein, putative / cell cycle protein [Arabidops... 62.4   62.4  8%    1e-09 65.85  NP_187927.1
probable cell cycle control protein; crooked neck-like protein... 62.4   62.4  8%    1e-09 65.85  BAB01413.1
unnamed protein product [Arabidopsis thaliana]                    62.4   62.4  8%    1e-09 72.97  VYS57206.1
hypothetical protein AXX17_AT3G13370 [Arabidopsis thaliana]       62.0   62.0  8%    2e-09 72.97  OAP04794.1
unnamed protein product [Arabidopsis thaliana]                    62.0   62.0  8%    2e-09 72.97  CAA0382256.1
crooked neck protein, putative / cell cycle protein [Arabidops... 57.4   57.4  15%   5e-08 45.21  NP_199411.1
crooked neck protein, putative / cell cycle protein [Arabidops... 57.4   57.4  15%   5e-08 45.21  NP_001329971.1
unnamed protein product [Arabidopsis thaliana]                    57.4   57.4  15%   5e-08 45.21  CAA0407888.1
hypothetical protein AXX17_AT5G44430 [Arabidopsis thaliana]       57.4   57.4  15%   5e-08 45.21  OAO94715.1
unnamed protein product [Arabidopsis thaliana]                    57.4   57.4  15%   5e-08 45.21  VYS69442.1
unnamed protein product [Arabidopsis thaliana]                    50.4   50.4  5%    9e-08 95.83  BAB09526.1


Alignments:
>unnamed protein product [Arabidopsis thaliana]
Sequence ID: CAA0385633.1 Length: 663
Range 1: 115 to 155

Score:76.3 bits(186), Expect:5e-14,
Method:Compositional matrix adjust.,
Identities:34/41(83%), Positives:36/41(87%), Gaps:0/41(0%)

Query  751  MKLWNKDVNHARNVWD*AVKILPHVDHFWYKYIHMEEILGN  629
             ++ NKDVNHARNVWD AVKILP VD FWYKYIHMEEILGN
Sbjct  115  FEMRNKDVNHARNVWDRAVKILPRVDQFWYKYIHMEEILGN  155


>Tetratricopeptide repeat (TPR)-like superfamily protein [Arabidopsis thaliana]
Sequence ID: NP_566944.1 Length: 413
>Tetratricopeptide repeat (TPR)-like superfamily protein [Arabidopsis thaliana]
Sequence ID: AEE78752.1 Length: 413
Range 1: 115 to 155

Score:73.6 bits(179), Expect:2e-13,
Method:Compositional matrix adjust.,
Identities:33/41(80%), Positives:35/41(85%), Gaps:0/41(0%)

Query  751  MKLWNKDVNHARNVWD*AVKILPHVDHFWYKYIHMEEILGN  629
             ++ NK VNHARNVWD AVKILP VD FWYKYIHMEEILGN
Sbjct  115  FEMRNKSVNHARNVWDRAVKILPRVDQFWYKYIHMEEILGN  155


>unnamed protein product [Arabidopsis thaliana]
Sequence ID: VYS60035.1 Length: 413
Range 1: 115 to 155

Score:73.6 bits(179), Expect:2e-13,
Method:Compositional matrix adjust.,
Identities:33/41(80%), Positives:35/41(85%), Gaps:0/41(0%)

Query  751  MKLWNKDVNHARNVWD*AVKILPHVDHFWYKYIHMEEILGN  629
             ++ NK VNHARNVWD AVKILP VD FWYKYIHMEEILGN
Sbjct  115  FEMRNKSVNHARNVWDRAVKILPRVDQFWYKYIHMEEILGN  155


>crooked neck-like protein [Arabidopsis thaliana]
Sequence ID: CAB62633.1 Length: 599
Range 1: 109 to 149

Score:73.9 bits(180), Expect:2e-13,
Method:Compositional matrix adjust.,
Identities:33/41(80%), Positives:35/41(85%), Gaps:0/41(0%)

Query  751  MKLWNKDVNHARNVWD*AVKILPHVDHFWYKYIHMEEILGN  629
             ++ NK VNHARNVWD AVKILP VD FWYKYIHMEEILGN
Sbjct  109  FEMRNKSVNHARNVWDRAVKILPRVDQFWYKYIHMEEILGN  149


>cell cycle control crn (crooked neck) protein-like [Arabidopsis thaliana]
Sequence ID: BAB10652.1 Length: 665
Range 1: 125 to 161

Score:65.1 bits(157), Expect:2e-10,
Method:Compositional matrix adjust.,
Identities:29/37(78%), Positives:30/37(81%), Gaps:0/37(0%)

Query  739  NKDVNHARNVWD*AVKILPHVDHFWYKYIHMEEILGN  629
            NK VN ARNVWD AV +LP VD  WYKYIHMEEILGN
Sbjct  125  NKFVNSARNVWDRAVTLLPRVDQLWYKYIHMEEILGN  161


>unnamed protein product [Arabidopsis thaliana]
Sequence ID: CAA0406845.1 Length: 705
Range 1: 139 to 175

Score:64.7 bits(156), Expect:2e-10,
Method:Compositional matrix adjust.,
Identities:29/37(78%), Positives:30/37(81%), Gaps:0/37(0%)

Query  739  NKDVNHARNVWD*AVKILPHVDHFWYKYIHMEEILGN  629
            NK VN ARNVWD AV +LP VD  WYKYIHMEEILGN
Sbjct  139  NKFVNSARNVWDRAVTLLPRVDQLWYKYIHMEEILGN  175


>cell cycle control crn (crooked neck) protein-like [Arabidopsis thaliana]
Sequence ID: AAN72051.1 Length: 705
Range 1: 139 to 175

Score:64.7 bits(156), Expect:2e-10,
Method:Compositional matrix adjust.,
Identities:29/37(78%), Positives:30/37(81%), Gaps:0/37(0%)

Query  739  NKDVNHARNVWD*AVKILPHVDHFWYKYIHMEEILGN  629
            NK VN ARNVWD AV +LP VD  WYKYIHMEEILGN
Sbjct  139  NKFVNSARNVWDRAVTLLPRVDQLWYKYIHMEEILGN  175


>crooked neck protein, putative / cell cycle protein [Arabidopsis thaliana]
Sequence ID: NP_198992.2 Length: 705
>crooked neck protein, putative / cell cycle protein [Arabidopsis thaliana]
Sequence ID: AED94726.1 Length: 705
Range 1: 139 to 175

Score:64.7 bits(156), Expect:2e-10,
Method:Compositional matrix adjust.,
Identities:29/37(78%), Positives:30/37(81%), Gaps:0/37(0%)

Query  739  NKDVNHARNVWD*AVKILPHVDHFWYKYIHMEEILGN  629
            NK VN ARNVWD AV +LP VD  WYKYIHMEEILGN
Sbjct  139  NKFVNSARNVWDRAVTLLPRVDQLWYKYIHMEEILGN  175


>hypothetical protein AXX17_AT5G39580 [Arabidopsis thaliana]
Sequence ID: OAO91098.1 Length: 705
>unnamed protein product [Arabidopsis thaliana]
Sequence ID: VYS68969.1 Length: 705
Range 1: 139 to 175

Score:64.7 bits(156), Expect:2e-10,
Method:Compositional matrix adjust.,
Identities:29/37(78%), Positives:30/37(81%), Gaps:0/37(0%)

Query  739  NKDVNHARNVWD*AVKILPHVDHFWYKYIHMEEILGN  629
            NK VN ARNVWD AV +LP VD  WYKYIHMEEILGN
Sbjct  139  NKFVNSARNVWDRAVTLLPRVDQLWYKYIHMEEILGN  175


>crooked neck protein, putative / cell cycle protein [Arabidopsis thaliana]
Sequence ID: NP_187927.1 Length: 657
>crooked neck protein, putative / cell cycle protein [Arabidopsis thaliana]
Sequence ID: AEE75311.1 Length: 657
Range 1: 100 to 140

Score:62.4 bits(150), Expect:1e-09,
Method:Compositional matrix adjust.,
Identities:27/41(66%), Positives:31/41(75%), Gaps:0/41(0%)

Query  751  MKLWNKDVNHARNVWD*AVKILPHVDHFWYKYIHMEEILGN  629
             ++ NK VN ARNVWD AV +LP VD  WYK+IHMEE LGN
Sbjct  100  FEMKNKSVNEARNVWDRAVSLLPRVDQLWYKFIHMEEKLGN  140


>probable cell cycle control protein; crooked neck-like protein [Arabidopsis thaliana]
Sequence ID: BAB01413.1 Length: 675
Range 1: 118 to 158

Score:62.4 bits(150), Expect:1e-09,
Method:Compositional matrix adjust.,
Identities:27/41(66%), Positives:31/41(75%), Gaps:0/41(0%)

Query  751  MKLWNKDVNHARNVWD*AVKILPHVDHFWYKYIHMEEILGN  629
             ++ NK VN ARNVWD AV +LP VD  WYK+IHMEE LGN
Sbjct  118  FEMKNKSVNEARNVWDRAVSLLPRVDQLWYKFIHMEEKLGN  158


>unnamed protein product [Arabidopsis thaliana]
Sequence ID: VYS57206.1 Length: 681
Range 1: 139 to 175

Score:62.4 bits(150), Expect:1e-09,
Method:Compositional matrix adjust.,
Identities:27/37(73%), Positives:29/37(78%), Gaps:0/37(0%)

Query  739  NKDVNHARNVWD*AVKILPHVDHFWYKYIHMEEILGN  629
            NK VN ARNVWD AV +LP VD  WYK+IHMEE LGN
Sbjct  139  NKSVNEARNVWDRAVSLLPRVDQLWYKFIHMEEKLGN  175


>hypothetical protein AXX17_AT3G13370 [Arabidopsis thaliana]
Sequence ID: OAP04794.1 Length: 692
Range 1: 139 to 175

Score:62.0 bits(149), Expect:2e-09,
Method:Compositional matrix adjust.,
Identities:27/37(73%), Positives:29/37(78%), Gaps:0/37(0%)

Query  739  NKDVNHARNVWD*AVKILPHVDHFWYKYIHMEEILGN  629
            NK VN ARNVWD AV +LP VD  WYK+IHMEE LGN
Sbjct  139  NKSVNEARNVWDRAVSLLPRVDQLWYKFIHMEEKLGN  175


>unnamed protein product [Arabidopsis thaliana]
Sequence ID: CAA0382256.1 Length: 692
Range 1: 139 to 175

Score:62.0 bits(149), Expect:2e-09,
Method:Compositional matrix adjust.,
Identities:27/37(73%), Positives:29/37(78%), Gaps:0/37(0%)

Query  739  NKDVNHARNVWD*AVKILPHVDHFWYKYIHMEEILGN  629
            NK VN ARNVWD AV +LP VD  WYK+IHMEE LGN
Sbjct  139  NKSVNEARNVWDRAVSLLPRVDQLWYKFIHMEEKLGN  175


>crooked neck protein, putative / cell cycle protein [Arabidopsis thaliana]
Sequence ID: NP_199411.1 Length: 673
>crooked neck protein, putative / cell cycle protein [Arabidopsis thaliana]
Sequence ID: AED95324.1 Length: 673 >CRN (crooked neck) protein [Arabidopsis thaliana]
Sequence ID: BAB08244.1 Length: 673
Range 1: 125 to 197

Score:57.4 bits(137), Expect:5e-08,
Method:Compositional matrix adjust.,
Identities:33/73(45%), Positives:44/73(60%), Gaps:1/73(1%)

Query  739  NKDVNHARNVWD*AVKILPHVDHFWYKYIHMEEILGNRRS*KD-G*RWKN*DFEIKQWKK  563
            NK VN+ARNVWD +V +LP VD  W KYI+MEE LGN    +    RW N   + K W
Sbjct  125  NKFVNNARNVWDRSVTLLPRVDQLWEKYIYMEEKLGNVTGARQIFERWMNWSPDQKAWLC  184

Query  562  MIKLEISKSNLKK  524
             IK E+  + +++
Sbjct  185  FIKFELRYNEIER  197


>crooked neck protein, putative / cell cycle protein [Arabidopsis thaliana]
Sequence ID: NP_001329971.1 Length: 683
>crooked neck protein, putative / cell cycle protein [Arabidopsis thaliana]
Sequence ID: ANM68198.1 Length: 683
Range 1: 135 to 207

Score:57.4 bits(137), Expect:5e-08,
Method:Compositional matrix adjust.,
Identities:33/73(45%), Positives:44/73(60%), Gaps:1/73(1%)

Query  739  NKDVNHARNVWD*AVKILPHVDHFWYKYIHMEEILGNRRS*KD-G*RWKN*DFEIKQWKK  563
            NK VN+ARNVWD +V +LP VD  W KYI+MEE LGN    +    RW N   + K W
Sbjct  135  NKFVNNARNVWDRSVTLLPRVDQLWEKYIYMEEKLGNVTGARQIFERWMNWSPDQKAWLC  194

Query  562  MIKLEISKSNLKK  524
             IK E+  + +++
Sbjct  195  FIKFELRYNEIER  207


>unnamed protein product [Arabidopsis thaliana]
Sequence ID: CAA0407888.1 Length: 683
Range 1: 135 to 207

Score:57.4 bits(137), Expect:5e-08,
Method:Compositional matrix adjust.,
Identities:33/73(45%), Positives:44/73(60%), Gaps:1/73(1%)

Query  739  NKDVNHARNVWD*AVKILPHVDHFWYKYIHMEEILGNRRS*KD-G*RWKN*DFEIKQWKK  563
            NK VN+ARNVWD +V +LP VD  W KYI+MEE LGN    +    RW N   + K W
Sbjct  135  NKFVNNARNVWDRSVTLLPRVDQLWEKYIYMEEKLGNVTGARQIFERWMNWSPDQKAWLC  194

Query  562  MIKLEISKSNLKK  524
             IK E+  + +++
Sbjct  195  FIKFELRYNEIER  207


>hypothetical protein AXX17_AT5G44430 [Arabidopsis thaliana]
Sequence ID: OAO94715.1 Length: 683
Range 1: 135 to 207

Score:57.4 bits(137), Expect:5e-08,
Method:Compositional matrix adjust.,
Identities:33/73(45%), Positives:44/73(60%), Gaps:1/73(1%)

Query  739  NKDVNHARNVWD*AVKILPHVDHFWYKYIHMEEILGNRRS*KD-G*RWKN*DFEIKQWKK  563
            NK VN+ARNVWD +V +LP VD  W KYI+MEE LGN    +    RW N   + K W
Sbjct  135  NKFVNNARNVWDRSVTLLPRVDQLWEKYIYMEEKLGNVTGARQIFERWMNWSPDQKAWLC  194

Query  562  MIKLEISKSNLKK  524
             IK E+  + +++
Sbjct  195  FIKFELRYNEIER  207


>unnamed protein product [Arabidopsis thaliana]
Sequence ID: VYS69442.1 Length: 683
Range 1: 135 to 207

Score:57.4 bits(137), Expect:5e-08,
Method:Compositional matrix adjust.,
Identities:33/73(45%), Positives:44/73(60%), Gaps:1/73(1%)

Query  739  NKDVNHARNVWD*AVKILPHVDHFWYKYIHMEEILGNRRS*KD-G*RWKN*DFEIKQWKK  563
            NK VN+ARNVWD +V +LP VD  W KYI+MEE LGN    +    RW N   + K W
Sbjct  135  NKFVNNARNVWDRSVTLLPRVDQLWEKYIYMEEKLGNVTGARQIFERWMNWSPDQKAWLC  194

Query  562  MIKLEISKSNLKK  524
             IK E+  + +++
Sbjct  195  FIKFELRYNEIER  207


>unnamed protein product [Arabidopsis thaliana]
Sequence ID: BAB09526.1 Length: 65
Range 1: 1 to 24

Score:50.4 bits(119), Expect:9e-08,
Method:Compositional matrix adjust.,
Identities:23/24(96%), Positives:23/24(95%), Gaps:0/24(0%)

Query  74  MDKFALEREMKGGGSVSFLYFSDY  3
           MDKFALEREMKGGGSVSFLYFS Y
Sbjct  1   MDKFALEREMKGGGSVSFLYFSGY  24


Query #284: XLOC_032481 Query ID: lcl|Query_30775 Length: 390

No significant similarity found.


Query #285: XLOC_032482 Query ID: lcl|Query_30776 Length: 857

No significant similarity found.


Query #286: XLOC_032483 Query ID: lcl|Query_30777 Length: 641

No significant similarity found.


Query #287: XLOC_032484 Query ID: lcl|Query_30778 Length: 700

No significant similarity found.


Query #288: XLOC_032485 Query ID: lcl|Query_30779 Length: 482

No significant similarity found.


Query #289: XLOC_032486 Query ID: lcl|Query_30780 Length: 871

No significant similarity found.


Query #290: XLOC_032487 Query ID: lcl|Query_30781 Length: 509

No significant similarity found.


Query #291: XLOC_032488 Query ID: lcl|Query_30782 Length: 851

No significant similarity found.


Query #292: XLOC_032489 Query ID: lcl|Query_30783 Length: 484

No significant similarity found.


Query #293: XLOC_032490 Query ID: lcl|Query_30784 Length: 578

No significant similarity found.


Query #294: XLOC_032491 Query ID: lcl|Query_30785 Length: 602

No significant similarity found.


Query #295: XLOC_032492 Query ID: lcl|Query_30786 Length: 1054

No significant similarity found.


Query #296: XLOC_032493 Query ID: lcl|Query_30787 Length: 501

No significant similarity found.


Query #297: XLOC_032494 Query ID: lcl|Query_30788 Length: 550

No significant similarity found.


Query #298: XLOC_032495 Query ID: lcl|Query_30789 Length: 774

No significant similarity found.


Query #299: XLOC_032496 Query ID: lcl|Query_30790 Length: 806

No significant similarity found.


Query #300: XLOC_032497ATTCCACATAACAGTATATCGTGCTATAACCACATTTAGTGGAATTAATGGTCATTGACCTAATATTCTCTCAATTTATAGCGACACACCAAATTATCCTTTTTTAATCTTTTAGATTTCTTCATGGTGCTGATTTTTAGTTCACTTTTATCCATAAGTTACTACTAAATTTTTCTGTTCTACGTACAAATGACACATAAGATTGATAAGCTTTTAGATTTATAGAAAGATGATACTTGCATCGTCAATATGGGAGGAGCAATGACTAATCGATGTAAATCCGACATCTTCTAATTTTTCGTCTAAATAAATTATTGCATTTTGATGTTTTTTTAGATTAGGAAAAAAAAAACCAAACTTTTGGTCCCTAATTTTCTCGAGATGTCCATAACAAGTTAAATCAGCCCTCTTCCCACCGTTGAGATTCAAACTCACGTACGATATTGAAGAACTTTTTCTCTAGGCGATCCCCAATTCAGCTACTTGAGGGGTTATAATTACTTTGACGTTTCTTGTTTGATTAACCCATAATTTGTTGTTAAAGCTTGTTGGATTTAAATTATTTTGGCCATACCCTAAAATATGGTTGGTCCTTGGTCCCCCTCATAAAAGGTTACAAGAATTTAAAGAGAAGTAGA Query ID: lcl|Query_30791 Length: 0

No significant similarity found.


Query #301: XLOC_032498 Query ID: lcl|Query_74289 Length: 851

No significant similarity found.


Query #302: XLOC_032499 Query ID: lcl|Query_74290 Length: 651

No significant similarity found.


Query #303: XLOC_032500 Query ID: lcl|Query_74291 Length: 281

No significant similarity found.


Query #304: XLOC_032501 Query ID: lcl|Query_74292 Length: 478

No significant similarity found.


Query #305: XLOC_032502 Query ID: lcl|Query_74293 Length: 711

No significant similarity found.


Query #306: XLOC_032503 Query ID: lcl|Query_74294 Length: 823

No significant similarity found.


Query #307: XLOC_032504 Query ID: lcl|Query_74295 Length: 802

No significant similarity found.


Query #308: XLOC_032505 Query ID: lcl|Query_74296 Length: 794

Sequences producing significant alignments:
                                                                  Max    Total Query   E   Per.
Description                                                       Score  Score cover Value Ident  Accession
Plant invertase/pectin methylesterase inhibitor superfamily...    97.4   97.4  18%   1e-23 100.00 NP_176006.1
unnamed protein product [Arabidopsis thaliana]                    82.8   154   26%   4e-18 100.00 BAB08248.1
hypothetical protein AXX17_AT4G13350 [Arabidopsis thaliana]       71.6   71.6  19%   2e-13 76.47  OAO98795.1
hypothetical protein AXX17_AT4G13350 [Arabidopsis thaliana]       71.6   71.6  19%   2e-13 76.47  OAO98794.1
FAM63A-like protein (DUF544) [Arabidopsis thaliana]               71.6   71.6  19%   2e-13 76.47  NP_567383.1
putative protein [Arabidopsis thaliana]                           71.2   71.2  19%   2e-13 76.47  CAB44324.1


Alignments:
>Plant invertase/pectin methylesterase inhibitor superfamily protein [Arabidopsis thaliana]
Sequence ID: OAO93173.1 Length: 232
>Plant invertase/pectin methylesterase inhibitor superfamily protein [Arabidopsis thaliana]
Sequence ID: AEE33342.1 Length: 232
Range 1: 184 to 232

Score:97.4 bits(241), Expect:1e-23,
Method:Compositional matrix adjust.,
Identities:49/49(100%), Positives:49/49(100%), Gaps:0/49(0%)

Query  296  ESLSSITMKSEDVNHDQLPSKQSGLETVRDVENASSSKKAIVDVTSSEA  442
            ESLSSITMKSEDVNHDQLPSKQSGLETVRDVENASSSKKAIVDVTSSEA
Sbjct  184  ESLSSITMKSEDVNHDQLPSKQSGLETVRDVENASSSKKAIVDVTSSEA  232


>unnamed protein product [Arabidopsis thaliana]
Sequence ID: BAB08248.1 Length: 238
Range 1: 1 to 40

Score:82.8 bits(203), Expect:4e-18,
Method:Compositional matrix adjust.,
Identities:40/40(100%), Positives:40/40(100%), Gaps:0/40(0%)

Query  471  MLGHSSMDNYASDEVTSTMAFLLLLAFSTSRTVSSPDCLE  352
            MLGHSSMDNYASDEVTSTMAFLLLLAFSTSRTVSSPDCLE
Sbjct  1    MLGHSSMDNYASDEVTSTMAFLLLLAFSTSRTVSSPDCLE  40


Range 2: 40 to 70

Score:71.2 bits(173), Expect:4e-14,
Method:Compositional matrix adjust.,
Identities:29/31(94%), Positives:31/31(100%), Gaps:0/31(0%)

Query  153  KEKEIKTMEKRVERENEEKALWFRGQFGNFD  61
            +E+EIKTMEKRVERENEEKALWFRGQFGNFD
Sbjct  40   EEREIKTMEKRVERENEEKALWFRGQFGNFD  70


>hypothetical protein AXX17_AT4G13350 [Arabidopsis thaliana]
Sequence ID: OAO98795.1 Length: 657
Range 1: 331 to 380

Score:71.6 bits(174), Expect:2e-13,
Method:Compositional matrix adjust.,
Identities:39/51(76%), Positives:44/51(86%), Gaps:1/51(1%)

Query  305  SSITMKSEDVNHDQLPSKQSGLETVRDVENASSSKKAIVDVTSSEA*LSID  457
            ++IT KSED+NHDQL SKQSG ET  DVEN SSSK+AIVDVTSSEA LS+D
Sbjct  331  TTITTKSEDLNHDQLSSKQSGGETACDVENVSSSKEAIVDVTSSEA-LSVD  380


>hypothetical protein AXX17_AT4G13350 [Arabidopsis thaliana]
Sequence ID: OAO98794.1 Length: 682
Range 1: 331 to 380

Score:71.6 bits(174), Expect:2e-13,
Method:Compositional matrix adjust.,
Identities:39/51(76%), Positives:44/51(86%), Gaps:1/51(1%)

Query  305  SSITMKSEDVNHDQLPSKQSGLETVRDVENASSSKKAIVDVTSSEA*LSID  457
            ++IT KSED+NHDQL SKQSG ET  DVEN SSSK+AIVDVTSSEA LS+D
Sbjct  331  TTITTKSEDLNHDQLSSKQSGGETACDVENVSSSKEAIVDVTSSEA-LSVD  380


>FAM63A-like protein (DUF544) [Arabidopsis thaliana]
Sequence ID: NP_567383.1 Length: 682
>unknown [Arabidopsis thaliana]
Sequence ID: AAM62819.1 Length: 682 >FAM63A-like protein (DUF544) [Arabidopsis thaliana]
Sequence ID: AEE83059.1 Length: 682 >unnamed protein product [Arabidopsis thaliana]
Sequence ID: CAA0394802.1 Length: 682 >unnamed protein product [Arabidopsis thaliana]
Sequence ID: VYS62360.1 Length: 682 >hypothetical protein [Arabidopsis thaliana]
Sequence ID: BAF01280.1 Length: 682
Range 1: 331 to 380

Score:71.6 bits(174), Expect:2e-13,
Method:Compositional matrix adjust.,
Identities:39/51(76%), Positives:44/51(86%), Gaps:1/51(1%)

Query  305  SSITMKSEDVNHDQLPSKQSGLETVRDVENASSSKKAIVDVTSSEA*LSID  457
            ++IT KSED+NHDQL SKQSG ET  DVEN SSSK+AIVDVTSSEA LS+D
Sbjct  331  TTITTKSEDLNHDQLSSKQSGGETACDVENVSSSKEAIVDVTSSEA-LSVD  380


>putative protein [Arabidopsis thaliana]
Sequence ID: CAB44324.1 Length: 626
>putative protein [Arabidopsis thaliana]
Sequence ID: CAB78229.1 Length: 626
Range 1: 331 to 380

Score:71.2 bits(173), Expect:2e-13,
Method:Compositional matrix adjust.,
Identities:39/51(76%), Positives:44/51(86%), Gaps:1/51(1%)

Query  305  SSITMKSEDVNHDQLPSKQSGLETVRDVENASSSKKAIVDVTSSEA*LSID  457
            ++IT KSED+NHDQL SKQSG ET  DVEN SSSK+AIVDVTSSEA LS+D
Sbjct  331  TTITTKSEDLNHDQLSSKQSGGETACDVENVSSSKEAIVDVTSSEA-LSVD  380


Query #309: XLOC_032506 Query ID: lcl|Query_74297 Length: 583

No significant similarity found.


Query #310: XLOC_032507 Query ID: lcl|Query_74298 Length: 780

No significant similarity found.


Query #311: XLOC_032508 Query ID: lcl|Query_74299 Length: 462

No significant similarity found.


Query #312: XLOC_032557 Query ID: lcl|Query_74300 Length: 481

No significant similarity found.
```
