## Supplemental Doc 3 for "Characterization of novel pollen-expressed transcripts reveals their potential roles in pollen heat stress response in *Arabidopsis thaliana*" (from another version of this preprint: version 2, 2020-08-26)

**XLOC_030751** genomic fragment

CTATGTTTCTTTTTTTTTTAGTTTTCCTATTTATTTAAGAGAAAAGTAAATATTTGTTGACACATGTCAACATCTGAACGATAGACTTGACGCGTGTCACAATCACACGAGTTAGTATTTTAAAAACCATATTTTATATAATAAGAAGATATATGAACTCATATTGTATTATAACATTACACAATACTCTAAACAAAATTAGACTTAGTTCATAAGATGCAATAAATGTCAATTACATAAACATTGGTTACTTTTGGAAAAAGAGAAGCCACCAATTAATTTTTGTTCCAAGATAATACGCGTACCAATGGTGTATTATATTTTCTTGCAAAGTATATCCCATCAATTCTGGCAGCCCATACGCGTAGCTGGCCACATCGAGGAATAAGTCTTGGTCCTGCTTTGTATACTTCAATGTCGTATGAATACGAATTGTTAGGTCCTTGCCTAAAAATACAATTCCATCTTGTTAAATTGGGTATTTCATCATGGAATCTAATGACAAATGGAGTGGCATTGAAGTTTAATTGTTTTACGCCAAGATTGTCATCTTCGGAACGACAATGATATTCGAGAATTTGACCAGGAGCGAGTTGATTAAGAATCTCGACTATATTTTTTATGCAAGCTTGATTTAGCCCAAAATACGTAATAGTAATCAAAATTACAAATACAGAATGGTTCATTTTTTGGATCTAAACCCTTTATTGAAGTAACAATGACATCAACATGTAGTATTTATAATTATTTTGTGTGGAAGACGTTTTTTCCTAAAATTACTAGATAAGCGTTTTATGGTTTTAGCTAAATCGAAAC

Reverse complement

**TCONS_00047695 (XLOC_030751 minus strand,** At5g36985)

GTTTCGATTTAGCTAAAACCATAAAACGCTTATCTAGTAATTTTAGGAAAAAACGTCTTCCACACAAAATAATTATAAATACTACATGTTGATGTCATTGTTACTTCAATAAAGGGTTTAGATCCAAAAA**ATG**AACCATTCTGTATTTGTAATTTTGATTACTATTACGTATTTTGGGCTAAATCAAGCTTGCATAAAAAATATAGTCGAGATTCTTAATCAACTCGCTCCTGGTCAAATTCTCGAATATCATTGTCGTTCCGAAGATGACAATCTTGGCGTAAAACAATTAAACTTCAATGCCACTCCATTTGTCATTAGATTCCATGATGAAATACCCAATTTAACAAGATGGAATTGTATTTTTAGGCAAGGACCTAACAATTCGTATTCATACGACATTGAAGTATACAAAGCAGGACCAAGACTTATTCCTCGATGTGGCCAGCTACGCGTATGGGCTGCCAGAATTGATGGGATATACTTTGCAAGAAAATATAATACACCATTGGTACGCGTATTATCTTGGAACAAAAATTAATTGGTGGCTTCTCTTTTTCCAAAAGTAACCAATGTTTATGTAATTGACATTTATTGCATCTTATGAACTAAGTCTAATTTTGTTTAGAGTATTGTGTAATGTTATAATACAATATGAGTTCATATATCTTCTTATTATATAAAATATGGTTTTTAAAATACTAACTCGTGTGATTGTGACACGCGTCAAGTCTATCGTTCAGATGTTGACATGTGTCAACAAATATTTACTTTTCTCTTAAATAAATAGGAAAACTAAAAAAAAAAGAAACATAG
